# Effects of true to life polyethylene terephthalate and polycaprolactone nanoparticles on macrophages under a repeated exposure mode

**DOI:** 10.1101/2025.11.08.687361

**Authors:** Véronique Collin-Faure, Aliro Villacorta, Hélène Diemer, Sarah Cianférani, Ricard Marcos, Alba Hernandez, Marie Carrière, Elisabeth Darrouzet, Thierry Rabilloud

## Abstract

Micro and nanoplastics are pollutants which concentration in different biotopes increases continuously over time, which poses the question of their potential effects on health. In animals, these micro and nanoplastics are recognized as particulate materials and thus handled by macrophages, either directly in the case of lung exposure, or after the particles have crossed the epithelial barriers in case of oral or dermal exposure. It is thus important to study the potential effects of micro and nanoplastics on macrophages. Most studies have used an experimental scheme in which the cells of interest are exposed to a single dose of plastics, and where the readout of the studied parameters is made immediately after exposure. However, this classical experimental scheme does not take into account the impact of biopersistence, nor the potential cellular adaptation that may take place when cells are exposed repeatedly to a low dose of plastics. We thus used a repeated exposure scheme, in order to better take into account these phenomena. Within this frame, we compared the macrophages responses to a persistent nanoplastic, i.e. true-to-life polyethylene terephthalate nanoparticles and to a biodegradable nanoplastic, i.e. polycaprolactone, by a combination of proteomic and targeted experiments. Our results show that under this repeated exposure scheme, the proteome changes were of a lesser extent than under the acute exposure mode, indicating cell adaptation. However, polyethylene terephthalate nanoparticles induced oxidative stress and a pro-inflammatory response, while polycaprolactone nanoparticles induced a depression of macrophages functions, indicating harmful effects even in the repeated exposure scheme.

## 1. Introduction

Because of their technical qualities and ease of use, plastics are produced nowadays in enormous quantities, close to 500Mt/year worldwide ^1^. Unfortunately, because of poor waste handling, an important amount of plastics is continuously released in the environment, the sole release into oceans being estimated to 10 Mt/year ^2^, leading to an estimated accumulation in the oceans of 150 Mt in 2025 ^3^. Although plastic pollution was first documented in marine environments ^4–7^, it has now been documented in marine sediments ^8,9^, freshwater environments ^10–12^ and terrestrial ones ^13,14^.

Part of the technical qualities of plastics lies in their chemical resistance, which however translates into very long half lives in the environment, amounting to decades ^15^. This long chemical half-life does not mean that plastics do not degrade. They do fragment first into microplastics (less than 1mm in size) and then into nanoplastics (less than 1µm in size). From knowledge gained on other type of particles, it is predicted that the smaller the plastic particles, the more easily they will cross the biological barriers and the heavier their interference with biological functions will be. This hypothesis has been documented for at least the intestinal barrier (e.g. in ^16^). In animal organisms, once the barriers have been crossed, a second line of defense comes into play, represented by the phagocytes. This cell types is encountered in invertebrates, from the worms’ coelomocytes to the insects’ hemocytes ^17^. It also occurs in vertebrates in the form of macrophages and neutrophils, and it has been demonstrated that these cells respond to nanoplastics ^18–22^.

In view of these adverse effects, the development of biodegradable plastics, which will not release such long-lasting particles, is a promising and highly active avenue of research. Among these biodegradable plastics and beside the ones based on polysaccharides, poly aliphatic hydroxyesters such as polylactide or poly hydroxy butyrate/valerate have been extensively proposed, as well as polycaprolactone (PCL), because of its attractive properties ^23^. PCL uses a single monomer, usually obtained from fossil sources but that can be biosourced ^24^, which is polymerized by ring opening, which allows a precise control of polymerization. The polymer combines a good degradability ^25^, e.g. though composting ^26,27^, or even in seawater ^28^, with a slow degradation in mammals, so that it has been extensively used in orthopedics and implants ^29–31^. However, its technical qualities have driven renewed interest for this polymer for applications in membranes ^32^ and in packaging ^33–38^.

In view of the relatively slow degradation of PCL in macrophages ^39^, compared for example to the faster degradation of polylactide ^40^, we decided to investigate the macrophage responses to PCL nanoparticles, but in a repeated exposure mode, and not in the acute exposure mode that is used in most nanotoxicology experiments on plastics (e.g. in ^20–22,39,40^). Repeated exposure is more representative of the everyday exposure that occurs in a plastic-contaminated environment, and it has been shown that repeated exposure reveals cellular responses to nanoplastics that are not shown in the acute exposure mode^41–43^.

In order to better investigate the responses to a biodegradable plastic, we decided to compare the response to PCL nanoparticles to the ones induced by a repeated exposure to a non-biodegradable plastic. To this purpose, we decided to use true-to-life poly(ethylene terephthalate) nanoparticles (PET), obtained by processing of plastic water bottles ^44,45^. Such plastic bottles have been shown to release particles upon normal use ^46–48^, so that the PET particles are an excellent proxy to true-to-life plastic exposure. Moreover, these particles have been used in repeated exposure schemes in other cell types ^43,49,50^.

We thus decided to investigate the responses of macrophages to PET and PCL nanoparticles under a repeated exposure scheme, using a combination of proteomic and targeted approaches, as previously done under acute exposure with the same PET ^51^ and PCL ^39^ nanoparticles, in order to gain a wide appraisal of the cellular responses to these plastics in the selected repeated exposure scheme.

## 2. Materials and Methods

### 2.1. Particles

The preparation and characterization of the particles have been described previously for the PET ^51^ and the PCL particles ^39^, as well as their labelling with the fluorescent dye Disperse Blue 14 (1,4-Bis(methylamino)anthraquinone, ABCR #AB177338, λ ex 640nm, λ em 685 nm) by the solvent swelling method ^52^. Briefly, a 10mg/ml solution of Disperse Blue 14 in tetrahydrofuran (THF) was prepared. 1% in volume of this solution and 10% THF were added to the nanoparticles suspension, and the mixture was agitated on a rotary wheel at room temperature for 30 minutes. Two volumes of water were then added and the particles collected by centrifugation (30 minutes at 15,000g). The pellet was then resuspended in water and the beads recollected by centrifugation. The final pellet was then resuspended in 20% ethanol. Both PET and PCL particles show an average size around 200 nm ^39,51^.

### 2.2. Cell culture and exposure to particles

The J774A.1 cell line (mouse macrophages) was purchased from European cell culture collection (Salisbury, UK). Cells were routinely propagated in DMEM supplemented with 10% fetal bovine serum (FBS) in non-adherent flasks (Cellstar flasks for suspension culture, Greiner Bio One, Les Ulis, France). For routine culture, the cells were seeded at 200,000 cells/ml and split two days later, with a cell density ranging from 800,000 to 1,000,000 cells/ml.

For exposure to plastic particles and to limit the effects of cell growth, cells were seeded at 500,000 cells/ml in 6 or 12 wells adherent plates in DMEM supplemented with 1% horse serum ^53^. The cells were let to settle and recover for 24 h, and then exposed to the selected nanoplastics at 10µg/ml and per day, with a medium renewal every two days. This exposure was carried out for 8 days, plus a final 24 hours period. Proteomic experiments were carried out in 6 well plates, and all the other experiments in 12 well plates. Cells were used at passage numbers from 5 to 15 post-reception from the repository. Cell viability was measured by the propidium iodide method ^54^, or with the SytoxGreen probe (Thermofisher S7020) using the protocol provided by the supplier.

The internalized dose was estimated as followed: cells cultured in 12 wells adherent plates, were exposed to fluorescent particles (PET or PCL) for 8 days as described previously (cumulated dose 80µg/ml). To measure the cells associated fluorescence, the medium was removed, the cell layer was then rinsed once with 1ml PBS and the PBS was removed. The cell layer was then lysed by the addition of 400µl of 10mM Hepes pH 7.5, and the cell lysate was collected. Its fluorescence was the measured using a DeNovix QFX instrument (excitation at 635 nm, emission collected in the 665-740 nm window). Untreated cell cultures of matched age were used to determine the cell autofluorescence background. In order to determine the reference value, the cumulated dose of the particles (i.e. 80µg/ml) was added into 400µl of 10mM Hepes pH 7.5, and the fluorescence measured as for the cell extracts. This protocol is by construction independent of cell division, as each well is considered as a whole.

### 2.3. Proteomics

Proteomics was carried out essentially as described previously ^39^., and will not be further detailed in this article.

### 2.4. Data analysis

For the global analysis of the protein abundances data, missing data were imputed with a low, non-null value. Proteins that were detected less than 3 times out of 5 in both groups were removed from the analysis. The complete abundance dataset was then analyzed by the PAST software ^55^.

Proteins were considered as significantly different if their p value in the Mann-Whitney U-test against control values was inferior to 0.05. No quantitative change threshold value was applied. The selected proteins were then submitted to pathway analysis using the DAVID tool ^56^, with a cutoff value set at a FDR of 0.25.

### 2.5. Mitochondrial transmembrane potential assay

The mitochondrial transmembrane potential assay was performed essentially as described previously ^57^. Rhodamine 123 (Rh123) was added to the cultures at an 80 nM final concentration (to avoid quenching ^58^), and the cultures were further incubated at 37°C for 30 minutes. At the end of this period, the cells were collected, washed in cold PBS containing 0.1% glucose, resuspended in PBS glucose and analyzed for the green fluorescence (excitation 488 nm emission 525nm) on a Melody flow cytometer. As a negative control, carbonyl cyanide 4-(trifluoromethoxy)phenylhydrazone (FCCP) was added at 5µM final concentration together with the Rh123 ^59^.

### 2.6. Assay for oxidative stress

For the oxidative stress assay, a protocol based on the oxidation of dihydrorhodamine 123 (DHR123) was used, essentially as described previously ^57^. After exposure to plastic beads as described in section 2.2, the cells were treated in PBS containing 500 ng/ml DHR123 for 20 minutes at 37°C. The cells were then harvested, washed in cold PBS containing 0.1% glucose, resuspended in PBS glucose and analyzed for the green fluorescence (same parameters as rhodamine 123) on a Melody flow cytometer. Menadione (applied on the cells for 2 hours prior to treatment with DHR123) was used as a positive control in a concentration range of 25-50µM.

### 2.7. Lysosomal assay

For the lysosomal function assay, the Lysosensor method was used, as described previously ^57^. After exposure to nanoplastics as described in section 2.2, the medium was removed, the cell layer was rinsed with complete culture medium and incubated with 1µM Lysosensor Green (Molecular Probes) diluted in warm (37°C) complete culture medium for 1 hour at 37°C. At the end of this period, the cells were collected, washed in cold PBS containing 0.1% glucose, resuspended in PBS glucose and analyzed for the green fluorescence (excitation 488 nm emission 540nm) on a Melody flow cytometer.

### 2.8. Phagocytosis assay

For this assay, the cells were first exposed to the PCL or PET nanoparticles as described in section 2.2. Twenty four hours after the final daily exposure to the plastics, the cells were then exposed to 0.5 µm latex beads (carboxylated surface, yellow green-labelled, from Polysciences excitation 488 nm emission 527/32 nm) for 3 hours. After this second exposure, the cells were collected, rinsed twice with PBS, and analyzed for the two fluorescences (green and red) on a Melody flow cytometer.

### 2.9. Cell surface markers

Cells were seeded into 12-well plates at a concentration of 500,000 cells/ml, and exposed to the nanoplastic particles as described in section 2.2. The day following the last addition of nanoplastics, the cells were harvested and washed in DMEM containing 3% FBS. For all labelled antibodies, the cells were treated with the fluorochrome-conjugated antibodies at the adequate dilution for 30 minutes on ice in DMEM-FBS in a final volume of 100µl. For collectin 12, the cells were first treated with the primary antibody at the adequate dilution for 30 minutes on ice in DMEM-FBS in a final volume of 100µl. The cells were then collected by centrifugation, resuspended in 100 µl DMEM.FBS and treated for 30 minutes on ice with the secondary antibody (Rabbit Anti-Goat igG, FITC conjugated, Invitrogen #31509, dilution 1/300)

In all cases, the cells were then washed with 2ml of PBS. The cell pellet was resuspended in 400 µl of PBS containing 1 µM Sytox blue for checking cell viability, and the suspension analyzed by flow cytometry on a Melody flow cytometer. First, live cells (Sytox blue-negative) were selected at 450 nm (excitation at 405 nm). The gated cells were then analyzed for the nanoplastics fluorescence (excitation 561nm emission 695 nm) and for the antibody fluorescence (excitation 488 nm emission 527 nm for FITC or A488 labelled antibodies or excitation 561 nm emission 582 nm for PE-conjugated antibodies). Isotypic control antibodies were used to compensate for non-specific binding.

The following antibodies were used:

Collectin-12 (Invitrogen #PA5-47456) dilution 1/6

CD86-FITC (BD-Pharmingen #553-691) dilution 1/25

CD204-A488 (Invitrogen #53-2046-82) dilution 1/25

PD-L1-A488 (BD-Pharmingen #566-864) dilution 1/25

TLR2-FITC (Invitrogen# 11-9021-82) dilution 1/25

TLR7-PE ( (BD-Pharmingen #565-557) dilution 1/50

In the experiments with TLR7, the cells were collected after exposure to the nanoplastics, washed with PBS, and then fixed and permeabilized with the BD cytofix-cytoperm kit (#554715). The cells were incubated with the anti-TLR7 antibody diluted to 1/50 (respectively) for 30 minutes on ice. The cells were then washed with the permeabilization buffer, resuspended in PBS and analyzed by flow cytometry as described above.

### 2.10. Cytokine release assays

Cells were first exposed to nanoplastics as described in section 2.2. At the end of this exposure period, i.e. 24 hours after the last addition of beads, the culture medium was collected and analyzed for proinflammatory cytokines. In half of the wells LPS (1 ng/ml) was added 18 hours before medium collection. Tumor necrosis factor (catalog number 558299, BD Biosciences, Le Pont de Claix, France), MCP1 (catalog number 558342, BD Biosciences, Le Pont de Claix, France)and interleukin 6 (IL-6) (catalog number 558301, BD Biosciences, Le Pont de Claix) levels were measured using the Cytometric Bead Array Mouse Inflammation Kit (catalog number 558266, BD Biosciences, Le Pont de Claix), and analyzed with FCAP Array software (3.0, BD Biosciences) according to the manufacturer’s instructions.

## 3. Results

### 3.1. Particle internalization

As the physico-chemical characteristics of the particles used have been described previously ^39,51^, we first focus on the total particle internalization during this repeated exposure scheme, using in-house labelled particles. The results (N=4) showed a very high internalization of the PET particles, close to quantitative (99±3%), while the internalization of the PCL particles was much lower (62±5%).

### 3.2. Global analysis of the proteomic results

The proteomic analysis led to the detection and quantification of 3328 proteins (**Table S1**). In order to evaluate the global extent of changes at the proteome scale, we performed a Principal Coordinate analysis, using the PAST software ^55^. The results, displayed on **Figure 1A**, showed an interpenetration of the three groups, indicating very moderate proteome changes. By contrast, the same analysis, performed on proteomic data collected after an acute exposure to the same particles ^39,51^, showed good group separation (**Figure 1B**) indicative of more profound proteome changes.

**Figure 1.**
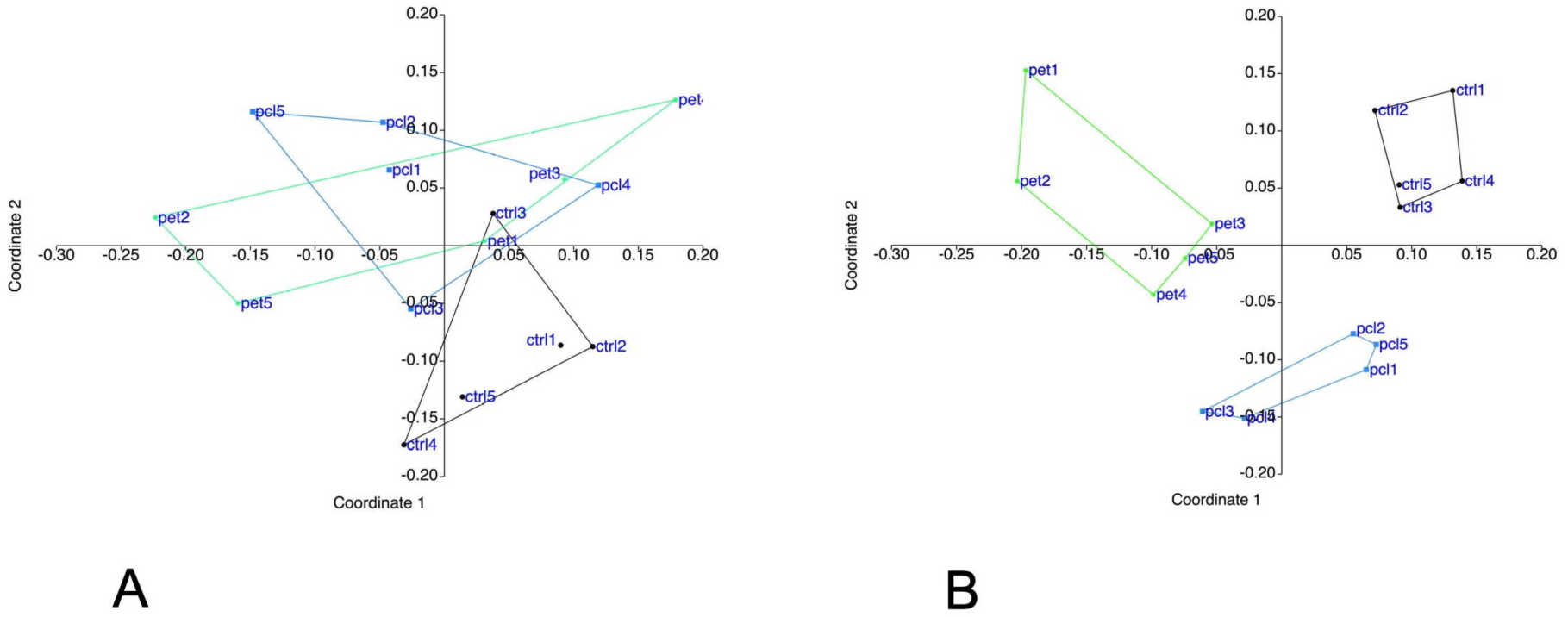
Global analysis of the proteomic data Panel A: The complete proteomic data table (3328 proteins) was analyzed by Principal Coordinates Analysis (Gower distance), using the PAST software. The results are represented as the X-Y diagram of the first two axes of the Principal Coordinate Analysis, representing 40% of the total variance. Eigenvalue scale. Panel B: The same analysis was performed on the datasets obtained in response to acute exposure to the same PET and PCL nanoparticles. the first two axes of the Principal Coordinate Analysis, representing 41% of the total variance. Eigenvalue scale. This representation allows to figure out how, at the global proteome scale, the samples are related to each other. Samples grouped in such a diagram indicate similar proteomes, and the larger the distance between samples are, the more dissimilar their respective proteomes.

Thus, the repeated exposure scheme induce less alteration of the proteome than an acute exposure to the same cumulated dose. This result was corroborated by the ANalysis Of SIMilarity test ^60^, which results are shown on **Table 1**.

**Table 1:**
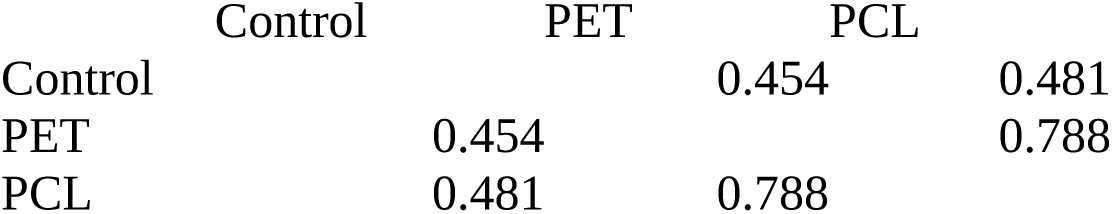
p-values of ANOSIM analysis in the repeated exposure scheme.

This analysis indicates that at the proteome level scale, the changes induced by the repeated exposure to PET and PCL nanoparticles were not statistically significant, and in fact close to the intra-group variability. Once again, this result was in sharp contrast of the one obtained after an acute exposure to the total cumulative dose, as shown on **Table 2**.

**Table 2:**
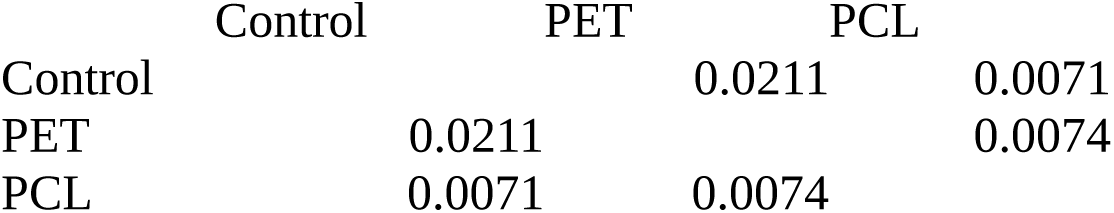
p-values of ANOSIM analysis in the acute exposure scheme.

Thus, the overall conclusion of this global proteome analysis was that repeated exposure induced moderate proteome changes, indicative of a progressive cell adaptation to the nanoplastics.

Nevertheless, it was still possible to select specific proteins which expression was significantly changed in response to the repeated exposure to the particles. To this purpose, we used the Mann-Whitney U test, with a cutoff at p<0.05. This process led to the selection of 254 proteins responding to the treatment with PET particles (**Table S2**) and of 282 proteins responding to the treatment with the PCL particles (**Table S3**). These selected proteins were then submitted to a pathway using the DAVID pathway analysis tool ^61^, and the results are displayed on **Tables S4 and S5**. The pathway analysis pointed out several cellular processes which were modulated upon treatment with plastic particles. Most were generic, e.g. mitochondria, lysosomes or endoplasmic reticulum.

### 3.3. Mitochondria

Mitochondrial proteins represented an abundant class among the proteins modulated in response to nanoplastics, with 83 proteins (**Table S6**). As a large collection of mitochondrial proteins in the particles-responsive proteins may indicate mitochondrial perturbations ^62^, we analyzed the transmembrane mitochondrial potential. The results, displayed in Figure 2, did not show any perturbation of the transmembrane mitochondrial potential in response to repeated exposure to either PET or PCL particles, indicating that the response observed at the proteome level is homeostatic.

**Figure 2:**
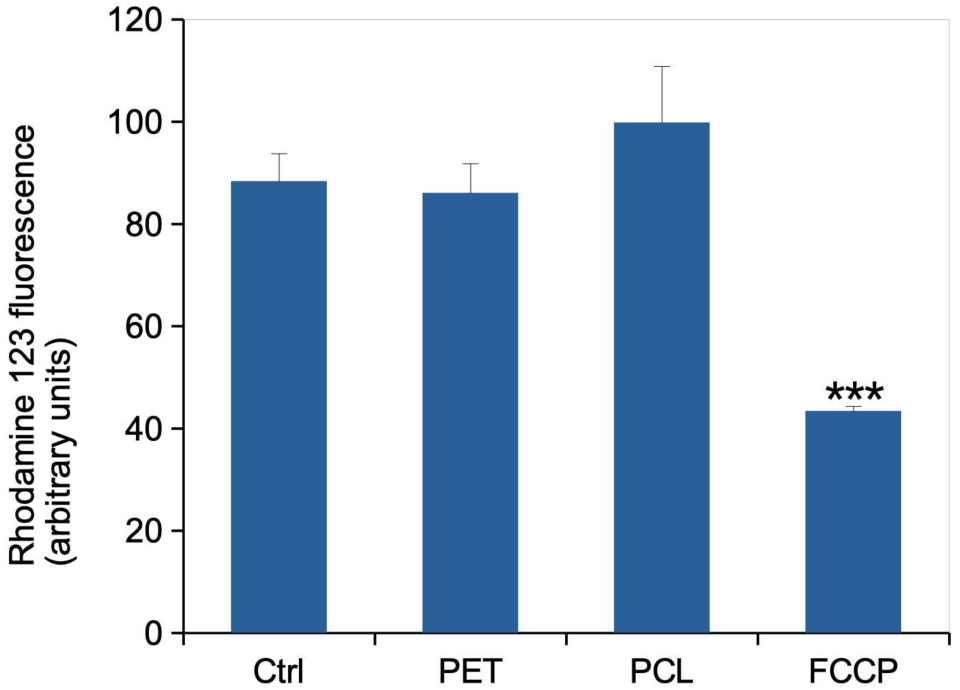
Mitochondrial transmembrane potential (rhodamine 123 method) All cells were positive for rhodamine 123 internalization in mitochondria, and the mean fluorescence is the displayed parameter. Results are displayed as mean± standard deviation (N=4). FCCP: carbonyl cyanide 4-(trifluoromethoxy)phenylhydrazone. Significance marks: ** p≤0.01; *** p≤0.001 (Student t-test method)

### 3.4. Oxidative stress

In our studies on different particles in a high dose, acute exposure mode, we observed that PET and PS increased the level of oxidative stress ^1,2^, while PCL decreased it ^2^. We thus investigated whether these responses also existed in response to a repeated exposure to a lower daily dose. The results, displayed in Figure 3, did not show any significant change in the level of oxidative stress in response to PCL particles. However, an increased oxidative stress level was observed in response to repeated exposure to PET particles.

**Figure 3:**
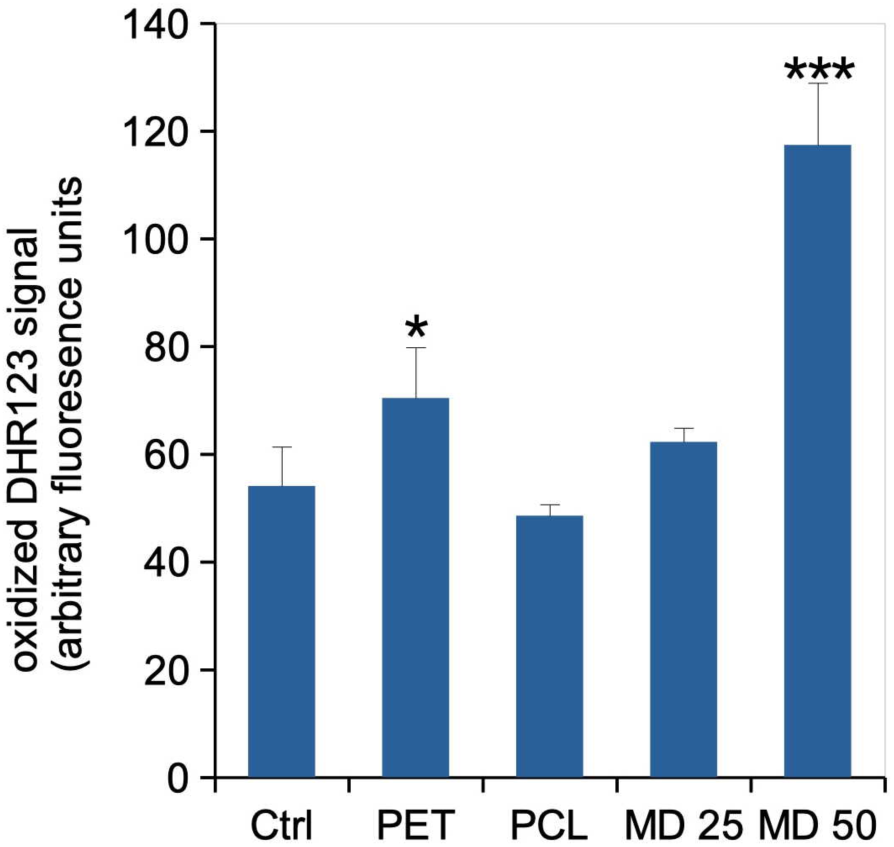
Cellular oxidative stress, measured with the dihydrorhodamine 123 (DHR123) indicator The cells were exposed to nanoplastics as described in section 2.2, and finally for 20 minutes to the DHR 123 probe. Menadione (25 or 50 µg/ml for 2 hours) was used as a positive oxidative stress control. Results are displayed as mean± standard deviation (N=4). Significance marks: * p≤0.05; *** p≤0.001 (Student t-test method). MD: Menadione.

### 3.5 Lysosomes

The proteomic screen led to a list of 26 lysosomal proteins which abundances were modulated in response to repeated exposure to either PET or PCL nanoplastics (**Table S7**). Furthermore, in our studies on different particles in a high dose, acute exposure mode, we observed that PET increased the lysosensor response ^51^, while PCL and PS did not ^39^. We thus investigated whether these responses also existed in response to a repeated exposure to a lower daily dose. The results, displayed in Figure 4, showed an insignificant increase in response to PET particles, and a significant one in response to PCL particles.

**Figure 4:**
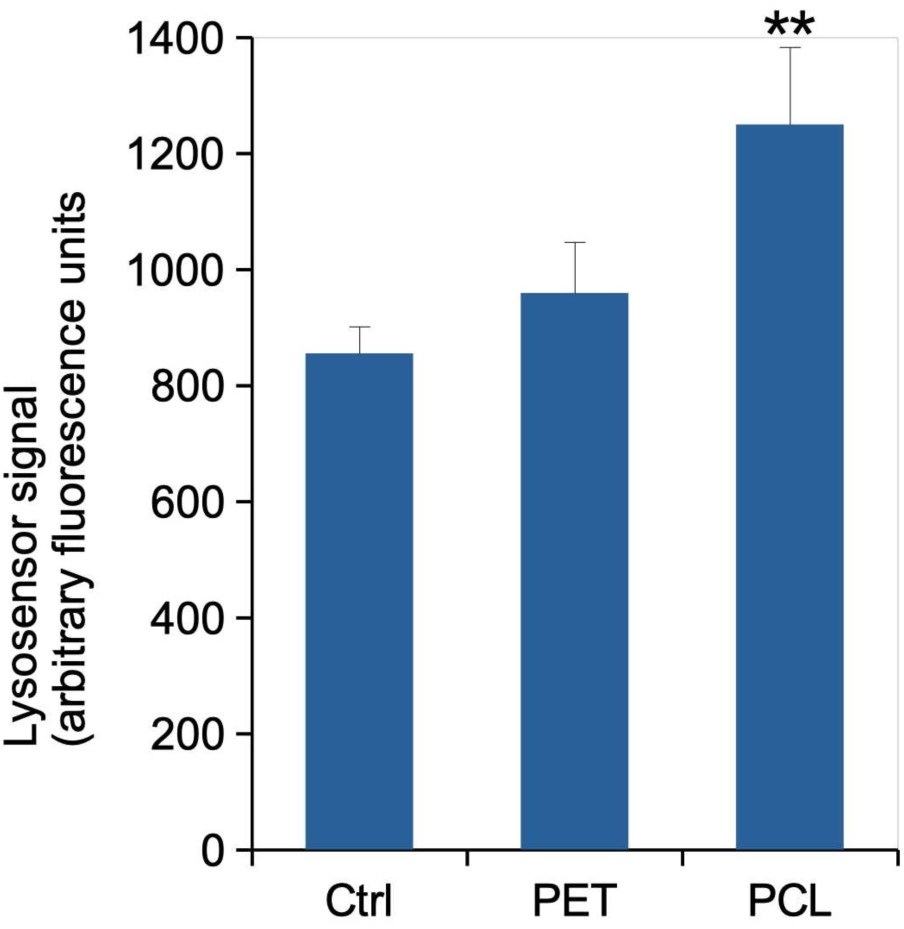
Lysosomal proton pumping (Lysosensor method). All cells were positive for lysosensor internalization in lysosomes, and the mean fluorescence is the displayed parameter. Results are displayed as mean± standard deviation (N=4). Significance marks: ** p≤0.01 (Student t-test method)

### 3.6 Phagocytosis

In a scheme of acute exposure to a high dose of nanoparticles, PET particles did not change the phagocytic capacities of macrophages, while PCL particles depresses phagocytosis ^39,51^. The situation was however different when the same particles were given to macrophages in a repeated exposure scheme. In this case, as shown in Figure 5, a repeated exposure to PET nanoparticles slightly but significantly depressed phagocytosis, while PCL nanoparticles did not show any influence on this specialized macrophage function.

**Figure 5.**
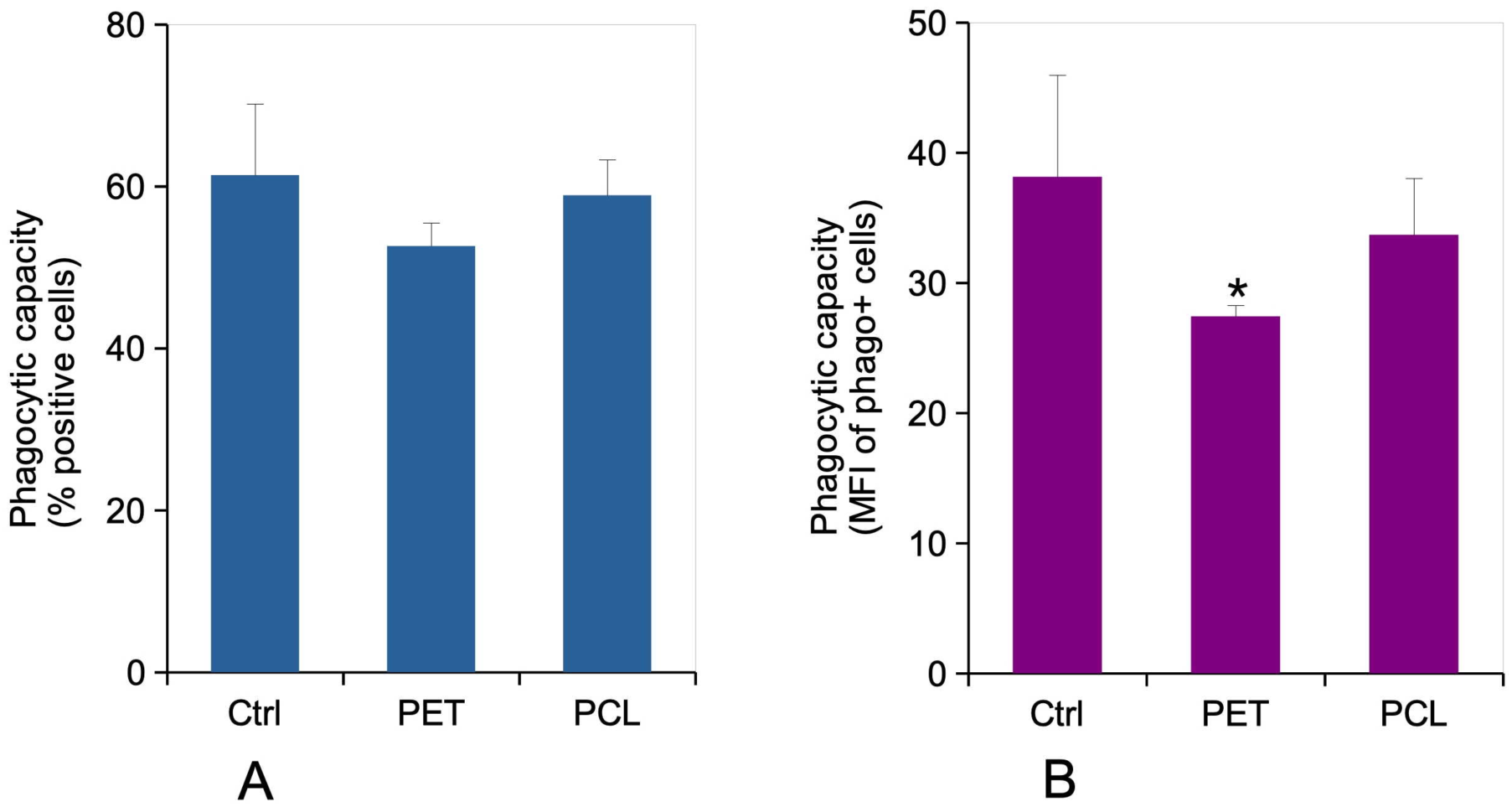
Phagocytic capacity The cells were exposed to nanoplastics as described in section 2.2. After removal of the plastics-containing cell culture medium, the cells were treated with green fluorophore labelled carboxylated polystyrene beads for 3 hours. Panel A: The percentage of green fluorescence-positive cells, indicating the percentage of cells able to internalize the test beads in 3 hours, is the displayed parameter. Results are displayed as mean± standard deviation (N=4). Significance marks: * p≤0.05 (Student t-test method)

### 3.7. Surface markers

The pathway analysis did not highlight immune response as modulated in response to a repeated exposure to PET or PCL nanoplastics. However, detailed analysis of the modulated proteins highlighted proteins implicated in the immune responses, which are listed in **Table S8**. We also included in this table proteins that we knew to be modulated in response to acute exposures, but that did not appear as such in the proteomic screen in response to repeated exposure. However, proteomics measures the total protein concentration, and not the one correctly folded and addressed to the relevant cell membrane, which is the useful parameter for membrane receptors. We thus measured this parameter using antibody labelling and flow cytometry, and the results are displayed on **Table 4**.

**Table 4:**
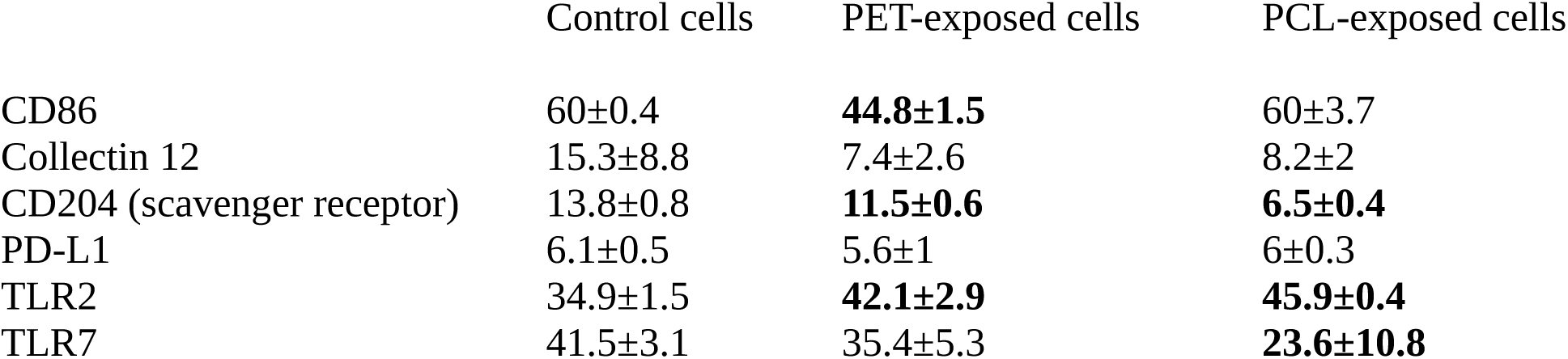
Assay of surface antigens by flow cytometry.

The results are expressed as mean fluorescence ± standard deviation (N=4) Bold figures indicate significant changes (Student T test, p<0.05)

Despite not being detected by the proteomic screen, several of these surface markers were indeed modulated in response to repeated exposure to PET or PCL nanoplastics.

### 3.8. Pro-inflammatory cytokine release

In a scheme of acute exposure to a high dose of nanoparticles, PET particles increased the release of TNF, while PCL particles depressed the release of both TNF and IL-6 after LPS stimulation ^1,2^. We thus investigated this parameter in response to a repeated exposure to the same particles. The results, displayed in Figure 6 showed that PET still showed an intrinsic pro-inflammatory effect (Figure 6A and 6C), and PCL an intrinsic anti-inflammatory effect (Figure 6A,C,E). When the interference of plastic exposure with LPS-induced inflammation was studied, PET and PCL decreased the release of IL-6 (Figure 6B), and PCL also decreased the release of TNF (Figure 6F). case in the acute exposure mode ^51^.

**Figure 6.**
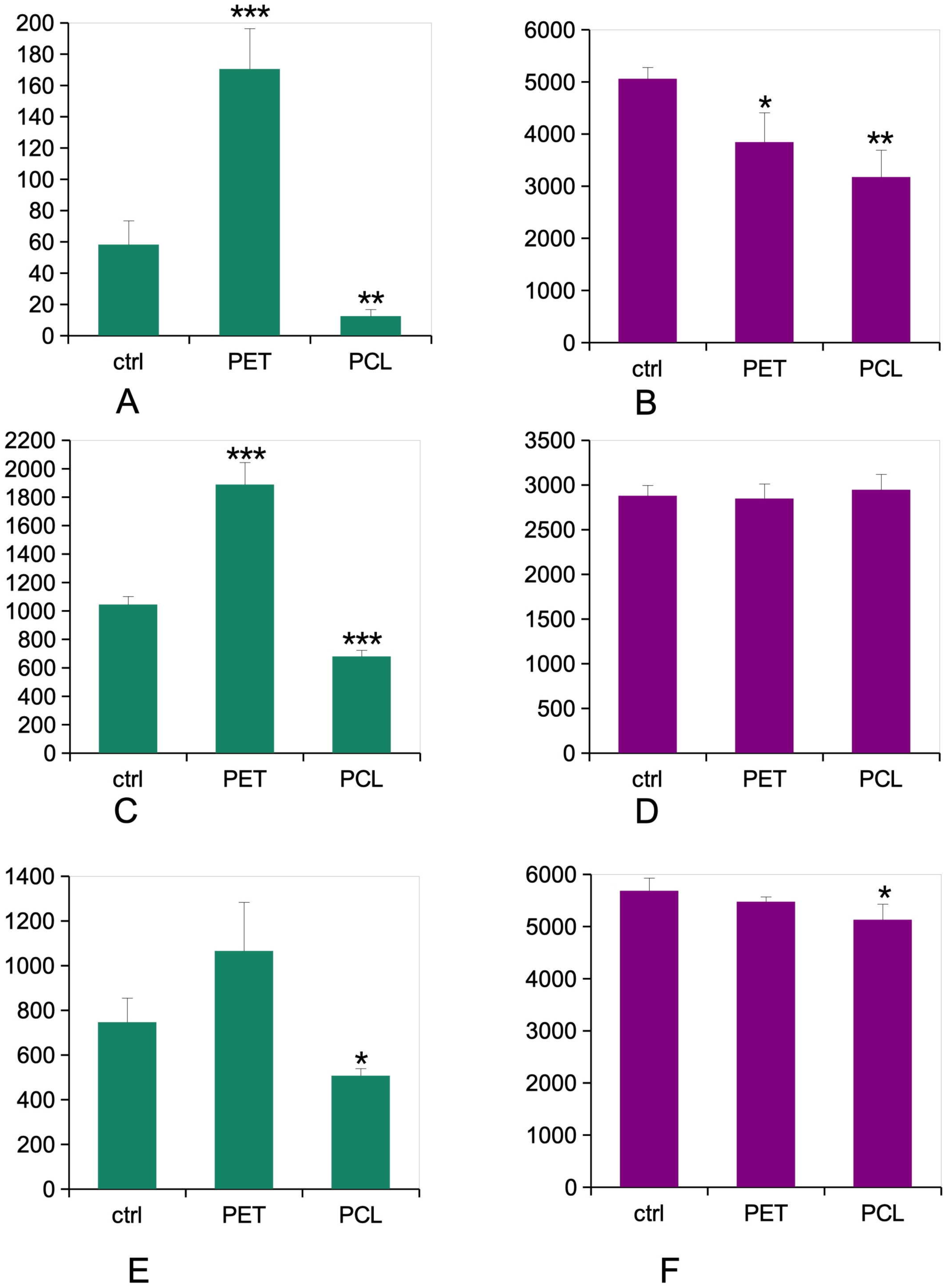

Regarding the pathogens recognition receptors, collectin 12 was decreased in response to both nanoplastics under the repeated exposure mode, but this decrease was not statistically significant, as opposed to our observations under acute exposure ^51^. Oppositely, TLR2 was significantly increased in response to both PET and PCL nanoparticles under the repeated exposure mode, while TLR7 was significantly decreased in response to PCL nanoparticles.

The scavenger receptor CD204 was significantly decreased in response to both nanoplastics, while it was unchanged in the acute exposure mode ^51^. CD204, also known as MSR-1, being strongly involved in absorption of toxic organic particles such as oxidized LDL ^77^, such a decrease may imply a lesser depurating function of plastic-treated macrophages. Moreover, CD204 is also a bacterial and viral receptor ^78^, so that its decrease may sign a lesser activity of macrophages in engulfing bacteria and viruses, which may not be compensated for by the increase in TLR2.

## 5. Conclusions

All in all, the repeated exposure regime induces a macrophage response that is fairly different from the one observed after acute exposure. While the general cell physiology parameters such as the mitochondrial transmembrane potential appear less altered by the repeated exposure regime, the oxidative stress level and the pro-inflammatory response induced by the PET nanoparticles appear higher in this exposure mode, probably because of the persistence of the particles in the cells, which leaves time for the effect to develop. Conversely, for PCL nanoparticles, which slowly dissolve and liberate the anti-inflammatory monomer 6-hydroxyhexanoic, an immunodepressive effect develops in the repeated exposure regime. Thus, widely different but still problematic immunological effects develop in the repeated exposure regime for both PET and PCL nanoparticles.

Following previous in vitro studies dealing with sustained/delayed effects of plastic particles on cells ^42,43,49,50,63^, this study also shows the necessity of carrying this type of studies and not only the acute exposure studies that are commonplace in *in vitro* toxicology, even for toxicants such as micro- or nano-plastics where persistence or oppositely biodegradation are key factors in the biological outcomes. This work also shows that biodegradability shall not be viewed as a panacea against the adverse effects of plastics. Release of particles during the intended period of use, as documented for different plastics (e.g. in ^46,79^), as well as through poor waste handling, liberating particles before the complete degradation of the plastics, can result into adverse effects.

## Acknowledgments

Thanks are due to the Biostudies database for hosting the non-proteomic data associated with this work, under the identifier S-BSST-2171 ^80^ Thanks are also due to the ProteomXchange consortium for hosting the proteomic data associated with this work, under the doi 10.6019/PXD049997 ^81^

## Author Contributions

TR designed and supervised the study. AV, RM, and AH generated and characterized the true-to-life PET nanoplastics. MC characterized the PCL nanoparticles. VCF, HD and SC performed the proteomic experiments, which were then analyzed by TR and ED. VCF performed and interpreted the flow cytometry experiments. TR drafted the initial version of the manuscript, which was completed and amended by all co-authors, who approved the final version of the manuscript.

