## Supplemental Table 1 for "Effects of true to life polyethylene terephthalate and polycaprolactone nanoparticles on macrophages under a repeated exposure mode"

Supplementary Table 1: list of proteins detected and quantified by the proteomic screen

| accession | gene_name | description | protein_set_scoverage | | MW | #specific_seq | #peptides | Ctrl1 rep | Ctrl5 rep | Ctrl4 rep | Ctrl3 rep | Ctrl2 rep | PET1 rep | PET5 rep | PET4 rep | PET3 rep | PET2 rep | PCL1 rep | PCL5 rep | PCL4 rep | PCL3 rep | PCL2 rep |
|---|---|---|---|---|---|---|---|---|---|---|---|---|---|---|---|---|---|---|---|---|---|---|
| P06238 | A2m | Alpha-2-macroglobulin Rattus norvegicus | 52.82 | 0.68 | 163785 | 1 | 1 | 2571648.5 | 1 | 1982401.88 | 808577.31 | 1 | 1013199.25 | 1216453.5 | 1120545.283 | 1097524 | 1155004.38 | 1759660 | 2182541 | 1734879 | 1478423.5 | 1991792.62 |
| P58742 | Aaas | Aladin Mus musculus | 312.2 | 21.61 | 59431 | 7 | 7 | 5700100 | 6476691.5 | 5992078 | 5249330.5 | 6063287 | 5450714 | 5841170.5 | 5929437.5 | 6197640 | 6054818 | 5844015 | 5293274.5 | 6424745.5 | 5803309 | 4928255 |
| Q9D2R0 | Aacs | Acetoacetyl-CoA synthetase Mus musculus | 628.09 | 23.66 | 75200 | 13 | 13 | 19207743 | 19185077 | 16988625 | 13469001 | 15923373 | 12639340 | 14612932 | 13654119 | 16553901 | 15990681 | 13710401 | 17783989 | 13510717 | 14955523 | 14849255 |
| P0C1X8 | Aak1 | AP2-associated protein kinase 1 Rattus norvegicus | 96.52 | 4.47 | 103761 | 2 | 2 | 410858.03 | 1 | 323195.53 | 1 | 310884.16 | 375156.69 | 411203.53 | 375989.06 | 455028.94 | 404344.555 | 328897.81 | 488275.5 | 481075.16 | 428982.665 | 417682.19 |
| Q8BGQ7 | Aars | Alanine--tRNA ligase, cytoplasmic Mus musculus | 2313.64 | 52.58 | 106909 | 31 | 32 | 75010617 | 80832313 | 81785641 | 98410385 | 88170257 | 79499833 | 81959881 | 76829409 | 78118417 | 94648721 | 88267337 | 81786065 | 84647705 | 80079969 | 94373345 |
| Q3THG9 | Aarsd1 | Alanyl-tRNA editing protein Aarsd1 Mus musculus | 299.59 | 27.18 | 44971 | 7 | 7 | 3747308 | 5152381 | 5212448 | 5635507 | 2850704.25 | 4617483 | 6086961 | 5946345 | 5498171 | 6366480 | 4396122 | 5146584 | 5368399.5 | 5231564 | 5198768.5 |
| Q9JKX4 | Aatf | Protein AATF Mus musculus | 219.63 | 9.13 | 59482 | 3 | 3 | 221213.81 | 370021.16 | 1 | 1 | 1 | 288928.28 | 1 | 1 | 1 | 1 | 235313.36 | 438999.94 | 231415.39 | 322051.66 | 538457.25 |
| P06795 | Abcb1b | Multidrug resistance protein 1B Mus musculus | 196.97 | 4.7 | 140994 | 3 | 3 | 1318353.75 | 907364.62 | 1376753.75 | 947343.5 | 625802.44 | 809881.5 | 580928.25 | 708097 | 696975.25 | 698970.5 | 941768.38 | 1225924.88 | 685851.5 | 505225.47 | 1109828.25 |
| Q61102 | Abcb7 | ATP-binding cassette sub-family B member 7, mitochondrial Mus musculus | 373.17 | 16.62 | 82581 | 2 | 6 | 5123448 | 2944719 | 4799254 | 3505946.25 | 3699019.5 | 4194061.5 | 4933149 | 4786500 | 4288263.5 | 5249030.5 | 3926306 | 4187827 | 4663284.5 | 4612658.5 | 3842482.75 |
| O35379 | Abcc1 | Multidrug resistance-associated protein 1 Mus musculus | 205.46 | 3.99 | 171185 | 4 | 4 | 1123700 | 1535258.5 | 1524385 | 1034228.31 | 1184643.88 | 1302136.25 | 1158030.12 | 1537593.75 | 2830235 | 1775848.25 | 2477462 | 1039836.12 | 1367875.12 | 1828197.5 | 1305154.5 |
| B2RX12 | Abcc3 | Canalicular multispecific organic anion transporter 2 Mus musculus | 264.86 | 5.38 | 169124 | 5 | 5 | 2378467.25 | 2236672.25 | 2379280.25 | 2060722.12 | 2223249.25 | 2155500.25 | 1983227.75 | 2221811 | 2908995 | 2065724 | 2231854.5 | 2160865.75 | 2183681.25 | 2439739.75 | 2108289 |
| D3ZHR2 | Abcd1 | ATP-binding cassette sub-family D member 1 Rattus norvegicus | 89.86 | 5.83 | 81938 | 2 | 2 | 1 | 1 | 613164.44 | 1 | 274918.53 | 730180.62 | 1349675.62 | 1 | 585955.31 | 1 | 459625.16 | 635499.75 | 539811.69 | 378621.03 | 798100.38 |
| P55096 | Abcd3 | ATP-binding cassette sub-family D member 3 Mus musculus | 45.76 | 1.82 | 75475 | 1 | 1 | 969135.44 | 753819.81 | 577534.38 | 775170.56 | 586447.5 | 859278.44 | 576717.31 | 1070231.88 | 732113.81 | 647938.75 | 471611.91 | 379563.16 | 757693.06 | 813575.44 | 615576.5 |
| P61222 | Abce1 | ATP-binding cassette sub-family E member 1 Mus musculus | 1117.73 | 41.9 | 67314 | 18 | 18 | 15265587 | 13586266 | 14239910 | 19967685 | 20255783 | 15471098 | 17346577 | 15409282 | 14921260 | 13513835 | 19229563 | 14325224 | 13635647 | 14371328 | 19345295 |
| Q6P542 | Abcf1 | ATP-binding cassette sub-family F member 1 Mus musculus | 464.31 | 19.12 | 94945 | 9 | 9 | 7467059 | 5819978.5 | 6286385.5 | 6161664 | 6544938.5 | 6664042.5 | 7772943 | 6478770 | 6717339 | 6960280 | 7536715.5 | 7737788 | 6782665 | 6396568 | 6845729.5 |
| Q99LE6 | Abcf2 | ATP-binding cassette sub-family F member 2 Mus musculus | 352.77 | 14.97 | 71782 | 8 | 8 | 2561787 | 3042396.75 | 3490268.5 | 2401754.25 | 2664650.5 | 2979684.5 | 4776007.5 | 3350205 | 3611228.75 | 3749946 | 3090189.25 | 3623357.25 | 4097384.75 | 2405138.25 | 4131764.25 |
| Q66H39 | Abcf3 | ATP-binding cassette sub-family F member 3 Rattus norvegicus | 49.48 | 2.96 | 79856 | 1 | 1 | 501458.97 | 405761.75 | 536086.56 | 779650.12 | 269166.16 | 238527.5 | 274571.72 | 1 | 1 | 364780.91 | 1 | 179937.77 | 1 | 545629.31 | 257873.44 |
| Q6AYT7 | Abhd12 | Monoacylglycerol lipase ABHD12 Rattus norvegicus 2 | 471.79 | 31.41 | 45296 | 9 | 9 | 8299984.5 | 5311613 | 9413133 | 7260250 | 5648225 | 8371796 | 10562007 | 8385347.5 | 8586765 | 8391826 | 7520391 | 11725091 | 7737445 | 11513198 | 9554713 |
| Q8VCR7 | Abhd14b | Protein ABHD14B Mus musculus | 102.13 | 15.24 | 22451 | 3 | 3 | 1861727.88 | 3702895 | 1522411.5 | 2582999 | 1558661.25 | 3145184 | 2135136.25 | 1694582.12 | 1229484.38 | 2190810.25 | 1420649.12 | 1658675.25 | 1910665.62 | 1702542.62 | 1139820 |
| Q8CBW3 | Abi1 | Abl interactor 1 Mus musculus | 85.4 | 4.99 | 52288 | 2 | 2 | 4001681.75 | 4286474.5 | 4502418 | 3655020.5 | 4132653.75 | 3932741 | 3846297.75 | 4393578 | 4507604.5 | 2948587.25 | 4471021 | 4392607 | 4157699.5 | 5057310.5 | 3814466.25 |
| Q4JIM5 | Abl2 | Tyrosine-protein kinase ABL2 Mus musculus | 43.38 | 1.52 | 128196 | 1 | 1 | 1 | 251990.64 | 195264.5 | 1 | 1 | 176600.6525 | 129267.35 | 164559.72 | 144373.2 | 268202.34 | 274011.22 | 167950.44 | 204638.825 | 177237.14 | 199356.5 |
| Q5SSL4 | Abr | Active breakpoint cluster region-related protein Mus musculus | 413.1 | 12.81 | 97667 | 8 | 8 | 3828630 | 4065722.5 | 4025891.75 | 1980865 | 4399927.5 | 5660986 | 3625264.25 | 4083371.75 | 7713215.5 | 3323895 | 6216541 | 4869163.5 | 4647166 | 6893983.5 | 6553381 |
| Q4KML4 | Abracl | Costars family protein ABRACL Mus musculus | 186.33 | 46.91 | 9030 | 3 | 3 | 29953261 | 21856413 | 18335557 | 21333597 | 19702327 | 21155317 | 17380759 | 20485889 | 22174239 | 16067721 | 19034755 | 19075671 | 18686881 | 21290243 | 17282871 |
| Q3TCJ1 | Abraxas2 | BRISC complex subunit Abraxas 2 Mus musculus | 49.2 | 2.89 | 46943 | 1 | 1 | 2015625 | 2341368 | 2138161.5 | 1699554 | 1894750.38 | 1653930.75 | 1803644.25 | 1574490.38 | 1780650 | 2010838 | 1713165.38 | 1651209 | 1774636.25 | 1798356 | 1480457 |
| Q921H8 | Acaa1a | 3-ketoacyl-CoA thiolase A, peroxisomal Mus musculus | 923.52 | 52.12 | 43953 | 13 | 13 | 42564881 | 40742409 | 43531681 | 41126677 | 34604249 | 39649481 | 42479985 | 44810045 | 40629481 | 41423605 | 43403857 | 40834881 | 45656185 | 43682189 | 40089721 |
| Q8BWT1 | Acaa2 | 3-ketoacyl-CoA thiolase, mitochondrial Mus musculus | 346.08 | 26.2 | 41830 | 6 | 6 | 8748761 | 11940587 | 12142917 | 9079196 | 9597698 | 8148865 | 10659839 | 12137217 | 11424172 | 9677157 | 10088767 | 8205334 | 8531937 | 9753223 | 5980469 |
| P11497 | Acaca | Acetyl-CoA carboxylase 1 Rattus norvegicus | 978.75 | 11.81 | 265194 | 17 | 22 | 16053941 | 14432279 | 18560953 | 15344241 | 16481721 | 18212811 | 20150013 | 14670331 | 24413653 | 18222867 | 21948717 | 23793103 | 19070543 | 17036307 | 23465953 |
| E9Q4Z2 | Acacb | Acetyl-CoA carboxylase 2 Mus musculus | 244.8 | 2.98 | 275750 | 1 | 6 | 2287776 | 2000558 | 2973958.5 | 2163551.5 | 1704186.88 | 1399991 | 2461307.75 | 1273598.38 | 6284896 | 2050882.88 | 3975914 | 4910972 | 2273073.25 | 2914604 | 3099833.75 |
| Q9D7B6 | Acad8 | Isobutyryl-CoA dehydrogenase, mitochondrial Mus musculus | 37.13 | 2.42 | 45020 | 1 | 1 | 1885928.75 | 2224785.5 | 1598112.25 | 1778057.62 | 1712586.12 | 2008989.88 | 1310819.12 | 1403083.88 | 1867244.88 | 1415810.75 | 1581461.88 | 1249996.25 | 1519860.75 | 1428049.62 | 1588770.62 |
| Q8JZN5 | Acad9 | Acyl-CoA dehydrogenase family member 9, mitochondrial Mus musculus | 741.86 | 28 | 68722 | 13 | 13 | 21458219 | 19486827 | 26590593 | 19012471 | 21718827 | 24948457 | 21685833 | 24312511 | 24872325 | 19183571 | 24177625 | 25952993 | 19680149 | 21587561 | 21516037 |
| P51174 | Acadl | Long-chain specific acyl-CoA dehydrogenase, mitochondrial Mus musculus | 970.14 | 50 | 47908 | 4 | 16 | 129661433 | 123162641 | 115800057 | 121847297 | 123761337 | 133138849 | 114699169 | 142751121 | 132941369 | 112445961 | 121655681 | 112682273 | 108313745 | 131440033 | 96865929 |
| P45952 | Acadm | Medium-chain specific acyl-CoA dehydrogenase, mitochondrial Mus musculus | 444.97 | 28.5 | 46481 | 9 | 9 | 42677457 | 50715201 | 46500937 | 45997517 | 47525865 | 43511933 | 43043197 | 45209525 | 44488141 | 43509069 | 41080921 | 40712041 | 44671013 | 41320329 | 36902633 |
| P15651 | Acads | Short-chain specific acyl-CoA dehydrogenase, mitochondrial Rattus norvegicus | 215.77 | 14.81 | 44765 | 5 | 5 | 4089614.5 | 4404529 | 4467067.5 | 4722528 | 3925899 | 3916745 | 4723415 | 4240356 | 4337095 | 4568409 | 4469548.5 | 4202721 | 4358307 | 4178074 | 3979667.75 |
| Q9DBL1 | Acadsb | Short/branched chain specific acyl-CoA dehydrogenase, mitochondrial Mus musculus | 542.63 | 33.8 | 47874 | 9 | 9 | 9718412 | 9999263 | 12876759 | 11010008 | 9727089 | 14303861 | 11936437 | 14167736 | 13978011 | 11570947 | 13984924 | 12747951 | 12073137 | 13843721 | 10698769 |
| P50544 | Acadvl | Very long-chain specific acyl-CoA dehydrogenase, mitochondrial Mus musculus | 610.36 | 27.9 | 70875 | 11 | 11 | 7397203.5 | 6852240 | 8378622.5 | 7202088.5 | 7427555 | 6896191 | 10267117 | 7898418.5 | 6292090 | 7979495 | 5824819 | 7568307.5 | 7628842 | 6266849 | 7732170 |
| Q5FVC7 | Acap2 | Arf-GAP with coiled-coil, ANK repeat and PH domain-containing protein 2 Rattus norveg | 88.33 | 2.47 | 87230 | 1 | 1 | 468502.62 | 649814.81 | 542981.44 | 733051.56 | 620654.19 | 508251.69 | 543199.94 | 1 | 1 | 676217.75 | 551682.44 | 477537.28 | 498394.88 | 407278.84 | 483723.36 |
| Q8QZT1 | Acat1 | Acetyl-CoA acetyltransferase, mitochondrial Mus musculus | 601.7 | 31.84 | 44816 | 10 | 10 | 21018019 | 20848719 | 21482667 | 24846375 | 24763999 | 24000857 | 39648097 | 22847431 | 22298747 | 32744299 | 23434125 | 25897753 | 26932491 | 20369291 | 28022563 |
| Q8CAY6 | Acat2 | Acetyl-CoA acetyltransferase, cytosolic Mus musculus | 467.61 | 27.46 | 41298 | 6 | 6 | 6614755.5 | 6744047 | 6972167.5 | 7411894 | 7119811.5 | 6692571.5 | 7820735 | 5158527 | 5290384.5 | 9215710 | 8191100 | 7704506.5 | 6899962.5 | 5972653 | 13751179 |
| Q8BMP6 | Acbd3 | Golgi resident protein GCP60 Mus musculus | 311.62 | 13.71 | 60181 | 4 | 4 | 2446929.5 | 2741295 | 4110402.75 | 2876475 | 2201998 | 2703105 | 3769329 | 2859294.75 | 2768780 | 3259249 | 2951486 | 4813264.5 | 3042259.75 | 1182302.88 | 4270609 |
| Q9D061 | Acbd6 | Acyl-CoA-binding domain-containing protein 6 Mus musculus | 54.8 | 4.26 | 30887 | 1 | 1 | 143866.42 | 238036.62 | 219430.64 | 136224.08 | 251276.25 | 248685.92 | 104656.03 | 182691.95 | 370626.06 | 180972.94 | 357960.59 | 153433.11 | 236857 | 253700.52 | 217659.5 |
| Q9JIX8 | Acin1 | Apoptotic chromatin condensation inducer in the nucleus Mus musculus | 469.93 | 8.07 | 150719 | 9 | 9 | 16880435 | 17403197 | 18043895 | 15951028 | 18141687 | 18578343 | 16515778 | 17003553 | 18387233 | 15189846 | 15858068 | 17111829 | 16325561 | 17217355 | 14696417 |
| Q91V92 | Acly | ATP-citrate synthase Mus musculus | 2898.74 | 53.99 | 119728 | 4 | 45 | 82236001 | 82837753 | 88651937 | 93847417 | 80978689 | 83758937 | 102994433 | 83957201 | 84866241 | 97974041 | 87634009 | 100401777 | 86688641 | 93853201 | 106846793 |
| P28271 | Aco1 | Cytoplasmic aconitate hydratase Mus musculus | 561.96 | 18.56 | 98126 | 10 | 10 | 14127157 | 14481343 | 16563087 | 16485249 | 16188815 | 15248672 | 15099980 | 15633938 | 14721148 | 14350107 | 12844197 | 15502669 | 15361059 | 15118525 | 14519971 |
| Q99KI0 | Aco2 | Aconitate hydratase, mitochondrial Mus musculus | 1715.58 | 40.38 | 85464 | 27 | 27 | 113500217 | 122642265 | 125108009 | 117830081 | 128722697 | 129978497 | 160286513 | 133078353 | 141122497 | 142741457 | 136772321 | 125233145 | 136051793 | 119969961 | 138064673 |
| P54987 | Acod1 | Cis-aconitate decarboxylase Mus musculus | 98.01 | 6.97 | 53758 | 2 | 2 | 516474.31 | 509054.03 | 870485.12 | 464057.28 | 567807.75 | 539561.19 | 691150.38 | 263412.94 | 955683.5 | 366607.25 | 659899.75 | 637476.88 | 822221.38 | 1588056.5 | 628332 |
| Q9CQR4 | Acot13 | Acyl-coenzyme A thioesterase 13 Mus musculus | 140.56 | 28.57 | 15183 | 3 | 3 | 2040353.5 | 1102507.12 | 3148655.5 | 2605679.25 | 984908 | 1720611.25 | 3585120.75 | 2784189.25 | 1637095.25 | 4193929.75 | 1205451.38 | 1936570.25 | 2836042.5 | 1913452.5 | 1717838.88 |
| O55171 | Acot2 | Acyl-coenzyme A thioesterase 2, mitochondrial Rattus norvegicus | 142.45 | 8.83 | 49701 | 3 | 3 | 550374.44 | 402008.53 | 423205.84 | 959802.75 | 493726.22 | 584808.75 | 1672619.75 | 436651.03 | 397434.41 | 1604013 | 834393.69 | 1329915.5 | 553110.31 | 475073.53 | 1668220 |
| Q91V12 | Acot7 | Cytosolic acyl coenzyme A thioester hydrolase Mus musculus | 440.58 | 23.88 | 42537 | 7 | 7 | 8168401 | 15032150 | 13087661 | 8471117 | 12175181 | 12546691 | 15173361 | 12640312 | 14555695 | 13625147 | 11284257 | 9785729 | 14685349 | 14196003 | 12959913 |
| Q9R0X4 | Acot9 | Acyl-coenzyme A thioesterase 9, mitochondrial Mus musculus | 207.23 | 13.67 | 50560 | 4 | 4 | 3850710 | 1346032 | 2549555.25 | 2736440.5 | 2932182 | 1842190.25 | 3738350 | 3609475.25 | 1544328.12 | 3595217.5 | 2916075.75 | 2067414.88 | 3945121.25 | 3704991.5 | 3809771 |
| Q9EPL9 | Acox3 | Peroxisomal acyl-coenzyme A oxidase 3 Mus musculus | 163.39 | 6.43 | 78404 | 2 | 3 | 2464755.25 | 2313931.75 | 2644338.25 | 1952754.62 | 2369746 | 1 | 1 | 2423189 | 2648813.5 | 2328251 | 2051216.38 | 2252051 | 2476068 | 2311456.25 | 2124280.25 |
| P41498 | Acp1 | Low molecular weight phosphotyrosine protein phosphatase Rattus norvegicus | 359.06 | 51.27 | 18152 | 5 | 5 | 20943549 | 22746611 | 20750335 | 23506131 | 23260161 | 21532829 | 21696753 | 22297525 | 21275747 | 23893903 | 24090413 | 20106139 | 22758851 | 23467669 | 20114045 |
| P20611 | Acp2 | Lysosomal acid phosphatase Rattus norvegicus | 136.81 | 6.38 | 48320 | 3 | 3 | 2490458.25 | 4108593.75 | 4197776.5 | 4352387.5 | 4202293.5 | 3564811.5 | 2431140.5 | 4353933.5 | 4572737 | 2834420.5 | 3036018.5 | 3367261 | 3948489.5 | 4281292.5 | 3402012 |
| Q8BP40 | Acp6 | Lysophosphatidic acid phosphatase type 6 Mus musculus | 216.89 | 13.64 | 47625 | 4 | 4 | 5040321 | 4781136.5 | 5236165.5 | 5986873 | 6641041 | 5829277.5 | 6505740.5 | 5926268.5 | 5512082.5 | 5772338 | 6205760 | 6282915.5 | 5197287.5 | 5484571.5 | 6044044.5 |
| Q8VCW8 | Acsf2 | Acyl-CoA synthetase family member 2, mitochondrial Mus musculus | 435.77 | 26.02 | 67951 | 8 | 8 | 6112308.5 | 6232736.5 | 7270555 | 10534029 | 6490749 | 6928523.5 | 6839775.5 | 9849699 | 8928291 | 5886367.5 | 5898951.5 | 6440000 | 6333306.5 | 7520961 | 10672875 |
| P41216 | Acsl1 | Long-chain-fatty-acid--CoA ligase 1 Mus musculus | 1095.66 | 40.63 | 77951 | 19 | 21 | 35063401 | 37848509 | 35577673 | 29520365 | 34174989 | 39867261 | 40307941 | 43095697 | 49797837 | 36277381 | 30323187 | 32020637 | 36572393 | 40145453 | 30011147 |
| Q9CZW4 | Acsl3 | Long-chain-fatty-acid--CoA ligase 3 Mus musculus | 602.7 | 22.36 | 80492 | 11 | 12 | 103249249 | 90838929 | 98982273 | 96452401 | 114946073 | 101720217 | 152736065 | 92533633 | 75627889 | 120659993 | 119966537 | 96649937 | 107708305 | 87359361 | 112732089 |
| Q9QUJ7 | Acsl4 | Long-chain-fatty-acid--CoA ligase 4 Mus musculus | 1179.65 | 41.63 | 79077 | 22 | 23 | 77413993 | 74544825 | 73041393 | 67446297 | 74989745 | 67852337 | 61157321 | 69063353 | 66458345 | 63639425 | 72232849 | 54891777 | 66980113 | 68631729 | 61568669 |
| Q8JZR0 | Acsl5 | Long-chain-fatty-acid--CoA ligase 5 Mus musculus | 568.47 | 25.04 | 76206 | 10 | 12 | 16369395 | 17501567 | 17539055 | 17835385 | 16363560 | 16946357 | 20474747 | 19975087 | 18550289 | 18982891 | 13986256 | 16427646 | 18183383 | 17549727 | 16317119 |
| Q9QXG4 | Acss2 | Acetyl-coenzyme A synthetase, cytoplasmic Mus musculus | 491.28 | 18.54 | 78862 | 8 | 8 | 11705761 | 10209376 | 11267837 | 12468933 | 11857124 | 14228726 | 16451247 | 11153476 | 10006216 | 16238296 | 12365202 | 12291376 | 12232099 | 11321986 | 15666322 |
| P60710 | Actb | Actin, cytoplasmic 1 Mus musculus | 4409.75 | 98.4 | 41737 | 2 | 61 | 8796781569 | 10125157377 | 9494925313 | 9269221377 | 9283770369 | 8538429441 | 8704195585 | 8662747137 | 9774555137 | 9442095105 | 9489776641 | 8805225473 | 9640908801 | 9386089473 | 8729010177 |
| Q8BFZ3 | Actbl2 | Beta-actin-like protein 2 Mus musculus | 1372.44 | 40.43 | 42004 | 1 | 23 | 3744656897 | 4347484673 | 3986426369 | 3925038081 | 3874814209 | 3589975041 | 3577766913 | 3736043521 | 4200019457 | 3744244993 | 3892407553 | 3716596225 | 4067629057 | 4025514497 | 3465602305 |
| P68033 | Actc1 | Actin, alpha cardiac muscle 1 Mus musculus | 1950.93 | 55.97 | 42019 | 4 | 35 | 6090285569 | 7046930433 | 6816580609 | 6644038657 | 6506530305 | 5832457217 | 6317768193 | 5989777921 | 6821425665 | 6894988289 | 6757610497 | 6363269633 | 6742809601 | 6426138113 | 6280207873 |
| P63259 | Actg1 | Actin, cytoplasmic 2 Rattus norvegicus | 4409.58 | 98.4 | 41793 | 2 | 61 | 8772901889 | 10096184321 | 9473689601 | 9240883201 | 9255208961 | 8518846977 | 8669237249 | 8644033537 | 9762530305 | 9421634561 | 9477673985 | 8789067777 | 9618493441 | 9367257089 | 8710659073 |
| Q9Z2N8 | Actl6a | Actin-like protein 6A Mus musculus | 638.6 | 42.19 | 47448 | 13 | 13 | 20609735 | 27996013 | 27881319 | 25040099 | 22246899 | 19478133 | 23934095 | 23910685 | 22975643 | 20409953 | 25592787 | 24188031 | 23036305 | 25460823 | 19105673 |
| Q7TPR4 | Actn1 | Alpha-actinin-1 Mus musculus | 2014.61 | 43.5 | 103068 | 1 | 34 | 93833825 | 94325033 | 103145209 | 95372721 | 84718417 | 91286257 | 96216657 | 102992185 | 103986561 | 86399809 | 96246665 | 103139593 | 92084545 | 103297537 | 97942057 |
| P57780 | Actn4 | Alpha-actinin-4 Mus musculus | 3053.32 | 63.93 | 104977 | 2 | 49 | 209436129 | 203842033 | 213859377 | 201181649 | 193270273 | 189955841 | 197074225 | 210276481 | 209417761 | 185271713 | 208642369 | 209698945 | 195022657 | 210785489 | 203274657 |
| Q9QZB7 | Actr10 | Actin-related protein 10 Mus musculus | 201.02 | 7.43 | 46208 | 2 | 2 | 1627365.25 | 1443089.5 | 1543676.75 | 1814677.25 | 1030554.06 | 1417333.5 | 1560400 | 1355979.12 | 1331342.75 | 1797201.12 | 1957993.38 | 1414599.12 | 1651427.38 | 1783187.5 | 1430318.5 |
| P61164 | Actr1a | Alpha-centractin Mus musculus | 481.85 | 35.64 | 42614 | 4 | 9 | 6828401.5 | 47553585 | 13629455 | 15522523 | 15123009 | 14040156 | 35470973 | 14983961 | 14921828 | 16274519 | 13731322 | 8242871 | 16614583 | 15110283 | 14046430 |
| Q8R5C5 | Actr1b | Beta-centractin Mus musculus | 411.77 | 23.67 | 42281 | 2 | 7 | 4891109.5 | 14206069 | 12028197 | 12383290 | 12416062 | 12844012 | 10664717 | 12019553 | 12127267 | 12829367 | 11228721 | 4777572 | 13132835 | 13262152 | 11246126 |
| P61161 | Actr2 | Actin-related protein 2 Mus musculus | 1276.89 | 57.36 | 44761 | 19 | 19 | 164446689 | 194105441 | 166601889 | 156775073 | 177940385 | 162424417 | 165946161 | 104754121 | 181018145 | 164536689 | 178418945 | 160258273 | 175460417 | 174143985 | 155152097 |
| Q4V7C7 | Actr3 | Actin-related protein 3 Rattus norvegicus | 1283.95 | 61.24 | 47357 | 20 | 20 | 192447953 | 232151825 | 219706193 | 217601153 | 234542769 | 219927665 | 217704257 | 220299921 | 211251537 | 215091489 | 264997937 | 221478609 | 231985041 | 232978161 | 231689601 |
| Q99JW2 | Acy1 | Aminoacylase-1 Mus musculus | 424.76 | 28.92 | 45781 | 7 | 7 | 7514963 | 4298089.5 | 6403944.5 | 6028198 | 5054992.5 | 6240584.5 | 5724649 | 6638113 | 5634249.5 | 5030270 | 6519693.5 | 6237289 | 5808592 | 6509311.5 | 3780838.5 |
| P03958 | Ada | Adenosine deaminase Mus musculus | 161.96 | 9.66 | 39992 | 3 | 3 | 4214858.5 | 4943317 | 4736053 | 3611837.25 | 4193373.5 | 3783621.5 | 3994147 | 4308828.5 | 4500475 | 3115153 | 3572425.5 | 3895219.5 | 4332480.5 | 5418278.5 | 3979522 |
| O88839 | Adam15 | Disintegrin and metalloproteinase domain-containing protein 15 Mus musculus | 102.67 | 2.08 | 92664 | 1 | 1 | 202693.86 | 264608.88 | 222039.39 | 219734.345 | 189595.25 | 214852.19 | 221888.36 | 266290.06 | 298931.56 | 250490.5425 | 384906.22 | 358725.75 | 189949.55 | 306247.22 | 314954.81 |
| Q9Z0F8 | Adam17 | Disintegrin and metalloproteinase domain-containing protein 17 Mus musculus | 243.27 | 4.47 | 93056 | 3 | 3 | 352922.03 | 434138.34 | 751878.5 | 450703.3825 | 263874.66 | 412453.28 | 836522.12 | 1102172.62 | 2053463 | 736048.5 | 995278.94 | 750296.12 | 132543.08 | 766799.56 | 1399384.75 |
| Q05910 | Adam8 | Disintegrin and metalloproteinase domain-containing protein 8 Mus musculus | 110.43 | 5.69 | 90046 | 3 | 3 | 1 | 1 | 1 | 1 | 1 | 1 | 1 | 226095.56 | 1 | 400122.12 | 1 | 1 | 316496.16 | 1 | 319461.28 |
| Q61072 | Adam9 | Disintegrin and metalloproteinase domain-containing protein 9 Mus musculus | 45.96 | 2.84 | 92079 | 1 | 1 | 559447.06 | 848215.62 | 617705.295 | 655406.69 | 407751.81 | 624698.94 | 806818.0475 | 650661.62 | 788164.88 | 1163746.75 | 469147.81 | 587000.06 | 637441.56 | 598369.31 | 649480.81 |
| Q8C9W3 | Adamts2 | A disintegrin and metalloproteinase with thrombospondin motifs 2 Mus musculus | 36.67 | 0.58 | 135299 | 1 | 1 | 4741734.5 | 6284738.5 | 7280784.5 | 6663445.5 | 6696178.5 | 7204461.5 | 8909308 | 5661507.5 | 5839923 | 8930979 | 7421133.5 | 6450391.5 | 6797673 | 5114457.5 | 7173918.5 |
| Q99MU3 | Adar | Double-stranded RNA-specific adenosine deaminase Mus musculus | 220.57 | 4.58 | 130447 | 4 | 4 | 236765 | 130874.45 | 1194792 | 874160.94 | 135664.2 | 242990.83 | 165915.91 | 760085.75 | 180372 | 720316.62 | 945249.75 | 82726.58 | 151415.16 | 735088.81 | 1029010.06 |
| P51829 | Adcy7 | Adenylate cyclase type 7 Mus musculus | 104.34 | 2.64 | 122708 | 2 | 2 | 879758.88 | 466861.75 | 1108726.38 | 486994.72 | 359808.03 | 469061.22 | 572168.44 | 1234908 | 1124735.12 | 423998.34 | 937802.5 | 871533.25 | 378466.59 | 1169370.25 | 678754.25 |
| Q63028 | Add1 | Alpha-adducin Rattus norvegicus | 35.3 | 3.67 | 80355 | 1 | 1 | 231378.59 | 1 | 188683.8 | 203929.55 | 1 | 1 | 117504.94 | 162324.22 | 1 | 214069.3 | 1 | 137980.59 | 194660.31 | 1 | 178518.92 |
| P28474 | Adh5 | Alcohol dehydrogenase class-3 Mus musculus | 837.02 | 56.15 | 39548 | 14 | 14 | 36729549 | 41145705 | 37752825 | 35846885 | 40345165 | 39064173 | 42104453 | 38225821 | 40197761 | 39989993 | 39847093 | 37549569 | 40054041 | 41322113 | 42303913 |
| P55264 | Adk | Adenosine kinase Mus musculus | 231.14 | 21.05 | 40149 | 2 | 4 | 1433594.25 | 1214920.5 | 2043164.25 | 1937322.12 | 1291554.38 | 1720331.75 | 3062530 | 1355738.25 | 1533008.62 | 3060196.5 | 2328143 | 1815017.62 | 1646242.75 | 1336069 | 3001759.75 |
| Q9JKL8 | Adnp | Activity-dependent neuroprotector homeobox protein Rattus norvegicus | 120.24 | 2.81 | 123626 | 3 | 3 | 1 | 1 | 1 | 996820.62 | 1 | 456210.28 | 425295.16 | 385961.38 | 1 | 1 | 1 | 1 | 1 | 1 | 1 |
| Q8VDL4 | Adpgk | ADP-dependent glucokinase Mus musculus | 209.99 | 11.29 | 53902 | 4 | 4 | 8361254.5 | 6984721.5 | 7357322.5 | 4871773 | 6078658 | 7104860 | 7545757 | 7750469.5 | 8325269 | 6881211 | 4021143.5 | 7909937 | 7714840 | 7243298.5 | 6796184 |
| P54923 | Adprh | [Protein ADP-ribosylarginine] hydrolase Mus musculus | 158.22 | 9.67 | 40068 | 3 | 3 | 4881424 | 3971885 | 4591478 | 4401517 | 4606919.5 | 3425866.25 | 3512414.75 | 4110159 | 4412743.5 | 3442471 | 4177360 | 3925895.75 | 3657698.25 | 4644210 | 3380407.25 |
| Q8CG72 | Adprhl2 | Poly(ADP-ribose) glycohydrolase ARH3 Mus musculus | 177.7 | 15.68 | 39414 | 4 | 4 | 3002238 | 4763269.5 | 4780709.5 | 4387344 | 4546301 | 3958845.5 | 5355005.5 | 5486425 | 4560545.5 | 4067111.25 | 5148211.5 | 4637607 | 4086839.25 | 4853875 | 5188380 |
| Q9JKV1 | Adrm1 | Proteasomal ubiquitin receptor ADRM1 Mus musculus | 226.99 | 16.46 | 42060 | 5 | 5 | 9169358 | 9070299 | 9524340 | 14480429 | 14783161 | 12781753 | 11772622 | 12278751 | 8608452 | 14529969 | 11013009 | 9476860 | 14290904 | 8509205 | 12016427 |
| P54822 | Adsl | Adenylosuccinate lyase Mus musculus | 813.72 | 48.76 | 54866 | 15 | 15 | 17095313 | 18313047 | 15103116 | 16259600 | 13355404 | 15921756 | 18899313 | 15659123 | 15999327 | 16014960 | 15703463 | 15662472 | 14920513 | 15426081 | 18813455 |
| P46664 | Adss | Adenylosuccinate synthetase isozyme 2 Mus musculus | 990.01 | 42.32 | 50021 | 16 | 17 | 42509193 | 49394993 | 54515937 | 40716057 | 39294245 | 47413097 | 39170577 | 49827565 | 50223845 | 44565313 | 40216657 | 46508585 | 45171361 | 45462009 | 48020797 |
| P28650 | Adssl1 | Adenylosuccinate synthetase isozyme 1 Mus musculus | 452.2 | 25.16 | 50254 | 8 | 8 | 24815345 | 19505815 | 7947449 | 16252192 | 5332461.5 | 8407743 | 7686831.5 | 10205971 | 9453597 | 6886983 | 8328268 | 7590516.5 | 8037311.5 | 26826711 | 5837711.5 |
| Q9QZQ1 | Afdn | Afadin Mus musculus | 78.26 | 0.99 | 206499 | 1 | 1 | 997290.12 | 1019074 | 1066013.75 | 898114.5 | 978138.25 | 1157515.12 | 1050873.25 | 1269727.38 | 1333742.5 | 968917.56 | 1104009.25 | 1086212.12 | 1301657.12 | 1336512.25 | 1016038.62 |
| Q920A7 | Afg3l1 | AFG3-like protein 1 Mus musculus | 223.41 | 8.11 | 87047 | 3 | 4 | 3144155 | 5008066.5 | 5348821.5 | 3231860.5 | 3044049.25 | 2142360.25 | 3425584 | 5233020.5 | 6611815 | 3958408.5 | 2861614.5 | 4175232.25 | 3117157.75 | 4300092 | 4393075 |
| Q8JZQ2 | Afg3l2 | AFG3-like protein 2 Mus musculus | 377.27 | 12.72 | 89519 | 8 | 9 | 4367963.5 | 4278519.5 | 4811836.5 | 5242098 | 4480939 | 5735902 | 5469554 | 6236984.5 | 5945194 | 5792793.5 | 4642165.5 | 4680066 | 5307019 | 4978489.5 | 4488273.5 |
| Q4KLH5 | Agfg1 | Arf-GAP domain and FG repeat-containing protein 1 Rattus norvegicus | 160.28 | 9.8 | 58170 | 3 | 3 | 862080.38 | 1442490.75 | 2389525.25 | 1366870.5 | 1249627.88 | 1930205.38 | 1574221.38 | 943838.12 | 757964.5 | 1246228.75 | 1136658 | 2051758.12 | 1364998 | 1730243.25 | 2031819 |
| Q80WC7 | Agfg2 | Arf-GAP domain and FG repeat-containing protein 2 Mus musculus | 105.3 | 5.64 | 48968 | 2 | 2 | 735089.94 | 1670095.5 | 1333213.25 | 1370309 | 1242813.25 | 1102802.5 | 1059594.12 | 1065337 | 1239383 | 1369986.38 | 835578.25 | 1268280.25 | 920155.88 | 258203.84 | 1347088.25 |
| Q9ESW4 | Agk | Acylglycerol kinase, mitochondrial Mus musculus | 114.69 | 6.65 | 46976 | 2 | 2 | 2610002 | 2282347 | 1911229.25 | 2118221.25 | 2412607.75 | 1778112.62 | 2005671.88 | 2282199.25 | 2162585 | 2907515.75 | 1826493.62 | 2031569.38 | 2451508.75 | 1987210.5 | 1837343.25 |
| Q8CJG0 | Ago2 | Protein argonaute-2 Mus musculus | 123.91 | 4.65 | 97304 | 3 | 3 | 376920.59 | 384765.5 | 519341.53 | 353858.5 | 449541.78 | 315670.16 | 534121.69 | 428432.78 | 730031.25 | 362310 | 1 | 493640.97 | 1 | 451319.06 | 411125.69 |
| Q8C0I1 | Agps | Alkyldihydroxyacetonephosphate synthase, peroxisomal Mus musculus | 624.79 | 23.26 | 71684 | 11 | 11 | 6235871 | 5000361 | 8535396 | 3113150 | 5289479.5 | 6221845 | 10641165 | 10973204 | 10437807 | 7975788 | 10978843 | 8863578 | 8071466 | 7456885 | 10407553 |
| Q8CJF7 | Ahctf1 | Protein ELYS Mus musculus | 111.96 | 1.52 | 247646 | 2 | 2 | 91999.9 | 180918.52 | 92771.87 | 1 | 1 | 1 | 1 | 1 | 73033.41 | 164434.98 | 1 | 1 | 1 | 1 | 1 |
| P50247 | Ahcy | Adenosylhomocysteinase Mus musculus | 1645.95 | 73.15 | 47688 | 26 | 27 | 261898465 | 220012497 | 216327809 | 185914257 | 190846721 | 191928417 | 216524401 | 198470993 | 233491201 | 206932417 | 242060545 | 221854369 | 211867169 | 220601617 | 218260897 |
| B5DFN2 | Ahcyl1 | S-adenosylhomocysteine hydrolase-like protein 1 Rattus norvegicus | 241.82 | 16.15 | 53753 | 3 | 6 | 5914869 | 8225328.5 | 8047324 | 7341394.5 | 8698729 | 9280869 | 10176526 | 5908079.5 | 8460177 | 9469762 | 6868791 | 7764813 | 8365127 | 7955541.5 | 6787916 |
| Q68FL4 | Ahcyl2 | Putative adenosylhomocysteinase 3 Mus musculus | 147.13 | 7.18 | 66899 | 1 | 4 | 5611331 | 5789497 | 6573405.5 | 5489033.5 | 5941449 | 6327441.5 | 8434446 | 5606238.5 | 5528567.5 | 6569804.5 | 6713568.5 | 5653532.5 | 6203136 | 4873598.5 | 6252244 |

| | | | | | | | | | | | | | | | | | | | | | | |
|---|---|---|---|---|---|---|---|---|---|---|---|---|---|---|---|---|---|---|---|---|---|---|
| Q8BK64 | Ahsa1 | Activator of 90 kDa heat shock protein ATPase homolog 1 Mus musculus | 710.49 | 57.4 | 38117 | 13 | 13 | 34326565 | 33961857 | 38453801 | 32439079 | 36210013 | 38930393 | 28871591 | 39068505 | 46166341 | 30772361 | 36783025 | 40812001 | 32756115 | 38396669 | 40054173 |
| O70200 | Aif1 | Allograft inflammatory factor 1 Mus musculus | 191.28 | 21.77 | 16911 | 2 | 2 | 6277422 | 5196575.5 | 3569797.5 | 3538031 | 3821591.25 | 2973521.5 | 2710058.75 | 3442092 | 4225986 | 3028569.75 | 2801122.25 | 3749154.5 | 4444987 | 3982692.5 | 2650226.5 |
| Q9Z0X1 | Aifm1 | Apoptosis-inducing factor 1, mitochondrial Mus musculus | 612.78 | 34.8 | 66765 | 13 | 13 | 16463537 | 16582204 | 19528453 | 20219293 | 23095575 | 20362615 | 18354087 | 20801145 | 19616457 | 19501151 | 16899433 | 19364091 | 17473465 | 19012971 | 18656051 |
| Q91VJ1 | Aim2 | Interferon-inducible protein AIM2 Mus musculus | 222.28 | 12.71 | 40155 | 4 | 4 | 3277398 | 2867726 | 3099296.5 | 3292299 | 2809503 | 3193824 | 3511077 | 3091330.75 | 1647589.88 | 2930478.5 | 3523602.5 | 3408283 | 1295894.75 | 3538926 | 2854124.5 |
| P31230 | Aimp1 | Aminoacyl tRNA synthase complex-interacting multifunctional protein 1 Mus musculus | 463.32 | 44.19 | 33997 | 8 | 8 | 7778655 | 8279096 | 8006657 | 9662687 | 7513502 | 6137935 | 10519636 | 5574385 | 7047870 | 8804933 | 7920547 | 8311564 | 8022749 | 7039005.5 | 9534025 |
| Q8R010 | Aimp2 | Aminoacyl tRNA synthase complex-interacting multifunctional protein 2 Mus musculus | 411.23 | 35.62 | 35378 | 5 | 5 | 2585745.75 | 3658902 | 2180424 | 2601406.5 | 3133163.75 | 3202677.5 | 5442961.5 | 2948986.5 | 2518666 | 3526309.5 | 3061857 | 2713090.5 | 3430759.25 | 3806553 | 3282496.5 |
| O08915 | Aip | AH receptor-interacting protein Mus musculus | 106.21 | 8.18 | 37605 | 2 | 2 | 3069253 | 4603496 | 4547992.5 | 2998881.25 | 3235235.25 | 4524630 | 3254764.25 | 3403683.75 | 4395634.5 | 3798911.5 | 3908950.25 | 4039634.5 | 4416085.5 | 4718111 | 3790716.5 |
| Q9WTP6 | Ak2 | Adenylate kinase 2, mitochondrial Mus musculus | 315.83 | 33.89 | 26469 | 6 | 6 | 20322113 | 21748331 | 21522219 | 22517561 | 21268597 | 20600765 | 23878439 | 23651653 | 22392623 | 23838995 | 20312133 | 19755853 | 26145533 | 24722225 | 17692919 |
| Q9WTP7 | Ak3 | GTP:AMP phosphotransferase AK3, mitochondrial Mus musculus | 211.67 | 25.11 | 25426 | 4 | 4 | 2491269.5 | 2715702 | 2521727.25 | 2693779.75 | 2454971.5 | 1306415.75 | 2221138 | 1804367.88 | 2516100.75 | 3011834.5 | 1129692.5 | 2135913.5 | 2132867.5 | 2220516.75 | 2380462 |
| O08715 | Akap1 | A-kinase anchor protein 1, mitochondrial Mus musculus | 43.88 | 1.52 | 92195 | 1 | 1 | 1 | 1 | 1 | 1 | 1 | 1 | 390539.28 | 1 | 484910.12 | 1 | 242898.72 | 1 | 1 | 1 | 1 |
| Q9DBR0 | Akap8 | A-kinase anchor protein 8 Mus musculus | 169.5 | 8.73 | 76294 | 3 | 3 | 1934461.5 | 3442657.5 | 3314414.5 | 4143002.5 | 4506964.5 | 1998681.62 | 3327024 | 3343058.5 | 3117308.25 | 2950946.75 | 3962735.75 | 3560206 | 3473614.75 | 3452441 | 2058032.75 |
| Q9JII6 | Akr1a1 | Alcohol dehydrogenase [NADP(+)] Mus musculus | 451.41 | 30.46 | 36587 | 9 | 9 | 17977653 | 19295517 | 19866239 | 15493695 | 17257193 | 17913013 | 19327225 | 17201579 | 18792581 | 16551147 | 16597063 | 18634157 | 19685827 | 17941361 | 15445709 |
| P45376 | Akr1b1 | Aldose reductase Mus musculus | 841.51 | 52.53 | 35732 | 15 | 15 | 190222113 | 212004353 | 199337377 | 196657073 | 209078065 | 200793185 | 192289489 | 201373953 | 205073585 | 195443137 | 216811233 | 184297313 | 216898993 | 209309857 | 168169489 |
| Q5RJP0 | Akr1b7 | Aldose reductase-related protein 1 Rattus norvegicus | 348.17 | 17.72 | 36122 | 1 | 6 | 36651325 | 37410433 | 31974077 | 29491129 | 35974133 | 34467585 | 30213069 | 37010093 | 39002105 | 25093509 | 33522491 | 31823087 | 35470809 | 39882353 | 26547281 |
| P21300 | Akr1b7 | Aldose reductase-related protein 1 Mus musculus | 262.98 | 15.82 | 35988 | 0 | 6 | 44295361 | 48995329 | 44048337 | 41327449 | 46377081 | 44065817 | 42811721 | 47901913 | 49089581 | 38550197 | 43476497 | 44225113 | 49395905 | 49637921 | 36971521 |
| P45377 | Akr1b8 | Aldose reductase-related protein 2 Mus musculus | 810.43 | 49.05 | 36121 | 6 | 13 | 58479109 | 60477413 | 54098949 | 51269453 | 57613033 | 57808637 | 53163465 | 61380401 | 61369273 | 46567685 | 57047833 | 55933925 | 61570769 | 63891225 | 47047365 |
| Q8CG45 | Akr7a2 | Aflatoxin B1 aldehyde reductase member 2 Rattus norvegicus | 61.95 | 4.09 | 40675 | 1 | 1 | 1 | 458293.44 | 694835.62 | 1 | 777704.19 | 702958.56 | 999241.5 | 774745.94 | 384596.72 | 859606.69 | 759441.56 | 911180.3275 | 857205.94 | 840697.56 | 1187376.25 |
| P31750 | Akt1 | RAC-alpha serine/threonine-protein kinase Mus musculus | 170.55 | 10.62 | 55707 | 2 | 3 | 1051679.38 | 943814.19 | 878722.9375 | 769563.62 | 749834.56 | 791098.44 | 839293.44 | 1528083.25 | 829340.75 | 805991.81 | 441918.84 | 1268817.75 | 977683.945 | 1407638.5 | 792360.69 |
| P47197 | Akt2 | RAC-beta serine/threonine-protein kinase Rattus norvegicus | 71.85 | 6.03 | 55543 | 0 | 2 | 645641.81 | 1 | 1 | 1 | 749834.56 | 1 | 1 | 724734.56 | 829340.75 | 1 | 441918.84 | 449421.84 | 1 | 422355.94 | 1 |
| P10518 | Alad | Delta-aminolevulinic acid dehydratase Mus musculus | 418.56 | 31.21 | 36024 | 6 | 7 | 7514570.5 | 8549166 | 8025639.5 | 7233266 | 7346623.5 | 7345902 | 7440667.5 | 8370005 | 7691856.5 | 7503848.5 | 7718128 | 6911619 | 7125579 | 8117753 | 7120672.5 |
| Q61490 | Alcam | CD166 antigen Mus musculus | 348.19 | 23.67 | 65092 | 8 | 8 | 3922109.75 | 3693753 | 4025636.5 | 1983142.12 | 5778949.5 | 5149710.5 | 5747331 | 2995682 | 4731855 | 4661006 | 3904770.5 | 3596569 | 4430866 | 4176208 | 2647710.25 |
| Q571I9 | Aldh16a1 | Aldehyde dehydrogenase family 16 member A1 Mus musculus | 328.47 | 11.47 | 84756 | 6 | 6 | 3665910 | 4166355.25 | 4120707 | 4882765 | 4007680 | 1966380.38 | 4657921.5 | 4491111.5 | 3024431.5 | 3570185.5 | 3850807.25 | 4867322.5 | 4895878 | 3568496.5 | 4376979.5 |
| Q9Z110 | Aldh18a1 | Delta-1-pyrroline-5-carboxylate synthase Mus musculus | 944.96 | 25.91 | 87266 | 16 | 17 | 27464813 | 31389645 | 29768825 | 29898957 | 27695143 | 34323477 | 30648901 | 27817197 | 36006093 | 31489571 | 31169295 | 29336437 | 23895557 | 24029511 | 25936149 |
| Q66HF8 | Aldh1b1 | Aldehyde dehydrogenase X, mitochondrial Rattus norvegicus | 111.29 | 4.82 | 57625 | 1 | 2 | 11603215 | 11684018 | 13051814 | 12502605 | 11462587 | 13491576 | 12992340 | 14037616 | 14561375 | 12347817 | 13110488 | 12577315 | 13281077 | 13556662 | 19708481 |
| Q8R0Y6 | Aldh1l1 | Cytosolic 10-formyltetrahydrofolate dehydrogenase Mus musculus | 426.66 | 15.08 | 98709 | 9 | 9 | 6059573.5 | 7205715.5 | 8514740 | 7488813 | 7148226 | 5474034 | 6413010.5 | 6343990 | 6629918.5 | 5236689.5 | 5049724 | 7208806 | 7871800.5 | 8178418 | 5388803 |
| P47738 | Aldh2 | Aldehyde dehydrogenase, mitochondrial Mus musculus | 1506.28 | 68.21 | 56538 | 24 | 25 | 182985393 | 204331585 | 209732513 | 200265169 | 210116641 | 191271025 | 209545857 | 225216913 | 228857569 | 194503729 | 212343073 | 207853137 | 227798145 | 214104417 | 213676385 |
| P47740 | Aldh3a2 | Fatty aldehyde dehydrogenase Mus musculus | 186.2 | 5.58 | 53971 | 2 | 2 | 1676653.38 | 1683196.62 | 1 | 486477.28 | 1 | 1 | 1 | 1 | 1 | 1196692.38 | 1078285.25 | 943794.4825 | 1033599.56 | 495197.12 | 1168096 |
| Q80VQ0 | Aldh3b1 | Aldehyde dehydrogenase family 3 member B1 Mus musculus | 105.23 | 8.55 | 52292 | 2 | 2 | 2891052.25 | 3489360.75 | 2962967.5 | 2642890.75 | 2990492.75 | 2874865.25 | 2193427.5 | 2756263.75 | 3004328 | 2437974.5 | 1855160.38 | 2087835.12 | 2403081.75 | 2925832.25 | 1889801.88 |
| Q9DBF1 | Aldh7a1 | Alpha-aminoadipic semialdehyde dehydrogenase Mus musculus | 42.14 | 2.6 | 58861 | 1 | 1 | 959145.5 | 1088013.62 | 726082.81 | 815029.56 | 992774.5 | 1080019.75 | 1072095.38 | 1169565.75 | 1134095 | 772771.38 | 888518.75 | 918887.75 | 1013396.19 | 1132162.75 | 1025874.94 |
| Q9JLJ2 | Aldh9a1 | 4-trimethylaminobutyraldehyde dehydrogenase Mus musculus | 1173.24 | 58.7 | 53515 | 20 | 21 | 53134621 | 59317857 | 56155481 | 64412857 | 62693641 | 56835877 | 63230073 | 54493965 | 57096013 | 60933033 | 57799745 | 51263457 | 54900193 | 56249469 | 57544345 |
| P05064 | Aldoa | Fructose-bisphosphate aldolase A Mus musculus | 2292.67 | 93.68 | 39356 | 27 | 36 | 1180237697 | 1044185217 | 1001171329 | 1153109889 | 1062884545 | 1200638337 | 1431839617 | 1166415361 | 1002550849 | 1276592769 | 1232825089 | 1143350401 | 1289505025 | 1190303873 | 995645825 |
| P05063 | Aldoc | Fructose-bisphosphate aldolase C Mus musculus | 1570.54 | 70.25 | 39395 | 16 | 23 | 264098593 | 255447873 | 239960209 | 228017905 | 262386721 | 321508961 | 260902945 | 295063201 | 302497985 | 227667473 | 270641345 | 251263441 | 238658609 | 326160929 | 211886577 |
| Q9DB25 | Alg5 | Dolichyl-phosphate beta-glucosyltransferase Mus musculus | 187.22 | 16.67 | 36791 | 4 | 4 | 7048862 | 4045596.5 | 8730081 | 7732013.5 | 4468782 | 7153639.5 | 8097790.5 | 8327683 | 8924620 | 4403396.5 | 3919209.25 | 7609129.5 | 4415455.5 | 6755473 | 8090145 |
| D3ZKD3 | Alkbh5 | RNA demethylase ALKBH5 Rattus norvegicus | 44.83 | 2.78 | 44453 | 1 | 1 | 770056.12 | 951099.19 | 902697.38 | 930684 | 958012.81 | 1288423.12 | 1763342.38 | 825632.19 | 801671.44 | 1171655.25 | 844340.94 | 1109016.62 | 982884.62 | 711236.88 | 1283230.62 |
| P48999 | Alox5 | Arachidonate 5-lipoxygenase Mus musculus | 772.07 | 31.01 | 77967 | 15 | 15 | 21994769 | 25299785 | 21351103 | 17280719 | 21516281 | 22085077 | 19985159 | 22293495 | 24923613 | 20071093 | 22566157 | 21830029 | 21766967 | 20838157 | 18276571 |
| P20291 | Alox5ap | Arachidonate 5-lipoxygenase-activating protein Rattus norvegicus | 38.16 | 4.97 | 18070 | 1 | 1 | 1 | 273593.47 | 1 | 1529866.12 | 199316.78 | 218265.8 | 5853626.5 | 1158138.88 | 187365.38 | 1571717.62 | 3324238 | 2523272.75 | 1541756.88 | 3160707.908 | 5253564 |
| O08583 | Alyref | THO complex subunit 4 Mus musculus | 452.53 | 40 | 26940 | 8 | 8 | 10586816 | 10303344 | 15216617 | 17258003 | 13529074 | 11968403 | 16365885 | 14091609 | 7535094 | 19209251 | 11514116 | 11997841 | 13896829 | 6196733 | 12067969 |
| Q8JZV7 | Amdhd2 | N-acetylglucosamine-6-phosphate deacetylase Mus musculus | 196.82 | 22.49 | 43501 | 5 | 5 | 6918675 | 8580279 | 6596870 | 8868889 | 8538026 | 7575070 | 4284778.5 | 6125598.5 | 6978516.5 | 10334745 | 7707855 | 6449142.5 | 8725626 | 6317035.5 | 7711004 |
| Q02356 | Ampd2 | AMP deaminase 2 Rattus norvegicus | 282.16 | 7.89 | 94787 | 6 | 6 | 4205408 | 3954117 | 3249520.5 | 5278274 | 4210056.5 | 4701920 | 2821730 | 2712324 | 3433593 | 5395316.5 | 5121619 | 3509451.5 | 4984255.5 | 2706736 | 4824482.5 |
| O08739 | Ampd3 | AMP deaminase 3 Mus musculus | 117.49 | 2.35 | 88652 | 2 | 2 | 3647009.5 | 1566312.5 | 1664332.25 | 1268097.25 | 1485964.62 | 1632760.5 | 2063198 | 2034629.38 | 2172845.25 | 1831941.5 | 1246675 | 1326819.5 | 1784477.75 | 1735553.5 | 2377186.75 |
| P53995 | Anapc1 | Anaphase-promoting complex subunit 1 Mus musculus | 85.35 | 1.7 | 215994 | 2 | 2 | 374076.56 | 272313.72 | 350240.84 | 205304.02 | 191318.83 | 231376.95 | 207422.94 | 346569.09 | 625956.31 | 265107.88 | 519731.34 | 403485.16 | 273123.59 | 356413.81 | 320576.88 |
| Q9WVM3 | Anapc7 | Anaphase-promoting complex subunit 7 Mus musculus | 46.32 | 2.65 | 63021 | 1 | 1 | 170881.94 | 225512.88 | 205361.45 | 196778.5375 | 185357.88 | 177195.38 | 165708.33 | 133887.28 | 147691.28 | 221631.55 | 148017.2 | 176537.95 | 238186.38 | 203239.66 | 156076.12 |
| Q810B6 | Ankfy1 | Rabankyrin-5 Mus musculus | 680.21 | 15.83 | 128653 | 11 | 11 | 7869792.5 | 13013437 | 8444439 | 12555536 | 9923451 | 10683901 | 7949605 | 14507235 | 13807257 | 8349459.5 | 16766907 | 9619489 | 8703377 | 9684171 | 13793938 |
| P58366 | Ankh | Progressive ankylosis protein homolog Rattus norvegicus | 85.84 | 4.27 | 54265 | 1 | 1 | 288447.91 | 143634.0975 | 98545.91 | 99877.84 | 87664.73 | 86453.02 | 1 | 76751.35 | 1 | 91991.62 | 78889.84 | 98150.19 | 109358.18 | 97868.2175 | 105074.66 |
| Q8BTI7 | Ankrd52 | Serine/threonine-protein phosphatase 6 regulatory ankyrin repeat subunit C Mus muscu | 72.83 | 1.21 | 115032 | 1 | 1 | 191753.88 | 388408.34 | 336735.25 | 275266.265 | 184167.59 | 260675.745 | 260288.14 | 291337.62 | 273367.19 | 217710.03 | 226587.22 | 176252.31 | 392126.38 | 288056.78 | 270755.6725 |
| Q8K298 | Anln | Anillin Mus musculus | 544.79 | 13.47 | 122794 | 10 | 10 | 5513382 | 4529574 | 5273436 | 6093720.5 | 4500061 | 4862777.5 | 5106853 | 4128478.5 | 6221053 | 4755466.5 | 5946740 | 4942632.5 | 3857615.5 | 5430092.5 | 5471853.5 |
| O35381 | Anp32a | Acidic leucine-rich nuclear phosphoprotein 32 family member A Mus musculus | 648.42 | 44.53 | 28538 | 9 | 11 | 65718145 | 61853977 | 59668621 | 48358549 | 57908813 | 54042785 | 70864657 | 58724569 | 47716621 | 67994921 | 64294397 | 65079153 | 65320485 | 62821549 | 78880537 |
| Q9EST5 | Anp32b | Acidic leucine-rich nuclear phosphoprotein 32 family member B Mus musculus | 354.38 | 27.21 | 31079 | 5 | 7 | 45747313 | 26284427 | 27068685 | 22621367 | 27982597 | 24173559 | 41308117 | 24952259 | 26429817 | 32020995 | 31394337 | 31902495 | 31099619 | 29436741 | 40127729 |
| P97822 | Anp32e | Acidic leucine-rich nuclear phosphoprotein 32 family member E Mus musculus | 639.73 | 30.38 | 29622 | 7 | 8 | 30915129 | 28389643 | 29572829 | 22445685 | 25878157 | 23046089 | 34795049 | 28374121 | 32637481 | 27289281 | 31980237 | 29797707 | 29763785 | 28071201 | 38081905 |
| P97449 | Anpep | Aminopeptidase N Mus musculus | 868.1 | 21.74 | 109651 | 10 | 14 | 13964333 | 13958208 | 17584101 | 11778133 | 13488800 | 11772538 | 10127049 | 11210214 | 9983079 | 7814818.5 | 11232727 | 9667924 | 12634146 | 14740973 | 10715277 |
| P10107 | Anxa1 | Annexin A1 Mus musculus | 1826.49 | 66.47 | 38734 | 25 | 27 | 236659969 | 249002929 | 250478305 | 247777201 | 248102913 | 238987201 | 253689697 | 237620001 | 230798369 | 239516465 | 290394721 | 232798593 | 258055921 | 246275857 | 214724129 |
| P97384 | Anxa11 | Annexin A11 Mus musculus | 445.24 | 18.69 | 54079 | 8 | 8 | 13249033 | 14049104 | 13320381 | 12341820 | 13008711 | 12048478 | 13904148 | 13049373 | 13981681 | 11995794 | 12242348 | 13440257 | 12907563 | 13243451 | 11639620 |
| P07356 | Anxa2 | Annexin A2 Mus musculus | 1632.99 | 70.8 | 38676 | 25 | 25 | 413457761 | 402877729 | 423519713 | 427536897 | 441378049 | 452286113 | 617554753 | 461282945 | 422670913 | 530866305 | 623050241 | 496078081 | 489559073 | 457095841 | 531130625 |
| O35639 | Anxa3 | Annexin A3 Mus musculus | 1289.29 | 65.94 | 36384 | 22 | 22 | 79483705 | 73001289 | 74065025 | 79753297 | 70614609 | 77069433 | 95019601 | 75583761 | 69191881 | 91936809 | 76330065 | 82631561 | 74143601 | 67705873 | 86767121 |
| P97429 | Anxa4 | Annexin A4 Mus musculus | 1416.22 | 68.97 | 35916 | 11 | 21 | 283375777 | 255470785 | 235954209 | 264655041 | 254395585 | 260850481 | 242834785 | 267951617 | 279258977 | 256457793 | 270299009 | 234139633 | 257308977 | 263742417 | 242927329 |
| P55260 | Anxa4 | Annexin A4 Rattus norvegicus | 674.85 | 29.47 | 35849 | 1 | 10 | 100536425 | 85344593 | 78703625 | 87442833 | 84842137 | 86861289 | 80692937 | 92692409 | 96522433 | 78821281 | 82798865 | 72949729 | 86266513 | 91522641 | 76502825 |
| P48036 | Anxa5 | Annexin A5 Mus musculus | 2513.52 | 81.19 | 35752 | 31 | 34 | 850601857 | 746683393 | 836200641 | 775538113 | 756466305 | 722939201 | 724072961 | 824630977 | 911030337 | 734993537 | 858684417 | 839202049 | 714561473 | 862061441 | 906411841 |
| P14824 | Anxa6 | Annexin A6 Mus musculus | 1282.66 | 47.99 | 75885 | 25 | 26 | 57874777 | 59606777 | 50453753 | 57954801 | 48983085 | 54463037 | 57021133 | 57677069 | 64262661 | 56056077 | 47556093 | 57899953 | 55571705 | 61373897 | 54025857 |
| Q07076 | Anxa7 | Annexin A7 Mus musculus | 685.25 | 27.21 | 49925 | 12 | 12 | 35527049 | 36895441 | 38392973 | 38771653 | 40532297 | 37622557 | 36212701 | 37532353 | 35429313 | 36600917 | 39602025 | 37719789 | 36347045 | 39082321 | 34578657 |
| O35643 | Ap1b1 | AP-1 complex subunit beta-1 Mus musculus | 2073.93 | 49.63 | 103935 | 20 | 34 | 43716193 | 42529389 | 50980005 | 37876845 | 44609745 | 55121121 | 50339689 | 52601265 | 61493421 | 48157125 | 57003153 | 63977649 | 48798821 | 53961233 | 61299533 |
| P22892 | Ap1g1 | AP-1 complex subunit gamma-1 Mus musculus | 662.83 | 20.92 | 91350 | 14 | 14 | 26613081 | 26423947 | 23144335 | 23785225 | 26812109 | 26024625 | 21151375 | 27115927 | 27479065 | 24659661 | 26139981 | 26147091 | 26417559 | 28388167 | 24411045 |
| O88512 | Ap1g2 | AP-1 complex subunit gamma-like 2 Mus musculus | 65.18 | 2.4 | 87863 | 2 | 2 | 887665.88 | 570473.915 | 515224.12 | 454511.97 | 424493.69 | 340971.25 | 1 | 1 | 1 | 618367 | 1 | 1 | 518186.19 | 497977.59 | 505188.5 |
| P35585 | Ap1m1 | AP-1 complex subunit mu-1 Mus musculus | 302.74 | 23.17 | 48543 | 7 | 7 | 6224573.5 | 7832745.5 | 8689140 | 5319143.5 | 2913590.5 | 7439850.5 | 6856398 | 5861928.5 | 6542044 | 6184570.5 | 7688444 | 7968838.5 | 5638298 | 6933908.5 | 9303891 |
| P61967 | Ap1s1 | AP-1 complex subunit sigma-1A Mus musculus | 210.85 | 22.15 | 18733 | 2 | 2 | 2613343.5 | 2705115 | 2649773.5 | 1826789.88 | 2470660.5 | 2283664.5 | 1931606.75 | 2947033.75 | 3027061 | 1428321.88 | 2828724 | 2559430.75 | 2551950.5 | 2940408.5 | 2165641 |
| P17426 | Ap2a1 | AP-2 complex subunit alpha-1 Mus musculus | 1066.92 | 33.27 | 107664 | 14 | 21 | 19275267 | 18982619 | 19978341 | 24179009 | 19944041 | 22767683 | 24242077 | 22833303 | 22902463 | 22360763 | 25962901 | 20697413 | 25549999 | 22699535 | 20742939 |
| P18484 | Ap2a2 | AP-2 complex subunit alpha-2 Rattus norvegicus | 821.51 | 22.6 | 104045 | 9 | 16 | 16922133 | 16935013 | 15711798 | 14436035 | 14473565 | 14512767 | 15736232 | 17646913 | 18082425 | 15473647 | 16976157 | 15426482 | 15416942 | 17459709 | 16185081 |
| P62944 | Ap2b1 | AP-2 complex subunit beta Rattus norvegicus | 1888.98 | 48.56 | 104553 | 21 | 35 | 62308241 | 63961745 | 70913601 | 56382481 | 64318025 | 75426113 | 65693937 | 72050961 | 84853321 | 64823637 | 76695273 | 76316321 | 66542965 | 71602969 | 70995681 |
| P84091 | Ap2m1 | AP-2 complex subunit mu Mus musculus | 464.25 | 28.05 | 49655 | 10 | 10 | 12989671 | 13649889 | 12941551 | 11748584 | 9434192 | 12193289 | 11750948 | 9709560 | 12964354 | 13680989 | 11730450 | 8819613 | 14162708 | 13215692 | 12581257 |
| P62743 | Ap2s1 | AP-2 complex subunit sigma Mus musculus | 148.16 | 21.13 | 17018 | 3 | 3 | 4508175 | 4206873 | 3162647 | 3444398 | 2829109 | 3734331 | 2936266.75 | 3645550.75 | 5749181.5 | 3631965 | 3644041 | 3285090.75 | 3575614 | 4087532.25 | 3489878 |
| Q9Z1T1 | Ap3b1 | AP-3 complex subunit beta-1 Mus musculus | 311.82 | 5.79 | 122740 | 6 | 6 | 10678527 | 9409184 | 8660827 | 8851153 | 10371617 | 8963745 | 6743306 | 7899023 | 8326370 | 7511278.5 | 8122873 | 7689817 | 8975975 | 8748051 | 8167061 |
| O54774 | Ap3d1 | AP-3 complex subunit delta-1 Mus musculus | 512.81 | 10.59 | 135081 | 9 | 9 | 8338149 | 9023508 | 8500252 | 6781809 | 7314837.5 | 8143129 | 6972247.5 | 8481893 | 9295818 | 7137738 | 7648257.5 | 9224034 | 8069774.5 | 8541226 | 9098076 |
| P53676 | Ap3m1 | AP-3 complex subunit mu-1 Rattus norvegicus | 218.03 | 16.99 | 46981 | 4 | 4 | 1528154.38 | 1988228.25 | 2611411.5 | 1744980.25 | 1936171.25 | 1876951.5 | 1292383.75 | 1848036.62 | 2250638 | 1900312.12 | 1705611 | 1965679 | 1415186.5 | 1950273.38 | 1345999.62 |
| Q9DCR2 | Ap3s1 | AP-3 complex subunit sigma-1 Mus musculus | 52.29 | 7.77 | 21732 | 1 | 1 | 1 | 1 | 1 | 1 | 1 | 1 | 612416 | 1 | 1 | 1 | 1 | 1 | 1 | 430480.53 | 728452.88 |
| Q80V94 | Ap4e1 | AP-4 complex subunit epsilon-1 Mus musculus | 78.97 | 1.78 | 124845 | 1 | 1 | 681580.38 | 454765.38 | 548853.5175 | 674298.69 | 384769.62 | 1 | 222189.72 | 450039.84 | 326010.31 | 1 | 1 | 322243.22 | 1 | 405152.5 | 315498.97 |
| Q3TAP4 | Ap5b1 | AP-5 complex subunit beta-1 Mus musculus | 89.99 | 3.31 | 94679 | 2 | 2 | 495568.12 | 610638.62 | 653603.31 | 594973.88 | 572988.12 | 526226.44 | 401804.34 | 575674.31 | 418292.03 | 452003.94 | 418735.5 | 502738.31 | 767162.56 | 538318.06 | 551607 |
| O88879 | Apaf1 | Apoptotic protease-activating factor 1 Mus musculus | 59.28 | 1.12 | 141003 | 1 | 1 | 410739.94 | 693278.06 | 611642.69 | 525197.38 | 571885.19 | 743338.06 | 430708.44 | 396215.53 | 495129.16 | 489195.06 | 1025989.94 | 509803.56 | 602517.62 | 598754.31 | 816694.69 |
| Q8R5A3 | Apbb1ip | Amyloid beta A4 precursor protein-binding family B member 1-interacting protein Mus m | 77.88 | 6.12 | 74319 | 2 | 2 | 596750.38 | 539546.88 | 1 | 1 | 1 | 656823.38 | 344331.12 | 521565.945 | 523884.28 | 561225 | 432311.56 | 1 | 424296.28 | 1 | 604253.19 |
| Q8R146 | Apeh | Acylamino-acid-releasing enzyme Mus musculus | 287.14 | 10.93 | 81581 | 5 | 5 | 7637083 | 8027463 | 7691495.5 | 5069089 | 8057225 | 7808023 | 6480799 | 8526049 | 9637132 | 5682103.5 | 7125105 | 8721167 | 8059879 | 8583783 | 7810223 |
| P28352 | Apex1 | DNA-(apurinic or apyrimidinic site) lyase Mus musculus | 595.7 | 40.69 | 35490 | 8 | 8 | 11758977 | 10707655 | 11214372 | 12466522 | 11296019 | 12177805 | 14195106 | 11479274 | 10509507 | 9054792 | 11942599 | 11418962 | 8121510.5 | 10445025 | 13795632 |
| O35841 | Api5 | Apoptosis inhibitor 5 Mus musculus | 768.84 | 43.06 | 56785 | 14 | 14 | 23142681 | 28603149 | 25478085 | 22739941 | 27038451 | 22853977 | 24430127 | 29294141 | 26946749 | 27843669 | 26202861 | 21837185 | 27564417 | 29018987 | 24522001 |
| Q9WVQ5 | Apip | Methylthioribulose-1-phosphate dehydratase Mus musculus | 35 | 6.22 | 26949 | 1 | 1 | 1 | 1187221.75 | 782819.5 | 1508583.25 | 1 | 631545.62 | 956909.44 | 1 | 1 | 343878.75 | 359569.38 | 1 | 815238.31 | 1 | 1 |
| P15943 | Aplp2 | Amyloid-like protein 2 Rattus norvegicus | 65.22 | 2.61 | 86883 | 1 | 1 | 281568.31 | 399158.03 | 118031.78 | 171600.78 | 122762.62 | 1 | 216480.14 | 90776.56 | 177170.2 | 1 | 202827.88 | 124032.35 | 290005.19 | 290598.12 | 106675.86 |
| Q9D7N9 | Apmap | Adipocyte plasma membrane-associated protein Mus musculus | 331.15 | 23.61 | 46434 | 5 | 5 | 2389094.5 | 2061793.88 | 2761905.75 | 2761008.75 | 2897959.75 | 2215951.5 | 1062412.38 | 2953991 | 2318955.25 | 2722312 | 1457125.75 | 2289474.5 | 3468349.5 | 3178003.5 | 2242692.25 |
| P51908 | Apobec1 | C->U-editing enzyme APOBEC-1 Mus musculus | 150.25 | 18.78 | 27522 | 1 | 3 | 4181783 | 3128322 | 3258482 | 3074789.75 | 3974909.5 | 3672350.5 | 1465117.25 | 2411577.5 | 2967318.25 | 2742101.25 | 4218480.5 | 2721056.25 | 2725524 | 3188807.5 | 3910395.25 |
| Q8VBT6 | Apobr | Apolipoprotein B receptor Mus musculus | 1003.48 | 23.89 | 102705 | 16 | 16 | 22938505 | 36570777 | 27218489 | 30706401 | 31003593 | 27281689 | 24215557 | 31592705 | 29532053 | 26602103 | 25542763 | 30403361 | 32004035 | 29774239 | 29522617 |
| Q9DCZ4 | Apoo | MICOS complex subunit Mic26 Mus musculus | 109.78 | 20.2 | 22604 | 2 | 2 | 571167.81 | 388203.75 | 801746.06 | 435004.59 | 431366.28 | 538724.69 | 606572.38 | 869747.38 | 1519030.88 | 883518.8325 | 536762.25 | 631475.69 | 362561.91 | 833082.19 | 642792.38 |
| Q78IK4 | Apool | MICOS complex subunit Mic27 Mus musculus | 389.35 | 33.96 | 29261 | 6 | 6 | 8955794 | 9403101 | 9080211 | 6970131 | 10111369 | 10401712 | 12222685 | 10010262 | 9923166 | 12218955 | 11985959 | 8942015 | 9415324 | 9401879 | 10025885 |
| Q8K3G9 | Appl2 | DCC-interacting protein 13-beta Mus musculus | 37.43 | 3.93 | 73854 | 1 | 1 | 140612.2 | 224415.69 | 157621.38 | 164640.83 | 230919.08 | 176270.98 | 135267.91 | 146267.05 | 171717.42 | 152073.33 | 261592.36 | 158359.36 | 236770.47 | 243048.45 | 167343.94 |
| P08030 | Aprt | Adenine phosphoribosyltransferase Mus musculus | 757.74 | 77.78 | 19724 | 10 | 11 | 70375441 | 67236081 | 71763953 | 76840985 | 74646585 | 69744681 | 74413337 | 67757361 | 65299469 | 78585449 | 72923281 | 71954289 | 72647057 | 67186913 | 82781761 |
| Q8CFQ3 | Aqr | RNA helicase aquarius Mus musculus | 96.52 | 2.5 | 170294 | 2 | 2 | 1 | 301020.62 | 305347.97 | 459881.59 | 1 | 335823.84 | 219919.83 | 487670.31 | 398017.76 | 548657.06 | 297689.75 | 361575.28 | 441242.47 | 223129.64 | 305554.78 |
| P04627 | Araf | Serine/threonine-protein kinase A-Raf Mus musculus | 95.75 | 4.64 | 67581 | 1 | 2 | 204934.8 | 156852.69 | 125078.04 | 328889.34 | 281851.28 | 151224.95 | 184152.72 | 211602.27 | 228109.78 | 244742.2 | 338932.56 | 229766.42 | 224797.92 | 189104.67 | 224131.05 |
| Q4LDD4 | Arap1 | Arf-GAP with Rho-GAP domain, ANK repeat and PH domain-containing protein 1 Mus m | 305.56 | 6.34 | 162276 | 6 | 6 | 4658516 | 4736144 | 1639542.5 | 5982300.5 | 5228927.5 | 4445239 | 2787607.5 | 3995588 | 3013153.75 | 3642768.75 | 4094981.75 | 4202068.5 | 3767929.75 | 4672734 | 2437721.5 |
| Q5XJY5 | Arcn1 | Coatomer subunit delta Mus musculus | 877.29 | 35.23 | 57229 | 16 | 16 | 19772421 | 21118951 | 21787121 | 21443305 | 21915279 | 21086959 | 25692955 | 20864117 | 20318203 | 23605525 | 21993439 | 22987913 | 21358515 | 21585625 | 25003069 |
| P84078 | Arf1 | ADP-ribosylation factor 1 Mus musculus | 1015.5 | 80.66 | 20697 | 3 | 16 | 139083505 | 125766337 | 118190753 | 113588457 | 106964153 | 115192753 | 102559345 | 136002273 | 152471473 | 101736161 | 111531409 | 125078089 | 104364377 | 140886993 | 125164697 |
| P84082 | Arf2 | ADP-ribosylation factor 2 Rattus norvegicus | 868.88 | 59.67 | 20746 | 1 | 13 | 113368641 | 100004065 | 92356241 | 84586289 | 82072665 | 88326457 | 80421793 | 107672361 | 123965273 | 81417129 | 84977977 | 100319441 | 82104737 | 111312657 | 99349025 |
| P61750 | Arf4 | ADP-ribosylation factor 4 Mus musculus | 735 | 60 | 20397 | 3 | 10 | 69729241 | 66907981 | 59064017 | 48535037 | 53583585 | 55590541 | 52021729 | 65755505 | 81582473 | 56630965 | 51637493 | 63377285 | 56494753 | 72634089 | 56437065 |
| P84083 | Arf5 | ADP-ribosylation factor 5 Rattus norvegicus | 792.64 | 65.56 | 20530 | 4 | 12 | 101121185 | 101248833 | 91550977 | 83105881 | 82470113 | 87986697 | 80175897 | 101091321 | 118263841 | 82238729 | 84167761 | 95804473 | 82481769 | 108785025 | 89195081 |
| P62331 | Arf6 | ADP-ribosylation factor 6 Mus musculus | 278.71 | 48 | 20082 | 5 | 5 | 9888003 | 7395230 | 6450047 | 6249971.5 | 7713709 | 8562729 | 6966621 | 8639470 | 9316416 | 6764912.5 | 7240679 | 7861416.5 | 7621024.5 | 8536791 | 8740473 |
| Q3MID3 | Arfgap2 | ADP-ribosylation factor GTPase-activating protein 2 Rattus norvegicus | 218.78 | 9.62 | 56556 | 4 | 4 | 1611204.88 | 3975532.5 | 2190533.5 | 465425.59 | 988052.31 | 1432197.25 | 1265287.38 | 3079709.5 | 250910.69 | 866432.81 | 1297828.88 | 2188001.75 | 2200807.5 | 677176.31 | 1980555 |
| D4A631 | Arfgef1 | Brefeldin A-inhibited guanine nucleotide-exchange protein 1 Rattus norvegicus | 206.15 | 2.65 | 207890 | 1 | 4 | 1543873 | 1527367.62 | 1536154.12 | 1412977 | 1764370.25 | 1658933.75 | 1279305.25 | 1453097.75 | 1909550.62 | 1259701.5 | 1277784.25 | 1608816.25 | 1425047.5 | 1537379.12 | 1310279.75 |
| A2A5R2 | Arfgef2 | Brefeldin A-inhibited guanine nucleotide-exchange protein 2 Mus musculus | 312.42 | 4.85 | 202240 | 3 | 6 | 2486530 | 2432411.5 | 2001258.38 | 2357806.25 | 2523400.25 | 2346854.25 | 2045441.12 | 2095549.88 | 2510827.5 | 2269214.5 | 2040980.38 | 2262210.25 | 2415327 | 2280477.75 | 1825929.12 |
| Q9JHU5 | Arfip1 | Arfaptin-1 Rattus norvegicus | 217.28 | 12.3 | 40779 | 3 | 3 | 4189904.25 | 2635163 | 3106042.25 | 3730031.75 | 5448298.5 | 3524568.75 | 1261589.88 | 5514972 | 2839374.5 | 3410929.5 | 1438950.38 | 3324222.5 | 3481790.5 | 6274663 | 3040812.75 |
| Q63055 | Arfrp1 | ADP-ribosylation factor-related protein 1 Rattus norvegicus 2 | 74.71 | 8.96 | 22659 | 1 | 1 | 609896.19 | 1 | 775940.44 | 1 | 896574.25 | 912373.62 | 428269.28 | 639798.38 | 765341.06 | 524996.12 | 834572.94 | 919559.56 | 711419.06 | 671008.06 | 858385.88 |
| Q3UL36 | Arglu1 | Arginine and glutamate-rich protein 1 Mus musculus | 38.25 | 2.95 | 32887 | 1 | 1 | 1 | 1 | 13492843 | 444879.12 | 370253.31 | 1 | 1 | 3421383.5 | 12908483 | 1 | 6711777.32 | 15849037 | 7841415 | 2713957.75 | 442699.53 |
| Q5FWK3 | Arhgap1 | Rho GTPase-activating protein 1 Mus musculus | 352.45 | 25.28 | 50411 | 8 | 8 | 7618287.5 | 5292056 | 5819092 | 5097715.5 | 5488070 | 5444264 | 5536548.5 | 6002909 | 5523277.5 | 5991380.5 | 6365420 | 5741169.5 | 7541197.5 | 6613854.5 | 5417336 |

| | | | | | | | | | | | | | | | | | | | | | | |
|---|---|---|---|---|---|---|---|---|---|---|---|---|---|---|---|---|---|---|---|---|---|---|
| Q3UIA2 | Arhgap17 | Rho GTPase-activating protein 17 Mus musculus | 444.6 | 14.89 | 92202 | 10 | 10 | 12156895 | 8757226 | 8017522 | 8870471 | 10846009 | 8406479 | 7135234 | 8053279 | 8126791 | 6930048 | 9586567 | 7796212.5 | 8925411 | 10199612 | 8748655 |
| Q8BYW1 | Arhgap25 | Rho GTPase-activating protein 25 Mus musculus | 636.68 | 23.61 | 73383 | 12 | 12 | 16615914 | 18968055 | 18833181 | 20017501 | 16429793 | 18103999 | 17795133 | 18004043 | 17896371 | 18711627 | 19051123 | 19705165 | 20088073 | 18120113 | 17334505 |
| Q640N3 | Arhgap30 | Rho GTPase-activating protein 30 Mus musculus | 311.04 | 5.99 | 120114 | 5 | 5 | 861896.06 | 966804 | 1010682.69 | 953408.6725 | 974251.94 | 918593.25 | 496853.69 | 1040884.56 | 835076.06 | 1702824 | 902714.31 | 1068083.88 | 669870.56 | 1127751.25 | 850157 |
| Q3TBD2 | Arhgap45 | Rho GTPase-activating protein 45 Mus musculus | 567.96 | 15.14 | 122902 | 11 | 11 | 9840119 | 10974965 | 11893690 | 12042698 | 12688360 | 10610252 | 15252583 | 12504563 | 10994698 | 13291527 | 13771200 | 11943539 | 10176933 | 11751056 | 12532616 |
| Q5XI73 | Arhgdia | Rho GDP-dissociation inhibitor 1 Rattus norvegicus | 883.41 | 80.39 | 23407 | 12 | 12 | 202342753 | 186281601 | 218119665 | 207268609 | 209968353 | 202380657 | 278526785 | 211458785 | 177658481 | 245643745 | 216567361 | 131556609 | 221175905 | 183573841 | 240978241 |
| Q61599 | Arhgdib | Rho GDP-dissociation inhibitor 2 Mus musculus | 530.49 | 71.5 | 22851 | 11 | 11 | 93330705 | 72605081 | 95821105 | 106086849 | 96515977 | 96419425 | 161940225 | 89602089 | 74450009 | 131120697 | 87446713 | 119971329 | 102847977 | 80142689 | 149698097 |
| Q61210 | Arhgef1 | Rho guanine nucleotide exchange factor 1 Mus musculus | 975.61 | 26.63 | 102805 | 18 | 18 | 14399199 | 18696363 | 17802819 | 19125075 | 15907942 | 20354721 | 17313389 | 16861017 | 19446019 | 19201095 | 15004285 | 17720505 | 20062863 | 19964235 | 18879257 |
| A2AWP8 | Arhgef10l | Rho guanine nucleotide exchange factor 10-like protein Mus musculus | 115.37 | 3.2 | 139987 | 3 | 3 | 1919286 | 1586151.38 | 1712004.5 | 2007021.5 | 1053128.5 | 1523692.38 | 1231562.88 | 223682.25 | 1761331.62 | 1588966.25 | 1896007.75 | 1996591 | 1883155.25 | 1584491.5 | 2623170.5 |
| Q60875 | Arhgef2 | Rho guanine nucleotide exchange factor 2 Mus musculus | 950.87 | 22.94 | 111974 | 18 | 18 | 21502987 | 18211187 | 19553369 | 20133037 | 22657689 | 18763819 | 18612121 | 17564711 | 16143507 | 22088847 | 22430087 | 19457757 | 19606231 | 17453007 | 22291915 |
| Q8K4I3 | Arhgef6 | Rho guanine nucleotide exchange factor 6 Mus musculus | 394.07 | 10.64 | 87051 | 6 | 7 | 8605478 | 16651154 | 9383427 | 8057653.5 | 8661356 | 8010575.5 | 7581453.5 | 7820426.5 | 9390173 | 9139196 | 8749298 | 7897695 | 7962568 | 7961265.5 | 7604135 |
| O55043 | Arhgef7 | Rho guanine nucleotide exchange factor 7 Rattus norvegicus | 409.57 | 17.18 | 73140 | 6 | 7 | 4854628 | 12482605 | 5101338 | 4127232 | 4401331 | 4840925.5 | 5147205 | 4162827 | 5072120 | 5107014 | 4147329.25 | 4718814 | 4563725 | 3806795.5 | 4369983 |
| A2BH40 | Arid1a | AT-rich interactive domain-containing protein 1A Mus musculus | 160.72 | 1.14 | 242092 | 0 | 2 | 27109421 | 27486317 | 30441325 | 29938315 | 29588977 | 25654037 | 28426233 | 25855039 | 25336797 | 24106743 | 28633063 | 22574985 | 27561691 | 27858853 | 19271655 |
| E9Q4N7 | Arid1b | AT-rich interactive domain-containing protein 1B Mus musculus | 145.35 | 1.43 | 237795 | 1 | 2 | 629789.69 | 1076270.62 | 663479.69 | 1036901.88 | 942483.75 | 971043.62 | 948614.44 | 808841.5 | 886893.19 | 903848.1875 | 960585.81 | 748142.69 | 791900.17 | 703575.06 | 755297.12 |
| E9Q7E2 | Arid2 | AT-rich interactive domain-containing protein 2 Mus musculus | 60.99 | 0.49 | 195988 | 1 | 1 | 1190090.12 | 779792.44 | 634920.94 | 854276.69 | 935487.62 | 840472.94 | 769822.31 | 570187.94 | 862552.06 | 847640.75 | 363091.84 | 800004.94 | 829700.62 | 664302.88 | 664275.07 |
| Q9Z1K5 | Arih1 | E3 ubiquitin-protein ligase ARIH1 Mus musculus | 132.77 | 7.39 | 64017 | 2 | 2 | 1302469.5 | 1893415.38 | 1967524.62 | 1277822.88 | 1766508.88 | 1361383.38 | 1117836 | 1757647.5 | 2719195.75 | 887204 | 2007927.75 | 1733061.5 | 1113160 | 1685939.88 | 1478844 |
| P61211 | Arl1 | ADP-ribosylation factor-like protein 1 Mus musculus | 382.62 | 52.49 | 20412 | 6 | 7 | 14524514 | 15820364 | 15911993 | 10523001 | 13318608 | 14476280 | 13439196 | 13410030 | 20727153 | 13475470 | 11812956 | 16001739 | 14585096 | 17153327 | 14951599 |
| O08697 | Arl2 | ADP-ribosylation factor-like protein 2 Rattus norvegicus | 383.51 | 45.65 | 20836 | 5 | 5 | 2996704 | 3156385.75 | 5255592.5 | 5050121.5 | 3702653.25 | 3734051 | 3828941.5 | 3749187.25 | 6236497.5 | 4069006 | 2780054.25 | 4700291 | 4580280 | 3180070.5 | 5324075 |
| Q9WUL7 | Arl3 | ADP-ribosylation factor-like protein 3 Mus musculus | 462.95 | 53.3 | 20487 | 8 | 8 | 5054397.5 | 5894785 | 5649855.5 | 4968932.5 | 5805500 | 6703813 | 6602306.5 | 7644625.5 | 5725963.5 | 5654499 | 5341239.5 | 4105115 | 5391750 | 5193307.5 | 6026990.5 |
| P61208 | Arl4c | ADP-ribosylation factor-like protein 4C Mus musculus | 41.78 | 7.81 | 21487 | 1 | 1 | 1230139.12 | 2216677.25 | 1874746.12 | 2059655.88 | 1853685.25 | 1400394.25 | 1460373.12 | 1482902.38 | 1751511.12 | 1854653.62 | 1402360.12 | 1627615.12 | 1408120.38 | 1669033.12 | 1814503.88 |
| Q8R5J9 | Arl6ip5 | PRA1 family protein 3 Mus musculus | 77.99 | 10.64 | 21558 | 2 | 2 | 9217537 | 2853742.5 | 2801725.25 | 2138155.5 | 2987978.5 | 1631919 | 1509586.25 | 2646058 | 3056150.75 | 2663004.25 | 3465081 | 2621700.5 | 3238105 | 3157022 | 3837860.25 |
| Q8VEH3 | Arl8a | ADP-ribosylation factor-like protein 8A Mus musculus | 362.06 | 44.62 | 21390 | 3 | 7 | 19094271 | 21402879 | 19415433 | 15555749 | 13841333 | 16782857 | 18825439 | 19703369 | 13938542 | 16222821 | 20623885 | 18662911 | 21103329 | 20534453 | 17764437 |
| Q66HA6 | Arl8b | ADP-ribosylation factor-like protein 8B Rattus norvegicus | 540.17 | 61.83 | 21539 | 6 | 10 | 34705193 | 28682611 | 34539589 | 33500075 | 34521117 | 33286639 | 37565069 | 36030121 | 38478121 | 36012605 | 35636813 | 34849265 | 37710021 | 37206641 | 26960449 |
| Q9D7A8 | Armc1 | Armadillo repeat-containing protein 1 Mus musculus | 57.15 | 9.57 | 31247 | 1 | 1 | 409928.34 | 566910.56 | 1 | 1 | 341695.69 | 380072.28 | 1 | 482771.81 | 1 | 1 | 340202.78 | 1 | 566819.81 | 333872.78 | 1 |
| Q9D0L7 | Armc10 | Armadillo repeat-containing protein 10 Mus musculus | 172.68 | 15.36 | 33311 | 3 | 3 | 1656080 | 1723679.12 | 1740246.5 | 1079363 | 1922319.38 | 1405192 | 1244279.62 | 2040289.88 | 2896997 | 1481595.62 | 2145202 | 1256589.88 | 1487177 | 2196148 | 1460184.62 |
| Q8BNU0 | Armc6 | Armadillo repeat-containing protein 6 Mus musculus | 134.17 | 8.33 | 50683 | 2 | 2 | 621959.12 | 564142.25 | 682003.25 | 615147.5 | 650408.81 | 816603.5 | 306071.31 | 724342.94 | 589406.485 | 510608.19 | 782154.5 | 375031.12 | 761217.44 | 593368.7875 | 455072.09 |
| Q9DBR3 | Armc8 | Armadillo repeat-containing protein 8 Mus musculus | 104.88 | 4.75 | 75364 | 2 | 2 | 1089267.25 | 789229.56 | 1345845.12 | 875397.12 | 936497.12 | 963148.75 | 1292819.12 | 1179779.88 | 796784.12 | 856849.56 | 1044881.31 | 1104187.38 | 1050806 | 842398.81 | 972192.94 |
| Q9WV32 | Arpc1b | Actin-related protein 2/3 complex subunit 1B Mus musculus | 931.68 | 55.65 | 41064 | 16 | 16 | 58040457 | 61409085 | 63811549 | 67494473 | 73469161 | 65445705 | 78073017 | 50087845 | 55039397 | 75084009 | 77510545 | 65589801 | 75336849 | 59851225 | 82514329 |
| Q9CVB6 | Arpc2 | Actin-related protein 2/3 complex subunit 2 Mus musculus | 1177.12 | 70.67 | 34357 | 19 | 19 | 127419329 | 119070625 | 129353473 | 118420425 | 118105409 | 122066385 | 105036097 | 132708713 | 146663745 | 106389665 | 136406401 | 142020545 | 114715761 | 142832033 | 133115209 |
| Q9JM76 | Arpc3 | Actin-related protein 2/3 complex subunit 3 Mus musculus | 481.08 | 53.37 | 20525 | 8 | 8 | 84300433 | 103555009 | 86298033 | 81326905 | 86582385 | 83992521 | 75467857 | 86834753 | 96797529 | 79602889 | 92912065 | 76014977 | 88421617 | 92808241 | 75356185 |
| P59999 | Arpc4 | Actin-related protein 2/3 complex subunit 4 Mus musculus | 400.77 | 54.17 | 19667 | 7 | 7 | 155915953 | 140423329 | 129911689 | 148521505 | 154074081 | 131829377 | 136309761 | 125166545 | 124687577 | 153481665 | 135947633 | 120669649 | 146090577 | 129732993 | 143611649 |
| Q9CPW4 | Arpc5 | Actin-related protein 2/3 complex subunit 5 Mus musculus | 544.13 | 58.94 | 16288 | 8 | 8 | 81309769 | 83109705 | 80807793 | 87037905 | 88914985 | 80124361 | 83064689 | 76747969 | 74075025 | 85783841 | 90077553 | 76819625 | 85419337 | 80516185 | 74665113 |
| P29066 | Arrb1 | Beta-arrestin-1 Rattus norvegicus | 199.58 | 15.31 | 47020 | 4 | 5 | 2347984.25 | 1851920.12 | 3963660.5 | 2899776.5 | 1999813 | 2863147.5 | 3317191.75 | 2461558.25 | 2402794.5 | 3195600 | 2880066.75 | 2565447.5 | 1993241 | 2792048.25 | 3881857.5 |
| P29067 | Arrb2 | Beta-arrestin-2 Rattus norvegicus | 455.84 | 20.73 | 46340 | 8 | 9 | 6769582 | 8025146 | 7476405 | 7378432 | 8299979 | 7813721 | 9443241 | 7666722 | 5831552 | 8431216 | 7064650.5 | 8412936 | 6619280 | 7438428 | 8513285 |
| P50428 | Arsa | Arylsulfatase A Mus musculus | 197.28 | 14.23 | 53748 | 3 | 4 | 12926354 | 3676284.5 | 3297498 | 3617777.5 | 5369760 | 4796202 | 5711402 | 4262608 | 5765558.5 | 6130536.5 | 4103650 | 3973085.75 | 4483086 | 4227171 | 3221848 |
| P50429 | Arsb | Arylsulfatase B Mus musculus | 676.72 | 24.16 | 59647 | 12 | 12 | 61991101 | 75576577 | 45951285 | 58174565 | 83004793 | 72145537 | 58180777 | 75767289 | 69533577 | 48830437 | 72146729 | 53941297 | 74034577 | 72124025 | 49592241 |
| Q91WU5 | As3mt | Arsenite methyltransferase Mus musculus | 119.12 | 8.51 | 41794 | 2 | 2 | 1717695.88 | 1549232.75 | 2006569.12 | 1131476.38 | 1260063.75 | 1313280.5 | 1426571.423 | 1584865.25 | 2132657.75 | 675482.19 | 1234242.62 | 1658469.88 | 918581.88 | 1613985 | 1239361 |
| Q9WV54 | Asah1 | Acid ceramidase Mus musculus | 647.15 | 34.01 | 44670 | 10 | 10 | 28765885 | 34030621 | 29372711 | 31656023 | 31882991 | 31057137 | 32931557 | 32294367 | 34239161 | 31782309 | 32115421 | 30675539 | 33614865 | 31853691 | 29890915 |
| Q9JHE3 | Asah2 | Neutral ceramidase Mus musculus | 37.1 | 1.72 | 83509 | 1 | 1 | 454990.69 | 401785 | 349317.94 | 207973.19 | 360640.59 | 396846.16 | 398794.94 | 422351.38 | 429175.72 | 329608.38 | 353700.34 | 345058.22 | 393525.12 | 337461.31 | 276504.16 |
| Q1AAU6 | Asap1 | Arf-GAP with SH3 domain, ANK repeat and PH domain-containing protein 1 Rattus norv | 459.92 | 9.53 | 127088 | 9 | 9 | 8820306 | 9953314 | 9387332 | 9091122 | 9597520 | 9544995 | 10366380 | 9029489 | 9568718 | 8646686 | 11066423 | 10552254 | 10286497 | 10128631 | 10680721 |
| Q9DAP7 | Asf1b | Histone chaperone ASF1B Mus musculus | 164.03 | 31.19 | 22464 | 3 | 3 | 249335.05 | 1 | 183284.09 | 330287.91 | 1 | 217639.86 | 569929 | 238677.11 | 280688.16 | 449893.12 | 317862.2975 | 232035.16 | 378078.94 | 488214.81 | 173120.28 |
| Q91X20 | Ash2l | Set1/Ash2 histone methyltransferase complex subunit ASH2 Mus musculus | 112.32 | 5.3 | 68250 | 2 | 2 | 690015.06 | 1 | 364462.91 | 1 | 613051.56 | 714846.12 | 1088333 | 796226.06 | 604197.88 | 560488.06 | 1613508 | 1584157.75 | 454052.47 | 722250 | 756632.12 |
| Q91YI0 | Asl | Argininosuccinate lyase Mus musculus | 528.51 | 28.23 | 51739 | 9 | 9 | 10089356 | 7535517.5 | 7922997.5 | 7927115 | 8766312 | 7199255 | 9289263 | 7549103.5 | 8148757 | 8473687 | 9394561 | 9595811 | 7033326.5 | 6675470 | 9356511 |
| O54984 | Asna1 | ATPase Asna1 Mus musculus | 491.22 | 48.56 | 38823 | 11 | 11 | 10520835 | 10712636 | 11106910 | 10759026 | 8052762 | 11138663 | 15411631 | 11149654 | 12599062 | 9894251 | 12256825 | 13523025 | 12745601 | 11721007 | 15265231 |
| Q61024 | Asns | Asparagine synthetase [glutamine-hydrolyzing] Mus musculus | 807.35 | 33.51 | 64283 | 14 | 14 | 22282757 | 16045791 | 22277965 | 22302875 | 22211675 | 30421359 | 24829973 | 23572991 | 23361801 | 23082309 | 32671895 | 23484009 | 20606761 | 21644909 | 33878257 |
| Q8BSY0 | Asph | Aspartyl/asparaginyl beta-hydroxylase Mus musculus | 704.88 | 24.16 | 83042 | 11 | 11 | 8928812 | 9671144 | 9140283 | 10347683 | 10067577 | 8499219 | 11044593 | 9041781 | 9980307 | 10403306 | 9030250 | 8619825 | 10198616 | 8208340 | 8399100 |
| Q8VBT9 | Aspscr1 | Tether containing UBX domain for GLUT4 Mus musculus | 141.82 | 6.73 | 59796 | 2 | 2 | 654962.44 | 539152.06 | 575547.265 | 693931.62 | 414142.94 | 1 | 389035.16 | 425997.88 | 395511.19 | 399742.31 | 1 | 348509.91 | 1 | 477888.97 | 603544.5 |
| Q8C0M9 | Asrgl1 | Isoaspartyl peptidase/L-asparaginase Mus musculus | 136.57 | 11.04 | 33950 | 3 | 3 | 5926281 | 9226601 | 9620158 | 3857887.75 | 3708439.25 | 8480114 | 7026979 | 9651371 | 9468516 | 6331902 | 7322392.5 | 6017175 | 6265015 | 8800451 | 5518763.5 |
| P16460 | Ass1 | Argininosuccinate synthase Mus musculus | 608.06 | 35.92 | 46584 | 11 | 11 | 13874908 | 10818543 | 12943893 | 12171462 | 10458345 | 14515763 | 17842701 | 14223831 | 11712321 | 13100612 | 13462029 | 12109435 | 10883541 | 13761638 | 15225647 |
| Q505J9 | Atad1 | ATPase family AAA domain-containing protein 1 Rattus norvegicus | 124.53 | 7.76 | 40717 | 2 | 2 | 1973895.75 | 2010580.62 | 2134381.75 | 2198934 | 2198041.25 | 1856553.62 | 1334367 | 1916852.75 | 2772469 | 1811794.62 | 2531929 | 1834967 | 1686575.75 | 2012126.75 | 1996360.5 |
| Q3KRE0 | Atad3 | ATPase family AAA domain-containing protein 3 Rattus norvegicus | 182.8 | 7.28 | 66759 | 3 | 3 | 1 | 1 | 153757.5 | 1 | 196889.88 | 254735.39 | 295225.88 | 1215284.38 | 277314.22 | 380743.91 | 288744.34 | 286751.28 | 231031.12 | 314241.22 | 455088.31 |
| Q60765 | Atf3 | Cyclic AMP-dependent transcription factor ATF-3 Mus musculus | 37.84 | 5.52 | 20722 | 1 | 1 | 136233.5975 | 121549.55 | 125725.25 | 121069.61 | 176589.98 | 93367.38 | 143561.59 | 122578.21 | 194032.61 | 115292.76 | 139997.33 | 91364.77 | 211305.41 | 157177.95 | 99572.75 |
| Q8C0J2 | Atg16l1 | Autophagy-related protein 16-1 Mus musculus | 37.68 | 4.61 | 68172 | 1 | 1 | 283115.12 | 277572.12 | 243052.09 | 295654.28 | 251957.73 | 245952.16 | 180505.14 | 279015.22 | 234166.92 | 297019.28 | 508323 | 315693 | 252249.78 | 252605.56 | 459719.69 |
| Q6AZ50 | Atg3 | Ubiquitin-like-conjugating enzyme ATG3 Rattus norvegicus | 117.78 | 10.51 | 35822 | 3 | 3 | 836774.06 | 1038730.56 | 2332083 | 2298163.25 | 2343778 | 2393637.25 | 2190133 | 2603733.5 | 2744377.5 | 2552920 | 2296206.25 | 2483532 | 2901585.5 | 999032.5 | 2449056.25 |
| Q8BGE6 | Atg4b | Cysteine protease ATG4B Mus musculus | 356.4 | 26.46 | 44375 | 7 | 7 | 5191828.5 | 3700420.25 | 4350060 | 3010067.5 | 3811216.75 | 5122335 | 4229345.5 | 3892771.5 | 5145357.5 | 4942341 | 4577149 | 4554911.5 | 3632259.5 | 4089643 | 4433458.5 |
| Q9D906 | Atg7 | Ubiquitin-like modifier-activating enzyme ATG7 Mus musculus | 879.05 | 30.95 | 77520 | 17 | 17 | 21356659 | 16975681 | 21445151 | 22598491 | 21669667 | 20748889 | 23379193 | 22031119 | 20777477 | 19290371 | 23502855 | 22678727 | 17832415 | 23813617 | 19719601 |
| Q9CWJ9 | Atic | Bifunctional purine biosynthesis protein PURH Mus musculus | 2297.18 | 75.68 | 64217 | 34 | 34 | 100501897 | 103995289 | 114436017 | 101709217 | 103917497 | 107134609 | 109766273 | 100694929 | 101800529 | 99483641 | 109387233 | 103311553 | 104208673 | 99582721 | 108660777 |
| Q6PA06 | Atl2 | Atlastin-2 Mus musculus | 92.44 | 4.12 | 66224 | 2 | 2 | 733836.25 | 899531.75 | 923835.75 | 877421.19 | 803702.88 | 972467.75 | 1475015.75 | 963623.56 | 894900.94 | 1103005.38 | 906268.88 | 1224530.5 | 1152944.62 | 1186608.25 | 902368.69 |
| Q91YH5 | Atl3 | Atlastin-3 Mus musculus | 719.84 | 34.01 | 60575 | 12 | 12 | 12180010 | 10808828 | 10887446 | 9888703 | 10379211 | 12343402 | 10396174 | 11914342 | 13762249 | 10167906 | 10891593 | 10601399 | 9346864 | 11263913 | 10221559 |
| O08997 | Atox1 | Copper transport protein ATOX1 Mus musculus | 342.18 | 88.24 | 7338 | 5 | 5 | 72652377 | 90571233 | 76901185 | 84529169 | 80775057 | 70803777 | 66235185 | 70460449 | 78487345 | 75845785 | 75851489 | 64893409 | 83123697 | 80838657 | 68297561 |
| P98197 | Atp11a | Probable phospholipid-transporting ATPase IH Mus musculus | 35.86 | 1.1 | 135502 | 1 | 1 | 1 | 1 | 216804.14 | 1 | 192745.98 | 1 | 1 | 1 | 266671.16 | 1 | 184822.95 | 293481.91 | 1 | 1 | 1 |
| Q9Z1W8 | Atp12a | Potassium-transporting ATPase alpha chain 2 Mus musculus | 170.42 | 2.71 | 114727 | 1 | 4 | 17240277 | 17120835 | 17445753 | 14583029 | 18905811 | 17289627 | 15093223 | 17531965 | 18836693 | 14755057 | 18857025 | 16331465 | 17359581 | 18265481 | 16729367 |
| Q9EPE9 | Atp13a1 | Manganese-transporting ATPase 13A1 Mus musculus | 294.17 | 9 | 132388 | 6 | 6 | 3277617 | 3065685.75 | 4283905 | 3475658.5 | 4203536 | 2474566 | 2997779.5 | 3451813 | 3325151 | 2347664.25 | 3963180.75 | 3785433.75 | 4284850 | 3696249.5 | 3962576 |
| Q8VDN2 | Atp1a1 | Sodium/potassium-transporting ATPase subunit alpha-1 Mus musculus | 2261.13 | 38.22 | 112982 | 22 | 35 | 233579809 | 261796497 | 257596817 | 243048401 | 255043793 | 239193121 | 235463025 | 237674049 | 260718497 | 250261585 | 257937313 | 236217665 | 260303953 | 230426769 | 240820913 |
| P06686 | Atp1a2 | Sodium/potassium-transporting ATPase subunit alpha-2 Rattus norvegicus | 950.36 | 15.1 | 112217 | 1 | 13 | 95076665 | 101328361 | 101793449 | 94733745 | 107726785 | 101363489 | 90027121 | 97736649 | 110475225 | 99766073 | 110204897 | 94060201 | 105178249 | 101189073 | 97880833 |
| P97370 | Atp1b3 | Sodium/potassium-transporting ATPase subunit beta-3 Mus musculus | 329.66 | 30.22 | 31776 | 5 | 5 | 14856685 | 11095817 | 12642517 | 9760679 | 10397499 | 11546070 | 9921414 | 12708262 | 11021067 | 9239713 | 11573889 | 10498268 | 10770833 | 11296063 | 8864323 |
| Q64578 | Atp2a1 | Sarcoplasmic/endoplasmic reticulum calcium ATPase 1 Rattus norvegicus | 625.01 | 12.98 | 109409 | 1 | 9 | 4560039.5 | 3859643 | 7743334 | 4583503 | 3654250.75 | 4812989.5 | 9044203 | 5744301.5 | 5078940 | 8007796.5 | 6780007.5 | 7323416 | 6540858.5 | 3794250.5 | 8419179 |
| O55143 | Atp2a2 | Sarcoplasmic/endoplasmic reticulum calcium ATPase 2 Mus musculus | 1778.63 | 35.06 | 114858 | 19 | 30 | 84705337 | 74889233 | 78532473 | 81246145 | 80584153 | 74211793 | 78800913 | 77574761 | 70837201 | 74358913 | 81667353 | 65258821 | 71248033 | 72375777 | 68934449 |
| Q64518 | Atp2a3 | Sarcoplasmic/endoplasmic reticulum calcium ATPase 3 Mus musculus | 740.84 | 15.9 | 113638 | 7 | 12 | 21217385 | 21617743 | 20382085 | 22246409 | 24183677 | 21881139 | 26024949 | 24156313 | 23034363 | 24962317 | 23945665 | 19828811 | 22295171 | 21749673 | 21179181 |
| G5E829 | Atp2b1 | Plasma membrane calcium-transporting ATPase 1 Mus musculus | 1028.14 | 18.03 | 134747 | 11 | 18 | 27450699 | 31212519 | 29367493 | 24706963 | 28031679 | 27508985 | 29637501 | 27770939 | 28687221 | 28764721 | 27536361 | 27577751 | 31252293 | 29117469 | 28338429 |
| Q64568 | Atp2b3 | Plasma membrane calcium-transporting ATPase 3 Rattus norvegicus | 544.51 | 7.15 | 138559 | 2 | 9 | 14151445 | 16483920 | 15763355 | 12696652 | 14428417 | 14029395 | 14469745 | 14294032 | 15635859 | 14701702 | 14190540 | 13559751 | 16395066 | 15724509 | 13715866 |
| Q03265 | Atp5f1a | ATP synthase subunit alpha, mitochondrial Mus musculus | 2133.34 | 60.94 | 59753 | 5 | 34 | 320112897 | 370741889 | 391404577 | 381308225 | 352192897 | 345301761 | 508392321 | 373155937 | 306088321 | 408230369 | 368013025 | 392308481 | 396195969 | 352835393 | 354202305 |
| P15999 | Atp5f1a | ATP synthase subunit alpha, mitochondrial Rattus norvegicus | 1877.48 | 54.25 | 59754 | 1 | 30 | 267228273 | 310851905 | 331039073 | 328174785 | 295759873 | 286685697 | 434455201 | 307517761 | 257606833 | 353191553 | 287215329 | 330597345 | 328253761 | 295125665 | 304283329 |
| P56480 | Atp5f1b | ATP synthase subunit beta, mitochondrial Mus musculus | 2406.32 | 78.26 | 56300 | 29 | 33 | 297902081 | 322207489 | 360714177 | 311538945 | 312757313 | 342934305 | 406174497 | 345349409 | 328851841 | 343275937 | 325838113 | 440164961 | 313371521 | 317626273 | 361723265 |
| Q91VR2 | Atp5f1c | ATP synthase subunit gamma, mitochondrial Mus musculus | 513.01 | 38.59 | 32886 | 10 | 10 | 29189741 | 35319389 | 32932073 | 34564013 | 34362513 | 31789083 | 30418841 | 31722565 | 27736379 | 28983405 | 34290293 | 25452151 | 30775957 | 31754009 | 27221585 |
| Q9D3D9 | Atp5f1d | ATP synthase subunit delta, mitochondrial Mus musculus | 205.66 | 49.4 | 17600 | 4 | 4 | 24280685 | 31734101 | 27633073 | 29550233 | 28133089 | 27721343 | 28333103 | 27242657 | 31518557 | 27988873 | 26115473 | 29310919 | 29554891 | 28393305 | 22510851 |
| O35143 | ATP5IF1 | ATPase inhibitor, mitochondrial Mus musculus | 45.13 | 8.49 | 12159 | 1 | 1 | 3372996.75 | 2822545.75 | 3568362 | 4602494 | 3475411.5 | 2664090.75 | 4331315 | 2383916.25 | 2342643.5 | 4543357 | 3201504.5 | 4427422.5 | 3900264.25 | 2636468.5 | 2748270.25 |
| P56383 | Atp5mc2 | ATP synthase F(0) complex subunit C2, mitochondrial Mus musculus | 111.41 | 21.23 | 15476 | 1 | 1 | 491922.41 | 4113024.5 | 2933016 | 1811611.5 | 2245393.25 | 349139.5 | 2485982.75 | 1813390.62 | 3958860 | 3662455.25 | 2739116 | 3293598 | 3000784.75 | 2978234.25 | 3714006.75 |
| Q78IK2 | Atp5md | Up-regulated during skeletal muscle growth protein 5 Mus musculus | 147.05 | 43.1 | 6382 | 2 | 2 | 5997268.5 | 5239435 | 4859546 | 5485673.5 | 6008910 | 5508034 | 2263053.5 | 5491307.5 | 5783245.5 | 5334869.5 | 5243256 | 4586493 | 5574884.5 | 5224262 | 4495753.5 |
| Q06185 | Atp5me | ATP synthase subunit e, mitochondrial Mus musculus | 106.08 | 42.25 | 8236 | 3 | 3 | 7695626 | 3353336 | 1131103.38 | 1549188.88 | 1145770.88 | 1291408.12 | 2675799 | 1471115 | 2598695.75 | 4024445.5 | 1601438.25 | 3603271.25 | 1144902 | 1866529 | 2481335.25 |
| P56135 | Atp5mf | ATP synthase subunit f, mitochondrial Mus musculus | 81.01 | 23.86 | 10344 | 1 | 2 | 17268877 | 15526217 | 13439054 | 11799988 | 15739141 | 15425487 | 14445295 | 14896276 | 17162229 | 14055054 | 12565972 | 12022511 | 15572601 | 13730737 | 12123355 |
| Q9CQQ7 | Atp5pb | ATP synthase F(0) complex subunit B1, mitochondrial Mus musculus | 357.22 | 29.3 | 28949 | 7 | 7 | 18217805 | 20069307 | 17556477 | 16688425 | 16813723 | 16899863 | 15286022 | 17264869 | 17103223 | 18504571 | 18155657 | 14062831 | 18422659 | 17473751 | 13093217 |
| Q9DCX2 | Atp5pd | ATP synthase subunit d, mitochondrial Mus musculus | 502.6 | 63.35 | 18749 | 9 | 9 | 27412769 | 28160661 | 28176317 | 28045433 | 23592771 | 29918719 | 22424197 | 29723903 | 26261209 | 20051059 | 24727429 | 23161897 | 22563941 | 28755925 | 20641569 |
| P97450 | Atp5pf | ATP synthase-coupling factor 6, mitochondrial Mus musculus | 111.26 | 17.59 | 12496 | 1 | 1 | 1423215.25 | 1208755.62 | 1075771.62 | 1035423.06 | 1000578.38 | 842888.81 | 597105.88 | 827233.88 | 864608.19 | 643821.56 | 591309.88 | 787930.12 | 747338 | 664091.12 | 644505.38 |
| Q9DB20 | Atp5po | ATP synthase subunit O, mitochondrial Mus musculus | 573.24 | 48.83 | 23364 | 8 | 9 | 16159396 | 16036384 | 16720265 | 16439538 | 16090287 | 16558936 | 14735394 | 14877993 | 16029947 | 12802812 | 14222713 | 17886153 | 17696207 | 16613318 | 11890727 |
| O54715 | Atp6ap1 | V-type proton ATPase subunit S1 Rattus norvegicus | 33.07 | 1.51 | 51123 | 1 | 1 | 2637324.75 | 2791397.5 | 3484690.5 | 3701668.75 | 3812607.75 | 3120753.25 | 2614943.75 | 3982357 | 2662193.25 | 3223519 | 2262195.5 | 3052354.25 | 3087130.25 | 2291379 | 2320108.75 |
| Q9Z1G4 | Atp6v0a1 | V-type proton ATPase 116 kDa subunit a isoform 1 Mus musculus | 262.69 | 9.06 | 96467 | 5 | 5 | 4060833.875 | 3957057 | 3542756 | 4797077 | 3946445.5 | 4623010 | 3999825 | 3724719.5 | 3754800.5 | 4109127.75 | 4686526 | 3677546 | 4819233 | 3650211 | 3785945 |
| P63081 | Atp6v0c | V-type proton ATPase 16 kDa proteolipid subunit Rattus norvegicus | 493.46 | 76.13 | 15808 | 5 | 8 | 33167599 | 61258881 | 39233713 | 32201433 | 44337745 | 42079037 | 52962921 | 47440393 | 53640217 | 42583365 | 42342781 | 41418193 | 60760873 | 59345001 | 38211841 |
| P51863 | Atp6v0d1 | V-type proton ATPase subunit d 1 Mus musculus | 952.93 | 54.13 | 40301 | 15 | 15 | 77478201 | 91544969 | 85966673 | 75775697 | 81929729 | 80536161 | 69743569 | 88457745 | 102318545 | 71652121 | 83118257 | 77920753 | 83770969 | 86352681 | 71237481 |
| P50516 | Atp6v1a | V-type proton ATPase catalytic subunit A Mus musculus | 2341.2 | 73.91 | 68326 | 34 | 35 | 142434705 | 139243073 | 154388161 | 126802617 | 127971937 | 133584481 | 156611505 | 133595289 | 135818369 | 139548545 | 137379185 | 140232641 | 142590881 | 135720433 | 151427889 |
| P62814 | Atp6v1b2 | V-type proton ATPase subunit B, brain isoform Mus musculus | 1684.59 | 76.13 | 56551 | 25 | 25 | 120825793 | 138188385 | 137582017 | 137719441 | 132128145 | 122381857 | 141374113 | 128505313 | 120940929 | 133262641 | 111528129 | 125736113 | 140566433 | 126654873 | 130266673 |
| Q5FVI6 | Atp6v1c1 | V-type proton ATPase subunit C 1 Rattus norvegicus | 974.91 | 43.98 | 43901 | 17 | 17 | 40574881 | 40408421 | 36351181 | 37687361 | 33457207 | 33817973 | 36484397 | 36915933 | 35789525 | 34642945 | 32005533 | 32905951 | 37586569 | 38171889 | 34558249 |
| P57746 | Atp6v1d | V-type proton ATPase subunit D Mus musculus | 183.41 | 17.41 | 28369 | 3 | 3 | 5983589 | 4213503 | 4272386.5 | 4684691 | 4494796.5 | 4503170.5 | 6257371 | 3932125 | 4081095.5 | 5282559.5 | 3925416.25 | 5521733.5 | 4262906.5 | 4104505 | 6234073 |
| P50518 | Atp6v1e1 | V-type proton ATPase subunit E 1 Mus musculus | 426.99 | 33.63 | 26157 | 6 | 6 | 28323443 | 31801433 | 27329297 | 39835665 | 36506273 | 22237769 | 34364157 | 23388009 | 14287708 | 38886657 | 34623533 | 19448889 | 28924509 | 23151577 | 30424403 |
| P50408 | Atp6v1f | V-type proton ATPase subunit F Rattus norvegicus | 255.94 | 45.38 | 13370 | 4 | 4 | 3539334.5 | 3892337.75 | 3070427.5 | 3382132.5 | 3904443 | 2945984.75 | 2783263.75 | 4118676.5 | 3652905 | 2723550.75 | 2229655 | 2588825 | 3274014.5 | 3535392.25 | 1955749.75 |
| Q9CR51 | Atp6v1g1 | V-type proton ATPase subunit G 1 Mus musculus | 204.02 | 36.44 | 13724 | 3 | 3 | 6386704.5 | 5091156 | 5424084.5 | 6782994 | 6561993 | 4975937.5 | 6630076.5 | 5241823.5 | 4710675.5 | 6084297.5 | 6598138 | 5796196.5 | 5662120.5 | 5521081 | 6025498.5 |
| Q8BVE3 | Atp6v1h | V-type proton ATPase subunit H Mus musculus | 696.59 | 32.3 | 55855 | 13 | 13 | 20321119 | 20710759 | 23600781 | 18563459 | 19201917 | 20135089 | 21577681 | 23176859 | 29077903 | 20791291 | 27810397 | 24554879 | 19549989 | 23056423 | 23805559 |
| Q64430 | Atp7a | Copper-transporting ATPase 1 Mus musculus | 158.3 | 2.88 | 161959 | 4 | 4 | 1918812.62 | 727557.19 | 1614132 | 1341063.62 | 1364779.5 | 1322772.88 | 1618093.75 | 1780776.62 | 1579250.25 | 1482090.38 | 1487820.62 | 1646720.25 | 1408533.25 | 1623504.88 | 1606581.25 |
| Q91YY4 | Atpaf2 | ATP synthase mitochondrial F1 complex assembly factor 2 Mus musculus | 40.45 | 3.81 | 33289 | 1 | 1 | 1 | 1 | 714317.5 | 1 | 1 | 989099.06 | 1 | 924172.25 | 1254300.88 | 1 | 1 | 1 | 1 | 895750.94 | 705407.12 |
| Q61687 | Atrx | Transcriptional regulator ATRX Mus musculus | 107.1 | 1.05 | 278587 | 2 | 2 | 626782.69 | 553771.25 | 402648.59 | 573780.31 | 663451.62 | 427559.62 | 419769.94 | 402060.78 | 584922.44 | 484968.34 | 541724.12 | 387320.34 | 427314.34 | 581454.69 | 393203.72 |
| P28658 | Atxn10 | Ataxin-10 Mus musculus | 760.79 | 45.47 | 53707 | 15 | 15 | 25570491 | 19160953 | 23684225 | 17806277 | 20947241 | 27277829 | 19889179 | 26847631 | 32449165 | 21204829 | 25472389 | 23532805 | 18547659 | 25689233 | 21609555 |
| Q7TQH0 | Atxn2l | Ataxin-2-like protein Mus musculus | 632.15 | 16.68 | 110649 | 13 | 13 | 9982069 | 8612443 | 12295653 | 11524145 | 10839567 | 11757244 | 14888793 | 9129724 | 10481496 | 14450275 | 12457846 | 12486555 | 12156531 | 8110220 | 13187891 |
| P70295 | Aup1 | Ancient ubiquitous protein 1 Mus musculus | 269.36 | 18.78 | 46121 | 5 | 5 | 2443452 | 1747838.5 | 2808141 | 2333366.25 | 2666960.5 | 2332861 | 2604836.25 | 2746299.75 | 2236720.5 | 2361930.25 | 1449527.75 | 2489578 | 2640177 | 2098460 | 2356172 |
| O55099 | Aurkb | Aurora kinase B Rattus norvegicus | 142.8 | 9.91 | 39234 | 2 | 3 | 291684.5 | 331943.88 | 768632.94 | 1160410.25 | 1127376.25 | 500183.75 | 425273.41 | 417932.53 | 706010.985 | 1480654.25 | 1021519.5 | 887615.94 | 372180.91 | 852758.025 | 1129715.75 |
| Q9D9K3 | Aven | Cell death regulator Aven Mus musculus | 50.07 | 6.43 | 37195 | 1 | 1 | 543086.62 | 1 | 417981.97 | 382421.25 | 1 | 1 | 1 | 1 | 501030.19 | 375564.22 | 410742.38 | 660335.12 | 1 | 401285.59 | 1 |
| P01887 | B2m | Beta-2-microglobulin Mus musculus | 47.53 | 8.4 | 13779 | 1 | 1 | 4676032.5 | 3371963 | 4980485.5 | 3501672.25 | 3830540.75 | 4607350.5 | 3469635.25 | 6130310.5 | 5188516.5 | 3055488 | 5912951.5 | 5629019 | 3219423.5 | 5698978 | 4435830.5 |

| | | | | | | | | | | | | | | | | | | | | | | |
|---|---|---|---|---|---|---|---|---|---|---|---|---|---|---|---|---|---|---|---|---|---|---|
| Q09200 | B4galnt1 | Beta-1,4 N-acetylgalactosaminyltransferase 1 Mus musculus | 165.3 | 12.95 | 59212 | 4 | 4 | 1801928 | 2613827.25 | 1044855.62 | 1320747.88 | 2539414 | 2137279.5 | 2540734 | 2353976.5 | 2691775.5 | 1763160.75 | 2816738.5 | 3623263.5 | 3182356.25 | 2389494 | 1545770.25 |
| P15535 | B4galt1 | Beta-1,4-galactosyltransferase 1 Mus musculus | 129.14 | 9.52 | 44411 | 3 | 3 | 595688.04 | 677965.44 | 605968 | 461883.34 | 636935.38 | 761654.62 | 611191.19 | 1035714.908 | 1989392.38 | 780621.44 | 1 | 655118.25 | 1 | 979601 | 1 |
| Q6P7Q1 | Babam2 | BRISC and BRCA1-A complex member 2 Rattus norvegicus | 141.76 | 9.92 | 43559 | 3 | 3 | 3014357.5 | 2838094 | 3680446.5 | 2320684 | 2456194.5 | 3400676 | 2762430 | 4285498.5 | 5212294 | 2218102.5 | 3677071 | 4304742.5 | 2226866 | 3557366 | 2316225.25 |
| Q6MG49 | Bag6 | Large proline-rich protein BAG6 Rattus norvegicus | 182.14 | 3.4 | 120012 | 3 | 3 | 5516473.5 | 5672206.5 | 6263784.5 | 4499107 | 6266172 | 6186944 | 4267088 | 6605824 | 6833656 | 4916137 | 5086526 | 5372989 | 4121672 | 6376343 | 4765759 |
| O08734 | Bak1 | Bcl-2 homologous antagonist/killer Mus musculus | 412.94 | 45.45 | 23295 | 8 | 8 | 24148415 | 20861775 | 21346057 | 19750739 | 19323359 | 21227583 | 15096963 | 20412599 | 26752557 | 19563267 | 21034099 | 17065945 | 18698693 | 20114255 | 18038281 |
| O54962 | Banf1 | Barrier-to-autointegration factor Mus musculus | 234.49 | 52.81 | 10103 | 4 | 4 | 31100997 | 37828185 | 33642749 | 36263769 | 34584713 | 36530777 | 34521289 | 35021469 | 38895429 | 30211003 | 28238129 | 32203767 | 36183381 | 34218081 | 29383213 |
| Q91XV3 | Basp1 | Brain acid soluble protein 1 Mus musculus | 926.1 | 77.43 | 22087 | 7 | 12 | 121687753 | 142960081 | 91223177 | 113236305 | 141292561 | 141975217 | 125391081 | 134325121 | 106876289 | 132395681 | 92391369 | 106971465 | 134536577 | 123852385 | 71478969 |
| Q07813 | Bax | Apoptosis regulator BAX Mus musculus | 635.21 | 63.02 | 21395 | 9 | 9 | 33075229 | 33938641 | 33393683 | 25691449 | 33127145 | 36354877 | 35049613 | 39079985 | 39443761 | 36925825 | 38119441 | 34537001 | 33858145 | 36521817 | 29488679 |
| Q9Z277 | Baz1b | Tyrosine-protein kinase BAZ1B Mus musculus | 327.78 | 6.63 | 170651 | 7 | 7 | 4002570.5 | 4994442 | 3172398.25 | 2495981.75 | 5464130 | 4203446 | 4255328 | 4187854.5 | 5199385 | 1418261.12 | 6574976 | 5294603 | 5057803 | 5065702 | 5509542.5 |
| Q9D287 | Bcas2 | Pre-mRNA-splicing factor SPF27 Mus musculus | 62.44 | 12.44 | 26131 | 1 | 1 | 1 | 1 | 164248.88 | 182089.86 | 155525.17 | 200908.58 | 1 | 165829.27 | 160975.48 | 1 | 144973.3 | 85800.32 | 136975.56 | 124844.4 | 123148.395 |
| P24288 | Bcat1 | Branched-chain-amino-acid aminotransferase, cytosolic Mus musculus | 189.22 | 9.59 | 42791 | 4 | 4 | 2924669 | 4449265.5 | 5201358 | 4306470 | 3904854 | 3750547 | 3364086.75 | 4155661.5 | 4322181 | 2906999.5 | 3494279.5 | 2456826 | 4132185.5 | 3065544 | 3984976 |
| O35855 | Bcat2 | Branched-chain-amino-acid aminotransferase, mitochondrial Mus musculus | 290.18 | 15.01 | 44127 | 5 | 5 | 23033639 | 21263059 | 21014819 | 22546623 | 23722181 | 22603343 | 22739415 | 21394163 | 22079177 | 21918281 | 20837241 | 19071875 | 21769841 | 20771775 | 19864535 |
| Q9CWI3 | Bccip | BRCA2 and CDKN1A-interacting protein Mus musculus | 151.58 | 15.19 | 35942 | 4 | 4 | 3967588 | 5860369 | 6255044.5 | 5587338.5 | 4560883.5 | 5505663 | 3834412.5 | 5498532.5 | 6799317 | 4155189 | 3906373 | 5534458 | 4111400 | 5281394 | 3910314.25 |
| P35738 | Bckdhb | 2-oxoisovalerate dehydrogenase subunit beta, mitochondrial Rattus norvegicus | 134.18 | 7.44 | 42823 | 2 | 2 | 957250.94 | 2317624.75 | 2008439.62 | 491190.06 | 352572.53 | 1812267.25 | 2123203 | 1667250.12 | 1904602.88 | 1406450.88 | 1661021.12 | 1410360.62 | 1949228.62 | 2046872.88 | 1926542.5 |
| O55028 | Bckdk | [3-methyl-2-oxobutanoate dehydrogenase [lipoamide]] kinase, mitochondrial Mus muscu | 123.32 | 11.41 | 46588 | 2 | 2 | 5335562 | 6367086 | 8666488 | 6777259 | 5026245 | 9499944 | 7600793.5 | 8711683 | 9167563 | 7695803.5 | 8097398.5 | 8480575 | 8272109 | 6295986 | 9204400 |
| Q9Z0H7 | Bcl10 | B-cell lymphoma/leukemia 10 Mus musculus | 119.69 | 22.75 | 25948 | 3 | 3 | 2468201.5 | 1522003.5 | 1450064.25 | 1656171.5 | 1788120 | 1274587.12 | 1576629.12 | 1311772.25 | 1476136.5 | 1486079.5 | 1922012.25 | 1557864.5 | 1633115.12 | 1556468.12 | 2139605 |
| P10417 | Bcl2 | Apoptosis regulator Bcl-2 Mus musculus | 37.3 | 4.24 | 26407 | 1 | 1 | 593319.06 | 778310.62 | 883003.56 | 1 | 1 | 579923.44 | 651006.12 | 1092372.38 | 1284822.25 | 491821.09 | 471439.19 | 820939.69 | 380369.16 | 1014814.5 | 785195.38 |
| P53563 | Bcl2l1 | Bcl-2-like protein 1 Rattus norvegicus | 116.49 | 8.15 | 26158 | 2 | 2 | 5620402.5 | 6564159 | 6171908.5 | 5036935 | 5291808 | 5153788.5 | 4285732 | 5906592 | 6587995 | 5672252 | 5151579 | 5708094 | 5730709.5 | 6096719 | 4708042.5 |
| P59017 | Bcl2l13 | Bcl-2-like protein 13 Mus musculus | 266.18 | 20.97 | 46719 | 5 | 5 | 5133233 | 7492595.5 | 7480295.5 | 8898548 | 7648284 | 6646305 | 8527362 | 10156359 | 9639056 | 10363053 | 9757421 | 9335088 | 10523231 | 9261940 | 10132683 |
| Q8K019 | Bclaf1 | Bcl-2-associated transcription factor 1 Mus musculus | 262.86 | 7.73 | 106002 | 6 | 6 | 5567837.5 | 5830390 | 5469708.5 | 6010429 | 6357267 | 6366613 | 7911666.5 | 5064015 | 5185794 | 6202368 | 7020352 | 5961194.5 | 6438839.5 | 4763376.5 | 5758986.5 |
| Q9CZP5 | Bcs1l | Mitochondrial chaperone BCS1 Mus musculus | 72.75 | 4.55 | 47406 | 1 | 1 | 279154.44 | 219089.17 | 206571.38 | 316904.25 | 264576.97 | 266265.97 | 281046.59 | 318335.81 | 259052.16 | 360377.41 | 247102.77 | 300244.59 | 265862.16 | 340096.34 | 298588.69 |
| P70444 | Bid | BH3-interacting domain death agonist Mus musculus | 68.36 | 9.74 | 21952 | 1 | 1 | 463153.19 | 284290.16 | 331945.22 | 315233.22 | 250286.28 | 366953.31 | 234012.69 | 312146.16 | 405817.81 | 350804.91 | 1 | 365787.16 | 312786.78 | 385747.09 | 1 |
| O08539 | Bin1 | Myc box-dependent-interacting protein 1 Mus musculus | 172.38 | 8.33 | 64470 | 2 | 3 | 1466632.25 | 1737278.88 | 1662260.75 | 1233827.25 | 1523102 | 1642246.25 | 1849649.38 | 1543089.62 | 1540212 | 1217612 | 1560822.38 | 1771671.75 | 1626294.5 | 1407271.5 | 1601768.5 |
| D3Z6Q9 | Bin2 | Bridging integrator 2 Mus musculus | 608.75 | 23.11 | 52554 | 8 | 9 | 13699195 | 11867651 | 12105952 | 13864309 | 13672661 | 11204565 | 11586169 | 10732576 | 10461511 | 11952553 | 12138047 | 10699399 | 11307664 | 9192905 | 11981107 |
| O88738 | Birc6 | Baculoviral IAP repeat-containing protein 6 Mus musculus | 831.91 | 5.28 | 532170 | 17 | 17 | 8297570.5 | 7470425 | 10081143 | 8263433.5 | 8246352 | 8090700 | 8647933 | 9016095 | 9602021 | 8134092.5 | 9173411 | 9379356 | 7106891.5 | 8693463 | 9262435 |
| Q8R016 | Blmh | Bleomycin hydrolase Mus musculus | 606.62 | 38.68 | 52511 | 2 | 12 | 17496483 | 15199557 | 15461817 | 15561043 | 17376735 | 17954097 | 15439263 | 12276567 | 14592746 | 10819339 | 19144445 | 13519559 | 14982298 | 16975097 | 17681341 |
| Q32WR5 | Bloc1s2 | Biogenesis of lysosome-related organelles complex-1 subunit 2 Rattus norvegicus | 36.14 | 7.75 | 16067 | 1 | 1 | 363234.62 | 1367884.5 | 340128.53 | 1090036.25 | 1005862.62 | 947351.19 | 748227.12 | 945434.31 | 1084635.88 | 1026057.62 | 518647.62 | 902954.56 | 1091311.25 | 827445.88 | 615047.12 |
| Q9CY64 | Blvra | Biliverdin reductase A Mus musculus | 753.23 | 58.64 | 33525 | 14 | 14 | 48784881 | 54583561 | 49228573 | 50434457 | 51295057 | 43970165 | 42104349 | 46433065 | 47243597 | 43225813 | 43782461 | 44680997 | 49135057 | 50856097 | 44507869 |
| Q923D2 | Blvrb | Flavin reductase (NADPH) Mus musculus | 516.99 | 60.19 | 22197 | 7 | 7 | 5689220 | 4773408 | 5004091 | 6257829 | 6131784 | 6852737 | 10259370 | 6199030 | 4261356.5 | 6354247.5 | 6197547.5 | 5678764.5 | 7560742.5 | 6740745 | 9289422 |
| Q8BXK4 | Bmt2 | S-adenosylmethionine sensor upstream of mTORC1 Mus musculus | 54.97 | 6.2 | 45952 | 1 | 1 | 148636.78 | 1 | 131433.78 | 228973.48 | 1 | 148685.02 | 140021.59 | 169349.61 | 178010.19 | 220318.52 | 242110.44 | 1 | 1 | 211277.23 | 1 |
| Q9D8S9 | Bola1 | BolA-like protein 1 Mus musculus | 85.03 | 21.17 | 14379 | 2 | 2 | 1910952.5 | 2213255 | 3035279 | 2462605.75 | 2425786.25 | 2413228 | 2388363.75 | 2449249.5 | 2299519.75 | 3104795 | 2458966.5 | 3019076.75 | 2491676.25 | 1992886.75 | 2140885.5 |
| P97452 | Bop1 | Ribosome biogenesis protein BOP1 Mus musculus | 260.95 | 10.79 | 82546 | 6 | 6 | 3946396.25 | 3694792.25 | 3855895 | 5074552 | 4697104 | 5503907.5 | 3801408.25 | 3293194 | 5023749 | 4343331 | 3391596 | 5558527 | 3182980 | 3025073.5 | 5881608 |
| Q8R164 | Bphl | Valacyclovir hydrolase Mus musculus | 116.8 | 11.68 | 32851 | 2 | 2 | 1100669.62 | 1138351.88 | 1481812.12 | 1607256.25 | 1239352.62 | 1250229.38 | 1505743.5 | 1886589.88 | 1692261.62 | 1054427 | 1442351.5 | 1258219.38 | 1410385.88 | 1406981.62 | 1180202 |
| Q9Z0S1 | Bpnt1 | 3'(2'),5'-bisphosphate nucleotidase 1 Mus musculus | 546.68 | 58.12 | 33196 | 5 | 12 | 13379334 | 13102381 | 11183731 | 12615153 | 13661309 | 12665043 | 13770296 | 12216773 | 13289735 | 13422182 | 12665637 | 11492543 | 12808331 | 13672009 | 11146133 |
| Q9ESU6 | Brd4 | Bromodomain-containing protein 4 Mus musculus | 43.71 | 1.43 | 155895 | 1 | 1 | 1527573 | 2074683.5 | 1545319 | 1 | 1 | 1 | 2001574.75 | 2492448.5 | 1669212.12 | 1 | 1691999.62 | 1803715.75 | 2112018.5 | 523180.69 | 1423174.88 |
| O88665 | Brd7 | Bromodomain-containing protein 7 Mus musculus | 65.44 | 4.15 | 74000 | 1 | 1 | 339867.565 | 310889.31 | 243446.92 | 365756.91 | 439377.12 | 322164.38 | 358658.84 | 342707.25 | 254828.58 | 338806.75 | 275992.03 | 1 | 274329.47 | 1 | 405631.97 |
| Q8BXV2 | Bri3bp | BRI3-binding protein Mus musculus | 109.82 | 7.51 | 28263 | 1 | 1 | 2298517.25 | 1 | 1171718.75 | 1 | 1322235.62 | 1042225.5 | 1107042.62 | 1194544 | 1399156 | 988520.06 | 1572364.12 | 1432993.38 | 1310059.5 | 968954.5 | 1108839.88 |
| Q4QQT6 | Brix1 | Ribosome biogenesis protein BRX1 homolog Rattus norvegicus | 271.25 | 17.9 | 41115 | 5 | 5 | 5540502 | 3599404.5 | 4631790.5 | 4308019 | 3612766.5 | 4653097 | 4167821.5 | 4393741 | 4449852.5 | 4142765.5 | 4360702 | 4590161 | 3700418.75 | 3943062.25 | 4256531.5 |
| Q91VR8 | Brk1 | Protein BRICK1 Mus musculus | 81.48 | 24 | 8761 | 2 | 2 | 1986134 | 1097114.5 | 2865366.5 | 1 | 1 | 1714889.75 | 791778.06 | 2712222.75 | 3832234 | 2262781.14 | 2080473.25 | 2623655.25 | 648225.88 | 3323578.25 | 1604953.38 |
| Q8K2Q7 | Brox | BRO1 domain-containing protein BROX Mus musculus | 362.44 | 25.79 | 46202 | 7 | 7 | 8301968 | 8458051 | 6718905 | 6004193.5 | 7874387.5 | 6876888.5 | 4914679.5 | 6022614 | 8239421.5 | 6559057 | 6015872 | 6593707 | 7464560 | 7463138.5 | 5467633 |
| Q80Y55 | Bsdc1 | BSD domain-containing protein 1 Mus musculus | 97.89 | 7.96 | 46953 | 2 | 2 | 926742.06 | 675796.88 | 724778.12 | 788749.06 | 798811.62 | 869940.19 | 699637.31 | 1651080.62 | 1511218.75 | 1583081.25 | 932425.4375 | 1492542.12 | 803018.94 | 913933.88 | 520206.81 |
| P18572 | Bsg | Basigin Mus musculus | 39.32 | 2.57 | 42445 | 1 | 1 | 1494309.62 | 1713495.38 | 1650802.25 | 1894312.5 | 1921786.12 | 2357860.75 | 2189130 | 1807753.75 | 1848912.75 | 1919691 | 1853990.88 | 2140935.25 | 2016074.25 | 1541446.88 | 1703023 |
| Q8R2Q8 | Bst2 | Bone marrow stromal antigen 2 Mus musculus | 110.88 | 11.63 | 19152 | 2 | 2 | 3549342 | 3271166.5 | 2628263 | 2198618.5 | 3423167 | 2915756 | 2751605 | 2980969.5 | 2292952 | 1142204.88 | 2646589.5 | 2503504.5 | 3338661.25 | 3167388 | 1145713.75 |
| Q64152 | Btf3 | Transcription factor BTF3 Mus musculus | 345.33 | 49.51 | 22031 | 5 | 6 | 8559690 | 3963968 | 4071065.75 | 5078866.5 | 3670445.75 | 3959851.75 | 4180962.25 | 4518402.5 | 3192492.25 | 4502075.5 | 2498636 | 3957996.75 | 4239217.5 | 3883990.25 | 3558071.75 |
| P35991 | Btk | Tyrosine-protein kinase BTK Mus musculus | 360.86 | 14.42 | 76437 | 7 | 7 | 2571734 | 1760990.5 | 4117530.5 | 2402241.5 | 2565336 | 4168168.25 | 5168041 | 4172703.5 | 1475512.62 | 1841801.75 | 2250203 | 1819691.5 | 2420179 | 2086457.38 | 2004940.5 |
| Q9WVA3 | Bub3 | Mitotic checkpoint protein BUB3 Mus musculus | 438.52 | 28.83 | 36955 | 7 | 7 | 16847627 | 19373095 | 17989345 | 18040573 | 16922063 | 15482078 | 17457343 | 15816368 | 16721081 | 16299055 | 17017861 | 17207475 | 16927135 | 14871439 | 19596597 |
| Q9CY21 | Bud23 | Probable 18S rRNA (guanine-N(7))-methyltransferase Mus musculus | 139.18 | 18.51 | 31587 | 3 | 3 | 975878.31 | 1687088.25 | 1467269 | 2788204 | 1119058.38 | 2372433 | 1856668.62 | 1247467 | 1960862.25 | 2879951.5 | 2104962.5 | 2446815.5 | 2524105.75 | 2317561.25 | 2547542 |
| O70454 | Bud31 | Protein BUD31 homolog Rattus norvegicus | 142.59 | 26.39 | 17000 | 3 | 3 | 3593911.5 | 3148453.5 | 2838716.5 | 1610258 | 2934452.25 | 1766646.75 | 2731381 | 2959914 | 3436079 | 3284922.75 | 2870740.75 | 2727642 | 1678955.75 | 3306195.5 | 2661695.5 |
| O54825 | Bysl | Bystin Mus musculus | 320.47 | 23.39 | 49784 | 7 | 7 | 6040633 | 3295103.25 | 2628441 | 3875066 | 4431506 | 3538979 | 3068835 | 3359036.25 | 4142788.5 | 4305323.5 | 1947090.12 | 2439854.75 | 2276910.75 | 3299198 | 4656492.5 |
| Q6P7P5 | Bzw1 | Basic leucine zipper and W2 domain-containing protein 1 Rattus norvegicus | 1012.89 | 61.34 | 48043 | 17 | 19 | 33573389 | 24747669 | 28850221 | 19980147 | 21328787 | 27725795 | 24143761 | 31217277 | 35622505 | 21127741 | 29968815 | 30689653 | 22027525 | 33085307 | 27758795 |
| Q2L4X1 | Bzw2 | Basic leucine zipper and W2 domain-containing protein 2 Mus musculus molossinus (J | 388.29 | 30.07 | 48063 | 7 | 9 | 7297814 | 4181495.5 | 6675814 | 3399781 | 4155829.5 | 5645589.5 | 5142063 | 7112829.5 | 7489221.5 | 4309045.5 | 5984271 | 6519428.5 | 4624041 | 6294790 | 5255935 |
| P98086 | C1qa | Complement C1q subcomponent subunit A Mus musculus | 50.02 | 6.12 | 25974 | 1 | 1 | 476973.56 | 391320.94 | 672807 | 879280.31 | 652380.44 | 298784.91 | 198500.78 | 298704.5 | 1 | 1 | 521637.09 | 649038 | 600793.12 | 348516.56 | 827070.19 |
| O35658 | C1qbp | Complement component 1 Q subcomponent-binding protein, mitochondrial Mus muscul | 427.26 | 43.53 | 31013 | 1 | 7 | 9895254 | 7132176.5 | 11802255 | 8187858 | 7418910.5 | 10760771 | 10335239 | 14878015 | 14816353 | 4759105.5 | 15472397 | 13583487 | 5098283 | 13144663 | 12258319 |
| Q80X80 | C2cd2l | Phospholipid transfer protein C2CD2L Mus musculus | 49.59 | 1.42 | 76329 | 1 | 1 | 1 | 1 | 349341.56 | 1 | 1 | 567739.62 | 1 | 1 | 645406.5 | 486975.19 | 290710.75 | 411528.545 | 483127.28 | 428528.12 | 443748.03 |
| P30993 | C5ar1 | C5a anaphylatoxin chemotactic receptor 1 Mus musculus | 90.77 | 6.84 | 39023 | 2 | 2 | 3853467 | 2171287.75 | 1685147 | 1996552.75 | 1890232 | 1317881.25 | 597943.56 | 1553283.12 | 1622435.75 | 1719334.75 | 2097651.25 | 1523115.88 | 1648563 | 2039875.88 | 1795117.25 |
| Q9D6N1 | Ca13 | Carbonic anhydrase 13 Mus musculus | 57.94 | 3.44 | 29522 | 1 | 1 | 2512639.25 | 3071649 | 2929762.75 | 2469199.5 | 2384382.75 | 2649599.25 | 2623670.5 | 3037292.5 | 3498025 | 2036705.75 | 2497723.595 | 2346112.5 | 2405546.25 | 3213676.75 | 2025558.88 |
| P00920 | Ca2 | Carbonic anhydrase 2 Mus musculus | 72.87 | 6.15 | 29033 | 1 | 1 | 995448.44 | 967557.69 | 745657 | 1006444.56 | 713991.19 | 903311.88 | 943092.75 | 1015276.12 | 916517.88 | 1041080.44 | 811227.62 | 964170.19 | 1033448.94 | 1170009 | 976034.62 |
| Q06138 | Cab39 | Calcium-binding protein 39 Mus musculus | 538.05 | 40.76 | 39843 | 11 | 12 | 19173705 | 12546711 | 12553784 | 12871691 | 8612880 | 17420629 | 13684938 | 13765623 | 16163784 | 16468150 | 15854407 | 13466406 | 11545555 | 14923713 | 17455559 |
| Q9DB16 | Cab39l | Calcium-binding protein 39-like Mus musculus | 83.15 | 5.34 | 39106 | 1 | 2 | 6080585.5 | 1 | 3140046.25 | 3207864.25 | 1 | 4062455.25 | 3988101.75 | 3676119.75 | 3664192.25 | 3518566.25 | 3525090.5 | 3365404.5 | 2988237 | 3013240.5 | 4534050 |
| Q9D424 | Cabyr | Calcium-binding tyrosine phosphorylation-regulated protein Mus musculus | 35.11 | 1.55 | 48333 | 1 | 1 | 3739951.5 | 4223158.5 | 3593227.25 | 3195507.25 | 3261948.75 | 3745839.5 | 3693080.25 | 3536663.25 | 4642472.5 | 3427979.5 | 3578298.75 | 3268752 | 3737386.25 | 3644280.5 | 3202986 |
| Q9CXW3 | Cacybp | Calcyclin-binding protein Mus musculus | 666.09 | 55.46 | 26510 | 11 | 11 | 22912845 | 22584781 | 26027987 | 19970155 | 21790039 | 25564707 | 31313195 | 22161011 | 21552883 | 23828985 | 25206621 | 24530917 | 24759933 | 21750691 | 25116251 |
| B2RQC6 | Cad | CAD protein Mus musculus | 1956.76 | 23.28 | 243238 | 37 | 37 | 42515117 | 45114417 | 50362805 | 46463789 | 44529117 | 45534573 | 38687993 | 36884033 | 44202581 | 38371297 | 45592093 | 38869269 | 38199497 | 44217401 | 45097709 |
| Q6AYP5 | Cadm1 | Cell adhesion molecule 1 Rattus norvegicus | 137 | 7.77 | 51854 | 2 | 2 | 2619958 | 3106014.75 | 2897103 | 2986280 | 2788312.25 | 2495120.75 | 2669904.25 | 3329516.25 | 3177209.75 | 2340254.25 | 3143258 | 3083361.5 | 3048868.75 | 2751343.75 | 2577975 |
| Q8VEC4 | Calhm2 | Calcium homeostasis modulator protein 2 Mus musculus 2 | 162.2 | 12.69 | 35823 | 3 | 3 | 1478173.12 | 1066539 | 1002386.44 | 524569.94 | 550972.19 | 570792.44 | 583806.88 | 1219253.38 | 1122452 | 565563.38 | 813863.62 | 1144653 | 1081233.5 | 628991.12 | 1068434 |
| P0DP26 | Calm1 | Calmodulin-1 Mus musculus | 661.71 | 80.54 | 16838 | 10 | 10 | 72509633 | 55863917 | 78980857 | 56340881 | 50930209 | 50189925 | 34234653 | 90442793 | 73208569 | 32686221 | 72122585 | 78409361 | 41185929 | 85293665 | 57948505 |
| P14211 | Calr | Calreticulin Mus musculus | 1301.77 | 59.62 | 47995 | 20 | 20 | 245965841 | 256566625 | 267784321 | 231734545 | 298205889 | 295970689 | 298608225 | 275262529 | 328718593 | 287326497 | 348832897 | 298924193 | 276418017 | 264746129 | 322402145 |
| O35887 | Calu | Calumenin Mus musculus | 548.33 | 47.94 | 37064 | 6 | 9 | 10071065 | 11226470 | 13939179 | 8559093 | 8033591 | 12951333 | 10628127 | 16121594 | 17144035 | 9203413 | 13093165 | 13318388 | 8649179 | 17121541 | 10359545 |
| Q63450 | Camk1 | Calcium/calmodulin-dependent protein kinase type 1 Rattus norvegicus | 188.81 | 10.16 | 41638 | 3 | 3 | 7956965 | 7125809 | 9297724 | 6216543 | 6378720.5 | 7394079 | 5263892.5 | 8434912 | 10842397 | 5439172 | 8339173 | 8647926 | 5542149 | 8159801 | 7460712 |
| Q6PHZ2 | Camk2d | Calcium/calmodulin-dependent protein kinase type II subunit delta Mus musculus | 490.44 | 26.45 | 56369 | 9 | 9 | 15552478 | 16226495 | 16876161 | 17452865 | 16448511 | 18309389 | 19223837 | 19063947 | 17683485 | 18653835 | 17858675 | 18811947 | 17461829 | 17797345 | 16200679 |
| P97536 | Cand1 | Cullin-associated NEDD8-dissociated protein 1 Rattus norvegicus | 1278.81 | 27.56 | 136362 | 24 | 24 | 60754897 | 57666085 | 56663609 | 50113409 | 60987517 | 59684573 | 51267197 | 60649361 | 66314077 | 52410937 | 64982537 | 52660245 | 60776229 | 63438629 | 58893969 |
| P35564 | Canx | Calnexin Mus musculus | 1226.02 | 49.58 | 67278 | 22 | 22 | 75944481 | 74517137 | 79112257 | 75547153 | 73482825 | 85744721 | 80736617 | 88423929 | 85699313 | 77040265 | 76696785 | 74075913 | 74686977 | 68860737 | 71764617 |
| P40124 | Cap1 | Adenylyl cyclase-associated protein 1 Mus musculus | 2231.67 | 70.25 | 51565 | 33 | 35 | 294169825 | 366957793 | 344019201 | 316606305 | 327385057 | 310588033 | 356050465 | 299629889 | 308219809 | 345013441 | 421149665 | 353205793 | 323675745 | 342178209 | 344971105 |
| P24452 | Capg | Macrophage-capping protein Mus musculus | 892.52 | 42.05 | 39240 | 1 | 13 | 390912129 | 435303329 | 312020865 | 343049249 | 445382593 | 260103345 | 380038849 | 477285473 | 349605569 | 532456417 | 271012065 | 313269601 | 529188673 | 464419905 | 505034913 |
| O35350 | Capn1 | Calpain-1 catalytic subunit Mus musculus | 559.77 | 18.65 | 82106 | 11 | 11 | 11923398 | 14869476 | 14141708 | 9304230 | 10484122 | 13192513 | 9547905 | 14753802 | 15585427 | 7468395.5 | 14162557 | 13375793 | 9137855 | 15645729 | 11490787 |
| O08529 | Capn2 | Calpain-2 catalytic subunit Mus musculus | 1130.69 | 34 | 79872 | 20 | 20 | 52366493 | 53023433 | 44892601 | 43010409 | 48304345 | 46590413 | 32719333 | 47120221 | 51938553 | 36381841 | 48413517 | 41974169 | 44859629 | 54005689 | 42972089 |
| O88456 | Capns1 | Calpain small subunit 1 Mus musculus | 587.5 | 59.48 | 28463 | 3 | 9 | 30998365 | 34028041 | 29501311 | 25698957 | 30787103 | 29111647 | 25618295 | 34026945 | 36323853 | 25965585 | 31356803 | 31513435 | 33781949 | 36024481 | 30014119 |
| Q60865 | Caprin1 | Caprin-1 Mus musculus | 867.1 | 27.72 | 78169 | 15 | 15 | 24154185 | 14004299 | 20487127 | 14336851 | 13727249 | 14572649 | 17441801 | 22206715 | 23198325 | 14826765 | 24641229 | 25355385 | 14433168 | 21133613 | 21886347 |
| P47753 | Capza1 | F-actin-capping protein subunit alpha-1 Mus musculus | 528 | 45.1 | 32940 | 8 | 10 | 59613385 | 60653597 | 55100237 | 56731757 | 63947313 | 62831041 | 51029169 | 63436597 | 65223509 | 52240865 | 53708869 | 54743813 | 53087337 | 66336829 | 50539093 |
| P47754 | Capza2 | F-actin-capping protein subunit alpha-2 Mus musculus | 760.1 | 64.34 | 32967 | 12 | 14 | 74794169 | 70675769 | 65121949 | 68851193 | 73661457 | 70041817 | 61051357 | 71039505 | 75849177 | 63154005 | 66933741 | 63085953 | 66332569 | 75219657 | 63689569 |
| Q5XI32 | Capzb | F-actin-capping protein subunit beta Rattus norvegicus | 845.61 | 65.44 | 30629 | 17 | 17 | 66935225 | 84005313 | 81562849 | 80085769 | 77732337 | 74701937 | 87304465 | 75726729 | 74844001 | 90024209 | 87995113 | 78690265 | 84879505 | 73655241 | 81258329 |
| Q9D1I2 | Card19 | Caspase recruitment domain-containing protein 19 Mus musculus | 46.92 | 4.37 | 20936 | 1 | 1 | 1020654.75 | 886030.19 | 868942.94 | 1367625.75 | 960020.12 | 942037.62 | 1238984.38 | 1025950.25 | 1364863.38 | 1132064.5 | 954374.75 | 985588 | 946033.56 | 1130156.5 | 1004038.203 |
| A2AIV8 | Card9 | Caspase recruitment domain-containing protein 9 Mus musculus | 126.62 | 3.54 | 62462 | 2 | 2 | 722139.81 | 972047 | 650519.81 | 1374586.12 | 546785.94 | 936601.5 | 578629.94 | 521143.38 | 560857.75 | 619926.56 | 363362.03 | 695627.5 | 1024411 | 487891.72 | 1675815.38 |
| Q4AE70 | Carm1 | Histone-arginine methyltransferase CARM1 Rattus norvegicus | 134.61 | 7.53 | 70341 | 3 | 3 | 2199049.75 | 2187363.25 | 2090592 | 1574659.12 | 914684.31 | 2110373.25 | 1596006.75 | 2311830.5 | 2150537.5 | 1919616.25 | 2048898.88 | 1629998.62 | 1999617.62 | 2399885 | 1956064 |
| Q9ER72 | Cars | Cysteine--tRNA ligase, cytoplasmic Mus musculus | 955.15 | 23.83 | 94860 | 17 | 17 | 18081165 | 15996308 | 18579007 | 17411201 | 12776451 | 17954483 | 21522679 | 17874929 | 20080811 | 20891403 | 23135531 | 21884095 | 16534584 | 18105631 | 23338735 |
| P29452 | Casp1 | Caspase-1 Mus musculus | 890.3 | 41.29 | 45640 | 16 | 16 | 26517867 | 27996515 | 28144805 | 26615815 | 29227735 | 23771153 | 22518755 | 23031809 | 24661715 | 23420683 | 30273639 | 24538399 | 27380405 | 26345641 | 18624521 |
| P29594 | Casp2 | Caspase-2 Mus musculus | 45.19 | 1.99 | 50661 | 1 | 1 | 778569.94 | 877849.38 | 870675.62 | 927981.62 | 863769.14 | 1 | 1 | 1 | 1 | 1 | 1 | 1 | 798172.12 | 831248.62 | 1 |
| P70677 | Casp3 | Caspase-3 Mus musculus | 315.97 | 29.6 | 31475 | 6 | 6 | 5906910.5 | 9678122 | 8811055 | 7282491 | 7355221.5 | 7349961 | 6514181 | 6465164 | 10788589 | 7791142 | 7677589 | 8035325 | 7426661.5 | 7913837.5 | 6537415 |
| O08738 | Casp6 | Caspase-6 Mus musculus | 100.48 | 6.88 | 31595 | 2 | 2 | 993009.38 | 1089150 | 1270771.62 | 714426.56 | 856172.25 | 847482.44 | 610609 | 1083207.38 | 1519684.38 | 725428.88 | 889403.12 | 1184241.88 | 728637.5 | 977670.5 | 820173.12 |
| P97864 | Casp7 | Caspase-7 Mus musculus | 38.54 | 3.3 | 34061 | 1 | 1 | 1 | 1 | 760315.75 | 1 | 640585.56 | 527867.38 | 641343.44 | 1 | 1 | 519957.53 | 1 | 1 | 487802.44 | 1 | 603302 |
| O89110 | Casp8 | Caspase-8 Mus musculus | 381.44 | 21.25 | 55357 | 8 | 9 | 8308729 | 14283832 | 12380746 | 12562698 | 13577395 | 11753753 | 11663902 | 12142726 | 14438283 | 13382517 | 13182705 | 11391081 | 12106071 | 13016689 | 12490636 |
| Q8C3Q9 | Casp9 | Caspase-9 Mus musculus | 182.09 | 11.67 | 49979 | 3 | 3 | 553626.38 | 1743199.5 | 139710.88 | 1245217.12 | 232351.16 | 1059394.62 | 790243.94 | 1647220.12 | 1118817.12 | 1215646.5 | 1499264.25 | 1104650.25 | 160771.83 | 1111471.5 | 1042863 |
| P51125 | Cast | Calpastatin Mus musculus | 360.36 | 15.36 | 84922 | 6 | 6 | 3787703.5 | 7911193.5 | 6578158 | 8220052.5 | 8366228 | 7441052.5 | 6966715 | 8512169 | 8783079 | 9111046 | 7183649 | 7985759 | 8728322 | 7720816.5 | 6249779.5 |
| P24270 | Cat | Catalase Mus musculus | 326.05 | 19.54 | 59795 | 6 | 6 | 4729484 | 5277336 | 7071070 | 5557493 | 5415771 | 6422386 | 8746106 | 5541012 | 5957669 | 7354645 | 6035146.5 | 5590136.5 | 5082671 | 5331161 | 5469019 |
| Q08024 | Cbfb | Core-binding factor subunit beta Mus musculus | 275.05 | 29.41 | 22030 | 3 | 3 | 4640339 | 6989548 | 4756424 | 4919753.5 | 5366629.5 | 4971501 | 4755779 | 6134031.5 | 5781500 | 4367793 | 5164268 | 4771800.5 | 3954016 | 5719144.5 | 4927406 |
| P22682 | Cbl | E3 ubiquitin-protein ligase CBL Mus musculus | 166.92 | 6.24 | 100564 | 4 | 4 | 3931353.5 | 4262439 | 887598.12 | 3733659 | 2774012 | 3124947.5 | 1468757 | 3207611 | 1675033.12 | 2343453 | 3444746 | 706312.5 | 3253701.25 | 2221325.75 | 2400949.25 |
| Q7TS56 | Cbr4 | Carbonyl reductase family member 4 Rattus norvegicus | 52.99 | 8.05 | 25287 | 1 | 1 | 277254.19 | 499812.47 | 559444.06 | 939096.25 | 859095.06 | 876849.75 | 971040.38 | 718326.69 | 774877.06 | 730320.75 | 608718 | 629002.25 | 786712.38 | 629668.12 | 785542.62 |
| Q8VEH6 | Cbwd1 | COBW domain-containing protein 1 Mus musculus | 132.73 | 9.16 | 43772 | 2 | 2 | 1518167.38 | 361844.88 | 1746521.25 | 1475068.88 | 1537275.12 | 2163748.25 | 2115234.75 | 2072216 | 1924196.12 | 1723247.12 | 1667754.62 | 1989867.75 | 1655800.62 | 2028631.25 | 1834892.62 |
| P83917 | Cbx1 | Chromobox protein homolog 1 Mus musculus | 368.92 | 45.95 | 21418 | 4 | 5 | 12186138 | 15126752 | 13201249 | 14069240 | 18252925 | 15903009 | 18624297 | 14372421 | 14558727 | 15781975 | 18141863 | 17037069 | 17883601 | 14945407 | 18166131 |
| P23198 | Cbx3 | Chromobox protein homolog 3 Mus musculus | 440.56 | 36.61 | 20855 | 5 | 7 | 36266237 | 42719905 | 35782081 | 40401865 | 45056341 | 41024453 | 44086485 | 41533629 | 40831349 | 45330669 | 46874357 | 45421365 | 45622977 | 40619385 | 46081829 |
| Q61686 | Cbx5 | Chromobox protein homolog 5 Mus musculus | 318.98 | 34.55 | 22186 | 5 | 6 | 11663887 | 14353037 | 11469198 | 11127766 | 14267771 | 13880597 | 12221624 | 10113260 | 14519076 | 11520891 | 14552556 | 11355186 | 12382409 | 12467597 | 13891751 |
| Q8K1A6 | Cc2d1a | Coiled-coil and C2 domain-containing protein 1A Mus musculus | 60.43 | 2.12 | 103698 | 1 | 1 | 1 | 1 | 1 | 769127.75 | 1 | 1 | 1 | 1 | 1 | 1 | 1 | 420600.22 | 1 | 1 | 461096.28 |
| Q8BRN9 | Cc2d1b | Coiled-coil and C2 domain-containing protein 1B Mus musculus | 177.26 | 11.32 | 93091 | 4 | 4 | 554515.06 | 907815.56 | 706672 | 1 | 740409.44 | 390996.19 | 607596.75 | 511895.12 | 1474714.5 | 661852.56 | 584963.44 | 624676 | 1494025.62 | 1 | 1 |
| Q8VDP4 | Ccar2 | Cell cycle and apoptosis regulator protein 2 Mus musculus | 257.02 | 8.68 | 103002 | 6 | 6 | 3319470 | 11913621 | 11904172 | 11383513 | 14229631 | 3710310.5 | 4622790 | 12251844 | 11912807 | 6387558 | 5169592 | 10174696 | 6254702.5 | 6665706 | 8075860 |
| Q8VE99 | Ccdc115 | Coiled-coil domain-containing protein 115 Mus musculus | 153.41 | 28.33 | 19743 | 3 | 3 | 887209.81 | 1163544.75 | 1355171.88 | 128817.3 | 939700.88 | 1088262.5 | 1787985.88 | 1155782.25 | 1334884.313 | 1307506.62 | 1362519.12 | 1273140.62 | 1229142 | 668144.88 | 1349445.38 |
| Q8R344 | Ccdc12 | Coiled-coil domain-containing protein 12 Mus musculus | 61.88 | 13.86 | 18891 | 1 | 1 | 1 | 1 | 1042977.25 | 877351.94 | 973050.62 | 1019331.75 | 418509.22 | 1 | 764587.25 | 1 | 697000.19 | 928446.75 | 989338.56 | 1041891.38 | 366025.12 |

| | | | | | | | | | | | | | | | | | | | | | | |
|---|---|---|---|---|---|---|---|---|---|---|---|---|---|---|---|---|---|---|---|---|---|---|
| Q9JIG7 | Ccdc22 | Coiled-coil domain-containing protein 22 Mus musculus | 209.62 | 13.24 | 70844 | 5 | 5 | 1427041.5 | 1870454.5 | 1686675.5 | 1998219.88 | 1917824 | 1568226.5 | 1573200.75 | 1411708.5 | 1591877.12 | 1737734.5 | 1683565.25 | 1580312.75 | 1699545.5 | 1375035.88 | 1443956.12 |
| Q9CR29 | Ccdc43 | Coiled-coil domain-containing protein 43 Mus musculus | 166.47 | 21.62 | 25049 | 3 | 3 | 2385759.25 | 2739219.5 | 2618146 | 2775880.25 | 2504314.75 | 2400350 | 2280018.5 | 2254367.5 | 2326717 | 3088481.5 | 2336251.5 | 2514121.5 | 2095371.75 | 2703854 | 2310800.25 |
| Q5U2X6 | Ccdc47 | Coiled-coil domain-containing protein 47 Rattus norvegicus 2 | 236.1 | 11.18 | 55733 | 5 | 5 | 5901036 | 6423451 | 7078342 | 6889855 | 2853142 | 4221791.5 | 5098454 | 7355603.5 | 5905429 | 5167719 | 3336585.5 | 4484224.5 | 5540563.5 | 5261082 | 4693778 |
| Q3ZAV0 | Ccdc60 | Coiled-coil domain-containing protein 60 Rattus norvegicus 2 | 39.47 | 1.48 | 62503 | 1 | 1 | 5209145 | 4348809 | 4012661 | 4671758 | 4392970.5 | 3957477 | 4857380 | 4869659.5 | 3962270.75 | 5670215.5 | 4271791 | 4023363.25 | 4258667.5 | 4059747.75 | 4090122.75 |
| Q5XIB5 | Ccdc86 | Coiled-coil domain-containing protein 86 Rattus norvegicus | 94.35 | 5.87 | 38585 | 2 | 2 | 2337335.25 | 1 | 1 | 826935 | 293555.72 | 252870.42 | 1 | 1 | 1 | 1 | 354800.59 | 1 | 327370.97 | 1 | 1 |
| Q4QRL3 | Ccdc88b | Coiled-coil domain-containing protein 88B Mus musculus | 136.3 | 2.5 | 166608 | 3 | 3 | 1218128.12 | 1376291.75 | 953901 | 1402982.25 | 1371815.5 | 1273097.75 | 1056586.88 | 1186405.5 | 1034238.31 | 977266 | 1437043.38 | 1062576 | 1155518.5 | 1127953.25 | 1166499.5 |
| Q5BJT7 | Ccdc93 | Coiled-coil domain-containing protein 93 Rattus norvegicus | 150.33 | 5.25 | 72636 | 3 | 3 | 3673638 | 4200331.5 | 2638253 | 2660034.25 | 2721032.5 | 3002259 | 2921148.25 | 3131827 | 3629392.5 | 3982411.75 | 3400241 | 3582706 | 2504681 | 2460640.25 | 2197373.5 |
| P51943 | Ccna2 | Cyclin-A2 Mus musculus | 36.96 | 3.32 | 47269 | 1 | 1 | 1 | 1 | 1 | 1 | 1 | 1 | 468130.16 | 581772.12 | 446328.03 | 1 | 639892.56 | 612440.62 | 550589.0225 | 517462.91 | 432560 |
| P24860 | Ccnb1 | G2/mitotic-specific cyclin-B1 Mus musculus | 75.11 | 4.88 | 48052 | 1 | 2 | 1 | 1 | 210056.78 | 1 | 1 | 305435.56 | 494806.69 | 343590.62 | 217477.61 | 356642.62 | 147209.69 | 347259.97 | 278458.2025 | 229709.59 | 389653.56 |
| P30276 | Ccnb2 | G2/mitotic-specific cyclin-B2 Mus musculus | 62.46 | 5.53 | 45383 | 1 | 2 | 1 | 1 | 210056.78 | 1 | 1 | 305435.56 | 261999.73 | 285388.88 | 217477.61 | 356642.62 | 147209.69 | 347259.97 | 1 | 1 | 389653.56 |
| P25322 | Ccnd1 | G1/S-specific cyclin-D1 Mus musculus | 116.65 | 8.14 | 33429 | 2 | 2 | 815802.12 | 537644.81 | 819930.62 | 446154.72 | 909177.38 | 510132 | 509525 | 933918.94 | 631376.06 | 672466.69 | 814362.31 | 496010.81 | 441316.75 | 1099973.25 | 947522 |
| Q8BGU5 | Ccny | Cyclin-Y Mus musculus | 98.72 | 8.21 | 39395 | 2 | 2 | 1668393.38 | 500108.31 | 1841246 | 1178046.38 | 1366125.75 | 1329987.25 | 1161933.88 | 1626322.12 | 2549601 | 1317235.75 | 1382303 | 1708856.62 | 2032148.88 | 401293.06 | 1270865.75 |
| Q9WU84 | Ccs | Copper chaperone for superoxide dismutase Mus musculus | 129.26 | 18.25 | 28912 | 2 | 2 | 1 | 405529.62 | 485801.06 | 1 | 1 | 1475405.11 | 2229266.5 | 1904709.75 | 1395607.38 | 372036.81 | 1500236.12 | 2651408 | 1660056.75 | 1752098.88 | 2375369 |
| P80314 | Cct2 | T-complex protein 1 subunit beta Mus musculus | 2345.441 | 73.08 | 57477 | 6 | 36 | 94758409 | 77278281 | 87927633 | 78477073 | 75809265 | 84827305 | 97647521 | 85464417 | 90562857 | 77292081 | 90450233 | 96904513 | 87655137 | 87510897 | 87865409 |
| P80318 | Cct3 | T-complex protein 1 subunit gamma Mus musculus | 1900.41 | 63.85 | 60630 | 32 | 33 | 188583281 | 213317041 | 224270465 | 219631633 | 215753985 | 196627393 | 213892497 | 209378929 | 208034785 | 221623713 | 210979217 | 195333617 | 210602017 | 208326497 | 206280545 |
| P80315 | Cct4 | T-complex protein 1 subunit delta Mus musculus | 2634.18 | 79.59 | 58066 | 36 | 39 | 203020705 | 232607457 | 233029761 | 218289569 | 202955649 | 222519809 | 229473617 | 230219809 | 245931569 | 226837953 | 217327265 | 247289185 | 217269361 | 226739441 | 208579521 |
| P80316 | Cct5 | T-complex protein 1 subunit epsilon Mus musculus | 1763.8 | 72.09 | 59624 | 30 | 30 | 126087761 | 135700049 | 141273249 | 132033305 | 129718777 | 129163249 | 144301457 | 135568529 | 140617585 | 142770625 | 145140705 | 134388081 | 140882017 | 145534289 | 133286529 |
| P80317 | Cct6a | T-complex protein 1 subunit zeta Mus musculus | 1564.93 | 57.44 | 58004 | 21 | 25 | 103082353 | 93190521 | 98589273 | 101548561 | 99895985 | 102450881 | 102881321 | 101232185 | 101203097 | 102456425 | 94064641 | 98925769 | 100736849 | 105983409 | 94274689 |
| Q61390 | Cct6b | T-complex protein 1 subunit zeta-2 Mus musculus | 253.78 | 11.49 | 58185 | 1 | 5 | 5120923 | 3767310 | 2864682 | 3987979.25 | 4153049.25 | 3511858.25 | 6514843 | 3229087 | 3870451.75 | 3456077.5 | 4078119 | 4494495.5 | 3232733 | 4580291 | 3532152.5 |
| P80313 | Cct7 | T-complex protein 1 subunit eta Mus musculus | 1864.45 | 72.24 | 59652 | 29 | 30 | 96133929 | 99549185 | 108503177 | 99808177 | 103498409 | 102276249 | 118680257 | 98758401 | 93342009 | 108990921 | 103003433 | 102350153 | 101190657 | 96536337 | 107738153 |
| P42932 | Cct8 | T-complex protein 1 subunit theta Mus musculus | 1978.12 | 66.24 | 59555 | 30 | 31 | 155005009 | 161322865 | 171043905 | 172423009 | 167592417 | 165151233 | 186819297 | 160251233 | 165127265 | 186107505 | 181490849 | 174691825 | 168512225 | 160097809 | 185293409 |
| P10810 | Cd14 | Monocyte differentiation antigen CD14 Mus musculus | 919.15 | 51.37 | 39204 | 13 | 13 | 46938701 | 51451925 | 49639101 | 46202521 | 44246909 | 34420477 | 35725377 | 42861973 | 38397501 | 40717569 | 40970305 | 48022557 | 48638241 | 49530917 | 40180361 |
| Q62192 | Cd180 | CD180 antigen Mus musculus | 843.69 | 28.59 | 74302 | 14 | 14 | 24539871 | 22443247 | 26022701 | 18453029 | 21320899 | 25811383 | 15314250 | 26346741 | 22741627 | 14089276 | 25664217 | 23542269 | 17068397 | 22729963 | 22852101 |
| Q9JLQ0 | Cd2ap | CD2-associated protein Mus musculus | 209.12 | 8.32 | 70450 | 4 | 4 | 2313159 | 1819108.5 | 2543226 | 1921423.62 | 2484053.5 | 1836580.75 | 2916917 | 2798201.75 | 3530756.5 | 3071759 | 1726977.75 | 2534956.25 | 2006711.25 | 2525750.75 | 2532662.75 |
| Q9CWK3 | Cd2bp2 | CD2 antigen cytoplasmic tail-binding protein 2 Mus musculus | 42.89 | 4.09 | 37694 | 1 | 1 | 924238.69 | 937526.25 | 700660.44 | 880414.75 | 959233.62 | 897650.69 | 761269.38 | 659756.19 | 753454.1125 | 695140.19 | 753146.94 | 645520.31 | 669303.595 | 654311.94 | 624235.19 |
| P15379 | Cd44 | CD44 antigen Mus musculus | 248.12 | 4.5 | 85617 | 4 | 4 | 21419985 | 27528795 | 25653127 | 22303749 | 26399949 | 26802497 | 22083039 | 29159913 | 28790141 | 22330665 | 25931949 | 25968417 | 29225595 | 25730689 | 23934493 |
| Q61735 | Cd47 | Leukocyte surface antigen CD47 Mus musculus | 204.21 | 11.22 | 33098 | 3 | 3 | 10764981 | 8533328 | 10018729 | 7033180 | 9425581 | 9404223 | 8113775 | 8918659 | 10482838 | 8005250 | 10844465 | 9175887 | 10076893 | 9295707 | 8340802 |
| P13379 | Cd5 | T-cell surface glycoprotein CD5 Mus musculus | 303.12 | 14.98 | 53849 | 5 | 5 | 2857282.5 | 4351480.5 | 2455905 | 2650475 | 2625699.25 | 4578047.5 | 5763710 | 4574634.5 | 3744935.75 | 5272260 | 3112701 | 3295489.75 | 3840225.5 | 3432162.5 | 2096972.75 |
| P31996 | Cd68 | Macrosialin Mus musculus | 141.59 | 13.5 | 34818 | 3 | 3 | 100387481 | 94790417 | 93696321 | 89821873 | 111325049 | 114962001 | 76916633 | 112623721 | 130320105 | 88327553 | 108855865 | 97327905 | 92050929 | 75931745 | 95495497 |
| P21855 | Cd72 | B-cell differentiation antigen CD72 Mus musculus | 200.92 | 13.28 | 40347 | 4 | 4 | 5472007 | 4998917 | 4320119.5 | 4838255 | 4878189.5 | 6957509.5 | 8683093 | 5973143 | 5982663 | 7385373 | 5618317 | 5174573.5 | 5244746 | 2656689.75 | 5098806.5 |
| P42082 | Cd86 | T-lymphocyte activation antigen CD86 Mus musculus | 46.91 | 4.53 | 34666 | 1 | 1 | 3228302 | 3338046.5 | 3090599.25 | 3026812.25 | 3151998.25 | 3583497.5 | 3383173.25 | 3663295.5 | 3080445.5 | 4008486.25 | 3696449 | 3576952.5 | 3739713.5 | 3068128 | 3175096.25 |
| P40240 | Cd9 | CD9 antigen Mus musculus | 82.53 | 11.06 | 25258 | 1 | 1 | 6655790.5 | 1 | 1 | 1 | 993260.69 | 1500925.12 | 1266282.12 | 1233344.5 | 1493229.25 | 1187592.5 | 1376572.25 | 1852455.75 | 1593315.625 | 913668 | 2230566.5 |
| Q8CII2 | Cdc123 | Cell division cycle protein 123 homolog Mus musculus | 166.21 | 9.82 | 38816 | 3 | 3 | 1019178.25 | 1426347.5 | 1078103.25 | 1218294.12 | 1018506.25 | 1196883.38 | 990248.12 | 1300476.25 | 1726969.75 | 1036555.62 | 1147144.25 | 505744.53 | 1118304.5 | 1290516.25 | 1331382.88 |
| Q8BGZ4 | Cdc23 | Cell division cycle protein 23 homolog Mus musculus | 100.45 | 4.02 | 68562 | 2 | 2 | 821238.69 | 1 | 1 | 312803.72 | 1 | 672225 | 1 | 561528.19 | 968073.56 | 1 | 560648.31 | 346945.72 | 329994.34 | 636959.12 | 280680.28 |
| Q8CFI2 | Cdc34 | Ubiquitin-conjugating enzyme E2 R1 Mus musculus | 49.06 | 11.91 | 26622 | 1 | 1 | 190175.52 | 354373 | 278904.62 | 179926.92 | 421873.56 | 1 | 209250.27 | 236215.31 | 177532.34 | 1 | 280699.75 | 166489.39 | 357942.47 | 281000.94 | 355681.03 |
| Q61081 | Cdc37 | Hsp90 co-chaperone Cdc37 Mus musculus | 512.12 | 25.86 | 44593 | 11 | 11 | 17853611 | 10750638 | 13867500 | 10214349 | 13448626 | 12670193 | 10875309 | 12792623 | 13436296 | 12052632 | 12288389 | 14649525 | 9450775 | 14185729 | 16160097 |
| P60766 | Cdc42 | Cell division control protein 42 homolog Mus musculus | 537.62 | 58.64 | 21259 | 8 | 9 | 41102253 | 34127069 | 39225869 | 42724749 | 36411677 | 36304377 | 46954521 | 38592845 | 34726221 | 44603321 | 44012897 | 40023745 | 36792909 | 34039105 | 44265829 |
| Q6A068 | Cdc5l | Cell division cycle 5-like protein Mus musculus | 774.22 | 25.44 | 92190 | 13 | 13 | 4205941.5 | 5390162.5 | 6357427.5 | 8175404.5 | 6828637 | 5878039.5 | 4982103.5 | 3800629.25 | 5974310 | 8951197 | 6967803.5 | 5874476.5 | 8420012 | 4134554.75 | 5099095 |
| Q4V8C8 | Cdc73 | Parafibromin Rattus norvegicus | 210.79 | 10.36 | 60603 | 5 | 5 | 1646665.25 | 613052.5 | 676812.5 | 1749981.25 | 758705.88 | 587972.19 | 1338722.75 | 1349336 | 1223693 | 840511.62 | 700854.62 | 849176 | 569705.81 | 1368341.75 | 1340526.88 |
| Q8VDP6 | Cdipt | CDP-diacylglycerol--inositol 3-phosphatidyltransferase Mus musculus | 291.04 | 37.56 | 23599 | 6 | 6 | 2878746.75 | 3684822.25 | 2994393.5 | 3307712.5 | 3192957 | 4436421 | 3031577.25 | 3021622.25 | 3893270.75 | 3402266 | 2941816.25 | 4142004.25 | 3694992.75 | 3636912 | 3545299.75 |
| P11440 | Cdk1 | Cyclin-dependent kinase 1 Mus musculus | 766.05 | 56.9 | 34107 | 13 | 14 | 9423399 | 6054676 | 7544436 | 6785699 | 5225055.5 | 6205774 | 9395870 | 5963741 | 5128043 | 4114015.5 | 7521132.5 | 7610936 | 8940263 | 6968811.5 | 7393586.5 |
| Q04735 | Cdk16 | Cyclin-dependent kinase 16 Mus musculus | 78.35 | 4.03 | 55914 | 1 | 2 | 450709.94 | 1 | 1 | 342831.16 | 155028.75 | 234218.73 | 473308.47 | 173605.03 | 341042.69 | 299750.41 | 450888.38 | 324475.19 | 1 | 429666.56 | 1 |
| Q04899 | Cdk18 | Cyclin-dependent kinase 18 Mus musculus | 295.61 | 17.07 | 51848 | 5 | 6 | 565470.75 | 616130.44 | 1319615.12 | 2423496 | 1317503.75 | 985417.12 | 1434984.5 | 1262761.88 | 235029.81 | 960308.38 | 1302395.5 | 956160.81 | 1114199.75 | 2129673.5 | 1195698.12 |
| P97377 | Cdk2 | Cyclin-dependent kinase 2 Mus musculus | 199.38 | 15.32 | 38978 | 1 | 4 | 1 | 337874.97 | 1 | 602628.19 | 1 | 1 | 1056240.38 | 316369.53 | 769794.25 | 1 | 536944.62 | 582501 | 1031977 | 453672.06 | 1879581.5 |
| O35207 | Cdk2ap1 | Cyclin-dependent kinase 2-associated protein 1 Mus musculus | 71.07 | 11.4 | 12354 | 1 | 1 | 1 | 1 | 141132.53 | 1 | 1 | 1 | 1 | 1 | 192075.73 | 1 | 136996.48 | 208208.06 | 108913.55 | 179882.38 | 132550.14 |
| P30285 | Cdk4 | Cyclin-dependent kinase 4 Mus musculus | 162.21 | 15.84 | 33751 | 3 | 3 | 1 | 357160.56 | 1 | 1937852.62 | 1 | 1 | 353266.78 | 388521.81 | 525629.19 | 1323695.38 | 1 | 334801.97 | 1 | 1188078.12 | 577344.75 |
| P49615 | Cdk5 | Cyclin-dependent-like kinase 5 Mus musculus | 107.15 | 10.62 | 33288 | 3 | 3 | 1243612.25 | 2242653.25 | 2584870.25 | 1929038.88 | 1953487.5 | 2275031.25 | 1848816 | 2539387.5 | 3012231.5 | 1796389.25 | 2821363.5 | 2708601.25 | 1817099.88 | 2645613.75 | 2520568.5 |
| Q99LM2 | Cdk5rap3 | CDK5 regulatory subunit-associated protein 3 Mus musculus | 146.32 | 7.55 | 56991 | 3 | 3 | 594501.25 | 551663.5 | 323095.75 | 164270.67 | 355088.12 | 148698.25 | 151323.56 | 353578.44 | 514214.91 | 511843.94 | 433278.31 | 295025.12 | 371755.12 | 119710.28 | 457603.94 |
| Q64261 | Cdk6 | Cyclin-dependent kinase 6 Mus musculus | 130.25 | 13.8 | 37028 | 3 | 3 | 602926.81 | 1661433.5 | 1175220 | 1501017 | 1249175.5 | 1533261.75 | 1711656.5 | 1082547 | 1117393.12 | 1938986 | 1528475.62 | 1125727.5 | 1210103.5 | 1118522.5 | 1239548.75 |
| Q641Z4 | Cdk9 | Cyclin-dependent kinase 9 Rattus norvegicus | 93.84 | 5.91 | 42762 | 2 | 2 | 294274.34 | 389637.75 | 429396.91 | 272575.97 | 373023.84 | 381494.16 | 204534.19 | 308365.565 | 363202.97 | 284230.94 | 292125.03 | 322174.5275 | 254780.86 | 385414.41 | 356377.81 |
| Q5U2X0 | Cdkn2aip | CDKN2A-interacting protein Rattus norvegicus | 34.6 | 1.93 | 60744 | 1 | 1 | 1137211.38 | 1 | 1566931.25 | 392017.84 | 1 | 1 | 1 | 1 | 1 | 1181075.25 | 1977603.12 | 1 | 1 | 1 | 1 |
| Q4VAA2 | Cdv3 | Protein CDV3 Mus musculus | 199.71 | 18.86 | 29729 | 3 | 3 | 7609962 | 8281187 | 6559194 | 7483821 | 9106882 | 7662372 | 6339039 | 6603798 | 6706385.5 | 7265611.5 | 8499153 | 5955907 | 7746962 | 6966769.5 | 5158444.5 |
| P28659 | Celf1 | CUGBP Elav-like family member 1 Mus musculus | 137.04 | 7.2 | 52107 | 2 | 3 | 989301.38 | 1403307.62 | 1312929.62 | 1368515.25 | 1008869 | 1701497.25 | 1258698 | 1083183 | 1352295.25 | 1417806.62 | 1472879.25 | 1540089.38 | 1240553.88 | 1315772.62 | 1277296 |
| Q792H5 | Celf2 | CUGBP Elav-like family member 2 Rattus norvegicus | 180.57 | 7.68 | 54271 | 2 | 3 | 957990.44 | 317687.5 | 1052966.25 | 816434.69 | 648280.06 | 1 | 981057.44 | 972921 | 918519.56 | 865328.81 | 1127249 | 1089862 | 1 | 1 | 1422637.5 |
| O88278 | Celsr3 | Cadherin EGF LAG seven-pass G-type receptor 3 Rattus norvegicus | 36.49 | 0.24 | 359355 | 1 | 1 | 3292740.75 | 3573454.5 | 4433192.5 | 2411884.5 | 3863846.25 | 3342885.5 | 1859105.88 | 4135015.5 | 8123628.5 | 1828634.38 | 4499519.5 | 3562533.25 | 2494928 | 4181365 | 2942666 |
| Q6A065 | Cep170 | Centrosomal protein of 170 kDa Mus musculus | 176.73 | 2.9 | 175050 | 4 | 4 | 6052194 | 4725799 | 4774246 | 6748361 | 4813490.5 | 4377784 | 4907023 | 3880492.5 | 3632213 | 4531489.5 | 4629409.5 | 2634284.5 | 3921717.5 | 3425801 | 4409769 |
| Q8BT07 | Cep55 | Centrosomal protein of 55 kDa Mus musculus | 89.24 | 4.55 | 53930 | 2 | 2 | 551833.94 | 1 | 682923.56 | 548555.31 | 1 | 478084.59 | 617216.62 | 547963.44 | 474553.62 | 919010.62 | 508967.88 | 515330.56 | 414074.88 | 639035.19 | 738743.44 |
| Q8BGS7 | Cept1 | Choline/ethanolaminephosphotransferase 1 Mus musculus | 135.24 | 8.89 | 46434 | 2 | 2 | 1634994 | 960700 | 1007702.62 | 882793.5 | 814427.62 | 939307.81 | 803171.75 | 688238.38 | 1043942.31 | 865466.69 | 1295202.5 | 911142.5 | 732999.94 | 764523.12 | 1325973 |
| Q924Z4 | Cers2 | Ceramide synthase 2 Mus musculus | 107.02 | 9.47 | 45024 | 2 | 2 | 4171508.5 | 1 | 1 | 1 | 1 | 1 | 1 | 313929.97 | 1 | 1 | 1 | 1 | 1 | 1 | 487471.22 |
| Q9D6J1 | Cers4 | Ceramide synthase 4 Mus musculus | 33.15 | 2.04 | 46017 | 1 | 1 | 1075401 | 1 | 1 | 1 | 1 | 432925.12 | 1 | 331484.16 | 339554.12 | 1 | 154257.45 | 280276.22 | 274672.5 | 266547.59 | 243938.44 |
| Q9D6K9 | Cers5 | Ceramide synthase 5 Mus musculus | 111.7 | 9.18 | 48167 | 2 | 2 | 834003.69 | 202209.67 | 186993.09 | 234875.84 | 311980.56 | 297585.72 | 176795.89 | 294363.97 | 176657.67 | 177449.12 | 210805.16 | 186945.55 | 238950.91 | 180294.47 | 201061.39 |
| Q8BTU1 | Cfap20 | Cilia- and flagella-associated protein 20 Mus musculus | 173.2 | 30.05 | 22748 | 4 | 4 | 7629927 | 8499859 | 8663983 | 7950704 | 6956509.5 | 7696530.5 | 4048041 | 7618430.5 | 7600208.5 | 5050671 | 575085.44 | 647639 | 5598790 | 6021284.5 | 4250360 |
| O88271 | Cfdp1 | Craniofacial development protein 1 Mus musculus | 215.9 | 21.02 | 32921 | 4 | 4 | 1691867.12 | 2441219.5 | 1857860.38 | 1772106.75 | 2289406 | 2278071 | 2304883.25 | 2290741.5 | 2125647.75 | 2897988 | 1485850.62 | 1861238.38 | 2743837.75 | 1908105.5 | 2950761.75 |
| P18760 | Cfl1 | Cofilin-1 Mus musculus | 813.63 | 74.7 | 18560 | 12 | 12 | 337236609 | 319565633 | 351056481 | 363054849 | 393487073 | 359553601 | 419760225 | 344835713 | 310789825 | 285014689 | 428539617 | 402376417 | 405216449 | 351821217 | 374839681 |
| Q9CRB9 | Chchd3 | MICOS complex subunit Mic19 Mus musculus | 238.76 | 18.94 | 26335 | 4 | 4 | 2915369 | 2642379.5 | 3237089.5 | 3782201.75 | 2855165.5 | 2639357 | 5250266 | 3008453.75 | 2633257.75 | 3928207.5 | 2910578.25 | 3825746 | 3442245 | 2398326 | 3436393.5 |
| Q5BJN5 | Chchd4 | Mitochondrial intermembrane space import and assembly protein 40 Rattus norvegicus | 91.49 | 26.62 | 15467 | 2 | 2 | 2378723 | 2261497.75 | 1914399.5 | 1950681.75 | 2899819.5 | 2820971 | 1126189.62 | 1279980.12 | 3400368.5 | 2307370 | 2385990 | 2332402.5 | 2990516.5 | 2435793.5 | 2329266.25 |
| Q91VN4 | Chchd6 | MICOS complex subunit Mic25 Mus musculus | 145.23 | 15.02 | 29852 | 3 | 3 | 1233463 | 1436226.88 | 1608062.88 | 1344892.88 | 1678401.88 | 1599635.25 | 1795223.5 | 1357081.88 | 1370131.62 | 2398057 | 2028535.5 | 1668546.88 | 1310520.62 | 1136523.62 | 1454281.12 |
| Q6PDQ2 | Chd4 | Chromodomain-helicase-DNA-binding protein 4 Mus musculus | 1228.67 | 15.87 | 217751 | 26 | 26 | 23176529 | 30768121 | 24352573 | 28578991 | 32957677 | 31086831 | 32592141 | 29978555 | 28002133 | 32507199 | 28572775 | 31629719 | 32698383 | 30630317 | 30941943 |
| O35280 | Chek1 | Serine/threonine-protein kinase Chk1 Mus musculus | 45.24 | 2.31 | 54381 | 1 | 1 | 1 | 1 | 1 | 1 | 538319.56 | 1 | 1 | 1 | 1 | 462890.22 | 419317.97 | 1 | 1 | 1 | 467609.06 |
| Q8CGZ0 | Cherp | Calcium homeostasis endoplasmic reticulum protein Mus musculus | 102.33 | 5.02 | 106169 | 3 | 3 | 1 | 1 | 1 | 1 | 1 | 1 | 495221.84 | 1 | 552290.31 | 874965.19 | 505553.69 | 337668.66 | 929510 | 647254.5725 | 816285.94 |
| P37727 | Chm | Rab proteins geranylgeranyltransferase component A 1 Rattus norvegicus | 127.88 | 4.31 | 72480 | 2 | 2 | 1589408.5 | 2299672 | 2029950.25 | 1617285.5 | 2542880.5 | 1518350 | 1985588 | 2458126 | 2678121 | 1552655.75 | 1394277 | 2028313.5 | 2018981.38 | 623289.94 | 1808154 |
| Q8CGS4 | Chmp3 | Charged multivesicular body protein 3 Rattus norvegicus | 88 | 8.52 | 25062 | 2 | 2 | 1 | 1 | 315959.31 | 1 | 278388.94 | 1 | 1 | 1 | 543802.69 | 309685.88 | 303834.75 | 626819 | 216329.8 | 430905.22 | 365441.81 |
| Q9D8B3 | Chmp4b | Charged multivesicular body protein 4b Mus musculus | 189.84 | 29.46 | 24936 | 5 | 5 | 3567356.25 | 3260036 | 2346466.25 | 3154089 | 2301949.25 | 2267600 | 5147316.5 | 4209074 | 2902757.5 | 4660558 | 2327044.25 | 2697765 | 3425674.5 | 2927495.5 | 2389628.25 |
| P0C0A3 | Chmp6 | Charged multivesicular body protein 6 Mus musculus | 47.93 | 6.5 | 23416 | 1 | 1 | 843311.44 | 977564.75 | 696011 | 936585.69 | 910120.06 | 783512.44 | 793820.06 | 835219.06 | 699506.94 | 820796.31 | 878863.25 | 730981.25 | 776854.69 | 836100.56 | 631598.31 |
| Q8R1T1 | Chmp7 | Charged multivesicular body protein 7 Mus musculus | 129.92 | 3.77 | 50633 | 2 | 2 | 587070.62 | 493761 | 444649.38 | 485101.81 | 473849.34 | 850190.38 | 634091.69 | 506364.22 | 532246.81 | 628442.25 | 731968.5 | 490311.44 | 686575.38 | 645324.25 | 457355.75 |
| D4A4T9 | Chordc1 | Cysteine and histidine-rich domain-containing protein 1 Rattus norvegicus | 436.58 | 35.65 | 37362 | 9 | 9 | 8176075 | 9282825 | 8180874.5 | 8925010 | 7551701 | 6095291.5 | 14304673 | 6295618.5 | 12419075 | 12940437 | 12714709 | 12742396 | 12036225 | 10196495 | 13519816 |
| P61022 | Chp1 | Calcineurin B homologous protein 1 Mus musculus | 69.3 | 6.15 | 22432 | 1 | 1 | 1712638 | 1428324.25 | 1986900.5 | 1183351 | 1516923 | 1655633.75 | 1642654.75 | 2448606.75 | 1754979.12 | 1747941.88 | 1294993.38 | 1920870.75 | 1342364.88 | 1873732.75 | 1384756 |
| Q9CY57 | Chtop | Chromatin target of PRMT1 protein Mus musculus | 78.43 | 5.22 | 26585 | 1 | 1 | 1 | 1 | 1 | 1 | 1 | 421872.91 | 1 | 1 | 295603.19 | 423144.03 | 468894.88 | 1 | 562923.38 | 548541.56 | 1 |
| Q60680 | Chuk | Inhibitor of nuclear factor kappa-B kinase subunit alpha Mus musculus | 68.26 | 3.36 | 84729 | 2 | 2 | 713013.5 | 800140.5 | 876958.75 | 758293.38 | 912225.06 | 835647.38 | 491359.62 | 784971.44 | 1293367.25 | 589711.25 | 528863.81 | 972727.62 | 701691.88 | 1342971.12 | 1245617.12 |
| Q9DCL2 | Ciao2a | Cytosolic iron-sulfur assembly component 2A Mus musculus | 43.48 | 5.62 | 18418 | 1 | 1 | 1704677.88 | 1674099.25 | 1883725.62 | 1286717 | 1468428.25 | 1667530.75 | 1018046.81 | 1684478.38 | 1915061.5 | 1327734.25 | 1519053.5 | 1262041 | 1648564 | 1630927.12 | 1010492.38 |
| Q8WTY4 | Ciapin1 | Anamorsin Mus musculus | 383.78 | 35.28 | 33429 | 7 | 7 | 4608548.5 | 8163353 | 5240400 | 7114874.5 | 7039459.5 | 7018846 | 8929018 | 5413015.5 | 6662300.5 | 7526856 | 9319681 | 7154783 | 6787567 | 7930821 | 7025837.5 |
| Q8BWY9 | Cip2a | Protein CIP2A Mus musculus | 52.33 | 1.54 | 102043 | 1 | 1 | 1 | 1 | 1 | 1 | 1 | 1 | 1 | 1 | 1 | 1 | 1 | 1 | 309564.5 | 1 | 1 |
| B0K020 | Cisd1 | CDGSH iron-sulfur domain-containing protein 1 Rattus norvegicus | 36.33 | 8.33 | 12097 | 1 | 1 | 1284328.875 | 1110574 | 1254126.12 | 1242726.5 | 1529888.88 | 1536594.38 | 1916388 | 973976.75 | 1097885.12 | 1924455.75 | 1561597.25 | 1649093.12 | 1410617.5 | 887616.81 | 1572790.75 |
| Q9CQB5 | Cisd2 | CDGSH iron-sulfur domain-containing protein 2 Mus musculus | 172.63 | 31.11 | 15242 | 4 | 4 | 7221406 | 4986387.5 | 5483813 | 8125508 | 5127212 | 6416161 | 9141583 | 4850144.5 | 4877805.5 | 8518269 | 7055127.5 | 6871820.5 | 8038968 | 6467616 | 6590943.5 |
| Q8BMK4 | Ckap4 | Cytoskeleton-associated protein 4 Mus musculus | 932.77 | 36.7 | 63692 | 15 | 15 | 21448823 | 17194699 | 19210087 | 15689124 | 16096324 | 22998951 | 26209203 | 25156477 | 20380719 | 22363581 | 20912823 | 20589509 | 18392507 | 20472573 | 17722387 |
| A2AGT5 | Ckap5 | Cytoskeleton-associated protein 5 Mus musculus | 888.94 | 11.71 | 225635 | 20 | 20 | 21181283 | 23680455 | 23635451 | 20457251 | 20914393 | 20445475 | 23942101 | 22410595 | 21930225 | 22766885 | 20030259 | 22147115 | 26418687 | 20899243 | 22427379 |
| Q04447 | Ckb | Creatine kinase B-type Mus musculus | 1693.04 | 67.72 | 42713 | 20 | 23 | 255307153 | 251038737 | 223875297 | 278372481 | 251370417 | 255638561 | 317101633 | 230963697 | 280863681 | 287091457 | 324837697 | 258508865 | 229898945 | 233064625 | 264287777 |
| P00564 | Ckm | Creatine kinase M-type Rattus norvegicus | 47.42 | 8.4 | 43045 | 1 | 1 | 2756509.75 | 1012554.81 | 1432916.62 | 2873567.75 | 3550010.25 | 2603503.75 | 3093986.75 | 1745810.62 | 900814.5 | 993026.62 | 2979174.25 | 754220.25 | 2319419 | 2084017.12 | 2594048.5 |
| Q80TV8 | Clasp1 | CLIP-associating protein 1 Mus musculus | 136.78 | 1.95 | 169227 | 1 | 3 | 739252.69 | 1063765 | 701905.12 | 591517.31 | 617944 | 788326.12 | 2417013.75 | 575177.62 | 729653 | 972022.12 | 1039842.06 | 659468.44 | 731603.5 | 580221.19 | 565493.88 |
| Q8BRT1 | Clasp2 | CLIP-associating protein 2 Mus musculus | 128.81 | 2.41 | 140739 | 1 | 3 | 739252.69 | 715702 | 453115.66 | 591517.31 | 617944 | 788326.12 | 517136.31 | 575177.62 | 729653 | 972022.12 | 1039842.06 | 489309.19 | 731603.5 | 580221.19 | 565493.88 |
| Q99LI2 | Clcc1 | Chloride channel CLIC-like protein 1 Mus musculus | 183.63 | 8.91 | 60621 | 3 | 3 | 1968337.25 | 1 | 571174.94 | 272969.69 | 1 | 749656.81 | 720467.69 | 736804.62 | 728879.75 | 441331.06 | 1048033.69 | 705264.5475 | 426461.25 | 240185 | 1106378.25 |
| P49300 | Clec10a | C-type lectin domain family 10 member A Mus musculus | 187.5 | 15.46 | 34596 | 4 | 4 | 2979940.25 | 2545870.25 | 3094110.5 | 3825382.25 | 2324808 | 3152211.5 | 2432490.25 | 3704485.75 | 3274285.25 | 1676765 | 2922276.75 | 4269314.5 | 2057123 | 2312394 | 2367037.5 |
| Q504P2 | Clec12a | C-type lectin domain family 12 member A Mus musculus | 58.83 | 3.37 | 30757 | 1 | 1 | 833108.38 | 804176.25 | 742673.62 | 603270.94 | 734008.75 | 1079318.88 | 798274.75 | 777697.06 | 821750.06 | 610533.56 | 657914.75 | 528790.19 | 830357.56 | 870315.75 | 659172.06 |
| Q6MG61 | Clic1 | Chloride intracellular channel protein 1 Rattus norvegicus | 1123.47 | 62.66 | 26981 | 14 | 14 | 202635617 | 232196769 | 213661681 | 198470033 | 209748017 | 217069617 | 224759153 | 229866289 | 248582001 | 214097857 | 208410321 | 223322369 | 225111281 | 227987697 | 216177409 |
| Q9QYB1 | Clic4 | Chloride intracellular channel protein 4 Mus musculus | 1131.61 | 74.31 | 28729 | 17 | 17 | 97964481 | 99478145 | 87956129 | 100850505 | 100354753 | 121194929 | 137273777 | 113730889 | 105239289 | 123683441 | 115262889 | 114896017 | 109952089 | 98826713 | 118439625 |
| Q99KN9 | Clint1 | Clathrin interactor 1 Mus musculus | 269.06 | 10.14 | 68513 | 5 | 5 | 10063867 | 10178283 | 9416835 | 8592088 | 9400185 | 10706768 | 9988097 | 9416449 | 11487594 | 8532107 | 10419606 | 9166081 | 9187862 | 9592605 | 9568208 |
| Q922J3 | Clip1 | CAP-Gly domain-containing linker protein 1 Mus musculus | 279.64 | 5.1 | 155814 | 5 | 5 | 3655231 | 1922632.5 | 3088183 | 4579470 | 4091953.25 | 3768759.75 | 5090205.5 | 1852150.25 | 1235377.12 | 3417225.5 | 4736918.5 | 1728942 | 2397249 | 1806043.5 | 3168269 |
| Q3UMW8 | Cln5 | Ceroid-lipofuscinosis neuronal protein 5 homolog Mus musculus | 100.62 | 4.69 | 39329 | 2 | 2 | 2946472.75 | 2793477.5 | 2899517.25 | 1730103.12 | 2395201 | 2712768 | 1703323.75 | 2603544 | 2516738.5 | 1771205.5 | 1916783 | 1880345.5 | 2367174.5 | 2870337.75 | 2000952.25 |
| Q8VBZ3 | Clptm1 | Cleft lip and palate transmembrane protein 1 homolog Mus musculus | 260.95 | 11.3 | 75291 | 4 | 4 | 8212460 | 8418120 | 5859314 | 5419005.5 | 6493413.5 | 6894831 | 4384326 | 7811002 | 6670826.5 | 3275917.25 | 8572105 | 7428446.5 | 6667761 | 8133376.5 | 8293119 |
| Q5U2U0 | Clpx | ATP-dependent Clp protease ATP-binding subunit clpX-like, mitochondrial Rattus norve | 222.6 | 10.43 | 69208 | 5 | 5 | 3114605.75 | 1965507.62 | 24007401 | 3122183.25 | 43430297 | 2039697.88 | 1756833 | 25015375 | 2284610.25 | 18799201 | 1837992.75 | 2636343 | 21770633 | 21242117 | 2178286.5 |
| O08585 | Clta | Clathrin light chain A Mus musculus | 123.19 | 10.64 | 25604 | 3 | 3 | 22239505 | 23305617 | 20524421 | 27492887 | 23438983 | 21142393 | 16876155 | 20501821 | 19461057 | 20894157 | 21101131 | 17924757 | 21634389 | 18912855 | 16271817 |
| Q68FD5 | Cltc | Clathrin heavy chain 1 Mus musculus | 5817.88 | 65.13 | 191557 | 85 | 85 | 464104705 | 492453857 | 482883425 | 470813473 | 474982721 | 448120993 | 412333281 | 476912961 | 509476097 | 428730433 | 455720513 | 464393409 | 468304065 | 490648257 | 448803425 |
| Q5SW19 | Cluh | Clustered mitochondria protein homolog Mus musculus | 502.78 | 11.03 | 148068 | 11 | 11 | 15011846 | 13132903 | 10006947 | 8579896 | 11412827 | 10926138 | 8422665 | 11820719 | 10552517 | 9760839 | 9512103 | 11493381 | 11406711 | 13172009 | 10395945 |
| Q8R4N0 | Clybl | Citramalyl-CoA lyase, mitochondrial Mus musculus | 129.86 | 14.79 | 37549 | 3 | 3 | 1999024.75 | 2276726 | 1732721.88 | 2577930.25 | 2283561.5 | 2651648.5 | 1992235.88 | 2966676.5 | 1958625.88 | 1632718 | 1968423.5 | 1698699.75 | 2229714.5 | 2171438 | 1665389 |
| Q99KK2 | Cmas | N-acylneuraminate cytidylyltransferase Mus musculus | 181.28 | 11.11 | 48058 | 4 | 4 | 11565397 | 13408209 | 12874419 | 9874251 | 13388289 | 13956426 | 13551037 | 14111877 | 14843570 | 13556021 | 13995903 | 12372027 | 11551751 | 13045272 | 12435074 |

| | | | | | | | | | | | | | | | | | | | | | | |
|---|---|---|---|---|---|---|---|---|---|---|---|---|---|---|---|---|---|---|---|---|---|---|
| Q9DBP5 | Cmpk1 | UMP-CMP kinase Mus musculus | 367.8 | 47.45 | 22165 | 7 | 7 | 13344170 | 12639166 | 15350142 | 12876946 | 14379416 | 15713997 | 15412135 | 14119387 | 12724581 | 14734828 | 13776251 | 15746605 | 13733987 | 15123623 | 18368211 |
| Q9DBC3 | Cmtr1 | Cap-specific mRNA (nucleoside-2'-O-)-methyltransferase 1 Mus musculus | 100.11 | 3.94 | 95676 | 3 | 3 | 1037354.94 | 2110121.5 | 936033.56 | 1068002.62 | 2170214.5 | 1792237.62 | 1970576.5 | 2208744 | 2169860.5 | 1839199.25 | 2052750.38 | 1837077.75 | 2382091.25 | 2338384 | 1728479.5 |
| P62634 | Cnbp | Cellular nucleic acid-binding protein Rattus norvegicus | 251.06 | 28.25 | 19463 | 4 | 4 | 7434644.5 | 13141701 | 11193080 | 14584421 | 11707286 | 9156525 | 13541223 | 9240622 | 9738617 | 12472703 | 12875909 | 9329858 | 12219937 | 11970521 | 12655523 |
| Q9D1A2 | Cndp2 | Cytosolic non-specific dipeptidase Mus musculus | 1509.08 | 60.84 | 52767 | 24 | 24 | 35378917 | 43228145 | 46832133 | 35846397 | 39982001 | 43498125 | 50321773 | 44239025 | 40840721 | 42656857 | 41241309 | 42470777 | 46303813 | 41103869 | 46332089 |
| Q9CX13 | Cnih4 | Protein cornichon homolog 4 Mus musculus | 67.89 | 14.39 | 16089 | 1 | 1 | 4690036 | 1326770.5 | 1530951.62 | 1344254.12 | 2211565.25 | 2451989.5 | 1678696.12 | 1314442.12 | 2370321.5 | 2282287.5 | 1831817.38 | 2634500 | 2739810 | 2749720.75 | 2865808.5 |
| Q6ZQ08 | Cnot1 | CCR4-NOT transcription complex subunit 1 Mus musculus | 298.98 | 4.46 | 266808 | 8 | 8 | 3597935.5 | 3165644.5 | 4145893 | 3276575.5 | 2317906.5 | 4083148.25 | 3232053 | 1909268.88 | 3812421.5 | 3372527 | 3166106.5 | 2238976.5 | 3282498.25 | 3744798.25 | 2171689.5 |
| Q5XIA4 | Cnot10 | CCR4-NOT transcription complex subunit 10 Rattus norvegicus 2 | 60.17 | 1.61 | 81819 | 1 | 1 | 716241.38 | 1 | 325269.28 | 645010.31 | 1 | 1 | 1 | 402336.25 | 711983.94 | 1 | 195637.77 | 492007.47 | 1 | 1 | 1 |
| Q8K0V4 | Cnot3 | CCR4-NOT transcription complex subunit 3 Mus musculus | 97.76 | 4.13 | 81946 | 2 | 2 | 575017.19 | 608880.31 | 571949.31 | 805101.12 | 757929 | 564979.38 | 589178.06 | 615387.25 | 643450.56 | 737831.69 | 720266 | 607145.75 | 504422.5 | 609518 | 1038052.69 |
| Q60809 | Cnot7 | CCR4-NOT transcription complex subunit 7 Mus musculus | 78.56 | 10.18 | 32718 | 2 | 2 | 1 | 452534.75 | 1 | 1 | 321391.88 | 1 | 1 | 505843.53 | 588084.94 | 1 | 219800.81 | 1 | 1 | 1 | 1 |
| Q5PQL2 | Cnot9 | CCR4-NOT transcription complex subunit 9 Rattus norvegicus | 132.57 | 11.71 | 33601 | 3 | 3 | 3470911 | 3178055.75 | 3712563 | 2878176.25 | 2229217 | 2499309 | 2646518.75 | 2901227.5 | 3308661.5 | 1913739.38 | 2582194 | 2981296.5 | 2707550 | 3225950.5 | 2605230.5 |
| P13233 | Cnp | 2',3'-cyclic-nucleotide 3'-phosphodiesterase Rattus norvegicus | 99.23 | 5.71 | 47268 | 2 | 2 | 177527.52 | 200313.7825 | 131076.45 | 256356.52 | 236294.64 | 89591.42 | 1 | 139433.39 | 174410.73 | 1 | 185504.78 | 152486.38 | 175437.8975 | 217315.27 | 146445.16 |
| Q9QXT0 | Cnpy2 | Protein canopy homolog 2 Mus musculus | 147.19 | 15.93 | 20767 | 2 | 2 | 2663602.5 | 2918795.5 | 2321279.5 | 2802940.5 | 2482477.5 | 2093529.62 | 2550972.25 | 2076847 | 2359377.5 | 2287966.25 | 2098434.5 | 2131443.25 | 2587944.75 | 1867885.38 | 1932708.25 |
| Q8BQ47 | Cnpy4 | Protein canopy homolog 4 Mus musculus | 35.44 | 4.9 | 28094 | 1 | 1 | 926013.4675 | 1050913.62 | 921976.56 | 817126.19 | 914037.5 | 978668.56 | 827407.19 | 923918.19 | 1030919.12 | 859698.25 | 948631.25 | 874085.81 | 803403.19 | 785820.25 | 948833.19 |
| Q5M7A7 | Cnrip1 | CB1 cannabinoid receptor-interacting protein 1 Rattus norvegicus | 57.64 | 9.76 | 18658 | 1 | 1 | 1062242.12 | 911571.5 | 1075807 | 917962.69 | 1316742.88 | 988307 | 928874.06 | 990646.56 | 937221.12 | 1032549.19 | 813023.19 | 785935.88 | 1091944 | 1019285.88 | 951804 |
| Q9DBL7 | Coasy | Bifunctional coenzyme A synthase Mus musculus | 180.57 | 12.79 | 62023 | 4 | 4 | 2029152.75 | 1589304.88 | 1651493.75 | 1497238.62 | 1222557.25 | 1567954.5 | 1743748.25 | 1878980.5 | 1779693.75 | 1488796.88 | 1487389.12 | 1738369.75 | 1373536.75 | 1740378.25 | 1831103.25 |
| Q9Z160 | Cog1 | Conserved oligomeric Golgi complex subunit 1 Mus musculus | 108.17 | 5.2 | 109050 | 3 | 3 | 1624671.5 | 1343315.75 | 1836370.12 | 1212280.12 | 1640043.62 | 1557877.5 | 1489945 | 1251667.88 | 1440379.12 | 1704305.75 | 1171968.5 | 1439937.88 | 1374985.88 | 1744033.38 | 1634463.25 |
| Q8C0L8 | Cog5 | Conserved oligomeric Golgi complex subunit 5 Mus musculus | 61.13 | 1.69 | 91391 | 1 | 1 | 666407.69 | 692205.75 | 561824.38 | 637569.38 | 589091.31 | 654213.69 | 664085.06 | 695587.56 | 535117.06 | 520011.09 | 568244.31 | 515894.5 | 727507.38 | 591793.06 | 555166.5 |
| Q3UM29 | Cog7 | Conserved oligomeric Golgi complex subunit 7 Mus musculus | 39.89 | 1.3 | 86073 | 1 | 1 | 877942.88 | 1048416.69 | 953760.94 | 772125.38 | 767377.81 | 993434.88 | 678240.62 | 832997.31 | 682597 | 904059.81 | 861147.81 | 891898.12 | 941321.25 | 870238.44 | 698000.5 |
| Q5SU73 | Coil | Coilin Mus musculus | 68.24 | 1.75 | 61917 | 1 | 1 | 716566.94 | 651259.75 | 686701.88 | 586638.19 | 793094.94 | 598069.31 | 683476 | 757092.19 | 781290.81 | 853089.12 | 618671.19 | 840528.06 | 664239.19 | 609101.88 | 592901.19 |
| Q9EQG9 | Col4a3bp | Collagen type IV alpha-3-binding protein Mus musculus | 226.77 | 10.26 | 71111 | 4 | 4 | 2020628 | 2719257.5 | 2112373.5 | 2053642.25 | 2452292 | 1923834.5 | 2054849.88 | 1328938.75 | 1909756.12 | 2044130.5 | 1933136.38 | 1072150.25 | 1880008.25 | 1940364.75 | 2142726.75 |
| Q8K4Q8 | Colec12 | Collectin-12 Mus musculus | 136.96 | 4.72 | 81304 | 2 | 2 | 1318988.75 | 1573271.25 | 1555674 | 927166.12 | 1725151.75 | 520393.94 | 787278.25 | 1630755.75 | 1744940.88 | 841760.38 | 1413186.75 | 832050.94 | 1832645.75 | 1385599.5 | 1024414 |
| Q8K297 | Colgalt1 | Procollagen galactosyltransferase 1 Mus musculus | 390.19 | 14.26 | 71061 | 8 | 8 | 5236550 | 5642999.5 | 5347797 | 5270898 | 5695256 | 4515568 | 4265825.5 | 5897949 | 5759001 | 7925002 | 5607499.5 | 5186760 | 4951342.5 | 4279729 | 6769379 |
| Q8BXC6 | Commd2 | COMM domain-containing protein 2 Mus musculus | 74.28 | 9.55 | 22848 | 2 | 2 | 2578403.25 | 2830499.25 | 2162274.75 | 2516614.75 | 2383693 | 1939588.75 | 1954253.25 | 2307488.5 | 2412509.5 | 1878735.5 | 2128163.75 | 2031448.12 | 2356999.25 | 2231211.75 | 1805655.25 |
| Q63829 | Commd3 | COMM domain-containing protein 3 Mus musculus | 218.1 | 22.05 | 22037 | 3 | 3 | 2526894.25 | 2162907.75 | 1817149.75 | 2571232.75 | 2248228.75 | 1859113.38 | 2333731 | 2237888 | 1823959.62 | 2273172.5 | 1464592.5 | 1790764.75 | 2088478.25 | 2439046.5 | 1672263.12 |
| Q9CQ02 | Commd4 | COMM domain-containing protein 4 Mus musculus | 99.52 | 17.59 | 21860 | 2 | 2 | 740026.56 | 614819 | 679925.75 | 539464.88 | 352466.81 | 512521.88 | 510310.41 | 699039.25 | 884181.06 | 331869.06 | 440089.22 | 642170 | 230866.75 | 728569.5 | 665594.75 |
| O88587 | Comt | Catechol O-methyltransferase Mus musculus | 409.03 | 37.74 | 29486 | 6 | 6 | 5544021 | 1821514.25 | 5037485 | 2181069 | 3859265.25 | 3225419 | 4194231 | 5515521 | 4545618 | 3702485.5 | 4255539 | 8430321 | 3678202.25 | 6803183 | 7034018.5 |
| Q8BIG7 | Comtd1 | Catechol O-methyltransferase domain-containing protein 1 Mus musculus | 557.36 | 41.22 | 28961 | 8 | 8 | 9628291 | 9743043 | 5514307.5 | 7986243 | 8284010 | 9144441 | 7482636 | 9171358 | 7921749 | 8738834 | 9938354 | 5728339 | 5193563 | 8440867 | 6488015.5 |
| Q8CIE6 | Copa | Coatomer subunit alpha Mus musculus | 2029.95 | 35.7 | 138432 | 37 | 37 | 80718177 | 78354401 | 77292553 | 71220417 | 76961153 | 79194657 | 73278201 | 77046705 | 84613137 | 73552569 | 76565417 | 76319881 | 71827425 | 80540321 | 84920737 |
| Q9JIF7 | Copb1 | Coatomer subunit beta Mus musculus | 1995.78 | 51.52 | 107066 | 35 | 35 | 56270101 | 53143837 | 54010401 | 48913977 | 52834477 | 51810437 | 53092741 | 51704717 | 60258221 | 51135037 | 53093949 | 48010037 | 53983849 | 54916357 | 51301253 |
| O35142 | Copb2 | Coatomer subunit beta' Rattus norvegicus | 1590.56 | 45.3 | 102551 | 29 | 29 | 65436313 | 74778705 | 72073345 | 71345665 | 73445985 | 71039689 | 60871901 | 70680129 | 79331473 | 67044377 | 74298033 | 72689201 | 68504417 | 74806177 | 70209881 |
| O89079 | Cope | Coatomer subunit epsilon Mus musculus | 679.2 | 45.45 | 34567 | 8 | 8 | 11416745 | 11867459 | 14669194 | 13050814 | 13300484 | 10700177 | 10110886 | 15609230 | 17998975 | 8903029 | 16868167 | 17178601 | 11666756 | 16864133 | 16680097 |
| Q9QZE5 | Copg1 | Coatomer subunit gamma-1 Mus musculus | 1278.92 | 42.33 | 97513 | 21 | 24 | 51265409 | 77387665 | 41395341 | 85211601 | 50153717 | 56067825 | 37443277 | 58178521 | 49806437 | 66764369 | 56251489 | 48980529 | 64215313 | 51575097 | 53271109 |
| Q9QXK3 | Copg2 | Coatomer subunit gamma-2 Mus musculus | 814.5 | 24.91 | 97681 | 12 | 15 | 40872573 | 61695881 | 28478757 | 73457185 | 41367229 | 46436433 | 26866997 | 49068777 | 35344265 | 58326049 | 47306193 | 37361617 | 52196069 | 39140489 | 39185521 |
| P61202 | Cops2 | COP9 signalosome complex subunit 2 Mus musculus | 427.81 | 25.51 | 51597 | 8 | 8 | 8871354 | 6822554 | 6067290 | 6486620.5 | 7420980.5 | 7066041.5 | 6817873 | 7915646 | 8681293 | 6311201.5 | 8859106 | 2806318 | 6373399.5 | 8243563.5 | 7115360.5 |
| O88543 | Cops3 | COP9 signalosome complex subunit 3 Mus musculus | 504.81 | 34.28 | 47832 | 10 | 10 | 13723129 | 12723765 | 11742243 | 12190972 | 12781533 | 12980569 | 11143552 | 12915070 | 13446033 | 13730878 | 14511005 | 11353115 | 13532867 | 10607540 | 12300288 |
| O88544 | Cops4 | COP9 signalosome complex subunit 4 Mus musculus | 650.88 | 33.99 | 46285 | 10 | 10 | 10610030 | 11319453 | 7651990 | 10390833 | 7450855 | 8820430 | 10195669 | 11576322 | 10077891 | 9007566 | 7217851 | 9653577 | 10214240 | 11219613 | 7927790 |
| O35864 | Cops5 | COP9 signalosome complex subunit 5 Mus musculus | 418.43 | 38.02 | 37549 | 9 | 9 | 12679536 | 13750784 | 13470223 | 12646730 | 11731257 | 10608725 | 7829446 | 13039632 | 13118319 | 11296933 | 11207419 | 12988509 | 11319573 | 13863181 | 8497076 |
| O88545 | Cops6 | COP9 signalosome complex subunit 6 Mus musculus | 83.87 | 9.57 | 35880 | 3 | 3 | 2011342.5 | 1936682.5 | 1757356.12 | 3251360 | 1788448.25 | 2573876 | 2890908 | 3501998 | 1696260.88 | 1822330 | 1838152.75 | 1559838.5 | 1948901.75 | 2015174.5 | 2180998.5 |
| Q9CZ04 | Cops7a | COP9 signalosome complex subunit 7a Mus musculus | 330.49 | 26.91 | 30224 | 5 | 5 | 1254091.88 | 881626.88 | 919515.62 | 940309.25 | 818783 | 1380102.25 | 1873357.5 | 1754560.88 | 782097.81 | 1803143.62 | 1045542 | 896291.25 | 1371584.88 | 1829811.62 | 1290439.75 |
| Q8BV13 | Cops7b | COP9 signalosome complex subunit 7b Mus musculus | 179.7 | 21.97 | 29689 | 4 | 4 | 1596074.75 | 1832160.75 | 2051897 | 1354905 | 1326802.75 | 1317764.75 | 1676015.38 | 1692496.62 | 2022012.62 | 1308041.38 | 1548799 | 1628493.75 | 1580274 | 1614473.25 | 1386344.25 |
| Q8VBV7 | Cops8 | COP9 signalosome complex subunit 8 Mus musculus | 155.15 | 22.97 | 23256 | 3 | 3 | 2836882 | 2869992.25 | 3055049.25 | 3014558.75 | 3912004 | 2919547 | 3429061.75 | 3606386.5 | 619240.56 | 3960748.5 | 2647963.75 | 4155796.5 | 3283151.5 | 3106287.5 | 3642048 |
| P61924 | Copz1 | Coatomer subunit zeta-1 Mus musculus | 326.15 | 51.41 | 20198 | 6 | 6 | 16257596 | 18866305 | 17206581 | 14601937 | 16798483 | 15719721 | 16267027 | 16044157 | 17661787 | 17031241 | 20058663 | 17477923 | 18168995 | 16735595 | 16905717 |
| P97478 | Coq7 | 5-demethoxyubiquinone hydroxylase, mitochondrial Mus musculus | 38 | 11.98 | 24042 | 1 | 1 | 170601.22 | 236639.27 | 152764.27 | 219756.44 | 170123.34 | 188320.38 | 159451.39 | 143349 | 177849.27 | 179436.06 | 161255.92 | 131202.88 | 175576.02 | 176588.16 | 152174.16 |
| Q6AY19 | Coq8b | Atypical kinase COQ8B, mitochondrial Rattus norvegicus | 34.75 | 1.89 | 58905 | 1 | 1 | 3658946.25 | 2610860.5 | 3624262.25 | 2453319 | 2308136 | 3122294.75 | 1484536.5 | 3026347.75 | 4630792 | 1396193 | 2588370 | 3509550 | 2122035 | 3363389.5 | 2638324.75 |
| O89053 | Coro1a | Coronin-1A Mus musculus | 1296.8 | 52.49 | 50989 | 20 | 21 | 234062945 | 239528401 | 267821985 | 303063969 | 299673473 | 274462945 | 392930081 | 215968305 | 188315041 | 341581313 | 341929089 | 286137185 | 293614209 | 213820881 | 404891553 |
| O89046 | Coro1b | Coronin-1B Rattus norvegicus | 128.19 | 4.96 | 53845 | 2 | 2 | 4789922 | 5263920 | 4542603 | 5334586 | 5267367 | 5036823 | 5725405 | 4090273.5 | 4164493.25 | 5340210 | 4516006 | 4831767.5 | 5094722 | 4705358.5 | 4522239 |
| Q9WUM4 | Coro1c | Coronin-1C Mus musculus | 900.89 | 43.25 | 53121 | 15 | 15 | 52956961 | 61785905 | 62616461 | 66189629 | 63500001 | 54755209 | 65506401 | 51478685 | 50907389 | 63424729 | 60985085 | 55652749 | 62228333 | 53847573 | 61115345 |
| Q9D2V7 | Coro7 | Coronin-7 Mus musculus | 719.38 | 33.41 | 100812 | 11 | 15 | 4761044 | 6848584.5 | 5650407 | 9400334 | 6425346.5 | 9674301 | 8186941 | 7588760.5 | 4289209.5 | 7051412.5 | 8253200 | 8946238 | 7225950 | 4578020 | 8238515 |
| Q9CQI6 | Cotl1 | Coactosin-like protein Mus musculus | 560.68 | 72.54 | 15944 | 11 | 11 | 82886673 | 79513065 | 77089809 | 91332793 | 108169353 | 89964905 | 79047993 | 69378473 | 59627641 | 93908465 | 110373593 | 85674705 | 91483193 | 79269257 | 101166713 |
| Q8BJ03 | Cox15 | Cytochrome c oxidase assembly protein COX15 homolog Mus musculus | 40.99 | 4.36 | 45853 | 1 | 1 | 422081.69 | 1 | 1 | 1 | 1 | 1 | 1 | 1 | 317211.31 | 1 | 1 | 1 | 311172.72 | 392891.16 | 1 |
| P56394 | Cox17 | Cytochrome c oxidase copper chaperone Mus musculus | 73.75 | 25.4 | 6784 | 1 | 1 | 853887.655 | 675136.25 | 976623.56 | 849466.06 | 914324.75 | 855314.12 | 2054855.5 | 967971.62 | 655813.06 | 1501825.12 | 1287886.12 | 1304602.12 | 1429794.5 | 798838 | 1858521.38 |
| Q9D7J4 | Cox20 | Cytochrome c oxidase assembly protein COX20, mitochondrial Mus musculus | 54.77 | 19.66 | 13163 | 1 | 1 | 399966.47 | 385341.97 | 316094.22 | 304034.22 | 344977.81 | 319861.88 | 183493.38 | 310951.38 | 273197.81 | 267268.06 | 491083.31 | 171187.28 | 223700.59 | 317257.81 | 598643.44 |
| P19783 | Cox4i1 | Cytochrome c oxidase subunit 4 isoform 1, mitochondrial Mus musculus | 215.7 | 30.77 | 19530 | 4 | 4 | 30285453 | 33025669 | 27630631 | 26359977 | 29911717 | 27798569 | 29864621 | 27529413 | 28965551 | 30185513 | 28858003 | 27037465 | 31557257 | 28329717 | 26301727 |
| P11240 | Cox5a | Cytochrome c oxidase subunit 5A, mitochondrial Rattus norvegicus | 288.69 | 38.36 | 16130 | 6 | 6 | 40878529 | 47620297 | 40817437 | 37513889 | 38771337 | 35279721 | 32005975 | 43548297 | 45468133 | 36913221 | 32122139 | 33837197 | 40824825 | 39007549 | 26683819 |
| P19536 | Cox5b | Cytochrome c oxidase subunit 5B, mitochondrial Mus musculus | 53.61 | 9.38 | 13813 | 1 | 1 | 7728957.5 | 8080394.5 | 9121866 | 9044157 | 8073188.5 | 7338581.5 | 8735322 | 7970701 | 6629896.5 | 8879628 | 8488965 | 7177254 | 7377040 | 6574613.5 | 7532560 |
| P56391 | Cox6b1 | Cytochrome c oxidase subunit 6B1 Mus musculus | 43.9 | 16.28 | 10071 | 1 | 1 | 2872200.25 | 4056435 | 2684032.25 | 2218423 | 3453565.75 | 3315658.5 | 2082919.5 | 3002068.25 | 2778321 | 2236878.5 | 1599410.5 | 2284663.75 | 2967242.75 | 2994007 | 1899282.38 |
| Q9CPQ1 | Cox6c | Cytochrome c oxidase subunit 6C Mus musculus | 67.57 | 19.74 | 8469 | 2 | 2 | 5674408 | 3656424 | 3796538 | 4099267.5 | 3276160.75 | 3247602 | 5114170.5 | 4025301.5 | 3138692.75 | 4965557.5 | 3346820.5 | 4056323 | 4487959.5 | 3961546.5 | 2805402.75 |
| P48771 | Cox7a2 | Cytochrome c oxidase subunit 7A2, mitochondrial Mus musculus | 55.83 | 12.05 | 9291 | 1 | 1 | 2024143.12 | 2778264.75 | 2558337.5 | 2066369.38 | 1934297.38 | 2112556.25 | 1667956 | 2436352.5 | 2390178.25 | 2201500 | 1749434.12 | 1992818.38 | 2259024.5 | 1891163.88 | 1984608.25 |
| O89001 | Cpd | Carboxypeptidase D Mus musculus | 285.86 | 6.25 | 152406 | 7 | 7 | 5301483 | 4425193.5 | 3982986.5 | 4140849 | 5125316.5 | 6938925 | 6277641 | 7419425.5 | 6260566.5 | 5917052 | 4396314.5 | 5032049 | 5245310 | 5387845.5 | 4855307.5 |
| Q8C166 | Cpne1 | Copine-1 Mus musculus | 608.36 | 28.36 | 58887 | 10 | 11 | 17995381 | 17383457 | 16692100 | 21040703 | 19500999 | 20580851 | 20763909 | 20206547 | 19734429 | 23906471 | 21675361 | 21673537 | 19350763 | 22531635 | 22290513 |
| Q8BT60 | Cpne3 | Copine-3 Mus musculus | 333.64 | 19.14 | 59585 | 5 | 7 | 12764879 | 9280576 | 9306238 | 10076015 | 10355059 | 9608716 | 9783219 | 10459169 | 11224380 | 11099819 | 9261786 | 9675414 | 11312369 | 9959538 | 8854409 |
| Q9DC53 | Cpne8 | Copine-8 Mus musculus | 166.02 | 6.93 | 64667 | 2 | 3 | 1 | 1 | 208111.88 | 619525.88 | 1 | 683007 | 1 | 230027.75 | 251022.44 | 1 | 138747.45 | 819394.5 | 438623.9325 | 265689.97 | 530663.81 |
| P36552 | Cpox | Oxygen-dependent coproporphyrinogen-III oxidase, mitochondrial Mus musculus | 149.92 | 7.22 | 49715 | 3 | 3 | 1503168 | 1347360.5 | 1329728.25 | 1128550.25 | 1129803.62 | 1433957.12 | 3423644.25 | 3157619 | 1434446 | 3432024 | 3203623.25 | 3139364 | 2982791.25 | 1181355.12 | 1388669.5 |
| Q9WVJ3 | Cpq | Carboxypeptidase Q Mus musculus | 159.27 | 9.79 | 51813 | 3 | 3 | 4235704.5 | 5433789.5 | 3963128 | 4633763.5 | 4864550 | 3552695.75 | 3248951.75 | 3824109 | 4722842 | 3420599 | 3064033 | 3378491 | 4432056.5 | 3971907 | 2850809 |
| Q9EPU4 | Cpsf1 | Cleavage and polyadenylation specificity factor subunit 1 Mus musculus | 280.91 | 4.86 | 160818 | 6 | 6 | 3494121.75 | 2602098.25 | 2750698.5 | 3393683.5 | 3468244.5 | 3238063 | 3605759.25 | 3163875.25 | 2665237.25 | 3587764 | 2536933.5 | 3004875.25 | 2928182.5 | 2829640.75 | 3061903.5 |
| O35218 | Cpsf2 | Cleavage and polyadenylation specificity factor subunit 2 Mus musculus | 347.87 | 11.76 | 88383 | 6 | 6 | 2244283.75 | 3929254 | 3872361.25 | 2699875 | 3522009 | 1838706.38 | 3539890.5 | 2565132 | 2905061 | 2193618.5 | 3349618.5 | 3001149.5 | 3153425.75 | 3471344.75 | 3295727.5 |
| Q9QXK7 | Cpsf3 | Cleavage and polyadenylation specificity factor subunit 3 Mus musculus | 176.93 | 8.48 | 77505 | 4 | 4 | 2630602.25 | 2300776.5 | 1728431 | 2361033.5 | 2241506 | 2666005 | 3166372 | 2808851 | 2102760.25 | 2430932.5 | 2508215 | 2727896 | 2275732.25 | 2906532.25 | 2637613.25 |
| Q6NVF9 | Cpsf6 | Cleavage and polyadenylation specificity factor subunit 6 Mus musculus | 453.08 | 17.06 | 59153 | 7 | 7 | 17312125 | 18308605 | 15433369 | 15300309 | 17152403 | 13926566 | 13411282 | 13718029 | 18235747 | 13430121 | 16637393 | 14411625 | 14533537 | 15553277 | 13672073 |
| Q5XI29 | Cpsf7 | Cleavage and polyadenylation specificity factor subunit 7 Rattus norvegicus | 210.07 | 13.42 | 51073 | 5 | 5 | 3155922 | 3872699 | 3390567.5 | 3668620.5 | 3314346.5 | 2595608.5 | 3553217 | 3594615.5 | 3212444 | 3165935 | 2491291.25 | 3390882.5 | 3014212 | 3489903 | 2268425 |
| P32198 | Cpt1a | Carnitine O-palmitoyltransferase 1, liver isoform Rattus norvegicus | 39.87 | 2.2 | 88125 | 1 | 1 | 220662.52 | 263728.84 | 292934.12 | 1 | 1 | 294054.88 | 203736.91 | 365982.03 | 570653.56 | 218630.44 | 239993.28 | 269995 | 123269.76 | 273285.09 | 263551.5 |
| P52825 | Cpt2 | Carnitine O-palmitoyltransferase 2, mitochondrial Mus musculus | 194.82 | 10.49 | 73981 | 5 | 5 | 9623966 | 7989849 | 7172153.5 | 8663070 | 8746708 | 8962298 | 7196045 | 7507622 | 7925918 | 7633857 | 6946224.5 | 8008253.5 | 6701052 | 7814660 | 4652081.5 |
| P15337 | Creb1 | Cyclic AMP-responsive element-binding protein 1 Rattus norvegicus | 88.21 | 10.56 | 36633 | 2 | 2 | 1094556.12 | 1386957.88 | 1462360.62 | 1320651.38 | 1367049.38 | 1524968.78 | 2491635.25 | 1068306 | 1147572.12 | 1392361.75 | 1324102.88 | 1247206.12 | 1149882.88 | 1031765.81 | 1188239.423 |
| O88668 | Creg1 | Protein CREG1 Mus musculus | 112.73 | 13.18 | 24452 | 2 | 2 | 5879620 | 4904086 | 3288679.75 | 5032750 | 5737065 | 6132261.5 | 4399349 | 5841980 | 4668494 | 4652700.5 | 4497737.5 | 4576369.5 | 4955055 | 5150101 | 4158699.5 |
| Q4V7F2 | Creld1 | Cysteine-rich with EGF-like domain protein 1 Rattus norvegicus | 85.57 | 7.38 | 45697 | 2 | 2 | 1567646.38 | 1949247.25 | 1451439.5 | 1601468.5 | 2347912.5 | 2633014.75 | 1459401.25 | 1702218.25 | 1955275.38 | 1530741.62 | 1789298.25 | 1415723.88 | 1092303.62 | 1779298.25 | 1628175.75 |
| Q9CYA0 | Creld2 | Cysteine-rich with EGF-like domain protein 2 Mus musculus | 139.69 | 7.43 | 38220 | 2 | 2 | 1209143.38 | 1328185.25 | 1600187 | 1328698.88 | 1718282 | 1681052.62 | 1128549.88 | 1396220.75 | 1543679.25 | 1150499 | 1532474.12 | 990312.19 | 1324098 | 1480360.62 | 1262380.5 |
| P63254 | Crip1 | Cysteine-rich protein 1 Mus musculus | 51.61 | 12.99 | 8550 | 1 | 1 | 1382662.62 | 567465.25 | 805599.19 | 1115996.62 | 734611.19 | 1 | 1384036.62 | 523232.53 | 595983.81 | 1305874.38 | 840987.38 | 853167.81 | 806300.56 | 729188.5 | 1275978.62 |
| Q63768 | Crk | Adapter molecule crk Rattus norvegicus | 200.61 | 14.8 | 33845 | 4 | 4 | 5765189 | 3860508.5 | 3913847.25 | 6410894 | 6449763 | 4316430.5 | 3676538.5 | 3943834 | 2637846.5 | 3663227.5 | 5093099 | 3668453 | 3967423.75 | 4366578.5 | 3285581.5 |
| Q9Z2L7 | Crlf3 | Cytokine receptor-like factor 3 Mus musculus | 141.76 | 10.86 | 49559 | 3 | 3 | 970408.38 | 1690071.75 | 2738421.25 | 1509731.62 | 1445621.5 | 1119275.62 | 2594349 | 1597194 | 2169990.25 | 1870202.218 | 2610753 | 2409065 | 1059650.88 | 2426997.25 | 2598059.75 |
| P63154 | Crnkl1 | Crooked neck-like protein 1 Mus musculus | 108.38 | 4.49 | 83416 | 3 | 3 | 2482272.75 | 2071717.75 | 2441203.75 | 3190848 | 2019391.75 | 2310142.5 | 1256227.75 | 2066481.25 | 2559921 | 1654012.5 | 1472084.5 | 1935470 | 2119509.25 | 1949526.88 | 1801299.75 |
| Q99KP3 | Cryl1 | Lambda-crystallin homolog Mus musculus | 305.99 | 25.39 | 35209 | 5 | 5 | 4758469.5 | 4669841 | 5362881.5 | 5612916 | 4733707 | 4952750 | 3994264.5 | 5966513 | 4993233 | 4645804.5 | 4110123.75 | 4914654 | 4413522.5 | 4859697 | 4633234 |
| P47199 | Cryz | Quinone oxidoreductase Mus musculus | 44.42 | 6.65 | 35269 | 1 | 1 | 1067594.25 | 1308112.12 | 1488117.25 | 1360542.88 | 1056024.75 | 1550064.12 | 1889349.38 | 1563990.12 | 1663548.75 | 1596370.88 | 2305991.75 | 1844169 | 1825380.5 | 1225050.12 | 2296096.75 |
| Q921W4 | Cryzl1 | Quinone oxidoreductase-like protein 1 Mus musculus | 83.52 | 10.06 | 38725 | 2 | 2 | 1 | 1 | 1485640.38 | 1012541 | 968118.94 | 1 | 1234969.25 | 1632423.38 | 1 | 1152267.62 | 1781154.62 | 2248605.75 | 1326648 | 1720535.968 | 1525735.5 |
| Q9CZU6 | Cs | Citrate synthase, mitochondrial Mus musculus | 782.65 | 42.67 | 51737 | 15 | 15 | 86699297 | 81383289 | 77836641 | 70827313 | 76641065 | 81424785 | 96928641 | 76114769 | 75742705 | 88418297 | 77682241 | 72050673 | 79281577 | 67767689 | 81686177 |
| P18395 | Csde1 | Cold shock domain-containing protein E1 Rattus norvegicus | 860.82 | 25.56 | 88895 | 18 | 18 | 26972421 | 24937707 | 24642163 | 25598829 | 21724057 | 22477435 | 23705071 | 26044269 | 26832715 | 23867501 | 25169655 | 23465053 | 21964621 | 22399883 | 21344965 |
| Q9ERK4 | Cse1l | Exportin-2 Mus musculus | 1738.81 | 37.59 | 110455 | 28 | 28 | 55513865 | 41289213 | 38080361 | 32701209 | 33126635 | 35957885 | 29483291 | 46621157 | 49940689 | 33417469 | 44366357 | 41208409 | 40182609 | 49548657 | 36589793 |
| P09581 | Csf1r | Macrophage colony-stimulating factor 1 receptor Mus musculus | 356.57 | 12.28 | 109179 | 7 | 8 | 10909409 | 11710867 | 10370802 | 7828121 | 10516187 | 11529666 | 11224981 | 10624267 | 10667697 | 9636939 | 10605659 | 10688475 | 11735289 | 11399131 | 9540347 |
| P32577 | Csk | Tyrosine-protein kinase CSK Rattus norvegicus | 339.91 | 15.78 | 50746 | 6 | 6 | 9015732 | 8588454 | 9982495 | 8774781 | 9298991 | 9314983 | 10910678 | 8904305 | 9067794 | 10544830 | 10863379 | 10913110 | 10406570 | 8939033 | 11029062 |
| P48729 | CSNK1A1 | Casein kinase I isoform alpha Homo sapiens | 86.86 | 9.2 | 38915 | 2 | 2 | 536957.25 | 1 | 1 | 1 | 1 | 205548.03 | 509977.53 | 1 | 1 | 332800.56 | 259250.8575 | 265576.84 | 161625.03 | 286007.22 | 323794.34 |
| Q06486 | Csnk1d | Casein kinase I isoform delta Rattus norvegicus | 162.38 | 9.4 | 47316 | 3 | 3 | 408847.41 | 930260.62 | 637895.75 | 582409.04 | 352632.38 | 768601.06 | 487697.84 | 777727.88 | 828873.88 | 801460.62 | 775535.62 | 383308.12 | 848936.12 | 673128.62 | 795773.25 |
| P19139 | Csnk2a1 | Casein kinase II subunit alpha Rattus norvegicus | 464.53 | 30.18 | 45073 | 9 | 9 | 18715861 | 14206223 | 12980470 | 13937643 | 13187150 | 12014377 | 15129864 | 15502479 | 16753336 | 15973507 | 15065627 | 14996028 | 15530108 | 14203491 | 17933531 |
| O54833 | Csnk2a2 | Casein kinase II subunit alpha' Mus musculus | 259.18 | 19.71 | 41215 | 5 | 5 | 4948351 | 3063600.5 | 4773701.5 | 3121328 | 3421565.5 | 3834383.25 | 2883361.75 | 4558894.5 | 4546221 | 3064577.25 | 4025516.75 | 4418320.5 | 3295765 | 4148148 | 4899870.5 |
| P67871 | Csnk2b | Casein kinase II subunit beta Mus musculus | 83.12 | 15.81 | 24942 | 2 | 2 | 3663886.5 | 4403108 | 3072081.25 | 3583049.5 | 4103227 | 3873420.75 | 3206806.75 | 4034069.25 | 4636860.5 | 3822581 | 3808751.75 | 3119350 | 4046850 | 4338823 | 3127441 |
| Q99388 | Csprs | Component of Sp100-rs Mus musculus | 91.15 | 8.65 | 24061 | 1 | 2 | 2141751.5 | 2036986.12 | 2158798.5 | 1540366.88 | 1584585.62 | 1841822.88 | 1266251.12 | 2185337.5 | 2387486.75 | 1095718.12 | 1623784.62 | 1730275.88 | 1179876 | 2183827.25 | 1396544.75 |
| P97315 | Csrp1 | Cysteine and glycine-rich protein 1 Mus musculus | 258.97 | 33.16 | 20583 | 5 | 5 | 1776077.75 | 1699707.62 | 1975467.25 | 2826553.5 | 1726693.5 | 1652419.25 | 5300689 | 1946070 | 1346030.88 | 4318256 | 3604607.5 | 1659365 | 3798009 | 1805884 | 4083338.75 |
| P14841 | Cst3 | Cystatin-C Rattus norvegicus | 37.83 | 7.86 | 15437 | 1 | 1 | 1246088.62 | 894169.31 | 927652.62 | 1088590.5 | 1185652.5 | 915518.19 | 1546501.62 | 997251.06 | 888087.69 | 1174474.5 | 750245.69 | 698831.44 | 783306.94 | 1269855.75 | 768149.75 |
| Q62426 | Cstb | Cystatin-B Mus musculus | 145.46 | 57.14 | 11046 | 3 | 3 | 7521179 | 7137837.5 | 6904083 | 12055766 | 6465746.5 | 3764908.75 | 5119400 | 4618857 | 3382433.5 | 4055859 | 4358063 | 4224369.5 | 5296863.5 | 7141643.5 | 5777587 |
| Q5BJQ6 | Cstf1 | Cleavage stimulation factor subunit 1 Rattus norvegicus | 93.52 | 3.71 | 48382 | 1 | 1 | 928786.5625 | 930250.25 | 920181.12 | 1003143.5 | 861571.38 | 946692.25 | 952646.56 | 865457.88 | 935715 | 929423.69 | 1280309.12 | 1072393.25 | 1041519.94 | 1193535.38 | 703514.31 |
| Q8BIQ5 | Cstf2 | Cleavage stimulation factor subunit 2 Mus musculus | 167.32 | 14.31 | 61341 | 4 | 4 | 1416987.5 | 738162.56 | 917080 | 2133209 | 1123908.62 | 1245420.5 | 1068155.25 | 1580760.75 | 1253530.75 | 1324903.62 | 1088404.62 | 1871115.75 | 2278308.25 | 884681.31 | 2350538.75 |
| Q99LI7 | Cstf3 | Cleavage stimulation factor subunit 3 Mus musculus | 372.6 | 13.81 | 82877 | 7 | 7 | 2502103 | 2949152.25 | 3310061.75 | 3082721.25 | 2948490 | 2783401.5 | 4172868.75 | 3038672.25 | 2925210.25 | 3400340.25 | 2917828.25 | 4128653.25 | 3488288.5 | 2805115 | 4029529.75 |
| Q8R311 | Ctage5 | Endoplasmic reticulum export factor CTAGE5 Mus musculus | 84.57 | 3.47 | 87718 | 2 | 2 | 606834.88 | 705294.5 | 929798.62 | 774351.5 | 855478 | 665345.56 | 870676.56 | 840798.75 | 605560.56 | 859189.25 | 730664.81 | 689362.38 | 696332.5 | 735150.25 | 837402.69 |
| O88712 | Ctbp1 | C-terminal-binding protein 1 Mus musculus | 357.62 | 21.54 | 47745 | 4 | 7 | 6084126.5 | 6867470.5 | 7400646 | 7924728 | 7334426.5 | 4653719 | 10668258 | 6802740 | 7925996 | 10158226 | 10567935 | 9932028 | 8020798 | 7461330 | 11083005 |
| P56546 | Ctbp2 | C-terminal-binding protein 2 Mus musculus | 495.05 | 36.63 | 48957 | 9 | 12 | 14222561 | 12440207 | 12064810 | 16314195 | 15451092 | 10794748 | 17882825 | 13722684 | 13085454 | 14781814 | 15716719 | 12685787 | 16125885 | 14783167 | 17610053 |
| Q8R242 | Ctbs | Di-N-acetylchitobiase Mus musculus | 84.47 | 8.2 | 41295 | 2 | 2 | 2802038 | 3222664.25 | 2995471.75 | 958706.62 | 2494720.155 | 2453756 | 700382.81 | 2772028 | 3115422.75 | 693103.06 | 1790968.5 | 2099055 | 2673635 | 2642120.5 | 2186283.5 |
| Q5SUQ9 | Ctc1 | CST complex subunit CTC1 Mus musculus | 89.28 | 3.3 | 134028 | 2 | 2 | 1 | 386601.38 | 1 | 315971.44 | 479609.97 | 250207.22 | 470622.81 | 490987.12 | 649958.06 | 546377.81 | 800005.69 | 628332.25 | 299729.72 | 319988.97 | 675966.75 |
| Q61164 | Ctcf | Transcriptional repressor CTCF Mus musculus | 123.49 | 3.67 | 83745 | 2 | 2 | 589231.62 | 706246.56 | 1244668.5 | 667464.88 | 1033430.69 | 945641 | 1042374.62 | 1069930.88 | 820117.88 | 1533062.75 | 1735759.88 | 1759515 | 635508.12 | 1084373.75 | 1257604.5 |

| | | | | | | | | | | | | | | | | | | | | | | |
|---|---|---|---|---|---|---|---|---|---|---|---|---|---|---|---|---|---|---|---|---|---|---|
| Q8VCN5 | Cth | Cystathionine gamma-lyase Mus musculus | 688.3 | 43.47 | 43567 | 13 | 13 | 41636849 | 40732541 | 49172553 | 51631009 | 46581893 | 48239193 | 44108457 | 45834553 | 48384565 | 50668917 | 51399573 | 39141849 | 39288621 | 37906089 | 45108657 |
| Q9CWL8 | Ctnnbl1 | Beta-catenin-like protein 1 Mus musculus | 289.25 | 13.32 | 64980 | 6 | 6 | 1876150.25 | 2011940.12 | 2942662.5 | 2278631.5 | 3450300 | 2306937.5 | 1767519 | 2900048.25 | 2350361.25 | 2553605.25 | 1629395.75 | 1998030.62 | 1832747.75 | 3420499.5 | 3004257 |
| P30999 | Ctnnd1 | Catenin delta-1 Mus musculus | 503.48 | 16.63 | 104925 | 10 | 10 | 3632788.5 | 5114385.5 | 6365331 | 7988858 | 5750182 | 6690540 | 6047871.5 | 4291262.5 | 5077226.5 | 6312568 | 4913604 | 5969753.5 | 6232151.5 | 4585982.5 | 5829989.5 |
| P70698 | Ctps1 | CTP synthase 1 Mus musculus | 717.68 | 27.41 | 66682 | 11 | 12 | 11293901 | 12770289 | 11429613 | 12607571 | 10775793 | 11497500 | 13185413 | 11892990 | 11944892 | 12215035 | 10013294 | 11457242 | 11958294 | 12335984 | 12041189 |
| P70303 | Ctps2 | CTP synthase 2 Mus musculus | 80.46 | 3.75 | 65514 | 1 | 2 | 881614.19 | 1296648.12 | 1375734.75 | 1357951.38 | 534595.62 | 1020651.88 | 1047655.75 | 1302147.12 | 2097673 | 1096010.5 | 1164075.62 | 1101520.75 | 1169386.62 | 1283433.5 | 930744.38 |
| P16675 | Ctsa | Lysosomal protective protein Mus musculus | 1103.64 | 36.71 | 53844 | 14 | 16 | 144273681 | 156101457 | 125186585 | 127415905 | 147836225 | 160673233 | 121686489 | 148432561 | 169038641 | 123056625 | 144753041 | 117974913 | 139963601 | 159121777 | 125056673 |
| P10605 | Ctsb | Cathepsin B Mus musculus | 1317.47 | 67.26 | 37280 | 19 | 20 | 334932353 | 395148609 | 352905249 | 324442689 | 350109057 | 331361185 | 246288065 | 367721953 | 402474017 | 266142913 | 431975041 | 299859201 | 357314625 | 370529249 | 287160737 |
| P97821 | Ctsc | Dipeptidyl peptidase 1 Mus musculus | 595.33 | 16.67 | 52376 | 7 | 7 | 49376273 | 52577253 | 45175817 | 45652129 | 52748553 | 44029841 | 30403867 | 40893277 | 42000561 | 30968901 | 39618785 | 36447381 | 45088625 | 42878961 | 35434073 |
| P18242 | Ctsd | Cathepsin D Mus musculus | 1308.71 | 65.37 | 44954 | 18 | 24 | 406401761 | 489922369 | 425776257 | 393210113 | 412787073 | 439375361 | 364178593 | 459516065 | 465317025 | 362879137 | 369210369 | 370707905 | 455374593 | 474837761 | 330233249 |
| P49935 | Ctsh | Pro-cathepsin H Mus musculus | 121.9 | 8.41 | 37170 | 2 | 2 | 1658226.88 | 4813508.5 | 3965152 | 3522042 | 4001618.5 | 2868396.5 | 2988841 | 3019015 | 3674507.25 | 2376675.5 | 1568366.38 | 3049570 | 4054391.25 | 3879002.75 | 3066697.5 |
| P06797 | Ctsl | Cathepsin L1 Mus musculus | 294.45 | 37.72 | 37547 | 6 | 6 | 13943462 | 17121151 | 17095199 | 10881181 | 14913645 | 14542510 | 11980905 | 16239980 | 19070405 | 9666820 | 16255895 | 15210764 | 15113552 | 17594797 | 14229834 |
| O70370 | Ctss | Cathepsin S Mus musculus | 809.73 | 49.12 | 38475 | 9 | 10 | 238743409 | 309175649 | 274737185 | 216478641 | 242877905 | 184963473 | 123772145 | 204258961 | 256792401 | 144348049 | 159074049 | 197928353 | 230172657 | 257985185 | 186827841 |
| Q9WUU7 | Ctsz | Cathepsin Z Mus musculus | 745.53 | 45.75 | 33996 | 12 | 12 | 342145953 | 436895265 | 362891745 | 375083137 | 406212577 | 357058721 | 307675553 | 399239489 | 438017537 | 313883777 | 408088449 | 320584641 | 406117377 | 398229409 | 298976737 |
| Q99LJ0 | Cttnbp2nl | CTTNBP2 N-terminal-like protein Mus musculus | 157.93 | 6.58 | 69841 | 2 | 2 | 370077.53 | 532828.12 | 236185.23 | 429336.66 | 462252.47 | 511236.41 | 694464.06 | 548655.62 | 254511.7 | 397691.88 | 615491.38 | 569580.5 | 427895.41 | 574348.12 | 383872.66 |
| Q3U308 | Ctu2 | Cytoplasmic tRNA 2-thiolation protein 2 Mus musculus | 54.55 | 3.7 | 56105 | 1 | 1 | 1 | 1 | 1 | 1 | 270970.25 | 234649.09 | 270152.84 | 1 | 1 | 1 | 324683.66 | 1 | 1 | 1 | 391401.56 |
| Q9WTX6 | Cul1 | Cullin-1 Mus musculus | 508.56 | 14.69 | 89692 | 10 | 10 | 9356674 | 8197263 | 7475768 | 8358356 | 10032447 | 10148699 | 6894816 | 10197417 | 9848283 | 8506364 | 12365513 | 10492369 | 9581891 | 10079986 | 12049857 |
| Q9D4H8 | Cul2 | Cullin-2 Mus musculus | 222.18 | 7.92 | 86877 | 5 | 5 | 1236850.5 | 1862539.88 | 1368809.25 | 1538803.75 | 1462571.5 | 1604546.5 | 2215151.5 | 2744167.5 | 2896369.25 | 1995367.25 | 1995650.75 | 2344885.75 | 1822770.25 | 2042566.5 | 2058204.75 |
| B5DF89 | Cul3 | Cullin-3 Rattus norvegicus | 415.77 | 13.41 | 88932 | 7 | 7 | 4959384.5 | 5196358 | 5075169.5 | 4650876 | 4922087 | 6304337.5 | 3692358 | 5129359 | 6231962 | 3682604.25 | 4069174.25 | 6223634 | 4924344.5 | 5770253 | 4353925 |
| Q3TCH7 | Cul4a | Cullin-4A Mus musculus | 193.37 | 5.67 | 87753 | 2 | 4 | 2419670.5 | 2496802.75 | 2800189 | 2487509 | 2312425.5 | 2753193.5 | 2618633.25 | 2697458.75 | 3130263 | 2110428.75 | 2470030.25 | 2471970.5 | 2726756.25 | 2439749.25 | 2348795.5 |
| A2A432 | Cul4b | Cullin-4B Mus musculus | 489.3 | 12.06 | 110699 | 7 | 9 | 5683032 | 3562498 | 5907558 | 3791890.75 | 3234073 | 2940669.75 | 2811338.75 | 5717354 | 6757709.5 | 3451041.25 | 5220386 | 5732512.5 | 4025807 | 5262784.5 | 6240267 |
| Q9D5V5 | Cul5 | Cullin-5 Mus musculus | 37.15 | 1.28 | 90974 | 1 | 1 | 257088.89 | 1 | 1 | 1 | 455063.47 | 344192.38 | 385769.25 | 1 | 1 | 397345.88 | 360900.53 | 422130.25 | 395248.81 | 283160.56 | 512461.69 |
| Q6MGD0 | Cuta | Protein CutA Rattus norvegicus | 181.04 | 29.94 | 18659 | 4 | 4 | 7819751 | 10969266 | 9721237 | 9869074 | 8752632 | 8195076 | 8559804 | 9581994 | 10986253 | 8194060 | 6261889.5 | 9033719 | 9763132 | 9994875 | 5777850 |
| Q5BJP2 | Cwc15 | Spliceosome-associated protein CWC15 homolog Rattus norvegicus | 173.56 | 16.16 | 26638 | 3 | 3 | 585857.06 | 3540086 | 2168444.25 | 2685335 | 3972490.25 | 3156038.5 | 2512284 | 3160186.25 | 2258822 | 3475213.75 | 3704789.75 | 3032779.5 | 2264693 | 2974401 | 3900220.5 |
| Q8C5N3 | Cwc22 | Pre-mRNA-splicing factor CWC22 homolog Mus musculus | 319.04 | 9.69 | 104773 | 6 | 6 | 3895319.5 | 4046651.25 | 1822862 | 3242381 | 2941762.5 | 2707296.5 | 1375473.75 | 3370121 | 4480906 | 2720466 | 1434224 | 3394846.5 | 2492222.5 | 3680913 | 1879319.5 |
| Q8CI33 | Cwf19l1 | CWF19-like protein 1 Mus musculus | 37.67 | 2.23 | 60193 | 1 | 1 | 1 | 1 | 1 | 1 | 1 | 1 | 1 | 1 | 1 | 1 | 466385.22 | 572857.725 | 713951.5 | 632285.62 | 478808.56 |
| Q9CQX2 | Cyb5b | Cytochrome b5 type B Mus musculus | 328.72 | 58.22 | 16318 | 5 | 5 | 13836063 | 8341799.5 | 10466041 | 13242618 | 11826971 | 12768556 | 8256498 | 9949392 | 9156796 | 10919889 | 9949127 | 9609423 | 11025258 | 10307903 | 7338011 |
| Q9DB73 | Cyb5r1 | NADH-cytochrome b5 reductase 1 Mus musculus | 274.09 | 25.57 | 34135 | 6 | 6 | 3272358.25 | 3996759 | 3589742.5 | 2439730 | 2806576.75 | 3718205.75 | 5921362 | 4036487.75 | 2860762 | 3411060 | 4072120.25 | 4000148.5 | 2377853.5 | 2918590.5 | 5111571.5 |
| Q9DCN2 | Cyb5r3 | NADH-cytochrome b5 reductase 3 Mus musculus | 705.95 | 45.18 | 34128 | 11 | 11 | 30700057 | 30305555 | 27908413 | 27498421 | 27467849 | 30559657 | 36595873 | 31887037 | 25538523 | 34970977 | 25337421 | 28651017 | 30294609 | 26959211 | 28986053 |
| Q61462 | Cyba | Cytochrome b-245 light chain Mus musculus | 136.89 | 7.29 | 20748 | 3 | 3 | 10207428 | 9629593 | 6990939 | 6998513.5 | 9168180 | 9030436 | 8567360 | 7124284 | 9498274 | 9023436 | 9287508 | 7798262 | 8459555 | 7243880 | 8370889.5 |
| Q61093 | Cybb | Cytochrome b-245 heavy chain Mus musculus | 666.11 | 30 | 65305 | 12 | 12 | 17238975 | 9279005 | 8931859 | 7187517 | 11210569 | 13629930 | 9403499 | 11130708 | 13779002 | 9778411 | 9081845 | 8245167.5 | 8042340 | 12008591 | 10933860 |
| Q3TYS2 | Cybc1 | Cytochrome b-245 chaperone 1 Mus musculus | 113.78 | 14.97 | 20921 | 3 | 3 | 4608574 | 2390832 | 2792405 | 1635330.25 | 2396028.5 | 2591553.25 | 2684360.5 | 2163934 | 2623659.75 | 2414485.25 | 3140217.25 | 2549667.5 | 2325239 | 2522696.5 | 3733787.5 |
| Q9D0M3 | Cyc1 | Cytochrome c1, heme protein, mitochondrial Mus musculus | 406.16 | 52.62 | 35328 | 9 | 9 | 15720952 | 12139899 | 12626407 | 10599710 | 13279112 | 14583336 | 12521733 | 16594440 | 15395325 | 10821960 | 13122790 | 15634693 | 14391525 | 12864305 | 12675628 |
| P62897 | Cycs | Cytochrome c, somatic Mus musculus | 296.2 | 55.24 | 11605 | 6 | 6 | 24296731 | 23385439 | 24839441 | 24829385 | 23277525 | 20800083 | 29902589 | 20825545 | 20168049 | 26616501 | 21760507 | 21760787 | 24166387 | 20784477 | 23410113 |
| Q7TMB8 | Cyfip1 | Cytoplasmic FMR1-interacting protein 1 Mus musculus | 1945.15 | 33.2 | 145241 | 22 | 37 | 53762173 | 46518805 | 46904561 | 42789213 | 44979477 | 49911569 | 49160321 | 51569785 | 58897345 | 50120113 | 60785341 | 55906693 | 49154529 | 55226257 | 59098337 |
| Q5SQX6 | Cyfip2 | Cytoplasmic FMR1-interacting protein 2 Mus musculus | 846.05 | 16.04 | 145659 | 3 | 17 | 14203615 | 13260923 | 13543792 | 11617989 | 9311673 | 12377857 | 19457649 | 15262736 | 17000729 | 17117897 | 19430573 | 16604685 | 13043795 | 15223705 | 20191151 |
| Q8BKE6 | Cyp20a1 | Cytochrome P450 20A1 Mus musculus | 218.71 | 17.32 | 52149 | 5 | 5 | 4086177.5 | 3415466 | 3399480.25 | 3047147 | 2857976.5 | 3315346.25 | 2728064.5 | 3322913.5 | 3827579 | 2499923 | 3207513.75 | 3448242.75 | 2746650.25 | 3553343 | 3473510 |
| Q9DBG1 | Cyp27a1 | Sterol 26-hydroxylase, mitochondrial Mus musculus | 148.06 | 9.19 | 60720 | 3 | 3 | 1309922.5 | 1143625.12 | 882316.56 | 749493 | 1021339.295 | 1101887.88 | 1119589.5 | 1261152.38 | 1203362.25 | 671702.25 | 1411813.38 | 1067840 | 1108913.12 | 1180143.62 | 1111370.62 |
| Q64654 | Cyp51a1 | Lanosterol 14-alpha demethylase Rattus norvegicus | 316.36 | 19.09 | 56707 | 7 | 7 | 3869052.75 | 4539243.5 | 4142854 | 2842192 | 3333263.5 | 2904624.5 | 2871803 | 3582716.75 | 3353921.5 | 2578145.25 | 3171969.5 | 3918709.25 | 3394481 | 4178745.75 | 3292459 |
| Q80YW0 | Cyth4 | Cytohesin-4 Mus musculus | 232.17 | 12.21 | 45285 | 4 | 4 | 5024233 | 4180476 | 3568252.5 | 3380212.5 | 4476086 | 4136396.5 | 828881 | 3963041 | 3871953.5 | 3091558.5 | 4973013.5 | 4854769 | 2102547.75 | 2235672.5 | 3378019.25 |
| P98078 | Dab2 | Disabled homolog 2 Mus musculus | 1579.37 | 45.04 | 82312 | 9 | 22 | 33908749 | 31361015 | 43650805 | 33180885 | 30821723 | 30817559 | 43306921 | 40136981 | 35062329 | 31336175 | 44797533 | 46747857 | 37451257 | 35115013 | 43923221 |
| P61804 | Dad1 | Dolichyl-diphosphooligosaccharide--protein glycosyltransferase subunit DAD1 Mus mus | 166.75 | 28.32 | 12497 | 3 | 3 | 13215215 | 13330451 | 11556073 | 10955833 | 15067723 | 13719991 | 12542960 | 11417371 | 13379887 | 14853337 | 14246395 | 11779221 | 14760191 | 13245293 | 12322441 |
| Q9ER88 | Dap3 | 28S ribosomal protein S29, mitochondrial Mus musculus | 264.87 | 20.46 | 44699 | 5 | 5 | 3156793 | 4620741.5 | 4832820.5 | 2470616 | 4198174.5 | 3616362 | 4180373.5 | 4495172.5 | 4356438 | 3824935.75 | 3456196.75 | 4105077 | 3654106.75 | 4007355.5 | 3256915.5 |
| Q922B2 | Dars | Aspartate--tRNA ligase, cytoplasmic Mus musculus | 1050.3 | 42.51 | 57147 | 19 | 19 | 66303153 | 75627097 | 70327633 | 64611905 | 65696493 | 69791009 | 60393625 | 73637657 | 82591473 | 63456029 | 70487153 | 72640617 | 66463701 | 70335593 | 63753177 |
| Q8BIP0 | Dars2 | Aspartate--tRNA ligase, mitochondrial Mus musculus | 111.11 | 4.59 | 74102 | 3 | 3 | 2521803.75 | 4048034 | 3894850.25 | 2972283.5 | 4086272.75 | 3788285.5 | 4259238 | 4226312.5 | 3972648.5 | 4551036.5 | 3939805 | 3834079 | 4065305.75 | 2394650.25 | 2891477 |
| Q9JII5 | Dazap1 | DAZ-associated protein 1 Mus musculus | 250.56 | 15.27 | 43214 | 4 | 4 | 3752942 | 5246539 | 3905766 | 5174724 | 5566712 | 5877685.5 | 5111768 | 5022576 | 4424411 | 3149179.75 | 4850857.5 | 4744109 | 5222840 | 4985551 | 4696101.5 |
| P11030 | Dbi | Acyl-CoA-binding protein Rattus norvegicus | 144.97 | 39.08 | 10027 | 2 | 2 | 8975738 | 7477482 | 6957561 | 8622616 | 7808447.5 | 6320285 | 6360749 | 6302452 | 4650732.5 | 8723353 | 5720608 | 6389145.5 | 7048027 | 6013892.5 | 5792070.5 |
| Q62418 | Dbnl | Drebrin-like protein Mus musculus | 587.36 | 24.08 | 48700 | 7 | 7 | 8094816.5 | 7775421.5 | 11942636 | 11969308 | 11536735 | 7922527 | 14366889 | 8314026.5 | 7863387.5 | 11779187 | 12030110 | 11235085 | 11142327 | 8241393 | 15493847 |
| Q923B1 | Dbr1 | Lariat debranching enzyme Mus musculus | 53.22 | 1.82 | 62289 | 1 | 1 | 1 | 244158.84 | 1 | 1 | 196371.23 | 322609.66 | 211480.2 | 185808 | 181101.11 | 233421.28 | 242298.66 | 223303.38 | 104564.19 | 177529.66 | 131258.75 |
| P53395 | Dbt | Lipoamide acyltransferase component of branched-chain alpha-keto acid dehydrogenas | 201.85 | 10.79 | 53247 | 4 | 4 | 934585.25 | 1303108.5 | 1277103.75 | 1273488.12 | 1362557.25 | 1402114.75 | 2359273.5 | 1484669.5 | 1433422.25 | 1512868 | 812452.94 | 1735044.25 | 1600530.12 | 1197393.5 | 1292173.75 |
| Q80TR8 | Dcaf1 | DDB1- and CUL4-associated factor 1 Mus musculus | 36.68 | 0.93 | 168931 | 1 | 1 | 116573.26 | 298595.81 | 200191.31 | 193877.98 | 140019.59 | 175267.56 | 153276.98 | 248155.2 | 265309.38 | 233612.88 | 134621.59 | 242754.61 | 266542.81 | 202670.2 | 196846.58 |
| Q5U2M6 | Dcaf8 | DDB1- and CUL4-associated factor 8 Rattus norvegicus | 36.78 | 1.35 | 66156 | 1 | 1 | 1 | 1 | 724706.62 | 1 | 296630.31 | 522535.88 | 487690.56 | 480638.1325 | 514366.5 | 397959.59 | 378581.66 | 479846.97 | 302832.34 | 521968.69 | 652189.62 |
| Q8BHC4 | Dcakd | Dephospho-CoA kinase domain-containing protein Mus musculus | 283.83 | 28.57 | 26476 | 6 | 6 | 5708384 | 4926203 | 4425020.5 | 4603862.5 | 3977113.5 | 4392350 | 5871942 | 5148057 | 3875011 | 3638414 | 5901837.5 | 5575547 | 4503115 | 4943977.5 | 5187808 |
| Q9DAR7 | Dcps | m7GpppX diphosphatase Mus musculus | 651.43 | 49.41 | 38988 | 13 | 13 | 12611214 | 10769439 | 11305334 | 11576916 | 13026258 | 11389765 | 11851803 | 10725545 | 10052820 | 11827936 | 11820401 | 11102310 | 11047740 | 9228161 | 11838747 |
| O08788 | Dctn1 | Dynactin subunit 1 Mus musculus | 395.37 | 9.76 | 141676 | 7 | 7 | 4209534 | 4921418 | 3515108.75 | 6853147 | 3822319.25 | 4966408 | 7151055 | 4241875.5 | 4242304 | 5786411 | 5601043.5 | 4639834 | 6152584.5 | 4213460.5 | 5436047.5 |
| Q99KJ8 | Dctn2 | Dynactin subunit 2 Mus musculus | 634.94 | 33.58 | 44117 | 11 | 11 | 22103775 | 23893517 | 21376959 | 24522623 | 24590845 | 23480173 | 26147221 | 25535555 | 20601097 | 26570927 | 22190939 | 23247911 | 21770877 | 23510899 | 20517621 |
| Q8CBY8 | Dctn4 | Dynactin subunit 4 Mus musculus | 116.78 | 5.78 | 53057 | 2 | 2 | 871438.31 | 877763.31 | 756759.69 | 956049.19 | 691533.44 | 777387.81 | 1120749 | 825161.69 | 613058.19 | 935667.31 | 758879.12 | 793917.94 | 816963.38 | 808261.62 | 907143.88 |
| Q9QZ73 | Dcun1d1 | DCN1-like protein 1 Mus musculus | 250.03 | 22.78 | 30097 | 4 | 4 | 3379584.25 | 2601892.75 | 2882740.75 | 3204482.25 | 2563609 | 3085217.75 | 3040334.25 | 2825890.25 | 3358710.5 | 3400748.75 | 2750840.5 | 2520839.5 | 3489474 | 2862288.5 | 2423609 |
| Q91X52 | Dcxr | L-xylulose reductase Mus musculus | 65.94 | 7.38 | 25746 | 2 | 2 | 134480689 | 89065553 | 74447881 | 112225169 | 119284305 | 70548753 | 53814041 | 66494233 | 38887289 | 62245973 | 104602489 | 16328652 | 57226097 | 66423661 | 48135453 |
| Q3U1J4 | Ddb1 | DNA damage-binding protein 1 Mus musculus | 2069.98 | 37.02 | 126853 | 5 | 37 | 74617521 | 78232241 | 78385473 | 74483321 | 76977025 | 77040681 | 80769745 | 78072993 | 90512401 | 76988841 | 82673193 | 78795521 | 70437657 | 72983609 | 76892625 |
| A2ADY9 | Ddi2 | Protein DDI1 homolog 2 Mus musculus | 288.58 | 18.8 | 44591 | 6 | 6 | 4650878 | 6068306 | 3721573 | 7267678 | 6312279.5 | 7031025 | 6584114 | 6845273.5 | 4785412 | 5239715.5 | 7177758 | 5902449 | 6593332.5 | 2156080.75 | 7160855.5 |
| O54734 | Ddost | Dolichyl-diphosphooligosaccharide--protein glycosyltransferase 48 kDa subunit Mus mu | 751.18 | 41.5 | 49028 | 13 | 13 | 519012865 | 34175413 | 129967369 | 173564145 | 191119313 | 36950333 | 47775645 | 38516305 | 37457721 | 42157145 | 33270161 | 39863357 | 37480813 | 91220017 | 37500993 |
| Q80WW9 | Ddrgk1 | DDRGK domain-containing protein 1 Mus musculus | 215.13 | 15.24 | 35977 | 3 | 3 | 2937053.5 | 3908093.75 | 3401647.25 | 2747720.25 | 3207205.75 | 3283543 | 2817227 | 3996858.5 | 4104785.5 | 2176598 | 2667373.5 | 3100155.25 | 3537014.75 | 3633007 | 2483953.25 |
| O35215 | Ddt | D-dopachrome decarboxylase Mus musculus | 52.72 | 9.32 | 13077 | 1 | 1 | 1 | 1 | 1 | 1 | 1 | 1 | 945238.5 | 1 | 1 | 1 | 1 | 1 | 877235.5 | 803039.25 | 758834.38 |
| Q91VR5 | Ddx1 | ATP-dependent RNA helicase DDX1 Mus musculus | 1054.07 | 34.32 | 82500 | 19 | 19 | 21494465 | 23311527 | 25019387 | 23063183 | 23753523 | 24373035 | 25186791 | 24691755 | 24103487 | 25796981 | 24154545 | 26848607 | 24054643 | 26628389 | 24637981 |
| Q80Y44 | Ddx10 | Probable ATP-dependent RNA helicase DDX10 Mus musculus | 110.36 | 3.2 | 100739 | 2 | 2 | 197709.77 | 182579.72 | 181611.52 | 224758.7 | 201889.14 | 225850.38 | 190651.06 | 208296.83 | 181243.81 | 254921.55 | 180653.33 | 302809.41 | 175936.33 | 154443.97 | 205760.86 |
| Q501J6 | Ddx17 | Probable ATP-dependent RNA helicase DDX17 Mus musculus | 957.64 | 31.54 | 72399 | 11 | 18 | 150077089 | 155918529 | 158525361 | 155182561 | 152922897 | 131302857 | 131145177 | 134895601 | 144248145 | 136351825 | 151269873 | 142305521 | 149909057 | 144687137 | 144611137 |
| Q8K363 | Ddx18 | ATP-dependent RNA helicase DDX18 Mus musculus | 1186.96 | 41.67 | 74181 | 20 | 20 | 17321211 | 17827805 | 16888109 | 16197246 | 11728795 | 17019201 | 15665945 | 15807677 | 15850639 | 16653373 | 17510295 | 17725025 | 16358788 | 15912138 | 15812109 |
| Q61655 | Ddx19a | ATP-dependent RNA helicase DDX19A Mus musculus | 416.28 | 23.43 | 53933 | 8 | 8 | 2878807 | 5348518 | 3112134 | 3867302.25 | 2950477 | 2648263 | 6967067 | 4194725 | 1641331.88 | 5153946.5 | 2563688.5 | 4010998.5 | 3600573.5 | 2911607.5 | 4383994.5 |
| Q9JJY4 | Ddx20 | Probable ATP-dependent RNA helicase DDX20 Mus musculus | 286.83 | 10.18 | 91710 | 6 | 6 | 5664927 | 6365350 | 1952999.12 | 6582219.5 | 5223471.5 | 4179188.75 | 955105.31 | 4910710.5 | 1127188.38 | 4598897.5 | 4531271 | 1612949.5 | 1389516 | 4395739.5 | 2442092 |
| Q9JIK5 | Ddx21 | Nucleolar RNA helicase 2 Mus musculus | 2053.9 | 54.76 | 93551 | 35 | 36 | 42154801 | 34022433 | 42058417 | 42927625 | 38194141 | 37060365 | 45113349 | 35940161 | 31665247 | 40163193 | 44285029 | 48608133 | 38488365 | 35962553 | 47258577 |
| Q9ESV0 | Ddx24 | ATP-dependent RNA helicase DDX24 Mus musculus | 345.21 | 10.74 | 96429 | 6 | 6 | 1468147.25 | 1989461.12 | 877275.38 | 1869901.25 | 1698180.5 | 542305.38 | 336972.75 | 565333.94 | 152117.78 | 1528880.75 | 1621332 | 1017331.44 | 643574.25 | 619783.88 | 1416769.75 |
| Q8VDW0 | Ddx39a | ATP-dependent RNA helicase DDX39A Mus musculus | 1355.7 | 69.32 | 49067 | 12 | 26 | 165816881 | 171790321 | 166252737 | 161726881 | 158953009 | 160027921 | 159381505 | 157467329 | 166698257 | 155496977 | 144335345 | 159304817 | 160838737 | 169610881 | 152561489 |
| Q63413 | Ddx39b | Spliceosome RNA helicase Ddx39b Rattus norvegicus 2 | 1524.45 | 73.83 | 49035 | 12 | 25 | 190927009 | 204810945 | 194309809 | 190996753 | 184235201 | 184215873 | 185416257 | 193178705 | 198341281 | 181412033 | 166506209 | 187616801 | 188813265 | 199545361 | 180550801 |
| Q62167 | Ddx3x | ATP-dependent RNA helicase DDX3X Mus musculus | 1544.52 | 48.79 | 73101 | 8 | 29 | 87973793 | 79139657 | 82385113 | 69691041 | 81013609 | 76875201 | 71658313 | 62282857 | 73474161 | 68232705 | 86649953 | 76894889 | 68378337 | 76194657 | 86889361 |
| Q62095 | Ddx3y | ATP-dependent RNA helicase DDX3Y Mus musculus | 1104.92 | 32.98 | 73428 | 2 | 23 | 77379209 | 69522481 | 72049065 | 60994129 | 71025609 | 67367865 | 58533945 | 53458897 | 64835241 | 57428789 | 75097913 | 66131121 | 57115589 | 67263249 | 72751697 |
| Q91VN6 | Ddx41 | Probable ATP-dependent RNA helicase DDX41 Mus musculus | 42.28 | 3.22 | 69820 | 1 | 1 | 1 | 1 | 962122.94 | 1 | 1 | 1193580.62 | 1 | 1 | 1770226.75 | 1 | 1 | 1 | 1 | 1013869.25 | 852927.75 |
| Q569Z5 | Ddx46 | Probable ATP-dependent RNA helicase DDX46 Mus musculus | 802.91 | 19.96 | 117448 | 17 | 17 | 19793049 | 18057853 | 20214551 | 16679657 | 19095113 | 17260405 | 23237461 | 19670713 | 17363569 | 18393571 | 20213449 | 21337833 | 18493913 | 19093335 | 19939303 |
| Q9CWX9 | Ddx47 | Probable ATP-dependent RNA helicase DDX47 Mus musculus | 375.31 | 22.2 | 50639 | 8 | 8 | 6133848 | 7564842 | 7843512 | 5960981 | 5154378.5 | 7758665 | 7277506 | 9126638 | 8453459 | 5529613 | 6041178 | 7248882 | 6293231.5 | 8859403 | 5491866 |
| Q4FZF3 | Ddx49 | Probable ATP-dependent RNA helicase DDX49 Mus musculus | 177.44 | 10.62 | 54094 | 3 | 3 | 804309.06 | 680725.81 | 1172160.75 | 642922.56 | 722912.12 | 869432 | 645084.75 | 733971.5 | 960636.31 | 689108.06 | 977025.62 | 982701.94 | 357246.91 | 1267582.38 | 474060 |
| Q61656 | Ddx5 | Probable ATP-dependent RNA helicase DDX5 Mus musculus | 1563.06 | 58.63 | 69290 | 22 | 29 | 225700161 | 233248081 | 244238129 | 239267329 | 231313041 | 209811377 | 204388465 | 201991633 | 228882321 | 210188881 | 241180161 | 219286065 | 218777841 | 218221761 | 217496545 |
| Q8K301 | Ddx52 | Probable ATP-dependent RNA helicase DDX52 Mus musculus | 150.01 | 8.19 | 67474 | 3 | 3 | 343005.78 | 337891.59 | 532897.12 | 330994.75 | 334590.16 | 296209.47 | 290149.09 | 349136.16 | 448818.09 | 432221.47 | 258045.5 | 324417.69 | 373828.44 | 334900.72 | 347076.72 |
| Q8K4L0 | Ddx54 | ATP-dependent RNA helicase DDX54 Mus musculus | 179.67 | 7.67 | 97748 | 5 | 5 | 1234306.88 | 602764.81 | 1249032 | 793462.38 | 979783.06 | 916264.31 | 1048350.5 | 942991.25 | 926111.62 | 691292.06 | 1018422.75 | 1090981 | 1045604.25 | 983050.88 | 1217710 |
| Q9D0R4 | Ddx56 | Probable ATP-dependent RNA helicase DDX56 Mus musculus | 178.1 | 9.34 | 61212 | 3 | 3 | 932268.62 | 817630.69 | 536774.25 | 1242635.12 | 1438026.62 | 1170881 | 737305.56 | 778880.25 | 877724.0475 | 823829.38 | 1258359 | 329583.75 | 571693.62 | 677364.94 | 690369.44 |
| Q66HG7 | Ddx59 | Probable ATP-dependent RNA helicase DDX59 Rattus norvegicus | 37.42 | 1.87 | 65066 | 1 | 1 | 175512.03 | 251675.5 | 279745.94 | 249485.97 | 195653.36 | 224255.92 | 1 | 216203.02 | 1 | 1 | 305113.75 | 147372.41 | 215540.9475 | 186545.83 | 223131.8 |
| P54823 | Ddx6 | Probable ATP-dependent RNA helicase DDX6 Mus musculus | 907.65 | 44.1 | 54192 | 13 | 13 | 17343113 | 18286627 | 16789805 | 17062831 | 17349095 | 16915557 | 19420463 | 14952106 | 15565606 | 19430723 | 16917825 | 18354501 | 18794103 | 17192743 | 16612632 |
| Q9CQ62 | Decr1 | 2,4-dienoyl-CoA reductase, mitochondrial Mus musculus | 266.22 | 26.57 | 36214 | 5 | 5 | 9110959 | 3718888 | 3367780.5 | 9925201 | 10073742 | 11239210 | 9472981 | 10148875 | 5347762 | 8170669.5 | 8985094 | 7597503 | 8599847 | 9895916 | 6769474.5 |
| Q8C2K1 | Def6 | Differentially expressed in FDCP 6 Mus musculus | 45.02 | 1.59 | 73454 | 1 | 1 | 79740.24 | 1 | 40264.1 | 58635.16 | 1 | 35110.97 | 119539.7 | 1 | 1 | 63210.22 | 52460.83 | 34276.04 | 1 | 1 | 54860.75 |
| Q6AXS3 | Dek | Protein DEK Rattus norvegicus | 458.83 | 24.07 | 42892 | 9 | 9 | 24901535 | 25517797 | 23843257 | 21857401 | 26834323 | 23380493 | 23307915 | 22175649 | 25220345 | 21764353 | 26807761 | 22602929 | 23493213 | 22805665 | 23142007 |
| Q3U1Y4 | Dennd4b | DENN domain-containing protein 4B Mus musculus | 37.34 | 0.87 | 164740 | 1 | 1 | 364688.84 | 196452.06 | 1 | 1 | 1 | 321428.03 | 1 | 1 | 1 | 404778.59 | 1 | 1 | 1 | 1 | 1 |
| A6H8H2 | Dennd4c | DENN domain-containing protein 4C Mus musculus | 427.89 | 6.56 | 211458 | 8 | 8 | 3449451.25 | 4322360.5 | 5971060.5 | 4559859 | 6918938 | 5570263 | 6279434 | 6413485 | 7249285 | 6695763.5 | 5748112.5 | 5513608 | 5576692 | 6227656.5 | 6118767 |
| Q9CQJ6 | Denr | Density-regulated protein Mus musculus | 212.91 | 19.19 | 22166 | 3 | 3 | 4609877 | 5230410 | 5181511.5 | 4923461 | 4933535 | 3900525 | 4577298 | 4487641 | 4550900.5 | 4414574.5 | 5179689 | 3995056 | 4462624 | 4221531.5 | 4472574 |
| Q91YP3 | Dera | Deoxyribose-phosphate aldolase Mus musculus | 437.65 | 32.08 | 34975 | 7 | 7 | 4085484.5 | 3181956 | 3918847.75 | 3109663.5 | 4620571.5 | 3823992.25 | 4241371.5 | 4527921 | 4115076 | 4434956.5 | 5059218 | 4667641 | 3283676.25 | 2824061 | 4573306.5 |
| O08560 | Dgkz | Diacylglycerol kinase zeta Rattus norvegicus | 71.91 | 4.41 | 104009 | 2 | 2 | 1274328.25 | 1207170.12 | 1073381.62 | 1266015.88 | 1295138.75 | 1484053.5 | 1665077.12 | 1293081.75 | 1490501 | 1164524.38 | 1226352.12 | 1426831 | 1216745.38 | 1265337 | 1331210.75 |
| Q8BH86 | Dglucy | D-glutamate cyclase, mitochondrial Mus musculus | 46.52 | 3.24 | 66366 | 1 | 1 | 783220.88 | 994559.88 | 892727.81 | 801676.81 | 900332.75 | 1051776.62 | 663309.56 | 877795.56 | 924246.25 | 796783.81 | 1106387 | 863730.25 | 947165.88 | 1116352.38 | 888823.75 |
| O88455 | Dhcr7 | 7-dehydrocholesterol reductase Mus musculus | 210.85 | 11.25 | 53919 | 4 | 4 | 4165302.5 | 652550.19 | 2017097.62 | 731162.75 | 1030888.75 | 1859448.25 | 575362.19 | 2281899.5 | 3678798.25 | 1266704.12 | 872595.31 | 1695753.62 | 844492.06 | 2041316.38 | 2379486.5 |
| Q9DBB8 | Dhdh | Trans-1,2-dihydrobenzene-1,2-diol dehydrogenase Mus musculus | 253.19 | 17.42 | 36301 | 4 | 4 | 4518272.5 | 3689742.5 | 4388964 | 3267605 | 3540505.25 | 4037920 | 4836099.5 | 4748797.5 | 4233262.5 | 6999435.5 | 4626121 | 5341414 | 3522813.5 | 4341920.5 | 5339389.5 |
| P00375 | Dhfr | Dihydrofolate reductase Mus musculus | 279.43 | 32.62 | 21606 | 5 | 5 | 14760133 | 15929109 | 14222924 | 13578961 | 14830744 | 14333998 | 12836034 | 15574199 | 14370444 | 7733765 | 14851590 | 14521323 | 14082929 | 14949848 | 11421259 |
| O35435 | Dhodh | Dihydroorotate dehydrogenase (quinone), mitochondrial Mus musculus | 93.26 | 8.35 | 42700 | 2 | 2 | 858975 | 905831.12 | 701636.5 | 959163.44 | 692961.5 | 881605.62 | 715146.31 | 683328.94 | 623604.19 | 725292.38 | 655397.69 | 522686.34 | 840175.38 | 743386.88 | 692607.38 |
| Q3TXU5 | Dhps | Deoxyhypusine synthase Mus musculus | 118.73 | 8.94 | 40642 | 2 | 2 | 323303.9 | 239570.72 | 328548.91 | 490637.78 | 234458.19 | 375425.47 | 202078.66 | 366851.91 | 474078.41 | 354608.6125 | 240974.52 | 403675.22 | 513869.28 | 314125.66 | 258034.06 |
| Q99L04 | Dhrs1 | Dehydrogenase/reductase SDR family member 1 Mus musculus | 449.48 | 31.31 | 34005 | 8 | 8 | 9228698 | 9496024 | 10085423 | 8401514 | 7871312 | 9786225 | 9617604 | 10768845 | 9678623 | 11349676 | 9324894 | 4661921.5 | 8594044 | 7676005 | 11351115 |
| Q99LB2 | Dhrs4 | Dehydrogenase/reductase SDR family member 4 Mus musculus | 158.01 | 26.16 | 29885 | 5 | 5 | 1410262.12 | 1028558.19 | 1760978.75 | 1354389.12 | 1388547.045 | 729860.12 | 2051849.75 | 1150848.25 | 1011418.12 | 1481677.75 | 1234317.75 | 1716028.5 | 1241311 | 730374.19 | 653561.12 |
| Q9CXR1 | Dhrs7 | Dehydrogenase/reductase SDR family member 7 Mus musculus | 173.06 | 13.91 | 38167 | 4 | 4 | 7001172.5 | 3615721.75 | 5859879 | 5189765 | 5816470 | 5181083.5 | 5034834 | 4137078.75 | 5368822.5 | 5896325 | 5361671.5 | 5140534 | 3481968 | 4397987 | 6592918 |
| Q99J47 | Dhrs7b | Dehydrogenase/reductase SDR family member 7B Mus musculus | 94.12 | 9.29 | 34987 | 2 | 2 | 1416543.75 | 1785989.88 | 1364105.88 | 1098790 | 1191268.75 | 1514119.5 | 1481832.38 | 1068151.25 | 1151558.75 | 1307921.5 | 1317672.38 | 1476677.5 | 1360893.75 | 1206912.38 | 1434426.12 |
| O35286 | Dhx15 | Pre-mRNA-splicing factor ATP-dependent RNA helicase DHX15 Mus musculus | 1227.15 | 33.08 | 91007 | 21 | 23 | 40992317 | 41301829 | 43735069 | 41636641 | 43165537 | 44596437 | 44462501 | 42507173 | 43727801 | 43284265 | 46653053 | 44389185 | 43071265 | 43825093 | 43951017 |
| Q6PGC1 | Dhx29 | ATP-dependent RNA helicase DHX29 Mus musculus | 81.86 | 1.61 | 153975 | 2 | 2 | 517866.44 | 265201.62 | 577928.75 | 253528.73 | 222110.31 | 509119.8575 | 399515 | 632660.06 | 668181.12 | 336123.25 | 566747.25 | 547106.44 | 344068.47 | 568414.5 | 506584.165 |

| | | | | | | | | | | | | | | | | | | | | | | |
|---|---|---|---|---|---|---|---|---|---|---|---|---|---|---|---|---|---|---|---|---|---|---|
| Q5BJS0 | Dhx30 | ATP-dependent RNA helicase DHX30 Rattus norvegicus | 417.71 | 7.2 | 133997 | 7 | 7 | 1912891 | 1417401.5 | 1832725.5 | 1846246.5 | 1779634.38 | 1786599.38 | 2021214.75 | 1498898.5 | 952998.12 | 2228655.75 | 2076461.12 | 1752084.75 | 1211854.75 | 1901684.62 | 1072036.75 |
| D4A2Z8 | Dhx36 | ATP-dependent DNA/RNA helicase DHX36 Rattus norvegicus | 261.69 | 7.4 | 113843 | 6 | 6 | 1602403.38 | 2228548.25 | 2125017.75 | 1845307 | 3894404.25 | 996273.81 | 1602195.75 | 2874390 | 1254084.38 | 1337136.88 | 1814774 | 1301755.88 | 1799316 | 1070283.62 | 1206505.25 |
| Q99J87 | Dhx58 | Probable ATP-dependent RNA helicase DHX58 Mus musculus | 65.33 | 1.62 | 76709 | 1 | 1 | 189459.12 | 153219.42 | 177991.3725 | 204719.97 | 164566.98 | 198482.25 | 212434.3 | 1 | 1 | 167838.06 | 149855.61 | 137130.72 | 145173.77 | 226606.05 | 319270.03 |
| A2A4P0 | Dhx8 | ATP-dependent RNA helicase DHX8 Mus musculus | 221.25 | 5.95 | 142572 | 5 | 5 | 1605208.75 | 1430641 | 1840368.5 | 1847155.38 | 1598596.25 | 1594515.75 | 1842691.5 | 1588289.38 | 2081833.38 | 1611962.75 | 1066993.12 | 2337044 | 1442831.38 | 1799979.75 | 1523982.25 |
| O70133 | Dhx9 | ATP-dependent RNA helicase A Mus musculus | 2773.33 | 43.12 | 149475 | 48 | 49 | 86166737 | 101898305 | 103340321 | 102476721 | 98230921 | 90741225 | 88853865 | 90048753 | 97773281 | 99210001 | 115657433 | 100752433 | 91750809 | 96470289 | 102910809 |
| F1M775 | Diaph1 | Protein diaphanous homolog 1 Rattus norvegicus | 716.21 | 15.57 | 140409 | 14 | 14 | 8559447 | 9502858 | 9598082 | 8899180 | 9757611 | 10790464 | 8604671 | 9252780 | 9687399 | 8292077 | 9851643 | 6757847 | 7761040 | 10986880 | 9488571 |
| Q8R418 | Dicer1 | Endoribonuclease Dicer Mus musculus | 128.52 | 3.08 | 216821 | 3 | 3 | 385942.12 | 259771.64 | 133663.34 | 624994.94 | 668523.62 | 259510.45 | 134457.02 | 670631.69 | 901235.12 | 290624.22 | 610493.88 | 439252.62 | 714862.25 | 818728.94 | 443291.12 |
| Q5M9G7 | Diexf | Digestive organ expansion factor homolog Rattus norvegicus | 160.09 | 4.59 | 87825 | 3 | 3 | 1 | 1 | 1 | 1 | 380879.97 | 1 | 264339.38 | 1 | 1 | 1 | 295015.78 | 524641.725 | 644858 | 786307.12 | 372386 |
| Q9D0D4 | Dimt1 | Probable dimethyladenosine transferase Mus musculus | 65.46 | 6.39 | 35274 | 2 | 2 | 2119810.25 | 1234545.88 | 3318215.5 | 1956084.72 | 1151767.25 | 1248771.25 | 1168478.5 | 1393652.75 | 1665552.88 | 2065020 | 3018185 | 2811874.5 | 810347.69 | 1259767.12 | 1290689.5 |
| Q3UH60 | Dip2b | Disco-interacting protein 2 homolog B Mus musculus | 436.25 | 7.24 | 171126 | 8 | 8 | 6769005 | 8648224 | 4753545 | 4431788.5 | 5067007 | 4762376 | 5531545.5 | 4683440.5 | 5372906 | 5708679 | 7362909 | 5239071.5 | 4269295 | 4426398 | 5607628 |
| Q9CSH3 | Dis3 | Exosome complex exonuclease RRP44 Mus musculus | 293.29 | 6.58 | 108838 | 5 | 5 | 7311804 | 4772028 | 5428749 | 5526741 | 5278218.5 | 3589113.5 | 5657455 | 3963166.75 | 4347398.5 | 6336550 | 10231491 | 5497096 | 3900234.5 | 5255286 | 7093523 |
| Q8CI75 | Dis3l2 | DIS3-like exonuclease 2 Mus musculus | 134.43 | 4.94 | 97775 | 3 | 3 | 1995034.5 | 1161081.88 | 1202536.38 | 1251003 | 1456505 | 1992600.25 | 1111291.25 | 2468764.5 | 2183435.25 | 1414187.62 | 506186.19 | 1026304.75 | 1423609.88 | 1215661.5 | 1403222 |
| Q9ESX5 | Dkc1 | H/ACA ribonucleoprotein complex subunit DKC1 Mus musculus | 643.3 | 30.84 | 57402 | 11 | 11 | 15481263 | 14396183 | 14286213 | 14283651 | 16759178 | 13539379 | 14231537 | 13626513 | 14275714 | 14695176 | 12761067 | 14904217 | 10622242 | 14806882 | 14497303 |
| Q8BMF4 | Dlat | Dihydrolipoyllysine-residue acetyltransferase component of pyruvate dehydrogenase cor | 709.5 | 31 | 67942 | 13 | 13 | 31919889 | 36952193 | 38732433 | 35680857 | 43382769 | 41310493 | 43577193 | 40824181 | 48855377 | 36362241 | 33291621 | 35398509 | 33457267 | 37747617 | 37035637 |
| O08749 | Dld | Dihydrolipoyl dehydrogenase, mitochondrial Mus musculus | 1010.55 | 45.38 | 54272 | 16 | 16 | 91068521 | 71589561 | 59574589 | 75143865 | 65895429 | 75847385 | 62355905 | 87209569 | 82844945 | 58102833 | 59013273 | 61097677 | 64878841 | 101063865 | 50467141 |
| B1AZP2 | Dlgap4 | Disks large-associated protein 4 Mus musculus | 100.25 | 2.62 | 108037 | 2 | 2 | 8189513.5 | 6492710 | 3329314 | 7200943 | 2492961.5 | 4159553 | 4904876 | 3977050 | 4014644.25 | 5787117 | 4138012 | 3511905 | 3841282 | 4286428.5 | 1449119.12 |
| Q01205 | Dlst | Dihydrolipoyllysine-residue succinyltransferase component of 2-oxoglutarate dehydroger | 615.58 | 33.48 | 48925 | 11 | 11 | 51557229 | 38689277 | 32494921 | 46522977 | 32387917 | 38520697 | 35296321 | 51536341 | 39557953 | 23115613 | 27032197 | 31331313 | 33709145 | 53435333 | 21449719 |
| P63036 | Dnaja1 | DnaJ homolog subfamily A member 1 Rattus norvegicus | 389.08 | 28.97 | 44868 | 7 | 7 | 5205532.5 | 4361426.5 | 5415119 | 5853004 | 4832383 | 5403787 | 5368215 | 5800223.5 | 5588741 | 5267306.5 | 6560593.5 | 6260800 | 4215096 | 5634190.5 | 6126166.5 |
| Q9QYJ0 | Dnaja2 | DnaJ homolog subfamily A member 2 Mus musculus | 666.3 | 48.79 | 45746 | 12 | 12 | 29684813 | 9392349 | 19706347 | 29303685 | 10326661 | 12412809 | 12681225 | 9835568 | 16294845 | 9972949 | 12739244 | 12039708 | 8959426 | 9026058 | 14424864 |
| Q99M87 | Dnaja3 | DnaJ homolog subfamily A member 3, mitochondrial Mus musculus | 95.77 | 8.12 | 52443 | 2 | 2 | 739663.94 | 749693.12 | 618731.38 | 834531.69 | 774624.56 | 696421.62 | 1487545.12 | 627053.69 | 682695.06 | 1262379.12 | 561176.38 | 846832 | 823406.81 | 585208.5 | 1029535 |
| Q9QYJ3 | Dnajb1 | DnaJ homolog subfamily B member 1 Mus musculus | 165.74 | 15 | 38167 | 5 | 5 | 4165890 | 3318580 | 3386349.25 | 3622478 | 3842204.5 | 2761575.5 | 3933247.5 | 3940806.75 | 3422719.25 | 4121723 | 4008319.5 | 4863904 | 5816166 | 5807165 | 5173635 |
| Q6TUG0 | Dnajb11 | DnaJ homolog subfamily B member 11 Rattus norvegicus | 321.83 | 19.55 | 40495 | 5 | 5 | 7270898.5 | 7374524.5 | 9912708 | 8706725 | 7944480 | 8736350 | 10738571 | 8924369 | 7291184 | 11332374 | 8656601 | 9698720 | 9586673 | 5506079.5 | 10359231 |
| O54946 | Dnajb6 | DnaJ homolog subfamily B member 6 Mus musculus | 48.66 | 3.56 | 39807 | 1 | 1 | 1401260.25 | 534497.81 | 799267.75 | 530233.31 | 1341338.5 | 667211.12 | 902567.62 | 952076 | 320206.88 | 708256.5 | 1138417.5 | 542119.75 | 721930.2175 | 606069.31 | 601114.31 |
| Q498R3 | Dnajc10 | DnaJ homolog subfamily C member 10 Rattus norvegicus | 77.08 | 2.52 | 90747 | 2 | 2 | 1558368.5 | 1356446.38 | 959210.19 | 1223846.38 | 1740151.12 | 1297684.233 | 195029.3 | 1786443 | 1701749.75 | 1507514.88 | 120921.12 | 191882.98 | 1132643.25 | 1445420.38 | 722716.9325 |
| Q5U458 | Dnajc11 | DnaJ homolog subfamily C member 11 Mus musculus | 68.93 | 6.08 | 63233 | 2 | 2 | 1756696.12 | 1877092.38 | 1830327.75 | 1543130.25 | 1021202.5 | 1634770.25 | 1593001.25 | 1571996.75 | 1842113.38 | 1330537.75 | 1001178.19 | 1537887 | 1137273.5 | 1425073.88 | 1235046.38 |
| Q9CQV7 | Dnajc19 | Mitochondrial import inner membrane translocase subunit TIM14 Mus musculus | 72.39 | 16.38 | 12437 | 1 | 1 | 2629887.25 | 1677089.62 | 1913949.5 | 2179990.75 | 1895540.75 | 1 | 1 | 1842547.12 | 1650783.62 | 2388106.75 | 1495065 | 1363410.75 | 2213421 | 2335724 | 1485132.5 |
| P54103 | Dnajc2 | DnaJ homolog subfamily C member 2 Mus musculus | 218.77 | 10.79 | 71722 | 5 | 5 | 2841132.5 | 2349083.75 | 3566257 | 3432205 | 3433569.5 | 3412009.25 | 2886638.5 | 4139736 | 4469408.5 | 3990827.25 | 4947841.5 | 2875952.5 | 3605293.5 | 2452718.75 | 2412716 |
| Q91YW3 | Dnajc3 | DnaJ homolog subfamily C member 3 Mus musculus | 326.8 | 15.67 | 57464 | 5 | 5 | 2656988.75 | 1071912 | 1458179.25 | 816552.88 | 1249762.5 | 735783.94 | 1313776.62 | 864397.19 | 1122786.75 | 1726949.38 | 831773 | 731729.69 | 1469003 | 1400662.62 | 1100087.62 |
| P60904 | Dnajc5 | DnaJ homolog subfamily C member 5 Mus musculus | 99.44 | 19.7 | 22101 | 2 | 2 | 2099868.25 | 1786901.88 | 1646291.75 | 1272206.5 | 1521797.38 | 1779029.75 | 1369170.25 | 1800175.88 | 1917789.75 | 1347160 | 1194039.75 | 1273004.38 | 1583518.5 | 1704126.38 | 1238720 |
| Q9QYI3 | Dnajc7 | DnaJ homolog subfamily C member 7 Mus musculus | 490.69 | 20.65 | 56476 | 8 | 8 | 12213347 | 12484708 | 14398115 | 11520088 | 11159544 | 10149566 | 10438176 | 11193330 | 12147035 | 11382222 | 12117433 | 12955840 | 10040374 | 13163616 | 12592617 |
| Q642C0 | Dnajc8 | DnaJ homolog subfamily C member 8 Rattus norvegicus 2 | 265.66 | 25.69 | 29813 | 6 | 6 | 11509153 | 7377620 | 6784950.5 | 9237301 | 10761839 | 9721004 | 9915423 | 8044927 | 6519385.5 | 10496287 | 9404443 | 10775951 | 10933352 | 8728448 | 7561952.5 |
| P21704 | Dnase1 | Deoxyribonuclease-1 Rattus norvegicus 2 | 35.75 | 2.46 | 32064 | 1 | 1 | 1642476 | 1399387.75 | 1698094.88 | 1988958.5 | 1682229.283 | 1742235.62 | 1522354.5 | 1601454.25 | 1434646.75 | 1311081.38 | 1623620.59 | 1798499.12 | 1825984.5 | 1401982.12 | 1468016.62 |
| P56542 | Dnase2 | Deoxyribonuclease-2-alpha Mus musculus | 215.95 | 23.8 | 38811 | 5 | 5 | 4207111 | 2816676.5 | 3643136.5 | 2673681.75 | 3293379.25 | 3762286 | 2731003 | 1262468 | 3730327.5 | 1437286.75 | 3789854.5 | 3732551.5 | 2405606.5 | 4119827.5 | 3202700 |
| P21575 | Dnm1 | Dynamin-1 Rattus norvegicus | 204.22 | 5.56 | 97295 | 1 | 5 | 5910187 | 5974341.5 | 4994817 | 5374599.5 | 5425708 | 4874912.5 | 4518357.5 | 4186290.25 | 4259415.5 | 5206855 | 3825795.75 | 4161140.75 | 5243458.5 | 4622964 | 4545649 |
| O35303 | Dnm1l | Dynamin-1-like protein Rattus norvegicus | 547.71 | 20.93 | 83908 | 11 | 11 | 8967139 | 7922948.5 | 9526005 | 7493573.5 | 6815973 | 6957055.5 | 6624463 | 9561986 | 7941688 | 5846445.5 | 8132781 | 7720258 | 6705387.5 | 8602376 | 7842387 |
| P39052 | Dnm2 | Dynamin-2 Rattus norvegicus | 1083.22 | 34.25 | 98230 | 15 | 22 | 24288561 | 25117165 | 23822793 | 25118757 | 24548741 | 24891017 | 23704319 | 25028799 | 27451965 | 26330753 | 21960011 | 24182271 | 24575167 | 24427085 | 26298017 |
| Q08877 | Dnm3 | Dynamin-3 Rattus norvegicus | 280.16 | 8.63 | 97914 | 2 | 7 | 7508758 | 5974341.5 | 6584546.5 | 6278341.5 | 6236007 | 6190637.5 | 5187748 | 5874033.5 | 6239044 | 5206855 | 5566173 | 6240221 | 5477093 | 6483449 | 4658788 |
| Q6TXD4 | Dnmbp | Dynamin-binding protein Mus musculus | 52.96 | 0.95 | 177277 | 1 | 1 | 159029.23 | 210863.73 | 187377.3 | 206897.78 | 180768.41 | 156227.78 | 158001.81 | 205828.19 | 242682.48 | 171737.3 | 253042.3 | 201979.98 | 203125.77 | 202077 | 232749.38 |
| P13864 | Dnmt1 | DNA (cytosine-5)-methyltransferase 1 Mus musculus | 1939.1 | 31.23 | 183189 | 20 | 36 | 39514585 | 29406143 | 31782447 | 43598445 | 37225265 | 32986203 | 30753991 | 33685505 | 25704207 | 31908715 | 30259665 | 25156389 | 28911847 | 31834239 | 35917869 |
| O88508 | Dnmt3a | DNA (cytosine-5)-methyltransferase 3A Mus musculus | 410.78 | 9.03 | 101672 | 7 | 7 | 7736163 | 7329320.5 | 7615047 | 8064499 | 7192452 | 6461072 | 4780640.5 | 7554471 | 8592235 | 7697444 | 6696447.5 | 8069507.5 | 6903971.5 | 8875811 | 8320791 |
| Q9Z2W0 | Dnpep | Aspartyl aminopeptidase Mus musculus | 473.59 | 34.25 | 52207 | 9 | 9 | 10101427 | 10563212 | 10241226 | 12578738 | 13703730 | 12802727 | 16119935 | 13877441 | 10408147 | 11666191 | 11512340 | 11185051 | 13407879 | 11963949 | 13982898 |
| Q8R2M2 | Dnttip2 | Deoxynucleotidyltransferase terminal-interacting protein 2 Mus musculus | 328.7 | 10.29 | 84277 | 6 | 6 | 1357707.25 | 1842853.75 | 756072 | 1547577 | 2191822.5 | 655513.19 | 366730.88 | 922301.88 | 820393.75 | 1623295 | 869142 | 742590.44 | 1435173.75 | 598913.25 | 1399097.62 |
| Q8BZN6 | Dock10 | Dedicator of cytokinesis protein 10 Mus musculus | 444.44 | 7.35 | 245758 | 10 | 10 | 5128567.5 | 5359183.5 | 5712585 | 5539528 | 5490950.5 | 5044509.5 | 3796692.5 | 5807967 | 6991778.5 | 5087212 | 4997829.5 | 5547793.5 | 5581743 | 5616482.5 | 5649757 |
| Q8C3J5 | Dock2 | Dedicator of cytokinesis protein 2 Mus musculus | 1397.04 | 22.26 | 211704 | 30 | 30 | 46099521 | 35438617 | 30045473 | 35338525 | 37959521 | 38929545 | 28959695 | 34419265 | 40649505 | 35698297 | 34504689 | 34685565 | 37125417 | 38336749 | 37562957 |
| Q8R1A4 | Dock7 | Dedicator of cytokinesis protein 7 Mus musculus | 78.76 | 1.46 | 241438 | 2 | 2 | 285566.44 | 297161.34 | 223013.19 | 1 | 1 | 283616.91 | 245760.7 | 204564.11 | 216376.92 | 237579.66 | 115367.21 | 263736.81 | 283089.41 | 143592 | 201446.3575 |
| Q8C147 | Dock8 | Dedicator of cytokinesis protein 8 Mus musculus | 692.25 | 8.52 | 238978 | 13 | 13 | 8653259 | 7572209 | 10662542 | 6787040 | 9179315 | 8963780 | 7819363.5 | 10120269 | 5724227 | 7254306 | 11294072 | 7822561 | 7661420 | 6846361 | 8319963.5 |
| Q99LN9 | Dohh | Deoxyhypusine hydroxylase Mus musculus | 35.57 | 6.29 | 32905 | 1 | 1 | 893574.75 | 1 | 1 | 1 | 1 | 1 | 1 | 1 | 1 | 1 | 1 | 1 | 369119.69 | 1 | 1 |
| P97465 | Dok1 | Docking protein 1 Mus musculus | 54.25 | 1.87 | 52452 | 1 | 1 | 1090736.38 | 982401.06 | 880917 | 1042535.5 | 1077710.5 | 850192.94 | 1025531.38 | 714807.94 | 771414.12 | 1208695 | 1104135.12 | 886617.38 | 957747.19 | 700932.62 | 1152789.25 |
| O70469 | Dok2 | Docking protein 2 Mus musculus | 120.68 | 6.8 | 45522 | 2 | 2 | 238499.89 | 1 | 1 | 328955.28 | 202227.56 | 1 | 1 | 862929.81 | 193373.97 | 145840.78 | 146853.17 | 1 | 203684.92 | 1 | 1 |
| Q61103 | Dpf2 | Zinc finger protein ubi-d4 Mus musculus | 145.11 | 6.39 | 44230 | 2 | 2 | 1463768.38 | 2731879.75 | 2694503.5 | 3174368 | 2388616 | 2224094.75 | 2744191.25 | 2400221.75 | 2697031 | 2686084 | 1831115 | 2661648 | 2602844 | 2958816.25 | 2746716 |
| Q9CWQ0 | Dph5 | Diphthine methyl ester synthase Mus musculus | 94.73 | 12.81 | 31218 | 2 | 2 | 1 | 444648.53 | 1 | 446650.34 | 803049.88 | 501548.41 | 366273.47 | 245534.3 | 362036.88 | 728808.94 | 208603.12 | 186628.73 | 676338.69 | 463575.91 | 383786.6125 |
| O70152 | Dpm1 | Dolichol-phosphate mannosyltransferase subunit 1 Mus musculus | 248.54 | 28.85 | 29175 | 5 | 5 | 6037562 | 7843319 | 3448376.5 | 3115702.25 | 5243636.5 | 5256171 | 4993951.5 | 6496517.5 | 7099908.5 | 4581021.5 | 5492992.5 | 5633431.5 | 2761424 | 6658735 | 5590401 |
| Q99KK7 | Dpp3 | Dipeptidyl peptidase 3 Mus musculus | 1139.97 | 36.99 | 82898 | 8 | 18 | 32399055 | 34278557 | 35120525 | 27244561 | 30640621 | 34138369 | 29615605 | 35494393 | 36990057 | 28395965 | 33961593 | 32059081 | 30663241 | 35449509 | 36141853 |
| Q9ET22 | Dpp7 | Dipeptidyl peptidase 2 Mus musculus | 242.86 | 13.44 | 56254 | 5 | 5 | 2962367 | 3716168.5 | 3367190.5 | 2330683.25 | 2784412 | 2824017.25 | 2879851.75 | 3065073.25 | 3059115 | 2665075.5 | 3909999.5 | 2901683.25 | 3295310.75 | 2992284.5 | 2708386.5 |
| Q8BVG4 | Dpp9 | Dipeptidyl peptidase 9 Mus musculus | 203.67 | 9.05 | 98001 | 5 | 5 | 1704558.75 | 1444616.25 | 1742400.5 | 946576 | 1329590.38 | 1148624.62 | 1064910.5 | 2446403.5 | 1719019.38 | 1008262.94 | 1295868.88 | 1909159.5 | 1092074.25 | 1146647.88 | 1404310 |
| Q99LT0 | Dpy30 | Protein dpy-30 homolog Mus musculus | 102.07 | 36.36 | 11213 | 2 | 2 | 4752878 | 7616383.5 | 6300841 | 6682824 | 5467161 | 5067646 | 5942710.5 | 4978094 | 5609830 | 6906103.5 | 6014397.5 | 5879500 | 5886779 | 3802336.75 | 2777971.75 |
| O08553 | Dpysl2 | Dihydropyrimidinase-related protein 2 Mus musculus | 1856.42 | 76.57 | 62278 | 28 | 30 | 191156849 | 225539953 | 227241681 | 212314257 | 229917153 | 217187729 | 229418545 | 205574497 | 218439697 | 216871297 | 246787457 | 234991649 | 241025569 | 225158913 | 232815281 |
| A0JPP1 | Drap1 | Dr1-associated corepressor Rattus norvegicus | 134.49 | 11.22 | 22326 | 3 | 3 | 3956198.25 | 4866696.5 | 3782367.25 | 4190249 | 4447282.5 | 4290098.5 | 4213034.5 | 4491935 | 4542670 | 3702688.75 | 3812719 | 3870091 | 3871978.75 | 4148859.25 | 3184178 |
| P32233 | Drg1 | Developmentally-regulated GTP-binding protein 1 Mus musculus | 409.81 | 28.34 | 40512 | 7 | 8 | 5193407.5 | 4370919 | 6439510.5 | 5552117 | 4935575 | 6503560 | 6693040.5 | 6088467.5 | 3171660 | 5716288.5 | 6128820.5 | 6605359 | 5868650 | 7169103.5 | 5566609.5 |
| Q9QXB9 | Drg2 | Developmentally-regulated GTP-binding protein 2 Mus musculus | 390.09 | 23.9 | 40718 | 6 | 7 | 5568224 | 3879384.75 | 5953378 | 6886031 | 6015587 | 6698829 | 6781650 | 6461302.5 | 6436981.5 | 6818936 | 7631782 | 6374659.5 | 6165341.5 | 5513332.5 | 7616607 |
| O35075 | Dscr3 | Down syndrome critical region protein 3 homolog Mus musculus | 181.98 | 18.86 | 32970 | 4 | 4 | 3226952.5 | 2074669 | 2602682 | 3277601 | 2343939.5 | 3417482 | 3546488.5 | 2750092 | 3267153 | 2515547.5 | 2864005.75 | 2698015 | 3306750.5 | 3625134 | 1749658.62 |
| O35902 | Dsg3 | Desmoglein-3 Mus musculus | 39.5 | 0.7 | 107889 | 1 | 1 | 1 | 1 | 661985.75 | 550804 | 511728.59 | 479302.78 | 938811.44 | 289229.41 | 348067.16 | 794589.94 | 553927.12 | 667113.44 | 559986 | 652168.015 | 827645.5 |
| Q9R0P5 | Dstn | Destrin Mus musculus | 489.98 | 52.12 | 18522 | 7 | 7 | 29927785 | 31798485 | 33788209 | 31869721 | 32292155 | 33226775 | 28372427 | 28077435 | 26973557 | 27633817 | 34233281 | 30639047 | 28537335 | 28868759 | 31430413 |
| Q9DD18 | Dtd1 | D-aminoacyl-tRNA deacylase 1 Mus musculus | 52.97 | 7.18 | 23384 | 1 | 1 | 1673924 | 1824673.25 | 1704369.62 | 2156659 | 1926094.25 | 1796980.25 | 1962014.62 | 2065641.905 | 2302953 | 2200619.75 | 1982123.563 | 2065957 | 2092495.5 | 2119282.5 | 1650759.25 |
| Q3UIR3 | Dtx3l | E3 ubiquitin-protein ligase DTX3L Mus musculus | 64.55 | 2.14 | 83044 | 1 | 1 | 569658.75 | 527961.69 | 302946.41 | 348627.94 | 294791.12 | 378019.66 | 313629.59 | 369992.97 | 432832.88 | 247330.14 | 299648.62 | 380748.06 | 268053.94 | 407158.47 | 317304.94 |
| P97930 | Dtymk | Thymidylate kinase Mus musculus | 165.99 | 22.64 | 23915 | 3 | 3 | 2683525.5 | 3438320 | 3873143.25 | 3131515 | 3096032 | 2960943.75 | 2594649.25 | 3357529 | 3886438.5 | 3083866.5 | 3537131.5 | 2674802.25 | 3094574.5 | 3875648 | 3376953.5 |
| Q3KRC5 | Dus3l | tRNA-dihydrouridine(47) synthase [NAD(P)(+)]-like Rattus norvegicus | 142.11 | 6.88 | 71540 | 3 | 3 | 157501.19 | 827886.69 | 1165373.12 | 570771.88 | 584694.12 | 694548.5 | 499607.56 | 749834.25 | 797824.31 | 551254.06 | 1006208.5 | 763718 | 835026.62 | 1300165.5 | 885182.5 |
| Q9D7X3 | Dusp3 | Dual specificity protein phosphatase 3 Mus musculus | 502.78 | 50.81 | 20472 | 7 | 7 | 19808213 | 22795277 | 21355283 | 21772333 | 21776871 | 21922753 | 20370689 | 19384411 | 19447423 | 20835371 | 18704197 | 19281205 | 21246511 | 21740795 | 18597505 |
| P70583 | Dut | Deoxyuridine 5'-triphosphate nucleotidohydrolase Rattus norvegicus | 229.27 | 21.46 | 22003 | 4 | 4 | 2970678.75 | 3781558.5 | 4836467.5 | 4905225 | 5730771 | 4131204.5 | 5490524 | 4540580 | 4179694.75 | 5334950 | 6615971 | 5946253.5 | 5018719.5 | 2805847 | 7033411 |
| Q9JHU4 | Dync1h1 | Cytoplasmic dynein 1 heavy chain 1 Mus musculus | 7019.131 | 36.95 | 532045 | 129 | 129 | 286282433 | 273644129 | 270429345 | 240687649 | 255813489 | 259577105 | 260107233 | 256712417 | 294523169 | 255507041 | 271238721 | 272258913 | 264283425 | 271438081 | 281013889 |
| O88487 | Dync1i2 | Cytoplasmic dynein 1 intermediate chain 2 Mus musculus | 631.77 | 21.73 | 68394 | 7 | 7 | 7223076.5 | 7905716.5 | 8210516.5 | 5927793.5 | 8112328 | 8590854 | 7130686.5 | 9842209 | 8877212 | 9682635 | 7664622 | 10056371 | 8562511 | 8290870 | 10915903 |
| Q8R1Q8 | Dync1li1 | Cytoplasmic dynein 1 light intermediate chain 1 Mus musculus | 358.85 | 18.74 | 56614 | 7 | 7 | 14092548 | 11484200 | 11401878 | 11801322 | 12836591 | 12880294 | 13140464 | 13876277 | 12994403 | 12357918 | 15156407 | 13410214 | 10925107 | 13178313 | 14458305 |
| Q62698 | Dync1li2 | Cytoplasmic dynein 1 light intermediate chain 2 Rattus norvegicus | 282.53 | 18.51 | 54745 | 5 | 5 | 1057448 | 1081162.62 | 400488.09 | 996846.9275 | 1448289 | 386691.59 | 204054.31 | 1321379.38 | 887251.81 | 216532.69 | 760371.06 | 463536.19 | 321691.41 | 885684.88 | 881868.56 |
| P63168 | Dynll1 | Dynein light chain 1, cytoplasmic Mus musculus | 183.83 | 49.44 | 10366 | 1 | 3 | 8300341 | 7898261.5 | 5338728 | 5613036.5 | 7264544 | 6406377 | 6545873 | 6043722 | 7526831.5 | 4744572 | 7646279 | 6226542.5 | 6974217 | 6745477 | 4279864 |
| Q78P75 | Dynll2 | Dynein light chain 2, cytoplasmic Rattus norvegicus | 121.15 | 49.44 | 10350 | 1 | 3 | 6247390.5 | 5720783.5 | 3357348.5 | 4144981.5 | 5320865 | 4424558 | 4867762 | 4459442 | 5183772 | 3982791 | 5494330 | 3884522.25 | 5178203.5 | 4954675 | 2929464.25 |
| P51807 | Dynlt1 | Dynein light chain Tctex-type 1 Mus musculus | 216.52 | 44.25 | 12483 | 3 | 3 | 1041992.75 | 870806.56 | 694207.62 | 601861 | 634501.62 | 811508.88 | 911727.75 | 859014.81 | 551336.56 | 657037.88 | 1022827.69 | 839592 | 700293.81 | 954660.88 | 1081444.12 |
| Q9ESD7 | Dysf | Dysferlin Mus musculus | 123.45 | 1 | 237911 | 1 | 2 | 4066897.25 | 4145274.25 | 4969147.5 | 2661205.75 | 3306966.5 | 3418708.75 | 3164014.25 | 4456238 | 5154119.5 | 3042744.5 | 2582214 | 4393231.5 | 3955927.5 | 4181697.5 | 3078481 |
| Q9D903 | Ebna1bp2 | Probable rRNA-processing protein EBP2 Mus musculus E | 179.36 | 17.97 | 34703 | 4 | 4 | 1567059.38 | 2113400.5 | 1280972 | 2690933 | 1567930.5 | 3626022 | 2448969.25 | 2046143.25 | 2400951.5 | 3971941 | 2342895.5 | 1259683.75 | 3169222 | 2328240 | 1800673.12 |
| Q9CS74 | Ecd | Protein ecdysoneless homolog Mus musculus | 72.85 | 5.77 | 71710 | 2 | 2 | 1 | 161375.7 | 160322.73 | 190806.33 | 1 | 1 | 1 | 156042.27 | 1 | 1 | 1 | 1 | 187768.22 | 1 | 1 |
| O35459 | Ech1 | Delta(3,5)-Delta(2,4)-dienoyl-CoA isomerase, mitochondrial Mus musculus | 237.98 | 19.27 | 36118 | 5 | 5 | 4140307.5 | 3485194 | 2630151.5 | 2344078.5 | 5104339 | 2891698.5 | 3243883.5 | 2950035.5 | 4287979 | 2559261 | 4452054 | 2593523 | 2775075.5 | 3067560.5 | 2647048 |
| Q9D9V3 | Echdc1 | Ethylmalonyl-CoA decarboxylase Mus musculus | 151.89 | 17.7 | 35467 | 3 | 4 | 3508413.25 | 3236497 | 2768885.75 | 3074386.75 | 3010218.25 | 3427880.25 | 2954424 | 3361480.5 | 3563428 | 3601682 | 3594795.25 | 3229923.5 | 2868494.75 | 3425319.25 | 2460279.5 |
| Q8BH95 | Echs1 | Enoyl-CoA hydratase, mitochondrial Mus musculus | 221.95 | 18.28 | 31474 | 4 | 4 | 1069732 | 682670.5 | 2071879.5 | 1335962.62 | 1448323 | 1385298 | 3960613.25 | 1196995.38 | 813357.69 | 3407155.5 | 2293513.5 | 2440284.25 | 1585113.38 | 829855.81 | 3390665.75 |
| P42125 | Eci1 | Enoyl-CoA delta isomerase 1, mitochondrial Mus musculus | 457.5 | 34.26 | 32250 | 6 | 6 | 7640934 | 8610438 | 9263151 | 7298505 | 8384398 | 9668596 | 7458483 | 9175449 | 7307647 | 8854411 | 8421346 | 7781673 | 6858285.5 | 8717936 | 9350134 |
| Q9WUR2 | Eci2 | Enoyl-CoA delta isomerase 2, mitochondrial Mus musculus | 237.6 | 12.53 | 43268 | 4 | 4 | 8570729 | 2119102.5 | 1910965.75 | 2013180 | 2101377 | 1800238.25 | 2108965.5 | 8902391 | 1767095.88 | 2222092.5 | 1899727.38 | 1784926.25 | 2122903 | 1951040 | 2027026.5 |
| Q6PDI5 | Ecpas | Proteasome adapter and scaffold protein ECM29 Mus musculus | 1066.03 | 16.47 | 203703 | 19 | 19 | 12597425 | 11245704 | 10650420 | 10602916 | 12927743 | 13011791 | 9728648 | 11641369 | 12438392 | 9839442 | 11013252 | 9910179 | 11244616 | 12993304 | 9935898 |
| Q8K2D3 | Edc3 | Enhancer of mRNA-decapping protein 3 Mus musculus | 81.51 | 7.87 | 55957 | 2 | 2 | 176901.52 | 250268.17 | 273306.28 | 347917.59 | 300374.81 | 396691.78 | 243509.27 | 207054.94 | 228587.41 | 192292.22 | 235788.55 | 231783.98 | 338142.91 | 239338.28 | 261263.43 |
| Q3UJB9 | Edc4 | Enhancer of mRNA-decapping protein 4 Mus musculus | 374.94 | 9.39 | 152484 | 1 | 7 | 4353501 | 4966455 | 6571259 | 5920117.5 | 6745177.5 | 6231074 | 5474740 | 6368606.5 | 5819670.5 | 7306529.5 | 5291176.5 | 6800293 | 6204443 | 5090312 | 4410522 |
| Q8BL66 | Eea1 | Early endosome antigen 1 Mus musculus | 405.09 | 8.5 | 160915 | 8 | 8 | 1834322.75 | 1916873.25 | 1880146.88 | 3753542.75 | 2587111 | 3757032 | 5312151 | 1879134.25 | 2514839 | 2714621.25 | 2298934.25 | 1870276.25 | 2330425 | 2057744.75 | 1945638.88 |
| Q921E6 | Eed | Polycomb protein EED Mus musculus | 113.96 | 7.71 | 50198 | 3 | 3 | 1085072.12 | 1522864.88 | 1627659 | 2256247.5 | 2388143.5 | 1166829.12 | 2072834.75 | 1979911 | 1505900.25 | 2414290.25 | 2568860 | 1573355.25 | 1207893.88 | 2143724.25 | 2572625 |
| P10126 | Eef1a1 | Elongation factor 1-alpha 1 Mus musculus | 2132.89 | 61.26 | 50114 | 17 | 36 | 1027456385 | 965531201 | 1188962817 | 1044751617 | 929421057 | 988552065 | 1438577793 | 1027633089 | 995068481 | 1166322049 | 1165979649 | 1250625665 | 1024875009 | 963097409 | 1265033857 |
| P62631 | Eef1a2 | Elongation factor 1-alpha 2 Mus musculus | 988.28 | 29.16 | 50454 | 1 | 17 | 684303553 | 651505345 | 811386049 | 745664257 | 637148673 | 670874561 | 1088766465 | 671818177 | 618791489 | 887173825 | 806739777 | 872346433 | 727632193 | 648798337 | 904016513 |
| Q9D853 | Eef1akmt2 | EEF1A lysine methyltransferase 2 Mus musculus | 47.95 | 8.61 | 26864 | 1 | 1 | 1 | 202524.69 | 209280.45 | 1 | 1 | 1 | 170089.61 | 165186.06 | 1 | 156562.53 | 1 | 167999.19 | 1 | 125584.28 | 163269.14 |
| O70251 | Eef1b | Elongation factor 1-beta Mus musculus | 753.58 | 66.22 | 24694 | 10 | 12 | 36619877 | 39156001 | 42035241 | 37366705 | 33599133 | 38946589 | 40622309 | 43436565 | 39191461 | 38810393 | 40232689 | 38505809 | 36716033 | 37612369 | 39460729 |
| P57776 | Eef1d | Elongation factor 1-delta Mus musculus | 933.53 | 56.94 | 31293 | 12 | 15 | 102198297 | 121863249 | 108618625 | 112663921 | 113851521 | 107410745 | 132228961 | 110301513 | 114136105 | 122816041 | 121483361 | 113338233 | 111315841 | 107603321 | 109266657 |
| Q9D1M4 | Eef1e1 | Eukaryotic translation elongation factor 1 epsilon-1 Mus musculus | 98.65 | 20.11 | 19859 | 2 | 2 | 1216115.75 | 895468.0925 | 645879.75 | 441789.62 | 1278087.25 | 1062562.88 | 506088.62 | 490350.12 | 1983323 | 1276212.38 | 1913167.25 | 1425502.38 | 2052571.38 | 1163893.38 | 1987580.25 |
| Q9D8N0 | Eef1g | Elongation factor 1-gamma Mus musculus | 1580.78 | 77.8 | 50061 | 26 | 26 | 137479553 | 131329681 | 126536641 | 116026641 | 130042561 | 145175553 | 125069761 | 137537889 | 140070833 | 125265865 | 133138569 | 124911513 | 128539353 | 138842449 | 128232649 |
| P58252 | Eef2 | Elongation factor 2 Mus musculus | 3344.97 | 61.54 | 95314 | 50 | 52 | 326634849 | 330409569 | 354644641 | 332265505 | 309318529 | 333630721 | 392710305 | 367695649 | 369891297 | 342406049 | 362267553 | 376556641 | 344632257 | 381164993 | 371968961 |
| Q9JHW4 | Eefsec | Selenocysteine-specific elongation factor Mus musculus | 318.03 | 16.64 | 63539 | 7 | 7 | 1463852.38 | 1441641.12 | 1455018.88 | 2027843 | 1809071 | 2110254.25 | 1441660.25 | 2718685.75 | 1214587.75 | 1924854.75 | 2705174.25 | 2840779.5 | 1270309.5 | 856788.88 | 3077397.25 |
| Q3TGW2 | Eepd1 | Endonuclease/exonuclease/phosphatase family domain-containing protein 1 Mus musc | 41.23 | 2.64 | 62953 | 1 | 1 | 884563.31 | 910401.94 | 1 | 646673 | 1 | 749273.62 | 660041 | 650920.725 | 865283.06 | 329085.22 | 1683334.88 | 1045418.06 | 959576.62 | 958430.62 | 1021241.69 |
| Q9D8Y0 | Efhd2 | EF-hand domain-containing protein D2 Mus musculus | 850.28 | 68.33 | 26791 | 1 | 16 | 111222113 | 112214137 | 105129297 | 120351553 | 124314817 | 115744489 | 101407225 | 111051489 | 102849449 | 118708673 | 124463697 | 123864825 | 90048897 | 115767697 | 114112865 |
| Q8C0D5 | Efl1 | Elongation factor-like GTPase 1 Mus musculus | 96.08 | 2.13 | 125777 | 2 | 2 | 948018.19 | 1185063 | 1237860.62 | 1375599 | 1284588.88 | 1496708.25 | 1295633.38 | 1478575 | 1517997.12 | 1101325 | 1046779.06 | 1241035.25 | 1392874 | 1337582.75 | 1145200.5 |
| Q8BG67 | Efr3a | Protein EFR3 homolog A Mus musculus | 102.1 | 3.91 | 92613 | 2 | 2 | 246584.97 | 2561251 | 1414644.75 | 1100180.145 | 178239.86 | 164717.58 | 782717.81 | 224224.3 | 1772948 | 736151.9225 | 1464518.38 | 1758094.88 | 1545032.25 | 1796551 | 1819811.38 |
| O08810 | Eftud2 | 116 kDa U5 small nuclear ribonucleoprotein component Mus musculus | 1673.83 | 45.93 | 109361 | 31 | 31 | 31971007 | 28137881 | 35297185 | 29333603 | 27991733 | 29894177 | 33773073 | 31691963 | 36217833 | 29174101 | 34100213 | 41556513 | 31580147 | 35340201 | 37975081 |
| Q99MS7 | Ehbp1l1 | EH domain-binding protein 1-like protein 1 Mus musculus | 330.96 | 5.19 | 184834 | 6 | 6 | 3124244.5 | 2300516.5 | 3008927.75 | 2818594.25 | 3847507.25 | 2962062 | 3271362.5 | 2618166 | 2106421 | 2718013 | 3340761 | 2530082.5 | 2839797.25 | 2621632.75 | 2158993.25 |
| Q641Z6 | Ehd1 | EH domain-containing protein 1 Rattus norvegicus | 1135.41 | 48.69 | 60603 | 9 | 21 | 37865081 | 28337299 | 30381685 | 28628741 | 27820769 | 27437673 | 42763981 | 28406469 | 26365889 | 35112617 | 26820525 | 33403451 | 33717177 | 31412245 | 37053689 |

| | | | | | | | | | | | | | | | | | | | | | | |
|---|---|---|---|---|---|---|---|---|---|---|---|---|---|---|---|---|---|---|---|---|---|---|
| Q8R491 | Ehd3 | EH domain-containing protein 3 Rattus norvegicus | 724.66 | 27.29 | 60791 | 1 | 15 | 19499203 | 14338282 | 16051792 | 15090915 | 13285443 | 14660328 | 22835395 | 13591563 | 13684031 | 19995409 | 15443583 | 16791457 | 17943557 | 15560443 | 20912135 |
| Q9EQP2 | Ehd4 | EH domain-containing protein 4 Mus musculus | 1173.64 | 47.13 | 61481 | 17 | 21 | 45898105 | 35515837 | 35702145 | 34853389 | 35500921 | 33009973 | 45246945 | 34140721 | 33530283 | 41459077 | 32046611 | 38263273 | 40481617 | 33393349 | 42143453 |
| P48024 | Eif1 | Eukaryotic translation initiation factor 1 Mus musculus | 234.65 | 53.1 | 12747 | 4 | 4 | 931350.94 | 396710.88 | 481420.78 | 522596.165 | 280902.06 | 400860.41 | 971968.25 | 625156.19 | 620325.75 | 654577.65 | 813558.25 | 978712.19 | 233438.59 | 883032.94 | 757308.62 |
| Q60872 | Eif1a | Eukaryotic translation initiation factor 1A Mus musculus | 297.5 | 29.17 | 16502 | 1 | 4 | 6098777.5 | 3301234.25 | 4047254 | 4697406.5 | 5013342 | 4301072.5 | 4110551 | 3769832.5 | 4211102.5 | 4396386.5 | 5005828 | 4159851.5 | 3879382.5 | 4021641.5 | 5689492 |
| Q8BMJ3 | Eif1ax | Eukaryotic translation initiation factor 1A, X-chromosomal Mus musculus 2 | 343.49 | 36.11 | 16460 | 2 | 5 | 9468905 | 5668160 | 6419049.5 | 7456208 | 7460835 | 7488428 | 7470624 | 6295926.5 | 6273260.5 | 7307352.5 | 7668800 | 6993119 | 6426501 | 6493853 | 8742148 |
| Q8BJW6 | Eif2a | Eukaryotic translation initiation factor 2A Mus musculus | 567.33 | 28.92 | 64403 | 11 | 11 | 5282611 | 4372255 | 6031035 | 5546082.5 | 4430541 | 5659145 | 5064962.5 | 4471462.5 | 4841509.5 | 4872396 | 5623518 | 6987787.5 | 5400350 | 4394674 | 6246345 |
| Q03963 | Eif2ak2 | Interferon-induced, double-stranded RNA-activated protein kinase Mus musculus | 387.12 | 17.09 | 58280 | 6 | 6 | 6568350 | 3811978 | 3449479.75 | 2326365 | 2465058.5 | 3296991.75 | 2494083 | 4076407 | 3000736.5 | 4648989 | 3080305.5 | 4837162.5 | 4293698 | 3604824.5 | 7976602.5 |
| Q99LC8 | Eif2b1 | Translation initiation factor eIF-2B subunit alpha Mus musculus | 206.58 | 13.11 | 33816 | 3 | 3 | 1076993.88 | 3882521.25 | 3815415 | 2441191 | 2117378 | 3265333.5 | 1104364.25 | 3937776.75 | 7279476.5 | 2015577.62 | 5459045 | 5313450.5 | 1746876 | 5607129.5 | 3027706 |
| Q62818 | Eif2b2 | Translation initiation factor eIF-2B subunit beta Rattus norvegicus 2 | 93.04 | 8.55 | 38875 | 2 | 2 | 551711.5 | 161822.95 | 189998.22 | 226018.19 | 151747.2 | 175272.55 | 262965.91 | 144395.56 | 127176.88 | 220051.83 | 207945.77 | 184101.28 | 180050.5275 | 141882.67 | 186272.39 |
| P70541 | Eif2b3 | Translation initiation factor eIF-2B subunit gamma Rattus norvegicus 2 | 184.38 | 11.06 | 50435 | 4 | 4 | 3885131 | 5173464 | 5457932 | 4769345.5 | 4171754 | 5390274 | 3565980 | 5631591.5 | 6388286.5 | 5088536 | 4932376 | 5920931.5 | 4634867 | 3647511 | 5084026 |
| Q61749 | Eif2b4 | Translation initiation factor eIF-2B subunit delta Mus musculus | 270.95 | 14.31 | 57624 | 5 | 5 | 5332604 | 5539814 | 7310752 | 5345673 | 5296609 | 5617039 | 6195340 | 6820249.5 | 6444198 | 6324946.5 | 5998558 | 6740134 | 5325196 | 6405422 | 5202983 |
| Q8CHW4 | Eif2b5 | Translation initiation factor eIF-2B subunit epsilon Mus musculus | 252.21 | 9.07 | 80086 | 5 | 5 | 6108872 | 4428276.5 | 6308202 | 4816150 | 5639807 | 5866734 | 5794174 | 5443265.5 | 5973258.5 | 5321509 | 5800871.5 | 5699700 | 5761342.5 | 6922453 | 5500154.5 |
| P68101 | Eif2s1 | Eukaryotic translation initiation factor 2 subunit 1 Rattus norvegicus | 1198.18 | 70.79 | 36108 | 19 | 20 | 74835921 | 83887617 | 85171801 | 80947633 | 85725169 | 85271433 | 85907377 | 80851425 | 93575089 | 80214745 | 98936641 | 88981849 | 81685833 | 82156537 | 93923137 |
| Q99L45 | Eif2s2 | Eukaryotic translation initiation factor 2 subunit 2 Mus musculus | 736.98 | 45.92 | 38092 | 14 | 14 | 13474387 | 6245353 | 8476924 | 8165452.5 | 4590240.5 | 7355509 | 9964107 | 8826696 | 8531469 | 11753798 | 8560900 | 8389542 | 10310648 | 8516247 | 12785249 |
| Q9Z0N1 | Eif2s3x | Eukaryotic translation initiation factor 2 subunit 3, X-linked Mus musculus | 1176.57 | 54.66 | 51065 | 19 | 20 | 34462809 | 33809809 | 35290821 | 40723009 | 31704717 | 38866845 | 51969633 | 37268349 | 34357225 | 32357897 | 36793153 | 31942313 | 36324557 | 35252305 | 38485625 |
| P23116 | Eif3a | Eukaryotic translation initiation factor 3 subunit A Mus musculus | 1322.97 | 25.89 | 161936 | 29 | 29 | 57096769 | 44178701 | 37261657 | 41998193 | 48662077 | 46227633 | 29425167 | 43891617 | 49490617 | 41581273 | 44310521 | 38802973 | 41567941 | 45613777 | 38278745 |
| Q8JZQ9 | Eif3b | Eukaryotic translation initiation factor 3 subunit B Mus musculus | 1398.48 | 48.19 | 91370 | 27 | 27 | 26909591 | 23923837 | 28357913 | 21258719 | 20898801 | 26769833 | 24161721 | 27105581 | 30318959 | 25451925 | 25304511 | 26317189 | 21919213 | 23826597 | 26733553 |
| B5DFC8 | Eif3c | Eukaryotic translation initiation factor 3 subunit C Rattus norvegicus | 1050.91 | 23.05 | 105435 | 20 | 20 | 42405489 | 36620485 | 39037785 | 35756045 | 41643897 | 36601593 | 39569709 | 39994857 | 40314185 | 40431857 | 42894501 | 37810717 | 38046477 | 40243373 | 41155313 |
| O70194 | Eif3d | Eukaryotic translation initiation factor 3 subunit D Mus musculus | 736.8 | 40.69 | 63989 | 13 | 13 | 16592148 | 18260059 | 19410627 | 16976993 | 19256145 | 20410293 | 20258221 | 19063727 | 18957449 | 19752059 | 17639235 | 18010525 | 19842167 | 18509211 | 23299001 |
| P60229 | Eif3e | Eukaryotic translation initiation factor 3 subunit E Mus musculus | 1219.69 | 54.16 | 52221 | 20 | 20 | 40375681 | 33781045 | 46782873 | 37196785 | 31846987 | 33984965 | 38077677 | 42057857 | 49353237 | 30718929 | 43975813 | 42882821 | 32564673 | 43991585 | 43163793 |
| Q9DCH4 | Eif3f | Eukaryotic translation initiation factor 3 subunit F Mus musculus | 896.39 | 48.75 | 37984 | 13 | 13 | 35004189 | 38736741 | 47615741 | 29747607 | 28078943 | 38742265 | 36153973 | 45540765 | 46461821 | 35263725 | 45732289 | 55116485 | 30381789 | 44319993 | 48688649 |
| Q9Z1D1 | Eif3g | Eukaryotic translation initiation factor 3 subunit G Mus musculus | 445.95 | 36.88 | 35638 | 9 | 9 | 22536869 | 21529325 | 26223417 | 25183975 | 24896031 | 21765389 | 26085885 | 21855323 | 19425323 | 26051749 | 33148325 | 24714665 | 24064869 | 20628575 | 24795611 |
| Q91WK2 | Eif3h | Eukaryotic translation initiation factor 3 subunit H Mus musculus | 774.68 | 52.27 | 39832 | 13 | 13 | 28118545 | 27135455 | 24421703 | 25477219 | 31439245 | 26874373 | 24836099 | 22525113 | 22313347 | 28470427 | 32545599 | 24099647 | 27389517 | 24818091 | 26313037 |
| Q9QZD9 | Eif3i | Eukaryotic translation initiation factor 3 subunit I Mus musculus | 737.39 | 64.92 | 36461 | 15 | 15 | 13739393 | 9488356 | 13646486 | 11548385 | 12774445 | 14819657 | 12234821 | 15855371 | 12510104 | 10751065 | 14650276 | 14085490 | 12254731 | 15380618 | 13957963 |
| Q3UGC7 | Eif3j1 | Eukaryotic translation initiation factor 3 subunit J-A Mus musculus 2 | 397.51 | 39.85 | 29344 | 8 | 8 | 6638919.5 | 8447682 | 5934762 | 7575206 | 8514462 | 10026749 | 5726272 | 7388883 | 9551444 | 7515288.5 | 6886103 | 7254439.5 | 7178577 | 9621770 | 6187941 |
| Q9DBZ5 | Eif3k | Eukaryotic translation initiation factor 3 subunit K Mus musculus | 282.58 | 27.98 | 25087 | 4 | 4 | 11773104 | 10850116 | 12716353 | 6303225 | 11711519 | 11754002 | 10123177 | 10323354 | 13251058 | 10306643 | 13232828 | 12300806 | 10901514 | 11319676 | 13540218 |
| Q8QZY1 | Eif3l | Eukaryotic translation initiation factor 3 subunit L Mus musculus | 989.73 | 37.06 | 66613 | 16 | 17 | 25309131 | 20410773 | 20383457 | 17605967 | 19905763 | 20526443 | 16227024 | 23336109 | 25245127 | 17901027 | 21726811 | 21957955 | 19809955 | 23108807 | 23350099 |
| Q99JX4 | Eif3m | Eukaryotic translation initiation factor 3 subunit M Mus musculus | 811.31 | 53.21 | 42517 | 12 | 12 | 15678228 | 13660635 | 15983906 | 13005503 | 10542719 | 17678875 | 13486972 | 10482984 | 15103219 | 12687666 | 14356876 | 16393093 | 15741886 | 14577091 | 16834057 |
| P60843 | Eif4a1 | Eukaryotic initiation factor 4A-I Mus musculus | 2206.6 | 83.25 | 46154 | 18 | 32 | 325793025 | 335565217 | 354217793 | 289204417 | 307334721 | 306750753 | 283940225 | 323638081 | 332415969 | 299011905 | 361690049 | 333512097 | 303993729 | 348110881 | 302615649 |
| P10630 | Eif4a2 | Eukaryotic initiation factor 4A-II Mus musculus | 938.42 | 33.42 | 46402 | 1 | 15 | 216110977 | 216853345 | 236527617 | 198028017 | 208413969 | 203532609 | 186717873 | 208324737 | 209725761 | 202729345 | 248980817 | 216053537 | 203146689 | 225434529 | 196969921 |
| Q91VC3 | Eif4a3 | Eukaryotic initiation factor 4A-III Mus musculus | 1315.96 | 59.61 | 46840 | 20 | 23 | 60105225 | 72170793 | 70375913 | 67179833 | 72956833 | 69072761 | 73364409 | 62472885 | 69620057 | 73335769 | 73965753 | 72591009 | 70213089 | 69209537 | 70662769 |
| Q8BGD9 | Eif4b | Eukaryotic translation initiation factor 4B Mus musculus | 1121.46 | 37.97 | 68840 | 19 | 19 | 26457303 | 19061721 | 21141321 | 25850855 | 22335831 | 28704847 | 23133677 | 27179369 | 22415391 | 25357341 | 26836741 | 26421615 | 30109869 | 22604137 | 26024005 |
| P63073 | Eif4e | Eukaryotic translation initiation factor 4E Mus musculus | 221.31 | 21.66 | 25053 | 5 | 5 | 10787272 | 7963477 | 8493354 | 12022545 | 8465217 | 10306722 | 12576000 | 10346052 | 8138502 | 10042454 | 7239332 | 7948714.5 | 9489239 | 9803756 | 10247599 |
| Q8BMB3 | Eif4e2 | Eukaryotic translation initiation factor 4E type 2 Mus musculus | 34.37 | 4.49 | 28263 | 1 | 1 | 1081350.5 | 1431085.12 | 1438545.88 | 1158269.62 | 1095246.75 | 1178353.25 | 1273885.03 | 1370204.25 | 1288272.5 | 1258710.12 | 782422.31 | 1036210.31 | 1122945.88 | 1250634 | 1100586.38 |
| Q6NZJ6 | Eif4g1 | Eukaryotic translation initiation factor 4 gamma 1 Mus musculus | 2441.32 | 30.88 | 176077 | 36 | 39 | 60170577 | 60680209 | 52271645 | 45380969 | 48350205 | 52738113 | 56043501 | 59534393 | 63853229 | 57414761 | 61569945 | 57067569 | 56454181 | 55904253 | 54736857 |
| Q62448 | Eif4g2 | Eukaryotic translation initiation factor 4 gamma 2 Mus musculus | 1138.33 | 36.09 | 102106 | 24 | 24 | 22325795 | 16805373 | 20646151 | 14888943 | 16771886 | 17749511 | 18253047 | 21478015 | 24822875 | 16455199 | 18324057 | 24579501 | 16154436 | 23850783 | 19968585 |
| Q80XI3 | Eif4g3 | Eukaryotic translation initiation factor 4 gamma 3 Mus musculus | 492.37 | 6.84 | 174890 | 7 | 10 | 14518640 | 13511615 | 14310582 | 11453613 | 10705974 | 13791869 | 12223135 | 14172137 | 16341624 | 10955109 | 12385858 | 13383316 | 11344409 | 15355313 | 12224529 |
| Q5XI72 | Eif4h | Eukaryotic translation initiation factor 4H Rattus norvegicus | 652.42 | 49.6 | 27324 | 11 | 11 | 23070013 | 32763969 | 22664043 | 25098147 | 26287501 | 25202943 | 26674013 | 23756941 | 31044705 | 26956265 | 26160005 | 22415163 | 25208691 | 24601369 | 22721013 |
| P59325 | Eif5 | Eukaryotic translation initiation factor 5 Mus musculus | 290.59 | 13.29 | 48968 | 5 | 5 | 10179051 | 8915562 | 9307121 | 9392106 | 10261104 | 11499940 | 9794546 | 10299813 | 7875977 | 10524850 | 9100531 | 9866274 | 10595963 | 11794707 | 8731362 |
| P63242 | Eif5a | Eukaryotic translation initiation factor 5A-1 Mus musculus | 1045.08 | 88.31 | 16832 | 14 | 15 | 125870945 | 127178625 | 152176417 | 127524241 | 88316369 | 138939345 | 155347121 | 151038017 | 174460145 | 139928065 | 188930113 | 149585169 | 92411873 | 157447409 | 156220897 |
| Q05D44 | Eif5b | Eukaryotic translation initiation factor 5B Mus musculus | 1257.04 | 32.24 | 137616 | 25 | 25 | 27146141 | 22656091 | 19270283 | 24558887 | 22207337 | 22595049 | 26485489 | 30984479 | 27806255 | 26389447 | 23986833 | 29153983 | 25724687 | 25954267 | 30827891 |
| O55135 | Eif6 | Eukaryotic translation initiation factor 6 Mus musculus | 504.25 | 46.53 | 26511 | 8 | 8 | 16486477 | 17500049 | 18771293 | 19021457 | 16594559 | 18406735 | 18447323 | 15667871 | 19947929 | 19243585 | 19222411 | 20159489 | 19510487 | 18996821 | 17033205 |
| Q80Y81 | Elac2 | Zinc phosphodiesterase ELAC protein 2 Mus musculus | 254.86 | 8.66 | 92719 | 6 | 6 | 6394434 | 6164484.5 | 5424469 | 6997979 | 6631873 | 5494917.5 | 5500029.5 | 6378114 | 6597293.5 | 6237981 | 6476727 | 5659564 | 6748754 | 4736724 | 5931182.5 |
| B5DF91 | Elavl1 | ELAV-like protein 1 Rattus norvegicus | 600.85 | 38.65 | 36169 | 10 | 10 | 24938427 | 24757395 | 27992295 | 29491109 | 24272397 | 23461669 | 26615533 | 24146117 | 24182269 | 28232797 | 27622481 | 25527005 | 25301949 | 23170825 | 25364941 |
| O08856 | Ell | RNA polymerase II elongation factor ELL Mus musculus | 67.85 | 1.83 | 67146 | 1 | 1 | 285321.78 | 228550.61 | 198114.03 | 274333.495 | 385347.56 | 261922 | 1 | 281062.69 | 1 | 1 | 347768.81 | 1 | 171179.67 | 207318.39 | 1 |
| Q8BPU7 | Elmo1 | Engulfment and cell motility protein 1 Mus musculus | 636.57 | 24.9 | 83936 | 12 | 13 | 12739519 | 14630181 | 15591896 | 15240503 | 16312340 | 12947941 | 12989691 | 13452189 | 14303376 | 14878008 | 17440023 | 13871717 | 14943385 | 15084368 | 13161749 |
| Q8BHL5 | Elmo2 | Engulfment and cell motility protein 2 Mus musculus | 164.3 | 7.38 | 83887 | 3 | 4 | 2126161.25 | 1153252.12 | 1210852.62 | 2955921 | 1700594.5 | 1919486.25 | 2230292.75 | 2122116.5 | 1688135.5 | 1825678.5 | 1618496.12 | 2449509.75 | 2488600.75 | 2687477 | 1472673.25 |
| Q8CB77 | Eloa | Elongin-A Mus musculus | 105.93 | 5.3 | 87161 | 2 | 2 | 861514 | 854246.44 | 685421.5 | 931225.75 | 914140.5 | 878217.56 | 742036.12 | 739629 | 865681.44 | 625875.06 | 901842.25 | 781796 | 920560.62 | 830336.81 | 853877.38 |
| P62869 | Elob | Elongin-B Mus musculus | 420.08 | 59.32 | 13170 | 8 | 8 | 40089149 | 39224705 | 40423361 | 46266561 | 39398197 | 37587689 | 43538977 | 39705173 | 35892905 | 43679421 | 39733733 | 38047089 | 41740989 | 36190745 | 34442481 |
| P83940 | Eloc | Elongin-C Mus musculus | 207.44 | 30.36 | 12473 | 2 | 2 | 3720203.5 | 3612083.5 | 3997336 | 4232823 | 4573965 | 4880331.5 | 5640385 | 4332781 | 3288922.5 | 6379224.5 | 4664807.5 | 4821539 | 4504654 | 3931125.5 | 6074387 |
| Q7TT37 | Elp1 | Elongator complex protein 1 Mus musculus | 267.45 | 4.65 | 149584 | 5 | 5 | 2316355 | 1619828.5 | 1629935.12 | 1721657.25 | 2597709.5 | 2420220.75 | 2322106.75 | 2087143.75 | 2786238 | 2703746.5 | 1400302.88 | 1484372 | 1134585.25 | 1689847 | 2752083 |
| Q9CZX0 | Elp3 | Elongator complex protein 3 Mus musculus | 259.32 | 11.88 | 62385 | 5 | 5 | 950490.5 | 482626.75 | 1815457.38 | 1871549.75 | 1444335.88 | 662905.94 | 3350875.5 | 717361.5 | 1266033.5 | 1784258.5 | 1868791.5 | 1829042.88 | 1352502.62 | 1492717 | 3105797.75 |
| Q8C7X2 | Emc1 | ER membrane protein complex subunit 1 Mus musculus | 706.36 | 19.36 | 111605 | 12 | 12 | 11590910 | 9152788 | 9775971 | 9215133 | 8609014 | 12716892 | 12560624 | 10386655 | 11325880 | 11244435 | 10678737 | 11651833 | 10988752 | 11440125 | 9343595 |
| Q5U2V8 | Emc3 | ER membrane protein complex subunit 3 Rattus norvegicus | 153.74 | 14.18 | 29980 | 3 | 3 | 3914306 | 5538289 | 4979377.5 | 4787448 | 3768954.75 | 5020724 | 5271960.5 | 5001991 | 5173018 | 6195509 | 3865886.75 | 3866610.5 | 4550641 | 3922141.5 | 3847495.25 |
| Q9CZX9 | Emc4 | ER membrane protein complex subunit 4 Mus musculus | 53.73 | 8.74 | 20117 | 1 | 1 | 3121233 | 3703782.5 | 3606992.75 | 4100907.5 | 3399092.25 | 3726315.5 | 3168380.5 | 3552956.75 | 3091991.5 | 3497489.5 | 3015585 | 2461886.25 | 3068528.25 | 3331299.75 | 3229043 |
| O70378 | Emc8 | ER membrane protein complex subunit 8 Mus musculus | 272.81 | 26.09 | 23348 | 4 | 4 | 6220320 | 7126992 | 6707797 | 6752939.5 | 5990222 | 6409999 | 6840782 | 6983173.5 | 7335282.5 | 7050405 | 6284305 | 6644389.5 | 6862570 | 7857440 | 5745389 |
| O08579 | Emd | Emerin Mus musculus | 347.14 | 41.7 | 29436 | 7 | 7 | 4667484.5 | 6607882.5 | 5862455 | 8795026 | 4948861 | 6970468 | 7700050 | 6324215 | 5600790 | 6980001.5 | 5660783 | 5403940 | 6324380 | 5109461.5 | 5491339 |
| O35130 | Emg1 | Ribosomal RNA small subunit methyltransferase NEP1 Mus musculus | 399.82 | 33.61 | 26974 | 6 | 6 | 3260497 | 2907007 | 4997991.5 | 2031453.5 | 3379251.5 | 3585765 | 6716789 | 4448584 | 3973134.5 | 3230620.75 | 3756508 | 7346980 | 2591314.5 | 3985720.75 | 3798895.5 |
| Q6P6T4 | Eml2 | Echinoderm microtubule-associated protein-like 2 Rattus norvegicus | 151.15 | 7.7 | 70711 | 3 | 3 | 5018700.938 | 6989340 | 7348600 | 2637560.75 | 3099303 | 9533779 | 8244659 | 8465422 | 8236636 | 9192467 | 10269151 | 3617734.5 | 8022556 | 7789995.5 | 8097664.5 |
| Q8VC03 | Eml3 | Echinoderm microtubule-associated protein-like 3 Mus musculus | 41.79 | 1.78 | 95695 | 1 | 1 | 1 | 1 | 320697.59 | 1 | 275340.22 | 229971.88 | 542201.38 | 1 | 218796.33 | 1 | 311464.5 | 332518.34 | 358659.03 | 352938.695 | 409112.91 |
| Q3UMY5 | Eml4 | Echinoderm microtubule-associated protein-like 4 Mus musculus | 82.13 | 1.62 | 110027 | 1 | 1 | 1 | 1 | 1 | 1 | 1 | 1337647.12 | 868349 | 530229.88 | 575253.5 | 713995 | 719144.62 | 913039.44 | 572256.31 | 1 | 1 |
| Q8BX80 | Engase | Cytosolic endo-beta-N-acetylglucosaminidase Mus musculus | 38.33 | 2.45 | 82945 | 1 | 1 | 630382.69 | 406771.66 | 582075.31 | 577889.12 | 318679.78 | 543110.18 | 357745.59 | 336047.69 | 584101.75 | 894545.69 | 345782.12 | 357100.56 | 374015.09 | 430166.84 | 432859 |
| P17182 | Eno1 | Alpha-enolase Mus musculus | 2394.93 | 77.42 | 47141 | 24 | 32 | 483027841 | 544320321 | 569124609 | 442800673 | 497630657 | 598201857 | 853236929 | 634970049 | 668755073 | 674073537 | 642606273 | 680968833 | 657382657 | 604103617 | 606306369 |
| P17183 | Eno2 | Gamma-enolase Mus musculus | 645.04 | 23.73 | 47297 | 2 | 7 | 73958097 | 73393017 | 84455273 | 71131409 | 67966833 | 76439761 | 153707633 | 81093729 | 60248977 | 121620017 | 89566849 | 96576849 | 94393841 | 67226737 | 93338161 |
| P15429 | Eno3 | Beta-enolase Rattus norvegicus | 565.86 | 29.26 | 47014 | 3 | 7 | 47131237 | 45882161 | 56982809 | 44746569 | 45084993 | 47866073 | 91130753 | 52101669 | 39132217 | 72189625 | 58181897 | 60887513 | 57517257 | 42066841 | 58038405 |
| Q8BGB7 | Enoph1 | Enolase-phosphatase E1 Mus musculus | 101.97 | 5.45 | 28600 | 2 | 2 | 1674472 | 514303.97 | 1643080.12 | 1983792.75 | 2291924 | 2070084.38 | 1718757.38 | 1427574.88 | 1425810.75 | 1797253.5 | 1309571.5 | 1739078.75 | 1891359.5 | 662238.81 | 533726.12 |
| Q8BTJ4 | Enpp4 | Bis(5'-adenosyl)-triphosphatase enpp4 Mus musculus | 99.91 | 4.82 | 51611 | 2 | 2 | 985076.69 | 1300517.25 | 1556378.12 | 1119300 | 1032830.94 | 1056971.88 | 892532.19 | 1048898.5 | 2425360 | 977536.94 | 1048328.31 | 870920.31 | 919050.75 | 1244061.75 | 1006300.12 |
| Q5NDL0 | Eogt | EGF domain-specific O-linked N-acetylglucosamine transferase Rattus norvegicus | 98.17 | 3.42 | 61524 | 2 | 2 | 1099925 | 2217043 | 2163710.25 | 1962838.38 | 1994900.62 | 2020897.25 | 2067710 | 2223503.5 | 2794494.75 | 2179595.25 | 2201223 | 2105901 | 2017428.25 | 2088845.25 | 1738973 |
| P97481 | Epas1 | Endothelial PAS domain-containing protein 1 Mus musculus | 34.03 | 0.8 | 96712 | 1 | 1 | 2291132.25 | 1687931.88 | 1177006.25 | 2503481.5 | 2628611.75 | 1435919.25 | 1274467.38 | 1163758.75 | 488025.5 | 1864217.5 | 1543764.75 | 459869.75 | 1435132.62 | 876038 | 885240.25 |
| P48193 | Epb41 | Protein 4.1 Mus musculus | 115.17 | 5.48 | 95911 | 3 | 3 | 2396828.5 | 1270234.38 | 2874250.5 | 3088874.75 | 3092124 | 2966239.5 | 3283475.75 | 2538733.25 | 2341838 | 3518524.5 | 2832361.75 | 2944574 | 2764398.75 | 2869861.75 | 3355180 |
| O70318 | Epb41l2 | Band 4.1-like protein 2 Mus musculus | 413.11 | 12.55 | 109940 | 8 | 8 | 7989059 | 4522140.5 | 5149627 | 7183709 | 9050957 | 7656614 | 9611260 | 7559136.5 | 8147104.5 | 8128090 | 8863043 | 6324543 | 5654183 | 6558867.5 | 7503347.5 |
| P58107 | EPPK1 | Epiplakin Homo sapiens OX=9606 | 213.95 | 2.95 | 555658 | 1 | 4 | 20477909 | 21831199 | 18154501 | 22418725 | 20974463 | 20832709 | 17785021 | 21375441 | 21082905 | 16647991 | 23774137 | 19904593 | 20237329 | 21370937 | 17507863 |
| Q8CGC7 | Eprs | Bifunctional glutamate/proline--tRNA ligase Mus musculus | 3129.3 | 44.97 | 170079 | 53 | 53 | 86093945 | 90823649 | 85946025 | 96304833 | 92856417 | 95480777 | 106109473 | 88575537 | 85943505 | 104969521 | 92361825 | 84959945 | 89311993 | 86354233 | 101776009 |
| P42567 | Eps15 | Epidermal growth factor receptor substrate 15 Mus musculus | 88.53 | 3.34 | 98471 | 2 | 2 | 781225.5 | 753682.5 | 836381.38 | 758778.5 | 593401 | 595565.94 | 649961.88 | 1215903.62 | 633807.12 | 634025.25 | 1034338.81 | 1366345.5 | 716639.56 | 871609.25 | 585669.06 |
| Q60902 | Eps15l1 | Epidermal growth factor receptor substrate 15-like 1 Mus musculus | 416.16 | 10.8 | 99309 | 8 | 8 | 1579996.25 | 631869.94 | 4871119.5 | 6067258.5 | 1889298.5 | 1351709.12 | 2146588.5 | 1364203.5 | 7695402 | 1933885.5 | 2841930.5 | 5903150 | 1777556.38 | 2074027.62 | 7951277 |
| Q08509 | Eps8 | Epidermal growth factor receptor kinase substrate 8 Mus musculus | 139.73 | 5.24 | 91737 | 2 | 2 | 1 | 1 | 1 | 1 | 1 | 1 | 1 | 1 | 1 | 1 | 616116.81 | 777872.88 | 975023.06 | 673069.81 | 760520.64 |
| Q9EQH2 | Erap1 | Endoplasmic reticulum aminopeptidase 1 Mus musculus | 946.18 | 26.34 | 106599 | 20 | 20 | 32018785 | 26761759 | 35973577 | 28325499 | 29424585 | 32323711 | 30123449 | 29919069 | 32171071 | 29950013 | 30651929 | 30985947 | 29246071 | 28702713 | 30900265 |
| Q80TH2 | Erbin | Erbin Mus musculus | 190.2 | 3.99 | 157248 | 3 | 3 | 1450685.75 | 1594659.12 | 1688859.75 | 1646393.25 | 1693545.5 | 1715269.5 | 1374004.75 | 1638350.88 | 1455976.88 | 1390422.25 | 1746605.88 | 1880624 | 1197821.62 | 1433319.75 | 1896968.38 |
| P07903 | Ercc1 | DNA excision repair protein ERCC-1 Mus musculus | 57.37 | 7.05 | 32910 | 1 | 1 | 1 | 1 | 1 | 101706.07 | 117672.57 | 1 | 1 | 99952.3 | 1 | 1 | 1 | 107693.48 | 1 | 1 | 1 |
| Q8BHK9 | Ercc6l | DNA excision repair protein ERCC-6-like Mus musculus | 45.2 | 1.21 | 138854 | 1 | 1 | 221364.56 | 403636.19 | 266241.91 | 396379.38 | 415197.25 | 332338.28 | 347575.88 | 358863.12 | 309778.41 | 351088.38 | 365093.75 | 415418.75 | 282510.72 | 322701.59 | 406109.44 |
| Q9ERY9 | Erg28 | Probable ergosterol biosynthetic protein 28 Mus musculus | 72.04 | 9.29 | 15806 | 1 | 1 | 666625.88 | 1048970.25 | 532570.44 | 500417.12 | 608880.25 | 799483.25 | 721442 | 567459.31 | 526238.69 | 651213.62 | 865679.19 | 670380.56 | 681331.69 | 554797.25 | 591001.38 |
| Q9DC16 | Ergic1 | Endoplasmic reticulum-Golgi intermediate compartment protein 1 Mus musculus | 77.85 | 6.21 | 32562 | 2 | 2 | 3226603 | 3379903.5 | 3210803.25 | 3267271.5 | 3166848 | 3181764 | 3601628.5 | 3769343 | 3422206.5 | 3596184 | 2691115 | 3083740.25 | 3087197 | 3004658.25 | 2982803.5 |
| P84089 | Erh | Enhancer of rudimentary homolog Mus musculus | 247.72 | 45.19 | 12259 | 4 | 4 | 9151626 | 11230285 | 8897056 | 8986972 | 10046686 | 8500464 | 7166663 | 9157656 | 9922256 | 8503789 | 9081167 | 6445727 | 7422902 | 8507233 | 7052146 |
| Q7TMF2 | Eri1 | 3'-5' exoribonuclease 1 Mus musculus | 105.39 | 6.96 | 39493 | 2 | 2 | 3691419 | 4465231 | 4722021 | 4387705 | 4198916 | 3952562.75 | 3683769.5 | 4741760 | 4936630 | 3690399 | 4219767 | 3485125 | 3810747.5 | 4126091.75 | 3600163.25 |
| Q8BFZ9 | Erlin2 | Erlin-2 Mus musculus | 206.7 | 17.65 | 37873 | 5 | 5 | 1397666 | 1118907 | 1771453.12 | 1481336.12 | 724465.38 | 954483.69 | 959106.56 | 1444425 | 1825291.5 | 1040618.31 | 834856.06 | 1751576.75 | 1062440.88 | 1056594.62 | 1506642.75 |
| Q3UVK0 | Ermp1 | Endoplasmic reticulum metallopeptidase 1 Mus musculus | 81.45 | 3.67 | 100148 | 2 | 2 | 177502.88 | 426273.2925 | 204563.97 | 918256.38 | 404769.94 | 486296.81 | 443030.97 | 105731.16 | 379656.03 | 914494.69 | 371607.2525 | 117564.33 | 837586.88 | 209819.55 | 321458.25 |
| Q8R180 | Ero1a | ERO1-like protein alpha Mus musculus | 1208.87 | 53.66 | 54084 | 17 | 19 | 34275897 | 37585565 | 33318769 | 30588371 | 32604637 | 46392561 | 59067661 | 50951521 | 56834589 | 49306309 | 41812017 | 39553205 | 38353937 | 44429225 | 42118393 |
| Q8R2E9 | Ero1b | ERO1-like protein beta Mus musculus | 314.36 | 12.85 | 53518 | 3 | 5 | 6508726 | 6918732 | 7956128 | 5295514.5 | 5025839.5 | 9151859 | 12265912 | 9667431 | 12705076 | 8577916 | 7859587.5 | 7979355.5 | 6369195 | 8053586 | 7467939.5 |
| P57759 | Erp29 | Endoplasmic reticulum resident protein 29 Mus musculus | 421.21 | 42.75 | 28823 | 7 | 7 | 23509187 | 25599647 | 25032001 | 19475835 | 23330189 | 20610617 | 20067849 | 26469735 | 27931169 | 20734189 | 22253321 | 24299801 | 23471831 | 25400501 | 21393193 |
| Q9D1Q6 | Erp44 | Endoplasmic reticulum resident protein 44 Mus musculus | 170.88 | 12.32 | 46853 | 4 | 4 | 759549.19 | 1013525.75 | 4785617.5 | 711205.12 | 1054735.25 | 4530434 | 781554.12 | 5324436.5 | 6084133.5 | 871411.88 | 517135.97 | 5130856.5 | 1195439.25 | 5490764 | 5307954 |
| Q9R0P3 | Esd | S-formylglutathione hydrolase Mus musculus | 964.57 | 64.89 | 31320 | 13 | 14 | 83114041 | 96321745 | 83802617 | 68926065 | 83589385 | 99679953 | 89278369 | 103742641 | 102332025 | 82160497 | 82338473 | 72737153 | 95881633 | 95968265 | 70264337 |
| Q3V1V3 | Esf1 | ESF1 homolog Mus musculus | 43.06 | 1.42 | 98048 | 1 | 1 | 295701.62 | 221579.8 | 296168.06 | 246615.015 | 173010.58 | 211978.69 | 308541.88 | 312044.56 | 452822.75 | 215109.88 | 306219.19 | 391339.5 | 285005.38 | 381102.28 | 420882.66 |
| P60330 | Espl1 | Separin Mus musculus | 109.06 | 1.56 | 233035 | 3 | 3 | 392002.09 | 188101.06 | 4667197.5 | 7043474.5 | 270140.88 | 191356.69 | 902061.25 | 412073.5 | 3235102.25 | 161634.69 | 152641.97 | 1997731.5 | 167668.16 | 3678977.5 | 6184648 |
| Q3U7R1 | Esyt1 | Extended synaptotagmin-1 Mus musculus | 848.32 | 17.12 | 121554 | 14 | 14 | 10172406 | 12263189 | 11941009 | 13940559 | 12000911 | 11330781 | 11736290 | 13624962 | 13633776 | 12142643 | 8972478 | 11915535 | 12454079 | 10988209 | 10303387 |
| Q3TZZ7 | Esyt2 | Extended synaptotagmin-2 Mus musculus | 119.37 | 4.5 | 94139 | 2 | 2 | 1064953.5 | 1177318.88 | 1586444.38 | 1074387.38 | 1557714.5 | 1514774.62 | 1509527.88 | 1875835.25 | 1541051.75 | 1869469 | 1393590.62 | 1603252.62 | 1513588 | 1278637.75 | 1203392 |
| Q5U2Q7 | Etf1 | Eukaryotic peptide chain release factor subunit 1 Rattus norvegicus | 1018.21 | 59.5 | 49031 | 18 | 18 | 16706434 | 18603379 | 22570753 | 18915315 | 16865905 | 21865413 | 19680507 | 24615825 | 30455587 | 19475593 | 20928809 | 23151867 | 19234061 | 25609017 | 23104641 |
| Q99LC5 | Etfa | Electron transfer flavoprotein subunit alpha, mitochondrial Mus musculus | 782.9 | 54.65 | 35009 | 13 | 13 | 57741413 | 66087065 | 70525881 | 65579277 | 65035497 | 59722145 | 83542521 | 66542325 | 63184201 | 66033765 | 59543613 | 63048341 | 67361441 | 63281569 | 52688829 |
| Q68FU3 | Etfb | Electron transfer flavoprotein subunit beta Rattus norvegicus | 567.95 | 36.86 | 27687 | 1 | 11 | 47981317 | 51292781 | 52961961 | 57274581 | 57576025 | 54098497 | 55713341 | 54497509 | 52514121 | 56474905 | 51194041 | 47670333 | 54683733 | 52780329 | 43025713 |
| Q9DCM0 | Ethe1 | Persulfide dioxygenase ETHE1, mitochondrial Mus musculus | 166.01 | 24.41 | 27739 | 4 | 4 | 8303762.5 | 9284230 | 9074427 | 10790023 | 10753311 | 8005515 | 8971851 | 7274087 | 6298631 | 8053064 | 9034530 | 6495023 | 8276262.5 | 8036088.5 | 8133424 |
| P97360 | Etv6 | Transcription factor ETV6 Mus musculus | 149.74 | 7.84 | 56406 | 3 | 3 | 1 | 119053.66 | 201921.03 | 1 | 1 | 1 | 428947 | 1 | 146286.77 | 1 | 1 | 1 | 258410.89 | 1 | 85393.64 |
| Q8VD58 | Evi2b | Protein EVI2B Mus musculus | 39.26 | 2.48 | 47993 | 1 | 1 | 3684902.25 | 2832548.5 | 2309091.5 | 1882541 | 2129549.25 | 1985683.75 | 2133556.938 | 2478218 | 2743180.25 | 1327145.75 | 1955819.25 | 2050267.12 | 1903019.5 | 2631792.5 | 1310739 |
| Q61545 | Ewsr1 | RNA-binding protein EWS Mus musculus | 292.15 | 12.21 | 68462 | 6 | 6 | 9790247 | 11386937 | 9336300 | 11701854 | 11464452 | 8871031 | 9649932 | 7776542.5 | 8169679.5 | 8879421 | 10322543 | 9094059 | 9754542 | 9174569 | 8572034 |
| Q9D4H1 | Exoc2 | Exocyst complex component 2 Mus musculus | 65.3 | 2.06 | 103959 | 2 | 2 | 1514566.12 | 1450794.75 | 1546134.88 | 1230913.12 | 1554094.12 | 1182017.38 | 1295753.62 | 1375678.283 | 1722235.88 | 1302706.25 | 1655269.25 | 1376527.62 | 1281562.88 | 1216487.75 | 1312652.12 |
| O35382 | Exoc4 | Exocyst complex component 4 Mus musculus | 290.01 | 6.77 | 110545 | 5 | 5 | 1848533.75 | 2479471 | 2428685 | 2457477 | 1848064.5 | 2303435.5 | 2152696.75 | 2243815 | 2951396.5 | 2159452 | 2567044.5 | 2606946.5 | 1732859 | 2425875.5 | 1782962.25 |
| Q3TPX4 | Exoc5 | Exocyst complex component 5 Mus musculus | 39.09 | 1.27 | 81738 | 1 | 1 | 1582412.25 | 1 | 1420503.88 | 415890.53 | 1 | 984522.88 | 1 | 1615828.25 | 1846380.5 | 1 | 935079.94 | 1271338.25 | 381862.66 | 1574968.62 | 1454480.12 |

| | | | | | | | | | | | | | | | | | | | | | | |
|---|---|---|---|---|---|---|---|---|---|---|---|---|---|---|---|---|---|---|---|---|---|---|
| O54923 | Exoc6 | Exocyst complex component 6 Rattus norvegicus | 100.95 | 2.49 | 93178 | 2 | 2 | 610581.38 | 603179.94 | 577174.06 | 650635.94 | 571622.25 | 613465.5 | 657084.12 | 525294.56 | 569305.19 | 696878.56 | 582763.69 | 474384.19 | 549699.44 | 473388.62 | 643432.62 |
| A6H5Z3 | Exoc6b | Exocyst complex component 6B Mus musculus | 104.09 | 3.95 | 94130 | 2 | 2 | 650293.88 | 556235.75 | 483403 | 618612.25 | 364234.78 | 510866.69 | 473100.03 | 635152.88 | 556686.88 | 458151.59 | 650885.75 | 512464.09 | 438386.78 | 628145.25 | 504904.19 |
| O54924 | Exoc8 | Exocyst complex component 8 Rattus norvegicus | 39.47 | 2.23 | 81043 | 1 | 1 | 1680718 | 2204113.75 | 1868724.5 | 1831553.12 | 1860770.5 | 1780783.12 | 1816859.25 | 1680740.62 | 1433218.25 | 1869701 | 1627611.38 | 1515096.25 | 1721683.12 | 1598812 | 1366646.75 |
| P56960 | Exosc10 | Exosome component 10 Mus musculus | 395.79 | 11.61 | 100942 | 8 | 8 | 3946711 | 5951535.5 | 5056039.5 | 4645242.5 | 5054771.5 | 5241410 | 3431025.25 | 4606383 | 4735302.5 | 3811241.75 | 3484003 | 5984884 | 3951719 | 5260951 | 5017781 |
| Q8VBV3 | Exosc2 | Exosome complex component RRP4 Mus musculus | 199.32 | 21.84 | 32632 | 4 | 4 | 2140369 | 2067023.62 | 1843501.5 | 4046490 | 2557314.25 | 1341222.62 | 2260987 | 1405346.12 | 1807680.75 | 4979916 | 1438546.5 | 1624044.88 | 1918860.25 | 5022327 | 1230731 |
| Q7TQK4 | Exosc3 | Exosome complex component RRP40 Mus musculus | 126.95 | 12.41 | 29546 | 2 | 2 | 1306568 | 1349430.5 | 2213209.5 | 1403089.38 | 1568074.345 | 1923700 | 1706774 | 1609940.88 | 846345.81 | 1650254.75 | 1910955.75 | 1831344.25 | 1466975.25 | 2017272 | 1780688.88 |
| Q921I9 | Exosc4 | Exosome complex component RRP41 Mus musculus | 60.6 | 4.9 | 26250 | 1 | 1 | 1 | 1 | 1 | 1 | 354221.69 | 1 | 876910.44 | 1 | 291807.81 | 1 | 345001.19 | 601768.56 | 465370.28 | 378441.03 | 625084.06 |
| Q9CRA8 | Exosc5 | Exosome complex component RRP46 Mus musculus | 145.76 | 13.62 | 25194 | 3 | 3 | 4624671.5 | 5615735 | 5979488 | 4829062.5 | 4547397 | 4046922 | 6653946 | 4802192 | 5532450 | 5144125 | 4008857 | 6667982 | 5440666 | 4778224 | 4595480 |
| Q9D0M0 | Exosc7 | Exosome complex exonuclease RRP42 Mus musculus | 187.06 | 19.24 | 31825 | 4 | 4 | 2964501.75 | 3029343.5 | 2762999.5 | 2791264 | 1334217 | 2347566 | 2355381 | 2641961.25 | 3016254.5 | 2749992 | 2958484.75 | 2777387 | 2713242.5 | 2877870.5 | 2462789 |
| Q9D753 | Exosc8 | Exosome complex component RRP43 Mus musculus | 213.44 | 24.28 | 29949 | 4 | 4 | 2567726 | 4718768 | 5453596.5 | 5592656.5 | 5231408.5 | 4948565 | 6167405 | 4014770.5 | 5895199 | 5011290.5 | 5382764.5 | 5547431.5 | 5792462.5 | 4226594.5 | 5693148 |
| Q9JHI7 | Exosc9 | Exosome complex component RRP45 Mus musculus | 282.82 | 13.47 | 48937 | 4 | 4 | 4550806 | 6657117 | 4695681 | 6797694 | 5475161 | 6594016 | 7012315 | 6058328 | 5227190.5 | 6112529 | 4636340 | 4795645 | 7038900 | 4965650 | 6256091 |
| P26040 | Ezr | Ezrin Mus musculus | 1125.14 | 34.13 | 69407 | 9 | 21 | 422243009 | 401172545 | 355826241 | 385307809 | 379793089 | 404867105 | 421201345 | 422890017 | 380058273 | 391309505 | 337121857 | 366715585 | 363425537 | 427346689 | 331877473 |
| P04117 | Fabp4 | Fatty acid-binding protein, adipocyte Mus musculus | 270.09 | 51.52 | 14650 | 6 | 6 | 14000153 | 14500083 | 15950010 | 6045960 | 11049351 | 7806180 | 5398586 | 15886175 | 21164209 | 10199754 | 7480256 | 7733770 | 13360821 | 16961701 | 10715658 |
| Q05816 | Fabp5 | Fatty acid-binding protein 5 Mus musculus | 1005.52 | 79.26 | 15137 | 13 | 16 | 367597665 | 362926977 | 390861889 | 341166817 | 309307073 | 341351137 | 282154497 | 353476769 | 386754945 | 286768897 | 362900289 | 391429281 | 341515041 | 413924353 | 319789505 |
| Q61160 | Fadd | FAS-associated death domain protein Mus musculus | 155.35 | 17.56 | 22960 | 2 | 2 | 948964.56 | 794998.44 | 736284 | 765240.405 | 580714.62 | 634150.38 | 719814.44 | 767128.19 | 785754.25 | 861784.81 | 678515.38 | 614329.06 | 663417.19 | 764970 | 575692.81 |
| Q920L1 | Fads1 | Fatty acid desaturase 1 Mus musculus | 88.65 | 6.26 | 52323 | 2 | 2 | 535070.75 | 1 | 1 | 1 | 1 | 133491.52 | 1 | 1 | 1 | 141469 | 109925.62 | 94175.37 | 115857.58 | 112273.94 | 262167.12 |
| Q9Z0R9 | Fads2 | Fatty acid desaturase 2 Mus musculus | 123.95 | 6.76 | 52387 | 2 | 3 | 392607.81 | 1 | 371367.59 | 1 | 1 | 1 | 1 | 1 | 447989.84 | 1 | 309193.62 | 338681.75 | 1 | 313293.09 | 1 |
| P54731 | Faf1 | FAS-associated factor 1 Mus musculus | 138.61 | 5.7 | 73863 | 3 | 3 | 1324567.75 | 794858.38 | 1910030 | 1 | 661915.69 | 953464.62 | 1498141.62 | 1951429.38 | 1785968 | 639239.44 | 1694422.75 | 1318934.12 | 672417.94 | 1245402.12 | 1709104.5 |
| Q3TDN2 | Faf2 | FAS-associated factor 2 Mus musculus | 150.93 | 12.58 | 52471 | 3 | 3 | 1774138.75 | 1 | 1315747.25 | 1 | 1 | 1088093.5 | 1945971.5 | 1467639.88 | 1403694.5 | 1756710.25 | 426513.56 | 921990 | 1083758.39 | 1188050.25 | 1798479.75 |
| P25093 | Fah | Fumarylacetoacetase Rattus norvegicus | 142.45 | 14.8 | 45976 | 3 | 3 | 633424.75 | 1208979.5 | 1571140 | 1002720 | 718923.38 | 1626286 | 1824892.75 | 1586687.25 | 668764.81 | 1346865.25 | 1257514.62 | 1371398.62 | 1474103.38 | 1605132 | 2093907.62 |
| Q9WUD8 | Faim | Fas apoptotic inhibitory molecule 1 Mus musculus | 108.3 | 13.97 | 20201 | 2 | 2 | 1509374.88 | 1934835.75 | 721436.44 | 948193.38 | 1248292.75 | 2372160.5 | 853865.31 | 1202087.38 | 2213676.5 | 1660447.423 | 519324.12 | 1164984.67 | 1874881 | 1711302.62 | 554430.94 |
| Q8VE88 | Fam114a2 | Protein FAM114A2 Mus musculus | 121.26 | 6.04 | 54045 | 2 | 2 | 701795.5 | 651817.5 | 774680.12 | 421943 | 1631702.25 | 1505522.75 | 779538.19 | 1185947.25 | 1807686.38 | 1273667.38 | 845165.88 | 1028720.88 | 876661.12 | 731601.44 | 1591447 |
| Q6A0A9 | FAM120A | Constitutive coactivator of PPAR-gamma-like protein 1 Mus musculus | 274.76 | 6.65 | 121646 | 6 | 6 | 3361158 | 3630206 | 3036079.75 | 1367072.12 | 3035716 | 2000739.25 | 2268727.75 | 2501100 | 5024286 | 5322940 | 1845198.62 | 2037186.62 | 5444205 | 4669540 | 2179477 |
| Q3UW53 | Fam129a | Protein Niban Mus musculus | 1269.04 | 33.26 | 120649 | 23 | 23 | 37346441 | 32697601 | 38217685 | 38209249 | 31493591 | 38220337 | 36828413 | 37287573 | 35379089 | 37781097 | 36834809 | 38067629 | 34627305 | 35039709 | 39653557 |
| Q8R1F1 | Fam129b | Niban-like protein 1 Mus musculus | 513.98 | 19.36 | 84819 | 9 | 9 | 21481605 | 20756937 | 21639297 | 18641639 | 18686853 | 19548591 | 21482379 | 22030755 | 21892651 | 22717279 | 19625161 | 19666543 | 15003076 | 19499501 | 23531539 |
| Q4QQV3 | Fam162a | Protein FAM162A Rattus norvegicus | 35.31 | 4.52 | 17827 | 1 | 1 | 421356.06 | 463096.21 | 430423.62 | 530424.94 | 470180.22 | 1 | 968590.5 | 1 | 398163.28 | 563122.62 | 510870.88 | 467323.25 | 564802.62 | 390943.72 | 520981.56 |
| Q91WE2 | Fam192a | PSME3-interacting protein Mus musculus | 49.63 | 8.66 | 28702 | 2 | 2 | 1037927.298 | 1135491.5 | 760348.69 | 1142106.25 | 1113762.75 | 1075547.75 | 709672.44 | 1054961 | 900327.5 | 868825.38 | 1079521 | 937730.88 | 984169.235 | 1008652.56 | 910772.5 |
| Q8CGI1 | Fam193a | Protein FAM193A Mus musculus | 35.38 | 0.81 | 136714 | 1 | 1 | 4412990 | 2779078.25 | 2425787 | 4287065.5 | 4312787 | 2320984.5 | 2365210.75 | 2306710.75 | 2495928.313 | 2990807.25 | 3153871.5 | 965172.31 | 1854834.88 | 2244710 | 2275938.5 |
| Q91VU0 | Fam3c | Protein FAM3C Mus musculus | 272.59 | 24.23 | 24753 | 4 | 4 | 3161076.75 | 3497232.25 | 3286509.25 | 3563666.75 | 3167611.75 | 3266087.5 | 2717533 | 3816119.5 | 2984135.5 | 3008745.5 | 3425647 | 3307697.75 | 3493411.75 | 3220939.25 | 2798750.5 |
| Q8BHZ0 | Fam49a | Protein FAM49A Mus musculus | 265.66 | 15.79 | 37343 | 2 | 4 | 20813073 | 23976357 | 20775189 | 20002189 | 23508867 | 21569331 | 19628199 | 22338541 | 22986499 | 22940925 | 21453365 | 22052221 | 24210941 | 24292779 | 19358503 |
| Q921M7 | Fam49b | Protein FAM49B Mus musculus | 1135.29 | 73.46 | 36776 | 17 | 19 | 52385545 | 53019641 | 50129689 | 44368365 | 50591877 | 49663761 | 50360925 | 52564653 | 54341853 | 46083357 | 46881857 | 49657953 | 51431165 | 56273785 | 46680197 |
| Q9WV03 | Fam50a | Protein FAM50A Mus musculus | 96.5 | 7.96 | 40252 | 2 | 2 | 1302013.25 | 465316.88 | 1157272.75 | 1587247.62 | 536215.12 | 848905.19 | 1925207.25 | 1444827.75 | 1338505.75 | 633321.19 | 1405985.25 | 1521014.25 | 1430157.75 | 1212088.5 | 2176871.75 |
| Q3TJZ6 | Fam98a | Protein FAM98A Mus musculus | 109.07 | 10.1 | 55055 | 2 | 3 | 1 | 323133.53 | 773351.38 | 482654.88 | 1 | 369119.59 | 727159.25 | 717876.88 | 529659.75 | 276255.41 | 1 | 642997.12 | 1 | 532882.12 | 491987.69 |
| Q80VD1 | Fam98b | Protein FAM98B Mus musculus | 231.79 | 15.85 | 45349 | 4 | 5 | 2202856.75 | 6076433 | 6850369 | 2981043 | 4320574 | 5830100 | 4917365 | 5785931 | 6818055 | 4628249 | 4268370.5 | 6223020 | 4234901 | 6097159 | 5720902 |
| Q922J9 | Far1 | Fatty acyl-CoA reductase 1 Mus musculus | 493.86 | 24.27 | 59435 | 9 | 9 | 18782985 | 12180763 | 12503131 | 11126202 | 14845694 | 13199593 | 8077743.5 | 12365055 | 15235503 | 10326685 | 12100120 | 11008039 | 11178495 | 14314301 | 11242265 |
| Q8C0C7 | Farsa | Phenylalanine--tRNA ligase alpha subunit Mus musculus | 594.17 | 23.03 | 57599 | 4 | 11 | 7553203 | 9834617 | 6850016 | 8593985 | 9612251 | 8935149 | 9539320 | 6943509 | 7844542 | 6482970 | 7331553.5 | 5957245.5 | 7453859 | 9357354 | 6170206.5 |
| Q9WUA2 | Farsb | Phenylalanine--tRNA ligase beta subunit Mus musculus | 868.51 | 39.56 | 65697 | 20 | 20 | 38717701 | 36137033 | 39857009 | 33997365 | 34118893 | 36924053 | 40073825 | 37317797 | 38360809 | 36570585 | 29922631 | 34844569 | 36405273 | 39079501 | 37811329 |
| P19096 | Fasn | Fatty acid synthase Mus musculus | 6246.8 | 61.86 | 272428 | 51 | 103 | 346178113 | 337950113 | 356189569 | 357735041 | 323721473 | 327054241 | 320900865 | 282894625 | 316224033 | 317165665 | 366675169 | 358927713 | 327721153 | 350089505 | 382199969 |
| P62862 | Fau | 40S ribosomal protein S30 Mus musculus | 94.78 | 18.64 | 6648 | 2 | 2 | 2374680.75 | 2146666.25 | 4830429 | 3816281.25 | 4727740 | 4324942 | 9719509 | 3234647.25 | 2939858.5 | 8485817 | 6693220 | 8018881.5 | 6764168 | 3156220 | 9703033 |
| P22509 | Fbl | rRNA 2'-O-methyltransferase fibrillarin Rattus norvegicus | 758.15 | 51.68 | 34222 | 9 | 13 | 11956617 | 12822240 | 16228307 | 13784236 | 12760894 | 13181497 | 20823661 | 13239790 | 11874036 | 17723375 | 15883009 | 17930829 | 14596052 | 12232495 | 19683323 |
| Q80WS3 | Fbll1 | rRNA/tRNA 2'-O-methyltransferase fibrillarin-like protein 1 Mus musculus | 193.75 | 11.46 | 33339 | 1 | 3 | 1879900.25 | 1368595.75 | 2122934.75 | 1937742.38 | 1605991 | 1864271.62 | 4867352 | 1611543.62 | 1087538.75 | 3330552.75 | 2259709.25 | 3231078.5 | 1793277.5 | 1494584.12 | 4436144.5 |
| Q9CZV8 | Fbxl20 | F-box/LRR-repeat protein 20 Mus musculus | 134.55 | 4.59 | 48396 | 2 | 2 | 1667723.12 | 1688105 | 1393354.75 | 845205.38 | 1029334.38 | 1666217.88 | 1867442 | 1473351.88 | 1500564 | 1028016.31 | 991737.62 | 822416.75 | 813035.88 | 1404302.38 | 1211586 |
| Q78JE5 | Fbxo22 | F-box only protein 22 Mus musculus | 226.06 | 24.38 | 44203 | 5 | 5 | 5727194 | 6290344.5 | 3759569.75 | 5147191.5 | 5312682.5 | 3455177.5 | 5062270.5 | 6014210.5 | 4049164 | 3716560.75 | 3376224.5 | 5038292 | 3654351.75 | 5644988 | 2838616 |
| Q9QZN4 | Fbxo6 | F-box only protein 6 Mus musculus | 34.09 | 3.39 | 34493 | 1 | 1 | 323568.19 | 250572.7 | 270163.19 | 281705.925 | 282519.62 | 310744.53 | 174462.64 | 277887.44 | 422050.25 | 296286.215 | 261319.94 | 345043.72 | 315860.445 | 349806.53 | 307271.59 |
| P20491 | Fcer1g | High affinity immunoglobulin epsilon receptor subunit gamma Mus musculus | 102.69 | 15.12 | 9652 | 2 | 2 | 3695508.25 | 2696233.25 | 4694493.5 | 3697796.5 | 5004032.5 | 3191050.25 | 3484394.75 | 2782449.25 | 2394742 | 2963159.063 | 3888945 | 2825169.25 | 4002238.25 | 383381.44 | 4494791 |
| P26151 | Fcgr1 | High affinity immunoglobulin gamma Fc receptor I Mus musculus | 275.57 | 22.77 | 44888 | 6 | 6 | 10544918 | 9299983 | 11888508 | 9437379 | 9035087 | 7002304.5 | 8227308 | 6693892.5 | 8077838.5 | 7228624 | 8291136 | 11055223 | 9099139 | 10037064 | 8133480 |
| P08101 | Fcgr2 | Low affinity immunoglobulin gamma Fc region receptor II Mus musculus | 176.94 | 11.21 | 36695 | 2 | 3 | 10047039 | 11144037 | 9987515 | 11168614 | 10237842 | 10315674 | 10276124 | 9847845 | 9114443 | 9896937 | 11139311 | 10876050 | 11913535 | 8923250 | 9255644 |
| P08508 | Fcgr3 | Low affinity immunoglobulin gamma Fc region receptor III Mus musculus | 190.29 | 16.09 | 30036 | 2 | 3 | 8633707 | 10154023 | 8342685.5 | 11564095 | 10968568 | 10210729 | 9930411 | 9121871 | 7859605 | 9624853 | 9541746 | 7276502 | 11037044 | 9024543 | 8448317 |
| A0A0B4 | Fcgr4 | Low affinity immunoglobulin gamma Fc region receptor IV Mus musculus | 45.23 | 4.02 | 28398 | 1 | 1 | 2495273 | 2113805.5 | 2117607.5 | 2473083 | 2382982.25 | 1806752 | 1644024.25 | 1732195.12 | 1702798.62 | 2170689.5 | 1926141.75 | 1631899.75 | 2058816.12 | 1699792 | 1765408.12 |
| D3ZYR1 | Fcho2 | F-BAR domain only protein 2 Rattus norvegicus | 115.95 | 3.83 | 88712 | 2 | 2 | 830894.94 | 883108.62 | 884735.75 | 744802.12 | 843027.44 | 1072914.5 | 966837.38 | 1172649.12 | 1471196.38 | 685448 | 970138.62 | 947084.31 | 987039.62 | 1045488.12 | 991401.75 |
| P53798 | Fdft1 | Squalene synthase Mus musculus | 184 | 9.86 | 48154 | 3 | 3 | 221825.56 | 1 | 1 | 316029.34 | 1 | 1 | 1 | 1 | 1 | 1 | 221829.25 | 1 | 1 | 1 | 182980.41 |
| Q920E5 | Fdps | Farnesyl pyrophosphate synthase Mus musculus | 1360.53 | 65.44 | 40582 | 20 | 21 | 184978865 | 157284449 | 182328657 | 169005761 | 156573297 | 180643905 | 180883649 | 169329393 | 171165217 | 171301041 | 192430577 | 203757601 | 162155457 | 188879729 | 209346849 |
| Q61578 | Fdxr | NADPH:adrenodoxin oxidoreductase, mitochondrial Mus musculus | 42.92 | 2.43 | 54202 | 1 | 1 | 1531694.25 | 1348532.75 | 1561898.88 | 1238214 | 2042713 | 1560948.5 | 1250240.38 | 1761743.88 | 2032807.75 | 1044475 | 1303655.25 | 1473437 | 1452312.75 | 1507527 | 2950264.75 |
| P39749 | Fen1 | Flap endonuclease 1 Mus musculus | 507.06 | 29.1 | 42315 | 9 | 9 | 20376619 | 16223644 | 14952076 | 17508535 | 15983569 | 17210517 | 16706858 | 17572557 | 15253387 | 17002533 | 17015587 | 16750761 | 14694451 | 17945981 | 18151991 |
| Q8K1B8 | Fermt3 | Fermitin family homolog 3 Mus musculus | 1846.33 | 60.15 | 75635 | 29 | 29 | 128422081 | 140474417 | 129359985 | 123999265 | 132227425 | 124964481 | 107730505 | 126575273 | 145321681 | 116872145 | 127136233 | 127050737 | 127812177 | 136235073 | 117121833 |
| P16879 | Fes | Tyrosine-protein kinase Fes/Fps Mus musculus | 35.95 | 1.46 | 93779 | 1 | 1 | 2004771.88 | 1651227.75 | 1695138.62 | 1868781.12 | 1642265.75 | 869185.25 | 1557708.753 | 1608757.88 | 2010727.88 | 1742164 | 395239 | 1599948 | 1653506.62 | 1560964.75 | 1514985 |
| O88842 | Fgd3 | FYVE, RhoGEF and PH domain-containing protein 3 Mus musculus | 113.28 | 4.09 | 80624 | 2 | 2 | 373792.97 | 382110 | 424591.03 | 406523.38 | 442048.72 | 481925.34 | 272183.06 | 382563.97 | 396512.03 | 320130 | 577629.31 | 337087.66 | 482339.78 | 549618.31 | 486668.765 |
| P97807 | Fh | Fumarate hydratase, mitochondrial Mus musculus | 1200.89 | 50.3 | 54357 | 17 | 19 | 86103625 | 83282137 | 88677953 | 80313417 | 82207745 | 89665089 | 109420321 | 96921553 | 86762569 | 88193537 | 81870185 | 91613777 | 85385977 | 86667217 | 86248009 |
| Q9R059 | Fhl3 | Four and a half LIM domains protein 3 Mus musculus | 227.87 | 19.03 | 31795 | 4 | 4 | 6045332.5 | 10425108 | 4754440 | 8569628 | 8681190 | 7539521.5 | 9198893 | 7985081.5 | 7342059 | 7648437.5 | 8517477 | 4763013 | 8788201 | 8287186 | 8775005 |
| Q6P9Q4 | Fhod1 | FH1/FH2 domain-containing protein 1 Mus musculus | 213.66 | 6.43 | 129600 | 5 | 5 | 3240982 | 2549733 | 2016947.75 | 2310462 | 2904079.5 | 3798339.5 | 3323964.25 | 2542040.25 | 3687376 | 2698747 | 3415920 | 2275540.5 | 2384818.75 | 3298350.75 | 3217131.25 |
| Q5U317 | Fip1l1 | Pre-mRNA 3'-end-processing factor FIP1 Rattus norvegicus | 224.01 | 13.62 | 60188 | 5 | 5 | 6427216 | 425029.22 | 277373.94 | 324814.94 | 306673.88 | 486592.31 | 1264669.12 | 765019 | 1126768.5 | 867372.5 | 472067.19 | 1926519.25 | 1065602.25 | 487219.66 | 2060065.5 |
| P84817 | Fis1 | Mitochondrial fission 1 protein Rattus norvegicus | 381.05 | 60.53 | 16995 | 6 | 6 | 14938037 | 9948465 | 9113640 | 8535140 | 8707892 | 7611154 | 7030542.5 | 6185563 | 9838641 | 7676634.5 | 7941659.5 | 8234960 | 9345029 | 9073769 | 4173943 |
| Q6P9Q6 | Fkbp15 | FK506-binding protein 15 Mus musculus | 71.65 | 0.9 | 132961 | 1 | 1 | 850937.69 | 875626.44 | 835598.31 | 831313.94 | 1455444.62 | 858840.94 | 520773.69 | 698489.5 | 775362.19 | 1003612.25 | 1051147.5 | 642920.19 | 931934.38 | 893426.56 | 844183.94 |
| P26883 | Fkbp1a | Peptidyl-prolyl cis-trans isomerase FKBP1A Mus musculus | 89.35 | 12.96 | 11923 | 1 | 1 | 3188589.5 | 2845916.5 | 2171562.25 | 3069237.5 | 3164370 | 3074279.5 | 4756014 | 2991823.25 | 2328316.25 | 3433569 | 2680988.5 | 3315011 | 3467787.25 | 2690670.75 | 2842459.75 |
| P45878 | Fkbp2 | Peptidyl-prolyl cis-trans isomerase FKBP2 Mus musculus | 125.57 | 19.29 | 15344 | 2 | 2 | 3244866 | 3025755.5 | 3779809.5 | 4243410.5 | 2996151 | 2733578.75 | 2386004.5 | 2459459.25 | 2125663 | 2787699.75 | 2336236.5 | 2201685.5 | 2678028.75 | 2236693.25 | 1730822.25 |
| Q62446 | Fkbp3 | Peptidyl-prolyl cis-trans isomerase FKBP3 Mus musculus | 46.87 | 4.91 | 25148 | 1 | 1 | 1944479.88 | 1016109.56 | 1316950 | 1529155.88 | 1503591.5 | 1491718 | 1411647.5 | 1262794.5 | 1056103.5 | 1625300.25 | 1308129.5 | 1113982.5 | 993797.88 | 1154138.25 | 1509092.88 |
| P30416 | Fkbp4 | Peptidyl-prolyl cis-trans isomerase FKBP4 Mus musculus | 1578 | 71.4 | 51572 | 27 | 27 | 63051041 | 57052389 | 60547837 | 56181509 | 58160157 | 67333433 | 68149817 | 60666165 | 66362621 | 56937637 | 63708801 | 64043349 | 54857621 | 59009137 | 62486377 |
| Q64378 | Fkbp5 | Peptidyl-prolyl cis-trans isomerase FKBP5 Mus musculus | 386.09 | 20.18 | 50966 | 8 | 8 | 6534787 | 8083692 | 9353432 | 8812209 | 9785943 | 8093821 | 10808657 | 8750878 | 8318993 | 10718577 | 10298433 | 9518801 | 9127921 | 7834670 | 9439170 |
| O35465 | Fkbp8 | Peptidyl-prolyl cis-trans isomerase FKBP8 Mus musculus | 522.65 | 29.1 | 43529 | 8 | 8 | 13277937 | 14322264 | 15320881 | 9608378 | 11283122 | 13820792 | 14712469 | 10054988 | 14991922 | 16717014 | 13952921 | 13789933 | 14064385 | 13250356 | 13503496 |
| Q8QZS3 | Flcn | Folliculin Mus musculus | 43.05 | 2.25 | 64327 | 1 | 1 | 235372.88 | 227347.08 | 1 | 1 | 210333.64 | 162474.89 | 188774.22 | 192436.62 | 294409 | 242190.56 | 256815.69 | 278214.16 | 217110.31 | 252109.97 | 251062.5325 |
| Q9JJ28 | Flii | Protein flightless-1 homolog Mus musculus | 1236.46 | 25.96 | 144803 | 23 | 23 | 33758693 | 35097469 | 24368847 | 33923405 | 33465589 | 32317555 | 29413665 | 31802795 | 34178101 | 26246255 | 32484221 | 36278693 | 27562447 | 33648729 | 35598589 |
| Q8BTM8 | Flna | Filamin-A Mus musculus | 5048.24 | 49.87 | 281222 | 80 | 84 | 369365921 | 351985377 | 418238977 | 354114593 | 340416193 | 337599777 | 334478689 | 328578017 | 331724321 | 332812321 | 304110753 | 310819073 | 362230049 | 334443937 | 349253377 |
| Q80X90 | Flnb | Filamin-B Mus musculus | 556.33 | 5.88 | 277825 | 8 | 11 | 16415002 | 14864654 | 17498361 | 15034736 | 14546203 | 18010651 | 13273053 | 15534447 | 14791514 | 14630182 | 14718791 | 13525170 | 14407211 | 14585579 | 15547450 |
| Q60634 | Flot2 | Flotillin-2 Mus musculus | 245.6 | 10.28 | 47038 | 4 | 4 | 765938.69 | 1848567.12 | 2336999.75 | 1 | 1162812.25 | 2029125.12 | 1032645.69 | 2425122 | 3293469.75 | 1397779.88 | 1983869.62 | 2583230.75 | 1996804.5 | 2044655.5 | 1680937 |
| Q9JL26 | Fmnl1 | Formin-like protein 1 Mus musculus | 1753.47 | 41.04 | 122060 | 30 | 35 | 57177841 | 58380137 | 60175849 | 68217953 | 54049305 | 55358689 | 63206233 | 58703593 | 47989625 | 64073273 | 72004369 | 57731141 | 57363857 | 50847497 | 61242297 |
| A2APV2 | Fmnl2 | Formin-like protein 2 Mus musculus | 296.73 | 6.45 | 123101 | 0 | 7 | 8938427 | 9728446 | 8357934 | 9296369 | 7466358.5 | 7636926.5 | 10669830 | 8158807.5 | 8583556 | 9918209 | 7501978 | 7150295 | 7096853 | 8122109 | 7521136.5 |
| Q6ZPF4 | Fmnl3 | Formin-like protein 3 Mus musculus | 324.93 | 8.27 | 117169 | 2 | 8 | 4148227 | 4034978.5 | 2735316 | 3048927 | 3106976 | 3535639 | 6457932.5 | 3596133 | 3762953.5 | 5133449.5 | 2458102.5 | 2692054 | 2487650.5 | 3821910 | 3486428 |
| P35922 | Fmr1 | Synaptic functional regulator FMR1 Mus musculus | 128.12 | 3.58 | 68989 | 1 | 2 | 8126656.5 | 8517745 | 8133283 | 8277533 | 8584711 | 9212033 | 9449282 | 9479262 | 9120482 | 9621187 | 9348197 | 8808661 | 8166096.5 | 7797563.5 | 7339911.5 |
| Q80TY0 | Fnbp1 | Formin-binding protein 1 Mus musculus | 797.37 | 28.57 | 71344 | 16 | 16 | 21962303 | 26344899 | 22767065 | 27794309 | 26876757 | 21370501 | 24292207 | 22858141 | 24516189 | 23674537 | 24718989 | 22715929 | 24276855 | 25655871 | 22542077 |
| Q2HWF0 | Fnbp1l | Formin-binding protein 1-like Rattus norvegicus | 101.73 | 3.47 | 69974 | 1 | 1 | 250212.36 | 1 | 169275.83 | 1 | 157250.34 | 1 | 1 | 1 | 1 | 183285.53 | 152306.12 | 179624.0475 | 167648.44 | 225191.75 | 173349.88 |
| Q6NWW9 | Fndc3b | Fibronectin type III domain-containing protein 3B Mus musculus | 137.45 | 2.57 | 132764 | 2 | 2 | 239133.61 | 324202 | 728307.06 | 279746.22 | 794567 | 846021.44 | 961582.19 | 321376.16 | 327865.84 | 908318.38 | 943348.5 | 898717.5 | 793548.56 | 775130.12 | 885799.94 |
| Q8K385 | FRRS1 | Ferric-chelate reductase 1 Mus musculus | 280.32 | 16.05 | 66047 | 5 | 5 | 7680971 | 8136465 | 7587437.5 | 6404462 | 7100279.5 | 7277554 | 4835877 | 7123532 | 9535531 | 5451019 | 6963862 | 7690502.5 | 6941023 | 7880678 | 5606483.5 |
| A2ARZ3 | Fsip2 | Fibrous sheath-interacting protein 2 Mus musculus | 40.85 | 0.1 | 784857 | 1 | 1 | 2031017 | 1698953.5 | 2305389.75 | 2006768.25 | 2115784.25 | 2226417 | 2532949.25 | 2846962.25 | 1817552.5 | 2299334.5 | 2099180.25 | 1992462.12 | 2320365 | 1693386 | 1847227.25 |
| P09528 | Fth1 | Ferritin heavy chain Mus musculus | 284.6 | 49.45 | 21067 | 7 | 7 | 30465959 | 35178017 | 27363913 | 23865913 | 23612213 | 36197889 | 28883209 | 36758805 | 42761513 | 27617529 | 25029359 | 21444361 | 28673671 | 33833061 | 18572271 |
| P29391 | Ftl1 | Ferritin light chain 1 Mus musculus | 527.69 | 57.38 | 20802 | 9 | 9 | 79946201 | 83928089 | 69717713 | 67122257 | 64559313 | 90455649 | 82218393 | 95403641 | 104411329 | 76863825 | 73641625 | 63990453 | 78471425 | 86271505 | 51925149 |
| Q8BGW1 | Fto | Alpha-ketoglutarate-dependent dioxygenase FTO Mus musculus | 118.81 | 6.57 | 58007 | 2 | 2 | 1 | 1 | 1 | 225302.11 | 1 | 1 | 1 | 1 | 1 | 1 | 1 | 1 | 177542.48 | 1 | 1 |
| Q9DBE9 | Ftsj3 | pre-rRNA processing protein FTSJ3 Mus musculus | 724.5 | 26.73 | 95532 | 13 | 13 | 4936288 | 2077780 | 1827505.12 | 3731672.25 | 2251585.5 | 5179969.5 | 525862 | 1789109.38 | 6400560 | 1934751.62 | 2988682 | 4139859.25 | 4371777.5 | 2612680.5 | 5883916.5 |
| Q32PX7 | Fubp1 | Far upstream element-binding protein 1 Rattus norvegicus | 758.75 | 29.26 | 67197 | 14 | 16 | 19452977 | 18649261 | 23757205 | 27023873 | 18323221 | 20067799 | 28658355 | 20464381 | 20218657 | 27100969 | 21849653 | 22538085 | 23320043 | 20848319 | 22997721 |
| Q9D6K8 | Fundc2 | FUN14 domain-containing protein 2 Mus musculus | 190.78 | 21.85 | 16564 | 3 | 3 | 5720753 | 6413602 | 5358784 | 1250962.12 | 5218878 | 4724162 | 4030340 | 5455043.5 | 4968696.5 | 4016456 | 5324705.5 | 4625639 | 4514197 | 3362009.25 | 4576155 |
| Q8R2K1 | Fuom | Fucose mutarotase Mus musculus | 126.66 | 17.65 | 16805 | 2 | 2 | 2262723.25 | 1758226.5 | 1265212.5 | 2424451 | 1251164.12 | 1606712.25 | 851718.69 | 2254243 | 1892225.38 | 1179163 | 1057864.5 | 1246393.62 | 1317505.12 | 2623719.5 | 1001794.38 |
| P56959 | Fus | RNA-binding protein FUS Mus musculus | 626.84 | 23.55 | 52673 | 10 | 10 | 35919305 | 40310081 | 39968925 | 42177397 | 43356985 | 38540873 | 40392353 | 34170813 | 35698389 | 43992073 | 41817881 | 42019377 | 43318897 | 35677777 | 49283045 |
| D3ZYW7 | Fxn | Frataxin, mitochondrial Rattus norvegicus | 48.28 | 5.77 | 23066 | 1 | 1 | 1470573.38 | 2018505.62 | 1671755.12 | 1573553.88 | 1436761.75 | 1670100.38 | 1613356.62 | 1867548 | 1853375.25 | 2004898.12 | 1579372.88 | 1365065 | 1990195.88 | 1625313.5 | 1294303.25 |
| Q61584 | Fxr1 | Fragile X mental retardation syndrome-related protein 1 Mus musculus | 897.56 | 29.69 | 76222 | 14 | 15 | 22631109 | 24512787 | 24441133 | 22195725 | 23071475 | 22917873 | 29483919 | 26284441 | 25207907 | 25685509 | 26416303 | 27366233 | 22737593 | 24062611 | 23474909 |
| Q9WVR4 | Fxr2 | Fragile X mental retardation syndrome-related protein 2 Mus musculus | 228.17 | 7.88 | 73743 | 3 | 4 | 8126656.5 | 8517745 | 8133283 | 8277533 | 8584711 | 9212033 | 9449282 | 9479262 | 9120482 | 9621187 | 9348197 | 8808661 | 8166096.5 | 7797563.5 | 7339911.5 |
| P97808 | Fxyd5 | FXYD domain-containing ion transport regulator 5 Mus musculus | 74.01 | 20.79 | 19454 | 2 | 2 | 700896.44 | 1 | 1 | 1 | 1 | 907385.12 | 617907.25 | 1491177.88 | 1603172.88 | 1154910.783 | 415702.38 | 576209.25 | 487871.72 | 749220.69 | 557251.01 |
| O35601 | Fyb1 | FYN-binding protein 1 Mus musculus | 454.81 | 14.53 | 90055 | 9 | 9 | 6926826 | 6698229 | 6649610.5 | 11833141 | 7970181 | 8003091 | 11744939 | 8316097 | 7888062 | 8358087 | 6361505.5 | 7770686 | 7938316.5 | 8849062 | 9231896 |
| P97855 | G3bp1 | Ras GTPase-activating protein-binding protein 1 Mus musculus | 846.24 | 35.91 | 51829 | 13 | 15 | 15862112 | 18706691 | 18623369 | 16756893 | 19503099 | 18675159 | 19491787 | 19694309 | 20723101 | 21274439 | 22027169 | 19947529 | 20443743 | 17310813 | 22847643 |
| P97379 | G3bp2 | Ras GTPase-activating protein-binding protein 2 Mus musculus | 256.45 | 11 | 54088 | 3 | 5 | 4602877.5 | 6003054.5 | 5904047.5 | 5445513 | 6130925 | 6844776.5 | 4102425 | 5902282.5 | 7081814.5 | 5990039 | 4051858 | 4278537 | 3644684 | 3627956 | 5791589.5 |
| Q00612 | G6pdx | Glucose-6-phosphate 1-dehydrogenase X Mus musculus | 1454.99 | 55.73 | 59263 | 27 | 27 | 97138265 | 93448481 | 95200425 | 96327689 | 96271081 | 100704065 | 112964753 | 92459241 | 91550201 | 103857017 | 97132793 | 96747545 | 95439857 | 102528985 | 108798145 |
| P70699 | Gaa | Lysosomal alpha-glucosidase Mus musculus | 690.87 | 23.08 | 106248 | 14 | 14 | 22272737 | 15396345 | 14726017 | 16850859 | 17535857 | 21450991 | 16185986 | 26091477 | 20216563 | 18204197 | 16909467 | 15191492 | 18420553 | 23435181 | 16154390 |
| P60517 | Gabarap | Gamma-aminobutyric acid receptor-associated protein Rattus norvegicus | 148.57 | 20.51 | 13918 | 1 | 2 | 3734132.75 | 3583941 | 5045262 | 4634650.5 | 4044040.75 | 4921501.5 | 5303447 | 3625458.5 | 3550099.75 | 5988633 | 4856262.5 | 4884021 | 4525588 | 3588337.5 | 4956313 |
| P60521 | Gabarapl2 | Gamma-aminobutyric acid receptor-associated protein-like 2 Mus musculus | 127.07 | 27.35 | 13667 | 2 | 3 | 3766432.5 | 3512370 | 5199525 | 4685455.5 | 4001487.25 | 4855082.5 | 5273703 | 3778479 | 3639987 | 5992077 | 4897004 | 4666593 | 4332217.5 | 3510511.5 | 4961268.5 |
| Q00422 | Gabpa | GA-binding protein alpha chain Mus musculus | 221.16 | 14.32 | 51344 | 5 | 5 | 4205862 | 4537628.5 | 3042829.75 | 3104090.75 | 2691167.5 | 4670722.5 | 4212723 | 2982249 | 4454709 | 4132986.75 | 2694177.75 | 4254841.5 | 4043727.5 | 3366174.75 | 3343081 |
| Q99KY4 | Gak | Cyclin-G-associated kinase Mus musculus | 198.88 | 5.21 | 143641 | 3 | 3 | 3257501.5 | 3764614 | 4303446 | 2971749 | 2496956 | 2708532 | 2988175.25 | 2273786.75 | 2212100 | 3467559.75 | 2517772.5 | 2623324.5 | 2736226.25 | 3214867.75 | 2618565 |
| P54818 | Galc | Galactocerebrosidase Mus musculus | 84.13 | 5.12 | 77256 | 2 | 2 | 1318176.12 | 1087074.12 | 921299.69 | 1000666.94 | 1183955 | 927410.56 | 926585.81 | 869747.06 | 1235706.25 | 841447.5 | 1086247.12 | 962972.5 | 980100.56 | 1061034 | 883762.94 |

| | | | | | | | | | | | | | | | | | | | | | | |
|---|---|---|---|---|---|---|---|---|---|---|---|---|---|---|---|---|---|---|---|---|---|---|
| P18645 | Gale | UDP-glucose 4-epimerase Rattus norvegicus | 126.2 | 8.65 | 38225 | 2 | 2 | 3551099.5 | 3361979.5 | 3645733.75 | 2802935.5 | 3505035 | 2760169.5 | 3260696 | 3379881.25 | 3853772.5 | 2937617.25 | 1072974 | 1057378.75 | 3252877.5 | 4029308 | 1129410 |
| Q9R0N0 | Galk1 | Galactokinase Mus musculus | 453.75 | 37.5 | 42295 | 9 | 10 | 8279408 | 12743603 | 18287175 | 10402457 | 10066241 | 20618689 | 16862377 | 18866037 | 20161931 | 22500399 | 19804265 | 19574217 | 20998037 | 19180293 | 18496455 |
| Q68FH4 | Galk2 | N-acetylgalactosamine kinase Mus musculus | 220.02 | 12.01 | 50503 | 4 | 4 | 4180592.5 | 5116891 | 4586589 | 5857616 | 5836480.5 | 4321879.5 | 4042732 | 5445908.5 | 6263936.5 | 3996993.75 | 5308883.5 | 3202763.75 | 3786082 | 5159760.5 | 4683753 |
| Q571E4 | Galns | N-acetylgalactosamine-6-sulfatase Mus musculus | 394.08 | 15.96 | 57673 | 6 | 7 | 10118437 | 11670897 | 9301575 | 9488543 | 10695998 | 11631975 | 10061961 | 10598719 | 14706933 | 11390054 | 9149415 | 6884509 | 9569795 | 9887627 | 8146229 |
| Q6PB93 | Galnt2 | Polypeptide N-acetylgalactosaminyltransferase 2 Mus musculus | 160.96 | 8.6 | 64514 | 4 | 4 | 1872842.25 | 1691759.5 | 2217570.5 | 1075653 | 390742.75 | 1893106 | 1759264.5 | 1999265.75 | 2779638.25 | 1215669.38 | 1971153 | 2002717.5 | 1314922.75 | 2013406.75 | 1874899.75 |
| Q80VA0 | Galnt7 | N-acetylgalactosaminyltransferase 7 Mus musculus | 365.49 | 18.11 | 75419 | 8 | 8 | 15136037 | 13011606 | 9584421 | 10479327 | 12797417 | 8539775 | 10284578 | 10612573 | 12581861 | 10078771 | 13210443 | 10486660 | 11719754 | 13229709 | 11176722 |
| Q8BHN3 | Ganab | Neutral alpha-glucosidase AB Mus musculus | 1370.82 | 41.74 | 106911 | 29 | 29 | 34234589 | 29828711 | 32994331 | 32246355 | 28628169 | 32806983 | 44377493 | 37427353 | 36029913 | 34980285 | 34698565 | 36054397 | 29361417 | 31399247 | 38813193 |
| Q8BVW0 | Ganc | Neutral alpha-glucosidase C Mus musculus | 41.45 | 1.45 | 102008 | 1 | 1 | 494299.16 | 529243.2125 | 528985.5 | 519233.81 | 574454.38 | 473868.41 | 328260.09 | 540940.88 | 461901.97 | 504538.5 | 469166.12 | 312593.34 | 384612.78 | 439569.41 | 346164.09 |
| P16858 | Gapdh | Glyceraldehyde-3-phosphate dehydrogenase Mus musculus | 2270.76 | 87.69 | 35810 | 29 | 36 | 1044268161 | 1000019969 | 1184233217 | 1225996161 | 1173118081 | 1245029633 | 2175302657 | 1257351297 | 918552705 | 1680648449 | 1555629825 | 1456819201 | 1350194817 | 1194443137 | 1688264577 |
| Q6PAR5 | Gapvd1 | GTPase-activating protein and VPS9 domain-containing protein 1 Mus musculus | 286.74 | 5.56 | 162402 | 7 | 7 | 1588914.12 | 3550225 | 3559893.75 | 4762644.5 | 3431529 | 3901695.25 | 2516369.5 | 4648640 | 5244888 | 5217527.5 | 4136505 | 2685292.75 | 3056649.5 | 4600724 | 3879038 |
| Q6AYA1 | Gar1 | H/ACA ribonucleoprotein complex subunit 1 Rattus norvegicus | 60.84 | 7.96 | 23011 | 2 | 2 | 3150930.5 | 2862889.313 | 1571096.25 | 3286665 | 3442865.5 | 1 | 1974114.75 | 1 | 1 | 1964938 | 2954199.75 | 1237376.62 | 1381899.5 | 1805204.53 | 1647342.25 |
| Q9CZD3 | Gars | Glycine--tRNA ligase Mus musculus | 1395.78 | 37.59 | 81878 | 3 | 22 | 96797529 | 94607849 | 95651737 | 99651545 | 93605913 | 107853665 | 118991417 | 105272225 | 107549889 | 113072337 | 106628337 | 102042625 | 91966089 | 99328257 | 117757241 |
| Q64737 | Gart | Trifunctional purine biosynthetic protein adenosine-3 Mus musculus | 1922.33 | 47.03 | 107503 | 30 | 31 | 21127445 | 21301797 | 22663411 | 23803385 | 23066641 | 24720621 | 25629605 | 24188119 | 22984761 | 22852515 | 25633621 | 21006767 | 26567891 | 21485295 | 29178757 |
| O55148 | Gas7 | Growth arrest-specific protein 7 Rattus norvegicus | 296.82 | 18.48 | 48275 | 6 | 6 | 8409904 | 8834103 | 6357651.5 | 10475651 | 11646223 | 6778479.5 | 6669258.5 | 6897031 | 6849718.5 | 6814416.5 | 8566804 | 9112002 | 8190583 | 8242139.5 | 8526238 |
| Q99JT1 | Gatb | Glutamyl-tRNA(Gln) amidotransferase subunit B, mitochondrial Mus musculus | 112.23 | 8.98 | 62118 | 2 | 2 | 1 | 240130.56 | 210578.31 | 187402.14 | 1 | 1 | 1 | 234005.34 | 255652.23 | 234993.34 | 1 | 392024.38 | 1 | 208856.53 | 289961.69 |
| Q8BFQ8 | Gatd1 | Glutamine amidotransferase-like class 1 domain-containing protein 1 Mus musculus | 232.11 | 30 | 23277 | 4 | 4 | 3404697.5 | 3548624.75 | 3877741.25 | 3405962.75 | 3242598.75 | 2896195 | 3343239.25 | 4276396.5 | 4953308.5 | 3022099 | 3734045.5 | 4494037.5 | 4260417 | 3614464.75 | 4216592 |
| P50442 | Gatm | Glycine amidinotransferase, mitochondrial Rattus norvegicus | 650.58 | 30.97 | 48242 | 10 | 10 | 44178961 | 47179373 | 42780881 | 40331505 | 48363341 | 51043637 | 48570473 | 50060309 | 51739009 | 47313949 | 49021369 | 42606877 | 49612941 | 46641313 | 41424745 |
| P17439 | Gba | Glucosylceramidase Mus musculus | 288.44 | 20.39 | 57622 | 6 | 6 | 7411030.5 | 6799296.5 | 6604925 | 8099719.5 | 6469604 | 5614887 | 4592642.5 | 6353095 | 6260019 | 5126661 | 5678315 | 6319695 | 6561909.5 | 7467879.5 | 4540328 |
| Q9D6Y9 | Gbe1 | 1,4-alpha-glucan-branching enzyme Mus musculus | 38.75 | 1.85 | 80364 | 1 | 1 | 1 | 1 | 1 | 1 | 1 | 237746.94 | 1304682.75 | 1 | 1 | 1 | 746103 | 790873.69 | 1263161.75 | 905875.265 | 823362.62 |
| Q9Z0E6 | Gbp2 | Guanylate-binding protein 2 Mus musculus | 328.57 | 16.3 | 66739 | 6 | 6 | 4043399.75 | 4796837 | 4765077 | 1587843.38 | 3894907.25 | 2766148 | 1475815.5 | 2725365 | 2767971.25 | 3609875.25 | 3101153.5 | 3270625.5 | 5350110.5 | 3725803 | 3394939.25 |
| Q61107 | Gbp4 | Guanylate-binding protein 4 Mus musculus | 569.72 | 26.13 | 70801 | 11 | 11 | 8514483 | 7450088 | 5133596 | 6443512 | 6430460 | 6631421 | 4583984 | 7573968.5 | 6604644 | 4558294 | 4360159.5 | 5280020 | 6765151.5 | 8085754 | 3890224.5 |
| O88986 | Gcat | 2-amino-3-ketobutyrate coenzyme A ligase, mitochondrial Mus musculus | 461.88 | 32.45 | 44931 | 7 | 7 | 2071480 | 2891570.25 | 3612637.25 | 2995040 | 3543770.25 | 3903265.25 | 4600048.5 | 3026401.5 | 2725379.75 | 4193372 | 3232584.75 | 4734198 | 4339858.5 | 2717244 | 4107851.25 |
| Q60759 | Gcdh | Glutaryl-CoA dehydrogenase, mitochondrial Mus musculus | 98.2 | 5.94 | 48606 | 2 | 2 | 1620535.62 | 1467741.88 | 1832715 | 1938497.88 | 1648733.38 | 1710278.62 | 1672455 | 1684310.75 | 1981961.38 | 1662749.25 | 1410592.25 | 1458489.62 | 1553409.62 | 1924093.62 | 1648273 |
| P97494 | Gclc | Glutamate--cysteine ligase catalytic subunit Mus musculus | 391.57 | 16.01 | 72571 | 6 | 6 | 1696302.5 | 3405893.5 | 3341330.25 | 3184514.5 | 3020486.75 | 3209728.5 | 3440805.5 | 3596549.5 | 4001922.75 | 3194735.5 | 2001469.88 | 3263487 | 3150558 | 3764336.75 | 3349231 |
| O09172 | Gclm | Glutamate--cysteine ligase regulatory subunit Mus musculus | 278.8 | 20.8 | 30535 | 4 | 4 | 4865610.5 | 4399010.5 | 6606841 | 2449319.75 | 4602144 | 6931210 | 5264398.5 | 6058782 | 6247513 | 3463600 | 4740659 | 4696512 | 4608425 | 5174415.5 | 4375673 |
| E9PVA8 | Gcn1 | eIF-2-alpha kinase activator GCN1 Mus musculus | 1253.66 | 11.08 | 293021 | 22 | 22 | 36018305 | 32823859 | 29905395 | 29609555 | 28048527 | 28314321 | 27525149 | 33992697 | 31922247 | 31891951 | 29441559 | 29630261 | 30489065 | 28208529 | 23325223 |
| Q91WK5 | Gcsh | Glycine cleavage system H protein, mitochondrial Mus musculus | 76.94 | 11.76 | 18637 | 1 | 1 | 567171.12 | 689098.62 | 795584.44 | 518279.56 | 615326.06 | 549736.25 | 507363.09 | 1066791.5 | 1042368.62 | 737249.19 | 602749.31 | 801113.38 | 703122.88 | 1072501.12 | 503969.47 |
| Q66H63 | Gdap2 | Ganglioside-induced differentiation-associated-protein 2 Rattus norvegicus | 38.01 | 1.61 | 56130 | 1 | 1 | 831176.88 | 720556.62 | 932246.88 | 1131572.75 | 758783.62 | 828674.31 | 763993.94 | 1119029.88 | 1 | 884930.5 | 346248.09 | 654188.38 | 1075016.88 | 975127.94 | 133509.84 |
| P50396 | Gdi1 | Rab GDP dissociation inhibitor alpha Mus musculus | 868.4 | 51.68 | 50522 | 9 | 14 | 52040957 | 48112345 | 39622377 | 44421417 | 52836105 | 49634369 | 48333477 | 49984137 | 46549521 | 58990509 | 48698881 | 47736161 | 53900105 | 53903453 | 50401873 |
| P50399 | Gdi2 | Rab GDP dissociation inhibitor beta Rattus norvegicus | 2045.24 | 74.38 | 50537 | 22 | 29 | 167608433 | 167957473 | 152872913 | 156400529 | 174835505 | 170658961 | 145581153 | 180769025 | 171308737 | 142153057 | 174200241 | 175019025 | 175844961 | 193122001 | 171391649 |
| Q9CQQ4 | Gemin2 | Gem-associated protein 2 Mus musculus 2 | 46.47 | 6.32 | 30441 | 1 | 1 | 1 | 1 | 1 | 131221.45 | 120477.13 | 1 | 110548.72 | 1 | 1 | 114871.26 | 111263.9975 | 72709.15 | 163863.7 | 109020.41 | 99462.73 |
| Q8BX17 | Gemin5 | Gem-associated protein 5 Mus musculus | 193.13 | 4.53 | 166592 | 5 | 5 | 3853023 | 3625824.5 | 3026013.5 | 2197521.5 | 2166582.25 | 1794819.62 | 1488040.12 | 2048674.12 | 2254236.25 | 1237340.5 | 2442654.25 | 2744502 | 2132943.75 | 2721516.75 | 2527636.75 |
| Q9D1H7 | Get4 | Golgi to ER traffic protein 4 homolog Mus musculus | 133.38 | 12.23 | 36525 | 3 | 3 | 2727767 | 2513847.5 | 2044945.5 | 2748242.75 | 2697812.5 | 1962030.25 | 2441285 | 2295243 | 1941175.62 | 1515364 | 1950727.5 | 869177.31 | 1803638.38 | 1665114.12 | 798530.88 |
| Q8K0D5 | Gfm1 | Elongation factor G, mitochondrial Mus musculus | 547.2 | 18.24 | 83550 | 3 | 10 | 9691852 | 9473548 | 10245069 | 6776132.5 | 8243651 | 11049715 | 10321173 | 12335506 | 11627431 | 6221459.5 | 8275499.5 | 10272118 | 6633757.5 | 11113708 | 9157451 |
| Q5BJP6 | Gfm2 | Ribosome-releasing factor 2, mitochondrial Rattus norvegicus | 47.55 | 1.67 | 85914 | 1 | 1 | 1 | 1 | 807486.94 | 879959.81 | 820097.69 | 731483.25 | 1 | 1040560.06 | 1 | 1 | 802185.5 | 670719 | 1 | 1 | 665571.81 |
| P47856 | Gfpt1 | Glutamine--fructose-6-phosphate aminotransferase [isomerizing] 1 Mus musculus | 178.28 | 8.32 | 78539 | 4 | 4 | 2224536 | 2953629 | 2589416 | 2316178.25 | 1497132 | 4924349.5 | 2269611.5 | 1754391.12 | 5016555 | 2781050.75 | 1337674.75 | 2626145.75 | 2314025.5 | 2007729 | 4344852 |
| Q8R0H9 | Gga1 | ADP-ribosylation factor-binding protein GGA1 Mus musculus | 122.02 | 6.61 | 69972 | 2 | 2 | 1854207 | 395725.62 | 1687098.5 | 1813656.25 | 1705823.5 | 1896028.12 | 2043798.25 | 1800032.5 | 1993087.62 | 1994358 | 2359629 | 1110457.88 | 1996401 | 698788.19 | 1927901.88 |
| Q6P5E6 | Gga2 | ADP-ribosylation factor-binding protein GGA2 Mus musculus | 51.11 | 4.48 | 66049 | 1 | 1 | 392778.47 | 599482.06 | 234197.67 | 447510.44 | 418492.16 | 339010.34 | 338027.125 | 330671.72 | 293665.38 | 388761.06 | 345763.56 | 282729.09 | 388835.5 | 271210.44 | 321227.31 |
| Q91VC9 | Ghitm | Growth hormone-inducible transmembrane protein Mus musculus | 181.47 | 16.76 | 37275 | 3 | 3 | 806760.25 | 3824930.5 | 4245632 | 4440761.5 | 4651806.5 | 2740670.75 | 3033131.5 | 1755733.5 | 1356863 | 2163241.75 | 4249665 | 2022930.25 | 1911453.75 | 734840.19 | 3485608 |
| Q9D7M1 | Gid8 | Glucose-induced degradation protein 8 homolog Mus musculus | 290.44 | 39.04 | 26779 | 5 | 5 | 2313184.25 | 3789896.5 | 3269249.5 | 3287851.75 | 3370374.5 | 2820376.5 | 3278349.5 | 3776084.5 | 4103018.75 | 3141153.5 | 954631.81 | 3889975 | 1884187.88 | 2437781.5 | 3444703 |
| Q6Y7W8 | Gigyf2 | GRB10-interacting GYF protein 2 Mus musculus | 228.63 | 5.65 | 149193 | 4 | 4 | 1181891.62 | 1094963.62 | 1509011.12 | 950411.12 | 983237.94 | 1681937.75 | 1565021.62 | 1586243.38 | 1407920.25 | 1456707.12 | 1357287 | 1923634.62 | 1535261.25 | 1057392.62 | 1491057.25 |
| Q68FF6 | Git1 | ARF GTPase-activating protein GIT1 Mus musculus | 82.2 | 1.82 | 85300 | 1 | 1 | 333864.03 | 464910.84 | 461032.47 | 310569.19 | 351643.22 | 452244.66 | 343256.88 | 444519.88 | 345536.03 | 301131.97 | 483546.34 | 432345.44 | 318503.28 | 471889.34 | 558169.62 |
| Q9JLQ2 | Git2 | ARF GTPase-activating protein GIT2 Mus musculus | 138.18 | 8.47 | 78766 | 3 | 3 | 300345.81 | 1919752.5 | 2231588.5 | 1810593.75 | 763491.31 | 1596283.38 | 693042.44 | 1699196.62 | 1909083.88 | 1700497.62 | 1322771.38 | 1444466.38 | 1656987.25 | 1440727.62 | 1272020 |
| Q64516 | Gk | Glycerol kinase Mus musculus | 418.09 | 20.57 | 61227 | 9 | 9 | 10587397 | 8673071 | 11610506 | 10183176 | 8658168 | 10350327 | 11201164 | 11971258 | 10224195 | 10740105 | 12078468 | 10335938 | 9777244 | 9246751 | 11075413 |
| P51569 | Gla | Alpha-galactosidase A Mus musculus | 791.73 | 39.38 | 47643 | 13 | 13 | 43265073 | 44987593 | 41021769 | 42075301 | 41287025 | 42653097 | 31188279 | 47063273 | 57836257 | 36876929 | 44437157 | 38004461 | 44125853 | 51136909 | 32988711 |
| P23780 | Glb1 | Beta-galactosidase Mus musculus | 782.19 | 24.57 | 73121 | 15 | 15 | 74988033 | 85227161 | 72027969 | 63749433 | 80690641 | 76143793 | 62905153 | 92426633 | 117766625 | 63614121 | 67545505 | 64589477 | 80704329 | 84875273 | 61483973 |
| Q61543 | Glg1 | Golgi apparatus protein 1 Mus musculus | 1636.4 | 32.26 | 133734 | 29 | 29 | 55488997 | 55751933 | 47964597 | 49339425 | 53261433 | 49642653 | 44011717 | 48636885 | 54343069 | 47131301 | 49781385 | 42701093 | 44232921 | 51945065 | 41103889 |
| Q9CWG1 | Glipr1 | Glioma pathogenesis-related protein 1 Mus musculus | 61.17 | 5.49 | 29129 | 1 | 1 | 1234218.25 | 1205148.25 | 1037347 | 1332321.75 | 1437550.5 | 1373508.38 | 1607569.25 | 1585905 | 1448201.62 | 1643199.25 | 1112794.62 | 1265565.38 | 1540564.75 | 1173762.62 | 1273699.5 |
| Q8BZM1 | Glmn | Glomulin Mus musculus | 192.62 | 6.21 | 67756 | 3 | 3 | 1630325.62 | 1510563.5 | 1626378.5 | 1526176 | 1512363.75 | 1475318.62 | 1100170 | 1547344.88 | 2231597.75 | 1558771 | 2072671.12 | 1359994.88 | 1423631.62 | 1540201.12 | 1106956.5 |
| Q9JHJ3 | Glmp | Glycosylated lysosomal membrane protein Mus musculus | 115.5 | 8.17 | 43804 | 2 | 2 | 390392.56 | 375639.5 | 895242.81 | 521804.69 | 836234.56 | 958240.62 | 2126805 | 926883.62 | 458883.94 | 1312384.88 | 845479.44 | 1227055.38 | 1280147 | 784451.62 | 2288071.75 |
| Q9CPU0 | Glo1 | Lactoylglutathione lyase Mus musculus | 441.84 | 46.74 | 20810 | 8 | 8 | 22031245 | 26920293 | 19755439 | 26164569 | 24276729 | 24253805 | 25177897 | 25576985 | 22727533 | 24131243 | 24639033 | 21893513 | 25781595 | 27852507 | 21437923 |
| Q9CPV4 | Glod4 | Glyoxalase domain-containing protein 4 Mus musculus | 817.04 | 60.4 | 33317 | 7 | 15 | 21218533 | 106618553 | 18382133 | 23281503 | 22404561 | 22825069 | 24242641 | 22493611 | 19602363 | 22265163 | 23638009 | 19477655 | 22789289 | 20351217 | 20093557 |
| Q9QUH0 | Glrx | Glutaredoxin-1 Mus musculus | 296.6 | 57.94 | 11871 | 4 | 4 | 14472304 | 17170877 | 22626497 | 12931024 | 14314939 | 18085289 | 20391531 | 22745715 | 20590049 | 12089333 | 18478683 | 16925849 | 16670385 | 28263493 | 17962181 |
| Q9CQM9 | Glrx3 | Glutaredoxin-3 Mus musculus | 666.96 | 43.62 | 37778 | 10 | 10 | 17375581 | 19209147 | 16288185 | 17734797 | 16590306 | 17243779 | 14340225 | 18202681 | 15719387 | 15947154 | 13080662 | 17061831 | 17889985 | 20685445 | 17007329 |
| Q80Y14 | Glrx5 | Glutaredoxin-related protein 5, mitochondrial Mus musculus | 164.67 | 20.39 | 16292 | 2 | 2 | 1583419.75 | 2376964 | 2317822.25 | 1911952.12 | 2597586.25 | 2227268.75 | 2361748.75 | 2270081.75 | 2160512.25 | 2199084.75 | 2303763.5 | 1853979.12 | 3016741.75 | 1793927.12 | 1767005.25 |
| D3Z7P3 | Gls | Glutaminase kidney isoform, mitochondrial Mus musculus | 705.94 | 29.08 | 73964 | 12 | 12 | 6084751 | 7923345.5 | 7132730 | 7813474 | 7803654.5 | 8542609 | 8769195 | 8983828 | 7749613 | 9890612 | 7788914 | 7623708 | 9743663 | 7672603 | 8728381 |
| B0BNM9 | GLTP | Glycolipid transfer protein Rattus norvegicus | 82.99 | 12.44 | 23704 | 2 | 2 | 947167.69 | 1288210.75 | 1221304.5 | 960443 | 1010822.81 | 1477777.62 | 1170716.5 | 1262669.88 | 1931614.88 | 1016682.88 | 871094.88 | 1206812.38 | 887346.94 | 1350765.62 | 1290825.75 |
| Q8K0R6 | Gltpd2 | Glycolipid transfer protein domain-containing protein 2 Mus musculus | 40.18 | 2.18 | 34453 | 1 | 1 | 2788235.5 | 248642.72 | 1 | 1 | 1 | 1 | 1 | 176211.44 | 1154215.12 | 1 | 1990554.75 | 1 | 1257170.25 | 1 | 2223108.5 |
| P26443 | Glud1 | Glutamate dehydrogenase 1, mitochondrial Mus musculus | 1417.03 | 53.76 | 61337 | 23 | 23 | 107670049 | 126157753 | 121263281 | 120967529 | 120317089 | 106721513 | 113684425 | 108947361 | 116462561 | 109764065 | 108862121 | 109005201 | 114537665 | 106262337 | 109737569 |
| Q5RKH0 | Glyr1 | Putative oxidoreductase GLYR1 Rattus norvegicus | 449 | 22.64 | 60421 | 7 | 7 | 5601259 | 7051058 | 7290744 | 7394831 | 9028924 | 6383805 | 7130860 | 6318178 | 8035091 | 7903793 | 6926281 | 6951681 | 7662134.5 | 7976978 | 6858112 |
| Q8K0C9 | Gmds | GDP-mannose 4,6 dehydratase Mus musculus | 354.99 | 23.12 | 41985 | 6 | 6 | 4954164.5 | 4834941 | 4326470.5 | 4401193.5 | 2836879.5 | 3669405.5 | 3431922.75 | 4684058.5 | 4421251.5 | 2871299 | 3394362.5 | 4213258.5 | 2587778.5 | 5528840 | 2690700 |
| Q63228 | Gmfb | Glia maturation factor beta Rattus norvegicus | 78.36 | 14.08 | 16736 | 1 | 2 | 4264387.5 | 3684344 | 5403790.5 | 4835011.5 | 5390039 | 4357091 | 10088639 | 3968566.25 | 2860754 | 7676126.5 | 6828725 | 6445672 | 5872738 | 2813663 | 8116768 |
| Q9ERL7 | Gmfg | Glia maturation factor gamma Mus musculus | 275.54 | 42.96 | 16748 | 5 | 6 | 7088348 | 9215614 | 7856842 | 10454353 | 12165899 | 10707248 | 17249705 | 6858734 | 5152323 | 16383408 | 11761759 | 9210099 | 12815019 | 6619094 | 11586667 |
| Q6PGG2 | Gmip | GEM-interacting protein Mus musculus | 250.95 | 8.75 | 107545 | 5 | 5 | 892273.06 | 1753771.12 | 1248583.75 | 1306761.62 | 944341.38 | 879177.38 | 1424241.38 | 1090178.25 | 1705523 | 1727357.5 | 1362505.75 | 2057642 | 1882085 | 1585554 | 1297705.25 |
| Q5XIC1 | Gmppa | Mannose-1-phosphate guanyltransferase alpha Rattus norvegicus | 92.41 | 6.43 | 46191 | 2 | 2 | 932103.75 | 38146285 | 693062.75 | 947676.38 | 50922325 | 1142363.12 | 1076920.62 | 846637.69 | 15172502 | 36367505 | 791339.62 | 9057393 | 27613139 | 928139.69 | 887704.88 |
| Q8BTZ7 | Gmppb | Mannose-1-phosphate guanyltransferase beta Mus musculus | 49.39 | 5 | 39917 | 1 | 1 | 4203369.5 | 4833290.5 | 4242727.5 | 4922349.5 | 5575047 | 3862472.25 | 3928683.5 | 3214383.25 | 2715428.75 | 3238080.25 | 1932733.88 | 2753327.47 | 3486299.25 | 2734631.75 | 2859645 |
| Q9DCZ1 | Gmpr | GMP reductase 1 Mus musculus | 192.79 | 8.99 | 37482 | 3 | 3 | 473300.62 | 958105.69 | 422140.38 | 943952 | 788612.88 | 154929.7 | 1057614.75 | 313945.53 | 524207.34 | 972661.88 | 1048538.94 | 444334.47 | 617711.62 | 551879.25 | 538216.25 |
| Q3THK7 | Gmps | GMP synthase [glutamine-hydrolyzing] Mus musculus | 1328.99 | 52.96 | 76723 | 23 | 23 | 29054957 | 30467919 | 32455597 | 32417665 | 32113261 | 33047503 | 30707157 | 33540379 | 34450685 | 33989437 | 30774319 | 33087417 | 33700305 | 33046579 | 33386651 |
| P21278 | Gna11 | Guanine nucleotide-binding protein subunit alpha-11 Mus musculus | 127.59 | 12.53 | 42024 | 1 | 3 | 1113086.62 | 816279.62 | 1206552.12 | 547198.38 | 527918.75 | 206681.34 | 282732.41 | 1245116.88 | 2346198.25 | 218111.86 | 1221309.5 | 1369644.5 | 213190.22 | 1488294.5 | 1010572.25 |
| P27601 | Gna13 | Guanine nucleotide-binding protein subunit alpha-13 Mus musculus | 378.98 | 22.81 | 44055 | 8 | 8 | 10219065 | 8161165 | 10680955 | 9740980 | 8717010 | 9889819 | 11651850 | 10416244 | 9958825 | 9087974 | 9976496 | 11548651 | 8961835 | 10443546 | 12707727 |
| P08752 | Gnai2 | Guanine nucleotide-binding protein G(i) subunit alpha-2 Mus musculus | 1594.85 | 67.89 | 40489 | 14 | 24 | 152412081 | 163942833 | 159485921 | 144590897 | 157207153 | 136298513 | 143087809 | 148418689 | 160018001 | 139544929 | 161640625 | 153185617 | 149727505 | 155522337 | 157470865 |
| P08753 | Gnai3 | Guanine nucleotide-binding protein G(k) subunit alpha Rattus norvegicus | 832.77 | 39.55 | 40522 | 4 | 12 | 30806729 | 33407441 | 33833005 | 25643085 | 31113389 | 30915059 | 26867787 | 31004269 | 36649917 | 24930329 | 34825313 | 33942897 | 30012353 | 34370793 | 36675097 |
| P21279 | Gnaq | Guanine nucleotide-binding protein G(q) subunit alpha Mus musculus | 358.17 | 24.51 | 42158 | 5 | 7 | 8880789 | 7985632 | 7864490.5 | 7197769.5 | 7928388 | 6318870 | 5087286 | 8081899.5 | 9233068 | 5616002.5 | 7527219.5 | 7639748 | 7547661.5 | 8350683.5 | 7938202 |
| P63094 | Gnas | Guanine nucleotide-binding protein G(s) subunit alpha isoforms short Mus musculus | 581.65 | 32.74 | 45664 | 10 | 11 | 9744239 | 12054281 | 10497814 | 10054858 | 11257588 | 12496204 | 10420075 | 12586772 | 15850714 | 12384327 | 12089701 | 10426759 | 11432931 | 12930429 | 12073455 |
| P54311 | Gnb1 | Guanine nucleotide-binding protein G(I)/G(S)/G(T) subunit beta-1 Rattus norvegicus | 705.63 | 50.59 | 37377 | 6 | 11 | 16494165 | 15670456 | 22188921 | 18765769 | 17749507 | 17683841 | 32903249 | 15350149 | 15196245 | 25660469 | 22624977 | 25201357 | 20230219 | 16156957 | 29334797 |
| P54313 | Gnb2 | Guanine nucleotide-binding protein G(I)/G(S)/G(T) subunit beta-2 Rattus norvegicus | 1056.32 | 63.53 | 37331 | 10 | 16 | 34841069 | 42208965 | 42039529 | 43463409 | 41996961 | 39691461 | 55842413 | 33753221 | 37081169 | 43652153 | 48606953 | 43302285 | 45054689 | 38390569 | 50009505 |
| P63213 | Gng2 | Guanine nucleotide-binding protein G(I)/G(S)/G(O) subunit gamma-2 Mus musculus | 83.93 | 39.44 | 7850 | 2 | 2 | 6892465 | 7611842 | 7428047 | 8924962 | 8298338.5 | 6617675 | 2847632.25 | 6250014.5 | 6781411.5 | 6216025 | 2560274 | 5558617 | 6667112 | 6446683 | 5136033 |
| P36916 | Gnl1 | Guanine nucleotide-binding protein-like 1 Mus musculus | 343.07 | 13.84 | 68771 | 5 | 5 | 2113483.5 | 3079817 | 3214730.5 | 2743833.25 | 4014801.5 | 3584622.25 | 3328813.75 | 3434436.75 | 4032114.25 | 3490115 | 2792895.25 | 2957653.75 | 3607424.5 | 3155429 | 3260612.25 |
| Q8CI11 | Gnl3 | Guanine nucleotide-binding protein-like 3 Mus musculus | 624.34 | 30.86 | 60786 | 12 | 12 | 6893120 | 6732897.5 | 10253097 | 8987618 | 5074815 | 6770998.5 | 11491380 | 10521610 | 10845042 | 6757273 | 13520740 | 6508590 | 9952386 | 10991560 | 7886043.5 |
| P98192 | Gnpat | Dihydroxyacetone phosphate acyltransferase Mus musculus | 70.04 | 2.36 | 76870 | 1 | 1 | 1 | 1 | 1 | 3075630.75 | 1 | 2185523.25 | 1 | 2462773 | 1 | 1 | 1088069.88 | 857306.06 | 1 | 1 | 2938933.5 |
| O88958 | Gnpda1 | Glucosamine-6-phosphate isomerase 1 Mus musculus | 160 | 16.26 | 32549 | 3 | 3 | 2575737.25 | 2873447.75 | 1285031.5 | 3602059.5 | 2907078 | 724108.62 | 2573007.25 | 2441519.5 | 3223147.75 | 1869679.5 | 2778069.75 | 3162842 | 2899888.5 | 2616059.75 | 3648050.5 |
| Q9JK38 | Gnpnat1 | Glucosamine 6-phosphate N-acetyltransferase Mus musculus | 111.98 | 15.76 | 20791 | 2 | 2 | 3336251.5 | 4078587.5 | 3908066.5 | 3978580.75 | 3855449.5 | 3341340 | 3143098.75 | 4049880 | 4586516.5 | 3101829.5 | 1927799.38 | 3427954.75 | 3517890 | 3961272.75 | 2852602.5 |
| Q8BFR4 | Gns | N-acetylglucosamine-6-sulfatase Mus musculus | 715.07 | 25.55 | 61175 | 13 | 13 | 53642101 | 56073937 | 50804413 | 59516449 | 58488501 | 48506885 | 41555033 | 59460501 | 51741617 | 43485393 | 44590681 | 51149161 | 54836793 | 55892289 | 38394673 |
| Q62839 | Golga2 | Golgin subfamily A member 2 Rattus norvegicus | 39.4 | 1 | 112872 | 1 | 1 | 347358.12 | 1 | 1 | 433134.72 | 551018.94 | 495304.16 | 1 | 413311.12 | 1 | 1 | 499611.88 | 1 | 451188.59 | 1 | 500692.06 |
| Q91VW5 | Golga4 | Golgin subfamily A member 4 Mus musculus | 59.57 | 0.58 | 257564 | 1 | 1 | 421708.03 | 476552.31 | 481143.72 | 511592.06 | 283701.88 | 435750.66 | 373394.34 | 533776.31 | 519779.53 | 465675.21 | 715942.5 | 408396.62 | 476649.72 | 410674.72 | 381674.75 |
| Q9CRA5 | Golph3 | Golgi phosphoprotein 3 Mus musculus | 161.82 | 9.73 | 33752 | 1 | 2 | 1233853.5 | 1361809.75 | 1235296.25 | 1315822.62 | 1022313.88 | 1217766.75 | 1085917.38 | 1164543.5 | 1487399.5 | 1299309 | 854062.06 | 1113622.88 | 1077656.25 | 1409345.5 | 1174953.25 |
| Q66H74 | Golph3l | Golgi phosphoprotein 3-like Rattus norvegicus | 155.56 | 8.77 | 32922 | 1 | 2 | 1976959.75 | 1936485 | 1743979.62 | 1889268.88 | 1744210.38 | 1673180.38 | 1338916.62 | 1678767.38 | 1943423.12 | 1702710.88 | 1468599 | 1514606.5 | 1601376.62 | 2102103 | 1383754.25 |
| Q9CR60 | Golt1b | Vesicle transport protein GOT1B Mus musculus | 130.62 | 10.87 | 15422 | 2 | 2 | 5513042 | 824664.62 | 806548.38 | 417443.72 | 390140.72 | 774840.5 | 767014 | 824755.44 | 1165281 | 374228.62 | 1026950.75 | 1014705 | 1276231.75 | 1043878.88 | 1300704.62 |
| Q99JX3 | Gorasp2 | Golgi reassembly-stacking protein 2 Mus musculus | 329.11 | 17.29 | 47038 | 5 | 5 | 4681539.5 | 4907770 | 5190959 | 6039763 | 5307992.5 | 4772086 | 5946581 | 4646035 | 5038584.5 | 5693793.5 | 5041090 | 3933318 | 5859188 | 5056585 | 5271223.5 |
| O88630 | Gosr1 | Golgi SNAP receptor complex member 1 Mus musculus | 47.82 | 5.2 | 28489 | 1 | 1 | 1441950.25 | 1730465.5 | 1363458 | 1220558.88 | 1553179.25 | 1464977.62 | 1281978.5 | 1442635.88 | 1680316 | 1131663.75 | 1502136.38 | 1257673.5 | 1331581.38 | 1506102.62 | 1601095.62 |
| P05201 | Got1 | Aspartate aminotransferase, cytoplasmic Mus musculus | 1339.22 | 63.44 | 46248 | 23 | 23 | 183289009 | 128224865 | 113330681 | 157477169 | 121211169 | 150253857 | 115374849 | 178882561 | 153455889 | 119146561 | 111692433 | 109156545 | 123135593 | 196295681 | 99476625 |
| P05202 | Got2 | Aspartate aminotransferase, mitochondrial Mus musculus | 1388.24 | 53.72 | 47411 | 23 | 23 | 312332705 | 298579585 | 324323841 | 323117889 | 290100417 | 289323009 | 364089185 | 330122113 | 305897729 | 326775009 | 270427905 | 286015873 | 290590497 | 317479617 | 280530113 |
| Q3ULJ0 | Gpd1l | Glycerol-3-phosphate dehydrogenase 1-like protein Mus musculus | 280.65 | 27.64 | 38226 | 7 | 7 | 2543968 | 2371350.5 | 3832269.25 | 988359.56 | 161670.52 | 1 | 2902751.75 | 1258495.5 | 3227046 | 1304971.5 | 1128687 | 1611747.75 | 3196916.5 | 2124222.25 | 2923386.25 |
| Q64521 | Gpd2 | Glycerol-3-phosphate dehydrogenase, mitochondrial Mus musculus | 536.62 | 22.28 | 80954 | 12 | 12 | 12265822 | 14680938 | 14065567 | 13584454 | 12169963 | 14921857 | 11084359 | 16472784 | 17411339 | 14225572 | 15291831 | 14163200 | 14163894 | 14038509 | 11772838 |
| P06745 | Gpi | Glucose-6-phosphate isomerase Mus musculus | 1639.38 | 58.6 | 62767 | 14 | 27 | 122799385 | 132587617 | 124140889 | 107792553 | 137788401 | 138885697 | 170301729 | 129185553 | 126804361 | 144078337 | 158679569 | 153559217 | 155513953 | 143906881 | 177933345 |
| Q99P91 | Gpnmb | Transmembrane glycoprotein NMB Mus musculus | 299.24 | 10.45 | 63676 | 6 | 6 | 15079978 | 18625543 | 14805663 | 15678684 | 18425977 | 12073892 | 9368046 | 14159889 | 13001834 | 9741060 | 20380285 | 19490113 | 17591201 | 17466241 | 19633593 |
| Q8CIM5 | Gpr84 | G-protein coupled receptor 84 Mus musculus | 320.45 | 27.02 | 43717 | 5 | 5 | 7807401 | 6131564 | 5569723 | 5427608.5 | 7374241 | 4714073 | 4009133.5 | 3872539 | 3643870.25 | 5941217 | 6648972 | 6603202 | 8071299 | 6868107 | 7713785.5 |
| P97834 | Gps1 | COP9 signalosome complex subunit 1 Rattus norvegicus | 423.14 | 24.42 | 53428 | 8 | 8 | 8377154.5 | 7295562.5 | 8448687 | 5767675.5 | 6388505.5 | 7760879 | 6109803.5 | 8882763 | 9606779 | 5939322 | 6951846.5 | 7693885.5 | 5729428.5 | 8564640 | 6480232 |
| P11352 | Gpx1 | Glutathione peroxidase 1 Mus musculus | 216.24 | 29.85 | 22329 | 4 | 4 | 12420509 | 9940401 | 8472454 | 11677418 | 11073843 | 10476687 | 8025136 | 10776045 | 9971315 | 8808347 | 9751047 | 11111080 | 10904863 | 12894579 | 8314123 |
| O70325 | Gpx4 | Phospholipid hydroperoxide glutathione peroxidase Mus musculus | 178.09 | 22.34 | 22229 | 4 | 4 | 12119297 | 14274557 | 13153134 | 13247991 | 12999605 | 13173246 | 11025923 | 10684127 | 14783681 | 12352489 | 11352050 | 11366790 | 11901251 | 12928428 | 10071254 |
| Q60631 | Grb2 | Growth factor receptor-bound protein 2 Mus musculus | 359.46 | 40.55 | 25238 | 7 | 7 | 19001541 | 20050225 | 18111553 | 11778371 | 17586897 | 19985509 | 20088131 | 19693401 | 19162577 | 19979609 | 17475993 | 18582545 | 19443053 | 20592803 | 16711871 |
| O35127 | Grcc10 | Protein C10 Mus musculus | 116.25 | 28.57 | 13194 | 2 | 2 | 493293.09 | 2113838.5 | 1189737.88 | 1739117.5 | 1862298.5 | 1587771.12 | 1557739.25 | 1806334.5 | 2049901.25 | 2056606.62 | 1072813.25 | 1447472.12 | 2211989.75 | 1832882.5 | 1282012.25 |
| Q91Z53 | Grhpr | Glyoxylate reductase/hydroxypyruvate reductase Mus musculus | 301.1 | 23.17 | 35329 | 6 | 6 | 4643523 | 5200505 | 4224973.5 | 4792153.5 | 4646698.5 | 5630431 | 6908860 | 5576433 | 5530829 | 6749450.5 | 5653425 | 4668840.5 | 5520226.5 | 5481964 | 4573659 |
| P26817 | Grk2 | Beta-adrenergic receptor kinase 1 Rattus norvegicus | 607.39 | 22.06 | 79785 | 13 | 13 | 6677560 | 5539719 | 6658146 | 4423462.5 | 6012117 | 6107881.5 | 7284262.5 | 6462795 | 7050721 | 6743517 | 7137021 | 6389029 | 6840420.5 | 6258227 | 7091894 |
| P28798 | Grn | Granulins Mus musculus | 564.23 | 22.07 | 63458 | 11 | 11 | 36123945 | 42283901 | 28707123 | 37969009 | 44100657 | 38868637 | 32891329 | 28197115 | 32443453 | 36313209 | 34847877 | 24416157 | 36574161 | 28908663 | 31348339 |

| | | | | | | | | | | | | | | | | | | | | | | |
|---|---|---|---|---|---|---|---|---|---|---|---|---|---|---|---|---|---|---|---|---|---|---|
| P97576 | Grpel1 | GrpE protein homolog 1, mitochondrial Rattus norvegicus | 223.31 | 25.35 | 24297 | 5 | 5 | 266657.03 | 958875.81 | 245341.73 | 1302648 | 955945.62 | 1517999.25 | 1688386.75 | 2520586.75 | 1998680.38 | 1031222.06 | 2168800.75 | 1620054 | 1082695.12 | 1430473.75 | 1375657.88 |
| Q8C5Q4 | Grsf1 | G-rich sequence factor 1 Mus musculus | 124.36 | 7.1 | 53076 | 2 | 2 | 1513430.5 | 265656.28 | 1716902 | 1505736 | 1397569.5 | 1432488 | 1673428 | 1339349.5 | 1480447.595 | 1476524.88 | 1267754.62 | 1119227.38 | 746208.31 | 1287794 | 1137763.75 |
| Q810D6 | Grwd1 | Glutamate-rich WD repeat-containing protein 1 Mus musculus | 220.73 | 12.56 | 49224 | 4 | 4 | 4084838.5 | 3332763 | 4120762 | 4366309 | 3994580.5 | 3127738.5 | 3298426 | 3635051.75 | 3549815 | 4126948.75 | 3315716.75 | 2887585.75 | 2307649.25 | 2685089.5 | 3251797 |
| Q9D8T2 | Gsdmdc1 | Gasdermin-D Mus musculus | 559.62 | 28.54 | 53238 | 10 | 10 | 30094597 | 28334321 | 28243353 | 24313145 | 26067663 | 30521217 | 24894347 | 28751411 | 29289421 | 22903531 | 32733341 | 28418891 | 27261977 | 29902879 | 27124061 |
| Q9Z2D3 | Gsdme | Gasdermin-E Mus musculus | 44.84 | 1.76 | 56631 | 1 | 1 | 643205.62 | 1 | 1 | 554476.19 | 1 | 1 | 1 | 714700.62 | 747052.25 | 523819.34 | 502362.53 | 505167.9875 | 653805.88 | 534976.38 | 329527.16 |
| Q2NL51 | Gsk3a | Glycogen synthase kinase-3 alpha Mus musculus | 114.63 | 4.69 | 51661 | 2 | 2 | 964385.38 | 694561.5 | 677590.38 | 743043.72 | 635637.62 | 651583.5 | 956683.19 | 1131174.75 | 1003989.56 | 826870.81 | 963242.38 | 802298.69 | 927427.75 | 600413.38 | 941740 |
| P18266 | Gsk3b | Glycogen synthase kinase-3 beta Rattus norvegicus | 49.49 | 6.9 | 46742 | 1 | 1 | 1 | 1 | 1 | 1 | 1 | 1 | 247538.27 | 1 | 820449.69 | 1 | 849122.12 | 1 | 1 | 1 | 1 |
| P13020 | Gsn | Gelsolin Mus musculus | 2023.14 | 54.87 | 85942 | 32 | 32 | 165182961 | 175577073 | 172119601 | 185502081 | 203728641 | 194231937 | 237573841 | 173074801 | 147152257 | 223159841 | 192610385 | 181898545 | 210186529 | 166437057 | 191626593 |
| Q8R050 | Gspt1 | Eukaryotic peptide chain release factor GTP-binding subunit ERF3A Mus musculus | 500.2 | 23.58 | 68625 | 9 | 9 | 6721178 | 6278470.5 | 6931352.5 | 7236142.5 | 4968875.5 | 7260557 | 8310847.5 | 6694693.5 | 5605776 | 6699829 | 6729612 | 8300189 | 5864187 | 5378073 | 7308069.5 |
| P47791 | Gsr | Glutathione reductase, mitochondrial Mus musculus | 691.09 | 30 | 53663 | 6 | 10 | 17212551 | 19611767 | 20513933 | 22004241 | 14095437 | 17398835 | 25047231 | 18289837 | 17702669 | 19923231 | 14123536 | 16297658 | 18971231 | 16006961 | 18235947 |
| P46413 | Gss | Glutathione synthetase Rattus norvegicus | 181.99 | 11.81 | 52345 | 4 | 4 | 4631200 | 4455283.5 | 5560903 | 5407622.5 | 5190456 | 5471811 | 5748432 | 5361140 | 5647647 | 5308959 | 4488885 | 5710236 | 4918991 | 4792910 | 5871404 |
| Q9DCM2 | Gstk1 | Glutathione S-transferase kappa 1 Mus musculus | 135.54 | 17.7 | 25704 | 3 | 3 | 1352608.5 | 796224.12 | 2325049 | 1762967.5 | 1370246.5 | 1922043.62 | 1670296 | 2021718.12 | 2984498 | 1772225.75 | 1812234.75 | 2170429.25 | 2360169.75 | 1058263 | 877343 |
| P10649 | Gstm1 | Glutathione S-transferase Mu 1 Mus musculus | 515.64 | 46.33 | 25970 | 8 | 9 | 12996580 | 10876059 | 17243077 | 11134448 | 11213759 | 15879021 | 21021607 | 19326637 | 17726963 | 18719055 | 17621615 | 16564218 | 18239183 | 10463180 | 16615743 |
| P48774 | Gstm5 | Glutathione S-transferase Mu 5 Mus musculus | 485.22 | 42.41 | 26635 | 7 | 8 | 14716581 | 17360573 | 24001869 | 14632357 | 17022199 | 23542993 | 23026677 | 23533709 | 26855253 | 22874713 | 22735393 | 21970951 | 23919865 | 17518349 | 20419113 |
| O09131 | Gsto1 | Glutathione S-transferase omega-1 Mus musculus | 225.92 | 20 | 27498 | 4 | 4 | 5040239 | 5463971 | 4953207.5 | 6277726 | 5098597 | 5274245 | 5203755.5 | 5297099 | 4818789.5 | 4311095.5 | 5521331 | 4547335 | 5328864 | 5044568 | 4788869 |
| P62915 | Gtf2b | Transcription initiation factor IIB Mus musculus | 257.34 | 18.99 | 34819 | 4 | 4 | 4488791 | 4869705.5 | 4240542 | 4620558 | 4386188.5 | 3949320.25 | 5433354 | 3923074.5 | 4302549.5 | 4055099 | 3863495.5 | 4806823 | 4643622 | 4051112.5 | 4398301 |
| Q9D0D5 | Gtf2e1 | General transcription factor IIE subunit 1 Mus musculus | 48.41 | 2.95 | 49593 | 1 | 1 | 1 | 445364.44 | 705151.12 | 506598.72 | 746931.44 | 513161.03 | 514593.69 | 829191.12 | 671877.88 | 871985.44 | 881547.94 | 747339.1975 | 453918.03 | 574935.94 | 1078954.88 |
| Q3THK3 | Gtf2f1 | General transcription factor IIF subunit 1 Mus musculus | 116.06 | 6.5 | 57241 | 2 | 2 | 1 | 1 | 590315.31 | 583785.19 | 1 | 1 | 1 | 2096143.75 | 1 | 1 | 489224.88 | 631193.62 | 2501478 | 1 | 1 |
| Q9DBA9 | Gtf2h1 | General transcription factor IIH subunit 1 Mus musculus | 48.22 | 2.38 | 61851 | 1 | 1 | 295063.1625 | 436338.5 | 267319.59 | 244863.11 | 231731.45 | 1 | 325825.31 | 414770.09 | 370496.5 | 375118.44 | 332338.19 | 317130.91 | 341350.34 | 338608.75 | 332357.0475 |
| Q9ESZ8 | Gtf2i | General transcription factor II-I Mus musculus | 383.49 | 15.43 | 112265 | 9 | 9 | 5702867 | 6604590.5 | 2522804 | 4596535.5 | 4069722.75 | 4563235 | 6423831 | 5349554.5 | 3206069.25 | 2377974 | 4733386 | 5073938.5 | 5252178.5 | 3185601.5 | 4194861 |
| Q8BMQ2 | Gtf3c4 | General transcription factor 3C polypeptide 4 Mus musculus | 48.54 | 2.33 | 91613 | 1 | 1 | 148779.95 | 95456.84 | 237425.92 | 144857.19 | 140392.62 | 167282.11 | 143460.28 | 260613.52 | 213879.7 | 126914.77 | 325874.72 | 262468.22 | 119153.37 | 194153.3 | 300000.94 |
| D2XV59 | Gtpbp1 | GTP-binding protein 1 Rattus norvegicus | 447.47 | 17.94 | 72489 | 8 | 8 | 5781637 | 3632949 | 5440118 | 4444974 | 5251820 | 4740340 | 3277514 | 4521078 | 3950525.5 | 4272981 | 4884853.5 | 4630603.5 | 4281313.5 | 4306110.5 | 3989681.5 |
| Q99ME9 | Gtpbp4 | Nucleolar GTP-binding protein 1 Mus musculus | 640.33 | 24.76 | 74113 | 13 | 13 | 4191265.5 | 1856079.5 | 4486196.5 | 2227758 | 2976197 | 2564551 | 2557827.25 | 3449579 | 4609742 | 1444376.38 | 4343091 | 4132430.5 | 813368.44 | 3342798 | 3365009.5 |
| P51840 | Gucy2e | Guanylyl cyclase GC-E Rattus norvegicus | 39.88 | 0.63 | 120801 | 1 | 1 | 770574 | 2047297.25 | 2362873.5 | 881445.12 | 876466.25 | 670419.75 | 2665794.25 | 1188630.62 | 1550959 | 1290964 | 1408668.62 | 1178519 | 1147723.25 | 968616.88 | 2364044.5 |
| Q64520 | Guk1 | Guanylate kinase Mus musculus | 98.74 | 14.14 | 21918 | 2 | 2 | 1425565.25 | 676793.38 | 1614261.88 | 1302717 | 1146173 | 393378.22 | 1165077.88 | 1352256 | 1215089.75 | 1031450.463 | 398133.81 | 1054761.25 | 958329.905 | 800724.81 | 1579699.75 |
| P12265 | Gusb | Beta-glucuronidase Mus musculus | 1304.21 | 44.44 | 74195 | 23 | 23 | 175019537 | 175104497 | 143682769 | 165715329 | 203578257 | 175363633 | 128900353 | 178704321 | 207572161 | 138225473 | 182598705 | 155683953 | 180804289 | 194134065 | 140565345 |
| Q3UHH8 | Gxylt1 | Glucoside xylosyltransferase 1 Mus musculus | 38.58 | 1.98 | 46491 | 1 | 1 | 385412.75 | 306356.34 | 428203.12 | 275930.5 | 1 | 1 | 251090.55 | 340300.09 | 480972.94 | 1 | 292698 | 1 | 172461.53 | 382266.72 | 1 |
| O08730 | Gyg1 | Glycogenin-1 Rattus norvegicus | 45.97 | 4.8 | 37378 | 1 | 1 | 3535684.75 | 3298503.5 | 3479268.125 | 3422287 | 3660597.25 | 1947706.38 | 1722916 | 2134305.25 | 1502266 | 1826798.408 | 2593475.25 | 1 | 1821876.38 | 2305709.25 | 1 |
| A2RRU1 | Gys1 | Glycogen [starch] synthase, muscle Rattus norvegicus | 64.7 | 1.9 | 84072 | 1 | 1 | 1 | 1617416.25 | 1889975.62 | 1317772.88 | 1 | 1 | 1016133.44 | 1584812.75 | 1350702.62 | 1530891.88 | 1 | 1 | 1442553.12 | 1324068 | 1434602 |
| P10922 | H1f0 | Histone H1.0 Mus musculus | 264.71 | 27.84 | 20861 | 6 | 6 | 6077676 | 5028575.5 | 5421710 | 8196977 | 6596668 | 4233503 | 9748551 | 4566301 | 3708041 | 7525276.5 | 5628881.5 | 6832011.5 | 6470930.5 | 4716845.5 | 6704440.5 |
| P01899 | H2-D1 | H-2 class I histocompatibility antigen, D-B alpha chain Mus musculus | 465.83 | 27.62 | 40836 | 0 | 9 | 94416209 | 100057817 | 97557457 | 82101145 | 98633273 | 106523505 | 97726049 | 108189201 | 105722705 | 89255881 | 90579161 | 76131473 | 88611705 | 95725089 | 75268665 |
| P01900 | H2-D1 | H-2 class I histocompatibility antigen, D-D alpha chain Mus musculus | 821.73 | 49.86 | 41110 | 8 | 16 | 141752129 | 156688449 | 144794481 | 127365297 | 153014273 | 158033153 | 139347969 | 162621041 | 163585745 | 131058081 | 136702513 | 112499961 | 134238097 | 147095569 | 116736857 |
| P14426 | H2-D1 | H-2 class I histocompatibility antigen, D-K alpha chain Mus musculus | 228.67 | 9.67 | 40620 | 0 | 4 | 76935561 | 88768785 | 78007345 | 74458721 | 84181505 | 86945089 | 77843313 | 85997273 | 84113057 | 76259217 | 71099593 | 57321649 | 74426801 | 73839521 | 57002289 |
| P01901 | H2-K1 | H-2 class I histocompatibility antigen, K-B alpha chain Mus musculus | 226.74 | 9.21 | 41302 | 0 | 5 | 69263969 | 77587809 | 70712513 | 67576433 | 76025153 | 78781385 | 73622025 | 78075777 | 74587649 | 68217969 | 65218909 | 52335877 | 68947729 | 66865181 | 53157721 |
| P01902 | H2-K1 | H-2 class I histocompatibility antigen, K-D alpha chain Mus musculus | 881.15 | 45.38 | 41490 | 10 | 16 | 96228993 | 98687489 | 100711857 | 77917329 | 95276449 | 98550681 | 94516249 | 101870425 | 105671121 | 82956337 | 86830865 | 77886273 | 90235753 | 93520233 | 77253201 |
| P01897 | H2-L | H-2 class I histocompatibility antigen, L-D alpha chain Mus musculus | 610.29 | 35.08 | 40711 | 1 | 12 | 97031153 | 107556897 | 103497809 | 84638721 | 106488073 | 114043049 | 105055457 | 115330993 | 109606201 | 96686297 | 96454665 | 78049673 | 95787193 | 98383169 | 80840073 |
| P01898 | H2-Q10 | H-2 class I histocompatibility antigen, Q10 alpha chain Mus musculus | 224.71 | 12 | 37251 | 0 | 5 | 31418471 | 41678173 | 36969873 | 29887583 | 37888525 | 41232549 | 45399737 | 40259705 | 32900509 | 39223229 | 31417237 | 24765473 | 37589321 | 26469861 | 29469625 |
| P06339 | H2-T23 | H-2 class I histocompatibility antigen, D-37 alpha chain Mus musculus | 236.87 | 10.64 | 40875 | 0 | 4 | 49192085 | 52357897 | 48947249 | 45976805 | 54361713 | 52769841 | 42708557 | 53685701 | 53456621 | 46247405 | 45304393 | 34500033 | 46781485 | 49921501 | 35725277 |
| P27661 | H2afx | Histone H2AX Mus musculus | 360.81 | 52.45 | 15143 | 0 | 5 | 229847233 | 215684609 | 251903953 | 234172321 | 248695969 | 240948321 | 416188929 | 198347537 | 153276977 | 352058145 | 264705425 | 302898689 | 281522305 | 171165985 | 316179137 |
| Q9QZQ8 | H2afy | Core histone macro-H2A.1 Mus musculus | 565.73 | 37.1 | 39735 | 10 | 10 | 13904306 | 16380042 | 16779923 | 13020004 | 19447685 | 13119343 | 20072915 | 12063269 | 15070696 | 19705031 | 18987819 | 13684424 | 19114057 | 15392023 | 19861025 |
| P0C0S6 | H2afz | Histone H2A.Z Mus musculus | 293.14 | 51.56 | 13553 | 3 | 5 | 235438561 | 223822273 | 266783489 | 242506881 | 251473025 | 254224577 | 436384961 | 208609665 | 160403489 | 379175553 | 274845537 | 317225601 | 289355841 | 180449297 | 341544065 |
| P02301 | H3f3c | Histone H3.3C Mus musculus | 368.36 | 52.21 | 15315 | 1 | 7 | 1011221633 | 887928705 | 971219457 | 1076199681 | 1066173505 | 968052097 | 1446038017 | 902914817 | 754275009 | 1323941889 | 1148690945 | 1067180161 | 1041177985 | 799666049 | 1246896897 |
| Q9D3B1 | Hacd2 | Very-long-chain (3R)-3-hydroxyacyl-CoA dehydratase 2 Mus musculus | 41.48 | 4.72 | 28402 | 1 | 1 | 1500543.62 | 370147.94 | 374076.28 | 305479.94 | 522270.91 | 1 | 1259901.75 | 294010.91 | 554551.56 | 289141.97 | 446637.09 | 332330.94 | 343535.03 | 246601.19 | 838013.06 |
| Q8K2C9 | Hacd3 | Very-long-chain (3R)-3-hydroxyacyl-CoA dehydratase 3 Mus musculus | 41.54 | 2.76 | 43131 | 1 | 1 | 1488737.12 | 437472.41 | 527302.81 | 264417.53 | 471759.06 | 482385.06 | 526097.31 | 1 | 1 | 1 | 510620.22 | 385671.94 | 560481 | 399707.22 | 464120.095 |
| Q61425 | Hadh | Hydroxyacyl-coenzyme A dehydrogenase, mitochondrial Mus musculus | 375.62 | 36.94 | 34464 | 7 | 7 | 12153773 | 15731794 | 13339585 | 13572125 | 14204149 | 11863714 | 11563783 | 15220319 | 15992069 | 11053046 | 14072256 | 13319234 | 15200637 | 13583478 | 14136048 |
| Q8BMS1 | Hadha | Trifunctional enzyme subunit alpha, mitochondrial Mus musculus | 1110.22 | 38.66 | 82670 | 19 | 19 | 28916769 | 24112179 | 27524993 | 26538753 | 25760241 | 30190747 | 31342299 | 28258399 | 26115371 | 28527293 | 30012017 | 30177529 | 28270761 | 28579523 | 30398627 |
| Q99JY0 | Hadhb | Trifunctional enzyme subunit beta, mitochondrial Mus musculus | 447.59 | 21.26 | 51386 | 9 | 9 | 18308545 | 19479009 | 20810149 | 20087425 | 20413189 | 19933737 | 26016787 | 36131593 | 18140233 | 21573145 | 19880055 | 22507683 | 21786859 | 18430429 | 22407761 |
| O35952 | Hagh | Hydroxyacylglutathione hydrolase, mitochondrial Rattus norvegicus | 57.78 | 6.15 | 34109 | 2 | 2 | 2686735.5 | 851342.94 | 3501160.25 | 1167546.12 | 1244698.62 | 2223414 | 1 | 1307123 | 4466503 | 1 | 1073410.62 | 2746226 | 2397209.53 | 3348371 | 2420830.5 |
| Q61035 | Hars | Histidine--tRNA ligase, cytoplasmic Mus musculus | 1010.52 | 48.33 | 57432 | 18 | 18 | 31521269 | 29842935 | 31278805 | 29822403 | 29324697 | 29989725 | 28242949 | 31432905 | 30279047 | 26553185 | 29834977 | 29368205 | 29426127 | 32500875 | 29734329 |
| Q8BY71 | Hat1 | Histone acetyltransferase type B catalytic subunit Mus musculus | 340.62 | 24.52 | 49278 | 7 | 7 | 5537363.5 | 4287962 | 3822267.25 | 4729448 | 4515848 | 4057220 | 1954393.75 | 3876311.75 | 3755265.5 | 3967787 | 4574253 | 3241018.25 | 3910476 | 4411754 | 3500725 |
| Q8BFT2 | Haus4 | HAUS augmin-like complex subunit 4 Mus musculus | 44.95 | 7.16 | 42197 | 1 | 1 | 304402.09 | 227842.42 | 302441.635 | 330933.56 | 346588.47 | 281396.22 | 292532.25 | 215648.11 | 277896.41 | 379687.56 | 263514.59 | 402875.53 | 271421.62 | 325295.75 | 477407.69 |
| Q9D786 | Haus5 | HAUS augmin-like complex subunit 5 Mus musculus | 107.36 | 6.79 | 69572 | 2 | 2 | 127841.55 | 262327.03 | 127895.41 | 165718.055 | 144808.23 | 416976.62 | 130306.16 | 233597.45 | 137262.42 | 229535.6625 | 392406.91 | 246834.97 | 134621.83 | 208944.19 | 203448.88 |
| Q69ZS7 | Hbs1l | HBS1-like protein Mus musculus | 76.01 | 3.67 | 75100 | 2 | 2 | 1 | 1 | 289044.81 | 283889.75 | 273839.38 | 349980.62 | 494486.41 | 339248.19 | 214008.78 | 446696.28 | 301638.1625 | 376395.28 | 240662.44 | 217509.52 | 371985.41 |
| P53702 | Hccs | Cytochrome c-type heme lyase Mus musculus | 71.53 | 7.35 | 30978 | 2 | 2 | 1115353 | 972654.62 | 1113247.25 | 1247881.25 | 1202632.75 | 1181589.88 | 1473398.38 | 1269184.62 | 1223265.25 | 1349905.88 | 1302357.88 | 1262175.25 | 1057132.12 | 1054712.12 | 1125473.38 |
| Q61191 | Hcfc1 | Host cell factor 1 Mus musculus | 644.41 | 11.34 | 210437 | 13 | 13 | 6203405.5 | 7394230.5 | 6269429.5 | 9585636 | 6893013.5 | 9165134 | 11147231 | 6064694 | 7675142 | 7925351 | 8292571.5 | 10413817 | 7774157 | 7928043.5 | 10836943 |
| P08103 | Hck | Tyrosine-protein kinase HCK Mus musculus | 202.92 | 13.17 | 59129 | 5 | 5 | 2724987.5 | 2864456 | 4521957 | 3100563.25 | 4244812 | 2857781.5 | 4255704 | 3621235.5 | 2881448 | 3775973 | 3148658.5 | 4152833.5 | 4886950 | 3987049.5 | 4422588 |
| P49710 | Hcls1 | Hematopoietic lineage cell-specific protein Mus musculus | 412.3 | 18.52 | 54240 | 8 | 8 | 11973537 | 10518004 | 12753000 | 12874864 | 16029502 | 14171697 | 22207715 | 10392699 | 13497092 | 22402993 | 17043595 | 15938959 | 15011619 | 14308212 | 18826651 |
| O09106 | Hdac1 | Histone deacetylase 1 Mus musculus | 633.27 | 35.48 | 55075 | 8 | 11 | 32608535 | 33157919 | 33158403 | 27421663 | 30621875 | 32839003 | 30629997 | 32066247 | 33704225 | 31695809 | 37174757 | 28729575 | 31760911 | 36673133 | 31083321 |
| P70288 | Hdac2 | Histone deacetylase 2 Mus musculus | 439.68 | 24.59 | 55302 | 5 | 8 | 20816513 | 19463119 | 18366981 | 16977519 | 21372809 | 19516629 | 17492069 | 18679619 | 19437815 | 18551453 | 20910693 | 16843299 | 19062107 | 20688051 | 17724539 |
| O88895 | Hdac3 | Histone deacetylase 3 Mus musculus | 72.08 | 4.01 | 48364 | 1 | 1 | 1348115.283 | 1613038.25 | 1429792.62 | 1183903.88 | 1165726.38 | 1449475 | 1297644.12 | 1 | 1 | 1287826.62 | 1151718.75 | 1468579.38 | 1271146.5 | 1480091.75 | 1342884.095 |
| P51859 | Hdgf | Hepatoma-derived growth factor Mus musculus | 448.32 | 48.95 | 26269 | 8 | 9 | 21098055 | 19599647 | 19918309 | 19440521 | 21249851 | 20570847 | 22549187 | 19100109 | 20613719 | 22956593 | 20856213 | 19938679 | 19590403 | 19793621 | 20038781 |
| Q3UMU9 | Hdgfl2 | Hepatoma-derived growth factor-related protein 2 Mus musculus | 88.95 | 3.14 | 74291 | 1 | 2 | 2615269.5 | 2156333 | 1839871.12 | 2171031.75 | 2839698.5 | 3147356.75 | 3288310.75 | 2141365 | 2267782.25 | 2499846.5 | 2954854.5 | 2147174 | 2162495.75 | 2386677 | 2338571.75 |
| Q3UGR5 | Hdhd2 | Haloacid dehalogenase-like hydrolase domain-containing protein 2 Mus musculus | 106.24 | 11.2 | 28730 | 2 | 2 | 1601858.88 | 1578865.75 | 1937501.25 | 1620819.12 | 1657198.5 | 1647318.12 | 1373273.5 | 1766775.25 | 1773282.75 | 689624.62 | 973225.62 | 1721742.5 | 1388190.88 | 2078054.75 | 1502458.12 |
| Q91WM2 | Hdhd5 | Haloacid dehalogenase-like hydrolase domain-containing 5 Mus musculus | 44.63 | 3.82 | 46306 | 1 | 1 | 360665.69 | 272300.16 | 435080.22 | 430365.12 | 235908.73 | 210557.94 | 317251.5625 | 374954.53 | 296181.44 | 387312.34 | 228527.11 | 258961.95 | 240702.23 | 477941.91 | 242304.7 |
| Q8VDJ3 | Hdlbp | Vigilin Mus musculus | 1592.19 | 29.02 | 141743 | 29 | 29 | 23232933 | 23506171 | 26037487 | 24295201 | 24547027 | 24931553 | 28839067 | 22101341 | 22768469 | 26323009 | 23505615 | 29655737 | 27935707 | 22415557 | 26550909 |
| Q8BQM4 | Heatr3 | HEAT repeat-containing protein 3 Mus musculus | 210.68 | 5.45 | 74305 | 3 | 3 | 2602077.75 | 1358947.62 | 2857003 | 2405509.25 | 1953410.5 | 2457395.25 | 3193491.5 | 2336385.5 | 2866733 | 3192725.5 | 2230861.5 | 3148713 | 2356304.25 | 2161334.5 | 3119875.5 |
| Q5PRF0 | Heatr5a | HEAT repeat-containing protein 5A Mus musculus | 55.06 | 0.59 | 219893 | 1 | 1 | 460746.09 | 1 | 373829 | 1 | 428216.5 | 403292 | 470544.12 | 417859.34 | 337188.84 | 413121.69 | 442978.56 | 529496.5 | 272320.62 | 471846.41 | 317337.28 |
| Q8C547 | Heatr5b | HEAT repeat-containing protein 5B Mus musculus | 63.18 | 0.82 | 224319 | 1 | 1 | 156976.59 | 135018.84 | 194960.61 | 217384.62 | 220957.33 | 216906.12 | 1 | 117873.15 | 117067.97 | 339472.38 | 341191.56 | 230596.47 | 148383.48 | 151197.41 | 217842.23 |
| A1EC95 | Heatr6 | HEAT repeat-containing protein 6 Rattus norvegicus | 82.91 | 2.16 | 136505 | 2 | 2 | 577690.69 | 396189.25 | 377223.16 | 446741.7975 | 435864.09 | 336233.69 | 409940.88 | 184043.7 | 387461.69 | 329419.99 | 409901.69 | 372334.91 | 1 | 410050.22 | 1 |
| Q69ZR2 | Hectd1 | E3 ubiquitin-protein ligase HECTD1 Mus musculus | 121 | 0.92 | 290086 | 2 | 2 | 278612.47 | 408957.69 | 258623.62 | 419540.75 | 365158.09 | 370289.53 | 333006.97 | 403438.81 | 321762.34 | 251312.8 | 297954.38 | 190343.58 | 311312.59 | 437433.47 | 345636.88 |
| Q6NVF4 | Helb | DNA helicase B Mus musculus | 40.58 | 0.74 | 121471 | 1 | 1 | 1 | 1 | 1 | 1309828.38 | 1 | 1 | 1 | 1 | 1017141.62 | 1 | 1 | 1 | 1014127.94 | 318803.34 | 1 |
| Q60848 | Hells | Lymphocyte-specific helicase Mus musculus | 625.19 | 15.71 | 95126 | 11 | 11 | 1720837.25 | 1543517.75 | 4692735.5 | 2017514.5 | 3456545 | 2275962.5 | 2491175.25 | 4761937.5 | 2462349.5 | 2057577.38 | 3931594 | 3166227.5 | 3120466.5 | 2843659.5 | 2158779 |
| E9QAM5 | Helz2 | Helicase with zinc finger domain 2 Mus musculus | 151.11 | 2.07 | 331563 | 4 | 4 | 379286.59 | 418748.47 | 384684.03 | 307553.22 | 292926.03 | 293149.34 | 201396.16 | 244313.59 | 324090.06 | 282640.19 | 320770.19 | 243937.83 | 375661.94 | 338183.62 | 333301.53 |
| P29416 | Hexa | Beta-hexosaminidase subunit alpha Mus musculus | 1086.12 | 42.23 | 60613 | 6 | 16 | 70220577 | 94596145 | 76982145 | 71568745 | 87501009 | 78313537 | 60454793 | 81979377 | 99432481 | 58229813 | 67001037 | 62307633 | 74563985 | 79399921 | 59802089 |
| P20060 | Hexb | Beta-hexosaminidase subunit beta Mus musculus | 738.72 | 39.37 | 61116 | 16 | 16 | 55997421 | 70052937 | 50176317 | 45176281 | 61938809 | 55889289 | 43850065 | 57115761 | 69450785 | 40957413 | 53826109 | 42018397 | 52498357 | 55664817 | 40687709 |
| Q99L13 | Hibadh | 3-hydroxyisobutyrate dehydrogenase, mitochondrial Mus musculus | 245.21 | 23.28 | 35440 | 5 | 5 | 2553083.25 | 2951650 | 1218244.12 | 2513776 | 3179769.5 | 3988897.5 | 3606878.5 | 3251169.75 | 2736060.5 | 3410391.5 | 3168675.25 | 3434962.25 | 2676077 | 3338556.25 | 3332386.75 |
| Q8QZS1 | Hibch | 3-hydroxyisobutyryl-CoA hydrolase, mitochondrial Mus musculus | 168.89 | 11.43 | 43038 | 3 | 3 | 2631392.25 | 2949682.5 | 3708862.5 | 2917451.5 | 3732863.25 | 3353308.5 | 3889585.25 | 3188299.5 | 2379954 | 3261591 | 3827087.25 | 3100509.75 | 1457143.12 | 2972237.5 | 1649377 |
| Q8VH49 | Higd1a | HIG1 domain family member 1A, mitochondrial Rattus norvegicus | 147.95 | 43.01 | 10301 | 2 | 2 | 813655.38 | 698692.56 | 782062.25 | 630587.57 | 227940.09 | 454788.72 | 1423300.12 | 315909.84 | 852475.88 | 1226544.25 | 1112542.75 | 941054.12 | 982619.75 | 923053 | 1025185.62 |
| P62959 | Hint1 | Histidine triad nucleotide-binding protein 1 Rattus norvegicus | 163.67 | 35.71 | 13777 | 3 | 3 | 41841437 | 48005725 | 47729601 | 49812909 | 48736601 | 39706669 | 35915009 | 35364945 | 39126501 | 41288473 | 44455905 | 38582081 | 40950265 | 40046777 | 41511153 |
| Q9D0S9 | Hint2 | Histidine triad nucleotide-binding protein 2, mitochondrial Mus musculus | 42.21 | 7.98 | 17320 | 1 | 1 | 1147041.25 | 1143952.62 | 1336661.75 | 1596866.62 | 1333003.38 | 1301646.25 | 1642832.62 | 1062027.88 | 789315.06 | 1552086.38 | 1539400.62 | 1192096 | 1191092.88 | 1141174.88 | 1272734.12 |
| Q8VD75 | Hip1 | Huntingtin-interacting protein 1 Mus musculus | 61.07 | 2.24 | 115202 | 2 | 2 | 2968139.75 | 1731633.38 | 1329137.38 | 3062914 | 3098671 | 1750194.5 | 1280531.62 | 1627824.12 | 1419510 | 1735044.62 | 2640273.5 | 1057913.5 | 1423392.88 | 1238416.88 | 1965899.75 |
| Q8BLH7 | Hirip3 | HIRA-interacting protein 3 Mus musculus | 142.01 | 3.66 | 65215 | 2 | 2 | 1 | 1 | 796076.69 | 711181.25 | 668247.88 | 614640.75 | 1 | 1 | 1 | 1 | 1 | 1 | 1 | 1 | 1 |
| P43275 | Hist1h1a | Histone H1.1 Mus musculus | 314.39 | 34.27 | 21785 | 4 | 6 | 73412177 | 52340465 | 53922981 | 63098833 | 49039541 | 45982625 | 88751593 | 45512233 | 35454069 | 74361785 | 47571789 | 64881785 | 51726421 | 42724569 | 50612733 |
| P43276 | Hist1h1b | Histone H1.5 Mus musculus | 292.83 | 22.42 | 22576 | 6 | 7 | 10660957 | 10001917 | 10339258 | 11442148 | 11154373 | 9314321 | 28856991 | 7924046.5 | 5515137 | 18491059 | 11495119 | 15400771 | 11996155 | 6722517.5 | 15368650 |
| P15864 | Hist1h1c | Histone H1.2 Mus musculus | 549.35 | 33.02 | 21267 | 3 | 10 | 142196337 | 107721441 | 120461889 | 129245409 | 101859425 | 92562705 | 236431921 | 96377281 | 70695801 | 175893825 | 107192745 | 159488769 | 115721065 | 85714409 | 127220857 |
| P43277 | Hist1h1d | Histone H1.3 Mus musculus | 504 | 24.43 | 22100 | 1 | 9 | 146072417 | 110155473 | 120200137 | 130880401 | 102216593 | 94418497 | 232164193 | 96125537 | 71094097 | 175083153 | 106144937 | 160894001 | 117784729 | 87160961 | 124962609 |
| P43274 | Hist1h1e | Histone H1.4 Mus musculus | 498.91 | 24.66 | 21977 | 2 | 8 | 113203889 | 80511689 | 90889553 | 98874025 | 73935537 | 69911681 | 179102049 | 72794777 | 55018817 | 131625001 | 80277257 | 121000721 | 88172297 | 66683589 | 95076457 |
| P15865 | Hist1h1e | Histone H1.4 Rattus norvegicus | 455.21 | 24.66 | 21987 | 0 | 8 | 130091425 | 95335217 | 108519433 | 117127905 | 89543865 | 82498361 | 211094145 | 86272097 | 63692361 | 158871969 | 95139081 | 144206113 | 103487953 | 77394097 | 113634145 |
| C0HKE1 | Hist1h2ab | Histone H2A type 1-B Mus musculus | 507.43 | 63.85 | 14135 | 1 | 7 | 231118113 | 216707329 | 253135185 | 234484961 | 250703665 | 241540289 | 418052897 | 198347537 | 153276977 | 356129665 | 266821457 | 304693825 | 282344737 | 171908705 | 327526049 |
| P10853 | Hist1h2bf | Histone H2B type 1-F/J/L Mus musculus | 1153.51 | 55.56 | 13936 | 2 | 18 | 2311764993 | 2088850177 | 1691605505 | 1780771073 | 1772783873 | 1610477569 | 1906436609 | 1738711041 | 1691146497 | 1909646337 | 1569978369 | 1570050305 | 1788572545 | 1684066561 | 1457320833 |
| P10854 | Hist1h2bm | Histone H2B type 1-M Mus musculus | 1109.6 | 55.56 | 13936 | 2 | 17 | 2307947009 | 2087647233 | 1690756609 | 1778428161 | 1768388353 | 1610089217 | 1905449985 | 1739111297 | 1691761153 | 1908359169 | 1571199233 | 1570955009 | 1787085057 | 1683226625 | 1458342017 |
| P68433 | Hist1h3a | Histone H3.1 Mus musculus | 547.58 | 52.21 | 15404 | 1 | 9 | 1012020993 | 888395521 | 972652545 | 1078272513 | 1066560641 | 968103041 | 1448623489 | 904576257 | 755075969 | 1325148801 | 1148890241 | 1066481409 | 1041598337 | 800282177 | 1247006593 |
| P84228 | Hist1h3b | Histone H3.2 Mus musculus | 513.57 | 52.94 | 15388 | 2 | 9 | 1010180225 | 887055361 | 970322305 | 1074411521 | 1064539265 | 966933505 | 1444682497 | 902012609 | 753903041 | 1321981697 | 1145983233 | 1065630209 | 1039458369 | 798487489 | 1243607809 |
| P62804 | Hist1h4b | Histone H4 Rattus norvegicus | 1131.95 | 58.25 | 11367 | 13 | 16 | 4833988609 | 2000589441 | 2137045889 | 1928947969 | 2121537153 | 1852347521 | 3072746241 | 1785708161 | 1916487937 | 2491089409 | 2064886913 | 2338077441 | 2004844161 | 1566010881 | 2545311745 |
| P0CC09 | Hist2h2aa3 | Histone H2A type 2-A Rattus norvegicus | 545.2 | 58.46 | 14095 | 0 | 7 | 232415809 | 218299201 | 257881473 | 244310337 | 255039889 | 247027233 | 428097793 | 202304321 | 154510881 | 363479649 | 272474945 | 307209025 | 291313857 | 172298785 | 335497665 |
| Q64522 | Hist2h2ab | Histone H2A type 2-B Mus musculus | 404.79 | 52.31 | 14013 | 0 | 5 | 65926153 | 43979121 | 106511801 | 92464297 | 88569009 | 71695153 | 201706385 | 56995697 | 40260193 | 177605121 | 120466177 | 143526273 | 94711625 | 39261457 | 180337281 |
| P17710 | Hk1 | Hexokinase-1 Mus musculus | 1409.73 | 30.6 | 108303 | 4 | 24 | 51614469 | 58252017 | 56549513 | 52917917 | 53608305 | 54690393 | 68166025 | 50993377 | 59741697 | 64231697 | 58671129 | 52278085 | 62176313 | 61808137 | 55813221 |
| O08528 | Hk2 | Hexokinase-2 Mus musculus | 1259.89 | 33.04 | 102535 | 18 | 23 | 36978961 | 40303117 | 41025301 | 33905165 | 36865553 | 38589873 | 41798797 | 42206665 | 50227873 | 40269713 | 47619449 | 47321605 | 45048685 | 53544405 | 39995781 |
| Q3TRM8 | Hk3 | Hexokinase-3 Mus musculus | 2099.18 | 46.85 | 100101 | 13 | 30 | 111634473 | 134472689 | 119574729 | 111517449 | 118247505 | 113514369 | 97629161 | 107615713 | 128792977 | 103279529 | 102817673 | 91471761 | 109456137 | 117566929 | 102365321 |
| Q91W97 | Hkdc1 | Putative hexokinase HKDC1 Mus musculus | 161.04 | 3.72 | 102259 | 1 | 4 | 11155605 | 13182875 | 13107003 | 10640257 | 11549691 | 10032299 | 12581892 | 8560592 | 15824990 | 14295275 | 15035771 | 8725641 | 11762737 | 16091545 | 11556207 |
| Q9D8V0 | Hm13 | Minor histocompatibility antigen H13 Mus musculus | 278.34 | 16.4 | 41748 | 6 | 6 | 17344729 | 18118161 | 16397292 | 14235864 | 17362193 | 19565511 | 15328159 | 19829249 | 24215603 | 16713429 | 16557207 | 16362697 | 17990509 | 19856825 | 14327526 |
| P22907 | Hmbs | Porphobilinogen deaminase Mus musculus | 319.45 | 25.21 | 39344 | 5 | 5 | 3290961.25 | 4298848 | 4512083 | 5402619.5 | 3872265.75 | 4390029 | 4899198 | 3930963.5 | 3802384.5 | 4472235.5 | 6137781 | 4019055 | 5243241 | 5615385 | 4745918.5 |
| P17095 | Hmga1 | High mobility group protein HMG-I/HMG-Y Mus musculus | 45.7 | 14.95 | 11614 | 1 | 1 | 911566.44 | 658399.25 | 938192.88 | 1175260.38 | 868148.69 | 911430.56 | 1726134.62 | 794135.62 | 437855.12 | 1517757.5 | 1135327 | 1265885.25 | 1063065.62 | 631716.44 | 999041.75 |
| P63158 | Hmgb1 | High mobility group protein B1 Mus musculus | 423.23 | 31.16 | 24894 | 6 | 7 | 27314087 | 22562343 | 23011071 | 23944357 | 31560505 | 31711323 | 24264609 | 23324497 | 19152905 | 25259847 | 25702389 | 26216779 | 21889881 | 28056389 | 16233772 |
| P30681 | Hmgb2 | High mobility group protein B2 Mus musculus | 509.12 | 29.05 | 24162 | 7 | 8 | 50777745 | 51925705 | 49958501 | 67947057 | 60783049 | 47880225 | 57428913 | 44423669 | 44436457 | 55767085 | 58868365 | 52152869 | 53545777 | 48453549 | 51332097 |

| | | | | | | | | | | | | | | | | | | | | | | |
|---|---|---|---|---|---|---|---|---|---|---|---|---|---|---|---|---|---|---|---|---|---|---|
| O54879 | Hmgb3 | High mobility group protein B3 Mus musculus | 107.64 | 13 | 23010 | 2 | 2 | 5262773.5 | 4866769.5 | 4656933 | 5142224 | 4846237 | 4609895 | 5785606 | 4329589 | 4245591.5 | 6036752 | 4540466.5 | 5082046 | 4738132.5 | 4276461 | 4743816.5 |
| P38060 | Hmgcl | Hydroxymethylglutaryl-CoA lyase, mitochondrial Mus musculus | 605.33 | 46.77 | 34239 | 10 | 10 | 20361335 | 25321399 | 25609585 | 17527379 | 23998657 | 24151177 | 26952119 | 26359047 | 29174529 | 23257205 | 26392333 | 24490017 | 23535615 | 28464439 | 23486301 |
| Q8JZK9 | Hmgcs1 | Hydroxymethylglutaryl-CoA synthase, cytoplasmic Mus musculus | 1358.43 | 53.27 | 57569 | 20 | 21 | 7472597 | 6713622 | 4069837.25 | 6253870.5 | 3350192 | 5307542.5 | 5670626 | 5640735.5 | 4240923.5 | 4400313 | 5840303.5 | 10136161 | 7317831.5 | 5878091 | 6567162.5 |
| Q9JL35 | Hmgn5 | High mobility group nucleosome-binding domain-containing protein 5 Mus musculus | 301.6 | 20.2 | 45344 | 6 | 6 | 7709765 | 8175025.5 | 9029629 | 7347658.5 | 10840799 | 9143321 | 10911669 | 7823442 | 9594441 | 11176072 | 12263837 | 10541215 | 10100637 | 7955312 | 11485959 |
| O70252 | Hmox2 | Heme oxygenase 2 Mus musculus | 417.53 | 38.1 | 35739 | 8 | 8 | 16528945 | 14790119 | 13183153 | 13672824 | 13986250 | 13914089 | 14693762 | 14683882 | 15387238 | 14228400 | 11878030 | 13513858 | 15199283 | 14847767 | 12221312 |
| Q9CX86 | Hnrnpa0 | Heterogeneous nuclear ribonucleoprotein A0 Mus musculus | 501.93 | 39.02 | 30530 | 8 | 8 | 18543561 | 23033749 | 22235577 | 24538709 | 22462929 | 20821383 | 21236255 | 21251393 | 23715599 | 16401795 | 18367701 | 25082201 | 22768091 | 21823891 | 22469321 |
| P04256 | Hnrnpa1 | Heterogeneous nuclear ribonucleoprotein A1 Rattus norvegicus | 1393.54 | 63.12 | 34212 | 15 | 19 | 25362331 | 29835313 | 38211317 | 30269097 | 30000265 | 33470355 | 49008969 | 30481521 | 30277051 | 40022609 | 36155109 | 43227885 | 34156825 | 30239915 | 48934757 |
| O88569 | Hnrnpa2b1 | Heterogeneous nuclear ribonucleoproteins A2/B1 Mus musculus | 1699.02 | 82.44 | 37403 | 25 | 25 | 139062753 | 131963097 | 176265057 | 137647809 | 127125321 | 129625969 | 163249809 | 157676961 | 171461825 | 143328529 | 158683633 | 195172529 | 119438833 | 144109569 | 186049601 |
| Q6URK4 | Hnrnpa3 | Heterogeneous nuclear ribonucleoprotein A3 Rattus norvegicus | 1355.72 | 48.55 | 39652 | 19 | 22 | 85827065 | 91923281 | 122147929 | 86692369 | 85959385 | 100825849 | 107075601 | 104524113 | 103717497 | 80775897 | 113221873 | 123668337 | 80524297 | 94478849 | 131802769 |
| Q99020 | Hnrnpab | Heterogeneous nuclear ribonucleoprotein A/B Mus musculus | 498.94 | 30.53 | 30831 | 8 | 9 | 23145561 | 25227909 | 26947213 | 24505421 | 26309121 | 24365885 | 25705569 | 23548265 | 23950799 | 25640345 | 27191403 | 24747429 | 24290659 | 22894053 | 26418743 |
| Q9Z204 | Hnrnpc | Heterogeneous nuclear ribonucleoproteins C1/C2 Mus musculus | 783.56 | 44.41 | 34385 | 1 | 13 | 90037665 | 95079089 | 95737849 | 93441361 | 108013793 | 99759897 | 102539561 | 98453489 | 100802489 | 100930041 | 98281265 | 103226081 | 101844505 | 92177529 | 97418665 |
| G3V9R8 | Hnrnpc | Heterogeneous nuclear ribonucleoprotein C Rattus norvegicus | 820.63 | 42.28 | 32857 | 1 | 13 | 91587977 | 96398145 | 97646769 | 95248209 | 110042449 | 101914593 | 103844113 | 100243545 | 102639345 | 102661545 | 100316409 | 104899041 | 103599873 | 93875337 | 98851721 |
| Q60668 | Hnrnpd | Heterogeneous nuclear ribonucleoprotein D0 Mus musculus | 495.15 | 25.63 | 38354 | 8 | 9 | 8594504 | 3998791.75 | 7767368.5 | 10836933 | 13714794 | 5972197 | 14910621 | 9610851 | 11714325 | 3720790.75 | 14433071 | 8708193 | 8413449 | 4727682.5 | 16694497 |
| Q3SWU3 | Hnrnpdl | Heterogeneous nuclear ribonucleoprotein D-like Rattus norvegicus | 135.9 | 16.46 | 35294 | 1 | 3 | 1 | 1 | 1 | 1 | 1 | 276821.53 | 1 | 1 | 1 | 596256.38 | 150960.83 | 1 | 412400.88 | 1 | 980294.38 |
| Q794E4 | Hnrnpf | Heterogeneous nuclear ribonucleoprotein F Rattus norvegicus | 935.53 | 54.46 | 45730 | 12 | 14 | 85709761 | 91869393 | 90898321 | 96375137 | 105867153 | 98073497 | 80818729 | 86310537 | 86725825 | 85619297 | 105057201 | 88012897 | 92238961 | 86229089 | 83986177 |
| O35737 | Hnrnph1 | Heterogeneous nuclear ribonucleoprotein H Mus musculus | 740.41 | 37.86 | 49199 | 6 | 12 | 74856865 | 82025825 | 78275569 | 89255617 | 91103553 | 78034145 | 68957505 | 73093281 | 74768993 | 73139289 | 93659633 | 73521057 | 82234961 | 74462849 | 74172009 |
| P70333 | Hnrnph2 | Heterogeneous nuclear ribonucleoprotein H2 Mus musculus | 596.54 | 32.74 | 49280 | 5 | 10 | 36482773 | 41749225 | 36871281 | 45399625 | 43615345 | 40553481 | 39274945 | 38920409 | 39345725 | 38804693 | 43249569 | 37316145 | 42844709 | 38140009 | 39533081 |
| P61979 | Hnrnpk | Heterogeneous nuclear ribonucleoprotein K Mus musculus | 1469.61 | 63.28 | 50976 | 23 | 23 | 144774369 | 158442625 | 157376049 | 163918337 | 162241809 | 159020609 | 188531905 | 152755585 | 145326065 | 179618657 | 162076433 | 171983041 | 167463217 | 151159793 | 181583473 |
| F1LQ48 | Hnrnpl | Heterogeneous nuclear ribonucleoprotein L Rattus norvegicus | 1366.59 | 67.09 | 67903 | 22 | 22 | 71784449 | 80882889 | 91346017 | 94987377 | 99295065 | 95737865 | 102456825 | 88620113 | 75528633 | 97771385 | 99839713 | 96328649 | 90911593 | 83901129 | 102355505 |
| Q921F4 | Hnrnpll | Heterogeneous nuclear ribonucleoprotein L-like Mus musculus | 554.52 | 26.23 | 64125 | 10 | 10 | 9680591 | 9633316 | 9131778 | 11410217 | 11651424 | 11658162 | 11457154 | 9811304 | 11973249 | 10573882 | 11572613 | 10755157 | 10489374 | 9993745 | 10451047 |
| Q9D0E1 | Hnrnpm | Heterogeneous nuclear ribonucleoprotein M Mus musculus | 1663.1 | 47.74 | 77649 | 27 | 28 | 85412833 | 88315073 | 105917417 | 107523809 | 106907113 | 99774585 | 150062913 | 82396225 | 76234897 | 145199073 | 115945865 | 125473313 | 111533817 | 82012609 | 140987681 |
| Q6IMY8 | Hnrnpu | Heterogeneous nuclear ribonucleoprotein U Rattus norvegicus | 2107.2 | 43.36 | 87732 | 31 | 32 | 164142273 | 185724625 | 181767889 | 187177761 | 182851089 | 164252401 | 196289889 | 166572289 | 190372049 | 177537633 | 194068001 | 187995361 | 184475761 | 176844817 | 187421905 |
| Q8VDM6 | Hnrnpul1 | Heterogeneous nuclear ribonucleoprotein U-like protein 1 Mus musculus | 398.51 | 17.46 | 96002 | 10 | 10 | 7594204 | 6369281 | 7597508 | 6940959 | 8571770 | 7095325.5 | 7540809 | 7084354 | 8861188 | 8340372.5 | 8735565 | 9193046 | 7414766 | 6905519 | 10081141 |
| Q00PI9 | Hnrnpul2 | Heterogeneous nuclear ribonucleoprotein U-like protein 2 Mus musculus | 568.93 | 17.58 | 84940 | 11 | 11 | 33132101 | 38198973 | 37533853 | 38153349 | 38778877 | 38370445 | 42770853 | 37274301 | 39841169 | 40285745 | 40348509 | 39058881 | 37562377 | 34970985 | 41011009 |
| Q8BUK6 | Hook3 | Protein Hook homolog 3 Mus musculus | 50.94 | 1.95 | 83218 | 1 | 1 | 250239.77 | 1 | 429937.62 | 1 | 1 | 1 | 1 | 274501.09 | 225404.53 | 1 | 671321.38 | 452964.84 | 497625.04 | 324865.88 | 541348.06 |
| Q3TEA8 | Hp1bp3 | Heterochromatin protein 1-binding protein 3 Mus musculus | 499.96 | 22.74 | 60867 | 9 | 9 | 15570894 | 17605471 | 16048955 | 19989929 | 18067201 | 16248513 | 18855963 | 16518037 | 15093464 | 18633601 | 18009843 | 16421454 | 17039755 | 16413559 | 17372133 |
| P62748 | Hpcal1 | Hippocalcin-like protein 1 Mus musculus | 128.53 | 16.06 | 22338 | 3 | 3 | 4420884 | 4650699 | 4425670 | 4190072 | 3879160 | 3963280 | 3402654.75 | 4417117 | 4776370 | 2205019.5 | 4023814.5 | 3702266.5 | 3687280 | 4264167.5 | 3734666.25 |
| Q8CFE2 | Hpf1 | Histone PARylation factor 1 Mus musculus | 195.1 | 15.32 | 39291 | 4 | 4 | 5105827 | 4907595.5 | 4066558 | 4904229 | 3923714.5 | 4778080.5 | 4991999 | 4392863 | 6537378 | 5025074.5 | 4846862.5 | 4686426 | 3763629 | 4926338.5 | 4824448 |
| P00493 | Hprt1 | Hypoxanthine-guanine phosphoribosyltransferase Mus musculus | 592.38 | 69.72 | 24570 | 12 | 12 | 37369653 | 39280897 | 42360637 | 37940021 | 36330717 | 37246881 | 41820009 | 38475141 | 40300653 | 39482077 | 40178777 | 43371929 | 40503681 | 42225049 | 39637569 |
| P59438 | Hps5 | Hermansky-Pudlak syndrome 5 protein homolog Mus musculus | 54.34 | 0.98 | 126337 | 1 | 1 | 1293994 | 977232.12 | 679642.94 | 1212638.62 | 1405209.38 | 752711.75 | 785273.75 | 664310 | 373976.38 | 977916.69 | 910365.5 | 354279.59 | 643137.06 | 433899.06 | 407823.03 |
| Q3TC93 | Hs1bp3 | HCLS1-binding protein 3 Mus musculus | 94.78 | 4.3 | 43691 | 1 | 1 | 1310085.62 | 1358001.75 | 1127502 | 1406968.38 | 1162737.12 | 1416550.75 | 1072403.75 | 1308767.88 | 1172401.88 | 1586099 | 1109239.62 | 1408298.5 | 1239864.62 | 1325506.38 | 1072009.5 |
| Q8R3H7 | Hs2st1 | Heparan sulfate 2-O-sulfotransferase 1 Mus musculus | 117.24 | 7.3 | 41813 | 2 | 2 | 2687547.75 | 1947724.62 | 1716276.5 | 1655924.12 | 1704163.75 | 1865927.62 | 2327658.75 | 1971137.62 | 1783764.12 | 1639515.5 | 1215139.5 | 1330773 | 1867140.88 | 1644459.88 | 1397714.12 |
| O08756 | Hsd17b10 | 3-hydroxyacyl-CoA dehydrogenase type-2 Mus musculus | 712.23 | 55.94 | 27419 | 5 | 9 | 23563111 | 25149027 | 29836307 | 30778667 | 26688145 | 26866877 | 33807909 | 29184769 | 27810249 | 33590381 | 20551707 | 31646097 | 31119893 | 26050409 | 31020495 |
| Q9EQ06 | Hsd17b11 | Estradiol 17-beta-dehydrogenase 11 Mus musculus | 68.04 | 11.74 | 32881 | 2 | 2 | 1640220.5 | 380058.09 | 923520 | 395099.84 | 510289 | 1172236.62 | 323346.09 | 370941.03 | 981021 | 410101.31 | 421643.47 | 580205.88 | 1167470.62 | 379013.19 | 1479086 |
| O70503 | Hsd17b12 | Very-long-chain 3-oxoacyl-CoA reductase Mus musculus | 1048.91 | 69.55 | 34742 | 17 | 17 | 29122711 | 30014685 | 31073283 | 28494255 | 27393423 | 30780749 | 28262161 | 32888229 | 33651777 | 30015011 | 26881951 | 28951479 | 27385445 | 30659465 | 28261717 |
| P51660 | Hsd17b4 | Peroxisomal multifunctional enzyme type 2 Mus musculus | 696.87 | 21.77 | 79482 | 11 | 11 | 15125930 | 10977580 | 14185545 | 13229414 | 12428735 | 11087934 | 15566326 | 12932371 | 13764974 | 13202561 | 11791403 | 11765314 | 11965546 | 13722366 | 12054153 |
| O88736 | Hsd17b7 | 3-keto-steroid reductase Mus musculus | 381.96 | 26.65 | 37317 | 7 | 7 | 4350245.5 | 3922892.75 | 4406959.5 | 3599696 | 3864512 | 4603924 | 4587904.5 | 4547565 | 5614308.5 | 4163731.5 | 3848698 | 4297725.5 | 3673317.5 | 3056918.25 | 4277964 |
| P50171 | Hsd17b8 | Estradiol 17-beta-dehydrogenase 8 Mus musculus | 194.79 | 18.15 | 26588 | 4 | 4 | 3826551 | 2408823.25 | 2803414 | 4490665.5 | 4305340 | 4241100.5 | 6590151.5 | 5075408.5 | 1638375.75 | 6003174.5 | 4486120 | 4360714 | 4935447 | 3725829 | 4600641.5 |
| Q8BTX9 | Hsdl1 | Inactive hydroxysteroid dehydrogenase-like protein 1 Mus musculus | 138.33 | 10.91 | 36868 | 2 | 2 | 1532226.12 | 1417680.75 | 1644205.25 | 1829637.88 | 1761696.75 | 1673343 | 1052542.88 | 1711213.88 | 1486577.12 | 1580726 | 1290970.25 | 1422520.62 | 1241677.12 | 1402199.5 | 1404548.25 |
| P07901 | Hsp90aa1 | Heat shock protein HSP 90-alpha Mus musculus | 2808.68 | 62.62 | 84788 | 28 | 46 | 416174593 | 466700545 | 429501857 | 418725409 | 478100257 | 500390401 | 465011041 | 493861025 | 477139297 | 470457889 | 514531585 | 451577217 | 483024193 | 480610753 | 476956513 |
| P11499 | Hsp90ab1 | Heat shock protein HSP 90-beta Mus musculus | 4068.121 | 76.38 | 83281 | 38 | 61 | 858858113 | 907080065 | 860898369 | 795965633 | 908185857 | 972307969 | 886827201 | 960210113 | 966483137 | 887882817 | 1011949569 | 938138369 | 905636929 | 960531841 | 927077569 |
| P08113 | Hsp90b1 | Endoplasmin Mus musculus | 3099.08 | 61.97 | 92476 | 44 | 46 | 438865441 | 435713089 | 428330785 | 391069985 | 457535489 | 524561441 | 436578017 | 501669633 | 531080865 | 431629921 | 476389889 | 476054881 | 458124705 | 469964161 | 461589377 |
| Q99M31 | Hspa14 | Heat shock 70 kDa protein 14 Mus musculus | 296.23 | 19.84 | 54650 | 7 | 7 | 3661207 | 4327122.5 | 3964697.75 | 2309327.25 | 5929704.5 | 3198346.25 | 5860888.5 | 3645877.5 | 4225383.5 | 6144375 | 3508753 | 5512146 | 4943221 | 5490742.5 | 6632613 |
| Q61316 | Hspa4 | Heat shock 70 kDa protein 4 Mus musculus | 2190.29 | 54.7 | 94133 | 5 | 33 | 144136369 | 161532769 | 164268225 | 142548609 | 158958481 | 152200961 | 160432737 | 160890753 | 172204625 | 153135265 | 162813457 | 162755393 | 161379745 | 162702193 | 149296833 |
| P48722 | Hspa4l | Heat shock 70 kDa protein 4L Mus musculus | 219.27 | 5.37 | 94382 | 1 | 3 | 5077805.5 | 7223141 | 6410148 | 5993005.5 | 6154040 | 6344784 | 7020845 | 6000031.5 | 7195492 | 6472335.5 | 7457999.5 | 6722868.5 | 7076841 | 6790422.5 | 6069588.5 |
| P06761 | Hspa5 | Endoplasmic reticulum chaperone BiP Rattus norvegicus | 2885.9 | 64.53 | 72347 | 34 | 40 | 943839105 | 1021198273 | 1015736385 | 966251521 | 977109313 | 977872705 | 958973057 | 1025025409 | 1018912385 | 950681665 | 943086849 | 943308161 | 975120769 | 976470401 | 842926977 |
| P63017 | Hspa8 | Heat shock cognate 71 kDa protein Mus musculus | 3579.499 | 77.86 | 70871 | 44 | 52 | 1801361665 | 1869525377 | 1957913473 | 1742636801 | 1764108289 | 1764362113 | 1740668801 | 1687456641 | 1893132033 | 1660502785 | 1771001345 | 1750775937 | 1835773441 | 1818041601 | 1576494977 |
| P38647 | Hspa9 | Stress-70 protein, mitochondrial Mus musculus | 2182.07 | 55.67 | 73461 | 31 | 32 | 183261889 | 181052913 | 198643089 | 186770193 | 185384353 | 203123617 | 195938625 | 207787377 | 206125681 | 202956081 | 179096545 | 182381313 | 181658849 | 189346785 | 180384081 |
| Q99P31 | Hspbp1 | Hsp70-binding protein 1 Mus musculus | 572.21 | 45.66 | 39167 | 10 | 10 | 14451627 | 14545725 | 16243478 | 15873172 | 14678513 | 14436924 | 13359110 | 13688211 | 14888887 | 15159259 | 15188917 | 14686078 | 13728458 | 15443687 | 16303059 |
| P63038 | Hspd1 | 60 kDa heat shock protein, mitochondrial Mus musculus | 3463.88 | 79.06 | 60955 | 43 | 47 | 544798337 | 529123073 | 613124481 | 569205889 | 537038273 | 576978177 | 605455681 | 612508929 | 565886145 | 599854913 | 613673473 | 589687809 | 553880385 | 551822081 | 536390689 |
| P26772 | Hspe1 | 10 kDa heat shock protein, mitochondrial Rattus norvegicus | 215.66 | 42.16 | 10902 | 4 | 4 | 18476301 | 19934239 | 19126895 | 21050439 | 16966677 | 16631221 | 22463085 | 16221974 | 14268253 | 20142719 | 12452855 | 15622031 | 18734659 | 13708221 | 10608190 |
| Q61699 | Hsph1 | Heat shock protein 105 kDa Mus musculus | 1053.25 | 31.12 | 96407 | 17 | 18 | 13979423 | 14701681 | 15538737 | 12002855 | 13341430 | 13245895 | 12824343 | 12819097 | 11668779 | 10081248 | 12718045 | 12109680 | 12317414 | 15305376 | 11462091 |
| Q9Z2G9 | Htatip2 | Oxidoreductase HTATIP2 Mus musculus | 209.49 | 17.77 | 26870 | 4 | 4 | 2660624.25 | 2843695.25 | 3325615.5 | 1404470 | 1155086.5 | 4310785.5 | 2001897.75 | 2399211.5 | 4184203.75 | 2184273.25 | 2993701.5 | 2433354.75 | 2797603 | 2389265.5 | 5469331.5 |
| Q9JIY5 | Htra2 | Serine protease HTRA2, mitochondrial Mus musculus | 220.09 | 17.69 | 49348 | 4 | 5 | 747603.06 | 685581.5 | 941821.56 | 487902.75 | 834827.5 | 666170.06 | 1195577.38 | 1 | 1 | 1108165.38 | 1111960.25 | 412649.44 | 935251.3575 | 1324903.62 | 891492.12 |
| P42859 | Htt | Huntingtin Mus musculus | 156.62 | 1.44 | 344690 | 4 | 4 | 6380583 | 4782427 | 4312055.5 | 4736919.5 | 5056132.5 | 5586866.5 | 2967366 | 5387019.5 | 5100559.5 | 4992050.5 | 4412990 | 3987716.25 | 4927762.5 | 5697757 | 3262723.25 |
| Q7TMY8 | Huwe1 | E3 ubiquitin-protein ligase HUWE1 Mus musculus | 998.72 | 7.86 | 482635 | 22 | 22 | 5177047 | 6848453.5 | 9441940 | 6243555.5 | 7194109 | 7150260.5 | 10531864 | 7019647 | 7065946.5 | 9711518 | 6881576.5 | 8967636 | 7442226 | 5685894 | 12944219 |
| Q8R1F5 | Hyi | Putative hydroxypyruvate isomerase Mus musculus | 282.88 | 24.55 | 30449 | 6 | 6 | 9398018 | 1449733.62 | 3642734.5 | 9416271 | 8546109 | 8625603 | 6326005 | 8670142 | 4074959.5 | 6888944.5 | 7234481.5 | 4808112 | 7806293 | 8710827 | 3770172 |
| Q9JKR6 | Hyou1 | Hypoxia up-regulated protein 1 Mus musculus | 2199.98 | 52.25 | 111181 | 37 | 37 | 68673689 | 74321409 | 74723737 | 70327585 | 73010289 | 88632897 | 82681649 | 94504201 | 89198097 | 76154657 | 68467225 | 82716937 | 69306761 | 66171125 | 72108873 |
| P03975 | Iap | IgE-binding protein Mus musculus V=1 | 590.01 | 23.16 | 62747 | 11 | 11 | 21785451 | 19728165 | 25046163 | 23513657 | 20108189 | 32126171 | 28558429 | 29440323 | 33373827 | 30037465 | 20853213 | 19034379 | 22356197 | 25871409 | 17249657 |
| Q8BU30 | Iars | Isoleucine--tRNA ligase, cytoplasmic Mus musculus | 622.99 | 13.55 | 144271 | 14 | 14 | 49474901 | 37118825 | 34908689 | 42228605 | 41116593 | 34525333 | 33683565 | 33059561 | 32200303 | 37844045 | 35971697 | 28782593 | 34869073 | 32809721 | 37057957 |
| Q8BIJ6 | Iars2 | Isoleucine--tRNA ligase, mitochondrial Mus musculus | 743.05 | 18.08 | 112804 | 13 | 13 | 19898681 | 25412257 | 23707031 | 20511649 | 23922073 | 26989709 | 25192323 | 24251637 | 25861863 | 26335175 | 23928361 | 24041737 | 24183341 | 21685737 | 25062351 |
| P13597 | Icam1 | Intercellular adhesion molecule 1 Mus musculus | 99.99 | 5.21 | 58844 | 2 | 2 | 1823847.25 | 1870921.75 | 1795167 | 1391329.25 | 1687625.75 | 1635972 | 2135166 | 1523345.88 | 1875223.12 | 1858107.75 | 1657985.75 | 1649515.88 | 1976383.62 | 2017826 | 1460441 |
| Q9JHR7 | Ide | Insulin-degrading enzyme Mus musculus | 413.9 | 9.72 | 117772 | 8 | 8 | 8222862 | 8821802 | 7243275 | 7681324 | 8548060 | 8964223 | 9437800 | 8204150 | 9594612 | 7138709 | 8303432.5 | 9423829 | 7724405 | 8054311 | 10041813 |
| O88844 | Idh1 | Isocitrate dehydrogenase [NADP] cytoplasmic Mus musculus | 1107.41 | 61.11 | 46674 | 19 | 20 | 63572689 | 55378865 | 71228601 | 62620189 | 52962105 | 57754361 | 69523769 | 60322349 | 58673097 | 60329849 | 62336681 | 67694745 | 62432041 | 64520305 | 69659129 |
| P54071 | Idh2 | Isocitrate dehydrogenase [NADP], mitochondrial Mus musculus | 1170.59 | 48.67 | 50906 | 19 | 21 | 72911833 | 74086593 | 74553665 | 72508329 | 70330505 | 70055553 | 87279513 | 74324025 | 73208593 | 71378273 | 75242825 | 75743465 | 69600001 | 69514977 | 74884697 |
| Q9D6R2 | Idh3a | Isocitrate dehydrogenase [NAD] subunit alpha, mitochondrial Mus musculus | 714.15 | 36.61 | 39639 | 12 | 12 | 108897041 | 108297993 | 103560273 | 101560265 | 102674825 | 113234601 | 121617921 | 112086977 | 106419665 | 104113385 | 109436849 | 101661617 | 108760113 | 111546185 | 97408057 |
| Q68FX0 | Idh3B | Isocitrate dehydrogenase [NAD] subunit beta, mitochondrial Rattus norvegicus | 357.97 | 25.71 | 42354 | 8 | 8 | 10234452 | 8873762 | 12745161 | 10257745 | 10238926 | 12115038 | 13074467 | 10247927 | 11901176 | 10541052 | 10802603 | 12018092 | 11136303 | 11289421 | 11044996 |
| P41565 | Idh3g | Isocitrate dehydrogenase [NAD] subunit gamma 1, mitochondrial Rattus norvegicus | 253.8 | 14.76 | 42851 | 6 | 6 | 16545307 | 13982288 | 16847583 | 16369147 | 13910443 | 19518973 | 17887521 | 16911499 | 16665583 | 16975001 | 21632737 | 17221929 | 17784893 | 16977761 | 18943285 |
| P58044 | Idi1 | Isopentenyl-diphosphate Delta-isomerase 1 Mus musculus | 580.47 | 37.89 | 26289 | 8 | 9 | 20077199 | 17077927 | 16589274 | 13619267 | 15197891 | 15606099 | 11648359 | 16225040 | 22795907 | 16820877 | 18968971 | 21449259 | 14207392 | 19356619 | 17131521 |
| Q9CR20 | Ier3ip1 | Immediate early response 3-interacting protein 1 Mus musculus =3 | 90.03 | 34.15 | 9017 | 2 | 2 | 1776517.62 | 1800477.88 | 1646131.38 | 1235866.38 | 1670221.38 | 1713906.62 | 1606007.88 | 1452064.5 | 1679178.5 | 1425714 | 1502382.75 | 1596586.75 | 1641275.38 | 1643529.62 | 1488419.62 |
| Q9R002 | Ifi202 | Interferon-activable protein 202 Mus musculus | 144.04 | 6.29 | 50466 | 2 | 2 | 2326020.25 | 2754549.75 | 2657673.25 | 1744767.5 | 1884176.88 | 1 | 1 | 2016734.5 | 2534400.25 | 1522665.25 | 1 | 1 | 1537492.38 | 1744321 | 1 |
| O35368 | Ifi203 | Interferon-activable protein 203 Mus musculus | 141.11 | 12.25 | 46300 | 2 | 3 | 3073621 | 4515976 | 3747676 | 3639146.25 | 3943940.5 | 2360474.5 | 3509265.5 | 3246721 | 3493934.5 | 1750590.88 | 3571588 | 3532776.75 | 3254557 | 3817383.75 | 3491074 |
| P0DOV2 | Ifi204 | Interferon-activable protein 204 Mus musculus | 675.02 | 27.46 | 69438 | 8 | 12 | 18012943 | 19144417 | 19639325 | 15222543 | 14087636 | 13049272 | 12466191 | 14117468 | 21483111 | 11129984 | 15463110 | 18022501 | 14927233 | 18685525 | 14927077 |
| Q9ESY9 | Ifi30 | Gamma-interferon-inducible lysosomal thiol reductase Mus musculus | 276.9 | 20.97 | 27784 | 4 | 4 | 9177378 | 12348510 | 9824344 | 9560805 | 11264979 | 9639017 | 8303887.5 | 5916286 | 9244666 | 8536427 | 13671677 | 9403076 | 9785586 | 10742749 | 9047784 |
| Q9D8C4 | Ifi35 | Interferon-induced 35 kDa protein homolog Mus musculus | 309.27 | 27.62 | 31876 | 6 | 6 | 1691151.5 | 2825326 | 1568373.88 | 2018846 | 2413499.75 | 1613606 | 2607730.75 | 1368141.38 | 2245816.75 | 1720195.25 | 2206540 | 2721759.75 | 2215842 | 2156578.5 | 2642599.5 |
| Q8BV66 | Ifi44 | Interferon-induced protein 44 Mus musculus | 78.42 | 5.92 | 47852 | 2 | 2 | 6552790.5 | 6363975 | 5733456 | 3897284.25 | 5024364 | 3923444.75 | 3860262.75 | 3941201 | 3923596.5 | 3422009.5 | 4342920 | 4489787.5 | 5628904 | 4878678.5 | 3820342.5 |
| Q9BDB7 | Ifi44l | Interferon-induced protein 44-like Mus musculus 2 | 231.32 | 12.98 | 49680 | 4 | 4 | 2844332.75 | 2146209.25 | 2052692.5 | 1848826.62 | 1618923.62 | 1314626.75 | 2480184.5 | 1574834.12 | 1739018.12 | 1437857.62 | 1829716 | 1786888.5 | 2272691 | 2051923.25 | 2314744 |
| Q8R5F7 | Ifih1 | Interferon-induced helicase C domain-containing protein 1 Mus musculus | 51.61 | 1.17 | 115971 | 1 | 1 | 303777.95 | 307292.41 | 403792.19 | 272384.78 | 231642.42 | 326796.78 | 284678.09 | 414699.91 | 394320.38 | 234971.73 | 574946.56 | 304170.81 | 235302.61 | 533458.12 | 389385.81 |
| Q99J93 | Ifitm2 | Interferon-induced transmembrane protein 2 Mus musculus | 146.22 | 16.67 | 15743 | 1 | 2 | 53026101 | 38082793 | 41511925 | 25423989 | 37172865 | 36482909 | 23732905 | 35572777 | 39805953 | 30830155 | 34303925 | 31042549 | 35815145 | 33455283 | 25317629 |
| Q9CQW9 | Ifitm3 | Interferon-induced transmembrane protein 3 Mus musculus | 539.03 | 48.91 | 14954 | 4 | 6 | 65032261 | 45017053 | 49339585 | 30601089 | 45552461 | 46177077 | 28895671 | 41664737 | 45953789 | 37614017 | 41467737 | 37230409 | 43990749 | 39364349 | 31506813 |
| P19182 | Ifrd1 | Interferon-related developmental regulator 1 Mus musculus | 141.02 | 5.35 | 49935 | 2 | 2 | 1 | 1 | 1 | 1 | 285720.09 | 1 | 1 | 1 | 1 | 1 | 342989.8075 | 258396.19 | 269131.66 | 481769.69 | 362661.69 |
| Q61249 | Igbp1 | Immunoglobulin-binding protein 1 Mus musculus | 307.62 | 18.24 | 38971 | 4 | 4 | 2160710 | 1080741.12 | 1455760 | 1554181.88 | 508005 | 281279.53 | 240805.83 | 1181360.88 | 1003010.75 | 1288775.75 | 1188648.25 | 1677909 | 2012978 | 1742024 | 711073.62 |
| Q9CPN8 | Igf2bp3 | Insulin-like growth factor 2 mRNA-binding protein 3 Mus musculus | 714.89 | 26.6 | 63575 | 11 | 11 | 11325107 | 11967692 | 12592103 | 11890570 | 14149274 | 11982937 | 14698317 | 12683299 | 10624519 | 12326338 | 11911167 | 13669237 | 13273649 | 11699370 | 13608521 |
| Q66HG8 | Ik | Protein Red Rattus norvegicus | 222.1 | 12.57 | 65588 | 5 | 5 | 5506375 | 4898198 | 2074125.12 | 4769928.5 | 4593217 | 5296089 | 5252405 | 4978019.5 | 5363349 | 4945519.5 | 5997052.5 | 4831717 | 5060582.5 | 5132638 | 4916553 |
| O88351 | Ikbkb | Inhibitor of nuclear factor kappa-B kinase subunit beta Mus musculus | 264.42 | 10.7 | 86690 | 5 | 5 | 9063181 | 2698539.25 | 8796161 | 2351809.5 | 2684371 | 8194339 | 2305185.5 | 8446505 | 10273911 | 6370558.5 | 10484828 | 9883095 | 1663427.38 | 8980895 | 8325549 |
| Q9R0T8 | Ikbke | Inhibitor of nuclear factor kappa-B kinase subunit epsilon Mus musculus | 202.04 | 6.83 | 80953 | 4 | 4 | 498817.88 | 1032041.25 | 1727046.75 | 2295590.75 | 1765663.25 | 1553224.5 | 1073502.75 | 1354425.25 | 2484573.75 | 1847502.12 | 1645857.62 | 1378011 | 1616792.5 | 1064119.5 | 713010.44 |
| Q9CXY6 | Ilf2 | Interleukin enhancer-binding factor 2 Mus musculus | 732.59 | 53.85 | 43062 | 14 | 14 | 20563567 | 19345431 | 20165027 | 20896637 | 20034229 | 21838131 | 15069050 | 20344179 | 22783701 | 20825677 | 20730559 | 18120875 | 18366059 | 18931381 | 18650325 |
| Q9Z1X4 | Ilf3 | Interleukin enhancer-binding factor 3 Mus musculus | 511.49 | 16.82 | 96021 | 10 | 11 | 6328511.5 | 6775481 | 7170851 | 6997951.5 | 7813943 | 7416571 | 6665427.5 | 7387566.5 | 6929435.5 | 7364557.5 | 6039111 | 6770238 | 7516968 | 6429702 | 6826411.5 |
| O55222 | Ilk | Integrin-linked protein kinase Mus musculus | 80.71 | 6.42 | 51373 | 2 | 2 | 2229950.25 | 2900513 | 2954583 | 2837169.25 | 3166686 | 2396546.25 | 3616420.5 | 2672500 | 3333894.5 | 3177164.5 | 2748003.5 | 2919836.75 | 3339225 | 2507001.25 | 3065287.75 |
| Q8R0F6 | Ilkap | Integrin-linked kinase-associated serine/threonine phosphatase 2C Mus musculus | 135.03 | 10.97 | 42774 | 3 | 3 | 1029290.25 | 1227866.88 | 1092892.12 | 1115054.88 | 891648.75 | 999663.75 | 1561678.62 | 1180186.5 | 1344525.12 | 1438719 | 1795422.25 | 1404302.75 | 1251686.88 | 1146389.12 | 1594840.5 |
| Q8CAQ8 | Immt | MICOS complex subunit Mic60 Mus musculus | 1666.42 | 45.18 | 83900 | 28 | 28 | 63464085 | 66969681 | 64434625 | 74027753 | 63271897 | 62337029 | 83471025 | 67653649 | 69099129 | 73465521 | 68807257 | 62074529 | 71004913 | 63277761 | 65428041 |
| O55023 | Impa1 | Inositol monophosphatase 1 Mus musculus | 611.74 | 46.93 | 30436 | 8 | 12 | 42729101 | 42525937 | 44979457 | 46504489 | 46628441 | 42121909 | 45067201 | 40793273 | 45422717 | 47971209 | 47030681 | 43702249 | 45122157 | 43491937 | 45894761 |
| Q8CIN7 | Impa2 | Inositol monophosphatase 2 Rattus norvegicus | 138.48 | 13.45 | 31796 | 3 | 3 | 3346350 | 2725572.75 | 1079522.75 | 2314741.75 | 3517454.5 | 2107666 | 2141215.75 | 3830807 | 3294819 | 2087530.5 | 2538839.5 | 3297847 | 3701151.75 | 2485698 | 1234047.75 |
| D3ZLZ7 | Impdh1 | Inosine-5'-monophosphate dehydrogenase 1 Rattus norvegicus 3 | 186.18 | 7.2 | 55293 | 3 | 4 | 10691699 | 7636193.5 | 8596957 | 10345150 | 9403159 | 10061851 | 10390848 | 10352499 | 7415078.5 | 11247505 | 10768299 | 8331152 | 9896100 | 9673842 | 9014182 |
| P24547 | Impdh2 | Inosine-5'-monophosphate dehydrogenase 2 Mus musculus | 1273.46 | 44.94 | 55815 | 18 | 19 | 76242937 | 59049025 | 62302537 | 75801265 | 75943457 | 82305945 | 71997513 | 77186657 | 69361825 | 77354129 | 72451945 | 71918625 | 71668425 | 82006233 | 61758429 |
| Q0GNC1 | Inf2 | Inverted formin-2 Mus musculus | 432.48 | 8.25 | 138560 | 8 | 8 | 1647709.12 | 1456264.75 | 1339991.88 | 1237981.5 | 1380480 | 1829130.75 | 1885823 | 2223330.5 | 2042381 | 1897470.62 | 1680831.38 | 2652184.5 | 2136495 | 2325219.75 | 2560672.25 |
| P49442 | Inpp1 | Inositol polyphosphate 1-phosphatase Mus musculus | 89.5 | 6.82 | 43346 | 2 | 2 | 3260557 | 2083278.62 | 2319977 | 1820683.38 | 3456455.5 | 3471099.25 | 1968263.38 | 3246999 | 2109267.25 | 2198920.25 | 2028144.25 | 2363486.75 | 4330527.5 | 3313900.5 | 2169513.75 |
| Q8K337 | Inpp5b | Type II inositol 1,4,5-trisphosphate 5-phosphatase Mus musculus | 49.91 | 2.22 | 112762 | 1 | 1 | 286034.19 | 270461.94 | 1 | 1 | 1 | 193351.47 | 164193.8675 | 172080.98 | 175723.83 | 115619.19 | 240072.64 | 180790.86 | 206491.86 | 156464.41 | 195954.9425 |
| Q9ES52 | Inpp5d | Phosphatidylinositol 3,4,5-trisphosphate 5-phosphatase 1 Mus musculus | 1433.81 | 30.81 | 133542 | 27 | 29 | 33906385 | 34887021 | 39670337 | 38651161 | 41056853 | 36341189 | 41920941 | 34823173 | 34998789 | 41492421 | 44510701 | 41028013 | 41814689 | 34505129 | 44060497 |
| Q6P549 | Inppl1 | Phosphatidylinositol 3,4,5-trisphosphate 5-phosphatase 2 Mus musculus | 580.63 | 13.37 | 138973 | 11 | 13 | 10057740 | 8966307 | 9686605 | 9075967 | 7102800 | 9230877 | 9330438 | 7206574.5 | 9301967 | 10750849 | 10561061 | 9660663 | 8539417 | 6747587.5 | 10331949 |
| Q6P4S8 | Ints1 | Integrator complex subunit 1 Mus musculus | 39.05 | 0.91 | 245168 | 1 | 1 | 181085.66 | 288791.91 | 246775.52 | 1 | 318217.59 | 288312.88 | 262578.97 | 218283.77 | 259448.88 | 257156.125 | 364139.12 | 181178.69 | 224392.91 | 300517.28 | 267557 |
| Q3MHC2 | Ints11 | Integrator complex subunit 11 Rattus norvegicus 2 | 43.91 | 1.67 | 67841 | 1 | 1 | 974125.75 | 675339.88 | 783002.31 | 834236.25 | 706802.12 | 683132.38 | 775600.56 | 749491.81 | 714013.94 | 768205.44 | 691523.12 | 825723.12 | 770311.31 | 698363.44 | 804314.06 |
| Q68FR3 | Ints12 | Integrator complex subunit 12 Rattus norvegicus | 54.09 | 3.04 | 48477 | 1 | 1 | 1 | 1 | 1 | 1 | 1 | 1 | 1 | 1 | 1 | 1 | 1 | 208647.5 | 1 | 1 | 1 |
| Q8QZV7 | IntS13 | Integrator complex subunit 13 Mus musculus | 38.84 | 1.5 | 82780 | 1 | 1 | 1 | 1232785.62 | 1174522.25 | 1 | 1 | 1077236.12 | 1 | 1202119.38 | 943212.81 | 971062.06 | 1206571.88 | 665577.06 | 1102311.88 | 887036.56 | 687624.44 |
| Q66H58 | Ints14 | Integrator complex subunit 14 Rattus norvegicus 2 | 62.67 | 2.33 | 57144 | 1 | 1 | 247919.2 | 239601.92 | 277329.53 | 306764.06 | 318210.34 | 292951.66 | 318388.22 | 314561.53 | 350557.41 | 228554.59 | 286080.94 | 435300.53 | 158567.58 | 274709.72 | 339944.09 |

| | | | | | | | | | | | | | | | | | | | | | | |
|---|---|---|---|---|---|---|---|---|---|---|---|---|---|---|---|---|---|---|---|---|---|---|
| Q7TPD0 | Ints3 | Integrator complex subunit 3 Mus musculus | 52.77 | 1.63 | 117938 | 1 | 1 | 278796.5 | 139191.3 | 172355.92 | 147970.53 | 151160.78 | 1 | 1 | 1 | 217773.11 | 1 | 306172 | 1 | 1 | 1 | 1 |
| Q8CIM8 | Ints4 | Integrator complex subunit 4 Mus musculus | 226.28 | 7.57 | 108193 | 4 | 4 | 867053.75 | 534346.56 | 1268611.25 | 409629.25 | 935553.06 | 673290.38 | 719734.19 | 906167.25 | 972654.38 | 432687.16 | 1720347 | 578673.88 | 420905.97 | 1012101.12 | 784397.62 |
| Q7TQK1 | Ints7 | Integrator complex subunit 7 Mus musculus | 162.44 | 6 | 106861 | 4 | 4 | 744211.305 | 459033.56 | 468421.53 | 1178950.25 | 870439.88 | 384318.97 | 1049578.25 | 777661.38 | 710967.62 | 1006322.88 | 750275.88 | 964036.25 | 560252.81 | 752050.31 | 1295763.75 |
| Q80V86 | Ints8 | Integrator complex subunit 8 Mus musculus | 34.38 | 0.7 | 113353 | 1 | 1 | 931067.31 | 809143.56 | 771522.56 | 846315.19 | 706830.19 | 961879.62 | 509927.06 | 616791.12 | 881921.31 | 713690.44 | 855227.56 | 795989.69 | 826372.81 | 806654.31 | 716276.75 |
| Q8K114 | Ints9 | Integrator complex subunit 9 Mus musculus | 39.47 | 3.19 | 74078 | 1 | 1 | 404984 | 477826.62 | 556368.56 | 373949.28 | 446336.06 | 437459.03 | 404162.06 | 457520.03 | 588679.25 | 471955.0925 | 321814.06 | 442936.81 | 393073.53 | 459112.78 | 475979.12 |
| Q8K2V6 | Ipo11 | Importin-11 Mus musculus | 136.31 | 5.85 | 112416 | 3 | 3 | 1769204.38 | 847586.31 | 1007741.88 | 182755.36 | 1513397.75 | 1390356 | 727747.12 | 494801.16 | 1350326.75 | 706998.19 | 1406877.88 | 712491.56 | 1011145.69 | 1195178.88 | 568544.38 |
| Q8VI75 | Ipo4 | Importin-4 Mus musculus | 259.13 | 6.65 | 119275 | 6 | 7 | 2748288.5 | 2278374.25 | 3183473.75 | 2987209.5 | 3082576.75 | 2634927.25 | 2358494.5 | 2696956 | 2615069.25 | 2838891.75 | 2477404.5 | 2681924 | 2663751.5 | 2837902.5 | 2564580.5 |
| Q8BKC5 | Ipo5 | Importin-5 Mus musculus | 1521.09 | 37.65 | 123591 | 28 | 28 | 48631161 | 48013341 | 51696005 | 32527311 | 41215461 | 46163733 | 44476613 | 49711777 | 52218445 | 46878661 | 53441677 | 52977557 | 44401093 | 51660825 | 46518005 |
| Q9EPL8 | Ipo7 | Importin-7 Mus musculus | 638.96 | 15.9 | 119486 | 11 | 12 | 21407035 | 12272100 | 17098311 | 10782678 | 9871825 | 12840737 | 9969355 | 20682309 | 24742109 | 9693170 | 13021430 | 15923448 | 10266249 | 20567949 | 12649033 |
| Q7TMY7 | Ipo8 | Importin-8 Mus musculus | 113.41 | 4.16 | 117078 | 2 | 3 | 222764.27 | 164700.03 | 2676387 | 134075.33 | 156044.33 | 107184.09 | 1 | 3487555.5 | 3065259.25 | 127487.95 | 120593.69 | 2167256.25 | 179611.88 | 3413050 | 142118.19 |
| Q91YE6 | Ipo9 | Importin-9 Mus musculus | 1449.63 | 33.72 | 116052 | 22 | 22 | 43797889 | 48878905 | 45943877 | 43513609 | 41303201 | 42467081 | 36107033 | 50419781 | 54053473 | 40134745 | 59078193 | 51752917 | 45524673 | 55024733 | 49172077 |
| Q9JKF1 | Iqgap1 | Ras GTPase-activating-like protein IQGAP1 Mus musculus | 6063.399 | 68.56 | 188742 | 90 | 96 | 519050561 | 488924513 | 508137121 | 496845985 | 463244961 | 484879105 | 564117761 | 515419329 | 514671393 | 502135009 | 536942913 | 547687681 | 493947617 | 500108257 | 597754369 |
| Q3UQ44 | Iqgap2 | Ras GTPase-activating-like protein IQGAP2 Mus musculus | 196.71 | 2.6 | 180528 | 1 | 5 | 22371009 | 24136241 | 22486175 | 24295237 | 22065695 | 21492253 | 29924151 | 21181037 | 20584471 | 26895545 | 27185649 | 25627359 | 26377093 | 23646155 | 30108579 |
| Q8R0S2 | Iqsec1 | IQ motif and SEC7 domain-containing protein 1 Mus musculus | 137.24 | 4.37 | 108015 | 3 | 3 | 1 | 289479.97 | 171411.81 | 1 | 147244.22 | 130430.73 | 1 | 169546.06 | 130048.69 | 1 | 238410.58 | 128315.45 | 171445.0625 | 174276.75 | 144777.47 |
| Q8R4K2 | Irak4 | Interleukin-1 receptor-associated kinase 4 Mus musculus | 74.58 | 3.7 | 50872 | 1 | 1 | 1060406.25 | 1101917 | 1002067.94 | 1146826.38 | 1106412.25 | 878716.38 | 1016870 | 1011746.06 | 947959.81 | 941204.88 | 1032768.5 | 1008588.38 | 1044315.12 | 1044499.31 | 784247.56 |
| E9Q1P8 | Irf2bp2 | Interferon regulatory factor 2-binding protein 2 Mus musculus | 121.44 | 7.19 | 59292 | 1 | 2 | 376504.16 | 340499.75 | 560696.69 | 29403.25 | 326775.9625 | 172124.7 | 511121.22 | 497689.59 | 381056.47 | 694325.94 | 190683.78 | 583130.38 | 528780 | 486550.62 | 629357 |
| Q5EIC4 | Irf2bpl | Interferon regulatory factor 2-binding protein-like Rattus norvegicus | 79.29 | 4.73 | 81496 | 1 | 2 | 886338.5 | 1033053.44 | 1089360.38 | 573580.88 | 509385.41 | 755249.88 | 1070262.62 | 1349956.25 | 1210404 | 1296108.75 | 769383.75 | 1135222.5 | 1106890.5 | 1133067.62 | 1167742.38 |
| P56477 | Irf5 | Interferon regulatory factor 5 Mus musculus | 469.74 | 21.33 | 56005 | 8 | 8 | 6195636 | 4725676.5 | 6614123.5 | 5685785.5 | 5408696 | 6617064 | 4813781.5 | 7065826 | 6369581.5 | 5236516 | 5406708 | 6193405.5 | 4283413 | 6160161.5 | 5824319 |
| Q60766 | Irgm1 | Immunity-related GTPase family M protein 1 Mus musculus | 382.54 | 19.07 | 46552 | 7 | 8 | 6445768.5 | 4892167.5 | 4781422.5 | 3869175 | 4389789 | 3603796.25 | 3929282.25 | 4063158.5 | 4858913 | 3849770 | 4091486 | 3651811.75 | 3988766.75 | 4910448.5 | 4166172.25 |
| Q64339 | Isg15 | Ubiquitin-like protein ISG15 Mus musculus | 59.95 | 11.18 | 17898 | 1 | 1 | 239879.86 | 383989.53 | 247970.45 | 363461.97 | 266334.69 | 467434.235 | 285057.97 | 935167.31 | 338083.44 | 311428.22 | 335288.44 | 144675.77 | 304836.81 | 387612.03 | 377774.41 |
| Q6I7R3 | Isoc1 | Isochorismatase domain-containing protein 1 Rattus norvegicus | 171.97 | 14.81 | 32002 | 3 | 3 | 3014118 | 2495339.5 | 2088693 | 479990.72 | 3078048.75 | 2776553.5 | 2269232 | 2544937.5 | 2818104.25 | 2617664.75 | 2896524 | 2239708.5 | 2743494.75 | 2037279.38 | 2860258 |
| P85094 | Isoc2a | Isochorismatase domain-containing protein 2A Mus musculus | 237.74 | 40.78 | 22417 | 4 | 4 | 3103277.25 | 2942515 | 2963688.25 | 2998752.25 | 3032113.75 | 3710737.25 | 4344897 | 4017708.5 | 2830442 | 3537400.75 | 3563840.5 | 3177828.5 | 3866942 | 2891957 | 4809830 |
| Q568Z6 | Ist1 | IST1 homolog Rattus norvegicus | 147.61 | 9.29 | 39942 | 3 | 3 | 1452078.5 | 824591 | 902055.94 | 1459026 | 1172317.5 | 666752.25 | 1087487.12 | 1530563.25 | 1371854.62 | 552077.69 | 1151781.75 | 796130.5 | 1242578.5 | 1269047.75 | 1190753.75 |
| Q9JHU9 | Isyna1 | Inositol-3-phosphate synthase 1 Mus musculus | 663.52 | 29.26 | 60932 | 11 | 11 | 19801961 | 23093807 | 24332087 | 22423971 | 21288923 | 21773973 | 21971611 | 22245865 | 23901375 | 18483153 | 20496993 | 25413549 | 25201799 | 26896711 | 25714635 |
| Q00651 | Itga4 | Integrin alpha-4 Mus musculus | 78.48 | 2.12 | 115696 | 2 | 2 | 1 | 1 | 758519.5 | 1 | 265363.81 | 239309.19 | 1 | 1 | 1 | 1 | 1 | 1 | 1 | 1 | 1 |
| P11688 | Itga5 | Integrin alpha-5 Mus musculus | 466.22 | 15.76 | 115043 | 10 | 10 | 5198483 | 3406540.75 | 7189628 | 4701444 | 4655489.5 | 6731099 | 10346844 | 6844839 | 6121590.5 | 7368993 | 4685426.5 | 5342822.5 | 5904770.5 | 5756229 | 7288680.5 |
| P05555 | Itgam | Integrin alpha-M Mus musculus | 1606.42 | 30.01 | 127481 | 24 | 24 | 68837145 | 72955153 | 78229753 | 61495005 | 73197409 | 59872513 | 67063117 | 64562041 | 64152969 | 60429577 | 67800809 | 69144889 | 71654849 | 72700105 | 63264653 |
| P09055 | Itgb1 | Integrin beta-1 Mus musculus | 902.36 | 27.19 | 88231 | 16 | 16 | 32506807 | 38435029 | 33373171 | 29439243 | 34020877 | 36107433 | 34103225 | 35279805 | 37318577 | 30563649 | 38189357 | 30102507 | 33182785 | 36734729 | 31689991 |
| P11835 | Itgb2 | Integrin beta-2 Mus musculus | 1755.53 | 44.75 | 85026 | 27 | 27 | 89636257 | 102390457 | 115078817 | 77016329 | 99660521 | 98875409 | 84958177 | 86078857 | 96993961 | 76166649 | 100868017 | 75187985 | 89682945 | 94862361 | 80953585 |
| O70309 | Itgb5 | Integrin beta-5 Mus musculus | 72.13 | 2.88 | 87909 | 2 | 2 | 1 | 1 | 1 | 1 | 1 | 117823.55 | 1 | 133851.56 | 138843.19 | 1 | 97349.78 | 107863.11 | 1 | 1 | 107525.8 |
| O89051 | Itm2b | Integral membrane protein 2B Mus musculus | 242.83 | 19.55 | 30260 | 4 | 4 | 3923463.5 | 4806815 | 4322810 | 3662523.25 | 2545112 | 4504029.5 | 4087874.5 | 4989060 | 6301094 | 3439098.25 | 3363689.5 | 4677718.5 | 3523362.5 | 4643131 | 3403569.25 |
| Q5PQL7 | Itm2c | Integral membrane protein 2C Rattus norvegicus | 105.92 | 11.52 | 30480 | 2 | 2 | 1782204.5 | 1484172.5 | 1676717.88 | 1326225.5 | 1185751 | 2260105.5 | 1728229 | 2192703.75 | 2326372 | 1278359.25 | 1250860.5 | 1330951.25 | 1283212.5 | 1758725.25 | 1141780.12 |
| Q9D892 | Itpa | Inosine triphosphate pyrophosphatase Mus musculus | 303.95 | 44.44 | 21897 | 6 | 6 | 11202732 | 6558981 | 11430209 | 10363730 | 9197700 | 9908752 | 10321356 | 9305027 | 6231827.5 | 6546965 | 7362660.5 | 10351330 | 10891811 | 9881259 | 10256157 |
| P11881 | Itpr1 | Inositol 1,4,5-trisphosphate receptor type 1 Mus musculus | 137.18 | 1.2 | 313167 | 1 | 3 | 2044397 | 2403410 | 2190753.75 | 2525277.75 | 1417940.88 | 2822627.25 | 2070549 | 2169437 | 2689122 | 2119557.5 | 2152588.25 | 2149111.25 | 1326175.38 | 1729214 | 2155706.75 |
| P29995 | Itpr2 | Inositol 1,4,5-trisphosphate receptor type 2 Rattus norvegicus | 202.91 | 1.93 | 307058 | 3 | 5 | 2451430.75 | 2791530.5 | 2356905.5 | 2536544.75 | 1545694 | 2881485.5 | 2378699.5 | 2615078.5 | 2770134 | 2583782 | 2397446.75 | 2243037.25 | 1777685.5 | 2098917.75 | 2258102 |
| P70227 | Itpr3 | Inositol 1,4,5-trisphosphate receptor type 3 Mus musculus | 131.55 | 1.91 | 304275 | 4 | 4 | 639147.94 | 1556119.5 | 1574866.25 | 1596880.12 | 1718204 | 1977254.38 | 1644122.38 | 2018316.5 | 1746773.75 | 1735237.62 | 3644302.75 | 1499567 | 1519330.5 | 1491310.5 | 1839541.25 |
| Q9JHI5 | Ivd | Isovaleryl-CoA dehydrogenase, mitochondrial Mus musculus | 286 | 17.22 | 46325 | 7 | 7 | 9670996 | 6238688 | 11463977 | 12403107 | 9045478 | 5656052.5 | 12947514 | 10004865 | 10238923 | 12224995 | 6144863 | 11338339 | 11112762 | 6369512.5 | 10201539 |
| Q4KM64 | Jagn1 | Protein jagunal homolog 1 Rattus norvegicus | 63.67 | 6.56 | 21102 | 1 | 1 | 1 | 1 | 1 | 1 | 1954519.75 | 1568794.75 | 751355.94 | 1422553.62 | 295775.28 | 1009619.898 | 401370.41 | 47676.22 | 1894798.75 | 369881.97 | 678431.8375 |
| P52332 | Jak1 | Tyrosine-protein kinase JAK1 Mus musculus | 169.62 | 4.94 | 133367 | 3 | 3 | 1758193.25 | 1617794.75 | 2500729.5 | 1903785.5 | 2008187.25 | 417229.25 | 1864966.25 | 2045325 | 2062455 | 689220.38 | 2282524.75 | 1856582.12 | 2220681.25 | 2355015.5 | 2361027.5 |
| P97825 | Jpt1 | Jupiter microtubule associated homolog 1 Mus musculus | 106.39 | 27.27 | 16081 | 2 | 2 | 2924789.25 | 4087840.5 | 3059245.5 | 4315181 | 3478096 | 2223103.5 | 2421793.5 | 2452645.5 | 2560232 | 3651883.75 | 887516.75 | 2085826.38 | 2894699.25 | 2233320.75 | 816898.06 |
| Q5BK20 | Jpt2 | Jupiter microtubule associated homolog 2 Rattus norvegicus | 105.31 | 18.42 | 20070 | 2 | 2 | 1159687.88 | 1111391.12 | 1737516.38 | 1378039.75 | 1613009.25 | 1017053.81 | 2687015.25 | 1100289.12 | 542921.06 | 1500170 | 1453064 | 1129826 | 1328611.75 | 855160.62 | 1899130.62 |
| Q02257 | Jup | Junction plakoglobin Mus musculus | 66.82 | 2.42 | 81801 | 1 | 1 | 681373.81 | 1 | 1 | 1 | 1 | 1 | 1 | 168941.64 | 1 | 1 | 1 | 1 | 206498.7 | 1 | 1 |
| Q99MN1 | Kars | Lysine--tRNA ligase Mus musculus | 801.4 | 33.45 | 67840 | 4 | 16 | 25529641 | 27604621 | 26149621 | 23147309 | 25424887 | 27076435 | 28545063 | 28187125 | 30812489 | 27797987 | 30966819 | 28971159 | 25493319 | 25335507 | 28560621 |
| Q5SVQ0 | Kat7 | Histone acetyltransferase KAT7 Mus musculus | 181.99 | 7.01 | 70641 | 3 | 3 | 2823160.75 | 1656111.88 | 1134877.5 | 950151 | 1963719.5 | 3009285 | 1390419.5 | 1259412.12 | 1295718.25 | 900977.75 | 2159809.5 | 1166540.38 | 1638566.5 | 1332219.75 | 2433752.5 |
| Q80UY2 | Kcmf1 | E3 ubiquitin-protein ligase KCMF1 Mus musculus | 169.46 | 8.14 | 41791 | 2 | 2 | 424839.94 | 440292.12 | 223873.81 | 451242.31 | 465944.56 | 199894.31 | 405224.78 | 255714.39 | 515153.78 | 381608 | 408686.88 | 309017.06 | 354848.31 | 250810.95 | 228390.55 |
| P62482 | Kcnab2 | Voltage-gated potassium channel subunit beta-2 Mus musculus | 379.23 | 35.97 | 41021 | 7 | 7 | 11593159 | 12929697 | 12474758 | 13290734 | 11773923 | 11195759 | 12852774 | 12725711 | 12060590 | 11924525 | 11029746 | 10607653 | 11235901 | 11154022 | 11206591 |
| Q6WVG3 | Kctd12 | BTB/POZ domain-containing protein KCTD12 Mus musculus | 512.69 | 36.39 | 35892 | 10 | 10 | 14306049 | 15819779 | 15026211 | 17265059 | 23246457 | 14858101 | 11815032 | 14306410 | 15243832 | 14383070 | 20399825 | 16458715 | 15110258 | 15611471 | 16566945 |
| Q9JHP7 | Kdelc1 | KDEL motif-containing protein 1 Mus musculus | 240.67 | 12.35 | 57985 | 5 | 5 | 1223272.62 | 1045887.25 | 1499118.38 | 367626.06 | 736394.88 | 760424.75 | 1062921.25 | 1358616.5 | 1604047 | 895302.56 | 1408395 | 1754977.12 | 631958.62 | 1280740.75 | 1182166.62 |
| Q569A6 | Kdelr1 | ER lumen protein-retaining receptor 1 Rattus norvegicus 2 | 97.28 | 10.38 | 24546 | 2 | 2 | 1144506.25 | 213300.92 | 162754.42 | 189118.08 | 427419.9175 | 1 | 1 | 1 | 125173.32 | 228693.59 | 257044.1575 | 354419.88 | 174475.75 | 360029.91 | 139251.09 |
| Q6ZQ88 | Kdm1a | Lysine-specific histone demethylase 1A Mus musculus | 155.68 | 5.16 | 92851 | 2 | 2 | 1145119.88 | 987750.56 | 1188586.62 | 1060773.375 | 921636.44 | 1216239.25 | 1167466.38 | 1330090.62 | 1169766.25 | 1003112.06 | 1073926 | 963921.25 | 708871.75 | 1150950.25 | 1144090.88 |
| P59997 | Kdm2a | Lysine-specific demethylase 2A Mus musculus | 161.2 | 2.76 | 132680 | 3 | 3 | 248617.81 | 1953887 | 2227117 | 1956581.5 | 3216276.75 | 2690116.5 | 3182731.75 | 2281076 | 2863007.25 | 1855299.75 | 1744461.38 | 2732460 | 3370199 | 2785649.5 | 2462160.25 |
| Q3UXZ9 | Kdm5a | Lysine-specific demethylase 5A Mus musculus | 47.87 | 0.77 | 192216 | 1 | 1 | 227078.34 | 255711.59 | 206055.94 | 242048.64 | 279348.69 | 196326.45 | 161289.11 | 292725.75 | 257782.3 | 181820.17 | 180838.3 | 195585.7 | 144043.59 | 268312.97 | 223733.31 |
| P41230 | Kdm5c | Lysine-specific demethylase 5C Mus musculus | 40.19 | 0.84 | 175313 | 1 | 1 | 105584.94 | 244402.84 | 183996.55 | 201710.8 | 210575.95 | 115698.74 | 92458.02 | 118737.34 | 114577.02 | 173868.47 | 178486.28 | 149903.62 | 186735.84 | 222474.06 | 153017.89 |
| Q6GV12 | Kdsr | 3-ketodihydrosphingosine reductase Mus musculus | 65.45 | 4.22 | 35956 | 1 | 1 | 1429564 | 1541790.62 | 1477920.38 | 1612317.12 | 1591729.62 | 1142529.62 | 1437055.188 | 1231357 | 1707083.88 | 1667250.25 | 942023.81 | 1137130.25 | 1547917.75 | 1543999.12 | 1258465.62 |
| Q60749 | Khdrbs1 | KH domain-containing, RNA-binding, signal transduction-associated protein 1 Mus musc | 156.47 | 16.48 | 48371 | 4 | 4 | 1929417.88 | 1149907.12 | 2048198 | 1872508.62 | 2247208.25 | 1966188 | 4700122 | 1943447.75 | 1437056.25 | 2634694 | 2135774 | 2756601.25 | 2056985.25 | 1601015.38 | 3757785 |
| Q3U0V1 | Khsrp | Far upstream element-binding protein 2 Mus musculus | 1332.31 | 48.93 | 76775 | 20 | 22 | 56492473 | 69930305 | 58396933 | 74231513 | 77922577 | 56382345 | 73145153 | 59536605 | 63281065 | 76340769 | 83107057 | 67387905 | 76535313 | 60973245 | 69253337 |
| Q6P9P6 | Kif11 | Kinesin-like protein KIF11 Mus musculus | 575.89 | 14.16 | 118027 | 9 | 9 | 1801478.25 | 1844002.5 | 1682134.75 | 1739286 | 1683549 | 2451006.5 | 2595642.75 | 2073151 | 1864301.5 | 2058784.5 | 1944497.75 | 2636322.5 | 2014056 | 2490137.25 | 2025372.5 |
| Q6P9L6 | Kif15 | Kinesin-like protein KIF15 Mus musculus | 215.27 | 3.53 | 160120 | 3 | 3 | 613811.88 | 450786.91 | 756072.19 | 419258.38 | 885964.5 | 1138113.62 | 695290.75 | 665664.12 | 714826.44 | 1127235.12 | 1048778.88 | 1160518.25 | 932265.19 | 1201514.38 | 1151807.75 |
| O35071 | Kif1c | Kinesin-like protein KIF1C Mus musculus | 63.83 | 1.18 | 122434 | 1 | 1 | 125842.36 | 103534.19 | 129607.06 | 111424.1 | 289267.75 | 182013.83 | 139674.75 | 184975.47 | 205745.11 | 149609.98 | 167668.61 | 150167.09 | 258670 | 175057.23 | 141959.58 |
| P97329 | Kif20a | Kinesin-like protein KIF20A Mus musculus | 57.07 | 3.16 | 99876 | 1 | 1 | 1 | 1 | 1 | 384765.12 | 260453.38 | 1 | 1 | 1 | 1 | 267649.09 | 131148.83 | 135673.23 | 182690.0375 | 194064.97 | 269873.12 |
| F1M5N7 | Kif21b | Kinesin-like protein KIF21B Rattus norvegicus | 236.75 | 5.2 | 182563 | 5 | 5 | 1035331.31 | 1890741 | 1367951.38 | 1164323.38 | 1275484.25 | 1136797.75 | 1446129.88 | 1210439.75 | 1566810 | 1295998 | 1287631.25 | 1420333.25 | 1007559.31 | 1528406.75 | 1412974.75 |
| Q3V300 | Kif22 | Kinesin-like protein KIF22 Mus musculus | 78.78 | 3.94 | 73190 | 2 | 2 | 1265744.75 | 1064443.25 | 957997.25 | 1238799.25 | 931019.81 | 794203.44 | 776615.88 | 968571.38 | 919149.44 | 734712 | 256059.19 | 425881.59 | 417362.6175 | 313512.31 | 673997.38 |
| E9Q5G3 | Kif23 | Kinesin-like protein KIF23 Mus musculus | 46.24 | 1.26 | 108776 | 1 | 1 | 896212.19 | 1619307 | 1253198.38 | 1764848.75 | 588226.38 | 982458.81 | 1148011 | 1156760.12 | 1362857.25 | 1162521.795 | 687108.25 | 1109488.685 | 1330163.12 | 1322606.12 | 1098077.25 |
| P28740 | Kif2a | Kinesin-like protein KIF2A Mus musculus | 102.84 | 2.7 | 79756 | 2 | 2 | 573731.31 | 1145016 | 1255727.5 | 935530.19 | 599947.31 | 739086 | 797987.19 | 1236618 | 852078.81 | 621762.31 | 1052326.75 | 1290619 | 562968.62 | 1334671 | 606596.12 |
| Q922S8 | Kif2c | Kinesin-like protein KIF2C Mus musculus | 95.48 | 4.72 | 81085 | 2 | 2 | 242709.95 | 1 | 275403.44 | 1 | 402230.97 | 641076.12 | 495552.03 | 824558.69 | 536093.19 | 496042.81 | 886416.38 | 885364.88 | 568168.5 | 839148.69 | 272486.38 |
| Q61771 | Kif3b | Kinesin-like protein KIF3B Mus musculus | 116.12 | 2.54 | 85288 | 1 | 1 | 2063001 | 2194726 | 1 | 1 | 1314586.12 | 2236246.25 | 2088933.38 | 1799780.5 | 3190120.5 | 2328770.158 | 3012843 | 1250017.75 | 1523129.62 | 2788953 | 2618484.5 |
| P33174 | Kif4 | Chromosome-associated kinesin KIF4 Mus musculus | 181.14 | 3.66 | 139519 | 4 | 4 | 2610901.75 | 2644056.5 | 2799358.5 | 1277284.25 | 2421715.5 | 1690237.25 | 2004129.5 | 1998186.5 | 2342025.75 | 1567213.38 | 1986585.25 | 910432.12 | 1607991.5 | 2021669.25 | 2125684.75 |
| P33175 | Kif5a | Kinesin heavy chain isoform 5A Mus musculus | 188.16 | 4.58 | 117019 | 0 | 4 | 3830728.5 | 2069237.5 | 2084377.88 | 1940073.25 | 2864564 | 2081468.38 | 3136549.5 | 2016880.5 | 1503613.25 | 3018181.5 | 2306496.5 | 2793225.5 | 2174731 | 1621242.75 | 1501071.38 |
| Q2PQA9 | Kif5b | Kinesin-1 heavy chain Rattus norvegicus | 993.88 | 23.36 | 109531 | 13 | 17 | 10117376 | 11674566 | 10884703 | 12310351 | 9799467 | 12104807 | 15252587 | 11680337 | 12620121 | 12048692 | 11850295 | 12001167 | 10322054 | 9823792 | 14142792 |
| P28738 | Kif5c | Kinesin heavy chain isoform 5C Mus musculus | 193.65 | 5.13 | 109275 | 1 | 4 | 1634748.62 | 1525816.5 | 2336881.75 | 1455427.75 | 1495552.12 | 2146269.75 | 2491676.5 | 2663655 | 2093496.62 | 1519343.25 | 1628108.62 | 2689723.5 | 1717549.75 | 2006546.25 | 2130530.5 |
| P70188 | Kifap3 | Kinesin-associated protein 3 Mus musculus | 232.68 | 5.8 | 91291 | 3 | 3 | 1491804.12 | 1501712.5 | 1419731.25 | 1250890.5 | 1425476 | 1625760.75 | 1426412.38 | 1774483 | 1699261 | 1560379.75 | 1216867.25 | 1421363.5 | 965687.56 | 1567311.25 | 1537212.62 |
| O88447 | Klc1 | Kinesin light chain 1 Mus musculus | 38.25 | 2.4 | 61450 | 1 | 1 | 804088.31 | 841914.5 | 922713.69 | 1119057.12 | 1045303.44 | 805650.25 | 985688.5775 | 955759.94 | 953064.62 | 1228279.5 | 914829.25 | 983760.12 | 909027.94 | 988359.81 | 986009.62 |
| Q921I2 | Klhdc4 | Kelch domain-containing protein 4 Mus musculus 2 | 52.98 | 2.57 | 64860 | 1 | 1 | 1290601.62 | 1177804.25 | 1553057.88 | 1303753.75 | 1331525 | 1186300.75 | 934673.69 | 1011991.88 | 1093696.12 | 1246541.75 | 1430117.75 | 1322820.25 | 1199798.88 | 1135465.5 | 1473135.38 |
| Q8C3Y4 | Kntc1 | Kinetochore-associated protein 1 Mus musculus | 106.39 | 1.63 | 250358 | 3 | 3 | 1642317.75 | 1803738.75 | 1552117.5 | 1696947.75 | 1526644.5 | 1469792.25 | 1137607 | 1726752.75 | 1686255.75 | 1766130.75 | 1373876.75 | 1253331.25 | 1363236.25 | 1336416.62 | 1132174.62 |
| P83953 | Kpna1 | Importin subunit alpha-5 Rattus norvegicus | 477.59 | 22.68 | 60137 | 3 | 7 | 4530387.5 | 4801344.5 | 4995068.5 | 4916343 | 3771598.75 | 4669032 | 5391840 | 4673739 | 5062438 | 4909408.5 | 5698086 | 4799731 | 5412978 | 5630760 | 6130915.5 |
| P52293 | Kpna2 | Importin subunit alpha-1 Mus musculus | 1192.77 | 48.77 | 57928 | 17 | 17 | 17248655 | 15881089 | 20675819 | 17715973 | 15642924 | 19244429 | 16398655 | 19726255 | 19123321 | 18890589 | 19595865 | 22154393 | 19340411 | 17033393 | 18748125 |
| O35344 | Kpna3 | Importin subunit alpha-4 Mus musculus | 574.8 | 27.45 | 57773 | 9 | 10 | 4867773 | 11178238 | 5797838.5 | 4495336 | 7245831 | 11884251 | 10734435 | 10022787 | 11952106 | 10560396 | 4340798 | 9237996 | 12659903 | 11707558 | 10227558 |
| O35343 | Kpna4 | Importin subunit alpha-3 Mus musculus | 520.35 | 26.1 | 57923 | 7 | 8 | 7532582 | 9383363 | 8000332 | 6291915 | 9114079 | 8328200.5 | 9957676 | 11073465 | 10662448 | 5345584 | 9601172 | 8058248 | 7022709.5 | 10534779 | 8294521 |
| Q56R16 | Kpna5 | Importin subunit alpha-6 Rattus norvegicus | 233.82 | 9.89 | 60299 | 1 | 5 | 4009986.5 | 4253468.5 | 4582332 | 4451326 | 2915368.25 | 3943884 | 9248177 | 4270956 | 4658801.5 | 4276679.5 | 4870079.5 | 8669782 | 4918613 | 5006213.5 | 4926133 |
| O35345 | Kpna6 | Importin subunit alpha-7 Mus musculus | 324.87 | 13.99 | 59964 | 2 | 5 | 4916037.5 | 3161914.75 | 3009315.75 | 5540338.5 | 1484844.38 | 7513101.5 | 6013622.5 | 2787555.25 | 6255277 | 4864986 | 5655620 | 3176839.5 | 5648979 | 5168717.5 | 6277001.5 |
| P70168 | Kpnb1 | Importin subunit beta-1 Mus musculus | 1892.39 | 55.25 | 97184 | 30 | 31 | 89736977 | 92410529 | 92801505 | 78001705 | 70889225 | 94317601 | 76582969 | 98008273 | 114190881 | 82001681 | 107822257 | 106825793 | 80468561 | 98115201 | 92251809 |
| Q8BGA5 | Krr1 | KRR1 small subunit processome component homolog Mus musculus | 104.57 | 7.89 | 43538 | 3 | 3 | 6586508 | 7482851.5 | 6580950 | 8051483 | 7085955.5 | 5609794 | 6059442.5 | 6982498.5 | 7120762.5 | 7176398 | 7432474.5 | 6977231 | 6772283 | 7441396 | 5836662 |
| Q61595 | Ktn1 | Kinectin Mus musculus | 162.01 | 3.47 | 152592 | 3 | 3 | 362564.12 | 347393.81 | 616992.12 | 217837.8 | 567984.19 | 452370.25 | 612005.81 | 724182.81 | 520119.25 | 840537.62 | 1057530.88 | 619798.88 | 572777.88 | 426352.47 | 610888.25 |
| Q8BZT9 | Lacc1 | Laccase domain-containing protein 1 Mus musculus | 54.75 | 3.95 | 47514 | 1 | 1 | 1 | 311313.31 | 280255.16 | 1 | 407837.97 | 204503.05 | 242696.11 | 229735.52 | 241878.66 | 291090.69 | 228031.075 | 211301.77 | 257644.12 | 223004.33 | 220174.08 |
| Q9EP89 | Lactb | Serine beta-lactamase-like protein LACTB, mitochondrial Mus musculus | 233.63 | 10.89 | 60705 | 5 | 5 | 3384683 | 3194884 | 2470866 | 3067446 | 1985797.75 | 3470037.25 | 2204035 | 2886756 | 2604449.25 | 2783187.5 | 3149065 | 2522639.5 | 2521967 | 3012823 | 2511450.5 |
| Q99KR3 | Lactb2 | Endoribonuclease LACTB2 Mus musculus | 137.57 | 11.46 | 32754 | 3 | 3 | 3368817.5 | 2992152.5 | 3056815.25 | 3435738 | 3577917 | 3806317.25 | 3899327 | 3062497.25 | 3133776 | 4400828.5 | 3753587.75 | 3184125.5 | 3549928.5 | 3257435 | 3829579.5 |
| P11438 | Lamp1 | Lysosome-associated membrane glycoprotein 1 Mus musculus | 568.22 | 28.33 | 43865 | 8 | 8 | 43631817 | 55507593 | 51127957 | 45247497 | 51616885 | 49241773 | 38503721 | 58262329 | 73058577 | 38721973 | 47512193 | 61312385 | 46322093 | 57319001 | 48128241 |
| P17047 | Lamp2 | Lysosome-associated membrane glycoprotein 2 Mus musculus | 263.8 | 12.29 | 45681 | 5 | 5 | 142638177 | 104922305 | 105360225 | 135854225 | 149493297 | 146733633 | 117929505 | 173657649 | 128253185 | 105396145 | 107065889 | 134515185 | 132805169 | 149650673 | 117587425 |
| Q6P791 | Lamtor1 | Ragulator complex protein LAMTOR1 Rattus norvegicus | 96.67 | 16.15 | 17721 | 2 | 2 | 3387687.5 | 3668629.5 | 2649623.5 | 3601528.25 | 3677311 | 3142082.5 | 3104773 | 3496276.5 | 2956564 | 3404064 | 2949646 | 2748305 | 3210982 | 2806524 | 1841028.75 |
| Q9JHS3 | Lamtor2 | Ragulator complex protein LAMTOR2 Mus musculus | 327.95 | 75.2 | 13480 | 5 | 5 | 3061190 | 3380003 | 2833678 | 3459445 | 3677292 | 2742901 | 3116119 | 3494620.5 | 2205079 | 3036534 | 3076171.75 | 2802734 | 3314180.25 | 3202927.25 | 2806877.5 |
| O88653 | Lamtor3 | Ragulator complex protein LAMTOR3 Mus musculus | 54.18 | 8.87 | 13553 | 1 | 1 | 332034.62 | 235556.84 | 417342.41 | 1 | 1 | 374260.66 | 279995.5 | 450683.44 | 862488.25 | 491856.9625 | 449411.56 | 401068.88 | 423667.36 | 437494.56 | 406694.44 |
| Q9D1L9 | Lamtor5 | Ragulator complex protein LAMTOR5 Mus musculus | 51.75 | 21.98 | 9642 | 1 | 1 | 2614597.25 | 2651509.75 | 3639202.75 | 2723047 | 3201065.75 | 2881602.25 | 1526875.75 | 2855783.75 | 2257623.75 | 2555237.25 | 2749118.25 | 2294180.75 | 2660481 | 2713110 | 2851787.5 |
| O89112 | Lancl1 | LanC-like protein 1 Mus musculus | 192.41 | 13.28 | 45341 | 4 | 4 | 2849887.5 | 2832787.5 | 2081036 | 2611689.75 | 2867080 | 2747133.75 | 2304092.25 | 3018422 | 4008795 | 3198304.5 | 2656598.5 | 2543225 | 2676268 | 3103280 | 2644585 |
| Q9CPY7 | Lap3 | Cytosol aminopeptidase Mus musculus | 787.76 | 39.5 | 56141 | 15 | 15 | 15427193 | 19944985 | 18782535 | 19058895 | 12815697 | 21893189 | 20372905 | 22657309 | 22645337 | 15088258 | 19578207 | 17564857 | 15951646 | 20069951 | 17617479 |
| Q6ZQ58 | Larp1 | La-related protein 1 Mus musculus | 512.21 | 12.5 | 121125 | 10 | 10 | 2039866.5 | 3495226.75 | 3501751.75 | 3309802.25 | 3028093.25 | 3369430.25 | 7568093 | 2056106.25 | 2352232 | 2470837 | 3547621.25 | 3619632 | 3068506.75 | 2532374.75 | 4482533.5 |
| Q8BWW4 | Larp4 | La-related protein 4 Mus musculus | 347.74 | 10.29 | 79763 | 6 | 6 | 5036020 | 4904411 | 5728868 | 5108949 | 6015560 | 2512211 | 4206557.5 | 5405228 | 5914432.5 | 4528595 | 3486999.75 | 5571416.5 | 6233404 | 3552225.5 | 5227052.5 |
| Q6A0A2 | Larp4b | La-related protein 4B Mus musculus | 316.71 | 11.2 | 81627 | 6 | 6 | 832142.81 | 3606668.5 | 2519778.75 | 3920381 | 2112063.25 | 3619525.75 | 2041931 | 3747437.5 | 2960972 | 1824049 | 3450534.5 | 1689881.75 | 3454085.5 | 2479531.75 | 1814593.5 |
| Q05CL8 | Larp7 | La-related protein 7 Mus musculus | 297.48 | 11.4 | 64802 | 5 | 5 | 10513111 | 11670235 | 7440189.5 | 11271201 | 11376124 | 9821228 | 5042094.5 | 6675311 | 8297351 | 5860961 | 6760591.5 | 5860076 | 7100140 | 7035883 | 6525869 |
| Q8BMJ2 | Lars | Leucine--tRNA ligase, cytoplasmic Mus musculus | 1924.83 | 41.17 | 134192 | 41 | 41 | 46871145 | 43986337 | 46065385 | 49140165 | 46022913 | 42356333 | 52942797 | 41958525 | 47097969 | 51880185 | 44187517 | 47754353 | 45113633 | 45964793 | 47566769 |
| A2BE28 | Las1l | Ribosomal biogenesis protein LAS1L Mus musculus | 58.04 | 2.19 | 89414 | 1 | 1 | 697141.19 | 724645.56 | 741313.75 | 604094.31 | 491069.44 | 695221.62 | 684222.5175 | 703625.19 | 838405.88 | 499637.38 | 342790.69 | 384772.88 | 671505.5 | 649520.81 | 302934.84 |
| Q61792 | Lasp1 | LIM and SH3 domain protein 1 Mus musculus | 418.55 | 28.9 | 29994 | 8 | 8 | 11757164 | 14238055 | 13827363 | 17997865 | 12548104 | 11233263 | 25912285 | 12255803 | 10470048 | 12864840 | 14693720 | 15893484 | 15020059 | 12232229 | 20577869 |
| Q3U9G9 | Lbr | Lamin-B receptor Mus musculus | 538.78 | 18.21 | 71440 | 8 | 9 | 20571297 | 4351733 | 7152046 | 7971330 | 8114032.5 | 7387927 | 10733897 | 5828516 | 4907955.5 | 9568443 | 7711392 | 7775682 | 6254605 | 6010312 | 10137497 |
| Q6P4Z6 | Lcmt1 | Leucine carboxyl methyltransferase 1 Rattus norvegicus | 44.66 | 2.71 | 38330 | 1 | 1 | 2115761.75 | 2183631.25 | 1710164.75 | 2253409.25 | 1937010.12 | 1561821.62 | 1717521 | 1571419.62 | 1810043.12 | 1668137.88 | 1979706.62 | 1382952.62 | 1519326.25 | 1621406.12 | 1396004.88 |
| Q61233 | Lcp1 | Plastin-2 Mus musculus | 4110.059 | 87.88 | 70149 | 48 | 62 | 1483341441 | 1456825729 | 1599929345 | 1449602433 | 1517976193 | 1574370177 | 1592926593 | 1567863937 | 1712278529 | 1415732481 | 1862544897 | 1695475073 | 1503721217 | 1615226625 | 1941363969 |
| Q60787 | Lcp2 | Lymphocyte cytosolic protein 2 Mus musculus | 350.12 | 15.95 | 60238 | 7 | 7 | 4554089.5 | 3082293.5 | 3843718.5 | 1894505.88 | 2389610.75 | 3158772.5 | 2183729 | 3602366.75 | 1461835.75 | 2694111.25 | 2554740.25 | 4185799.25 | 3875051 | 3816590.5 | 2983308.75 |

| | | | | | | | | | | | | | | | | | | | | | | |
|---|---|---|---|---|---|---|---|---|---|---|---|---|---|---|---|---|---|---|---|---|---|---|
| P06151 | Ldha | L-lactate dehydrogenase A chain Mus musculus | 2215.19 | 87.05 | 36499 | 28 | 32 | 712083585 | 734669377 | 670777793 | 591029249 | 691721537 | 752190401 | 913085825 | 777360193 | 746621441 | 770667137 | 801903489 | 738947969 | 793248129 | 807796097 | 718574017 |
| P35951 | Ldlr | Low-density lipoprotein receptor Mus musculus | 247.14 | 7.31 | 94947 | 4 | 5 | 503156.88 | 299449.81 | 499312.5 | 751841.75 | 937581.75 | 693843.62 | 622486.94 | 1170847.5 | 1171703.12 | 595550.19 | 1455178.88 | 569894.94 | 913648.94 | 1184749 | 454581.44 |
| Q8C142 | Ldlrap1 | Low density lipoprotein receptor adapter protein 1 Mus musculus | 154.1 | 12.34 | 33975 | 3 | 3 | 5834719.5 | 5932507 | 2432658.75 | 5441983 | 3378121.5 | 3378363.5 | 2663696 | 3545232.75 | 2461191.75 | 4571573.5 | 5508212 | 2611380.5 | 3170693.25 | 6088487 | 2118960.5 |
| Q6DVA0 | Lemd2 | LEM domain-containing protein 2 Mus musculus | 81.54 | 5.09 | 57507 | 2 | 2 | 1213773.5 | 1456737.88 | 1518883.25 | 1223020.12 | 1783008.38 | 1874080.12 | 1507388.12 | 1960973.12 | 2167348.75 | 1208565.5 | 1497947.88 | 1355763.62 | 1941146.75 | 1746860.5 | 1584796.5 |
| Q5XJE5 | Leo1 | RNA polymerase-associated protein LEO1 Mus musculus | 98.94 | 3.45 | 75597 | 2 | 2 | 1584213.25 | 626570.31 | 2074630.12 | 2103116 | 1275064.25 | 754076.75 | 2350653 | 1667391.12 | 1851574.12 | 2568776.5 | 1002574.19 | 1484759.5 | 686192.88 | 764710.25 | 1940066 |
| Q9Z2I0 | Letm1 | Mitochondrial proton/calcium exchanger protein Mus musculus | 228.53 | 15.85 | 82989 | 5 | 5 | 3292706 | 3563616.75 | 3094787.5 | 3588668 | 3040835.5 | 3198759.5 | 3309884.5 | 3057995.75 | 5691692.5 | 3724923 | 4088417 | 3863391.5 | 3183221 | 3399827.25 | 2935341 |
| P16045 | Lgals1 | Galectin-1 Mus musculus | 1008.62 | 92.59 | 14866 | 12 | 14 | 395518401 | 477600097 | 483867105 | 452777025 | 462985857 | 405213185 | 316142049 | 397866241 | 473231105 | 371457825 | 506359361 | 425563297 | 463646049 | 463614785 | 405597025 |
| P16110 | Lgals3 | Galectin-3 Mus musculus | 516.11 | 36.36 | 27515 | 8 | 8 | 66420117 | 61348193 | 74391713 | 82619921 | 79175249 | 64093365 | 96575201 | 49991193 | 48349465 | 86532737 | 91066545 | 87302841 | 79313913 | 59218301 | 112149009 |
| Q07797 | Lgals3bp | Galectin-3-binding protein Mus musculus | 1156.02 | 39.34 | 64491 | 16 | 16 | 84722809 | 85872353 | 84740609 | 60414577 | 75841993 | 71856209 | 53194921 | 77907057 | 77459025 | 56634241 | 70820697 | 70486865 | 85650745 | 88942289 | 62509033 |
| O89017 | Lgmn | Legumain Mus musculus | 339.48 | 14.48 | 49373 | 5 | 5 | 16174174 | 13847101 | 17326819 | 8485222 | 10875123 | 15247561 | 9462480 | 17787485 | 18325745 | 7163404.5 | 15980343 | 15853225 | 9234612 | 18835327 | 15434863 |
| Q5I0D5 | Lhpp | Phospholysine phosphohistidine inorganic pyrophosphate phosphatase Rattus norvegic | 149.53 | 11.85 | 29190 | 2 | 2 | 1550441.62 | 1435424.5 | 365567.44 | 2014820.5 | 1277171.75 | 1502475.5 | 1769603.5 | 2025260.88 | 1379704.62 | 1902545 | 1888473 | 1876137.25 | 1682065.62 | 2016141.62 | 2029006.5 |
| P37913 | Lig1 | DNA ligase 1 Mus musculus | 1179.01 | 34.83 | 102290 | 23 | 23 | 14698591 | 15500032 | 14990569 | 18926393 | 14219613 | 14175713 | 19752621 | 17834565 | 19594231 | 17353997 | 18596265 | 14896536 | 15324205 | 17963829 | 18795285 |
| P97386 | Lig3 | DNA ligase 3 Mus musculus | 132.74 | 3.45 | 113072 | 3 | 3 | 2243806 | 2093747.75 | 2771492.25 | 2356689.5 | 2377674.5 | 2534370.5 | 2337223.25 | 2531481.75 | 2724523.25 | 2071867.5 | 1633149 | 2316506.5 | 2234707.75 | 2515064.75 | 2408449 |
| Q64281 | Lilrb4 | Leukocyte immunoglobulin-like receptor subfamily B member 4 Mus musculus | 272.17 | 22.09 | 37544 | 4 | 4 | 6278929.5 | 6655734.5 | 7452466.5 | 7018348 | 6715694 | 8366322.5 | 4856848.5 | 6735425 | 9364690 | 6715971.5 | 6034132 | 6145590 | 6781667.5 | 6146472 | 6380980 |
| Q4KM31 | Limd2 | LIM domain-containing protein 2 Rattus norvegicus | 61.9 | 13.28 | 14236 | 1 | 1 | 5087711 | 5415087 | 5407738 | 5114712 | 5454524 | 4938950 | 6209732 | 4666135 | 5014229 | 5947500 | 5208784.5 | 5087711 | 6254985 | 4174494 | 4872308.5 |
| Q99JW4 | Lims1 | LIM and senescent cell antigen-like-containing domain protein 1 Mus musculus | 64.65 | 4.62 | 37240 | 1 | 1 | 807918.62 | 615479.19 | 607873.88 | 693736.845 | 743675.69 | 801625.5 | 575379.56 | 693712.69 | 644530.06 | 850770.12 | 803774.19 | 553709.94 | 735566.38 | 815529.5 | 779837.69 |
| Q9Z0M5 | Lipa | Lysosomal acid lipase/cholesteryl ester hydrolase Mus musculus | 683.19 | 22.17 | 45325 | 9 | 9 | 14897863 | 18597939 | 18244205 | 13282737 | 12877767 | 27761769 | 31115139 | 26800337 | 32122465 | 31368847 | 17982701 | 19490887 | 18888273 | 18895353 | 19370957 |
| P54310 | Lipe | Hormone-sensitive lipase Mus musculus | 253.27 | 10.01 | 83348 | 4 | 4 | 1120625 | 1076144.75 | 1234195.38 | 588111.56 | 545402.5 | 1144806 | 1121520.75 | 1118750.5 | 848954.69 | 1142410 | 1249371.5 | 1122087.62 | 779465.12 | 564097.19 | 1645758 |
| Q9D0F3 | Lman1 | Protein ERGIC-53 Mus musculus | 395.53 | 26.5 | 57789 | 8 | 8 | 23873111 | 23560533 | 23336553 | 21160401 | 25816227 | 26915873 | 23712867 | 26165949 | 31886691 | 26150517 | 21671843 | 21498739 | 23594359 | 23723969 | 22130595 |
| Q9DBH5 | Lman2 | Vesicular integral-membrane protein VIP36 Mus musculus | 187.23 | 18.99 | 40430 | 4 | 4 | 3604307 | 3640636.5 | 3198043.5 | 2183465.5 | 2834169.5 | 3741705.25 | 2478054.75 | 3487530.75 | 2948068.25 | 2418696.25 | 2640474 | 3363993 | 2939694 | 2474587.5 | 4081264.5 |
| P48678 | Lmna | Prelamin-A/C Mus musculus | 2521.97 | 56.54 | 74238 | 39 | 42 | 281224001 | 298698145 | 292395201 | 342851937 | 302216961 | 270821345 | 380018401 | 277044801 | 246558609 | 326586177 | 303433889 | 308485121 | 314678337 | 273462913 | 305799169 |
| P14733 | Lmnb1 | Lamin-B1 Mus musculus | 2033.55 | 62.24 | 66786 | 29 | 34 | 87113625 | 89518945 | 92807689 | 108647713 | 102776625 | 96088481 | 109271009 | 86508297 | 83600553 | 104807889 | 112470993 | 105516449 | 93097553 | 81573209 | 110355601 |
| P21619 | Lmnb2 | Lamin-B2 Mus musculus | 713.07 | 25.84 | 67318 | 10 | 15 | 24706799 | 24913981 | 25952291 | 25501765 | 26472293 | 24499965 | 34936837 | 24816461 | 21623161 | 30114117 | 30167031 | 28365905 | 25112213 | 21642931 | 39568461 |
| Q8C129 | Lnpep | Leucyl-cystinyl aminopeptidase Mus musculus | 682.9 | 13.95 | 117304 | 11 | 11 | 12268318 | 9276440 | 12960425 | 9956491 | 8948255 | 10722078 | 14453362 | 13929013 | 17828809 | 10107477 | 17887525 | 17407457 | 8531183 | 16262555 | 13889243 |
| Q8CGK3 | Lonp1 | Lon protease homolog, mitochondrial Mus musculus | 1072.35 | 34.04 | 105843 | 20 | 20 | 28760953 | 28925937 | 29730303 | 31896783 | 30825573 | 31245865 | 40242765 | 34838405 | 34046585 | 40484405 | 30877559 | 34808485 | 33005507 | 30805621 | 38701497 |
| Q3TFD2 | Lpcat1 | Lysophosphatidylcholine acyltransferase 1 Mus musculus | 222.91 | 8.61 | 59744 | 4 | 4 | 1168222 | 1306953.75 | 1139208.25 | 1965804.25 | 1955921.25 | 1083786.5 | 993420.88 | 1529426.25 | 2043105.75 | 840210.31 | 2205769.5 | 1568746.25 | 1500747.25 | 936966 | 802018.31 |
| Q8BYI6 | Lpcat2 | Lysophosphatidylcholine acyltransferase 2 Mus musculus | 480.64 | 21.14 | 60254 | 6 | 10 | 2924251.5 | 3248657.25 | 5680712 | 3541806 | 3749135.5 | 6322366 | 8250237 | 6910561 | 8601355 | 5416486 | 6808583.5 | 3752962.5 | 3571998.75 | 6074669 | 7310826.5 |
| P11152 | Lpl | Lipoprotein lipase Mus musculus | 177.45 | 11.18 | 53109 | 3 | 3 | 308996.12 | 678906.56 | 317656.5 | 1969860 | 818854.795 | 620808.94 | 2749662.25 | 767345.88 | 891031.44 | 2884472.75 | 212032.45 | 802949.25 | 223508.33 | 978973 | 653630.75 |
| Q99N69 | Lpxn | Leupaxin Mus musculus | 542.15 | 26.94 | 43478 | 7 | 7 | 6798333.5 | 7566228.5 | 7889414 | 7975122 | 4547158 | 5996903.5 | 9496647 | 5826585.5 | 4832742 | 6216927 | 9940547 | 7260504 | 6040794.5 | 7053347 | 11085497 |
| Q8BVU0 | Lrch3 | Leucine-rich repeat and calponin homology domain-containing protein 3 Mus musculus | 35.82 | 2.44 | 86341 | 1 | 1 | 1 | 1 | 1 | 1 | 101635.62 | 107581.6 | 97683.2725 | 67982.48 | 99053.8 | 116115.21 | 1 | 1 | 132145.11 | 119021.93 | 99527.57 |
| Q921G6 | Lrch4 | Leucine-rich repeat and calponin homology domain-containing protein 4 Mus musculus | 192.74 | 8.97 | 73163 | 3 | 3 | 1620449 | 1876541.88 | 1666251.5 | 1657480 | 2064907.25 | 1568965.75 | 965997 | 1402861.38 | 1866565.25 | 1355490.38 | 2545334.75 | 1721160.75 | 1838216.5 | 999013.5 | 1882735 |
| Q60664 | Lrmp | Lymphoid-restricted membrane protein Mus musculus | 110.8 | 5.57 | 59588 | 2 | 2 | 2122874.5 | 1019856.94 | 720089.81 | 1669346 | 1802042.38 | 1354095.75 | 1957892.5 | 862300.25 | 1448675.75 | 1506286 | 1262793.25 | 829329.19 | 1061678.88 | 699036.38 | 1744186.25 |
| Q91ZX7 | Lrp1 | Prolow-density lipoprotein receptor-related protein 1 Mus musculus | 3428.96 | 18.13 | 504742 | 59 | 60 | 98539633 | 125103761 | 100120561 | 105283761 | 133641889 | 131665225 | 90854833 | 104063769 | 131390953 | 93237441 | 120847897 | 103781393 | 123592905 | 118607953 | 108138017 |
| Q9JI18 | Lrp1b | Low-density lipoprotein receptor-related protein 1B Mus musculus | 63.99 | 0.37 | 513634 | 1 | 2 | 1296615.25 | 2340284.75 | 1373193 | 1505938.5 | 1582396.62 | 1585748.38 | 977069.94 | 1601982.88 | 1748821.12 | 1159434.5 | 1647113.5 | 1536476.25 | 1644818.38 | 1357106.5 | 1285874.38 |
| P55302 | Lrpap1 | Alpha-2-macroglobulin receptor-associated protein Mus musculus | 371.87 | 25.56 | 42215 | 7 | 7 | 25035591 | 17206813 | 19915209 | 19133793 | 15211021 | 18384359 | 24317251 | 18857117 | 17127353 | 20067993 | 18601155 | 22421001 | 19116123 | 18663137 | 20380763 |
| Q6PB66 | Lrpprc | Leucine-rich PPR motif-containing protein, mitochondrial Mus musculus | 1840.37 | 30.17 | 156615 | 24 | 33 | 51690861 | 44126069 | 48019153 | 43048493 | 46688893 | 48795297 | 43093713 | 48349237 | 54101913 | 41167825 | 44418717 | 48779501 | 45850917 | 49823777 | 49792369 |
| Q8K1T1 | Lrrc25 | Leucine-rich repeat-containing protein 25 Mus musculus | 53.64 | 6.06 | 32673 | 1 | 1 | 654861.69 | 776330.31 | 764671.89 | 721633 | 905862.56 | 616043.38 | 1 | 1 | 1 | 1 | 547019.38 | 1 | 1401490.25 | 1 | 1 |
| Q9CRC8 | Lrrc40 | Leucine-rich repeat-containing protein 40 Mus musculus | 346.08 | 14.78 | 68076 | 7 | 7 | 6767478 | 4537993 | 5081819.5 | 4743589 | 5067823 | 6695875 | 4831228.5 | 5137521 | 5411015 | 5220018 | 5494859.5 | 4474955 | 5784632 | 6571072.5 | 5458749 |
| Q8K1C9 | Lrrc41 | Leucine-rich repeat-containing protein 41 Mus musculus | 136.54 | 7.56 | 88180 | 3 | 3 | 496309.31 | 682914.62 | 789214 | 456527 | 560264.12 | 828169 | 623893 | 737603.12 | 732572.19 | 588050.94 | 902477.12 | 754585.44 | 628512.25 | 865161.62 | 909014 |
| Q505F5 | Lrrc47 | Leucine-rich repeat-containing protein 47 Mus musculus | 188.49 | 11.88 | 63590 | 4 | 4 | 2665587 | 2880447.25 | 2403833.75 | 2991008 | 3062402.5 | 1663320.75 | 2378221 | 2316510.5 | 2246629.75 | 2719847.5 | 3119483.5 | 2201012 | 1437302.25 | 2387049.75 | 2628460.25 |
| Q5RJR8 | Lrrc59 | Leucine-rich repeat-containing protein 59 Rattus norvegicus | 560.67 | 38.44 | 34869 | 10 | 10 | 61149985 | 65145753 | 58488453 | 57033969 | 60603913 | 60562609 | 57170417 | 62258201 | 68720601 | 55715657 | 67155601 | 52864757 | 60232253 | 66278485 | 54208469 |
| Q498T9 | Lrrc8c | Volume-regulated anion channel subunit LRRC8C Rattus norvegicus | 181.54 | 3.86 | 92463 | 2 | 2 | 285118.25 | 180492.56 | 295830.62 | 227927.3775 | 150268.08 | 772115.88 | 275593.34 | 342910.34 | 415650.625 | 271982.94 | 235665.42 | 847373.25 | 373225.41 | 1374425 | 695306.62 |
| Q3UZ39 | Lrrfip1 | Leucine-rich repeat flightless-interacting protein 1 Mus musculus | 1055.96 | 27.43 | 79249 | 16 | 16 | 25806603 | 32912419 | 25704533 | 31371381 | 31016885 | 24962047 | 27270123 | 27349463 | 33053467 | 26947037 | 35492709 | 29659365 | 30361193 | 28945731 | 27595037 |
| Q3UM18 | Lsg1 | Large subunit GTPase 1 homolog Mus musculus | 67.9 | 4.66 | 73157 | 1 | 1 | 676685.31 | 163861.2 | 1 | 1 | 222588.08 | 154766.78 | 213101.69 | 1 | 143683.97 | 1 | 137299.28 | 461602.09 | 232901.4325 | 94343.41 | 238360.95 |
| Q9D0R8 | Lsm12 | Protein LSM12 homolog Mus musculus | 170.35 | 27.69 | 21701 | 3 | 3 | 1417764.38 | 1503190.063 | 1301414.62 | 1666060.5 | 1627520.75 | 1497767.75 | 2477655.5 | 1375910.88 | 1220670.88 | 1895278 | 1542624 | 1658640.88 | 1518284.62 | 1378380.12 | 1955409.75 |
| Q8K2F8 | Lsm14a | Protein LSM14 homolog A Mus musculus | 113.18 | 6.28 | 50546 | 2 | 2 | 507717.91 | 651041.44 | 616954.69 | 1233059.25 | 1107622.38 | 515815 | 742213.38 | 1249375.25 | 382303.84 | 908220.38 | 806417.69 | 679606.38 | 589282.38 | 426634.78 | 932195 |
| O35900 | Lsm2 | U6 snRNA-associated Sm-like protein LSm2 Mus musculus | 229.45 | 60 | 10835 | 4 | 4 | 2667687 | 1314288.12 | 682846.94 | 1468977.62 | 1897637.25 | 2336430 | 1299943.62 | 2123173.75 | 1378127.12 | 1603576.12 | 2898852.75 | 1957306.38 | 1514792.62 | 2816348 | 1466035.5 |
| P62311 | Lsm3 | U6 snRNA-associated Sm-like protein LSm3 Mus musculus | 139.07 | 32.35 | 11845 | 2 | 2 | 3110765 | 3487201.5 | 3687708 | 4149744 | 5675498.5 | 4669788.5 | 5263869 | 2655713 | 2542697.5 | 5126910 | 4488397.5 | 4994771 | 4753767.5 | 2939550.25 | 7358085 |
| Q9QXA5 | Lsm4 | U6 snRNA-associated Sm-like protein LSm4 Mus musculus | 83.69 | 12.41 | 15076 | 2 | 2 | 1314896.5 | 2189684 | 1335778.5 | 1566682.25 | 1819093.5 | 1502105.75 | 1389878.38 | 1476767.5 | 1758030 | 1558347.88 | 1503328.12 | 1329077.38 | 1635586.38 | 2013875.62 | 1471586.5 |
| P62322 | Lsm5 | U6 snRNA-associated Sm-like protein LSm5 Mus musculus | 81.46 | 20.88 | 9937 | 1 | 1 | 3710096.25 | 3849165.75 | 3430881.5 | 4940065.5 | 4260182.5 | 4123623.5 | 3157772.5 | 4201783 | 3559079.25 | 3386109.5 | 3628095.75 | 2972256.5 | 4452601 | 4332595 | 3346046.25 |
| P62313 | Lsm6 | U6 snRNA-associated Sm-like protein LSm6 Mus musculus | 43.15 | 10 | 9128 | 1 | 1 | 3327701 | 2948293.25 | 2895213.25 | 3063587.25 | 2961613 | 2456458.75 | 2321062 | 2609329.25 | 2972809.5 | 2594495 | 2895601 | 836077.5 | 2468198.25 | 2749669.5 | 1986840.5 |
| Q9CQQ8 | Lsm7 | U6 snRNA-associated Sm-like protein LSm7 Mus musculus | 125.58 | 33.01 | 11636 | 3 | 3 | 4030209.75 | 3583573.75 | 2910486.75 | 4427214.5 | 3855994.25 | 3735659.25 | 2688837.5 | 3420580.25 | 2778893.5 | 3008065.5 | 3042774 | 2444941 | 3359525.5 | 3448306.25 | 2409257.5 |
| Q6ZWM4 | Lsm8 | U6 snRNA-associated Sm-like protein LSm8 Mus musculus | 357.2 | 82.29 | 10403 | 5 | 5 | 6014043.5 | 6108049 | 5574844.5 | 6910368 | 7500382 | 7593925.5 | 6100028 | 5961951.5 | 6473045 | 6400047 | 7108621.5 | 6568560.5 | 6788524.5 | 6565878 | 5950896.5 |
| P19973 | Lsp1 | Lymphocyte-specific protein 1 Mus musculus | 677.48 | 42.12 | 36714 | 9 | 9 | 91496321 | 96268945 | 91651585 | 113484225 | 131125113 | 95352073 | 97737697 | 87151985 | 79754553 | 106197057 | 119629729 | 106418313 | 112033569 | 85546257 | 110875337 |
| Q8BLN5 | Lss | Lanosterol synthase Mus musculus | 321.46 | 8.59 | 83141 | 2 | 5 | 5062840 | 5013338 | 4277994 | 4248439 | 3634731.75 | 4813564 | 4106688.25 | 4263366 | 4383103 | 4426795 | 4633614 | 4301500 | 3047628.5 | 3825047 | 6018478 |
| P24527 | Lta4h | Leukotriene A-4 hydrolase Mus musculus | 476.73 | 19.15 | 69051 | 9 | 9 | 11273067 | 12678380 | 11299983 | 12328483 | 8893505 | 14190301 | 11719074 | 12639149 | 13363240 | 12204603 | 14242503 | 12547616 | 12097333 | 14218033 | 12877881 |
| Q6A009 | Ltn1 | E3 ubiquitin-protein ligase listerin Mus musculus | 141.11 | 1.08 | 198921 | 2 | 2 | 173121.45 | 418949.69 | 139976.73 | 341836.88 | 295224.5 | 222691.17 | 289265.34 | 1 | 1 | 462804.16 | 122668.54 | 93998.7 | 398643.56 | 90088.16 | 68093.45 |
| Q68FR7 | Ltv1 | Protein LTV1 homolog Rattus norvegicus | 48.27 | 4.04 | 53562 | 1 | 1 | 1 | 1 | 1 | 483846.47 | 1 | 736129.25 | 1 | 1 | 1 | 1 | 651492.88 | 1 | 1 | 1 | 635842.94 |
| Q9CYI4 | Luc7l | Putative RNA-binding protein Luc7-like 1 Mus musculus | 229.78 | 15.36 | 43934 | 3 | 6 | 10329964 | 8477223 | 9781321 | 10140205 | 10196263 | 10342648 | 10979835 | 10596477 | 9499153 | 10606157 | 12706463 | 10599553 | 9801411 | 11600265 | 10492187 |
| Q7TNC4 | Luc7l2 | Putative RNA-binding protein Luc7-like 2 Mus musculus | 395.75 | 20.15 | 46583 | 5 | 8 | 23241649 | 21309971 | 22017795 | 25316113 | 23186323 | 22341151 | 24637431 | 22769285 | 21351455 | 23420365 | 26719651 | 22633501 | 21928989 | 25433357 | 23285625 |
| Q5SUF2 | Luc7l3 | Luc7-like protein 3 Mus musculus | 640.75 | 26.39 | 51450 | 9 | 10 | 9580275 | 9362646 | 12781082 | 11444175 | 8638739 | 4937541.5 | 14347786 | 8059469.5 | 4103360.75 | 10753866 | 12834958 | 11138591 | 10079067 | 5770129.5 | 12664612 |
| P70202 | Lxn | Latexin Mus musculus | 92.6 | 9.46 | 25492 | 2 | 2 | 4335441.5 | 5021072 | 4927896 | 4757538.5 | 4461408.5 | 3962391.5 | 2843572.75 | 4782727 | 5097369.5 | 3215122 | 4308377.5 | 3982099.25 | 3797591 | 4557076 | 3241313.25 |
| O88188 | Ly86 | Lymphocyte antigen 86 Mus musculus | 309.35 | 43.83 | 17811 | 4 | 4 | 5974965.5 | 6362964.5 | 5429174.5 | 3877232.75 | 5239046 | 4882296 | 3042052.75 | 5188622 | 5083450.5 | 3376694 | 2768598.5 | 3170351.75 | 5883196 | 4471177 | 3075815.5 |
| Q08288 | Lyar | Cell growth-regulating nucleolar protein Mus musculus | 79.37 | 3.61 | 43736 | 1 | 1 | 84915.59 | 120043.96 | 108362.73 | 126396.12 | 193602.12 | 94887.19 | 183297.22 | 153773.86 | 185671.25 | 92254.12 | 151694.23 | 115081.44 | 237334.23 | 152822.61 | 140310.89 |
| P25911 | Lyn | Tyrosine-protein kinase Lyn Mus musculus | 639.8 | 34.18 | 58812 | 12 | 12 | 19735451 | 21564789 | 19844963 | 14761444 | 15744967 | 18977243 | 15757230 | 17581703 | 19196379 | 17499357 | 19015887 | 20999477 | 18140959 | 18221981 | 18564133 |
| P97823 | Lypla1 | Acyl-protein thioesterase 1 Mus musculus | 226.94 | 26.52 | 24688 | 4 | 4 | 4853141 | 6326202.5 | 5610991.5 | 5826329 | 6182496 | 4488624.5 | 5833022 | 5198583.5 | 4552704.5 | 5999974.5 | 3847012 | 5343115 | 5657525.5 | 5751681.5 | 5267131 |
| Q9QYL8 | Lypla2 | Acyl-protein thioesterase 2 Rattus norvegicus | 40.87 | 10.82 | 24807 | 1 | 1 | 875456.56 | 942451.81 | 740642.75 | 577799.25 | 760094.81 | 957185.1575 | 1123751.25 | 1057006.75 | 944334.38 | 703648.25 | 928682.81 | 945914.44 | 1022898.44 | 880852.88 | 1012071 |
| P08905 | Lyz2 | Lysozyme C-2 Mus musculus | 403.81 | 49.32 | 16689 | 4 | 4 | 23830993 | 30157265 | 25073081 | 23394253 | 22318589 | 17470505 | 13516613 | 18438205 | 21674717 | 15363785 | 16017241 | 18982533 | 25560087 | 23386109 | 17933897 |
| P24668 | M6pr | Cation-dependent mannose-6-phosphate receptor Mus musculus | 423.18 | 32.37 | 31172 | 6 | 6 | 27936077 | 31874237 | 26059501 | 30118059 | 28856205 | 31352989 | 28344109 | 31856833 | 31613069 | 28279981 | 24333985 | 28129995 | 30647217 | 27031043 | 21025123 |
| Q9QXZ0 | Macf1 | Microtubule-actin cross-linking factor 1 Mus musculus | 1182.11 | 4.98 | 831878 | 26 | 27 | 6852718.5 | 8346064 | 8966120 | 7128024 | 6447014 | 10441765 | 8321477.5 | 9335056 | 10645233 | 8126683.5 | 8852429 | 9248541 | 6501426.5 | 7553945 | 9086503 |
| Q9Z1B5 | Mad2l1 | Mitotic spindle assembly checkpoint protein MAD2A Mus musculus | 341.8 | 37.56 | 23598 | 6 | 6 | 3833004.5 | 7963356 | 6602414 | 5019572.5 | 4244114.5 | 3856596 | 4480660 | 7155411 | 6776612 | 4166848 | 4513406 | 3578923.25 | 4662156.5 | 7070666 | 4750830.5 |
| Q80U28 | Madd | MAP kinase-activating death domain protein Mus musculus | 55.4 | 1.52 | 175180 | 1 | 1 | 351095.78 | 223346.52 | 1 | 1 | 187522.52 | 1 | 203251.45 | 1 | 143324.2 | 246358.84 | 365296.38 | 206720.75 | 260997.18 | 233663.67 | 238307.92 |
| P61327 | Magoh | Protein mago nashi homolog Mus musculus | 278.44 | 56.85 | 17164 | 6 | 6 | 5254852 | 2559180.25 | 2357303.5 | 2722990 | 3714786.5 | 2911401.5 | 3099627.5 | 4123094 | 5735171.5 | 3579199.5 | 2871302.5 | 4228562 | 2997796.75 | 4619422.5 | 4629659 |
| Q6AY04 | Maip1 | m-AAA protease-interacting protein 1, mitochondrial Rattus norvegicus | 91.79 | 8.59 | 33090 | 2 | 2 | 1 | 1 | 1380012.25 | 1500286.62 | 1859372.12 | 1177076.62 | 1243934.75 | 1296111.12 | 1170632.88 | 1612460.88 | 404690.81 | 1265788 | 1373282.62 | 1030137.19 | 1405724.12 |
| Q8BGS0 | Mak16 | Protein MAK16 homolog Mus musculus | 150.13 | 11.49 | 35185 | 3 | 3 | 1338627.62 | 1080735.5 | 1124333.88 | 1422618.5 | 1444412.88 | 1871758.75 | 1235420.88 | 1037698 | 1022798.38 | 995313.62 | 2002784.5 | 1756488.88 | 1778153.38 | 801522.38 | 1563788.75 |
| P27046 | Man2a1 | Alpha-mannosidase 2 Mus musculus | 74.78 | 2.96 | 131631 | 2 | 2 | 245755.61 | 444731.75 | 384437.62 | 267502.25 | 468527.28 | 383370.44 | 180055.95 | 422658.38 | 348174.31 | 364240.38 | 417303.38 | 419317.78 | 420751.94 | 132917.03 | 803808.56 |
| O09159 | Man2b1 | Lysosomal alpha-mannosidase Mus musculus | 636.26 | 14.61 | 114648 | 10 | 10 | 29760767 | 29134727 | 24821189 | 32504589 | 31066853 | 23837261 | 17736253 | 27431651 | 22412547 | 18264985 | 23631789 | 21328147 | 27183449 | 28367575 | 20803459 |
| O54782 | Man2b2 | Epididymis-specific alpha-mannosidase Mus musculus | 370.99 | 8.74 | 115610 | 6 | 6 | 6902012 | 8631566 | 8239087.5 | 8540969 | 7448676 | 7527293 | 7016130 | 8440931 | 8982213 | 7580344 | 7163155 | 8005399.5 | 7525883 | 8054745 | 8100779 |
| Q91W89 | Man2c1 | Alpha-mannosidase 2C1 Mus musculus | 310.28 | 6.93 | 115688 | 5 | 5 | 3701658.25 | 4677847 | 3995086.5 | 4081363.25 | 4436063 | 4874156.5 | 5025830 | 4367494 | 5233921 | 4216562 | 5097562 | 3336773 | 4301826.5 | 4618849.5 | 5153943.5 |
| Q8K2I4 | Manba | Beta-mannosidase Mus musculus | 358.22 | 8.99 | 100831 | 6 | 6 | 9049945 | 13534204 | 10444386 | 8421119 | 12710343 | 9371352 | 5955927.5 | 11636815 | 14556287 | 7898340.5 | 12307220 | 7599927.5 | 11703797 | 12665702 | 7650581 |
| P0C5H9 | Manf | Mesencephalic astrocyte-derived neurotrophic factor Rattus norvegicus | 263.78 | 21.23 | 20388 | 5 | 5 | 13466765 | 12135204 | 10767937 | 10744876 | 10496057 | 9700052 | 6837870 | 9948804 | 10329289 | 8493842 | 8759529 | 8539654 | 8978007 | 9614908 | 6948340 |
| P21396 | Maoa | Amine oxidase [flavin-containing] A Rattus norvegicus | 54.51 | 3.8 | 59508 | 1 | 1 | 241152.31 | 184421.59 | 172652.61 | 1 | 1 | 216185.11 | 1 | 236086.14 | 158976.73 | 1 | 235672.33 | 179693.86 | 223936.5 | 209716.8175 | 199564.58 |
| Q8C052 | Map1s | Microtubule-associated protein 1S Mus musculus | 322.18 | 8.74 | 102939 | 7 | 7 | 3955355.25 | 2762875.25 | 4707181 | 4585850.5 | 5361405.5 | 2606583 | 4223812 | 2702834.25 | 2646373.5 | 5154245 | 3620877.25 | 3360472.5 | 3240853.75 | 3247316.25 | 3752345.75 |
| P31938 | Map2k1 | Dual specificity mitogen-activated protein kinase kinase 1 Mus musculus | 779.04 | 53.18 | 43474 | 10 | 14 | 19489445 | 34046405 | 26518319 | 17445085 | 18984441 | 24755697 | 35263649 | 35652461 | 36735221 | 19353849 | 28237287 | 35084861 | 32213949 | 33160409 | 32380597 |
| Q63932 | Map2k2 | Dual specificity mitogen-activated protein kinase kinase 2 Mus musculus | 544.34 | 33.92 | 44402 | 6 | 11 | 14525667 | 18130511 | 18250865 | 9743858 | 14086204 | 20031855 | 16930211 | 21343161 | 21209413 | 11911940 | 22156575 | 18961217 | 16341217 | 19477371 | 17907089 |
| O09110 | Map2k3 | Dual specificity mitogen-activated protein kinase kinase 3 Mus musculus | 181.02 | 14.7 | 39296 | 3 | 3 | 3462949.75 | 3640004.5 | 5152590 | 4161518.5 | 3491682 | 4699073.5 | 3566088 | 4918088.5 | 5088987 | 4503666 | 4200446 | 3913613.5 | 3884897.5 | 3936264 | 4374870 |
| P47809 | Map2k4 | Dual specificity mitogen-activated protein kinase kinase 4 Mus musculus | 37.09 | 4.28 | 44114 | 1 | 1 | 1 | 1 | 1 | 132612.8 | 1 | 96134.67 | 1 | 1 | 1 | 1 | 107762.04 | 125785.77 | 1 | 126635.97 | 1 |
| Q9ESL4 | Map3k20 | Mitogen-activated protein kinase kinase kinase 20 Mus musculus | 282.33 | 7.48 | 91720 | 5 | 5 | 9457455 | 9591159 | 8883040 | 8947703 | 9302907 | 10555577 | 10188621 | 10224819 | 9568499 | 9639521 | 8029131.5 | 8647950 | 8369205.5 | 8882327 | 8455357 |
| P27546 | Map4 | Microtubule-associated protein 4 Mus musculus | 897.26 | 23.91 | 117429 | 19 | 19 | 24230473 | 27092593 | 23207729 | 27445529 | 28212669 | 29225759 | 29463713 | 29460409 | 26927997 | 31233855 | 28264557 | 30898617 | 27405479 | 26615099 | 29703025 |
| P70218 | Map4k1 | Mitogen-activated protein kinase kinase kinase kinase 1 Mus musculus | 156.07 | 5.44 | 91535 | 3 | 3 | 354441.78 | 1 | 330072.12 | 1 | 1 | 238257.89 | 247555.53 | 1 | 1 | 1 | 370741 | 1 | 226848.86 | 262881.91 | 1 |
| P97820 | Map4k4 | Mitogen-activated protein kinase kinase kinase kinase 4 Mus musculus | 71.94 | 2.68 | 140602 | 1 | 1 | 1 | 1 | 1 | 1 | 1 | 1 | 129742.57 | 117059.59 | 1 | 1 | 1 | 147775.83 | 1 | 1 | 1 |
| A2AJI0 | Map7d1 | MAP7 domain-containing protein 1 Mus musculus | 112.83 | 5.79 | 93276 | 3 | 3 | 6545858 | 7890116 | 2292030.5 | 6643053.5 | 6494837.5 | 5762882.5 | 2126584 | 5326539.5 | 4907235 | 2341340.5 | 2827592.25 | 4182535 | 6241000.5 | 6769984.5 | 2738138.5 |
| P63085 | Mapk1 | Mitogen-activated protein kinase 1 Mus musculus | 341.41 | 25.7 | 41276 | 4 | 7 | 18330447 | 19120885 | 17701097 | 17579577 | 20027901 | 18795445 | 22135505 | 14682778 | 18394687 | 21483125 | 22976027 | 19604241 | 19391817 | 18676875 | 20956637 |
| P47811 | Mapk14 | Mitogen-activated protein kinase 14 Mus musculus | 583.22 | 39.17 | 41287 | 11 | 11 | 13121958 | 13253919 | 12043329 | 12279388 | 13826778 | 13503179 | 10447452 | 13629205 | 13391221 | 10692225 | 14125568 | 12686770 | 12731494 | 13605951 | 13122195 |
| Q8BH93 | Mapk1ip1l | MAPK-interacting and spindle-stabilizing protein-like Mus musculus PE | 36.11 | 10.33 | 23885 | 1 | 1 | 1 | 1 | 1522731.12 | 1 | 1938282.88 | 1566761 | 1231215.25 | 1426725.88 | 1686760.62 | 1440120.62 | 1259871.88 | 1281799.38 | 1898361.5 | 1416176.535 | 1224673.38 |
| P21708 | Mapk3 | Mitogen-activated protein kinase 3 Rattus norvegicus | 285.39 | 25.53 | 43081 | 3 | 6 | 14046033 | 13884697 | 13244656 | 12927679 | 14114616 | 14595198 | 18821231 | 14158102 | 13997421 | 16445258 | 18213003 | 14374974 | 15016812 | 13967979 | 16860421 |
| P49186 | Mapk9 | Mitogen-activated protein kinase 9 Rattus norvegicus | 83.81 | 8.04 | 48017 | 2 | 2 | 570272.56 | 933954.75 | 656750.62 | 349013.44 | 688350.19 | 1029000.19 | 1085672 | 736413.5 | 342536.03 | 843902.12 | 832114 | 776053.25 | 843688.88 | 733105.12 | 968941.25 |
| P49138 | Mapkapk2 | MAP kinase-activated protein kinase 2 Mus musculus | 193.8 | 11.66 | 44050 | 5 | 5 | 5646235 | 7711157 | 6409378 | 4354192.5 | 5777333.5 | 8800377 | 6391827 | 5932323 | 5144389 | 5818148.5 | 5760440.5 | 5108907 | 5220928 | 4651759 | 5553625 |
| Q61166 | Mapre1 | Microtubule-associated protein RP/EB family member 1 Mus musculus | 569.78 | 41.42 | 30016 | 9 | 9 | 16125955 | 14248805 | 17313273 | 15565018 | 14982298 | 16276526 | 19021509 | 15987542 | 14104061 | 17398791 | 16833921 | 19591521 | 14102004 | 14088633 | 20052059 |
| Q3B8Q0 | Mapre2 | Microtubule-associated protein RP/EB family member 2 Rattus norvegicus | 65.24 | 3.68 | 36988 | 1 | 1 | 494036.25 | 466571.81 | 1 | 1 | 1 | 629652.31 | 734220.25 | 1 | 1 | 226570.38 | 381266.59 | 700935.81 | 1 | 1 | 1 |
| Q922Q1 | Marc2 | Mitochondrial amidoxime reducing component 2 Mus musculus | 363.39 | 26.04 | 38194 | 3 | 8 | 7982832 | 7891377.5 | 7739494 | 9074475 | 10240457 | 11299854 | 11273633 | 9314421 | 10555654 | 10828539 | 11373523 | 8398520 | 9625687 | 5908090 | 12796351 |
| Q3KNM2 | March5 | E3 ubiquitin-protein ligase MARCH5 Mus musculus | 89.65 | 13.67 | 31232 | 3 | 3 | 209912.09 | 1351820.25 | 220500.03 | 316599.56 | 313799.12 | 1654514.25 | 414318.12 | 1423890.12 | 94616.84 | 393573.12 | 1803902.5 | 162175.61 | 335661.53 | 1834112 | 2601277.75 |
| P26645 | Marcks | Myristoylated alanine-rich C-kinase substrate Mus musculus | 463.97 | 34.63 | 29661 | 7 | 7 | 12324777 | 14816291 | 15684935 | 17136421 | 13898977 | 13251401 | 15585615 | 12905209 | 12067596 | 15032479 | 17262151 | 14308515 | 16701567 | 11746071 | 16029996 |
| P28667 | Marcksl1 | MARCKS-related protein Mus musculus | 205.08 | 34.5 | 20165 | 4 | 4 | 9196065 | 8834829 | 8280734.5 | 8980619 | 9934519 | 9885404 | 10909023 | 9531033 | 10259880 | 10419569 | 10909821 | 8958838 | 11949703 | 9352347 | 9023325 |
| Q68FL6 | Mars | Methionine--tRNA ligase, cytoplasmic Mus musculus | 1471.44 | 44.79 | 101431 | 28 | 28 | 33976821 | 34399789 | 34109697 | 36394849 | 35676701 | 34540673 | 39304693 | 33299379 | 45140517 | 34955665 | 45581285 | 39532729 | 33952749 | 49106417 | 40426305 |

| | | | | | | | | | | | | | | | | | | | | | | |
|---|---|---|---|---|---|---|---|---|---|---|---|---|---|---|---|---|---|---|---|---|---|---|
| Q3THS6 | Mat2a | S-adenosylmethionine synthase isoform type-2 Mus musculus | 436.56 | 25.06 | 43689 | 2 | 9 | 25768187 | 26025637 | 27081829 | 21670393 | 25990649 | 22923229 | 18715961 | 21935389 | 22757025 | 22994443 | 27119191 | 25795261 | 24792169 | 22357549 | 23757319 |
| Q5U2R0 | Mat2b | Methionine adenosyltransferase 2 subunit beta Rattus norvegicus | 217.94 | 20.66 | 37375 | 4 | 4 | 7180173.5 | 7486520 | 7835277.5 | 8525633 | 7883800.5 | 6610390.5 | 8010135.5 | 6433476 | 7441368.5 | 8134600.5 | 7547166 | 8461697 | 8604052 | 8399129 | 7352428 |
| Q8K310 | Matr3 | Matrin-3 Mus musculus | 768.83 | 23.4 | 94630 | 13 | 13 | 13998072 | 13357893 | 12686952 | 13257240 | 14324118 | 12190444 | 16045667 | 13828481 | 13780901 | 14444488 | 12504033 | 12692393 | 12517168 | 12534885 | 12869282 |
| Q9D2X5 | Mau2 | MAU2 chromatid cohesion factor homolog Mus musculus | 45.49 | 1.45 | 69579 | 1 | 1 | 741195.69 | 486170.12 | 475540.56 | 651269.19 | 563693.38 | 515148.78 | 290366.75 | 583169.81 | 482183.38 | 396249.03 | 378729.22 | 500849.69 | 440087.09 | 498872.09 | 400296.31 |
| Q8VCF0 | Mavs | Mitochondrial antiviral-signaling protein Mus musculus | 442.01 | 21.27 | 53399 | 7 | 7 | 6068584 | 6987083 | 7016185.5 | 7826510.5 | 7408904.5 | 7279218.5 | 9546327 | 6432463 | 6107639 | 9448465 | 7774293.5 | 7189405 | 8505017 | 5776088 | 6965839.5 |
| Q9Z2D8 | Mbd3 | Methyl-CpG-binding domain protein 3 Mus musculus | 273.07 | 16.84 | 32168 | 5 | 5 | 4428769 | 4878586 | 4117598.5 | 4551651.5 | 4654179 | 5030441 | 4463523.5 | 4713647.5 | 4981041 | 4617727 | 4773880.5 | 4009404.25 | 4306938.5 | 4662281 | 4163779 |
| Q9JKP5 | Mbnl1 | Muscleblind-like protein 1 Mus musculus | 210.35 | 21.99 | 36976 | 4 | 4 | 7373277 | 7182238 | 8920765 | 10352111 | 7573455 | 7303777 | 9941333 | 5609857 | 5581010 | 8807922 | 8955787 | 7408966 | 8269470 | 6650192 | 8950141 |
| Q8CHK3 | Mboat7 | Lysophospholipid acyltransferase 7 Mus musculus | 112.25 | 7.19 | 53436 | 2 | 2 | 1874685.25 | 1911352.25 | 1763730.75 | 1283265.38 | 1490587.75 | 2119007.25 | 1784092.25 | 2117795 | 2197439.75 | 1577961.25 | 2136073.5 | 1788512.75 | 1498588.75 | 2159222.25 | 2423544.25 |
| Q8R3F5 | Mcat | Malonyl-CoA-acyl carrier protein transacylase, mitochondrial Mus musculus | 70.75 | 3.15 | 41928 | 1 | 1 | 263039.78 | 643665.5 | 685530.56 | 664288.5 | 560193.69 | 641982.19 | 537005.19 | 746076.94 | 295090.34 | 463961.97 | 722373.31 | 556132.19 | 671250.56 | 656426.94 | 696316.69 |
| Q99MR8 | Mccc1 | Methylcrotonoyl-CoA carboxylase subunit alpha, mitochondrial Mus musculus | 38.88 | 1.12 | 79344 | 1 | 1 | 1006580.12 | 1059265.62 | 795484.06 | 1220062.38 | 1020348.045 | 808319.88 | 968058.12 | 831676.88 | 1043036.19 | 1087660.88 | 675009.5 | 962002 | 907706.94 | 872648.56 | 874487.75 |
| P97287 | Mcl1 | Induced myeloid leukemia cell differentiation protein Mcl-1 homolog Mus musculus | 133.72 | 10.27 | 35217 | 2 | 2 | 1736595.25 | 1185432.5 | 784035.94 | 1118408.38 | 1162852.75 | 1286865.12 | 1024680.06 | 1259993.62 | 894616.69 | 865470.5 | 1175052.75 | 1107212.88 | 1334884.5 | 1352414.12 | 1234958.88 |
| P97310 | Mcm2 | DNA replication licensing factor MCM2 Mus musculus | 1917.27 | 46.57 | 102078 | 31 | 31 | 51710697 | 47636637 | 44773761 | 45241849 | 44152637 | 41239689 | 47016289 | 41910509 | 38273897 | 45575573 | 42672317 | 47244137 | 45432245 | 43149689 | 45808573 |
| P25206 | Mcm3 | DNA replication licensing factor MCM3 Mus musculus | 2118.5 | 59.11 | 91546 | 35 | 35 | 71992825 | 70460033 | 64243505 | 77191561 | 76612785 | 63882489 | 72516785 | 64645861 | 60041437 | 72097969 | 69825841 | 61873005 | 76039449 | 67244313 | 71158849 |
| P49717 | Mcm4 | DNA replication licensing factor MCM4 Mus musculus | 1408.25 | 42.58 | 96736 | 24 | 25 | 22723659 | 21678979 | 24885785 | 23017115 | 20432631 | 20134069 | 26444739 | 22435757 | 18286059 | 24864963 | 20701159 | 27618729 | 21847619 | 19988679 | 27590979 |
| P49718 | Mcm5 | DNA replication licensing factor MCM5 Mus musculus | 1617.39 | 52.8 | 82343 | 29 | 30 | 45386225 | 52188905 | 47732161 | 51570645 | 48091649 | 38566941 | 48509325 | 43889333 | 43948809 | 52127081 | 47361213 | 56365357 | 50431813 | 44853341 | 52264749 |
| P97311 | Mcm6 | DNA replication licensing factor MCM6 Mus musculus | 1859.34 | 44.7 | 92867 | 32 | 32 | 40374621 | 38405533 | 37149001 | 44161857 | 42873825 | 38194229 | 33903773 | 40212669 | 41255377 | 39472101 | 47902229 | 41409045 | 34854225 | 40874269 | 46066469 |
| Q61881 | Mcm7 | DNA replication licensing factor MCM7 Mus musculus | 1286.23 | 44.37 | 81211 | 22 | 22 | 33899465 | 33035453 | 32675285 | 34860397 | 31904891 | 30800413 | 33701697 | 32243081 | 30027863 | 31454177 | 30305339 | 33345529 | 32425317 | 30066031 | 28753209 |
| Q8R3C0 | Mcmbp | Mini-chromosome maintenance complex-binding protein Mus musculus | 462.89 | 16.98 | 72891 | 8 | 8 | 4546606.5 | 6107745 | 5097431.5 | 7207752.5 | 5396642 | 5107400.5 | 6610795.5 | 5932968.5 | 5866298 | 5785574 | 7564571 | 5714445 | 5006997 | 6173168 | 5693410 |
| Q4G009 | Mcts1 | Malignant T-cell-amplified sequence 1 Rattus norvegicus | 140.67 | 21.43 | 20550 | 2 | 2 | 1085782 | 917313.94 | 2002678.5 | 2162327.5 | 2437907.25 | 2194036 | 1912864.75 | 2456157 | 1957547.5 | 1957741.75 | 1588543.12 | 1772070.12 | 1851604.25 | 2102364.5 | 1622186.62 |
| Q3UMR5 | Mcu | Calcium uniporter protein, mitochondrial Mus musculus | 55.29 | 2.86 | 39682 | 1 | 1 | 1657174.38 | 1464395 | 2021857.12 | 1750472.5 | 1484662.75 | 1388039.25 | 5483325.5 | 1323816.25 | 1561929.12 | 3400979.5 | 2097781 | 658677.62 | 1553278.5 | 1132349.5 | 655920.88 |
| Q5PSV9 | Mdc1 | Mediator of DNA damage checkpoint protein 1 Mus musculus | 144.43 | 3.4 | 184670 | 3 | 3 | 1 | 1665963.38 | 1 | 1 | 2225106.5 | 1910796.5 | 2069446.88 | 1856462.88 | 1882883.12 | 1929897.345 | 2735882.5 | 1806760 | 1923563.25 | 2556360 | 1835662.88 |
| P14152 | Mdh1 | Malate dehydrogenase, cytoplasmic Mus musculus | 893.88 | 62.28 | 36511 | 15 | 15 | 66041913 | 69056273 | 74115449 | 65314393 | 67860985 | 74122481 | 103824353 | 73630985 | 69764073 | 67989105 | 93644617 | 91153769 | 62989953 | 64318209 | 91381633 |
| P08249 | Mdh2 | Malate dehydrogenase, mitochondrial Mus musculus | 1155.56 | 63.02 | 35611 | 17 | 18 | 362337793 | 384085889 | 367957345 | 372896545 | 337987393 | 334660097 | 346496481 | 392144545 | 403736833 | 332904993 | 336973505 | 303844001 | 353237185 | 410576417 | 283495201 |
| Q9D967 | Mdp1 | Magnesium-dependent phosphatase 1 Mus musculus | 152.52 | 21.95 | 18582 | 3 | 3 | 1781096.62 | 1341321.88 | 991375.06 | 1564949.12 | 1129446.88 | 1080433.88 | 2101787.5 | 2108498.5 | 1558929.88 | 1408847.38 | 1188682.88 | 1353881.12 | 2533164 | 1837832.62 | 850866.56 |
| P06801 | Me1 | NADP-dependent malic enzyme Mus musculus | 247.32 | 15.03 | 63954 | 4 | 4 | 589037.31 | 1014610 | 448303.91 | 355041.78 | 263235.97 | 620213.75 | 1471164.12 | 813567.38 | 429723.19 | 307183.78 | 717052.19 | 349280 | 377819.69 | 643062.75 | 1029452.56 |
| Q99KE1 | Me2 | NAD-dependent malic enzyme, mitochondrial Mus musculus | 842.78 | 47.88 | 65799 | 16 | 16 | 6508553 | 10857120 | 9220604 | 8049399 | 11346127 | 14347749 | 13381621 | 10554337 | 7233176 | 13447375 | 10335303 | 10858512 | 13479356 | 8707930 | 10047550 |
| Q9DCS3 | Mecr | Enoyl-[acyl-carrier-protein] reductase, mitochondrial Mus musculus | 190.64 | 23.59 | 40343 | 3 | 3 | 200994.44 | 270153.69 | 361610.19 | 359813.31 | 298142.9075 | 358415.09 | 1234498.5 | 264972.34 | 181679.12 | 1584490.75 | 607389.19 | 478681.69 | 329424.97 | 441094.78 | 1150039.75 |
| Q9CQ39 | Med21 | Mediator of RNA polymerase II transcription subunit 21 Mus musculus | 43.56 | 13.89 | 15588 | 1 | 1 | 146402.66 | 1 | 197672.39 | 1 | 190558.34 | 1 | 1 | 1 | 184221.69 | 213335.39 | 228802.25 | 161102.7 | 178865.42 | 1 | 1 |
| Q4QQR9 | Memo1 | Protein MEMO1 Rattus norvegicus | 200.91 | 18.18 | 33679 | 4 | 4 | 2324965.25 | 3121353.75 | 2687123.5 | 2386020.5 | 2087837.75 | 2145313.25 | 2557009.25 | 2071562.5 | 2349659.5 | 2423985.75 | 2741985.5 | 3616283.25 | 2242895.5 | 2551637 | 2849168.25 |
| O88559 | Men1 | Menin Mus musculus | 33.31 | 2.29 | 67501 | 1 | 1 | 36176.1 | 378646.75 | 1 | 1 | 1 | 63135.54 | 33572.28 | 90057.27 | 1 | 1 | 56412.84 | 59713.78 | 1 | 1 | 1 |
| Q9ERE7 | Mesd | LRP chaperone MESD Mus musculus | 345.63 | 37.5 | 25207 | 6 | 6 | 5726722.5 | 7060420 | 5772009 | 6588132.5 | 5376967 | 5685062 | 7328977.5 | 5816082 | 5125051 | 6661092 | 4396081 | 4654013.5 | 6014859.5 | 4511861 | 4771166 |
| Q8BP48 | Metap1 | Methionine aminopeptidase 1 Mus musculus | 271.74 | 17.1 | 43221 | 5 | 5 | 6111351 | 6590463 | 6052901 | 6348026 | 6844733.5 | 7734556 | 7960192 | 6672260.5 | 6492444 | 9490515 | 6599505 | 3907432.5 | 7658152 | 7004021.5 | 6814766 |
| O08663 | Metap2 | Methionine aminopeptidase 2 Mus musculus | 500.59 | 32.01 | 52922 | 10 | 10 | 3759914.25 | 7570577.5 | 4777828 | 7124577 | 2888116.25 | 4925495 | 5168707 | 5685972 | 10475768 | 2980386.5 | 7342398 | 4162866 | 4168419 | 4340205.5 | 5608252 |
| Q9Z120 | Mettl1 | tRNA (guanine-N(7)-)-methyltransferase Mus musculus | 67.78 | 5.6 | 30603 | 1 | 1 | 251424.14 | 278458.44 | 217846.14 | 404747.72 | 347269.28 | 246294.27 | 280955.31 | 195280.88 | 253386.12 | 243016.19 | 224517.19 | 271732.84 | 332107.72 | 225146.83 | 272056.31 |
| Q91YR5 | Mettl13 | Methyltransferase-like protein 13 Mus musculus | 177.6 | 5.73 | 78757 | 4 | 4 | 2987178 | 2296236.25 | 1540658.75 | 2430628.5 | 2493596.75 | 2108257.5 | 2355615.5 | 2088975.62 | 1762051.62 | 2290956.25 | 1443906.25 | 1469632 | 1957274 | 2345634.25 | 784545.25 |
| Q9CQG2 | Mettl16 | U6 small nuclear RNA (adenine-(43)-N(6))-methyltransferase Mus musculus | 192.29 | 9.22 | 62341 | 4 | 4 | 31932109 | 2204096.5 | 2618429.75 | 24724479 | 35056745 | 20351865 | 2890761.5 | 21059399 | 1611873.75 | 1918542.12 | 3487777.5 | 2133760 | 2324965.75 | 18818737 | 2153133.25 |
| Q8BMK1 | Mettl2 | Methyltransferase-like protein 2 Mus musculus 2 | 37.87 | 5.14 | 43912 | 1 | 1 | 354146.12 | 277974.16 | 1 | 1 | 396939 | 271496.81 | 1 | 1 | 1 | 1 | 1 | 1 | 276989.19 | 1 | 309382.31 |
| Q9DCS2 | Mettl26 | Methyltransferase-like 26 Mus musculus | 98.55 | 8.82 | 22687 | 1 | 1 | 385797.88 | 276300.4525 | 205587.38 | 275702.25 | 238114.3 | 398930.5 | 316907.19 | 469006.5 | 305341.56 | 1 | 416980.22 | 476534.72 | 291105.16 | 381253.78 | 352190.41 |
| P21956 | Mfge8 | Lactadherin Mus musculus | 401.72 | 23.11 | 51241 | 8 | 8 | 25970557 | 26130477 | 26266839 | 20592657 | 23680475 | 22919561 | 16553943 | 22179799 | 24858511 | 21689687 | 21856813 | 21454067 | 22331245 | 20124009 | 21959535 |
| Q811U4 | Mfn1 | Mitofusin-1 Mus musculus | 55.91 | 1.89 | 83726 | 1 | 1 | 438864.47 | 544852.44 | 673544.94 | 489100.62 | 551190.5 | 686973.5 | 744649.38 | 669053.12 | 767150.31 | 793130.75 | 608298.38 | 691769.12 | 651370 | 453173.53 | 751486.69 |
| Q02527 | Mgat3 | Beta-1,4-mannosyl-glycoprotein 4-beta-N-acetylglucosaminyltransferase Rattus norvegi | 52.47 | 1.3 | 62022 | 1 | 1 | 7478302.5 | 7832926.5 | 4383691.5 | 6138261 | 8969417 | 5192263 | 4447244 | 5483558 | 3172323 | 3684922 | 3852163 | 1 | 3743604.25 | 4723743 | 1 |
| Q9CPU4 | Mgst3 | Microsomal glutathione S-transferase 3 Mus musculus | 221.11 | 33.99 | 16958 | 3 | 3 | 9168685 | 7486690 | 8288514 | 6290236 | 7887625 | 11045711 | 8713239 | 9111533 | 11396111 | 10221440 | 12560580 | 8997930 | 7340395 | 7246861 | 9107664 |
| Q8BI84 | Mia3 | Transport and Golgi organization protein 1 homolog Mus musculus | 39.75 | 0.62 | 213675 | 1 | 1 | 1082089.38 | 883635.25 | 987032.12 | 1040188.593 | 1207997.62 | 1105673.38 | 1110250.188 | 1066333 | 1068194.62 | 1200799.75 | 866228.88 | 917563.19 | 942392.38 | 736970.12 | 1023616.06 |
| Q8R404 | Mic13 | MICOS complex subunit MIC13 Mus musculus | 246.72 | 68.91 | 13373 | 4 | 5 | 8285664 | 10155471 | 7339599 | 7039126.5 | 8109818.5 | 9119759 | 7970191.5 | 9567863 | 10537282 | 7163515 | 7716787.5 | 7123086 | 9093156 | 9140297 | 5202812 |
| Q8VDP3 | Mical1 | [F-actin]-monooxygenase MICAL1 Mus musculus | 145.43 | 6.11 | 116785 | 3 | 3 | 1849722 | 1880653.12 | 1756609.62 | 2122856.25 | 2245656.75 | 2028367.75 | 2749129.75 | 1927275.5 | 469583.66 | 2346413.75 | 2063351.62 | 1673999.12 | 1672303 | 1603599.12 | 2057392.62 |
| Q6P6Q9 | Micu1 | Calcium uptake protein 1, mitochondrial Rattus norvegicus | 66.62 | 3.14 | 54195 | 1 | 1 | 344107.8825 | 471585.47 | 205662.03 | 347466.62 | 351717.41 | 549101.94 | 1 | 381531.91 | 347559.66 | 1 | 341184.91 | 339641.31 | 304711.6925 | 165603.14 | 372417.41 |
| P34884 | Mif | Macrophage migration inhibitory factor Mus musculus | 354.63 | 83.48 | 12504 | 5 | 5 | 53402745 | 43973705 | 51153873 | 36193417 | 43227489 | 56815617 | 46628553 | 66941141 | 62565797 | 34336081 | 105319169 | 51559233 | 40566277 | 68917201 | 48404857 |
| Q3TB92 | Milr1 | Allergin-1 Mus musculus | 71.87 | 5.69 | 27366 | 1 | 1 | 221939.36 | 275979.75 | 322224.44 | 265192 | 387710.78 | 350095.56 | 225470.28 | 367820.69 | 431001.47 | 268989.25 | 600115.06 | 314987.16 | 347028.81 | 340935.12 | 442903.75 |
| Q76LS9 | Mindy1 | Ubiquitin carboxyl-terminal hydrolase MINDY-1 Mus musculus | 79.69 | 4.49 | 51226 | 1 | 1 | 580975.62 | 1 | 568249.25 | 719290.12 | 1 | 582233.38 | 703725.88 | 690074.38 | 1 | 1 | 563645.31 | 446902.47 | 518899.44 | 544714.94 | 1 |
| Q9Z2L6 | Minpp1 | Multiple inositol polyphosphate phosphatase 1 Mus musculus | 92.8 | 6.44 | 54537 | 2 | 2 | 257190.27 | 1 | 1 | 1 | 1 | 1 | 365241.66 | 396947.47 | 197719.47 | 1 | 269899.19 | 236273.56 | 1 | 1 | 1 |
| E9PVX6 | Mki67 | Proliferation marker protein Ki-67 Mus musculus | 3338.53 | 27.7 | 350864 | 60 | 60 | 194457617 | 64824269 | 67822953 | 70583001 | 81777945 | 58333197 | 84126649 | 55133081 | 42261441 | 71884745 | 35683229 | 43975813 | 57443013 | 36264141 | 59601305 |
| O08605 | Mknk1 | MAP kinase-interacting serine/threonine-protein kinase 1 Mus musculus | 48.96 | 6.32 | 47915 | 1 | 1 | 296722.34 | 372066.1075 | 349093.56 | 498455.59 | 343992.94 | 511366.53 | 432717.53 | 325888.56 | 243706.64 | 317589.69 | 1175411.38 | 1 | 408323.94 | 726518.94 | 460942.31 |
| Q6ZQI3 | Mlec | Malectin Mus musculus | 238.37 | 16.49 | 32342 | 4 | 4 | 7046681.5 | 6176772.5 | 7601901 | 7534347.5 | 7374707 | 7299998 | 7641296 | 7809415.5 | 7927231 | 7339636 | 7884431.5 | 6805549 | 6738093 | 6871492.5 | 7697427.5 |
| P97679 | Mlh1 | DNA mismatch repair protein Mlh1 Rattus norvegicus | 86.38 | 5.28 | 84540 | 2 | 2 | 1 | 1 | 1 | 1 | 1 | 1 | 200922.08 | 1 | 117705.29 | 148997.25 | 303186.66 | 1 | 1 | 1 | 1 |
| P34960 | Mmp12 | Macrophage metalloelastase Mus musculus | 180.84 | 8.67 | 54971 | 4 | 4 | 3640142.5 | 664269.56 | 2515680.75 | 1 | 2652994.75 | 9242780 | 7025381.5 | 7287591 | 7860636 | 4319993.5 | 2635937.25 | 772832.56 | 3587987.5 | 4561804.5 | 3832573.5 |
| Q9D071 | Mms19 | MMS19 nucleotide excision repair protein homolog Mus musculus | 833.62 | 19.98 | 113089 | 16 | 16 | 10553330 | 9221958 | 10853849 | 9209429 | 9126669 | 9742415 | 6968886 | 9922578 | 10819175 | 8296782.5 | 11743134 | 9127397 | 8585987 | 11628490 | 9991583 |
| P0DOV1 | Mnda | Interferon-activable protein 205-B Mus musculus | 59.12 | 2.82 | 46977 | 1 | 1 | 1493655.88 | 1912190.38 | 1427573.62 | 1756698.25 | 1660475.75 | 1210737.75 | 1551060.12 | 1608413.12 | 1409332.5 | 1121738.5 | 856138.88 | 1074790.88 | 1283166.38 | 1 | 1 |
| D0QMC3 | Mndal | Myeloid cell nuclear differentiation antigen-like protein Mus musculus | 513.11 | 22.68 | 60526 | 6 | 11 | 21679509 | 23776097 | 24524045 | 21039025 | 22592671 | 25138805 | 16432444 | 19662113 | 25045779 | 16369926 | 20291685 | 21405621 | 18382235 | 29141609 | 19207615 |
| Q3T1J9 | Mob1a | MOB kinase activator 1A Rattus norvegicus | 133.11 | 13.89 | 25080 | 3 | 3 | 15805977 | 9417855 | 8342740.5 | 8268693 | 9284386 | 8516825 | 9384810 | 10312929 | 9336459 | 9482547 | 7404905 | 8067255 | 9006929 | 8927687 | 9927971 |
| Q8VI63 | Mob2 | MOB kinase activator 2 Mus musculus | 122.93 | 10.64 | 26851 | 3 | 3 | 1377899.12 | 1714181.12 | 1378204.62 | 1764960.88 | 1905178.12 | 1517040.25 | 1297537.38 | 1447131.25 | 1614476.75 | 1327483.12 | 1425693.12 | 1401620.12 | 1414090.25 | 1656007.5 | 1352261.75 |
| A2BDX3 | Mocs3 | Adenylyltransferase and sulfurtransferase MOCS3 Mus musculus | 96.99 | 6.74 | 49375 | 3 | 3 | 688121.12 | 1 | 1 | 1 | 443013.94 | 299968.59 | 1 | 498892.06 | 1 | 1 | 1 | 402425.06 | 1 | 1 | 561757.62 |
| Q80UM7 | Mogs | Mannosyl-oligosaccharide glucosidase Mus musculus | 693.4 | 23.26 | 91831 | 6 | 12 | 7922013 | 5397955 | 5427015 | 6176937.5 | 7854608 | 7084843 | 7770154.5 | 6361268.5 | 7237773 | 7770962.5 | 6425466.5 | 5793141 | 6932134 | 5485247 | 4880962 |
| Q80TL7 | Mon2 | Protein MON2 homolog Mus musculus | 269.9 | 4.66 | 189078 | 6 | 6 | 1295220.62 | 1792681.75 | 1658617.25 | 700139.25 | 525037.75 | 414465.31 | 1500135.88 | 2031197.75 | 2007654.62 | 1642909.12 | 512913.81 | 1677216 | 1990791.88 | 1744468 | 1476472.62 |
| P23249 | Mov10 | Putative helicase MOV-10 Mus musculus | 410.31 | 9.76 | 113583 | 8 | 8 | 6950960 | 6710272 | 6152657.5 | 4463334 | 5575883 | 6229301 | 5949671.5 | 7663310.5 | 4792296.5 | 4446301 | 4205508 | 3757924 | 7266387 | 4960285.5 | 3994688.25 |
| P63030 | Mpc1 | Mitochondrial pyruvate carrier 1 Mus musculus | 79.86 | 19.27 | 12455 | 1 | 1 | 487731.34 | 372120.62 | 303557.84 | 549422.75 | 332989.03 | 684115.81 | 351809.12 | 545433.12 | 383697.88 | 313133.59 | 589057.12 | 300158.16 | 443527.03 | 492225.72 | 453787.78 |
| P38718 | Mpc2 | Mitochondrial pyruvate carrier 2 Rattus norvegicus | 117.89 | 28.35 | 14258 | 3 | 3 | 3960276.25 | 6464934 | 5513718 | 3025832.5 | 6765437 | 4823519.5 | 5626982 | 7071609 | 7485481 | 6063197 | 4316270 | 4855503.5 | 5723050 | 5704676 | 4712549.5 |
| Q9R0Q9 | Mpdu1 | Mannose-P-dolichol utilization defect 1 protein Mus musculus | 159.91 | 12.55 | 26498 | 3 | 3 | 5326098.5 | 4257626.5 | 4327888 | 3015309 | 4579706 | 4681526 | 2852262 | 4501807.5 | 5469042 | 3522867.25 | 4155700.75 | 4377900 | 5887792 | 5008082 | 4414582 |
| A1L314 | Mpeg1 | Macrophage-expressed gene 1 protein Mus musculus | 1083.6 | 44.04 | 78390 | 18 | 18 | 51348273 | 40907589 | 51192469 | 39580849 | 48383313 | 43501053 | 36561421 | 32908427 | 45002225 | 41012809 | 59094585 | 54659457 | 48381089 | 40596129 | 55119353 |
| Q810V0 | Mphosph10 | U3 small nucleolar ribonucleoprotein protein MPP10 Mus musculus | 118.73 | 10.28 | 78735 | 3 | 3 | 854143.12 | 794846.12 | 663416.75 | 879502.38 | 1050323.12 | 1063610.25 | 1179874.12 | 919820.75 | 981474.94 | 915290.19 | 986946.5025 | 1018628.94 | 1024030 | 909479.38 | 995647.69 |
| Q9D1Q1 | Mphosph6 | M-phase phosphoprotein 6 Mus musculus | 39.11 | 6.83 | 19086 | 1 | 1 | 1656604.75 | 1630591 | 1144415.12 | 2229464 | 1933099.5 | 1710344 | 1572283.88 | 1359665.12 | 1591341.5 | 1971318.25 | 1758521.88 | 1706928.38 | 1937863.62 | 1444112.38 | 1398851.88 |
| P70290 | Mpp1 | 55 kDa erythrocyte membrane protein Mus musculus | 329.81 | 19.53 | 52227 | 6 | 6 | 6443307 | 4687410 | 8043545 | 4995342.5 | 4394527.5 | 5472213 | 5539893 | 5376055.5 | 5633835.5 | 3734910 | 5405972 | 6746709 | 5115590 | 5375714 | 6141201.5 |
| Q9JLB0 | Mpp6 | MAGUK p55 subfamily member 6 Mus musculus | 317.28 | 15.37 | 62631 | 6 | 6 | 5821362.5 | 6624546.5 | 6686558 | 4830265.5 | 5605394 | 6285822 | 6255780 | 6780115 | 6603355 | 6432459.5 | 6603153 | 5986424 | 6271730 | 6631751.5 | 4821817 |
| Q99J99 | Mpst | 3-mercaptopyruvate sulfurtransferase Mus musculus | 83.11 | 13.13 | 33097 | 2 | 2 | 1035780.94 | 1259697.62 | 1375035.5 | 965799.88 | 2324861.25 | 1728159 | 2077730.5 | 1079785 | 1743364.75 | 906486.12 | 981048.31 | 1095510.75 | 422835.97 | 1081370.75 | 1477096.5 |
| Q61216 | Mre11 | Double-strand break repair protein MRE11 Mus musculus | 259.71 | 9.92 | 80223 | 6 | 6 | 598519.19 | 4096804.5 | 4784435 | 2311033.5 | 1564075.12 | 3189718.5 | 3554439 | 4163245 | 3717477.75 | 3377057.75 | 6498223 | 3543446.75 | 3247432.5 | 4071206 | 3126020 |
| Q9DAT2 | Mrgbp | MRG/MORF4L-binding protein Mus musculus | 47.29 | 6.37 | 22436 | 1 | 1 | 802670.06 | 979215.6225 | 1053447.12 | 1030878.19 | 1029867.12 | 1 | 1 | 871928.31 | 787295.5 | 1 | 773234.75 | 1110343.38 | 810500.25 | 685922 | 845000.095 |
| Q9CQT1 | Mri1 | Methylthioribose-1-phosphate isomerase Mus musculus | 494.63 | 38.75 | 39411 | 9 | 9 | 6321781.5 | 6129971 | 6594098 | 5600923.5 | 6837747 | 7896112 | 6479865 | 7192331 | 8182895 | 9877010 | 8995960 | 6502941 | 7852937 | 6185218 | 5944828 |
| Q99N96 | Mrpl1 | 39S ribosomal protein L1, mitochondrial Mus musculus | 45.19 | 6.25 | 37597 | 1 | 1 | 979944.81 | 1 | 659672 | 1 | 1 | 853443 | 1 | 1 | 1 | 776193.38 | 442790.72 | 459405.59 | 942534.75 | 647606.25 | 623084.3275 |
| Q3TBW2 | Mrpl10 | 39S ribosomal protein L10, mitochondrial Mus musculus | 145.13 | 16.03 | 29396 | 3 | 3 | 7641554 | 8895426 | 1781394.5 | 9099606 | 2036918.5 | 1668218.88 | 1954193.62 | 1819033.12 | 1807074.12 | 1573250.38 | 6118583.5 | 1444522.88 | 2194353.25 | 2396348.75 | 1703930.38 |
| Q7M0E7 | Mrpl14 | 39S ribosomal protein L14, mitochondrial Rattus norvegicus | 71.54 | 6.9 | 15913 | 1 | 1 | 632822.62 | 389845.28 | 377724.06 | 928022.75 | 734948.62 | 435415.09 | 532290.88 | 350100.28 | 461436.6575 | 527940.38 | 567333.06 | 1 | 334776.44 | 1 | 428715.12 |
| Q9CPR5 | Mrpl15 | 39S ribosomal protein L15, mitochondrial Mus musculus | 45.13 | 3.05 | 33542 | 1 | 1 | 894098.5 | 766796 | 1178934 | 936892.38 | 944180.22 | 891385.62 | 1158516.5 | 953509.4375 | 810194.38 | 953941.25 | 737478.69 | 1218189.38 | 1019444.06 | 873498.69 | 1210646 |
| Q9D338 | Mrpl19 | 39S ribosomal protein L19, mitochondrial Mus musculus | 107.4 | 10.62 | 33578 | 3 | 3 | 1592018.25 | 2688175.25 | 3359624.5 | 4051172.75 | 3064025 | 2945403.5 | 3582468 | 2888656 | 2926600 | 2337007.5 | 3604516 | 3482047.75 | 3411375.5 | 630010.25 | 3443400.5 |
| Q9CQL4 | Mrpl20 | 39S ribosomal protein L20, mitochondrial Mus musculus | 44.48 | 7.38 | 17595 | 1 | 1 | 784133 | 1138434 | 711862.38 | 820748 | 988449.62 | 807455.56 | 432574 | 419399.38 | 775752.62 | 964691.5 | 474564.16 | 767937.62 | 1247984.5 | 621086.81 | 803246.88 |
| Q9D1N9 | Mrpl21 | 39S ribosomal protein L21, mitochondrial Mus musculus | 138.58 | 17.7 | 23366 | 3 | 3 | 1 | 1 | 717395.31 | 1 | 239972.38 | 1 | 989860 | 246164.58 | 1 | 918198.88 | 403308.34 | 335380.66 | 573892.44 | 312881.44 | 1039761 |
| Q9D1B9 | Mrpl28 | 39S ribosomal protein L28, mitochondrial Mus musculus | 111.93 | 18.68 | 30170 | 3 | 3 | 1 | 323581.31 | 1 | 1 | 1 | 1 | 1 | 247868.5 | 1 | 1 | 1 | 237007.3 | 1 | 1 | 1 |
| Q6AXT0 | Mrpl37 | 39S ribosomal protein L37, mitochondrial Rattus norvegicus 2 | 106.56 | 7.09 | 48366 | 2 | 2 | 1101481.5 | 817284.75 | 1022061.88 | 1101236.25 | 898417.94 | 1058787.75 | 993076.19 | 995783.12 | 1057020.25 | 1026661.38 | 946023.19 | 1118219.88 | 864697.88 | 822143.19 | 999798.25 |
| Q9JKF7 | Mrpl39 | 39S ribosomal protein L39, mitochondrial Mus musculus | 79.52 | 8.04 | 38549 | 2 | 2 | 2685010.063 | 2745866 | 1424881.75 | 3128495.75 | 3440796.75 | 3176980.5 | 1304408.25 | 3134471.5 | 2943509 | 3076609.5 | 384961.59 | 1949055.62 | 412743.28 | 1640935.5 | 1677848 |
| Q9CY73 | Mrpl44 | 39S ribosomal protein L44, mitochondrial Mus musculus | 254.42 | 15.02 | 37527 | 3 | 3 | 1087725.88 | 1012119 | 1242485.88 | 1022207.38 | 918183.12 | 1000681.44 | 1059136.12 | 1086006.88 | 1040056.38 | 1271801 | 1001467.31 | 1110797 | 813456 | 1128601.88 | 1042115.12 |
| Q9D0Q7 | Mrpl45 | 39S ribosomal protein L45, mitochondrial Mus musculus | 110.26 | 15.03 | 35411 | 3 | 3 | 2053782.88 | 1848439.12 | 2141995.25 | 1893388.25 | 2197622.5 | 2317783.5 | 4101546 | 1598053.38 | 1565322.25 | 3084533.5 | 2222800 | 2173797.25 | 2020370.25 | 1462094 | 3146124.5 |
| Q9EQI8 | Mrpl46 | 39S ribosomal protein L46, mitochondrial Mus musculus | 139.41 | 14.13 | 32132 | 3 | 3 | 861975.12 | 601137.44 | 545947.44 | 716169.155 | 855616.62 | 1018796.25 | 806903.5 | 944854.56 | 1143185.75 | 909091 | 609543.5 | 872724.38 | 693577.38 | 775235.19 | 863920.81 |
| Q9CPW3 | Mrpl54 | 39S ribosomal protein L54, mitochondrial Mus musculus | 56.12 | 13.33 | 15422 | 1 | 1 | 695990 | 584164.38 | 978873.12 | 794012.12 | 825405.12 | 1 | 1 | 1 | 663800.56 | 1 | 1 | 1 | 1 | 1 | 928411.44 |
| Q9CPX7 | Mrps16 | 28S ribosomal protein S16, mitochondrial Mus musculus | 38.09 | 5.93 | 15192 | 1 | 1 | 1601795.12 | 1679947.62 | 1821793.75 | 2012624.5 | 1772623.25 | 1265519.12 | 1292718.12 | 1539882.25 | 1866163.12 | 1924255 | 1616489 | 1371493.5 | 1597628 | 1363025.62 | 1617792.38 |
| Q9CQE3 | Mrps17 | 28S ribosomal protein S17, mitochondrial Mus musculus | 32.7 | 7.5 | 13382 | 1 | 1 | 596581.88 | 571504.94 | 569459.94 | 384478.53 | 563171 | 563863.19 | 862836.19 | 567870.62 | 438414.75 | 1 | 455557.34 | 781190.94 | 624149.56 | 653230.81 | 708543.12 |
| Q924T2 | Mrps2 | 28S ribosomal protein S2, mitochondrial Mus musculus | 210.62 | 23.02 | 32313 | 4 | 4 | 3232446 | 2209466.5 | 3335009 | 2988228.25 | 3097873.75 | 3085847.75 | 1759312.38 | 3309275.75 | 4721167 | 3777705.25 | 3645021.75 | 3324964 | 3483462.25 | 3036574.75 | 2999895.75 |
| P58059 | Mrps21 | 28S ribosomal protein S21, mitochondrial Mus musculus | 91.71 | 14.94 | 10561 | 1 | 1 | 1 | 1 | 1 | 1 | 1 | 1 | 6103944.5 | 1 | 1 | 1 | 4720752 | 4724733.5 | 4083026.89 | 482241.06 | 6404381 |
| Q9CXW2 | Mrps22 | 28S ribosomal protein S22, mitochondrial Mus musculus | 97.09 | 6.41 | 41192 | 2 | 2 | 1 | 471504.78 | 1 | 1 | 1741433.75 | 903099 | 869618.6075 | 1482049.12 | 529526.69 | 563799.62 | 1 | 1 | 439971.75 | 1 | 1 |
| Q8VE22 | Mrps23 | 28S ribosomal protein S23, mitochondrial Mus musculus | 212.44 | 24.86 | 20348 | 4 | 4 | 5079033 | 5106976.5 | 6020982 | 4458610.5 | 5111542 | 4671242 | 4978623 | 4932307 | 4941034 | 5571765.5 | 4894941.5 | 5323363 | 4971036 | 5604231 | 4751768 |
| Q9CQV5 | Mrps24 | 28S ribosomal protein S24, mitochondrial Mus musculus | 100.12 | 11.98 | 18901 | 2 | 2 | 3103318 | 3143296 | 2894432 | 2772794.25 | 3048753.25 | 3277156 | 2771649.5 | 1704159 | 2068711.25 | 1282162.75 | 2735666.25 | 2718346.5 | 3038648.5 | 3170506 | 2656759.5 |
| Q4QR80 | Mrps25 | 28S ribosomal protein S25, mitochondrial Rattus norvegicus 2 | 48.31 | 7.6 | 19775 | 1 | 1 | 551060.31 | 297166.72 | 379615.66 | 630213.75 | 412936.53 | 494020.22 | 570622.38 | 416734.5 | 390915.66 | 703123.56 | 470500 | 385005.75 | 489437.94 | 443061.94 | 244322.8 |
| Q8BK72 | Mrps27 | 28S ribosomal protein S27, mitochondrial Mus musculus | 225.09 | 14.22 | 47779 | 5 | 5 | 4435851 | 4147179.25 | 6525800 | 4556549 | 3804033.25 | 4244422.5 | 4000516 | 6534715.5 | 7128624 | 4838162 | 5665711 | 6864255 | 4656785.5 | 6378647 | 5731083 |
| Q9CY16 | Mrps28 | 28S ribosomal protein S28, mitochondrial Mus musculus | 201.01 | 23.12 | 20520 | 3 | 3 | 8898625 | 3262783.75 | 3165366.75 | 2974766.75 | 2653114 | 10015390 | 2524615.25 | 2946718.5 | 3108333 | 2797092.5 | 2972559.75 | 3056227 | 2392439 | 2540127.75 | 2592871.5 |
| Q9D0G0 | Mrps30 | 28S ribosomal protein S30, mitochondrial Mus musculus | 214.45 | 16.52 | 49939 | 4 | 4 | 1011809.5 | 96723.46 | 1106397.88 | 1256107.75 | 1277453.75 | 2111868.25 | 2451351.5 | 1135040.25 | 1273020 | 2426968.5 | 2553890.25 | 1687515.62 | 2055870 | 1117331.88 | 2717801 |
| Q61733 | Mrps31 | 28S ribosomal protein S31, mitochondrial Mus musculus | 148.48 | 13.54 | 43881 | 3 | 3 | 26679185 | 36153341 | 27488103 | 34474009 | 34572117 | 31374403 | 15706679 | 29199931 | 36221877 | 24155491 | 29000823 | 23182697 | 32606915 | 33083319 | 16407108 |
| Q9JIK9 | Mrps34 | 28S ribosomal protein S34, mitochondrial Mus musculus | 76.26 | 6.88 | 25827 | 2 | 2 | 3645397.25 | 1378870.12 | 1469048.5 | 1533717.12 | 1899317.62 | 1447089.75 | 1785502.75 | 1641718.75 | 1593488.62 | 1802907.12 | 1100248.75 | 1400770.12 | 1677640.75 | 1310673.75 | 1754472 |
| Q8BJZ4 | Mrps35 | 28S ribosomal protein S35, mitochondrial Mus musculus | 77.65 | 12.5 | 35975 | 2 | 2 | 1252758.38 | 385422.41 | 657607.44 | 394956.56 | 497457.38 | 423761.47 | 505323.34 | 327513.75 | 395735.72 | 1152955.12 | 776251.25 | 385034.62 | 367649.56 | 650117.94 | 255985.55 |
| P58064 | Mrps6 | 28S ribosomal protein S6, mitochondrial Mus musculus | 44.2 | 6.4 | 14309 | 1 | 1 | 925275.12 | 452644.59 | 603942.81 | 514467.69 | 467362.88 | 738712.19 | 624573.445 | 628269.38 | 478109.09 | 653203.12 | 241103.5 | 824089.69 | 591464.19 | 567668.31 | 546933.06 |

| | | | | | | | | | | | | | | | | | | | | | | |
|---|---|---|---|---|---|---|---|---|---|---|---|---|---|---|---|---|---|---|---|---|---|---|
| Q5I0K8 | Mrps7 | 28S ribosomal protein S7, mitochondrial Rattus norvegicus | 71.51 | 4.96 | 28197 | 1 | 1 | 2196894.25 | 593319.44 | 1789828.5 | 3287881 | 1966980.798 | 749228.88 | 2454863.25 | 1904289.25 | 1432760.25 | 2076166.5 | 1 | 1449954.38 | 1 | 1 | 1426669.5 |
| Q9D7N3 | Mrps9 | 28S ribosomal protein S9, mitochondrial Mus musculus | 133.67 | 10.77 | 44929 | 3 | 3 | 1 | 1 | 191513.38 | 311392.88 | 357242.94 | 1 | 1 | 1 | 1 | 474847.38 | 1 | 214387.03 | 1 | 1 | 205243.77 |
| Q9D0I8 | Mrto4 | mRNA turnover protein 4 homolog Mus musculus | 345.49 | 31.8 | 27546 | 8 | 8 | 10086518 | 9320718 | 10442178 | 9928135 | 9072290 | 9172751 | 10028885 | 9756648 | 10683085 | 9057641 | 11541089 | 10531672 | 9014316 | 10378425 | 9800875 |
| P43247 | Msh2 | DNA mismatch repair protein Msh2 Mus musculus | 918.75 | 24.17 | 104151 | 15 | 15 | 10034431 | 10094448 | 12280420 | 10476568 | 9570140 | 9845860 | 12409941 | 11511225 | 11980744 | 10485193 | 9982683 | 11195778 | 11557960 | 11805912 | 12145347 |
| P54276 | Msh6 | DNA mismatch repair protein Msh6 Mus musculus | 430.13 | 7.44 | 151084 | 9 | 9 | 9817449 | 10534568 | 16664263 | 7030455 | 16395957 | 9071337 | 6040997 | 14691134 | 17489601 | 7865646 | 13439553 | 13120059 | 10961298 | 13835095 | 8963809 |
| Q9CRA4 | Msmo1 | Methylsterol monooxygenase 1 Mus musculus | 141.9 | 17.75 | 34772 | 1 | 3 | 3628856.25 | 3325793.5 | 2443600.5 | 1321450.25 | 2322797.75 | 3540707 | 3196534.75 | 2819581.75 | 4042601.75 | 2393734 | 3608496 | 2763530.75 | 2743506.5 | 3654358.5 | 2931577.25 |
| P26041 | Msn | Moesin Mus musculus | 3037.55 | 68.11 | 67767 | 34 | 50 | 691954433 | 676819137 | 608385089 | 702829697 | 649914689 | 671759041 | 778371073 | 704165889 | 625331969 | 738729025 | 601272897 | 643699009 | 646492481 | 690489921 | 611907265 |
| P30204 | Msr1 | Macrophage scavenger receptor types I and II Mus musculus | 227.23 | 9.17 | 50170 | 5 | 5 | 29196489 | 30667877 | 38901625 | 33723585 | 36951481 | 31965879 | 21496055 | 27872259 | 33922577 | 26876545 | 38371281 | 34472853 | 31199197 | 33114189 | 32142261 |
| Q9D6Y7 | Msra | Mitochondrial peptide methionine sulfoxide reductase Mus musculus | 86.64 | 6.87 | 25988 | 1 | 1 | 1688829.12 | 1673941.5 | 1439020.12 | 1893916.5 | 1607827.38 | 1459937.62 | 1791646.5 | 1410751.25 | 1255386.5 | 1911761.25 | 1888425.62 | 1778847.88 | 1643723.38 | 1408810.88 | 1675769.38 |
| Q2YDW2 | Msto1 | Protein misato homolog 1 Mus musculus | 144.39 | 8.45 | 61230 | 3 | 3 | 717175.69 | 994928.44 | 1171781.75 | 1458062.75 | 1224273.62 | 883674.31 | 1213271.75 | 1148199.38 | 1215096.62 | 1146711.62 | 969233.25 | 1252001.38 | 918501.12 | 1172062.5 | 1079078 |
| P02802 | Mt1 | Metallothionein-1 Mus musculus | 193.08 | 68.85 | 6018 | 3 | 3 | 1 | 3184470.75 | 1 | 1963729.25 | 533792.25 | 1452133.805 | 3113598.5 | 856847.38 | 496307.09 | 1341782.25 | 1047314.94 | 350843.34 | 2305579.5 | 6726602.5 | 1755515.38 |
| P02798 | Mt2 | Metallothionein-2 Mus musculus | 54.76 | 19.67 | 6115 | 1 | 1 | 1 | 1 | 1 | 1 | 1 | 1 | 1188942.62 | 1 | 1 | 1 | 1 | 1 | 1110625.12 | 1980507.12 | 1 |
| Q62599 | Mta1 | Metastasis-associated protein MTA1 Rattus norvegicus | 167.11 | 5.69 | 79412 | 1 | 3 | 6018245 | 5941622.5 | 5369196.5 | 6008476 | 7029515 | 5826439 | 5461351.5 | 4928652 | 6201638.5 | 5625673 | 5382556 | 5434864 | 5748372 | 5998656 | 5641112 |
| Q9R190 | Mta2 | Metastasis-associated protein MTA2 Mus musculus | 489.42 | 19.61 | 75030 | 9 | 11 | 11753179 | 11028814 | 10616998 | 12457280 | 13647856 | 12654588 | 15627111 | 10619115 | 12510635 | 14581495 | 13904392 | 13097851 | 14276408 | 12763800 | 9760561 |
| Q924K8 | Mta3 | Metastasis-associated protein MTA3 Mus musculus | 159.11 | 5.92 | 67077 | 0 | 3 | 8610102 | 7772625 | 7776590 | 9174893 | 9826137 | 7865443 | 7981966 | 7194526.5 | 8442561 | 8285850.5 | 8212292.5 | 7520600 | 8291024 | 8305427 | 2262336 |
| Q9CQ65 | Mtap | S-methyl-5'-thioadenosine phosphorylase Mus musculus | 475.76 | 42.05 | 31062 | 8 | 8 | 10984400 | 16676826 | 17895295 | 18858059 | 17826605 | 18817905 | 19822821 | 16933043 | 17379365 | 20272975 | 14052065 | 17109991 | 18156673 | 16734811 | 17138587 |
| Q791T5 | Mtch1 | Mitochondrial carrier homolog 1 Mus musculus | 140.74 | 15.42 | 41565 | 4 | 4 | 272478.12 | 1 | 1 | 1 | 1 | 438768.72 | 478555.44 | 1 | 1 | 832767.88 | 538508.19 | 594802.81 | 1 | 1 | 882335.62 |
| Q791V5 | Mtch2 | Mitochondrial carrier homolog 2 Mus musculus | 539.84 | 49.17 | 33499 | 11 | 11 | 20859653 | 18690925 | 19395263 | 19571011 | 20258059 | 26268839 | 24911621 | 26405663 | 20723793 | 24487557 | 20665377 | 19517699 | 24262715 | 22680453 | 20860165 |
| P00405 | Mtco2 | Cytochrome c oxidase subunit 2 Mus musculus | 271.03 | 26.43 | 25976 | 5 | 5 | 66056129 | 80171209 | 69072009 | 67014545 | 75373865 | 71435993 | 61583457 | 77417961 | 84754945 | 74872145 | 67592657 | 72247049 | 85413145 | 76410585 | 67923617 |
| Q80WJ7 | Mtdh | Protein LYRIC Mus musculus | 728.98 | 26.08 | 63846 | 11 | 11 | 11561751 | 14092343 | 12164339 | 11791865 | 11385665 | 14338540 | 15994349 | 14363140 | 16077521 | 13664296 | 13644385 | 12956419 | 13812068 | 10856245 | 13924995 |
| Q8BVN4 | Mterf4 | Transcription termination factor 4, mitochondrial Mus musculus 2 | 35.3 | 2.02 | 40249 | 1 | 1 | 4459182.5 | 4166899.75 | 4839473.5 | 4991558.5 | 4571420.5 | 4377548.5 | 5855536.5 | 3777747 | 4073868 | 5146128 | 5249510.5 | 3882162.25 | 4561914 | 3693646.75 | 4894913.5 |
| Q5XII9 | Mtfr1l | Mitochondrial fission regulator 1-like Rattus norvegicus | 38.76 | 5.88 | 31730 | 1 | 1 | 267904.22 | 287358.22 | 237853.92 | 502640.09 | 459358.84 | 318848.12 | 443248.19 | 346046.75 | 483711.38 | 540012.44 | 524999.69 | 484571.81 | 399810.41 | 294965.91 | 336118.53 |
| Q922D8 | Mthfd1 | C-1-tetrahydrofolate synthase, cytoplasmic Mus musculus | 979.69 | 27.38 | 101200 | 20 | 20 | 32226977 | 32587449 | 33619313 | 35887057 | 31192873 | 36066481 | 32808281 | 36769237 | 36545053 | 35955041 | 32689741 | 37854033 | 32517277 | 36351813 | 32547945 |
| Q3V3R1 | Mthfd1l | Monofunctional C1-tetrahydrofolate synthase, mitochondrial Mus musculus | 1183.16 | 30.3 | 105729 | 21 | 21 | 21410699 | 23157723 | 30254647 | 22551873 | 23307525 | 27294911 | 39552053 | 23736281 | 27575867 | 35881437 | 30535925 | 30565063 | 28135147 | 17682691 | 35271153 |
| P18155 | Mthfd2 | Bifunctional methylenetetrahydrofolate dehydrogenase/cyclohydrolase, mitochondrial M | 324.3 | 38.86 | 37863 | 7 | 7 | 6193325.5 | 5235505 | 6414935 | 1609984.25 | 5457327.5 | 5951863 | 6734630.5 | 5845368.5 | 1879967.75 | 6568827.5 | 4665446 | 1898103 | 4504863.5 | 4826602.5 | 3847455.25 |
| Q9D110 | Mthfs | 5-formyltetrahydrofolate cyclo-ligase Mus musculus | 79.25 | 10.34 | 23202 | 1 | 1 | 825248.19 | 586814.81 | 853625.56 | 654623.62 | 573649.5 | 651745.25 | 362416.28 | 858110.75 | 461058.34 | 600639.25 | 1130968.25 | 793545.06 | 408923.91 | 687912.38 | 932309.81 |
| Q8VEL2 | Mtmr14 | Myotubularin-related protein 14 Mus musculus | 95.82 | 4.32 | 72443 | 2 | 2 | 1152797.75 | 907174.81 | 900426.94 | 1559847 | 1803581.5 | 953258.75 | 640350.81 | 756780.44 | 1150351.5 | 1498338.75 | 880187.19 | 1233279.5 | 1538195.25 | 904430.44 | 1488916.75 |
| Q5PQT2 | Mtmr3 | Myotubularin-related protein 3 Rattus norvegicus | 38.67 | 1.42 | 133500 | 1 | 1 | 147283.08 | 1 | 218126.02 | 206626.69 | 1 | 129225.39 | 1 | 142201.52 | 203858.91 | 1 | 83251.91 | 144207.64 | 126496.165 | 158820.64 | 119704.47 |
| P03911 | Mtnd4 | NADH-ubiquinone oxidoreductase chain 4 Mus musculus | 73.08 | 6.1 | 51882 | 2 | 2 | 2704138.25 | 1376480.75 | 574861.81 | 1463478.828 | 1198434.5 | 1566246.62 | 871606 | 810858.62 | 936965.56 | 323681.34 | 1114182.38 | 1239144 | 1453201.38 | 1700520.38 | 1191121.25 |
| P03921 | Mtnd5 | NADH-ubiquinone oxidoreductase chain 5 Mus musculus | 87.32 | 5.44 | 68475 | 2 | 2 | 696661.12 | 653871.69 | 411395.28 | 362957.81 | 502774.44 | 532770.94 | 288829.19 | 438108 | 487623.97 | 691814.19 | 391861.72 | 426498.38 | 444295.12 | 461585.19 | 393269.16 |
| P42346 | Mtor | Serine/threonine-protein kinase mTOR Rattus norvegicus | 173.43 | 1.8 | 288794 | 3 | 3 | 373131.56 | 360144.03 | 170056.17 | 984476.19 | 700564.44 | 435611.38 | 797848.44 | 588460.38 | 414648.69 | 532946.5 | 418622.06 | 679473.31 | 761185.44 | 629255.31 | 559204.31 |
| P62774 | Mtpn | Myotrophin Mus musculus | 212.54 | 43.22 | 12861 | 3 | 3 | 12414931 | 9254644 | 8558970 | 11747155 | 10568561 | 16165451 | 18051709 | 13240505 | 10138163 | 17026961 | 14373694 | 14042488 | 13171149 | 13982410 | 14974792 |
| Q9CZU3 | Mtrex | Exosome RNA helicase MTR4 Mus musculus | 704.57 | 17.5 | 117636 | 13 | 13 | 11266927 | 13654202 | 12180767 | 12814407 | 12901595 | 12328153 | 11354130 | 11957718 | 14201503 | 12295121 | 11427858 | 13452144 | 12865590 | 12701359 | 11112904 |
| Q8R1S4 | Mtss1 | Metastasis suppressor protein 1 Mus musculus | 496.42 | 18.45 | 82408 | 10 | 10 | 9374076 | 7320747 | 12592615 | 9704686 | 9922731 | 11249242 | 14471201 | 11002817 | 6368768 | 13305278 | 10031719 | 12152684 | 12040544 | 6114211 | 11538070 |
| P47802 | Mtx1 | Metaxin-1 Mus musculus | 233.22 | 16.72 | 35624 | 5 | 5 | 2503474.25 | 1896446.12 | 1906389 | 5213587 | 1850215.25 | 2466114.75 | 1544003.25 | 2197440.25 | 2293766.5 | 1583859.5 | 1468828 | 1649796.25 | 944485.19 | 2388024.25 | 5136908 |
| O88441 | Mtx2 | Metaxin-2 Mus musculus | 363.42 | 51.71 | 29758 | 6 | 6 | 11953773 | 15446370 | 16026865 | 13889297 | 12873955 | 14631909 | 13939845 | 18824101 | 13615977 | 10107697 | 15717619 | 15175725 | 12913724 | 13155752 | 14257678 |
| P16332 | Mut | Methylmalonyl-CoA mutase, mitochondrial Mus musculus | 326.95 | 15.11 | 82844 | 7 | 7 | 7811386 | 8878732 | 7843282 | 8798362 | 8054289.5 | 9480040 | 9308007 | 9888978 | 9556689 | 10542777 | 5362669.5 | 7747002 | 7738866.5 | 9155077 | 7098422 |
| Q99JF5 | Mvd | Diphosphomevalonate decarboxylase Mus musculus | 633.68 | 33.42 | 44072 | 11 | 11 | 4716165 | 2736008 | 3696750 | 3064832.5 | 2196102 | 5317029 | 5740591 | 3615557.75 | 3545672 | 5091995 | 5095548.5 | 4096635.5 | 3180471.5 | 3533714.75 | 4571157.5 |
| Q9R008 | Mvk | Mevalonate kinase Mus musculus | 98.03 | 7.85 | 41877 | 2 | 2 | 1 | 1 | 159690.31 | 145431.55 | 1 | 1 | 72010.27 | 1 | 1 | 1 | 131975.05 | 93893.99 | 95965.9975 | 83132.19 | 74862.76 |
| Q9EQK5 | Mvp | Major vault protein Mus musculus | 1618.74 | 49.13 | 95924 | 26 | 26 | 34631369 | 49552661 | 51757525 | 35835677 | 33339521 | 48483757 | 59953017 | 49242329 | 46167037 | 59462853 | 50196069 | 49707045 | 52084957 | 46827289 | 53554565 |
| O35682 | Myadm | Myeloid-associated differentiation marker Mus musculus | 142.12 | 16.25 | 35285 | 3 | 3 | 1 | 400232.06 | 1 | 628709.5 | 532174.56 | 1 | 978064.12 | 921647.62 | 1 | 988146.12 | 661894.69 | 863061.44 | 631438.06 | 820105.0475 | 1124026 |
| Q7TPV4 | Mybbp1a | Myb-binding protein 1A Mus musculus | 1507.34 | 28.5 | 152037 | 21 | 27 | 54774545 | 29585349 | 29353129 | 34995205 | 34132297 | 35005301 | 27255643 | 34150005 | 28435289 | 32564989 | 35526153 | 31045971 | 34012405 | 31362745 | 33035979 |
| P22366 | Myd88 | Myeloid differentiation primary response protein MyD88 Mus musculus | 127.41 | 15.54 | 33753 | 3 | 3 | 241400.28 | 381308.16 | 334468.31 | 323696.16 | 173338.16 | 675455.5 | 338819.88 | 748254.38 | 648797.44 | 146790.27 | 437649.84 | 326204.19 | 500238.94 | 724273.5 | 385349.03 |
| Q9CPT4 | Mydgf | Myeloid-derived growth factor Mus musculus | 164.89 | 21.69 | 17982 | 4 | 4 | 11454445 | 12826983 | 12146643 | 12124269 | 12093198 | 12994965 | 9486807 | 12052926 | 14675054 | 10846009 | 10593973 | 10177813 | 10693323 | 13319679 | 8674148 |
| Q8C854 | Myef2 | Myelin expression factor 2 Mus musculus | 368.43 | 17.26 | 63295 | 8 | 8 | 12451083 | 10399837 | 13744827 | 13478114 | 12110127 | 12343095 | 19700153 | 8007901 | 10526763 | 17143715 | 16479902 | 15093848 | 16291723 | 9874145 | 18097647 |
| Q9JK81 | Myg1 | UPF0160 protein MYG1, mitochondrial Mus musculus | 541.26 | 36.32 | 42723 | 11 | 11 | 17665041 | 19092641 | 16112001 | 19558717 | 18558771 | 15172226 | 14119496 | 16615068 | 14502936 | 13141444 | 11896425 | 12494022 | 16402564 | 17723461 | 13808548 |
| Q61879 | Myh10 | Myosin-10 Mus musculus | 1341.1 | 16.3 | 228996 | 8 | 24 | 74988881 | 60605669 | 69193473 | 77776465 | 65900389 | 69059809 | 86483169 | 67327729 | 60804973 | 77094769 | 90852697 | 79134369 | 67801033 | 68004025 | 88711889 |
| O08638 | Myh11 | Myosin-11 Mus musculus | 687.13 | 5.93 | 227028 | 0 | 12 | 73974265 | 58429205 | 65075965 | 70564665 | 50668577 | 61408245 | 68093041 | 61273837 | 48171845 | 74556745 | 71069481 | 64397861 | 61788985 | 57310161 | 79841241 |
| Q6URW6 | Myh14 | Myosin-14 Mus musculus | 411.8 | 4.15 | 228586 | 1 | 7 | 43131049 | 32586251 | 38244089 | 42434093 | 36154369 | 35196869 | 51431373 | 36383325 | 28295087 | 46652965 | 40019805 | 40153841 | 39097297 | 33659197 | 50548921 |
| Q8VDD5 | Myh9 | Myosin-9 Mus musculus | 7949.92 | 59.54 | 226372 | 96 | 119 | 543778561 | 529906881 | 549097857 | 587732737 | 528381601 | 517495297 | 649327425 | 520414305 | 495540129 | 604882497 | 708237185 | 588931585 | 555899713 | 548900289 | 635551873 |
| Q3THE2 | Myl12b | Myosin regulatory light chain 12B Mus musculus | 507.79 | 57.56 | 19779 | 1 | 8 | 51397537 | 54624149 | 54150541 | 53190337 | 52433781 | 50693841 | 43056745 | 61615581 | 64047113 | 41761297 | 64054729 | 60438733 | 45674753 | 61436193 | 53196045 |
| P16409 | Myl3 | Myosin light chain 3 Rattus norvegicus | 152.53 | 12 | 22156 | 1 | 2 | 49731337 | 46274817 | 55362829 | 41102137 | 40091953 | 43833953 | 34905141 | 58699341 | 61523741 | 29426739 | 54996253 | 50806057 | 32071827 | 61523129 | 45168173 |
| Q60605 | Myl6 | Myosin light polypeptide 6 Mus musculus | 692.56 | 66.23 | 16930 | 9 | 10 | 247381857 | 287915105 | 259841569 | 272482433 | 250616641 | 247928753 | 215454673 | 270071713 | 271766785 | 210760689 | 263635601 | 252097921 | 247731697 | 285546561 | 211839985 |
| Q9CQ19 | Myl9 | Myosin regulatory light polypeptide 9 Mus musculus | 357.35 | 45.35 | 19854 | 1 | 6 | 26152073 | 28341347 | 28584151 | 26103673 | 24852313 | 23432933 | 23581819 | 32437901 | 30725433 | 18637333 | 31501957 | 28654913 | 23204717 | 32794059 | 27094915 |
| Q9JMH9 | Myo18a | Unconventional myosin-XVIIIa Mus musculus | 490.17 | 6.59 | 232755 | 9 | 9 | 4793741 | 6160377.5 | 3045286.25 | 4735160.5 | 3423998.25 | 4580609 | 5992669 | 4113594 | 2869131.25 | 5575406.5 | 4456787.5 | 4444380.5 | 3791967 | 4646501.5 | 6960737 |
| Q9WTI7 | Myo1c | Unconventional myosin-Ic Mus musculus | 1457.81 | 27.66 | 121944 | 23 | 24 | 35383717 | 19853017 | 14692594 | 18217151 | 19301913 | 18123587 | 18115915 | 20734559 | 22049653 | 16426158 | 15553814 | 14160003 | 18898529 | 19240839 | 17995841 |
| E9Q634 | Myo1e | Unconventional myosin-Ie Mus musculus | 1357.31 | 32.25 | 126818 | 21 | 26 | 33731409 | 33920177 | 35435257 | 31804043 | 30204529 | 34960029 | 45135253 | 36068857 | 37696557 | 37441933 | 36075349 | 37220225 | 30034521 | 31792161 | 37623081 |
| P70248 | Myo1f | Unconventional myosin-If Mus musculus | 612.94 | 12.47 | 125947 | 5 | 10 | 17446385 | 18365493 | 20330939 | 17379969 | 17404041 | 17881721 | 21518267 | 21414247 | 20865487 | 17954833 | 20501909 | 19532381 | 18227569 | 20962405 | 22259079 |
| Q5SUA5 | Myo1g | Unconventional myosin-Ig Mus musculus | 345.99 | 9.47 | 117227 | 7 | 7 | 5781062.5 | 4118383 | 3759806.75 | 2363663.5 | 3330902.5 | 4019308.5 | 2260571.5 | 3411717.5 | 5383044 | 1074628.88 | 2376298.75 | 3072088.75 | 2738005.75 | 3986157.25 | 3177312 |
| Q99104 | Myo5a | Unconventional myosin-Va Mus musculus | 229.53 | 3.72 | 215538 | 5 | 5 | 1932483.25 | 1126682.25 | 1334531.75 | 1824382.25 | 737221.5 | 2085048.38 | 1712326.62 | 2045467.25 | 1514152.75 | 1337784.25 | 1769727.88 | 2318938.25 | 1174769.12 | 1908650.62 | 1173686.5 |
| Q63358 | Myo9b | Unconventional myosin-IXb Rattus norvegicus | 384.83 | 5.35 | 225037 | 7 | 7 | 1958592.38 | 2189105.75 | 2147990.75 | 1461775.75 | 2002667.5 | 2843152 | 1636450.62 | 3239462.25 | 2390385.25 | 1580061 | 2036236.25 | 1947830.12 | 2930629.25 | 1594579.25 | 1846895.38 |
| Q69ZN7 | Myof | Myoferlin Mus musculus | 889.94 | 12.65 | 233324 | 18 | 19 | 19943977 | 22640817 | 24982759 | 14619662 | 16671501 | 24429891 | 13748687 | 24146349 | 31444101 | 16480558 | 18455535 | 17997463 | 19048553 | 19375953 | 20479159 |
| Q80UM3 | Naa15 | N-alpha-acetyltransferase 15, NatA auxiliary subunit Mus musculus | 805.53 | 21.62 | 100961 | 17 | 17 | 26797527 | 30386665 | 28246201 | 25291909 | 28832999 | 30307547 | 24758117 | 30610747 | 35741469 | 25954925 | 28514317 | 25664019 | 27661367 | 31208631 | 27780277 |
| Q8BWZ3 | Naa25 | N-alpha-acetyltransferase 25, NatB auxiliary subunit Mus musculus | 298.5 | 10.29 | 111708 | 7 | 7 | 7524617.5 | 7141157 | 8268304.5 | 6840852 | 6667260 | 6287126.5 | 5543761.5 | 7184932.5 | 9530668 | 6552742 | 7970114.5 | 7910983 | 6300373.5 | 8743658 | 7564413 |
| Q8VE10 | Naa40 | N-alpha-acetyltransferase 40 Mus musculus | 76.46 | 12.24 | 27229 | 2 | 2 | 1771650.12 | 1859755.5 | 1967480.38 | 1627302.75 | 1980099.25 | 1868312.75 | 1476187.5 | 2100908 | 2435407.75 | 1622323.75 | 1851461.5 | 1942988.62 | 1864574.5 | 1690895.25 | 1935354.25 |
| Q6PGB6 | Naa50 | N-alpha-acetyltransferase 50 Mus musculus | 413.5 | 46.15 | 19414 | 9 | 9 | 8352918.5 | 7574797.5 | 10420285 | 6676337 | 6330412 | 7159823 | 7510619 | 10088599 | 10222564 | 5967632.5 | 9599028 | 11221631 | 6091327 | 8832491 | 10311465 |
| Q9D7V9 | Naaa | N-acylethanolamine-hydrolyzing acid amidase Mus musculus | 120.6 | 6.63 | 40075 | 2 | 2 | 1150586.12 | 1279250.5 | 1134581.25 | 614297.88 | 1006296.25 | 1043552.62 | 806134.69 | 1127589.12 | 1054890 | 936892.12 | 1243839.12 | 906577.81 | 1053686.62 | 981037.31 | 707908.62 |
| Q3SWT1 | Nabp2 | SOSS complex subunit B1 Rattus norvegicus | 155.87 | 17.06 | 22377 | 2 | 2 | 590464.69 | 490507.22 | 419391.28 | 516070.09 | 181658.59 | 607642.44 | 377223.66 | 358981.44 | 298463.41 | 450616.94 | 276430.44 | 335512.44 | 422607.88 | 336463.16 | 352699.06 |
| P70670 | Naca | Nascent polypeptide-associated complex subunit alpha, muscle-specific form Mus musc | 450.4 | 3.2 | 220499 | 6 | 6 | 16618589 | 2569047.5 | 20120981 | 21405269 | 23384415 | 21061647 | 27509769 | 20355949 | 19788319 | 26388099 | 24760885 | 27221315 | 24861289 | 20408983 | 29466805 |
| P58058 | Nadk | NAD kinase Mus musculus | 117.21 | 7.29 | 48597 | 2 | 2 | 1 | 528761.81 | 1 | 859106.5 | 634041.38 | 741606.69 | 1285249.5 | 786674.69 | 655745.25 | 761251.25 | 572016.38 | 1 | 1 | 810098.19 | 877215.12 |
| Q8VBW6 | Nae1 | NEDD8-activating enzyme E1 regulatory subunit Mus musculus | 551.95 | 32.21 | 60274 | 10 | 10 | 1727273 | 6163646 | 5096017.5 | 7515592.5 | 7532802 | 7891399 | 7319317 | 15850521 | 6879275 | 6958896 | 9237315 | 7704216 | 6289983 | 8334077 | 8297414.5 |
| Q9QWR8 | Naga | Alpha-N-acetylgalactosaminidase Mus musculus | 613.16 | 36.63 | 47235 | 12 | 12 | 29794001 | 21989381 | 21418603 | 26626067 | 23937645 | 23951231 | 24366701 | 32317445 | 23852023 | 24589385 | 26402205 | 22897709 | 25825721 | 35017021 | 18893269 |
| Q9QZ08 | Nagk | N-acetyl-D-glucosamine kinase Mus musculus | 217.37 | 14.29 | 37268 | 5 | 5 | 8062789 | 6713548 | 6884579 | 6825881.5 | 7194295 | 6200336.5 | 4328567 | 7022307.5 | 8154922 | 4084050.5 | 6073564 | 6354214 | 5897616.5 | 6994232.5 | 5720382.5 |
| Q8BJ48 | Nagpa | N-acetylglucosamine-1-phosphodiester alpha-N-acetylglucosaminidase Mus musculus | 98.94 | 3.09 | 56044 | 1 | 1 | 2048881.38 | 2216414.75 | 2235319.5 | 1364360.88 | 1683257.25 | 1401361.38 | 1499742.25 | 1553627.88 | 1776869.75 | 1557900.315 | 1569706.25 | 1973821 | 2124168.75 | 1588607.62 | 2054517.88 |
| Q9JIB3 | Naip7 | Baculoviral IAP repeat-containing protein 1g Mus musculus | 48.44 | 1 | 159664 | 1 | 1 | 238930.83 | 261196.81 | 289199.44 | 216950.89 | 175438.23 | 225246.94 | 208195.7 | 415120.66 | 280713.47 | 240604.67 | 363072.94 | 232289.33 | 176995.45 | 310773.72 | 303314.53 |
| Q99KQ4 | Nampt | Nicotinamide phosphoribosyltransferase Mus musculus | 731.17 | 32.38 | 55447 | 13 | 13 | 30820605 | 24699147 | 27517631 | 22974717 | 25876253 | 26737619 | 31666981 | 31383607 | 28360365 | 27479149 | 24159849 | 27318265 | 29055137 | 30535261 | 28077785 |
| Q5M969 | Nanp | N-acylneuraminate-9-phosphatase Rattus norvegicus | 83.18 | 7.26 | 27741 | 1 | 1 | 564400.44 | 1 | 1 | 1 | 388258.62 | 1 | 1 | 1 | 1 | 618128.44 | 461203.91 | 388203.91 | 1 | 1 | 719529.44 |
| Q99J77 | Nans | Sialic acid synthase Mus musculus | 469.57 | 40.11 | 40024 | 9 | 9 | 13509025 | 10797360 | 11397695 | 12177869 | 13164238 | 11711229 | 15495305 | 10982580 | 11423910 | 15235578 | 12778183 | 12103586 | 12374812 | 13181250 | 12085517 |
| P28656 | Nap1l1 | Nucleosome assembly protein 1-like 1 Mus musculus | 477.44 | 29.67 | 45345 | 7 | 8 | 19945157 | 16459026 | 16837247 | 16041722 | 14483890 | 18343643 | 18327599 | 21418165 | 23223669 | 19615713 | 20760789 | 24900105 | 16099575 | 24397691 | 24128023 |
| Q78ZA7 | Nap1l4 | Nucleosome assembly protein 1-like 4 Mus musculus | 735.31 | 49.6 | 42679 | 11 | 12 | 20299347 | 22217325 | 19984163 | 19435575 | 21151159 | 22721809 | 28214701 | 22337447 | 20613611 | 21809107 | 20069511 | 22927211 | 21072621 | 22228017 | 26141353 |
| Q5PPG6 | Nap1l5 | Nucleosome assembly protein 1-like 5 Rattus norvegicus 2 | 35.42 | 4.52 | 16858 | 1 | 1 | 2030543 | 1892679.25 | 1967458.75 | 2355257 | 1713276.25 | 1562184.62 | 1458416.5 | 1537442.88 | 1685425.25 | 1486442.25 | 1721551.62 | 1277105.88 | 1430691.25 | 1837019.5 | 1297264.25 |
| P54921 | Napa | Alpha-soluble NSF attachment protein Rattus norvegicus | 501.16 | 38.31 | 33193 | 9 | 9 | 20515501 | 19961555 | 22465731 | 19491621 | 22112343 | 20292081 | 22808813 | 22449261 | 23533235 | 22529017 | 20221521 | 19799385 | 21441867 | 22332789 | 20625117 |
| Q9CWZ7 | Napg | Gamma-soluble NSF attachment protein Mus musculus | 301.23 | 20.83 | 34732 | 6 | 6 | 14751396 | 14937638 | 14208314 | 11512289 | 12807202 | 13285316 | 14091431 | 18414487 | 13377137 | 11714127 | 12146953 | 14137447 | 12179302 | 14055446 | 12034240 |
| O09043 | Napsa | Napsin-A Mus musculus | 217.39 | 18.62 | 45544 | 4 | 4 | 8752844 | 10914111 | 8979887 | 9367902 | 8667265 | 7870043 | 8075980 | 9791338 | 8612572 | 8252161 | 6165996.5 | 6857027 | 8400214 | 10566349 | 5748153 |
| Q8BP47 | Nars | Asparagine--tRNA ligase, cytoplasmic Mus musculus | 1235.19 | 42.04 | 64279 | 19 | 19 | 57360417 | 58800577 | 63115161 | 62486093 | 61635505 | 63478877 | 68530929 | 62602749 | 65115601 | 59085489 | 56784737 | 63482305 | 54749661 | 57111721 | 78140361 |
| Q99MD9 | Nasp | Nuclear autoantigenic sperm protein Mus musculus | 853.85 | 28.85 | 83954 | 12 | 12 | 17193899 | 22030449 | 23544573 | 18471017 | 22064153 | 20320479 | 22374751 | 22680553 | 23143349 | 19786163 | 24232521 | 22993941 | 20763949 | 22707957 | 23510937 |
| Q8K224 | Nat10 | RNA cytidine acetyltransferase Mus musculus | 588.29 | 16.41 | 115419 | 11 | 11 | 3630593.75 | 2518550 | 3124602.75 | 3726393 | 2927654 | 3551784 | 2561862 | 3104384 | 2660527.5 | 3372636.25 | 3408210.25 | 3235300.25 | 3973482 | 2090993 | 2707353 |
| Q9CZ42 | Naxd | ATP-dependent (S)-NAD(P)H-hydrate dehydratase Mus musculus | 95.41 | 8.45 | 36717 | 2 | 2 | 1031333.31 | 1133381.38 | 1491334.25 | 1585236.38 | 1723380.75 | 1371223.38 | 1732870.38 | 1181604.25 | 1024400.12 | 1974308.75 | 1244064.5 | 1288382.5 | 1830541.38 | 1028861.44 | 1825941.25 |
| Q9EPN1 | Nbea | Neurobeachin Mus musculus | 40.57 | 0.41 | 326743 | 1 | 1 | 1 | 403126.84 | 412973.84 | 1 | 154410.02 | 1 | 1 | 1 | 536142.75 | 1 | 1 | 160964.72 | 1 | 171126.81 | 1 |
| Q6ZQA0 | Nbeal2 | Neurobeachin-like protein 2 Mus musculus | 204.87 | 2.59 | 301924 | 5 | 5 | 552293.81 | 632431.62 | 866172.62 | 536646.12 | 296409.5 | 336494.69 | 653408.88 | 424745.75 | 515609.56 | 575197.88 | 572131.25 | 574300.69 | 1203880 | 513091.25 | 715850.7975 |
| Q8K2Z4 | Ncapd2 | Condensin complex subunit 1 Mus musculus | 831.87 | 17.17 | 155665 | 17 | 17 | 16534728 | 14165364 | 16248981 | 12173057 | 12275724 | 14590014 | 12342837 | 15563107 | 16540992 | 11817647 | 17475097 | 16834961 | 12288043 | 16557811 | 14413225 |
| Q8C156 | Ncaph | Condensin complex subunit 2 Mus musculus | 335.07 | 11.22 | 82303 | 5 | 5 | 1761884.62 | 2871634 | 1722349.75 | 1184029.75 | 2470834.25 | 2855149 | 1579366.88 | 2555982.5 | 1919802 | 1544288.62 | 3432220.75 | 1879194.38 | 2715397.75 | 3369783.5 | 1654734 |
| Q3UYV9 | Ncbp1 | Nuclear cap-binding protein subunit 1 Mus musculus | 480.77 | 13.04 | 91927 | 7 | 7 | 5706985 | 4680056 | 3885938.75 | 3584168 | 3357126 | 4697062 | 3687778.75 | 4720909 | 5424302 | 3314329 | 4622160 | 4568405 | 4671207.5 | 4789098 | 4617401.5 |
| Q8BLF1 | Nceh1 | Neutral cholesterol ester hydrolase 1 Mus musculus | 759.72 | 42.16 | 45740 | 11 | 11 | 39716793 | 42137821 | 51028613 | 47627873 | 41299869 | 39123021 | 46520077 | 42684525 | 39880349 | 43449409 | 42859117 | 48522205 | 45966045 | 37418073 | 57594229 |
| Q09014 | Ncf1 | Neutrophil cytosol factor 1 Mus musculus | 735.9 | 43.33 | 44667 | 13 | 13 | 33010849 | 37826905 | 34606425 | 33224639 | 42468509 | 37518105 | 33192217 | 36535069 | 42572861 | 34807405 | 44271641 | 38403769 | 40613373 | 37520145 | 39310641 |
| O70145 | Ncf2 | Neutrophil cytosol factor 2 Mus musculus | 1100.57 | 50.48 | 59485 | 21 | 21 | 69790921 | 61107409 | 54740737 | 56351425 | 58903945 | 54505757 | 50419097 | 53201425 | 50333133 | 43332877 | 58715913 | 54120005 | 58673849 | 55727033 | 53169381 |
| P97369 | Ncf4 | Neutrophil cytosol factor 4 Mus musculus | 1302.8 | 59.88 | 38707 | 19 | 19 | 81977657 | 88707193 | 93146001 | 82076441 | 86537425 | 78501305 | 64930977 | 83264225 | 90215337 | 69265785 | 84280529 | 86067985 | 77158385 | 91006489 | 79301585 |
| Q8K1X4 | Nckap1l | Nck-associated protein 1-like Mus musculus | 1623.37 | 32.72 | 128905 | 25 | 26 | 41108977 | 46357361 | 44054669 | 39256133 | 43858401 | 43130249 | 40746561 | 47510421 | 54999697 | 43869125 | 44223705 | 48275745 | 43024349 | 43062145 | 46780773 |
| P09405 | Ncl | Nucleolin Mus musculus | 2657.22 | 41.58 | 76723 | 41 | 43 | 512448801 | 493881793 | 545296833 | 505632737 | 517477665 | 522771777 | 505673281 | 590702081 | 596260609 | 534518209 | 627965249 | 647741057 | 518995009 | 544230337 | 680891457 |
| Q5XIA1 | Ncln | Nicalin Rattus norvegicus | 196.73 | 7.28 | 62992 | 4 | 4 | 5398326.5 | 5630011.5 | 5604782 | 4915900.5 | 7233276.5 | 4679726.5 | 5718171 | 6838591.5 | 6555756.5 | 6540872.5 | 5064213.5 | 4589628.5 | 6329046.5 | 5409497 | 3790156.25 |
| P57716 | Ncstn | Nicastrin Mus musculus | 874.54 | 25.99 | 78492 | 13 | 13 | 66070841 | 61829253 | 59012321 | 56884313 | 61278481 | 52296193 | 41820505 | 59430529 | 67205633 | 38452045 | 49731573 | 49054261 | 53749849 | 58862705 | 41054609 |
| Q8VCB1 | Ndc1 | Nucleoporin NDC1 Mus musculus | 108.12 | 6.24 | 75409 | 2 | 2 | 1 | 569035.88 | 1 | 1142505 | 474074.88 | 1 | 419277.38 | 533280.44 | 1197301.88 | 1 | 445003.78 | 670941.19 | 1665263.5 | 1225471.75 | 704155.88 |
| Q78PB6 | Ndel1 | Nuclear distribution protein nudE-like 1 Rattus norvegicus | 70.68 | 3.48 | 38366 | 1 | 1 | 413963 | 358324.19 | 535885.44 | 692874.75 | 438339.22 | 419005.88 | 659004.06 | 343841.88 | 545952.56 | 645122.69 | 726294.69 | 459239.41 | 396372.41 | 443903.34 | 426555.59 |
| Q62433 | Ndrg1 | Protein NDRG1 Mus musculus | 402.01 | 32.74 | 43009 | 7 | 7 | 12050821 | 17884593 | 12918770 | 12185809 | 17734777 | 14966141 | 24916441 | 18337219 | 15743440 | 19967781 | 20175771 | 18640775 | 19921097 | 20909529 | 20497007 |
| Q6AYR2 | Ndrg3 | Protein NDRG3 Rattus norvegicus | 61.03 | 5.07 | 41507 | 1 | 1 | 1 | 1 | 1 | 1 | 1 | 351664.03 | 541392.81 | 489861.38 | 462837.0475 | 468429.97 | 1 | 1 | 576094 | 510622.81 | 1 |

| | | | | | | | | | | | | | | | | | | | | | | |
|---|---|---|---|---|---|---|---|---|---|---|---|---|---|---|---|---|---|---|---|---|---|---|
| Q99LC3 | Ndufa10 | NADH dehydrogenase [ubiquinone] 1 alpha subcomplex subunit 10, mitochondrial Mus | 384.06 | 29.01 | 40603 | 6 | 6 | 7593042.5 | 6444211 | 6788890 | 5793231.5 | 6718288 | 6801233 | 4951883 | 6716888.5 | 6738045 | 4700996 | 6431244.5 | 6337336 | 6084051.5 | 7522786 | 6514136.5 |
| Q9D8B4 | Ndufa11 | NADH dehydrogenase [ubiquinone] 1 alpha subcomplex subunit 11 Mus musculus | 59.19 | 7.8 | 14982 | 1 | 1 | 1637960.25 | 1386063.25 | 1465539.38 | 1019381.25 | 1312374.5 | 1261003.88 | 1034737.88 | 1456406.38 | 1712311.12 | 1092315.12 | 1105244.12 | 1478673.75 | 1323287.75 | 1500728.5 | 1044742.69 |
| Q7TMF3 | Ndufa12 | NADH dehydrogenase [ubiquinone] 1 alpha subcomplex subunit 12 Mus musculus | 139.85 | 17.93 | 17086 | 3 | 3 | 1921479.5 | 1833003.38 | 851830.31 | 1647268.38 | 1864334.75 | 710522.06 | 985509.25 | 1734850 | 1527220.75 | 1666251.75 | 1737052 | 1446581.88 | 2042878.75 | 2036926.5 | 1270189.62 |
| Q9ERS2 | Ndufa13 | NADH dehydrogenase [ubiquinone] 1 alpha subcomplex subunit 13 Mus musculus | 88.83 | 16.67 | 16860 | 2 | 2 | 4614131 | 4919397 | 3511846 | 3350242.75 | 3415066.5 | 1912128.62 | 4083673 | 4159653.5 | 4371077 | 3481777 | 3021598 | 3553918.5 | 4058047.5 | 4709128 | 2808167 |
| Q9CQ75 | Ndufa2 | NADH dehydrogenase [ubiquinone] 1 alpha subcomplex subunit 2 Mus musculus | 84.03 | 14.14 | 10916 | 1 | 1 | 2463771.5 | 3071521.25 | 2938854.5 | 3102648.5 | 3003064.75 | 2451165 | 2893777.75 | 2619348.25 | 2506425 | 3222139.5 | 2713392.75 | 2767822 | 2940098 | 2975330.25 | 2536009.25 |
| Q62425 | Ndufa4 | Cytochrome c oxidase subunit NDUFA4 Mus musculus | 180.38 | 68.29 | 9327 | 4 | 4 | 24869155 | 20107309 | 17811985 | 16011653 | 17451605 | 16824603 | 15451265 | 17895747 | 18815299 | 14645603 | 13871896 | 15618972 | 17945335 | 16440467 | 12764875 |
| Q9CPP6 | Ndufa5 | NADH dehydrogenase [ubiquinone] 1 alpha subcomplex subunit 5 Mus musculus | 64.98 | 17.24 | 13360 | 1 | 1 | 703088.19 | 303784.12 | 667029.88 | 1 | 1 | 558158.88 | 284163.41 | 602679.19 | 1097706.12 | 276389.69 | 309743.38 | 812541.56 | 324640.56 | 849835.38 | 431696.38 |
| Q9DCJ5 | Ndufa8 | NADH dehydrogenase [ubiquinone] 1 alpha subcomplex subunit 8 Mus musculus | 115.78 | 24.42 | 19992 | 3 | 3 | 2939968 | 2200855.5 | 2275720 | 2544502.25 | 2041394.75 | 931514.25 | 1775843.75 | 2104614.5 | 1826504.25 | 2335788.5 | 2490212.5 | 1941957.5 | 2234249.5 | 1918498.75 | 1687831.25 |
| Q9DC69 | Ndufa9 | NADH dehydrogenase [ubiquinone] 1 alpha subcomplex subunit 9, mitochondrial Mus m | 551 | 36.87 | 42525 | 9 | 9 | 17553387 | 7885362 | 8357411 | 12999701 | 12867800 | 4230188.5 | 9092818 | 8788841 | 8189734 | 13074165 | 8390773 | 7531643.5 | 13428258 | 8079425.5 | 12684996 |
| Q9DCS9 | Ndufb10 | NADH dehydrogenase [ubiquinone] 1 beta subcomplex subunit 10 Mus musculus | 237.4 | 31.25 | 21024 | 4 | 4 | 3483242.5 | 3497644.5 | 3301506 | 3235374 | 4771064 | 2778629 | 862451.06 | 2527288.75 | 3343823.25 | 3880597 | 3610570.5 | 3130773 | 2979130.25 | 3448124.5 | 2279444 |
| O09111 | Ndufb11 | NADH dehydrogenase [ubiquinone] 1 beta subcomplex subunit 11, mitochondrial Mus m | 37.18 | 11.26 | 17444 | 1 | 1 | 1 | 1 | 1 | 1 | 835302.44 | 1 | 1 | 1 | 1 | 1 | 1103796.88 | 543035.06 | 1302110 | 1082863.485 | 1382512 |
| Q9CQZ6 | Ndufb3 | NADH dehydrogenase [ubiquinone] 1 beta subcomplex subunit 3 Mus musculus | 49.58 | 16.35 | 11692 | 1 | 1 | 478849.84 | 939127.62 | 1228604 | 1 | 1 | 1 | 1 | 248453.27 | 1222054.62 | 1 | 666415.88 | 960678.81 | 1 | 362576.53 | 1 |
| Q9CQH3 | Ndufb5 | NADH dehydrogenase [ubiquinone] 1 beta subcomplex subunit 5, mitochondrial Mus mu | 95.5 | 12.7 | 21710 | 2 | 2 | 1 | 1 | 1 | 1 | 1 | 1 | 1 | 332189.25 | 1 | 1 | 1 | 1 | 1 | 1 | 1 |
| Q9CQ54 | Ndufc2 | NADH dehydrogenase [ubiquinone] 1 subunit C2 Mus musculus | 41.04 | 10.83 | 14164 | 1 | 1 | 1530846.62 | 1188782.38 | 1155049.5 | 983285.12 | 1004454.69 | 1613328.25 | 665314.38 | 1182471.12 | 1202890.62 | 860145.12 | 945709.75 | 969569.81 | 917689.19 | 975707.5 | 838160.94 |
| Q91VD9 | Ndufs1 | NADH-ubiquinone oxidoreductase 75 kDa subunit, mitochondrial Mus musculus | 822.8 | 32.6 | 79777 | 15 | 15 | 17999677 | 19507047 | 19924165 | 19055877 | 18308039 | 16905187 | 22426259 | 18905651 | 19791435 | 20570611 | 16016000 | 18989651 | 19521053 | 18470229 | 20553465 |
| Q91WD5 | Ndufs2 | NADH dehydrogenase [ubiquinone] iron-sulfur protein 2, mitochondrial Mus musculus | 481.97 | 19.65 | 52626 | 7 | 7 | 12808319 | 15790347 | 14714339 | 14247455 | 13583423 | 13839916 | 11661773 | 15733393 | 16733032 | 13484361 | 12844472 | 15228914 | 10523319 | 14241449 | 13010371 |
| Q9DCT2 | Ndufs3 | NADH dehydrogenase [ubiquinone] iron-sulfur protein 3, mitochondrial Mus musculus | 441.05 | 33.08 | 30149 | 7 | 7 | 11549762 | 10014481 | 10929866 | 10903163 | 10815180 | 10859460 | 8732876 | 11599679 | 11499027 | 10191845 | 8231620 | 9858302 | 8792021 | 10124517 | 8114924.5 |
| Q9DC70 | Ndufs7 | NADH dehydrogenase [ubiquinone] iron-sulfur protein 7, mitochondrial Mus musculus | 360.71 | 30.8 | 24683 | 6 | 6 | 9863442 | 10096206 | 9947674 | 9556809 | 11060647 | 10005982 | 11495370 | 10159458 | 10186697 | 10813249 | 11193240 | 10473590 | 11453582 | 9965572 | 11056017 |
| Q8K3J1 | Ndufs8 | NADH dehydrogenase [ubiquinone] iron-sulfur protein 8, mitochondrial Mus musculus | 221.95 | 17.92 | 24038 | 3 | 3 | 1898490.5 | 2204512 | 1916932.75 | 1833128.62 | 2253961.75 | 1837062 | 1291086.38 | 696583.12 | 2433068.5 | 829704.31 | 1263139.75 | 2050221.25 | 1710265.5 | 2211888 | 1838653.25 |
| Q91YT0 | Ndufv1 | NADH dehydrogenase [ubiquinone] flavoprotein 1, mitochondrial Mus musculus | 482.5 | 37.5 | 50834 | 10 | 10 | 14304968 | 9603685 | 7987856.5 | 10541915 | 11027295 | 12456700 | 8248384.5 | 5738244 | 11164302 | 9171232 | 7566310 | 8408803 | 11585326 | 11095663 | 7658936 |
| Q9D6J6 | Ndufv2 | NADH dehydrogenase [ubiquinone] flavoprotein 2, mitochondrial Mus musculus | 83.11 | 8.47 | 27285 | 2 | 2 | 2964811.5 | 3032189.75 | 2951593.938 | 2531769.5 | 3277605 | 2173704.25 | 2603983.25 | 2922741.5 | 3545565.25 | 3146395.5 | 1922862 | 2776270 | 2898458 | 2620104 | 2446905.5 |
| Q9CR95 | Necap1 | Adaptin ear-binding coat-associated protein 1 Mus musculus | 227.34 | 26.55 | 29639 | 5 | 5 | 4157405.25 | 4732588.5 | 5920900 | 4974133.5 | 2267945.25 | 5529031 | 8082748 | 6744798 | 4729301.5 | 5864685 | 2674031.5 | 6517323.5 | 3879512 | 5358924 | 5973501.5 |
| Q9D1J1 | Necap2 | Adaptin ear-binding coat-associated protein 2 Mus musculus | 270.8 | 31.95 | 28598 | 2 | 5 | 3710688 | 4054495.5 | 3515519.5 | 3933436.5 | 4115200 | 5116532 | 5677140.5 | 4120635.25 | 3991092.5 | 5271131 | 4688011 | 5291598 | 4013848.5 | 4674000 | 5776335.5 |
| P29595 | Nedd8 | NEDD8 Mus musculus | 143.7 | 34.57 | 8972 | 3 | 3 | 2820473.75 | 2314128.5 | 2063765 | 2557613.5 | 2498568.5 | 2160452.25 | 3627895.5 | 1858144.25 | 1665597.25 | 3059152.5 | 2056326.62 | 2876290.5 | 2822677 | 2114297.5 | 3126590.5 |
| D3ZBE5 | Nek7 | Serine/threonine-protein kinase Nek7 Rattus norvegicus | 34.07 | 2.65 | 34529 | 1 | 1 | 1883261.25 | 1644848.5 | 1576378 | 1386857.75 | 1375839.62 | 1239585.62 | 2042190.12 | 1504383.88 | 1286564.12 | 1875685.75 | 1737880.75 | 1809556.75 | 1380434.5 | 1523666.12 | 1806219 |
| Q8K1R7 | Nek9 | Serine/threonine-protein kinase Nek9 Mus musculus | 255 | 5.89 | 107143 | 5 | 5 | 12113090 | 7071253 | 9067258 | 10640010 | 10020693 | 8333070 | 16405935 | 7649468.5 | 4961077 | 13459911 | 11522827 | 5414227 | 8086280 | 2936357.25 | 9766945 |
| Q8C4Y3 | Nelfb | Negative elongation factor B Mus musculus | 75.63 | 5.34 | 65636 | 2 | 2 | 1081040.62 | 867708.44 | 1432506.62 | 402737.62 | 398659.06 | 834390.25 | 562943.69 | 1936995.38 | 1684132.12 | 711364.94 | 1295936.62 | 1864630.62 | 501085.78 | 1743841.88 | 1647410 |
| Q922L6 | Nelfcd | Negative elongation factor D Mus musculus | 107.55 | 5.25 | 66276 | 2 | 2 | 1230443.75 | 1123769.88 | 1206099 | 1379372.75 | 1355226.75 | 1138841.88 | 1923525 | 1390953 | 1189703.38 | 1083839 | 1362231.25 | 1542039.75 | 1271144.5 | 1454931.5 | 1676711.38 |
| Q8CCP0 | Nemf | Nuclear export mediator factor Nemf Mus musculus | 38.68 | 0.66 | 121188 | 1 | 1 | 484501.22 | 500323.765 | 477963.78 | 596490 | 442340.06 | 418161.22 | 1379068.75 | 1 | 1 | 758709.38 | 1 | 770774.88 | 1 | 1 | 925531.19 |
| Q6P5H2 | Nes | Nestin Mus musculus | 2639.19 | 34.39 | 207124 | 46 | 46 | 79465009 | 89528201 | 88990681 | 93163257 | 81493441 | 85294497 | 114059977 | 87703145 | 84792841 | 92678281 | 90391873 | 88822529 | 84090057 | 80549513 | 97401273 |
| O35657 | Neu1 | Sialidase-1 Mus musculus | 37.63 | 2.2 | 44591 | 1 | 1 | 1 | 1 | 1 | 1 | 1 | 301409.75 | 326632.44 | 286419.97 | 171354.48 | 399850.75 | 1 | 1 | 331133.06 | 1 | 1 |
| P97526 | Nf1 | Neurofibromin Rattus norvegicus | 65.62 | 0.78 | 317083 | 1 | 1 | 279746.625 | 297530.44 | 277005.28 | 275340.78 | 269110 | 187287.83 | 225938.3875 | 238583.58 | 238894.75 | 238987.39 | 242400.69 | 227861.19 | 1 | 1 | 222089.89 |
| Q60591 | Nfatc2 | Nuclear factor of activated T-cells, cytoplasmic 2 Mus musculus | 39.21 | 0.97 | 100020 | 1 | 1 | 668746.62 | 1 | 763157.81 | 837551.31 | 1001204.44 | 589921.38 | 828593 | 544047.5 | 482650.06 | 915513.62 | 740621.31 | 1046204.44 | 862761.31 | 672489.38 | 903337.69 |
| P25799 | Nfkb1 | Nuclear factor NF-kappa-B p105 subunit Mus musculus | 417.44 | 10.71 | 105615 | 9 | 9 | 7976543 | 6919470.5 | 7398418 | 7659858.5 | 7483320 | 6989426 | 7913558 | 6895425 | 6919832 | 7030083 | 6369846 | 7230730 | 7209320 | 7103661 | 7140083 |
| Q9WTK5 | Nfkb2 | Nuclear factor NF-kappa-B p100 subunit Mus musculus | 402.2 | 13.35 | 96832 | 8 | 8 | 1745666.38 | 1787161 | 1787656.12 | 1938950.5 | 2539147.5 | 2193523 | 2537529.75 | 2386991.75 | 2369237.5 | 2981077.75 | 2386242.5 | 2772368.5 | 2397434.25 | 2467902.75 | 3206902.5 |
| O54910 | Nfkbie | NF-kappa-B inhibitor epsilon Mus musculus | 66.86 | 3.3 | 39200 | 1 | 1 | 547031.81 | 576346.38 | 607238.94 | 826548.88 | 678006.31 | 744547.75 | 564732.88 | 867905.69 | 869913.12 | 796918.19 | 871475.31 | 767873.88 | 804633.62 | 749348.88 | 593695.81 |
| Q9QZ23 | Nfu1 | NFU1 iron-sulfur cluster scaffold homolog, mitochondrial Mus musculus | 172.78 | 17.25 | 28568 | 3 | 3 | 7487796 | 4663643 | 8237416 | 8452325 | 9131911 | 9425617 | 8330752.5 | 9055488 | 7921005 | 7775651 | 7743966 | 7533619 | 8858501 | 8868390 | 6782584 |
| P63139 | Nfyb | Nuclear transcription factor Y subunit beta Mus musculus | 53.87 | 10.14 | 22787 | 1 | 1 | 130565.93 | 226263.86 | 158491.7 | 181244.34 | 240914.52 | 177173.08 | 107263.48 | 214744.92 | 121122.02 | 202445.47 | 212577.47 | 170715.7 | 183144.55 | 226353.12 | 138469.7 |
| P70353 | Nfyc | Nuclear transcription factor Y subunit gamma Mus musculus | 100.51 | 7.46 | 37254 | 2 | 2 | 483421.32 | 322263.41 | 645538.25 | 392668.31 | 573215.31 | 506464 | 606847.45 | 736892.12 | 511578.06 | 672455.62 | 495310.69 | 779248 | 381297.72 | 803853.5 | 525476.38 |
| Q9DB96 | Ngdn | Neuroguidin Mus musculus | 67.49 | 5.71 | 35659 | 1 | 1 | 402678.41 | 253262.73 | 370786.69 | 345428.28 | 240595.23 | 291265.47 | 299457.06 | 247946.31 | 307271.545 | 390417.34 | 253856.44 | 393611.56 | 1 | 1 | 1 |
| Q8BZW8 | Nhlrc2 | NHL repeat-containing protein 2 Mus musculus | 38.12 | 1.66 | 78430 | 1 | 1 | 716599.81 | 273726.16 | 923782.62 | 975982.38 | 940498.81 | 806059.12 | 1339776.62 | 378431.88 | 629871.81 | 1098909.88 | 1132421.5 | 715103.44 | 967853 | 618888.94 | 905200.44 |
| Q8CCH2 | Nhlrc3 | NHL repeat-containing protein 3 Mus musculus | 485.16 | 52.74 | 38195 | 11 | 11 | 7638655 | 13080141 | 8901204 | 6809373.5 | 12766574 | 13166230 | 11268390 | 15742675 | 15222999 | 9950221 | 9858394 | 11584825 | 11103207 | 13737367 | 8658439 |
| Q9CRB2 | Nhp2 | H/ACA ribonucleoprotein complex subunit 2 Mus musculus | 136.73 | 16.99 | 17247 | 2 | 2 | 5380567 | 5823086.5 | 6158144 | 6858105 | 7214916 | 5642747 | 5602543.5 | 5549286 | 6540669.5 | 5882747 | 6385178.5 | 5405367.5 | 5834645.5 | 6153697.5 | 5704277 |
| Q9EQ80 | Nif3l1 | NIF3-like protein 1 Mus musculus | 202.23 | 17.29 | 41746 | 5 | 5 | 6159853.5 | 6868709 | 6970456 | 6814075 | 6238669.5 | 5311910 | 10643473 | 8370599 | 7484893 | 5661077 | 6751440 | 6154888.5 | 6799010 | 6959680 | 9658381 |
| Q91VE6 | Nifk | MKI67 FHA domain-interacting nucleolar phosphoprotein Mus musculus | 390.61 | 28.71 | 36265 | 7 | 7 | 3278893.5 | 2901934.5 | 5127383 | 4733415.5 | 4422989 | 1223862.5 | 8594990 | 4372007 | 4467610 | 6725061.5 | 3932597.75 | 5661116 | 2246650.75 | 3618053.5 | 3527652.75 |
| Q9CXK8 | Nip7 | 60S ribosome subunit biogenesis protein NIP7 homolog Mus musculus | 196.89 | 42.78 | 20452 | 5 | 5 | 6615078 | 5026366.5 | 4666064 | 5472882.5 | 4653561 | 4572171 | 4518198 | 5583236.5 | 9942044 | 3949102.5 | 5736600.5 | 4711905 | 4269444 | 5104197 | 5707637 |
| Q6KCD5 | Nipbl | Nipped-B-like protein Mus musculus | 106.75 | 0.93 | 315450 | 2 | 2 | 214160.67 | 1076155.88 | 1031369 | 1035851.31 | 914168.38 | 1017996.38 | 753780.75 | 895719.88 | 972484.62 | 798929.06 | 1181723.38 | 885187 | 906511.81 | 948122.94 | 851002.75 |
| O55125 | Nipsnap1 | Protein NipSnap homolog 1 Mus musculus | 49.61 | 3.17 | 33363 | 1 | 1 | 6229544 | 3744931.5 | 4046890.5 | 3863978.75 | 4515974.5 | 4165455.25 | 5076316.5 | 4642726 | 4009423.25 | 4489069 | 4864970.5 | 3672686.25 | 4207127.5 | 4216884.5 | 3914595 |
| O55126 | Nipsnap2 | Protein NipSnap homolog 2 Mus musculus | 94.84 | 9.96 | 32933 | 2 | 2 | 524071.69 | 492649.66 | 356549.66 | 453230.065 | 439649.25 | 589478.25 | 955871.38 | 431857.34 | 323601.44 | 860322.5 | 655657 | 759500.25 | 597890.62 | 740721.0925 | 949836.5 |
| Q9CQE1 | Nipsnap3b | Protein NipSnap homolog 3B Mus musculus | 128.67 | 13.77 | 28308 | 2 | 2 | 2797086 | 2111008.5 | 2202724 | 737958.81 | 2399881.5 | 2206846.5 | 2177944 | 2542619.25 | 3140485.25 | 1709531.38 | 565099.56 | 2299737.75 | 1711953 | 1879874.38 | 1638837.38 |
| Q80TM9 | Nisch | Nischarin Mus musculus | 422.25 | 5.15 | 175012 | 6 | 6 | 7754023 | 3385398 | 4322361.5 | 3333334.5 | 4967568 | 5242093 | 3102097 | 4595772 | 5939310 | 3361108.25 | 4129667 | 4509234 | 3746739.25 | 5323169 | 4665986 |
| Q8VDK1 | Nit1 | Deaminated glutathione amidase Mus musculus | 177.25 | 15.17 | 35705 | 3 | 3 | 2300366 | 2724879.5 | 3690259.5 | 3415774 | 2825136.25 | 3189676.5 | 3246968.5 | 3533508.5 | 2066809.75 | 2993412.75 | 2700977.5 | 3039626.25 | 2553111.75 | 3839204.75 | 3123026.5 |
| Q9JHW2 | Nit2 | Omega-amidase NIT2 Mus musculus | 414.17 | 32.61 | 30502 | 6 | 6 | 6482804 | 8996017 | 8459023 | 7126254 | 7951580 | 6811954 | 6462520.5 | 7136231 | 8327779 | 7220967 | 5782137.5 | 6794974 | 7805556.5 | 7756086 | 6487116 |
| P42676 | Nln | Neurolysin, mitochondrial Rattus norvegicus | 83.57 | 2.41 | 80254 | 2 | 2 | 1119613.5 | 946692.56 | 1929110.25 | 1672242.62 | 2221845 | 2175493.75 | 1933695.12 | 1855616.25 | 1803579.06 | 1249511.12 | 1192774.88 | 1760938.5 | 1282646.12 | 831545.06 | 1396165.5 |
| Q3TL44 | Nlrx1 | NLR family member X1 Mus musculus | 82.6 | 4.31 | 107831 | 2 | 2 | 673997.12 | 496556.88 | 627119 | 480419.31 | 634754.94 | 618632.88 | 333649.84 | 663953.88 | 615891.88 | 391222.25 | 763992.38 | 434099.59 | 455540.72 | 713802.19 | 594334.56 |
| Q99L48 | Nmd3 | 60S ribosomal export protein NMD3 Mus musculus | 165.05 | 9.54 | 57611 | 4 | 4 | 1698565 | 695013.06 | 1551777 | 491514.81 | 1111073 | 634840.5 | 1183239 | 561466.5 | 1909896.5 | 1445049.75 | 2306669.25 | 1207846.5 | 1224708.12 | 1207423.75 | 1438283.12 |
| P15532 | Nme1 | Nucleoside diphosphate kinase A Mus musculus | 480.54 | 73.03 | 17208 | 4 | 10 | 104322881 | 79151873 | 83748209 | 89986673 | 72114777 | 80399505 | 100144769 | 84830153 | 76839393 | 89852321 | 86376473 | 91046009 | 82882913 | 90800129 | 84105393 |
| Q01768 | Nme2 | Nucleoside diphosphate kinase B Mus musculus | 594.75 | 69.08 | 17363 | 3 | 10 | 146289873 | 108808457 | 121069241 | 123355457 | 99568065 | 107215089 | 135732465 | 116316593 | 103129881 | 120267193 | 116167641 | 123370689 | 112983713 | 123055897 | 116140353 |
| O35309 | Nmi | N-myc-interactor Mus musculus | 100.72 | 9.55 | 35236 | 2 | 2 | 2499665.5 | 2235807.25 | 2187643.75 | 2451689.75 | 2832923.5 | 1962217.38 | 1871991.88 | 2256231.5 | 2300748.25 | 2354640.25 | 2607871.25 | 2009405.5 | 2229253 | 2124551.25 | 1859865.88 |
| Q8K2T1 | Nmral1 | NmrA-like family domain-containing protein 1 Mus musculus | 327.72 | 27.18 | 34376 | 6 | 6 | 3122672.75 | 3768937.75 | 2600572 | 2868376 | 2902405.25 | 3169794 | 3823665.5 | 2854111 | 2751747.25 | 3819788.25 | 3928778.5 | 3694927 | 3322461.5 | 3341236.5 | 4270827.5 |
| O70310 | Nmt1 | Glycylpeptide N-tetradecanoyltransferase 1 Mus musculus | 804.42 | 38.51 | 56888 | 13 | 16 | 16814465 | 22363301 | 20927843 | 25617065 | 22983845 | 24918231 | 28951975 | 23902739 | 23509805 | 24261393 | 23504503 | 20954049 | 26520071 | 22495585 | 23504083 |
| O70311 | Nmt2 | Glycylpeptide N-tetradecanoyltransferase 2 Mus musculus | 225.93 | 10.02 | 60484 | 1 | 4 | 3513524.25 | 2774711.5 | 3552664 | 3528102 | 1920932.75 | 2632756 | 5154028 | 3579479 | 4059112 | 2933658.25 | 3169997 | 3654971.25 | 2791952.5 | 2806930.25 | 3931645 |
| Q61941 | Nnt | NAD(P) transhydrogenase, mitochondrial Mus musculus | 251.93 | 9.3 | 113838 | 7 | 7 | 5226674.5 | 5245070.5 | 7385412.5 | 7033046 | 9416817 | 8570942 | 7190501.5 | 6771302 | 4171281.5 | 7620573 | 6364900 | 7060352.5 | 6812786 | 5902545.5 | 7480722 |
| Q8BW10 | Nob1 | RNA-binding protein NOB1 Mus musculus | 112.9 | 7.94 | 45464 | 2 | 2 | 1 | 683194.5 | 1 | 1 | 1 | 1 | 1087016.5 | 1 | 523258.44 | 945681.44 | 348804.44 | 601615.08 | 535577.56 | 544064.38 | 978013.94 |
| Q8BJW5 | Nol11 | Nucleolar protein 11 Mus musculus | 286.94 | 10.24 | 80818 | 7 | 7 | 924443.62 | 2229335.5 | 2206846.5 | 2595892.5 | 2166788.25 | 1915659.25 | 991698.81 | 1505748.5 | 1531597.25 | 1858821.75 | 2476039.5 | 1422641.12 | 1907142.5 | 4910670.5 | 3745971 |
| Q8R5K4 | Nol6 | Nucleolar protein 6 Mus musculus | 141.63 | 4.08 | 129228 | 3 | 3 | 1330696.12 | 620115 | 258588.98 | 663051.7525 | 442806.91 | 654050.94 | 1477070.25 | 929194.2325 | 256767.12 | 1328888.62 | 321701.59 | 1138634.38 | 1194841.5 | 728821.38 | 532033.19 |
| Q3TZX8 | Nol9 | Polynucleotide 5'-hydroxyl-kinase NOL9 Mus musculus | 169.82 | 5.6 | 80840 | 3 | 3 | 628201.56 | 809801.6325 | 462534.03 | 1214701.25 | 933769.69 | 553225.69 | 348564.53 | 763891.5 | 2240042.75 | 976431.1175 | 656102.75 | 1335781.5 | 1173301.72 | 2170610.25 | 530712.38 |
| E9Q5C9 | Nolc1 | Nucleolar and coiled-body phosphoprotein 1 Mus musculus | 137.62 | 3.85 | 73698 | 3 | 3 | 74356.43 | 100289.9025 | 93275.03 | 72647.51 | 160880.64 | 1 | 1 | 1 | 1 | 1 | 108274.23 | 83402.91 | 1 | 1 | 1 |
| Q6GQT9 | Nomo1 | Nodal modulator 1 Mus musculus | 1342.23 | 33.69 | 133420 | 24 | 24 | 23861947 | 19982323 | 21988367 | 22763577 | 23571889 | 23468467 | 26281543 | 22974421 | 23995043 | 22851763 | 24331701 | 24177179 | 22455121 | 19701735 | 24886353 |
| Q99K48 | Nono | Non-POU domain-containing octamer-binding protein Mus musculus | 914.56 | 49.68 | 54541 | 15 | 16 | 23923795 | 25871525 | 28505361 | 29361659 | 24664091 | 27525685 | 40620805 | 25282673 | 21751725 | 36998117 | 33223803 | 32609763 | 26786301 | 17910305 | 40476989 |
| Q8R3N1 | Nop14 | Nucleolar protein 14 Mus musculus | 132.46 | 5.12 | 98769 | 3 | 3 | 1 | 399382.25 | 674298.75 | 271218.31 | 1 | 540614.31 | 523731.19 | 899769 | 730366.69 | 266057.78 | 835031.19 | 695569.06 | 184953.36 | 514895.97 | 537156.88 |
| Q9CPT5 | Nop16 | Nucleolar protein 16 Mus musculus | 79.68 | 8.99 | 21139 | 1 | 1 | 1 | 397197.81 | 604668.38 | 1 | 1 | 1 | 535261.38 | 485831.75 | 699981.62 | 1 | 322435.16 | 1 | 1 | 1 | 819655.44 |
| Q922K7 | Nop2 | Probable 28S rRNA (cytosine-C(5))-methyltransferase Mus musculus | 669.24 | 21.56 | 86752 | 13 | 13 | 10487542 | 7249033.5 | 16366221 | 8456655 | 8513229 | 9560548 | 15076805 | 9044955 | 12136524 | 10064253 | 13221681 | 14078812 | 11090011 | 9837487 | 11679268 |
| Q9D6Z1 | Nop56 | Nucleolar protein 56 Mus musculus | 1414.75 | 56.9 | 64464 | 24 | 24 | 47372017 | 40001381 | 43229321 | 44106425 | 45624717 | 41190553 | 43130237 | 41638401 | 43126309 | 42308357 | 44240957 | 42843873 | 41490201 | 43401693 | 41333813 |
| Q6DFW4 | Nop58 | Nucleolar protein 58 Mus musculus | 988 | 39.18 | 60343 | 15 | 17 | 29170077 | 31379703 | 29777683 | 27070727 | 30653845 | 28950653 | 28540093 | 28893901 | 30724005 | 27942231 | 28753737 | 30311065 | 27306191 | 29074853 | 30000673 |
| Q8BMC4 | Nop9 | Nucleolar protein 9 Mus musculus | 231.92 | 6.6 | 70046 | 3 | 3 | 639804.75 | 481929.97 | 787109.12 | 1003767.75 | 918819.75 | 1145871.12 | 704524.62 | 746982.75 | 846280.62 | 846693.25 | 774538.69 | 298746.25 | 781442 | 813957.12 | 1122892.25 |
| O35604 | Npc1 | NPC intracellular cholesterol transporter 1 Mus musculus | 240.09 | 5.56 | 142885 | 5 | 5 | 5466884 | 8042086 | 6392956.5 | 5789944.5 | 7449742 | 6926567.5 | 5366535 | 7151685.5 | 7385734 | 5888651 | 6856844 | 6492027 | 7591264 | 6734765 | 6376106.5 |
| Q9Z0J0 | Npc2 | NPC intracellular cholesterol transporter 2 Mus musculus | 223.6 | 41.61 | 16442 | 5 | 5 | 44724541 | 60019929 | 47149653 | 40513769 | 46260973 | 46060909 | 33179089 | 50040265 | 47737569 | 33892921 | 35437833 | 37040825 | 45005085 | 51096793 | 29559257 |
| Q6NSR8 | Npepl1 | Probable aminopeptidase NPEPL1 Mus musculus | 394.99 | 16.6 | 55940 | 6 | 6 | 22768453 | 16636672 | 17058027 | 19636999 | 14535225 | 13633430 | 15695921 | 15452034 | 10848403 | 16799283 | 15665205 | 12052456 | 15221070 | 15339371 | 16592221 |
| Q11011 | Npepps | Puromycin-sensitive aminopeptidase Mus musculus | 879.35 | 30.11 | 103325 | 19 | 19 | 29901503 | 28245597 | 28708541 | 28451621 | 28245683 | 26880889 | 30208221 | 31141367 | 31482041 | 28227957 | 27422149 | 30216155 | 26631845 | 35377165 | 28681247 |
| P60670 | Nploc4 | Nuclear protein localization protein 4 homolog Mus musculus | 561.66 | 28.45 | 68017 | 9 | 9 | 3993311 | 3392168 | 6051331.5 | 2780395.5 | 3892635.5 | 3746189.75 | 5350393.5 | 3236005.5 | 5738956.5 | 3792856 | 3680780.75 | 6732320.5 | 4088052 | 6090354.5 | 8054620 |
| P13084 | Npm1 | Nucleophosmin Rattus norvegicus | 979.92 | 45.21 | 32560 | 12 | 14 | 136147089 | 112855337 | 132390833 | 127735353 | 137749089 | 138508577 | 165334129 | 115679241 | 107704865 | 154920337 | 157767649 | 148101473 | 139003233 | 106333985 | 168231569 |
| Q9CPP0 | Npm3 | Nucleoplasmin-3 Mus musculus | 214.55 | 25.71 | 19023 | 3 | 3 | 3219045 | 2339045.5 | 4039733.25 | 4012519.25 | 4656113 | 4588377 | 4902138 | 4957080 | 2431888.25 | 4922930.5 | 3461670.5 | 4236091 | 4118785.25 | 2832540.25 | 6327449 |
| P97300 | Nptn | Neuroplastin Mus musculus | 130.03 | 6.3 | 44373 | 2 | 2 | 3599201 | 3667696.75 | 3625200 | 4168693 | 4074775 | 3769316.25 | 4979328.5 | 3636639.75 | 3319186 | 4586678.5 | 3786982.5 | 4104769.5 | 3871936.5 | 3115849.25 | 3619425.5 |
| Q9JI75 | Nqo2 | Ribosyldihydronicotinamide dehydrogenase [quinone] Mus musculus | 120.65 | 14.72 | 26248 | 2 | 2 | 2998852.75 | 3186597 | 3126613.75 | 3019940.5 | 2777386.5 | 2605594 | 2444238.75 | 3168671 | 3939255.25 | 2996649.75 | 2475991.5 | 2613862.5 | 3148455 | 3283391.25 | 1898284.5 |
| P08556 | Nras | GTPase NRas Mus musculus | 188.98 | 25.4 | 21199 | 3 | 3 | 4121496.5 | 3810453.25 | 5570454 | 5352785 | 6521370 | 6865825 | 5243836 | 4183389.5 | 5622708 | 6640284.5 | 5512254.5 | 6032394.5 | 6268911.5 | 5416485.5 | 6317581 |
| Q8BHG1 | Nrdc | Nardilysin Mus musculus | 374.9 | 10.51 | 132891 | 9 | 9 | 10176996 | 8305207 | 7807701.5 | 7826306.5 | 7628328 | 8810374 | 6166838 | 8166184.5 | 8672748 | 6439079 | 7781659.5 | 7894457.5 | 7374458 | 9319447 | 7808285.5 |
| P97333 | Nrp1 | Neuropilin-1 Mus musculus | 105.47 | 3.58 | 103000 | 2 | 2 | 2158467 | 1629235.62 | 1624815.75 | 1570022 | 1405104.25 | 702659 | 1167366.5 | 1577390.88 | 696628.5 | 1222015.88 | 1256722.88 | 1540028.25 | 1073999.75 | 1566660.38 | 1148377 |
| Q9R1J0 | Nsdhl | Sterol-4-alpha-carboxylate 3-dehydrogenase, decarboxylating Mus musculus | 330.15 | 25.97 | 40686 | 6 | 6 | 4963761 | 2714816.5 | 4315897 | 3331419.5 | 3430446 | 3590332 | 6767532.5 | 3682288 | 5342688.5 | 6153319.5 | 4070346.25 | 6350070.5 | 5142092 | 4998720 | 6530081 |
| P46460 | Nsf | Vesicle-fusing ATPase Mus musculus | 1487.1 | 46.24 | 82613 | 28 | 28 | 28648385 | 35985257 | 34471449 | 30676035 | 35588877 | 33337239 | 33229173 | 30541329 | 34299277 | 32067423 | 35412481 | 36123613 | 37951761 | 32193571 | 40382369 |
| O35987 | Nsfl1c | NSFL1 cofactor p47 Rattus norvegicus | 667.58 | 40.54 | 40680 | 11 | 11 | 18535299 | 21998747 | 17681253 | 24993347 | 20707339 | 21670761 | 23565567 | 18596343 | 18309145 | 24248651 | 22804835 | 21975409 | 22582219 | 19491455 | 19295937 |
| Q91VT1 | Nsmce2 | E3 SUMO-protein ligase NSE2 Mus musculus 2 | 75.68 | 6.07 | 28231 | 1 | 1 | 1 | 1 | 507294.69 | 1 | 1 | 1 | 1 | 459184.53 | 1 | 1 | 319715 | 755956.25 | 1 | 1 | 1 |
| Q4FZU3 | Nsrp1 | Nuclear speckle splicing regulatory protein 1 Rattus norvegicus | 36.96 | 1.83 | 64180 | 1 | 1 | 90277.63 | 2875351.25 | 2368752.5 | 3279576.75 | 3486250 | 1791109.5 | 4049998.25 | 1858533.62 | 1952125.438 | 108860.38 | 2107446.75 | 1572255.935 | 2351855.75 | 877208.12 | 952513.12 |
| Q1HFZ0 | Nsun2 | tRNA (cytosine(34)-C(5))-methyltransferase Mus musculus | 549.78 | 19.68 | 85452 | 10 | 10 | 11462024 | 11726053 | 11252619 | 10511617 | 9920910 | 11628313 | 10329161 | 12595110 | 12219544 | 8276973.5 | 10743725 | 10790789 | 9402578 | 12348867 | 10670124 |
| Q9JM14 | Nt5c | 5'(3')-deoxyribonucleotidase, cytosolic type Mus musculus | 350.22 | 38.5 | 23076 | 4 | 4 | 4898428.5 | 5409124.5 | 4442569 | 6602330.5 | 4849104 | 4737544 | 6396700 | 2317035.25 | 1786012 | 10299191 | 3865551.75 | 6273737.5 | 4740240.5 | 5183124.5 | 6202576.5 |
| Q3V1L4 | Nt5c2 | Cytosolic purine 5'-nucleotidase Mus musculus | 785.37 | 36.07 | 64809 | 15 | 15 | 11805436 | 15177108 | 12163181 | 14185695 | 11898906 | 12346851 | 13854295 | 13807298 | 14803553 | 12153173 | 12537989 | 11550763 | 15391181 | 12817752 | 11694165 |
| Q3UFY7 | Nt5c3b | 7-methylguanosine phosphate-specific 5'-nucleotidase Mus musculus | 172.13 | 19.67 | 34425 | 3 | 3 | 1345667.12 | 1863601.25 | 2155446.75 | 689296.62 | 1895672 | 1381866.38 | 801201.31 | 2029958 | 2179464.5 | 1505715.5 | 1707610.5 | 1843541 | 1792320.5 | 2381118.5 | 1842512.25 |
| Q5BJX0 | Ntmt1 | N-terminal Xaa-Pro-Lys N-methyltransferase 1 Rattus norvegicus | 53.68 | 3.14 | 25464 | 1 | 1 | 826081 | 663944.56 | 877264.38 | 470766.31 | 690390.19 | 797174.38 | 693422.88 | 719456.56 | 633617 | 741302.25 | 593296.38 | 651003 | 793579.5 | 726929.31 | 573393.88 |
| Q9CQA9 | Ntpcr | Cancer-related nucleoside-triphosphatase homolog Mus musculus | 144.1 | 28.95 | 20667 | 3 | 3 | 7992914.5 | 5703867 | 2309010 | 4058435.5 | 3470464.25 | 6469517.5 | 2532949.75 | 2024818.5 | 3334803.5 | 4582399 | 3450272.75 | 1793821.25 | 3409893.5 | 4444392.5 | 4900685 |
| P54729 | Nub1 | NEDD8 ultimate buster 1 Mus musculus | 132.86 | 7.98 | 70307 | 3 | 3 | 999581.62 | 1611360.38 | 839202.38 | 806820.19 | 554152.69 | 651337.38 | 710417.75 | 504647.88 | 922005.25 | 896291.12 | 542686.62 | 1532032.38 | 579398.88 | 546158.12 | 744005.31 |
| Q9R061 | Nubp2 | Cytosolic Fe-S cluster assembly factor NUBP2 Mus musculus | 133.66 | 14.91 | 29518 | 3 | 3 | 3554955 | 4566682 | 2580230 | 3886354 | 3901968 | 3609415.5 | 4212867.5 | 3450824.5 | 3914734.25 | 4105364 | 3407690.5 | 4222438.5 | 4619389 | 3941809 | 4197473 |
| Q02819 | Nucb1 | Nucleobindin-1 Mus musculus | 158.35 | 10.68 | 53409 | 3 | 3 | 1698301.12 | 1853747.88 | 1118186.75 | 1884950.25 | 1270880.88 | 1525191.12 | 1263804.88 | 1666508.62 | 1348658.25 | 1174699.62 | 971685.19 | 1657074 | 1470935.75 | 1040539.69 | 1244949.62 |
| P81117 | Nucb2 | Nucleobindin-2 Mus musculus | 207.66 | 14.05 | 50304 | 4 | 4 | 4508465 | 4875450.5 | 4607830.5 | 4960994 | 4038016.25 | 4188479.5 | 3623639.75 | 5539452.5 | 5410798 | 4471655.5 | 3441778.25 | 4161369 | 3847329 | 4682715 | 3807748.5 |
| Q80XU3 | Nucks1 | Nuclear ubiquitous casein and cyclin-dependent kinase substrate 1 Mus musculus | 140.91 | 10.26 | 26313 | 2 | 2 | 1549337.88 | 5748996 | 4899099 | 5975176 | 8229839.5 | 6780911 | 9814120 | 5693306 | 4263365 | 9881570 | 7689289 | 6360804 | 7678111 | 4428735.5 | 9083165 |
| O35685 | Nudc | Nuclear migration protein nudC Mus musculus | 1046.91 | 67.47 | 38358 | 18 | 18 | 52287017 | 53520913 | 51342013 | 55752165 | 56364201 | 54704721 | 57963593 | 53523305 | 50976525 | 59632629 | 55688989 | 55485249 | 53687465 | 52251397 | 58793921 |

| | | | | | | | | | | | | | | | | | | | | | | |
|---|---|---|---|---|---|---|---|---|---|---|---|---|---|---|---|---|---|---|---|---|---|---|
| Q6PIP5 | Nudcd1 | NudC domain-containing protein 1 Mus musculus | 391 | 24.4 | 66705 | 9 | 9 | 8369140.5 | 8550534 | 7894536 | 8618849 | 8807447 | 9861959 | 7856768 | 9653839 | 11885193 | 9191013 | 10208554 | 9337844 | 9079144 | 10043479 | 8939247 |
| Q5M823 | Nudcd2 | NudC domain-containing protein 2 Rattus norvegicus 2 | 177.9 | 24.2 | 17674 | 3 | 3 | 2781873.25 | 2213616.75 | 1811639.62 | 2710745.5 | 2033663.25 | 1933194.62 | 3412648.75 | 2105429.25 | 1288694.75 | 3130715 | 1695644.5 | 2039195.38 | 2084303.5 | 2213279 | 2009113 |
| Q8JZU0 | Nudt13 | Nucleoside diphosphate-linked moiety X motif 13 Mus musculus | 50.78 | 5.68 | 39137 | 1 | 1 | 508560.72 | 513306.47 | 533769.44 | 528764.38 | 458402.59 | 679118.94 | 517676.28 | 542857.7275 | 524007.66 | 450628.03 | 527319.6775 | 451798.75 | 578128.81 | 606787.62 | 472563.53 |
| Q8VHN8 | Nudt16l1 | Tudor-interacting repair regulator protein Mus musculus | 111.47 | 19.43 | 23414 | 3 | 3 | 1841913.25 | 1 | 1 | 1 | 254077.17 | 1 | 1 | 2093287.12 | 200399.22 | 1 | 1 | 1329214.38 | 312267.69 | 1 | 303622.91 |
| Q3U2V3 | Nudt18 | 8-oxo-dGDP phosphatase NUDT18 Mus musculus | 59.15 | 4.95 | 35694 | 1 | 1 | 211769.22 | 261989.75 | 238622.64 | 305746.12 | 292658.69 | 260973.78 | 331649.06 | 256088.72 | 221630.56 | 315054.47 | 429770.25 | 269225.69 | 258976.72 | 290095.19 | 279858.19 |
| P56380 | Nudt2 | Bis(5'-nucleosyl)-tetraphosphatase [asymmetrical] Mus musculus | 172.09 | 27.89 | 16989 | 3 | 3 | 1725483.5 | 1933229.5 | 3011968 | 2932848.5 | 2588678.25 | 2818607.75 | 1849077.25 | 3474031 | 4175410.75 | 776465.12 | 592150.38 | 1465661.62 | 946737.12 | 1576744.75 | 1336226 |
| Q4KM65 | Nudt21 | Cleavage and polyadenylation specificity factor subunit 5 Rattus norvegicus 2 | 385.6 | 49.78 | 26240 | 8 | 8 | 738469.69 | 646067.06 | 523869.62 | 985249.62 | 627260.62 | 674868.75 | 1238725 | 471981.5 | 258316.53 | 919491.88 | 479637.12 | 354107.44 | 904196.75 | 380106.69 | 449486.72 |
| Q9JKX6 | Nudt5 | ADP-sugar pyrophosphatase Mus musculus | 128.9 | 23.85 | 23984 | 3 | 3 | 2263312.75 | 1435845.38 | 2279520.75 | 2242799 | 1905312.38 | 1512001.25 | 4133290 | 1418981.75 | 1277584.62 | 3002416.5 | 1725992.12 | 2819403 | 1131754.5 | 1094980.62 | 2909580.25 |
| E9Q7G0 | Numa1 | Nuclear mitotic apparatus protein 1 Mus musculus | 2416.61 | 29.89 | 235630 | 47 | 47 | 27765579 | 32704075 | 33247279 | 38427785 | 36773177 | 32430267 | 40411201 | 28603351 | 26605165 | 37896457 | 37796969 | 33288285 | 34782521 | 34402145 | 32521427 |
| Q8BH74 | Nup107 | Nuclear pore complex protein Nup107 Mus musculus | 111.85 | 3.02 | 106717 | 2 | 2 | 1156890.25 | 880226.56 | 1242359.88 | 986539.31 | 916754.88 | 769362.25 | 1042908.19 | 1129777.75 | 1359340 | 972787.88 | 1273165 | 1403019.12 | 923053.19 | 1400165.5 | 1359088.12 |
| Q8R0G9 | Nup133 | Nuclear pore complex protein Nup133 Mus musculus | 1105.75 | 22.16 | 128620 | 18 | 18 | 5994157 | 6022322 | 6925766.5 | 7015285.5 | 5036463.5 | 8355805.5 | 5718366.5 | 7383221.5 | 5229202.5 | 5836591 | 7556981 | 7754657.5 | 4439237 | 5812145 | 4904007.5 |
| P49791 | Nup153 | Nuclear pore complex protein Nup153 Rattus norvegicus | 155.16 | 3.54 | 152824 | 3 | 3 | 705399.83 | 598635.19 | 886343.19 | 637318.19 | 699302.75 | 1 | 674306.81 | 478154.47 | 1 | 749682.5 | 573911.69 | 892431.88 | 672566.55 | 589266.25 | 634656.38 |
| Q99P88 | Nup155 | Nuclear pore complex protein Nup155 Mus musculus | 1193.77 | 24.51 | 155118 | 22 | 22 | 10530650 | 10842067 | 9080624 | 8302901 | 10888363 | 11208939 | 14921269 | 10425586 | 17211931 | 12820143 | 13202617 | 9852421 | 10717960 | 9905343 | 12792303 |
| Q9Z0W3 | Nup160 | Nuclear pore complex protein Nup160 Mus musculus | 297.61 | 6.28 | 158232 | 6 | 6 | 2887493.25 | 2270639 | 2146823 | 1754315 | 2257415.5 | 2927658 | 1924138.5 | 2468241.75 | 3488354.5 | 1832460.12 | 3159370 | 2384735.25 | 1858952 | 3311424.5 | 2646075 |
| Q6ZQH8 | Nup188 | Nucleoporin NUP188 homolog Mus musculus | 42.66 | 0.97 | 196696 | 1 | 1 | 194343.38 | 175196.77 | 140924.03 | 174023.38 | 229176.86 | 204735.78 | 110213.91 | 184442.98 | 164814.94 | 221957.36 | 93120.43 | 215520.14 | 190570.16 | 144969.94 | 124634.96 |
| Q9QY81 | Nup210 | Nuclear pore membrane glycoprotein 210 Mus musculus | 293.1 | 4.88 | 204101 | 7 | 7 | 1168368.62 | 2198284 | 1950679.5 | 1867510 | 787036.12 | 2681813.5 | 1823803.12 | 2858954 | 3270294 | 1352636 | 2718654.25 | 2801125.5 | 1917068.88 | 2969995.5 | 2132215.75 |
| Q80U93 | Nup214 | Nuclear pore complex protein Nup214 Mus musculus | 531.86 | 9.16 | 212979 | 10 | 10 | 3388357.25 | 7711486.5 | 7344560 | 8551104 | 7801118 | 3403761.5 | 9021114 | 3927719.5 | 2254545.75 | 7058766.5 | 7477798.5 | 3951108.5 | 5122566.5 | 6318791 | 6712382 |
| Q8R4R6 | Nup35 | Nucleoporin NUP35 Mus musculus | 173.28 | 14.15 | 34786 | 3 | 3 | 1 | 1 | 670254.44 | 1409240.25 | 1 | 1 | 863630.75 | 1 | 1 | 625737.06 | 478814.31 | 485495.41 | 858573.25 | 275389.56 | 837048.44 |
| Q9CWU9 | Nup37 | Nucleoporin Nup37 Mus musculus | 165.38 | 13.5 | 36732 | 3 | 3 | 1842334.75 | 2400950.25 | 686927.44 | 2705120 | 1737793 | 1990318 | 1695664 | 2663448.75 | 1667483.5 | 1872044.75 | 1866941.5 | 2029530.88 | 1764553.5 | 1646891.5 | 697873.56 |
| P59235 | Nup43 | Nucleoporin Nup43 Mus musculus | 92.14 | 8.16 | 41990 | 2 | 2 | 1128446.5 | 328673.97 | 1697451.62 | 595154.75 | 1537837.5 | 1338765 | 1752172.5 | 1398278.515 | 835077.31 | 1667099.25 | 343287 | 1411126.75 | 1092298 | 373790.59 | 1493386.88 |
| Q9JIH2 | Nup50 | Nuclear pore complex protein Nup50 Mus musculus | 343.54 | 19.96 | 49485 | 7 | 7 | 2637541 | 1693044.5 | 1724740.5 | 2573843.5 | 1836018.38 | 2835484.25 | 3147674 | 2449124.5 | 2067631.25 | 2902921.25 | 2470205 | 3214510 | 1359574 | 2910446.75 | 1609295.75 |
| Q8R332 | Nup58 | Nucleoporin p58/p45 Mus musculus | 42.75 | 2.21 | 59445 | 1 | 1 | 559396.3575 | 383305.03 | 709591.94 | 662437.12 | 482251.34 | 669442.56 | 1 | 1 | 1 | 1 | 523897.66 | 683773.62 | 601787.75 | 628717.3975 | 705410.56 |
| P17955 | Nup62 | Nuclear pore glycoprotein p62 Rattus norvegicus | 55.66 | 5.33 | 53397 | 2 | 2 | 589935.69 | 521881.25 | 1020462.38 | 638039.3225 | 419877.97 | 294917.81 | 808836.6625 | 418255.84 | 2164195 | 357978 | 700693.44 | 1555371.75 | 333623.12 | 548934.12 | 693244.69 |
| Q8R480 | Nup85 | Nuclear pore complex protein Nup85 Mus musculus | 83.37 | 2.9 | 74776 | 1 | 2 | 1118061.62 | 1210350.38 | 1055577.515 | 1357544.5 | 536353.56 | 1405816.62 | 1556512.88 | 1040785.25 | 1225537.88 | 1378871.5 | 943633 | 901835.12 | 1295156.5 | 1069749.75 | 886878.31 |
| O08658 | Nup88 | Nuclear pore complex protein Nup88 Rattus norvegicus | 98.15 | 4.45 | 83585 | 2 | 2 | 839509.62 | 807910.88 | 751638.9375 | 702566.31 | 656568.94 | 606684.25 | 349864.03 | 537066.6175 | 765492.94 | 426225.25 | 743247.31 | 655471.19 | 790606.06 | 942822.38 | 665930.94 |
| Q8BJ71 | Nup93 | Nuclear pore complex protein Nup93 Mus musculus | 1117.96 | 38.83 | 93281 | 22 | 22 | 15696478 | 18930093 | 18347425 | 14754703 | 15892601 | 17157047 | 15980099 | 16263409 | 18069197 | 18215011 | 17278179 | 17488529 | 17057045 | 20476405 | 20189101 |
| Q6PFD9 | Nup98 | Nuclear pore complex protein Nup98-Nup96 Mus musculus | 440.96 | 10.63 | 197241 | 4 | 11 | 4648943.5 | 4615951 | 4380564 | 3515120 | 4242840.5 | 3545705.5 | 2422928.25 | 4295546.5 | 2437759.75 | 4109813.5 | 4489076 | 3441909 | 4060102.75 | 3065047 | 2618882.5 |
| P61971 | Nutf2 | Nuclear transport factor 2 Mus musculus | 220.68 | 55.12 | 14478 | 4 | 4 | 3060826 | 4045924.75 | 3370719.75 | 4297897.5 | 3546473.5 | 4536013.5 | 2909065 | 3620175.75 | 3421231 | 3590200.5 | 2583586.25 | 2918857 | 3316110 | 3650829 | 2486308 |
| Q9DBY8 | Nvl | Nuclear valosin-containing protein-like Mus musculus | 211.68 | 7.25 | 94476 | 4 | 4 | 5006893 | 3699745.25 | 4004343 | 4415970 | 4322784.5 | 2172377.5 | 2996748.25 | 2547878.25 | 2998211.375 | 4275841.5 | 2058915.62 | 1517011.75 | 2354571.5 | 1917346.75 | 2530980 |
| Q99JX7 | Nxf1 | Nuclear RNA export factor 1 Mus musculus | 419.97 | 21.68 | 70300 | 8 | 8 | 4546236.5 | 3721399 | 4519531 | 4359245 | 3964913.25 | 4661964.5 | 6090408.5 | 2428079 | 4491090 | 4142605.75 | 4737069.5 | 4648029.5 | 4092033.75 | 4226561.5 | 5190956 |
| P11928 | Oas1a | 2'-5'-oligoadenylate synthase 1A Mus musculus | 477.04 | 27.79 | 42429 | 7 | 7 | 6047583.5 | 6469717 | 5685870 | 2889836.75 | 4769320.5 | 4831525 | 2330317 | 4603861 | 3942358 | 4617247 | 6050117 | 2535689 | 5767860.5 | 5507856.5 | 5430937.5 |
| Q8VI93 | Oas3 | 2'-5'-oligoadenylate synthase 3 Mus musculus | 1016.07 | 27.15 | 126333 | 21 | 21 | 15974175 | 14548404 | 17765663 | 11573377 | 15456911 | 14809910 | 14963656 | 19179459 | 18673015 | 13459611 | 16832143 | 14683896 | 15168237 | 18568099 | 18276549 |
| Q8VI94 | Oasl1 | 2'-5'-oligoadenylate synthase-like protein 1 Mus musculus | 173.91 | 10.37 | 59088 | 4 | 4 | 902312 | 652955.69 | 787475.69 | 421891.06 | 521197.62 | 589562.88 | 304173.94 | 960994.12 | 1474480.88 | 1 | 288143.56 | 566720.75 | 130630.88 | 1261727.25 | 550358.75 |
| P29758 | Oat | Ornithine aminotransferase, mitochondrial Mus musculus | 1084.71 | 50.11 | 48355 | 16 | 16 | 54754557 | 48940553 | 49442789 | 54602601 | 45663557 | 57831221 | 70576329 | 64418657 | 55301041 | 62190553 | 52630437 | 51418689 | 53731941 | 60485337 | 52549909 |
| Q9CRD0 | Ociad1 | OCIA domain-containing protein 1 Mus musculus | 177.45 | 19.43 | 27610 | 1 | 3 | 1188468.12 | 1515241.5 | 1298614.5 | 1118878 | 1385194 | 1301042.38 | 1468559 | 1236961.62 | 1815408 | 1175793.75 | 975841.19 | 1071489.62 | 1592451.75 | 1251039.5 | 1303759.38 |
| Q4PJX1 | Odr4 | Protein odr-4 homolog Mus musculus | 49.47 | 3.13 | 49998 | 1 | 1 | 927952.94 | 1150725.38 | 1484045.62 | 1079645.25 | 1102688.75 | 1118702 | 1 | 1118720.38 | 1262821.88 | 1 | 1220948.125 | 1206858.88 | 1284041.62 | 1105604.75 | 1287287.25 |
| Q9EQQ9 | Oga | Protein O-GlcNAcase Mus musculus | 122.1 | 7.64 | 103162 | 3 | 3 | 1154215.38 | 1077223 | 1109541.12 | 977999.25 | 985560.69 | 1232354.62 | 1280929 | 1256184.88 | 1035038.31 | 1139215.88 | 868202.44 | 1113611.5 | 1346341.62 | 1009973.25 | 1130429.12 |
| Q60597 | Ogdh | 2-oxoglutarate dehydrogenase, mitochondrial Mus musculus | 1003.28 | 23.66 | 116449 | 18 | 18 | 27483485 | 22267293 | 26240773 | 22031441 | 23816069 | 22920967 | 23483645 | 25327473 | 28691315 | 21516827 | 23627197 | 21401137 | 21360585 | 22779611 | 18422675 |
| Q3U0K8 | Ogfod1 | Prolyl 3-hydroxylase OGFOD1 Mus musculus | 117.14 | 9.54 | 62734 | 3 | 3 | 247794.14 | 409292.31 | 1993545.75 | 931927.94 | 465156.06 | 449778 | 468681.53 | 534931.12 | 856140.69 | 577382.835 | 698236.94 | 643731.06 | 1281734.12 | 618716.38 | 507579.69 |
| Q99PG2 | Ogfr | Opioid growth factor receptor Mus musculus | 388.28 | 21.48 | 70679 | 7 | 7 | 2979361.75 | 4873649 | 4787515 | 4835608 | 3108700 | 4248837 | 3790761.25 | 3660421 | 2651587.75 | 4447385 | 3317407.75 | 3211440 | 4269712 | 3997655.25 | 1432560.88 |
| Q8CGY8 | Ogt | UDP-N-acetylglucosamine--peptide N-acetylglucosaminyltransferase 110 kDa subunit M | 734.49 | 17.69 | 116952 | 13 | 13 | 6992855.5 | 9327094 | 9089789 | 8556394 | 11275407 | 9388019 | 11491333 | 7741297 | 6920084.5 | 9413489 | 9496299 | 8813815 | 10957500 | 10430153 | 9631309 |
| Q9CZ30 | Ola1 | Obg-like ATPase 1 Mus musculus | 538.06 | 31.31 | 44730 | 8 | 8 | 24990359 | 24290319 | 25150643 | 25384167 | 26113907 | 25695397 | 24605877 | 26068797 | 25882065 | 23488253 | 24036801 | 29305233 | 23570971 | 27963487 | 26558397 |
| P58281 | Opa1 | Dynamin-like 120 kDa protein, mitochondrial Mus musculus | 176.73 | 5.73 | 111339 | 3 | 3 | 2297043.75 | 2142200.5 | 617528.69 | 1406200.75 | 1987594 | 531941.69 | 1869250.25 | 2251691.75 | 2534416.75 | 293830.22 | 2293628.5 | 1968197.12 | 1817539.25 | 715799.94 | 1932447 |
| Q60862 | Orc2 | Origin recognition complex subunit 2 Mus musculus | 41.47 | 4.17 | 65893 | 1 | 1 | 356100.53 | 312731.78 | 203458.05 | 250421.69 | 165191.72 | 260256.34 | 121894.66 | 169976.51 | 159479.66 | 138275.38 | 301507.31 | 208785.97 | 120572.63 | 1 | 1 |
| Q5RKH6 | Os9 | Protein OS-9 Rattus norvegicus | 35.79 | 1.95 | 75365 | 1 | 1 | 625222.5 | 872828.5 | 781162.19 | 778855.0775 | 836207.12 | 946487.44 | 744357.56 | 919867.75 | 827172.88 | 499158.34 | 850905.19 | 637520.5 | 663420.38 | 664853 | 702940.5 |
| Q3B7Z2 | Osbp | Oxysterol-binding protein 1 Mus musculus | 323 | 13.29 | 88797 | 7 | 7 | 3191671 | 4634675.5 | 6035773 | 4057666.5 | 6499011.5 | 4001157 | 4824222.5 | 5412131.5 | 5561892 | 6069501 | 4861591 | 4061826 | 5164109 | 6094600.5 | 5560279 |
| Q8CI95 | Osbpl11 | Oxysterol-binding protein-related protein 11 Mus musculus | 125.9 | 5.73 | 83629 | 3 | 3 | 903306.31 | 1133303.38 | 865538.25 | 1084914.75 | 1070425.62 | 1139047.75 | 995253.19 | 1064255.12 | 1097870.62 | 881657.44 | 1054775.5 | 1001150.94 | 588499.81 | 825300 | 1069269.25 |
| Q8BX94 | Osbpl2 | Oxysterol-binding protein-related protein 2 Mus musculus | 84.85 | 6.82 | 55384 | 2 | 2 | 1280324.328 | 819243.31 | 1299020.62 | 942533.5 | 2060499.88 | 2067582.88 | 1991421.12 | 1942878.88 | 2183089.5 | 978767.5 | 1113728.5 | 2108249.25 | 943885.5 | 788640 | 2187270 |
| B9EJ86 | Osbpl8 | Oxysterol-binding protein-related protein 8 Mus musculus | 956.37 | 31.05 | 101269 | 18 | 18 | 42724825 | 33936841 | 33793157 | 26560865 | 40318105 | 41059165 | 31879655 | 39496781 | 45168441 | 33284369 | 40238825 | 39287437 | 36398821 | 37470917 | 48169385 |
| Q8BWU5 | Osgep | Probable tRNA N6-adenosine threonylcarbamoyltransferase Mus musculus | 196.87 | 13.43 | 36301 | 3 | 3 | 1212880.38 | 1635722.25 | 1653178.75 | 1559167 | 1533169.75 | 1197989 | 1785449.25 | 1572288 | 1436694.62 | 1714689.12 | 1676663.25 | 1996758.25 | 1753964.5 | 1336730.62 | 1592864 |
| Q62422 | Ostf1 | Osteoclast-stimulating factor 1 Mus musculus | 340.08 | 38.6 | 23783 | 6 | 6 | 18533831 | 17938783 | 16404530 | 14253617 | 17516889 | 19915213 | 16136219 | 17177319 | 19781357 | 9703479 | 18634321 | 17942941 | 17338113 | 19906097 | 15524966 |
| B2RYG6 | Otub1 | Ubiquitin thioesterase OTUB1 Rattus norvegicus | 504.51 | 49.82 | 31270 | 9 | 9 | 21379525 | 21825843 | 20062337 | 18928377 | 18055105 | 21235103 | 19863461 | 21856049 | 25222781 | 17053281 | 22888203 | 21627797 | 18989853 | 21608079 | 21019205 |
| B2RRE7 | Otud4 | OTU domain-containing protein 4 Mus musculus | 52.92 | 0.72 | 123055 | 1 | 1 | 577512.62 | 870309.88 | 664846.75 | 937472.62 | 694400.5 | 600392.06 | 608496.75 | 655632.88 | 818856.94 | 563718 | 465032.44 | 607788.25 | 872435.19 | 789505 | 591964.06 |
| Q8K2H2 | Otud6b | Deubiquitinase OTUD6B Mus musculus | 37.55 | 4.42 | 33758 | 1 | 1 | 1063783.563 | 913113.69 | 923101.06 | 1267940.75 | 1150978.75 | 1037561.44 | 992476.94 | 1172265 | 1191219 | 1392712.62 | 813182.3425 | 982274.19 | 1223442 | 352720.56 | 694292.62 |
| Q9D7E3 | Ovca2 | Esterase OVCA2 Mus musculus | 38.44 | 4.89 | 24246 | 1 | 1 | 1623532.033 | 1819773.25 | 1587083.38 | 1572344.25 | 1514927.25 | 1448689.62 | 1595412.25 | 1368589.25 | 2130469.25 | 1433900.88 | 1198952.5 | 1299781.75 | 1843657.88 | 1722358 | 1154386.25 |
| Q9D0K2 | Oxct1 | Succinyl-CoA:3-ketoacid coenzyme A transferase 1, mitochondrial Mus musculus | 1174.91 | 43.65 | 55989 | 15 | 15 | 38869481 | 50020709 | 50447477 | 51833617 | 52585401 | 52766841 | 68385017 | 55093925 | 47794729 | 55525261 | 56438569 | 59848025 | 58012089 | 53772453 | 59956445 |
| Q6P9R2 | Oxsr1 | Serine/threonine-protein kinase OSR1 Mus musculus | 314.35 | 15.56 | 58214 | 6 | 6 | 4060466.75 | 2940360 | 4116421.5 | 1372253.25 | 2363157.5 | 2087592 | 2222373.5 | 4344826.5 | 4259801 | 1849769.88 | 3017637.25 | 3376015.5 | 779900.12 | 3744790.5 | 2170686 |
| P54001 | P4ha1 | Prolyl 4-hydroxylase subunit alpha-1 Rattus norvegicus | 390.79 | 18.54 | 60898 | 8 | 8 | 2903602.75 | 507346.91 | 1275662 | 2642445 | 1580002.75 | 1140461.5 | 2965084.5 | 386696.91 | 1657427.5 | 2126334.5 | 1361249.5 | 1183066.12 | 1223822.25 | 823065 | 3131280.5 |
| P09103 | P4hb | Protein disulfide-isomerase Mus musculus | 1769.22 | 55.6 | 57058 | 28 | 28 | 136559073 | 139142049 | 125258673 | 127324713 | 149719969 | 171607425 | 190946033 | 169422401 | 153140801 | 160991313 | 152096065 | 148470625 | 157284705 | 157814065 | 142346193 |
| P50580 | Pa2g4 | Proliferation-associated protein 2G4 Mus musculus | 935.58 | 59.64 | 43699 | 18 | 18 | 38409153 | 32099041 | 39336401 | 40108437 | 35228189 | 34776045 | 49786393 | 37154529 | 30424767 | 40848153 | 39254601 | 39664289 | 35903549 | 34279729 | 41604873 |
| P29341 | Pabpc1 | Polyadenylate-binding protein 1 Mus musculus | 1665.87 | 60.69 | 70671 | 30 | 31 | 76494185 | 81910409 | 88746609 | 87175105 | 83188929 | 77496993 | 93428769 | 74256033 | 72538153 | 87669593 | 80428529 | 89638033 | 89925289 | 78300353 | 95442609 |
| Q8CCS6 | Pabpn1 | Polyadenylate-binding protein 2 Mus musculus | 218.4 | 27.15 | 32297 | 4 | 4 | 10433612 | 11457711 | 10696737 | 11764491 | 12181833 | 11124757 | 12334383 | 10160416 | 10751481 | 11601816 | 11075137 | 11515323 | 10852823 | 10268235 | 10467551 |
| Q9WVE8 | Pacsin2 | Protein kinase C and casein kinase substrate in neurons protein 2 Mus musculus | 267.12 | 16.87 | 55833 | 7 | 7 | 8856287 | 9064631 | 9657616 | 8117228 | 8317619 | 8573510 | 7203483.5 | 9104658 | 11845181 | 6537586.5 | 10330099 | 10643504 | 9105320 | 9973478 | 8826263 |
| Q4V886 | Paf1 | RNA polymerase II-associated factor 1 homolog Rattus norvegicus | 43.27 | 2.06 | 60546 | 1 | 1 | 287980.97 | 360996.91 | 289336.97 | 1 | 1 | 230679.84 | 1 | 1 | 386678.69 | 1 | 214166.59 | 254879.05 | 1 | 1 | 173699.89 |
| P63004 | Pafah1b1 | Platelet-activating factor acetylhydrolase IB subunit alpha Rattus norvegicus | 532.77 | 29.51 | 46670 | 11 | 11 | 15892587 | 9029093 | 15389069 | 14904659 | 13506808 | 11345749 | 18517927 | 12950877 | 14054746 | 15623649 | 16239371 | 15034300 | 15066133 | 16122273 | 15994743 |
| O35264 | Pafah1b2 | Platelet-activating factor acetylhydrolase IB subunit beta Rattus norvegicus | 136.06 | 12.23 | 25581 | 2 | 2 | 2320830.5 | 1351602 | 1813642.62 | 1926399 | 1 | 2195508.75 | 415517.91 | 2458073.25 | 2419237.25 | 952650.38 | 1954607.88 | 2044550.62 | 946815.94 | 2541495.25 | 2985759.5 |
| Q9DCL9 | Paics | Multifunctional protein ADE2 Mus musculus | 1061.37 | 48.71 | 47006 | 18 | 18 | 18406545 | 19401191 | 24465001 | 22549427 | 24582029 | 24542641 | 22948017 | 22450863 | 26748589 | 25406857 | 23463287 | 24475931 | 23016847 | 20707239 | 25519977 |
| Q8VE62 | Paip1 | Polyadenylate-binding protein-interacting protein 1 Mus musculus | 38.81 | 2.75 | 45702 | 1 | 1 | 1277745.5 | 1062720.25 | 1153127.12 | 1012264.56 | 391925.97 | 1039625.81 | 1159275.12 | 1311095.75 | 1239593.5 | 777507.81 | 611136.9525 | 611287.19 | 1081176.62 | 346153.22 | 405930.78 |
| Q9DCE5 | Pak1ip1 | p21-activated protein kinase-interacting protein 1 Mus musculus | 269.41 | 20.42 | 42116 | 6 | 6 | 1412074.38 | 1034778.94 | 1321317.75 | 1364280.5 | 1623268.38 | 920557.06 | 845591.5 | 1101438.5 | 905271.62 | 740721.88 | 1813598.25 | 1273017.08 | 854659.44 | 988373.75 | 1435436.88 |
| Q8CIN4 | Pak2 | Serine/threonine-protein kinase PAK 2 Mus musculus | 756.6 | 32.82 | 57930 | 9 | 13 | 13375337 | 8896386 | 13045911 | 10489531 | 11003372 | 12705217 | 16869867 | 11866644 | 10544881 | 13444614 | 11490451 | 12366475 | 14048644 | 13029368 | 14224669 |
| Q61036 | Pak3 | Serine/threonine-protein kinase PAK 3 Mus musculus | 343.94 | 15.56 | 62398 | 2 | 6 | 3598529.5 | 3919888.25 | 3638312.25 | 3730203 | 3620463.5 | 3185629.75 | 4874399.5 | 2933372.75 | 2850413.5 | 4545257.5 | 2481436.5 | 2866627.5 | 3621907.5 | 3208091.5 | 3669238 |
| Q9ET54 | Palld | Palladin Mus musculus | 79.78 | 3.12 | 152131 | 2 | 2 | 1 | 1 | 210994.59 | 212372.02 | 108996.18 | 128144.14 | 694002.62 | 174479.31 | 321622.69 | 289864.69 | 222202.81 | 281508.31 | 170849.36 | 123434.37 | 558698.56 |
| Q8K4K6 | Pank1 | Pantothenate kinase 1 Mus musculus | 50.91 | 1.64 | 60091 | 1 | 1 | 681330.06 | 668368.5 | 897217.38 | 723368.625 | 646558.56 | 837769.81 | 680345.12 | 806273.38 | 947516.31 | 649281.62 | 727113.955 | 655626.94 | 624376 | 800419.69 | 828033.19 |
| Q80YV4 | Pank4 | Pantothenate kinase 4 Mus musculus | 83.69 | 5.12 | 91522 | 2 | 2 | 390686.81 | 509115.09 | 599090.44 | 1 | 1 | 439694.22 | 629152.12 | 1 | 405789.91 | 1 | 361896 | 695100.56 | 452683.5 | 703679.12 | 180657.38 |
| Q60967 | Papss1 | Bifunctional 3'-phosphoadenosine 5'-phosphosulfate synthase 1 Mus musculus | 327.46 | 17.31 | 70794 | 7 | 7 | 3402638.5 | 5502268 | 5422769.5 | 4365005.5 | 5463396.5 | 5982951 | 4171174.75 | 4228502.5 | 5981984.5 | 4809961.5 | 3276638 | 6364839 | 3619013.25 | 5405406 | 4455478.5 |
| Q99LX0 | Park7 | Protein/nucleic acid deglycase DJ-1 Mus musculus | 623.23 | 76.19 | 20021 | 10 | 10 | 39469889 | 48986005 | 47944301 | 48110437 | 49732665 | 47677957 | 67264553 | 47584717 | 48625261 | 58814809 | 49599509 | 52057473 | 54521921 | 45797925 | 57046509 |
| Q5XJY4 | Parl | Presenilins-associated rhomboid-like protein, mitochondrial Mus musculus | 131.53 | 10.08 | 41964 | 2 | 2 | 1511375.75 | 853155.31 | 1231294.75 | 373937.16 | 1110664.88 | 1413692 | 1339057.62 | 1035115.12 | 1285429.12 | 1407576.75 | 1783960.62 | 1481441.75 | 1879075.62 | 1154811.38 | 1942707.25 |
| Q8VDG3 | Parn | Poly(A)-specific ribonuclease PARN Mus musculus | 55.84 | 2.88 | 71559 | 1 | 1 | 317543.69 | 526220.81 | 517402.84 | 359931.91 | 392415.41 | 527879.94 | 530086 | 496750.56 | 232928.75 | 372171.31 | 340027.5 | 443583.44 | 327056.38 | 476874.78 | 458423.91 |
| P11103 | Parp1 | Poly [ADP-ribose] polymerase 1 Mus musculus | 1111.99 | 23 | 113100 | 4 | 17 | 23729741 | 29371337 | 31731749 | 23119535 | 26062539 | 24353781 | 33466111 | 25617167 | 27191097 | 25637837 | 27368585 | 31991491 | 30740585 | 23403573 | 32178659 |
| Q8BZ20 | Parp12 | Poly [ADP-ribose] polymerase 12 Mus musculus | 125.95 | 5.2 | 79917 | 2 | 2 | 1 | 1 | 1 | 1 | 1 | 1 | 685821.56 | 1020994.44 | 1 | 1 | 1034420.62 | 816406.5 | 1 | 1 | 677478.56 |
| Q2EMV9 | Parp14 | Poly [ADP-ribose] polymerase 14 Mus musculus | 321.01 | 6.05 | 203802 | 7 | 7 | 2628801.75 | 1012310.38 | 2278531 | 1326686.62 | 1773543.5 | 1411212.75 | 1589171.75 | 2147582.5 | 2613716.75 | 1150361.38 | 1535348 | 2159403.25 | 1779776.5 | 1673615.25 | 1324245.62 |
| Q9ES46 | Parvb | Beta-parvin Mus musculus | 161.49 | 15.34 | 41669 | 4 | 4 | 429852 | 248073.16 | 193111.55 | 316781.56 | 451728.94 | 649424.62 | 128458.06 | 367902.47 | 910582.88 | 339792.75 | 778507.31 | 153416.16 | 395393.2825 | 279835 | 369814.66 |
| Q9ERD8 | Parvg | Gamma-parvin Mus musculus | 81.35 | 9.06 | 37603 | 2 | 2 | 2437660.25 | 1708778.12 | 1937751.75 | 1990288.88 | 2465646.5 | 1812238.75 | 5801343 | 2055059.62 | 1446262.88 | 3322678.5 | 2890433.5 | 3352810 | 1967724.62 | 2706140.25 | 4157164 |
| Q9D0B6 | Pbdc1 | Protein PBDC1 Mus musculus | 279.99 | 30.3 | 22223 | 5 | 5 | 9506809 | 10912057 | 10215348 | 11892442 | 9902948 | 11083385 | 8001520.5 | 11449819 | 13504393 | 9497466 | 8185025 | 11243184 | 8844554 | 10386651 | 8644418 |
| Q9JJ78 | Pbk | Lymphokine-activated killer T-cell-originated protein kinase Mus musculus | 44.52 | 3.94 | 36745 | 1 | 1 | 480654.28 | 856266.69 | 746374.75 | 786979.43 | 1064622 | 1 | 522000.34 | 473305.5 | 800807.62 | 1 | 1 | 451763.31 | 776399.44 | 570921.38 | 1 |
| Q8BSQ9 | Pbrm1 | Protein polybromo-1 Mus musculus | 69.92 | 1.1 | 187188 | 1 | 1 | 245477.16 | 297227.59 | 250973.81 | 203114.95 | 210653.03 | 230997.92 | 218546.47 | 360508.69 | 167828.61 | 296740.84 | 187910.98 | 352866.56 | 205166 | 283573 | 247391.77 |
| Q3TVI8 | Pbxip1 | Pre-B-cell leukemia transcription factor-interacting protein 1 Mus musculus | 354.97 | 8.8 | 81164 | 5 | 5 | 7381188.5 | 3296212.5 | 4588834.5 | 5065322 | 6231069.5 | 6550962 | 3272191 | 4400231 | 6832232 | 4895641 | 4681923.5 | 4559900 | 4436049 | 4429099 | 4253132 |
| P60335 | Pcbp1 | Poly(rC)-binding protein 1 Mus musculus | 1103.32 | 72.19 | 37498 | 12 | 16 | 126769065 | 134962737 | 139234417 | 141957777 | 141860001 | 134030345 | 162725537 | 123957129 | 115921097 | 150171969 | 142516529 | 130767921 | 138743745 | 120799473 | 153149585 |
| Q61990 | Pcbp2 | Poly(rC)-binding protein 2 Mus musculus | 925.26 | 72.65 | 38222 | 13 | 16 | 81268449 | 84730753 | 87683425 | 85525457 | 86003377 | 85641201 | 98829601 | 85797289 | 84396657 | 91557745 | 85617737 | 87460529 | 87217729 | 79557625 | 89456113 |
| Q91ZA3 | Pcca | Propionyl-CoA carboxylase alpha chain, mitochondrial Mus musculus | 284.25 | 11.05 | 79922 | 5 | 5 | 3053064 | 3140877.5 | 3337323 | 3037756.5 | 3906972.5 | 3092826.25 | 3604976.25 | 3933884 | 3788351.5 | 3806554.75 | 2797969.75 | 2887171 | 3605320 | 2225906 | 2628975.5 |
| Q99MN9 | Pccb | Propionyl-CoA carboxylase beta chain, mitochondrial Mus musculus | 479.37 | 22.18 | 58409 | 8 | 8 | 10446075 | 9421685 | 8901953 | 10852820 | 9400124 | 10429771 | 12174366 | 11066493 | 8598584 | 10226291 | 7567857 | 10712431 | 8551633 | 9077052 | 9414729 |
| Q8BFV2 | Pcid2 | PCI domain-containing protein 2 Mus musculus | 113.78 | 8.77 | 46132 | 2 | 2 | 474814.62 | 199473.06 | 396730.5275 | 452961.81 | 459672.62 | 1 | 1 | 1 | 1 | 1 | 215465.2 | 132868.91 | 215523.865 | 266739.91 | 247021.44 |
| P59114 | Pcif1 | Phosphorylated CTD-interacting factor 1 Mus musculus | 85.52 | 4.11 | 80504 | 2 | 2 | 453812.1875 | 478226.53 | 461242.25 | 408870.56 | 466909.41 | 580768.25 | 415200.75 | 518692.765 | 527837.94 | 550964.12 | 496264.84 | 322581.84 | 201567.19 | 448907.84 | 476627.97 |
| Q8BH04 | Pck2 | Phosphoenolpyruvate carboxykinase [GTP], mitochondrial Mus musculus | 1132.38 | 40.47 | 70528 | 19 | 19 | 44999569 | 51270049 | 49064105 | 55250529 | 52849193 | 48753889 | 58527389 | 53439625 | 54435153 | 59236605 | 44582397 | 47537873 | 50137173 | 47702097 | 54090617 |
| P17918 | Pcna | Proliferating cell nuclear antigen Mus musculus | 1091 | 85.06 | 28785 | 16 | 17 | 197016353 | 229116033 | 214982929 | 200755601 | 222280225 | 197876129 | 173251697 | 203184577 | 226266177 | 194732721 | 234645121 | 217275521 | 238552545 | 228498561 | 211049873 |
| Q9CQF9 | Pcyox1 | Prenylcysteine oxidase Mus musculus | 382.06 | 23.76 | 56495 | 7 | 7 | 15368321 | 16023936 | 15011778 | 10549521 | 14543288 | 14747302 | 12421434 | 16639123 | 18194537 | 12806388 | 12610264 | 14112996 | 13992239 | 16243966 | 12700599 |
| Q8C7K6 | Pcyox1l | Prenylcysteine oxidase-like Mus musculus | 121.79 | 6.87 | 54875 | 3 | 3 | 4274521 | 1098993.75 | 7082154 | 5048351 | 3787326 | 3970868 | 4964274 | 4390816 | 558014.62 | 5236973.5 | 3014650 | 2222518 | 1221117.25 | 2413953.75 | 1716722.5 |
| Q922E4 | Pcyt2 | Ethanolamine-phosphate cytidylyltransferase Mus musculus | 202.85 | 12.62 | 45235 | 4 | 4 | 13041319 | 10575002 | 12158069 | 11861909 | 9189696 | 11370640 | 12197076 | 9411295 | 10880722 | 11579171 | 12578533 | 14121734 | 9226862 | 11209390 | 13220027 |
| Q3UHX2 | Pdap1 | 28 kDa heat- and acid-stable phosphoprotein Mus musculus | 275.21 | 33.15 | 20605 | 5 | 5 | 5561523.5 | 6001897.5 | 3730673.5 | 6465843.5 | 5753828 | 4777719 | 5764064.5 | 4793592.5 | 4065350 | 6378303.5 | 4877971 | 4946721 | 5915897.5 | 3045336 | 4346222.5 |
| Q6NX65 | Pdcd10 | Programmed cell death protein 10 Rattus norvegicus 2 | 102.98 | 10.48 | 24355 | 2 | 2 | 5014375 | 1446141.75 | 3302033.75 | 1264186.88 | 2664207.75 | 3335885.5 | 3847736 | 3312089.5 | 1368679.12 | 3639177 | 1292267.12 | 3291349.5 | 3824410.5 | 3349758.25 | 1371853 |
| Q6NS46 | Pdcd11 | Protein RRP5 homolog Mus musculus | 1027.78 | 18.37 | 207779 | 21 | 21 | 11391401 | 10210932 | 14812261 | 10521552 | 6393869 | 15496319 | 15429837 | 13808677 | 15560133 | 12267632 | 18562757 | 16093093 | 13425174 | 8873012 | 13902959 |
| P56812 | Pdcd5 | Programmed cell death protein 5 Mus musculus | 122.84 | 19.84 | 14275 | 2 | 2 | 2714823 | 1559986 | 2663629.25 | 3307334.5 | 2622942.5 | 2687395 | 3136329 | 3029677 | 2446550.25 | 3265645.25 | 2257954.5 | 2628888.5 | 2903420 | 2445540.75 | 1704927.88 |
| P12815 | Pdcd6 | Programmed cell death protein 6 Mus musculus | 331.17 | 47.12 | 21867 | 6 | 6 | 5503205 | 6544410 | 5117478.5 | 4584561 | 7744203 | 5939343 | 4472012 | 5010839.5 | 4405043.5 | 3720162.25 | 6762976 | 4633161.5 | 4541607 | 4169054.25 | 5135505 |
| Q9WU78 | Pdcd6ip | Programmed cell death 6-interacting protein Mus musculus | 1562.22 | 41.2 | 96024 | 25 | 25 | 45678109 | 52402553 | 50640681 | 49440153 | 49380501 | 48651941 | 55650861 | 48960941 | 48965413 | 51615701 | 49081457 | 52271469 | 47856049 | 48654301 | 53462165 |
| Q8BVF2 | Pdcl3 | Phosducin-like protein 3 Mus musculus | 303.36 | 32.08 | 27581 | 6 | 6 | 4534774 | 4207844.5 | 4094438 | 4482828.5 | 4854951 | 5804317 | 4771687 | 4488003.5 | 3847716.75 | 5077696.5 | 5193918 | 5541845 | 4838930 | 4356271 | 4702395 |

| | | | | | | | | | | | | | | | | | | | | | | |
|---|---|---|---|---|---|---|---|---|---|---|---|---|---|---|---|---|---|---|---|---|---|---|
| Q3TIU4 | Pde12 | 2',5'-phosphodiesterase 12 Mus musculus | 249.54 | 9.38 | 67514 | 4 | 4 | 3170236 | 3794616.75 | 3080938.25 | 3403213 | 2253006.5 | 3399504.5 | 3503783.5 | 3018901.25 | 3810431.5 | 3401960 | 3380251 | 3160529.5 | 3744623 | 3525713.5 | 3200394.25 |
| O55057 | Pde6d | Retinal rod rhodopsin-sensitive cGMP 3',5'-cyclic phosphodiesterase subunit delta Mus | 41.04 | 8 | 17348 | 1 | 1 | 598764.94 | 946548.19 | 794202.75 | 870121.38 | 681159.19 | 802037.38 | 676463.69 | 954305.94 | 980659.81 | 778993.75 | 493388.72 | 879536.75 | 754458.12 | 975874.31 | 617405.5 |
| Q9D051 | Pdhb | Pyruvate dehydrogenase E1 component subunit beta, mitochondrial Mus musculus | 459.56 | 37.88 | 38937 | 10 | 10 | 11917652 | 15018408 | 13561663 | 12633189 | 14306156 | 11905609 | 16736098 | 14493065 | 13732016 | 16171836 | 12452775 | 14224782 | 14042631 | 12736000 | 13153180 |
| Q8BKZ9 | Pdhx | Pyruvate dehydrogenase protein X component, mitochondrial Mus musculus | 164.94 | 8.38 | 53999 | 4 | 4 | 5800042.5 | 6961759.5 | 6824678 | 5991619 | 6342486 | 6496799 | 6415422.5 | 6777571.5 | 7296752 | 5855310 | 5176232 | 5586012 | 6445381 | 5677110.5 | 3986162.5 |
| P27773 | Pdia3 | Protein disulfide-isomerase A3 Mus musculus | 1767.9 | 60.2 | 56678 | 30 | 30 | 254216289 | 217690513 | 247481441 | 249726273 | 236587809 | 240429697 | 311219137 | 264819489 | 234764929 | 269871681 | 263540177 | 274125697 | 240323441 | 242135521 | 268809057 |
| P08003 | Pdia4 | Protein disulfide-isomerase A4 Mus musculus | 1869.14 | 53.61 | 71982 | 32 | 32 | 53638141 | 41823233 | 54634405 | 44436601 | 43733833 | 50848017 | 64155573 | 57014573 | 52986641 | 50704353 | 55308045 | 60637233 | 45910577 | 55917929 | 62759317 |
| Q922R8 | Pdia6 | Protein disulfide-isomerase A6 Mus musculus | 940.22 | 42.05 | 48100 | 15 | 15 | 100966177 | 86687601 | 92713761 | 90096721 | 83267161 | 96363057 | 141952177 | 103985865 | 86219113 | 113482169 | 113697969 | 108514425 | 93133545 | 92450225 | 99606369 |
| Q63065 | Pdk1 | [Pyruvate dehydrogenase (acetyl-transferring)] kinase isozyme 1, mitochondrial Rattus r | 190.83 | 11.52 | 49081 | 3 | 3 | 2626979.5 | 3395204 | 1657055.38 | 2282470.5 | 1776084.12 | 3586100.5 | 3640269.5 | 3950429 | 3601029 | 3254742 | 2454559.5 | 2917510.5 | 3570620.5 | 3743815.75 | 2720999.5 |
| Q922H2 | Pdk3 | [Pyruvate dehydrogenase (acetyl-transferring)] kinase isozyme 3, mitochondrial Mus mu | 309.05 | 13.49 | 47923 | 4 | 4 | 4694207 | 4650144 | 4087162.75 | 3807998.25 | 4096631 | 4182480.25 | 4416773 | 5227752 | 4265907.5 | 4275591.5 | 1851385.25 | 4009467 | 4798117 | 2721006.75 | 2279022 |
| Q8CI51 | Pdlim5 | PDZ and LIM domain protein 5 Mus musculus | 353.4 | 16.41 | 63299 | 7 | 7 | 2167191 | 1759710.75 | 2945875.5 | 1207917.75 | 1192635.38 | 1377900.38 | 5103851 | 1475628.25 | 1591000.12 | 2721514.25 | 1790508 | 2610265 | 1342065 | 1541228.38 | 3823051 |
| A4L9P7 | Pds5a | Sister chromatid cohesion protein PDS5 homolog A Rattus norvegicus | 296.95 | 4.95 | 150286 | 4 | 5 | 3646718.5 | 2701575 | 2707299.5 | 2592853 | 3371456.25 | 3518119.5 | 2819983.5 | 2868081 | 3077248 | 3003082.5 | 3587469.75 | 3112363 | 2387251.5 | 2477733.5 | 4136025.25 |
| Q4VA53 | Pds5b | Sister chromatid cohesion protein PDS5 homolog B Mus musculus | 121.11 | 1.66 | 164419 | 1 | 2 | 1654523.75 | 1661654.88 | 1050859.38 | 1462088.25 | 2150609.5 | 1805046.88 | 1114708.5 | 1451093 | 1298351.75 | 1066333.75 | 1540426.62 | 932759.06 | 1411877.38 | 1152293.5 | 1549837.88 |
| Q99K01 | Pdxdc1 | Pyridoxal-dependent decarboxylase domain-containing protein 1 Mus musculus | 546.27 | 19.31 | 87335 | 11 | 11 | 5451270.5 | 4473760.5 | 5188518 | 6987475 | 8213314 | 7161334.5 | 6952363 | 5694870.5 | 5344612.5 | 7398951 | 5620555 | 5149570.5 | 6429734 | 6043642.5 | 7496766 |
| Q8K183 | Pdxk | Pyridoxal kinase Mus musculus | 664.85 | 28.53 | 35015 | 9 | 11 | 15056884 | 15631310 | 21250323 | 17656895 | 14409173 | 15400815 | 16286347 | 15696088 | 17043277 | 13467835 | 18192925 | 19914269 | 13277689 | 17190093 | 21917431 |
| Q62048 | Pea15 | Astrocytic phosphoprotein PEA-15 Mus musculus | 285.53 | 56.15 | 15054 | 6 | 6 | 1709723.25 | 27973147 | 20755053 | 16688222 | 19577303 | 22993537 | 17957279 | 20255787 | 17088371 | 19629989 | 1871706.75 | 2063850.88 | 736474.62 | 21995275 | 22223667 |
| Q69Z38 | Peak1 | Inactive tyrosine-protein kinase PEAK1 Mus musculus | 91.36 | 1.56 | 191097 | 2 | 2 | 693472.88 | 723881.19 | 1 | 1 | 534121.5 | 752258.44 | 412462.81 | 700686.7175 | 915729.06 | 722296.56 | 521593.75 | 619113.38 | 596712.19 | 1 | 1 |
| P70296 | Pebp1 | Phosphatidylethanolamine-binding protein 1 Mus musculus | 293.22 | 44.39 | 20830 | 5 | 5 | 9264044 | 9414791 | 8561913 | 9974959 | 11958726 | 9556189 | 10389513 | 8852047 | 10011603 | 9036119 | 9144397 | 9028508 | 12260143 | 8064141.5 | 15467518 |
| Q99MZ7 | Pecr | Peroxisomal trans-2-enoyl-CoA reductase Mus musculus | 78.69 | 2.97 | 32410 | 1 | 1 | 1798545.25 | 1351021.75 | 1489071.88 | 1 | 1326310.88 | 1226157.75 | 1690157 | 1520593.5 | 1279635.5 | 1824094.88 | 1674253.75 | 1602167.38 | 1174060.75 | 1164959.12 | 1375250.25 |
| Q8BFY6 | Pef1 | Peflin Mus musculus | 177.9 | 17.45 | 29228 | 4 | 4 | 685249.44 | 1989509 | 3559292.5 | 3504781.5 | 3390456 | 1875611.5 | 2788645.5 | 4517974.5 | 3886296.25 | 3845501.5 | 981306.38 | 4281820 | 3628056.75 | 5092175 | 3503236 |
| Q5XIP1 | Pelo | Protein pelota homolog Rattus norvegicus | 47.91 | 4.68 | 43407 | 1 | 1 | 472490.75 | 613910.62 | 1 | 1 | 286769.41 | 570445 | 224521.73 | 1 | 1 | 1 | 485744.81 | 565199.3825 | 610615.81 | 738342 | 426094.91 |
| Q9DBD5 | Pelp1 | Proline-, glutamic acid- and leucine-rich protein 1 Mus musculus | 331.84 | 4.81 | 118069 | 5 | 5 | 3186774 | 4325926 | 4950494 | 5275342.5 | 3836460.5 | 3901770.5 | 2983257 | 3777625.25 | 6271977 | 3527321 | 2781888.25 | 3746554 | 3666092.75 | 5601257 | 1388681.5 |
| Q11136 | Pepd | Xaa-Pro dipeptidase Mus musculus | 711.8 | 34.48 | 55029 | 15 | 15 | 33498953 | 32399651 | 30950619 | 29717185 | 25378425 | 25806581 | 24168087 | 32270591 | 30796269 | 19764191 | 27278177 | 26117735 | 28496295 | 35173077 | 23716789 |
| Q9EQ61 | Pes1 | Pescadillo homolog Mus musculus | 723.77 | 29.45 | 67796 | 16 | 16 | 9817133 | 8869289 | 7449325 | 7575934 | 7898886.5 | 9236557 | 5917568.5 | 9474148 | 8752657 | 7414217.5 | 9799547 | 7813963.5 | 7954650.5 | 9817062 | 7916840 |
| Q8VCI5 | Pex19 | Peroxisomal biogenesis factor 19 Mus musculus | 53.67 | 9.7 | 32733 | 1 | 1 | 170118.98 | 1 | 228473.72 | 1 | 181924.33 | 280168.62 | 177072.7 | 1 | 1 | 389645.22 | 138085.31 | 1 | 325639.84 | 177225.8 | 1 |
| Q5SUR0 | Pfas | Phosphoribosylformylglycinamidine synthase Mus musculus | 1320.05 | 22.51 | 144629 | 19 | 19 | 22435121 | 30871269 | 25488663 | 34036005 | 26040129 | 28697383 | 31782669 | 25659697 | 20927573 | 30960595 | 28387881 | 22996063 | 35949649 | 28792621 | 29439479 |
| Q9CWM4 | Pfdn1 | Prefoldin subunit 1 Mus musculus | 61.53 | 9.02 | 14255 | 1 | 1 | 2782856.25 | 2875558.25 | 2645507 | 3131745.75 | 2722431 | 2910236.5 | 2586206 | 2909724.5 | 1865114.25 | 3186527 | 2148616.75 | 2516925.25 | 2740792 | 2574439 | 1983105.62 |
| B0BN18 | Pfdn2 | Prefoldin subunit 2 Rattus norvegicus | 233.75 | 29.22 | 16580 | 4 | 4 | 12981779 | 15286921 | 14376549 | 16331211 | 13527823 | 13015427 | 11610633 | 13775427 | 13256202 | 12146410 | 11573621 | 12970065 | 13288693 | 13611364 | 10168105 |
| Q9WU28 | Pfdn5 | Prefoldin subunit 5 Mus musculus | 227.71 | 33.12 | 17356 | 3 | 3 | 2941656 | 1710702.12 | 3679333.5 | 1357294.38 | 1450175 | 2322226 | 2433837.75 | 3704707.5 | 4438666 | 1171198.5 | 4991814.5 | 3471042.25 | 1247512.38 | 3918660.5 | 3077188.5 |
| Q03958 | Pfdn6 | Prefoldin subunit 6 Mus musculus | 58.42 | 9.45 | 14455 | 1 | 1 | 1665957 | 1816728.12 | 1824478.53 | 2297962.75 | 1517266.25 | 1382403.25 | 1530436.88 | 1854146.88 | 1597139.065 | 1621569.25 | 1187155.38 | 1751231.38 | 1405580.38 | 1418356 | 933757.81 |
| P12382 | Pfkl | ATP-dependent 6-phosphofructokinase, liver type Mus musculus | 576.84 | 22.82 | 85360 | 10 | 12 | 19641895 | 16978521 | 16034875 | 13188097 | 19513529 | 21386235 | 20890561 | 20718255 | 22054261 | 18848263 | 19522317 | 17954289 | 17573799 | 20557623 | 18941789 |
| P47857 | Pfkm | ATP-dependent 6-phosphofructokinase, muscle type Mus musculus | 106.5 | 3.85 | 85269 | 2 | 3 | 1968725.25 | 1618983.75 | 1741868.75 | 1884924 | 2017922.5 | 1977161.62 | 1760814.5 | 1691696 | 1979585 | 1846418.5 | 1541518.75 | 1748068.25 | 1828702.75 | 2024954.5 | 1681966.5 |
| Q9WUA3 | Pfkp | ATP-dependent 6-phosphofructokinase, platelet type Mus musculus | 1812.23 | 54.97 | 85455 | 20 | 27 | 62324449 | 64117569 | 62208097 | 54761193 | 61724577 | 73752281 | 87211129 | 74364329 | 80531273 | 73031857 | 75789449 | 65038041 | 67673449 | 74192025 | 70099769 |
| P62962 | Pfn1 | Profilin-1 Mus musculus | 1060.25 | 94.29 | 14957 | 13 | 17 | 698839361 | 817273217 | 818383169 | 799125441 | 853652097 | 748273857 | 755138561 | 766024833 | 765827137 | 775051841 | 767106433 | 803243521 | 861669633 | 850744385 | 661077761 |
| P25113 | Pgam1 | Phosphoglycerate mutase 1 Rattus norvegicus | 1137.31 | 74.02 | 28832 | 15 | 16 | 160273809 | 158861713 | 135597505 | 130547473 | 140905953 | 185438881 | 210870913 | 170879073 | 152729809 | 166426593 | 168983073 | 153981697 | 170122593 | 155666209 | 161602097 |
| Q9DCD0 | Pgd | 6-phosphogluconate dehydrogenase, decarboxylating Mus musculus | 1812.8 | 69.57 | 53247 | 26 | 27 | 151253345 | 164736529 | 175545713 | 136175393 | 153375793 | 150455393 | 202020865 | 177817105 | 194957041 | 174516961 | 194816513 | 191027361 | 169827873 | 183155201 | 198447505 |
| Q8BP56 | Pgghg | Protein-glucosylgalactosylhydroxylysine glucosidase Mus musculus | 52.99 | 2.9 | 76488 | 1 | 1 | 323060.22 | 299136.78 | 216324.42 | 289850.87 | 320882.06 | 262628.84 | 1 | 396586.28 | 341662.59 | 1 | 261123.61 | 261400.8725 | 215417.02 | 327375.91 | 241686.95 |
| P09411 | Pgk1 | Phosphoglycerate kinase 1 Mus musculus | 2512.98 | 79.14 | 44550 | 26 | 36 | 841446529 | 935717121 | 922474369 | 795992001 | 906236481 | 1038448385 | 1169212673 | 984401217 | 1019757825 | 785129473 | 1041379649 | 927721537 | 1083609985 | 1050507585 | 913426945 |
| P09041 | Pgk2 | Phosphoglycerate kinase 2 Mus musculus | 505 | 17.03 | 44853 | 1 | 8 | 44837493 | 39106101 | 49576689 | 51804593 | 54221653 | 61544201 | 89815801 | 49908245 | 38082497 | 71106385 | 57031521 | 62745489 | 59985309 | 42624509 | 69505985 |
| Q9CQ60 | Pgls | 6-phosphogluconolactonase Mus musculus | 557.62 | 59.53 | 27254 | 8 | 8 | 49917609 | 63425085 | 49683365 | 18717755 | 50748305 | 48310433 | 46220497 | 53606933 | 48243793 | 61155649 | 53576193 | 57382545 | 55254961 | 51660417 | 46420601 |
| Q9D0F9 | Pgm1 | Phosphoglucomutase-1 Mus musculus | 1225.89 | 48.04 | 61418 | 18 | 18 | 24706457 | 27459615 | 26923959 | 22012243 | 24741713 | 28689809 | 32885623 | 30358749 | 32106349 | 26712269 | 30071991 | 29115397 | 28329307 | 30982065 | 27291501 |
| Q7TSV4 | Pgm2 | Phosphoglucomutase-2 Mus musculus | 85 | 7.74 | 68748 | 2 | 2 | 2201647.75 | 1594923.38 | 2156999.25 | 2122491.75 | 2083694 | 1835397 | 908060.5 | 1710602.38 | 1409135.62 | 1465798.875 | 559778.5 | 1733968.75 | 1853111.38 | 1709271.75 | 2048442.38 |
| Q9CYR6 | Pgm3 | Phosphoacetylglucosamine mutase Mus musculus | 58.22 | 2.58 | 59453 | 1 | 1 | 2303533 | 2253810.5 | 1989791.62 | 2480269.25 | 2497793.5 | 2364001.5 | 2141933 | 2449218.25 | 2056747 | 2434888.25 | 1587321.5 | 1774059.38 | 2460277.5 | 2464856.75 | 2199000.25 |
| Q8CHP8 | Pgp | Glycerol-3-phosphate phosphatase Mus musculus | 426.81 | 41.74 | 34541 | 9 | 9 | 19739845 | 10070202 | 22113225 | 7777014.5 | 9715953 | 8857305 | 10877085 | 9956388 | 16607823 | 25137255 | 21083985 | 19601761 | 10978472 | 9395607 | 24186333 |
| O55022 | Pgrmc1 | Membrane-associated progesterone receptor component 1 Mus musculus | 61.64 | 7.18 | 21694 | 1 | 1 | 2714451.25 | 2481273 | 3081889.75 | 3019477.25 | 3099087 | 2957255.5 | 3143471.75 | 2840541 | 2822718.25 | 3088255 | 2101849.5 | 2826110.25 | 3047052.25 | 2915827.5 | 2450738 |
| Q80UU9 | Pgrmc2 | Membrane-associated progesterone receptor component 2 Mus musculus | 254.4 | 21.66 | 23334 | 3 | 3 | 3782386.25 | 3627643.5 | 3018508.25 | 3366951 | 3852169.25 | 3638149.25 | 5009959.5 | 4183041.25 | 3090780 | 3506060 | 2404666.75 | 3611190.75 | 4002143.5 | 3821570.5 | 3256571.25 |
| Q8BHF7 | Pgs1 | CDP-diacylglycerol--glycerol-3-phosphate 3-phosphatidyltransferase, mitochondrial Mus | 83.58 | 5.61 | 62489 | 2 | 2 | 679683.38 | 826181.75 | 691918.06 | 720012.625 | 682267.31 | 619187.5925 | 537546.5 | 656831.31 | 806267.56 | 476105 | 481501.72 | 1 | 619129.12 | 1 | 447023.5 |
| Q63068 | Phax | Phosphorylated adapter RNA export protein Rattus norvegicus | 131.34 | 5.71 | 43166 | 2 | 2 | 2160803.25 | 578286.62 | 1998157 | 710178.44 | 825671 | 1755255.5 | 783618.81 | 1276202 | 1337462.12 | 935399.5 | 631017.94 | 1778799.25 | 1928691.62 | 2422027.75 | 1003025.19 |
| P67778 | Phb | Prohibitin Mus musculus | 1423.85 | 81.62 | 29820 | 24 | 24 | 189354209 | 196653537 | 199541985 | 203101233 | 193904209 | 182458689 | 204987361 | 204365249 | 201918465 | 208902513 | 219182881 | 176241921 | 197794753 | 194013793 | 179348433 |
| O35129 | Phb2 | Prohibitin-2 Mus musculus | 910.13 | 53.18 | 33296 | 14 | 14 | 28280971 | 23675733 | 30372779 | 34590209 | 32233809 | 30741135 | 43965793 | 24006799 | 19113323 | 40497125 | 34712641 | 32182311 | 30683603 | 21488097 | 39415617 |
| Q8CHP6 | Phc3 | Polyhomeotic-like protein 3 Mus musculus | 41.81 | 1.33 | 105353 | 1 | 1 | 696666.44 | 526710.75 | 724392.88 | 424297.03 | 641194.19 | 611984.75 | 471048.72 | 730451.06 | 742972.62 | 502038.03 | 451680.56 | 605714.88 | 404022.44 | 1018380.31 | 495107.97 |
| A6H5X4 | Phf11 | PHD finger protein 11 Mus musculus | 71.1 | 10.39 | 37728 | 2 | 2 | 1 | 197548.05 | 936138.56 | 288671.03 | 1 | 630002.485 | 996176.5 | 254274.73 | 236154.33 | 1033404.38 | 724201.38 | 1 | 1 | 1 | 1078695.5 |
| P83870 | Phf5a | PHD finger-like domain-containing protein 5A Mus musculus | 325.89 | 47.27 | 12405 | 5 | 5 | 1582985.75 | 1990887.62 | 836719.88 | 1843662.25 | 2040853.38 | 1806416.12 | 855587.5 | 819317.88 | 1827957.88 | 796110 | 2510374.25 | 791392.31 | 1780945.5 | 2223900 | 2334701.5 |
| Q9DAK9 | Phpt1 | 14 kDa phosphohistidine phosphatase Mus musculus | 42.6 | 15.32 | 13997 | 1 | 1 | 505115.25 | 469146.16 | 1 | 480579.22 | 1 | 473992.305 | 472966.38 | 501259.78 | 424152.12 | 497590.94 | 514511.56 | 451532.03 | 1 | 473896.09 | 377637.66 |
| Q2TBE6 | Pi4k2a | Phosphatidylinositol 4-kinase type 2-alpha Mus musculus | 169.01 | 6.26 | 54258 | 3 | 3 | 2433654.75 | 1953824.25 | 2286650 | 2070547.25 | 2256479 | 1964162 | 1641800 | 2090900.25 | 1991144.25 | 1854194.25 | 1619286.5 | 1906954.25 | 1762642.5 | 1953715.38 | 2187275.5 |
| E9Q3L2 | Pi4ka | Phosphatidylinositol 4-kinase alpha Mus musculus | 125.61 | 1.9 | 237042 | 2 | 2 | 1 | 1 | 134357.41 | 1 | 1 | 152290.88 | 144313.59 | 156126.62 | 240124.73 | 173213.955 | 315343.44 | 310312.59 | 138144.89 | 1 | 161621.23 |
| Q7M6Y3 | Picalm | Phosphatidylinositol-binding clathrin assembly protein Mus musculus | 827.62 | 35.3 | 71543 | 1 | 16 | 25045929 | 29878061 | 27825441 | 24341813 | 26191467 | 27421983 | 25896903 | 29508549 | 31852605 | 27655693 | 28359787 | 26586489 | 28228931 | 30963481 | 18745827 |
| Q9CXY9 | Pigk | GPI-anchor transamidase Mus musculus | 98.46 | 5.82 | 44895 | 2 | 2 | 2596389.5 | 2604529.5 | 2888618.5 | 2876564 | 2786547.75 | 2933369.75 | 2748335 | 2954459 | 3114773.75 | 3042729 | 2533728.5 | 1891810.5 | 2006862.38 | 2601223.75 | 2926023 |
| Q9R1S3 | Pign | GPI ethanolamine phosphate transferase 1 Mus musculus | 162.1 | 4.3 | 105045 | 3 | 3 | 2485611.5 | 1528267.25 | 2648687.5 | 1693881.25 | 1762883.62 | 2018195.75 | 3033967 | 2122479 | 3358870.75 | 2178967 | 2207121 | 1884504.75 | 2717090.5 | 2980519.5 | 1502473.5 |
| Q6PD26 | Pigs | GPI transamidase component PIG-S Mus musculus | 284.3 | 12.97 | 61711 | 5 | 5 | 2806845.25 | 2095252.75 | 1947335.75 | 2793377.5 | 2648203 | 3023653 | 2797078 | 2713289.5 | 3039603 | 2713424.5 | 2018872.75 | 2493429.75 | 2706039.75 | 2561288.75 | 2671496 |
| Q8BXQ2 | Pigt | GPI transamidase component PIG-T Mus musculus | 90.4 | 5.5 | 65705 | 2 | 2 | 1827111.88 | 2020421.38 | 1231352.25 | 1495157.5 | 1825833.12 | 1833093 | 1357298.62 | 1562264.38 | 1841414.75 | 1628401.12 | 2088555.12 | 1123268.62 | 1810981.5 | 1482783.12 | 1620518.88 |
| Q8CHJ1 | Pigu | Phosphatidylinositol glycan anchor biosynthesis class U protein Rattus norvegicus | 55.11 | 3.91 | 49953 | 1 | 1 | 728036.62 | 1 | 1 | 1 | 1 | 302314.78 | 275532.12 | 257988.7 | 1 | 1 | 375064 | 1 | 1 | 1 | 262145.12 |
| Q9EQ32 | Pik3ap1 | Phosphoinositide 3-kinase adapter protein 1 Mus musculus | 769.93 | 23.06 | 90928 | 14 | 14 | 16486035 | 22704965 | 18916209 | 17843853 | 24338759 | 17378197 | 17102133 | 20054475 | 27634521 | 13959258 | 26033513 | 24045835 | 20438115 | 26597849 | 20869101 |
| Q8BTI9 | Pik3cb | Phosphatidylinositol 4,5-bisphosphate 3-kinase catalytic subunit beta isoform Mus musc | 54.26 | 1.5 | 121711 | 1 | 1 | 528539.31 | 442406.34 | 1 | 1 | 1 | 700645.56 | 1 | 697379.31 | 691684.5 | 1 | 558780.56 | 583713.38 | 424283.34 | 940180.88 | 614179.69 |
| Q9JHG7 | Pik3cg | Phosphatidylinositol 4,5-bisphosphate 3-kinase catalytic subunit gamma isoform Mus mu | 51.14 | 0.82 | 126400 | 1 | 1 | 939349.38 | 814353.44 | 1030546.19 | 1136400.25 | 1166521 | 811272.69 | 590112.06 | 736915.25 | 925851.88 | 690882.62 | 700723.75 | 894396.81 | 883107.5 | 835333.25 | 956282.25 |
| P26450 | Pik3r1 | Phosphatidylinositol 3-kinase regulatory subunit alpha Mus musculus | 44.51 | 1.66 | 83517 | 1 | 1 | 983763.25 | 863371.75 | 797035.06 | 946502.44 | 897668.125 | 953726.31 | 826178.56 | 932935.31 | 687935.88 | 891840.94 | 859455.5 | 959882.06 | 1040639.19 | 746381.5 | 1184095 |
| P0C0R5 | Pik3r4 | Phosphoinositide 3-kinase regulatory subunit 4 Rattus norvegicus 3 | 197.14 | 3.83 | 152447 | 4 | 4 | 3335696 | 2340216 | 3167534.25 | 2253984.75 | 3034508 | 2819406.75 | 3010763.5 | 3258840.25 | 3044306 | 2187434.5 | 2727116.5 | 3115201 | 2923615.5 | 3083591.25 | 2836881.5 |
| Q9QUR7 | Pin1 | Peptidyl-prolyl cis-trans isomerase NIMA-interacting 1 Mus musculus | 200.2 | 29.7 | 18370 | 3 | 3 | 4130590 | 5457612.5 | 5450382 | 3411387 | 5057563 | 4617024 | 3665084.25 | 4275717 | 4757846.5 | 4430607.5 | 4198293 | 4565831.5 | 5507673 | 4517691.5 | 5170160.5 |
| O70172 | Pip4k2a | Phosphatidylinositol 5-phosphate 4-kinase type-2 alpha Mus musculus | 133.09 | 9.14 | 46152 | 2 | 3 | 1924650.88 | 2897249.5 | 3319645.5 | 2115855.75 | 2787880.5 | 2529179 | 2064898.25 | 3775024 | 4157457 | 2107762.25 | 1814586.25 | 3225287.5 | 1847329.5 | 3816774.25 | 2787624.75 |
| O88377 | Pip4k2b | Phosphatidylinositol 5-phosphate 4-kinase type-2 beta Rattus norvegicus | 74.26 | 6.25 | 47264 | 1 | 2 | 2861611.25 | 3089132 | 3489054 | 2074659 | 1855538.5 | 2971298.25 | 2622971 | 3534754 | 3296179.5 | 1924916.38 | 2746778.75 | 2468917.25 | 3150129.25 | 2855068 | 2907325.5 |
| D3ZSI8 | Pip5k1a | Phosphatidylinositol 4-phosphate 5-kinase type-1 alpha Rattus norvegicus | 50.39 | 2.56 | 60605 | 1 | 1 | 1118438.75 | 1 | 1222851.75 | 891078.56 | 1 | 932422.81 | 1 | 811022.12 | 889182.81 | 1 | 1 | 1131298.75 | 1064961.75 | 1086167.88 | 1 |
| P16446 | Pitpna | Phosphatidylinositol transfer protein alpha isoform Rattus norvegicus | 535.86 | 45.02 | 31907 | 10 | 10 | 12655302 | 10579984 | 11744957 | 8875902 | 8547021 | 10711872 | 9562581 | 8863426 | 14078211 | 7666662.5 | 12961364 | 13508403 | 8527246 | 13721389 | 13989385 |
| P53811 | Pitpnb | Phosphatidylinositol transfer protein beta isoform Mus musculus | 267.34 | 33.21 | 31487 | 7 | 7 | 4186602 | 3557856 | 3335846 | 3566810 | 4621709.5 | 2527672 | 5778245.5 | 5328751 | 4341744 | 4356838 | 3907344.5 | 5409830 | 3556324.25 | 3733740.5 | 5506708.5 |
| Q8K411 | Pitrm1 | Presequence protease, mitochondrial Mus musculus | 392.16 | 12.64 | 117372 | 8 | 8 | 7504507 | 9185966 | 9678748 | 8046197 | 8562303 | 7538425 | 9263572 | 9977348 | 9112857 | 8088358 | 9939912 | 9032827 | 8581021 | 8477862 | 6613596.5 |
| P52480 | Pkm | Pyruvate kinase PKM Mus musculus | 3772.789 | 77.02 | 57845 | 13 | 56 | 1353953409 | 1675794817 | 1641204481 | 1688209025 | 1613781761 | 1724068737 | 2174259713 | 1714558721 | 1596637569 | 1949092737 | 1848550913 | 1820358273 | 1680236033 | 1617761921 | 2053767425 |
| P70268 | Pkn1 | Serine/threonine-protein kinase N1 Mus musculus | 434.14 | 13.11 | 104411 | 9 | 9 | 3849104.75 | 3042281.75 | 3887365.75 | 4210186 | 2088987.25 | 1772087.5 | 4412052 | 3391952.5 | 2648734 | 3953073 | 3231235 | 4280587 | 3393062 | 2985815.5 | 3916131.5 |
| Q8VEB4 | Pla2g15 | Group XV phospholipase A2 Mus musculus | 229.78 | 17.96 | 47307 | 6 | 6 | 9589357 | 8330239 | 7938845.5 | 8285196 | 8603706 | 6980135 | 7484498.5 | 9988698 | 8190934 | 7648682 | 6550774 | 6992050 | 8497103 | 10025213 | 6184650.5 |
| P47713 | Pla2g4a | Cytosolic phospholipase A2 Mus musculus | 977.38 | 35.16 | 85222 | 19 | 19 | 30164811 | 22980913 | 25101831 | 18467209 | 23935063 | 25609927 | 37088273 | 26936509 | 26041617 | 28118429 | 27520659 | 29129017 | 25722145 | 24098527 | 36919637 |
| P27612 | Plaa | Phospholipase A-2-activating protein Mus musculus | 763.86 | 26.45 | 87221 | 14 | 14 | 6908505 | 8042811 | 8340288 | 9604593 | 11049423 | 11503360 | 11235243 | 11187209 | 8149116 | 13860019 | 12879563 | 12492611 | 11326172 | 9749513 | 10956633 |
| Q3TCN2 | Plbd2 | Putative phospholipase B-like 2 Mus musculus | 514.5 | 21.72 | 66289 | 10 | 10 | 10900258 | 13685235 | 8361534 | 13256259 | 15092123 | 15266032 | 13058971 | 15292061 | 12682149 | 11809541 | 9894554 | 12179119 | 14370970 | 14162646 | 10129503 |
| P51432 | Plcb3 | 1-phosphatidylinositol 4,5-bisphosphate phosphodiesterase beta-3 Mus musculus | 235.68 | 7.54 | 139487 | 5 | 5 | 870095.25 | 1536200 | 1617768.75 | 974266.44 | 1494747.25 | 2052958.38 | 2355656.75 | 1938502.5 | 1166802.75 | 2317917.75 | 2077326 | 1854785.5 | 1573709.75 | 2019036.62 | 2411529.5 |
| Q8CIH5 | Plcg2 | 1-phosphatidylinositol 4,5-bisphosphate phosphodiesterase gamma-2 Mus musculus | 905.48 | 20.32 | 147592 | 20 | 20 | 19160359 | 20443325 | 19328189 | 19770709 | 21158481 | 19989227 | 20750949 | 19838119 | 21604703 | 18449055 | 24070763 | 20393471 | 20709883 | 20599931 | 21991593 |
| O35405 | Pld3 | Phospholipase D3 Mus musculus | 139.88 | 11.27 | 54389 | 3 | 3 | 9169067 | 7480470.5 | 6721049 | 9324024 | 8638230 | 5449954.5 | 5937753.5 | 6070406.5 | 4910840 | 6251654 | 6166208 | 3916503.25 | 5830383 | 5306693 | 4008033 |
| Q8BG07 | Pld4 | Phospholipase D4 Mus musculus | 1299.01 | 42.94 | 56154 | 16 | 19 | 217774625 | 283700417 | 237069425 | 208314785 | 226738369 | 195780433 | 147569569 | 212401057 | 248510593 | 170943409 | 175277409 | 176572001 | 217984177 | 226305905 | 134701857 |
| Q9QXS1 | Plec | Plectin Mus musculus | 10489.382 | 46.9 | 534188 | 183 | 188 | 545612481 | 526952289 | 559298369 | 612632001 | 553254721 | 511777441 | 555931905 | 538400257 | 507956833 | 526475041 | 571951233 | 553687937 | 538793921 | 530393473 | 527144097 |
| Q9JHK5 | Plek | Pleckstrin Mus musculus | 743.29 | 43.14 | 39901 | 11 | 11 | 18507219 | 18363019 | 16011034 | 13170645 | 19258467 | 20944603 | 17638267 | 20701035 | 21860819 | 16437377 | 20468499 | 19960031 | 18381253 | 20538555 | 18891369 |
| Q9ERS5 | Plekha2 | Pleckstrin homology domain-containing family A member 2 Mus musculus | 146.21 | 8.94 | 47380 | 3 | 3 | 1054817.12 | 738206.06 | 254992.8 | 786830.62 | 783559.5 | 1666901.25 | 1384205.25 | 1880943 | 778793.5 | 1070753 | 739442.62 | 889524.25 | 2129786.5 | 836233.44 | 1651463 |
| Q8K124 | Plekho2 | Pleckstrin homology domain-containing family O member 2 Mus musculus | 766.48 | 46.87 | 53872 | 13 | 13 | 17818069 | 14481283 | 20974085 | 19805351 | 20332395 | 18973539 | 24121361 | 18876175 | 12854926 | 22297671 | 21667189 | 21806961 | 21331027 | 16788137 | 22995727 |
| P43883 | Plin2 | Perilipin-2 Mus musculus | 38.59 | 3.76 | 46646 | 1 | 1 | 1331561.88 | 1921657.12 | 2026767.25 | 1684054.12 | 1694100.12 | 1281032.88 | 1414913.5 | 1548605.38 | 1822210.62 | 1405844.12 | 1271895.5 | 1028179.5 | 1673374.62 | 1635060.25 | 1130297.38 |
| Q9DBG5 | Plin3 | Perilipin-3 Mus musculus | 899.51 | 55.38 | 47262 | 13 | 14 | 63572549 | 62403309 | 57810949 | 82654337 | 80980849 | 59821193 | 66896361 | 59893113 | 45320541 | 78346441 | 66281445 | 43557129 | 66549169 | 55152493 | 53599833 |
| Q07832 | Plk1 | Serine/threonine-protein kinase PLK1 Mus musculus | 176.32 | 8.13 | 68301 | 4 | 4 | 661800.31 | 1368742.25 | 1096273.25 | 1421582.62 | 593047.19 | 535434.81 | 285001.69 | 573119.75 | 1437363.75 | 649158.5 | 258292.77 | 596445.75 | 699260 | 1 | 1 |
| Q9R0E2 | Plod1 | Procollagen-lysine,2-oxoglutarate 5-dioxygenase 1 Mus musculus | 218.15 | 8.79 | 83595 | 5 | 5 | 1740708.88 | 996182.12 | 2236154.5 | 505680.28 | 1145521.25 | 1213544.5 | 1967818 | 2080649.38 | 2748213.5 | 1073571 | 2368922.75 | 2646561.25 | 1286637.5 | 2373125.75 | 2656902.5 |
| Q9R0E1 | Plod3 | Multifunctional procollagen lysine hydroxylase and glycosyltransferase LH3 Mus muscul | 424.41 | 14.98 | 84922 | 8 | 8 | 5364478 | 5032839.5 | 5109065.5 | 4433972 | 4825268 | 5005546.5 | 4407895 | 3857017.75 | 2831055 | 4861467 | 5855996 | 4540805.5 | 3819171 | 3991320.5 | 4360407 |
| Q9Z2Y8 | Plpbp | Pyridoxal phosphate homeostasis protein Mus musculus | 97.09 | 8.03 | 30049 | 2 | 2 | 2219025.25 | 2299979.75 | 2153731.75 | 2477237.75 | 2008060.75 | 2101538 | 1939842.75 | 1925310.62 | 2005135.62 | 2053876.38 | 1700573.5 | 2400256.75 | 2018431.38 | 2054112.38 | 2065341.62 |
| Q922V4 | Plrg1 | Pleiotropic regulator 1 Mus musculus | 336.99 | 16.37 | 56938 | 6 | 6 | 1673796.25 | 1381166.25 | 1512445.25 | 1609265 | 1577880.62 | 1759510.25 | 1639479 | 1497589.38 | 1423158.5 | 1301887.75 | 1439209.5 | 1343377.88 | 1516354.38 | 2754262.5 | 1990755.75 |
| Q3V0K9 | Pls1 | Plastin-1 Mus musculus | 239.73 | 7.46 | 70408 | 0 | 4 | 162932801 | 139718385 | 169041841 | 120631785 | 120178753 | 148299553 | 126135921 | 171022801 | 202128273 | 98668169 | 191923057 | 157791553 | 111919833 | 171208593 | 160788561 |
| Q99K51 | Pls3 | Plastin-3 Mus musculus | 1757.78 | 62.38 | 70742 | 23 | 32 | 428140929 | 438180577 | 453587393 | 434619361 | 434747297 | 478048801 | 420662689 | 478435649 | 552829761 | 413355105 | 521652705 | 468438081 | 413847201 | 495582721 | 486345697 |
| P58195 | Plscr1 | Phospholipid scramblase 1 Rattus norvegicus | 58.81 | 5.97 | 36711 | 1 | 1 | 1147784.62 | 1143446 | 1155475.38 | 1194112.88 | 1016785.81 | 1064462.12 | 1093437.88 | 1341492.12 | 907372.69 | 761047.12 | 938408.12 | 1109438.75 | 1211447.5 | 1002894 | 833585.88 |
| Q6QBQ4 | Plscr3 | Phospholipid scramblase 3 Rattus norvegicus 2 | 49.15 | 2.7 | 31646 | 1 | 1 | 1329259.75 | 887402.75 | 908221.69 | 739300.56 | 984196.69 | 922290.31 | 804533.69 | 1090882.12 | 1094262.38 | 1051713.25 | 775149.88 | 741093.38 | 1006034.56 | 780889.44 | 820545.06 |
| P70206 | Plxna1 | Plexin-A1 Mus musculus | 409.96 | 6.12 | 211099 | 7 | 10 | 7789652.5 | 11109825 | 8503231 | 8246097.5 | 9001116 | 9596887 | 6376457.5 | 10837247 | 5927987 | 9208995 | 7481957.5 | 3697516.75 | 7450620.5 | 9742844 | 6114407.5 |
| Q80UG2 | Plxna4 | Plexin-A4 Mus musculus | 152.08 | 2.06 | 212560 | 1 | 4 | 202236.52 | 669311.75 | 676408.44 | 395640.09 | 662095 | 759506.31 | 803732.62 | 736717.69 | 854433.12 | 2423596 | 524748.25 | 182676.22 | 179451.16 | 757093.31 | 733826.5 |
| B2RXS4 | Plxnb2 | Plexin-B2 Mus musculus | 1544.57 | 23.45 | 206230 | 27 | 28 | 38733833 | 38123869 | 35281401 | 38462425 | 39450841 | 36691541 | 42548861 | 34656321 | 37957737 | 36720509 | 41330781 | 38536137 | 41337077 | 38475149 | 42693209 |
| Q9QZC2 | Plxnc1 | Plexin-C1 Mus musculus | 80.51 | 1.59 | 176474 | 1 | 2 | 2130857.5 | 1942698.88 | 1995050.25 | 1645297.38 | 1675762.75 | 1954429.12 | 1529058.12 | 2174283.25 | 2249409.5 | 1154507.5 | 1663315.38 | 1847124.88 | 1538220.62 | 1818346.62 | 1529587.5 |
| Q3UH93 | Plxnd1 | Plexin-D1 Mus musculus | 82.31 | 0.73 | 211608 | 1 | 2 | 2130857.5 | 1942698.88 | 1995050.25 | 1645297.38 | 1675762.75 | 1954429.12 | 1529058.12 | 2174283.25 | 2249409.5 | 1154507.5 | 1663315.38 | 1847124.88 | 1538220.62 | 1818346.62 | 1529587.5 |
| Q60953 | Pml | Protein PML Mus musculus | 405.72 | 12.54 | 98242 | 8 | 8 | 3727383 | 4943247.5 | 3920596 | 5544937 | 3840300.75 | 3578635 | 4202713.5 | 3007930.75 | 4751786 | 3360276 | 5139249.5 | 4635943.5 | 5108696.5 | 4323795 | 4640382 |
| O35621 | Pmm1 | Phosphomannomutase 1 Mus musculus | 37.88 | 2.67 | 29775 | 1 | 1 | 1099842.12 | 1270457.62 | 1381751 | 1062134.88 | 1327062.62 | 1064005.38 | 1138212.5 | 1090881.38 | 1201482.5 | 1147649.38 | 1302059.62 | 1537385.88 | 1349640.5 | 1077071.5 | 1367066.25 |

| | | | | | | | | | | | | | | | | | | | | | | |
|---|---|---|---|---|---|---|---|---|---|---|---|---|---|---|---|---|---|---|---|---|---|---|
| Q9CXT8 | Pmpcb | Mitochondrial-processing peptidase subunit beta Mus musculus | 236.7 | 9.2 | 54614 | 3 | 3 | 1601344.25 | 572129.62 | 954106.81 | 815037.75 | 929149.25 | 657776.31 | 1110399.38 | 979980.62 | 891062.44 | 758345.56 | 761407.5 | 957623 | 944748.12 | 1 | 756437.81 |
| Q9D1G2 | Pmvk | Phosphomevalonate kinase Mus musculus | 364.17 | 40.62 | 21916 | 7 | 7 | 4786072 | 6624294 | 4326060 | 4172114.5 | 3573549.75 | 6383203 | 6053788 | 6576488.5 | 3624684.5 | 6840131.5 | 4334692 | 5889277.5 | 5568996 | 3786223.75 | 5309931 |
| Q9JLV6 | Pnkp | Bifunctional polynucleotide phosphatase/kinase Mus musculus | 286.05 | 21.84 | 57223 | 6 | 6 | 3381179.5 | 3531104.75 | 2664787.75 | 3616525 | 3762678 | 3852496.5 | 3276492.5 | 3587662.5 | 2813416.5 | 3204723.25 | 3831070 | 4151723 | 3650293.75 | 3796615.5 | 2813064.75 |
| O35691 | Pnn | Pinin Mus musculus | 87.17 | 3.03 | 82436 | 2 | 2 | 1 | 1 | 1 | 1 | 1 | 1 | 200145.16 | 1 | 1 | 1 | 1 | 1 | 171576.14 | 1 | 1 |
| Q9CPS7 | Pno1 | RNA-binding protein PNO1 Mus musculus | 360.37 | 35.08 | 27454 | 6 | 6 | 10794129 | 9796901 | 10321737 | 12206626 | 9797798 | 11691622 | 13629114 | 9557030 | 8180444.5 | 13363577 | 11588766 | 9381720 | 9075680 | 8555680 | 10659260 |
| P23492 | Pnp | Purine nucleoside phosphorylase Mus musculus | 568.02 | 53.63 | 32277 | 9 | 9 | 49712993 | 48014693 | 53075109 | 46328725 | 46769449 | 40124613 | 43401909 | 44552413 | 42753189 | 38955417 | 34036297 | 42497253 | 48795501 | 45479489 | 38081717 |
| O88794 | Pnpo | Pyridoxine-5'-phosphate oxidase Rattus norvegicus | 78.54 | 7.28 | 30184 | 1 | 1 | 211857.48 | 289780 | 1 | 1 | 1 | 172118.52 | 211852.53 | 1 | 1 | 1 | 187327.12 | 166892.05 | 1 | 164162.2 | 1 |
| Q8K1R3 | Pnpt1 | Polyribonucleotide nucleotidyltransferase 1, mitochondrial Mus musculus | 362.7 | 10.47 | 85683 | 6 | 6 | 2268576.25 | 3983668 | 4151390.5 | 3165388 | 2837620.5 | 3872550.5 | 4669460.5 | 3778381.25 | 5569446.5 | 4313570.5 | 4826252 | 3832493 | 4130779 | 3738899.25 | 4371092 |
| Q91ZW2 | Pofut1 | GDP-fucose protein O-fucosyltransferase 1 Mus musculus | 161.58 | 6.87 | 44688 | 2 | 2 | 1080820.38 | 1082729.38 | 1334043.38 | 1367417.38 | 994716.12 | 958509.88 | 956801.19 | 1479884.5 | 1230814 | 1182593.5 | 1152870.25 | 1178409.25 | 854106.12 | 1242265.5 | 937270.44 |
| Q8VHI3 | Pofut2 | GDP-fucose protein O-fucosyltransferase 2 Mus musculus | 123.34 | 8.39 | 49429 | 3 | 3 | 2027957.62 | 2636437 | 2116389.75 | 2418792.5 | 1831347.38 | 2009690 | 2210819.5 | 2282802.5 | 2166920.5 | 2231635.75 | 1332081 | 776206.69 | 2291060 | 2369112.5 | 1663111 |
| Q8BYB9 | Poglut1 | Protein O-glucosyltransferase 1 Mus musculus | 63.4 | 3.57 | 46379 | 1 | 1 | 1357154.38 | 1038875.62 | 1063874.25 | 1 | 1 | 1150196.62 | 868994.94 | 757421.88 | 1459919.38 | 1 | 1127027.12 | 754216.38 | 941880.25 | 968884 | 1136035 |
| P33609 | Pola1 | DNA polymerase alpha catalytic subunit Mus musculus | 298.12 | 7.24 | 167340 | 3 | 7 | 2384784.75 | 1858212.75 | 2085828 | 1297634.75 | 2496580.5 | 1095820 | 2477432 | 1287277.25 | 1443388.38 | 1441670.62 | 2754721 | 2541151 | 2528388.25 | 1152414.25 | 2661813.75 |
| P33611 | Pola2 | DNA polymerase alpha subunit B Mus musculus | 373.46 | 19.17 | 66214 | 8 | 8 | 1202134.5 | 3810813.75 | 2276942.5 | 2836926.5 | 4987065.5 | 1306454.88 | 1556825.75 | 1314285.5 | 3515866 | 1533746.38 | 1229496.88 | 1578022.12 | 1400075.25 | 1990131.12 | 1746928.62 |
| P52431 | Pold1 | DNA polymerase delta catalytic subunit Mus musculus | 851.72 | 17.1 | 123790 | 15 | 15 | 5683090 | 6021531.5 | 7041082.5 | 6819180.5 | 6743368.5 | 5405067 | 8472224 | 5986467.5 | 5149754 | 7915330 | 6366652 | 7262877.5 | 6756669 | 5661533.5 | 6906650 |
| O35654 | Pold2 | DNA polymerase delta subunit 2 Mus musculus | 164.16 | 5.97 | 51355 | 2 | 2 | 2079043.75 | 1584234 | 1449829.75 | 2223484 | 2042590 | 1809852.25 | 1567421.75 | 2060777.88 | 1888824.38 | 1665696.5 | 1525260 | 2007112.88 | 1996999.62 | 1692187.62 | 1659710 |
| Q91VA6 | Poldip2 | Polymerase delta-interacting protein 2 Mus musculus | 97.95 | 6.52 | 41870 | 2 | 2 | 2267955.75 | 2765633.5 | 2291774 | 2530238.5 | 2195065 | 2072105.5 | 1879094.88 | 2798380.5 | 2955383 | 2185600 | 2052285.75 | 2575010.5 | 1857353.5 | 2398288.75 | 2278924.5 |
| Q8BG81 | Poldip3 | Polymerase delta-interacting protein 3 Mus musculus | 135.29 | 6.43 | 46132 | 2 | 2 | 2322907.75 | 2237060 | 4153206.25 | 2973884.75 | 3496466.25 | 2944985.5 | 6080661.5 | 3013433.75 | 2103621.25 | 4533623.5 | 4257183 | 5858234.5 | 4463919 | 2155423.75 | 5649421.5 |
| O35134 | Polr1a | DNA-directed RNA polymerase I subunit RPA1 Mus musculus | 294.49 | 6.52 | 194110 | 7 | 7 | 1208313.25 | 1723323 | 1655271 | 1210774.88 | 1437643.12 | 1011945.62 | 768896.75 | 2046092.88 | 2723908.75 | 1047883.5 | 1903639.88 | 1915562 | 1134429.25 | 2503339.5 | 2232029 |
| O54888 | Polr1b | DNA-directed RNA polymerase I subunit RPA2 Rattus norvegicus | 35.31 | 1.06 | 127719 | 1 | 1 | 301151.81 | 472892.16 | 363816.28 | 373456.12 | 304530.62 | 244496.84 | 242155.75 | 378090.06 | 415501 | 353803.25 | 346943.16 | 337212.56 | 295013.03 | 400451.09 | 284846.34 |
| P52432 | Polr1c | DNA-directed RNA polymerases I and III subunit RPAC1 Mus musculus | 80.67 | 4.62 | 39107 | 1 | 1 | 649597.62 | 1 | 563999.06 | 1 | 1 | 620335.62 | 502883.19 | 548365.94 | 1 | 1 | 617286.94 | 592371.88 | 535361.76 | 421041.53 | 510746.69 |
| P08775 | Polr2a | DNA-directed RNA polymerase II subunit RPB1 Mus musculus | 859.94 | 10.81 | 217176 | 15 | 15 | 6130495 | 7017554.5 | 4406641 | 8638785 | 8580928 | 6533809.5 | 7471256 | 8019568.5 | 3536499.75 | 10527657 | 8033026.5 | 5859756 | 7050041 | 6317866 | 8217921.5 |
| Q8CFI7 | Polr2b | DNA-directed RNA polymerase II subunit RPB2 Mus musculus | 503.31 | 9.54 | 133911 | 10 | 10 | 3887331.5 | 5512474 | 3498497.25 | 5503975.5 | 6842985 | 4133664.5 | 4821762 | 5866831 | 3904039.5 | 5273989 | 7268254 | 5781915.5 | 5413182 | 5674370.5 | 6720699 |
| P97760 | Polr2c | DNA-directed RNA polymerase II subunit RPB3 Mus musculus | 247.91 | 16 | 31443 | 5 | 5 | 3377675 | 3345010.5 | 3336368.5 | 2728157.25 | 3007289.25 | 3336210.25 | 2814097 | 3037456.5 | 4321200 | 2464641.25 | 3553106.75 | 2986552.5 | 2431494.75 | 3771939.5 | 2687842.25 |
| B0BNE2 | Polr2e | DNA-directed RNA polymerases I, II, and III subunit RPABC1 Rattus norvegicus 2 | 83.11 | 11.43 | 24570 | 2 | 2 | 824166.38 | 435735 | 690125.94 | 834303.69 | 666298.69 | 543037.31 | 463638.88 | 540177.1325 | 465547.09 | 688485.25 | 548231.31 | 538262.475 | 571203.5 | 617202.25 | 416412.84 |
| O88828 | Polr2f | DNA-directed RNA polymerases I, II, and III subunit RPABC2 Rattus norvegicus | 52.49 | 5.51 | 14464 | 1 | 1 | 1859274.38 | 1941991.62 | 1347851.62 | 1806857.25 | 1563165.38 | 1641512.25 | 2328712.75 | 1763425.62 | 1876140.62 | 1820919.88 | 1845306.5 | 1819397.12 | 1802090.25 | 1798562 | 1673582.62 |
| P62488 | Polr2g | DNA-directed RNA polymerase II subunit RPB7 Mus musculus | 117.01 | 18.6 | 19294 | 3 | 3 | 1126125.38 | 1104469.75 | 1446462.62 | 1017018.19 | 1332020.12 | 1273120.75 | 2262775.5 | 1338252.88 | 1448461.12 | 1228221.75 | 1230474 | 1501509 | 1764226 | 1312997.25 | 2064090.5 |
| Q923G2 | Polr2h | DNA-directed RNA polymerases I, II, and III subunit RPABC3 Mus musculus | 267.07 | 30 | 17143 | 4 | 4 | 9638787 | 11448442 | 13317813 | 9702410 | 8062018.5 | 12334288 | 9061301 | 13160747 | 13396754 | 6207312.5 | 16457284 | 13565696 | 9190343 | 12049604 | 14564152 |
| P60898 | Polr2i | DNA-directed RNA polymerase II subunit RPB9 Mus musculus | 35.06 | 18.4 | 14523 | 1 | 1 | 1 | 561935.12 | 363359.94 | 1 | 424748.72 | 373344.25 | 475057.53 | 528277.19 | 399007.88 | 302682.22 | 618882.94 | 452240.47 | 1 | 363016.72 | 1 |
| Q62086 | Pon2 | Serum paraoxonase/arylesterase 2 Mus musculus | 270.17 | 19.49 | 39617 | 4 | 4 | 952454.44 | 618924.5 | 1425951.25 | 957423.25 | 898176.5 | 529091.25 | 1699005.12 | 688367.44 | 1333435.62 | 807906.94 | 1195335.75 | 1760084.12 | 656201.25 | 1317961.5 | 1944991.62 |
| Q62087 | Pon3 | Serum paraoxonase/lactonase 3 Mus musculus | 312.35 | 30.79 | 39351 | 6 | 6 | 4878769 | 3702919.25 | 4530359.5 | 4513536.5 | 6223161.5 | 3863432.5 | 1519230.25 | 4437053 | 1230345.5 | 5547435 | 5838655 | 4814623.5 | 4081482.25 | 3949868.25 | 6349205.5 |
| P37040 | Por | NADPH--cytochrome P450 reductase Mus musculus | 626.12 | 21.98 | 77044 | 10 | 10 | 10977568 | 12653331 | 11925998 | 10276973 | 11639826 | 12489812 | 11812693 | 15083091 | 12795495 | 11978495 | 10945005 | 12699791 | 10345222 | 13800639 | 10680938 |
| P25425 | Pou2f1 | POU domain, class 2, transcription factor 1 Mus musculus | 168.61 | 4.68 | 79547 | 3 | 3 | 2128809.5 | 3422702 | 927902 | 2532669.5 | 3173457.5 | 1972434.5 | 3686545 | 3341982.25 | 2325571.5 | 3795193 | 3319169 | 3351820.5 | 2021177.25 | 2366648.25 | 2953800 |
| Q9D819 | Ppa1 | Inorganic pyrophosphatase Mus musculus | 459.65 | 31.83 | 32667 | 8 | 8 | 13805321 | 17046017 | 17311405 | 13757168 | 13618210 | 15374375 | 17351365 | 18794359 | 19440063 | 14713495 | 20427483 | 18493195 | 15220213 | 16956569 | 14634169 |
| Q91VM9 | Ppa2 | Inorganic pyrophosphatase 2, mitochondrial Mus musculus | 328 | 26.36 | 38115 | 7 | 7 | 2332932.75 | 2222790 | 4204944 | 4992291.5 | 5508218 | 6111630.5 | 5159551 | 3097356.5 | 6571249 | 4214813.5 | 5841555 | 4848789 | 5385730.5 | 3325636 | 4035865.75 |
| Q91YU8 | Ppan | Suppressor of SWI4 1 homolog Mus musculus | 194.17 | 10 | 52756 | 4 | 4 | 1447403.38 | 906248.94 | 1049341.12 | 1373492.62 | 1488146.75 | 1493199.12 | 839895.06 | 2719543.5 | 1724863.25 | 2123574 | 2719189.75 | 2263799.75 | 1928529.25 | 1750653.25 | 1591125.62 |
| O35711 | Ppfibp2 | Liprin-beta-2 Mus musculus | 103.44 | 1.7 | 98753 | 1 | 1 | 1006869.12 | 749892.81 | 1093282.75 | 949830.06 | 933834.31 | 925331.75 | 782226.12 | 1646475.38 | 1282011.38 | 1105367.12 | 736792.38 | 815194.5 | 1024037.94 | 913935.2675 | 1079716.25 |
| P17742 | Ppia | Peptidyl-prolyl cis-trans isomerase A Mus musculus | 1083.22 | 79.27 | 17971 | 14 | 16 | 258784449 | 231428353 | 278078177 | 284652577 | 258668129 | 236810417 | 341449217 | 218678721 | 189301361 | 299569761 | 305611713 | 278770369 | 282352833 | 232542465 | 327449121 |
| P24369 | Ppib | Peptidyl-prolyl cis-trans isomerase B Mus musculus | 527.2 | 44.44 | 23713 | 8 | 8 | 71991793 | 64528973 | 64954889 | 56645589 | 54968821 | 57889409 | 64909681 | 71535641 | 65447317 | 55802705 | 51641881 | 58293209 | 58536237 | 62307401 | 53517733 |
| Q9CR16 | Ppid | Peptidyl-prolyl cis-trans isomerase D Mus musculus | 600.07 | 36.76 | 40743 | 12 | 12 | 9842316 | 10942507 | 10216312 | 8024192.5 | 8889505 | 9258254 | 14609218 | 11570972 | 15736516 | 11319548 | 9939344 | 14422290 | 11010841 | 12876103 | 10794071 |
| A2AR02 | Ppig | Peptidyl-prolyl cis-trans isomerase G Mus musculus | 102.79 | 3.19 | 88325 | 2 | 2 | 3784205 | 657225 | 569312 | 728397.94 | 826942.56 | 3348497.75 | 1307730.75 | 3000965.75 | 2790466 | 469128.78 | 474611 | 1975152.11 | 3587722.5 | 3245617.5 | 592657.44 |
| Q9D868 | Ppih | Peptidyl-prolyl cis-trans isomerase H Mus musculus | 248.13 | 29.26 | 20464 | 5 | 5 | 5652242 | 4064936 | 3795146.5 | 6647986.5 | 6445592 | 5567255.5 | 4984241 | 4934285.5 | 5213492.5 | 5678158 | 5431382 | 5095928 | 5673935 | 5364411 | 3985583.75 |
| Q9D0W5 | Ppil1 | Peptidyl-prolyl cis-trans isomerase-like 1 Mus musculus | 52.84 | 20.48 | 18237 | 1 | 1 | 288334.03 | 1 | 1 | 1 | 1 | 299879.09 | 1 | 406092.41 | 277877.41 | 1 | 1 | 1 | 1 | 1 | 1 |
| Q9CXG3 | Ppil4 | Peptidyl-prolyl cis-trans isomerase-like 4 Mus musculus | 117.62 | 7.72 | 57230 | 2 | 2 | 1875297 | 1 | 1724944 | 1 | 1 | 1688223.5 | 688413.19 | 2116294.5 | 1479205.143 | 1423889.38 | 1376245.25 | 1 | 1841550.12 | 1 | 1726755.75 |
| Q6ZQB6 | Ppip5k2 | Inositol hexakisphosphate and diphosphoinositol-pentakisphosphate kinase 2 Mus musc | 141.43 | 4.07 | 128427 | 3 | 3 | 1313955.12 | 1195377.88 | 756056.56 | 1340971.5 | 1055065.38 | 558345.19 | 984502.94 | 717093.25 | 233355.45 | 968674.94 | 895340.44 | 684509.46 | 769825.56 | 609484.75 | 463387.09 |
| P20650 | Ppm1a | Protein phosphatase 1A Rattus norvegicus | 46.33 | 2.88 | 42417 | 1 | 1 | 833024.5 | 879418.62 | 771367.88 | 770505.12 | 910806.38 | 984325.56 | 688370.31 | 693092.94 | 1030403.69 | 706622.62 | 862421.75 | 757548.81 | 875772.69 | 897596.75 | 806253.38 |
| P36993 | Ppm1b | Protein phosphatase 1B Mus musculus | 61.6 | 4.62 | 42795 | 1 | 1 | 1816424 | 2651372.5 | 2343631.75 | 1847520.25 | 1682665.88 | 2060318.38 | 2258622.25 | 2572896 | 2665262.25 | 2023392.62 | 2342689.5 | 2116949.5 | 2278418.5 | 2178062.75 | 1690667 |
| Q61074 | Ppm1g | Protein phosphatase 1G Mus musculus | 525.34 | 27.12 | 58728 | 10 | 10 | 11635980 | 11763391 | 11682340 | 6516474 | 12106905 | 13112960 | 9422892 | 14270834 | 19195739 | 10274364 | 15948641 | 14860905 | 10249938 | 14076596 | 13110192 |
| Q4FZT2 | Ppme1 | Protein phosphatase methylesterase 1 Rattus norvegicus | 146.88 | 6.99 | 42316 | 3 | 3 | 6184713.5 | 4729126 | 4370266 | 5407718 | 6564517 | 5121067 | 5132259 | 4989349 | 3349357 | 5544782 | 3402395.5 | 2781138.25 | 4867165 | 4711431.5 | 3815242 |
| P51175 | Ppox | Protoporphyrinogen oxidase Mus musculus | 93.2 | 6.92 | 50871 | 2 | 2 | 2312565.25 | 1977656.25 | 2023392.38 | 2096344 | 2418702 | 1777451 | 2660926 | 1685796.62 | 1592522.88 | 2099865.5 | 1930988.12 | 2303973.75 | 1683330.88 | 1597731 | 2095537.5 |
| P62137 | Ppp1ca | Serine/threonine-protein phosphatase PP1-alpha catalytic subunit Mus musculus | 955.57 | 61.21 | 37540 | 1 | 16 | 63125605 | 63142109 | 58299197 | 55401437 | 64901245 | 68946385 | 55425297 | 62027181 | 65971677 | 51589969 | 67399193 | 58162829 | 57972549 | 65687545 | 66829873 |
| P62141 | Ppp1cb | Serine/threonine-protein phosphatase PP1-beta catalytic subunit Mus musculus | 849.14 | 51.68 | 37187 | 4 | 14 | 56258513 | 56350577 | 49453221 | 48099637 | 56901533 | 61581837 | 48083065 | 53705617 | 58286869 | 45376337 | 56752857 | 51396033 | 51865017 | 57301149 | 58830221 |
| P63087 | Ppp1cc | Serine/threonine-protein phosphatase PP1-gamma catalytic subunit Mus musculus | 961.19 | 60.06 | 36984 | 2 | 15 | 57093889 | 55505417 | 52111693 | 50224805 | 59528773 | 63690129 | 52116997 | 55848905 | 59293277 | 47377985 | 60164209 | 55033821 | 52135033 | 59790361 | 63190345 |
| O55000 | Ppp1r10 | Serine/threonine-protein phosphatase 1 regulatory subunit 10 Rattus norvegicus | 39.52 | 2.06 | 92828 | 1 | 1 | 1 | 870048.5 | 553815.44 | 1 | 540611.88 | 1 | 1 | 1 | 795859.12 | 819652.44 | 1 | 418635.66 | 1 | 1 | 1 |
| Q9DBR7 | Ppp1r12a | Protein phosphatase 1 regulatory subunit 12A Mus musculus | 147.14 | 3.98 | 114996 | 3 | 3 | 276169.25 | 1491922 | 1230581.12 | 2161822.5 | 1647309.25 | 1880612.25 | 3149995.5 | 1488532.88 | 1669691.5 | 2647385 | 2565769.75 | 2031912.12 | 2079481 | 1167766.38 | 2287275.25 |
| Q62084 | Ppp1r14b | Protein phosphatase 1 regulatory subunit 14B Mus musculus | 138.95 | 9.52 | 15957 | 2 | 2 | 378832.88 | 1017930.19 | 1128509.12 | 1069769.12 | 832537.12 | 1052005.5 | 496462.91 | 1139953.25 | 1451926.5 | 923257.19 | 1171968.25 | 1057029.62 | 710671.12 | 1029127.5 | 800321.94 |
| Q8BQ30 | Ppp1r18 | Phostensin Mus musculus | 211.23 | 8.59 | 65630 | 5 | 5 | 3270208.75 | 1535491.12 | 1069069.25 | 1639436.5 | 877134.12 | 1345487 | 1797612.5 | 4880640.5 | 1701602 | 1677963.75 | 2236930.5 | 1028403 | 322190.22 | 1658876.5 | 1730036 |
| Q3TDD9 | Ppp1r21 | Protein phosphatase 1 regulatory subunit 21 Mus musculus | 137.28 | 4.1 | 88338 | 2 | 2 | 530425.69 | 357747.75 | 1 | 779400.19 | 1 | 329235.72 | 540345.19 | 1 | 189917.33 | 1 | 1 | 686716.25 | 1 | 545839.25 | 549226.44 |
| Q3UM45 | Ppp1r7 | Protein phosphatase 1 regulatory subunit 7 Mus musculus | 1010.14 | 52.35 | 41292 | 14 | 14 | 20763125 | 19350099 | 17294293 | 19419697 | 24535133 | 22337813 | 21183739 | 26121829 | 28048631 | 23384607 | 23332757 | 20504381 | 24725103 | 23053595 | 23564041 |
| O35274 | Ppp1r9b | Neurabin-2 Rattus norvegicus | 171.3 | 7.22 | 89646 | 3 | 3 | 1191231 | 1580052.38 | 1734393 | 1338055.75 | 1314881.75 | 1264759.12 | 986733.06 | 1326602.62 | 1497092.38 | 1172949.75 | 1174366.62 | 1707832.75 | 1291252.12 | 1132314.12 | 1266323.75 |
| P63330 | Ppp2ca | Serine/threonine-protein phosphatase 2A catalytic subunit alpha isoform Mus musculus | 455.31 | 41.75 | 35608 | 1 | 9 | 32261029 | 34238525 | 30054383 | 29710217 | 36659589 | 39146817 | 31896409 | 31368545 | 35761877 | 26893875 | 30669379 | 28676943 | 33948805 | 31658273 | 32695577 |
| P62715 | Ppp2cb | Serine/threonine-protein phosphatase 2A catalytic subunit beta isoform Mus musculus | 444.64 | 40.13 | 35575 | 1 | 9 | 31010917 | 33454095 | 29690579 | 33054857 | 36666785 | 37964353 | 31743009 | 29933285 | 34595661 | 30575491 | 30589815 | 27777251 | 33756873 | 35401893 | 32403585 |
| Q76MZ3 | Ppp2r1a | Serine/threonine-protein phosphatase 2A 65 kDa regulatory subunit A alpha isoform Mu | 2107.18 | 64.35 | 65323 | 23 | 33 | 114746873 | 128048809 | 126684097 | 118607377 | 116083617 | 136471233 | 125694593 | 135473073 | 141894609 | 124516929 | 141427409 | 131924945 | 127061265 | 137160801 | 137223073 |
| Q4QQT4 | Ppp2r1b | Serine/threonine-protein phosphatase 2A 65 kDa regulatory subunit A beta isoform Ratt | 933.86 | 29.78 | 66006 | 4 | 13 | 39195481 | 41542317 | 44276041 | 40827941 | 41004665 | 48668225 | 39258677 | 46801641 | 48819097 | 43445749 | 49151237 | 44037349 | 44210489 | 45610553 | 48188337 |
| P36876 | Ppp2r2a | Serine/threonine-protein phosphatase 2A 55 kDa regulatory subunit B alpha isoform Rat | 271.45 | 13.2 | 51678 | 5 | 5 | 8231431 | 8330970 | 11728833 | 8788771 | 6324596 | 8178441 | 10886505 | 11011589 | 10072828 | 11722037 | 11258904 | 12492355 | 10089824 | 11274214 | 12144532 |
| Q6PD03 | Ppp2r5a | Serine/threonine-protein phosphatase 2A 56 kDa regulatory subunit alpha isoform Mus | 286.95 | 14.81 | 56347 | 5 | 6 | 5057821.5 | 4250047 | 5595199 | 5685191 | 7293131.5 | 5643824 | 5178254.5 | 6266589 | 6111413 | 6156669.5 | 7152029.5 | 5757738.5 | 4035691.5 | 4761174.5 | 3885083.5 |
| Q60996 | Ppp2r5c | Serine/threonine-protein phosphatase 2A 56 kDa regulatory subunit gamma isoform Mu | 205.03 | 9.73 | 60824 | 2 | 3 | 824246.5 | 524290.88 | 517954.97 | 545492.62 | 430880.75 | 574337.56 | 509591.66 | 477658.16 | 667976.31 | 566702.38 | 623053.5 | 589412.06 | 453726.41 | 751720.5 | 678368.94 |
| Q61151 | Ppp2r5e | Serine/threonine-protein phosphatase 2A 56 kDa regulatory subunit epsilon isoform Mus | 110.81 | 5.14 | 54713 | 0 | 2 | 1416328.25 | 1445775.12 | 1328024.88 | 1152912.12 | 1212347.38 | 1357267.88 | 1330850.12 | 1406270.62 | 1427182.5 | 1258735.88 | 1299022.75 | 1134677.5 | 1140540.25 | 1522133.12 | 1251530.5 |
| P63328 | Ppp3ca | Serine/threonine-protein phosphatase 2B catalytic subunit alpha isoform Mus musculus | 428.71 | 18.43 | 58644 | 3 | 8 | 7832728.5 | 7283644 | 10544078 | 8440826 | 9449262 | 9008021 | 9309907 | 7368214.5 | 8222637.5 | 10425691 | 10026029 | 9414173 | 9462170 | 7472819.5 | 10168744 |
| P20651 | Ppp3cb | Serine/threonine-protein phosphatase 2B catalytic subunit beta isoform Rattus norvegicu | 280.43 | 12.95 | 59113 | 1 | 6 | 7411415.5 | 7283644 | 10116448 | 8440826 | 9009060 | 8698148 | 8311306.5 | 7368214.5 | 8222637.5 | 9318263 | 9636245 | 9414173 | 8951857 | 7472819.5 | 9011505 |
| P63100 | Ppp3r1 | Calcineurin subunit B type 1 Rattus norvegicus | 207.88 | 52.35 | 19300 | 4 | 4 | 2545973.25 | 4231836 | 4503000 | 3813856.5 | 2342452.5 | 3845834 | 4138260.5 | 5017452 | 3134483.25 | 3497762 | 4216934 | 3853289 | 4457455 | 3896966.75 | 2913643.5 |
| P97470 | Ppp4c | Serine/threonine-protein phosphatase 4 catalytic subunit Mus musculus | 253.62 | 18.89 | 35080 | 3 | 5 | 8024741 | 9617601 | 7312884 | 9913145 | 8796322 | 8430583 | 7384979.5 | 8099691 | 8839269 | 7002512 | 8365353.5 | 7004892 | 8502909 | 9713556 | 6512654.5 |
| Q8K2V1 | Ppp4r1 | Serine/threonine-protein phosphatase 4 regulatory subunit 1 Mus musculus | 405.36 | 11.25 | 106298 | 7 | 7 | 4969056 | 3926590 | 3827186 | 4019834.5 | 3875424.5 | 4199255 | 2724534 | 3786100.75 | 3857741.5 | 3793389.25 | 3788582.25 | 3708579 | 3609772.25 | 4375936 | 3473000.5 |
| Q0VGB7 | Ppp4r2 | Serine/threonine-protein phosphatase 4 regulatory subunit 2 Mus musculus | 177.66 | 10.55 | 46479 | 3 | 3 | 1850140 | 2034461.25 | 2903052.75 | 1955833 | 2421902 | 797446.75 | 3237861.5 | 2491572.5 | 3306604 | 3330071.25 | 3062334 | 2896851.5 | 3428952 | 3049706.25 | 1773749 |
| Q6P2K6 | Ppp4r3a | Serine/threonine-protein phosphatase 4 regulatory subunit 3A Mus musculus | 170.73 | 7.2 | 93842 | 3 | 3 | 727461.94 | 249066.88 | 549090.38 | 178199.16 | 939538.94 | 245749.95 | 127423.92 | 445817.72 | 890609.19 | 427400.195 | 347935.59 | 838248.88 | 645792.75 | 374856.03 | 423733.94 |
| Q922R5 | Ppp4r3b | Serine/threonine-protein phosphatase 4 regulatory subunit 3B Mus musculus | 152.42 | 4.15 | 93934 | 2 | 2 | 282221.12 | 310789.46 | 234770.5 | 304347.28 | 421818.94 | 246671.14 | 331703.59 | 328561.34 | 446356.94 | 289513.69 | 255934.25 | 329967.94 | 296528.205 | 303993.41 | 296217.22 |
| Q60676 | Ppp5c | Serine/threonine-protein phosphatase 5 Mus musculus | 317.02 | 13.63 | 56877 | 5 | 5 | 7287939.5 | 7789658 | 7772254 | 6768153.5 | 7353698 | 6919388.5 | 5880583 | 7301741 | 7956025 | 5868248.5 | 6344223 | 6607450.5 | 7216182.5 | 7829835 | 6409882 |
| Q64620 | Ppp6c | Serine/threonine-protein phosphatase 6 catalytic subunit Rattus norvegicus | 410.62 | 31.48 | 35159 | 7 | 7 | 7714178.5 | 8356844 | 6788874 | 8338708 | 7995899.5 | 8915446 | 8472667 | 7886204.5 | 9033417 | 7314402.5 | 6281929 | 5670804 | 8161770 | 7967846.5 | 7411298 |
| Q7TSI3 | Ppp6r1 | Serine/threonine-protein phosphatase 6 regulatory subunit 1 Mus musculus | 258.19 | 7.71 | 94527 | 4 | 4 | 3986228.5 | 3452987.5 | 3294930.5 | 3856874.5 | 3947366 | 3690305.5 | 3830286.25 | 3611517.5 | 3888571 | 3874904.5 | 4276198.5 | 3651668 | 4029664 | 3483268.5 | 3537296.5 |
| Q922D4 | Ppp6r3 | Serine/threonine-protein phosphatase 6 regulatory subunit 3 Mus musculus | 372.73 | 10.31 | 94653 | 7 | 7 | 7959086 | 7394944 | 7224796 | 6713954.5 | 7296041 | 7281385 | 5699610 | 7336018 | 8738812 | 6198966.5 | 6778997 | 6601205.5 | 5360688 | 7317629 | 5969491.5 |
| O88531 | Ppt1 | Palmitoyl-protein thioesterase 1 Mus musculus | 361.35 | 27.45 | 34490 | 6 | 6 | 23402251 | 21343829 | 19147169 | 20901289 | 22389045 | 15715685 | 7344088 | 27392117 | 23994811 | 12082441 | 19651949 | 12995618 | 21592907 | 14847686 | 10042230 |
| Q8CEC6 | Ppwd1 | Peptidylprolyl isomerase domain and WD repeat-containing protein 1 Mus musculus | 52.69 | 2.01 | 73385 | 1 | 1 | 6784737.5 | 14874537 | 46376205 | 30773133 | 69166489 | 46328325 | 83625465 | 9730022 | 10543374 | 89583481 | 80103761 | 49236205 | 54468749 | 7522870 | 90981481 |
| P35700 | Prdx1 | Peroxiredoxin-1 Mus musculus | 1304.28 | 89.45 | 22176 | 8 | 23 | 943166273 | 1082265217 | 1051464513 | 721838849 | 938903553 | 958195841 | 875788353 | 1052827329 | 1100930689 | 838971969 | 944323713 | 878618049 | 999233793 | 1050074561 | 745564289 |
| Q61171 | Prdx2 | Peroxiredoxin-2 Mus musculus | 761.15 | 60.1 | 21779 | 13 | 13 | 193823601 | 219943553 | 203400849 | 203886273 | 214063009 | 200751393 | 177464673 | 203374625 | 234636385 | 197124465 | 206974625 | 189187441 | 207282145 | 212356817 | 177755761 |
| P20108 | Prdx3 | Thioredoxin-dependent peroxide reductase, mitochondrial Mus musculus | 403.04 | 42.8 | 28127 | 3 | 8 | 35121993 | 37198285 | 39623681 | 31474173 | 32779713 | 34929449 | 30423497 | 44716305 | 46675673 | 31070857 | 31228437 | 37341493 | 33389731 | 37633449 | 27593643 |
| O08807 | Prdx4 | Peroxiredoxin-4 Mus musculus | 864.57 | 63.87 | 31053 | 14 | 16 | 217877649 | 262339073 | 235947489 | 166877601 | 216095537 | 219068817 | 187843281 | 253784273 | 289052481 | 182908561 | 200529169 | 191766497 | 232361249 | 260797521 | 152626785 |
| P99029 | Prdx5 | Peroxiredoxin-5, mitochondrial Mus musculus | 774.76 | 57.14 | 21897 | 12 | 12 | 56091017 | 48160737 | 47277485 | 38873749 | 49548785 | 61708961 | 68934545 | 64003477 | 47615481 | 59412049 | 49820665 | 47589333 | 59442349 | 53751717 | 45146181 |
| O08709 | Prdx6 | Peroxiredoxin-6 Mus musculus | 1011.65 | 80.36 | 24871 | 17 | 17 | 76961673 | 86185241 | 90438033 | 75179377 | 87438225 | 95907689 | 90753481 | 94788385 | 93994225 | 78945737 | 100493825 | 87899361 | 83954257 | 88914793 | 84407977 |
| Q9WUQ2 | Preb | Prolactin regulatory element-binding protein Mus musculus | 186.51 | 11.75 | 45437 | 3 | 3 | 1280340.5 | 1837462.88 | 1958280.5 | 1251712.12 | 460748.56 | 1559344.5 | 1235186.25 | 1625002.62 | 1858771.62 | 2065298.75 | 845142.62 | 1384834.12 | 1497056.62 | 1257312.88 | 1153972.88 |
| Q9QUR6 | Prep | Prolyl endopeptidase Mus musculus | 1197.57 | 38.59 | 80752 | 19 | 19 | 24552079 | 25021749 | 26155509 | 23544407 | 22921967 | 21927601 | 26460453 | 24265533 | 31616683 | 24314531 | 27246803 | 27897549 | 22311927 | 27608513 | 26582129 |
| Q69ZK0 | Prex1 | Phosphatidylinositol 3,4,5-trisphosphate-dependent Rac exchanger 1 protein Mus musc | 1388.41 | 22.42 | 184935 | 27 | 27 | 26847303 | 27958801 | 27497355 | 24238315 | 29390133 | 24354819 | 27414825 | 30366721 | 29688279 | 26298881 | 28874389 | 27887683 | 27619745 | 30202313 | 26521631 |
| P20664 | Prim1 | DNA primase small subunit Mus musculus | 79.63 | 6 | 49295 | 2 | 2 | 1491189.25 | 1334787.75 | 1393445.62 | 1285053.62 | 1336268.62 | 693750.69 | 1397605.75 | 970380.69 | 1479323 | 1273168.38 | 1446610.38 | 1347115 | 562054.19 | 1617413.75 | 1519465.25 |
| P33610 | Prim2 | DNA primase large subunit Mus musculus | 281.84 | 12.48 | 58408 | 5 | 5 | 2131409.25 | 2714766.25 | 3903885 | 2777066.25 | 1657883.38 | 3064174.75 | 2638930.5 | 3749950 | 3264352.5 | 2443136.75 | 4738721 | 3269441.5 | 2593775.25 | 2419439.5 | 3458527.25 |
| P54645 | Prkaa1 | 5'-AMP-activated protein kinase catalytic subunit alpha-1 Rattus norvegicus | 131.28 | 6.44 | 63973 | 3 | 3 | 1389563.25 | 1004741 | 1438128.75 | 1325978 | 1516875 | 1388945.25 | 1844298 | 1059585.88 | 1266091.38 | 1747111.25 | 1872544 | 1482560.25 | 1745537.25 | 1528098.75 | 1764208.5 |
| P05132 | Prkaca | cAMP-dependent protein kinase catalytic subunit alpha Mus musculus | 381.87 | 26.5 | 40571 | 2 | 8 | 24790171 | 23126309 | 21086561 | 19925703 | 21379051 | 25180197 | 24480627 | 25415005 | 24839729 | 22893679 | 23939923 | 23890895 | 22760153 | 24746317 | 22745525 |
| P68181 | Prkacb | cAMP-dependent protein kinase catalytic subunit beta Mus musculus | 394.28 | 26.5 | 40708 | 2 | 8 | 23713019 | 22095197 | 19357943 | 18849219 | 21044219 | 24429861 | 22865175 | 23631851 | 23870901 | 20608445 | 22065061 | 23050943 | 21032291 | 24043785 | 21237135 |
| O54950 | Prkag1 | 5'-AMP-activated protein kinase subunit gamma-1 Mus musculus | 591.44 | 43.03 | 37520 | 9 | 9 | 3652924.25 | 3171194.5 | 3296764.75 | 3395168.25 | 1629812 | 4296376.5 | 3277884 | 4921022.5 | 5332472.5 | 4491335.5 | 4700770.5 | 2627611.75 | 3793732 | 4659817 | 5149325.5 |
| Q9DBC7 | Prkar1a | cAMP-dependent protein kinase type I-alpha regulatory subunit Mus musculus | 631.42 | 34.91 | 43185 | 11 | 11 | 20935051 | 23816355 | 21509539 | 23857553 | 24367271 | 19492985 | 20858967 | 21995307 | 20642907 | 21458645 | 25810685 | 19861993 | 23097477 | 22228725 | 20516323 |
| P12368 | Prkar2a | cAMP-dependent protein kinase type II-alpha regulatory subunit Rattus norvegicus | 337.12 | 20.95 | 45480 | 6 | 6 | 5819845.5 | 6378244.5 | 3912883.5 | 5561034 | 8573565 | 8615870 | 8338015.5 | 5684000 | 6032389.5 | 7653203 | 9010062 | 8371977 | 8368564.5 | 8181497 | 8117033.5 |
| P68403 | Prkcb | Protein kinase C beta type Rattus norvegicus | 192.2 | 6.11 | 76751 | 3 | 4 | 629502.56 | 477210.38 | 1 | 1 | 284359.34 | 1 | 305410.91 | 1 | 1 | 1 | 1 | 1 | 1 | 432548.56 | 1 |
| P28867 | Prkcd | Protein kinase C delta type Mus musculus | 396.02 | 13.2 | 77547 | 7 | 8 | 9568542 | 12952222 | 10283728 | 12139951 | 14702367 | 10037143 | 12280448 | 9296543 | 9719068 | 13183679 | 12097124 | 10897304 | 12222587 | 10423347 | 13862661 |
| O08795 | Prkcsh | Glucosidase 2 subunit beta Mus musculus | 423.96 | 19.39 | 58793 | 8 | 8 | 14769977 | 14824747 | 16255006 | 15607839 | 14695123 | 15093752 | 16962517 | 15263621 | 15543984 | 16714175 | 15302437 | 13976107 | 14199965 | 14540262 | 12272433 |
| Q5XIS9 | Prkd2 | Serine/threonine-protein kinase D2 Rattus norvegicus | 73.48 | 1.26 | 96497 | 1 | 1 | 1 | 143415.03 | 1 | 1 | 144991.11 | 161520.64 | 153071.89 | 237913.2 | 235054.06 | 145472.48 | 220041.52 | 193733.435 | 142557.83 | 218713.11 | 193621.28 |
| Q63009 | Prmt1 | Protein arginine N-methyltransferase 1 Rattus norvegicus | 1101.34 | 53.54 | 40522 | 20 | 20 | 48532225 | 47985949 | 46430077 | 42264709 | 45721829 | 40695521 | 28322159 | 41056333 | 45250313 | 31501509 | 35086825 | 35684169 | 35647897 | 44480921 | 35413465 |
| Q8CIG8 | Prmt5 | Protein arginine N-methyltransferase 5 Mus musculus | 264.02 | 12.24 | 72680 | 6 | 6 | 5985871 | 4608271 | 5556597 | 2380798 | 3578270 | 4291364.5 | 4521112.5 | 2713782.25 | 3214155.5 | 2192918.75 | 4089212 | 5308928 | 2478735.25 | 3100281 | 3960148.5 |
| Q64695 | Procr | Endothelial protein C receptor Mus musculus | 40.87 | 7.02 | 27189 | 1 | 1 | 793269.44 | 716483.94 | 656806.94 | 634748.1975 | 372432.47 | 919169.81 | 584449.44 | 1005127.44 | 852544.12 | 739452.25 | 708580.06 | 850846.81 | 762320.62 | 876379 | 429103.12 |

| | | | | | | | | | | | | | | | | | | | | | | |
|---|---|---|---|---|---|---|---|---|---|---|---|---|---|---|---|---|---|---|---|---|---|---|
| Q9D820 | Prorsd1 | Prolyl-tRNA synthetase associated domain-containing protein 1 Mus musculus | 74.67 | 10.06 | 18862 | 2 | 2 | 615482.5 | 1214442.62 | 849631.19 | 1042563.75 | 827395.31 | 846980.06 | 1 | 785814.44 | 1009813.12 | 660652.155 | 1080762 | 716558.06 | 1 | 864257.5 | 1 |
| Q99KP6 | Prpf19 | Pre-mRNA-processing factor 19 Mus musculus | 682.12 | 52.18 | 55239 | 13 | 13 | 22918815 | 21627957 | 27461579 | 24921621 | 24320841 | 26642835 | 29978067 | 24468527 | 25520125 | 24892495 | 28712487 | 29329833 | 26174671 | 23695363 | 31568425 |
| Q922U1 | Prpf3 | U4/U6 small nuclear ribonucleoprotein Prp3 Mus musculus | 117.93 | 4.83 | 77455 | 3 | 3 | 761582 | 645401.25 | 668652.5 | 694886.12 | 692630.4675 | 878189.62 | 944834.75 | 593266.75 | 532077.19 | 737092.0775 | 537024.69 | 771841.31 | 788096.12 | 654809.25 | 687942.8425 |
| Q8CCF0 | Prpf31 | U4/U6 small nuclear ribonucleoprotein Prp31 Mus musculus | 664.09 | 31.66 | 55430 | 11 | 11 | 10524550 | 9749807 | 9692875 | 8772612 | 8067656.5 | 9848536 | 8292537 | 9831371 | 11840226 | 9509913 | 12552029 | 8437800 | 9397418 | 12630679 | 11060948 |
| Q4FK66 | Prpf38a | Pre-mRNA-splicing factor 38A Mus musculus | 220.03 | 12.82 | 37437 | 4 | 4 | 3209023.5 | 3982061 | 3610993.25 | 3744178.25 | 4512680 | 4135696 | 4012286 | 4006234.25 | 4363598 | 3766379.5 | 3762819.5 | 3547055.25 | 4133751.75 | 3531624 | 3644941.5 |
| Q6AXY7 | Prpf38b | Pre-mRNA-splicing factor 38B Rattus norvegicus | 94.67 | 5.35 | 63932 | 2 | 2 | 1203714.38 | 1538512.62 | 1099557.25 | 1420370 | 1767775.12 | 1457849 | 1392986.12 | 1317267 | 1248132.5 | 1584808.5 | 1568630.62 | 1058446.62 | 1436315.25 | 1661562 | 1267982.25 |
| Q9DAW6 | Prpf4 | U4/U6 small nuclear ribonucleoprotein Prp4 Mus musculus | 103.33 | 4.99 | 58370 | 2 | 2 | 3025687.75 | 505579.22 | 3164263.25 | 3665880.5 | 3719378.5 | 331177.34 | 3092026.75 | 3162237.75 | 1188638.75 | 3180538 | 2502166 | 1256467 | 3464063 | 2773851.5 | 2542603.75 |
| Q9R1C7 | Prpf40a | Pre-mRNA-processing factor 40 homolog A Mus musculus | 288.99 | 10.18 | 108481 | 6 | 6 | 7514857 | 7282676.5 | 4758313 | 5327646.5 | 7648230 | 6668801.5 | 7264723 | 7856353.5 | 9614325 | 5456056.5 | 8189809 | 7708784.5 | 7328389 | 6969762.5 | 6789990 |
| Q5RKH1 | Prpf4b | Serine/threonine-protein kinase PRP4 homolog Rattus norvegicus | 51.07 | 1.69 | 117007 | 1 | 1 | 1424606.88 | 1689704.62 | 1048682.38 | 1515060.25 | 1387228.25 | 1146401 | 830589 | 1064548.62 | 1440622.75 | 1162901.75 | 841158.12 | 543436.88 | 1413520.12 | 973611.88 | 782595.94 |
| A1A5S1 | Prpf6 | Pre-mRNA-processing factor 6 Rattus norvegicus | 217.37 | 4.89 | 106764 | 6 | 6 | 1923015.5 | 4360011.5 | 4236407 | 2191759 | 2376259.5 | 2919618.5 | 3261218.5 | 3972309 | 3542511 | 3757666.5 | 3428335 | 3143214 | 4173383.5 | 4459431 | 2916248 |
| Q99PV0 | Prpf8 | Pre-mRNA-processing-splicing factor 8 Mus musculus | 1434.24 | 18.03 | 273616 | 29 | 29 | 31754929 | 29173377 | 25986165 | 24888961 | 31501925 | 30290443 | 17482775 | 21225451 | 33987981 | 24562371 | 36510345 | 26148149 | 25184441 | 29099357 | 27607725 |
| P60892 | Prps1 | Ribose-phosphate pyrophosphokinase 1 Rattus norvegicus | 753.02 | 41.82 | 34834 | 5 | 12 | 13199419 | 12918571 | 13263252 | 13187058 | 13574412 | 14846805 | 14590116 | 12678001 | 12778328 | 13122595 | 17515087 | 14124436 | 12222706 | 13588345 | 14381525 |
| Q9CS42 | Prps2 | Ribose-phosphate pyrophosphokinase 2 Mus musculus | 653.17 | 39.31 | 34786 | 3 | 10 | 10548243 | 10978101 | 10697736 | 11481259 | 13096030 | 13377699 | 13415596 | 10520777 | 10430280 | 12091653 | 15463045 | 11645007 | 12096508 | 10994697 | 11701136 |
| Q9D0M1 | Prpsap1 | Phosphoribosyl pyrophosphate synthase-associated protein 1 Mus musculus | 446.08 | 27.25 | 39432 | 6 | 8 | 10095758 | 8078865 | 5478888 | 6610907.5 | 7423608 | 8964591 | 9267661 | 8221745 | 6921202.5 | 7080429 | 8021567 | 6849039.5 | 7542565 | 8763459 | 7213427 |
| Q8R574 | Prpsap2 | Phosphoribosyl pyrophosphate synthase-associated protein 2 Mus musculus | 388.39 | 24.39 | 40881 | 5 | 7 | 7194530 | 6435119 | 5972740.5 | 6093794 | 6516345 | 7039867 | 6577612.5 | 7174980 | 7030698 | 5975576.5 | 6218749 | 6067642 | 6605254.5 | 6907699 | 5968958 |
| Q3T1I4 | Prrc1 | Protein PRRC1 Rattus norvegicus | 295.8 | 27.31 | 46295 | 6 | 6 | 2347264 | 5160084 | 4681518 | 4292462 | 3467514.75 | 3175674.5 | 4499048 | 4846456.5 | 5315431.5 | 2996516.5 | 3546854.25 | 4046140.5 | 3915721 | 4400546.5 | 3216468.5 |
| Q3TLH4 | Prrc2c | Protein PRRC2C Mus musculus | 602.04 | 6.01 | 310892 | 11 | 11 | 1514673.25 | 2235882 | 2596029 | 2538311.25 | 1600206.5 | 1019815.31 | 3827010 | 742232.19 | 1786509.62 | 3308577.5 | 4606957 | 3289167 | 2921892.25 | 2193540 | 3430478 |
| Q61207 | Psap | Prosaposin Mus musculus | 500.34 | 16.34 | 61422 | 8 | 9 | 62608617 | 85619153 | 61576009 | 67607481 | 62274225 | 57890369 | 50920657 | 60608353 | 58004545 | 61684089 | 49026825 | 53208489 | 63861249 | 60159913 | 39871305 |
| Q99K85 | Psat1 | Phosphoserine aminotransferase Mus musculus | 1147.95 | 60 | 40473 | 18 | 18 | 65698977 | 68842881 | 75055409 | 73179913 | 70061217 | 74374121 | 82046257 | 75086745 | 77025009 | 78450913 | 70546609 | 76939481 | 65017617 | 69432281 | 84725393 |
| O88777 | Psen2 | Presenilin-2 Rattus norvegicus | 101.35 | 3.35 | 50051 | 1 | 1 | 1112437.88 | 1977275.62 | 1674955.25 | 1653055.25 | 1574831.88 | 1467197.12 | 1218371.88 | 1660993.88 | 1497099.5 | 1730824.38 | 1783517.38 | 1750324.25 | 1528952.12 | 1742912.25 | 1347876.38 |
| Q99JF8 | Psip1 | PC4 and SFRS1-interacting protein Mus musculus | 460.55 | 19.89 | 59697 | 8 | 9 | 12897072 | 12902274 | 8302025.5 | 13204524 | 13383367 | 13210175 | 16973495 | 13176537 | 9890811 | 16218694 | 14189571 | 13559573 | 13771264 | 12785392 | 14988838 |
| Q9R1P4 | Psma1 | Proteasome subunit alpha type-1 Mus musculus | 521.4 | 60.46 | 29547 | 12 | 12 | 31595289 | 35210129 | 31054659 | 36645581 | 35170449 | 33022625 | 30388365 | 33461167 | 30956035 | 32170853 | 31537723 | 30059445 | 33707921 | 32201673 | 30202039 |
| P17220 | Psma2 | Proteasome subunit alpha type-2 Rattus norvegicus | 631.85 | 46.15 | 25927 | 9 | 9 | 5367493 | 3770205.5 | 4416430 | 4643863.5 | 5352430 | 4511849.5 | 8736249 | 3593253.25 | 3821175.75 | 7530334.5 | 4721331 | 6072753.5 | 5195541 | 4120800 | 6378772 |
| O70435 | Psma3 | Proteasome subunit alpha type-3 Mus musculus | 525.17 | 34.51 | 28405 | 8 | 8 | 46703641 | 45811433 | 40112433 | 37888277 | 41449493 | 42375601 | 35972329 | 43279397 | 47110881 | 35909113 | 40462545 | 40920281 | 40363965 | 43540297 | 34579001 |
| P21670 | Psma4 | Proteasome subunit alpha type-4 Rattus norvegicus | 623.23 | 52.87 | 29498 | 9 | 9 | 23731827 | 22714291 | 22937429 | 19437545 | 20412123 | 21530549 | 17594749 | 23660381 | 26163075 | 16283758 | 21584183 | 21095757 | 18845487 | 23855371 | 20469281 |
| Q9Z2U1 | Psma5 | Proteasome subunit alpha type-5 Mus musculus | 628.62 | 65.98 | 26411 | 10 | 10 | 48418773 | 54426885 | 61356933 | 50844089 | 53689413 | 54951185 | 50443937 | 54582761 | 60188785 | 49441741 | 56022701 | 56810769 | 50048133 | 57594309 | 53610281 |
| Q9QUM9 | Psma6 | Proteasome subunit alpha type-6 Mus musculus | 601.56 | 59.35 | 27372 | 10 | 10 | 40426513 | 36778433 | 43377193 | 41162473 | 38554765 | 41697081 | 56388525 | 37738249 | 33595761 | 47771261 | 50397181 | 44536305 | 43779805 | 38813057 | 44323369 |
| Q9Z2U0 | Psma7 | Proteasome subunit alpha type-7 Mus musculus | 662.54 | 51.61 | 27855 | 11 | 11 | 48832961 | 37826333 | 51889441 | 36299857 | 35391637 | 41619001 | 46928113 | 46266865 | 45569249 | 41902809 | 50638649 | 52977277 | 36241369 | 44737745 | 57327933 |
| O09061 | Psmb1 | Proteasome subunit beta type-1 Mus musculus | 889.38 | 65.42 | 26372 | 11 | 13 | 44223393 | 44306617 | 46898557 | 46310325 | 47877153 | 44547173 | 45034729 | 43387329 | 43009189 | 46373913 | 51612761 | 45344029 | 46367753 | 43292409 | 40877133 |
| O35955 | Psmb10 | Proteasome subunit beta type-10 Mus musculus | 263.8 | 21.25 | 29063 | 4 | 4 | 4053446 | 4426572 | 5276907.5 | 5294959 | 3176399 | 4656366 | 5913327 | 4066431.5 | 3414086.5 | 5696757 | 5048151 | 5126815.5 | 5229559.5 | 3605173.75 | 5591885.5 |
| Q9R1P3 | Psmb2 | Proteasome subunit beta type-2 Mus musculus | 564.71 | 64.18 | 22906 | 10 | 10 | 25188259 | 23431355 | 23791241 | 22826801 | 21087685 | 23770749 | 21359665 | 22817479 | 30952149 | 23923517 | 19654127 | 21194407 | 24122603 | 30999695 | 28066205 |
| Q9R1P1 | Psmb3 | Proteasome subunit beta type-3 Mus musculus | 493.69 | 53.17 | 22965 | 8 | 8 | 38088821 | 39792409 | 41585749 | 38853289 | 41475689 | 42663561 | 43969641 | 40239129 | 43768621 | 44703805 | 42588357 | 41988505 | 45713765 | 41178841 | 42573357 |
| P99026 | Psmb4 | Proteasome subunit beta type-4 Mus musculus | 390.14 | 37.12 | 29116 | 6 | 6 | 28099033 | 25629421 | 30363549 | 29061781 | 29095897 | 28179021 | 30972817 | 27523893 | 29235857 | 32255615 | 29321575 | 29704207 | 28865173 | 27351103 | 26683009 |
| Q60692 | Psmb6 | Proteasome subunit beta type-6 Mus musculus | 201.31 | 26.89 | 25379 | 4 | 4 | 10038085 | 11115873 | 7537159 | 8827584 | 10774531 | 8785147 | 7036649 | 7497808 | 10464204 | 6686046 | 9557131 | 7699830.5 | 8418009 | 11424003 | 7038564 |
| P70195 | Psmb7 | Proteasome subunit beta type-7 Mus musculus | 151.7 | 28.16 | 29891 | 4 | 4 | 4540134 | 3425151.5 | 3747979.25 | 3307585.75 | 3136085.75 | 3354645 | 6652334 | 3035939.5 | 3324130.75 | 6219445 | 3948413.25 | 3590511.75 | 3548869.75 | 3184951.75 | 3281334 |
| P28063 | Psmb8 | Proteasome subunit beta type-8 Mus musculus | 763.5 | 51.81 | 30260 | 11 | 11 | 27842901 | 29829893 | 29364637 | 24339501 | 27690625 | 27703267 | 30708893 | 28118943 | 28526273 | 28270823 | 33534149 | 28482445 | 29449441 | 24041749 | 30564633 |
| O35522 | Psmb9 | Proteasome subunit beta type-9 Mus musculus bactrianus | 260.75 | 34.25 | 23427 | 5 | 5 | 23776365 | 27161923 | 22946583 | 26063835 | 24925917 | 20846527 | 22049807 | 21615975 | 21897407 | 22490425 | 22223953 | 19638735 | 25946293 | 24033761 | 19326683 |
| P62192 | Psmc1 | 26S proteasome regulatory subunit 4 Mus musculus | 766.86 | 32.5 | 49185 | 10 | 12 | 14150257 | 12966606 | 15130525 | 12603510 | 12033492 | 12846303 | 15426095 | 14296621 | 12750314 | 14644984 | 14585228 | 15224443 | 14018320 | 12878628 | 15900432 |
| P46471 | Psmc2 | 26S proteasome regulatory subunit 7 Mus musculus | 1108.77 | 47.34 | 48648 | 17 | 17 | 27022051 | 24684971 | 23998349 | 28707055 | 25778543 | 27546665 | 32506967 | 21629127 | 24078985 | 30224319 | 27506867 | 25101701 | 25334869 | 25881917 | 29900575 |
| O88685 | Psmc3 | 26S proteasome regulatory subunit 6A Mus musculus | 1011.96 | 48.87 | 49549 | 16 | 16 | 29903755 | 30740665 | 30914379 | 34134981 | 32561789 | 31713101 | 36560713 | 31284645 | 32597799 | 36314149 | 37198785 | 33246609 | 32556369 | 30235531 | 33638421 |
| P54775 | Psmc4 | 26S proteasome regulatory subunit 6B Mus musculus | 1029.62 | 51.44 | 47408 | 16 | 17 | 19170535 | 17510731 | 16672966 | 14593041 | 15335300 | 18418825 | 19538051 | 20611535 | 18087327 | 12820755 | 23665193 | 19226619 | 17482703 | 17981765 | 22371985 |
| P62196 | Psmc5 | 26S proteasome regulatory subunit 8 Mus musculus | 930.07 | 47.78 | 45626 | 15 | 17 | 20966437 | 18491385 | 16955559 | 21492891 | 19581869 | 19661965 | 20929459 | 19542395 | 14644315 | 20782883 | 19844875 | 20807527 | 19620125 | 18920231 | 19320231 |
| P62334 | Psmc6 | 26S proteasome regulatory subunit 10B Mus musculus | 719.43 | 44.47 | 44173 | 15 | 15 | 22500033 | 21118061 | 23835547 | 24038083 | 20199929 | 23137295 | 23980195 | 20330471 | 21310263 | 18416593 | 20036223 | 22865879 | 21661487 | 22128807 | 21982073 |
| Q3TXS7 | Psmd1 | 26S proteasome non-ATPase regulatory subunit 1 Mus musculus | 1808.57 | 42.29 | 105730 | 26 | 28 | 46912189 | 51100169 | 50941269 | 59514185 | 53150733 | 58020177 | 59441737 | 55986841 | 58868753 | 58399465 | 51532649 | 55687457 | 53551669 | 54945981 | 52742013 |
| Q9Z2X2 | Psmd10 | 26S proteasome non-ATPase regulatory subunit 10 Mus musculus | 42.58 | 3.46 | 25084 | 1 | 1 | 1447422.5 | 1131062.5 | 1309101.75 | 1804776.25 | 1462486.5 | 1399569.12 | 1846210.12 | 1467233.5 | 1229676.62 | 1590755.12 | 1469448.25 | 1325979.5 | 1441440.25 | 1401701 | 887516.88 |
| F1LMZ8 | Psmd11 | 26S proteasome non-ATPase regulatory subunit 11 Rattus norvegicus | 1109.24 | 57.58 | 47464 | 19 | 19 | 45383285 | 36889545 | 33363937 | 32518591 | 34964637 | 33118371 | 42080521 | 35544201 | 34715937 | 34547017 | 31258237 | 35079037 | 33268581 | 36434157 | 39269121 |
| Q9D8W5 | Psmd12 | 26S proteasome non-ATPase regulatory subunit 12 Mus musculus | 1211.41 | 50.88 | 52895 | 19 | 20 | 52846409 | 55731561 | 53422661 | 46323533 | 49097925 | 53630717 | 50394361 | 60490153 | 61712669 | 47976273 | 52744325 | 54581133 | 49205925 | 58145681 | 49325049 |
| Q9WVJ2 | Psmd13 | 26S proteasome non-ATPase regulatory subunit 13 Mus musculus | 918.77 | 47.87 | 42809 | 17 | 17 | 45466909 | 42412457 | 37190153 | 40671525 | 38362953 | 38866533 | 32126231 | 39512293 | 45080521 | 37336421 | 39669213 | 36723645 | 37274429 | 43617293 | 32171065 |
| O35593 | Psmd14 | 26S proteasome non-ATPase regulatory subunit 14 Mus musculus | 392.99 | 38.06 | 34577 | 6 | 6 | 17339677 | 21239563 | 20101507 | 21653543 | 20161839 | 20380545 | 20575307 | 22081643 | 23212209 | 20174833 | 15135055 | 19840455 | 19683951 | 23193875 | 17174365 |
| Q8VDM4 | Psmd2 | 26S proteasome non-ATPase regulatory subunit 2 Mus musculus | 1800.66 | 47.8 | 100203 | 29 | 29 | 66501597 | 65012813 | 71737081 | 67672377 | 68472497 | 69628657 | 75036825 | 74853001 | 77078833 | 76089073 | 74211785 | 67134289 | 69674009 | 69574081 | 69056593 |
| P14685 | Psmd3 | 26S proteasome non-ATPase regulatory subunit 3 Mus musculus | 444.5 | 22.45 | 60718 | 9 | 9 | 16284630 | 7235685 | 4872775 | 6756865 | 9649383 | 7423300 | 7099787.5 | 6783511 | 6904869.5 | 6293767.5 | 8629677 | 4852735 | 7196402 | 8272240 | 9794190 |
| O35226 | Psmd4 | 26S proteasome non-ATPase regulatory subunit 4 Mus musculus | 501.55 | 28.72 | 40704 | 10 | 10 | 14012863 | 12021024 | 13135014 | 11817732 | 13702150 | 12422613 | 12818471 | 12727047 | 12619238 | 11531887 | 15015264 | 12941744 | 15008648 | 13524925 | 11577807 |
| Q8BJY1 | Psmd5 | 26S proteasome non-ATPase regulatory subunit 5 Mus musculus | 674.42 | 36.11 | 55972 | 14 | 14 | 7831379 | 8856142 | 9746756 | 10063931 | 8296208 | 8653305 | 11622876 | 10485882 | 11587257 | 9591136 | 10416021 | 10316296 | 8695538 | 10285460 | 10845950 |
| Q99JI4 | Psmd6 | 26S proteasome non-ATPase regulatory subunit 6 Mus musculus | 1081.49 | 61.95 | 45536 | 20 | 20 | 35138045 | 31516525 | 36492601 | 30299551 | 27870467 | 32982151 | 30841693 | 34913929 | 41049277 | 33126151 | 30579901 | 31486229 | 27866617 | 33979817 | 33906901 |
| P26516 | Psmd7 | 26S proteasome non-ATPase regulatory subunit 7 Mus musculus | 747.02 | 57.63 | 36540 | 13 | 13 | 20427787 | 14682115 | 18734853 | 13634627 | 15109045 | 21202217 | 20499759 | 20417823 | 20886733 | 15956409 | 19255897 | 20435945 | 18662419 | 19885185 | 20165889 |
| Q9CX56 | Psmd8 | 26S proteasome non-ATPase regulatory subunit 8 Mus musculus | 305.13 | 24.93 | 39930 | 7 | 7 | 7137359 | 8078915.5 | 8560070 | 7215239 | 6351252 | 8870267 | 7240834 | 9250976 | 9693083 | 6478867 | 7433696 | 7799549.5 | 8356902 | 9578237 | 8095244.5 |
| P97371 | Psme1 | Proteasome activator complex subunit 1 Mus musculus | 975.63 | 57.03 | 28673 | 15 | 15 | 89540657 | 95525945 | 95758017 | 75130945 | 81183945 | 77432225 | 65626649 | 90223281 | 96883505 | 55299213 | 81488961 | 79773993 | 68114489 | 95421921 | 72298025 |
| P97372 | Psme2 | Proteasome activator complex subunit 2 Mus musculus | 1003.36 | 56.07 | 27057 | 13 | 13 | 44316197 | 46731149 | 47717777 | 34232265 | 37201365 | 41080369 | 38288165 | 47612025 | 53822141 | 33134359 | 51732081 | 48846381 | 41113289 | 52623105 | 40767733 |
| P61290 | Psme3 | Proteasome activator complex subunit 3 Mus musculus | 587.29 | 41.73 | 29506 | 10 | 10 | 9824825 | 10620431 | 10491492 | 10125554 | 10774689 | 10111493 | 8752079 | 9348315 | 8149677 | 9397495 | 8986351 | 9282449 | 10676023 | 10458923 | 8910849 |
| Q8BHL8 | Psmf1 | Proteasome inhibitor PI31 subunit Mus musculus | 73.24 | 4.43 | 29664 | 1 | 1 | 1691224.25 | 1649087.38 | 1543821.88 | 1740047.75 | 1757861.75 | 1619504.75 | 1629131.75 | 1686997.88 | 1565297.88 | 2014497.88 | 1874670 | 1579742.88 | 1726252.25 | 1267617.38 | 1386040.88 |
| Q9JK23 | Psmg1 | Proteasome assembly chaperone 1 Mus musculus | 251.7 | 17.65 | 33104 | 4 | 4 | 6532307 | 6510476 | 5634383 | 6581602 | 6723938.5 | 6521783.5 | 6925000.5 | 6837767 | 6550259.5 | 6009874.5 | 5947568 | 6022981.5 | 6418891 | 6876816 | 6032895.5 |
| Q9EST4 | Psmg2 | Proteasome assembly chaperone 2 Mus musculus | 103.54 | 3.03 | 29524 | 2 | 2 | 1 | 347121.22 | 1 | 1 | 296036.81 | 1 | 864768.44 | 1 | 383271.09 | 647174.75 | 489631.44 | 600059 | 1 | 1 | 713264.94 |
| Q9CZH3 | Psmg3 | Proteasome assembly chaperone 3 Mus musculus | 106.03 | 22.95 | 13331 | 2 | 2 | 1786032.75 | 853473.06 | 1653060 | 1845699 | 4105884 | 1865804 | 1618780 | 1328066.38 | 1434259.25 | 1885574.25 | 1507362.25 | 1891405 | 1835965 | 1570923.5 | 1759070.5 |
| P0C7N9 | Psmg4 | Proteasome assembly chaperone 4 Mus musculus | 80.71 | 33.33 | 13974 | 2 | 2 | 709616.09 | 422771.12 | 248700.12 | 1575617 | 591376.12 | 540769.44 | 356613.75 | 352448.88 | 438808.7375 | 505402.88 | 394481.53 | 290394 | 1294059.12 | 357856.34 | 416538.81 |
| Q4KLH4 | Pspc1 | Paraspeckle component 1 Rattus norvegicus | 180.35 | 9 | 58760 | 2 | 3 | 935184 | 1933783 | 1416285.88 | 1579770.065 | 2033827.38 | 1753243.5 | 1757022.75 | 634264.25 | 2039394.12 | 1439202.25 | 1472555.88 | 1375788.88 | 1314308.88 | 1630062.88 | 1280527.25 |
| P97814 | Pstpip1 | Proline-serine-threonine phosphatase-interacting protein 1 Mus musculus | 282.97 | 17.11 | 47590 | 5 | 5 | 9298495 | 9274206 | 8585177 | 9588877 | 8833164 | 7979274.5 | 6909640 | 7803951 | 8235725 | 7891581 | 8050923.5 | 6541285 | 8315439 | 8028290 | 6663570 |
| P17225 | Ptbp1 | Polypyrimidine tract-binding protein 1 Mus musculus | 1433.44 | 61.86 | 56478 | 3 | 18 | 36129277 | 46671741 | 39793857 | 46020513 | 52002501 | 58203237 | 50407449 | 45924985 | 41225441 | 55155829 | 49510073 | 49313849 | 52045905 | 38655081 | 58904225 |
| Q8BHD7 | Ptbp3 | Polypyrimidine tract-binding protein 3 Mus musculus | 640.14 | 43.98 | 56701 | 8 | 11 | 5983338 | 4179773.25 | 2239316.75 | 5139728 | 7018103 | 5657670 | 8404461 | 5390723 | 5041107.5 | 3787759.75 | 4094367 | 5768920 | 6877242.5 | 4780394.5 | 8601088 |
| Q14C51 | Ptcd3 | Pentatricopeptide repeat domain-containing protein 3, mitochondrial Mus musculus | 41.2 | 1.17 | 77796 | 1 | 1 | 3577212.75 | 1079706.12 | 1003975.44 | 1367289.62 | 1257993.88 | 1242714.5 | 1026440.75 | 1544739.38 | 1473160.38 | 1015439.81 | 1086720.5 | 1102436.12 | 1005738.56 | 1698376 | 839811.94 |
| Q60866 | Pter | Phosphotriesterase-related protein Mus musculus | 148.09 | 11.46 | 39218 | 3 | 3 | 3970088 | 3982317.5 | 4329619 | 4762867 | 3923348.5 | 4664138 | 4918193 | 4239480.5 | 3915235.5 | 4579753 | 4481563.5 | 4206794 | 3947258 | 4444093 | 4412808 |
| P83868 | Ptges3 | Prostaglandin E synthase 3 Rattus norvegicus | 173.34 | 26.25 | 18721 | 3 | 3 | 17011015 | 13231800 | 9545441 | 14601500 | 16514794 | 14582362 | 11462211 | 13124046 | 12282417 | 10070159 | 16438310 | 11708492 | 13570177 | 11803669 | 12535289 |
| Q91YR9 | Ptgr1 | Prostaglandin reductase 1 Mus musculus | 711.79 | 57.75 | 35560 | 13 | 13 | 30446173 | 43041713 | 39012065 | 37214841 | 37579401 | 34058449 | 33369465 | 37481161 | 37256065 | 34186161 | 34449169 | 28422047 | 40287389 | 40317081 | 30583057 |
| Q8VDQ1 | Ptgr2 | Prostaglandin reductase 2 Mus musculus | 218.17 | 15.95 | 38015 | 4 | 4 | 5464668 | 5582771 | 5610796.5 | 5416513 | 6787951 | 5651682 | 4994673 | 4618614 | 4654487.5 | 4985868 | 6444029.5 | 4985827.5 | 6032444.5 | 5751722.5 | 5624075.5 |
| P22437 | Ptgs1 | Prostaglandin G/H synthase 1 Mus musculus | 354.16 | 15.12 | 69042 | 6 | 7 | 318926.88 | 3227523.5 | 2729618 | 6842449 | 5433438.5 | 3693015.25 | 8007340 | 4256412 | 2639490.25 | 6504527.5 | 3947107.25 | 5041612.5 | 3913665.75 | 4822849.5 | 6306760 |
| Q9QVP9 | Ptk2b | Protein-tyrosine kinase 2-beta Mus musculus | 949.64 | 28.25 | 115794 | 20 | 20 | 28455413 | 22882949 | 22717637 | 23024557 | 27615605 | 25209293 | 29832521 | 30859177 | 32050053 | 23046845 | 28831157 | 25656069 | 22287801 | 30755293 | 23496483 |
| P26350 | Ptma | Prothymosin alpha Mus musculus | 518.26 | 36.04 | 12254 | 6 | 6 | 136869921 | 163779153 | 148286337 | 148546609 | 136402113 | 126140729 | 165949233 | 156914545 | 156770737 | 145221489 | 185128273 | 168499521 | 140097473 | 126146577 | 159937185 |
| P04550 | Ptms | Parathymosin Rattus norvegicus | 208.88 | 22.55 | 11559 | 3 | 3 | 39655745 | 50878625 | 34129413 | 44393737 | 47642841 | 32824741 | 34044149 | 34531041 | 36336933 | 39699049 | 37535217 | 36624265 | 42700593 | 37788865 | 27152449 |
| Q5U2W6 | Ptov1 | Prostate tumor-overexpressed gene 1 protein homolog Rattus norvegicus | 51.38 | 4.57 | 46849 | 1 | 1 | 159473.75 | 243645.39 | 307737.06 | 193803.83 | 183177.56 | 221902.98 | 236345.89 | 169205.84 | 173310.44 | 158939.56 | 237393.94 | 165916.92 | 177473.16 | 228011.88 | 250561.28 |
| P58389 | Ptpa | Serine/threonine-protein phosphatase 2A activator Mus musculus | 348.32 | 35.29 | 36710 | 6 | 6 | 10686629 | 10124990 | 9324665 | 7714597.5 | 8110287 | 9650231 | 8335903 | 10535254 | 10525963 | 7323362.5 | 10397893 | 10464449 | 8750284 | 10306647 | 9889179 |
| P35821 | Ptpn1 | Tyrosine-protein phosphatase non-receptor type 1 Mus musculus | 193.39 | 12.5 | 49593 | 5 | 5 | 5496886 | 6492162 | 4846930.5 | 5256600 | 5298270.5 | 5718734.5 | 5655759.125 | 5985242.5 | 5908588.5 | 5010471 | 5091198.5 | 5081913 | 5500860 | 6615809 | 5719012 |
| P35831 | Ptpn12 | Tyrosine-protein phosphatase non-receptor type 12 Mus musculus | 122.42 | 5.29 | 86526 | 3 | 3 | 1075914.88 | 1664086.5 | 2275661 | 2266905.5 | 2073114 | 2333059 | 1919338.88 | 2365919.75 | 1507573.88 | 2157460.75 | 1599590.62 | 2399366.5 | 2304979 | 2533805 | 1469217.12 |
| Q6PB44 | Ptpn23 | Tyrosine-protein phosphatase non-receptor type 23 Mus musculus | 40.18 | 0.71 | 185216 | 1 | 1 | 1 | 1 | 1313953.12 | 1 | 1 | 1 | 1410873.25 | 1 | 1 | 1 | 1 | 1 | 1 | 1 | 1 |
| P29351 | Ptpn6 | Tyrosine-protein phosphatase non-receptor type 6 Mus musculus | 2297.35 | 72.61 | 67559 | 36 | 38 | 126555201 | 130331697 | 140119553 | 135952289 | 137699537 | 132791625 | 170589937 | 131545329 | 136400545 | 141682897 | 152898225 | 147387585 | 131766217 | 131598225 | 159898321 |
| P49445 | Ptpn7 | Tyrosine-protein phosphatase non-receptor type 7 Rattus norvegicus | 46.59 | 3.62 | 40314 | 1 | 1 | 504673.72 | 502326.03 | 530789.44 | 704985.31 | 605939.31 | 414063.91 | 501825.22 | 396904.84 | 315470.19 | 510820.41 | 613955.81 | 515263.69 | 472780.59 | 502754.09 | 582398.19 |
| O35239 | Ptpn9 | Tyrosine-protein phosphatase non-receptor type 9 Mus musculus | 70.43 | 3.71 | 67970 | 2 | 2 | 564572 | 206183.41 | 1 | 1 | 335159.47 | 441312.0225 | 664540 | 374838.72 | 331779.59 | 394089.78 | 247905.98 | 338145.06 | 1 | 1 | 454419.56 |
| P18052 | Ptpra | Receptor-type tyrosine-protein phosphatase alpha Mus musculus | 153.81 | 4.7 | 93698 | 3 | 3 | 3998869.5 | 4070362.5 | 4259567.5 | 3541227 | 3619625 | 3966731 | 5193270 | 3860530 | 3336141 | 5108891 | 4141171.25 | 3452435.5 | 4099288.25 | 3214092.75 | 3127349.5 |
| P06800 | Ptprc | Receptor-type tyrosine-protein phosphatase C Mus musculus | 2183.95 | 32.71 | 144837 | 36 | 37 | 137451825 | 142186913 | 141481121 | 140101329 | 153271585 | 142148673 | 142630129 | 147533457 | 156084049 | 138566753 | 152226401 | 155517457 | 142221921 | 134267633 | 157059569 |
| B2GV87 | Ptpre | Receptor-type tyrosine-protein phosphatase epsilon Rattus norvegicus | 155.66 | 5.87 | 80734 | 3 | 3 | 1086241.88 | 1589133.38 | 1746180.5 | 1015236.94 | 1454266.12 | 1595609.62 | 1510047.75 | 1708245 | 1811719.75 | 1221773.88 | 143847.23 | 2005437.5 | 1493401.75 | 1295198.5 | 1432576.88 |
| E9Q612 | Ptpro | Receptor-type tyrosine-protein phosphatase O Mus musculus | 40.97 | 0.9 | 138589 | 1 | 1 | 1243094 | 1322124.5 | 897329 | 1212953 | 1481179.75 | 1414661.25 | 1129347.62 | 1041716.44 | 875006.19 | 1137118 | 1168129.12 | 1178105.38 | 1246654.25 | 1121494.38 | 1007905.25 |
| Q8R2Y8 | Ptrh2 | Peptidyl-tRNA hydrolase 2, mitochondrial Mus musculus | 138.26 | 27.62 | 19527 | 3 | 3 | 3005910.5 | 2955586 | 3802728.5 | 3109108.5 | 4159630 | 4066211.5 | 3041505 | 3222833.75 | 3259444.5 | 2823532.5 | 4025072.5 | 3201273.75 | 3635496.5 | 3563348.5 | 3936388.75 |
| Q8R143 | Pttg1ip | Pituitary tumor-transforming gene 1 protein-interacting protein Mus musculus | 68.09 | 8.62 | 19965 | 2 | 2 | 3739812 | 3049899.5 | 2303842.5 | 1109900.88 | 2465014.75 | 2933901.5 | 818443.31 | 2567969.5 | 2642475.25 | 1592065.75 | 1694182.75 | 1355375.25 | 1740640.25 | 2697098 | 954992.81 |
| Q3UEB3 | Puf60 | Poly(U)-binding-splicing factor PUF60 Mus musculus | 835.78 | 34.75 | 60249 | 13 | 13 | 21663497 | 19814313 | 20259581 | 17078499 | 16589848 | 17813015 | 26472347 | 18406417 | 16265367 | 21007323 | 22687497 | 23267315 | 16360435 | 21364075 | 21370243 |
| Q8BKS9 | Pum3 | Pumilio homolog 3 Mus musculus | 315.32 | 12.06 | 72800 | 5 | 5 | 2330401.25 | 988184.69 | 4035540.75 | 1167950.75 | 1099484.5 | 1547791.12 | 1170958.75 | 509302 | 1530249.5 | 192235.98 | 2521669.25 | 1600978.62 | 1072614.38 | 404857.75 | 2231860.5 |
| P42669 | Pura | Transcriptional activator protein Pur-alpha Mus musculus | 268.63 | 20.87 | 34884 | 4 | 4 | 1150833.62 | 905254.06 | 1 | 1301468.38 | 2129022.25 | 1386218.75 | 1145850.218 | 1501154 | 677249.12 | 1018779 | 1068913.983 | 375355.56 | 1379645.25 | 1783052.62 | 737602.5 |
| O35295 | Purb | Transcriptional activator protein Pur-beta Mus musculus | 225.25 | 24.07 | 33901 | 5 | 5 | 3858990.5 | 4825599 | 3841536.75 | 4611350 | 4764654.5 | 5250101 | 6905895.5 | 5837604 | 3114281.5 | 4755829.5 | 6213444.5 | 5672742 | 4670730.5 | 4834612.5 | 8436056 |
| Q91VU7 | Pus7 | Pseudouridylate synthase 7 homolog Mus musculus | 171.23 | 9.7 | 74793 | 4 | 4 | 994688.25 | 618611.925 | 941911.62 | 281371.38 | 256476.45 | 1 | 1116127.5 | 758627.25 | 1 | 946415.38 | 716554.88 | 425006.31 | 669099.94 | 736657.5 | 435819.16 |
| Q99LL5 | Pwp1 | Periodic tryptophan protein 1 homolog Mus musculus | 159.94 | 11.98 | 55587 | 3 | 3 | 1304749.25 | 1121091.5 | 1550567.5 | 1523970.5 | 1444675.25 | 923845.62 | 1271006.25 | 1389861.88 | 1003989.19 | 1814135.12 | 978552 | 1903119.25 | 1 | 1 | 1877999.25 |
| Q8BU03 | Pwp2 | Periodic tryptophan protein 2 homolog Mus musculus | 190.18 | 6.42 | 102910 | 4 | 4 | 966987.56 | 924193.19 | 874700.38 | 771334.69 | 928563.06 | 922506.06 | 898877.94 | 1021896.19 | 1004003.62 | 1838072.5 | 752125.56 | 1380612.5 | 875357.5 | 768983.31 | 1517670.12 |
| Q8VI36 | Pxn | Paxillin Mus musculus | 80.45 | 9.98 | 64476 | 2 | 2 | 132730.3 | 1 | 1 | 1 | 1 | 228257.44 | 224912.44 | 178405.8 | 174848.36 | 201606.01 | 1 | 1 | 226889.34 | 1 | 228691.73 |
| Q9EPB4 | Pycard | Apoptosis-associated speck-like protein containing a CARD Mus musculus | 455.27 | 56.99 | 21459 | 8 | 8 | 16844279 | 12940189 | 15560411 | 14470844 | 12986655 | 14783743 | 16385940 | 16556210 | 12455200 | 11825238 | 15214829 | 18091293 | 13785192 | 13885901 | 14293007 |
| Q922Q4 | Pycr2 | Pyrroline-5-carboxylate reductase 2 Mus musculus | 704.07 | 35.31 | 33659 | 10 | 10 | 18715853 | 16591343 | 17587775 | 21074971 | 21293397 | 20108017 | 27966041 | 18560263 | 17765955 | 23566789 | 21217779 | 17820171 | 21349573 | 18398891 | 18840713 |
| P53534 | Pygb | Glycogen phosphorylase, brain form Rattus norvegicus | 997.77 | 23.39 | 96174 | 13 | 16 | 38852877 | 40796329 | 42306413 | 36641517 | 39003997 | 40018929 | 37348593 | 41387297 | 44803833 | 38795773 | 40348505 | 39404781 | 37251281 | 41502369 | 39944345 |
| P09811 | Pygl | Glycogen phosphorylase, liver form Rattus norvegicus | 237.34 | 7.53 | 97483 | 1 | 4 | 9630024 | 9077226 | 9885653 | 9628375 | 10200595 | 10177779 | 8804851 | 9910746 | 10932997 | 10825972 | 9989645 | 9071025 | 9772832 | 9729255 | 10277826 |
| Q8BML9 | Qars | Glutamine--tRNA ligase Mus musculus | 641.94 | 21.03 | 87677 | 14 | 14 | 10923693 | 12416317 | 13178927 | 11004910 | 11090575 | 12922905 | 9599288 | 12675586 | 11377319 | 12610415 | 14536939 | 13243258 | 11439175 | 13074809 | 14414360 |
| Q8BVI4 | Qdpr | Dihydropteridine reductase Mus musculus | 342.47 | 33.61 | 25570 | 5 | 5 | 5157324.5 | 5930228.5 | 6655475 | 5754099.5 | 5579340 | 5382977 | 8050064 | 6065327.5 | 5019559 | 7423363 | 7350944.5 | 7264231 | 5823841 | 6298136 | 6584812 |

| | | | | | | | | | | | | | | | | | | | | | | |
|---|---|---|---|---|---|---|---|---|---|---|---|---|---|---|---|---|---|---|---|---|---|---|
| Q91XU1 | Qki | Protein quaking Rattus norvegicus | 307.42 | 29.62 | 37643 | 8 | 8 | 12454134 | 12073374 | 11243338 | 12401428 | 11836760 | 11752992 | 12612518 | 12272901 | 11578730 | 11881159 | 11450831 | 11060240 | 11941747 | 12150130 | 10659613 |
| Q91X91 | Qprt | Nicotinate-nucleotide pyrophosphorylase [carboxylating] Mus musculus | 248.55 | 22.41 | 31530 | 5 | 5 | 703643.69 | 3376261.25 | 2422194.5 | 1380101.25 | 676490.81 | 1643066.62 | 844171.88 | 2360696 | 299485.44 | 341783.44 | 1962863.75 | 451082.31 | 832059.44 | 2531833 | 673953.81 |
| B8ZXI1 | Qtrt2 | Queuine tRNA-ribosyltransferase accessory subunit 2 Mus musculus | 36.87 | 3.86 | 46288 | 1 | 1 | 1786591.12 | 1844135.25 | 664873.75 | 1860417 | 1881407.38 | 1960697.88 | 1496493 | 1537258.5 | 1348760.5 | 497518.25 | 1316937.62 | 1 | 1471475.25 | 1348192.62 | 1309158.88 |
| P61027 | Rab10 | Ras-related protein Rab-10 Mus musculus | 490.94 | 42 | 22541 | 7 | 10 | 21850323 | 21533451 | 21389139 | 20575261 | 19195639 | 22414771 | 24549345 | 22130397 | 24093405 | 22802987 | 22346657 | 21915347 | 20294859 | 21344905 | 23429885 |
| O35509 | Rab11b | Ras-related protein Rab-11B Rattus norvegicus | 556.61 | 50.92 | 24489 | 11 | 11 | 69784809 | 61384121 | 58891449 | 50684601 | 51708893 | 57576489 | 59140133 | 63768661 | 66859153 | 54703369 | 51251041 | 58935909 | 53962693 | 57336233 | 54003417 |
| Q9D620 | Rab11fip1 | Rab11 family-interacting protein 1 Mus musculus | 96.99 | 5.43 | 70684 | 2 | 2 | 1 | 1 | 188517.3 | 1 | 1 | 163756.91 | 129644.3 | 192197.28 | 1 | 1 | 1 | 1 | 1 | 1 | 1 |
| P35283 | Rab12 | Ras-related protein Rab-12 Mus musculus | 149.63 | 12.76 | 27329 | 2 | 3 | 1353021.5 | 618446.38 | 1338756.5 | 849985.31 | 917115.62 | 745929.19 | 848737.38 | 1170509.25 | 1477587.75 | 541788.5 | 1187288.5 | 1639938.75 | 777616.38 | 1429961 | 1266699.12 |
| Q91V41 | Rab14 | Ras-related protein Rab-14 Mus musculus | 815.37 | 80 | 23897 | 12 | 14 | 25779797 | 25251989 | 26092057 | 26823629 | 27031935 | 26280155 | 29128297 | 25126925 | 28368967 | 28239017 | 30238873 | 27247043 | 27063093 | 26129909 | 30110257 |
| P35293 | Rab18 | Ras-related protein Rab-18 Mus musculus | 320.7 | 41.26 | 23035 | 6 | 6 | 4780978 | 4271044.5 | 4749120 | 3416822.5 | 2344066.5 | 3726898 | 3753096 | 5050686 | 5726793.5 | 3217098.25 | 3595877.5 | 3547333 | 2923707.25 | 4960284 | 5248573 |
| P62821 | Rab1A | Ras-related protein Rab-1A Mus musculus | 874.43 | 68.78 | 22678 | 7 | 15 | 50089177 | 53785033 | 55181625 | 40476249 | 45167837 | 46451489 | 50056553 | 54818921 | 60102953 | 44725789 | 52823241 | 52510601 | 48496985 | 55065801 | 50308645 |
| P10536 | Rab1b | Ras-related protein Rab-1B Rattus norvegicus | 837.06 | 72.14 | 22163 | 6 | 13 | 40798405 | 45746005 | 45516297 | 37083493 | 36079633 | 34850977 | 45926961 | 46113413 | 50992997 | 42262421 | 40286489 | 43919973 | 41154561 | 44668453 | 44462585 |
| P35282 | Rab21 | Ras-related protein Rab-21 Mus musculus | 264.32 | 25.23 | 24106 | 4 | 4 | 2766645.25 | 3741388 | 4337648 | 2008819 | 3630291.25 | 4127224.5 | 3723402 | 5249527.5 | 3754618 | 3857296.5 | 4177080.25 | 4253510.5 | 2528999 | 3541165 | 4264561.5 |
| P35285 | Rab22a | Ras-related protein Rab-22A Mus musculus | 90.91 | 12.89 | 21802 | 1 | 2 | 1856008.5 | 2247866 | 2327043.5 | 2879868 | 526297.94 | 2320947.25 | 1952677.25 | 2379402.75 | 1782504.75 | 3169204.25 | 2713845.875 | 2593341.25 | 2828733.25 | 2633957.5 | 2799351.5 |
| P35290 | Rab24 | Ras-related protein Rab-24 Mus musculus | 35.96 | 6.9 | 23144 | 1 | 1 | 577564.44 | 582131.75 | 1717788.25 | 934448.88 | 599634.31 | 1 | 539703 | 2569173.5 | 1 | 1 | 1565944.75 | 934012.31 | 524753.75 | 730555.44 | 938816.5625 |
| P23640 | Rab27a | Ras-related protein Rab-27A Rattus norvegicus | 40.34 | 9.5 | 25068 | 1 | 1 | 1 | 134758.64 | 1 | 235721.61 | 1 | 97772.61 | 112160.51 | 118793.05 | 100866.88 | 131209.5 | 115457.44 | 136726.52 | 1 | 140313.5 | 116278.74 |
| P05712 | Rab2a | Ras-related protein Rab-2A Rattus norvegicus | 527.07 | 46.7 | 23536 | 9 | 9 | 11523637 | 10334534 | 11464839 | 8168102.5 | 9611938 | 10864496 | 9293991 | 8674693 | 13868620 | 9993659 | 10887408 | 10587430 | 8508561 | 11514820 | 9053906 |
| Q921E2 | Rab31 | Ras-related protein Rab-31 Mus musculus | 240.24 | 24.74 | 21331 | 3 | 4 | 6154674.5 | 6432402 | 7534732 | 7578820 | 3406910 | 4372826.5 | 6656660 | 7182610.5 | 6933237 | 7693797 | 4692328 | 7140588 | 7175379 | 6791934 | 6916402 |
| Q9CZE3 | Rab32 | Ras-related protein Rab-32 Mus musculus | 80.73 | 10.76 | 25068 | 2 | 2 | 4747686 | 2531041 | 2179624.25 | 2626370.75 | 1742248.25 | 1804326.75 | 328220.88 | 2684827 | 3330051 | 2198127.25 | 725567 | 2404381 | 2218022 | 2448133.5 | 1427903.62 |
| Q5U316 | Rab35 | Ras-related protein Rab-35 Rattus norvegicus | 318.15 | 32.84 | 23025 | 4 | 5 | 8934905 | 7395600 | 6895737 | 6688692 | 7213246 | 6667119 | 6149368 | 7119379 | 9365408 | 7217007 | 6157352 | 6507396.5 | 7305264 | 8001960 | 6371037 |
| P35276 | Rab3d | Ras-related protein Rab-3D Mus musculus | 229.91 | 14.16 | 24416 | 1 | 4 | 19148383 | 13234569 | 13854055 | 15892462 | 13911381 | 17122511 | 10266790 | 14227222 | 16109384 | 13343683 | 16237969 | 10400051 | 12284448 | 14250737 | 12772075 |
| Q80UJ7 | Rab3gap1 | Rab3 GTPase-activating protein catalytic subunit Mus musculus | 267.42 | 8.87 | 110198 | 6 | 6 | 4529544 | 4017258 | 3924811 | 3890960.75 | 3313279.25 | 4674397.5 | 2838800.75 | 4381949 | 5222209 | 4453786 | 3099251 | 4148622.5 | 4077519.25 | 3670789 | 2913982.25 |
| Q8BMG7 | Rab3gap2 | Rab3 GTPase-activating protein non-catalytic subunit Mus musculus | 516.99 | 10.61 | 152535 | 11 | 11 | 3521419.5 | 2475348.75 | 3809159.5 | 3572298.75 | 4909346 | 3155280 | 3186837.5 | 3330440.75 | 4186848.5 | 5978675 | 3863926.5 | 5191364 | 2854715.5 | 2656678.5 | 3151664 |
| Q8VDV3 | Rab3il1 | Guanine nucleotide exchange factor for Rab-3A Mus musculus | 42.02 | 5.22 | 42713 | 1 | 1 | 243104.86 | 142138.69 | 184855.16 | 275147.28 | 229194.58 | 198883.97 | 95166.98 | 162331.33 | 1 | 1 | 329281.5 | 279193 | 255428.31 | 171762.42 | 258916.3075 |
| Q53B90 | Rab43 | Ras-related protein Rab-43 Rattus norvegicus | 119.83 | 11.9 | 23229 | 1 | 2 | 723732.56 | 825376.44 | 648969.88 | 710570.33 | 644202.44 | 539916.81 | 560097.69 | 828195.44 | 693280.5 | 467389.06 | 729306.88 | 767082.88 | 597814.56 | 622019.94 | 648381.44 |
| P51146 | Rab4b | Ras-related protein Rab-4B Rattus norvegicus | 426.26 | 46.48 | 23629 | 6 | 7 | 3815972 | 3995803.75 | 3681148.5 | 4220570.5 | 4985689 | 4284792 | 3313503 | 4740453 | 4014088.25 | 5207389.5 | 5072805 | 5258936 | 3760072.25 | 3700416 | 4653293 |
| Q9CQD1 | Rab5a | Ras-related protein Rab-5A Mus musculus | 424.65 | 48.84 | 23599 | 5 | 7 | 14677289 | 14337149 | 14606747 | 13195073 | 13316318 | 12811042 | 11685133 | 15368421 | 22230985 | 12885306 | 12961747 | 15183495 | 11924106 | 15485415 | 13511973 |
| P61021 | Rab5b | Ras-related protein Rab-5B Mus musculus | 329.18 | 32.09 | 23707 | 3 | 5 | 14392134 | 12708080 | 13514980 | 10427497 | 12971203 | 12305253 | 10986209 | 13317440 | 20278487 | 10389993 | 12063124 | 13096990 | 9492832 | 13533747 | 10572046 |
| P35278 | Rab5c | Ras-related protein Rab-5C Mus musculus | 630.37 | 67.13 | 23413 | 7 | 9 | 41195041 | 37879833 | 39473465 | 38617997 | 43166841 | 39663121 | 34993433 | 38275005 | 46313745 | 35486381 | 33935101 | 37302553 | 39968557 | 40097969 | 34411789 |
| P35279 | Rab6a | Ras-related protein Rab-6A Mus musculus | 401.88 | 42.31 | 23590 | 7 | 8 | 1844885.25 | 528157 | 3234779.25 | 1951627.75 | 1326213.5 | 1864997.75 | 4840913.5 | 3054175.5 | 3750136.75 | 3071151 | 1815067.5 | 1909401.75 | 2670387.5 | 1683272.62 | 4194085.75 |
| P09527 | Rab7a | Ras-related protein Rab-7a Rattus norvegicus | 1025.46 | 68.12 | 23504 | 14 | 15 | 97731393 | 109525561 | 103554041 | 103969705 | 107382417 | 102180417 | 89586025 | 105958585 | 107245825 | 97257081 | 83996561 | 95928697 | 102199545 | 103322585 | 97441481 |
| Q8VEA8 | Rab7b | Ras-related protein Rab-7b Mus musculus | 62.26 | 7.54 | 22502 | 1 | 1 | 1112938.12 | 1039819.06 | 913769.69 | 477229.94 | 848847.12 | 1072858.5 | 891576.75 | 925611.19 | 1203883.12 | 1023482.39 | 983631.19 | 952729.11 | 951635.75 | 1089111.75 | 786537.75 |
| P35280 | Rab8a | Ras-related protein Rab-8A Rattus norvegicus | 596.15 | 51.69 | 23668 | 5 | 11 | 5161799 | 6839662 | 3150111.75 | 7603442 | 4447727 | 5223365 | 5487628.5 | 5999380.5 | 7240934 | 6774634 | 6333389.5 | 5316387 | 5490535 | 7844269.5 | 4379080.5 |
| P61028 | Rab8b | Ras-related protein Rab-8B Mus musculus | 643.59 | 59.42 | 23603 | 6 | 12 | 35356293 | 35358469 | 28363077 | 31688473 | 30676647 | 30444615 | 26692663 | 29812517 | 36229945 | 30202529 | 29517471 | 26809577 | 32108909 | 32395159 | 28891997 |
| Q99P75 | Rab9a | Ras-related protein Rab-9A Rattus norvegicus | 123.28 | 23.38 | 22896 | 3 | 3 | 1211371.12 | 2999493.5 | 4264398.5 | 3507301.5 | 2999012.5 | 2637220 | 3949582 | 3735852.75 | 3832931.5 | 3229463 | 2940606.75 | 3488690.5 | 1533251.88 | 3221526.5 | 1483139.75 |
| O35550 | Rabep1 | Rab GTPase-binding effector protein 1 Rattus norvegicus | 57.62 | 2.2 | 99428 | 1 | 1 | 643260.19 | 581520.31 | 724240.25 | 902920.62 | 808063.19 | 639264.5 | 501295.69 | 671163.62 | 543435.44 | 611807.62 | 661475.5 | 612623.38 | 580553.88 | 733464.88 | 576701.31 |
| A2AWA9 | Rabgap1 | Rab GTPase-activating protein 1 Mus musculus | 81.95 | 2.54 | 120798 | 2 | 2 | 935754.94 | 1237422.44 | 801623.69 | 920566.38 | 2291744.75 | 1 | 680578.38 | 1 | 1 | 842574.5 | 603713.38 | 1708585 | 966373.25 | 1009624.048 | 759824.56 |
| Q9JHK4 | Rabggta | Geranylgeranyl transferase type-2 subunit alpha Mus musculus | 99.73 | 7.41 | 64989 | 2 | 2 | 1014871.25 | 1558044.5 | 930421.31 | 1962574 | 1197761.5 | 692981.06 | 1228663.25 | 787764 | 925654.44 | 1424792.5 | 422641.88 | 769033.5 | 1775010.75 | 311886.75 | 611318.56 |
| Q5U3K5 | Rabl6 | Rab-like protein 6 Mus musculus | 58.35 | 1.52 | 79831 | 1 | 1 | 532287.44 | 572339.12 | 348638.31 | 373884.62 | 453624.84 | 476133.06 | 522774.25 | 494525.56 | 474796.62 | 390854 | 396220.41 | 343881.25 | 446448.09 | 510138.69 | 483000 |
| P63001 | Rac1 | Ras-related C3 botulinum toxin substrate 1 Mus musculus | 502.89 | 51.04 | 21450 | 4 | 8 | 117916473 | 124658545 | 104534513 | 105572913 | 111204041 | 95567065 | 105353017 | 95020897 | 105875465 | 100663809 | 105636841 | 91149673 | 113064169 | 110298529 | 90888913 |
| Q05144 | Rac2 | Ras-related C3 botulinum toxin substrate 2 Mus musculus | 504.12 | 51.04 | 21441 | 4 | 8 | 157320433 | 173500753 | 153848577 | 153689553 | 156357969 | 137894273 | 140611185 | 141572081 | 154037185 | 144666257 | 131843929 | 123000297 | 162315857 | 161514865 | 122822481 |
| Q9WVM1 | Racgap1 | Rac GTPase-activating protein 1 Mus musculus | 125.57 | 6.05 | 70158 | 3 | 3 | 888202.88 | 1045679.06 | 1100534 | 830627.69 | 1006828.88 | 1258749.88 | 1087599.75 | 1310430.25 | 1376474.62 | 785016.5 | 1537023.25 | 1264069.75 | 1197067.62 | 1196775 | 758526.94 |
| P63245 | Rack1 | Receptor of activated protein C kinase 1 Rattus norvegicus | 1368.58 | 86.75 | 35077 | 20 | 20 | 148829329 | 154126033 | 163496273 | 146850577 | 149176609 | 157570561 | 172445329 | 158052033 | 168914849 | 158741809 | 167660721 | 160151057 | 153476337 | 160341793 | 165804849 |
| Q61550 | Rad21 | Double-strand-break repair protein rad21 homolog Mus musculus | 172.78 | 9.13 | 72083 | 4 | 4 | 2132063.25 | 1589362.88 | 1646797 | 3037397.5 | 2517096 | 1929230.25 | 1687557.25 | 1551177 | 1066532.5 | 1054067.25 | 2019613.25 | 1525110.12 | 2220672.75 | 1904101.25 | 2418089 |
| P54726 | Rad23a | UV excision repair protein RAD23 homolog A Mus musculus | 123.61 | 8.26 | 39706 | 1 | 1 | 1758432 | 1334628.5 | 1559897.5 | 1908512 | 1431107.25 | 1534476.88 | 1693224.88 | 1793258.25 | 1126778.88 | 1819964.12 | 3114062.5 | 1644214.5 | 1353191.75 | 1427210.62 | 1916857.75 |
| P54728 | Rad23b | UV excision repair protein RAD23 homolog B Mus musculus | 179.3 | 11.54 | 43513 | 3 | 3 | 2478165.5 | 1950238.75 | 3278661.5 | 2996743.5 | 2192811.25 | 3035778 | 3267833 | 2978861.5 | 2063570.5 | 2240879.5 | 2007333.25 | 2126395 | 2770447 | 3433534.5 | 3011628 |
| P70388 | Rad50 | DNA repair protein RAD50 Mus musculus | 453.79 | 9.15 | 153488 | 9 | 9 | 3588007.25 | 3754498 | 5015912 | 4149389.75 | 4204187 | 3520238 | 5515793.5 | 4038270.25 | 3762428.5 | 5249799.5 | 3924181.5 | 4446630.5 | 4129331.5 | 3617528.75 | 4801443 |
| Q3SWS8 | Rae1 | mRNA export factor Rattus norvegicus | 158.18 | 14.13 | 40919 | 4 | 4 | 2581383.5 | 444001.88 | 1662393.38 | 2962797 | 2986501 | 2529761.75 | 985918.12 | 1592652.88 | 2185719.5 | 3727858 | 2235835.905 | 2350539.75 | 2411150.25 | 1900743.12 | 2280910.5 |
| O08602 | Raet1a | Retinoic acid early-inducible protein 1-alpha Mus musculus | 51.58 | 4.74 | 28616 | 1 | 1 | 2361884.5 | 2118501.75 | 1474918.38 | 1908892.25 | 1955331.5 | 2096614 | 1980076.658 | 2036199.5 | 2200479.75 | 1587013.38 | 1853108.88 | 1408053.88 | 1814551.88 | 2156027 | 1302800.88 |
| P11345 | Raf1 | RAF proto-oncogene serine/threonine-protein kinase Rattus norvegicus | 81.69 | 3.55 | 72928 | 1 | 2 | 1449602.75 | 1954012.88 | 1695082.62 | 1398639.25 | 1985536.25 | 1702077.25 | 1347063.75 | 1562572.62 | 1990556.12 | 2022789.25 | 1601741.5 | 1517448.25 | 1923005.12 | 1627259.75 | 1557308.62 |
| P63321 | Rala | Ras-related protein Ral-A Mus musculus | 343.61 | 40.29 | 23553 | 1 | 6 | 4510105.5 | 3817739.25 | 4884840 | 4392306.5 | 5976085 | 5002385 | 5690717.5 | 5368738 | 4496577.5 | 4736729 | 4890178 | 5507657.5 | 5161133.5 | 5217241 | 6311813 |
| Q9JIW9 | Ralb | Ras-related protein Ral-B Mus musculus | 396.86 | 50.97 | 23349 | 3 | 8 | 5540295 | 4588644.5 | 5215243 | 3503016 | 5817572.5 | 5520307 | 6592695.5 | 4781339.5 | 2973128.5 | 5687505.5 | 5896857.5 | 4572626 | 5569053 | 5547903.5 | 7090303 |
| A3KGS3 | Ralgapa2 | Ral GTPase-activating protein subunit alpha-2 Mus musculus | 100.01 | 1.28 | 210288 | 2 | 2 | 750213.25 | 618278.19 | 617803.38 | 1975379.62 | 214315.11 | 1237959.75 | 838077.88 | 846027.81 | 1124343 | 208244.23 | 849946.38 | 185024.78 | 218650.72 | 916990.25 | 542653.0325 |
| Q8BQZ4 | Ralgapb | Ral GTPase-activating protein subunit beta Mus musculus | 219.53 | 4.45 | 165200 | 4 | 4 | 2820274.5 | 1817084.12 | 1522997.75 | 2239717.5 | 1341277.5 | 2203928 | 2731557.75 | 2833176.75 | 2087429.62 | 2544330.75 | 3885782.5 | 2521724.5 | 2174848.75 | 2570440.25 | 2327749 |
| Q64012 | Raly | RNA-binding protein Raly Mus musculus | 537.27 | 32.37 | 33188 | 11 | 11 | 13281002 | 14878608 | 14140905 | 15923953 | 16666228 | 13704714 | 14973231 | 11122777 | 13328677 | 15420375 | 21732941 | 15784088 | 13854414 | 11387357 | 20958663 |
| P62827 | Ran | GTP-binding nuclear protein Ran Mus musculus | 787.85 | 57.87 | 24423 | 13 | 13 | 25982069 | 24143423 | 22978157 | 25446659 | 33917089 | 31762579 | 46462697 | 24334681 | 21410821 | 36919797 | 34570865 | 28659241 | 30700889 | 25121837 | 44584205 |
| P34022 | Ranbp1 | Ran-specific GTPase-activating protein Mus musculus | 205.91 | 20.69 | 23596 | 5 | 5 | 6620613.5 | 7181381 | 2928177 | 7263486.5 | 6782964 | 6040965 | 6445848 | 5677277 | 5359052 | 7369050 | 6401442 | 6082846.5 | 6668476 | 5636021.5 | 7332721 |
| Q6VN19 | Ranbp10 | Ran-binding protein 10 Mus musculus | 57.29 | 2.26 | 67188 | 1 | 1 | 394196.38 | 405803.62 | 443727.66 | 964522.69 | 901931.25 | 845884.94 | 1046312.25 | 653085.06 | 603937.88 | 1253675.62 | 1105480.12 | 823594.81 | 853795.88 | 958109.2975 | 1049566.38 |
| Q9ERU9 | Ranbp2 | E3 SUMO-protein ligase RanBP2 Mus musculus | 1516.77 | 13.72 | 341121 | 27 | 27 | 11768863 | 10494381 | 14115749 | 10526775 | 10692732 | 10162623 | 12214579 | 14892794 | 12020124 | 12672151 | 11786133 | 12964330 | 8298740 | 10087925 | 11600552 |
| Q9CT10 | Ranbp3 | Ran-binding protein 3 Mus musculus | 397.34 | 18.53 | 52573 | 6 | 6 | 8239164.5 | 6848450 | 5475961.5 | 6928614.5 | 10246101 | 6784169 | 10626560 | 6338119 | 3567941.75 | 8594131 | 6346961 | 6368646 | 7293615.5 | 6695310 | 7709434 |
| P46061 | Rangap1 | Ran GTPase-activating protein 1 Mus musculus | 1367.74 | 45.16 | 63531 | 25 | 26 | 20220831 | 21796129 | 20268741 | 17210879 | 16143941 | 15422770 | 21051931 | 26411297 | 27670637 | 18415669 | 25619179 | 23381051 | 18590947 | 24592497 | 28129085 |
| P62835 | Rap1a | Ras-related protein Rap-1A Mus musculus | 974.52 | 76.63 | 20987 | 4 | 14 | 121260257 | 124766001 | 110734497 | 116421289 | 112651857 | 122432673 | 106912601 | 120162361 | 130077433 | 119087377 | 109165841 | 108909337 | 117251601 | 125596633 | 106013641 |
| Q99JI6 | Rap1b | Ras-related protein Rap-1b Mus musculus | 1023.91 | 80.43 | 20825 | 5 | 16 | 136927169 | 141910849 | 128245809 | 131854817 | 126793345 | 138092017 | 119998257 | 136780609 | 147930049 | 132150849 | 120950049 | 121767537 | 132533417 | 142390945 | 117884001 |
| P61226 | Rap2b | Ras-related protein Rap-2b Mus musculus | 358.41 | 48.09 | 20504 | 2 | 6 | 10758406 | 11178974 | 12366614 | 11726889 | 9633748 | 10184477 | 9530767 | 12335845 | 12131833 | 11248148 | 10932527 | 12480315 | 10994007 | 11912981 | 11147952 |
| Q8BU31 | Rap2c | Ras-related protein Rap-2c Mus musculus | 277.87 | 38.25 | 20745 | 1 | 5 | 10418436 | 17213223 | 9644153 | 13419739 | 15866324 | 8818359 | 7634529 | 16103499 | 18533381 | 8444860 | 8833030 | 9513481 | 8518815 | 14344186 | 8866649 |
| F1M386 | Rapgef2 | Rap guanine nucleotide exchange factor 2 Rattus norvegicus | 41.99 | 0.87 | 166388 | 1 | 1 | 480017.22 | 612132.38 | 555573.75 | 308647.25 | 472058.34 | 605064.5 | 475549.34 | 768776.88 | 704809.19 | 406267.56 | 988363.81 | 693867.12 | 770892.1225 | 672832.25 | 728505.31 |
| Q9D0I9 | Rars | Arginine--tRNA ligase, cytoplasmic Mus musculus | 1781.69 | 47.12 | 75674 | 29 | 31 | 118466857 | 109909561 | 121923481 | 111019025 | 108712305 | 114475465 | 113479345 | 119297329 | 130067081 | 115751649 | 111493513 | 118582313 | 111626177 | 115423321 | 119873993 |
| Q3U186 | Rars2 | Probable arginine--tRNA ligase, mitochondrial Mus musculus | 190.86 | 4.84 | 65336 | 2 | 2 | 1508299.5 | 1395321.88 | 1814915.5 | 1228651.75 | 1364387 | 1577085.5 | 1418283 | 1749385.25 | 1818606.25 | 1245689 | 1975537.25 | 1627694.25 | 1636825.25 | 1616103 | 1798420.75 |
| Q60790 | Rasa3 | Ras GTPase-activating protein 3 Mus musculus | 189.07 | 4.68 | 95987 | 4 | 4 | 4543903 | 5110698.5 | 5025591 | 4775877 | 5900853 | 4988440.5 | 4181027 | 4639062 | 5875388.5 | 4837963 | 4694122 | 4711689.5 | 5006582 | 5458027.5 | 4839378.5 |
| Q6PFQ7 | Rasa4 | Ras GTPase-activating protein 4 Mus musculus | 72.49 | 4.49 | 90060 | 2 | 2 | 458248.4375 | 514896.69 | 410078.59 | 465511.75 | 442506.72 | 496946.9675 | 572664.62 | 489084.72 | 379529.47 | 546509.06 | 673999.12 | 420512.44 | 561427.19 | 594783.2825 | 723194.38 |
| Q8C2K5 | Rasal3 | RAS protein activator like-3 Mus musculus | 178.56 | 5.67 | 114782 | 4 | 4 | 836467.88 | 783370.44 | 924220 | 756868.38 | 873788.38 | 947397.19 | 888160.69 | 1117444.75 | 1018222.19 | 903517.38 | 846915.88 | 1057478.25 | 909133.25 | 1014630.69 | 878119.38 |
| Q8CB96 | Rassf4 | Ras association domain-containing protein 4 Mus musculus | 195.53 | 11.8 | 36934 | 3 | 3 | 1276281.25 | 966730 | 1242547.62 | 1017789.06 | 960419.12 | 1103710.38 | 1914337.5 | 1092949.25 | 1120694 | 1174117.75 | 1337999.88 | 1125515.75 | 932151.88 | 1192824.12 | 1387405.5 |
| Q9CW46 | Raver1 | Ribonucleoprotein PTB-binding 1 Mus musculus | 387.71 | 21.66 | 79382 | 8 | 8 | 1 | 541926 | 1 | 1 | 1 | 1 | 1253157.5 | 1 | 424953.62 | 836435.06 | 334614.47 | 519089.34 | 451210.28 | 1 | 1 |
| Q60972 | Rbbp4 | Histone-binding protein RBBP4 Mus musculus | 390.27 | 22.35 | 47656 | 3 | 7 | 26910333 | 26352515 | 26713375 | 27903791 | 29951127 | 26322243 | 25654647 | 25885757 | 25857219 | 24366521 | 26589383 | 26965031 | 26018345 | 26632221 | 24660207 |
| Q8BX09 | Rbbp5 | Retinoblastoma-binding protein 5 Mus musculus | 57.48 | 1.67 | 59098 | 1 | 1 | 1099447.25 | 1230452.38 | 1205071 | 1385181.62 | 1310407.38 | 1110323.62 | 755671.19 | 1103980.75 | 1415510.5 | 1023861.12 | 1105930.5 | 948955.94 | 908911.62 | 1153794.5 | 879509.94 |
| Q60973 | Rbbp7 | Histone-binding protein RBBP7 Mus musculus | 335.63 | 22.12 | 47790 | 2 | 6 | 21178625 | 20790347 | 20901695 | 22812941 | 24752673 | 21053733 | 19667089 | 20613129 | 19419183 | 19703325 | 20016581 | 20420863 | 21444059 | 21649737 | 19451945 |
| Q64701 | Rbl1 | Retinoblastoma-like protein 1 Mus musculus | 76.44 | 1.69 | 119456 | 2 | 2 | 853589.38 | 760856.25 | 1096405 | 1326140 | 1091439.62 | 831950.81 | 715228.31 | 856842.62 | 1071018.75 | 1197898.25 | 1162736.5 | 1421939.12 | 1388330 | 818282.56 | 1320663.88 |
| Q8C2Q3 | Rbm14 | RNA-binding protein 14 Mus musculus | 435.13 | 22.57 | 69449 | 10 | 10 | 6665431.5 | 7423135.5 | 7817220.5 | 10178391 | 8662953 | 8167562.5 | 9929490 | 6395138.5 | 6674108 | 9383995 | 7909004.5 | 8277538 | 7266681 | 6687522 | 8520953 |
| Q0VBL3 | Rbm15 | RNA-binding protein 15 Mus musculus | 179.31 | 5.2 | 105722 | 4 | 4 | 1143650.88 | 1253638.5 | 1074182.25 | 1482568.12 | 1370062.12 | 1652140.75 | 1114756.75 | 988258.5 | 650327.94 | 1688307.5 | 745620.5 | 1226603.75 | 644566.81 | 366201.38 | 997105.62 |
| Q8JZX4 | Rbm17 | Splicing factor 45 Mus musculus | 142.07 | 8.15 | 45304 | 3 | 3 | 1852686.5 | 2012469.12 | 872493.62 | 2370725 | 1001566.62 | 1592328 | 2151658 | 1650437.75 | 1518296.25 | 2241120 | 1636705.5 | 1854599.25 | 1963344 | 1512040.38 | 2273276.5 |
| Q8R3C6 | Rbm19 | Probable RNA-binding protein 19 Mus musculus | 102.13 | 2.94 | 106083 | 2 | 2 | 769180.94 | 791832.655 | 761084.06 | 633465 | 1003600.62 | 323088.91 | 1381611.5 | 762661.56 | 600763.31 | 1383542.75 | 298855.38 | 592030.25 | 718205.56 | 633411.5 | 1129407.25 |
| Q4V7D7 | Rbm22 | Pre-mRNA-splicing factor RBM22 Rattus norvegicus | 79.83 | 4.52 | 46864 | 1 | 1 | 480776.72 | 1 | 393129.44 | 1 | 1 | 316757.66 | 192209.34 | 1 | 1 | 478856.88 | 174057.83 | 1 | 1 | 1 | 1 |
| B2RY56 | Rbm25 | RNA-binding protein 25 Mus musculus | 328.76 | 10.26 | 99552 | 6 | 6 | 3141130.75 | 2363907 | 3301986.25 | 2883145.25 | 3357609 | 3022378.25 | 3912238.25 | 3022294.5 | 3389697 | 2544197 | 3713508.25 | 3944888.25 | 2523660.75 | 2986907.5 | 4919040 |
| Q8CGC6 | Rbm28 | RNA-binding protein 28 Mus musculus | 440.11 | 13.87 | 84214 | 9 | 9 | 3444275 | 3734736.5 | 787139.31 | 3987158 | 445939.22 | 4349352 | 5835207 | 1057811.25 | 3911269 | 4912077 | 6249065.5 | 4920380.5 | 4835301.5 | 3772427 | 5794008 |
| O89086 | Rbm3 | RNA-binding protein 3 Mus musculus | 263.18 | 48.37 | 16605 | 4 | 4 | 16201317 | 14056728 | 17451029 | 17496437 | 18559707 | 13890065 | 13601522 | 13944243 | 11691299 | 13727713 | 18621125 | 17886511 | 16596929 | 15088548 | 14502953 |
| Q8CSL7 | Rbm34 | RNA-binding protein 34 Mus musculus | 36.19 | 2.67 | 41326 | 1 | 1 | 980492.62 | 577497 | 764433.94 | 691917.8125 | 445247.69 | 405755.06 | 438014.31 | 560746.88 | 612809.19 | 504331.36 | 463326.19 | 680196.94 | 422184.69 | 623050.38 | 558984.25 |
| Q8VH51 | Rbm39 | RNA-binding protein 39 Mus musculus | 850.65 | 34.91 | 59407 | 13 | 13 | 18583955 | 20376495 | 26137809 | 25612119 | 27527821 | 25573469 | 26615987 | 23718035 | 23904429 | 28816969 | 27248661 | 30683947 | 27805369 | 21589877 | 32234967 |
| Q91V81 | Rbm42 | RNA-binding protein 42 Mus musculus | 71.11 | 4.43 | 49875 | 2 | 2 | 1 | 1 | 360364.38 | 1 | 1 | 1 | 374205.16 | 326169.34 | 1 | 355660.69 | 203473.89 | 479164.47 | 1 | 366593.12 | 1 |
| Q91WT8 | Rbm47 | RNA-binding protein 47 Mus musculus | 43.03 | 3.39 | 64062 | 1 | 1 | 1 | 1 | 1 | 521404.94 | 318446.12 | 404823.5075 | 460456.84 | 391070.22 | 401344.56 | 366422.41 | 1 | 345397.97 | 1 | 416913.41 | 380784.75 |
| Q9CQT2 | Rbm7 | RNA-binding protein 7 Mus musculus | 51.72 | 6.79 | 30148 | 1 | 1 | 316903.72 | 1 | 1 | 1 | 1 | 1 | 1 | 1 | 1 | 1 | 223666.56 | 150289.25 | 314505.44 | 332629.22 | 1 |
| Q27W01 | Rbm8a | RNA-binding protein 8A Rattus norvegicus | 202.12 | 47.7 | 19889 | 4 | 4 | 7136055.5 | 6256920 | 6194536 | 5901224 | 6468162 | 7497923 | 5195970 | 7295695.5 | 9527012 | 5192030 | 7798030.5 | 7875885.5 | 5893921 | 6370579 | 6605081 |
| Q5PQP1 | Rbms1 | RNA-binding motif, single-stranded-interacting protein 1 Rattus norvegicus | 48.7 | 3.72 | 44068 | 1 | 1 | 1 | 2622672.75 | 2923624 | 1 | 2065368.88 | 2454787.5 | 2039895.5 | 2789830.75 | 3092668.75 | 2145763.5 | 1807975.88 | 1 | 2290104.25 | 1 | 1 |
| Q9WV02 | Rbmx | RNA-binding motif protein, X chromosome Mus musculus | 463.19 | 28.64 | 42301 | 0 | 11 | 14676479 | 14162876 | 11357965 | 15574562 | 17912887 | 10981879 | 14325791 | 9301412 | 7653447 | 22323999 | 14372431 | 13163348 | 17858497 | 8607787 | 19722267 |
| Q91VM5 | Rbmxl1 | RNA binding motif protein, X-linked-like-1 Mus musculus 2 | 628.15 | 36.08 | 42162 | 3 | 13 | 18824915 | 19790105 | 16963171 | 22123673 | 25133949 | 16680383 | 20710401 | 13902353 | 12680507 | 29398027 | 21769685 | 19425931 | 24579197 | 13342895 | 27712957 |
| P31266 | Rbpj | Recombining binding protein suppressor of hairless Mus musculus | 391.96 | 26.62 | 58537 | 8 | 8 | 25027323 | 23856861 | 19681669 | 24367335 | 24816885 | 23362747 | 18315733 | 20762575 | 25411561 | 19522855 | 25879023 | 18683823 | 23607915 | 25657393 | 20682863 |
| P62878 | Rbx1 | E3 ubiquitin-protein ligase RBX1 Mus musculus | 86.98 | 16.67 | 12274 | 1 | 1 | 4225062 | 4060234.75 | 3408895.25 | 3481492.75 | 4201550.5 | 3826355.25 | 3447901.25 | 3394183.5 | 3603653.25 | 3947607.5 | 3910858.25 | 2999090.5 | 3889049.25 | 3475926 | 2891245.5 |
| Q8VE37 | Rcc1 | Regulator of chromosome condensation Mus musculus | 336.85 | 21.62 | 44931 | 6 | 6 | 7333107 | 6421106 | 8114995.5 | 9350778 | 9880416 | 8505862 | 10267985 | 7451200 | 5936945 | 9177313 | 7980707 | 9072841 | 8215369 | 6739434 | 9064699 |
| Q8BK67 | Rcc2 | Protein RCC2 Mus musculus | 1140.43 | 57.31 | 55983 | 20 | 20 | 50144673 | 56313645 | 56713385 | 67145321 | 62236345 | 54125845 | 63677577 | 51396229 | 52525537 | 61529021 | 56150097 | 54932181 | 55577369 | 51634513 | 61196949 |
| Q9JJT0 | Rcl1 | RNA 3'-terminal phosphate cyclase-like protein Mus musculus | 110.28 | 6.17 | 40841 | 2 | 2 | 1951306.25 | 2081479 | 1325929.38 | 1148578.38 | 1117398.88 | 1923563.88 | 1585251.5 | 2281750.25 | 2620964.5 | 1518559.88 | 1952014.75 | 1855172.88 | 1858352 | 2148795 | 2077960.5 |
| Q8BP92 | Rcn2 | Reticulocalbin-2 Mus musculus | 534.81 | 36.56 | 37271 | 8 | 8 | 8223233.5 | 9635146 | 8256234.5 | 4061390 | 8449753 | 9847711 | 3066376.5 | 10115649 | 11026869 | 7073551.5 | 8089672 | 8689658 | 7080369.5 | 8707970 | 7249414.5 |
| Q3UZA1 | Rcsd1 | CapZ-interacting protein Mus musculus | 372.28 | 25.97 | 44114 | 7 | 7 | 22690795 | 6848788.5 | 5966656.5 | 6237873.5 | 8090971 | 6572137 | 6538259.5 | 5351002.5 | 7262668 | 6899219.5 | 7809546.5 | 5604206 | 7340904 | 6229603 | 5765836 |
| Q9QYF1 | Rdh11 | Retinol dehydrogenase 11 Mus musculus | 215.49 | 19.3 | 35148 | 5 | 5 | 6818532.5 | 5668518 | 6806729 | 6052273.5 | 5862307.5 | 5827885 | 9511445 | 5974274.5 | 5977569 | 8294025 | 7874501 | 6588527 | 6713272.5 | 6140294 | 6673557.5 |
| Q8CEE7 | Rdh13 | Retinol dehydrogenase 13 Mus musculus | 46.64 | 5.69 | 36464 | 1 | 1 | 229952.3425 | 182689.22 | 281971.94 | 185605.27 | 269542.94 | 207024.06 | 131408.25 | 117791.98 | 160295.2 | 212378.61 | 215455.91 | 173741.12 | 222101.95 | 194574.27 | 365609.78 |
| P26043 | Rdx | Radixin Mus musculus | 1301.23 | 39.97 | 68543 | 10 | 24 | 451788545 | 429860481 | 379326657 | 426065889 | 406720193 | 426244961 | 452088705 | 445550593 | 398309697 | 425842273 | 359942529 | 390315617 | 390523169 | 449967841 | 353532449 |
| Q9Z129 | Recql | ATP-dependent DNA helicase Q1 Mus musculus | 37.86 | 1.7 | 72484 | 1 | 1 | 649946.69 | 684190.44 | 933678 | 810436.69 | 956424.56 | 860565.69 | 1028686.06 | 679077.69 | 990592.56 | 808598.62 | 1074095.5 | 797234.19 | 990740.31 | 890674.94 | 803021.69 |
| B2RZ37 | Reep5 | Receptor expression-enhancing protein 5 Rattus norvegicus | 61.03 | 5.29 | 21431 | 1 | 1 | 7432805 | 3583975.75 | 3614411 | 3200738.75 | 4175538.75 | 5864029 | 3971444.75 | 4307912 | 4131722 | 3879152.5 | 4818361 | 4067905 | 4754617 | 4561645 | 2894322.75 |

| | | | | | | | | | | | | | | | | | | | | | | |
|---|---|---|---|---|---|---|---|---|---|---|---|---|---|---|---|---|---|---|---|---|---|---|
| P15307 | Rel | Proto-oncogene c-Rel Mus musculus | 41.15 | 1.87 | 64960 | 1 | 1 | 1 | 490613.88 | 1 | 1 | 1 | 1 | 1 | 1 | 167662.66 | 1 | 1 | 1 | 243901.69 | 631775.06 | 1 |
| Q04207 | Rela | Transcription factor p65 Mus musculus | 82.81 | 8.93 | 60212 | 2 | 2 | 211088.44 | 1 | 145839.23 | 194205.16 | 1 | 1 | 237981.23 | 1 | 1 | 1 | 1 | 1 | 236170.19 | 1 | 1 |
| Q148V7 | Relch | RAB11-binding protein RELCH Mus musculus | 98.82 | 1.97 | 134587 | 2 | 2 | 423175.78 | 438693.66 | 405497.84 | 502120.75 | 370819.09 | 449272.81 | 333030.41 | 402050.06 | 753523.56 | 1 | 411732.06 | 595088.19 | 431041.28 | 454126.12 | 548173.12 |
| P82343 | Renbp | N-acylglucosamine 2-epimerase Mus musculus | 372.96 | 21.16 | 49771 | 9 | 9 | 27482973 | 22888361 | 22671903 | 21481823 | 21501491 | 21375173 | 19257167 | 21012849 | 24845457 | 21451099 | 23156149 | 19332399 | 20099689 | 23410829 | 20957783 |
| Q5FVM3 | Retreg1 | Reticulophagy regulator 1 Rattus norvegicus | 70.42 | 2.5 | 52782 | 1 | 1 | 1 | 635444.75 | 1 | 477877.72 | 390176.22 | 210491.86 | 367324.25 | 1 | 912323.5 | 1 | 605764.38 | 443446.28 | 517017.22 | 540874.69 | 1 |
| Q3MHU5 | Retreg2 | Reticulophagy regulator 2 Rattus norvegicus | 58.32 | 1.48 | 57659 | 1 | 1 | 1 | 1 | 999634.12 | 1 | 862420.62 | 684649.19 | 1015803.81 | 838793.31 | 853547.44 | 764859.56 | 1 | 638174 | 1014498.06 | 1 | 1 |
| Q64FW2 | Retsat | All-trans-retinol 13,14-reductase Mus musculus | 284.96 | 14.29 | 67334 | 5 | 5 | 2062565.88 | 16080405 | 1844958.62 | 1249876.62 | 25397993 | 12776418 | 9363789 | 11372773 | 1056313.88 | 1201594.75 | 13935841 | 848959.25 | 511906.28 | 10714771 | 9660419 |
| Q5U1X1 | Rexo2 | Oligoribonuclease, mitochondrial Rattus norvegicus | 127.73 | 10.97 | 26752 | 2 | 2 | 4250126 | 4397954 | 4659792 | 4377616 | 4009168.25 | 4392658.5 | 4159544.25 | 4074488 | 4911412 | 5119558 | 5213442 | 4237713.5 | 4566582 | 4337712 | 4034555.75 |
| P35601 | Rfc1 | Replication factor C subunit 1 Mus musculus | 223.36 | 5.48 | 125985 | 4 | 4 | 1754153.25 | 1955838.12 | 1638564.38 | 2245255 | 1607385 | 1727797.75 | 1766827.12 | 1911690.12 | 1281196 | 2143486.5 | 1427513.62 | 2209484 | 1761824.12 | 625199.62 | 2275399.25 |
| Q9WUK4 | Rfc2 | Replication factor C subunit 2 Mus musculus | 214.45 | 16.91 | 38725 | 4 | 4 | 1973131.25 | 1699930.5 | 2027881.75 | 4414021 | 4219553 | 4186890.75 | 3861770 | 3896181.5 | 1701488.25 | 4637150.5 | 4281910 | 3819789.75 | 4538027 | 3166624 | 3969675 |
| Q8R323 | Rfc3 | Replication factor C subunit 3 Mus musculus | 208.7 | 11.24 | 40526 | 4 | 4 | 3114739.25 | 2834600.5 | 2661804 | 1520486.12 | 3500486 | 4026629 | 1649905.62 | 4811175 | 3375425.5 | 1723176.38 | 4520938.5 | 4174300 | 2019658 | 4784001 | 4158869.5 |
| Q99J62 | Rfc4 | Replication factor C subunit 4 Mus musculus | 248.48 | 19.51 | 39867 | 5 | 5 | 6632325 | 10782153 | 7629577 | 9015264 | 9698691 | 8308206.5 | 8393353 | 8876423 | 9170841 | 8581029 | 9629446 | 7065782 | 8245974 | 8740152 | 8267560 |
| Q9D0F6 | Rfc5 | Replication factor C subunit 5 Mus musculus | 257.05 | 27.14 | 38096 | 6 | 6 | 3489320.75 | 3808726.5 | 3205226.5 | 3610095 | 3384961.5 | 4443581.5 | 4075791.75 | 3680420 | 4294434 | 3690032.5 | 3499960.5 | 3566589.5 | 3502885.25 | 3953832.25 | 3816098.5 |
| Q8CFV9 | Rfk | Riboflavin kinase Mus musculus | 168.51 | 13.55 | 17437 | 2 | 2 | 1046277.62 | 1499406.25 | 1439349.75 | 909639.38 | 1100302.12 | 1045189.94 | 1097617.5 | 900917.56 | 1166422.62 | 975469.25 | 2316704 | 1501384.88 | 1496463.5 | 1589010.5 | 1068606.5 |
| Q6A0D4 | Rftn1 | Raftlin Mus musculus | 128.03 | 8.12 | 61537 | 3 | 3 | 2560086.25 | 167247.33 | 1422655.25 | 2004712.62 | 2072418 | 1552348 | 2031222.88 | 397576.44 | 1554954 | 1740403.25 | 2531314.75 | 1667287.12 | 2384392 | 1687138.25 | 1973929 |
| Q9CQE5 | Rgs10 | Regulator of G-protein signaling 10 Mus musculus | 92.63 | 9.39 | 21151 | 1 | 1 | 1260703.12 | 1249538.62 | 880824.56 | 927223.12 | 954264.5 | 867003.25 | 745056.81 | 1024196.69 | 994259.12 | 806851.94 | 700866.62 | 1035998.19 | 950607.56 | 930174.31 | 775032.12 |
| O70521 | Rgs19 | Regulator of G-protein signaling 19 Rattus norvegicus | 39.26 | 6.02 | 24738 | 1 | 1 | 1425710 | 1221182 | 1282710.75 | 1493605.38 | 1355802.033 | 375430.56 | 1058403.12 | 1323065.62 | 1498147.88 | 1294972.88 | 1168665.38 | 1102665.5 | 500619.91 | 1107105.12 | 716740.19 |
| Q4V8F3 | Rhbdd1 | Rhomboid-related protein 4 Rattus norvegicus 2 | 55.79 | 4.43 | 35879 | 1 | 1 | 1 | 1 | 1 | 446310.66 | 277795.25 | 406301.25 | 406245.5775 | 319164.12 | 332071.88 | 567445.06 | 1 | 265738.19 | 1 | 633639.12 | 408249.84 |
| Q921J2 | Rheb | GTP-binding protein Rheb Mus musculus | 214.44 | 28.26 | 20451 | 5 | 5 | 3202839 | 3763590 | 2978251.25 | 2856487.5 | 1729558.62 | 2353985.5 | 3469082 | 3119570 | 3682997.5 | 1324314.38 | 2235366.5 | 2953689.5 | 3189473.75 | 3435904.5 | 2765222 |
| P61589 | Rhoa | Transforming protein RhoA Rattus norvegicus | 759.83 | 65.8 | 21782 | 3 | 12 | 78086985 | 48889153 | 73282449 | 55225093 | 51837897 | 57090557 | 66209089 | 87462769 | 84399873 | 83363089 | 61011009 | 59348113 | 52820857 | 50665997 | 86960377 |
| P62746 | Rhob | Rho-related GTP-binding protein RhoB Mus musculus | 196.09 | 25.51 | 22123 | 1 | 3 | 30660373 | 5411296.5 | 30118673 | 6870403 | 5860114.5 | 6898794.5 | 6109198 | 38755713 | 42755717 | 31485449 | 5407387.5 | 10696951 | 4992457 | 7209405.5 | 29284121 |
| Q62159 | Rhoc | Rho-related GTP-binding protein RhoC Mus musculus | 747.29 | 65.8 | 22006 | 3 | 12 | 72336065 | 43518089 | 68619497 | 49710129 | 46555773 | 52899601 | 57888069 | 84052865 | 82017537 | 77854025 | 55186869 | 54676885 | 46732309 | 46798201 | 81588713 |
| P84096 | Rhog | Rho-related GTP-binding protein RhoG Mus musculus | 660.33 | 53.93 | 21309 | 8 | 9 | 17318869 | 18855237 | 15860312 | 18122857 | 17035473 | 15680418 | 19320863 | 16437198 | 15090949 | 18750607 | 13213663 | 15438392 | 18235173 | 16008265 | 15766782 |
| Q8BG51 | Rhot1 | Mitochondrial Rho GTPase 1 Mus musculus | 186.76 | 6.02 | 72242 | 3 | 4 | 334383.47 | 1 | 295852.75 | 214383.38 | 1 | 1 | 1 | 1 | 1 | 347244.47 | 278279.97 | 1 | 1 | 1 | 1 |
| Q7TSA0 | Rhot2 | Mitochondrial Rho GTPase 2 Rattus norvegicus | 193.54 | 6.27 | 69153 | 2 | 3 | 501147.69 | 595937.88 | 646770.06 | 642362.8775 | 825595.88 | 708936.81 | 649272.12 | 558124.88 | 552189.75 | 676384.25 | 727759.31 | 650175 | 1255702.38 | 692343.12 | 704415.38 |
| P52760 | Rida | 2-iminobutanoate/2-iminopropanoate deaminase Mus musculus | 102.6 | 17.78 | 14255 | 2 | 2 | 1087577.62 | 1302952.12 | 983554.19 | 1607621.75 | 397875.16 | 1122672.25 | 1224834.38 | 837864.06 | 418449.12 | 1164916.12 | 1199731.25 | 352588.66 | 1008776.25 | 509523.34 | 1183579.75 |
| Q6PR54 | Rif1 | Telomere-associated protein RIF1 Mus musculus | 189.22 | 2.32 | 266228 | 3 | 3 | 1624920.38 | 2046007.75 | 1606687.25 | 2072905.25 | 1487789.88 | 1001058 | 1176537.25 | 1620655 | 144852.36 | 1636714.5 | 1451509.5 | 1388940 | 1297931.25 | 1229656.5 | 1019543 |
| Q9QZL0 | Ripk3 | Receptor-interacting serine/threonine-protein kinase 3 Mus musculus | 404.16 | 23.25 | 53322 | 7 | 7 | 13043554 | 12830318 | 15241674 | 3231674.75 | 4065368.25 | 16699065 | 10890657 | 10462004 | 10355861 | 12224462 | 17378813 | 13891309 | 14380917 | 11366492 | 15962486 |
| P13832 | Rlc-a | Myosin regulatory light chain RLC-A Rattus norvegicus | 681.86 | 79.65 | 19895 | 4 | 11 | 52345889 | 55833101 | 55399397 | 55234433 | 52837229 | 53142969 | 44929145 | 61821953 | 64594777 | 43356293 | 65792821 | 60819301 | 47637445 | 62375613 | 53993949 |
| Q9CWY8 | Rnaseh2a | Ribonuclease H2 subunit A Mus musculus | 54.23 | 5.32 | 33513 | 1 | 1 | 453792.31 | 325170.91 | 500514.62 | 436523.62 | 482824.31 | 642599.88 | 724128.38 | 392136.34 | 305759.91 | 581576.62 | 568938.06 | 526288.62 | 485672.84 | 512037.69 | 940967.25 |
| Q5DTM8 | Rnf20 | E3 ubiquitin-protein ligase BRE1A Mus musculus | 98.04 | 2.77 | 113520 | 2 | 2 | 1 | 161327.36 | 247613.83 | 1 | 1 | 134158.55 | 1 | 1 | 180217.69 | 1 | 197194.86 | 158214.34 | 164946.7125 | 161657.27 | 142720.38 |
| E9Q555 | Rnf213 | E3 ubiquitin-protein ligase RNF213 Mus musculus | 2031.58 | 11.9 | 584784 | 43 | 43 | 45362837 | 41682497 | 32155981 | 35600493 | 38076313 | 42968309 | 36043049 | 40133861 | 39902261 | 31733005 | 39891225 | 34840109 | 41812917 | 42241293 | 34479877 |
| Q91VI7 | Rnh1 | Ribonuclease inhibitor Mus musculus | 2471.85 | 90.79 | 49816 | 29 | 29 | 331530049 | 365746049 | 323439425 | 375143137 | 344204865 | 323917121 | 317745985 | 309386913 | 333260769 | 327228065 | 362858209 | 304973185 | 341939009 | 350689345 | 344746913 |
| Q5U2U7 | Rnmt | mRNA cap guanine-N7 methyltransferase Rattus norvegicus | 147.61 | 8.03 | 52817 | 3 | 3 | 3920096.75 | 3663730.25 | 3442175.5 | 3841588.75 | 4448384.5 | 4452551.5 | 4622124 | 3840538.5 | 4473065 | 4046351.75 | 4370883.5 | 4252909 | 3824983 | 3161144.5 | 4462742 |
| Q8VCT3 | Rnpep | Aminopeptidase B Mus musculus | 1563.82 | 54.46 | 72416 | 26 | 26 | 62735149 | 67485457 | 71716025 | 64039325 | 62866457 | 57524073 | 57695817 | 59065049 | 69829625 | 55437421 | 64643705 | 66692065 | 61147713 | 64651405 | 67744161 |
| P70336 | Rock2 | Rho-associated protein kinase 2 Mus musculus | 116.68 | 2.67 | 160586 | 3 | 3 | 918762.94 | 1955630 | 1393872.88 | 1431697.12 | 2111055.5 | 1382735.75 | 1887184.5 | 1156124.88 | 867515.31 | 1360896 | 1051650 | 1762333.25 | 1892385.25 | 950562.94 | 572681.56 |
| P60603 | Romo1 | Reactive oxygen species modulator 1 Mus musculus | 113.24 | 51.9 | 8183 | 2 | 2 | 507002.25 | 705462.4375 | 827640.75 | 435317 | 1051889.75 | 815735.31 | 1781674.62 | 299183.12 | 1065605.75 | 1291565.5 | 754983.5 | 801054.56 | 1060207 | 1059147.88 | 915947.12 |
| Q63132 | Ros1 | Proto-oncogene tyrosine-protein kinase ROS Rattus norvegicus | 71.5 | 0.64 | 261656 | 1 | 2 | 2297278.75 | 1518774.38 | 1538482.75 | 466991.91 | 1850511.62 | 1893897.25 | 1350944.5 | 1429691.25 | 1331146.75 | 1438125.25 | 1878847.88 | 1365556.75 | 1782837.62 | 1633359.88 | 1488303.5 |
| Q9EPK2 | Rp2 | Protein XRP2 Mus musculus | 108.05 | 5.19 | 39377 | 2 | 2 | 1473764.12 | 1049215.62 | 1728861.25 | 1075393 | 1417357.62 | 1856085.38 | 1456314.88 | 1783323.75 | 2431413 | 1501541.5 | 1631701 | 2039006.38 | 1195231.62 | 1918258.25 | 1496012.5 |
| Q8VEE4 | Rpa1 | Replication protein A 70 kDa DNA-binding subunit Mus musculus | 839.18 | 26.16 | 69037 | 15 | 15 | 17327145 | 21230773 | 24112441 | 18150669 | 19528737 | 19884861 | 22522549 | 18085569 | 22019681 | 20047625 | 17289625 | 24726707 | 18546091 | 22156217 | 27076595 |
| Q62193 | Rpa2 | Replication protein A 32 kDa subunit Mus musculus | 182.28 | 21.11 | 29718 | 4 | 4 | 8180673.5 | 5909737 | 7714308 | 10611604 | 9788269 | 8675204 | 5567929 | 4898674 | 8233217.5 | 9580599 | 9486183 | 8335501 | 9156307 | 9212617 | 4686138 |
| Q9CQ71 | Rpa3 | Replication protein A 14 kDa subunit Mus musculus | 118.83 | 32.23 | 13584 | 3 | 3 | 2110414.75 | 2540167 | 2299456 | 2579842.75 | 2428874.5 | 2836435.75 | 2218502 | 3013007.5 | 2446049.5 | 2344943.5 | 2953038.5 | 2876081.75 | 2697634.5 | 2461365.25 | 2223710.5 |
| Q80TE0 | Rpap1 | RNA polymerase II-associated protein 1 Mus musculus | 44.06 | 0.92 | 155270 | 1 | 1 | 394524.53 | 429404.41 | 408864.81 | 426645.22 | 414859.7425 | 302784.22 | 426448.12 | 452765.31 | 377381.41 | 389844.765 | 347041.75 | 273627.06 | 403163.72 | 281051.38 | 326220.9775 |
| Q9D706 | Rpap3 | RNA polymerase II-associated protein 3 Mus musculus | 165.13 | 6.82 | 74096 | 3 | 3 | 1504026.88 | 1642477.12 | 1195072.5 | 1533197.62 | 233922.06 | 1032450.06 | 1259067.25 | 1019960.31 | 1234934.5 | 1136603.03 | 1004412 | 1140427.88 | 1181591.62 | 981534.62 | 1247824.75 |
| Q8VEE0 | Rpe | Ribulose-phosphate 3-epimerase Mus musculus | 281.86 | 36.84 | 24945 | 4 | 4 | 2575407.75 | 2132401 | 2770585 | 2216684 | 2219400 | 2113905.5 | 1052885.12 | 2521034.75 | 1488886.88 | 2219893.25 | 2394489.25 | 2138468.25 | 2346291 | 2248811.5 | 2136879 |
| Q9JJ80 | Rpf2 | Ribosome production factor 2 homolog Mus musculus | 333.42 | 27.45 | 35364 | 6 | 6 | 1278550 | 1473422 | 1305544.5 | 2019469.38 | 2440933.5 | 1119897 | 1536298.25 | 1132022 | 403525.84 | 1684512.5 | 1177737.88 | 918642.62 | 1304206.75 | 424626.34 | 1059695.25 |
| P47968 | Rpia | Ribose-5-phosphate isomerase Mus musculus | 282.23 | 28.38 | 32451 | 7 | 7 | 9771853 | 5273146.5 | 4866029.5 | 9437076 | 9593231 | 8881825 | 7279951.5 | 8193318 | 5066853.5 | 8017896.5 | 7878348 | 5999900 | 8385844.5 | 10257803 | 5696205 |
| Q6PDV7 | Rpl10 | 60S ribosomal protein L10 Rattus norvegicus | 504.9 | 38.79 | 24604 | 8 | 8 | 19296765 | 18216813 | 22525409 | 20898781 | 20913821 | 21972029 | 32218533 | 19237945 | 17132767 | 24961167 | 26646309 | 24515643 | 22567541 | 19389811 | 30953643 |
| P62907 | Rpl10a | 60S ribosomal protein L10a Rattus norvegicus | 740.84 | 58.53 | 24831 | 13 | 13 | 14175484 | 14322817 | 21082419 | 16076777 | 13926860 | 15101351 | 40840713 | 12160530 | 11819636 | 24396513 | 17271203 | 20355781 | 16585845 | 13579405 | 29260435 |
| P62914 | Rpl11 | 60S ribosomal protein L11 Rattus norvegicus | 577.37 | 62.92 | 20252 | 11 | 11 | 32958047 | 26836899 | 41043341 | 41170781 | 33928033 | 32391117 | 62819001 | 28624455 | 26568899 | 48695645 | 45120653 | 40010161 | 35455953 | 27708787 | 55462493 |
| P35979 | Rpl12 | 60S ribosomal protein L12 Mus musculus | 490.93 | 51.52 | 17805 | 7 | 7 | 42948813 | 36512005 | 40066069 | 40067593 | 38795725 | 37422149 | 52392757 | 35904721 | 38438297 | 34963069 | 41662925 | 35419969 | 33502421 | 36809837 | 37795141 |
| P41123 | Rpl13 | 60S ribosomal protein L13 Rattus norvegicus | 265.56 | 25.59 | 24309 | 5 | 5 | 9368862 | 6131949 | 9752551 | 10905688 | 8065040 | 7144798 | 20510413 | 3934294.75 | 4948538 | 14529443 | 11164476 | 12190231 | 8515282 | 6595691 | 17212695 |
| P19253 | Rpl13a | 60S ribosomal protein L13a Mus musculus | 449.35 | 33.99 | 23464 | 8 | 9 | 19012237 | 11100493 | 13220825 | 13650727 | 4404996 | 9509334 | 20575023 | 11688402 | 8275985.5 | 14297702 | 10764090 | 14038974 | 11222795 | 9265260 | 12414583 |
| Q9CR57 | Rpl14 | 60S ribosomal protein L14 Mus musculus | 266.01 | 25.81 | 23564 | 5 | 5 | 48757397 | 41464161 | 49147209 | 50989849 | 40586785 | 38935113 | 58988657 | 42632645 | 38162553 | 44745221 | 43687833 | 48242529 | 44680781 | 45807781 | 45207057 |
| P61314 | Rpl15 | 60S ribosomal protein L15 Rattus norvegicus | 608.47 | 45.1 | 24146 | 11 | 11 | 18648701 | 13333976 | 11521345 | 9381727 | 11246411 | 11552257 | 24372187 | 12413661 | 16842217 | 11432087 | 12216952 | 16177580 | 11613767 | 12333368 | 27749391 |
| P24049 | Rpl17 | 60S ribosomal protein L17 Rattus norvegicus | 292.67 | 30.98 | 21397 | 5 | 5 | 17367001 | 13188777 | 15337442 | 14684893 | 12363700 | 13778587 | 17930597 | 11656540 | 12987141 | 14324509 | 13085410 | 13766002 | 12135532 | 12418609 | 18667537 |
| P35980 | Rpl18 | 60S ribosomal protein L18 Mus musculus | 527.2 | 49.47 | 21645 | 10 | 10 | 41226201 | 23076019 | 20713081 | 23666485 | 27125053 | 22478797 | 31737905 | 19234575 | 91232753 | 29210037 | 21710493 | 51869121 | 23770759 | 22388305 | 25645271 |
| P62717 | Rpl18a | 60S ribosomal protein L18a Mus musculus | 352.19 | 33.52 | 20732 | 8 | 8 | 27571991 | 17590765 | 18432631 | 25090469 | 25397873 | 24264855 | 30609589 | 19750757 | 17651859 | 22963537 | 23534117 | 24765649 | 16859349 | 22577725 | 17642381 |
| P84099 | Rpl19 | 60S ribosomal protein L19 Mus musculus | 308.5 | 18.37 | 23466 | 5 | 5 | 2841524.75 | 3748496.25 | 4027587.5 | 3882654.5 | 4441050 | 4021494.5 | 6342369 | 3186891.25 | 3519376 | 4765551 | 3947168.5 | 3940416.75 | 4561912.5 | 3492383.5 | 7387925 |
| O09167 | Rpl21 | 60S ribosomal protein L21 Mus musculus | 244.37 | 32.5 | 18562 | 5 | 5 | 23035485 | 21861553 | 28818509 | 33698633 | 30063827 | 25679281 | 34221825 | 21005701 | 22506087 | 33525783 | 35825073 | 28758655 | 30133053 | 21659331 | 34602029 |
| P67984 | Rpl22 | 60S ribosomal protein L22 Mus musculus | 387.03 | 56.25 | 14759 | 7 | 7 | 35659789 | 27045523 | 32735147 | 24856975 | 31397327 | 32970473 | 53560437 | 31822643 | 31620405 | 43437709 | 43142553 | 42582445 | 34805049 | 29670977 | 53513533 |
| Q9D7S7 | Rpl22l1 | 60S ribosomal protein L22-like 1 Mus musculus | 252.54 | 45.08 | 14467 | 4 | 4 | 8332675.5 | 4641697 | 4296884 | 4652856 | 5900927 | 5429573 | 8922058 | 5315496.5 | 4570625.5 | 6852798 | 5925064.5 | 7226938 | 5896403 | 5109605.5 | 8961221 |
| P62830 | Rpl23 | 60S ribosomal protein L23 Mus musculus | 397.75 | 62.86 | 14865 | 8 | 8 | 78422417 | 81408617 | 92665329 | 89340329 | 84629801 | 78916377 | 110813473 | 72096737 | 72524937 | 98905969 | 86677041 | 91332889 | 83215601 | 70138841 | 97485537 |
| P62751 | Rpl23a | 60S ribosomal protein L23a Mus musculus | 345.1 | 50.64 | 17695 | 8 | 8 | 12156759 | 9817307 | 7802661.5 | 17052715 | 8816283 | 8550817 | 10360273 | 5741490 | 6209967 | 14357269 | 7412881.5 | 10270523 | 10305301 | 7659735 | 7312365 |
| P83732 | Rpl24 | 60S ribosomal protein L24 Rattus norvegicus | 279.62 | 32.48 | 17779 | 5 | 5 | 7935524.5 | 6400114 | 8622589 | 8188637.5 | 6570155.5 | 6320483.5 | 15558642 | 5839484 | 5557059.5 | 12029572 | 9597618 | 12638157 | 8093770 | 5680647 | 14804111 |
| P61255 | Rpl26 | 60S ribosomal protein L26 Mus musculus | 467.96 | 41.38 | 17258 | 9 | 9 | 16771569 | 15726351 | 22666187 | 5770501 | 6629578 | 20333821 | 13983333 | 21262351 | 5617879 | 18937537 | 26155813 | 25552627 | 17811199 | 19100779 | 14184162 |
| P61354 | Rpl27 | 60S ribosomal protein L27 Rattus norvegicus | 213.29 | 36.03 | 15798 | 4 | 4 | 1205200.12 | 1757168.703 | 2642859.5 | 2145694.5 | 1034920.69 | 1576396.5 | 5963299 | 1226097.88 | 676850.38 | 3502949 | 2697874.25 | 3534017 | 2454153 | 1028978.88 | 9013954 |
| P14115 | Rpl27a | 60S ribosomal protein L27a Mus musculus | 371 | 33.11 | 16605 | 7 | 7 | 40037089 | 42905105 | 45102537 | 44705289 | 43530089 | 39790725 | 44296481 | 40801053 | 41524761 | 42693853 | 27935301 | 38935337 | 39817385 | 39647617 | 43217513 |
| P41105 | Rpl28 | 60S ribosomal protein L28 Mus musculus | 337.58 | 32.85 | 15733 | 6 | 6 | 7195704.5 | 6425701 | 9476703 | 11688524 | 6312725 | 9828943 | 17083885 | 7686481 | 5394699.5 | 13781724 | 11810211 | 11905365 | 7830762 | 2551491.25 | 13473737 |
| P47915 | Rpl29 | 60S ribosomal protein L29 Mus musculus | 86.32 | 11.25 | 17587 | 2 | 2 | 21712929 | 26435781 | 36960413 | 33214665 | 36677345 | 28011065 | 37558697 | 22528227 | 23595109 | 37783049 | 38789809 | 31339737 | 36228953 | 21367175 | 35594549 |
| P27659 | Rpl3 | 60S ribosomal protein L3 Mus musculus | 1434.59 | 52.85 | 46110 | 22 | 24 | 23059385 | 17892201 | 22967791 | 31215469 | 29197835 | 17477883 | 39619201 | 19949205 | 10257610 | 33484371 | 27584969 | 28585413 | 18226883 | 15793561 | 41797073 |
| P62889 | Rpl30 | 60S ribosomal protein L30 Mus musculus | 490.07 | 59.13 | 12784 | 7 | 7 | 33088765 | 25803993 | 29098033 | 50881621 | 34721301 | 38623125 | 46873493 | 40704961 | 26187595 | 46584225 | 46520137 | 35321081 | 33563161 | 39302953 | 47214309 |
| P62900 | Rpl31 | 60S ribosomal protein L31 Mus musculus | 186.39 | 30.4 | 14463 | 4 | 4 | 4916891 | 3206677.5 | 4019639.5 | 4328282 | 3507651.75 | 6910504 | 7026098.5 | 3283880.75 | 3131420 | 4980630 | 9654187 | 6896194.5 | 5224235 | 3718093.5 | 4511923.5 |
| P62911 | Rpl32 | 60S ribosomal protein L32 Mus musculus | 393.86 | 40 | 15860 | 7 | 7 | 5341341.5 | 3066487.5 | 6492211.5 | 7782269 | 5765616 | 2751628.25 | 12862469 | 2907666.75 | 1679074.12 | 9639253 | 6046159.5 | 4302067.5 | 3028866 | 4773719 | 9234658 |
| P11250 | Rpl34 | 60S ribosomal protein L34 Rattus norvegicus | 82.51 | 12.82 | 13507 | 2 | 2 | 575018.31 | 1489885.5 | 1470175.75 | 2556895.25 | 1522993.703 | 1362897.25 | 4728556.5 | 1509290.25 | 1152320.12 | 2858015.25 | 1721481.12 | 2187604.5 | 1652867.75 | 1613878 | 2311341.5 |
| P17078 | Rpl35 | 60S ribosomal protein L35 Rattus norvegicus | 190.89 | 21.95 | 14553 | 4 | 4 | 1 | 646843.44 | 667667.94 | 1050045.75 | 1 | 1 | 496964.25 | 1 | 558708.88 | 1 | 445393.47 | 310857.94 | 583513.94 | 981563.56 | 1855557.38 |
| O55142 | Rpl35a | 60S ribosomal protein L35a Mus musculus | 114.96 | 20.91 | 12554 | 3 | 3 | 8128167.5 | 4272391 | 4715124.5 | 7664477 | 4816295.5 | 5077534 | 6694694 | 5955059 | 4644370 | 6461525.5 | 5144044 | 4639142.5 | 5901367 | 5897883.5 | 5334414.5 |
| P39032 | Rpl36 | 60S ribosomal protein L36 Rattus norvegicus | 105.28 | 19.05 | 12268 | 1 | 2 | 1348309.25 | 1191118.5 | 2931916.5 | 1890920.88 | 1829570 | 2083938.75 | 6313029.5 | 1461888.5 | 1275342.75 | 4085827.25 | 3511817.75 | 4115853.5 | 2312828.25 | 3813123 | 5311992.5 |
| P47964 | Rpl36 | 60S ribosomal protein L36 Mus musculus | 82.39 | 18.1 | 12216 | 1 | 2 | 4759367 | 2821063 | 5226443 | 4375304.5 | 4081034.75 | 4128766.5 | 10414738 | 3131678.5 | 2985654.5 | 7677518 | 6348020 | 7418321 | 4509123 | 1729251.38 | 8549767 |
| P61514 | Rpl37a | 60S ribosomal protein L37a Mus musculus | 169.49 | 41.3 | 10275 | 3 | 3 | 632743.88 | 1557243.75 | 1481535.38 | 558657.25 | 1394191.38 | 1733586.88 | 2303326 | 1922794.47 | 1738659.75 | 1915605.25 | 1619224.38 | 1998351.62 | 1404028 | 603507.44 | 3049755 |
| P62892 | Rpl39 | 60S ribosomal protein L39 Mus musculus | 35.58 | 19.61 | 6407 | 1 | 1 | 2724630.5 | 2267559 | 2507127.75 | 2371130.75 | 2485049.75 | 2759560.5 | 4202510 | 1712064.88 | 1793877.88 | 4094096.25 | 4266793.5 | 3344361.25 | 3003319.5 | 1495774.62 | 5443239 |
| Q9D8E6 | Rpl4 | 60S ribosomal protein L4 Mus musculus | 1226.3 | 46.54 | 47154 | 20 | 20 | 68622793 | 50576013 | 61833941 | 60178089 | 56715809 | 59906541 | 88179833 | 53309465 | 50483829 | 74490841 | 63058797 | 69573545 | 62210837 | 60132545 | 81110209 |
| P47962 | Rpl5 | 60S ribosomal protein L5 Mus musculus | 654.47 | 50.51 | 34401 | 12 | 12 | 58611145 | 60605537 | 58989681 | 61306309 | 64149569 | 57577105 | 57768605 | 52878429 | 61290753 | 66535249 | 67167281 | 61491397 | 62066365 | 57926233 | 66994089 |
| P47911 | Rpl6 | 60S ribosomal protein L6 Mus musculus | 687.47 | 44.59 | 33510 | 14 | 14 | 26706893 | 18230993 | 25584587 | 25728261 | 23738183 | 20898361 | 34123193 | 15081187 | 17518947 | 28551437 | 20210969 | 27780793 | 24569283 | 19577685 | 25847939 |
| P14148 | Rpl7 | 60S ribosomal protein L7 Mus musculus | 1284.82 | 58.15 | 31420 | 23 | 25 | 82860465 | 55623253 | 44417761 | 54578425 | 63209765 | 61654605 | 105821257 | 58171325 | 50201937 | 72673937 | 46812161 | 69125457 | 61241513 | 52836777 | 82603553 |
| P12970 | Rpl7a | 60S ribosomal protein L7a Mus musculus | 828.14 | 49.62 | 29977 | 15 | 15 | 42082725 | 27784061 | 32498789 | 45993913 | 39162697 | 31883767 | 53472793 | 28622759 | 22725499 | 43177393 | 35466385 | 36808793 | 30476059 | 28264605 | 38219429 |
| Q9D8M4 | Rpl7l1 | 60S ribosomal protein L7-like 1 Mus musculus | 317.61 | 41.06 | 28544 | 8 | 8 | 1269463.12 | 1459463.75 | 1023326.69 | 886850 | 715230.25 | 777957.56 | 1052258.25 | 779273.38 | 845428.94 | 774166.62 | 791402.12 | 1112552.75 | 689357 | 735516.19 | 732823.62 |
| P62918 | Rpl8 | 60S ribosomal protein L8 Mus musculus | 396.23 | 21.79 | 28025 | 7 | 7 | 7948369 | 6989144 | 12437464 | 14813648 | 12839873 | 11135084 | 26963695 | 10865650 | 4508191 | 18474297 | 16019143 | 14478083 | 8789271 | 8642984 | 21485319 |
| P51410 | Rpl9 | 60S ribosomal protein L9 Mus musculus | 1059.24 | 73.44 | 21881 | 15 | 15 | 46457813 | 29655009 | 42141465 | 32684757 | 33831709 | 35156725 | 55821573 | 29382697 | 31054791 | 39880397 | 39746805 | 41618837 | 35398657 | 29279941 | 58422621 |
| P14869 | Rplp0 | 60S acidic ribosomal protein P0 Mus musculus | 871.61 | 54.57 | 34216 | 13 | 13 | 218328369 | 246103585 | 248938705 | 213388145 | 213936561 | 238142145 | 213199537 | 246778913 | 262035329 | 207752385 | 214872289 | 235594433 | 223709553 | 239935905 | 198255665 |
| P47955 | Rplp1 | 60S acidic ribosomal protein P1 Mus musculus | 340.33 | 80.7 | 11475 | 5 | 5 | 82563521 | 83746257 | 79999273 | 84038497 | 79610681 | 93513113 | 67857705 | 97977641 | 86858305 | 75537401 | 73590609 | 78058921 | 79294353 | 93381305 | 72483689 |
| P99027 | Rplp2 | 60S acidic ribosomal protein P2 Mus musculus | 500.73 | 71.3 | 11651 | 6 | 6 | 5981313 | 4798400 | 7109557.5 | 4406071 | 4282963 | 6010403.5 | 5264679 | 6556234 | 5273114.5 | 4716227.5 | 6821492 | 5594071 | 4110059 | 6749691 | 5210276 |
| Q91YQ5 | Rpn1 | Dolichyl-diphosphooligosaccharide--protein glycosyltransferase subunit 1 Mus musculus | 1447.2 | 51.15 | 68528 | 24 | 24 | 61956117 | 57415889 | 67091837 | 56183701 | 55390873 | 66095485 | 78847553 | 69433105 | 63415133 | 59248505 | 60346057 | 62895293 | 58590545 | 59161693 | 60384317 |
| Q9DBG6 | Rpn2 | Dolichyl-diphosphooligosaccharide--protein glycosyltransferase subunit 2 Mus musculus | 1455.44 | 47.7 | 69063 | 21 | 21 | 33773069 | 32964589 | 43071273 | 40588133 | 38263321 | 38895321 | 32684789 | 40279549 | 53789453 | 32076787 | 36331013 | 39648713 | 29488843 | 37463605 | 44135733 |
| O88796 | Rpp30 | Ribonuclease P protein subunit p30 Mus musculus | 248.8 | 21.64 | 29473 | 4 | 4 | 3400004 | 2658113.25 | 3338017.5 | 2545187 | 3049643.25 | 2267332.75 | 3375255.25 | 2591961.25 | 2748170 | 3454770 | 2469392.5 | 3129970.75 | 2891311.75 | 2426507.5 | 3712520.5 |
| Q9CSU0 | Rprd1b | Regulation of nuclear pre-mRNA domain-containing protein 1B Mus musculus | 228.51 | 25.46 | 36884 | 6 | 6 | 2211920.5 | 4216523 | 1878051.5 | 4398867 | 4087899 | 1676696.5 | 3378257 | 3745260.5 | 3068342.5 | 3587579.25 | 1529555.75 | 3395875 | 3946135.75 | 4355827.5 | 3102931.25 |
| P63325 | Rps10 | 40S ribosomal protein S10 Mus musculus | 322.71 | 32.73 | 18916 | 5 | 5 | 5408785 | 3677941.5 | 4942438 | 4542353 | 4919256.5 | 2963363.5 | 5269617 | 3412983 | 3513871.75 | 5029783.5 | 4047942.5 | 4721007.5 | 3601844.25 | 3473414.5 | 5491057 |
| P62281 | Rps11 | 40S ribosomal protein S11 Mus musculus | 365.91 | 37.34 | 18431 | 8 | 8 | 24565721 | 29397857 | 39459553 | 38108433 | 37294617 | 32741717 | 48825761 | 25799055 | 29215759 | 47113645 | 41843021 | 40229401 | 37294697 | 26017803 | 50929449 |
| P63323 | Rps12 | 40S ribosomal protein S12 Mus musculus | 395.05 | 59.09 | 14525 | 6 | 7 | 32113307 | 32623473 | 36488805 | 37181425 | 36133601 | 32742535 | 33883633 | 29287857 | 31044137 | 34051113 | 38188721 | 32732387 | 30299389 | 31864519 | 35821125 |
| P62278 | Rps13 | 40S ribosomal protein S13 Rattus norvegicus | 488.7 | 47.02 | 17222 | 10 | 10 | 50209561 | 26923841 | 31766423 | 28352915 | 23271535 | 25429359 | 43275153 | 29026755 | 26846813 | 31558969 | 27368721 | 31476201 | 27310469 | 25320473 | 33339145 |
| P13471 | Rps14 | 40S ribosomal protein S14 Rattus norvegicus | 499.57 | 65.56 | 16259 | 8 | 8 | 24958129 | 28438065 | 32976529 | 30938233 | 32210651 | 28294079 | 39396773 | 23937073 | 23385117 | 34603441 | 32805181 | 30095877 | 31309807 | 24243559 | 38954005 |
| P62843 | Rps15 | 40S ribosomal protein S15 Mus musculus | 308.52 | 36.55 | 17040 | 3 | 5 | 5904843 | 3562673.5 | 6260171 | 3366951.75 | 3147139.5 | 5051790 | 6836052 | 3104899.75 | 2509436.25 | 4568525 | 3965820 | 6344303 | 3750933.5 | 5385886 | 5236898 |
| P62245 | Rps15a | 40S ribosomal protein S15a Mus musculus | 446.73 | 64.62 | 14840 | 8 | 8 | 80841425 | 61381577 | 69723017 | 67028045 | 58379629 | 61354985 | 93297001 | 60282961 | 55907153 | 77730249 | 69710105 | 69974337 | 66286721 | 60002981 | 71965673 |
| P14131 | Rps16 | 40S ribosomal protein S16 Mus musculus | 608.94 | 62.33 | 16445 | 13 | 13 | 96580201 | 62943681 | 77330849 | 81807889 | 73851537 | 55399113 | 108574569 | 69644473 | 52961857 | 97552889 | 84003817 | 87268553 | 73604577 | 61297389 | 91860393 |
| P63276 | Rps17 | 40S ribosomal protein S17 Mus musculus | 574.73 | 65.93 | 15524 | 9 | 9 | 11185217 | 9556584 | 12078351 | 11516377 | 10479789 | 10228023 | 15932201 | 10043385 | 10066535 | 12904672 | 12195917 | 15590023 | 9534679 | 11718882 | 16956417 |

| | | | | | | | | | | | | | | | | | | | | | | |
|---|---|---|---|---|---|---|---|---|---|---|---|---|---|---|---|---|---|---|---|---|---|---|
| P62270 | Rps18 | 40S ribosomal protein S18 Mus musculus | 546.05 | 58.55 | 17719 | 11 | 11 | 21092969 | 16196550 | 24059113 | 21873423 | 18565451 | 13297044 | 44838337 | 16648285 | 11378733 | 29805579 | 19699589 | 25008583 | 20719507 | 15404607 | 31912733 |
| P17074 | Rps19 | 40S ribosomal protein S19 Rattus norvegicus | 377.39 | 35.17 | 16085 | 7 | 7 | 25781753 | 23033861 | 20522993 | 22798165 | 21703593 | 13058609 | 22001325 | 19951247 | 18184585 | 23158705 | 16848813 | 18612305 | 20105417 | 20242365 | 17059563 |
| P25444 | Rps2 | 40S ribosomal protein S2 Mus musculus | 707.16 | 39.25 | 31231 | 13 | 13 | 38287237 | 37984777 | 32127023 | 37984445 | 31504255 | 36845561 | 52979953 | 36590373 | 37282665 | 38887105 | 28481991 | 34778573 | 34579305 | 36565161 | 42988033 |
| P60867 | Rps20 | 40S ribosomal protein S20 Mus musculus | 86.11 | 19.33 | 13373 | 2 | 2 | 4084804.5 | 2527764 | 3224031.75 | 4230608 | 3628275 | 3282660.5 | 8428385 | 3085640.25 | 2595923.5 | 3304713.25 | 3409858 | 4220137 | 3312480.75 | 3433110.5 | 7158708 |
| P05765 | Rps21 | 40S ribosomal protein S21 Rattus norvegicus | 259.31 | 57.83 | 9127 | 4 | 4 | 18322141 | 16647120 | 20788409 | 23679899 | 23835877 | 20379323 | 25676239 | 17923493 | 15511365 | 26609681 | 24521735 | 22110769 | 21946145 | 17832425 | 26331073 |
| P62267 | Rps23 | 40S ribosomal protein S23 Mus musculus | 265.94 | 41.96 | 15808 | 4 | 4 | 2338638.5 | 443509.69 | 2379563.5 | 2421914.5 | 933587.94 | 1652806.25 | 4117675 | 1810234.75 | 990653.62 | 1925144.75 | 1990295.25 | 1663897.12 | 1056521.88 | 1779003.75 | 2091862.5 |
| P62849 | Rps24 | 40S ribosomal protein S24 Mus musculus | 342.18 | 35.34 | 15423 | 5 | 6 | 21076983 | 11508957 | 13300076 | 16061965 | 12982588 | 12132090 | 22236473 | 12765705 | 10935915 | 17753339 | 10546821 | 14910199 | 15008945 | 12300426 | 20649941 |
| P62852 | Rps25 | 40S ribosomal protein S25 Mus musculus | 258.58 | 27.2 | 13742 | 5 | 5 | 122681505 | 131242185 | 122062353 | 114899049 | 110873033 | 98731705 | 112699505 | 109716065 | 121508337 | 116524537 | 112746241 | 111687641 | 113042825 | 122593625 | 103206369 |
| P62855 | Rps26 | 40S ribosomal protein S26 Mus musculus | 134.38 | 20.87 | 13015 | 2 | 2 | 4817003.5 | 3925846.25 | 6285140 | 6238324 | 6075653.5 | 4842034.5 | 9130802 | 4996170 | 4373917.5 | 8693813 | 5470297.5 | 7381696 | 6281331 | 5337551 | 9693419 |
| P62982 | Rps27a | Ubiquitin-40S ribosomal protein S27a Rattus norvegicus | 749.14 | 59.62 | 17951 | 12 | 12 | 311169857 | 368934945 | 305579585 | 356052225 | 354830369 | 340621505 | 402782753 | 297669249 | 320640737 | 440585313 | 340055425 | 279667937 | 336813217 | 278657857 | 288982081 |
| P24051 | Rps27l | 40S ribosomal protein S27-like Rattus norvegicus | 163.95 | 26.19 | 9477 | 3 | 3 | 1373994 | 717258.62 | 1610744.88 | 2520221.75 | 1822715 | 1111135.75 | 7268634 | 867114.5 | 710638.5 | 4421258 | 3057214.5 | 3033914.5 | 2097809.75 | 629464.5 | 5905415.5 |
| P62858 | Rps28 | 40S ribosomal protein S28 Mus musculus | 111.9 | 30.43 | 7841 | 2 | 2 | 22935545 | 19365419 | 21594133 | 26559035 | 24158677 | 22622721 | 21911397 | 21767921 | 19012813 | 25382685 | 29610751 | 23762099 | 23560389 | 22051459 | 20818151 |
| P62908 | Rps3 | 40S ribosomal protein S3 Mus musculus | 1028.02 | 74.07 | 26674 | 20 | 20 | 197884705 | 163660033 | 167921617 | 181191601 | 179688833 | 171180849 | 183089665 | 165362993 | 157015985 | 185191361 | 167703777 | 174812129 | 168811505 | 172657729 | 182811297 |
| P49242 | Rps3a | 40S ribosomal protein S3a Rattus norvegicus | 740.66 | 45.83 | 29945 | 13 | 13 | 113750105 | 112958609 | 114948289 | 118346137 | 109531193 | 101607961 | 115720113 | 97707385 | 102285449 | 119121521 | 107497841 | 110747673 | 108324145 | 89856257 | 105194505 |
| P62702 | Rps4x | 40S ribosomal protein S4, X isoform Mus musculus | 909.89 | 59.7 | 29598 | 17 | 17 | 111978305 | 91429985 | 114099345 | 111929257 | 95494737 | 93838873 | 163233633 | 100524521 | 89369801 | 123487777 | 115651129 | 118552337 | 106064169 | 99054905 | 130369353 |
| P97461 | Rps5 | 40S ribosomal protein S5 Mus musculus | 837 | 55.88 | 22889 | 13 | 13 | 65677265 | 68578081 | 71479009 | 69414161 | 76406825 | 68157289 | 75840953 | 60549517 | 58962681 | 73934241 | 63476377 | 63132409 | 62625005 | 63256313 | 58274537 |
| P62754 | Rps6 | 40S ribosomal protein S6 Mus musculus | 498.01 | 31.33 | 28681 | 8 | 8 | 31294497 | 32319381 | 26106187 | 33848281 | 29148607 | 27574125 | 24452037 | 25081713 | 28347789 | 27514545 | 27458877 | 25960007 | 27811073 | 27614385 | 21543841 |
| P18654 | Rps6ka3 | Ribosomal protein S6 kinase alpha-3 Mus musculus | 192.51 | 5.27 | 83694 | 2 | 4 | 2134561.5 | 1042357 | 1728925.38 | 765146.88 | 1434409.5 | 1946144.5 | 681329.31 | 858061.25 | 2125074.5 | 2125302.25 | 1564352.5 | 1881015 | 1457119.75 | 1743161.75 | 1945697.12 |
| P62082 | Rps7 | 40S ribosomal protein S7 Mus musculus | 430.09 | 40.72 | 22127 | 9 | 9 | 30112991 | 31808681 | 27780311 | 29600301 | 27330341 | 24722817 | 30645545 | 26318529 | 24116005 | 30292645 | 20324865 | 28862171 | 28840269 | 27128669 | 24115957 |
| P62242 | Rps8 | 40S ribosomal protein S8 Mus musculus | 677.39 | 50 | 24205 | 11 | 11 | 155997169 | 147219825 | 142502753 | 145336257 | 133317769 | 129331801 | 188689905 | 144652305 | 137097249 | 158827857 | 143184065 | 154701873 | 149239233 | 144948865 | 175906369 |
| P29314 | Rps9 | 40S ribosomal protein S9 Rattus norvegicus | 615.29 | 52.06 | 22591 | 14 | 14 | 177725201 | 71877393 | 87435513 | 51817165 | 70549825 | 67165985 | 134455489 | 80595489 | 74650577 | 92767361 | 79846825 | 84190097 | 77243017 | 80163257 | 108922433 |
| P14206 | Rpsa | 40S ribosomal protein SA Mus musculus | 1142.58 | 61.36 | 32838 | 15 | 17 | 75961185 | 67448417 | 76124585 | 82350089 | 72655481 | 81130297 | 113641601 | 76696385 | 60930221 | 91338273 | 79563113 | 86441985 | 80257993 | 58691109 | 92183993 |
| Q8K4Q0 | Rptor | Regulatory-associated protein of mTOR Mus musculus | 117.3 | 2.17 | 149471 | 2 | 2 | 1317334 | 1095099.75 | 1190953.5 | 589439.56 | 1241394 | 1312076.12 | 921427.38 | 1075679.88 | 1555969.12 | 1314960.5 | 1651691.75 | 1051108.5 | 1103982.75 | 1315226 | 1252477.5 |
| Q63486 | Rraga | Ras-related GTP-binding protein A Rattus norvegicus | 415.81 | 34.5 | 36566 | 7 | 7 | 4424732 | 7776330.5 | 7697051.5 | 6519489 | 6985390 | 5666951 | 6443223.5 | 6367376.5 | 6860332 | 6895289 | 7008107 | 6016036.5 | 6871838 | 6888019 | 4315153.5 |
| Q99K70 | Rragc | Ras-related GTP-binding protein C Mus musculus | 579.47 | 33.67 | 44121 | 10 | 10 | 30362663 | 37790445 | 35462301 | 25095525 | 24505407 | 31452585 | 20631563 | 38130613 | 43545557 | 23279883 | 29492501 | 32837263 | 29664377 | 33961769 | 26879457 |
| D3Z8L7 | Rras | Ras-related protein R-Ras Rattus norvegicus | 235.96 | 28.9 | 23909 | 5 | 5 | 5490828.5 | 9192823 | 5410132.5 | 5420434.5 | 6203214 | 11037345 | 10429717 | 12269221 | 12232379 | 10866950 | 7416872 | 8608885 | 8146214 | 8678971 | 7332298.5 |
| Q99PL5 | Rrbp1 | Ribosome-binding protein 1 Mus musculus | 1522.1 | 24.05 | 172879 | 29 | 29 | 50452633 | 47263069 | 54767909 | 48342281 | 48508321 | 44804677 | 49755257 | 47865281 | 44731509 | 46037853 | 52697145 | 55662805 | 48889173 | 53441061 | 49761749 |
| P07742 | Rrm1 | Ribonucleoside-diphosphate reductase large subunit Mus musculus | 647.75 | 26.64 | 90210 | 14 | 14 | 18246387 | 12284210 | 5937449 | 16716314 | 16616034 | 9535447 | 4218572.5 | 4721790 | 7043884.5 | 4911893.5 | 5650169 | 4992719 | 5052815.5 | 8698067 | 9365249 |
| P11157 | Rrm2 | Ribonucleoside-diphosphate reductase subunit M2 Mus musculus | 298.68 | 14.36 | 45096 | 4 | 4 | 4144637.5 | 3025292.5 | 5096769.5 | 3848701.5 | 3711027.5 | 4687451.5 | 2232711 | 5038131 | 3665417 | 3868701 | 4074120 | 4326796 | 4586293.5 | 3736405 | 4675557.5 |
| P56183 | Rrp1 | Ribosomal RNA processing protein 1 homolog A Mus musculus | 284.55 | 21.26 | 54777 | 5 | 5 | 2734320.75 | 2070033.75 | 3952067.5 | 3494916.75 | 1842751.38 | 2609223.5 | 1195677.25 | 3179994.25 | 8007906 | 984107.38 | 2903160.75 | 2617141.5 | 2566642.5 | 2562727.75 | 2456662.5 |
| Q6P5B0 | Rrp12 | RRP12-like protein Mus musculus | 114.21 | 2.63 | 143131 | 2 | 2 | 756523.94 | 144750.52 | 304117.66 | 425822.5375 | 497898.03 | 504293.75 | 510356.6375 | 647105.62 | 469632.06 | 420395.12 | 322834.72 | 396259.47 | 312019.28 | 325565.94 | 308559.59 |
| Q5M947 | Rrp15 | RRP15-like protein Rattus norvegicus | 68.1 | 6.43 | 30926 | 2 | 2 | 771086.56 | 1 | 843622.56 | 1 | 1 | 1 | 1 | 1 | 1 | 292496.78 | 1 | 1 | 1 | 195609.31 | 1 |
| Q91YK2 | Rrp1b | Ribosomal RNA processing protein 1 homolog B Mus musculus | 268.17 | 10.22 | 80582 | 5 | 5 | 5406735 | 5591422.5 | 6015206 | 4887104 | 5657125 | 5466688 | 5306124.5 | 5151011 | 4506822.5 | 5585854.5 | 5893757 | 5400276.5 | 5062272.5 | 5013775 | 5297409.5 |
| Q9DB85 | Rrp8 | Ribosomal RNA-processing protein 8 Mus musculus | 46.41 | 3.28 | 51066 | 1 | 1 | 759697 | 1 | 742256.19 | 1 | 794526.88 | 573842.62 | 1 | 551482.88 | 592495.5 | 1 | 855662.38 | 768368.06 | 640464.5 | 789603.875 | 893920.56 |
| Q91WM3 | Rrp9 | U3 small nucleolar RNA-interacting protein 2 Mus musculus | 86.69 | 6.95 | 52107 | 2 | 2 | 408973.44 | 324433.41 | 385185.69 | 323987.12 | 362741.88 | 350116.75 | 335863.38 | 334129.97 | 274777.81 | 355674.91 | 491611.28 | 236473.34 | 303495.03 | 415717.66 | 408555.75 |
| Q9CYH6 | Rrs1 | Ribosome biogenesis regulatory protein homolog Mus musculus | 233.57 | 16.16 | 41552 | 5 | 5 | 649089.12 | 891159 | 1558807.5 | 415914.56 | 497779.84 | 1091770.88 | 978960.4925 | 885689.56 | 1598360.5 | 340021.03 | 990652.88 | 921217.25 | 607019.94 | 1053143.25 | 744572.19 |
| Q8BVY0 | Rsl1d1 | Ribosomal L1 domain-containing protein 1 Mus musculus | 603.76 | 22.79 | 50421 | 9 | 9 | 37531885 | 21401785 | 28499665 | 50338025 | 40785601 | 21256345 | 52828565 | 22149901 | 8003401.5 | 40106913 | 37764845 | 11189997 | 22271705 | 18651323 | 30034529 |
| Q6P6G7 | Rsl24d1 | Probable ribosome biogenesis protein RLP24 Rattus norvegicus | 72.87 | 11.66 | 19653 | 2 | 2 | 612981.56 | 618317 | 2277953.25 | 1096974 | 815260.75 | 1776942.25 | 725475.5 | 1873370.88 | 1952748 | 797489.19 | 530418.69 | 487960.19 | 945391.62 | 603459.88 | 838615.75 |
| Q01730 | Rsu1 | Ras suppressor protein 1 Mus musculus | 361.67 | 45.85 | 31550 | 7 | 7 | 9417672 | 10143515 | 9792885 | 10924646 | 8617757 | 8766041 | 10656497 | 11248521 | 8152309.5 | 8982613 | 8705945 | 10210407 | 9587488 | 10077393 | 9714795 |
| Q9D7H3 | RtcA | RNA 3'-terminal phosphate cyclase Mus musculus | 274.63 | 17.21 | 39254 | 4 | 4 | 2739914.75 | 2188632.5 | 2828345.75 | 2486478.5 | 1789373.12 | 2073584.75 | 3769622.5 | 2877978.5 | 2410350 | 2781736.5 | 2768993.25 | 2931411.25 | 1969552.88 | 2524566.5 | 3503022.25 |
| Q6AYT3 | Rtcb | tRNA-splicing ligase RtcB homolog Rattus norvegicus | 579.08 | 28.71 | 55249 | 12 | 12 | 15347944 | 16400637 | 20071873 | 16082753 | 16561951 | 15380314 | 20431461 | 18819553 | 17399801 | 16663386 | 18801921 | 21035597 | 19608581 | 18290285 | 21672957 |
| Q3T1J8 | Rtfdc1 | Protein RTF2 homolog Rattus norvegicus 2 | 40.84 | 3.27 | 33812 | 1 | 1 | 663241.5 | 444095.25 | 635196.69 | 287354.75 | 386546.75 | 697869.38 | 549651.38 | 594218.31 | 746330.56 | 279902.72 | 536012.19 | 709177.44 | 497214.03 | 658513.75 | 683971.75 |
| Q9ES97 | Rtn3 | Reticulon-3 Mus musculus | 186.54 | 6.54 | 103879 | 3 | 3 | 7813480 | 4670133 | 4226623.5 | 3397479.75 | 4866151 | 6120038.5 | 5051828.5 | 4849366 | 6529556.5 | 4775475.5 | 5907406 | 5318843 | 3971695.75 | 5267054.5 | 4484266.5 |
| Q99P72 | Rtn4 | Reticulon-4 Mus musculus | 984.33 | 25.13 | 126613 | 15 | 15 | 62349229 | 58925973 | 62964929 | 54204753 | 69060793 | 73949689 | 78835825 | 72101969 | 69173321 | 78174937 | 76448497 | 76274977 | 69393881 | 63865777 | 76015553 |
| Q9CQE8 | RTRAF | RNA transcription, translation and transport factor protein Mus musculus | 513.98 | 40.57 | 28152 | 8 | 8 | 10452018 | 7446543.5 | 8612863 | 5779640.5 | 6114199 | 8829911 | 4587251 | 8851881 | 11123328 | 5691044.5 | 8777928 | 9154152 | 5000510 | 5529457.5 | 6755909 |
| Q8R4Y8 | Rttn | Rotatin Mus musculus | 37.11 | 0.31 | 248489 | 1 | 1 | 1948464.75 | 1 | 1000575.56 | 658666.75 | 1 | 241182.8 | 5618045.5 | 621784.56 | 2003355.31 | 1532408.38 | 648713.5 | 1580382.75 | 542479.62 | 409325.09 | 2557746 |
| Q8BIJ7 | Rufy1 | RUN and FYVE domain-containing protein 1 Mus musculus | 408.16 | 14.19 | 80376 | 9 | 9 | 7251219 | 8579220 | 7904312.5 | 9282424 | 8046382 | 7573953 | 7824689.5 | 7853511 | 7303923 | 8667405 | 9030163 | 8075134.5 | 8600797 | 4091493 | 7923236.5 |
| P60122 | Ruvbl1 | RuvB-like 1 Mus musculus | 878.56 | 42.76 | 50214 | 15 | 15 | 34250041 | 33843885 | 35951393 | 33710441 | 32442131 | 31914625 | 36454393 | 34351917 | 40955901 | 34444449 | 32980035 | 32842621 | 31407479 | 34245669 | 33998749 |
| Q9WTM5 | Ruvbl2 | RuvB-like 2 Mus musculus | 1156.47 | 54.64 | 51113 | 21 | 21 | 159727153 | 116194353 | 119336737 | 173795409 | 156698625 | 109794737 | 136161889 | 102835385 | 67733913 | 153292465 | 139777217 | 71474217 | 107363233 | 89141105 | 108427185 |
| Q9CQK7 | Rwdd1 | RWD domain-containing protein 1 Mus musculus | 94.49 | 14.4 | 27785 | 2 | 2 | 419287.75 | 1 | 464638 | 1 | 491980.84 | 549059.81 | 1 | 1 | 620763.94 | 452754.69 | 1 | 395958.72 | 631264.81 | 1 | 460009.66 |
| Q8CCI5 | Rybp | RING1 and YY1-binding protein Mus musculus | 79.92 | 18.86 | 24777 | 2 | 2 | 285203.12 | 581921.25 | 569095.38 | 500089.66 | 575833.75 | 538703.75 | 676379.56 | 256069.75 | 273930.22 | 566106.12 | 708080.12 | 593571.69 | 488233.69 | 465996.03 | 686449.75 |
| P56565 | S100a1 | Protein S100-A1 Mus musculus | 101.41 | 19.15 | 10505 | 2 | 2 | 967942.06 | 1518875.62 | 417689.59 | 1440948.75 | 1295038.62 | 1308662 | 1055602.38 | 425716.47 | 589707.38 | 1386539.75 | 547152.75 | 727761.62 | 1537787.12 | 895658.38 | 586546.31 |
| P08207 | S100a10 | Protein S100-A10 Mus musculus | 143.58 | 27.84 | 11186 | 2 | 2 | 19057881 | 19975681 | 17181357 | 18627387 | 24212561 | 24788229 | 16170709 | 21301541 | 20854801 | 13801921 | 16347381 | 16251079 | 18277677 | 25650941 | 11135357 |
| P50543 | S100a11 | Protein S100-A11 Mus musculus | 300.34 | 54.08 | 11083 | 4 | 5 | 8823502 | 10569310 | 9505049 | 9836684 | 10115149 | 11567734 | 8819007 | 10503759 | 11680303 | 11295280 | 11203014 | 8981044 | 12278790 | 11297112 | 9069191 |
| P07091 | S100a4 | Protein S100-A4 Mus musculus | 540 | 84.16 | 11721 | 7 | 9 | 342641921 | 365499169 | 282173601 | 455036193 | 395784161 | 305730529 | 240651697 | 243941521 | 241786849 | 256404145 | 387054721 | 286390977 | 356148481 | 383670561 | 255797857 |
| P05964 | S100a6 | Protein S100-A6 Rattus norvegicus | 373.2 | 64.04 | 10035 | 5 | 5 | 147885137 | 153946993 | 109438433 | 153265377 | 135110529 | 130598289 | 117486065 | 120629009 | 97620825 | 132451969 | 108503449 | 87248649 | 130402329 | 117599337 | 84515473 |
| Q9D2C2 | Saal1 | Protein SAAL1 Mus musculus | 228.47 | 10.55 | 52769 | 4 | 4 | 145683.84 | 853157.88 | 868802.88 | 552912.88 | 495066.44 | 443192.5 | 876225.25 | 186438.02 | 182700.55 | 282972.03 | 691631.75 | 371825.88 | 777005.25 | 232151.12 | 284088.75 |
| Q9EP69 | Sacm1l | Phosphatidylinositide phosphatase SAC1 Mus musculus | 310.02 | 14.48 | 66944 | 6 | 6 | 4200392 | 4747371.5 | 4502654.5 | 5046367 | 4558957 | 4593182.5 | 4332263.5 | 4768535.5 | 6247812 | 4207126 | 5765025 | 4072136.5 | 2861735.5 | 5248997.5 | 4560538.5 |
| Q9R1T2 | Sae1 | SUMO-activating enzyme subunit 1 Mus musculus | 764.91 | 47.14 | 38620 | 13 | 13 | 17726315 | 19295161 | 16435100 | 20360735 | 21586009 | 18134305 | 12540303 | 18431221 | 12122507 | 19195353 | 20974161 | 17139447 | 20918775 | 19708653 | 16638579 |
| D3YXK2 | Safb | Scaffold attachment factor B1 Mus musculus | 378.03 | 15.58 | 105104 | 4 | 8 | 18522413 | 19448347 | 18781235 | 17714511 | 19606143 | 18638445 | 19660857 | 19846767 | 24859657 | 19590445 | 21014193 | 20537213 | 17869799 | 18324619 | 17581825 |
| Q80YR5 | Safb2 | Scaffold attachment factor B2 Mus musculus | 411.82 | 10.8 | 111838 | 4 | 7 | 14648402 | 16451889 | 15362969 | 13104848 | 15903931 | 15253180 | 14560295 | 16047743 | 21428155 | 13648321 | 16317720 | 17051331 | 13230141 | 15126912 | 15613664 |
| Q0VE29 | Samd12 | Sterile alpha motif domain-containing protein 12 Mus musculus 2 | 34.63 | 6.21 | 18245 | 1 | 1 | 1921470.25 | 2018619.25 | 463416.31 | 1535382.733 | 1738025.12 | 1304731.75 | 1007372.56 | 1470577.5 | 1184298.89 | 954513.75 | 1145692.62 | 837247.25 | 1 | 1633194.12 | 1 |
| Q60710 | Samhd1 | Deoxynucleoside triphosphate triphosphohydrolase SAMHD1 Mus musculus | 1087.99 | 36.93 | 75893 | 18 | 19 | 79472193 | 78731289 | 68233169 | 67630641 | 75887353 | 67221225 | 66274661 | 74948001 | 82125209 | 59066997 | 83260425 | 81396009 | 75062257 | 84048809 | 83737897 |
| Q8BGH2 | Samm50 | Sorting and assembly machinery component 50 homolog Mus musculus | 769.28 | 41.58 | 51864 | 15 | 15 | 21734131 | 19847863 | 20459641 | 19857669 | 19958101 | 22979163 | 21053505 | 21147901 | 22422585 | 23522843 | 20830119 | 20976719 | 21553759 | 21648367 | 22887605 |
| O55128 | Sap18 | Histone deacetylase complex subunit SAP18 Mus musculus | 126.72 | 15.03 | 17595 | 2 | 2 | 1 | 1 | 331385.66 | 533466.81 | 1 | 1001582.75 | 461195.81 | 468020.06 | 1 | 1 | 801331.44 | 1 | 1 | 1 | 347247.19 |
| P36536 | Sar1a | GTP-binding protein SAR1a Mus musculus | 240.6 | 27.27 | 22371 | 2 | 3 | 2159137.5 | 1953319.5 | 1877033.5 | 1812069.75 | 1819013.25 | 2092246.5 | 1496555 | 2033484.38 | 1786829.75 | 1491120.5 | 1478593.25 | 1353969 | 1677708 | 1984249.38 | 1 |
| Q5HZY2 | Sar1b | GTP-binding protein SAR1b Rattus norvegicus | 295.25 | 29.8 | 22410 | 3 | 4 | 3482802 | 3752000.25 | 5363574 | 3161496.75 | 2763001.25 | 4725783 | 3171140 | 5058654.5 | 5515585 | 3034021.25 | 3789060 | 3318430 | 3435573 | 5522411 | 4652247 |
| Q498U4 | Sarnp | SAP domain-containing ribonucleoprotein Rattus norvegicus | 306.32 | 27.62 | 23605 | 5 | 5 | 3018168.75 | 4519868 | 2776125.25 | 5417837.5 | 2201666 | 2964328.5 | 10402167 | 2938577.5 | 3137358.75 | 8494605 | 2720610 | 4270714 | 3647784.75 | 3021876.5 | 8284282.5 |
| P26638 | Sars | Serine--tRNA ligase, cytoplasmic Mus musculus | 1343.12 | 51.56 | 58389 | 22 | 22 | 93568409 | 104996937 | 91261817 | 111538577 | 105797465 | 106350121 | 115956369 | 96024313 | 92389777 | 113551769 | 114903161 | 94205977 | 99625553 | 94083561 | 114623897 |
| Q9JJL8 | Sars2 | Serine--tRNA ligase, mitochondrial Mus musculus | 63.79 | 2.32 | 58316 | 1 | 1 | 1031626.44 | 1129053.38 | 2041283.12 | 1310794.25 | 1255277 | 1135310.12 | 996734.88 | 1297059.5 | 1254403 | 1094267 | 783215.38 | 1372080.38 | 1312003.88 | 1288894.5 | 888822.44 |
| Q5XIW8 | Sart1 | U4/U6.U5 tri-snRNP-associated protein 1 Rattus norvegicus | 126.17 | 4.09 | 91011 | 3 | 3 | 1022317.44 | 1042419.88 | 1167091.5 | 1258104.25 | 1175796.25 | 1229363.62 | 2252825.5 | 1240864 | 1284431.5 | 1898648 | 1657308.75 | 1626504.62 | 1166248.62 | 947230.75 | 1661527.25 |
| Q9JLI8 | Sart3 | Squamous cell carcinoma antigen recognized by T-cells 3 Mus musculus | 375.27 | 8.73 | 109619 | 7 | 7 | 5469114.5 | 5766772.5 | 5798954 | 5408690 | 4815226.5 | 4488576 | 5218169 | 4841246.5 | 4900946 | 5873959 | 5797944 | 4074403.75 | 4307432 | 5432483.5 | 4543464.5 |
| P59808 | Sash1 | SAM and SH3 domain-containing protein 1 Mus musculus | 67.6 | 0.89 | 135591 | 1 | 1 | 1415810 | 1346030.88 | 1071407 | 1438931.88 | 1253963 | 1292953.75 | 1574047 | 1316084.12 | 1463555.75 | 1394348.38 | 1452332.12 | 1056617.75 | 1220500 | 1108360.12 | 1093164.25 |
| P70122 | Sbds | Ribosome maturation protein SBDS Mus musculus | 52.49 | 4 | 28781 | 1 | 1 | 468094.38 | 261346.23 | 200839.12 | 131399.03 | 657034.5 | 529655.88 | 1 | 451828.5 | 1 | 1 | 174153.56 | 332977.44 | 1 | 767806.19 | 1 |
| Q6ZPE2 | Sbf1 | Myotubularin-related protein 5 Mus musculus | 110.26 | 1.55 | 208693 | 2 | 2 | 560052.06 | 577469.31 | 622520.06 | 691935.44 | 894105.12 | 930743.81 | 594777.69 | 748315.06 | 568180 | 1038379.12 | 640258.06 | 789420.25 | 624922.38 | 797414.12 | 750716.75 |
| Q63623 | Scaf8 | Protein SCAF8 Rattus norvegicus | 74.17 | 0.95 | 139504 | 1 | 1 | 3052153.75 | 2777303.75 | 2723363 | 2300131 | 2893278.5 | 2873564.25 | 1828263.62 | 2984410.25 | 3082506.5 | 2294988 | 1808447.38 | 2245673.25 | 2516986.25 | 2886650.25 | 1817127.62 |
| O35114 | Scarb2 | Lysosome membrane protein 2 Mus musculus | 295.67 | 14.85 | 54044 | 5 | 5 | 3792335.75 | 3064972 | 4783295 | 2932860 | 2375041.5 | 5984264 | 2572157.25 | 9275039 | 8410833 | 2821439.75 | 7242045 | 6889365 | 2531352.5 | 3676426 | 6615578 |
| Q8R127 | Sccpdh | Saccharopine dehydrogenase-like oxidoreductase Mus musculus | 159.27 | 10.49 | 47129 | 3 | 3 | 1243900.75 | 2271185.75 | 1727978.5 | 1769493 | 1637801.5 | 1709623.25 | 2418504.5 | 1819070.38 | 1580104.12 | 1821500.12 | 1300250.12 | 1455996.38 | 1789188.5 | 1622517.75 | 1548424.38 |
| Q8BRF7 | Scfd1 | Sec1 family domain-containing protein 1 Mus musculus | 512.72 | 21.91 | 72323 | 9 | 9 | 7423727 | 9705450 | 8513180 | 5661733.5 | 9249032 | 9386353 | 10200310 | 10102412 | 9367615 | 8835872 | 9754659 | 6301885 | 7666374.5 | 6228955 | 8859304 |
| Q9JLI6 | Scly | Selenocysteine lyase Mus musculus | 238.14 | 13.43 | 47174 | 5 | 5 | 6722862.5 | 6215261 | 6757017 | 5159280 | 7493572 | 4867310 | 6720086.5 | 3727194.75 | 5572226 | 7169341 | 7529892.5 | 6406845 | 5056374 | 5465251.5 | 4075367 |
| P32020 | Scp2 | Non-specific lipid-transfer protein Mus musculus | 321.76 | 16.09 | 59126 | 6 | 6 | 12118544 | 14167560 | 10426317 | 9385977 | 12512601 | 11678144 | 13028001 | 11318397 | 11095413 | 11757495 | 11314534 | 10780570 | 13608964 | 12416604 | 9138405 |
| Q920A5 | Scpep1 | Retinoid-inducible serine carboxypeptidase Mus musculus | 619.39 | 28.76 | 50965 | 11 | 13 | 64574505 | 79703097 | 73722761 | 67582617 | 66291773 | 74839105 | 58413009 | 82795249 | 81054241 | 62996741 | 58051809 | 65807773 | 75398865 | 79624153 | 57425889 |
| Q80U72 | Scrib | Protein scribble homolog Mus musculus | 108.41 | 3.47 | 174059 | 3 | 3 | 243420.61 | 164917.41 | 227873.67 | 222660.3 | 177761.56 | 1 | 339582.41 | 219570.06 | 1 | 258466.03 | 208865.64 | 327367.06 | 256213.23 | 172898.14 | 223411.83 |
| Q8CFE4 | Scyl2 | SCY1-like protein 2 Mus musculus | 80.93 | 2.69 | 103317 | 2 | 2 | 185693.36 | 540642.88 | 685600.88 | 205727.7 | 206164.47 | 597248.5 | 255295.67 | 1101223.5 | 1218457.12 | 206922.92 | 720390.12 | 973893.62 | 283078.81 | 854196.25 | 770933.62 |
| Q9DBQ7 | Scyl3 | Protein-associating with the carboxyl-terminal domain of ezrin Mus musculus | 100.97 | 6.53 | 81333 | 3 | 3 | 1147515.5 | 1362701 | 1369663.38 | 1772628 | 651327.56 | 932057 | 1022021.12 | 1112259.25 | 1387206.5 | 1249644.38 | 870453.94 | 1078728 | 1307787 | 1342724 | 1046273.62 |
| O08992 | Sdcbp | Syntenin-1 Mus musculus | 423.87 | 53.18 | 32379 | 7 | 7 | 1300222.12 | 1207490.5 | 2344484 | 801476.5 | 1769486.5 | 1932388.88 | 3215072 | 1519113.38 | 1112105.38 | 3408814 | 2434432 | 2705793.25 | 2373426.25 | 1328876 | 3378984 |
| Q9ESP1 | Sdf2l1 | Stromal cell-derived factor 2-like protein 1 Mus musculus | 69.61 | 13.57 | 23648 | 2 | 2 | 685499.3125 | 737664.38 | 493002.62 | 688225.5 | 823104.75 | 1260629 | 1777144 | 1 | 1 | 1 | 494808.47 | 2000686.25 | 475358.44 | 584000.62 | 973912.19 |
| Q8K2B3 | Sdha | Succinate dehydrogenase [ubiquinone] flavoprotein subunit, mitochondrial Mus musculu | 994.47 | 38.86 | 72585 | 16 | 16 | 52981733 | 55452269 | 54265109 | 67279817 | 65987317 | 60513309 | 71457033 | 56273513 | 55917593 | 66756813 | 68518249 | 60083713 | 63532485 | 58953057 | 62867629 |
| Q9CQA3 | Sdhb | Succinate dehydrogenase [ubiquinone] iron-sulfur subunit, mitochondrial Mus musculus | 438.18 | 30.5 | 31814 | 8 | 8 | 44796633 | 49837093 | 50325585 | 50812549 | 49363261 | 41455277 | 58671445 | 53031077 | 56065157 | 51527645 | 54888665 | 49934385 | 47705673 | 51334309 | 52418457 |
| Q9CZB0 | Sdhc | Succinate dehydrogenase cytochrome b560 subunit, mitochondrial Mus musculus | 167.48 | 21.89 | 18382 | 3 | 3 | 12820827 | 11249965 | 9982293 | 8338896 | 10829868 | 12264037 | 9319235 | 11650094 | 13994507 | 8502263 | 8661667 | 8844897 | 9398869 | 11597479 | 8803520 |
| Q9CXV1 | Sdhd | Succinate dehydrogenase [ubiquinone] cytochrome b small subunit, mitochondrial Mus | 82.47 | 16.35 | 17014 | 1 | 1 | 2303064.75 | 1947185.12 | 1287627.62 | 1616036.25 | 1565868.62 | 1784257.62 | 946340.81 | 1329562.62 | 928632.25 | 1572531.5 | 934084.25 | 731724.19 | 1033517.31 | 1302246 | 1174623 |
| P42667 | Sec11a | Signal peptidase complex catalytic subunit SEC11A Rattus norvegicus 2 | 126.39 | 10.61 | 20599 | 2 | 2 | 1012932.69 | 1468945.88 | 1810254.12 | 783664.69 | 1064958.88 | 1129913.75 | 857695.62 | 1145088 | 832212.62 | 828176.44 | 689583.62 | 685292.94 | 1365365.12 | 742732.12 | 762588.75 |
| Q5XFW8 | Sec13 | Protein SEC13 homolog Rattus norvegicus | 301.91 | 23.91 | 35548 | 5 | 5 | 6235783 | 7814998.5 | 8158002 | 5373641 | 7645573 | 9151619 | 7808382 | 7049820 | 8690486 | 7286746 | 7132029 | 6163055 | 6908753 | 7332959 | 8234748 |
| O08547 | Sec22b | Vesicle-trafficking protein SEC22b Mus musculus | 415.75 | 43.72 | 24741 | 8 | 8 | 5180140 | 3469864 | 3923791 | 4471955 | 4959938 | 4032170 | 4798745 | 3993187 | 3506067.5 | 5204595 | 4952113 | 4319432.5 | 3824658 | 3687140.5 | 4244183 |
| Q01405 | Sec23a | Protein transport protein Sec23A Mus musculus | 237.9 | 6.93 | 86162 | 2 | 4 | 5033725 | 2407798.75 | 4261623 | 2926656 | 4742403 | 4947660.5 | 4081634.75 | 4734281 | 2606148.75 | 3177244.5 | 3477855.5 | 5243938.5 | 2352268.5 | 4074583 | 2741295 |
| Q9D662 | Sec23b | Protein transport protein Sec23B Mus musculus | 799.74 | 32.46 | 86437 | 12 | 14 | 7470714 | 8951778 | 9549329 | 8320190.5 | 8321831 | 7492965 | 11040161 | 7908860 | 8388824 | 9786457 | 9431752 | 10746352 | 8946931 | 9210559 | 11158122 |
| Q6NZC7 | Sec23ip | SEC23-interacting protein Mus musculus | 128.94 | 4.11 | 110780 | 3 | 3 | 1 | 1 | 1 | 636808.69 | 668890.06 | 587912.75 | 624330.12 | 401792.38 | 458002.12 | 813772.69 | 536200 | 601152.44 | 677581.06 | 758952.62 | 795702.31 |
| Q3U2P1 | Sec24a | Protein transport protein Sec24A Mus musculus | 45.44 | 1.38 | 118782 | 1 | 1 | 1 | 1 | 1 | 1 | 1 | 1 | 1 | 1 | 1 | 1 | 318074.5 | 1 | 1 | 247933.58 | 243367.59 |
| Q3UPL0 | Sec31a | Protein transport protein Sec31A Mus musculus | 938.73 | 21.38 | 133569 | 18 | 18 | 14539925 | 15577787 | 15663304 | 13616549 | 14152855 | 14833278 | 15090087 | 16356816 | 19330721 | 15194078 | 18416195 | 17659845 | 14135043 | 16158387 | 16556284 |
| P61620 | Sec61a1 | Protein transport protein Sec61 subunit alpha isoform 1 Mus musculus | 425.25 | 24.58 | 52265 | 6 | 6 | 15066201 | 10202939 | 5386791 | 9174268 | 5527858 | 7350473 | 5912462 | 9188252 | 7822223 | 9863161 | 8964027 | 7621081 | 9812327 | 9207623 | 6592578.5 |
| Q9CQS8 | Sec61b | Protein transport protein Sec61 subunit beta Mus musculus | 122.98 | 26.04 | 9958 | 2 | 2 | 12811087 | 15127483 | 16301788 | 15294817 | 15382770 | 16022591 | 16788525 | 13774515 | 15814754 | 17857737 | 17071121 | 14378607 | 15610267 | 12978260 | 14112879 |
| P60060 | Sec61g | Protein transport protein Sec61 subunit gamma Mus musculus 3 | 49.17 | 17.65 | 7741 | 1 | 1 | 1013166.935 | 940403.62 | 1254208.25 | 832944.81 | 1025111.06 | 984734.12 | 1383385.12 | 1081144.38 | 1001019.38 | 1425719.62 | 1 | 1 | 1141201.62 | 930074.06 | 1 |
| Q8BU14 | Sec62 | Translocation protein SEC62 Mus musculus | 41.11 | 2.51 | 45581 | 1 | 1 | 2574255 | 2461246.25 | 2742003.5 | 2221585.5 | 3078395.25 | 2764114.75 | 2672945.5 | 2267111.25 | 2822207.75 | 3086892.75 | 1074914.62 | 2608327.25 | 2915313.75 | 2542399.75 | 2817198.25 |
| Q6A098 | Secisbp2l | Selenocysteine insertion sequence-binding protein 2-like Mus musculus | 49.94 | 1.93 | 119641 | 1 | 1 | 123743.5275 | 161187.5 | 95700.15 | 109622.68 | 128463.78 | 110926.4775 | 120930.13 | 99025.23 | 153159.39 | 70591.16 | 1 | 99853.72 | 120482.49 | 167033.58 | 1 |
| Q8R2U0 | Seh1l | Nucleoporin SEH1 Mus musculus | 77.55 | 5.28 | 39775 | 2 | 2 | 795350.62 | 860282.765 | 863834.31 | 920956.69 | 860989.44 | 584581.75 | 1645547.5 | 803193.12 | 677919.62 | 1505455.75 | 999941.81 | 317386.34 | 829985.75 | 641349.31 | 1358253.88 |

| | | | | | | | | | | | | | | | | | | | | | | |
|---|---|---|---|---|---|---|---|---|---|---|---|---|---|---|---|---|---|---|---|---|---|---|
| Q9Z2G6 | Sel1l | Protein sel-1 homolog 1 Mus musculus | 180.16 | 7.22 | 88340 | 3 | 3 | 659071.56 | 842661.31 | 786070.31 | 617422.25 | 558371.38 | 733101 | 569973.38 | 769432.88 | 881951.31 | 528611.56 | 618094.31 | 670241.75 | 451161.19 | 800923.69 | 818309.19 |
| Q923V8 | Selenof | Selenoprotein F Rattus norvegicus | 196.66 | 41.36 | 17822 | 4 | 4 | 706844.62 | 1173329.88 | 792476.62 | 955051 | 1068103.25 | 1340912.38 | 1366017 | 1037170.44 | 1315911.25 | 1261245.75 | 1136084.75 | 916536.12 | 1073166.75 | 1364991.38 | 1157744.75 |
| O09126 | Sema4d | Semaphorin-4D Mus musculus | 229.88 | 7.08 | 95640 | 5 | 5 | 4311965 | 3744597.75 | 2528419 | 3849758.5 | 2527764.5 | 3540151 | 2227539.25 | 2855496 | 2407455 | 3447965.5 | 4867582.5 | 2169156 | 3548767 | 1809004.62 | 2951847.75 |
| Q8BH69 | Sephs1 | Selenide, water dikinase 1 Mus musculus | 124.17 | 9.95 | 42907 | 4 | 4 | 4561529.5 | 3079394.75 | 4759814.5 | 3105929.25 | 3595903.75 | 3596215.25 | 2734172 | 3437457 | 6957712 | 2196949 | 3990442.75 | 4998822.5 | 3252063 | 5207463 | 2862915 |
| B3GNI6 | Sept11 | Septin-11 Rattus norvegicus | 426.09 | 30.16 | 49695 | 6 | 10 | 2556646.75 | 2712279.5 | 3552162.25 | 4299992.5 | 4791268.5 | 4037406 | 4426391 | 3081886 | 3193402.5 | 4043102.25 | 3337563 | 4207519.5 | 3822486.5 | 3631260 | 4168002 |
| P42208 | Sept2 | Septin-2 Mus musculus | 642.11 | 43.21 | 41526 | 9 | 10 | 28462507 | 28598655 | 27987833 | 26868779 | 28410479 | 32278057 | 31145361 | 30028825 | 29847553 | 29637537 | 26870447 | 29317097 | 26165293 | 27561039 | 28092311 |
| Q9R1T4 | Sept6 | Septin-6 Mus musculus | 173.51 | 8.53 | 49620 | 1 | 4 | 1726269.25 | 1790439.25 | 1607049.5 | 1807783.62 | 2788831.75 | 2086441.88 | 2118532.75 | 1837648.38 | 1920277.75 | 2040481.5 | 1298919.12 | 1949446.62 | 1863291.62 | 1838939.62 | 2132534.75 |
| O55131 | Sept7 | Septin-7 Mus musculus | 811.74 | 41.06 | 50550 | 15 | 16 | 12041432 | 10079734 | 12569328 | 13203832 | 12218833 | 11956940 | 13118102 | 13342762 | 12357618 | 13350972 | 14086022 | 13749328 | 12739399 | 11394393 | 12658205 |
| Q8CHH9 | Sept8 | Septin-8 Mus musculus | 389.48 | 14.45 | 49812 | 4 | 6 | 1973114.75 | 2295610.5 | 2710563.5 | 2638609.5 | 2549811.25 | 2279521.75 | 2369875.5 | 2271432.5 | 2286127.25 | 2577795 | 2295327 | 2363730.5 | 2385478.75 | 2210245 | 2337062 |
| Q80UG5 | Sept9 | Septin-9 Mus musculus | 247.66 | 10.81 | 65575 | 5 | 5 | 13375406 | 13316225 | 14692731 | 15236556 | 12774570 | 11895209 | 16317031 | 13971242 | 13639912 | 13573063 | 11957314 | 13839541 | 12339729 | 12194129 | 13673095 |
| Q6AXS5 | Serbp1 | Plasminogen activator inhibitor 1 RNA-binding protein Rattus norvegicus | 661.45 | 36.36 | 44754 | 11 | 11 | 24421401 | 20042011 | 22517981 | 26039509 | 20794507 | 22446361 | 34296809 | 18213463 | 15269111 | 31015259 | 21959089 | 23392705 | 17674361 | 15527697 | 25071765 |
| Q9QZI9 | Serinc3 | Serine incorporator 3 Mus musculus | 59.3 | 2.54 | 52623 | 1 | 1 | 1522499.25 | 1 | 1 | 1 | 1 | 413860.06 | 391812.62 | 1 | 1 | 299865.12 | 1 | 323974.75 | 639092.62 | 1 | 1 |
| Q9DCQ7 | Serpina1f | Alpha-1-antitrypsin 1-6 Mus musculus | 33.93 | 1.95 | 46337 | 1 | 1 | 5145926 | 4528943.5 | 5142068 | 3530996.75 | 3343327.5 | 2619945.25 | 2006220.5 | 3350871.5 | 3546238.5 | 2204783 | 2894394.5 | 3639189.5 | 3290525.25 | 4034838.75 | 2574036.25 |
| Q4G075 | Serpinb1a | Leukocyte elastase inhibitor A Rattus norvegicus | 69.24 | 2.64 | 42727 | 1 | 1 | 1023972.69 | 1064173.5 | 1096896.38 | 1305444.12 | 1008452.94 | 1294906.5 | 1288696.62 | 1336356.75 | 1678912.5 | 1365857.38 | 847812.5 | 1097065.88 | 1041159.5 | 1034437.62 | 859220.75 |
| Q60854 | Serpinb6 | Serpin B6 Mus musculus | 1224.84 | 60.85 | 42599 | 18 | 19 | 83696937 | 106701513 | 88571385 | 97106305 | 93216201 | 82281505 | 82543825 | 88433337 | 87140977 | 80464817 | 95602673 | 85191641 | 100108713 | 98665201 | 79883217 |
| O08800 | Serpinb8 | Serpin B8 Mus musculus | 234.19 | 15.51 | 42150 | 3 | 3 | 2481567 | 2146444 | 1905547 | 2350871.75 | 1693934.5 | 1737286.12 | 1129401.75 | 1478502.5 | 1597049.38 | 1587338.75 | 996161 | 749999 | 3080298.75 | 2348123.25 | 2155532.25 |
| P32261 | Serpinc1 | Antithrombin-III Mus musculus | 315.63 | 15.05 | 52004 | 6 | 6 | 29093393 | 14399019 | 18221479 | 21083791 | 18452405 | 16267651 | 11063163 | 19723605 | 14302457 | 12624853 | 21934445 | 24116121 | 17251047 | 19069079 | 18769401 |
| Q63945 | Set | Protein SET Rattus norvegicus | 612.54 | 43.94 | 33406 | 9 | 10 | 34842169 | 30783585 | 37072669 | 26052119 | 28159197 | 31624009 | 36621593 | 36604709 | 38291477 | 34933497 | 38876497 | 46800985 | 31705861 | 34280037 | 50837605 |
| Q91WC0 | Setd3 | Histone-lysine N-methyltransferase setd3 Mus musculus | 208.66 | 11.28 | 67176 | 4 | 4 | 1710877 | 1393691.75 | 1837505.38 | 1575329.5 | 1700180.5 | 1365570.5 | 1842663.38 | 1751277.5 | 1641582.88 | 1472501.5 | 2612402.25 | 1561957.25 | 1783547.12 | 1562352.38 | 1678190.5 |
| Q8VHL1 | Setd7 | Histone-lysine N-methyltransferase SETD7 Mus musculus | 88.82 | 4.92 | 40506 | 1 | 1 | 1 | 1 | 1 | 1 | 1980008.75 | 1828391.62 | 1 | 1 | 1653080.5 | 1 | 1 | 995159.38 | 1 | 1306751.5 | 1 |
| Q64213 | Sf1 | Splicing factor 1 Mus musculus | 595.69 | 24.81 | 70403 | 11 | 11 | 19301177 | 19982713 | 18181747 | 23425367 | 21831543 | 15248152 | 18593895 | 19307299 | 20155971 | 21583181 | 20572007 | 18441991 | 21091913 | 18487137 | 17791963 |
| Q8K4Z5 | Sf3a1 | Splicing factor 3A subunit 1 Mus musculus | 1022.27 | 32.11 | 88545 | 19 | 19 | 30420623 | 25910533 | 30410029 | 30604865 | 32499881 | 31520633 | 33847185 | 29505725 | 25029291 | 29680073 | 26389287 | 32480201 | 28922655 | 27658267 | 31252329 |
| Q62203 | Sf3a2 | Splicing factor 3A subunit 2 Mus musculus | 165.61 | 16.21 | 49911 | 4 | 4 | 1577524 | 1475952 | 1110295.5 | 1313441.25 | 1468282.25 | 2210893.25 | 1533661.5 | 1676481.75 | 1284987.62 | 2605487.75 | 2410131.5 | 1157259.5 | 2194460.75 | 1101543.88 | 2364881.75 |
| Q9D554 | Sf3a3 | Splicing factor 3A subunit 3 Mus musculus | 868.71 | 38.72 | 58842 | 14 | 14 | 18947025 | 19965373 | 17937693 | 16690158 | 19102867 | 19732329 | 20906475 | 19693309 | 22981763 | 18912499 | 21286517 | 20089519 | 19252769 | 20745159 | 22759827 |
| Q99NB9 | Sf3b1 | Splicing factor 3B subunit 1 Mus musculus | 985.69 | 23.54 | 145816 | 21 | 21 | 34263065 | 14947607 | 13554555 | 16902905 | 15601250 | 12576204 | 11827910 | 12929530 | 12649411 | 16618713 | 15566746 | 12770150 | 12598195 | 12158316 | 12078774 |
| Q921M3 | Sf3b3 | Splicing factor 3B subunit 3 Mus musculus | 2181.02 | 39.19 | 135550 | 34 | 35 | 37008477 | 37335373 | 42581665 | 40881613 | 40728669 | 39165621 | 43409825 | 41608809 | 41599789 | 45229281 | 40042325 | 43880781 | 41567261 | 40873617 | 43552005 |
| Q6AYL5 | Sf3b4 | Splicing factor 3B subunit 4 Rattus norvegicus | 415.15 | 20.99 | 44356 | 6 | 6 | 8127862 | 8459824 | 6111461.5 | 7344115 | 7829358.5 | 7180527 | 5630800.5 | 7968479 | 6288229 | 7392701.5 | 8783575 | 7213539 | 7471256 | 8095367.5 | 7661741 |
| P59708 | Sf3b6 | Splicing factor 3B subunit 6 Mus musculus | 106.7 | 20 | 14585 | 2 | 2 | 2154996.75 | 2299903.25 | 2438491.25 | 2262488.5 | 1923720 | 1649243.5 | 1623238.25 | 2362472.25 | 2412293 | 1753803.88 | 1848293.38 | 2430997.75 | 2314165.75 | 2071862.75 | 1830272.5 |
| O70456 | Sfn | 14-3-3 protein sigma Mus musculus | 284.71 | 16.13 | 27706 | 1 | 6 | 144261889 | 165393505 | 152082865 | 155488065 | 144588449 | 150192193 | 157130337 | 156185361 | 207688289 | 162917537 | 166139729 | 168053473 | 154000993 | 160814129 | 142794129 |
| Q8VIJ6 | Sfpq | Splicing factor, proline- and glutamine-rich Mus musculus | 993.57 | 27.18 | 75442 | 16 | 16 | 14007516 | 11468027 | 12631364 | 16690725 | 12777351 | 11117236 | 21063405 | 10467364 | 9786328 | 18019475 | 11817000 | 18647049 | 11749400 | 8648924 | 20058349 |
| Q99JR1 | Sfxn1 | Sideroflexin-1 Mus musculus | 434.94 | 31.99 | 35649 | 5 | 6 | 8041927 | 7853453 | 7753863 | 8344489 | 7582172.5 | 8136862.5 | 8068985 | 8361450 | 6902357 | 9079661 | 8915815 | 7339004 | 8974630 | 7099116 | 8670337 |
| Q91V61 | Sfxn3 | Sideroflexin-3 Mus musculus | 202.32 | 18.38 | 35406 | 2 | 3 | 1603607.75 | 1896116 | 1578090.88 | 1786388.12 | 1732143 | 1762444.12 | 1382771.88 | 1981592.75 | 1461174.38 | 1581889.75 | 1932007.62 | 1407369.75 | 1575332.12 | 1696302.88 | 1758414.88 |
| Q8R4V0 | Sgk3 | Serine/threonine-protein kinase Sgk3 Rattus norvegicus | 43 | 2.42 | 57171 | 1 | 1 | 1 | 512506.91 | 1 | 1 | 1 | 488454.25 | 476689.12 | 450544.6925 | 454654.34 | 382381.06 | 393533.41 | 402744.75 | 1 | 1 | 373694.22 |
| Q8R0X7 | Sgpl1 | Sphingosine-1-phosphate lyase 1 Mus musculus | 342.72 | 16.55 | 63677 | 7 | 7 | 12433941 | 14669872 | 14132386 | 12871850 | 14855282 | 13504681 | 11171279 | 15493274 | 16331848 | 13248790 | 13255136 | 12947465 | 15255812 | 15273946 | 12744781 |
| O70593 | Sgta | Small glutamine-rich tetratricopeptide repeat-containing protein alpha Rattus norvegicus | 63.96 | 4.14 | 34157 | 1 | 1 | 1241965.5 | 1529314.5 | 1433334.38 | 1016860.31 | 1864432.75 | 1116288.75 | 753631.06 | 1130791.88 | 2294098 | 960887.5 | 1459850.5 | 1508405.62 | 1383349 | 1474200.88 | 1023197.62 |
| Q9JJU8 | Sh3bgrl | SH3 domain-binding glutamic acid-rich-like protein Mus musculus | 383.94 | 56.14 | 12811 | 7 | 7 | 55023249 | 54861409 | 54446401 | 59335037 | 60166041 | 50005341 | 43702705 | 49228841 | 49986949 | 45807265 | 53488293 | 50770393 | 51868257 | 51506133 | 43117437 |
| Q8BG73 | Sh3bgrl2 | SH3 domain-binding glutamic acid-rich-like protein 2 Mus musculus | 138.63 | 38.32 | 12255 | 3 | 3 | 5735677 | 5209213.5 | 6878262 | 4435783 | 4723495 | 5795696 | 4982274 | 6481742 | 7866784.5 | 3184320 | 4128500 | 5168880 | 4624642 | 7677230.5 | 5458539 |
| Q91VW3 | Sh3bgrl3 | SH3 domain-binding glutamic acid-rich-like protein 3 Mus musculus | 237.33 | 66.67 | 10477 | 4 | 4 | 8359976 | 8215580.5 | 10558641 | 12215968 | 9039162 | 8872921 | 10580030 | 11185343 | 6743043.5 | 11459385 | 8895375 | 10805440 | 9733337 | 8160925.5 | 10663913 |
| P55194 | Sh3bp1 | SH3 domain-binding protein 1 Mus musculus | 366.42 | 14.71 | 74172 | 6 | 6 | 4435971.5 | 4594058 | 3631084 | 4682397 | 3457839 | 4334369 | 4896577.5 | 4223900.5 | 4083624.5 | 4478935.5 | 4523543.5 | 4819995 | 4779207.5 | 4729102.5 | 4572197.5 |
| O35964 | Sh3gl1 | Endophilin-A2 Rattus norvegicus | 262.24 | 17.93 | 41492 | 6 | 6 | 2962311.5 | 1598904.12 | 3551249 | 2889640 | 1 | 2450617 | 2296145.5 | 2736332 | 2893410 | 2224062 | 3010198.25 | 3218377.5 | 2476151 | 2379493 | 2965732 |
| Q6AYE2 | Sh3glb1 | Endophilin-B1 Rattus norvegicus | 215.15 | 10.68 | 40787 | 3 | 3 | 6654449 | 7580770 | 5955345.5 | 7714775 | 7210518 | 6026126 | 5893195.5 | 5893043 | 5585304.5 | 6307689 | 5398864 | 5289601.5 | 6323193 | 5579678 | 4803519 |
| A2AAY5 | Sh3pxd2b | SH3 and PX domain-containing protein 2B Mus musculus | 87.58 | 1.87 | 101517 | 2 | 2 | 861233.5 | 1130739.38 | 1630043.75 | 1116311 | 1239085.88 | 924107 | 1404769.25 | 1053168.75 | 812525.62 | 1113140.12 | 844695.94 | 1098276.12 | 1238174.75 | 876797.44 | 972267.31 |
| P50431 | Shmt1 | Serine hydroxymethyltransferase, cytosolic Mus musculus | 184.35 | 8.16 | 52601 | 2 | 3 | 10176252 | 9464960 | 11348258 | 11411757 | 8762703 | 6860973 | 14751377 | 11142271 | 9838443 | 13501829 | 8173301 | 11727814 | 8679267 | 9864336 | 12158698 |
| Q9CZN7 | Shmt2 | Serine hydroxymethyltransferase, mitochondrial Mus musculus | 1282.55 | 57.54 | 55759 | 23 | 24 | 110442577 | 112967289 | 116171249 | 138613057 | 121700257 | 103043881 | 124117649 | 104289353 | 111176185 | 128823521 | 106630577 | 105782385 | 104379561 | 99759425 | 112781897 |
| Q8K2Q9 | Shtn1 | Shootin-1 Mus musculus | 453.89 | 12.84 | 71343 | 9 | 9 | 18959481 | 21447931 | 19827315 | 19525633 | 19291207 | 17314035 | 16918841 | 18066455 | 19119765 | 17418537 | 19144227 | 17340565 | 18011889 | 20242727 | 15561636 |
| P70665 | Siae | Sialate O-acetylesterase Mus musculus | 186.47 | 7.76 | 60775 | 3 | 3 | 3026954.5 | 6427991 | 5872064.5 | 4669695 | 5355039 | 4488697 | 760011.62 | 6277894 | 5648157.5 | 2467600 | 5038678.5 | 5362460 | 3235274.25 | 5618768 | 3876460.5 |
| O55242 | Sigmar1 | Sigma non-opioid intracellular receptor 1 Mus musculus | 126.45 | 13.9 | 25250 | 2 | 2 | 1644043 | 888876.69 | 952398.69 | 788086.75 | 1070364 | 662997.69 | 747085.19 | 912535.81 | 732123.62 | 778910.88 | 673738.19 | 441712.53 | 522908.47 | 675793.81 | 635142.12 |
| Q6P6S4 | Sil1 | Nucleotide exchange factor SIL1 Rattus norvegicus | 140.87 | 6.45 | 52350 | 3 | 3 | 2155357 | 341537.28 | 1373500.62 | 1334709.12 | 1289616.75 | 2665482.5 | 2068998.88 | 2240426 | 1860893.25 | 2282855.75 | 1863360.62 | 1992335.75 | 1529215.5 | 1685061 | 1271360.12 |
| Q60520 | Sin3a | Paired amphipathic helix protein Sin3a Mus musculus | 186.69 | 4.47 | 145088 | 4 | 4 | 842738.75 | 1095125.75 | 844350.25 | 316112.09 | 396585.84 | 680971.19 | 1093173 | 675921.94 | 775473 | 790957.5 | 577425.88 | 357492.62 | 239757.97 | 383293.84 | 896251.62 |
| Q8C8M1 | Sinhcaf | SIN3-HDAC complex-associated factor Mus musculus | 45.19 | 5.88 | 24829 | 1 | 1 | 193652.61 | 151803.62 | 1 | 311316.62 | 145616.91 | 118383.45 | 137383.42 | 240184.55 | 135893.34 | 162515.42 | 183373.88 | 127560.11 | 151872.88 | 241478.09 | 280828.72 |
| P46062 | Sipa1 | Signal-induced proliferation-associated protein 1 Mus musculus | 144.33 | 4.53 | 112066 | 2 | 2 | 2053289.38 | 2238001 | 2395500 | 2068664.88 | 2462241.25 | 2007359.75 | 1828882.5 | 1950199.5 | 2163299.5 | 1848355.5 | 2374772.5 | 2100890.5 | 1674338.5 | 1975010.75 | 2292782.75 |
| P97797 | Sirpa | Tyrosine-protein phosphatase non-receptor type substrate 1 Mus musculus | 389.84 | 23 | 56425 | 7 | 7 | 23094845 | 21341977 | 20353763 | 19189805 | 20057461 | 18328419 | 18098739 | 16482364 | 15807549 | 19767037 | 19855515 | 16491931 | 20869047 | 17731829 | 16703818 |
| Q5RJQ4 | Sirt2 | NAD-dependent protein deacetylase sirtuin-2 Rattus norvegicus | 136.95 | 7.14 | 39319 | 2 | 2 | 1776509.38 | 1368802.88 | 1590799.25 | 1262741.88 | 1485338.25 | 1612311.188 | 1585086.12 | 1567058.75 | 1536164.5 | 1760935.38 | 1154806 | 1490253.5 | 1697473.75 | 1538355.38 | 1552792.12 |
| Q8R216 | Sirt4 | NAD-dependent protein lipoamidase sirtuin-4, mitochondrial Mus musculus | 51.98 | 3.6 | 37525 | 1 | 1 | 472245.59 | 448417.03 | 421806.75 | 477248.72 | 454929.5225 | 456790.41 | 511965.12 | 479036.81 | 571783.94 | 456906.81 | 235662.91 | 386782.44 | 370353.0325 | 426299.44 | 432667.34 |
| Q3UND0 | Skap2 | Src kinase-associated phosphoprotein 2 Mus musculus | 634.12 | 60.34 | 40712 | 11 | 11 | 24933905 | 26948779 | 22605333 | 23218359 | 24837851 | 23824731 | 23520475 | 23378857 | 25105013 | 22391923 | 26693785 | 21488257 | 24590703 | 23606977 | 22319071 |
| Q6PEC4 | Skp1 | S-phase kinase-associated protein 1 Rattus norvegicus | 412.72 | 55.21 | 18672 | 6 | 6 | 12821218 | 15930318 | 12726561 | 13665122 | 12955385 | 14990764 | 9156320 | 15748193 | 21817427 | 10888639 | 13335001 | 13094884 | 10407639 | 13971800 | 10894756 |
| Q8CI08 | Slain2 | SLAIN motif-containing protein 2 Mus musculus | 48.31 | 2.07 | 62378 | 1 | 1 | 1065192.75 | 797507.75 | 527056.31 | 1317345.38 | 1272489.5 | 648170.75 | 492699.72 | 1 | 1 | 857958 | 792764.94 | 1 | 623578.38 | 1 | 1 |
| Q99MR3 | Slc12a9 | Solute carrier family 12 member 9 Mus musculus | 135.45 | 3.61 | 96330 | 3 | 3 | 1759231 | 1167908.75 | 481658.88 | 1132727.5 | 752233.5 | 1305876.25 | 477560.69 | 1293812.62 | 643966.19 | 728307.19 | 785976.88 | 675650.81 | 1081094.75 | 607229.56 | 692154.69 |
| Q8BPX9 | Slc15a3 | Solute carrier family 15 member 3 Mus musculus | 285.85 | 18.86 | 64051 | 6 | 6 | 9729724 | 9355371 | 5480519.5 | 7941604 | 8179974 | 7885469 | 9350377 | 4799762 | 7009899 | 9149539 | 6385173 | 5921354.5 | 7786643.5 | 5586840.5 | 9852246 |
| P53986 | Slc16a1 | Monocarboxylate transporter 1 Mus musculus | 323.79 | 14.81 | 53267 | 5 | 5 | 3190276 | 1698462.5 | 1684076 | 1281738.88 | 1902557.12 | 2604979.75 | 2350784.25 | 2373236.25 | 2485440.5 | 2502067.5 | 2205914.75 | 1497801.88 | 2107720 | 1910852.62 | 1720931.75 |
| P51912 | Slc1a5 | Neutral amino acid transporter B(0) Mus musculus | 742.2 | 29.29 | 58483 | 11 | 11 | 18431581 | 5509180.5 | 10296889 | 9211809 | 8687740 | 9875344 | 8862510 | 8021202 | 6748150 | 8765106 | 10050407 | 9311061 | 8282765.5 | 3985337 | 9952620 |
| P32089 | Slc25a1 | Tricarboxylate transport protein, mitochondrial Rattus norvegicus | 236.06 | 20.58 | 33835 | 6 | 6 | 14630337 | 11117192 | 12152416 | 11667673 | 15221807 | 14818658 | 16310409 | 11756346 | 12355765 | 15004025 | 20037829 | 14370519 | 14488103 | 14101738 | 15088024 |
| Q9QZD8 | Slc25a10 | Mitochondrial dicarboxylate carrier Mus musculus | 376.96 | 31.71 | 31715 | 6 | 6 | 7433726.5 | 6977366.5 | 5758274 | 7106157.5 | 6173945 | 5713402.5 | 10614308 | 7861584 | 6650987 | 6276371 | 5933855 | 7859627.5 | 5729062 | 5321244.5 | 8012081 |
| Q9CR62 | Slc25a11 | Mitochondrial 2-oxoglutarate/malate carrier protein Mus musculus | 803.98 | 63.06 | 34155 | 13 | 13 | 29607567 | 22388391 | 21583083 | 28706909 | 28475977 | 26152867 | 26658847 | 28771313 | 26309105 | 28259523 | 23405235 | 25493407 | 27606287 | 31150537 | 23051369 |
| Q8BH59 | Slc25a12 | Calcium-binding mitochondrial carrier protein Aralar1 Mus musculus | 1161.56 | 43.72 | 74570 | 17 | 20 | 36904181 | 39552901 | 41379253 | 35412145 | 39784721 | 47953093 | 42510873 | 40043901 | 41561633 | 44538293 | 39124877 | 41970797 | 50035101 | 48464373 | 41117585 |
| Q9QXX4 | Slc25a13 | Calcium-binding mitochondrial carrier protein Aralar2 Mus musculus | 1659.72 | 63.61 | 74467 | 26 | 29 | 62194497 | 61106117 | 65557537 | 57746685 | 64471141 | 81916873 | 79193121 | 74813577 | 69371073 | 76902177 | 73078401 | 71403545 | 77964905 | 74740009 | 72802361 |
| O70579 | Slc25a17 | Peroxisomal membrane protein PMP34 Mus musculus | 72.36 | 4.56 | 34413 | 1 | 1 | 245448.55 | 232337.67 | 208585.3775 | 152327.23 | 204228.06 | 267970.72 | 195969.05 | 173659.59 | 263921.31 | 300494.88 | 204372.53 | 212269.28 | 188301.03 | 1 | 1 |
| Q9D6M3 | Slc25a22 | Mitochondrial glutamate carrier 1 Mus musculus | 110.3 | 7.12 | 34670 | 2 | 2 | 1748356.12 | 2944769.25 | 2173978.25 | 3293245.5 | 2781396.75 | 834409.31 | 1821314 | 1891688.62 | 1848412.5 | 2106839.25 | 2054618.75 | 1987969.75 | 2473728 | 2728627.75 | 1295491.62 |
| Q8BMD8 | Slc25a24 | Calcium-binding mitochondrial carrier protein SCaMC-1 Mus musculus | 335.4 | 19.16 | 52902 | 7 | 7 | 4205191.5 | 3714077.75 | 3746114 | 4745642 | 3702689.75 | 3340369.5 | 5282649 | 3492725.5 | 3382206.75 | 5663385.5 | 3592489 | 3934415 | 3684080.25 | 3052181 | 3711321 |
| P16036 | Slc25a3 | Phosphate carrier protein, mitochondrial Rattus norvegicus | 713.67 | 44.66 | 39445 | 12 | 12 | 36062841 | 41107849 | 41221105 | 34584221 | 38753701 | 44266377 | 50259629 | 50509249 | 58528813 | 47451661 | 47089453 | 42631413 | 42631385 | 42273857 | 59066741 |
| P48962 | Slc25a4 | ADP/ATP translocase 1 Mus musculus | 1171.58 | 63.76 | 32904 | 8 | 20 | 324441665 | 356342497 | 317014433 | 262045585 | 319668129 | 338069601 | 312447553 | 350662273 | 383579681 | 309951841 | 302464481 | 294652673 | 328591585 | 356832001 | 278648705 |
| Q8CFJ7 | Slc25a45 | Solute carrier family 25 member 45 Mus musculus | 80.69 | 5.9 | 31690 | 1 | 1 | 555261.12 | 936192 | 254531.72 | 661330.19 | 674990.75 | 520780.69 | 523943.25 | 662670.38 | 684669.12 | 848316.81 | 556981.5 | 701911.75 | 573125.81 | 687386.38 | 648542.19 |
| P51881 | Slc25a5 | ADP/ATP translocase 2 Mus musculus | 1478.8 | 76.85 | 32931 | 13 | 25 | 436639297 | 474269697 | 425312257 | 362385185 | 427604641 | 453676257 | 420179361 | 461184961 | 504586081 | 416191745 | 406345473 | 390573825 | 428822433 | 463117473 | 376836705 |
| P97849 | Slc27a1 | Long-chain fatty acid transport protein 1 Rattus norvegicus | 37.03 | 2.01 | 71283 | 1 | 1 | 426460.14 | 441851.38 | 373752.81 | 447369.78 | 442866.59 | 518984.88 | 321941.38 | 582023.62 | 647557.94 | 519308.59 | 338933.53 | 420089.44 | 364902.84 | 484121.31 | 321980.22 |
| Q91VE0 | Slc27a4 | Long-chain fatty acid transport protein 4 Mus musculus | 46.27 | 1.4 | 72319 | 1 | 1 | 544022.06 | 804977.62 | 522594.25 | 785672 | 696072.31 | 554160.06 | 906434.69 | 613890.38 | 826047.56 | 717819.12 | 501555.31 | 747909.44 | 775323.31 | 633830.94 | 772583.81 |
| P11167 | Slc2a1 | Solute carrier family 2, facilitated glucose transporter member 1 Rattus norvegicus | 148.34 | 5.69 | 53963 | 4 | 4 | 2903824.5 | 1 | 326483.44 | 1 | 1 | 326671.69 | 410542.91 | 424807.62 | 267417.47 | 431274.53 | 303414.31 | 384290.94 | 565976.44 | 479015.91 | 355922.91 |
| Q9JKN1 | Slc30a7 | Zinc transporter 7 Mus musculus | 51.04 | 3.97 | 41790 | 1 | 1 | 1650122.5 | 1308745.75 | 1441584.25 | 1486542.75 | 1122267 | 1430953 | 1705152.88 | 1201334.25 | 1176084.62 | 1359613 | 1439341 | 1337348.38 | 1411829.25 | 1043521.69 | 1246917.75 |
| Q6AYY8 | Slc33a1 | Acetyl-coenzyme A transporter 1 Rattus norvegicus | 41.69 | 2.36 | 61101 | 1 | 1 | 473776.47 | 1 | 1 | 1 | 205475.41 | 133623.25 | 1 | 174119.39 | 142220.52 | 1 | 151880.48 | 198955.25 | 167400.97 | 200864.86 | 308464.72 |
| P97858 | Slc35b1 | Solute carrier family 35 member B1 Mus musculus | 46.8 | 3.11 | 35883 | 1 | 1 | 1833371.62 | 1034841.94 | 1278477.25 | 897085.38 | 978839.69 | 1194764.5 | 1042191.5 | 986210.31 | 1821536.5 | 1014378.62 | 1236847.75 | 977972.94 | 1055817 | 1105580.88 | 1038799.5 |
| A2AKQ0 | Slc35d1 | UDP-glucuronic acid/UDP-N-acetylgalactosamine transporter Mus musculus | 55.91 | 5.92 | 39272 | 1 | 1 | 238848.25 | 1 | 171245.02 | 1 | 1 | 268929.31 | 1 | 1 | 183434.09 | 243597.64 | 219619.8 | 1 | 158683.8 | 268710.84 | 1 |
| Q5RKH7 | Slc35f6 | Solute carrier family 35 member F6 Rattus norvegicus | 40.68 | 4.03 | 41092 | 1 | 1 | 1099326.5 | 1 | 1 | 1 | 1 | 1 | 1 | 1 | 1 | 1 | 1 | 1 | 1 | 423392.5 | 225399.61 |
| Q9WU81 | Slc37a2 | Glucose-6-phosphate exchanger SLC37A2 Mus musculus | 160.7 | 7.78 | 55073 | 3 | 3 | 6496954.5 | 1 | 431650.62 | 267736.12 | 386036.78 | 206913.72 | 1 | 731310.81 | 1006255.25 | 1 | 282832.72 | 417963.16 | 568821.75 | 502424.72 | 1152194.25 |
| Q8CFE6 | Slc38a2 | Sodium-coupled neutral amino acid transporter 2 Mus musculus | 70.86 | 3.77 | 55503 | 2 | 2 | 3341282.5 | 1337418.12 | 1624271.88 | 886244.25 | 842715.62 | 559096.81 | 274663.62 | 881824 | 1584325 | 635062.44 | 630351.62 | 1547584.5 | 942701.19 | 1708415 | 962752.56 |
| Q6P6S2 | Slc39a11 | Zinc transporter ZIP11 Rattus norvegicus | 78.44 | 6.57 | 34868 | 1 | 1 | 398985.34 | 461895.41 | 569950.69 | 472844.44 | 411936.06 | 472239.53 | 454645.47 | 513119.81 | 444122.31 | 537376.56 | 178761.19 | 483983.81 | 576074.94 | 507140.78 | 522796.56 |
| Q78IQ7 | Slc39a4 | Zinc transporter ZIP4 Mus musculus | 201.38 | 12.73 | 71063 | 4 | 4 | 2840990.5 | 2027986.25 | 4243827 | 1403588.5 | 2391140 | 5039159.5 | 1121060.75 | 4161590.5 | 3851729.5 | 2167843.75 | 4722965 | 2513411 | 1613344.88 | 2733570 | 2325183.25 |
| Q31125 | Slc39a7 | Zinc transporter SLC39A7 Mus musculus | 136.99 | 8.19 | 50657 | 2 | 2 | 1483161.62 | 1283374.12 | 1112795.5 | 1289551.25 | 1187442.88 | 1435898.62 | 1038287.88 | 1298566.38 | 1125391 | 1083028.75 | 1393585.38 | 882471.75 | 1431080.88 | 1205987.5 | 1481592.12 |
| P10852 | Slc3a2 | 4F2 cell-surface antigen heavy chain Mus musculus | 986.12 | 53.23 | 58337 | 19 | 19 | 41247057 | 18164911 | 19587141 | 18179335 | 20064393 | 15072141 | 17644853 | 18291085 | 23555641 | 18584575 | 19004211 | 16875387 | 18805683 | 18618715 | 20505067 |
| Q8BTY2 | Slc4a7 | Sodium bicarbonate cotransporter 3 Mus musculus | 198.97 | 3.77 | 116514 | 3 | 3 | 1 | 1 | 1 | 1 | 1 | 1 | 714774 | 1 | 1 | 955521 | 1 | 1 | 1 | 1 | 564693.5 |
| Q9CXK4 | Slc50a1 | Sugar transporter SWEET1 Mus musculus | 46.08 | 9.05 | 24649 | 1 | 1 | 1643345 | 1234161.75 | 1432607.12 | 1383773 | 1731784.12 | 1235421.25 | 1139877.25 | 1010332.81 | 1566378.75 | 1235383.5 | 2716432.25 | 1275449.88 | 1635241.62 | 1529133.5 | 1675788.88 |
| O35316 | Slc6a6 | Sodium- and chloride-dependent taurine transporter Mus musculus | 81.01 | 3.06 | 69856 | 2 | 2 | 11509354 | 10552174 | 11006932 | 8452274 | 9802037 | 9152196 | 7896457 | 10624392 | 13439038 | 8537096 | 9909297 | 9509162 | 9671916 | 11276758 | 8896263 |
| Q6P9P0 | Slf2 | SMC5-SMC6 complex localization factor protein 2 Mus musculus | 36.1 | 0.55 | 143970 | 1 | 1 | 1371724 | 1059111.62 | 392924.06 | 1025785.38 | 1202280.12 | 786973.88 | 890074 | 506874.47 | 187942.3 | 1417611 | 267475.47 | 270415.28 | 209157.97 | 560455.44 | 326876.04 |
| B1ARD6 | Slfn9 | Schlafen family member 9 Mus musculus | 404.65 | 14.62 | 103992 | 8 | 8 | 5376753.5 | 4439211 | 5863270 | 3434523.5 | 6125938.5 | 5010773 | 6210240.5 | 5443562.5 | 3116176 | 6764406 | 5174344.5 | 6270549 | 6808394 | 5405727 | 6136594 |
| O54988 | Slk | STE20-like serine/threonine-protein kinase Mus musculus | 646.06 | 19.22 | 141457 | 13 | 13 | 9749107 | 8270267 | 9828851 | 15975512 | 13573023 | 10305814 | 8646175 | 10167693 | 9855183 | 10394343 | 18602777 | 11309755 | 15831060 | 10277169 | 10288204 |
| Q8CH25 | Sltm | SAFB-like transcription modulator Mus musculus | 127.99 | 2.42 | 116919 | 2 | 2 | 1 | 3062424.25 | 983633.5 | 1 | 1 | 376670.97 | 978374.81 | 332876.91 | 657069.095 | 940353.69 | 724227.38 | 1 | 1 | 1 | 799085.75 |
| O54835 | Smad9 | Mothers against decapentaplegic homolog 9 Rattus norvegicus | 79.33 | 5.53 | 48992 | 2 | 2 | 1104051.5 | 1915360.25 | 1419825.5 | 1939111.5 | 1950017.12 | 1582745.75 | 2143182 | 1531190.38 | 1740760.75 | 2211408 | 1598751.62 | 1531796.12 | 1769353.75 | 1394530.12 | 1147222.38 |
| Q9R0P4 | Smap | Small acidic protein Mus musculus | 218.45 | 21.55 | 20046 | 2 | 2 | 1115714.75 | 1380716.62 | 1308635.5 | 1145949.62 | 1572490 | 1160891.5 | 1109969.25 | 1380888 | 1250769.62 | 1048353.75 | 1394012 | 1204558.88 | 1189907.25 | 1194580.12 | 1113883.88 |
| Q91VZ6 | Smap1 | Stromal membrane-associated protein 1 Mus musculus | 41.36 | 2.73 | 47660 | 1 | 1 | 818227.31 | 1 | 1 | 1 | 1035083.69 | 880978.94 | 1188068 | 581820.88 | 958232.0175 | 1182060.25 | 1126479.375 | 1292725.62 | 1158749.38 | 941863.62 | 1112578.88 |
| Q3TKT4 | Smarca4 | Transcription activator BRG1 Mus musculus | 440.52 | 6.51 | 181427 | 8 | 8 | 3384356.75 | 6823248.5 | 5306927.5 | 5596839.5 | 3889618.25 | 4380138.5 | 4700269.5 | 4791234.5 | 6701064.5 | 6214118 | 5985911 | 8436193 | 5997263 | 9370973 | 4125935 |
| Q91ZW3 | Smarca5 | SWI/SNF-related matrix-associated actin-dependent regulator of chromatin subfamily A | 813.84 | 19.7 | 121627 | 18 | 18 | 20324881 | 20998975 | 21987059 | 23634985 | 17373133 | 19190429 | 18493125 | 23390077 | 23270143 | 18816975 | 23200657 | 22983091 | 22493825 | 20680395 | 20355505 |
| Q04692 | Smarcad1 | SWI/SNF-related matrix-associated actin-dependent regulator of chromatin subfamily A | 34.57 | 1.57 | 116451 | 1 | 1 | 497805.5 | 992761.12 | 860097.06 | 855697.5 | 975915.81 | 880184.88 | 804493.81 | 712954.06 | 779477 | 773644.69 | 699692.5 | 739208.06 | 835894.94 | 805075.5 | 762439.19 |
| B4F769 | Smarcal1 | | 37.68 | 0.99 | 101212 | 1 | 1 | 401547.94 | 351870.12 | 358495.8575 | 324722.06 | 355843.31 | 330827.47 | 556132.88 | 528519.19 | 296367.19 | 373460.25 | 312523 | 1 | 1 | 315119.19 | 306350.97 |
| Q9Z0H3 | Smarcb1 | SWI/SNF-related matrix-associated actin-dependent regulator of chromatin subfamily B | 160.83 | 7.79 | 44141 | 2 | 2 | 3590110.5 | 3535351.5 | 4380336 | 4039877 | 4316787 | 3696420.5 | 4788009 | 4481514 | 4051924.75 | 2376257.5 | 4344430.5 | 4695593 | 3882050.75 | 1975395.5 | 4561969 |
| P97496 | Smarcc1 | SWI/SNF complex subunit SMARCC1 Mus musculus | 421.07 | 9.96 | 122890 | 4 | 7 | 15891359 | 19538943 | 17350991 | 13023304 | 10092841 | 10253921 | 11849424 | 19249305 | 16038435 | 17529679 | 13151010 | 12104360 | 13598117 | 13492665 | 12990746 |
| Q6PDG5 | Smarcc2 | SWI/SNF complex subunit SMARCC2 Mus musculus | 426.14 | 7.34 | 132604 | 4 | 7 | 10748632 | 12745533 | 11604058 | 12007481 | 11015288 | 10335093 | 11994309 | 12247537 | 12234576 | 12360744 | 11432980 | 11618662 | 12500186 | 11717522 | 10987760 |
| O54772 | Smarcd2 | SWI/SNF-related matrix-associated actin-dependent regulator of chromatin subfamily D | 169.1 | 9.23 | 59128 | 3 | 3 | 1 | 1 | 1 | 1 | 229647.23 | 228183.2 | 203706.75 | 1 | 1 | 252418.69 | 1 | 176650.62 | 186843.5 | 1 | 182890.69 |

| | | | | | | | | | | | | | | | | | | | | | | |
|---|---|---|---|---|---|---|---|---|---|---|---|---|---|---|---|---|---|---|---|---|---|---|
| O54941 | Smarce1 | SWI/SNF-related matrix-associated actin-dependent regulator of chromatin subfamily E | 151.4 | 6.81 | 46638 | 2 | 2 | 1753462.12 | 2170892.75 | 1625276.88 | 1852051.75 | 1706193.12 | 1580803.88 | 1626304.25 | 1641722 | 1456211.62 | 1871260 | 1710690.25 | 1530764.5 | 1582983.38 | 1416933.25 | 1670098.5 |
| Q9CU62 | Smc1a | Structural maintenance of chromosomes protein 1A Mus musculus | 939.73 | 18.82 | 143235 | 19 | 19 | 17392211 | 15854419 | 16932417 | 13097605 | 16627759 | 13922343 | 18165167 | 12893368 | 10068940 | 14342759 | 16865935 | 15261798 | 12760360 | 13546056 | 14950707 |
| Q8CG48 | Smc2 | Structural maintenance of chromosomes protein 2 Mus musculus | 1010.75 | 28.3 | 134239 | 26 | 26 | 13232687 | 10911837 | 18553215 | 14979910 | 15309549 | 15814105 | 16345951 | 13605809 | 13185461 | 14709001 | 21411825 | 15318227 | 14956270 | 11643139 | 13810782 |
| Q9CW03 | Smc3 | Structural maintenance of chromosomes protein 3 Mus musculus | 1357.97 | 28.02 | 141556 | 26 | 26 | 20230691 | 25036955 | 22632877 | 21715167 | 26981207 | 18700469 | 27582895 | 23303909 | 23447939 | 25359165 | 20663597 | 24897071 | 22886365 | 24417751 | 27315787 |
| Q8CG47 | Smc4 | Structural maintenance of chromosomes protein 4 Mus musculus | 1030.89 | 23.64 | 146895 | 22 | 22 | 11295095 | 13646517 | 16613988 | 17815375 | 16215852 | 17869107 | 17799831 | 15435929 | 12166426 | 18673385 | 15423082 | 17703205 | 16597849 | 15626738 | 17644709 |
| Q6P5D8 | Smchd1 | Structural maintenance of chromosomes flexible hinge domain-containing protein 1 Mus | 992.29 | 14.25 | 225648 | 18 | 18 | 7834954 | 11752611 | 10098537 | 10123781 | 8980470 | 9506679 | 9917977 | 9781662 | 11907380 | 10034468 | 10636811 | 10014218 | 8086303.5 | 11097412 | 7991707 |
| P58242 | Smpdl3b | Acid sphingomyelinase-like phosphodiesterase 3b Mus musculus | 269.42 | 18.64 | 51600 | 6 | 6 | 9478891 | 10678620 | 8368701 | 9966309 | 10576836 | 9799435 | 7365186 | 8832467 | 9138325 | 8167567 | 11375455 | 7512871.5 | 10695841 | 10698709 | 7413641 |
| P97355 | Sms | Spermine synthase Mus musculus | 332.14 | 28.96 | 41313 | 6 | 6 | 3529398.5 | 4052233.5 | 3624725.75 | 3494258.5 | 3224708.75 | 3609729.75 | 2850677.75 | 3752108.5 | 3769451.5 | 3980508 | 4003495 | 2960100 | 3473872.25 | 4381327.5 | 3016635.25 |
| Q3UKJ7 | Smu1 | WD40 repeat-containing protein SMU1 Mus musculus | 420.49 | 24.76 | 57544 | 9 | 9 | 5199344.5 | 3802362.75 | 4249785 | 5334716.5 | 5519688 | 3754138.5 | 9670383 | 3848529.5 | 3094762.5 | 8450751 | 3747299.5 | 3790369.75 | 5246956.5 | 4954863 | 4881902.5 |
| Q3TYX3 | Smyd5 | SET and MYND domain-containing protein 5 Mus musculus | 163.18 | 9.13 | 47095 | 3 | 3 | 280062.72 | 286163.16 | 144167.31 | 302051.06 | 416204.91 | 193432.28 | 279755 | 174116.12 | 107702.05 | 280062.72 | 166931.42 | 1 | 177169.7 | 166567.06 | 1 |
| O09044 | Snap23 | Synaptosomal-associated protein 23 Mus musculus | 89.9 | 6.67 | 23261 | 1 | 1 | 2179954.25 | 2663049.25 | 2291606.75 | 2343632.25 | 1942696.5 | 2121699.25 | 1809910.25 | 2704765.25 | 2287853.5 | 2034623.75 | 1416420.25 | 1816306.25 | 2151266.25 | 1559292.38 | 2093848.62 |
| Q9ERB0 | Snap29 | Synaptosomal-associated protein 29 Mus musculus | 43.16 | 4.23 | 29572 | 1 | 1 | 579252.31 | 476272.28 | 387013.06 | 668182.69 | 526233.5 | 568789.5 | 658646 | 519201.88 | 407108.59 | 471609.62 | 339853.62 | 533209.31 | 562031.88 | 502219.5 | 391313.09 |
| Q78PY7 | Snd1 | Staphylococcal nuclease domain-containing protein 1 Mus musculus | 2243.23 | 53.52 | 102088 | 38 | 38 | 97090297 | 95015009 | 104223953 | 101757113 | 98515169 | 98656353 | 115381873 | 97613633 | 98297313 | 111966649 | 99908673 | 98704009 | 98980921 | 93169625 | 100185441 |
| Q5RK19 | Snf8 | Vacuolar-sorting protein SNF8 Rattus norvegicus | 42.99 | 5.43 | 28884 | 1 | 1 | 789109.94 | 790146.25 | 834083.19 | 924989.38 | 875019.69 | 855687.81 | 1011078.25 | 925146.31 | 990464.62 | 1154387.12 | 1024107.56 | 1008168.12 | 940731.44 | 1049158 | 758774.81 |
| F1LNJ2 | Snrnp200 | U5 small nuclear ribonucleoprotein 200 kDa helicase Rattus norvegicus | 2586.93 | 31.09 | 244875 | 48 | 48 | 60487281 | 58840725 | 58822573 | 57602289 | 63683121 | 59416369 | 60649101 | 58118453 | 59599001 | 61317609 | 68487897 | 56173773 | 61217337 | 56299381 | 66872285 |
| Q6PE01 | Snrnp40 | U5 small nuclear ribonucleoprotein 40 kDa protein Mus musculus | 382.16 | 32.4 | 39276 | 5 | 5 | 1 | 1 | 1 | 1032575.38 | 1487945.12 | 1533760.5 | 1082958.88 | 1444757.25 | 937682.38 | 874475.88 | 1220282.5 | 1192435.12 | 1305440.75 | 989627.19 | 271608.22 |
| Q62376 | Snrnp70 | U1 small nuclear ribonucleoprotein 70 kDa Mus musculus | 408.36 | 17.86 | 51992 | 7 | 7 | 16101511 | 13816689 | 11579233 | 17936449 | 16424201 | 15368370 | 20124403 | 15401248 | 14705649 | 19778775 | 11211165 | 13900322 | 14366908 | 16992095 | 20386789 |
| Q62189 | Snrpa | U1 small nuclear ribonucleoprotein A Mus musculus | 295.14 | 31.71 | 31835 | 5 | 6 | 5356152.5 | 5656992.5 | 5636798.5 | 5084548 | 5502430 | 4659304.5 | 6160083 | 4622001.5 | 4284844 | 6169961.5 | 4777405.5 | 3970186.5 | 5356026 | 4360877.5 | 3275023.75 |
| P57784 | Snrpa1 | U2 small nuclear ribonucleoprotein A' Mus musculus | 577.45 | 45.88 | 28357 | 11 | 11 | 11704108 | 11399720 | 13774256 | 15603991 | 11045075 | 13217827 | 9615221 | 10708041 | 9503952 | 8811258 | 12770319 | 13404275 | 13147361 | 14974173 | 11273036 |
| P17136 | Snrpb | Small nuclear ribonucleoprotein-associated protein B Rattus norvegicus | 181.4 | 12.99 | 23656 | 3 | 3 | 6944501.5 | 3822040 | 5843892.5 | 7536809 | 5309032.5 | 4953571.5 | 9317841 | 4222938 | 4253762.5 | 7821286.5 | 6873010.5 | 6914666 | 5548760.5 | 4616082 | 7386406 |
| Q9CQI7 | Snrpb2 | U2 small nuclear ribonucleoprotein B'' Mus musculus | 187.01 | 20 | 25323 | 3 | 4 | 1173842.485 | 341499.44 | 337097.5 | 2369642 | 1647131 | 241212.42 | 2256475.25 | 1944115.25 | 980722.62 | 2530317.25 | 1937613.38 | 1803015.75 | 2059599.38 | 1918240.408 | 1872733.12 |
| D3ZCL3 | Snrpc | U1 small nuclear ribonucleoprotein C Rattus norvegicus | 75.42 | 7.55 | 17364 | 1 | 1 | 1886324.38 | 2063909.12 | 1915900.5 | 1998641.38 | 2116162.25 | 1994514.12 | 1863694.62 | 2002316.38 | 2240450 | 1818880 | 2087636.75 | 1729565.88 | 2322834.75 | 1901511.62 | 1747779.88 |
| P62315 | Snrpd1 | Small nuclear ribonucleoprotein Sm D1 Mus musculus | 246.73 | 54.62 | 13282 | 5 | 5 | 2063207 | 1520507.5 | 1185624.62 | 2076748.62 | 2390305.5 | 1664916.62 | 2782973.25 | 2007178.53 | 918802.75 | 2662021.5 | 1339019.75 | 989558.69 | 1932338.62 | 129753.21 | 135787.48 |
| P62317 | Snrpd2 | Small nuclear ribonucleoprotein Sm D2 Mus musculus | 221.84 | 35.59 | 13527 | 5 | 5 | 2880013.25 | 1414651.88 | 1495560 | 1183361.5 | 1269746 | 482199.47 | 4523459 | 1433560.62 | 413307.53 | 4065708.5 | 1226563.38 | 1232116.12 | 1188984.75 | 1841466.75 | 1627234.12 |
| P62320 | Snrpd3 | Small nuclear ribonucleoprotein Sm D3 Mus musculus | 88.47 | 15.08 | 13916 | 2 | 2 | 25269827 | 26480363 | 28734623 | 31722185 | 32007723 | 28900891 | 30066753 | 26421551 | 27757581 | 32257915 | 33056409 | 29996619 | 29876153 | 28967365 | 30615945 |
| P62305 | Snrpe | Small nuclear ribonucleoprotein E Mus musculus | 181.2 | 52.17 | 10804 | 3 | 4 | 24198429 | 32978761 | 34201445 | 29037191 | 30816033 | 32637403 | 31663135 | 32239501 | 34067849 | 34718553 | 24157711 | 32276881 | 32236641 | 27680299 | 21185729 |
| P62307 | Snrpf | Small nuclear ribonucleoprotein F Mus musculus | 229.71 | 83.72 | 9725 | 4 | 4 | 1863000 | 1003075.06 | 1714080 | 1687725.12 | 2597979.25 | 1847317.38 | 2266229.75 | 1715296.38 | 1498658.12 | 2543073.75 | 4344976 | 3024307.25 | 2534152 | 1451341.12 | 3997320.5 |
| Q61235 | Sntb2 | Beta-2-syntrophin Mus musculus | 71.99 | 1.73 | 56382 | 1 | 1 | 449934.72 | 439862.88 | 453626.09 | 498403.125 | 650188.81 | 517621.22 | 595283.94 | 543435.31 | 521238.44 | 564463.69 | 541896.5 | 489026.12 | 572175.25 | 395357.53 | 588093.5 |
| P55770 | Snu13 | NHP2-like protein 1 Rattus norvegicus | 235.11 | 55.47 | 14174 | 5 | 5 | 13444588 | 11597464 | 13798339 | 15383133 | 16392825 | 14155240 | 16461852 | 12113783 | 10953752 | 16297507 | 16616823 | 14767637 | 14972281 | 12426310 | 16897849 |
| Q9CSN1 | Snw1 | SNW domain-containing protein 1 Mus musculus | 345.85 | 15.11 | 61475 | 6 | 6 | 3172287.75 | 1872244.62 | 3028839.5 | 1570786.62 | 4597744 | 2365052.75 | 3503774.5 | 4188838.25 | 2645066.25 | 4701543 | 4095271.5 | 5957777 | 3685204.5 | 3033974 | 4211589 |
| Q99N27 | Snx1 | Sorting nexin-1 Rattus norvegicus | 741.77 | 30.65 | 59044 | 13 | 14 | 27401709 | 34500917 | 28031507 | 29407069 | 30222635 | 26990603 | 28737413 | 27796159 | 27752361 | 26914331 | 28999425 | 29811621 | 29440631 | 28691465 | 28774131 |
| O70493 | Snx12 | Sorting nexin-12 Mus musculus | 245.49 | 44.24 | 19116 | 4 | 5 | 7651322 | 6783016 | 6796568.5 | 5799132 | 7286726 | 7115620.5 | 7444176.5 | 6557141 | 6684842 | 6585914 | 3428679 | 7972359 | 6704743 | 5956166.5 | 5977960 |
| Q8BVL3 | Snx17 | Sorting nexin-17 Mus musculus | 151.8 | 13.62 | 52797 | 4 | 4 | 1532372 | 157143.75 | 918231.44 | 1128782.25 | 934132.36 | 651040.75 | 1136710.5 | 1 | 1 | 460132.5 | 702606.62 | 849984.38 | 642142.0125 | 890827.5 | 125149.55 |
| Q91ZR2 | Snx18 | Sorting nexin-18 Mus musculus | 131.78 | 6.35 | 67904 | 3 | 3 | 5560716.5 | 3640845.5 | 2257478.5 | 3922595.5 | 2254448 | 4546767 | 4577057 | 3631256 | 3827787 | 4530150 | 2706033.25 | 3958979.75 | 3542400.25 | 4056257.25 | 2295318.25 |
| Q9CWK8 | Snx2 | Sorting nexin-2 Mus musculus | 1058.94 | 48.36 | 58471 | 17 | 18 | 42139717 | 39019877 | 42667173 | 42373893 | 37328149 | 33036595 | 39527005 | 35774001 | 35753597 | 37002513 | 39356345 | 40050249 | 36835505 | 37219633 | 45065821 |
| Q5BK61 | Snx20 | Sorting nexin-20 Rattus norvegicus | 45.89 | 3.83 | 35723 | 1 | 1 | 781292.69 | 719573.88 | 749767.56 | 606141.81 | 766262.44 | 501851.78 | 613557.06 | 646920.75 | 816417.94 | 751198.75 | 555807.88 | 516052.69 | 586433 | 729454.69 | 591070.69 |
| Q3UHD6 | Snx27 | Sorting nexin-27 Mus musculus | 160.06 | 9.28 | 60989 | 3 | 3 | 1525009.533 | 1829758.5 | 187545 | 1828063.88 | 2254670.75 | 1 | 1711457.12 | 1 | 1 | 342474.53 | 356925.91 | 329185.94 | 226029.92 | 1 | 1 |
| Q9D3S3 | Snx29 | Sorting nexin-29 Mus musculus | 36.24 | 1.59 | 91387 | 1 | 1 | 258800.97 | 299483.16 | 236869.2 | 275548.41 | 215580.48 | 1 | 1 | 209849.48 | 1 | 318428.94 | 245953.83 | 297834.56 | 234013.33 | 260143.46 | 262772.12 |
| O70492 | Snx3 | Sorting nexin-3 Mus musculus | 283.42 | 48.15 | 18757 | 5 | 6 | 10380674 | 7210111.5 | 11032516 | 8297273.5 | 7993428 | 7922807 | 10022926 | 9591908 | 8885324 | 7614498 | 7474445.5 | 9355642 | 7680692 | 6775138 | 10080302 |
| Q8CE50 | Snx30 | Sorting nexin-30 Mus musculus | 44.73 | 4.35 | 49520 | 1 | 1 | 210270.38 | 292144.66 | 360198.59 | 512337.06 | 174994.08 | 386761.34 | 281801.06 | 285117.97 | 235457.56 | 339589.78 | 280059.25 | 333956.22 | 296077.72 | 326601.5 | 505373.91 |
| Q91YJ2 | Snx4 | Sorting nexin-4 Mus musculus | 69.29 | 2 | 51778 | 1 | 1 | 1887102.38 | 1511936.88 | 1589528.12 | 1633072.62 | 2072632.62 | 1750982.62 | 2174979 | 2049668.25 | 1845501.12 | 2065352.75 | 1891318.12 | 1816688.12 | 1867424.75 | 1802289.12 | 2147183 |
| Q9D8U8 | Snx5 | Sorting nexin-5 Mus musculus | 926.58 | 45.3 | 46797 | 18 | 18 | 50526221 | 47983113 | 50641529 | 45504009 | 45378405 | 40410321 | 48343549 | 45185685 | 49140021 | 48275905 | 46107381 | 55010561 | 48297721 | 52694105 | 51454677 |
| Q6P8X1 | Snx6 | Sorting nexin-6 Mus musculus | 205.46 | 15.02 | 46649 | 5 | 5 | 2909167.25 | 4147249.5 | 2350855.5 | 1733028.88 | 2891085.5 | 2868612 | 2731646.5 | 3512420.5 | 3880251 | 3027227.25 | 3202175.75 | 3454020.5 | 3074361.75 | 2576280.75 | 4050953 |
| Q9CY18 | Snx7 | Sorting nexin-7 Mus musculus | 41.25 | 3.1 | 45000 | 1 | 1 | 1 | 1220634.25 | 1 | 938071.06 | 1293163.12 | 1310768 | 845451 | 1304121.62 | 1421255.75 | 1019196.69 | 1286056.75 | 1199372.62 | 1022391.12 | 1010883.19 | 1012606.31 |
| Q8CFD4 | Snx8 | Sorting nexin-8 Mus musculus | 103.54 | 7.19 | 52059 | 2 | 2 | 301617.22 | 1 | 745870.19 | 1 | 1 | 1 | 187287.31 | 451999.19 | 692556 | 1 | 510891.94 | 441919.84 | 399621.5325 | 359726.38 | 285947.97 |
| Q91VH2 | Snx9 | Sorting nexin-9 Mus musculus | 103.12 | 7.73 | 66546 | 2 | 2 | 1294086.12 | 870328.25 | 1544029 | 670678.25 | 1034551 | 1383439.25 | 831367.06 | 1853200.38 | 2521905.25 | 518163.31 | 1600277.75 | 1759337.75 | 214338.11 | 1969890.25 | 1494574.88 |
| Q61263 | Soat1 | Sterol O-acyltransferase 1 Mus musculus | 297.96 | 13.7 | 63799 | 5 | 5 | 7986168 | 791642.75 | 2632665.5 | 2175211.5 | 1571731.5 | 2217528.25 | 4770988.5 | 1643008.75 | 2062320.62 | 3058683.25 | 2788912 | 2941640 | 1249419.12 | 2011052.62 | 4367954.5 |
| P08228 | Sod1 | Superoxide dismutase [Cu-Zn] Mus musculus | 269.37 | 32.47 | 15943 | 4 | 4 | 34171897 | 36139077 | 31668321 | 38968433 | 37639953 | 37189365 | 29335247 | 27587449 | 32672757 | 27979851 | 36082993 | 29931549 | 32774123 | 32032001 | 25136185 |
| P09671 | Sod2 | Superoxide dismutase [Mn], mitochondrial Mus musculus | 292.17 | 41.89 | 24603 | 5 | 5 | 24467249 | 28106105 | 30527805 | 22500363 | 25449799 | 28424509 | 36324853 | 28395011 | 27370107 | 30738281 | 24654713 | 27579045 | 31217657 | 25506093 | 26575503 |
| Q9QX47 | Son | Protein SON Mus musculus | 301.94 | 3.48 | 265651 | 5 | 5 | 6658958 | 6934572 | 5505999 | 6920907 | 7817023.5 | 6954433.5 | 5251586.5 | 8216305 | 9287555 | 7679912 | 7526965.5 | 6246482.5 | 7541070 | 7259849.5 | 7590765 |
| O35892 | Sp100 | Nuclear autoantigen Sp-100 Mus musculus | 189.49 | 9.96 | 54727 | 3 | 4 | 3439218.5 | 3207927 | 3216481.25 | 2612204 | 2956948.5 | 2875577.25 | 2323306.75 | 3101980.5 | 3566695.5 | 2169429.5 | 2708727 | 2768655.5 | 2335071 | 3295967.5 | 2218676.75 |
| Q8BVK9 | Sp110 | Sp110 nuclear body protein Mus musculus | 472.34 | 32.58 | 50140 | 10 | 10 | 9270450 | 7255536 | 8656968 | 7135867.5 | 8281462.5 | 7427609.5 | 9096913 | 8777348 | 8347409 | 8369341.5 | 6370892 | 7174590 | 7467874 | 7653144 | 7089843.5 |
| Q7TNE3 | Spag7 | Sperm-associated antigen 7 Mus musculus | 88.28 | 15.86 | 25921 | 2 | 2 | 615179.38 | 506796 | 641619.44 | 668591.81 | 904419.81 | 806096.69 | 855170.44 | 846231.7975 | 563813.44 | 1159846.62 | 853530.88 | 1 | 1186548.38 | 591864.88 | 1 |
| Q58A65 | Spag9 | C-Jun-amino-terminal kinase-interacting protein 4 Mus musculus | 403.93 | 9.01 | 146219 | 7 | 7 | 4373358.5 | 4123626 | 3886400 | 3164691 | 5128186 | 5395138.5 | 5981896 | 5632487.5 | 6745488 | 5882954 | 4880454 | 5267936.5 | 4656879 | 5219346.5 | 6306012.5 |
| Q8R1X6 | Spart | Spartin Mus musculus | 64.11 | 1.94 | 72655 | 1 | 1 | 678906.94 | 1005219.44 | 895302.44 | 923381.62 | 997698.25 | 828276.75 | 862186.31 | 815806.75 | 1153732.25 | 915000.515 | 1288291.5 | 762818.31 | 1013422 | 964442.81 | 780472.94 |
| B2RYN7 | Spast | Spastin Rattus norvegicus | 79.52 | 6.88 | 63022 | 2 | 2 | 533605.88 | 178711.44 | 314964.69 | 300102.91 | 403308.5 | 383760.47 | 634064.56 | 147850.72 | 456057.0625 | 658552.5 | 191973 | 298073.66 | 409075.5575 | 443500.88 | 702754.69 |
| Q0P557 | Spata18 | Mitochondria-eating protein Mus musculus | 40.01 | 1.49 | 60398 | 1 | 1 | 1319964.88 | 1498278.88 | 1609656 | 1620784.62 | 1431329 | 1818972.75 | 1199232.75 | 1450689.5 | 1219126.62 | 1152685.12 | 1527523.25 | 1197690.25 | 1408040.12 | 1236688.62 | 1232168.38 |
| Q9D083 | Spc24 | Kinetochore protein Spc24 Mus musculus | 155.91 | 21.89 | 23416 | 3 | 3 | 2456834.5 | 2157392 | 1952820.75 | 1715648.5 | 1469050.62 | 2080853 | 1713120.62 | 2191882 | 2391509 | 1082765.25 | 2300918.5 | 1889560.75 | 2007611.25 | 2046501.25 | 1628385.75 |
| Q3UA16 | Spc25 | Kinetochore protein Spc25 Mus musculus | 43.59 | 4.87 | 26457 | 1 | 1 | 871541.06 | 1022696.31 | 983653.56 | 1 | 1 | 680363.06 | 747459.06 | 894685.44 | 906906.12 | 639484.75 | 1 | 863997.69 | 818437.31 | 1025398.06 | 1 |
| Q9CYN2 | Spcs2 | Signal peptidase complex subunit 2 Mus musculus | 180.34 | 19.47 | 24978 | 4 | 4 | 2645786.75 | 1444518.75 | 2136541 | 1882667.62 | 1688463.88 | 2457252.5 | 1879292 | 2169125.75 | 2660976 | 2823115.75 | 1357379.5 | 1305659.25 | 1373662.12 | 2244487.75 | 2823982.5 |
| Q6ZWQ7 | Spcs3 | Signal peptidase complex subunit 3 Mus musculus | 177.23 | 21.67 | 20313 | 4 | 4 | 5424392.5 | 4938548 | 4468089 | 5067779.5 | 4013489.25 | 5586489.5 | 6127951 | 5763785.5 | 4470409 | 6896460.5 | 5932688 | 5307993.5 | 7108939.5 | 5921153.5 | 5555923 |
| Q5SXY1 | Specc1 | Cytospin-B Mus musculus | 51.34 | 0.75 | 118101 | 1 | 1 | 891141.69 | 461935.31 | 915271.12 | 938618.12 | 619028.31 | 641131.06 | 834230.69 | 751493.31 | 510935.91 | 872695.44 | 808946.44 | 718785.44 | 750268.5 | 758575.81 | 830860.88 |
| Q9JIA7 | Sphk2 | Sphingosine kinase 2 Mus musculus | 244.92 | 8.75 | 65618 | 4 | 4 | 3011618 | 3209972.75 | 3824089.25 | 3107739.5 | 2815345.25 | 3570180.75 | 3794087.5 | 3460189 | 2807926 | 3841815.25 | 2873797.25 | 3249709 | 3635611.25 | 2676648.5 | 3472879.5 |
| P17433 | Spi1 | Transcription factor PU.1 Mus musculus | 74.39 | 3.68 | 31349 | 1 | 1 | 860899.19 | 1514500.38 | 1234576.88 | 1313593.12 | 975769.31 | 1174973.88 | 1645928.12 | 1216650.75 | 1132840.12 | 1430092.38 | 2053088.62 | 1258194.12 | 1185470.12 | 1116332.62 | 1401185.25 |
| Q2YDU8 | Spns1 | Protein spinster homolog 1 Rattus norvegicus | 74.92 | 5.11 | 56796 | 1 | 1 | 1171658.5 | 1 | 739291.38 | 1 | 1 | 494769.07 | 434278.47 | 659957.12 | 192047.19 | 692793.5 | 733115.31 | 436933 | 948846.69 | 1 | 828618.56 |
| Q3UHX9 | Spout1 | Putative methyltransferase C9orf114 homolog Mus musculus | 38.88 | 3.12 | 42958 | 1 | 1 | 1 | 1 | 1 | 255840.88 | 198152.83 | 202722.91 | 183330.41 | 203758.58 | 207185.85 | 238931.5 | 207845.6975 | 253468.91 | 204644.19 | 183035.05 | 190234.64 |
| Q64105 | Spr | Sepiapterin reductase Mus musculus | 732.36 | 58.62 | 27883 | 11 | 11 | 20878269 | 23049057 | 24121913 | 26636331 | 23832011 | 22420449 | 26260093 | 19813623 | 17014745 | 21211791 | 25814971 | 21240213 | 27625725 | 24645077 | 24894549 |
| O88838 | Spsb2 | SPRY domain-containing SOCS box protein 2 Mus musculus | 35.47 | 2.65 | 28938 | 1 | 1 | 9523748 | 11776717 | 10869080 | 11468616 | 11094955 | 8139882 | 7965588.5 | 9188951 | 10026194 | 7675419 | 9446949 | 8060028 | 9073398 | 9557168 | 6493642.5 |
| P16086 | Sptan1 | Spectrin alpha chain, non-erythrocytic 1 Rattus norvegicus | 117.82 | 1.5 | 284637 | 2 | 2 | 360823.97 | 1043366.38 | 1750962.62 | 868981.36 | 320772.47 | 1549231 | 572942.06 | 2070602 | 2065275.12 | 1564512.545 | 577492.44 | 1821237.12 | 338633.78 | 614595.56 | 695723.19 |
| Q62261 | Sptbn1 | Spectrin beta chain, non-erythrocytic 1 Mus musculus | 195.59 | 2.54 | 274223 | 5 | 5 | 6039017 | 5244930 | 4114364.25 | 5571106.5 | 5208593.5 | 5353301 | 6212628 | 4807033.5 | 5122075.5 | 5337650 | 4469254 | 4163230.25 | 4761831 | 4625840.5 | 3981463.25 |
| O35704 | Sptlc1 | Serine palmitoyltransferase 1 Mus musculus | 160.62 | 6.13 | 52535 | 2 | 2 | 101991.68 | 467272.62 | 497187.44 | 432215.12 | 353916.41 | 490708.06 | 418573.09 | 273211.62 | 348416.03 | 599544.56 | 282055.31 | 589391.69 | 660751.25 | 1 | 223307.06 |
| P97363 | Sptlc2 | Serine palmitoyltransferase 2 Mus musculus | 89.65 | 6.79 | 62982 | 2 | 2 | 719878.12 | 1 | 1 | 1 | 1 | 166500.83 | 1 | 173508.3 | 1 | 1 | 364157.84 | 1 | 1 | 144940.17 | 1 |
| Q9R112 | Sqor | Sulfide:quinone oxidoreductase, mitochondrial Mus musculus | 758.22 | 35.11 | 50282 | 12 | 12 | 26959017 | 29870575 | 30165297 | 28299773 | 34691545 | 37028809 | 37249385 | 36522933 | 39867093 | 30743109 | 24611925 | 25713689 | 31846289 | 29247759 | 23953697 |
| Q64337 | Sqstm1 | Sequestosome-1 Mus musculus | 493.06 | 34.62 | 48163 | 8 | 10 | 9116029 | 7441562 | 8011082 | 6849564 | 4125496.5 | 7728917 | 6729773.5 | 5223859.5 | 7806434 | 6268631 | 8005020.5 | 7576888 | 5112239 | 7216354 | 8240949.5 |
| Q497V5 | Srbd1 | S1 RNA-binding domain-containing protein 1 Mus musculus | 286.78 | 9.67 | 110281 | 7 | 7 | 34034501 | 33414335 | 41727673 | 34932977 | 36857381 | 36849473 | 54394209 | 32874035 | 31674947 | 45106433 | 42786917 | 40655801 | 39871177 | 30967531 | 50341033 |
| D4A208 | Srgap2 | SLIT-ROBO Rho GTPase-activating protein 2 Rattus norvegicus | 536.55 | 12.04 | 120878 | 11 | 11 | 11569112 | 10469069 | 10177344 | 8966176 | 11575758 | 11621720 | 11192711 | 8807416 | 11039589 | 8122678.5 | 11170649 | 10440967 | 10534066 | 11615356 | 10947200 |
| Q812A2 | Srgap3 | SLIT-ROBO Rho GTPase-activating protein 3 Mus musculus | 77.63 | 1.09 | 124420 | 1 | 1 | 402724.81 | 569853.25 | 428571.34 | 488392.74 | 552421.56 | 557970.12 | 548108.12 | 657049.75 | 562640.81 | 401294.84 | 590623.88 | 492646.03 | 430247.47 | 571477.81 | 479149.22 |
| Q6P069 | Sri | Sorcin Mus musculus | 230.69 | 26.26 | 21627 | 5 | 5 | 6887902.5 | 8405801 | 7203348 | 8216794 | 8774899 | 7709453.5 | 8312001 | 7639512 | 8626642 | 8115344.5 | 8212954.5 | 6927790.5 | 8209790.5 | 7124886.5 | 7454649 |
| Q64674 | Srm | Spermidine synthase Mus musculus | 482.09 | 44.04 | 33995 | 10 | 10 | 10481149 | 15168779 | 11882889 | 14265563 | 16794055 | 14469739 | 14798526 | 13231351 | 14867361 | 17556669 | 15830995 | 15751329 | 15174613 | 14457142 | 18984675 |
| P16254 | Srp14 | Signal recognition particle 14 kDa protein Mus musculus | 157.1 | 33.64 | 12510 | 3 | 3 | 2109382.5 | 2303947.75 | 1127622.5 | 1105371.38 | 1661581.033 | 532284.5 | 556388 | 2024434.5 | 1865564.62 | 616516.94 | 1497073.88 | 1328739 | 1780868.75 | 1 | 1 |
| Q9D7A6 | Srp19 | Signal recognition particle 19 kDa protein Mus musculus | 108.91 | 25.69 | 16191 | 2 | 2 | 1186770.88 | 1212089.62 | 1358957.625 | 1583497.5 | 1453472.5 | 1143791.5 | 1319021 | 1660507.5 | 694376.12 | 1680315.38 | 1265413.12 | 632922.12 | 1415818.38 | 912731.12 | 1334309.25 |
| P14576 | Srp54 | Signal recognition particle 54 kDa protein Mus musculus | 252.93 | 15.67 | 55721 | 6 | 6 | 4842025.5 | 4733749 | 4221085.5 | 5124884 | 2779329.5 | 3824778.75 | 3265858 | 4832878 | 4855245 | 5805780 | 4307663 | 3889518.25 | 4039866 | 3178114 | 3330753.25 |
| Q8BMA6 | Srp68 | Signal recognition particle subunit SRP68 Mus musculus | 758.24 | 24 | 70574 | 12 | 12 | 8877004 | 9019523 | 9309933 | 9427344 | 8799103 | 7988861 | 7355755.5 | 9801037 | 11187711 | 8442404 | 7482645 | 9892884 | 8328576 | 10059029 | 8249697 |
| P49962 | Srp9 | Signal recognition particle 9 kDa protein Mus musculus | 77.08 | 25.58 | 10194 | 2 | 2 | 2654223 | 2354702.25 | 2011528.75 | 2710504.5 | 2749296.5 | 2827648 | 1972418.5 | 2157164 | 2124534.5 | 2188113 | 1927274.62 | 2538062.5 | 2257080 | 2605097 | 1747251.75 |
| O70551 | Srpk1 | SRSF protein kinase 1 Mus musculus | 500.77 | 19.44 | 73088 | 9 | 9 | 9392259 | 4168226.25 | 2440486.5 | 3807400.5 | 3431408 | 7392386 | 7162449.5 | 3607439.5 | 5403236.5 | 4271728 | 5242731.5 | 5123117 | 2521372 | 3130018 | 5004597 |
| Q9DBG7 | Srpra | Signal recognition particle receptor subunit alpha Mus musculus | 906.16 | 32.55 | 69623 | 14 | 14 | 14421949 | 13493169 | 12227201 | 20366015 | 15431319 | 16775071 | 15326319 | 22198569 | 13960493 | 17536191 | 15432146 | 18614519 | 13548887 | 21345041 | 16554725 |
| P47758 | Srprb | Signal recognition particle receptor subunit beta Mus musculus | 350.86 | 32.34 | 29579 | 7 | 7 | 1671979.75 | 4487028.5 | 3294738.25 | 1963707.12 | 1942231.12 | 4093212.25 | 4079337 | 1848443.5 | 1159354.62 | 4798218.5 | 2110333.5 | 3503961.5 | 2102975.25 | 1953313.5 | 3779778.75 |
| Q52KI8 | Srrm1 | Serine/arginine repetitive matrix protein 1 Mus musculus | 417.81 | 10.25 | 106862 | 7 | 7 | 4782300 | 5934505 | 7111439 | 6608492 | 6399699 | 9298970 | 8526461 | 5678670 | 5430090 | 8221419.5 | 7197931.5 | 6326018.5 | 7616430 | 6631429.5 | 8591877 |
| Q8BTI8 | Srrm2 | Serine/arginine repetitive matrix protein 2 Mus musculus | 514.83 | 6.51 | 294840 | 11 | 11 | 4423304.5 | 5107881.5 | 4832537 | 4396774 | 5783462.5 | 4384941.5 | 6452567 | 3747607.5 | 2612023.5 | 6066971 | 6230616 | 4131170 | 4899481 | 4474772.5 | 5609537 |
| Q99MR6 | Srrt | Serrate RNA effector molecule homolog Mus musculus | 568.12 | 19.09 | 100452 | 12 | 12 | 13330740 | 11962635 | 18398007 | 19085989 | 19595179 | 17454169 | 24222509 | 15635674 | 14457018 | 22528735 | 23228485 | 20243099 | 20051607 | 14582552 | 23158963 |
| Q6PDM2 | Srsf1 | Serine/arginine-rich splicing factor 1 Mus musculus | 549.92 | 37.5 | 27745 | 11 | 11 | 32474185 | 35429337 | 29109567 | 40848225 | 45081749 | 40261313 | 31053311 | 31209615 | 32287263 | 38936549 | 39893637 | 33566649 | 36997733 | 32607827 | 38751793 |
| Q9R0U0 | Srsf10 | Serine/arginine-rich splicing factor 10 Mus musculus | 316.32 | 23.28 | 31301 | 5 | 5 | 10842501 | 12178756 | 12920575 | 5026208 | 9880966 | 10106074 | 9160821 | 11216879 | 13434230 | 10211701 | 8861585 | 10404478 | 10018828 | 11815891 | 12046615 |
| Q62093 | Srsf2 | Serine/arginine-rich splicing factor 2 Mus musculus | 361.38 | 25.34 | 25476 | 5 | 5 | 14396855 | 13050013 | 14028079 | 16717478 | 16280948 | 13613581 | 22785755 | 12026737 | 11926190 | 19975673 | 17230823 | 18510477 | 16751402 | 12548917 | 18804525 |
| P84104 | Srsf3 | Serine/arginine-rich splicing factor 3 Mus musculus | 264.33 | 36.59 | 19330 | 5 | 6 | 30546295 | 35529941 | 32581633 | 41087673 | 38271949 | 32452187 | 35424729 | 29909785 | 28924671 | 38302485 | 37581481 | 33929489 | 38674505 | 34279997 | 36351977 |
| Q8VE97 | Srsf4 | Serine/arginine-rich splicing factor 4 Mus musculus | 302.3 | 13.29 | 55979 | 4 | 6 | 33271275 | 35565693 | 32026681 | 37520329 | 38076009 | 32410385 | 36387457 | 30215109 | 33392647 | 30811461 | 39838681 | 35401213 | 34030961 | 31322795 | 34931753 |
| O35326 | Srsf5 | Serine/arginine-rich splicing factor 5 Mus musculus | 351.71 | 25.28 | 30891 | 5 | 6 | 46996529 | 46404385 | 42676233 | 45301057 | 48428285 | 44589705 | 46142745 | 36188825 | 44134465 | 42469053 | 48416625 | 46624313 | 44848953 | 42497081 | 45752241 |
| G3V6S8 | Srsf6 | Serine/arginine-rich splicing factor 6 Rattus norvegicus | 448.16 | 24.78 | 39025 | 6 | 8 | 42628281 | 43954549 | 39842809 | 43961929 | 45972813 | 40950273 | 43017729 | 34712901 | 39242241 | 37338029 | 46556217 | 42892633 | 41991909 | 40144917 | 41434697 |
| Q8BL97 | Srsf7 | Serine/arginine-rich splicing factor 7 Mus musculus | 234.32 | 18.73 | 30818 | 4 | 5 | 11821619 | 11745668 | 12058998 | 17130155 | 15611909 | 14931284 | 16769953 | 12360847 | 11743849 | 18898549 | 16388515 | 14872394 | 15623872 | 11738557 | 17736361 |
| P32067 | Ssb | Lupus La protein homolog Mus musculus | 635.85 | 26.51 | 47756 | 11 | 11 | 12495275 | 10936265 | 11841657 | 11789481 | 11767524 | 11731943 | 12986463 | 11721502 | 11424622 | 10870523 | 13397356 | 12378793 | 12014759 | 12165197 | 14549800 |
| Q9CYR0 | Ssbp1 | Single-stranded DNA-binding protein, mitochondrial Mus musculus | 161.08 | 30.92 | 17319 | 4 | 4 | 12816508 | 15230558 | 12364125 | 13573183 | 14595897 | 11846412 | 12247835 | 13597120 | 15801031 | 11784428 | 13220862 | 11825093 | 12729901 | 13978837 | 9443620 |
| Q7TPJ0 | Ssr1 | Translocon-associated protein subunit alpha Rattus norvegicus | 155.14 | 10.66 | 35629 | 3 | 3 | 13274833 | 16414066 | 16455467 | 14157776 | 14268068 | 16032420 | 15022717 | 18373849 | 16630559 | 13989660 | 15613477 | 15420427 | 13765915 | 16069556 | 15270117 |
| Q08013 | Ssr3 | Translocon-associated protein subunit gamma Rattus norvegicus | 88.41 | 7.57 | 21064 | 1 | 1 | 5342460 | 4176789.5 | 4127466.5 | 3821924.25 | 3767802.25 | 4365391 | 3388211 | 4616557 | 3807040.5 | 4222170.5 | 3094529.5 | 3171599.5 | 3895573.75 | 3526537.5 | 3963467 |
| Q07984 | Ssr4 | Translocon-associated protein subunit delta Rattus norvegicus | 205.41 | 24.86 | 18980 | 3 | 3 | 5593567.5 | 3916806.5 | 3909770.5 | 3409753 | 3364499.25 | 4191618.25 | 4403748 | 4063969.5 | 2700429.25 | 4674254 | 1929585.12 | 3243683.75 | 3886765.25 | 3632479 | 2255370 |
| Q04931 | Ssrp1 | FACT complex subunit SSRP1 Rattus norvegicus | 514.66 | 19.04 | 80914 | 9 | 9 | 8968763 | 8489271 | 8607678 | 9083119 | 9493513 | 9006370 | 7772367.5 | 8045508.5 | 11652471 | 8457424 | 9794556 | 9063816 | 8737096 | 9883990 | 9138341 |
| Q4KLK9 | Ssu72 | RNA polymerase II subunit A C-terminal domain phosphatase SSU72 Rattus norvegicus | 83.33 | 12.37 | 22544 | 2 | 2 | 1450481.88 | 1394365.25 | 2578613.75 | 1868535 | 1867524.38 | 2109091 | 2047064.62 | 2399185.5 | 2215824 | 1891024.5 | 2378389.5 | 1912992 | 619784.06 | 2032688 | 2098140 |

| | | | | | | | | | | | | | | | | | | | | | | |
|---|---|---|---|---|---|---|---|---|---|---|---|---|---|---|---|---|---|---|---|---|---|---|
| Q99L47 | St13 | Hsc70-interacting protein Mus musculus | 558.98 | 23.99 | 41656 | 11 | 11 | 29978735 | 32553649 | 36151301 | 31068037 | 31646495 | 31654867 | 33198385 | 32238313 | 33211789 | 31871445 | 34861705 | 32639123 | 31684001 | 33873993 | 31488875 |
| O88829 | St3gal5 | Lactosylceramide alpha-2,3-sialyltransferase Mus musculus | 64.94 | 5.56 | 47360 | 2 | 2 | 2107032.25 | 2109276.25 | 1447866.62 | 2348310.5 | 2417529.5 | 2289934.5 | 2279218.25 | 1071938 | 1033000.19 | 2375525.25 | 2315327.75 | 1004309.19 | 1176561.25 | 2146346 | 2525386 |
| Q9D3E6 | Stag1 | Cohesin subunit SA-1 Mus musculus | 290.82 | 6.04 | 144440 | 4 | 6 | 3265961.5 | 1683671.88 | 2003439.88 | 806558.44 | 1580293 | 1472806.88 | 1193658.25 | 2086036.12 | 4449971.5 | 1284281.38 | 2298417.5 | 1973223.88 | 1196207.12 | 2363116 | 1921665.12 |
| O35638 | Stag2 | Cohesin subunit SA-2 Mus musculus | 406.31 | 10.24 | 141281 | 7 | 9 | 6644299.5 | 3486620.25 | 3699485.5 | 2715289.75 | 3123931 | 3477900.25 | 2259712.25 | 4393074 | 7560112.5 | 2715264 | 4851134 | 4716778.5 | 2424407.25 | 4873907.5 | 3992234.25 |
| Q8R424 | Stambp | STAM-binding protein Rattus norvegicus 2 | 57.59 | 4.72 | 48512 | 1 | 1 | 1653348.88 | 1594908.75 | 1296538.12 | 1474115 | 1617054 | 1502972.25 | 1376798 | 1540588.25 | 1433840.88 | 1448905.62 | 1745278.25 | 1161965.75 | 1239486.75 | 1579390.38 | 1481028.62 |
| Q76N33 | Stambpl1 | AMSH-like protease Mus musculus | 56.9 | 4.59 | 49640 | 1 | 1 | 630870.81 | 1153767.12 | 1032593.88 | 1220138.12 | 1061599.25 | 1087762.38 | 1250633.25 | 767900.62 | 826805.38 | 1357939.62 | 978864.88 | 917366.25 | 1069859.88 | 839880.06 | 980438.12 |
| Q9JM90 | Stap1 | Signal-transducing adaptor protein 1 Mus musculus | 637.43 | 59.93 | 34628 | 13 | 13 | 17729511 | 12382445 | 13236752 | 13402185 | 15024507 | 21969469 | 22907261 | 20415201 | 17022513 | 22093031 | 24029665 | 22339825 | 17381839 | 18226125 | 24936277 |
| P42225 | Stat1 | Signal transducer and activator of transcription 1 Mus musculus | 1044.25 | 29.37 | 87197 | 17 | 17 | 27494325 | 28965477 | 27279907 | 21606003 | 25210429 | 22397969 | 22292283 | 25126355 | 24739785 | 20545685 | 24202561 | 24190333 | 28907261 | 28002423 | 23290905 |
| Q9WVL2 | Stat2 | Signal transducer and activator of transcription 2 Mus musculus | 153.72 | 3.47 | 105417 | 3 | 3 | 2279999.25 | 2238939.75 | 2558215.5 | 1865435.12 | 1575645.25 | 1662354.88 | 1336009 | 1722134.88 | 1930979.12 | 1187847.38 | 1414406.62 | 1709033.12 | 1693435.62 | 1945513.38 | 1506688.62 |
| P42227 | Stat3 | Signal transducer and activator of transcription 3 Mus musculus | 483.59 | 22.08 | 88054 | 10 | 10 | 3515281.25 | 2005150.62 | 2500997.5 | 2524098.25 | 2473764.25 | 2875416.5 | 2494043.5 | 1850583 | 1900709.12 | 2691929 | 2902942 | 3008125.25 | 3063253.75 | 3301005.75 | 2563130.75 |
| Q60864 | Stip1 | Stress-induced-phosphoprotein 1 Mus musculus | 1419.29 | 51.93 | 62582 | 25 | 25 | 92142401 | 98960561 | 114734121 | 106186937 | 98563025 | 92165889 | 118198745 | 99953553 | 106279353 | 109009281 | 105028009 | 99428609 | 102928289 | 106449769 | 95878193 |
| O55098 | Stk10 | Serine/threonine-protein kinase 10 Mus musculus | 380.67 | 8.9 | 111906 | 6 | 6 | 4882280.5 | 6873606 | 6064906.5 | 6973655 | 5283605.5 | 5114473.5 | 7686036 | 5740538.5 | 5651798.5 | 6227445 | 6794479 | 6064995 | 6020234.5 | 5871653 | 6016698 |
| Q3TAA7 | Stk11ip | Serine/threonine-protein kinase 11-interacting protein Mus musculus | 104.3 | 2.71 | 118005 | 2 | 2 | 1087222.38 | 435459.94 | 1472558 | 422537.34 | 1048523.12 | 1133528 | 1453642.25 | 1212984.25 | 556679.12 | 1129770.75 | 706297.12 | 1297364.12 | 398395.91 | 407612.69 | 1222280.88 |
| Q99KH8 | Stk24 | Serine/threonine-protein kinase 24 Mus musculus | 404.54 | 25.99 | 47954 | 8 | 8 | 3952477.25 | 5645742.5 | 5255120 | 3935718 | 3924994.5 | 6752330 | 7187371.5 | 7718677.5 | 3507326 | 5698509 | 8061818.5 | 6721213.5 | 6346914.5 | 5458301.5 | 7959442 |
| Q9JI10 | Stk3 | Serine/threonine-protein kinase 3 Mus musculus | 241.69 | 15.9 | 56855 | 2 | 6 | 11367116 | 10492509 | 11975225 | 7462452 | 8947741 | 9667442 | 10316663 | 10999652 | 13049112 | 9698013 | 12047759 | 9893994 | 8721953 | 11139011 | 9887034 |
| Q9JI11 | Stk4 | Serine/threonine-protein kinase 4 Mus musculus | 300.6 | 17.04 | 55541 | 2 | 6 | 6237303 | 5766168.5 | 6652365 | 5194270 | 5365338 | 5610066 | 5332667 | 5609299.5 | 6952883 | 4560973 | 7850775.5 | 5519828 | 4560803.5 | 5740619.5 | 5636116 |
| P13668 | Stmn1 | Stathmin Rattus norvegicus | 123.22 | 23.49 | 17288 | 3 | 3 | 2168176.25 | 1101631.5 | 1538180.38 | 3119678.75 | 1734427.75 | 1713161.38 | 1495789.25 | 1543892.62 | 988564.81 | 2036140.38 | 1707803.75 | 1364339.5 | 1270417.12 | 1833469 | 1413578.25 |
| P54116 | Stom | Erythrocyte band 7 integral membrane protein Mus musculus | 532.44 | 40.49 | 31375 | 9 | 9 | 100953953 | 109211873 | 100468361 | 87066729 | 103234697 | 101366065 | 93146657 | 97780105 | 113819601 | 99952625 | 98492209 | 106308529 | 109589281 | 109130353 | 91718833 |
| Q99JB2 | Stoml2 | Stomatin-like protein 2, mitochondrial Mus musculus | 587.54 | 39.94 | 38385 | 10 | 10 | 8086672 | 8972740 | 10520236 | 10774133 | 9481619 | 10429419 | 12962686 | 9814540 | 9640389 | 12486943 | 8561371 | 7068949 | 9174356 | 9436132 | 11170018 |
| Q5XIG8 | Strap | Serine-threonine kinase receptor-associated protein Rattus norvegicus | 777.83 | 55.14 | 38456 | 12 | 12 | 19361671 | 18032113 | 19995617 | 13686660 | 16922465 | 19698885 | 19090737 | 19875627 | 21519445 | 19021483 | 21654087 | 20386071 | 17907947 | 18157819 | 20160867 |
| Q8VIM6 | Strc | Stereocilin Mus musculus | 35.29 | 0.44 | 196347 | 1 | 1 | 722819.81 | 2210075.5 | 1922223.88 | 1858499.88 | 719493.31 | 2165312.75 | 4443522 | 2196245.5 | 2423662.75 | 3768064.25 | 3559237.75 | 2465075.75 | 3501635.25 | 3825082.5 | 3071432.75 |
| Q8C079 | Strip1 | Striatin-interacting protein 1 Mus musculus | 38.35 | 0.84 | 95585 | 1 | 1 | 1 | 475437 | 1 | 1 | 580265.19 | 496667.5 | 492188.4125 | 492710.53 | 606396.5 | 372979.12 | 563151.06 | 1 | 636094.5 | 557065.5 | 1 |
| O55106 | Strn | Striatin Mus musculus | 43.21 | 1.15 | 85966 | 1 | 1 | 783681.75 | 224602.83 | 983225.56 | 865151.06 | 966120.81 | 846069.12 | 1168787 | 973699.12 | 1023939.84 | 1107204.12 | 866763.7825 | 557104 | 940572.75 | 722223.88 | 1247154.5 |
| P15589 | Sts | Steryl-sulfatase Rattus norvegicus | 37.5 | 1.21 | 62679 | 1 | 1 | 18672391 | 18864409 | 19582717 | 18834287 | 19137203 | 18368079 | 19541797 | 17022287 | 20665637 | 18907025 | 20036181 | 17887673 | 20844767 | 19369047 | 17605733 |
| P46978 | Stt3a | Dolichyl-diphosphooligosaccharide--protein glycosyltransferase subunit STT3A Mus mu | 475.62 | 17.87 | 80598 | 11 | 11 | 21405637 | 17041797 | 20985511 | 19263519 | 18418625 | 15834267 | 17014527 | 15660461 | 19791207 | 14192629 | 15218139 | 13920015 | 15997305 | 20201897 | 13824605 |
| Q3TDQ1 | Stt3b | Dolichyl-diphosphooligosaccharide--protein glycosyltransferase subunit STT3B Mus mu | 160.21 | 4.25 | 93246 | 4 | 4 | 3941781 | 4491079 | 3968801 | 4955797 | 4669015 | 4310805 | 4417855.5 | 4636284.5 | 4111456 | 5044597 | 4921321 | 3816584.5 | 4508349 | 4278041.5 | 3805508.75 |
| Q9WUD1 | Stub1 | STIP1 homology and U box-containing protein 1 Mus musculus | 57.9 | 3.95 | 34909 | 1 | 1 | 3146873.188 | 3086832.25 | 2984497.5 | 3650552 | 2865611 | 2347790.25 | 2517792.875 | 2580080.75 | 2306820.25 | 2836480.25 | 2477411 | 2749622.5 | 2833288.75 | 2666649.75 | 2117652.5 |
| Q9ER00 | Stx12 | Syntaxin-12 Mus musculus | 159.31 | 14.23 | 31195 | 3 | 3 | 5299605 | 5022711 | 5399471.5 | 4769017 | 5899928 | 4867511 | 4145562.75 | 4446703 | 5096103 | 5503597 | 5038344.5 | 4641475 | 5584963.5 | 4957836 | 4170496 |
| Q8BVI5 | Stx16 | Syntaxin-16 Mus musculus | 173.78 | 9.82 | 37080 | 2 | 2 | 672622.56 | 793185.31 | 2133389.5 | 2124225.5 | 642382.19 | 2157544.75 | 1615870 | 2464491.5 | 2939275.5 | 533966.88 | 1048386.88 | 2160195.75 | 602571 | 2529208.5 | 2327919 |
| Q08849 | Stx3 | Syntaxin-3 Rattus norvegicus | 65.64 | 4.84 | 33257 | 1 | 1 | 1304755.5 | 1256390.12 | 1570297.62 | 1050472.62 | 1043461.81 | 1280934.75 | 882770.38 | 1609597.62 | 1557425.5 | 1017394.62 | 925126.75 | 1392238.5 | 953872.38 | 1335986.38 | 928389.75 |
| Q8K1E0 | Stx5 | Syntaxin-5 Mus musculus | 146.87 | 13.24 | 39713 | 3 | 3 | 1 | 1 | 1 | 1 | 502111.38 | 558502 | 1549535 | 729857.56 | 759368.19 | 332548.47 | 1 | 237673.48 | 538010.94 | 187900.88 | 1 |
| O70439 | Stx7 | Syntaxin-7 Mus musculus | 225.56 | 22.22 | 29821 | 4 | 4 | 6188589 | 9226391 | 10026963 | 10626933 | 9494574 | 8400626 | 9780594 | 8765448 | 8234069 | 10140139 | 9369934 | 8251309 | 9043073 | 9339129 | 7470413 |
| O88983 | Stx8 | Syntaxin-8 Mus musculus | 152.78 | 11.86 | 26925 | 2 | 2 | 1879497.563 | 1758616.75 | 2002885.88 | 1978806.12 | 1777681.5 | 1506068 | 1273243.62 | 1465819.25 | 1720422.5 | 1757657.62 | 1576225.25 | 522700.72 | 1897613.12 | 1489897.38 | 1259577 |
| O08599 | Stxbp1 | Syntaxin-binding protein 1 Mus musculus | 145.38 | 6.9 | 67569 | 3 | 3 | 2562417.75 | 2881951 | 2218400 | 2610813.75 | 2852319.5 | 2764546.25 | 2085540.12 | 3054068.5 | 2665402 | 1822894 | 2898295 | 921842.88 | 1371573.5 | 2264031 | 2197214.25 |
| Q64324 | Stxbp2 | Syntaxin-binding protein 2 Mus musculus | 695.14 | 24.28 | 66357 | 11 | 11 | 5440815 | 5271924 | 5078561 | 5142059.5 | 4795616 | 4492609.5 | 4539740 | 6139227 | 6691495 | 4662551 | 5192618 | 6173726 | 5227207.5 | 5299343 | 5753323.5 |
| Q60770 | Stxbp3 | Syntaxin-binding protein 3 Mus musculus | 239.34 | 14.19 | 67942 | 5 | 5 | 2474597 | 3777330.5 | 3009555 | 2729361 | 2934874.75 | 2740933.5 | 2975964.5 | 2984830 | 3371910 | 3213292 | 3327255.25 | 2814181 | 1754097.38 | 3153285.25 | 3453271 |
| Q5DQR4 | Stxbp5l | Syntaxin-binding protein 5-like Mus musculus | 36.53 | 0.93 | 131844 | 1 | 1 | 5538802.5 | 6822379.5 | 4978563.5 | 5871094 | 7652958.5 | 5255096 | 14494090 | 5008501.5 | 6862474 | 11678574 | 10519407 | 8299066.5 | 10844078 | 8508189 | 10373591 |
| P11031 | Sub1 | Activated RNA polymerase II transcriptional coactivator p15 Mus musculus | 271.01 | 44.88 | 14427 | 6 | 6 | 20297853 | 17595211 | 17189027 | 15575062 | 19474133 | 20345529 | 19064891 | 20591687 | 18573529 | 15399311 | 19699479 | 20126683 | 14358023 | 19573059 | 16823299 |
| Q9Z2I9 | Sucla2 | Succinate--CoA ligase [ADP-forming] subunit beta, mitochondrial Mus musculus | 456.06 | 29.37 | 50114 | 9 | 9 | 8295629.5 | 10424399 | 10797735 | 8882850 | 9556912 | 10075774 | 12508839 | 13778828 | 11953550 | 9170292 | 11457529 | 12496327 | 11051370 | 12591289 | 11729394 |
| Q9WUM5 | Suclg1 | Succinate--CoA ligase [ADP/GDP-forming] subunit alpha, mitochondrial Mus musculus | 308.19 | 23.12 | 36155 | 5 | 5 | 1708247.25 | 1856638 | 2603641.25 | 2963060.5 | 3241120.5 | 3401179 | 5673253 | 526358.19 | 466124.16 | 6004803 | 3435702 | 3428697.5 | 3211728 | 529758.56 | 5883449 |
| Q9Z2I8 | Suclg2 | Succinate--CoA ligase [GDP-forming] subunit beta, mitochondrial Mus musculus | 832.03 | 43.42 | 46840 | 14 | 14 | 46315337 | 40460253 | 48964709 | 41624685 | 43707373 | 53398785 | 41200969 | 55963313 | 61248589 | 37070281 | 54612129 | 52764033 | 38491477 | 56986657 | 45118721 |
| Q9CX34 | Sugt1 | Protein SGT1 homolog Mus musculus | 291.94 | 23.81 | 38159 | 6 | 6 | 6946320 | 7296044.5 | 11091518 | 8172529.5 | 8410025 | 7654505.5 | 7638505 | 9142288 | 10288957 | 8432627 | 9587723 | 9198531 | 7469486.5 | 9095881 | 10635051 |
| Q8R0F3 | Sumf1 | Formylglycine-generating enzyme Mus musculus | 57.92 | 4.03 | 40659 | 1 | 1 | 810070.88 | 982812.97 | 1167581 | 927408.75 | 1026191.25 | 686075.62 | 1299337.38 | 825096.62 | 930235.81 | 1090828.38 | 983567 | 990986.25 | 965604.75 | 819768.75 | 928052.81 |
| P63166 | Sumo1 | Small ubiquitin-related modifier 1 Mus musculus | 33.75 | 6.93 | 11557 | 1 | 1 | 2866870.75 | 2263323 | 2727641.75 | 2733389.5 | 2330636.75 | 2529495 | 4045481.25 | 2560367 | 2250176.25 | 3424269.5 | 2262142.25 | 2881433.25 | 2828226 | 2668008.75 | 2960613 |
| P61957 | Sumo2 | Small ubiquitin-related modifier 2 Mus musculus | 77.77 | 12.63 | 10871 | 1 | 1 | 17521405 | 2550764.75 | 3024832.5 | 8302931 | 18981269 | 9775257 | 4411687.5 | 2285951.75 | 2406698.5 | 11814443 | 11101485 | 2164145.5 | 3208033.75 | 3596990.75 | 7876416 |
| Q9D666 | Sun1 | SUN domain-containing protein 1 Mus musculus | 68.36 | 0.99 | 101976 | 1 | 1 | 1243723.62 | 930462.25 | 1211287.453 | 968380.06 | 1702583.88 | 1275486.38 | 1199882.38 | 1245976.88 | 1252881.75 | 964495.56 | 1314430.12 | 1168718.62 | 1175336.75 | 1280961.25 | 786723.69 |
| Q8BJS4 | Sun2 | SUN domain-containing protein 2 Mus musculus | 357.59 | 12.72 | 81605 | 7 | 7 | 7134746 | 7171136 | 6667814 | 8281610.5 | 8802811 | 7091866 | 7631134 | 6864571 | 6787869 | 8751663 | 9431669 | 7373396.5 | 7313194 | 7297480 | 8474897 |
| Q920B9 | Supt16h | FACT complex subunit SPT16 Mus musculus | 1362.52 | 30.47 | 119825 | 26 | 26 | 35326781 | 36160489 | 38503145 | 33197889 | 36850221 | 35338693 | 39147093 | 32659973 | 40558993 | 33202923 | 40544133 | 38777613 | 32968771 | 33802905 | 38773745 |
| O55201 | Supt5h | Transcription elongation factor SPT5 Mus musculus | 622.98 | 16.82 | 120664 | 13 | 13 | 6384110 | 5187356.5 | 4203389 | 3230445.25 | 5565550 | 4453712 | 2633916 | 3106731.5 | 3691993 | 4539777 | 3547915 | 5083947.5 | 3902555.5 | 5100100 | 5571018 |
| Q62383 | Supt6h | Transcription elongation factor SPT6 Mus musculus | 38.59 | 0.87 | 199086 | 1 | 1 | 815871 | 938990 | 839073.06 | 844556.06 | 902020 | 749238.56 | 607558.81 | 951418 | 918896.25 | 807035 | 923763.75 | 945723.69 | 816697.38 | 910033.25 | 758630.56 |
| Q64310 | Surf4 | Surfeit locus protein 4 Mus musculus | 274.41 | 21.19 | 30381 | 4 | 4 | 9366277 | 4119156.5 | 6308821 | 3941644.25 | 3997260 | 4508995.5 | 4310565 | 6117197 | 7402717.5 | 3557408 | 4570312.5 | 5972971 | 4579291.5 | 5098657 | 6151820 |
| P70279 | Surf6 | Surfeit locus protein 6 Mus musculus | 80.44 | 4.79 | 41235 | 2 | 2 | 1 | 1 | 371485.03 | 1 | 1 | 1 | 541889.56 | 421203.66 | 1 | 417635.28 | 374938.62 | 516285.59 | 460355.56 | 264122.03 | 686076 |
| Q80U70 | Suz12 | Polycomb protein Suz12 Mus musculus | 154.41 | 5.67 | 83026 | 3 | 3 | 1049436 | 949619.69 | 723083.44 | 868012.5 | 936224.38 | 1306857.5 | 1199797.25 | 520489.88 | 606293.88 | 939515.5 | 847004.88 | 774157.75 | 967783.75 | 716751 | 574349 |
| Q8K4L3 | Svil | Supervillin Mus musculus | 70.09 | 1.24 | 243162 | 2 | 2 | 903145.62 | 1424224.25 | 1116957.25 | 793862.38 | 1657934.75 | 901411.25 | 1789168 | 1688930.5 | 1683442.25 | 1541226.62 | 1073059.25 | 1020774.12 | 1651339.75 | 1545853.88 | 1034867.19 |
| Q6A028 | Swap70 | Switch-associated protein 70 Mus musculus | 826.42 | 35.21 | 68996 | 18 | 18 | 8438880 | 4376241.5 | 9912001 | 6593201 | 9192176 | 9780575 | 9011705 | 8434352 | 10127617 | 11306538 | 11334125 | 10442167 | 13070007 | 11563968 | 16333501 |
| Q80X82 | Sympk | Symplekin Mus musculus | 607.63 | 13.55 | 142284 | 12 | 12 | 6440025.5 | 5867272.5 | 7302222 | 5437608 | 5401326.5 | 6835220 | 4272746.5 | 6141338.5 | 9246548 | 6149009 | 6766833.5 | 6616016 | 4850413.5 | 7612429.5 | 6917543 |
| Q7TMK9 | Syncrip | Heterogeneous nuclear ribonucleoprotein Q Mus musculus | 1155.25 | 39.65 | 69633 | 19 | 19 | 40615889 | 36728261 | 36924769 | 44591941 | 42022185 | 40662693 | 43531909 | 38910841 | 42826529 | 45042593 | 43753389 | 45455261 | 39449641 | 41055433 | 46191989 |
| O55100 | Syngr1 | Synaptogyrin-1 Mus musculus | 38.68 | 5.13 | 25653 | 1 | 1 | 2450956.5 | 2020317 | 2084739.25 | 1717727.38 | 2355973 | 2252462.5 | 316088.19 | 1816375 | 2296093.75 | 2176585.75 | 1766132.38 | 2292465.75 | 2864408 | 1643943.12 | 2284268.25 |
| O55101 | Syngr2 | Synaptogyrin-2 Mus musculus 2 | 48.79 | 4.46 | 24778 | 1 | 1 | 6716908 | 5932428.5 | 6593905.5 | 6176990 | 6504500.5 | 7400437.5 | 5357093 | 6119711 | 7860009.5 | 6760877.5 | 9162380 | 7337439.5 | 7657008 | 6388194.5 | 7239333 |
| Q62910 | Synj1 | Synaptojanin-1 Rattus norvegicus | 57.38 | 0.76 | 172881 | 1 | 1 | 650035.81 | 819361.69 | 762494.56 | 645826.94 | 733179.38 | 780039.75 | 482878.59 | 734940.12 | 716394.19 | 501863.41 | 831700.44 | 605563.5 | 686236.62 | 687034.38 | 585716.25 |
| O09117 | Sypl1 | Synaptophysin-like protein 1 Mus musculus | 74.56 | 4.21 | 28899 | 2 | 2 | 7626975 | 7761680 | 7000726 | 6510608 | 7570838 | 7720568 | 7924660.5 | 7573445.5 | 7328807.5 | 4292792 | 7802798 | 7109643 | 7548428 | 6296194 | 7660582 |
| Q6Y685 | Tacc1 | Transforming acidic coiled-coil-containing protein 1 Mus musculus | 87.02 | 2.97 | 83952 | 2 | 2 | 1 | 1 | 1281975 | 209819.17 | 230212.3 | 249929.91 | 709251.5125 | 968461.94 | 1365721.62 | 252892.58 | 1220298.12 | 1136883.62 | 180982.06 | 886094.94 | 275156.97 |
| Q9JJ11 | Tacc3 | Transforming acidic coiled-coil-containing protein 3 Mus musculus | 41.96 | 2.06 | 70627 | 1 | 1 | 570164.88 | 683694.69 | 984348.31 | 634961.25 | 639077.06 | 713601.38 | 542661.94 | 672478.94 | 867654.5 | 716917.69 | 797539.75 | 1095959.75 | 864344.88 | 669858.06 | 781100.12 |
| B2RYT9 | Taco1 | Translational activator of cytochrome c oxidase 1 Rattus norvegicus | 72.6 | 6.1 | 32754 | 1 | 1 | 520645.19 | 917651.88 | 1258428.62 | 1122164.12 | 911669.06 | 1284125.88 | 433098.72 | 1047591.5 | 1052326.25 | 1219332.5 | 826150.62 | 1206542.62 | 942746.94 | 1088475.88 | 1278857.62 |
| Q8C092 | Taf5 | Transcription initiation factor TFIID subunit 5 Mus musculus | 48.69 | 1.37 | 87045 | 1 | 1 | 249290.66 | 221653.39 | 291923.91 | 275313.72 | 263304.69 | 308147.91 | 298303.97 | 266253.5025 | 191302.22 | 267259.91 | 363836.97 | 253867.36 | 221478.52 | 274619.5 | 213031.02 |
| Q91WQ5 | Taf5l | TAF5-like RNA polymerase II p300/CBP-associated factor-associated factor 65 kDa sub | 43.84 | 3.74 | 65911 | 1 | 1 | 196988.38 | 279299.28 | 218964.33 | 247291.38 | 293913.53 | 305316.56 | 206286.39 | 207516.0375 | 188603.42 | 129857.78 | 240755.28 | 255686.06 | 381888.06 | 297750.3575 | 312672.03 |
| Q9WVA4 | Tagln2 | Transgelin-2 Mus musculus | 1085.59 | 81.41 | 22395 | 15 | 17 | 360666753 | 431408257 | 389466945 | 432111265 | 416786625 | 406168129 | 398763649 | 415400961 | 436928961 | 432150465 | 398752417 | 372969793 | 381565665 | 372534113 | 360367649 |
| Q93092 | Taldo1 | Transaldolase Mus musculus | 1291.53 | 72.4 | 37387 | 24 | 24 | 313788705 | 345215713 | 352452129 | 280869825 | 309088321 | 308337665 | 324675841 | 356930977 | 355244193 | 290641665 | 342404641 | 339987457 | 320346721 | 332710913 | 289879361 |
| Q0VGY8 | Tanc1 | Protein TANC1 Mus musculus | 276.4 | 5.87 | 200805 | 6 | 6 | 1665739.12 | 1572150.38 | 1584758.25 | 1529621.75 | 1553410 | 2079376.75 | 2248288.5 | 2210395.5 | 1783772.38 | 2533140.5 | 1947109.12 | 2153689 | 2294152 | 2127409.5 | 2663115.25 |
| Q53UA7 | Taok3 | Serine/threonine-protein kinase TAO3 Rattus norvegicus | 226.72 | 5.57 | 105474 | 4 | 4 | 1636289.5 | 1070588.38 | 1438767.5 | 1499944.12 | 1010315 | 1181438.62 | 2298503.25 | 1430709 | 958852.62 | 1710735 | 1001445.88 | 1496270.12 | 1191015.12 | 1209211 | 1491950.25 |
| P21958 | Tap1 | Antigen peptide transporter 1 Mus musculus | 423.56 | 17.68 | 78864 | 9 | 9 | 10058543 | 10971870 | 10419274 | 7036405.5 | 7464756 | 8379706 | 9821185 | 10371351 | 9781903 | 9070415 | 6994086.5 | 8340282.5 | 9574261 | 10422529 | 8378264 |
| P36371 | Tap2 | Antigen peptide transporter 2 Mus musculus | 530.08 | 19.52 | 77445 | 9 | 9 | 14567948 | 13917693 | 10892413 | 10225757 | 12761393 | 12345364 | 11126403 | 12282652 | 13814503 | 11760215 | 10867383 | 6823816.5 | 12415366 | 13111455 | 6884573.5 |
| Q9R233 | Tapbp | Tapasin Mus musculus | 211.01 | 12.9 | 49736 | 4 | 4 | 11257769 | 11536198 | 11120231 | 6555127 | 10087860 | 9637459 | 11177781 | 10197578 | 8762071 | 9692128 | 8629771 | 7036440 | 11578260 | 9324442 | 6762948 |
| Q921F2 | Tardbp | TAR DNA-binding protein 43 Mus musculus | 930.72 | 50 | 44548 | 14 | 14 | 23175409 | 22328111 | 23927153 | 27226111 | 27859395 | 29521305 | 29482325 | 22002141 | 21513303 | 31092945 | 30423061 | 25118199 | 27087009 | 23185759 | 33674397 |
| Q9D0R2 | Tars | Threonine--tRNA ligase, cytoplasmic Mus musculus | 1599.79 | 46.12 | 83356 | 28 | 29 | 65506257 | 62482541 | 61105641 | 61218469 | 61199493 | 62353825 | 61020933 | 62215949 | 68411145 | 59597841 | 61003633 | 61387781 | 62498965 | 66520833 | 69408977 |
| Q3UQ84 | Tars2 | Threonine--tRNA ligase, mitochondrial Mus musculus | 118.35 | 3.6 | 81700 | 2 | 2 | 1062753.38 | 1229340 | 1103442.12 | 1291838.75 | 1461157 | 1365174.5 | 1188529.12 | 1461139.88 | 1414028.62 | 1304064 | 941526.12 | 1205765.88 | 984947.69 | 1273739.12 | 1024414.38 |
| Q8R3D1 | Tbc1d13 | TBC1 domain family member 13 Mus musculus | 45.93 | 2.5 | 46453 | 1 | 1 | 724563.75 | 774910.88 | 598706.19 | 716630.38 | 715621.12 | 926029.94 | 452823.25 | 574433.5 | 799547.06 | 638341.56 | 700286.56 | 642817.44 | 629012 | 1117873.12 | 496288.59 |
| Q9CXF4 | Tbc1d15 | TBC1 domain family member 15 Mus musculus | 470.21 | 14.9 | 76527 | 8 | 8 | 11482830 | 9404947 | 13581257 | 11388000 | 11701886 | 9675982 | 11807250 | 11340820 | 13708710 | 8165829.5 | 8943522 | 11351048 | 8809320 | 12591999 | 9355059 |
| Q8BYH7 | Tbc1d17 | TBC1 domain family member 17 Mus musculus | 69.78 | 1.55 | 72860 | 1 | 1 | 1826476 | 2320990 | 2133825.75 | 2061661.5 | 2286611.5 | 2271583 | 1311340.38 | 2229124.5 | 2493446.75 | 1844517.75 | 2145783 | 1830050.12 | 1856198.75 | 2059067.5 | 1866915.25 |
| Q5SVR0 | Tbc1d9b | TBC1 domain family member 9B Mus musculus | 165.54 | 3.09 | 141779 | 3 | 3 | 1177040.5 | 1080592.25 | 1238921.62 | 1190100.12 | 983897.62 | 1251904.75 | 1072114.62 | 1182771.25 | 1395886.88 | 1098644.88 | 762685.25 | 1062780.75 | 1233271.5 | 1233671.62 | 961897.75 |
| P48428 | Tbca | Tubulin-specific chaperone A Mus musculus | 224.12 | 56.48 | 12758 | 5 | 5 | 9131711 | 9397210 | 8753151 | 10509868 | 8785775 | 7585664 | 6202783 | 8991456 | 8507587 | 8926501 | 9005037 | 7408565.5 | 8686178 | 8077935.5 | 7330137 |
| Q9D1E6 | Tbcb | Tubulin-folding cofactor B Mus musculus | 287.78 | 32.79 | 27386 | 5 | 5 | 1688019.12 | 1699028.62 | 2197702.25 | 1859222.75 | 3468018.25 | 3319070 | 3338407 | 2372161.75 | 2746277.75 | 2065962.25 | 2705406.25 | 2897079 | 3475558 | 2689482.75 | 3690229.25 |
| Q8VCN9 | Tbcc | Tubulin-specific chaperone C Mus musculus | 120.93 | 8.8 | 38125 | 2 | 2 | 1930435.62 | 1688610.62 | 2308558.75 | 1972986.88 | 1508172.5 | 1285844.5 | 1225599 | 884210.94 | 1320738.88 | 2916509.5 | 1746553.62 | 1540185.12 | 1672979.5 | 1735233.5 | 4243309 |
| Q8BYA0 | Tbcd | Tubulin-specific chaperone D Mus musculus | 479.79 | 11.12 | 133321 | 10 | 10 | 9827232 | 10397821 | 10799128 | 11129129 | 9635791 | 11272850 | 8150864 | 9646606 | 12438733 | 8210696.5 | 9843651 | 10845919 | 9541308 | 12098717 | 9428615 |
| Q5FVQ9 | Tbce | Tubulin-specific chaperone E Rattus norvegicus | 38.82 | 2.1 | 59043 | 1 | 1 | 273605.12 | 1 | 202520.45 | 1 | 264504.56 | 278054.25 | 287940.38 | 258549.41 | 308550.22 | 283273.565 | 203796.41 | 149373.03 | 190033.5 | 214477.62 | 261524.84 |
| Q5PQJ7 | Tbcel | Tubulin-specific chaperone cofactor E-like protein Rattus norvegicus | 88.23 | 5.9 | 48046 | 1 | 1 | 602050.81 | 885528.81 | 499498.59 | 948888.75 | 915210.38 | 692415.25 | 454505.84 | 893922.75 | 759208.56 | 646952.62 | 498025.78 | 753073.88 | 446445.44 | 506574.03 | 1105205.25 |
| Q9QXE7 | Tbl1x | F-box-like/WD repeat-containing protein TBL1X Mus musculus | 149.35 | 5.5 | 56802 | 1 | 2 | 2227288 | 1777417.62 | 1661314.12 | 2389203.5 | 1955666.62 | 1457449.62 | 1323082.12 | 1617709.38 | 1014356.56 | 1444561.12 | 1691760 | 689089.94 | 1212120.62 | 1110346.62 | 676803 |
| Q8BHJ5 | Tbl1xr1 | F-box-like/WD repeat-containing protein TBL1XR1 Mus musculus | 213.14 | 9.14 | 55661 | 2 | 3 | 14106546 | 16720502 | 16898611 | 15335139 | 1277937 | 893155.56 | 1168404.25 | 18563871 | 22492433 | 1007102.31 | 2849797.5 | 1058262 | 666507.12 | 17569359 | 940471.25 |
| Q9R099 | Tbl2 | Transducin beta-like protein 2 Mus musculus | 77.91 | 5.88 | 49584 | 2 | 2 | 1479462 | 1627648.5 | 1688083.25 | 696873 | 1092593 | 755756.62 | 806844.75 | 1759795.75 | 607305.19 | 1891126.25 | 589999.38 | 797473.88 | 868704.38 | 1011439.25 | 1857822.5 |
| Q8C4J7 | Tbl3 | Transducin beta-like protein 3 Mus musculus | 352.15 | 13.61 | 88266 | 8 | 8 | 3707170.75 | 3182216.25 | 3525159.5 | 3772371 | 2670673.75 | 3312765.5 | 3581206 | 4231631.5 | 4162804.5 | 3312815.75 | 3638801.5 | 3440066.75 | 3401407.25 | 4006966 | 3239519 |
| Q91YM4 | Tbrg4 | FAST kinase domain-containing protein 4 Mus musculus | 130.32 | 7.62 | 71513 | 3 | 3 | 2510035 | 2642429.75 | 2381086.75 | 1631869.62 | 1641948 | 2452593.75 | 1612505.953 | 604672.5 | 2741559.5 | 651198.06 | 1398655.315 | 605356.12 | 2397238 | 2349604.5 | 242422.64 |
| P10711 | Tcea1 | Transcription elongation factor A protein 1 Mus musculus | 556.38 | 36.21 | 33880 | 2 | 9 | 8782650 | 10094231 | 9063652 | 10786020 | 8662249 | 6913421 | 9966594 | 9492569 | 8034122 | 11314221 | 6753447.5 | 11440706 | 9811065 | 8251111 | 11155971 |
| Q8CGF7 | Tcerg1 | Transcription elongation regulator 1 Mus musculus | 152.12 | 3.91 | 123788 | 3 | 3 | 4030092.5 | 1967734.62 | 1834887 | 1163685.75 | 743651.25 | 2168156.5 | 2726350.75 | 1908883.88 | 1433781.62 | 3089966 | 1833505 | 1888489 | 1791077.5 | 1978683.75 | 1947287.38 |
| O08784 | Tcof1 | Treacle protein Mus musculus | 455.56 | 13.94 | 135001 | 12 | 12 | 5144935.5 | 5856552 | 5091077.5 | 5333352.5 | 4983309.5 | 5410892 | 12252951 | 4404813 | 5207385 | 6410353 | 7727158.5 | 4639139 | 4886903 | 5513065 | 7493827 |
| P11983 | Tcp1 | T-complex protein 1 subunit alpha Mus musculus | 1961.24 | 70.14 | 60449 | 28 | 29 | 103142201 | 98834041 | 128821737 | 113434849 | 119067593 | 120087921 | 145437329 | 113149433 | 108312497 | 135172289 | 125451217 | 133218609 | 118899385 | 109676065 | 135243841 |
| Q4G056 | Tdp1 | Tyrosyl-DNA phosphodiesterase 1 Rattus norvegicus | 46.82 | 3.12 | 68632 | 1 | 1 | 1 | 1 | 1 | 1 | 1 | 1 | 1 | 1 | 297044 | 397036.84 | 409229.78 | 295515.09 | 1 | 1 | 386490.41 |
| Q64232 | Tecr | Very-long-chain enoyl-CoA reductase Rattus norvegicus | 109.64 | 6.82 | 36123 | 2 | 2 | 14777568 | 765068.69 | 857992 | 548268.62 | 1024155.94 | 1276692 | 1191148.75 | 860610.38 | 1461098.88 | 1002251.88 | 1155001.25 | 978747.5 | 1287770.62 | 988903.69 | 1419098.88 |
| Q5EAN7 | Terf2ip | Telomeric repeat-binding factor 2-interacting protein 1 Rattus norvegicus | 69.76 | 3.56 | 43418 | 1 | 1 | 303358.16 | 1 | 368227.56 | 1 | 1 | 357669.56 | 340004.56 | 401303.25 | 572864 | 417960.3425 | 478680.19 | 393511.78 | 1 | 455871.5 | 391174.84 |
| P47226 | Tes | Testin Mus musculus | 353.71 | 21.04 | 47983 | 8 | 8 | 3696221.75 | 4564285.5 | 4730642 | 6025362.5 | 4329370.5 | 6012790 | 7860529.5 | 4216560.5 | 4572421 | 6256796.5 | 5923914.5 | 7187647 | 4801883.5 | 4016422.5 | 5624385 |
| Q3URQ0 | Tex10 | Testis-expressed protein 10 Mus musculus | 40.31 | 1.4 | 105209 | 1 | 1 | 482972.75 | 307244.69 | 168606.17 | 225089.27 | 284859.88 | 290707.75 | 154352.08 | 276752.53 | 248410.47 | 230305.92 | 202727.47 | 163674.56 | 237493.2 | 338007.28 | 164695.41 |
| Q921I1 | Tf | Serotransferrin Mus musculus | 40.02 | 2.01 | 76724 | 1 | 1 | 307375.94 | 237959.55 | 224392.09 | 422393.12 | 246643.97 | 296060.03 | 317108 | 326695.4225 | 331302.72 | 362310.94 | 1 | 215861.28 | 1 | 235606.75 | 291837.56 |
| Q811P6 | Tfb1m | Dimethyladenosine transferase 1, mitochondrial Rattus norvegicus | 66.78 | 5.51 | 39181 | 2 | 2 | 800350.69 | 1495634.25 | 1176110.5 | 1251058.25 | 1301972.25 | 1471660.75 | 1512859 | 1951628.38 | 576315.44 | 501673.78 | 1150582.25 | 1387338 | 1335850.5 | 1547714.75 | 1230709.25 |
| Q9ERA6 | Tfip11 | Tuftelin-interacting protein 11 Mus musculus | 150.74 | 5.49 | 96305 | 3 | 3 | 272489.78 | 336598.19 | 352474.6175 | 357605.84 | 443204.66 | 366940.09 | 1388584.25 | 439833.31 | 395238.03 | 202218.86 | 432561.97 | 367157.0775 | 233450.78 | 400424.25 | 402191.31 |
| Q62351 | Tfrc | Transferrin receptor protein 1 Mus musculus | 939.45 | 30.54 | 85731 | 20 | 20 | 11530963 | 11633155 | 10900484 | 7395272.5 | 10115463 | 8576711 | 8159413.5 | 11486460 | 11872200 | 6156256 | 10086660 | 11244051 | 10993838 | 13388253 | 8401939 |
| Q91YX0 | Themis2 | Protein THEMIS2 Mus musculus | 451.31 | 22.32 | 74378 | 9 | 9 | 6519084.5 | 7443011.5 | 6576726.5 | 7534454 | 7404014.5 | 5142677 | 3751690 | 4517776.5 | 4762057.5 | 3096810 | 5101699 | 4559930.5 | 3806501.5 | 3769314.25 | 4837313 |

| | | | | | | | | | | | | | | | | | | | | | |
|---|---|---|---|---|---|---|---|---|---|---|---|---|---|---|---|---|---|---|---|---|---|
| P59924 | Thoc1 | THO complex subunit 1 Rattus norvegicus | 36.64 | 2.62 | 40243 | 1 | 1 | 1837730 | 1421757.88 | 2604596.75 | 1773867.813 | 1231386.62 | 1477927.25 | 940296.81 | 1593948.38 | 1974223.5 | 1496598.985 | 1329269.75 | 1761381 | 665528.44 | 2065170.25 | 1515509 |
| B1AZI6 | Thoc2 | THO complex subunit 2 Mus musculus | 570.07 | 9.28 | 182773 | 12 | 12 | 8149270.5 | 7828305.5 | 9587872 | 8968936 | 9977361 | 9896656 | 6210598.5 | 8303131 | 8084742 | 7049125 | 8239659 | 9345430 | 8345822.5 | 9296373 | 8459494 |
| Q8VE80 | Thoc3 | THO complex subunit 3 Mus musculus | 45.07 | 6.55 | 38738 | 1 | 1 | 210528.8425 | 270131.34 | 162510.39 | 237879.44 | 171594.2 | 1 | 177124.12 | 210987.75 | 166225.73 | 1 | 296710 | 243910.135 | 269618.22 | 174504.12 | 234808.2 |
| Q68FX7 | Thoc5 | THO complex subunit 5 homolog Rattus norvegicus | 48.69 | 1.03 | 78660 | 1 | 1 | 1197032.38 | 946366.56 | 1253918.25 | 743662.75 | 765300.56 | 873429.5 | 672751.56 | 809667.39 | 1141042.62 | 551445.88 | 1279751.25 | 1143033.12 | 1182953.093 | 1243712.12 | 1065315.88 |
| Q5U4D9 | Thoc6 | THO complex subunit 6 homolog Mus musculus | 114.37 | 10.56 | 37315 | 2 | 2 | 1086235.25 | 781938 | 1699729.5 | 1547681.75 | 1151054.75 | 895295.06 | 563872.5 | 1027862.25 | 691709.44 | 1355419 | 962372.62 | 1050282 | 676497.06 | 945531.06 | 1125191.75 |
| Q8C1A5 | Thop1 | Thimet oligopeptidase Mus musculus | 366.94 | 13.39 | 78026 | 9 | 9 | 11717284 | 10877380 | 10613722 | 12053381 | 12043315 | 11789105 | 10074378 | 11376863 | 13210187 | 11059504 | 11691063 | 9957071 | 11505736 | 11914489 | 11591025 |
| Q569Z6 | Thrap3 | Thyroid hormone receptor-associated protein 3 Mus musculus | 482.4 | 13.67 | 108178 | 10 | 10 | 3810779 | 3779500.5 | 3694478.75 | 2688826.75 | 3653554 | 3145846.5 | 4527348.5 | 2941184.25 | 2697955.5 | 5247651 | 3536816.25 | 4278294.5 | 4127388.5 | 2719459 | 4984430 |
| Q99J36 | Thumpd1 | THUMP domain-containing protein 1 Mus musculus | 246.32 | 23.14 | 38885 | 6 | 6 | 5087066 | 3713403.5 | 4923494 | 3482210.5 | 4537704 | 4337629 | 3465145.25 | 3561460.5 | 4201563 | 4359024 | 4731547 | 3585659.25 | 3288894.25 | 4462889.5 | 6526747 |
| P97770 | Thumpd3 | THUMP domain-containing protein 3 Mus musculus | 227.58 | 11.49 | 56431 | 5 | 5 | 899067.62 | 1041571.62 | 1029383.31 | 1288757.88 | 1037148.62 | 1052995 | 1857270.25 | 896109.19 | 1021293.94 | 1645488.75 | 1235171.25 | 1368186.25 | 1408643.62 | 1017139 | 1643561.62 |
| P52912 | Tia1 | Nucleolysin TIA-1 Mus musculus | 51.35 | 4.66 | 42800 | 1 | 1 | 329928.09 | 292098.72 | 249242.75 | 298583 | 292463.14 | 1 | 188017.73 | 1 | 1 | 331924.75 | 405246.81 | 1 | 272014.53 | 1 | 298345.25 |
| P70318 | Tial1 | Nucleolysin TIAR Mus musculus | 228.31 | 11.99 | 43389 | 4 | 4 | 6755246 | 3901110.25 | 2767010.5 | 3009831 | 4636751 | 3765651 | 4413943.5 | 3462739.75 | 4035390.5 | 3869400.5 | 4521019.5 | 3968794 | 3365234.5 | 4162907.75 | 4238766.5 |
| Q793I8 | Tifa | TRAF-interacting protein with FHA domain-containing protein A Mus musculus | 39.76 | 4.89 | 21560 | 1 | 1 | 559648.69 | 481834.75 | 630341.81 | 799239.31 | 653510.75 | 582405.81 | 843120.88 | 702981.56 | 631316.19 | 893154.62 | 532165.25 | 669609 | 670203.44 | 811587.5 | 782861.62 |
| Q8BZA9 | Tigar | Fructose-2,6-bisphosphatase TIGAR Mus musculus | 247.69 | 34.2 | 29191 | 5 | 5 | 3817089.5 | 4261817 | 4053701 | 3545411.5 | 4825893.5 | 4491076.5 | 3614555.5 | 4047236.5 | 4172443.5 | 5303476 | 3658969.5 | 3224379.5 | 4198461.5 | 4345671 | 5301178 |
| P62075 | Timm13 | Mitochondrial import inner membrane translocase subunit Tim13 Mus musculus | 36.2 | 14.74 | 10458 | 1 | 1 | 2119784.5 | 2271527.25 | 2447546.5 | 2781436.25 | 2072312.75 | 1758826 | 1823802.25 | 1746608 | 1605808 | 2113046.25 | 1606487.5 | 1664509.88 | 2017443.62 | 1194384.88 | 1110859.12 |
| O35092 | Timm17a | Mitochondrial import inner membrane translocase subunit Tim17-A Rattus norvegicus | 225.05 | 43.27 | 18038 | 3 | 3 | 1488920.75 | 1750135.12 | 1635881.25 | 5872844.5 | 1660234.25 | 2043096.12 | 1617769.75 | 2026209.5 | 2024882.38 | 1666908.25 | 1307253.62 | 2011417.12 | 1786844.5 | 1711404 | 1320164.62 |
| Q9Z0V7 | Timm17b | Mitochondrial import inner membrane translocase subunit Tim17-B Mus musculus | 97.86 | 29.65 | 18352 | 2 | 2 | 571825.06 | 364785.34 | 1 | 1 | 1 | 1 | 1804580.25 | 1 | 1 | 1245187.75 | 1554491.25 | 1 | 1305471.88 | 2182517.5 | 1 |
| Q9WTQ8 | Timm23 | Mitochondrial import inner membrane translocase subunit Tim23 Mus musculus | 208.48 | 37.32 | 21978 | 3 | 3 | 1655220 | 2294022.75 | 2154669.5 | 2537059 | 2370898.5 | 2511439.25 | 2424310.75 | 2349864 | 1993917.75 | 2060339.88 | 1764037 | 561292.5 | 2269700.5 | 2393850.25 | 1736563.5 |
| O35857 | Timm44 | Mitochondrial import inner membrane translocase subunit TIM44 Mus musculus | 644.41 | 25.44 | 51091 | 11 | 11 | 11188471 | 12907923 | 13961999 | 10798965 | 11944740 | 13031578 | 14903231 | 14342581 | 14710107 | 10635579 | 14037452 | 12757525 | 11339333 | 13921178 | 11464936 |
| Q9D880 | Timm50 | Mitochondrial import inner membrane translocase subunit TIM50 Mus musculus | 500.41 | 28.33 | 39776 | 8 | 8 | 21850333 | 21642433 | 23470217 | 21432963 | 19927959 | 22439989 | 22235993 | 22615759 | 24134467 | 22675727 | 22840037 | 24715841 | 20623819 | 21367929 | 22979923 |
| A2VCX1 | Tiprl | TIP41-like protein Rattus norvegicus | 88.43 | 6.64 | 31225 | 2 | 2 | 1010077.5 | 1185112.38 | 1003873.19 | 1027271.5 | 951998.25 | 1127915.62 | 1285338.38 | 1153446.62 | 1505938.12 | 1188953.5 | 821440.38 | 1140050.38 | 923500.38 | 1049318.88 | 1118792.5 |
| Q9Z0U1 | Tjp2 | Tight junction protein ZO-2 Mus musculus | 551.53 | 11.14 | 131280 | 10 | 10 | 4912769 | 5892577 | 5076197 | 6428088.5 | 5285090 | 7309216 | 10987118 | 6877278.5 | 7799566 | 8736109 | 7421016 | 5551806.5 | 5444674.5 | 6376130.5 | 5578498.5 |
| P04184 | Tk1 | Thymidine kinase, cytosolic Mus musculus | 211.99 | 19.74 | 25776 | 4 | 4 | 1585144.5 | 1742422.25 | 1060971.75 | 2080591.25 | 3288614.25 | 939617.25 | 1523118.5 | 2832684.5 | 1244693.12 | 2032342.5 | 902201.06 | 927999.75 | 1807973.25 | 1788207 | 872240.75 |
| P40142 | Tkt | Transketolase Mus musculus | 2163.61 | 67.74 | 67630 | 33 | 34 | 278338977 | 278360673 | 292927297 | 297235425 | 310469057 | 294017281 | 336577473 | 282155457 | 290506945 | 319185633 | 322402721 | 333478849 | 297733889 | 293938433 | 347451233 |
| P26039 | Tln1 | Talin-1 Mus musculus | 7638.56 | 62.42 | 269821 | 106 | 106 | 416044193 | 454960129 | 469737793 | 454685089 | 458625025 | 446006625 | 493456545 | 431872033 | 449273313 | 477256737 | 517623073 | 442095585 | 433020609 | 437770657 | 474966753 |
| Q6R5N8 | Tlr13 | Toll-like receptor 13 Mus musculus | 647.17 | 13.62 | 114443 | 9 | 9 | 12779083 | 9427389 | 12441244 | 11319293 | 11369952 | 9088357 | 7111226 | 9942213 | 11771393 | 7924610 | 10095746 | 11090743 | 9746947 | 10198689 | 9800169 |
| Q9QUN7 | Tlr2 | Toll-like receptor 2 Mus musculus | 571.43 | 19.01 | 89449 | 10 | 10 | 4176299.25 | 2433391 | 3841125 | 3986160.25 | 4094781.25 | 2837021 | 2115625.75 | 3126486.75 | 2309765 | 3566617.25 | 2210471.5 | 2219441.75 | 3385693.75 | 3173684.75 | 3141249.5 |
| Q99MB1 | Tlr3 | Toll-like receptor 3 Mus musculus | 130.23 | 4.64 | 103671 | 3 | 3 | 1 | 289401.53 | 257530.61 | 1 | 1 | 1 | 1 | 359711.91 | 467623.62 | 1 | 233424.7 | 299428.03 | 270601.1625 | 376826.84 | 172725.08 |
| P58681 | Tlr7 | Toll-like receptor 7 Mus musculus | 670.44 | 16.38 | 121837 | 14 | 14 | 9379617 | 8604142 | 9166527 | 10330100 | 8842673 | 7084208 | 7745215.5 | 8023693.5 | 7613674.5 | 9231597 | 10090106 | 11624013 | 8499927 | 9060215 | 12543507 |
| P58021 | Tm9sf2 | Transmembrane 9 superfamily member 2 Mus musculus | 278.51 | 9.37 | 75330 | 5 | 5 | 13855640 | 12156082 | 12090649 | 9590521 | 11324615 | 11261947 | 9247238 | 12940331 | 15473517 | 8904592 | 11874275 | 11825741 | 9817219 | 12888621 | 11845877 |
| Q9ET30 | Tm9sf3 | Transmembrane 9 superfamily member 3 Mus musculus | 260.78 | 14.65 | 67545 | 6 | 6 | 7030180 | 6193510.5 | 4196395.5 | 4461002.5 | 7014155 | 8680897 | 5177240 | 5348909 | 6398025 | 5213541 | 5083022 | 7886623 | 7982915 | 5762371 | 4544794.5 |
| Q9CR02 | Tma16 | Translation machinery-associated protein 16 Mus musculus | 97.78 | 17.19 | 25782 | 2 | 2 | 2082810.12 | 1679686.38 | 1689868.88 | 2088082.88 | 2318952.25 | 1 | 1882131.25 | 709378.19 | 1 | 1 | 1672875.62 | 1105107.5 | 2073011.38 | 746404.31 | 1399349.703 |
| Q7TQ69 | Tmc3 | Transmembrane channel-like protein 3 Mus musculus | 36.33 | 0.62 | 128691 | 1 | 1 | 1 | 236465.83 | 695763.5 | 361556 | 1 | 411150.22 | 2795173 | 437173.56 | 1151786.54 | 963649.38 | 649359.62 | 938158.88 | 520808.34 | 258791.66 | 2074597 |
| Q5I0H4 | Tmco1 | Calcium load-activated calcium channel Rattus norvegicus | 180.96 | 19.68 | 21175 | 3 | 3 | 3399643.25 | 2650346.5 | 3606003 | 3552383 | 5420269 | 5345404 | 5156370 | 3167469 | 3806158.75 | 5689660 | 5073249 | 4603659 | 4276126 | 3239956.25 | 6491190.5 |
| Q63584 | Tmed10 | Transmembrane emp24 domain-containing protein 10 Rattus norvegicus | 529.49 | 31.96 | 24858 | 6 | 6 | 40057721 | 39002757 | 39721865 | 38627953 | 39894673 | 43361621 | 36950153 | 42159661 | 48148473 | 35565925 | 39568193 | 34286705 | 39431869 | 38816513 | 31766693 |
| Q63524 | Tmed2 | Transmembrane emp24 domain-containing protein 2 Rattus norvegicus | 189.89 | 16.92 | 22733 | 3 | 3 | 3422113.75 | 3281659.5 | 3057820 | 2860088 | 4130505 | 3833309.25 | 3060001.5 | 3501323 | 3885015.75 | 3280067 | 4057153.75 | 3533265.75 | 3613287.5 | 3557531 | 2895905 |
| Q8R1V4 | Tmed4 | Transmembrane emp24 domain-containing protein 4 Mus musculus | 140.56 | 16.3 | 26022 | 2 | 3 | 2765311.5 | 5435591 | 6167950 | 5655069.5 | 2494016.5 | 3412021.5 | 7345674.5 | 4944779 | 2276392.25 | 4747698 | 2583307.75 | 2629604 | 5849579 | 4188806 | 2889706 |
| Q6AXN3 | Tmed5 | Transmembrane emp24 domain-containing protein 5 Rattus norvegicus | 163.24 | 10.04 | 26136 | 2 | 3 | 1766584.25 | 3235119.75 | 1965206.5 | 1636073.38 | 1891252 | 2396357 | 1883187.62 | 2375637.25 | 2178638.25 | 2201073 | 4154040.5 | 2063419.88 | 1847076.75 | 3723469.25 | 2658284 |
| D3ZTX0 | Tmed7 | Transmembrane emp24 domain-containing protein 7 Rattus norvegicus | 219.34 | 24.34 | 25479 | 4 | 4 | 3980779.5 | 4075977 | 4367660 | 3285284.25 | 4314586 | 4586953.5 | 2326771.25 | 4833457 | 6993833 | 3435006.25 | 4671310 | 4567411.5 | 3002524.5 | 4269741 | 4129274 |
| Q5I0E7 | Tmed9 | Transmembrane emp24 domain-containing protein 9 Rattus norvegicus | 225.67 | 20 | 27028 | 4 | 5 | 27894041 | 27263373 | 25066293 | 24434101 | 24617893 | 26948341 | 23535077 | 25655101 | 27429665 | 23196233 | 22401157 | 22268675 | 21987895 | 24936635 | 19726717 |
| Q8VC04 | Tmem106a | Transmembrane protein 106A Mus musculus | 138.31 | 14.56 | 29109 | 3 | 3 | 8038553.5 | 10927428 | 7606400.5 | 10944657 | 11518702 | 10527926 | 9443261 | 11144524 | 10712117 | 9079064 | 5618031 | 9445599 | 11357999 | 7643185 | 7495882 |
| Q6AYA5 | Tmem106b | Transmembrane protein 106B Rattus norvegicus | 52.41 | 5.82 | 31152 | 1 | 1 | 1 | 808104.06 | 1153426 | 1 | 1 | 1283176.25 | 1468533.88 | 1 | 1409474.75 | 1 | 984498.5 | 1 | 1 | 1133895.75 | 1038747.25 |
| Q3UBX0 | Tmem109 | Transmembrane protein 109 Mus musculus | 208.92 | 11.93 | 26306 | 3 | 3 | 11520050 | 8938495 | 7579169.5 | 7653542 | 8806654 | 9022244 | 7361057.5 | 9620886 | 10214941 | 8469681 | 6491364 | 7773021.5 | 9383861 | 8499415 | 6723520 |
| B0BN86 | Tmem11 | Transmembrane protein 11, mitochondrial Rattus norvegicus 2 | 39.37 | 4.21 | 21342 | 1 | 1 | 1 | 841602.56 | 1191889.12 | 755534.25 | 881622.19 | 1110277.25 | 878418.56 | 1512629.88 | 1728173.62 | 871786.75 | 1276877.75 | 1314066.12 | 922817.44 | 1309639.88 | 910818.19 |
| Q9D1R1 | Tmem126b | Complex I assembly factor TMEM126B, mitochondrial Mus musculus | 48.71 | 7.83 | 25444 | 1 | 1 | 452009.94 | 505386.88 | 705776.31 | 524030.79 | 432950.03 | 1 | 221344.66 | 597179.94 | 616770.69 | 1 | 451473.84 | 239190.86 | 432896.62 | 606434.25 | 432498.8925 |
| Q9CQN6 | Tmem14c | Transmembrane protein 14C Mus musculus | 340.79 | 92.11 | 11642 | 5 | 5 | 8133633 | 8748256 | 8877349 | 8262232 | 6547783.5 | 8909549 | 7213107 | 8735069 | 12139177 | 8142869.5 | 7961475.5 | 9064509 | 7747802 | 10660888 | 7872670 |
| Q9D938 | Tmem160 | Transmembrane protein 160 Mus musculus | 45.37 | 3.72 | 19587 | 1 | 1 | 1820692.62 | 2003869.12 | 1692980 | 1684022.88 | 1704514.62 | 1630150.12 | 1566000.62 | 1492078.75 | 1943340.75 | 1780485.38 | 1517364.38 | 1316195.25 | 1553558.38 | 1669387.75 | 1311380.88 |
| P52875 | Tmem165 | Transmembrane protein 165 Mus musculus | 128.09 | 13.31 | 34791 | 2 | 2 | 1156994.62 | 1776877 | 2666680.5 | 1616710.88 | 1752277.88 | 2378381 | 1560377.12 | 2366160.5 | 2824837 | 1203024 | 1637259.75 | 2603311.75 | 2490310.75 | 1938500.75 | 2540753 |
| Q9CR64 | Tmem167a | Protein kish-A Mus musculus | 45.37 | 12.5 | 8074 | 1 | 1 | 1144087.38 | 278626.41 | 697241.62 | 661018.47 | 524118.47 | 518566.97 | 510715.22 | 455602.81 | 565911 | 512699 | 672907.12 | 534568.5 | 620388 | 334776.5 | 742189.12 |
| Q5PQM0 | Tmem168 | Transmembrane protein 168 Rattus norvegicus | 168.83 | 7.32 | 79772 | 4 | 4 | 2888776 | 435973.72 | 1 | 1 | 387919.75 | 2047417 | 305151.16 | 1767174.25 | 806082.19 | 883403.75 | 1212000.38 | 432972.25 | 1 | 509725.25 | 1 |
| Q3TBT3 | Tmem173 | Stimulator of interferon genes protein Mus musculus | 106.37 | 11.9 | 42830 | 3 | 3 | 3873826.25 | 2567173 | 2623640.25 | 2622259.75 | 2671752.25 | 2014190.75 | 2866758.25 | 2512490 | 2507934 | 2173869.25 | 1850743.62 | 1777059.12 | 3383542.25 | 2476794.25 | 2707484 |
| Q9R1Q6 | Tmem176b | Transmembrane protein 176B Mus musculus | 199.52 | 10.65 | 28367 | 4 | 4 | 2148956.75 | 2091327.5 | 2594122 | 1331876.12 | 2838658 | 3180977.75 | 2480596.25 | 2615892.25 | 3622178.5 | 1715711.25 | 4051206.5 | 3053607.5 | 2538131.5 | 2677670 | 2823112 |
| Q9D771 | Tmem206 | Transmembrane protein 206 Mus musculus | 190.62 | 17.43 | 40178 | 4 | 4 | 224886.72 | 895148.5 | 1072906.62 | 520046.97 | 596969.31 | 577145.38 | 1124885.75 | 819921.56 | 791365.31 | 626720.75 | 844523.38 | 1323165.5 | 608327.5 | 832242.94 | 1437495.25 |
| A1L1L2 | Tmem214 | Transmembrane protein 214 Rattus norvegicus | 58.63 | 1.9 | 76566 | 1 | 1 | 145575.83 | 1 | 166379.84 | 1 | 1 | 167607.42 | 1 | 154167.31 | 270845.69 | 1 | 171508.94 | 187978.91 | 187508.0875 | 204431 | 186113.5 |
| Q8VEK0 | Tmem30a | Cell cycle control protein 50A Mus musculus | 42.17 | 4.67 | 41061 | 1 | 1 | 1 | 811879.38 | 1072021.5 | 1 | 523757.34 | 666997.44 | 602619.5 | 657832 | 891566.38 | 628587.12 | 763440.62 | 583781.31 | 563033.19 | 742355 | 766000.12 |
| Q9CR67 | Tmem33 | Transmembrane protein 33 Mus musculus | 222.6 | 19.84 | 28031 | 5 | 5 | 13066707 | 10189787 | 9696099 | 9721362 | 11396923 | 12123695 | 9309699 | 12247055 | 13516023 | 11616371 | 10164028 | 10491466 | 11284677 | 11950647 | 9735457 |
| Q9DAV9 | Tmem38b | Trimeric intracellular cation channel type B Mus musculus | 100.84 | 9.93 | 32640 | 2 | 2 | 2071083.38 | 1757264.38 | 2027588.38 | 1166918 | 1924872.12 | 707258.12 | 1763065.25 | 1941456 | 2662866.75 | 1936174 | 1255524.88 | 1697544 | 1823465.25 | 2185037 | 1914483.62 |
| Q9DBS1 | Tmem43 | Transmembrane protein 43 Mus musculus | 609.13 | 33.5 | 44783 | 11 | 11 | 9646623 | 8312537 | 7369328.5 | 7296842.5 | 10582217 | 11876304 | 12611603 | 7950107 | 7707695.5 | 10613247 | 9056261 | 8562108 | 10118799 | 8330701.5 | 8766651 |
| Q4VAE3 | Tmem65 | Transmembrane protein 65 Mus musculus | 36.81 | 4.7 | 24918 | 1 | 1 | 605327.5 | 993269 | 657702.12 | 560369.31 | 616304.94 | 753278.88 | 494299.16 | 665936.88 | 789535.81 | 554691.75 | 368151.53 | 500033.31 | 573944.81 | 741270.62 | 510765.03 |
| B9EKI3 | Tmf1 | TATA element modulatory factor Mus musculus | 49.99 | 1.37 | 121803 | 1 | 1 | 971443.31 | 1054957.62 | 913891.81 | 1043313.94 | 967069.44 | 1230952.25 | 924115.06 | 1134322.5 | 1262078.12 | 839566.31 | 879737.19 | 1094448.75 | 737850.19 | 1088220.62 | 967523.38 |
| P49813 | Tmod1 | Tropomodulin-1 Mus musculus | 76.86 | 7.52 | 40466 | 2 | 2 | 1126163.25 | 1262612.5 | 1341541.12 | 1705706.75 | 1437078.62 | 1431257.5 | 1514628.88 | 1192582.62 | 1128837.38 | 1169178 | 1147476.75 | 1067749.62 | 1476690 | 1121245.88 | 1203290.563 |
| Q9JHJ0 | Tmod3 | Tropomodulin-3 Mus musculus | 473.86 | 28.41 | 39503 | 8 | 8 | 9038000 | 9776202 | 8172756 | 9641553 | 9345298 | 10047767 | 10262123 | 8629653 | 8236387 | 11242734 | 9267182 | 10141959 | 11012855 | 7528836 | 8186914.5 |
| Q61029 | Tmpo | Lamina-associated polypeptide 2, isoforms beta/delta/epsilon/gamma Mus musculus | 870.19 | 43.58 | 50373 | 8 | 14 | 22853437 | 25097985 | 24252831 | 27683307 | 23396299 | 22366935 | 23885211 | 20599997 | 18926121 | 27132291 | 21269833 | 25795455 | 24930807 | 19033327 | 24158599 |
| Q61033 | Tmpo | Lamina-associated polypeptide 2, isoforms alpha/zeta Mus musculus | 726.2 | 27.85 | 75168 | 6 | 12 | 10506329 | 10680562 | 11460652 | 12180816 | 10627226 | 10639192 | 9933948 | 9755647 | 7554904.5 | 13514693 | 11244121 | 10091486 | 10846937 | 7279483 | 11938616 |
| Q8VBT0 | Tmx1 | Thioredoxin-related transmembrane protein 1 Mus musculus | 230.44 | 17.27 | 31396 | 5 | 5 | 12106276 | 12540248 | 11158372 | 10139955 | 10436282 | 12124070 | 10330554 | 13266288 | 15825128 | 10028213 | 12497911 | 10742115 | 10930295 | 13739903 | 10584543 |
| Q5XIK2 | Tmx2 | Thioredoxin-related transmembrane protein 2 Rattus norvegicus | 67.31 | 5.08 | 33866 | 1 | 1 | 290727.25 | 284959.56 | 230181.39 | 201141.98 | 298347.53 | 308454.22 | 214079.53 | 202978.12 | 177729.73 | 313262.97 | 164659.52 | 146422.12 | 230294.78 | 229968.8 | 165014.75 |
| Q8BXZ1 | Tmx3 | Protein disulfide-isomerase TMX3 Mus musculus | 404.66 | 19.3 | 51848 | 6 | 6 | 5363058 | 5698235 | 5681977.5 | 6452302.5 | 5437719.5 | 6346432.5 | 6310491 | 6350472 | 6484533.5 | 5718502 | 3760506.5 | 5398581 | 4248813 | 6190207 | 3496077 |
| Q8C0L0 | Tmx4 | Thioredoxin-related transmembrane protein 4 Mus musculus | 133.29 | 11.34 | 37131 | 3 | 3 | 2123360.25 | 2472592.25 | 2666886.75 | 2784210 | 2305559.25 | 2829305 | 3001017.25 | 3217444.5 | 3160461.75 | 3169392 | 2631051.25 | 2506664 | 2530072.25 | 2645566.5 | 1942877 |
| Q61333 | Tnfaip2 | Tumor necrosis factor alpha-induced protein 2 Mus musculus | 61.25 | 1.74 | 78102 | 1 | 1 | 404323.91 | 502148.03 | 584102.56 | 275638.88 | 300789.28 | 410517.12 | 378936.91 | 368920.06 | 511584.56 | 215448.28 | 853703.94 | 598794 | 371795.97 | 461625.34 | 628856.5 |
| Q921Z5 | Tnfaip8 | Tumor necrosis factor alpha-induced protein 8 Mus musculus | 48.55 | 10.1 | 22960 | 1 | 1 | 1 | 1 | 200955.91 | 1 | 1 | 212538.22 | 1 | 339603.62 | 195759.8 | 320066.66 | 1 | 1 | 1 | 179897.83 | 371903.75 |
| Q9D8Y7 | Tnfaip8l2 | Tumor necrosis factor alpha-induced protein 8-like protein 2 Mus musculus | 111.73 | 26.09 | 20615 | 3 | 3 | 474344.91 | 1228828.75 | 325598.09 | 1 | 1 | 1 | 824359.62 | 952847.81 | 234298.17 | 1 | 114838.09 | 562517.19 | 1 | 131830.33 | 1 |
| Q8BFY9 | Tnpo1 | Transportin-1 Mus musculus | 916.78 | 28.73 | 102357 | 13 | 18 | 39307329 | 40928837 | 39648933 | 39648505 | 41273549 | 45209493 | 39987253 | 46967865 | 48067793 | 38546409 | 41422205 | 40707365 | 38270489 | 48051985 | 41011189 |
| Q99LG2 | Tnpo2 | Transportin-2 Mus musculus | 428.93 | 10.6 | 100456 | 3 | 8 | 18682063 | 19357955 | 19835205 | 20231513 | 20493809 | 21667879 | 21239207 | 20934807 | 22627675 | 20314281 | 20325561 | 18779817 | 20385281 | 22349967 | 20408797 |
| Q6P2B1 | Tnpo3 | Transportin-3 Mus musculus | 427.89 | 17.12 | 104170 | 10 | 10 | 9316738 | 6364965.5 | 9056856 | 8481058 | 9776535 | 9983824 | 6875405 | 9064673 | 12351942 | 7281931.5 | 7654625.5 | 9752371 | 6136645.5 | 9268264 | 8666553 |
| Q5SSZ5 | Tns3 | Tensin-3 Mus musculus | 44.72 | 0.97 | 155589 | 1 | 1 | 596497.81 | 404810.47 | 723231.25 | 484870.19 | 789873 | 1 | 1016857.31 | 1 | 742882 | 833255.5 | 1 | 387263.41 | 1 | 1 | 1 |
| Q9D2E2 | Toe1 | Target of EGR1 protein 1 Mus musculus | 113.89 | 6.26 | 56871 | 3 | 3 | 289474.69 | 192869.33 | 215007.33 | 164737.58 | 1 | 184560.94 | 386413.84 | 338972.03 | 288333.84 | 1056405.75 | 261196.12 | 890978.12 | 405956.1825 | 294746.66 | 176903.83 |
| A2RUW1 | Tollip | Toll-interacting protein Rattus norvegicus | 127.83 | 8.39 | 30315 | 2 | 2 | 1371221.5 | 351292.06 | 1948126.5 | 483997.88 | 1203978.62 | 607098.25 | 2858348 | 853853.75 | 1486513.563 | 1626754.25 | 557377.12 | 1472439.25 | 537304.31 | 337443.03 | 1458811.38 |
| O88746 | Tom1 | Target of Myb protein 1 Mus musculus | 362.27 | 20.12 | 54325 | 6 | 6 | 5696313 | 5496928 | 5123093 | 6027098.5 | 5160333.5 | 5732918.5 | 6058093 | 5632882.5 | 5123059 | 6468515.5 | 4492056 | 5434134 | 6425549 | 5831767 | 5871981 |
| Q62760 | Tomm20 | Mitochondrial import receptor subunit TOM20 homolog Rattus norvegicus | 122.93 | 44.14 | 16284 | 3 | 3 | 288771.03 | 299915.03 | 654164.56 | 291334.91 | 182377.92 | 321020 | 223137.39 | 380418.81 | 661447.5 | 267247.06 | 358588.34 | 1048273.5 | 200187.25 | 518334.59 | 365377.56 |
| Q9CPQ3 | Tomm22 | Mitochondrial import receptor subunit TOM22 homolog Mus musculus | 247.17 | 42.25 | 15537 | 4 | 4 | 7626078 | 8556125 | 8900279 | 8178021 | 8377904.5 | 9541206 | 7787068 | 9433737 | 10958023 | 8177177 | 6281855.5 | 9007589 | 7795993.5 | 9019957 | 7760330.5 |
| Q9CYG7 | Tomm34 | Mitochondrial import receptor subunit TOM34 Mus musculus | 659.61 | 45.95 | 34278 | 11 | 11 | 12720540 | 14699377 | 15130840 | 12742153 | 14684081 | 15213853 | 14159056 | 15551760 | 16411531 | 15532522 | 15337081 | 14836868 | 14846759 | 14637847 | 14971549 |
| Q9QYA2 | Tomm40 | Mitochondrial import receptor subunit TOM40 homolog Mus musculus | 481.47 | 29.92 | 37895 | 7 | 7 | 7149887.5 | 8180150 | 8028069 | 8924805 | 9096925 | 9588007 | 9049245 | 8519370 | 9378713 | 9378623 | 7094507 | 9020132 | 8635037 | 8377884 | 10281954 |
| B1AXP6 | Tomm5 | Mitochondrial import receptor subunit TOM5 homolog Mus musculus | 38.98 | 19.61 | 6037 | 1 | 1 | 598000.94 | 803049.94 | 523047.84 | 456146.28 | 518794.09 | 718559.06 | 449265.47 | 725946.88 | 624454.38 | 656465.88 | 405318.81 | 517375.5 | 529965.5375 | 811684.56 | 385483.28 |
| Q9CZW5 | Tomm70 | Mitochondrial import receptor subunit TOM70 Mus musculus | 521.79 | 17.18 | 67590 | 10 | 10 | 26322325 | 23164519 | 18914515 | 23519605 | 21850621 | 20017483 | 22704781 | 22669715 | 20348947 | 21919279 | 22485569 | 18791571 | 19476237 | 20118605 | 18564623 |
| Q04750 | Top1 | DNA topoisomerase 1 Mus musculus | 448.38 | 16.3 | 90876 | 9 | 9 | 11616658 | 13068545 | 12492185 | 10531881 | 10801429 | 12196587 | 13236767 | 10963562 | 13116817 | 11022379 | 11347401 | 11477448 | 10487253 | 11379737 | 13653849 |
| Q01320 | Top2a | DNA topoisomerase 2-alpha Mus musculus | 2130.18 | 33.77 | 172790 | 34 | 41 | 33302683 | 28116785 | 34092885 | 32116889 | 35976841 | 32400713 | 28512015 | 32896829 | 34774341 | 33934049 | 39159097 | 35024309 | 34486673 | 30353901 | 33719861 |
| Q64511 | Top2b | DNA topoisomerase 2-beta Mus musculus | 581.87 | 9.49 | 181909 | 6 | 13 | 6199150.5 | 5327997 | 6893181 | 5923109 | 7973737 | 5986190 | 4981956.5 | 7540133 | 8251194 | 6226945.5 | 7022698.5 | 7929832 | 6481390 | 5326236 | 8473331 |
| Q68G38 | Tor1a | Torsin-1A Rattus norvegicus | 149.4 | 11.41 | 37949 | 3 | 3 | 2457221.25 | 1550796.75 | 1719114.25 | 1065484.25 | 1840050.75 | 1842331.62 | 1804756.5 | 1809780.88 | 2686083.75 | 1626934.38 | 1199671.75 | 2481156 | 110296.74 | 1605986.12 | 1795257.5 |
| Q921T2 | Tor1aip1 | Torsin-1A-interacting protein 1 Mus musculus | 570.9 | 23.7 | 66781 | 11 | 11 | 14347833 | 14843691 | 18382027 | 12886996 | 13369572 | 16260650 | 11878577 | 16332538 | 13633764 | 11275262 | 14593823 | 15074722 | 11929503 | 12902045 | 13651780 |
| Q9ER41 | Tor1b | Torsin-1B Mus musculus | 159.29 | 12.8 | 37818 | 3 | 3 | 1876690.88 | 2223194 | 1662219.12 | 2164013.5 | 2329247.75 | 1401242.75 | 2157024.75 | 3018589.25 | 2480806.75 | 2158016.5 | 2094168.62 | 2534087.75 | 1311082.5 | 1839306.38 | 1996608.88 |
| Q9ER38 | Tor3a | Torsin-3A Mus musculus | 189.48 | 15.06 | 43814 | 4 | 4 | 5560227.5 | 4338360 | 3494521.5 | 4100928 | 3502685 | 3472997.25 | 4359784 | 4388087.5 | 4317606.5 | 3941134 | 4417634.5 | 3964678.5 | 4141039 | 4661489 | 3663879 |
| Q8BU11 | Tox4 | TOX high mobility group box family member 4 Mus musculus | 36.05 | 1.62 | 65961 | 1 | 1 | 1 | 365459.31 | 1 | 1 | 233071.77 | 1 | 1 | 1 | 1 | 1 | 250821.56 | 1 | 1 | 456933.31 | 1 |
| Q62393 | Tpd52 | Tumor protein D52 Mus musculus | 484.65 | 43.3 | 24313 | 7 | 7 | 31624619 | 34161057 | 30067463 | 34338641 | 28192053 | 25716123 | 33483949 | 34657817 | 32759097 | 32659513 | 29332065 | 31887379 | 25926199 | 30479785 | 21438259 |
| Q6PCT3 | Tpd52l2 | Tumor protein D54 Rattus norvegicus | 368.03 | 35.45 | 23992 | 6 | 6 | 6379085 | 6059255.5 | 8893107 | 8007671 | 7538736 | 4694506 | 10348797 | 7788530 | 7349329 | 9416194 | 9286249 | 10091888 | 7952104.5 | 7184369.5 | 10839096 |
| P17751 | Tpi1 | Triosephosphate isomerase Mus musculus | 1395.28 | 72.91 | 32192 | 18 | 19 | 491184737 | 665689857 | 585128065 | 657117441 | 671526529 | 467263777 | 738749249 | 646810689 | 620562753 | 581032065 | 701603457 | 588272065 | 681600641 | 679248769 | 626882497 |
| P04692 | Tpm1 | Tropomyosin alpha-1 chain Rattus norvegicus | 483.89 | 17.96 | 32681 | 4 | 9 | 181939681 | 212132929 | 173293713 | 206825809 | 185180545 | 176263473 | 154770017 | 176321905 | 192776481 | 171913697 | 184480081 | 166221889 | 184463713 | 184117905 | 159219905 |
| P58774 | Tpm2 | Tropomyosin beta chain Mus musculus | 253.82 | 12.68 | 32837 | 0 | 5 | 186449921 | 213143153 | 175054945 | 208202273 | 187314593 | 180881825 | 154971393 | 178159969 | 195160673 | 171947073 | 187096369 | 167451729 | 184551809 | 186542049 | 161527297 |
| P21107 | Tpm3 | Tropomyosin alpha-3 chain Mus musculus | 643.1 | 32.63 | 32994 | 3 | 12 | 274021505 | 324433089 | 267948113 | 309771105 | 276667489 | 261179041 | 226744081 | 265845377 | 296999105 | 251781985 | 282090689 | 251969665 | 281866241 | 277943585 | 236680705 |
| Q6IRU2 | Tpm4 | Tropomyosin alpha-4 chain Mus musculus | 1222.06 | 58.87 | 28468 | 9 | 20 | 171244289 | 181799265 | 161074145 | 175204321 | 158968001 | 175631921 | 150522961 | 174925457 | 204026353 | 164365985 | 218563905 | 181938561 | 162059809 | 190834129 | 169790609 |
| O89023 | Tpp1 | Tripeptidyl-peptidase 1 Mus musculus | 494.53 | 28.29 | 61342 | 3 | 8 | 33809873 | 47168833 | 35362789 | 36958925 | 34351465 | 43300117 | 32882745 | 46825409 | 45889661 | 33317441 | 34461601 | 38624889 | 43918057 | 42330265 | 33193675 |
| Q64514 | Tpp2 | Tripeptidyl-peptidase 2 Mus musculus | 1755.26 | 37.56 | 139879 | 4 | 37 | 51876317 | 46468493 | 61087669 | 54042949 | 48974405 | 46019433 | 50874441 | 58008413 | 56560101 | 46547793 | 54847729 | 61124365 | 46612641 | 50481869 | 54532089 |
| F6ZDS4 | Tpr | Nucleoprotein TPR Mus musculus | 1444.52 | 16.58 | 273990 | 10 | 29 | 19221561 | 24636911 | 26501733 | 24298715 | 23207349 | 22710069 | 29210617 | 17239131 | 16057465 | 28786281 | 23903685 | 25767537 | 20567531 | 18959703 | 17944359 |
| A8WCF8 | Tprg1l | Tumor protein p63-regulated gene 1-like protein Rattus norvegicus | 38.68 | 6.77 | 29844 | 1 | 1 | 1653772.32 | 423464.59 | 650947.94 | 2443804.25 | 3096872.5 | 2699171.75 | 1 | 768015.38 | 1 | 1 | 1 | 1 | 2646117 | 1 | 1 |
| P63028 | Tpt1 | Translationally-controlled tumor protein Mus musculus | 499.76 | 75.58 | 19462 | 8 | 8 | 26796833 | 23246987 | 23427929 | 17855349 | 18320633 | 28193119 | 18759883 | 30924485 | 38294641 | 15317461 | 31877053 | 29666465 | 19677469 | 32515387 | 23161665 |
| A2APB8 | Tpx2 | Targeting protein for Xklp2 Mus musculus | 145.9 | 6.44 | 85894 | 3 | 3 | 882806.94 | 1429659.75 | 1757683.75 | 1648329.12 | 1584731.88 | 1255869.723 | 1294924.88 | 1097320.25 | 1301016.38 | 1330217.38 | 1637407.62 | 1445823.75 | 1379754.12 | 1 | 1467506.12 |
| Q6PFR5 | Tra2a | Transformer-2 protein homolog alpha Mus musculus | 181.14 | 14.59 | 32316 | 3 | 4 | 5919604 | 5434528 | 3995547.75 | 6059926 | 6816700.5 | 7097843 | 4701206 | 3917886.5 | 4424774 | 4883734.5 | 7054278 | 4403276 | 5743335 | 5169003 | 6051795 |

| | | | | | | | | | | | | | | | | | | | | | | |
|---|---|---|---|---|---|---|---|---|---|---|---|---|---|---|---|---|---|---|---|---|---|---|
| P62996 | Tra2b | Transformer-2 protein homolog beta Mus musculus | 413.94 | 23.96 | 33666 | 5 | 6 | 12482151 | 14761592 | 10882459 | 14648524 | 18349399 | 16965689 | 15747835 | 13915928 | 15037396 | 15767382 | 16017343 | 14509173 | 14733633 | 13820871 | 17360669 |
| Q3U0V2 | Tradd | Tumor necrosis factor receptor type 1-associated DEATH domain protein Mus musculus | 216.01 | 15.16 | 34577 | 3 | 3 | 1922995.5 | 3142418 | 1180384.12 | 1779870.62 | 1701181.12 | 2123301 | 1639157.25 | 1681309.62 | 952472 | 1528565.88 | 1503772.88 | 1823781 | 1691123.5 | 1822388.5 | 2122787.25 |
| Q5XI41 | Tram1 | Translocating chain-associated membrane protein 1 Rattus norvegicus | 186.08 | 7.75 | 43030 | 3 | 3 | 4125778.25 | 825884.38 | 1388401 | 732517.75 | 1169592.12 | 1046699.44 | 1490182.25 | 1022494.31 | 1934479 | 1374669.25 | 1745063 | 1334333.88 | 1358424.38 | 1134652 | 3036751.5 |
| Q9CQN1 | Trap1 | Heat shock protein 75 kDa, mitochondrial Mus musculus | 993.01 | 31.16 | 80209 | 6 | 16 | 16313866 | 14580007 | 20722707 | 15664015 | 18790687 | 19662863 | 21724385 | 19029629 | 17939025 | 16470504 | 16710225 | 19014901 | 18124879 | 16045781 | 19340297 |
| B2RXC1 | Trappc11 | Trafficking protein particle complex subunit 11 Mus musculus | 83.56 | 2.82 | 128396 | 2 | 2 | 935231.8125 | 401060.81 | 1186073.38 | 1523631.5 | 630161.56 | 1178386.12 | 279361.97 | 1 | 796706.56 | 1 | 1239281 | 771620.94 | 836210.4075 | 400939.69 | 933000 |
| Q8K2L8 | Trappc12 | Trafficking protein particle complex subunit 12 Mus musculus | 44.54 | 1.13 | 87694 | 1 | 1 | 927506.655 | 889102.12 | 1309256.88 | 725802.62 | 785865 | 755672 | 957190.31 | 1234793.12 | 1214743.88 | 700093.81 | 761434.19 | 978453.44 | 919606 | 968287.69 | 918634.94 |
| Q9CQA1 | Trappc5 | Trafficking protein particle complex subunit 5 Mus musculus | 84.5 | 16.49 | 20795 | 2 | 2 | 1020425.75 | 847211.19 | 917518.75 | 952143.19 | 691442.5 | 764730.5 | 820612.56 | 978466.38 | 967872.69 | 882920.5325 | 743914.62 | 896702.19 | 836484 | 1036085.69 | 814052.31 |
| Q9D289 | Trappc6b | Trafficking protein particle complex subunit 6B Mus musculus | 88.62 | 18.99 | 17936 | 2 | 2 | 1019057.25 | 429156.5 | 80090.02 | 604001.06 | 674858.69 | 669708.88 | 348098.34 | 777436.31 | 299540.5 | 371742.41 | 317636.56 | 623253.06 | 438841.81 | 784071.75 | 451859.94 |
| Q99NH8 | Trem2 | Triggering receptor expressed on myeloid cells 2 Mus musculus | 177.52 | 12.33 | 24527 | 2 | 2 | 1202371.345 | 1312040.38 | 979611.88 | 1306919.62 | 1210913.5 | 1239831.12 | 1248131.12 | 1158104.75 | 1106845.88 | 1188228.218 | 1248949.38 | 980318.44 | 1101413.12 | 1015033.12 | 1044359.25 |
| Q91XB0 | Trex1 | Three-prime repair exonuclease 1 Mus musculus | 159.45 | 14.01 | 33675 | 3 | 3 | 3858302.5 | 6591903 | 3002202.5 | 5408457.5 | 5267453.5 | 3568175 | 3237596.5 | 3974026.25 | 2732724.25 | 3301940 | 4098353.25 | 3569976 | 5439630 | 4218746.5 | 3847842 |
| Q8BVW3 | Trim14 | Tripartite motif-containing protein 14 Mus musculus | 71.01 | 7.27 | 49641 | 2 | 2 | 1259331.88 | 1518784.5 | 1706086.38 | 852161.62 | 1193682.75 | 142908.38 | 110709.77 | 1213810.12 | 1649694.25 | 895716.81 | 919215.12 | 1179231.12 | 1346947.88 | 1488999.62 | 803284.06 |
| Q61510 | Trim25 | E3 ubiquitin/ISG15 ligase TRIM25 Mus musculus | 42.96 | 1.89 | 71726 | 1 | 1 | 1870693.75 | 1544405.88 | 1901792.25 | 1853437.5 | 1749071.25 | 1716854.38 | 1605197.88 | 1723985.12 | 1649199.25 | 1760053.38 | 1399626.5 | 1408073.75 | 1631595 | 1201017 | 1578895.12 |
| P62603 | Trim26 | Tripartite motif-containing protein 26 Rattus norvegicus 3 | 48.08 | 3.51 | 62631 | 1 | 1 | 205442 | 153655.53 | 1 | 1 | 1 | 1 | 1 | 1 | 1 | 1 | 156318.23 | 144177.09 | 158514.8075 | 220710.89 | 112853.02 |
| Q62318 | Trim28 | Transcription intermediary factor 1-beta Mus musculus | 1250.12 | 35.13 | 88847 | 21 | 21 | 46268641 | 52740009 | 54350965 | 65436005 | 52466577 | 46254985 | 80848049 | 47581129 | 35971877 | 66085017 | 51817641 | 56997617 | 61504965 | 46437297 | 53943689 |
| Q8CH72 | Trim32 | E3 ubiquitin-protein ligase TRIM32 Mus musculus | 54 | 2.29 | 72057 | 1 | 1 | 622830.06 | 621233.69 | 698187.44 | 567710.69 | 2221752 | 602570.56 | 837378.88 | 623572.81 | 763708 | 872225.06 | 974830.19 | 670245.94 | 660891.69 | 787514 | 901792.31 |
| Q99PP7 | Trim33 | E3 ubiquitin-protein ligase TRIM33 Mus musculus | 54.25 | 0.96 | 123843 | 1 | 1 | 1 | 1 | 902001.06 | 630287.94 | 1 | 287052 | 630094.75 | 409762.59 | 461545.69 | 777305.75 | 1190662.75 | 408059.31 | 507616.69 | 835190.56 | 669469.31 |
| Q80VI1 | Trim56 | E3 ubiquitin-protein ligase TRIM56 Mus musculus | 269.54 | 12.4 | 79513 | 5 | 5 | 1026202.38 | 905165 | 1186880.5 | 710034.62 | 1074505 | 1437536.12 | 1079178.12 | 809746.88 | 1084979.5 | 883661.19 | 863091.25 | 758358.62 | 731299.31 | 955416.25 | 1016484.38 |
| Q99KW3 | Triobp | TRIO and F-actin-binding protein Mus musculus | 36.94 | 0.5 | 223368 | 1 | 1 | 1 | 500874.97 | 849372.25 | 1 | 634677.62 | 1 | 1 | 1 | 1 | 1 | 647036.38 | 1 | 408543.66 | 839493.81 | 1 |
| P97531 | Trip10 | Cdc42-interacting protein 4 Rattus norvegicus | 131.09 | 9.51 | 62799 | 3 | 3 | 648270.38 | 886000.44 | 1134428.38 | 1 | 247482.66 | 447203.88 | 1047728.75 | 1028414.62 | 812342.5 | 1283843.5 | 602178.81 | 1160920 | 925131.06 | 1185726.62 | 658808.38 |
| F1LP64 | Trip12 | E3 ubiquitin-protein ligase TRIP12 Rattus norvegicus | 719.57 | 10.32 | 223928 | 14 | 14 | 8402313 | 6882629.5 | 8042867.5 | 6013898 | 7340522.5 | 8652207 | 8060691.5 | 6681868.5 | 8546480 | 6691952 | 9581316 | 6647225.5 | 7751800.5 | 8519384 | 9154526 |
| Q3UA06 | Trip13 | Pachytene checkpoint protein 2 homolog Mus musculus | 95.9 | 5.32 | 48377 | 2 | 2 | 2079211.5 | 3025766.5 | 2844043 | 2085358.38 | 2357648.5 | 2633167 | 2339757.5 | 2611363.75 | 2797482 | 2129685.5 | 2344989 | 2590292.5 | 2369588.5 | 2603204.5 | 953637.06 |
| Q3TX08 | Trmt1 | tRNA (guanine(26)-N(2))-dimethyltransferase Mus musculus | 105.58 | 4.52 | 72350 | 2 | 2 | 1127776.625 | 1147970.75 | 996056.25 | 1157810.12 | 1209269.38 | 891402.88 | 1097255.12 | 1358401.62 | 986048.56 | 959129.81 | 1212774.25 | 1090761.78 | 1005780.12 | 1147032.75 | 997460 |
| A2RSY6 | Trmt1l | TRMT1-like protein Mus musculus | 359.46 | 13.05 | 80861 | 6 | 6 | 2862737 | 2160277 | 1737797.25 | 1551891.12 | 1599637.25 | 3001938.25 | 1805556.5 | 1902800.75 | 2266919.25 | 1693493.5 | 3316958.75 | 2038778 | 2386766 | 2861063.75 | 2776799.5 |
| Q8BNV1 | Trmt2a | tRNA (uracil-5-)-methyltransferase homolog A Mus musculus | 32.54 | 1.74 | 63301 | 1 | 1 | 1 | 1 | 1 | 1 | 1 | 1 | 1 | 1 | 1 | 1 | 1 | 1 | 1 | 1 | 405039.91 |
| Q9D0C4 | Trmt5 | tRNA (guanine(37)-N1)-methyltransferase Mus musculus | 42.14 | 1.8 | 56794 | 1 | 1 | 710043.5 | 916123.88 | 730682.31 | 818078.31 | 1 | 665808.25 | 708809.75 | 661148.19 | 729117.75 | 564157.31 | 859478.81 | 860005.62 | 782223.28 | 833239.19 | 576169.5 |
| Q8CE96 | Trmt6 | tRNA (adenine(58)-N(1))-methyltransferase non-catalytic subunit TRM6 Mus musculus | 243.1 | 10.87 | 55518 | 4 | 4 | 957405.56 | 714771.25 | 1303577.5 | 853337 | 957272.8275 | 1036518.06 | 926655.69 | 1408644.62 | 1379336.75 | 1187788.78 | 1154066.56 | 1105553.62 | 772360.5 | 1297437.62 | 1440914.5 |
| Q8K1J6 | Trnt1 | CCA tRNA nucleotidyltransferase 1, mitochondrial Mus musculus | 380.86 | 26.04 | 49895 | 9 | 9 | 10779536 | 9099076 | 9943824 | 8867748 | 11954073 | 10162082 | 9418134 | 9571325 | 9406133 | 10305651 | 9717292 | 7415869 | 7829989.5 | 8558168 | 10172693 |
| Q9WTR1 | Trpv2 | Transient receptor potential cation channel subfamily V member 2 Mus musculus | 274.13 | 8.86 | 85965 | 4 | 4 | 3648372.5 | 3049682.25 | 2692374.5 | 2306182 | 2552853 | 2758333.5 | 2207849.5 | 3348566 | 2993158.5 | 1816666.5 | 728022.81 | 1358563.62 | 2620795 | 3123710.5 | 3018895.5 |
| Q9CZR8 | Tsfm | Elongation factor Ts, mitochondrial Mus musculus | 443.4 | 48.77 | 35334 | 7 | 7 | 8348572 | 9932174 | 8412015 | 8252476.5 | 8658664 | 10108733 | 8790735 | 9035529 | 10271758 | 9999639 | 7242701 | 8429215 | 9112904 | 7243775.5 | 8881974 |
| Q61187 | Tsg101 | Tumor susceptibility gene 101 protein Mus musculus | 168 | 12.28 | 44124 | 4 | 4 | 3959817.5 | 3640241.25 | 4122413.25 | 3878653.75 | 3800367.5 | 3885619 | 3674698 | 3113393.5 | 3390150.5 | 4150553.25 | 3957939 | 3448351.75 | 3674526.25 | 3264377.75 | 3604940.25 |
| Q62348 | Tsn | Translin Mus musculus | 734.46 | 60.96 | 26201 | 11 | 11 | 48678913 | 39909309 | 40311953 | 36538065 | 36860081 | 36998537 | 33140749 | 36486845 | 44836417 | 36029929 | 37249589 | 35245501 | 35274565 | 37511161 | 35522029 |
| Q9QZE7 | Tsnax | Translin-associated protein X Mus musculus | 385.33 | 42.41 | 32926 | 5 | 8 | 2525334 | 1737998.12 | 2549686 | 2309082 | 2140633.75 | 2335023.75 | 2242932.25 | 2116349.75 | 1973180.62 | 1934555.25 | 1695069.25 | 1564361.88 | 1779394.5 | 1663456.62 | 1869812 |
| Q9CQ88 | Tspan31 | Tetraspanin-31 Mus musculus | 45.3 | 5.71 | 22694 | 1 | 1 | 1429178.5 | 1445725.88 | 1218858.12 | 964262.25 | 1071089.12 | 1582237.25 | 1245494.75 | 1859571.25 | 1487373 | 1337380.62 | 1569959 | 849500.5 | 1163436.88 | 1495029.5 | 1258513.88 |
| P50637 | Tspo | Translocator protein Mus musculus | 236.83 | 46.75 | 18841 | 4 | 4 | 26044009 | 6385051 | 5716660 | 5612279 | 11752173 | 13476809 | 5858911 | 8655651 | 7984624 | 8314511 | 9801099 | 8232562 | 10585081 | 10641561 | 9438609 |
| Q5SWD9 | Tsr1 | Pre-rRNA-processing protein TSR1 homolog Mus musculus | 513.92 | 15.69 | 92105 | 9 | 9 | 3070351 | 3919106.75 | 1639943 | 1836093.25 | 4685954 | 2305951.5 | 3320072.75 | 3946734.25 | 3550158 | 3806495.75 | 2779912.75 | 2943429.5 | 3461055.5 | 2576280 | 2251931.25 |
| Q5XIB1 | Tssc4 | Protein TSSC4 Rattus norvegicus | 39.93 | 3.15 | 33939 | 1 | 1 | 927600.94 | 1171855.62 | 1108860.62 | 1141664 | 1066645.38 | 1009403.38 | 1123632.658 | 1212365.88 | 1186011.25 | 1086750.12 | 840756.88 | 1112481.38 | 1301405.25 | 1130412 | 883463.81 |
| P23591 | Tsta3 | GDP-L-fucose synthase Mus musculus | 230.02 | 20.25 | 35878 | 5 | 5 | 7730363.5 | 8365780 | 7979294 | 8123320 | 8205173.5 | 8520995 | 8138888 | 8981063 | 6627575.5 | 7113156 | 6084183 | 6867531 | 7541159 | 7180785.5 | 7097822 |
| Q9D6K7 | Ttc33 | Tetratricopeptide repeat protein 33 Mus musculus | 78.75 | 4.96 | 29371 | 1 | 1 | 2230320.75 | 2738433.5 | 2508648.25 | 2295292.25 | 2051364 | 2263919.5 | 1875302.38 | 2261485.25 | 2375176.75 | 2205044.25 | 1886599.62 | 2640217.5 | 2131747 | 2267783 | 1889756.25 |
| A3KMP2 | Ttc38 | Tetratricopeptide repeat protein 38 Mus musculus | 56.29 | 1.94 | 52224 | 1 | 1 | 476321.41 | 291528.38 | 521739.12 | 191043.45 | 318513.09 | 429566.5 | 261028.22 | 379350.25 | 481358.66 | 387825.9075 | 367908.62 | 512819.75 | 263319.97 | 405867.69 | 390773.94 |
| Q5BK48 | Ttc5 | Tetratricopeptide repeat protein 5 Rattus norvegicus | 72.23 | 6.36 | 48793 | 2 | 2 | 1 | 412489.59 | 538074.75 | 284557.75 | 1 | 1 | 376552.22 | 437078.88 | 1 | 1 | 247425.3 | 317705.31 | 1 | 1 | 226945.83 |
| Q5NC05 | Ttf2 | Transcription termination factor 2 Mus musculus | 100.85 | 2.9 | 125530 | 3 | 3 | 1898142.75 | 1383248 | 1174040.62 | 1850015.38 | 1632152.38 | 1 | 1081068.25 | 1087629.25 | 259628.09 | 975710 | 1107200.25 | 1 | 695842.94 | 1 | 644047.88 |
| Q3UDE2 | Ttll12 | Tubulin--tyrosine ligase-like protein 12 Mus musculus | 493.99 | 25.67 | 74043 | 11 | 11 | 7946159.5 | 7170459.5 | 9636674 | 8998819 | 6736971 | 9495998 | 11256114 | 8186414.5 | 8854434 | 9160113 | 8907758 | 10485683 | 7385868.5 | 7633900 | 11921222 |
| P68369 | Tuba1a | Tubulin alpha-1A chain Mus musculus | 2138.28 | 75.39 | 50136 | 1 | 31 | 1277217537 | 1397301377 | 1323573889 | 1206727169 | 1333526913 | 1333091329 | 1180298113 | 1420576129 | 1496099585 | 1168114817 | 1320227329 | 1374636289 | 958299265 | 1496678401 | 1207250049 |
| P05213 | Tuba1b | Tubulin alpha-1B chain Mus musculus | 2208.86 | 75.39 | 50152 | 0 | 32 | 1412480385 | 1560608897 | 1517606529 | 1353083393 | 1480593025 | 1521045249 | 1308060033 | 1623391105 | 1695030145 | 1294858497 | 1514424833 | 1552707329 | 1100125313 | 1677692929 | 1382205697 |
| P68373 | Tuba1c | Tubulin alpha-1C chain Mus musculus | 2169.27 | 71.05 | 49909 | 1 | 31 | 1361132033 | 1494732033 | 1480577537 | 1323368193 | 1421069185 | 1459601153 | 1267645953 | 1558584961 | 1633488897 | 1261364097 | 1466869761 | 1498815745 | 1032444289 | 1618890625 | 1332514689 |
| P68368 | Tuba4a | Tubulin alpha-4A chain Mus musculus | 1677.86 | 64.96 | 49924 | 4 | 24 | 900939393 | 1041748737 | 1001077889 | 889242113 | 965179777 | 1029459713 | 850599169 | 1076236801 | 1123653121 | 826096385 | 1051284737 | 1048783553 | 582852993 | 1113703425 | 936016961 |
| Q3UX10 | Tubal3 | Tubulin alpha chain-like 3 Mus musculus 2 | 144.57 | 11.66 | 49988 | 1 | 3 | 18683813 | 24827005 | 17155457 | 15835367 | 20305229 | 23150319 | 30874673 | 18547001 | 28414065 | 23256077 | 32788669 | 20666781 | 25526539 | 30867749 | 24560469 |
| P85108 | Tubb2a | Tubulin beta-2A chain Rattus norvegicus | 2026.68 | 58.2 | 49907 | 1 | 29 | 824646721 | 1311703297 | 1105619969 | 1120288129 | 1217874817 | 1159565057 | 1090367361 | 1160523137 | 1206018689 | 1098114817 | 941171585 | 1080587777 | 1071531329 | 1220109697 | 965659457 |
| Q3KRE8 | Tubb2b | Tubulin beta-2B chain Rattus norvegicus | 2106.08 | 66.97 | 49953 | 1 | 30 | 827297537 | 1315153665 | 1108002561 | 1123333121 | 1220174081 | 1161881857 | 1090426625 | 1159963649 | 1205477505 | 1100893697 | 943384449 | 1082894465 | 1074262529 | 1222876289 | 965307137 |
| Q4QRB4 | Tubb3 | Tubulin beta-3 chain Rattus norvegicus | 1327.39 | 38.89 | 50419 | 2 | 20 | 581574337 | 1012644865 | 854428033 | 848915585 | 927739137 | 853709377 | 806254721 | 882082753 | 937389505 | 827936513 | 647552257 | 816861953 | 774003457 | 939601473 | 706780417 |
| Q9D6F9 | Tubb4a | Tubulin beta-4A chain Mus musculus | 1632.81 | 70.95 | 49586 | 1 | 24 | 580736193 | 981451585 | 832888961 | 809792129 | 898003841 | 862790529 | 823115457 | 864795585 | 909522433 | 813717633 | 628654849 | 814298113 | 750701185 | 903164993 | 725188737 |
| P68372 | Tubb4b | Tubulin beta-4B chain Mus musculus | 2411.1 | 82.92 | 49831 | 1 | 34 | 707777793 | 1169986817 | 995606273 | 1004987777 | 1098371969 | 1054605313 | 1010579841 | 1039106113 | 1075468289 | 1007670465 | 835204033 | 992485505 | 953682369 | 1087687553 | 912449537 |
| P69897 | Tubb5 | Tubulin beta-5 chain Rattus norvegicus | 2650.3 | 83.78 | 49671 | 6 | 36 | 1062513793 | 1632415745 | 1373111809 | 1368667521 | 1503917057 | 1468668161 | 1348673025 | 1443863681 | 1513320193 | 1364355329 | 1235014273 | 1357094529 | 1387411969 | 1530551937 | 1229987585 |
| Q922F4 | Tubb6 | Tubulin beta-6 chain Mus musculus | 1448.46 | 65.32 | 50090 | 9 | 23 | 417234241 | 575988545 | 517882305 | 509164769 | 566127745 | 530666369 | 512356641 | 538753921 | 551671041 | 512247489 | 501503617 | 517376737 | 413400961 | 553170305 | 455124225 |
| P83887 | Tubg1 | Tubulin gamma-1 chain Mus musculus | 151.76 | 15.3 | 51101 | 4 | 4 | 3125687.75 | 3614018 | 8423060 | 3376430 | 3839941 | 3680825.75 | 5155509 | 3765442.5 | 3393449 | 9810308 | 6246369 | 7468662 | 6498992 | 6857555 | 8527668 |
| Q921G8 | Tubgcp2 | Gamma-tubulin complex component 2 Mus musculus | 39.08 | 1.44 | 103223 | 1 | 1 | 1 | 577559.94 | 937959.19 | 1 | 1 | 645492 | 773157.3425 | 968051.31 | 793729.5 | 685356.56 | 628869 | 595931.19 | 806446.75 | 1 | 1 |
| Q8BFR5 | Tufm | Elongation factor Tu, mitochondrial Mus musculus | 527.02 | 37.61 | 49508 | 11 | 11 | 17981583 | 18943269 | 14903769 | 19271393 | 19213115 | 15812158 | 20790057 | 20019351 | 19616253 | 18989637 | 16306313 | 19349837 | 15150787 | 19070097 | 18909461 |
| Q91YR1 | Twf1 | Twinfilin-1 Mus musculus | 266 | 21.71 | 40079 | 5 | 5 | 10499071 | 10866777 | 11384323 | 9589999 | 11015919 | 9276069 | 9695669 | 9453937 | 11832501 | 8508754 | 7927132.5 | 9912957 | 12085080 | 9777834 | 9165444 |
| Q9Z0P5 | Twf2 | Twinfilin-2 Mus musculus | 574.02 | 44.13 | 39471 | 8 | 8 | 7669736 | 5994880.5 | 8146624 | 8317787.5 | 8789805 | 6732471 | 7481480.5 | 5931085 | 6895419 | 7781426 | 7954022.5 | 9507320 | 9844677 | 5805735 | 9629803 |
| Q6PAM1 | Txlna | Alpha-taxilin Mus musculus | 130 | 6.14 | 62369 | 2 | 2 | 1280284.75 | 1384573.938 | 1239255.75 | 1673974.75 | 1344780.5 | 939730.81 | 456411.59 | 1017399.88 | 1012583 | 1036303.62 | 1006096.12 | 1032015.56 | 1 | 1209596.62 | 1 |
| P10639 | Txn | Thioredoxin Mus musculus | 439.25 | 46.67 | 11675 | 5 | 5 | 103845633 | 105214497 | 117619593 | 56874173 | 76428097 | 104567849 | 80126113 | 129106577 | 159096945 | 53170537 | 127661457 | 120621985 | 70417849 | 138031569 | 113653241 |
| P97493 | Txn2 | Thioredoxin, mitochondrial Mus musculus | 73.23 | 9.04 | 18255 | 1 | 1 | 1 | 339997.91 | 244386.84 | 1 | 298630.62 | 1 | 231147.06 | 728198.06 | 1 | 1 | 531020.56 | 224749.58 | 1 | 695750.38 | 1 |
| Q498E0 | Txndc12 | Thioredoxin domain-containing protein 12 Rattus norvegicus | 126.59 | 24.71 | 19019 | 3 | 3 | 1352617.62 | 1492037.12 | 1199635.62 | 1423276.12 | 1626131.5 | 1582512.38 | 1539788.25 | 1395628 | 1327214.38 | 1679391 | 1224672.75 | 1022328.62 | 1734377.12 | 1351775.25 | 1057810.38 |
| Q9CQM5 | Txndc17 | Thioredoxin domain-containing protein 17 Mus musculus | 231.16 | 39.84 | 14015 | 4 | 4 | 10687913 | 11595468 | 12056245 | 11825727 | 11382406 | 11841683 | 15313184 | 11843380 | 11408493 | 13330705 | 11970653 | 13543410 | 13247875 | 12179062 | 14637917 |
| Q91W90 | Txndc5 | Thioredoxin domain-containing protein 5 Mus musculus | 531.49 | 30.22 | 46415 | 9 | 9 | 9467288 | 7339417.5 | 11260229 | 9304043 | 9124493 | 10217056 | 10585147 | 9854201 | 11333547 | 10164267 | 7441138.5 | 10936156 | 9660571 | 11199741 | 10527345 |
| Q8K581 | Txndc9 | Thioredoxin domain-containing protein 9 Rattus norvegicus 2 | 39.36 | 3.98 | 26272 | 1 | 1 | 314342.66 | 1 | 321456.78 | 432841.44 | 1 | 1 | 572175.81 | 334772.31 | 1 | 454566.03 | 520307.06 | 1 | 469282.22 | 361485.06 | 1 |
| Q5M7W1 | Txnip | Thioredoxin-interacting protein Rattus norvegicus | 47.22 | 3.05 | 44018 | 1 | 1 | 1007481.5 | 1452248.25 | 1143807.88 | 740502.25 | 934415.31 | 1306623.62 | 1373269.12 | 1543729.25 | 1121634.38 | 1064064.75 | 1775832 | 1301369.25 | 1546708 | 1302672.62 | 1168256.62 |
| Q8CDN6 | Txnl1 | Thioredoxin-like protein 1 Mus musculus | 487.49 | 48.79 | 32237 | 10 | 10 | 8772278 | 10316230 | 7470956 | 9537057 | 11095900 | 10134067 | 10158061 | 9657881 | 9626485 | 11083354 | 10391118 | 9622422 | 11174851 | 9275972 | 10518274 |
| Q9JMH6 | Txnrd1 | Thioredoxin reductase 1, cytoplasmic Mus musculus | 814.75 | 38.17 | 67084 | 15 | 15 | 34148489 | 29293857 | 37767077 | 30790565 | 24352639 | 32649083 | 32670545 | 43695917 | 43974541 | 29550933 | 37703449 | 38034945 | 29738045 | 40065657 | 35990057 |
| P07607 | Tyms | Thymidylate synthase Mus musculus | 152.05 | 11.73 | 34958 | 3 | 3 | 1808373.75 | 1584692.88 | 1165552.25 | 1332723 | 1349304 | 1213023.5 | 478287.38 | 1210181.88 | 1077251.5 | 1584227.75 | 1468450.25 | 1753491.12 | 1670725.75 | 1499411.62 | 1863635 |
| Q9D883 | U2af1 | Splicing factor U2AF 35 kDa subunit Mus musculus | 309.82 | 22.59 | 27815 | 5 | 5 | 11448645 | 9639451 | 12226973 | 11376870 | 11514768 | 11484760 | 13171850 | 10052024 | 8489219 | 12589071 | 10683995 | 12989195 | 10080796 | 10178822 | 13247418 |
| P26369 | U2af2 | Splicing factor U2AF 65 kDa subunit Mus musculus | 826.54 | 42.74 | 53517 | 12 | 12 | 25919089 | 30357559 | 35058081 | 34916045 | 29659301 | 28332425 | 33459787 | 35408569 | 40334525 | 36172917 | 36312177 | 42281941 | 34605001 | 35972701 | 37405653 |
| Q6NV83 | U2surp | U2 snRNP-associated SURP motif-containing protein Mus musculus | 254.9 | 7.77 | 118261 | 6 | 6 | 1312861.75 | 345402.53 | 1885097.88 | 1140783.75 | 522111.66 | 1259090.75 | 1146227.62 | 1064325.25 | 1407512.38 | 576476.38 | 1117281.88 | 1389533 | 1420393 | 982958 | 646759.75 |
| Q91YN5 | Uap1 | UDP-N-acetylhexosamine pyrophosphorylase Mus musculus | 129.05 | 4.41 | 58609 | 1 | 2 | 6990582.5 | 7103780 | 7469785.5 | 8474543 | 8029916.5 | 6740409.5 | 6641296.5 | 6318650.5 | 7082946.5 | 7396536 | 7611741 | 6770122.5 | 8029114.5 | 7221898 | 7015055 |
| Q3TW96 | Uap1l1 | UDP-N-acetylhexosamine pyrophosphorylase-like protein 1 Mus musculus | 793.49 | 36.09 | 56614 | 13 | 14 | 52067277 | 56779461 | 59052273 | 56250797 | 61191093 | 49473773 | 54484701 | 47004173 | 54581969 | 56645973 | 57049885 | 56047613 | 61579877 | 58761625 | 57923429 |
| Q02053 | Uba1 | Ubiquitin-like modifier-activating enzyme 1 Mus musculus | 2035.76 | 41.68 | 117809 | 29 | 29 | 87735785 | 96284081 | 84422105 | 98210033 | 97630369 | 92623881 | 96570225 | 94844281 | 91320577 | 89745713 | 98728753 | 92323809 | 96062081 | 94481489 | 97126369 |
| Q9Z1F9 | Uba2 | SUMO-activating enzyme subunit 2 Mus musculus | 1202.24 | 44.2 | 70569 | 18 | 18 | 33409885 | 41087249 | 28453777 | 27419885 | 30028393 | 35683721 | 32458669 | 37069721 | 38859401 | 31117107 | 37204417 | 36717225 | 35957381 | 36641593 | 35961137 |
| Q8C878 | Uba3 | NEDD8-activating enzyme E1 catalytic subunit Mus musculus | 520.74 | 36.58 | 51737 | 10 | 10 | 7718246 | 9037193 | 11423203 | 12034932 | 11194233 | 8146454.5 | 10520109 | 12469714 | 10656951 | 10582437 | 9968251 | 11533463 | 9301895 | 12012327 | 10010403 |
| Q8VE47 | Uba5 | Ubiquitin-like modifier-activating enzyme 5 Mus musculus | 207.24 | 16.13 | 44790 | 4 | 4 | 2713038.5 | 2294605.25 | 2945991.75 | 2004612.5 | 3471097.5 | 1941291 | 3685390.75 | 2373504 | 1925530.88 | 3072181 | 3076504 | 2600945.5 | 2126740 | 1799645.88 | 3795073.5 |
| Q8C7R4 | Uba6 | Ubiquitin-like modifier-activating enzyme 6 Mus musculus | 499.16 | 15.19 | 117966 | 11 | 11 | 8222249.5 | 3920951.25 | 5513448.5 | 4615173 | 5408250 | 8226422.5 | 7586854 | 5800503 | 7956004.5 | 4008184 | 8233572 | 7333471.5 | 6508544.5 | 6292510 | 7674551 |
| Q91VX2 | Ubap2 | Ubiquitin-associated protein 2 Mus musculus | 250.7 | 6.18 | 117966 | 5 | 5 | 1 | 1 | 1018405.44 | 1134142.88 | 1689859.38 | 1194491.25 | 1510741.5 | 1742725 | 1 | 1 | 1630513.62 | 1322991.88 | 1445380.12 | 2163855 | 1774518.88 |
| Q80X50 | Ubap2l | Ubiquitin-associated protein 2-like Mus musculus | 618.97 | 16.35 | 116799 | 11 | 11 | 21484491 | 20054767 | 21571233 | 20913437 | 22209131 | 22605747 | 23541005 | 20933659 | 19001039 | 15859269 | 22896645 | 20072547 | 20742169 | 20485641 | 20731697 |
| Q9D1C1 | Ube2c | Ubiquitin-conjugating enzyme E2 C Mus musculus | 111.75 | 17.32 | 19606 | 2 | 2 | 2829670.25 | 3431147.5 | 39140193 | 2226269.75 | 64193329 | 3178502.75 | 2604888.75 | 1943988.75 | 53412549 | 2936184 | 2459716.5 | 2356225 | 56944209 | 55807617 | 56079765 |
| P61078 | Ube2d3 | Ubiquitin-conjugating enzyme E2 D3 Rattus norvegicus 2 | 177.74 | 36.05 | 16687 | 3 | 3 | 11385062 | 11975149 | 10739178 | 10172582 | 11948288 | 12036330 | 11880317 | 10865994 | 11329213 | 11089898 | 12212648 | 11691495 | 11433167 | 11168481 | 12355805 |
| P63280 | Ube2i | SUMO-conjugating enzyme UBC9 Mus musculus | 286.27 | 53.8 | 18007 | 6 | 6 | 11402778 | 14158973 | 13540894 | 13210278 | 15421310 | 14525035 | 14256855 | 12134867 | 13236761 | 13465874 | 13353181 | 14180849 | 14195612 | 13653193 | 16858533 |
| Q9JJZ4 | Ube2j1 | Ubiquitin-conjugating enzyme E2 J1 Mus musculus | 128.33 | 11.01 | 34990 | 3 | 3 | 2193777.5 | 2154871.5 | 646798 | 3101341 | 3452234.25 | 2334088 | 1870255.75 | 850447.81 | 2297878.25 | 3702004.5 | 4018090.75 | 2130910 | 2272742.75 | 2432519.5 | 1876706 |
| P61087 | Ube2k | Ubiquitin-conjugating enzyme E2 K Mus musculus | 292.54 | 38.5 | 22407 | 5 | 5 | 12695037 | 12623145 | 12040264 | 11796697 | 11388548 | 12821945 | 13787842 | 11389553 | 11751147 | 12675031 | 13058544 | 7125433 | 12212328 | 12832077 | 12330295 |
| P68037 | Ube2l3 | Ubiquitin-conjugating enzyme E2 L3 Mus musculus | 392.22 | 59.09 | 17862 | 6 | 6 | 18667483 | 24371525 | 25664853 | 18493715 | 24950803 | 21550113 | 23888057 | 21199681 | 22249693 | 18249269 | 23789589 | 16995725 | 26518091 | 25965303 | 25384233 |
| P61082 | Ube2m | NEDD8-conjugating enzyme Ubc12 Mus musculus | 304.42 | 35.52 | 20900 | 6 | 6 | 15813014 | 15193225 | 18217401 | 15580575 | 21075753 | 21098215 | 16356279 | 21813355 | 22093951 | 16873581 | 12874356 | 14946075 | 16407887 | 19831733 | 20421671 |
| P61089 | Ube2n | Ubiquitin-conjugating enzyme E2 N Mus musculus | 301.24 | 56.58 | 17138 | 6 | 7 | 12204837 | 13243491 | 12766043 | 11743315 | 13561907 | 12879577 | 14163993 | 13374221 | 11479909 | 13373061 | 11536524 | 12264428 | 12678322 | 11731045 | 12367930 |
| Q6ZPJ3 | Ube2o | (E3-independent) E2 ubiquitin-conjugating enzyme UBE2O Mus musculus | 94.23 | 2.56 | 140834 | 2 | 2 | 855169.88 | 1104413.5 | 1068413.38 | 895552.6425 | 554213.81 | 704823.25 | 1183445.5 | 796826.31 | 832269.5 | 1455877 | 974041.12 | 612627.69 | 886475.265 | 1077055 | 882177.25 |
| Q7TSS2 | Ube2q1 | Ubiquitin-conjugating enzyme E2 Q1 Mus musculus | 159.62 | 8.77 | 46173 | 3 | 3 | 852529.38 | 358739.12 | 1268407.12 | 1 | 1 | 852839.56 | 702386.7825 | 515115.12 | 1266972.75 | 174619.7 | 570054.69 | 549850.81 | 735263.655 | 1051940.62 | 769208.5 |
| Q9CQ37 | Ube2t | Ubiquitin-conjugating enzyme E2 T Mus musculus | 110.46 | 9.31 | 22975 | 1 | 2 | 2891851 | 3155219.25 | 3087698.25 | 2853902.25 | 2370678 | 2808933.75 | 3673465.5 | 3152667 | 2596274 | 2987371.75 | 2978929.75 | 2667025.5 | 3195386.25 | 2669010.75 | 2803816.25 |
| Q9CZY3 | Ube2v1 | Ubiquitin-conjugating enzyme E2 variant 1 Mus musculus | 250.93 | 29.25 | 16355 | 4 | 4 | 18891277 | 19778333 | 22533873 | 21941629 | 18727319 | 17837437 | 25699129 | 19049917 | 18422301 | 23075607 | 22590255 | 21336949 | 22597991 | 20073357 | 22482157 |
| Q3B7D1 | Ube2z | Ubiquitin-conjugating enzyme E2 Z Rattus norvegicus | 89.35 | 8.71 | 38352 | 2 | 2 | 1019622.62 | 1025016.25 | 1030110.81 | 835277.12 | 1065174.25 | 1160690.25 | 1009227.62 | 872550.75 | 1008114.44 | 1120296.75 | 1247358.25 | 875441.75 | 1053782.5 | 1187342.75 | 908059.12 |
| O08759 | Ube3a | Ubiquitin-protein ligase E3A Mus musculus | 266.1 | 7.47 | 99819 | 5 | 5 | 27386849 | 20405337 | 25101607 | 27857019 | 29841137 | 33784813 | 36036953 | 32921333 | 21070223 | 36375017 | 39138753 | 34748021 | 38388857 | 27940439 | 29675299 |
| Q9ES00 | Ube4b | Ubiquitin conjugation factor E4 B Mus musculus | 102.79 | 4.01 | 133318 | 3 | 3 | 439971 | 962346 | 1530228.12 | 872385.12 | 728183.56 | 1265487.38 | 804700.25 | 1332053.5 | 1598718.88 | 939348.5 | 1753108 | 1543893.62 | 1069739.5 | 1625496 | 1515047 |
| Q9EPV8 | Ubl5 | Ubiquitin-like protein 5 Mus musculus | 38.38 | 12.33 | 8547 | 1 | 1 | 3824312.25 | 3188218.25 | 2980332.5 | 2408776.25 | 3759906.25 | 3431434.75 | 352473.09 | 3015636 | 2991068 | 1282030.62 | 3215392.25 | 2608279.735 | 3186429.25 | 3319840.25 | 711457.19 |
| Q99NB8 | Ubqln4 | Ubiquilin-4 Mus musculus | 147.67 | 11.58 | 63506 | 3 | 3 | 933609.69 | 1449453.75 | 1192275.25 | 987305.8925 | 373884.88 | 1218807.25 | 1050563.12 | 1324641 | 1214098.62 | 858728.75 | 609170.5 | 817898.75 | 691865.56 | 810400.75 | 1253247.25 |
| Q6WKZ8 | Ubr2 | E3 ubiquitin-protein ligase UBR2 Mus musculus | 84.28 | 0.74 | 199155 | 1 | 1 | 387065.75 | 498156.81 | 519618.09 | 286536.16 | 201287.05 | 388002.34 | 182311.23 | 406300.94 | 395944.56 | 238315.92 | 739426.44 | 496685.62 | 340681.28 | 602807.62 | 517013.47 |
| A2AN08 | Ubr4 | E3 ubiquitin-protein ligase UBR4 Mus musculus | 1210.58 | 6.12 | 572290 | 22 | 22 | 13464246 | 15326411 | 13744404 | 14036837 | 11869789 | 14024235 | 14150882 | 16705946 | 15258322 | 12945340 | 16425964 | 13937296 | 14578530 | 14820147 | 16833023 |
| Q62671 | Ubr5 | E3 ubiquitin-protein ligase UBR5 Rattus norvegicus | 126.4 | 1.97 | 308027 | 2 | 2 | 1 | 184980.03 | 406640.84 | 525496.25 | 1 | 1 | 1 | 1 | 1 | 1 | 1 | 388104.53 | 563721.06 | 583355.38 | 1 |
| P25976 | Ubtf | Nucleolar transcription factor 1 Mus musculus | 94.05 | 3.27 | 89509 | 2 | 2 | 1509451.88 | 1644149 | 1590948.5 | 1521410.88 | 2415347 | 1099718 | 1 | 1281186.25 | 1352014.25 | 1 | 1406312 | 1161178 | 1347687.38 | 1292350.5 | 2136781 |
| Q922Y1 | Ubxn1 | UBX domain-containing protein 1 Mus musculus | 390.34 | 34.34 | 33573 | 5 | 6 | 6506754.5 | 9303493 | 4879799 | 10758774 | 5061277.5 | 7275944 | 10473761 | 7380027.5 | 6640979 | 10870327 | 7457899.5 | 7958892.5 | 9438663 | 5191939.5 | 6533180 |
| Q8VCH8 | Ubxn4 | UBX domain-containing protein 4 Mus musculus | 238.39 | 12.85 | 56472 | 4 | 4 | 3544424.25 | 4576210.5 | 3864571 | 3838892.75 | 4312834.5 | 3625390.75 | 3007256.25 | 3877575.75 | 4725319 | 3060386.75 | 3172478.25 | 3415360 | 4167480.5 | 3798845.25 | 2514706.5 |

| | | | | | | | | | | | | | | | | | | | | | | |
|---|---|---|---|---|---|---|---|---|---|---|---|---|---|---|---|---|---|---|---|---|---|---|
| Q99PL6 | Ubxn6 | UBX domain-containing protein 6 Mus musculus | 84.89 | 6.33 | 49796 | 2 | 2 | 2164644 | 2293407.75 | 2197554 | 2322861 | 724293.75 | 782500.88 | 2169799 | 1476616.75 | 2456077 | 2199571.75 | 2182034.5 | 2077556.5 | 2617061.5 | 2740204.25 | 1961770.12 |
| Q9JKB1 | Uchl3 | Ubiquitin carboxyl-terminal hydrolase isozyme L3 Mus musculus | 234.9 | 23.04 | 26152 | 4 | 4 | 5858486 | 5441184 | 5181906 | 5494765 | 5473786.5 | 5540257 | 5545080.5 | 5733629 | 5860209 | 5148649 | 5358455 | 5310409 | 5871801 | 5511554 | 5128135 |
| Q9WUP7 | Uchl5 | Ubiquitin carboxyl-terminal hydrolase isozyme L5 Mus musculus | 816.93 | 66.26 | 37617 | 16 | 16 | 18753543 | 16516076 | 16795345 | 18152443 | 18655609 | 20488689 | 17931367 | 21754229 | 19801285 | 16999261 | 21258003 | 20479617 | 16306368 | 23009719 | 20647001 |
| Q3U1V6 | Uevld | Ubiquitin-conjugating enzyme E2 variant 3 Mus musculus | 110.15 | 14.01 | 51681 | 3 | 3 | 1330330 | 1469679 | 2752659.25 | 1350396.62 | 2624959.5 | 1463671.25 | 1323100.62 | 1491278.88 | 1829578 | 2097212 | 3483823 | 1422308.25 | 2201084.75 | 1529029.12 | 1428982.75 |
| Q9CR09 | Ufc1 | Ubiquitin-fold modifier-conjugating enzyme 1 Mus musculus | 173.85 | 31.14 | 19481 | 4 | 4 | 7331790 | 6625408.5 | 6249144.5 | 5932308 | 6208152.5 | 5879627 | 6233661 | 5425873 | 5544552.5 | 5861724 | 5403181 | 5793377.5 | 6144519.5 | 5187507 | 6461896.5 |
| P70362 | Ufd1 | Ubiquitin recognition factor in ER-associated degradation protein 1 Mus musculus | 206.76 | 19.54 | 34481 | 5 | 5 | 5358554 | 5680012 | 5781273.5 | 5793175 | 5958451 | 5806836 | 5966910 | 6251997 | 5413330 | 6577658 | 6416657 | 6496376.5 | 5738774 | 5433259 | 4997507.5 |
| Q8CCJ3 | Ufl1 | E3 UFM1-protein ligase 1 Mus musculus | 407.08 | 13.49 | 89520 | 6 | 6 | 3361500 | 3305349 | 3690832.75 | 3432519.25 | 1910361.5 | 3603082 | 3889765.5 | 3879137.75 | 3862947.75 | 3473269.5 | 2888599.75 | 3277545.25 | 3403040.25 | 3961524.75 | 2530876 |
| Q99K23 | Ufsp2 | Ufm1-specific protease 2 Mus musculus | 263.03 | 17.14 | 52515 | 6 | 6 | 3958302.5 | 4498152 | 4389915.5 | 5304419 | 4965032 | 5058198.5 | 5166905 | 5219383 | 4855213 | 4786572 | 5055517 | 4657555 | 4756276.5 | 4497647.5 | 4414571.5 |
| O70199 | Ugdh | UDP-glucose 6-dehydrogenase Rattus norvegicus | 46.3 | 1.83 | 54892 | 1 | 1 | 1 | 1 | 1 | 1 | 1 | 1 | 212528.34 | 1 | 1 | 1 | 216782.94 | 1 | 283815.22 | 1 | 316071.62 |
| Q6P5E4 | Uggt1 | UDP-glucose:glycoprotein glucosyltransferase 1 Mus musculus | 2310.6 | 40.04 | 176434 | 41 | 42 | 55591629 | 46955769 | 47187621 | 46302633 | 51739361 | 52291305 | 57080193 | 50508841 | 53292677 | 58953205 | 60166449 | 54993145 | 54448341 | 46498781 | 69873881 |
| Q91ZJ5 | Ugp2 | UTP--glucose-1-phosphate uridylyltransferase Mus musculus | 278.07 | 13.78 | 56979 | 6 | 6 | 1285799.88 | 5131077 | 4584418 | 6727989.5 | 7102984 | 4728760 | 6795957 | 5163417 | 3743640.25 | 6667090 | 6955679 | 5425594 | 6830269.5 | 4547190 | 6236832 |
| Q6ZQM8 | Ugt1a7c | UDP-glucuronosyltransferase 1-7C Mus musculus | 1044.94 | 45.01 | 59758 | 18 | 19 | 78058561 | 83750217 | 64516753 | 77695905 | 77941233 | 75049305 | 76013465 | 67671353 | 70664089 | 72452873 | 72873353 | 72334025 | 74089721 | 69563761 | 73259265 |
| Q8VDF2 | Uhrf1 | E3 ubiquitin-protein ligase UHRF1 Mus musculus | 849.41 | 25.45 | 88304 | 11 | 16 | 6293807.5 | 2417324 | 6369521.5 | 7107202 | 6277823 | 5259401 | 6041884.5 | 5630988 | 5863606 | 7775974 | 8583450 | 6415577 | 1995108.38 | 6907473 | 8393315 |
| P13439 | Umps | Uridine 5'-monophosphate synthase Mus musculus | 809.7 | 44.07 | 52292 | 15 | 15 | 22133903 | 23610109 | 22258945 | 21785751 | 18322199 | 20474699 | 19281133 | 19168951 | 22184687 | 17744081 | 24402877 | 21963853 | 20821373 | 19409333 | 20438413 |
| Q62885 | Unc119 | Protein unc-119 homolog A Rattus norvegicus | 61.98 | 5.42 | 27048 | 1 | 1 | 1 | 1 | 1 | 1 | 1 | 512629.56 | 562204.38 | 684030.78 | 601355.06 | 1059934.12 | 1 | 554497.62 | 552798.31 | 1 | 920076.31 |
| B2RUP2 | Unc13d | Protein unc-13 homolog D Mus musculus | 36.8 | 1.01 | 123119 | 1 | 1 | 247152.02 | 184416.53 | 91451.2 | 211817.7 | 86256.26 | 85754.12 | 1 | 61813.97 | 64886.64 | 177058.64 | 99040.52 | 99626.11 | 89339.3 | 50909.33 | 61351.48 |
| Q99KD5 | Unc45a | Protein unc-45 homolog A Mus musculus | 119.85 | 4.24 | 103447 | 2 | 2 | 1 | 1 | 271007.47 | 93428.48 | 1 | 1 | 63461.32 | 283832.47 | 228165.52 | 1 | 253575.97 | 355847.19 | 276605.485 | 295359.03 | 201639.75 |
| Q8VCW4 | Unc93b1 | Protein unc-93 homolog B1 Mus musculus | 358.12 | 13.21 | 66981 | 6 | 6 | 10856783 | 4375958.5 | 4493601 | 3998579 | 3573923 | 4026241.5 | 2974632.5 | 4067163.5 | 4143039.5 | 2546903 | 4990788.5 | 3384434.5 | 3822362 | 4357389 | 3677521.5 |
| Q9EPU0 | Upf1 | Regulator of nonsense transcripts 1 Mus musculus | 769.85 | 16.64 | 123967 | 15 | 15 | 4513688.5 | 4762802 | 4437975 | 5356068 | 5269039.5 | 4200129.5 | 7651897.5 | 4579007 | 3328904.75 | 5869676.5 | 7074408 | 7934471 | 5645362 | 4980795.5 | 7717048 |
| Q9CWU6 | Uqcc1 | Ubiquinol-cytochrome-c reductase complex assembly factor 1 Mus musculus | 139.84 | 10.51 | 34300 | 3 | 3 | 1099861.5 | 3819063.5 | 3298992.5 | 2966367.5 | 2181181.5 | 2830307.25 | 2495087.5 | 3273012.25 | 4283074 | 3163121 | 2657445.75 | 2348317.5 | 2459916.5 | 2756716 | 2650413.75 |
| Q8R1I1 | Uqcr10 | Cytochrome b-c1 complex subunit 9 Mus musculus | 74.73 | 37.5 | 7446 | 2 | 2 | 5445867 | 1762729.62 | 2465869.75 | 2581725.5 | 3023601 | 3298172.25 | 2996779.5 | 3234226 | 2592440.75 | 2500676 | 2275567 | 2365811.5 | 3080107.5 | 2677549.5 | 2086487.75 |
| Q9CZ13 | Uqcrc1 | Cytochrome b-c1 complex subunit 1, mitochondrial Mus musculus | 833 | 44.38 | 52852 | 13 | 13 | 18654481 | 21889973 | 21706489 | 20146351 | 20867265 | 20505363 | 25215809 | 19341009 | 20422713 | 20250713 | 21641561 | 20517481 | 20014281 | 20939845 | 22990273 |
| Q9DB77 | Uqcrc2 | Cytochrome b-c1 complex subunit 2, mitochondrial Mus musculus | 1118.98 | 68.65 | 48235 | 18 | 18 | 20296187 | 25128709 | 18761921 | 18604255 | 21652477 | 19747657 | 26874311 | 21775237 | 13422409 | 17233697 | 19085605 | 20628507 | 18869887 | 22422429 | 20070071 |
| Q9CR68 | Uqcrfs1 | Cytochrome b-c1 complex subunit Rieske, mitochondrial Mus musculus | 358.37 | 41.97 | 29368 | 7 | 7 | 8335855 | 9108856 | 12914313 | 9917850 | 10165819 | 10456101 | 15203493 | 9487886 | 7714672.5 | 11624945 | 10958289 | 12507911 | 10959190 | 9713235 | 12797079 |
| P99028 | Uqcrh | Cytochrome b-c1 complex subunit 6, mitochondrial Mus musculus | 131.91 | 28.09 | 10435 | 2 | 2 | 4904808 | 4955743.5 | 4000095 | 5226052 | 5368117 | 5346700 | 1799336 | 4713200 | 5052550 | 4055730.5 | 4753040.5 | 4188655.25 | 1881561.25 | 4728470 | 3699978 |
| Q7TQ16 | Uqcrq | Cytochrome b-c1 complex subunit 8 Rattus norvegicus | 94.57 | 24.39 | 9849 | 2 | 2 | 1 | 716689.88 | 1 | 224516.77 | 1 | 875421.19 | 899883.06 | 1153519.12 | 811709.42 | 318014.31 | 288866.69 | 1069416.75 | 164866.89 | 937533 | 356120.97 |
| P70697 | Urod | Uroporphyrinogen decarboxylase Mus musculus | 321.18 | 23.43 | 40692 | 5 | 5 | 5958787.5 | 5213655.5 | 4495623.5 | 5482164 | 5296505 | 5009652 | 6180548.5 | 5156727 | 4498654 | 6065050.5 | 4540251.5 | 4647923 | 4979196 | 5083291.5 | 5032150.5 |
| Q9Z1Z0 | Uso1 | General vesicular transport factor p115 Mus musculus | 717.45 | 20.44 | 106983 | 15 | 15 | 20782991 | 23835381 | 23481537 | 25961857 | 23067939 | 24239063 | 21799839 | 24950205 | 20539957 | 24216521 | 22623899 | 23707785 | 25329655 | 23614703 | 24006201 |
| P52479 | Usp10 | Ubiquitin carboxyl-terminal hydrolase 10 Mus musculus | 283.26 | 12.75 | 87022 | 7 | 7 | 4203533 | 2332308.75 | 3097966.25 | 4501043 | 2947893.5 | 3149260.25 | 2948637.5 | 3851496 | 3100160 | 4585553 | 4597848 | 3168345.25 | 3183294 | 4556701 | 3162682 |
| Q9JMA1 | Usp14 | Ubiquitin carboxyl-terminal hydrolase 14 Mus musculus | 1417.72 | 68.56 | 56002 | 25 | 25 | 40458277 | 38805877 | 41205489 | 44496385 | 43180869 | 43888957 | 37618793 | 43617537 | 47388085 | 43024573 | 41096165 | 40471605 | 43397197 | 42767325 | 40102201 |
| Q8R5H1 | Usp15 | Ubiquitin carboxyl-terminal hydrolase 15 Mus musculus | 480.37 | 15.7 | 112325 | 10 | 11 | 6104938.5 | 5670110 | 4913352.5 | 5859005.5 | 6201124 | 4678916 | 6040420.5 | 5608456 | 4542854.5 | 5453421.5 | 5921113 | 4732982.5 | 6049366.5 | 6007191 | 5369255 |
| Q9WTV6 | Usp18 | Ubl carboxyl-terminal hydrolase 18 Mus musculus | 76.99 | 6.25 | 42472 | 2 | 2 | 1 | 1 | 178631.36 | 182691.03 | 1 | 1 | 1 | 1 | 1 | 118559.18 | 216500.23 | 1 | 1 | 291493.12 | 1 |
| Q3UJD6 | Usp19 | Ubiquitin carboxyl-terminal hydrolase 19 Mus musculus | 226.22 | 4.34 | 150549 | 5 | 5 | 1141519.88 | 1723307.75 | 1526260.88 | 1577298.75 | 1645881.88 | 1335582.5 | 1530906.12 | 1440975 | 1646378.5 | 1762631.5 | 1472423.75 | 1495290.5 | 1984273.38 | 1461267 | 1695388.75 |
| B1AY13 | Usp24 | Ubiquitin carboxyl-terminal hydrolase 24 Mus musculus | 77.01 | 1.15 | 294001 | 2 | 2 | 663189.25 | 624075 | 769901.38 | 528914.75 | 505905.31 | 824687.88 | 524386.25 | 553394.69 | 741097.62 | 520136.41 | 969886.75 | 675529.62 | 577170.38 | 906336 | 646254.31 |
| P57080 | Usp25 | Ubiquitin carboxyl-terminal hydrolase 25 Mus musculus | 61.6 | 0.95 | 121420 | 1 | 1 | 1 | 242203.67 | 1 | 1 | 1 | 155820.23 | 180378.3 | 147644.61 | 147863.3 | 192990 | 158352.97 | 164367.58 | 191081.05 | 131156.78 | 176879.52 |
| Q3TIX9 | Usp39 | U4/U6.U5 tri-snRNP-associated protein 2 Mus musculus | 553.77 | 26.77 | 65146 | 11 | 11 | 8693761 | 10850620 | 9177515 | 7901195.5 | 11353966 | 11836285 | 9775705 | 12226249 | 13010987 | 7529925 | 12657305 | 9287495 | 8548933 | 11018430 | 9807441 |
| B2GUZ1 | Usp4 | Ubiquitin carboxyl-terminal hydrolase 4 Rattus norvegicus | 421.77 | 11.24 | 108373 | 7 | 8 | 8456999 | 7108444 | 6530091.5 | 5582978 | 6051841.5 | 5157251 | 6912875 | 7358370 | 8223301 | 6198349 | 8293381 | 7210355.5 | 7173106.5 | 7480441.5 | 6816135 |
| Q8BY87 | Usp47 | Ubiquitin carboxyl-terminal hydrolase 47 Mus musculus | 140.46 | 2.69 | 157455 | 3 | 3 | 602432.31 | 2089618.62 | 1939205.5 | 1105125.12 | 1603429 | 506494.31 | 1397834.12 | 2609616.5 | 2855845.5 | 462006.84 | 1607423.5 | 2230575 | 1490220.88 | 2051183 | 1831055.12 |
| P56399 | Usp5 | Ubiquitin carboxyl-terminal hydrolase 5 Mus musculus | 1665.19 | 45.8 | 95833 | 26 | 26 | 84498225 | 93753113 | 91397561 | 92476553 | 89759745 | 92308785 | 90908993 | 95772745 | 89853713 | 90120737 | 99733969 | 89964849 | 95548097 | 87319025 | 91420465 |
| Q4VSI4 | Usp7 | Ubiquitin carboxyl-terminal hydrolase 7 Rattus norvegicus | 219.57 | 4.81 | 128431 | 5 | 5 | 3975493.25 | 2977166 | 2808469.75 | 2560292.25 | 2880270 | 1792849.25 | 1709990.38 | 2924766 | 3849858.5 | 2092537 | 3280725.5 | 3376103.25 | 2517094 | 2752874.5 | 3284871.5 |
| P70398 | Usp9x | Probable ubiquitin carboxyl-terminal hydrolase FAF-X Mus musculus | 412.87 | 4.34 | 290711 | 8 | 8 | 2325340.5 | 3451192.75 | 3006313.5 | 3166280.75 | 2296593.5 | 2204140 | 4122456.25 | 2840018.5 | 3931250.25 | 2871579.5 | 2974099.5 | 3140165.25 | 2026919.12 | 3545290.75 | 3041875.75 |
| Q640M1 | Utp14a | U3 small nucleolar RNA-associated protein 14 homolog A Mus musculus | 67.13 | 4.3 | 87265 | 2 | 2 | 1 | 463575.97 | 1 | 1 | 594368.88 | 340364.66 | 652142.88 | 1 | 1 | 428953 | 1 | 1 | 355117.62 | 1 | 420919.56 |
| Q8C7V3 | Utp15 | U3 small nucleolar RNA-associated protein 15 homolog Mus musculus | 345.09 | 18.94 | 59375 | 7 | 7 | 2518481.5 | 2785008 | 2513311.5 | 2797119 | 2384890.75 | 2688292.5 | 2908984.25 | 2372962 | 2517565.25 | 2576680 | 2137697.75 | 2294734 | 2224503.5 | 3338185.5 | 2584093.25 |
| Q5SSI6 | Utp18 | U3 small nucleolar RNA-associated protein 18 homolog Mus musculus | 502.03 | 30.98 | 61218 | 9 | 10 | 7200306.5 | 6874788.5 | 6430860 | 7406476 | 8152441 | 6059425.5 | 5492762 | 6682784.5 | 6503827 | 6473492 | 6229336.5 | 6372983.5 | 7633244.5 | 5687722 | 5592385 |
| Q9JI13 | Utp3 | Something about silencing protein 10 Mus musculus | 271.13 | 13.43 | 53399 | 5 | 5 | 3654444.75 | 4159856 | 2575663 | 4289463 | 4867771 | 2891608.5 | 3117193.25 | 2570143.5 | 2026012.25 | 2324770 | 3239254 | 3173196.75 | 2751775.5 | 2337231.75 | 1556417.25 |
| Q8R2N2 | Utp4 | U3 small nucleolar RNA-associated protein 4 homolog Mus musculus | 271.32 | 11.95 | 76909 | 5 | 5 | 1 | 1 | 806678.25 | 891446 | 1 | 1 | 688351.75 | 1 | 1049417.88 | 1 | 952078.75 | 1 | 1 | 1032580.81 | 1 |
| Q8K245 | Uvrag | UV radiation resistance associated protein Mus musculus | 45.36 | 1.43 | 77525 | 1 | 1 | 1 | 1 | 1 | 1 | 1 | 1 | 367093.56 | 1 | 1 | 298949.31 | 277858.56 | 296255.69 | 331283.84 | 1 | 196855.39 |
| Q9WTZ0 | Uxt | Protein UXT Mus musculus | 45.66 | 7.01 | 18187 | 1 | 1 | 840198.69 | 989501.88 | 870619.56 | 887313.31 | 897253.56 | 923897.81 | 971356.62 | 1139656.88 | 1090172.75 | 1031271.015 | 919638.75 | 1059172 | 1111709.5 | 1228954.62 | 790788.12 |
| Q80WQ2 | Vac14 | Protein VAC14 homolog Mus musculus | 768.49 | 25.96 | 88048 | 14 | 14 | 13528564 | 10920689 | 10440508 | 9211076 | 10567934 | 11857268 | 11897888 | 13589852 | 14730631 | 12519382 | 15634950 | 13910067 | 10104660 | 12500781 | 14871010 |
| P63024 | Vamp3 | Vesicle-associated membrane protein 3 Mus musculus | 90.95 | 32.04 | 11480 | 2 | 2 | 1 | 1 | 1 | 1 | 411596.5 | 1 | 1 | 1 | 1 | 622543.19 | 1 | 566550.56 | 527985 | 1 | 647409.19 |
| O70404 | Vamp8 | Vesicle-associated membrane protein 8 Mus musculus | 63.54 | 11.88 | 11451 | 1 | 1 | 391275.59 | 425782.22 | 429650.78 | 268917.22 | 469063.53 | 300058.62 | 264817.34 | 434681.91 | 547931.19 | 355776.66 | 415594.72 | 391707.31 | 603102.31 | 450439.81 | 326859.75 |
| Q9WV55 | Vapa | Vesicle-associated membrane protein-associated protein A Mus musculus | 329.96 | 31.73 | 27855 | 6 | 7 | 10996689 | 11994211 | 14204713 | 13406575 | 13966539 | 15014039 | 17181543 | 11470686 | 12966744 | 17597597 | 14718941 | 14365209 | 13180647 | 10461143 | 19171521 |
| Q9QY76 | Vapb | Vesicle-associated membrane protein-associated protein B Mus musculus | 395.28 | 28.81 | 26946 | 7 | 8 | 7922319 | 4367373 | 9264227 | 8058532.5 | 7800590.5 | 7399091 | 11047254 | 7565865.5 | 8031529 | 9666180 | 8512676 | 9420571 | 8230127.5 | 4854481 | 11306076 |
| Q9Z1Q9 | Vars | Valine--tRNA ligase Mus musculus | 2525.76 | 51.46 | 140215 | 44 | 44 | 99511017 | 107113025 | 104057321 | 105556265 | 107404969 | 120400185 | 128984305 | 107190897 | 120652129 | 123631953 | 115033665 | 113963129 | 107953753 | 112612465 | 119659857 |
| Q9CZT5 | Vasn | Vasorin Mus musculus | 54.53 | 1.04 | 72261 | 1 | 1 | 3280807.5 | 3123268.5 | 3746866.5 | 3503898 | 3298635.5 | 3215103.25 | 3833396 | 3311548.5 | 3197055.75 | 3891698 | 3647834.75 | 3525021.25 | 3652496.5 | 3544358 | 3856691 |
| P70460 | Vasp | Vasodilator-stimulated phosphoprotein Mus musculus | 398.93 | 26.4 | 39667 | 8 | 8 | 7519228 | 6332024 | 9423862 | 10308521 | 9601369 | 8164956.5 | 16323068 | 7468682 | 5317822.5 | 13236192 | 11588755 | 10904721 | 8820969 | 5787937.5 | 13239809 |
| Q62465 | Vat1 | Synaptic vesicle membrane protein VAT-1 homolog Mus musculus | 635.9 | 35.47 | 43097 | 10 | 10 | 13013952 | 12600807 | 11770273 | 9058213 | 10924196 | 13991794 | 20890149 | 14604519 | 12760817 | 19295287 | 10335105 | 15816727 | 15712436 | 12519341 | 18737527 |
| P27870 | Vav1 | Proto-oncogene vav Mus musculus | 1530.61 | 33.96 | 98137 | 22 | 23 | 51367053 | 43368361 | 55340193 | 44861849 | 49595133 | 51060677 | 47615881 | 48192189 | 63439425 | 49550341 | 53254949 | 54831245 | 45299229 | 50411405 | 53312141 |
| Q9R0C8 | Vav3 | Guanine nucleotide exchange factor VAV3 Mus musculus | 464.12 | 12.28 | 97968 | 8 | 9 | 6905495.5 | 5524216 | 6153152.5 | 5147491 | 5953059.5 | 5867254.5 | 4289431.5 | 5379711.5 | 6103645 | 3824042.25 | 6529208 | 6377695 | 4933067 | 6681928 | 7422887 |
| P61759 | Vbp1 | Prefoldin subunit 3 Mus musculus | 364.86 | 38.27 | 22436 | 7 | 7 | 5948499.5 | 6263599 | 6214578.5 | 4969961 | 4569902 | 5409954.5 | 5684890 | 6930047.5 | 7104349 | 5394386.5 | 6995671 | 6913492 | 5233096 | 6720143.5 | 6627034.5 |
| Q64727 | Vcl | Vinculin Mus musculus | 958.92 | 20.17 | 116717 | 17 | 17 | 16008462 | 27164945 | 24175101 | 29153369 | 28499287 | 20311925 | 15019441 | 24319957 | 20771823 | 20612593 | 19855099 | 24432689 | 27519673 | 11911103 | 26105871 |
| P46462 | Vcp | Transitional endoplasmic reticulum ATPase Rattus norvegicus | 2628.17 | 61.41 | 89349 | 39 | 41 | 111624025 | 112881953 | 111558361 | 120316785 | 111253761 | 114293097 | 132438505 | 111936049 | 100151633 | 128674033 | 123225737 | 109222977 | 110847105 | 105294777 | 128338385 |
| Q8CDG3 | Vcpip1 | Deubiquitinating protein VCIP135 Mus musculus | 117.73 | 2.79 | 134503 | 2 | 2 | 1332798 | 1507742.5 | 1314535.25 | 1718037.88 | 1391673.38 | 1211282 | 1312199 | 1 | 1549483.12 | 1 | 1002000 | 1276699.38 | 1439071.5 | 625351.06 | 1085780.485 |
| Q9Z2L0 | Vdac1 | Voltage-dependent anion-selective channel protein 1 Rattus norvegicus | 1323.92 | 78.8 | 30756 | 17 | 17 | 90162177 | 100447313 | 85169633 | 88416833 | 102944129 | 102152713 | 106514865 | 99965057 | 109305313 | 108290153 | 103115545 | 98041697 | 97521505 | 97159993 | 102305809 |
| Q60930 | Vdac2 | Voltage-dependent anion-selective channel protein 2 Mus musculus | 1007.83 | 56.61 | 31733 | 13 | 14 | 155728449 | 168903905 | 177213537 | 146716865 | 154621441 | 168737985 | 182034017 | 169498321 | 195064977 | 167148433 | 162906609 | 165127041 | 151896993 | 169683681 | 173625105 |
| Q60931 | Vdac3 | Voltage-dependent anion-selective channel protein 3 Mus musculus | 600 | 38.87 | 30753 | 7 | 9 | 42390313 | 40488137 | 52823513 | 36550153 | 37206093 | 40660641 | 52767713 | 48665681 | 54000065 | 42258185 | 49403905 | 52054569 | 36906401 | 45240049 | 51967941 |
| P20152 | Vim | Vimentin Mus musculus | 3269.56 | 86.48 | 53688 | 43 | 51 | 2434394369 | 2796715521 | 2589878017 | 2831042049 | 2816222721 | 2746197761 | 2924476929 | 2644526337 | 2887560705 | 2946960385 | 3032498689 | 2863035393 | 3148944641 | 2916097025 | 3008812801 |
| Q91ZQ0 | Vmp1 | Vacuole membrane protein 1 Rattus norvegicus | 102.3 | 3.94 | 45901 | 2 | 2 | 1524172.88 | 1283855.75 | 1099408.88 | 1219152.3 | 969171.69 | 755324 | 2530842 | 386091.94 | 762935.88 | 1592055.25 | 1 | 1352837.62 | 1 | 465910.72 | 1724362.5 |
| Q91W86 | Vps11 | Vacuolar protein sorting-associated protein 11 homolog Mus musculus | 95.53 | 3.4 | 107719 | 2 | 2 | 682377.81 | 683623.4125 | 506056.03 | 893279.75 | 652780.06 | 474398.09 | 1 | 1266118.12 | 520507.59 | 1 | 912573.6075 | 1025154.12 | 643838.12 | 1223052.25 | 758249.94 |
| Q8BX70 | Vps13c | Vacuolar protein sorting-associated protein 13C Mus musculus | 3355.131 | 24.09 | 420089 | 63 | 63 | 95545865 | 88624833 | 98818273 | 82741225 | 90174377 | 92405145 | 86627073 | 92460161 | 104781521 | 82778697 | 93228057 | 93176241 | 80144625 | 98279753 | 92267385 |
| Q920Q4 | Vps16 | Vacuolar protein sorting-associated protein 16 homolog Mus musculus | 296.13 | 12.28 | 94928 | 6 | 6 | 3724955.25 | 4112053 | 4163690 | 3637693.5 | 3905642.25 | 3387699.75 | 1330288.5 | 3900121 | 4317352 | 3687960.5 | 3259410.5 | 3466263 | 3656434.5 | 4106688.75 | 3754109.75 |
| Q8R307 | Vps18 | Vacuolar protein sorting-associated protein 18 homolog Mus musculus | 117.84 | 2.88 | 110219 | 2 | 2 | 887620.56 | 373677.84 | 234487.78 | 376040.69 | 380077.09 | 451314.94 | 223941.53 | 356910.03 | 335028.56 | 235621.11 | 405721.47 | 281906.56 | 333010 | 594984.62 | 403905.6625 |
| P0C0A1 | Vps25 | Vacuolar protein-sorting-associated protein 25 Rattus norvegicus | 167.23 | 21.02 | 20762 | 3 | 3 | 4261923 | 3802196 | 3971357 | 2041096.75 | 3977827.5 | 4184368 | 2805504.75 | 3853647.5 | 3549586 | 3558352.25 | 3297007.25 | 2758480.5 | 3621714.5 | 3305001.75 | 3300750.5 |
| P40336 | Vps26a | Vacuolar protein sorting-associated protein 26A Mus musculus | 519.69 | 39.14 | 38114 | 9 | 10 | 11493229 | 8537860 | 10521782 | 8121111.5 | 10240659 | 8039809.5 | 12005433 | 10985765 | 7580499.5 | 7766420 | 10112917 | 12364090 | 14523108 | 9571574 | 9842798 |
| Q8C0E2 | Vps26b | Vacuolar protein sorting-associated protein 26B Mus musculus | 117.51 | 9.23 | 39125 | 2 | 3 | 1498711.75 | 8649769 | 8341549 | 5296583.5 | 5104544.5 | 6648860 | 4166084.5 | 6691383 | 5773318.5 | 6264169 | 4614847 | 5356272.5 | 6618503 | 6143924.5 | 4285269.5 |
| B5DEN9 | Vps28 | Vacuolar protein sorting-associated protein 28 homolog Rattus norvegicus | 182.21 | 17.54 | 26110 | 3 | 3 | 1357463.88 | 1802175 | 1675546.88 | 2291588 | 2502938 | 2733562 | 327481.75 | 2199390.75 | 2053661.75 | 1974719.5 | 2228359 | 2219100.5 | 1479656 | 2220711.75 | 734241.31 |
| Q9QZ88 | Vps29 | Vacuolar protein sorting-associated protein 29 Mus musculus | 150.31 | 29.12 | 20496 | 4 | 4 | 5598654 | 5453934 | 4981585.5 | 5349165 | 5998336 | 6395298 | 7404802 | 4871431 | 4794811.5 | 6094856 | 6671470 | 6621229 | 6835878.5 | 5448261 | 7631898.5 |
| Q63615 | Vps33a | Vacuolar protein sorting-associated protein 33A Rattus norvegicus | 219.52 | 11.56 | 67514 | 3 | 3 | 1307783.5 | 1303169.62 | 1088316.25 | 1534171 | 1176797.75 | 1148764 | 1030872.19 | 1246758.75 | 910919.31 | 1151826 | 1172046.25 | 632820.31 | 1312654.12 | 1103989.88 | 1207696.5 |
| Q9EQH3 | Vps35 | Vacuolar protein sorting-associated protein 35 Mus musculus | 1651.39 | 42.96 | 91713 | 27 | 27 | 77016625 | 67792897 | 79955761 | 60206765 | 65895489 | 72262449 | 56078133 | 80757641 | 91777169 | 52366025 | 76304297 | 78313417 | 54767777 | 78032481 | 76392625 |
| P0C0A2 | Vps36 | Vacuolar protein-sorting-associated protein 36 Rattus norvegicus | 251.35 | 15.8 | 43718 | 5 | 5 | 7320975.5 | 9481144 | 23444781 | 37198309 | 29594149 | 23050603 | 6284301 | 26574615 | 19985531 | 27304421 | 9732122 | 6938655 | 20947313 | 8140566 | 5684187.5 |
| Q8R105 | Vps37c | Vacuolar protein sorting-associated protein 37C Mus musculus | 92.67 | 9.09 | 38453 | 2 | 2 | 2901238.5 | 3828342.25 | 2644181.5 | 1177149.75 | 3265096 | 2442878 | 2130216.75 | 2546811 | 2640526 | 1631777 | 1957700.75 | 2040102.12 | 2782307 | 1019875.56 | 1966572 |
| Q8R5L3 | Vps39 | Vam6/Vps39-like protein Mus musculus | 146.49 | 4.29 | 101693 | 3 | 3 | 1434395.62 | 1657410.75 | 991179.44 | 1260462.12 | 1247210.38 | 2016975.25 | 902467.44 | 1810171.75 | 2095288.88 | 1799693.75 | 1388012.5 | 1650536.25 | 1924592 | 1561128 | 1670936.38 |
| Q5KU39 | Vps41 | Vacuolar protein sorting-associated protein 41 homolog Mus musculus | 114.77 | 5.28 | 98602 | 3 | 3 | 1764089.75 | 2877492 | 1975183 | 2186869 | 2298270.5 | 2139223.5 | 2036275.75 | 2113150.75 | 2754968.75 | 1851145.5 | 2011441 | 1952366.88 | 2669039.5 | 1968880 | 1846946.75 |
| P97390 | Vps45 | Vacuolar protein sorting-associated protein 45 Mus musculus | 201.05 | 11.05 | 65053 | 4 | 4 | 2015103 | 2130352.5 | 2321523 | 1991782.25 | 2260159 | 2491038 | 1984698.75 | 2417920.5 | 2470344 | 2213532 | 2278184 | 2349827 | 1995984 | 2372315.5 | 2139077 |
| Q793F9 | Vps4a | Vacuolar protein sorting-associated protein 4A Rattus norvegicus | 152.46 | 10.07 | 48907 | 1 | 4 | 3217811.5 | 2766266.25 | 2672727.5 | 2452150.75 | 3413551.5 | 3155098 | 4702636 | 2728847.5 | 2761848 | 4106585 | 2817565 | 3837685 | 3169802.75 | 3177269.75 | 4409573.5 |
| P46467 | Vps4b | Vacuolar protein sorting-associated protein 4B Mus musculus | 495.86 | 32.66 | 49419 | 6 | 9 | 7534549 | 8680039 | 6019704.5 | 6241002 | 7659982 | 9420079 | 10713117 | 6007973 | 7439550 | 9100958 | 10523410 | 8482718 | 8922350 | 6878884.5 | 7066618.5 |
| F1LSG8 | Vps50 | Syndetin Rattus norvegicus | 143.18 | 4.88 | 111188 | 3 | 3 | 1385889.5 | 1878717.5 | 1521972.62 | 1271391.25 | 1867205.25 | 1697695.25 | 1442847.75 | 1568617.88 | 1685859.5 | 1481516.75 | 1558856.75 | 1695417.5 | 1233650.38 | 1956223 | 1501356.75 |
| Q3UVL4 | Vps51 | Vacuolar protein sorting-associated protein 51 homolog Mus musculus | 329.73 | 12.02 | 86187 | 6 | 6 | 6871667 | 7778310.5 | 6939899 | 6797230 | 6580675 | 6668949 | 5379714 | 7284897 | 7661468 | 6227242 | 4818772 | 5928755 | 6399530 | 7035787 | 6024827 |
| O55166 | Vps52 | Vacuolar protein sorting-associated protein 52 homolog Rattus norvegicus | 370.9 | 15.35 | 82103 | 8 | 8 | 68504841 | 72353409 | 70649513 | 86977121 | 70330769 | 67046397 | 78290545 | 67961257 | 66560329 | 81271561 | 82029041 | 74027113 | 73776937 | 67348945 | 75856321 |
| Q8CCB4 | Vps53 | Vacuolar protein sorting-associated protein 53 homolog Mus musculus | 100.89 | 3.85 | 94423 | 2 | 2 | 999012 | 794355.5 | 1 | 1 | 1 | 877333.7475 | 824968.56 | 772101.31 | 985567.06 | 926698.06 | 643030.38 | 1 | 839505.44 | 1 | 830679.5 |
| Q80X41 | Vrk1 | Serine/threonine-protein kinase VRK1 Mus musculus | 289.35 | 19.32 | 49741 | 7 | 7 | 9471291 | 8664119 | 7830619 | 6107584 | 6991410.5 | 8022031 | 8245107 | 8559220 | 9488988 | 6520763 | 8086112 | 9223964 | 6660399 | 8661350 | 8153413 |
| Q8BN21 | Vrk2 | Serine/threonine-protein kinase VRK2 Mus musculus | 280.93 | 14.91 | 58119 | 5 | 5 | 5893602 | 4538886 | 3376686.5 | 4370146.5 | 4444076 | 4594383.5 | 4152334.25 | 4470281 | 2683222.75 | 3430349 | 2251766 | 2712444 | 3883148.25 | 4966665 | 2698838.75 |
| P29788 | Vtn | Vitronectin Mus musculus | 80.38 | 3.77 | 54849 | 1 | 1 | 690520.81 | 656204.25 | 617229.56 | 475170.97 | 465189.84 | 793610.56 | 1092978.12 | 653107 | 649661.81 | 753121.88 | 370059 | 305932.31 | 336121.8825 | 430688.81 | 237807.41 |
| Q99KC8 | Vwa5a | von Willebrand factor A domain-containing protein 5A Mus musculus | 1849.75 | 49.81 | 87143 | 29 | 29 | 141498321 | 159484497 | 143030481 | 122763457 | 156414961 | 124186081 | 108159329 | 123161337 | 153572161 | 138460561 | 146397905 | 135037745 | 147435889 | 154859409 | 113132265 |
| Q8CC88 | Vwa8 | von Willebrand factor A domain-containing protein 8 Mus musculus | 137.71 | 1.84 | 213421 | 3 | 3 | 1 | 1 | 226946.95 | 169567.12 | 137882.53 | 114980.92 | 156685.55 | 197874.97 | 155416.33 | 136439.17 | 169044.91 | 112105.79 | 198083.94 | 214992.7 | 157381.72 |
| Q65Z40 | Wapl | Wings apart-like protein homolog Mus musculus | 286.43 | 7.58 | 134070 | 5 | 5 | 1373054.38 | 1563977.25 | 1570771 | 1787935.88 | 1526152.38 | 1857571.88 | 1260868.75 | 1617831.5 | 1016509.12 | 2019379.75 | 2016278.25 | 1611247.88 | 1637796 | 1236420.5 | 1925407.75 |
| P32921 | Wars | Tryptophan--tRNA ligase, cytoplasmic Mus musculus | 620.24 | 36.8 | 54358 | 12 | 12 | 20430443 | 20311007 | 19161015 | 23268679 | 18740545 | 19718467 | 18531521 | 16484641 | 19825177 | 17253001 | 17789717 | 18628847 | 19343121 | 19991629 | 13638537 |
| P70315 | Was | Wiskott-Aldrich syndrome protein homolog Mus musculus | 336.07 | 13.65 | 54192 | 7 | 7 | 5165638.5 | 5253825 | 6108551.5 | 5458462.5 | 6272689 | 4487892 | 5052734.5 | 4846404 | 4716140.5 | 4673451 | 6086846 | 5760981 | 6565875 | 4640480 | 5520264 |
| Q8BH43 | Wasf2 | Wiskott-Aldrich syndrome protein family member 2 Mus musculus | 520.27 | 21.33 | 54074 | 9 | 9 | 7733303.5 | 8904244 | 9569390 | 7331849 | 11301141 | 10284164 | 11763803 | 8315744.5 | 11253333 | 12061459 | 13948751 | 10160071 | 12063375 | 10305633 | 11377461 |
| B2RYF7 | Washc1 | WASH complex subunit 1 Rattus norvegicus 2 | 232.58 | 15.37 | 51327 | 3 | 3 | 346658.91 | 438681.59 | 1 | 1 | 427266.47 | 504237.84 | 531138.62 | 380904.28 | 159510.78 | 147497.58 | 242161.72 | 139219.52 | 401006.66 | 390871 | 544262.38 |
| Q6PGL7 | Washc2 | WASH complex subunit 2 Mus musculus | 490.49 | 11.92 | 145311 | 7 | 7 | 4654708.5 | 4389613 | 3335635.25 | 4487746.5 | 4130611.5 | 4028121 | 3860768.25 | 5125362.5 | 4522024 | 3217640.25 | 4468570.5 | 3964872 | 3749218.25 | 4000250.75 | 3887162.25 |
| Q9CR27 | Washc3 | WASH complex subunit 3 Mus musculus | 78.7 | 7.22 | 21093 | 2 | 2 | 1253019.5 | 1583527.12 | 1490122.38 | 1421575.38 | 1757485.62 | 1655527.12 | 1353083.88 | 1798924.25 | 1438302 | 1487443.5 | 1403555.38 | 1359966 | 1569842.12 | 1446249.25 | 805823.94 |
| Q3UMB9 | Washc4 | WASH complex subunit 4 Mus musculus | 444.22 | 8.44 | 136370 | 7 | 7 | 6317588 | 4460619 | 6048707.5 | 3869675.75 | 5690689 | 5263284 | 4096481.25 | 6448896 | 6584553.5 | 4733543 | 4629424.5 | 5717986 | 4731866 | 5910541 | 5594249 |
| Q8C2E7 | Washc5 | WASH complex subunit 5 Mus musculus | 753.17 | 16.22 | 134110 | 13 | 13 | 15136523 | 15837017 | 13416941 | 14184977 | 13099698 | 12101872 | 12485802 | 16420640 | 17703983 | 12943288 | 14078891 | 12199118 | 14014450 | 15386567 | 15554678 |
| O08816 | Wasl | Neural Wiskott-Aldrich syndrome protein Rattus norvegicus | 38.63 | 4.59 | 54325 | 1 | 1 | 182711.09 | 1 | 1 | 1 | 221595.27 | 1 | 1 | 1 | 141900.66 | 200085.06 | 1 | 1 | 1 | 1 | 1 |

| | | | | | | | | | | | | | | | | | | | | | | |
|---|---|---|---|---|---|---|---|---|---|---|---|---|---|---|---|---|---|---|---|---|---|---|
| P59328 | Wdhd1 | WD repeat and HMG-box DNA-binding protein 1 Mus musculus | 158.15 | 4.03 | 124254 | 4 | 4 | 514364.81 | 1815379.12 | 523639 | 405847.94 | 554513.62 | 537546.12 | 662503.69 | 534961.56 | 647426.06 | 587409.75 | 1116014 | 551526 | 423999.28 | 471033.66 | 697052.12 |
| O88342 | Wdr1 | WD repeat-containing protein 1 Mus musculus | 1398.86 | 54.46 | 66407 | 22 | 22 | 92064257 | 82679289 | 89392897 | 99880233 | 90978177 | 84273321 | 78843457 | 79241313 | 83858249 | 95232401 | 92627553 | 91938761 | 95186313 | 82716689 | 85771993 |
| Q8K1X1 | Wdr11 | WD repeat-containing protein 11 Mus musculus | 375.65 | 9.32 | 135937 | 7 | 7 | 1746721.25 | 2654190.5 | 1646001.75 | 2812441 | 2961961.5 | 1955377.25 | 2086253.38 | 2719508.5 | 2230777 | 2175385.25 | 2686044.5 | 2767506 | 2943005 | 2816075 | 2865922 |
| Q9JJA4 | Wdr12 | Ribosome biogenesis protein WDR12 Mus musculus | 431.24 | 21.75 | 47347 | 6 | 6 | 3496997.75 | 1817594.12 | 2151153.25 | 2161436.5 | 1697828.5 | 2104478.25 | 1931527.38 | 2951268.5 | 5326280 | 1984307.5 | 2325240.25 | 3804059 | 2929128.5 | 3334911.75 | 3519312.75 |
| Q4VBE8 | Wdr18 | WD repeat-containing protein 18 Mus musculus | 108.02 | 9.05 | 47211 | 3 | 3 | 1 | 453416.16 | 309514.03 | 1 | 325002.22 | 1 | 646285.5 | 1 | 1 | 1 | 1 | 1 | 1 | 1 | 1 |
| Q8C6G8 | Wdr26 | WD repeat-containing protein 26 Mus musculus | 366.31 | 14.04 | 70543 | 6 | 6 | 3310922.25 | 2065627.88 | 3672164.5 | 3922655.5 | 4275283 | 3687136.5 | 4254889.5 | 3832485.5 | 3080745.5 | 3223011 | 4822341 | 3635488.75 | 3660766 | 3584416.25 | 4574874 |
| Q8BHB4 | Wdr3 | WD repeat-containing protein 3 Mus musculus | 417.79 | 10.62 | 105775 | 8 | 8 | 1887802.75 | 2171924.25 | 2491827.75 | 3237982 | 2894142.5 | 3494599.5 | 3987529.5 | 2149668 | 1652421 | 2117337.25 | 2840286 | 2510044.75 | 2183589.75 | 3633126 | 2796250 |
| Q6ZQL4 | Wdr43 | WD repeat-containing protein 43 Mus musculus | 536.53 | 21.57 | 75381 | 9 | 9 | 3795969 | 3502650.5 | 4467145 | 4847759.5 | 1830786.75 | 3710048 | 4319607.5 | 3890060.25 | 5051064 | 2254400.75 | 5698956 | 3634829.5 | 3892129 | 3637013.5 | 3808892.25 |
| Q6NVE8 | Wdr44 | WD repeat-containing protein 44 Mus musculus | 41.3 | 0.77 | 101555 | 1 | 1 | 3851419.75 | 3086079.75 | 2049291.12 | 3511173.75 | 4193344 | 2258744.75 | 2203582.5 | 2233053 | 1548110.5 | 2491681.75 | 2745036 | 928694 | 2054426.88 | 2293182 | 2118948 |
| Q9Z0H1 | Wdr46 | WD repeat-containing protein 46 Mus musculus | 233.7 | 11.09 | 69048 | 4 | 4 | 6926345.5 | 3481912 | 699087.38 | 5402370 | 4405157 | 5364512 | 828379.06 | 4283558 | 2570715 | 3493718.25 | 2154979.75 | 4388679 | 4033581.5 | 4959962 | 438193.09 |
| Q9ERF3 | Wdr61 | WD repeat-containing protein 61 Mus musculus | 345.45 | 28.52 | 33773 | 5 | 5 | 5545834.5 | 5051768 | 5856887.5 | 4464965 | 5519179 | 5936384.5 | 5153745 | 7148342 | 6942700 | 5738829 | 7427465 | 6948221 | 5304578.5 | 5788818.5 | 6239546 |
| Q920I9 | Wdr7 | WD repeat-containing protein 7 Mus musculus | 94.65 | 1.61 | 163450 | 2 | 2 | 1002719.735 | 1159235.5 | 904080.81 | 981449.44 | 966113.19 | 939328.44 | 744535.19 | 1000400.31 | 749268.19 | 714154.12 | 782606.38 | 1 | 1 | 905023.94 | 539627.19 |
| Q8VCG3 | Wdr74 | WD repeat-containing protein 74 Mus musculus | 171.13 | 15.36 | 42636 | 4 | 4 | 1 | 1 | 1 | 914682.19 | 551223.12 | 703320.405 | 586424.5 | 806778.75 | 534078.12 | 886000.25 | 1 | 634355.56 | 1 | 1 | 1 |
| Q3U821 | Wdr75 | WD repeat-containing protein 75 Mus musculus | 276.24 | 7.83 | 94037 | 5 | 5 | 1112476.62 | 1787282.12 | 2255014 | 1392409.75 | 1587204.88 | 1072125 | 1784262.5 | 1665839.88 | 1589466.38 | 1712178.62 | 1972825.75 | 1641123.38 | 1007283.38 | 1473469.62 | 1849289.12 |
| Q99J09 | Wdr77 | Methylosome protein 50 Mus musculus | 227.72 | 19.88 | 36943 | 4 | 4 | 482799.38 | 1696376.75 | 1950942.75 | 1685725.88 | 1953306.25 | 583816 | 2084632.75 | 1575378.25 | 1929636.5 | 2149346 | 813795.88 | 1760512 | 1538492 | 1649279.5 | 952543.88 |
| Q8BFQ4 | Wdr82 | WD repeat-containing protein 82 Mus musculus | 122.95 | 10.22 | 35079 | 2 | 2 | 1241876.62 | 1899788.25 | 2435892.25 | 1747820.75 | 1870588.5 | 1396507 | 2378175.25 | 2022044.38 | 2905090 | 1248211.88 | 1351515.88 | 2124019 | 2034812.12 | 2360761.25 | 2698842.5 |
| B2RYI0 | Wdr91 | WD repeat-containing protein 91 Rattus norvegicus | 81.86 | 3.21 | 83146 | 2 | 2 | 943704.44 | 1 | 1218338.5 | 1 | 1 | 625318.12 | 1330353.38 | 1 | 1 | 1190148.88 | 1087312.62 | 1456386 | 1152519.88 | 1 | 1443699.75 |
| O88286 | Wiz | Protein Wiz Mus musculus | 43.91 | 0.89 | 184291 | 1 | 1 | 1078049.38 | 1476510.38 | 1311159 | 844011.31 | 1269574.75 | 1043124.06 | 1 | 894587.81 | 1797914.75 | 1 | 424755.94 | 780148.81 | 620749.5 | 852027.94 | 669420.5475 |
| P83741 | Wnk1 | Serine/threonine-protein kinase WNK1 Mus musculus | 553.13 | 6.52 | 250934 | 10 | 12 | 110027521 | 83453697 | 72778305 | 98426177 | 83955393 | 62985649 | 59351965 | 63745725 | 49866589 | 74684753 | 78977457 | 30579749 | 51005705 | 59091017 | 56883345 |
| Q3UH66 | Wnk2 | Serine/threonine-protein kinase WNK2 Mus musculus | 130.28 | 1.44 | 227527 | 1 | 3 | 4534244.5 | 5505072 | 4700283 | 5202611 | 5731587 | 5604202 | 4548929 | 5740697.5 | 5597091 | 5980990.5 | 5384795.5 | 4269236 | 6059670 | 5663712.5 | 5220275 |
| Q8BZZ3 | Wwp1 | NEDD4-like E3 ubiquitin-protein ligase WWP1 Mus musculus | 51.43 | 1.31 | 104694 | 1 | 1 | 1038385.44 | 968847.12 | 736336.5 | 636559.19 | 642218.94 | 1138196.88 | 587091.31 | 815132.12 | 1119717.5 | 688653.81 | 665816.31 | 721537.69 | 778652.75 | 988245.56 | 777933.38 |
| Q9DBH0 | Wwp2 | NEDD4-like E3 ubiquitin-protein ligase WWP2 Mus musculus | 138.69 | 5.17 | 98761 | 3 | 3 | 794372.06 | 844796.31 | 1055113.12 | 1053161.5 | 897406.12 | 1015621.19 | 949523.38 | 1203763.88 | 1478663.88 | 1135712.62 | 1042498.62 | 1091671.25 | 1129693.62 | 1194279.5 | 1007118.5 |
| Q00519 | Xdh | Xanthine dehydrogenase/oxidase Mus musculus | 1074.64 | 22.32 | 146562 | 20 | 20 | 33818797 | 31264845 | 26359433 | 38434429 | 29072485 | 24892181 | 29188915 | 34786489 | 30194461 | 32564425 | 26748575 | 26489479 | 29321569 | 40964285 | 34345713 |
| Q6P1B1 | Xpnpep1 | Xaa-Pro aminopeptidase 1 Mus musculus | 787.32 | 36.92 | 69591 | 3 | 15 | 22986119 | 19389019 | 20236287 | 19305361 | 21173427 | 22984553 | 22079649 | 21018479 | 22256475 | 22265023 | 19511599 | 29267021 | 22291719 | 20099303 | 30903043 |
| B7ZMP1 | Xpnpep3 | Xaa-Pro aminopeptidase 3 Mus musculus | 67.36 | 3.36 | 56677 | 1 | 1 | 887883.5 | 1 | 836949.19 | 555046.56 | 692962.06 | 812988.31 | 828603.5 | 823061.88 | 1006657.31 | 723959.75 | 919560.81 | 696899.19 | 945615.94 | 661694.69 | 832359.19 |
| Q6P5F9 | Xpo1 | Exportin-1 Mus musculus | 1634.78 | 36.04 | 123093 | 31 | 31 | 67067733 | 63259401 | 52247017 | 53474477 | 65256521 | 66634925 | 43511073 | 71633977 | 85701241 | 56538101 | 69466689 | 61825485 | 59162297 | 73992657 | 59460549 |
| Q924C1 | Xpo5 | Exportin-5 Mus musculus | 817.62 | 14.62 | 136973 | 13 | 13 | 13229798 | 15525363 | 13483556 | 10659500 | 10972389 | 13140506 | 6741980.5 | 11488685 | 14845118 | 11591848 | 14049870 | 12841740 | 11422185 | 13486297 | 12465499 |
| Q9EPK7 | Xpo7 | Exportin-7 Mus musculus | 77.36 | 2.48 | 123810 | 2 | 2 | 763644.25 | 548928.75 | 446127.56 | 175356.64 | 436515.34 | 425870.42 | 148878.88 | 604726.62 | 691909.56 | 257966.62 | 299719.28 | 622878.38 | 462029.62 | 619458.44 | 517273.03 |
| Q9CRT8 | Xpot | Exportin-T Mus musculus | 443.92 | 11.53 | 109734 | 9 | 9 | 3612543 | 2686021 | 5183394 | 3536804.5 | 3199093.75 | 4168759 | 2774055.5 | 4942350 | 5573550 | 3255352 | 5113925.5 | 3811193.5 | 2683412 | 4814774.5 | 4700869 |
| Q60596 | Xrcc1 | DNA repair protein XRCC1 Mus musculus | 100.73 | 4.75 | 68971 | 2 | 2 | 1 | 1 | 1 | 1 | 326521.94 | 1 | 422946.03 | 1 | 253022.02 | 572903.75 | 1 | 326736.78 | 338712.19 | 1 | 1 |
| P27641 | Xrcc5 | X-ray repair cross-complementing protein 5 Mus musculus | 265.8 | 8.47 | 83057 | 5 | 5 | 4528747 | 4249620 | 5734083 | 3648882.75 | 3654808 | 4502568 | 4667109 | 5387361 | 5471079.5 | 4649770 | 4873184 | 4708862 | 3990443 | 5605610.5 | 5201351 |
| P23475 | Xrcc6 | X-ray repair cross-complementing protein 6 Mus musculus | 82.76 | 4.77 | 69484 | 2 | 2 | 1266195 | 1761616.38 | 1318388.88 | 1495142 | 1732070.88 | 1559999.88 | 1572573.283 | 1487805.75 | 1951526.88 | 1290960.62 | 1470536 | 1401939.88 | 1392979.88 | 1574607.75 | 1460015.878 |
| P97789 | Xrn1 | 5'-3' exoribonuclease 1 Mus musculus | 63.96 | 0.99 | 194307 | 1 | 1 | 1 | 1 | 240862.66 | 1 | 207752.36 | 217364.23 | 168171.09 | 268561.94 | 214867.67 | 217241.2325 | 243868.42 | 238338.66 | 1 | 198737.91 | 280446 |
| Q9DBR1 | Xrn2 | 5'-3' exoribonuclease 2 Mus musculus | 1106.68 | 27.87 | 108687 | 21 | 21 | 15612037 | 14278676 | 19510667 | 14982429 | 17092421 | 17991569 | 16889809 | 18759791 | 21033045 | 15071528 | 20653423 | 21080833 | 19314541 | 21936897 | 22489725 |
| Q91WQ3 | Yars | Tyrosine--tRNA ligase, cytoplasmic Mus musculus | 720.92 | 38.07 | 59105 | 16 | 16 | 28210869 | 21748569 | 27875767 | 28025875 | 24279579 | 25758391 | 31209769 | 23083709 | 26740377 | 28688625 | 26715365 | 30016169 | 23874199 | 24940741 | 30832493 |
| Q8BYL4 | Yars2 | Tyrosine--tRNA ligase, mitochondrial Mus musculus | 102.7 | 6.99 | 52597 | 2 | 2 | 449523.19 | 521184.19 | 537727.75 | 480602.28 | 523785.03 | 787554.62 | 576529.31 | 966171 | 644023.25 | 883885.38 | 554409.25 | 421773.62 | 492188.12 | 558398.06 | 759183.88 |
| P62960 | Ybx1 | Nuclease-sensitive element-binding protein 1 Mus musculus | 806.76 | 49.07 | 35730 | 7 | 10 | 12411681 | 8943158 | 10193305 | 12765116 | 12786760 | 7409431.5 | 12623945 | 5950567.5 | 5466759.5 | 13967870 | 11341111 | 15656913 | 9822615 | 6391223.5 | 16893201 |
| Q9JKB3 | Ybx3 | Y-box-binding protein 3 Mus musculus | 414.99 | 26.87 | 38814 | 3 | 6 | 2490044.5 | 1521259.38 | 2400492.22 | 2960451.75 | 2630213.25 | 395236.62 | 619650.88 | 1 | 1 | 2824585.75 | 1403937 | 3667065 | 1 | 1 | 2472342 |
| Q5M7T4 | Yipf4 | Protein YIPF4 Rattus norvegicus | 38.77 | 4.47 | 27283 | 1 | 1 | 1563630.25 | 1 | 392467.5 | 438158.66 | 1 | 1 | 1 | 633833.25 | 464424.94 | 1 | 1 | 1 | 327032.62 | 697180.19 | 1 |
| Q9CQW1 | Ykt6 | Synaptobrevin homolog YKT6 Mus musculus | 249.38 | 38.38 | 22314 | 6 | 6 | 4701246.5 | 5363432.5 | 5751318 | 3564797.5 | 4207009.5 | 4941291.5 | 5225771 | 6384797 | 6601969 | 3083021.25 | 5935799 | 6058508 | 3069045.5 | 5993260 | 6124572 |
| O88967 | Yme1l1 | ATP-dependent zinc metalloprotease YME1L1 Mus musculus | 165.06 | 8.81 | 80028 | 4 | 4 | 1 | 420507.09 | 493427 | 1 | 1 | 304346.81 | 385634.16 | 453419.97 | 405403.84 | 458317.88 | 242205.56 | 343850.53 | 416398.94 | 396094.56 | 349637.3975 |
| Q91YT7 | Ythdf2 | YTH domain-containing family protein 2 Mus musculus | 116.33 | 5.87 | 62280 | 2 | 3 | 1386122.75 | 836985.88 | 794061.81 | 935737.44 | 1361119.88 | 763902.12 | 807471.81 | 672922.56 | 733388.06 | 744421.1375 | 800132.44 | 888842.56 | 616520.44 | 929443.81 | 809574.69 |
| Q8BYK6 | Ythdf3 | YTH domain-containing family protein 3 Mus musculus | 215.77 | 8.72 | 63961 | 3 | 4 | 1038374.69 | 836985.88 | 794061.81 | 935737.44 | 1263717.88 | 763902.12 | 807471.81 | 565579.25 | 733388.06 | 717585.31 | 800132.44 | 715230.06 | 588295.12 | 839894.25 | 809574.69 |
| Q9CQV8 | Ywhab | 14-3-3 protein beta/alpha Mus musculus | 1360.37 | 73.17 | 28086 | 9 | 20 | 188957809 | 208764881 | 198768225 | 197832049 | 182322161 | 199231521 | 202878305 | 203882081 | 262777649 | 210340433 | 226072449 | 220333265 | 197025793 | 203466385 | 192485409 |
| P62259 | Ywhae | 14-3-3 protein epsilon Mus musculus | 1780.16 | 85.88 | 29174 | 26 | 30 | 469202977 | 506398337 | 465715905 | 483936257 | 451317441 | 476342017 | 473778625 | 502052065 | 568016193 | 471899489 | 503563745 | 506017633 | 461788865 | 509351457 | 446007777 |
| P61982 | Ywhag | 14-3-3 protein gamma Mus musculus | 1202.65 | 71.26 | 28303 | 11 | 19 | 212918113 | 242106865 | 219071857 | 219759761 | 206556465 | 220144369 | 239932945 | 221655041 | 277120065 | 228657601 | 237277665 | 242348065 | 221679553 | 225774273 | 214991009 |
| P68510 | Ywhah | 14-3-3 protein eta Mus musculus | 1279.82 | 60.98 | 28212 | 14 | 23 | 232024433 | 251203857 | 241545553 | 238275393 | 221811153 | 233217937 | 241562449 | 243044641 | 304326849 | 243777985 | 266048129 | 259141889 | 233215841 | 243425985 | 231755425 |
| P68254 | Ywhaq | 14-3-3 protein theta Mus musculus | 1359.98 | 73.47 | 27778 | 14 | 21 | 197620593 | 216977153 | 212601121 | 204379681 | 192531041 | 204647361 | 215985921 | 228252625 | 290007745 | 218751377 | 242305857 | 239982529 | 202162257 | 222999457 | 207387601 |
| P63101 | Ywhaz | 14-3-3 protein zeta/delta Mus musculus | 1487.54 | 78.37 | 27771 | 15 | 24 | 370861025 | 412231649 | 403401985 | 414648225 | 381436033 | 381194145 | 438805409 | 407026305 | 453680769 | 450459425 | 431656417 | 443676097 | 396828449 | 402120257 | 401630689 |
| P0C6P6 | Yy2 | Transcription factor YY2 Rattus norvegicus V=1 | 40.73 | 2.13 | 41805 | 1 | 1 | 670397.06 | 767132.5 | 757949.31 | 1078833.25 | 853574.81 | 721269.56 | 1069226.25 | 700152.19 | 704739.38 | 1150494.75 | 525686.56 | 745128.31 | 1092363.88 | 812341.75 | 1082315.75 |
| Q8BGC4 | Zadh2 | Prostaglandin reductase-3 Mus musculus | 63.86 | 7.69 | 40529 | 1 | 1 | 227485.3 | 342538.455 | 262807.34 | 575417.56 | 304443.62 | 237109.14 | 278108.91 | 291937.72 | 215249.05 | 536909.12 | 371284.5325 | 286830.28 | 655500.94 | 315529.16 | 227277.75 |
| Q505B7 | Zbtb8os | Protein archease Mus musculus | 47.66 | 5.95 | 19727 | 1 | 1 | 1287060.12 | 1329972.12 | 1224755.25 | 1403056.38 | 1149620.25 | 1227718.38 | 922835.88 | 1382794.5 | 1972079 | 1107142.38 | 811937.5 | 1053506.75 | 908428.81 | 1220635.12 | 902848.06 |
| Q6NZF1 | Zc3h11a | Zinc finger CCCH domain-containing protein 11A Mus musculus | 217.47 | 8.33 | 86492 | 5 | 5 | 1478486.5 | 1571028 | 1751738.25 | 1529590 | 1550828.5 | 439925.28 | 1228319 | 501807.97 | 1153117.12 | 1342855.25 | 1137574.62 | 1445505.75 | 1545351.75 | 605430.94 | 1186179.25 |
| Q3TIV5 | Zc3h15 | Zinc finger CCCH domain-containing protein 15 Mus musculus | 202.4 | 9.62 | 48327 | 3 | 3 | 379547.72 | 478608.06 | 597786.88 | 451137.66 | 644188.06 | 1178483.62 | 950049.62 | 682897.75 | 816810.62 | 648497.94 | 1054115.88 | 724461.31 | 645336.62 | 532725.44 | 1222010.12 |
| Q6ZPZ3 | Zc3h4 | Zinc finger CCCH domain-containing protein 4 Mus musculus | 56.5 | 1.23 | 140967 | 1 | 1 | 1 | 1 | 1 | 1 | 208738.22 | 1 | 160299.05 | 1 | 1 | 1 | 484093.34 | 1 | 168554.92 | 1 | 134188.2 |
| Q3UPF5 | Zc3hav1 | Zinc finger CCCH-type antiviral protein 1 Mus musculus | 33.78 | 1.48 | 106688 | 1 | 1 | 899400.81 | 874464.06 | 819844.75 | 934696.69 | 805721.5 | 737528.56 | 1146249.12 | 759289.25 | 699264.62 | 1082468.88 | 819385.69 | 758639.88 | 933364.81 | 1219575.12 | 994220.81 |
| Q80YV2 | Zc3hc1 | Nuclear-interacting partner of ALK Mus musculus | 80.12 | 3.79 | 55196 | 1 | 1 | 192793.98 | 1 | 213104.2 | 964038.62 | 65215.49 | 1 | 122768.12 | 181111.48 | 366158.19 | 1 | 142965.31 | 206343.72 | 1 | 193408.91 | 1 |
| Q9CYA6 | Zcchc8 | Zinc finger CCHC domain-containing protein 8 Mus musculus | 171.88 | 7.48 | 78026 | 3 | 3 | 1134459.75 | 1824514.62 | 2568045 | 2077077.5 | 2022760.75 | 1920274.25 | 1596319.12 | 2613807 | 1914622.12 | 1972629.75 | 2188873.5 | 1673856.25 | 2260453.5 | 1602111.25 | 2195209.5 |
| Q9D270 | Zdhhc21 | Probable palmitoyltransferase ZDHHC21 Mus musculus | 44.93 | 3.02 | 31331 | 1 | 1 | 429161.62 | 221667.58 | 216614.39 | 305444.67 | 354335.09 | 406610.19 | 246655.58 | 259089.28 | 280996.38 | 298337.8575 | 273284.88 | 222657.27 | 374556.38 | 259143.59 | 420269.12 |
| Q9Z2U2 | Zfp292 | Zinc finger protein 292 Mus musculus | 34.66 | 0.41 | 301048 | 1 | 1 | 923951.88 | 2765312 | 2547395 | 1599058 | 1505235.12 | 1128554.38 | 1181828 | 965151.19 | 2146554.25 | 836511.81 | 1 | 1 | 1081832.38 | 1 | 1 |
| Q9DB43 | Zfpl1 | Zinc finger protein-like 1 Mus musculus | 43.99 | 3.87 | 34155 | 1 | 1 | 368699.84 | 422595.25 | 443284.66 | 287951.34 | 471469.69 | 308148.88 | 320042.28 | 464991.94 | 664968.25 | 351438.59 | 393765.69 | 286881.16 | 432801.19 | 334756.16 | 254435.08 |
| O88532 | Zfr | Zinc finger RNA-binding protein Mus musculus | 109.18 | 2.79 | 116859 | 2 | 2 | 806900.19 | 1133684.25 | 1 | 1 | 1342708.88 | 1135409.75 | 953730.25 | 1211809 | 1196508.12 | 1124364.28 | 1814285.75 | 1589720.75 | 1260961.25 | 1051262.25 | 1 |
| Q3TXX3 | Zfyve27 | Protrudin Mus musculus | 53.67 | 3.37 | 46202 | 1 | 1 | 278513.91 | 1 | 330742.41 | 278902.81 | 1 | 305099.47 | 1 | 1 | 315205.84 | 400711.12 | 469674.09 | 304465.72 | 340971.375 | 359516.47 | 230229.22 |
| Q80W54 | Zmpste24 | CAAX prenyl protease 1 homolog Mus musculus | 40.98 | 2.32 | 54735 | 1 | 1 | 2383957.25 | 1678499.5 | 1610838.38 | 497427.59 | 1428656.25 | 1703202.62 | 1747835.5 | 1507223.5 | 1692495.968 | 1811722.25 | 1546261.75 | 1553818.75 | 1560685 | 1722636.5 | 1466675.88 |
| O88466 | Znf106 | Zinc finger protein 106 Mus musculus | 52.08 | 0.74 | 208964 | 1 | 1 | 1 | 168882.31 | 150078.77 | 1 | 204795.86 | 190128.83 | 174592.8175 | 145654.83 | 248670.41 | 113917.2 | 251768.58 | 1 | 123172.58 | 1 | 1 |
| Q9JMD0 | Znf207 | BUB3-interacting and GLEBS motif-containing protein ZNF207 Mus musculus | 167.13 | 6.67 | 52793 | 3 | 3 | 1021271.69 | 1589116.12 | 1309227 | 1383830.12 | 988423.06 | 1410846 | 1703649.75 | 798411.56 | 1826288.75 | 1887776.38 | 1180916.88 | 935107.75 | 828752.56 | 561604.44 | 2112165 |
| O88291 | Znf326 | DBIRD complex subunit ZNF326 Mus musculus | 87.14 | 2.76 | 65225 | 1 | 1 | 671484.12 | 1 | 434279.41 | 1 | 1 | 1 | 1029194.62 | 638941.75 | 1 | 569395.88 | 405542.31 | 562168.28 | 665541.19 | 545948.81 | 631640.81 |
| Q80YR4 | Znf598 | E3 ubiquitin-protein ligase ZNF598 Mus musculus | 36.77 | 1.54 | 99192 | 1 | 1 | 669236.88 | 730088.88 | 643516.69 | 1071744.75 | 744059.88 | 495694.12 | 682799.31 | 683198.31 | 474349.41 | 693498.19 | 686416.5 | 494240.09 | 442985.97 | 536930.94 | 613733.69 |
| Q91VY9 | Znf622 | Zinc finger protein 622 Mus musculus | 208.9 | 11.97 | 53451 | 4 | 4 | 1 | 293586.12 | 963740.31 | 971338.75 | 1 | 1 | 1584403.12 | 1 | 299052.12 | 908500.25 | 448399.41 | 608758.75 | 436256.44 | 397106.09 | 946375.12 |
| Q62384 | Zpr1 | Zinc finger protein ZPR1 Mus musculus | 87.28 | 4.58 | 50715 | 2 | 2 | 1537296.62 | 715885 | 1433982.25 | 958626.5 | 816974.56 | 1166884.5 | 1197394 | 1521117.88 | 1680227 | 753266.06 | 2687497.5 | 1540901.25 | 990624.75 | 1598998.12 | 1245680 |
| O35986 | Zranb2 | Zinc finger Ran-binding domain-containing protein 2 Rattus norvegicus | 55.42 | 5.45 | 37350 | 2 | 2 | 2275240 | 1912361.75 | 2048002.38 | 2818413.75 | 396779.72 | 2819589.25 | 4015935 | 3103283.75 | 2330255.5 | 3314549.5 | 3620474.25 | 2438190 | 3774589.75 | 2166895.25 | 3606103.25 |
| O54692 | Zw10 | Centromere/kinetochore protein zw10 homolog Mus musculus | 209.48 | 7.57 | 88063 | 4 | 4 | 1426157.75 | 666612 | 925775.31 | 897703.06 | 928143.38 | 1456184.88 | 739761.62 | 1320476.75 | 1915430 | 1380504.88 | 748105.5 | 832629.5 | 994234.94 | 1074728.25 | 1189612.25 |
| Q9CQU5 | Zwint | ZW10 interactor Mus musculus | 42.04 | 9.52 | 28713 | 1 | 1 | 315264.34 | 276110.19 | 346374.72 | 1 | 291008.81 | 372255.28 | 271654.38 | 595865.25 | 435728.6325 | 503139.62 | 315121.03 | 297309.62 | 302287.16 | 360445.25 | 343291.75 |
| Q62523 | Zyx | Zyxin Mus musculus | 239.13 | 14.36 | 60546 | 5 | 5 | 1166992.75 | 729891.44 | 1693928.548 | 2839388.5 | 2039441.5 | 1244043 | 3569779.5 | 1501020.38 | 928861.12 | 3097308 | 3094110 | 2159499 | 1812958.5 | 2156824.75 | 2498835 |
| Q5SSH7 | Zzef1 | Zinc finger ZZ-type and EF-hand domain-containing protein 1 Mus musculus | 95.71 | 1.54 | 328312 | 2 | 2 | 178799.81 | 368272.22 | 280991.78 | 1 | 1 | 272683.72 | 159895.34 | 233682.6325 | 259499.88 | 242651.59 | 206653.02 | 176736.53 | 145319.83 | 313113.97 | 284782.88 |
