## Supplemental Tables 2-5 for "Effects of true to life polyethylene terephthalate and polycaprolactone nanoparticles on macrophages under a repeated exposure mode"

Supplementary table 2: List of modulated proteins in response to PET particles, selected by the Mann Whitney U test (p<0.05)

| accession | gene_name | description | protein_set_s | coverage | MW | #specific_seq | #peptides | Ctrl1 rep | Ctrl5 rep | Ctrl4 rep | Ctrl3 rep | Ctrl2 rep | PET1 rep | PET5 rep | PET4 rep | PET3 rep | PET2 rep | PCL1 rep | PCL5 rep | PCL4 rep | PCL3 rep | PCL2 rep | U-PET | U-PCL | ratio PET | ratio PCL |
|---|---|---|---|---|---|---|---|---|---|---|---|---|---|---|---|---|---|---|---|---|---|---|---|---|---|---|
| Q99KI0 | Aco2 | Aconitate hydratase, mitochondrial Mus musculus | 1715.58 | 40.38 | 85464 | 27 | 27 | 113500217 | 122642265 | 125108009 | 117830081 | 128722697 | 129978497 | 160286513 | 133078353 | 141122497 | 142741457 | 136772321 | 125233145 | 136051793 | 119969961 | 138064673 | 0 | 4 | 1.163546419 | 1.079447786 |
| Q8JZQ2 | Afg3l2 | AFG3-like protein 2 Mus musculus | 377.27 | 12.72 | 89519 | 8 | 9 | 4367963.5 | 4278519.5 | 4811836.5 | 5242098 | 4480939 | 5735902 | 5469554 | 6236984.5 | 5945194 | 5792793.5 | 4642165.5 | 4680066 | 5307019 | 4978489.5 | 4488273.5 | 0 | 7 | 1.25878863 | 1.039456578 |
| P97449 | Anpep | Aminopeptidase N Mus musculus | 868.1 | 21.74 | 109651 | 10 | 14 | 13964333 | 13958208 | 17584101 | 11778133 | 13488800 | 11772538 | 10127049 | 11210214 | 9983079 | 7814818.5 | 11232727 | 9667924 | 12634146 | 14740973 | 10715277 | 0 | 5 | 0.719303759 | 0.833517976 |
| Q61024 | Asns | Asparagine synthetase [glutamine-hydrolyzing] Mus musculus | 807.35 | 33.51 | 64283 | 14 | 14 | 22282757 | 16045791 | 22277965 | 22302875 | 22211675 | 30421359 | 24829973 | 23572991 | 23361801 | 23082309 | 32671895 | 23484009 | 20606761 | 21644909 | 33878257 | 0 | 8 | 1.191658735 | 1.25841413 |
| Q3KRE0 | Atad3 | ATPase family AAA domain-containing protein 3 Rattus norvegicus | 182.8 | 7.28 | 66759 | 3 | 3 | 1 | 1 | 153757.5 | 1 | 196889.88 | 254735.39 | 295225.88 | 1215284.38 | 277314.22 | 380743.91 | 288744.34 | 286751.28 | 231031.12 | 314241.22 | 455088.31 | 0 | 0 | 6.910883085 | 4.494095429 |
| P97450 | Atp5pf | ATP synthase-coupling factor 6, mitochondrial Mus musculus | 111.26 | 17.59 | 12496 | 1 | 1 | 1423215.25 | 1208755.62 | 1075771.62 | 1035423.06 | 1000578.38 | 842888.81 | 597105.88 | 827233.88 | 864608.19 | 643821.56 | 591309.88 | 787930.12 | 747338 | 664091.12 | 644505.38 | 0 | 0 | 0.657351436 | 0.598072362 |
| Q07813 | Bax | Apoptosis regulator BAX Mus musculus | 635.21 | 63.02 | 21395 | 9 | 9 | 33075229 | 33938641 | 33393683 | 25691449 | 33127145 | 36354877 | 35049613 | 39079985 | 39443761 | 36925825 | 38119441 | 34537001 | 33858145 | 36521817 | 29488679 | 0 | 5 | 1.173513675 | 1.083522312 |
| Q9CY64 | Blvra | Biliverdin reductase A Mus musculus | 753.23 | 58.64 | 33525 | 14 | 14 | 48784881 | 54583561 | 49228573 | 50434457 | 51295057 | 43970165 | 42104349 | 46433065 | 47243597 | 43225813 | 43782461 | 44680997 | 49135057 | 50856097 | 44507869 | 0 | 4 | 0.876735077 | 0.915997564 |
| P98086 | C1qa | Complement C1q subcomponent subunit A Mus musculus | 50.02 | 6.12 | 25974 | 1 | 1 | 476973.56 | 391320.94 | 672807 | 879280.31 | 652380.44 | 298784.91 | 198500.78 | 298704.5 | 1 | 1 | 521637.09 | 649038 | 600793.12 | 348516.56 | 827070.19 | 0 | 10 | 0.259047764 | 0.959089809 |
| Q6PHZ2 | Camk2d | Calcium/calmodulin-dependent protein kinase type II subunit delta Mus musculus | 490.44 | 26.45 | 56369 | 9 | 9 | 15552478 | 16226495 | 16876161 | 17452865 | 16448511 | 18309389 | 19223837 | 19063947 | 17683485 | 18653835 | 17858675 | 18811947 | 17461829 | 17797345 | 16200679 | 0 | 4 | 1.125707627 | 1.067516965 |
| P29452 | Casp1 | Caspase-1 Mus musculus | 890.3 | 41.29 | 45640 | 16 | 16 | 26517867 | 27996515 | 28144805 | 26615815 | 29227735 | 23771153 | 22518755 | 23031809 | 24661715 | 23420683 | 30273639 | 24538399 | 27380405 | 26345641 | 18624521 | 0 | 7 | 0.847666389 | 0.918123409 |
| P29594 | Casp2 | Caspase-2 Mus musculus | 45.19 | 1.99 | 50661 | 1 | 1 | 778569.94 | 877849.38 | 870675.62 | 927981.62 | 863769.14 | 1 | 1 | 1 | 1 | 1 | 1 | 1 | 798172.12 | 831248.62 | 1 | 0 | 2 | 1.15772E-06 | 0.377282231 |
| P24860 | Ccnb1 | G2/mitotic-specific cyclin-B1 Mus musculus | 75.11 | 4.88 | 48052 | 1 | 2 | 1 | 1 | 210056.78 | 1 | 1 | 305435.56 | 494806.69 | 343590.62 | 217477.61 | 356642.62 | 147209.69 | 347259.97 | 278458.2025 | 229709.59 | 389653.56 | 0 | 1 | 8.178361996 | 6.628038859 |
| P30276 | Ccnb2 | G2/mitotic-specific cyclin-B2 Mus musculus | 62.46 | 5.53 | 45383 | 1 | 2 | 1 | 1 | 210056.78 | 1 | 1 | 305435.56 | 261999.73 | 285388.88 | 217477.61 | 356642.62 | 147209.69 | 347259.97 | 1 | 1 | 389653.56 | 0 | 7 | 6.793007243 | 4.208901919 |
| P10810 | Cd14 | Monocyte differentiation antigen CD14 Mus musculus | 919.15 | 51.37 | 39204 | 13 | 13 | 46938701 | 51451925 | 49639101 | 46202521 | 44246909 | 34420477 | 35725377 | 42861973 | 38397501 | 40717569 | 40970305 | 48022557 | 48638241 | 49530917 | 40180361 | 0 | 9 | 0.805617142 | 0.953300841 |
| P21855 | Cd72 | B-cell differentiation antigen CD72 Mus musculus | 200.92 | 13.28 | 40347 | 4 | 4 | 5472007 | 4998917 | 4320119.5 | 4838255 | 4878189.5 | 6957509.5 | 8683093 | 5973143 | 5982663 | 7385373 | 5618317 | 5174573.5 | 5244746 | 2656689.75 | 5098806.5 | 0 | 8 | 1.427391559 | 0.970851552 |
| Q9QYB1 | Clic4 | Chloride intracellular channel protein 4 Mus musculus | 1131.61 | 74.31 | 28729 | 17 | 17 | 97964481 | 99478145 | 87956129 | 100850505 | 100354753 | 121194929 | 137273777 | 113730889 | 105239289 | 123683441 | 115262889 | 114896017 | 109952089 | 98826713 | 118439625 | 0 | 3 | 1.235341898 | 1.145443355 |
| Q99KK2 | Cmas | N-acylneuraminate cytidylyltransferase Mus musculus | 181.28 | 11.11 | 48058 | 4 | 4 | 11565397 | 13408209 | 12874419 | 9874251 | 13388289 | 13956426 | 13551037 | 14111877 | 14843570 | 13556021 | 13995903 | 12372027 | 11551751 | 13045272 | 12435074 | 0 | 12 | 1.145774565 | 1.037464258 |
| O89001 | Cpd | Carboxypeptidase D Mus musculus | 285.86 | 6.25 | 152406 | 7 | 7 | 5301483 | 4425193.5 | 3982986.5 | 4140849 | 5125316.5 | 6938925 | 6277641 | 7419425.5 | 6260566.5 | 5917052 | 4396314.5 | 5032049 | 5245310 | 5387845.5 | 4855307.5 | 0 | 8 | 1.428179619 | 1.08448 |
| P47199 | Cryz | Quinone oxidoreductase Mus musculus | 44.42 | 6.65 | 35269 | 1 | 1 | 1067594.25 | 1308112.12 | 1488117.25 | 1360542.88 | 1056024.75 | 1550064.12 | 1889349.38 | 1563990.12 | 1663548.75 | 1596370.88 | 2305991.75 | 1844169 | 1825380.5 | 1225050.12 | 2296096.75 | 0 | 3 | 1.315733833 | 1.512117278 |
| Q62426 | Cstb | Cystatin-B Mus musculus | 145.46 | 57.14 | 11046 | 3 | 3 | 7521179 | 7137837.5 | 6904083 | 12055766 | 6465746.5 | 3764908.75 | 5119400 | 4618857 | 3382433.5 | 4055859 | 4358063 | 4224369.5 | 5296863.5 | 7141643.5 | 5777587 | 0 | 3 | 0.522431357 | 0.668548981 |
| P97821 | Ctsc | Dipeptidyl peptidase 1 Mus musculus | 595.33 | 16.67 | 52376 | 7 | 7 | 49376273 | 52577253 | 45175817 | 45652129 | 52748553 | 44029841 | 30403867 | 40893277 | 42000561 | 30968901 | 39618785 | 36447381 | 45088625 | 42878961 | 35434073 | 0 | 0 | 0.766897845 | 0.812396875 |
| P53395 | Dbt | Lipoamide acyltransferase component of branched-chain alpha-keto acid dehydrogenase complex, mitochondrial Mus musculu | 201.85 | 10.79 | 53247 | 4 | 4 | 934585.25 | 1303108.5 | 1277103.75 | 1273488.12 | 1362557.25 | 1402114.75 | 2359273.5 | 1484669.5 | 1433422.25 | 1512868 | 812452.94 | 1735044.25 | 1600530.12 | 1197393.5 | 1292173.75 | 0 | 11 | 1.331906565 | 1.079135771 |
| Q91X52 | Dcxr | L-xylulose reductase Mus musculus | 65.94 | 7.38 | 25746 | 2 | 2 | 134480689 | 89065553 | 74447881 | 112225169 | 119284305 | 70548753 | 53814041 | 66494233 | 38887289 | 62245973 | 104602489 | 16328652 | 57226097 | 66423661 | 48135453 | 0 | 2 | 0.551441559 | 0.552812773 |
| Q501J6 | Ddx17 | Probable ATP-dependent RNA helicase DDX17 Mus musculus | 957.64 | 31.54 | 72399 | 11 | 18 | 150077089 | 155918529 | 158525361 | 155182561 | 152922897 | 131302857 | 131145177 | 134895601 | 144248145 | 136351825 | 151269873 | 142305521 | 149909057 | 144687137 | 144611137 | 0 | 1 | 0.877453285 | 0.94843082 |
| Q9CQJ6 | Denr | Density-regulated protein Mus musculus | 212.91 | 19.19 | 22166 | 3 | 3 | 4609877 | 5230410 | 5181511.5 | 4923461 | 4933535 | 3900525 | 4577298 | 4487641 | 4550900.5 | 4414574.5 | 5179689 | 3995056 | 4462624 | 4221531.5 | 4472574 | 0 | 3 | 0.881511321 | 0.897610795 |
| Q6P6T4 | Eml2 | Echinoderm microtubule-associated protein-like 2 Rattus norvegicus | 151.15 | 7.7 | 70711 | 3 | 3 | 5018700.938 | 6989340 | 7348600 | 2637560.75 | 3099303 | 9533779 | 8244659 | 8465422 | 8236636 | 9192467 | 10269151 | 3617734.5 | 8022556 | 7789995.5 | 8097664.5 | 0 | 3 | 1.740409064 | 1.506250401 |
| Q3UMY5 | Eml4 | Echinoderm microtubule-associated protein-like 4 Mus musculus | 82.13 | 1.62 | 110027 | 1 | 1 | 1 | 1 | 1 | 1 | 1 | 1337647.12 | 868349 | 530229.88 | 575253.5 | 713995 | 719144.62 | 913039.44 | 572256.31 | 1 | 1 | 0 | 5 | 805094.9 | 440888.474 |
| P17182 | Eno1 | Alpha-enolase Mus musculus | 2394.93 | 77.42 | 47141 | 24 | 32 | 483027841 | 544320321 | 569124609 | 442800673 | 497630657 | 598201857 | 853236929 | 634970049 | 668755073 | 674073537 | 642606273 | 680968833 | 657382657 | 604103617 | 606306369 | 0 | 0 | 1.351741063 | 1.257977291 |
| Q8R180 | Ero1a | ERO1-like protein alpha Mus musculus | 1208.87 | 53.66 | 54084 | 17 | 19 | 34275897 | 37585565 | 33318769 | 30588371 | 32604637 | 46392561 | 59067661 | 50951521 | 56834589 | 49306309 | 41812017 | 39553205 | 38353937 | 44429225 | 42118393 | 0 | 0 | 1.559348995 | 1.225056774 |
| Q8R2E9 | Ero1b | ERO1-like protein beta Mus musculus | 314.36 | 12.85 | 53518 | 3 | 5 | 6508726 | 6918732 | 7956128 | 5295514.5 | 5025839.5 | 9151859 | 12265912 | 9667431 | 12705076 | 8577916 | 7859587.5 | 7979355.5 | 6369195 | 8053586 | 7467939.5 | 0 | 5 | 1.651736102 | 1.190024756 |
| P26151 | Fcgr1 | High affinity immunoglobulin gamma Fc receptor I Mus musculus | 275.57 | 22.77 | 44888 | 6 | 6 | 10544918 | 9299983 | 11888508 | 9437379 | 9035087 | 7002304.5 | 8227308 | 6693892.5 | 8077838.5 | 7228624 | 8291136 | 11055223 | 9099139 | 10037064 | 8133480 | 0 | 8 | 0.741546034 | 0.928497751 |
| P45878 | Fkbp2 | Peptidyl-prolyl cis-trans isomerase FKBP2 Mus musculus | 125.57 | 19.29 | 15344 | 2 | 2 | 3244866 | 3025755.5 | 3779809.5 | 4243410.5 | 2996151 | 2733578.75 | 2386004.5 | 2459459.25 | 2125663 | 2787699.75 | 2336236.5 | 2201685.5 | 2678028.75 | 2236693.25 | 1730822.25 | 0 | 0 | 0.722522306 | 0.646817299 |
| Q8BTM8 | Flna | Filamin-A Mus musculus | 5048.24 | 49.87 | 281222 | 80 | 84 | 369365921 | 351985377 | 418238977 | 354114593 | 340416193 | 337599777 | 334478689 | 328578017 | 331724321 | 332812321 | 304110753 | 310819073 | 362230049 | 334443937 | 349253377 | 0 | 4 | 0.907897063 | 0.905533023 |
| P35922 | Fmr1 | Synaptic functional regulator FMR1 Mus musculus | 128.12 | 3.58 | 68989 | 1 | 2 | 8126656.5 | 8517745 | 8133283 | 8277533 | 8584711 | 9212033 | 9449282 | 9479262 | 9120482 | 9621187 | 9348197 | 8808661 | 8166096.5 | 7797563.5 | 7339911.5 | 0 | 12 | 1.12589641 | 0.995689258 |
| Q9WVR4 | Fxr2 | Fragile X mental retardation syndrome-related protein 2 Mus musculus | 228.17 | 7.88 | 73743 | 3 | 4 | 8126656.5 | 8517745 | 8133283 | 8277533 | 8584711 | 9212033 | 9449282 | 9479262 | 9120482 | 9621187 | 9348197 | 8808661 | 8166096.5 | 7797563.5 | 7339911.5 | 0 | 12 | 1.12589641 | 0.995689258 |
| Q9CZD3 | Gars | Glycine--tRNA ligase Mus musculus | 1395.78 | 37.59 | 81878 | 3 | 22 | 96797529 | 94607849 | 95651737 | 99651545 | 93605913 | 107853665 | 118991417 | 105272225 | 107549889 | 113072337 | 106628337 | 102042625 | 91966089 | 99328257 | 117757241 | 0 | 6 | 1.150786514 | 1.077882242 |
| P17439 | Gba | Glucosylceramidase Mus musculus | 288.44 | 20.39 | 57622 | 6 | 6 | 7411030.5 | 6799296.5 | 6604925 | 8099719.5 | 6469604 | 5614887 | 4592642.5 | 6353095 | 6260019 | 5126661 | 5678315 | 6319695 | 6561909.5 | 7467879.5 | 4540328 | 0 | 5 | 0.789816017 | 0.863882824 |
| Q8BTZ7 | Gmppb | Mannose-1-phosphate guanyltransferase beta Mus musculus | 49.39 | 5 | 39917 | 1 | 1 | 4203369.5 | 4833290.5 | 4242727.5 | 4922349.5 | 5575047 | 3862472.25 | 3928683.5 | 3214383.25 | 2715428.75 | 3238080.25 | 1932733.88 | 2753327.47 | 3486299.25 | 2734631.75 | 2859645 | 0 | 0 | 0.713260801 | 0.578994928 |
| P63213 | Gng2 | Guanine nucleotide-binding protein G(I)/G(S)/G(O) subunit gamma-2 Mus musculus | 83.93 | 39.44 | 7850 | 2 | 2 | 6892465 | 7611842 | 7428047 | 8924962 | 8298338.5 | 6617675 | 2847632.25 | 6250014.5 | 6781411.5 | 6216025 | 2560274 | 5558617 | 6667112 | 6446683 | 5136033 | 0 | 0 | 0.733297875 | 0.673433233 |
| Q99P91 | Gpnmb | Transmembrane glycoprotein NMB Mus musculus | 299.24 | 10.45 | 63676 | 6 | 6 | 15079978 | 18625543 | 14805663 | 15678684 | 18425977 | 12073892 | 9368046 | 14159889 | 13001834 | 9741060 | 20380285 | 19490113 | 17591201 | 17466241 | 19633593 | 0 | 4 | 0.706217082 | 1.144591973 |
| Q91Z53 | Grhpr | Glyoxylate reductase/hydroxypyruvate reductase Mus musculus | 301.1 | 23.17 | 35329 | 6 | 6 | 4643523 | 5200505 | 4224973.5 | 4792153.5 | 4646698.5 | 5630431 | 6908860 | 5576433 | 5530829 | 6749450.5 | 5653425 | 4668840.5 | 5520226.5 | 5481964 | 4573659 | 0 | 6 | 1.293014843 | 1.101679275 |
| O08730 | Gyg1 | Glycogenin-1 Rattus norvegicus | 45.97 | 4.8 | 37378 | 1 | 1 | 3535684.75 | 3298503.5 | 3479268.125 | 3422287 | 3660597.25 | 1947706.38 | 1722916 | 2134305.25 | 1502266 | 1826798.408 | 2593475.25 | 1 | 1821876.38 | 2305709.25 | 1 | 0 | 0 | 0.525052494 | 0.386349234 |
| P62959 | Hint1 | Histidine triad nucleotide-binding protein 1 Rattus norvegicus | 163.67 | 35.71 | 13777 | 3 | 3 | 41841437 | 48005725 | 47729601 | 49812909 | 48736601 | 39706669 | 35915009 | 35364945 | 39126501 | 41288473 | 44455905 | 38582081 | 40950265 | 40046777 | 41511153 | 0 | 1 | 0.810590006 | 0.870492633 |
| Q9JKR6 | Hyou1 | Hypoxia up-regulated protein 1 Mus musculus | 2199.98 | 52.25 | 111181 | 37 | 37 | 68673689 | 74321409 | 74723737 | 70327585 | 73010289 | 88632897 | 82681649 | 94504201 | 89198097 | 76154657 | 68467225 | 82716937 | 69306761 | 66171125 | 72108873 | 0 | 8 | 1.194193295 | 0.993669172 |
| P03975 | Iap | IgE-binding protein Mus musculus V=1 | 590.01 | 23.16 | 62747 | 11 | 11 | 21785451 | 19728165 | 25046163 | 23513657 | 20108189 | 32126171 | 28558429 | 29440323 | 33373827 | 30037465 | 20853213 | 19034379 | 22356197 | 25871409 | 17249657 | 0 | 10 | 1.393482942 | 0.956283364 |
| Q0GNC1 | Inf2 | Inverted formin-2 Mus musculus | 432.48 | 8.25 | 138560 | 8 | 8 | 1647709.12 | 1456264.75 | 1339991.88 | 1237981.5 | 1380480 | 1829130.75 | 1885823 | 2223330.5 | 2042381 | 1897470.62 | 1680831.38 | 2652184.5 | 2136495 | 2325219.75 | 2560672.25 | 0 | 0 | 1.398688513 | 1.607861218 |
| Q6P9P6 | Kif11 | Kinesin-like protein KIF11 Mus musculus | 575.89 | 14.16 | 118027 | 9 | 9 | 1801478.25 | 1844002.5 | 1682134.75 | 1739286 | 1683549 | 2451006.5 | 2595642.75 | 2073151 | 1864301.5 | 2058784.5 | 1944497.75 | 2636322.5 | 2014056 | 2490137.25 | 2025372.5 | 0 | 0 | 1.261979169 | 1.269693029 |
| Q922S8 | Kif2c | Kinesin-like protein KIF2C Mus musculus | 95.48 | 4.72 | 81085 | 2 | 2 | 242709.95 | 1 | 275403.44 | 1 | 402230.97 | 641076.12 | 495552.03 | 824558.69 | 536093.19 | 496042.81 | 886416.38 | 885364.88 | 568168.5 | 839148.69 | 272486.38 | 0 | 2 | 3.252387329 | 3.750310731 |
| P06151 | Ldha | L-lactate dehydrogenase A chain Mus musculus | 2215.19 | 87.05 | 36499 | 28 | 32 | 712083585 | 734669377 | 670777793 | 591029249 | 691721537 | 752190401 | 913085825 | 777360193 | 746621441 | 770667137 | 801903489 | 738947969 | 793248129 | 807796097 | 718574017 | 0 | 1 | 1.164587388 | 1.135338252 |
| Q9Z0M5 | Lipa | Lysosomal acid lipase/cholesteryl ester hydrolase Mus musculus | 683.19 | 22.17 | 45325 | 9 | 9 | 14897863 | 18597939 | 18244205 | 13282737 | 12877767 | 27761769 | 31115139 | 26800337 | 32122465 | 31368847 | 17982701 | 19490887 | 18888273 | 18895353 | 19370957 | 0 | 2 | 1.914859801 | 1.214731069 |
| Q8K1T1 | Lrrc25 | Leucine-rich repeat-containing protein 25 Mus musculus | 53.64 | 6.06 | 32673 | 1 | 1 | 654861.69 | 776330.31 | 764671.89 | 721633 | 905862.56 | 616043.38 | 1 | 1 | 1 | 1 | 547019.38 | 1 | 1401490.25 | 1 | 1 | 0 | 5 | 0.161127246 | 0.50963365 |
| P08905 | Lyz2 | Lysozyme C-2 Mus musculus | 403.81 | 49.32 | 16689 | 4 | 4 | 23830993 | 30157265 | 25073081 | 23394253 | 22318589 | 17470505 | 13516613 | 18438205 | 21674717 | 15363785 | 16017241 | 18982533 | 25560087 | 23386109 | 17933897 | 0 | 5 | 0.692962473 | 0.816514011 |
| P0C5H9 | Manf | Mesencephalic astrocyte-derived neurotrophic factor Rattus norvegicus | 263.78 | 21.23 | 20388 | 5 | 5 | 13466765 | 12135204 | 10767937 | 10744876 | 10496057 | 9700052 | 6837870 | 9948804 | 10329289 | 8493842 | 8759529 | 8539654 | 8978007 | 9614908 | 6948340 | 0 | 0 | 0.786481464 | 0.743617672 |
| P34960 | Mmp12 | Macrophage metalloelastase Mus musculus | 180.84 | 8.67 | 54971 | 4 | 4 | 3640142.5 | 664269.56 | 2515680.75 | 1 | 2652994.75 | 9242780 | 7025381.5 | 7287591 | 7860636 | 4319993.5 | 2635937.25 | 772832.56 | 3587987.5 | 4561804.5 | 3832573.5 | 0 | 6 | 3.772410843 | 1.624721991 |
| P16332 | Mut | Methylmalonyl-CoA mutase, mitochondrial Mus musculus | 326.95 | 15.11 | 82844 | 7 | 7 | 7811386 | 8878732 | 7843282 | 8798362 | 8054289.5 | 9480040 | 9308007 | 9888978 | 9556689 | 10542777 | 5362669.5 | 7747002 | 7738866.5 | 9155077 | 7098422 | 0 | 5 | 1.178573196 | 0.896486513 |
| Q5PPG6 | Nap1l5 | Nucleosome assembly protein 1-like 5 Rattus norvegicus 2 | 35.42 | 4.52 | 16858 | 1 | 1 | 2030543 | 1892679.25 | 1967458.75 | 2355257 | 1713276.25 | 1562184.62 | 1458416.5 | 1537442.88 | 1685425.25 | 1486442.25 | 1721551.62 | 1277105.88 | 1430691.25 | 1837019.5 | 1297264.25 | 0 | 2 | 0.776156764 | 0.759460768 |
| O70145 | Ncf2 | Neutrophil cytosol factor 2 Mus musculus | 1100.57 | 50.48 | 59485 | 21 | 21 | 69790921 | 61107409 | 54740737 | 56351425 | 58903945 | 54505757 | 50419097 | 53201425 | 50333133 | 43332877 | 58715913 | 54120005 | 58673849 | 55727033 | 53169381 | 0 | 5 | 0.836812709 | 0.931908824 |
| Q6AYR2 | Ndrg3 | Protein NDRG3 Rattus norvegicus | 61.03 | 5.07 | 41507 | 1 | 1 | 1 | 1 | 1 | 1 | 1 | 351664.03 | 541392.81 | 489861.38 | 462837.0475 | 468429.97 | 1 | 1 | 576094 | 510622.81 | 1 | 0 | 7.5 | 462837.0475 | 217343.962 |
| O35657 | Neu1 | Sialidase-1 Mus musculus | 37.63 | 2.2 | 44591 | 1 | 1 | 1 | 1 | 1 | 1 | 1 | 301409.75 | 326632.44 | 286419.97 | 171354.48 | 399850.75 | 1 | 1 | 331133.06 | 1 | 1 | 0 | 10 | 297133.478 | 66227.412 |
| P97526 | Nf1 | Neurofibromin Rattus norvegicus | 65.62 | 0.78 | 317083 | 1 | 1 | 279746.625 | 297530.44 | 277005.28 | 275340.78 | 269110 | 187287.83 | 225938.3875 | 238583.58 | 238894.75 | 238987.39 | 242400.69 | 227861.19 | 1 | 1 | 222089.89 | 0 | 0 | 0.807653667 | 0.494986326 |
| E9Q5C9 | Nolc1 | Nucleolar and coiled-body phosphoprotein 1 Mus musculus | 137.62 | 3.85 | 73698 | 3 | 3 | 74356.43 | 100289.9025 | 93275.03 | 72647.51 | 160880.64 | 1 | 1 | 1 | 1 | 1 | 108274.23 | 83402.91 | 1 | 1 | 1 | 0 | 6 | 9.97109E-06 | 0.382252122 |
| P29758 | Oat | Ornithine aminotransferase, mitochondrial Mus musculus | 1084.71 | 50.11 | 48355 | 16 | 16 | 54754557 | 48940553 | 49442789 | 54602601 | 45663557 | 57831221 | 70576329 | 64418657 | 55301041 | 62190553 | 52630437 | 51418689 | 53731941 | 60485337 | 52549909 | 0 | 8 | 1.224596815 | 1.068713407 |
| P09103 | P4hb | Protein disulfide-isomerase Mus musculus | 1769.22 | 55.6 | 57058 | 28 | 28 | 136559073 | 139142049 | 125258673 | 127324713 | 149719969 | 171607425 | 190946033 | 169422401 | 153140801 | 160991313 | 152096065 | 148470625 | 157284705 | 157814065 | 142346193 | 0 | 2 | 1.247938622 | 1.118003905 |
| Q8BFV2 | Pcid2 | PCI domain-containing protein 2 Mus musculus | 113.78 | 8.77 | 46132 | 2 | 2 | 474814.62 | 199473.06 | 396730.5275 | 452961.81 | 459672.62 | 1 | 1 | 1 | 1 | 1 | 215465.2 | 132868.91 | 215523.865 | 266739.91 | 247021.44 | 0 | 4 | 2.5206E-06 | 0.543250015 |
| Q9WUA3 | Pfkp | ATP-dependent 6-phosphofructokinase, platelet type Mus musculus | 1812.23 | 54.97 | 85455 | 20 | 27 | 62324449 | 64117569 | 62208097 | 54761193 | 61724577 | 73752281 | 87211129 | 74364329 | 80531273 | 73031857 | 75789449 | 65038041 | 67673449 | 74192025 | 70099769 | 0 | 0 | 1.274484215 | 1.156182378 |
| E9Q3L2 | Pi4ka | Phosphatidylinositol 4-kinase alpha Mus musculus | 125.61 | 1.9 | 237042 | 2 | 2 | 1 | 1 | 134357.41 | 1 | 1 | 152290.88 | 144313.59 | 156126.62 | 240124.73 | 173213.955 | 315343.44 | 310312.59 | 138144.89 | 1 | 161621.23 | 0 | 3 | 6.445822316 | 6.887566527 |
| O35405 | Pld3 | Phospholipase D3 Mus musculus | 139.88 | 11.27 | 54389 | 3 | 3 | 9169067 | 7480470.5 | 6721049 | 9324024 | 8638230 | 5449954.5 | 5937753.5 | 6070406.5 | 4910840 | 6251654 | 6166208 | 3916503.25 | 5830383 | 5306693 | 4008033 | 0 | 0 | 0.692442333 | 0.610357767 |
| Q80UG2 | Plxna4 | Plexin-A4 Mus musculus | 152.08 | 2.06 | 212560 | 1 | 4 | 202236.52 | 669311.75 | 676408.44 | 395640.09 | 662095 | 759506.31 | 803732.62 | 736717.69 | 854433.12 | 2423596 | 524748.25 | 182676.22 | 179451.16 | 757093.31 | 733826.5 | 0 | 12 | 2.140692825 | 0.912539019 |
| P23492 | Pnp | Purine nucleoside phosphorylase Mus musculus | 568.02 | 53.63 | 32277 | 9 | 9 | 49712993 | 48014693 | 53075109 | 46328725 | 46769449 | 40124613 | 43401909 | 44552413 | 42753189 | 38955417 | 34036297 | 42497253 | 48795501 | 45479489 | 38081717 | 0 | 3 | 0.860134102 | 0.856455216 |
| Q5XIS9 | Prkd2 | Serine/threonine-protein kinase D2 Rattus norvegicus | 73.48 | 1.26 | 96497 | 1 | 1 | 1 | 143415.03 | 1 | 1 | 144991.11 | 161520.64 | 153071.89 | 237913.2 | 235054.06 | 145472.48 | 220041.52 | 193733.435 | 142557.83 | 218713.11 | 193621.28 | 0 | 2 | 3.235099519 | 3.3586563 |
| O35522 | Psmb9 | Proteasome subunit beta type-9 Mus musculus bactrianus | 260.75 | 34.25 | 23427 | 5 | 5 | 23776365 | 27161923 | 22946583 | 26063835 | 24925917 | 20846527 | 22049807 | 21615975 | 21897407 | 22490425 | 22223953 | 19638735 | 25946293 | 24033761 | 19326683 | 0 | 5 | 0.872075834 | 0.890248333 |
| P97814 | Pstpip1 | Proline-serine-threonine phosphatase-interacting protein 1 Mus musculus | 282.97 | 17.11 | 47590 | 5 | 5 | 9298495 | 9274206 | 8585177 | 9588877 | 8833164 | 7979274.5 | 6909640 | 7803951 | 8235725 | 7891581 | 8050923.5 | 6541285 | 8315439 | 8028290 | 6663570 | 0 | 0 | 0.851694614 | 0.824913873 |
| Q8VI36 | Pxn | Paxillin Mus musculus | 80.45 | 9.98 | 64476 | 2 | 2 | 132730.3 | 1 | 1 | 1 | 1 | 228257.44 | 224912.44 | 178405.8 | 174848.36 | 201606.01 | 1 | 1 | 226889.34 | 1 | 228691.73 | 0 | 9 | 7.594344868 | 3.432300995 |
| P61027 | Rab10 | Ras-related protein Rab-10 Mus musculus | 490.94 | 42 | 22541 | 7 | 10 | 21850323 | 21533451 | 21389139 | 20575261 | 19195639 | 22414771 | 24549345 | 22130397 | 24093405 | 22802987 | 22346657 | 21915347 | 20294859 | 21344905 | 23429885 | 0 | 7 | 1.109495643 | 1.04579745 |
| Q60972 | Rbbp4 | Histone-binding protein RBBP4 Mus musculus | 390.27 | 22.35 | 47656 | 3 | 7 | 26910333 | 26352515 | 26713375 | 27903791 | 29951127 | 26322243 | 25654647 | 25885757 | 25857219 | 24366521 | 26589383 | 26965031 | 26018345 | 26632221 | 24660207 | 0 | 5 | 0.92929933 | 0.949460231 |
| O89086 | Rbm3 | RNA-binding protein 3 Mus musculus | 263.18 | 48.37 | 16605 | 4 | 4 | 16201317 | 14056728 | 17451029 | 17496437 | 18559707 | 13890065 | 13601522 | 13944243 | 11691299 | 13727713 | 18621125 | 17886511 | 16596929 | 15088548 | 14502953 | 0 | 12 | 0.798121745 | 0.987236325 |
| P61514 | Rpl37a | 60S ribosomal protein L37a Mus musculus | 169.49 | 41.3 | 10275 | 3 | 3 | 632743.88 | 1557243.75 | 1481535.38 | 558657.25 | 1394191.38 | 1733586.88 | 2303326 | 1922794.47 | 1738659.75 | 1915605.25 | 1619224.38 | 1998351.62 | 1404028 | 603507.44 | 3049755 | 0 | 6 | 1.709341588 | 1.542370774 |
| D3Z8L7 | Rras | Ras-related protein R-Ras Rattus norvegicus | 235.96 | 28.9 | 23909 | 5 | 5 | 5490828.5 | 9192823 | 5410132.5 | 5420434.5 | 6203214 | 11037345 | 10429717 | 12269221 | 12232379 | 10866950 | 7416872 | 8608885 | 8146214 | 8678971 | 7332298.5 | 0 | 5 | 1.791936091 | 1.266913408 |
| Q99P72 | Rtn4 | Reticulon-4 Mus musculus | 984.33 | 25.13 | 126613 | 15 | 15 | 62349229 | 58925973 | 62964929 | 54204753 | 69060793 | 73949689 | 78835825 | 72101969 | 69173321 | 78174937 | 76448497 | 76274977 | 69393881 | 63865777 | 76015553 | 0 | 1 | 1.210500387 | 1.177209763 |
| P42208 | Sept2 | Septin-2 Mus musculus | 642.11 | 43.21 | 41526 | 9 | 10 | 28462507 | 28598655 | 27987833 | 26868779 | 28410479 | 32278057 | 31145361 | 30028825 | 29847553 | 29637537 | 26870447 | 29317097 | 26165293 | 27561039 | 28092311 | 0 | 9 | 1.089854179 | 0.983452612 |
| Q9JJU8 | Sh3bgrl | SH3 domain-binding glutamic acid-rich-like protein Mus musculus | 383.94 | 56.14 | 12811 | 7 | 7 | 55023249 | 54861409 | 54446401 | 59335037 | 60166041 | 50005341 | 43702705 | 49228841 | 49986949 | 45807265 | 53488293 | 50770393 | 51868257 | 51506133 | 43117437 | 0 | 0 | 0.841099614 | 0.883446518 |
| Q9QXX4 | Slc25a13 | Calcium-binding mitochondrial carrier protein Aralar2 Mus musculus | 1659.72 | 63.61 | 74467 | 26 | 29 | 62194497 | 61106117 | 65557537 | 57746685 | 64471141 | 81916873 | 79193121 | 74813577 | 69371073 | 76902177 | 73078401 | 71403545 | 77964905 | 74740009 | 72802361 | 0 | 0 | 1.228628532 | 1.189385386 |
| P16036 | Slc25a3 | Phosphate carrier protein, mitochondrial Rattus norvegicus | 713.67 | 44.66 | 39445 | 12 | 12 | 36062841 | 41107849 | 41221105 | 34584221 | 38753701 | 44266377 | 50259629 | 50509249 | 58528813 | 47451661 | 47089453 | 42631413 | 42631385 | 42273857 | 59066741 | 0 | 0 | 1.309216604 | 1.218866082 |
| Q58A65 | Spag9 | C-Jun-amino-terminal kinase-interacting protein 4 Mus musculus | 403.93 | 9.01 | 146219 | 7 | 7 | 4373358.5 | 4123626 | 3886400 | 3164691 | 5128186 | 5395138.5 | 5981896 | 5632487.5 | 6745488 | 5882954 | 4880454 | 5267936.5 | 4656879 | 5219346.5 | 6306012.5 | 0 | 2 | 1.433429539 | 1.27347144 |
| Q9WUD1 | Stub1 | STIP1 homology and U box-containing protein 1 Mus musculus | 57.9 | 3.95 | 34909 | 1 | 1 | 3146873.188 | 3086832.25 | 2984497.5 | 3650552 | 2865611 | 2347790.25 | 2517792.875 | 2580080.75 | 2306820.25 | 2836480.25 | 2477411 | 2749622.5 | 2833288.75 | 2666649.75 | 2117652.5 | 0 | 0 | 0.800093529 | 0.816342047 |
| O88983 | Stx8 | Syntaxin-8 Mus musculus | 152.78 | 11.86 | 26925 | 2 | 2 | 1879497.563 | 1758616.75 | 2002885.88 | 1978806.12 | 1777681.5 | 1506068 | 1273243.62 | 1465819.25 | 1720422.5 | 1757657.62 | 1576225.25 | 522700.72 | 1897613.12 | 1489897.38 | 1259577 | 0 | 3 | 0.82183783 | 0.717852856 |
| Q0VGY8 | Tanc1 | Protein TANC1 Mus musculus | 276.4 | 5.87 | 200805 | 6 | 6 | 1665739.12 | 1572150.38 | 1584758.25 | 1529621.75 | 1553410 | 2079376.75 | 2248288.5 | 2210395.5 | 1783772.38 | 2533140.5 | 1947109.12 | 2153689 | 2294152 | 2127409.5 | 2663115.25 | 0 | 0 | 1.373060169 | 1.414865714 |
| Q9QXE7 | Tbl1x | F-box-like/WD repeat-containing protein TBL1X Mus musculus | 149.35 | 5.5 | 56802 | 1 | 2 | 2227288 | 1777417.62 | 1661314.12 | 2389203.5 | 1955666.62 | 1457449.62 | 1323082.12 | 1617709.38 | 1014356.56 | 1444561.12 | 1691760 | 689089.94 | 1212120.62 | 1110346.62 | 676803 | 0 | 1 | 0.684969957 | 0.537426768 |
| Q91YX0 | Themis2 | Protein THEMIS2 Mus musculus | 451.31 | 22.32 | 74378 | 9 | 9 | 6519084.5 | 7443011.5 | 6576726.5 | 7534454 | 7404014.5 | 5142677 | 3751690 | 4517776.5 | 4762057.5 | 3096810 | 5101699 | 4559930.5 | 3806501.5 | 3769314.25 | 4837313 | 0 | 0 | 0.599566946 | 0.622222205 |
| Q9Z0U1 | Tjp2 | Tight junction protein ZO-2 Mus musculus | 551.53 | 11.14 | 131280 | 10 | 10 | 4912769 | 5892577 | 5076197 | 6428088.5 | 5285090 | 7309216 | 10987118 | 6877278.5 | 7799566 | 8736109 | 7421016 | 5551806.5 | 5444674.5 | 6376130.5 | 5578498.5 | 0 | 7 | 1.511495142 | 1.100649847 |
| Q8C0L0 | Tmx4 | Thioredoxin-related transmembrane protein 4 Mus musculus | 133.29 | 11.34 | 37131 | 3 | 3 | 2123360.25 | 2472592.25 | 2666886.75 | 2784210 | 2305559.25 | 2829305 | 3001017.25 | 3217444.5 | 3160461.75 | 3169392 | 2631051.25 | 2506664 | 2530072.25 | 2645566.5 | 1942877 | 0 | 12 | 1.244888519 | 0.992197802 |
| Q5NC05 | Ttf2 | Transcription termination factor 2 Mus musculus | 100.85 | 2.9 | 125530 | 3 | 3 | 1898142.75 | 1383248 | 1174040.62 | 1850015.38 | 1632152.38 | 1 | 1081068.25 | 1087629.25 | 259628.09 | 975710 | 1107200.25 | 1 | 695842.94 | 1 | 644047.88 | 0 | 0 | 0.428849648 | 0.30829134 |
| Q6PAM1 | Txlna | Alpha-taxilin Mus musculus | 130 | 6.14 | 62369 | 2 | 2 | 1280284.75 | 1384573.938 | 1239255.75 | 1673974.75 | 1344780.5 | 939730.81 | 456411.59 | 1017399.88 | 1012583 | 1036303.62 | 1006096.12 | 1032015.56 | 1 | 1209596.62 | 1 | 0 | 0 | 0.644592359 | 0.469127753 |
| P25976 | Ubtf | Nucleolar transcription factor 1 Mus musculus | 94.05 | 3.27 | 89509 | 2 | 2 | 1509451.88 | 1644149 | 1590948.5 | 1521410.88 | 2415347 | 1099718 | 1 | 1281186.25 | 1352014.25 | 1 | 1406312 | 1161178 | 1347687.38 | 1292350.5 | 2136781 | 0 | 4 | 0.429995206 | 0.845991123 |
| Q62885 | Unc119 | Protein unc-119 homolog A Rattus norvegicus | 61.98 | 5.42 | 27048 | 1 | 1 | 1 | 1 | 1 | 1 | 1 | 512629.56 | 562204.38 | 684030.78 | 601355.06 | 1059934.12 | 1 | 554497.62 | 552798.31 | 1 | 920076.31 | 0 | 5 | 684030.78 | 405474.848 |
| P70315 | Was | Wiskott-Aldrich syndrome protein homolog Mus musculus | 336.07 | 13.65 | 54192 | 7 | 7 | 5165638.5 | 5253825 | 6108551.5 | 5458462.5 | 6272689 | 4487892 | 5052734.5 | 4846404 | 4716140.5 | 4673451 | 6086846 | 5760981 | 6565875 | 4640480 | 5520264 | 0 | 11 | 0.841377328 | 1.011156716 |
| Q8BYL4 | Yars2 | Tyrosine--tRNA ligase, mitochondrial Mus musculus | 102.7 | 6.99 | 52597 | 2 | 2 | 449523.19 | 521184.19 | 537727.75 | 480602.28 | 523785.03 | 787554.62 | 576529.31 | 966171 | 644023.25 | 883885.38 | 554409.25 | 421773.62 | 492188.12 | 558398.06 | 759183.88 | 0 | 8 | 1.535390443 | 1.108694703 |
| Q6NZF1 | Zc3h11a | Zinc finger CCCH domain-containing protein 11A Mus musculus | 217.47 | 8.33 | 86492 | 5 | 5 | 1478486.5 | 1571028 | 1751738.25 | 1529590 | 1550828.5 | 439925.28 | 1228319 | 501807.97 | 1153117.12 | 1342855.25 | 1137574.62 | 1445505.75 | 1545351.75 | 605430.94 | 1186179.25 | 0 | 2 | 0.592009546 | 0.751115103 |
| Q3TIV5 | Zc3h15 | Zinc finger CCCH domain-containing protein 15 Mus musculus | 202.4 | 9.62 | 48327 | 3 | 3 | 379547.72 | 478608.06 | 597786.88 | 451137.66 | 644188.06 | 1178483.62 | 950049.62 | 682897.75 | 816810.62 | 648497.94 | 1054115.88 | 724461.31 | 645336.62 | 532725.44 | 1222010.12 | 0 | 2 | 1.676318957 | 1.637871344 |
| P41216 | Acsl1 | Long-chain-fatty-acid--CoA ligase 1 Mus musculus | 1095.66 | 40.63 | 77951 | 19 | 21 | 35063401 | 37848509 | 35577673 | 29520365 | 34174989 | 39867261 | 40307941 | 43095697 | 49797837 | 36277381 | 30323187 | 32020637 | 36572393 | 40145453 | 30011147 | 1 | 12 | 1.215821318 | 0.981925713 |
| P58366 | Ankh | Progressive ankylosis protein homolog Rattus norvegicus | 85.84 | 4.27 | 54265 | 1 | 1 | 288447.91 | 143634.0975 | 98545.91 | 99877.84 | 87664.73 | 86453.02 | 1 | 76751.35 | 1 | 91991.62 | 78889.84 | 98150.19 | 109358.18 | 97868.2175 | 105074.66 | 1 | 8 | 0.35534458 | 0.681371758 |
| Q80V94 | Ap4e1 | AP-4 complex subunit epsilon-1 Mus musculus | 78.97 | 1.78 | 124845 | 1 | 1 | 681580.38 | 454765.38 | 548853.5175 | 674298.69 | 384769.62 | 1 | 222189.72 | 450039.84 | 326010.31 | 1 | 1 | 322243.22 | 1 | 405152.5 | 315498.97 | 1 | 1 | 0.363755297 | 0.380027332 |
| Q8VBT9 | Aspscr1 | Tether containing UBX domain for GLUT4 Mus musculus | 141.82 | 6.73 | 59796 | 2 | 2 | 654962.44 | 539152.06 | 575547.265 | 693931.62 | 414142.94 | 1 | 389035.16 | 425997.88 | 395511.19 | 399742.31 | 1 | 348509.91 | 1 | 477888.97 | 603544.5 | 1 | 4 | 0.559567437 | 0.496899375 |
| D3Z6Q9 | Bin2 | Bridging integrator 2 Mus musculus | 608.75 | 23.11 | 52554 | 8 | 9 | 13699195 | 11867651 | 12105952 | 13864309 | 13672661 | 11204565 | 11586169 | 10732576 | 10461511 | 11952553 | 12138047 | 10699399 | 11307664 | 9192905 | 11981107 | 1 | 3 | 0.857806671 | 0.848325699 |
| P30993 | C5ar1 | C5a anaphylatoxin chemotactic receptor 1 Mus musculus | 90.77 | 6.84 | 39023 | 2 | 2 | 3853467 | 2171287.75 | 1685147 | 1996552.75 | 1890232 | 1317881.25 | 597943.56 | 1553283.12 | 1622435.75 | 1719334.75 | 2097651.25 | 1523115.88 | 1648563 | 2039875.88 | 1795117.25 | 1 | 7 | 0.587312456 | 0.785079709 |
| P35564 | Canx | Calnexin Mus musculus | 1226.02 | 49.58 | 67278 | 22 | 22 | 75944481 | 74517137 | 79112257 | 75547153 | 73482825 | 85744721 | 80736617 | 88423929 | 85699313 | 77040265 | 76696785 | 74075913 | 74686977 | 68860737 | 71764617 | 1 | 7 | 1.103118317 | 0.96693424 |
| Q8VEH6 | Cbwd1 | COBW domain-containing protein 1 Mus musculus | 132.73 | 9.16 | 43772 | 2 | 2 | 1518167.38 | 361844.88 | 1746521.25 | 1475068.88 | 1537275.12 | 2163748.25 | 2115234.75 | 2072216 | 1924196.12 | 1723247.12 | 1667754.62 | 1989867.75 | 1655800.62 | 2028631.25 | 1834892.62 | 1 | 2 | 1.506074216 | 1.382303988 |
| P13379 | Cd5 | T-cell surface glycoprotein CD5 Mus musculus | 303.12 | 14.98 | 53849 | 5 | 5 | 2857282.5 | 4351480.5 | 2455905 | 2650475 | 2625699.25 | 4578047.5 | 5763710 | 4574634.5 | 3744935.75 | 5272260 | 3112701 | 3295489.75 | 3840225.5 | 3432162.5 | 2096972.75 | 1 | 9 | 1.601890131 | 1.056001478 |
| Q8BMK4 | Ckap4 | Cytoskeleton-associated protein 4 Mus musculus | 932.77 | 36.7 | 63692 | 15 | 15 | 21448823 | 17194699 | 19210087 | 15689124 | 16096324 | 22998951 | 26209203 | 25156477 | 20380719 | 22363581 | 20912823 | 20589509 | 18392507 | 20472573 | 17722387 | 1 | 7 | 1.306449832 | 1.094275222 |
| Q9D4H8 | Cul2 | Cullin-2 Mus musculus | 222.18 | 7.92 | 86877 | 5 | 5 | 1236850.5 | 1862539.88 | 1368809.25 | 1538803.75 | 1462571.5 | 1604546.5 | 2215151.5 | 2744167.5 | 2896369.25 | 1995367.25 | 1995650.75 | 2344885.75 | 1822770.25 | 2042566.5 | 2058204.75 | 1 | 1 | 1.533635071 | 1.374118094 |
| Q61656 | Ddx5 | Probable ATP-dependent RNA helicase DDX5 Mus musculus | 1563.06 | 58.63 | 69290 | 22 | 29 | 225700161 | 233248081 | 244238129 | 239267329 | 231313041 | 209811377 | 204388465 | 201991633 | 228882321 | 210188881 | 241180161 | 219286065 | 218777841 | 218221761 | 217496545 | 1 | 4 | 0.899039511 | 0.949901147 |
| P21575 | Dnm1 | Dynamin-1 Rattus norvegicus | 204.22 | 5.56 | 97295 | 1 | 5 | 5910187 | 5974341.5 | 4994817 | 5374599.5 | 5425708 | 4874912.5 | 4518357.5 | 4186290.25 | 4259415.5 | 5206855 | 3825795.75 | 4161140.75 | 5243458.5 | 4622964 | 4545649 | 1 | 1 | 0.832591028 | 0.809222861 |
| Q9DCM0 | Ethe1 | Persulfide dioxygenase ETHE1, mitochondrial Mus musculus | 166.01 | 24.41 | 27739 | 4 | 4 | 8303762.5 | 9284230 | 9074427 | 10790023 | 10753311 | 8005515 | 8971851 | 7274087 | 6298631 | 8053064 | 9034530 | 6495023 | 8276262.5 | 8036088.5 | 8133424 | 1 | 1 | 0.800799598 | 0.829264664 |
| Q61545 | Ewsr1 | RNA-binding protein EWS Mus musculus | 292.15 | 12.21 | 68462 | 6 | 6 | 9790247 | 11386937 | 9336300 | 11701854 | 11464452 | 8871031 | 9649932 | 7776542.5 | 8169679.5 | 8879421 | 10322543 | 9094059 | 9754542 | 9174569 | 8572034 | 1 | 3 | 0.807503271 | 0.874030003 |
| P19096 | Fasn | Fatty acid synthase Mus musculus | 6246.8 | 61.86 | 272428 | 51 | 103 | 346178113 | 337950113 | 356189569 | 357735041 | 323721473 | 327054241 | 320900865 | 282894625 | 316224033 | 317165665 | 366675169 | 358927713 | 327721153 | 350089505 | 382199969 | 1 | 6 | 0.908504338 | 1.037077566 |
| P97808 | Fxyd5 | FXYD domain-containing ion transport regulator 5 Mus musculus | 74.01 | 20.79 | 19454 | 2 | 2 | 700896.44 | 1 | 1 | 1 | 1 | 907385.12 | 617907.25 | 1491177.88 | 1603172.88 | 1154910.783 | 415702.38 | 576209.25 | 487871.72 | 749220.69 | 557251.01 | 1 | 4 | 8.238764856 | 3.975250822 |
| Q9R0N0 | Galk1 | Galactokinase Mus musculus | 453.75 | 37.5 | 42295 | 9 | 10 | 8279408 | 12743603 | 18287175 | 10402457 | 10066241 | 20618689 | 16862377 | 18866037 | 20161931 | 22500399 | 19804265 | 19574217 | 20998037 | 19180293 | 18496455 | 1 | 0 | 1.65626098 | 1.640265934 |
| P50442 | Gatm | Glycine amidinotransferase, mitochondrial Rattus norvegicus | 650.58 | 30.97 | 48242 | 10 | 10 | 44178961 | 47179373 | 42780881 | 40331505 | 48363341 | 51043637 | 48570473 | 50060309 | 51739009 | 47313949 | 49021369 | 42606877 | 49612941 | 46641313 | 41424745 | 1 | 10 | 1.116199992 | 1.029049347 |
| Q9CWG1 | Glipr1 | Glioma pathogenesis-related protein 1 Mus musculus | 61.17 | 5.49 | 29129 | 1 | 1 | 1234218.25 | 1205148.25 | 1037347 | 1332321.75 | 1437550.5 | 1373508.38 | 1607569.25 | 1585905 | 1448201.62 | 1643199.25 | 1112794.62 | 1265565.38 | 1540564.75 | 1173762.62 | 1273699.5 | 1 | 12 | 1.226011105 | 1.019178656 |
| P97576 | Grpel1 | GrpE protein homolog 1, mitochondrial Rattus norvegicus | 223.31 | 25.35 | 24297 | 5 | 5 | 266657.03 | 958875.81 | 245341.73 | 1302648 | 955945.62 | 1517999.25 | 1688386.75 | 2520586.75 | 1998680.38 | 1031222.06 | 2168800.75 | 1620054 | 1082695.12 | 1430473.75 | 1375657.88 | 1 | 1 | 2.348022491 | 2.058653167 |
| P10649 | Gstm1 | Glutathione S-transferase Mu 1 Mus musculus | 515.64 | 46.33 | 25970 | 8 | 9 | 12996580 | 10876059 | 17243077 | 11134448 | 11213759 | 15879021 | 21021607 | 19326637 | 17726963 | 18719055 | 17621615 | 16564218 | 18239183 | 10463180 | 16615743 | 1 | 7 | 1.460251409 | 1.252742271 |
| O35737 | Hnrnph1 | Heterogeneous nuclear ribonucleoprotein H Mus musculus | 740.41 | 37.86 | 49199 | 6 | 12 | 74856865 | 82025825 | 78275569 | 89255617 | 91103553 | 78034145 | 68957505 | 73093281 | 74768993 | 73139289 | 93659633 | 73521057 | 82234961 | 74462849 | 74172009 | 1 | 8 | 0.88562642 | 0.957963448 |
| P38647 | Hspa9 | Stress-70 protein, mitochondrial Mus musculus | 2182.07 | 55.67 | 73461 | 31 | 32 | 183261889 | 181052913 | 198643089 | 186770193 | 185384353 | 203123617 | 195938625 | 207787377 | 206125681 | 202956081 | 179096545 | 182381313 | 181658849 | 189346785 | 180384081 | 1 | 6 | 1.08642698 | 0.976211562 |
| P41565 | Idh3g | Isocitrate dehydrogenase [NAD] subunit gamma 1, mitochondrial Rattus norvegicus | 253.8 | 14.76 | 42851 | 6 | 6 | 16545307 | 13982288 | 16847583 | 16369147 | 13910443 | 19518973 | 17887521 | 16911499 | 16665583 | 16975001 | 21632737 | 17221929 | 17784893 | 16977761 | 18943285 | 1 | 0 | 1.132687397 | 1.191950055 |
| P70227 | Itpr3 | Inositol 1,4,5-trisphosphate receptor type 3 Mus musculus | 131.55 | 1.91 | 304275 | 4 | 4 | 639147.94 | 1556119.5 | 1574866.25 | 1596880.12 | 1718204 | 1977254.38 | 1644122.38 | 2018316.5 | 1746773.75 | 1735237.62 | 3644302.75 | 1499567 | 1519330.5 | 1491310.5 | 1839541.25 | 1 | 12 | 1.287427553 | 1.41054972 |
| Q99MN1 | Kars | Lysine--tRNA ligase Mus musculus | 801.4 | 33.45 | 67840 | 4 | 16 | 25529641 | 27604621 | 26149621 | 23147309 | 25424887 | 27076435 | 28545063 | 28187125 | 30812489 | 27797987 | 30966819 | 28971159 | 25493319 | 25335507 | 28560621 | 1 | 7 | 1.113901663 | 1.089720771 |

| | | | | | | | | | | | | | | | | | | | | | | | | | |
|---|---|---|---|---|---|---|---|---|---|---|---|---|---|---|---|---|---|---|---|---|---|---|---|---|---|
| Q8CGK3 | Lonp1 | Lon protease homolog, mitochondrial Mus musculus | 1072.35 | 34.04 | 105843 | 20 | 20 | 28760953 | 28925937 | 29730303 | 31896783 | 30825573 | 31245865 | 40242765 | 34838405 | 34046585 | 40484405 | 30877559 | 34808485 | 33005507 | 30805621 | 38701497 | 1 | 3 | 1.204599496 | 1.120282232 |
| Q8BYI6 | Lpcat2 | Lysophosphatidylcholine acyltransferase 2 Mus musculus | 480.64 | 21.14 | 60254 | 6 | 10 | 2924251.5 | 3248657.25 | 5680712 | 3541806 | 3749135.5 | 6322366 | 8250237 | 6910561 | 8601355 | 5416486 | 6808583.5 | 3752962.5 | 3571998.75 | 6074669 | 7310826.5 | 1 | 3 | 1.854364939 | 1.437433768 |
| O09159 | Man2b1 | Lysosomal alpha-mannosidase Mus musculus | 636.26 | 14.61 | 114648 | 10 | 10 | 29760767 | 29134727 | 24821189 | 32504589 | 31066853 | 23837261 | 17736253 | 27431651 | 22412547 | 18264985 | 23631789 | 21328147 | 27183449 | 28367575 | 20803459 | 1 | 2 | 0.744681195 | 0.823653767 |
| Q9ESL4 | Map3k20 | Mitogen-activated protein kinase kinase kinase 20 Mus musculus | 282.33 | 7.48 | 91720 | 5 | 5 | 9457455 | 9591159 | 8883040 | 8947703 | 9302907 | 10555577 | 10188621 | 10224819 | 9568499 | 9639521 | 8029131.5 | 8647950 | 8369205.5 | 8882327 | 8455357 | 1 | 0 | 1.086500155 | 0.917754292 |
| Q922Q1 | Marc2 | Mitochondrial amidoxime reducing component 2 Mus musculus | 363.39 | 26.04 | 38194 | 3 | 8 | 7982832 | 7891377.5 | 7739494 | 9074475 | 10240457 | 11299854 | 11273633 | 9314421 | 10555654 | 10828539 | 11373523 | 8398520 | 9625687 | 5908090 | 12796351 | 1 | 8 | 1.240945592 | 1.12051479 |
| Q811U4 | Mfn1 | Mitofusin-1 Mus musculus | 55.91 | 1.89 | 83726 | 1 | 1 | 438864.47 | 544852.44 | 673544.94 | 489100.62 | 551190.5 | 686973.5 | 744649.38 | 669053.12 | 767150.31 | 793130.75 | 608298.38 | 691769.12 | 651370 | 453173.53 | 751486.69 | 1 | 6 | 1.357140008 | 1.169985448 |
| Q9DAT2 | Mrgbp | MRG/MORF4L-binding protein Mus musculus | 47.29 | 6.37 | 22436 | 1 | 1 | 802670.06 | 979215.6225 | 1053447.12 | 1030878.19 | 1029867.12 | 1 | 1 | 871928.31 | 787295.5 | 1 | 773234.75 | 1110343.38 | 810500.25 | 685922 | 845000.095 | 1 | 7 | 0.338888958 | 0.862935676 |
| Q9CPW3 | Mrpl54 | 39S ribosomal protein L54, mitochondrial Mus musculus | 56.12 | 13.33 | 15422 | 1 | 1 | 695990 | 584164.38 | 978873.12 | 794012.12 | 825405.12 | 1 | 1 | 1 | 663800.56 | 1 | 1 | 1 | 1 | 1 | 928411.44 | 1 | 4 | 0.171152254 | 0.239378282 |
| Q791V5 | Mtch2 | Mitochondrial carrier homolog 2 Mus musculus | 539.84 | 49.17 | 33499 | 11 | 11 | 20859653 | 18690925 | 19395263 | 19571011 | 20258059 | 26268839 | 24911621 | 26405663 | 20723793 | 24487557 | 20665377 | 19517699 | 24262715 | 22680453 | 20860165 | 1 | 4 | 1.243205099 | 1.093257467 |
| Q80WJ7 | Mtdh | Protein LYRIC Mus musculus | 728.98 | 26.08 | 63846 | 11 | 11 | 11561751 | 14092343 | 12164339 | 11791865 | 11385665 | 14338540 | 15994349 | 14363140 | 16077521 | 13664296 | 13644385 | 12956419 | 13812068 | 10856245 | 13924995 | 1 | 9 | 1.220373322 | 1.06882667 |
| Q9JK81 | Myg1 | UPF0160 protein MYG1, mitochondrial Mus musculus | 541.26 | 36.32 | 42723 | 11 | 11 | 17665041 | 19092641 | 16112001 | 19558717 | 18558771 | 15172226 | 14119496 | 16615068 | 14502936 | 13141444 | 11896425 | 12494022 | 16402564 | 17723461 | 13808548 | 1 | 3 | 0.808368578 | 0.794892502 |
| E9Q634 | Myo1e | Unconventional myosin-Ie Mus musculus | 1357.31 | 32.25 | 126818 | 21 | 26 | 33731409 | 33920177 | 35435257 | 31804043 | 30204529 | 34960029 | 45135253 | 36068857 | 37696557 | 37441933 | 36075349 | 37220225 | 30034521 | 31792161 | 37623081 | 1 | 9 | 1.158739805 | 1.046336369 |
| Q9DCE5 | Pak1ip1 | p21-activated protein kinase-interacting protein 1 Mus musculus | 269.41 | 20.42 | 42116 | 6 | 6 | 1412074.38 | 1034778.94 | 1321317.75 | 1364280.5 | 1623268.38 | 920557.06 | 845591.5 | 1101438.5 | 905271.62 | 740721.88 | 1813598.25 | 1273017.08 | 854659.44 | 988373.75 | 1435436.88 | 1 | 10 | 0.668112443 | 0.942177214 |
| Q63065 | Pdk1 | [Pyruvate dehydrogenase (acetyl-transferring)] kinase isozyme 1, mitochondrial Rattus norvegicus | 190.83 | 11.52 | 49081 | 3 | 3 | 2626979.5 | 3395204 | 1657055.38 | 2282470.5 | 1776084.12 | 3586100.5 | 3640269.5 | 3950429 | 3601029 | 3254742 | 2454559.5 | 2917510.5 | 3570620.5 | 3743815.75 | 2720999.5 | 1 | 4 | 1.536282778 | 1.312640723 |
| Q9JHG7 | Pik3cg | Phosphatidylinositol 4,5-bisphosphate 3-kinase catalytic subunit gamma isoform Mus musculus | 51.14 | 0.82 | 126400 | 1 | 1 | 939349.38 | 814353.44 | 1030546.19 | 1136400.25 | 1166521 | 811272.69 | 590112.06 | 736915.25 | 925851.88 | 690882.62 | 700723.75 | 894396.81 | 883107.5 | 835333.25 | 956282.25 | 1 | 5 | 0.738138161 | 0.83933569 |
| Q63009 | Prmt1 | Protein arginine N-methyltransferase 1 Rattus norvegicus | 1101.34 | 53.54 | 40522 | 20 | 20 | 48532225 | 47985949 | 46430077 | 42264709 | 45721829 | 40695521 | 28322159 | 41056333 | 45250313 | 31501509 | 35086825 | 35684169 | 35647897 | 44480921 | 35413465 | 1 | 1 | 0.808998228 | 0.806778735 |
| Q61207 | Psap | Prosaposin Mus musculus | 500.34 | 16.34 | 61422 | 8 | 9 | 62608617 | 85619153 | 61576009 | 67607481 | 62274225 | 57890369 | 50920657 | 60608353 | 58004545 | 61684089 | 49026825 | 53208489 | 63861249 | 60159913 | 39871305 | 1 | 3 | 0.851104995 | 0.783453497 |
| Q99K85 | Psat1 | Phosphoserine aminotransferase Mus musculus | 1147.95 | 60 | 40473 | 18 | 18 | 65698977 | 68842881 | 75055409 | 73179913 | 70061217 | 74374121 | 82046257 | 75086745 | 77025009 | 78450913 | 70546609 | 76939481 | 65017617 | 69432281 | 84725393 | 1 | 10 | 1.09677135 | 1.03917653 |
| Q8VDM4 | Psmd2 | 26S proteasome non-ATPase regulatory subunit 2 Mus musculus | 1800.66 | 47.8 | 100203 | 29 | 29 | 66501597 | 65012813 | 71737081 | 67672377 | 68472497 | 69628657 | 75036825 | 74853001 | 77078833 | 76089073 | 74211785 | 67134289 | 69674009 | 69574081 | 69056593 | 1 | 6 | 1.098085977 | 1.030213618 |
| P61290 | Psme3 | Proteasome activator complex subunit 3 Mus musculus | 587.29 | 41.73 | 29506 | 10 | 10 | 9824825 | 10620431 | 10491492 | 10125554 | 10774689 | 10111493 | 8752079 | 9348315 | 8149677 | 9397495 | 8986351 | 9282449 | 10676023 | 10458923 | 8910849 | 1 | 6 | 0.882749136 | 0.932048602 |
| A2AWA9 | Rabgap1 | Rab GTPase-activating protein 1 Mus musculus | 81.95 | 2.54 | 120798 | 2 | 2 | 935754.94 | 1237422.44 | 801623.69 | 920566.38 | 2291744.75 | 1 | 680578.38 | 1 | 1 | 842574.5 | 603713.38 | 1708585 | 966373.25 | 1009624.048 | 759824.56 | 1 | 10 | 0.246182036 | 0.815908953 |
| P07091 | S100a4 | Protein S100-A4 Mus musculus | 540 | 84.16 | 11721 | 7 | 9 | 342641921 | 365499169 | 282173601 | 455036193 | 395784161 | 305730529 | 240651697 | 243941521 | 241786849 | 256404145 | 387054721 | 286390977 | 356148481 | 383670561 | 255797857 | 1 | 9 | 0.699848034 | 0.906540018 |
| O08800 | Serpinb8 | Serpin B8 Mus musculus | 234.19 | 15.51 | 42150 | 3 | 3 | 2481567 | 2146444 | 1905547 | 2350871.75 | 1693934.5 | 1737286.12 | 1129401.75 | 1478502.5 | 1597049.38 | 1587338.75 | 996161 | 749999 | 3080298.75 | 2348123.25 | 2155532.25 | 1 | 11 | 0.711790436 | 0.881999715 |
| Q8K2Q9 | Shtn1 | Shootin-1 Mus musculus | 453.89 | 12.84 | 71343 | 9 | 9 | 18959481 | 21447931 | 19827315 | 19525633 | 19291207 | 17314035 | 16918841 | 18066455 | 19119765 | 17418537 | 19144227 | 17340565 | 18011889 | 20242727 | 15561636 | 1 | 5 | 0.896882661 | 0.911656895 |
| P97797 | Sirpa | Tyrosine-protein phosphatase non-receptor type substrate 1 Mus musculus | 389.84 | 23 | 56425 | 7 | 7 | 23094845 | 21341977 | 20353763 | 19189805 | 20057461 | 18328419 | 18098739 | 16482364 | 15807549 | 19767037 | 19855515 | 16491931 | 20869047 | 17731829 | 16703818 | 1 | 4 | 0.85049919 | 0.880949953 |
| Q8BH59 | Slc25a12 | Calcium-binding mitochondrial carrier protein Aralar1 Mus musculus | 1161.56 | 43.72 | 74570 | 17 | 20 | 36904181 | 39552901 | 41379253 | 35412145 | 39784721 | 47953093 | 42510873 | 40043901 | 41561633 | 44538293 | 39124877 | 41970797 | 50035101 | 48464373 | 41117585 | 1 | 4 | 1.122127136 | 1.143392597 |
| O88838 | Spsb2 | SPRY domain-containing SOCS box protein 2 Mus musculus | 35.47 | 2.65 | 28938 | 1 | 1 | 9523748 | 11776717 | 10869080 | 11468616 | 11094955 | 8139882 | 7965588.5 | 9188951 | 10026194 | 7675419 | 9446949 | 8060028 | 9073398 | 9557168 | 6493642.5 | 1 | 1 | 0.785557952 | 0.778891987 |
| Q9R112 | Sqor | Sulfide:quinone oxidoreductase, mitochondrial Mus musculus | 758.22 | 35.11 | 50282 | 12 | 12 | 26959017 | 29870575 | 30165297 | 28299773 | 34691545 | 37028809 | 37249385 | 36522933 | 39867093 | 30743109 | 24611925 | 25713689 | 31846289 | 29247759 | 23953697 | 1 | 6 | 1.209520079 | 0.902572055 |
| Q9JM90 | Stap1 | Signal-transducing adaptor protein 1 Mus musculus | 637.43 | 59.93 | 34628 | 13 | 13 | 17729511 | 12382445 | 13236752 | 13402185 | 15024507 | 21969469 | 22907261 | 20415201 | 17022513 | 22093031 | 24029665 | 22339825 | 17381839 | 18226125 | 24936277 | 1 | 1 | 1.454641493 | 1.489559529 |
| Q8K1E0 | Stx5 | Syntaxin-5 Mus musculus | 146.87 | 13.24 | 39713 | 3 | 3 | 1 | 1 | 1 | 1 | 502111.38 | 558502 | 1549535 | 729857.56 | 759368.19 | 332548.47 | 1 | 237673.48 | 538010.94 | 187900.88 | 1 | 1 | 8 | 7.826510353 | 1.919055537 |
| P62075 | Timm13 | Mitochondrial import inner membrane translocase subunit Tim13 Mus musculus | 36.2 | 14.74 | 10458 | 1 | 1 | 2119784.5 | 2271527.25 | 2447546.5 | 2781436.25 | 2072312.75 | 1758826 | 1823802.25 | 1746608 | 1605808 | 2113046.25 | 1606487.5 | 1664509.88 | 2017443.62 | 1194384.88 | 1110859.12 | 1 | 0 | 0.77383002 | 0.649443263 |
| Q99LG2 | Tnpo2 | Transportin-2 Mus musculus | 428.93 | 10.6 | 100456 | 3 | 8 | 18682063 | 19357955 | 19835205 | 20231513 | 20493809 | 21667879 | 21239207 | 20934807 | 22627675 | 20314281 | 20325561 | 18779817 | 20385281 | 22349967 | 20408797 | 1 | 7 | 1.082994511 | 1.037006672 |
| Q7TQ16 | Uqcrq | Cytochrome b-c1 complex subunit 8 Rattus norvegicus | 94.57 | 24.39 | 9849 | 2 | 2 | 1 | 716689.88 | 1 | 224516.77 | 1 | 875421.19 | 899883.06 | 1153519.12 | 811709.42 | 318014.31 | 288866.69 | 1069416.75 | 164866.89 | 937533 | 356120.97 | 1 | 4 | 4.31205428 | 2.992749065 |
| Q9Z1Q9 | Vars | Valine--tRNA ligase Mus musculus | 2525.76 | 51.46 | 140215 | 44 | 44 | 99511017 | 107113025 | 104057321 | 105556265 | 107404969 | 120400185 | 128984305 | 107190897 | 120652129 | 123631953 | 115033665 | 113963129 | 107953753 | 112612465 | 119659857 | 1 | 0 | 1.147461021 | 1.087044622 |
| Q62465 | Vat1 | Synaptic vesicle membrane protein VAT-1 homolog Mus musculus | 635.9 | 35.47 | 43097 | 10 | 10 | 13013952 | 12600807 | 11770273 | 9058213 | 10924196 | 13991794 | 20890149 | 14604519 | 12760817 | 19295287 | 10335105 | 15816727 | 15712436 | 12519341 | 18737527 | 1 | 6 | 1.421408461 | 1.274610384 |
| P83741 | Wnk1 | Serine/threonine-protein kinase WNK1 Mus musculus | 553.13 | 6.52 | 250934 | 10 | 12 | 110027521 | 83453697 | 72778305 | 98426177 | 83955393 | 62985649 | 59351965 | 63745725 | 49866589 | 74684753 | 78977457 | 30579749 | 51005705 | 59091017 | 56883345 | 1 | 1 | 0.692390166 | 0.61638864 |
| Q91YT7 | Ythdf2 | YTH domain-containing family protein 2 Mus musculus | 116.33 | 5.87 | 62280 | 2 | 3 | 1386122.75 | 836985.88 | 794061.81 | 935737.44 | 1361119.88 | 763902.12 | 807471.81 | 672922.56 | 733388.06 | 744421.1375 | 800132.44 | 888842.56 | 616520.44 | 929443.81 | 809574.69 | 1 | 6 | 0.70043023 | 0.761101395 |
| Q8BYK6 | Ythdf3 | YTH domain-containing family protein 3 Mus musculus | 215.77 | 8.72 | 63961 | 3 | 4 | 1038374.69 | 836985.88 | 794061.81 | 935737.44 | 1263717.88 | 763902.12 | 807471.81 | 565579.25 | 733388.06 | 717585.31 | 800132.44 | 715230.06 | 588295.12 | 839894.25 | 809574.69 | 1 | 4 | 0.736910387 | 0.770840179 |
| O35986 | Zranb2 | Zinc finger Ran-binding domain-containing protein 2 Rattus norvegicus | 55.42 | 5.45 | 37350 | 2 | 2 | 2275240 | 1912361.75 | 2048002.38 | 2818413.75 | 396779.72 | 2819589.25 | 4015935 | 3103283.75 | 2330255.5 | 3314549.5 | 3620474.25 | 2438190 | 3774589.75 | 2166895.25 | 3606103.25 | 1 | 3 | 1.648920404 | 1.651315916 |
| Q66H39 | Abcf3 | ATP-binding cassette sub-family F member 3 Rattus norvegicus | 49.48 | 2.96 | 79856 | 1 | 1 | 501458.97 | 405761.75 | 536086.56 | 779650.12 | 269166.16 | 238527.5 | 274571.72 | 1 | 1 | 364780.91 | 1 | 179937.77 | 1 | 545629.31 | 257873.44 | 2 | 4 | 0.352262682 | 0.394620289 |
| Q9DBL1 | Acadsb | Short/branched chain specific acyl-CoA dehydrogenase, mitochondrial Mus musculus | 542.63 | 33.8 | 47874 | 9 | 9 | 9718412 | 9999263 | 12876759 | 11010008 | 9727089 | 14303861 | 11936437 | 14167736 | 13978011 | 11570947 | 13984924 | 12747951 | 12073137 | 13843721 | 10698769 | 2 | 4 | 1.236735394 | 1.187824554 |
| Q9QUJ7 | Acsl4 | Long-chain-fatty-acid--CoA ligase 4 Mus musculus | 1179.65 | 41.63 | 79077 | 22 | 23 | 77413993 | 74544825 | 73041393 | 67446297 | 74989745 | 67852337 | 61157321 | 69063353 | 66458345 | 63639425 | 72232849 | 54891777 | 66980113 | 68631729 | 61568669 | 2 | 2 | 0.893136642 | 0.882616057 |
| O35643 | Ap1b1 | AP-1 complex subunit beta-1 Mus musculus | 2073.93 | 49.63 | 103935 | 20 | 34 | 43716193 | 42529389 | 50980005 | 37876845 | 44609745 | 55121121 | 50339689 | 52601265 | 61493421 | 48157125 | 57003153 | 63977649 | 48798821 | 53961233 | 61299533 | 2 | 1 | 1.218469657 | 1.297335418 |
| P62944 | Ap2b1 | AP-2 complex subunit beta Rattus norvegicus | 1888.98 | 48.56 | 104553 | 21 | 35 | 62308241 | 63961745 | 70913601 | 56382481 | 64318025 | 75426113 | 65693937 | 72050961 | 84853321 | 64823637 | 76695273 | 76316321 | 66542965 | 71602969 | 70995681 | 2 | 1 | 1.141447392 | 1.139261816 |
| Q9Z1T1 | Ap3b1 | AP-3 complex subunit beta-1 Mus musculus | 311.82 | 5.79 | 122740 | 6 | 6 | 10678527 | 9409184 | 8660827 | 8851153 | 10371617 | 8963745 | 6743306 | 7899023 | 8326370 | 7511278.5 | 8122873 | 7689817 | 8975975 | 8748051 | 8167061 | 2 | 3 | 0.82223571 | 0.86934834 |
| Q78IK4 | Apool | MICOS complex subunit Mic27 Mus musculus | 389.35 | 33.96 | 29261 | 6 | 6 | 8955794 | 9403101 | 9080211 | 6970131 | 10111369 | 10401712 | 12222685 | 10010262 | 9923166 | 12218955 | 11985959 | 8942015 | 9415324 | 9401879 | 10025885 | 2 | 8 | 1.230369146 | 1.117933166 |
| A2BH40 | Arid1a | AT-rich interactive domain-containing protein 1A Mus musculus | 160.72 | 1.14 | 242092 | 0 | 2 | 27109421 | 27486317 | 30441325 | 29938315 | 29588977 | 25654037 | 28426233 | 25855039 | 25336797 | 24106743 | 28633063 | 22574985 | 27561691 | 27858853 | 19271655 | 2 | 6 | 0.894956775 | 0.870894122 |
| Q6PA06 | Atl2 | Atlastin-2 Mus musculus | 92.44 | 4.12 | 66224 | 2 | 2 | 733836.25 | 899531.75 | 923835.75 | 877421.19 | 803702.88 | 972467.75 | 1475015.75 | 963623.56 | 894900.94 | 1103005.38 | 906268.88 | 1224530.5 | 1152944.62 | 1186608.25 | 902368.69 | 2 | 2 | 1.276214019 | 1.267651104 |
| P15535 | B4galt1 | Beta-1,4-galactosyltransferase 1 Mus musculus | 129.14 | 9.52 | 44411 | 3 | 3 | 595688.04 | 677965.44 | 605968 | 461883.34 | 636935.38 | 761654.62 | 611191.19 | 1035714.908 | 1989392.38 | 780621.44 | 1 | 655118.25 | 1 | 979601 | 1 | 2 | 9 | 1.738686759 | 0.548851795 |
| O55028 | Bckdk | [3-methyl-2-oxobutanoate dehydrogenase [lipoamide]] kinase, mitochondrial Mus musculus | 123.32 | 11.41 | 46588 | 2 | 2 | 5335562 | 6367086 | 8666488 | 6777259 | 5026245 | 9499944 | 7600793.5 | 8711683 | 9167563 | 7695803.5 | 8097398.5 | 8480575 | 8272109 | 6295986 | 9204400 | 2 | 6 | 1.326462081 | 1.254185808 |
| Q9WVA3 | Bub3 | Mitotic checkpoint protein BUB3 Mus musculus | 438.52 | 28.83 | 36955 | 7 | 7 | 16847627 | 19373095 | 17989345 | 18040573 | 16922063 | 15482078 | 17457343 | 15816368 | 16721081 | 16299055 | 17017861 | 17207475 | 16927135 | 14871439 | 19596597 | 2 | 11 | 0.917051096 | 0.960164985 |
| Q9WU84 | Ccs | Copper chaperone for superoxide dismutase Mus musculus | 129.26 | 18.25 | 28912 | 2 | 2 | 1 | 405529.62 | 485801.06 | 1 | 1 | 1475405.11 | 2229266.5 | 1904709.75 | 1395607.38 | 372036.81 | 1500236.12 | 2651408 | 1660056.75 | 1752098.88 | 2375369 | 2 | 0 | 8.276390442 | 11.15089553 |
| Q8R1T1 | Chmp7 | Charged multivesicular body protein 7 Mus musculus | 129.92 | 3.77 | 50633 | 2 | 2 | 587070.62 | 493761 | 444649.38 | 485101.81 | 473849.34 | 850190.38 | 634091.69 | 506364.22 | 532246.81 | 628442.25 | 731968.5 | 490311.44 | 686575.38 | 645324.25 | 457355.75 | 2 | 6 | 1.268432849 | 1.212162433 |
| O08585 | Clta | Clathrin light chain A Mus musculus | 123.19 | 10.64 | 25604 | 3 | 3 | 22239505 | 23305617 | 20524421 | 27492887 | 23438983 | 21142393 | 16876155 | 20501821 | 19461057 | 20894157 | 21101131 | 17924757 | 21634389 | 18912855 | 16271817 | 2 | 2 | 0.845080247 | 0.819177705 |
| P13233 | Cnp | 2',3'-cyclic-nucleotide 3'-phosphodiesterase Rattus norvegicus | 99.23 | 5.71 | 47268 | 2 | 2 | 177527.52 | 200313.7825 | 131076.45 | 256356.52 | 236294.64 | 89591.42 | 1 | 139433.39 | 174410.73 | 1 | 185504.78 | 152486.38 | 175437.8975 | 217315.27 | 146445.16 | 2 | 8 | 0.402805573 | 0.87581541 |
| P36552 | Cpox | Oxygen-dependent coproporphyrinogen-III oxidase, mitochondrial Mus musculus | 149.92 | 7.22 | 49715 | 3 | 3 | 1503168 | 1347360.5 | 1329728.25 | 1128550.25 | 1129803.62 | 1433957.12 | 3423644.25 | 3157619 | 1434446 | 3432024 | 3203623.25 | 3139364 | 2982791.25 | 1181355.12 | 1388669.5 | 2 | 4 | 2.000694114 | 1.847572997 |
| Q9ESV0 | Ddx24 | ATP-dependent RNA helicase DDX24 Mus musculus | 345.21 | 10.74 | 96429 | 6 | 6 | 1468147.25 | 1989461.12 | 877275.38 | 1869901.25 | 1698180.5 | 542305.38 | 336972.75 | 565333.94 | 152117.78 | 1528880.75 | 1621332 | 1017331.44 | 643574.25 | 619783.88 | 1416769.75 | 2 | 4 | 0.395498449 | 0.673012089 |
| Q62095 | Ddx3y | ATP-dependent RNA helicase DDX3Y Mus musculus | 1104.92 | 32.98 | 73428 | 2 | 23 | 77379209 | 69522481 | 72049065 | 60994129 | 71025609 | 67367865 | 58533945 | 53458897 | 64835241 | 57428789 | 75097913 | 66131121 | 57115589 | 67263249 | 72751697 | 2 | 10 | 0.859401981 | 0.964068421 |
| P58252 | Eef2 | Elongation factor 2 Mus musculus | 3344.97 | 61.54 | 95314 | 50 | 52 | 326634849 | 330409569 | 354644641 | 332265505 | 309318529 | 333630721 | 392710305 | 367695649 | 369891297 | 342406049 | 362267553 | 376556641 | 344632257 | 381164993 | 371968961 | 2 | 1 | 1.092580547 | 1.110881446 |
| A0A0B4 | Fcgr4 | Low affinity immunoglobulin gamma Fc region receptor IV Mus musculus | 45.23 | 4.02 | 28398 | 1 | 1 | 2495273 | 2113805.5 | 2117607.5 | 2473083 | 2382982.25 | 1806752 | 1644024.25 | 1732195.12 | 1702798.62 | 2170689.5 | 1926141.75 | 1631899.75 | 2058816.12 | 1699792 | 1765408.12 | 2 | 0 | 0.781891909 | 0.784101941 |
| P97807 | Fh | Fumarate hydratase, mitochondrial Mus musculus | 1200.89 | 50.3 | 54357 | 17 | 19 | 86103625 | 83282137 | 88677953 | 80313417 | 82207745 | 89665089 | 109420321 | 96921553 | 86762569 | 88193537 | 81870185 | 91613777 | 85385977 | 86667217 | 86248009 | 2 | 8 | 1.119781273 | 1.026630268 |
| Q8BHN3 | Ganab | Neutral alpha-glucosidase AB Mus musculus | 1370.82 | 41.74 | 106911 | 29 | 29 | 34234589 | 29828711 | 32994331 | 32246355 | 28628169 | 32806983 | 44377493 | 37427353 | 36029913 | 34980285 | 34698565 | 36054397 | 29361417 | 31399247 | 38813193 | 2 | 7 | 1.17532764 | 1.07848094 |
| Q6AYA1 | Gar1 | H/ACA ribonucleoprotein complex subunit 1 Rattus norvegicus | 60.84 | 7.96 | 23011 | 2 | 2 | 3150930.5 | 2862889.313 | 1571096.25 | 3286665 | 3442865.5 | 1 | 1974114.75 | 1 | 1 | 1964938 | 2954199.75 | 1237376.62 | 1381899.5 | 1805204.53 | 1647342.25 | 2 | 4 | 0.275180443 | 0.630553379 |
| Q8BX17 | Gemin5 | Gem-associated protein 5 Mus musculus | 193.13 | 4.53 | 166592 | 5 | 5 | 3853023 | 3625824.5 | 3026013.5 | 2197521.5 | 2166582.25 | 1794819.62 | 1488040.12 | 2048674.12 | 2254236.25 | 1237340.5 | 2442654.25 | 2744502 | 2132943.75 | 2721516.75 | 2527636.75 | 2 | 8 | 0.59339105 | 0.845334811 |
| Q9D1H7 | Get4 | Golgi to ER traffic protein 4 homolog Mus musculus | 133.38 | 12.23 | 36525 | 3 | 3 | 2727767 | 2513847.5 | 2044945.5 | 2748242.75 | 2697812.5 | 1962030.25 | 2441285 | 2295243 | 1941175.62 | 1515364 | 1950727.5 | 869177.31 | 1803638.38 | 1665114.12 | 798530.88 | 2 | 0 | 0.797565753 | 0.55661685 |
| Q8R0H9 | Gga1 | ADP-ribosylation factor-binding protein GGA1 Mus musculus | 122.02 | 6.61 | 69972 | 2 | 2 | 1854207 | 395725.62 | 1687098.5 | 1813656.25 | 1705823.5 | 1896028.12 | 2043798.25 | 1800032.5 | 1993087.62 | 1994358 | 2359629 | 1110457.88 | 1996401 | 698788.19 | 1927901.88 | 2 | 8 | 1.304538364 | 1.085384048 |
| Q6Y7W8 | Gigyf2 | GRB10-interacting GYF protein 2 Mus musculus | 228.63 | 5.65 | 149193 | 4 | 4 | 1181891.62 | 1094963.62 | 1509011.12 | 950411.12 | 983237.94 | 1681937.75 | 1565021.62 | 1586243.38 | 1407920.25 | 1456707.12 | 1357287 | 1923634.62 | 1535261.25 | 1057392.62 | 1491057.25 | 2 | 5 | 1.345888516 | 1.28763229 |
| Q8CIM5 | Gpr84 | G-protein coupled receptor 84 Mus musculus | 320.45 | 27.02 | 43717 | 5 | 5 | 7807401 | 6131564 | 5569723 | 5427608.5 | 7374241 | 4714073 | 4009133.5 | 3872539 | 3643870.25 | 5941217 | 6648972 | 6603202 | 8071299 | 6868107 | 7713785.5 | 2 | 7 | 0.686489129 | 1.111258688 |
| Q99L13 | Hibadh | 3-hydroxyisobutyrate dehydrogenase, mitochondrial Mus musculus | 245.21 | 23.28 | 35440 | 5 | 5 | 2553083.25 | 2951650 | 1218244.12 | 2513776 | 3179769.5 | 3988897.5 | 3606878.5 | 3251169.75 | 2736060.5 | 3410391.5 | 3168675.25 | 3434962.25 | 2676077 | 3338556.25 | 3332386.75 | 2 | 3 | 1.368611642 | 1.284631589 |
| O88736 | Hsd17b7 | 3-keto-steroid reductase Mus musculus | 381.96 | 26.65 | 37317 | 7 | 7 | 4350245.5 | 3922892.75 | 4406959.5 | 3599696 | 3864512 | 4603924 | 4587904.5 | 4547565 | 5614308.5 | 4163731.5 | 3848698 | 4297725.5 | 3673317.5 | 3056918.25 | 4277964 | 2 | 8 | 1.167448201 | 0.950870359 |
| Q8BU30 | Iars | Isoleucine--tRNA ligase, cytoplasmic Mus musculus | 622.99 | 13.55 | 144271 | 14 | 14 | 49474901 | 37118825 | 34908689 | 42228605 | 41116593 | 34525333 | 33683565 | 33059561 | 32200303 | 37844045 | 35971697 | 28782593 | 34869073 | 32809721 | 37057957 | 2 | 2 | 0.836293889 | 0.827400615 |
| Q8BIJ6 | Iars2 | Isoleucine--tRNA ligase, mitochondrial Mus musculus | 743.05 | 18.08 | 112804 | 13 | 13 | 19898681 | 25412257 | 23707031 | 20511649 | 23922073 | 26989709 | 25192323 | 24251637 | 25861863 | 26335175 | 23928361 | 24041737 | 24183341 | 21685737 | 25062351 | 2 | 7 | 1.13379277 | 1.048036622 |
| Q8R4K2 | Irak4 | Interleukin-1 receptor-associated kinase 4 Mus musculus | 74.58 | 3.7 | 50872 | 1 | 1 | 1060406.25 | 1101917 | 1002067.94 | 1146826.38 | 1106412.25 | 878716.38 | 1016870 | 1011746.06 | 947959.81 | 941204.88 | 1032768.5 | 1008588.38 | 1044315.12 | 1044499.31 | 784247.56 | 2 | 4 | 0.885349736 | 0.907116033 |
| Q3V300 | Kif22 | Kinesin-like protein KIF22 Mus musculus | 78.78 | 3.94 | 73190 | 2 | 2 | 1265744.75 | 1064443.25 | 957997.25 | 1238799.25 | 931019.81 | 794203.44 | 776615.88 | 968571.38 | 919149.44 | 734712 | 256059.19 | 425881.59 | 417362.6175 | 313512.31 | 673997.38 | 2 | 0 | 0.768275711 | 0.382339949 |
| P70188 | Kifap3 | Kinesin-associated protein 3 Mus musculus | 232.68 | 5.8 | 91291 | 3 | 3 | 1491804.12 | 1501712.5 | 1419731.25 | 1250890.5 | 1425476 | 1625760.75 | 1426412.38 | 1774483 | 1699261 | 1560379.75 | 1216867.25 | 1421363.5 | 965687.56 | 1567311.25 | 1537212.62 | 2 | 12 | 1.140583459 | 0.946235131 |
| Q921I2 | Klhdc4 | Kelch domain-containing protein 4 Mus musculus 2 | 52.98 | 2.57 | 64860 | 1 | 1 | 1290601.62 | 1177804.25 | 1553057.88 | 1303753.75 | 1331525 | 1186300.75 | 934673.69 | 1011991.88 | 1093696.12 | 1246541.75 | 1430117.75 | 1322820.25 | 1199798.88 | 1135465.5 | 1473135.38 | 2 | 12 | 0.822204583 | 0.985667954 |
| Q05CL8 | Larp7 | La-related protein 7 Mus musculus | 297.48 | 11.4 | 64802 | 5 | 5 | 10513111 | 11670235 | 7440189.5 | 11271201 | 11376124 | 9821228 | 5042094.5 | 6675311 | 8297351 | 5860961 | 6760591.5 | 5860076 | 7100140 | 7035883 | 6525869 | 2 | 0 | 0.682922476 | 0.636732573 |
| Q6P4Z6 | Lcmt1 | Leucine carboxyl methyltransferase 1 Rattus norvegicus | 44.66 | 2.71 | 38330 | 1 | 1 | 2115761.75 | 2183631.25 | 1710164.75 | 2253409.25 | 1937010.12 | 1561821.62 | 1717521 | 1571419.62 | 1810043.12 | 1668137.88 | 1979706.62 | 1382952.62 | 1519326.25 | 1621406.12 | 1396004.88 | 2 | 2 | 0.816564894 | 0.774452373 |
| Q9D0F3 | Lman1 | Protein ERGIC-53 Mus musculus | 395.53 | 26.5 | 57789 | 8 | 8 | 23873111 | 23560533 | 23336553 | 21160401 | 25816227 | 26915873 | 23712867 | 26165949 | 31886691 | 26150517 | 21671843 | 21498739 | 23594359 | 23723969 | 22130595 | 2 | 9 | 1.145100065 | 0.956454707 |
| Q505F5 | Lrrc47 | Leucine-rich repeat-containing protein 47 Mus musculus | 188.49 | 11.88 | 63590 | 4 | 4 | 2665587 | 2880447.25 | 2403833.75 | 2991008 | 3062402.5 | 1663320.75 | 2378221 | 2316510.5 | 2246629.75 | 2719847.5 | 3119483.5 | 2201012 | 1437302.25 | 2387049.75 | 2628460.25 | 2 | 6 | 0.808705583 | 0.840753667 |
| P21708 | Mapk3 | Mitogen-activated protein kinase 3 Rattus norvegicus | 285.39 | 25.53 | 43081 | 3 | 6 | 14046033 | 13884697 | 13244656 | 12927679 | 14114616 | 14595198 | 18821231 | 14158102 | 13997421 | 16445258 | 18213003 | 14374974 | 15016812 | 13967979 | 16860421 | 2 | 2 | 1.143650867 | 1.149748685 |
| P28667 | Marcksl1 | MARCKS-related protein Mus musculus | 205.08 | 34.5 | 20165 | 4 | 4 | 9196065 | 8834829 | 8280734.5 | 8980619 | 9934519 | 9885404 | 10909023 | 9531033 | 10259880 | 10419569 | 10909821 | 8958838 | 11949703 | 9352347 | 9023325 | 2 | 6 | 1.127759355 | 1.109830259 |
| Q9CPU4 | Mgst3 | Microsomal glutathione S-transferase 3 Mus musculus | 221.11 | 33.99 | 16958 | 3 | 3 | 9168685 | 7486690 | 8288514 | 6290236 | 7887625 | 11045711 | 8713239 | 9111533 | 11396111 | 10221440 | 12560580 | 8997930 | 7340395 | 7246861 | 9107664 | 2 | 10 | 1.290536185 | 1.156733275 |
| Q9CQT1 | Mri1 | Methylthioribose-1-phosphate isomerase Mus musculus | 494.63 | 38.75 | 39411 | 9 | 9 | 6321781.5 | 6129971 | 6594098 | 5600923.5 | 6837747 | 7896112 | 6479865 | 7192331 | 8182895 | 9877010 | 8995960 | 6502941 | 7852937 | 6185218 | 5944828 | 2 | 9 | 1.258657008 | 1.126962802 |
| Q9EQI8 | Mrpl46 | 39S ribosomal protein L46, mitochondrial Mus musculus | 139.41 | 14.13 | 32132 | 3 | 3 | 861975.12 | 601137.44 | 545947.44 | 716169.155 | 855616.62 | 1018796.25 | 806903.5 | 944854.56 | 1143185.75 | 909091 | 609543.5 | 872724.38 | 693577.38 | 775235.19 | 863920.81 | 2 | 8 | 1.346841323 | 1.065391111 |
| Q922D8 | Mthfd1 | C-1-tetrahydrofolate synthase, cytoplasmic Mus musculus | 979.69 | 27.38 | 101200 | 20 | 20 | 32226977 | 32587449 | 33619313 | 35887057 | 31192873 | 36066481 | 32808281 | 36769237 | 36545053 | 35955041 | 32689741 | 37854033 | 32517277 | 36351813 | 32547945 | 2 | 8 | 1.076310459 | 1.038952312 |
| Q9D1J1 | Necap2 | Adaptin ear-binding coat-associated protein 2 Mus musculus | 270.8 | 31.95 | 28598 | 2 | 5 | 3710688 | 4054495.5 | 3515519.5 | 3933436.5 | 4115200 | 5116532 | 5677140.5 | 4120635.25 | 3991092.5 | 5271131 | 4688011 | 5291598 | 4013848.5 | 4674000 | 5776335.5 | 2 | 2 | 1.250768618 | 1.264595358 |
| P97333 | Nrp1 | Neuropilin-1 Mus musculus | 105.47 | 3.58 | 103000 | 2 | 2 | 2158467 | 1629235.62 | 1624815.75 | 1570022 | 1405104.25 | 702659 | 1167366.5 | 1577390.88 | 696628.5 | 1222015.88 | 1256722.88 | 1540028.25 | 1073999.75 | 1566660.38 | 1148377 | 2 | 2 | 0.639757763 | 0.785177312 |
| Q9DBY8 | Nvl | Nuclear valosin-containing protein-like Mus musculus | 211.68 | 7.25 | 94476 | 4 | 4 | 5006893 | 3699745.25 | 4004343 | 4415970 | 4322784.5 | 2172377.5 | 2996748.25 | 2547878.25 | 2998211.375 | 4275841.5 | 2058915.62 | 1517011.75 | 2354571.5 | 1917346.75 | 2530980 | 2 | 0 | 0.69889238 | 0.483867295 |
| Q6NS46 | Pdcd11 | Protein RRP5 homolog Mus musculus | 1027.78 | 18.37 | 207779 | 21 | 21 | 11391401 | 10210932 | 14812261 | 10521552 | 6393869 | 15496319 | 15429837 | 13808677 | 15560133 | 12267632 | 18562757 | 16093093 | 13425174 | 8873012 | 13902959 | 2 | 6 | 1.360633369 | 1.328651323 |
| P12382 | Pfkl | ATP-dependent 6-phosphofructokinase, liver type Mus musculus | 576.84 | 22.82 | 85360 | 10 | 12 | 19641895 | 16978521 | 16034875 | 13188097 | 19513529 | 21386235 | 20890561 | 20718255 | 22054261 | 18848263 | 19522317 | 17954289 | 17573799 | 20557623 | 18941789 | 2 | 7 | 1.217213304 | 1.107699532 |
| P25113 | Pgam1 | Phosphoglycerate mutase 1 Rattus norvegicus | 1137.31 | 74.02 | 28832 | 15 | 16 | 160273809 | 158861713 | 135597505 | 130547473 | 140905953 | 185438881 | 210870913 | 170879073 | 152729809 | 166426593 | 168983073 | 153981697 | 170122593 | 155666209 | 161602097 | 2 | 4 | 1.220547788 | 1.115905792 |
| Q9D0F9 | Pgm1 | Phosphoglucomutase-1 Mus musculus | 1225.89 | 48.04 | 61418 | 18 | 18 | 24706457 | 27459615 | 26923959 | 22012243 | 24741713 | 28689809 | 32885623 | 30358749 | 32106349 | 26712269 | 30071991 | 29115397 | 28329307 | 30982065 | 27291501 | 2 | 1 | 1.197934066 | 1.158500016 |
| Q7TSV4 | Pgm2 | Phosphoglucomutase-2 Mus musculus | 85 | 7.74 | 68748 | 2 | 2 | 2201647.75 | 1594923.38 | 2156999.25 | 2122491.75 | 2083694 | 1835397 | 908060.5 | 1710602.38 | 1409135.62 | 1465798.875 | 559778.5 | 1733968.75 | 1853111.38 | 1709271.75 | 2048442.38 | 2 | 4 | 0.721375029 | 0.778027805 |
| Q9Z2Y8 | Plpbp | Pyridoxal phosphate homeostasis protein Mus musculus | 97.09 | 8.03 | 30049 | 2 | 2 | 2219025.25 | 2299979.75 | 2153731.75 | 2477237.75 | 2008060.75 | 2101538 | 1939842.75 | 1925310.62 | 2005135.62 | 2053876.38 | 1700573.5 | 2400256.75 | 2018431.38 | 2054112.38 | 2065341.62 | 2 | 7 | 0.898518704 | 0.917609185 |
| Q6ZQB6 | Ppip5k2 | Inositol hexakisphosphate and diphosphoinositol-pentakisphosphate kinase 2 Mus musculus | 141.43 | 4.07 | 128427 | 3 | 3 | 1313955.12 | 1195377.88 | 756056.56 | 1340971.5 | 1055065.38 | 558345.19 | 984502.94 | 717093.25 | 233355.45 | 968674.94 | 895340.44 | 684509.46 | 769825.56 | 609484.75 | 463387.09 | 2 | 2 | 0.611501678 | 0.604537979 |
| Q9R1P1 | Psmb3 | Proteasome subunit beta type-3 Mus musculus | 493.69 | 53.17 | 22965 | 8 | 8 | 38088821 | 39792409 | 41585749 | 38853289 | 41475689 | 42663561 | 43969641 | 40239129 | 43768621 | 44703805 | 42588357 | 41988505 | 45713765 | 41178841 | 42573357 | 2 | 2 | 1.077823397 | 1.071307089 |
| Q9QVP9 | Ptk2b | Protein-tyrosine kinase 2-beta Mus musculus | 949.64 | 28.25 | 115794 | 20 | 20 | 28455413 | 22882949 | 22717637 | 23024557 | 20719505 | 25209293 | 29832521 | 30859177 | 32050053 | 23046845 | 28831157 | 25656069 | 22287801 | 30755293 | 23496483 | 2 | 6 | 1.196925433 | 1.112281283 |
| P49445 | Ptpn7 | Tyrosine-protein phosphatase non-receptor type 7 Rattus norvegicus | 46.59 | 3.62 | 40314 | 1 | 1 | 504673.72 | 502326.03 | 530789.44 | 704985.31 | 605939.31 | 414063.91 | 501825.22 | 396904.84 | 315470.19 | 510820.41 | 613955.81 | 515263.69 | 472780.59 | 502754.09 | 582398.19 | 2 | 10 | 0.750894864 | 0.943286181 |
| Q05144 | Rac2 | Ras-related C3 botulinum toxin substrate 2 Mus musculus | 504.12 | 51.04 | 21441 | 4 | 8 | 157320433 | 173500753 | 153848577 | 153689553 | 156357969 | 137894273 | 140611185 | 141572081 | 154037185 | 144666257 | 131843929 | 123000297 | 162315857 | 161514865 | 122822481 | 2 | 8 | 0.904448657 | 0.882700606 |
| Q8C2K5 | Rasal3 | RAS protein activator like-3 Mus musculus | 178.56 | 5.67 | 114782 | 4 | 4 | 836467.88 | 783370.44 | 924220 | 756868.38 | 873788.38 | 947397.19 | 888160.69 | 1117444.75 | 1018222.19 | 903517.38 | 846915.88 | 1057478.25 | 909133.25 | 1014630.69 | 878119.38 | 2 | 4 | 1.16768261 | 1.127329018 |
| Q60973 | Rbbp7 | Histone-binding protein RBBP7 Mus musculus | 335.63 | 22.12 | 47790 | 2 | 6 | 21178625 | 20790347 | 20901695 | 22812941 | 24752673 | 21053733 | 19667089 | 20613129 | 19419183 | 19703325 | 20016581 | 20420863 | 21444059 | 21649737 | 19451945 | 2 | 6 | 0.909632759 | 0.932512251 |
| Q9D0F6 | Rfc5 | Replication factor C subunit 5 Mus musculus | 257.05 | 27.14 | 38096 | 6 | 6 | 3489320.75 | 3808726.5 | 3205226.5 | 3610095 | 3384961.5 | 4443581.5 | 4075791.75 | 3680420 | 4294434 | 3690032.5 | 3499960.5 | 3566589.5 | 3502885.25 | 3953832.25 | 3816098.5 | 2 | 6 | 1.153496331 | 1.048063772 |
| P07742 | Rrm1 | Ribonucleoside-diphosphate reductase large subunit Mus musculus | 647.75 | 26.64 | 90210 | 14 | 14 | 18246387 | 12284210 | 5937449 | 16716314 | 16616034 | 9535447 | 4218572.5 | 4721790 | 7043884.5 | 4911893.5 | 5650169 | 4992719 | 5052815.5 | 8698067 | 9365249 | 2 | 2 | 0.435980168 | 0.483650844 |
| Q8BGH2 | Samm50 | Sorting and assembly machinery component 50 homolog Mus musculus | 769.28 | 41.58 | 51864 | 15 | 15 | 21734131 | 19847863 | 20459641 | 19857669 | 19958101 | 22979163 | 21053505 | 21147901 | 22422585 | 23522843 | 20830119 | 20976719 | 21553759 | 21648367 | 22887605 | 2 | 4 | 1.09099576 | 1.059290378 |
| Q5XIW8 | Sart1 | U4/U6.U5 tri-snRNP-associated protein 1 Rattus norvegicus | 126.17 | 4.09 | 91011 | 3 | 3 | 1022317.44 | 1042419.88 | 1167091.5 | 1258104.25 | 1175796.25 | 1229363.62 | 2252825.5 | 1240864 | 1284431.5 | 1898648 | 1657308.75 | 1626504.62 | 1166248.62 | 947230.75 | 1661527.25 | 2 | 8 | 1.395430698 | 1.245880202 |
| Q923V8 | Selenof | Selenoprotein F Rattus norvegicus | 196.66 | 41.36 | 17822 | 4 | 4 | 706844.62 | 1173329.88 | 792476.62 | 955051 | 1068103.25 | 1340912.38 | 1366017 | 1037170.44 | 1315911.25 | 1261245.75 | 1136084.75 | 916536.12 | 1073166.75 | 1364991.38 | 1157744.75 | 2 | 6 | 1.346149664 | 1.20288711 |
| Q4G075 | Serpinb1a | Leukocyte elastase inhibitor A Rattus norvegicus | 69.24 | 2.64 | 42727 | 1 | 1 | 1023972.69 | 1064173.5 | 1096896.38 | 1305444.12 | 1008452.94 | 1294906.5 | 1288696.62 | 1336356.75 | 1678912.5 | 1365857.38 | 847812.5 | 1097065.88 | 1041159.5 | 1034437.62 | 859220.75 | 2 | 8 | 1.266558686 | 0.887388584 |
| Q60854 | Serpinb6 | Serpin B6 Mus musculus | 1224.84 | 60.85 | 42599 | 18 | 19 | 83696937 | 106701513 | 88571385 | 97106305 | 93216201 | 82281505 | 82543825 | 88433337 | 87140977 | 80464817 | 95602673 | 85191641 | 100108713 | 98665201 | 79883217 | 2 | 12 | 0.896806584 | 0.97903035 |
| Q6AYE2 | Sh3glb1 | Endophilin-B1 Rattus norvegicus | 215.15 | 10.68 | 40787 | 3 | 3 | 6654449 | 7580770 | 5955345.5 | 7714775 | 7210518 | 6026126 | 5893195.5 | 5893043 | 5585304.5 | 6307689 | 5398864 | 5289601.5 | 6323193 | 5579678 | 4803519 | 2 | 1 | 0.845924324 | 0.780127767 |
| O55242 | Sigmar1 | Sigma non-opioid intracellular receptor 1 Mus musculus | 126.45 | 13.9 | 25250 | 2 | 2 | 1644043 | 888876.69 | 952398.69 | 788086.75 | 1070364 | 662997.69 | 747085.19 | 912535.81 | 732123.62 | 778910.88 | 673738.19 | 441712.53 | 522908.47 | 675793.81 | 635142.12 | 2 | 0 | 0.717406216 | 0.5519129 |
| Q6P6S4 | Sil1 | Nucleotide exchange factor SIL1 Rattus norvegicus | 140.87 | 6.45 | 52350 | 3 | 3 | 2155357 | 341537.28 | 1373500.62 | 1334709.12 | 1289616.75 | 2665482.5 | 2068998.88 | 2240426 | 1860893.25 | 2282855.75 | 1863360.62 | 1992335.75 | 1529215.5 | 1685061 | 1271360.12 | 2 | 8 | 1.71195295 | 1.284325113 |
| P46062 | Sipa1 | Signal-induced proliferation-associated protein 1 Mus musculus | 144.33 | 4.53 | 112066 | 2 | 2 | 2053289.38 | 2238001 | 2395500 | 2068664.88 | 2462241.25 | 2007359.75 | 1828882.5 | 1950199.5 | 2163299.5 | 1848355.5 | 2374772.5 | 2100890.5 | 1674338.5 | 1975010.75 | 2292782.75 | 2 | 8 | 0.873449976 | 0.928692891 |
| Q5RK19 | Snf8 | Vacuolar-sorting protein SNF8 Rattus norvegicus | 42.99 | 5.43 | 28884 | 1 | 1 | 789109.94 | 790146.25 | 834083.19 | 924989.38 | 875019.69 | 855687.81 | 1011078.25 | 925146.31 | 990464.62 | 1154387.12 | 1024107.56 | 1008168.12 | 940731.44 | 1049158 | 758774.81 | 2 | 5 | 1.171696139 | 1.134712684 |
| Q9CWK8 | Snx2 | Sorting nexin-2 Mus musculus | 1058.94 | 48.36 | 58471 | 17 | 18 | 42139717 | 39019877 | 42667173 | 42373893 | 37328149 | 33036595 | 39527005 | 35774001 | 35753597 | 37002513 | 39356345 | 40050249 | 36835505 | 37219633 | 45065821 | 2 | 9 | 0.889769423 | 0.975427282 |
| Q8VIM6 | Strc | Stereocilin Mus musculus | 35.29 | 0.44 | 196347 | 1 | 1 | 722819.81 | 2210075.5 | 1922223.88 | 1858499.88 | 719493.31 | 2165312.75 | 4443522 | 2196245.5 | 2423662.75 | 3768064.25 | 3559237.75 | 2465075.75 | 3501635.25 | 3825082.5 | 3071432.75 | 2 | 0 | 2.017567673 | 2.209365762 |
| Q5FVQ9 | Tbce | Tubulin-specific chaperone E Rattus norvegicus | 38.82 | 2.1 | 59043 | 1 | 1 | 273605.12 | 1 | 202520.45 | 1 | 264504.56 | 278054.25 | 287940.38 | 258549.41 | 308550.22 | 283273.565 | 203796.41 | 149373.03 | 190033.5 | 214477.62 | 261524.84 | 2 | 12 | 1.912376965 | 1.376129064 |
| Q5EAN7 | Terf2ip | Telomeric repeat-binding factor 2-interacting protein 1 Rattus norvegicus | 69.76 | 3.56 | 43418 | 1 | 1 | 303358.16 | 1 | 368227.56 | 1 | 1 | 357669.56 | 340004.56 | 401303.25 | 572864 | 417960.3425 | 478680.19 | 393511.78 | 1 | 455871.5 | 391174.84 | 2 | 3.5 | 3.111728429 | 2.559958586 |
| A2VCX1 | Tiprl | TIP41-like protein Rattus norvegicus | 88.43 | 6.64 | 31225 | 2 | 2 | 1010077.5 | 1185112.38 | 1003873.19 | 1027271.5 | 951998.25 | 1127915.62 | 1285338.38 | 1153446.62 | 1505938.12 | 1188953.5 | 821440.38 | 1140050.38 | 923500.38 | 1049318.88 | 1118792.5 | 2 | 12 | 1.209190768 | 0.975816483 |
| Q9CR02 | Tma16 | Translation machinery-associated protein 16 Mus musculus | 97.78 | 17.19 | 25782 | 2 | 2 | 2082810.12 | 1679686.38 | 1689868.88 | 2088082.88 | 2318952.25 | 1 | 1882131.25 | 709378.19 | 1 | 1 | 1672875.62 | 1105107.5 | 2073011.38 | 746404.31 | 1399349.703 | 2 | 2 | 0.262846857 | 0.70965253 |
| Q7TQ69 | Tmc3 | Transmembrane channel-like protein 3 Mus musculus | 36.33 | 0.62 | 128691 | 1 | 1 | 1 | 236465.83 | 695763.5 | 361556 | 1 | 411150.22 | 2795173 | 437173.56 | 1151786.54 | 963649.38 | 649359.62 | 938158.88 | 520808.34 | 258791.66 | 2074597 | 2 | 4 | 4.451220511 | 3.433110989 |
| Q9CZR8 | Tsfm | Elongation factor Ts, mitochondrial Mus musculus | 443.4 | 48.77 | 35334 | 7 | 7 | 8348572 | 9932174 | 8412015 | 8252476.5 | 8658664 | 10108733 | 8790735 | 9035529 | 10271758 | 9999639 | 7242701 | 8429215 | 9112904 | 7243775.5 | 8881974 | 2 | 11 | 1.105552309 | 0.938231858 |
| Q9CR09 | Ufc1 | Ubiquitin-fold modifier-conjugating enzyme 1 Mus musculus | 173.85 | 31.14 | 19481 | 4 | 4 | 7331790 | 6625408.5 | 6249144.5 | 5932308 | 6208152.5 | 5879627 | 6233661 | 5425873 | 5544552.5 | 5861724 | 5403181 | 5793377.5 | 6144519.5 | 5187507 | 6461896.5 | 2 | 4 | 0.894846921 | 0.896239454 |
| Q8CCJ3 | Ufl1 | E3 UFM1-protein ligase 1 Mus musculus | 407.08 | 13.49 | 89520 | 6 | 6 | 3361500 | 3305349 | 3690832.75 | 3432519.25 | 1910361.5 | 3603082 | 3889765.5 | 3879137.75 | 3862947.75 | 3473269.5 | 2888599.75 | 3277545.25 | 3403040.25 | 3961524.75 | 2530876 | 2 | 11 | 1.191562563 | 1.022994304 |
| B2RUP2 | Unc13d | Protein unc-13 homolog D Mus musculus | 36.8 | 1.01 | 123119 | 1 | 1 | 247152.02 | 184416.53 | 91451.2 | 211817.7 | 86256.26 | 85754.12 | 1 | 61813.97 | 64886.64 | 177058.64 | 99040.52 | 99626.11 | 89339.3 | 50909.33 | 61351.48 | 2 | 5 | 0.474384793 | 0.487479974 |
| Q9JI13 | Utp3 | Something about silencing protein 10 Mus musculus | 271.13 | 13.43 | 53399 | 5 | 5 | 3654444.75 | 4159856 | 2575663 | 4289463 | 4867771 | 2891608.5 | 3117193.25 | 2570143.5 | 2026012.25 | 2324770 | 3239254 | 3173196.75 | 2751775.5 | 2337231.75 | 1556417.25 | 2 | 3 | 0.661461948 | 0.66801776 |
| Q9WTZ0 | Uxt | Protein UXT Mus musculus | 45.66 | 7.01 | 18187 | 1 | 1 | 840198.69 | 989501.88 | 870619.56 | 887313.31 | 897253.56 | 923897.81 | 971356.62 | 1139656.88 | 1090172.75 | 1031271.015 | 919638.75 | 1059172 | 1111709.5 | 1228954.62 | 790788.12 | 2 | 6 | 1.149717947 | 1.139440746 |

Supplementary table 2: List of modulated proteins in response to PCL particles, selected by the Mann Whitney U test (p<0.05)

| accession | gene_name | description | protein_set_scoverage | | MW | #specific_seq | #peptides | Ctrl1 rep | Ctrl5 rep | Ctrl4 rep | Ctrl3 rep | Ctrl2 rep | PET1 rep | PET5 rep | PET4 rep | PET3 rep | PET2 rep | PCL1 rep | PCL5 rep | PCL4 rep | PCL3 rep | PCL2 rep | U-PET | U-PCL | ratio PET | ratio PCL |
|---|---|---|---|---|---|---|---|---|---|---|---|---|---|---|---|---|---|---|---|---|---|---|---|---|---|---|
| Q5SSL4 | Abr | Active breakpoint cluster region-related protein Mus musculus | 413.1 | 12.81 | 97667 | 8 | 8 | 3828630 | 4065722.5 | 4025891.75 | 1980865 | 4399927.5 | 5660986 | 3625264.25 | 4083371.75 | 7713215.5 | 3323895 | 6216541 | 4869163.5 | 4647166 | 6893983.5 | 6553381 | 9 | 0 | 1.333625676 | 1.59445803 |
| Q9D7B6 | Acad8 | Isobutyryl-CoA dehydrogenase, mitochondrial Mus musculus | 37.13 | 2.42 | 45020 | 1 | 1 | 1885928.75 | 2224785.5 | 1598112.25 | 1778057.62 | 1712586.12 | 2008989.88 | 1310819.12 | 1403083.88 | 1867244.88 | 1415810.75 | 1581461.88 | 1249996.25 | 1519860.75 | 1428049.62 | 1588770.62 | 7 | 0 | 0.870261907 | 0.800930807 |
| Q9WTP7 | Ak3 | GTP:AMP phosphotransferase AK3, mitochondrial Mus musculus | 211.67 | 25.11 | 25426 | 4 | 4 | 2491269.5 | 2715702 | 2521727.25 | 2693779.75 | 2454971.5 | 1306415.75 | 2221138 | 1804367.88 | 2516100.75 | 3011834.5 | 1129692.5 | 2135913.5 | 2132867.5 | 2220516.75 | 2380462 | 7 | 0 | 0.843323552 | 0.776508723 |
| P20291 | Alox5ap | Arachidonate 5-lipoxygenase-activating protein Rattus norvegicus | 38.16 | 4.97 | 18070 | 1 | 1 | 1 | 273593.47 | 1 | 1529866.12 | 199316.78 | 218265.8 | 5853626.5 | 1158138.88 | 187365.38 | 1571717.62 | 3324238 | 2523272.75 | 1541756.88 | 3160707.908 | 5253564 | 6 | 0 | 4.48832198 | 7.890807977 |
| P07356 | Anxa2 | Annexin A2 Mus musculus | 1632.99 | 70.8 | 38676 | 25 | 25 | 413457761 | 402877729 | 423519713 | 427536897 | 441378049 | 452286113 | 617554753 | 461282945 | 422670913 | 530866305 | 623050241 | 496078081 | 489559073 | 457095841 | 531130625 | 3 | 0 | 1.178251233 | 1.231482655 |
| Q1AAU6 | Asap1 | Arf-GAP with SH3 domain, ANK repeat and PH domain-containing protein 1 Rattus norvegicus | 459.92 | 9.53 | 127088 | 9 | 9 | 8820306 | 9953314 | 9387332 | 9091122 | 9597520 | 9544995 | 10366380 | 9029489 | 9568718 | 8646686 | 11066423 | 10552254 | 10286497 | 10128631 | 10680721 | 12 | 0 | 1.006545927 | 1.1251864 |
| O54984 | Asna1 | ATPase Asna1 Mus musculus | 491.22 | 48.56 | 38823 | 11 | 11 | 10520835 | 10712636 | 11106910 | 10759026 | 8052762 | 11138663 | 15411631 | 11149654 | 12599062 | 9894251 | 12256825 | 13523025 | 12745601 | 11721007 | 15265231 | 4 | 0 | 1.176748947 | 1.280721625 |
| Q3KRE0 | Atad3 | ATPase family AAA domain-containing protein 3 Rattus norvegicus | 182.8 | 7.28 | 66759 | 3 | 3 | 1 | 1 | 153757.5 | 1 | 196889.88 | 254735.39 | 295225.88 | 1215284.38 | 277314.22 | 380743.91 | 288744.34 | 286751.28 | 231031.12 | 314241.22 | 455088.31 | 0 | 0 | 6.910883085 | 4.494095429 |
| P97450 | Atp5pf | ATP synthase-coupling factor 6, mitochondrial Mus musculus | 111.26 | 17.59 | 12496 | 1 | 1 | 1423215.25 | 1208755.62 | 1075771.62 | 1035423.06 | 1000578.38 | 842888.81 | 597105.88 | 827233.88 | 864608.19 | 643821.56 | 591309.88 | 787930.12 | 747338 | 664091.12 | 644505.38 | 0 | 0 | 0.657351436 | 0.598072362 |
| P59017 | Bcl2l13 | Bcl-2-like protein 13 Mus musculus | 266.18 | 20.97 | 46719 | 5 | 5 | 5133233 | 7492595.5 | 7480295.5 | 8898548 | 7648284 | 6646305 | 8527362 | 10156359 | 9639056 | 10363053 | 9757421 | 9335088 | 10523231 | 9261940 | 10132683 | 5 | 0 | 1.236793425 | 1.337146259 |
| P51943 | Ccna2 | Cyclin-A2 Mus musculus | 36.96 | 3.32 | 47269 | 1 | 1 | 1 | 1 | 1 | 1 | 1 | 1 | 468130.16 | 581772.12 | 446328.03 | 1 | 639892.56 | 612440.62 | 550589.0225 | 517462.91 | 432560 | 5 | 0 | 299246.462 | 550589.0225 |
| Q9WU84 | Ccs | Copper chaperone for superoxide dismutase Mus musculus | 129.26 | 18.25 | 28912 | 2 | 2 | 1 | 405529.62 | 485801.06 | 1 | 1 | 1475405.11 | 2229266.5 | 1904709.75 | 1395607.38 | 372036.81 | 1500236.12 | 2651408 | 1660056.75 | 1752098.88 | 2375369 | 2 | 0 | 8.276390442 | 11.15089553 |
| Q6A065 | Cep170 | Centrosomal protein of 170 kDa Mus musculus | 176.73 | 2.9 | 175050 | 4 | 4 | 6052194 | 4725799 | 4774246 | 6748361 | 4813490.5 | 4377784 | 4907023 | 3880492.5 | 3632213 | 4531489.5 | 4629409.5 | 2634284.5 | 3921717.5 | 3425801 | 4409769 | 3 | 0 | 0.786639036 | 0.701516486 |
| Q8BTU1 | Cfap20 | Cilia- and flagella-associated protein 20 Mus musculus | 173.2 | 30.05 | 22748 | 4 | 4 | 7629927 | 8499859 | 8663983 | 7950704 | 6956509.5 | 7696530.5 | 4048041 | 7618430.5 | 7600208.5 | 5050671 | 575085.44 | 647639 | 5598790 | 6021284.5 | 4250360 | 4 | 0 | 0.806375044 | 0.430547504 |
| Q8CGZ0 | Cherp | Calcium homeostasis endoplasmic reticulum protein Mus musculus | 102.33 | 5.02 | 106169 | 3 | 3 | 1 | 1 | 1 | 1 | 1 | 1 | 495221.84 | 1 | 552290.31 | 874965.19 | 505553.69 | 337668.66 | 929510 | 647254.5725 | 816285.94 | 5 | 0 | 384495.868 | 647254.5725 |
| D4A4T9 | Chordc1 | Cysteine and histidine-rich domain-containing protein 1 Rattus norvegicus | 436.58 | 35.65 | 37362 | 9 | 9 | 8176075 | 9282825 | 8180874.5 | 8925010 | 7551701 | 6095291.5 | 14304673 | 6295618.5 | 12419075 | 12940437 | 12714709 | 12742396 | 12036225 | 10196495 | 13519816 | 10 | 0 | 1.23597908 | 1.453341614 |
| P97821 | Ctsc | Dipeptidyl peptidase 1 Mus musculus | 595.33 | 16.67 | 52376 | 7 | 7 | 49376273 | 52577253 | 45175817 | 45652129 | 52748553 | 44029841 | 30403867 | 40893277 | 42000561 | 30968901 | 39618785 | 36447381 | 45088625 | 42878961 | 35434073 | 0 | 0 | 0.766897845 | 0.812396875 |
| Q8CI33 | Cwf19l1 | CWF19-like protein 1 Mus musculus | 37.67 | 2.23 | 60193 | 1 | 1 | 1 | 1 | 1 | 1 | 1 | 1 | 1 | 1 | 1 | 1 | 466385.22 | 572857.725 | 713951.5 | 632285.62 | 478808.56 | 12.5 | 0 | 1 | 572857.725 |
| Q3TGW2 | Eepd1 | Endonuclease/exonuclease/phosphatase family domain-containing protein 1 Mus musculus | 41.23 | 2.64 | 62953 | 1 | 1 | 884563.31 | 910401.94 | 1 | 646673 | 1 | 749273.62 | 660041 | 650920.725 | 865283.06 | 329085.22 | 1683334.88 | 1045418.06 | 959576.62 | 958430.62 | 1021241.69 | 11 | 0 | 1.332957886 | 2.321391069 |
| P17182 | Eno1 | Alpha-enolase Mus musculus | 2394.93 | 77.42 | 47141 | 24 | 32 | 483027841 | 544320321 | 569124609 | 442800673 | 497630657 | 598201857 | 853236929 | 634970049 | 668755073 | 674073537 | 642606273 | 680968833 | 657382657 | 604103617 | 606306369 | 0 | 0 | 1.351741063 | 1.257977291 |
| Q08509 | Eps8 | Epidermal growth factor receptor kinase substrate 8 Mus musculus | 139.73 | 5.24 | 91737 | 2 | 2 | 1 | 1 | 1 | 1 | 1 | 1 | 1 | 1 | 1 | 1 | 616116.81 | 777872.88 | 975023.06 | 673069.81 | 760520.64 | 12.5 | 0 | 1 | 760520.64 |
| Q9DC16 | Ergic1 | Endoplasmic reticulum-Golgi intermediate compartment protein 1 Mus musculus | 77.85 | 6.21 | 32562 | 2 | 2 | 3226603 | 3379903.5 | 3210803.25 | 3267271.5 | 3166848 | 3181764 | 3601628.5 | 3769343 | 3422206.5 | 3596184 | 2691115 | 3083740.25 | 3087197 | 3004658.25 | 2982803.5 | 4 | 0 | 1.081204965 | 0.91373588 |
| Q8R180 | Ero1a | ERO1-like protein alpha Mus musculus | 1208.87 | 53.66 | 54084 | 17 | 19 | 34275897 | 37585565 | 33318769 | 30588371 | 32604637 | 46392561 | 59067661 | 50951521 | 56834589 | 49306309 | 41812017 | 39553205 | 38353937 | 44429225 | 42118393 | 0 | 0 | 1.559348995 | 1.225056774 |
| A0A0B4 | Fcgr4 | Low affinity immunoglobulin gamma Fc region receptor IV Mus musculus | 45.23 | 4.02 | 28398 | 1 | 1 | 2495273 | 2113805.5 | 2117607.5 | 2473083 | 2382982.25 | 1806752 | 1644024.25 | 1732195.12 | 1702798.62 | 2170689.5 | 1926141.75 | 1631899.75 | 2058816.12 | 1699792 | 1765408.12 | 2 | 0 | 0.781891909 | 0.784101941 |
| D3ZYR1 | Fcho2 | F-BAR domain only protein 2 Rattus norvegicus | 115.95 | 3.83 | 88712 | 2 | 2 | 830894.94 | 883108.62 | 884735.75 | 744802.12 | 843027.44 | 1072914.5 | 966837.38 | 1172649.12 | 1471196.38 | 685448 | 970138.62 | 947084.31 | 987039.62 | 1045488.12 | 991401.75 | 5 | 0 | 1.282445255 | 1.180239135 |
| P45878 | Fkbp2 | Peptidyl-prolyl cis-trans isomerase FKBP2 Mus musculus | 125.57 | 19.29 | 15344 | 2 | 2 | 3244866 | 3025755.5 | 3779809.5 | 4243410.5 | 2996151 | 2733578.75 | 2386004.5 | 2459459.25 | 2125663 | 2787699.75 | 2336236.5 | 2201685.5 | 2678028.75 | 2236693.25 | 1730822.25 | 0 | 0 | 0.722522306 | 0.646817299 |
| Q9R0N0 | Galk1 | Galactokinase Mus musculus | 453.75 | 37.5 | 42295 | 9 | 10 | 8279408 | 12743603 | 18287175 | 10402457 | 10066241 | 20618689 | 16862377 | 18866037 | 20161931 | 22500399 | 19804265 | 19574217 | 20998037 | 19180293 | 18496455 | 1 | 0 | 1.65626098 | 1.640265934 |
| Q8BVW0 | Ganc | Neutral alpha-glucosidase C Mus musculus | 41.45 | 1.45 | 102008 | 1 | 1 | 494299.16 | 529243.2125 | 528985.5 | 519233.81 | 574454.38 | 473868.41 | 328260.09 | 540940.88 | 461901.97 | 504538.5 | 469166.12 | 312593.34 | 384612.78 | 439569.41 | 346164.09 | 5 | 0 | 0.872759365 | 0.737697034 |
| Q9D6Y9 | Gbe1 | 1,4-alpha-glucan-branching enzyme Mus musculus | 38.75 | 1.85 | 80364 | 1 | 1 | 1 | 1 | 1 | 1 | 1 | 237746.94 | 1304682.75 | 1 | 1 | 1 | 746103 | 790873.69 | 1263161.75 | 905875.265 | 823362.62 | 7.5 | 0 | 308486.538 | 905875.265 |
| Q9D1H7 | Get4 | Golgi to ER traffic protein 4 homolog Mus musculus | 133.38 | 12.23 | 36525 | 3 | 3 | 2727767 | 2513847.5 | 2044945.5 | 2748242.75 | 2697812.5 | 1962030.25 | 2441285 | 2295243 | 1941175.62 | 1515364 | 1950727.5 | 869177.31 | 1803638.38 | 1665114.12 | 798530.88 | 2 | 0 | 0.797565753 | 0.55661685 |
| Q8BTZ7 | Gmppb | Mannose-1-phosphate guanyltransferase beta Mus musculus | 49.39 | 5 | 39917 | 1 | 1 | 4203369.5 | 4833290.5 | 4242727.5 | 4922349.5 | 5575047 | 3862472.25 | 3928683.5 | 3214383.25 | 2715428.75 | 3238080.25 | 1932733.88 | 2753327.47 | 3486299.25 | 2734631.75 | 2859645 | 0 | 0 | 0.713260801 | 0.578994928 |
| P63213 | Gng2 | Guanine nucleotide-binding protein G(I)/G(S)/G(O) subunit gamma-2 Mus musculus | 83.93 | 39.44 | 7850 | 2 | 2 | 6892465 | 7611842 | 7428047 | 8924962 | 8298338.5 | 6617675 | 2847632.25 | 6250014.5 | 6781411.5 | 6216025 | 2560274 | 5558617 | 6667112 | 6446683 | 5136033 | 0 | 0 | 0.733297875 | 0.673433233 |
| P06745 | Gpi | Glucose-6-phosphate isomerase Mus musculus | 1639.38 | 58.6 | 62767 | 14 | 27 | 122799385 | 132587617 | 124140889 | 107792553 | 137788401 | 138885697 | 170301729 | 129185553 | 126804361 | 144078337 | 158679569 | 153559217 | 155513953 | 143906881 | 177933345 | 4 | 0 | 1.134611488 | 1.263128768 |
| Q810D6 | Grwd1 | Glutamate-rich WD repeat-containing protein 1 Mus musculus | 220.73 | 12.56 | 49224 | 4 | 4 | 4084838.5 | 3332763 | 4120762 | 4366309 | 3994580.5 | 3127738.5 | 3298426 | 3635051.75 | 3549815 | 4126948.75 | 3315716.75 | 2887585.75 | 2307649.25 | 2685089.5 | 3251797 | 6 | 0 | 0.89138924 | 0.726049277 |
| O08730 | Gyg1 | Glycogenin-1 Rattus norvegicus | 45.97 | 4.8 | 37378 | 1 | 1 | 3535684.75 | 3298503.5 | 3479268.125 | 3422287 | 3660597.25 | 1947706.38 | 1722916 | 2134305.25 | 1502266 | 1826798.408 | 2593475.25 | 1 | 1821876.38 | 2305709.25 | 1 | 0 | 0 | 0.525052494 | 0.386349234 |
| P14426 | H2-D1 | H-2 class I histocompatibility antigen, D-K alpha chain Mus musculus | 228.67 | 9.67 | 40620 | 0 | 4 | 76935561 | 88768785 | 78007345 | 74458721 | 84181505 | 86945089 | 77843313 | 85997273 | 84113057 | 76259217 | 71099593 | 57321649 | 74426801 | 73839521 | 57002289 | 11 | 0 | 1.021886393 | 0.829348237 |
| Q8VH49 | Higd1a | HIG1 domain family member 1A, mitochondrial Rattus norvegicus | 147.95 | 43.01 | 10301 | 2 | 2 | 813655.38 | 698692.56 | 782062.25 | 630587.57 | 227940.09 | 454788.72 | 1423300.12 | 315909.84 | 852475.88 | 1226544.25 | 1112542.75 | 941054.12 | 982619.75 | 923053 | 1025185.62 | 8 | 0 | 1.355249933 | 1.580892322 |
| P08113 | Hsp90b1 | Endoplasmin Mus musculus | 3099.08 | 61.97 | 92476 | 44 | 46 | 438865441 | 435713089 | 428330785 | 391069985 | 457535489 | 524561441 | 436578017 | 501669633 | 531080865 | 431629921 | 476389889 | 476054881 | 458124705 | 469964161 | 461589377 | 5 | 0 | 1.127354499 | 1.08859257 |
| P41565 | Idh3g | Isocitrate dehydrogenase [NAD] subunit gamma 1, mitochondrial Rattus norvegicus | 253.8 | 14.76 | 42851 | 6 | 6 | 16545307 | 13982288 | 16847583 | 16369147 | 13910443 | 19518973 | 17887521 | 16911499 | 16665583 | 16975001 | 21632737 | 17221929 | 17784893 | 16977761 | 18943285 | 1 | 0 | 1.132687397 | 1.191950055 |
| Q9R002 | Ifi202 | Interferon-activable protein 202 Mus musculus | 144.04 | 6.29 | 50466 | 2 | 2 | 2326020.25 | 2754549.75 | 2657673.25 | 1744767.5 | 1884176.88 | 1 | 1 | 2016734.5 | 2534400.25 | 1522665.25 | 1 | 1 | 1537492.38 | 1744321 | 1 | 5 | 0 | 0.534327593 | 0.288709616 |
| Q0GNC1 | Inf2 | Inverted formin-2 Mus musculus | 432.48 | 8.25 | 138560 | 8 | 8 | 1647709.12 | 1456264.75 | 1339991.88 | 1237981.5 | 1380480 | 1829130.75 | 1885823 | 2223330.5 | 2042381 | 1897470.62 | 1680831.38 | 2652184.5 | 2136495 | 2325219.75 | 2560672.25 | 0 | 0 | 1.398688513 | 1.607861218 |
| P97825 | Jpt1 | Jupiter microtubule associated homolog 1 Mus musculus | 106.39 | 27.27 | 16081 | 2 | 2 | 2924789.25 | 4087840.5 | 3059245.5 | 4315181 | 3478096 | 2223103.5 | 2421793.5 | 2452645.5 | 2560232 | 3651883.75 | 887516.75 | 2085826.38 | 2894699.25 | 2233320.75 | 816898.06 | 3 | 0 | 0.745006707 | 0.499198723 |
| P62482 | Kcnab2 | Voltage-gated potassium channel subunit beta-2 Mus musculus | 379.23 | 35.97 | 41021 | 7 | 7 | 11593159 | 12929697 | 12474758 | 13290734 | 11773923 | 11195759 | 12852774 | 12725711 | 12060590 | 11924525 | 11029746 | 10607653 | 11235901 | 11154022 | 11206591 | 10 | 0 | 0.979006376 | 0.889975699 |
| Q6P9P6 | Kif11 | Kinesin-like protein KIF11 Mus musculus | 575.89 | 14.16 | 118027 | 9 | 9 | 1801478.25 | 1844002.5 | 1682134.75 | 1739286 | 1683549 | 2451006.5 | 2595642.75 | 2073151 | 1864301.5 | 2058784.5 | 1944497.75 | 2636322.5 | 2014056 | 2490137.25 | 2025372.5 | 0 | 0 | 1.261979169 | 1.269693029 |
| Q6P9L6 | Kif15 | Kinesin-like protein KIF15 Mus musculus | 215.27 | 3.53 | 160120 | 3 | 3 | 613811.88 | 450786.91 | 756072.19 | 419258.38 | 885964.5 | 1138113.62 | 695290.75 | 665664.12 | 714826.44 | 1127235.12 | 1048778.88 | 1160518.25 | 932265.19 | 1201514.38 | 1151807.75 | 6 | 0 | 1.388764381 | 1.757860214 |
| Q3V300 | Kif22 | Kinesin-like protein KIF22 Mus musculus | 78.78 | 3.94 | 73190 | 2 | 2 | 1265744.75 | 1064443.25 | 957997.25 | 1238799.25 | 931019.81 | 794203.44 | 776615.88 | 968571.38 | 919149.44 | 734712 | 256059.19 | 425881.59 | 417362.6175 | 313512.31 | 673997.38 | 2 | 0 | 0.768275711 | 0.382339949 |
| Q8C3Y4 | Kntc1 | Kinetochore-associated protein 1 Mus musculus | 106.39 | 1.63 | 250358 | 3 | 3 | 1642317.75 | 1803738.75 | 1552117.5 | 1696947.75 | 1526644.5 | 1469792.25 | 1137607 | 1726752.75 | 1686255.75 | 1766130.75 | 1373876.75 | 1253331.25 | 1363236.25 | 1336416.62 | 1132174.62 | 11 | 0 | 0.94706396 | 0.785601937 |
| Q56R16 | Kpna5 | Importin subunit alpha-6 Rattus norvegicus | 233.82 | 9.89 | 60299 | 1 | 5 | 4009986.5 | 4253468.5 | 4582332 | 4451326 | 2915368.25 | 3943884 | 9248177 | 4270956 | 4658801.5 | 4276679.5 | 4870079.5 | 8669782 | 4918613 | 5006213.5 | 4926133 | 8 | 0 | 1.30604935 | 1.404618297 |
| Q05CL8 | Larp7 | La-related protein 7 Mus musculus | 297.48 | 11.4 | 64802 | 5 | 5 | 10513111 | 11670235 | 7440189.5 | 11271201 | 11376124 | 9821228 | 5042094.5 | 6675311 | 8297351 | 5860961 | 6760591.5 | 5860076 | 7100140 | 7035883 | 6525869 | 2 | 0 | 0.682922476 | 0.636732573 |
| P0C5H9 | Manf | Mesencephalic astrocyte-derived neurotrophic factor Rattus norvegicus | 263.78 | 21.23 | 20388 | 5 | 5 | 13466765 | 12135204 | 10767937 | 10744876 | 10496057 | 9700052 | 6837870 | 9948804 | 10329289 | 8493842 | 8759529 | 8539654 | 8978007 | 9614908 | 6948340 | 0 | 0 | 0.786481464 | 0.743617672 |
| Q9ESL4 | Map3k20 | Mitogen-activated protein kinase kinase kinase 20 Mus musculus | 282.33 | 7.48 | 91720 | 5 | 5 | 9457455 | 9591159 | 8883040 | 8947703 | 9302907 | 10555577 | 10188621 | 10224819 | 9568499 | 9639521 | 8029131.5 | 8647950 | 8369205.5 | 8882327 | 8455357 | 1 | 0 | 1.086500155 | 0.917754292 |
| E9PVX6 | Mki67 | Proliferation marker protein Ki-67 Mus musculus | 3338.53 | 27.7 | 350864 | 60 | 60 | 194457617 | 64824269 | 67822953 | 70583001 | 81777945 | 58333197 | 84126649 | 55133081 | 42261441 | 71884745 | 35683229 | 43975813 | 57443013 | 36264141 | 59601305 | 7 | 0 | 0.650180102 | 0.48588973 |
| P0DOV1 | Mnda | Interferon-activable protein 205-B Mus musculus | 59.12 | 2.82 | 46977 | 1 | 1 | 1493655.88 | 1912190.38 | 1427573.62 | 1756698.25 | 1660475.75 | 1210737.75 | 1551060.12 | 1608413.12 | 1409332.5 | 1121738.5 | 856138.88 | 1074790.88 | 1283166.38 | 1 | 1 | 4 | 0 | 0.836458816 | 0.389559611 |
| P58059 | Mrps21 | 28S ribosomal protein S21, mitochondrial Mus musculus | 91.71 | 14.94 | 10561 | 1 | 1 | 1 | 1 | 1 | 1 | 1 | 1 | 6103944.5 | 1 | 1 | 1 | 4720752 | 4724733.5 | 4083026.89 | 482241.06 | 6404381 | 10 | 0 | 1220789.7 | 4083026.89 |
| P62774 | Mtpn | Myotrophin Mus musculus | 212.54 | 43.22 | 12861 | 3 | 3 | 12414931 | 9254644 | 8558970 | 11747155 | 10568561 | 16165451 | 18051709 | 13240505 | 10138163 | 17026961 | 14373694 | 14042488 | 13171149 | 13982410 | 14974792 | 3 | 0 | 1.420189143 | 1.342573512 |
| O35682 | Myadm | Myeloid-associated differentiation marker Mus musculus | 142.12 | 16.25 | 35285 | 3 | 3 | 1 | 400232.06 | 1 | 628709.5 | 532174.56 | 1 | 978064.12 | 921647.62 | 1 | 988146.12 | 661894.69 | 863061.44 | 631438.06 | 820105.0475 | 1124026 | 8 | 0 | 1.849866338 | 2.626659178 |
| Q62433 | Ndrg1 | Protein NDRG1 Mus musculus | 402.01 | 32.74 | 43009 | 7 | 7 | 12050821 | 17884593 | 12918770 | 12185809 | 17734777 | 14966141 | 24916441 | 18337219 | 15743440 | 19967781 | 20175771 | 18640775 | 19921097 | 20909529 | 20497007 | 4 | 0 | 1.290708607 | 1.376083758 |
| Q9CQ54 | Ndufc2 | NADH dehydrogenase [ubiquinone] 1 subunit C2 Mus musculus | 41.04 | 10.83 | 14164 | 1 | 1 | 1530846.62 | 1188782.38 | 1155049.5 | 983285.12 | 1004454.69 | 1613328.25 | 665314.38 | 1182471.12 | 1202890.62 | 860145.12 | 945709.75 | 969569.81 | 917689.19 | 975707.5 | 838160.94 | 12 | 0 | 0.942298758 | 0.792648519 |
| P97526 | Nf1 | Neurofibromin Rattus norvegicus | 65.62 | 0.78 | 317083 | 1 | 1 | 279746.625 | 297530.44 | 277005.28 | 275340.78 | 269110 | 187287.83 | 225938.3875 | 238583.58 | 238894.75 | 238987.39 | 242400.69 | 227861.19 | 1 | 1 | 222089.89 | 0 | 0 | 0.807653667 | 0.494986326 |
| O55126 | Nipsnap2 | Protein NipSnap homolog 2 Mus musculus | 94.84 | 9.96 | 32933 | 2 | 2 | 524071.69 | 492649.66 | 356549.66 | 453230.065 | 439649.25 | 589478.25 | 955871.38 | 431857.34 | 323601.44 | 860322.5 | 655657 | 759500.25 | 597890.62 | 740721.0925 | 949836.5 | 9 | 0 | 1.394934341 | 1.63431588 |
| Q6PIP5 | Nudcd1 | NudC domain-containing protein 1 Mus musculus | 391 | 24.4 | 66705 | 9 | 9 | 8369140.5 | 8550534 | 7894536 | 8618849 | 8807447 | 9861959 | 7856768 | 9653839 | 11885193 | 9191013 | 10208554 | 9337844 | 9079144 | 10043479 | 8939247 | 5 | 0 | 1.14697422 | 1.127076163 |
| P56380 | Nudt2 | Bis(5'-nucleosyl)-tetraphosphatase [asymmetrical] Mus musculus | 172.09 | 27.89 | 16989 | 3 | 3 | 1725483.5 | 1933229.5 | 3011968 | 2932848.5 | 2588678.25 | 2818607.75 | 1849077.25 | 3474031 | 4175410.75 | 776465.12 | 592150.38 | 1465661.62 | 946737.12 | 1576744.75 | 1336226 | 11 | 0 | 1.073931165 | 0.485352611 |
| Q9DBY8 | Nvl | Nuclear valosin-containing protein-like Mus musculus | 211.68 | 7.25 | 94476 | 4 | 4 | 5006893 | 3699745.25 | 4004343 | 4415970 | 4322784.5 | 2172377.5 | 2996748.25 | 2547878.25 | 2998211.375 | 4275841.5 | 2058915.62 | 1517011.75 | 2354571.5 | 1917346.75 | 2530980 | 2 | 0 | 0.69889238 | 0.483867295 |
| Q9D0K2 | Oxct1 | Succinyl-CoA:3-ketoacid coenzyme A transferase 1, mitochondrial Mus musculus | 1174.91 | 43.65 | 55989 | 15 | 15 | 38869481 | 50020709 | 50447477 | 51833617 | 52585401 | 52766841 | 68385017 | 55093925 | 47794729 | 55525261 | 56438569 | 59848025 | 58012089 | 53772453 | 59956445 | 4 | 0 | 1.14690505 | 1.181619208 |
| Q9WUA3 | Pfkp | ATP-dependent 6-phosphofructokinase, platelet type Mus musculus | 1812.23 | 54.97 | 85455 | 20 | 27 | 62324449 | 64117569 | 62208097 | 54761193 | 61724577 | 73752281 | 87211129 | 74364329 | 80531273 | 73031857 | 75789449 | 65038041 | 67673449 | 74192025 | 70099769 | 0 | 0 | 1.274484215 | 1.156182378 |
| Q8BHF7 | Pgs1 | CDP-diacylglycerol--glycerol-3-phosphate 3-phosphatidyltransferase, mitochondrial Mus musculus | 83.58 | 5.61 | 62489 | 2 | 2 | 679683.38 | 826181.75 | 691918.06 | 720012.625 | 682267.31 | 619187.5925 | 537546.5 | 656831.31 | 806267.56 | 476105 | 481501.72 | 1 | 619129.12 | 1 | 447023.5 | 4 | 0 | 0.859967688 | 0.429897001 |
| O35405 | Pld3 | Phospholipase D3 Mus musculus | 139.88 | 11.27 | 54389 | 3 | 3 | 9169067 | 7480470.5 | 6721049 | 9324024 | 8638230 | 5449954.5 | 5937753.5 | 6070406.5 | 4910840 | 6251654 | 6166208 | 3916503.25 | 5830383 | 5306693 | 4008033 | 0 | 0 | 0.692442333 | 0.610357767 |
| Q91YU8 | Ppan | Suppressor of SWI4 1 homolog Mus musculus | 194.17 | 10 | 52756 | 4 | 4 | 1447403.38 | 906248.94 | 1049341.12 | 1373492.62 | 1488146.75 | 1493199.12 | 839895.06 | 2719543.5 | 1724863.25 | 2123574 | 2719189.75 | 2263799.75 | 1928529.25 | 1750653.25 | 1591125.62 | 5 | 0 | 1.420845435 | 1.636695706 |
| P97814 | Pstpip1 | Proline-serine-threonine phosphatase-interacting protein 1 Mus musculus | 282.97 | 17.11 | 47590 | 5 | 5 | 9298495 | 9274206 | 8585177 | 9588877 | 8833164 | 7979274.5 | 6909640 | 7803951 | 8235725 | 7891581 | 8050923.5 | 6541285 | 8315439 | 8028290 | 6663570 | 0 | 0 | 0.851694614 | 0.824913873 |
| F1M386 | Rapgef2 | Rap guanine nucleotide exchange factor 2 Rattus norvegicus | 41.99 | 0.87 | 166388 | 1 | 1 | 480017.22 | 612132.38 | 555573.75 | 308647.25 | 472058.34 | 605064.5 | 475549.34 | 768776.88 | 704809.19 | 406267.56 | 988363.81 | 693867.12 | 770892.1225 | 672832.25 | 728505.31 | 8 | 0 | 1.219087543 | 1.587223966 |
| O70521 | Rgs19 | Regulator of G-protein signaling 19 Rattus norvegicus | 39.26 | 6.02 | 24738 | 1 | 1 | 1425710 | 1221182 | 1282710.75 | 1493605.38 | 1355802.033 | 375430.56 | 1058403.12 | 1323065.62 | 1498147.88 | 1294972.88 | 1168665.38 | 1102665.5 | 500619.91 | 1107105.12 | 716740.19 | 9 | 0 | 0.818706556 | 0.677945008 |
| Q6PR54 | Rif1 | Telomere-associated protein RIF1 Mus musculus | 189.22 | 2.32 | 266228 | 3 | 3 | 1624920.38 | 2046007.75 | 1606687.25 | 2072905.25 | 1487789.88 | 1001058 | 1176537.25 | 1620655 | 144852.36 | 1636714.5 | 1451509.5 | 1388940 | 1297931.25 | 1229656.5 | 1019543 | 5 | 0 | 0.631321688 | 0.722715076 |
| P17074 | Rps19 | 40S ribosomal protein S19 Rattus norvegicus | 377.39 | 35.17 | 16085 | 7 | 7 | 25781753 | 23033861 | 20522993 | 22798165 | 21703593 | 13058609 | 22001325 | 19951247 | 18184585 | 23158705 | 16848813 | 18612305 | 20105417 | 20242365 | 17059563 | 6 | 0 | 0.846399878 | 0.81577798 |
| P97461 | Rps5 | 40S ribosomal protein S5 Mus musculus | 837 | 55.88 | 22889 | 13 | 13 | 65677265 | 68578081 | 71479009 | 69414161 | 76406825 | 68157289 | 75840953 | 60549517 | 58962681 | 73934241 | 63476377 | 63132409 | 62625005 | 63256313 | 58274537 | 9 | 0 | 0.959862194 | 0.883970757 |
| Q9JJU8 | Sh3bgrl | SH3 domain-binding glutamic acid-rich-like protein Mus musculus | 383.94 | 56.14 | 12811 | 7 | 7 | 55023249 | 54861409 | 54446401 | 59335037 | 60166041 | 50005341 | 43702705 | 49228841 | 49986949 | 45807265 | 53488293 | 50770393 | 51868257 | 51506133 | 43117437 | 0 | 0 | 0.841099614 | 0.883446518 |
| O55242 | Sigmar1 | Sigma non-opioid intracellular receptor 1 Mus musculus | 126.45 | 13.9 | 25250 | 2 | 2 | 1644043 | 888876.69 | 952398.69 | 788086.75 | 1070364 | 662997.69 | 747085.19 | 912535.81 | 732123.62 | 778910.88 | 673738.19 | 441712.53 | 522908.47 | 675793.81 | 635142.12 | 2 | 0 | 0.717406216 | 0.5519129 |
| Q9QXX4 | Slc25a13 | Calcium-binding mitochondrial carrier protein Aralar2 Mus musculus | 1659.72 | 63.61 | 74467 | 26 | 29 | 62194497 | 61106117 | 65557537 | 57746685 | 64471141 | 81916873 | 79193121 | 74813577 | 69371073 | 76902177 | 73078401 | 71403545 | 77964905 | 74740009 | 72802361 | 0 | 0 | 1.228628532 | 1.189385386 |
| P16036 | Slc25a3 | Phosphate carrier protein, mitochondrial Rattus norvegicus | 713.67 | 44.66 | 39445 | 12 | 12 | 36062841 | 41107849 | 41221105 | 34584221 | 38753701 | 44266377 | 50259629 | 50509249 | 58528813 | 47451661 | 47089453 | 42631413 | 42631385 | 42273857 | 59066741 | 0 | 0 | 1.309216604 | 1.218866082 |
| B4F769 | Smarcal1 | | 37.68 | 0.99 | 101212 | 1 | 1 | 401547.94 | 351870.12 | 358495.8575 | 324722.06 | 355843.31 | 330827.47 | 556132.88 | 528519.19 | 296367.19 | 373460.25 | 312523 | 1 | 1 | 315119.19 | 306350.97 | 10 | 0 | 1.163364617 | 0.521063293 |
| Q8VIM6 | Strc | Stereocilin Mus musculus | 35.29 | 0.44 | 196347 | 1 | 1 | 722819.81 | 2210075.5 | 1922223.88 | 1858499.88 | 719493.31 | 2165312.75 | 4443522 | 2196245.5 | 2423662.75 | 3768064.25 | 3559237.75 | 2465075.75 | 3501635.25 | 3825082.5 | 3071432.75 | 2 | 0 | 2.017567673 | 2.209365762 |
| Q9WUD1 | Stub1 | STIP1 homology and U box-containing protein 1 Mus musculus | 57.9 | 3.95 | 34909 | 1 | 1 | 3146873.188 | 3086832.25 | 2984497.5 | 3650552 | 2865611 | 2347790.25 | 2517792.875 | 2580080.75 | 2306820.25 | 2836480.25 | 2477411 | 2749622.5 | 2833288.75 | 2666649.75 | 2117652.5 | 0 | 0 | 0.800093529 | 0.816342047 |
| Q5DQR4 | Stxbp5l | Syntaxin-binding protein 5-like Mus musculus | 36.53 | 0.93 | 131844 | 1 | 1 | 5538802.5 | 6822379.5 | 4978563.5 | 5871094 | 7652958.5 | 5255096 | 14494090 | 5008501.5 | 6862474 | 11678574 | 10519407 | 8299066.5 | 10844078 | 8508189 | 10373591 | 9 | 0 | 1.40289719 | 1.572856701 |
| Q9Z2I9 | Sucla2 | Succinate--CoA ligase [ADP-forming] subunit beta, mitochondrial Mus musculus | 456.06 | 29.37 | 50114 | 9 | 9 | 8295629.5 | 10424399 | 10797735 | 8882850 | 9556912 | 10075774 | 12508839 | 13778828 | 11953550 | 9170292 | 11457529 | 12496327 | 11051370 | 12591289 | 11729394 | 5 | 0 | 1.198712452 | 1.237051086 |
| Q6A028 | Swap70 | Switch-associated protein 70 Mus musculus | 826.42 | 35.21 | 68996 | 18 | 18 | 8438880 | 4376241.5 | 9912001 | 6593201 | 9192176 | 9780575 | 9011705 | 8434352 | 10127617 | 11306538 | 11334125 | 10442167 | 13070007 | 11563968 | 16333501 | 6 | 0 | 1.263506333 | 1.629179327 |
| Q0VGY8 | Tanc1 | Protein TANC1 Mus musculus | 276.4 | 5.87 | 200805 | 6 | 6 | 1665739.12 | 1572150.38 | 1584758.25 | 1529621.75 | 1553410 | 2079376.75 | 2248288.5 | 2210395.5 | 1783772.38 | 2533140.5 | 1947109.12 | 2153689 | 2294152 | 2127409.5 | 2663115.25 | 0 | 0 | 1.373060169 | 1.414865714 |
| Q91YX0 | Themis2 | Protein THEMIS2 Mus musculus | 451.31 | 22.32 | 74378 | 9 | 9 | 6519084.5 | 7443011.5 | 6576726.5 | 7534454 | 7404014.5 | 5142677 | 3751690 | 4517776.5 | 4762057.5 | 3096810 | 5101699 | 4559930.5 | 3806501.5 | 3769314.25 | 4837313 | 0 | 0 | 0.599566946 | 0.622222205 |
| P62075 | Timm13 | Mitochondrial import inner membrane translocase subunit Tim13 Mus musculus | 36.2 | 14.74 | 10458 | 1 | 1 | 2119784.5 | 2271527.25 | 2447546.5 | 2781436.25 | 2072312.75 | 1758826 | 1823802.25 | 1746608 | 1605808 | 2113046.25 | 1606487.5 | 1664509.88 | 2017443.62 | 1194384.88 | 1110859.12 | 1 | 0 | 0.77383002 | 0.649443263 |
| Q9D938 | Tmem160 | Transmembrane protein 160 Mus musculus | 45.37 | 3.72 | 19587 | 1 | 1 | 1820692.62 | 2003869.12 | 1692980 | 1684022.88 | 1704514.62 | 1630150.12 | 1566000.62 | 1492078.75 | 1943340.75 | 1780485.38 | 1517364.38 | 1316195.25 | 1553558.38 | 1669387.75 | 1311380.88 | 7 | 0 | 0.944529618 | 0.827287344 |
| A1L1L2 | Tmem214 | Transmembrane protein 214 Rattus norvegicus | 58.63 | 1.9 | 76566 | 1 | 1 | 145575.83 | 1 | 166379.84 | 1 | 1 | 167607.42 | 1 | 154167.31 | 270845.69 | 1 | 171508.94 | 187978.91 | 187508.0875 | 204431 | 186113.5 | 8 | 0 | 1.89968248 | 3.00533541 |
| Q5SSZ5 | Tns3 | Tensin-3 Mus musculus | 44.72 | 0.97 | 155589 | 1 | 1 | 596497.81 | 404810.47 | 723231.25 | 484870.19 | 789873 | 1 | 1016857.31 | 1 | 742882 | 833255.5 | 1 | 387263.41 | 1 | 1 | 1 | 11 | 0 | 0.864538976 | 0.129120008 |
| P23591 | Tsta3 | GDP-L-fucose synthase Mus musculus | 230.02 | 20.25 | 35878 | 5 | 5 | 7730363.5 | 8365780 | 7979294 | 8123320 | 8205173.5 | 8520995 | 8138888 | 8981063 | 6627575.5 | 7113156 | 6084183 | 6867531 | 7541159 | 7180785.5 | 7097822 | 12 | 0 | 0.974699157 | 0.860596473 |
| Q5NC05 | Ttf2 | Transcription termination factor 2 Mus musculus | 100.85 | 2.9 | 125530 | 3 | 3 | 1898142.75 | 1383248 | 1174040.62 | 1850015.38 | 1632152.38 | 1 | 1081068.25 | 1087629.25 | 259628.09 | 975710 | 1107200.25 | 1 | 695842.94 | 1 | 644047.88 | 0 | 0 | 0.428849648 | 0.30829134 |
| Q6PAM1 | Txlna | Alpha-taxilin Mus musculus | 130 | 6.14 | 62369 | 2 | 2 | 1280284.75 | 1384573.938 | 1239255.75 | 1673974.75 | 1344780.5 | 939730.81 | 456411.59 | 1017399.88 | 1012583 | 1036303.62 | 1006096.12 | 1032015.56 | 1 | 1209596.62 | 1 | 0 | 0 | 0.644592359 | 0.469127753 |
| Q9Z1Q9 | Vars | Valine--tRNA ligase Mus musculus | 2525.76 | 51.46 | 140215 | 44 | 44 | 99511017 | 107113025 | 104057321 | 105556265 | 107404969 | 120400185 | 128984305 | 107190897 | 120652129 | 123631953 | 115033665 | 113963129 | 107953753 | 112612465 | 119659857 | 1 | 0 | 1.147461021 | 1.087044622 |
| P20152 | Vim | Vimentin Mus musculus | 3269.56 | 86.48 | 53688 | 43 | 51 | 2434394369 | 2796715521 | 2589878017 | 2831042049 | 2816222721 | 2746197761 | 2924476929 | 2644526337 | 2887560705 | 2946960385 | 3032498689 | 2863035393 | 3148944641 | 2916097025 | 3008812801 | 6 | 0 | 1.050598207 | 1.111457359 |
| P29788 | Vtn | Vitronectin Mus musculus | 80.38 | 3.77 | 54849 | 1 | 1 | 690520.81 | 656204.25 | 617229.56 | 475170.97 | 465189.84 | 793610.56 | 1092978.12 | 653107 | 649661.81 | 753121.88 | 370059 | 305932.31 | 336121.8825 | 430688.81 | 237807.41 | 4 | 0 | 1.357455643 | 0.578659396 |
| P0C1X8 | Aak1 | AP2-associated protein kinase 1 Rattus norvegicus | 96.52 | 4.47 | 103761 | 2 | 2 | 410858.03 | 1 | 323195.53 | 1 | 310884.16 | 375156.69 | 411203.53 | 375989.06 | 455028.94 | 404344.555 | 328897.81 | 488275.5 | 481075.16 | 428982.665 | 417682.19 | 3 | 1 | 1.934774549 | 2.052667043 |
| P11497 | Acaca | Acetyl-CoA carboxylase 1 Rattus norvegicus | 978.75 | 11.81 | 265194 | 17 | 22 | 16053941 | 14432279 | 18560953 | 15344241 | 16481721 | 18212811 | 20150013 | 14670331 | 24413653 | 18222867 | 21948717 | 23793103 | 19070543 | 17036307 | 23465953 | 6 | 1 | 1.182959891 | 1.302220113 |
| P45952 | Acadm | Medium-chain specific acyl-CoA dehydrogenase, mitochondrial Mus musculus | 444.97 | 28.5 | 46481 | 9 | 9 | 42677457 | 50715201 | 46500937 | 45997517 | 47525865 | 43511933 | 43043197 | 45209525 | 44488141 | 43509069 | 41080921 | 40712041 | 44671013 | 41320329 | 36902633 | 5 | 1 | 0.941499062 | 0.87691538 |
| Q8CG45 | Akr7a2 | Aflatoxin B1 aldehyde reductase member 2 Rattus norvegicus | 61.95 | 4.09 | 40675 | 1 | 1 | 1 | 458293.44 | 694835.62 | 1 | 777704.19 | 702958.56 | 999241.5 | 774745.94 | 384596.72 | 859606.69 | 759441.56 | 911180.3275 | 857205.94 | 840697.56 | 1187376.25 | 5 | 1 | 1.927222641 | 2.35954965 |
| Q66HF8 | Aldh1b1 | Aldehyde dehydrogenase X, mitochondrial Rattus norvegicus | 111.29 | 4.82 | 57625 | 1 | 2 | 11603215 | 11684018 | 13051814 | 12502605 | 11462587 | 13491576 | 12992340 | 14037616 | 14561375 | 12347817 | 13110488 | 12577315 | 13281077 | 13556662 | 19708481 | 3 | 1 | 1.118175523 | 1.197826624 |
| O35643 | Ap1b1 | AP-1 complex subunit beta-1 Mus musculus | 2073.93 | 49.63 | 103935 | 20 | 34 | 43716193 | 42529389 | 50980005 | 37876845 | 44609745 | 55121121 | 50339689 | 52601265 | 61493421 | 48157125 | 57003153 | 63977649 | 48798821 | 53961233 | 61299533 | 2 | 1 | 1.218469657 | 1.297335418 |
| P62944 | Ap2b1 | AP-2 complex subunit beta Rattus norvegicus | 1888.98 | 48.56 | 104553 | 21 | 35 | 62308241 | 63961745 | 70913601 | 56382481 | 64318025 | 75426113 | 65693937 | 72050961 | 84853321 | 64823637 | 76695273 | 76316321 | 66542965 | 71602969 | 70995681 | 2 | 1 | 1.141447392 | 1.139261816 |
| Q80V94 | Ap4e1 | AP-4 complex subunit epsilon-1 Mus musculus | 78.97 | 1.78 | 124845 | 1 | 1 | 681580.38 | 454765.38 | 548853.5175 | 674298.69 | 384769.62 | 1 | 222189.72 | 450039.84 | 326010.31 | 1 | 1 | 322243.22 | 1 | 405152.5 | 315498.97 | 1 | 1 | 0.363755297 | 0.380027332 |
| Q80X80 | C2cd2l | Phospholipid transfer protein C2CD2L Mus musculus | 49.59 | 1.42 | 76329 | 1 | 1 | 1 | 1 | 349341.56 | 1 | 1 | 567739.62 | 1 | 1 | 645406.5 | 486975.19 | 290710.75 | 411528.545 | 483127.28 | 428528.12 | 443748.03 | 6 | 1 | 4.86659487 | 5.889992491 |
| P24860 | Ccnb1 | G2/mitotic-specific cyclin-B1 Mus musculus | 75.11 | 4.88 | 48052 | 1 | 2 | 1 | 1 | 210056.78 | 1 | 1 | 305435.56 | 494806.69 | 343590.62 | 217477.61 | 356642.62 | 147209.69 | 347259.97 | 278458.2025 | 229709.59 | 389653.56 | 0 | 1 | 8.178361996 | 6.628038859 |
| Q9CWK3 | Cd2bp2 | CD2 antigen cytoplasmic tail-binding protein 2 Mus musculus | 42.89 | 4.09 | 37694 | 1 | 1 | 924238.69 | 937526.25 | 700660.44 | 880414.75 | 959233.62 | 897650.69 | 761269.38 | 659756.19 | 753454.1125 | 695140.19 | 753146.94 | 645520.31 | 669303.595 | 654311.94 | 624235.19 | 4 | 1 | 0.855794513 | 0.760213973 |
| Q921W4 | Cryzl1 | Quinone oxidoreductase-like protein 1 Mus musculus | 83.52 | 10.06 | 38725 | 2 | 2 | 1 | 1 | 1485640.38 | 1012541 | 968118.94 | 1 | 1234969.25 | 1632423.38 | 1 | 1152267.62 | 1781154.62 | 2248605.75 | 1326648 | 1720535.968 | 1525735.5 | 10 | 1 | 1.159639835 | 2.481803098 |
| O88712 | Ctbp1 | C-terminal-binding protein 1 Mus musculus | 357.62 | 21.54 | 47745 | 4 | 7 | 6084126.5 | 6867470.5 | 7400646 | 7924728 | 7334426.5 | 4653719 | 10668258 | 6802740 | 7925996 | 10158226 | 10567935 | 9932028 | 8020798 | 7461330 | 11083005 | 9 | 1 | 1.129103091 | 1.321630133 |
| Q9WTX6 | Cul1 | Cullin-1 Mus musculus | 508.56 | 14.69 | 89692 | 10 | 10 | 9356674 | 8197263 | 7475768 | 8358356 | 10032447 | 10148699 | 6894816 | 10197417 | 9848283 | 8506364 | 12365513 | 10492369 | 9581891 | 10079986 | 12049857 | 8 | 1 | 1.050093173 | 1.256770556 |
| Q9D4H8 | Cul2 | Cullin-2 Mus musculus | 222.18 | 7.92 | 86877 | 5 | 5 | 1236850.5 | 1862539.88 | 1368809.25 | 1538803.75 | 1462571.5 | 1604546.5 | 2215151.5 | 2744167.5 | 2896369.25 | 1995367.25 | 1995650.75 | 2344885.75 | 1822770.25 | 2042566.5 | 2058204.75 | 1 | 1 | 1.533635071 | 1.374118094 |
| Q7TMB8 | Cyfip1 | Cytoplasmic FMR1-interacting protein 1 Mus musculus | 1945.15 | 33.2 | 145241 | 22 | 37 | 53762173 | 46518805 | 46904561 | 42789213 | 44979477 | 49911569 | 49160321 | 51569785 | 58897345 | 50120113 | 60785341 | 55906693 | 49154529 | 55226257 | 59098337 | 4 | 1 | 1.105147731 | 1.192449943 |
| P11030 | Dbi | Acyl-CoA-binding protein Rattus norvegicus | 144.97 | 39.08 | 10027 | 2 | 2 | 8975738 | 7477482 | 6957561 | 8622616 | 7808447.5 | 6320285 | 6360749 | 6302452 | 4650732.5 | 8723353 | 5720608 | 6389145.5 | 7048027 | 6013892.5 | 5792070.5 | 4 | 1 | 0.812150439 | 0.777166416 |
| Q501J6 | Ddx17 | Probable ATP-dependent RNA helicase DDX17 Mus musculus | 957.64 | 31.54 | 72399 | 11 | 18 | 150077089 | 155918529 | 158525361 | 155182561 | 152922897 | 131302857 | 131145177 | 134895601 | 144248145 | 136351825 | 151269873 | 142305521 | 149909057 | 144687137 | 144611137 | 0 | 1 | 0.877453285 | 0.94843082 |
| Q9QYJ3 | Dnajb1 | DnaJ homolog subfamily B member 1 Mus musculus | 165.74 | 15 | 38167 | 5 | 5 | 4165890 | 3318580 | 3386349.25 | 3622478 | 3842204.5 | 2761575.5 | 3933247.5 | 3940806.75 | 3422719.25 | 4121723 | 4008319.5 | 4863904 | 5816166 | 5807165 | 5173635 | 11 | 1 | 0.991523016 | 1.399972024 |
| P21575 | Dnm1 | Dynamin-1 Rattus norvegicus | 204.22 | 5.56 | 97295 | 1 | 5 | 5910187 | 5974341.5 | 4994817 | 5374599.5 | 5425708 | 4874912.5 | 4518357.5 | 4186290.25 | 4259415.5 | 5206855 | 3825795.75 | 4161140.75 | 5243458.5 | 4622964 | 4545649 | 1 | 1 | 0.832591028 | 0.809222861 |
| P58252 | Eef2 | Elongation factor 2 Mus musculus | 3344.97 | 61.54 | 95314 | 50 | 52 | 326634849 | 330409569 | 354644641 | 332265505 | 309318529 | 333630721 | 392710305 | 367695649 | 369891297 | 342406049 | 362267553 | 376556641 | 344632257 | 381164993 | 371968961 | 2 | 1 | 1.092580547 | 1.110881446 |
| Q8VC03 | Eml3 | Echinoderm microtubule-associated protein-like 3 Mus musculus | 41.79 | 1.78 | 95695 | 1 | 1 | 1 | 1 | 320697.59 | 1 | 275340.22 | 229971.88 | 542201.38 | 1 | 218796.33 | 1 | 311464.5 | 332518.34 | 358659.03 | 352938.695 | 409112.91 | 11 | 1 | 1.662590167 | 2.960692364 |
| Q9DCM0 | Ethe1 | Persulfide dioxygenase ETHE1, mitochondrial Mus musculus | 166.01 | 24.41 | 27739 | 4 | 4 | 8303762.5 | 9284230 | 9074427 | 10790023 | 10753311 | 8005515 | 8971851 | 7274087 | 6298631 | 8053064 | 9034530 | 6495023 | 8276262.5 | 8036088.5 | 8133424 | 1 | 1 | 0.800799598 | 0.829264664 |
| O54924 | Exoc8 | Exocyst complex component 8 Rattus norvegicus | 39.47 | 2.23 | 81043 | 1 | 1 | 1680718 | 2204113.75 | 1868724.5 | 1831553.12 | 1860770.5 | 1780783.12 | 1816859.25 | 1680740.62 | 1433218.25 | 1869701 | 1627611.38 | 1515096.25 | 1721683.12 | 1598812 | 1366646.75 | 7 | 1 | 0.908470397 | 0.828916904 |
| Q921I9 | Exosc4 | Exosome complex component RRP41 Mus musculus | 60.6 | 4.9 | 26250 | 1 | 1 | 1 | 1 | 1 | 1 | 354221.69 | 1 | 876910.44 | 1 | 291807.81 | 1 | 345001.19 | 601768.56 | 465370.28 | 378441.03 | 625084.06 | 10 | 1 | 3.29936897 | 6.819565007 |
| P16858 | Gapdh | Glyceraldehyde-3-phosphate dehydrogenase Mus musculus | 2270.76 | 87.69 | 35810 | 29 | 36 | 1044268161 | 1000019969 | 1184233217 | 1225996161 | 1173118081 | 1245029633 | 2175302657 | 1257351297 | 918552705 | 1680648449 | 1555629825 | 1456819201 | 1350194817 | 1194443137 | 1688264577 | 5 | 1 | 1.293062535 | 1.287459261 |
| O70325 | Gpx4 | Phospholipid hydroperoxide glutathione peroxidase Mus musculus | 178.09 | 22.34 | 22229 | 4 | 4 | 12119297 | 14274557 | 13153134 | 13247991 | 12999605 | 13173246 | 11025923 | 10684127 | 14783681 | 12352489 | 11352050 | 11366790 | 11901251 | 12928428 | 10071254 | 9 | 1 | 0.942622663 | 0.875752524 |
| P97576 | Grpel1 | GrpE protein homolog 1, mitochondrial Rattus norvegicus | 223.31 | 25.35 | 24297 | 5 | 5 | 266657.03 | 958875.81 | 245341.73 | 1302648 | 955945.62 | 1517999.25 | 1688386.75 | 2520586.75 | 1998680.38 | 1031222.06 | 2168800.75 | 1620054 | 1082695.12 | 1430473.75 | 1375657.88 | 1 | 1 | 2.348022491 | 2.058653167 |
| P01901 | H2-K1 | H-2 class I histocompatibility antigen, K-B alpha chain Mus musculus | 226.74 | 9.21 | 41302 | 0 | 5 | 69263969 | 77587809 | 70712513 | 67576433 | 76025153 | 78781385 | 73622025 | 78075777 | 74587649 | 68217969 | 65218909 | 52335877 | 68947729 | 66865181 | 53157721 | 8 | 1 | 1.033555019 | 0.848710901 |
| P62959 | Hint1 | Histidine triad nucleotide-binding protein 1 Rattus norvegicus | 163.67 | 35.71 | 13777 | 3 | 3 | 41841437 | 48005725 | 47729601 | 49812909 | 48736601 | 39706669 | 35915009 | 35364945 | 39126501 | 41288473 | 44455905 | 38582081 | 40950265 | 40046777 | 41511153 | 0 | 1 | 0.810590006 | 0.870492633 |
| Q8BUK6 | Hook3 | Protein Hook homolog 3 Mus musculus | 50.94 | 1.95 | 83218 | 1 | 1 | 250239.77 | 1 | 429937.62 | 1 | 1 | 1 | 1 | 274501.09 | 225404.53 | 1 | 671321.38 | 452964.84 | 497625.04 | 324865.88 | 541348.06 | 11.5 | 1 | 0.734964764 | 3.658037245 |
| P59438 | Hps5 | Hermansky-Pudlak syndrome 5 protein homolog Mus musculus | 54.34 | 0.98 | 126337 | 1 | 1 | 1293994 | 977232.12 | 679642.94 | 1212638.62 | 1405209.38 | 752711.75 | 785273.75 | 664310 | 373976.38 | 977916.69 | 910365.5 | 354279.59 | 643137.06 | 433899.06 | 407823.03 | 4 | 1 | 0.638241902 | 0.493741056 |
| Q8BTX9 | Hsdl1 | Inactive hydroxysteroid dehydrogenase-like protein 1 Mus musculus | 138.33 | 10.91 | 36868 | 2 | 2 | 1532226.12 | 1417680.75 | 1644205.25 | 1829637.88 | 1761696.75 | 1673343 | 1052542.88 | 1711213.88 | 1486577.12 | 1580726 | 1290970.25 | 1422520.62 | 1241677.12 | 1402199.5 | 1404548.25 | 9 | 1 | 0.916798204 | 0.826090004 |
| Q8R0F6 | Ilkap | Integrin-linked kinase-associated serine/threonine phosphatase 2C Mus musculus | 135.03 | 10.97 | 42774 | 3 | 3 | 1029290.25 | 1227866.88 | 1092892.12 | 1115054.88 | 891648.75 | 999663.75 | 1561678.62 | 1180186.5 | 1344525.12 | 1438719 | 1795422.25 | 1404302.75 | 1251686.88 | 1146389.12 | 1594840.5 | 5 | 1 | 1.218046293 | 1.342724158 |
| P06151 | Ldha | L-lactate dehydrogenase A chain Mus musculus | 2215.19 | 87.05 | 36499 | 28 | 32 | 712083585 | 734669377 | 670777793 | 591029249 | 691721537 | 752190401 | 913085825 | 777360193 | 746621441 | 770667137 | 801903489 | 738947969 | 793248129 | 807796097 | 718574017 | 0 | 1 | 1.164587388 | 1.135338252 |
| P62313 | Lsm6 | U6 snRNA-associated Sm-like protein LSm6 Mus musculus | 43.15 | 10 | 9128 | 1 | 1 | 3327701 | 2948293.25 | 2895213.25 | 3063587.25 | 2961613 | 2456458.75 | 2321062 | 2609329.25 | 2972809.5 | 2594495 | 2895601 | 836077.5 | 2468198.25 | 2749669.5 | 1986840.5 | 3 | 1 | 0.852448468 | 0.719669209 |
| Q02527 | Mgat3 | Beta-1,4-mannosyl-glycoprotein 4-beta-N-acetylglucosaminyltransferase Rattus norvegicus | 52.47 | 1.3 | 62022 | 1 | 1 | 7478302.5 | 7832926.5 | 4383691.5 | 6138261 | 8969417 | 5192263 | 4447244 | 5483558 | 3172323 | 3684922 | 3852163 | 1 | 3743604.25 | 4723743 | 1 | 3 | 1 | 0.631570944 | 0.353982541 |
| P57716 | Ncstn | Nicastrin Mus musculus | 874.54 | 25.99 | 78492 | 13 | 13 | 66070841 | 61829253 | 59012321 | 56884313 | 61278481 | 52296193 | 41820505 | 59430529 | 67205633 | 38452045 | 49731573 | 49054261 | 53749849 | 58862705 | 41054609 | 7 | 1 | 0.849642637 | 0.827510691 |
| O09111 | Ndufb11 | NADH dehydrogenase [ubiquinone] 1 beta subcomplex subunit 11, mitochondrial Mus musculus | 37.18 | 11.26 | 17444 | 1 | 1 | 1 | 1 | 1 | 1 | 835302.44 | 1 | 1 | 1 | 1 | 1 | 1103796.88 | 543035.06 | 1302110 | 1082863.485 | 1382512 | 10 | 1 | 5.98583E-06 | 6.481833691 |
| Q9DCT2 | Ndufs3 | NADH dehydrogenase [ubiquinone] iron-sulfur protein 3, mitochondrial Mus musculus | 441.05 | 33.08 | 30149 | 7 | 7 | 11549762 | 10014481 | 10929866 | 10903163 | 10815180 | 10859460 | 8732876 | 11599679 | 11499027 | 10191845 | 8231620 | 9858302 | 8792021 | 10124517 | 8114924.5 | 12 | 1 | 0.975474915 | 0.832306653 |
| Q9DCD0 | Pgd | 6-phosphogluconate dehydrogenase, decarboxylating Mus musculus | 1812.8 | 69.57 | 53247 | 26 | 27 | 151253345 | 164736529 | 175545713 | 136175393 | 153375793 | 150455393 | 202020865 | 177817105 | 194957041 | 174516961 | 194816513 | 191027361 | 169827873 | 183155201 | 198447505 | 5 | 1 | 1.151942903 | 1.199962009 |
| Q9D0F9 | Pgm1 | Phosphoglucomutase-1 Mus musculus | 1225.89 | 48.04 | 61418 | 18 | 18 | 24706457 | 27459615 | 26923959 | 22012243 | 24741713 | 28689809 | 32885623 | 30358749 | 32106349 | 26712269 | 30071991 | 29115397 | 28329307 | 30982065 | 27291501 | 2 | 1 | 1.197934066 | 1.158500016 |
| P51432 | Plcb3 | 1-phosphatidylinositol 4,5-bisphosphate phosphodiesterase beta-3 Mus musculus | 235.68 | 7.54 | 139487 | 5 | 5 | 870095.25 | 1536200 | 1617768.75 | 974266.44 | 1494747.25 | 2052958.38 | 2355656.75 | 1938502.5 | 1166802.75 | 2317917.75 | 2077326 | 1854785.5 | 1573709.75 | 2019036.62 | 2411529.5 | 3 | 1 | 1.514203064 | 1.53030471 |
| Q76MZ3 | Ppp2r1a | Serine/threonine-protein phosphatase 2A 65 kDa regulatory subunit A alpha isoform Mus musculus | 2107.18 | 64.35 | 65323 | 23 | 33 | 114746873 | 128048809 | 126684097 | 118607377 | 116083617 | 136471233 | 125694593 | 135473073 | 141894609 | 124516929 | 141427409 | 131924945 | 127061265 | 137160801 | 137223073 | 4 | 1 | 1.099110494 | 1.116898604 |
| P54645 | Prkaa1 | 5'-AMP-activated protein kinase catalytic subunit alpha-1 Rattus norvegicus | 131.28 | 6.44 | 63973 | 3 | 3 | 1389563.25 | 1004741 | 1438128.75 | 1325978 | 1516875 | 1388945.25 | 1844298 | 1059585.88 | 1266091.38 | 1747111.25 | 1872544 | 1482560.25 | 1745537.25 | 1528098.75 | 1764208.5 | 11 | 1 | 1.094489698 | 1.257316728 |
| Q63009 | Prmt1 | Protein arginine N-methyltransferase 1 Rattus norvegicus | 1101.34 | 53.54 | 40522 | 20 | 20 | 48532225 | 47985949 | 46430077 | 42264709 | 45721829 | 40695521 | 28322159 | 41056333 | 45250313 | 31501509 | 35086825 | 35684169 | 35647897 | 44480921 | 35413465 | 1 | 1 | 0.808998228 | 0.806778735 |

| | | | | | | | | | | | | | | | | | | | | | | | | | | |
|---|---|---|---|---|---|---|---|---|---|---|---|---|---|---|---|---|---|---|---|---|---|---|---|---|---|---|
| Q8CGC6 | Rbm28 | RNA-binding protein 28 Mus musculus | 440.11 | 13.87 | 84214 | 9 | 9 | 3444275 | 3734736.5 | 787139.31 | 3987158 | 445939.22 | 4349352 | 5835207 | 1057811.25 | 3911269 | 4912077 | 6249065.5 | 4920380.5 | 4835301.5 | 3772427 | 5794008 | 4 | 1 | 1.618301062 | 2.062317202 |
| Q9CWY8 | Rnaseh2a | Ribonuclease H2 subunit A Mus musculus | 54.23 | 5.32 | 33513 | 1 | 1 | 453792.31 | 325170.91 | 500514.62 | 436523.62 | 482824.31 | 642599.88 | 724128.38 | 392136.34 | 305759.91 | 581576.62 | 568938.06 | 526288.62 | 485672.84 | 512037.69 | 940967.25 | 9 | 1 | 1.203461032 | 1.379783929 |
| Q80TE0 | Rpap1 | RNA polymerase II-associated protein 1 Mus musculus | 44.06 | 0.92 | 155270 | 1 | 1 | 394524.53 | 429404.41 | 408864.81 | 426645.22 | 414859.7425 | 302784.22 | 426448.12 | 452765.31 | 377381.41 | 389844.765 | 347041.75 | 273627.06 | 403163.72 | 281051.38 | 326220.9775 | 8 | 1 | 0.939702567 | 0.786340404 |
| Q9JJ80 | Rpf2 | Ribosome production factor 2 homolog Mus musculus | 333.42 | 27.45 | 35364 | 6 | 6 | 1278550 | 1473422 | 1305544.5 | 2019469.38 | 2440933.5 | 1119897 | 1536298.25 | 1132022 | 403525.84 | 1684512.5 | 1177737.88 | 918642.62 | 1304206.75 | 424626.34 | 1059695.25 | 6 | 1 | 0.689869829 | 0.573486156 |
| P39032 | Rpl36 | 60S ribosomal protein L36 Rattus norvegicus | 105.28 | 19.05 | 12268 | 1 | 2 | 1348309.25 | 1191118.5 | 2931916.5 | 1890920.88 | 1829570 | 2083938.75 | 6313029.5 | 1461888.5 | 1275342.75 | 4085827.25 | 3511817.75 | 4115853.5 | 2312828.25 | 3813123 | 5311992.5 | 8 | 1 | 1.655820251 | 2.074190271 |
| P49242 | Rps3a | 40S ribosomal protein S3a Rattus norvegicus | 740.66 | 45.83 | 29945 | 13 | 13 | 113750105 | 112958609 | 114948289 | 118346137 | 109531193 | 101607961 | 115720113 | 97707385 | 102285449 | 119121521 | 107497841 | 110747673 | 108324145 | 89856257 | 105194505 | 9 | 1 | 0.94189656 | 0.915871776 |
| Q99P72 | Rtn4 | Reticulon-4 Mus musculus | 984.33 | 25.13 | 126613 | 15 | 15 | 62349229 | 58925973 | 62964929 | 54204753 | 69060793 | 73949689 | 78835825 | 72101969 | 69173321 | 78174937 | 76448497 | 76274977 | 69393881 | 63865777 | 76015553 | 0 | 1 | 1.210500387 | 1.177209763 |
| Q9CXV1 | Sdhd | Succinate dehydrogenase [ubiquinone] cytochrome b small subunit, mitochondrial Mus musculus | 82.47 | 16.35 | 17014 | 1 | 1 | 2303064.75 | 1947185.12 | 1287627.62 | 1616036.25 | 1565868.62 | 1784257.62 | 946340.81 | 1329562.62 | 928632.25 | 1572531.5 | 934084.25 | 731724.19 | 1033517.31 | 1302246 | 1174623 | 6 | 1 | 0.752464285 | 0.593615131 |
| Q9D554 | Sf3a3 | Splicing factor 3A subunit 3 Mus musculus | 868.71 | 38.72 | 58842 | 14 | 14 | 18947025 | 19965373 | 17937693 | 16690158 | 19102867 | 19732329 | 20906475 | 19693309 | 22981763 | 18912499 | 21286517 | 20089519 | 19252769 | 20745159 | 22759827 | 5 | 1 | 1.103442753 | 1.124031612 |
| Q6AYE2 | Sh3glb1 | Endophilin-B1 Rattus norvegicus | 215.15 | 10.68 | 40787 | 3 | 3 | 6654449 | 7580770 | 5955345.5 | 7714775 | 7210518 | 6026126 | 5893195.5 | 5893043 | 5585304.5 | 6307689 | 5398864 | 5289601.5 | 6323193 | 5579678 | 4803519 | 2 | 1 | 0.845924324 | 0.780127767 |
| Q9CZN7 | Shmt2 | Serine hydroxymethyltransferase, mitochondrial Mus musculus | 1282.55 | 57.54 | 55759 | 23 | 24 | 110442577 | 112967289 | 116171249 | 138613057 | 121700257 | 103043881 | 124117649 | 104289353 | 111176185 | 128823521 | 106630577 | 105782385 | 104379561 | 99759425 | 112781897 | 9 | 1 | 0.952585257 | 0.882378331 |
| Q6P9P0 | Slf2 | SMC5-SMC6 complex localization factor protein 2 Mus musculus | 36.1 | 0.55 | 143970 | 1 | 1 | 1371724 | 1059111.62 | 392924.06 | 1025785.38 | 1202280.12 | 786973.88 | 890074 | 506874.47 | 187942.3 | 1417611 | 267475.47 | 270415.28 | 209157.97 | 560455.44 | 326876.04 | 8 | 1 | 0.750120108 | 0.323522715 |
| Q91VZ6 | Smap1 | Stromal membrane-associated protein 1 Mus musculus | 41.36 | 2.73 | 47660 | 1 | 1 | 818227.31 | 1 | 1 | 1 | 1035083.69 | 880978.94 | 1188068 | 581820.88 | 958232.0175 | 1182060.25 | 1126479.375 | 1292725.62 | 1158749.38 | 941863.62 | 1112578.88 | 4 | 1 | 2.585185288 | 3.039094765 |
| Q62189 | Snrpa | U1 small nuclear ribonucleoprotein A Mus musculus | 295.14 | 31.71 | 31835 | 5 | 6 | 5356152.5 | 5656992.5 | 5636798.5 | 5084548 | 5502430 | 4659304.5 | 6160083 | 4622001.5 | 4284844 | 6169961.5 | 4777405.5 | 3970186.5 | 5356026 | 4360877.5 | 3275023.75 | 10 | 1 | 0.950775384 | 0.798163598 |
| Q6ZWQ7 | Spcs3 | Signal peptidase complex subunit 3 Mus musculus | 177.23 | 21.67 | 20313 | 4 | 4 | 5424392.5 | 4938548 | 4468089 | 5067779.5 | 4013489.25 | 5586489.5 | 6127951 | 5763785.5 | 4470409 | 6896460.5 | 5932688 | 5307993.5 | 7108939.5 | 5921153.5 | 5555923 | 3 | 1 | 1.206287041 | 1.247337131 |
| O88838 | Spsb2 | SPRY domain-containing SOCS box protein 2 Mus musculus | 35.47 | 2.65 | 28938 | 1 | 1 | 9523748 | 11776717 | 10869080 | 11468616 | 11094955 | 8139882 | 7965588.5 | 9188951 | 10026194 | 7675419 | 9446949 | 8060028 | 9073398 | 9557168 | 6493642.5 | 1 | 1 | 0.785557952 | 0.778891987 |
| Q9JM90 | Stap1 | Signal-transducing adaptor protein 1 Mus musculus | 637.43 | 59.93 | 34628 | 13 | 13 | 17729511 | 12382445 | 13236752 | 13402185 | 15024507 | 21969469 | 22907261 | 20415201 | 17022513 | 22093031 | 24029665 | 22339825 | 17381839 | 18226125 | 24936277 | 1 | 1 | 1.454641493 | 1.489559529 |
| Q99KH8 | Stk24 | Serine/threonine-protein kinase 24 Mus musculus | 404.54 | 25.99 | 47954 | 8 | 8 | 3952477.25 | 5645742.5 | 5255120 | 3935718 | 3924994.5 | 6752330 | 7187371.5 | 7718677.5 | 3507326 | 5698509 | 8061818.5 | 6721213.5 | 6346914.5 | 5458301.5 | 7959442 | 5 | 1 | 1.358815841 | 1.520983117 |
| P70279 | Surf6 | Surfeit locus protein 6 Mus musculus | 80.44 | 4.79 | 41235 | 2 | 2 | 1 | 1 | 371485.03 | 1 | 1 | 1 | 541889.56 | 421203.66 | 1 | 417635.28 | 374938.62 | 516285.59 | 460355.56 | 264122.03 | 686076 | 6 | 1 | 3.71674636 | 6.196085521 |
| Q9QXE7 | Tbl1x | F-box-like/WD repeat-containing protein TBL1X Mus musculus | 149.35 | 5.5 | 56802 | 1 | 2 | 2227288 | 1777417.62 | 1661314.12 | 2389203.5 | 1955666.62 | 1457449.62 | 1323082.12 | 1617709.38 | 1014356.56 | 1444561.12 | 1691760 | 689089.94 | 1212120.62 | 1110346.62 | 676803 | 0 | 1 | 0.684969957 | 0.537426768 |
| Q9CQM5 | Txndc17 | Thioredoxin domain-containing protein 17 Mus musculus | 231.16 | 39.84 | 14015 | 4 | 4 | 10687913 | 11595468 | 12056245 | 11825727 | 11382406 | 11841683 | 15313184 | 11843380 | 11408493 | 13330705 | 11970653 | 13543410 | 13247875 | 12179062 | 14637917 | 5 | 1 | 1.107557377 | 1.139556399 |
| Q920I9 | Wdr7 | WD repeat-containing protein 7 Mus musculus | 94.65 | 1.61 | 163450 | 2 | 2 | 1002719.735 | 1159235.5 | 904080.81 | 981449.44 | 966113.19 | 939328.44 | 744535.19 | 1000400.31 | 749268.19 | 714154.12 | 782606.38 | 1 | 1 | 905023.94 | 539627.19 | 4 | 1 | 0.827287248 | 0.444243677 |
| O88286 | Wiz | Protein Wiz Mus musculus | 43.91 | 0.89 | 184291 | 1 | 1 | 1078049.38 | 1476510.38 | 1311159 | 844011.31 | 1269574.75 | 1043124.06 | 1 | 894587.81 | 1797914.75 | 1 | 424755.94 | 780148.81 | 620749.5 | 852027.94 | 669420.5475 | 7 | 1 | 0.624759689 | 0.559781252 |
| P83741 | Wnk1 | Serine/threonine-protein kinase WNK1 Mus musculus | 553.13 | 6.52 | 250934 | 10 | 12 | 110027521 | 83453697 | 72778305 | 98426177 | 83955393 | 62985649 | 59351965 | 63745725 | 49866589 | 74684753 | 78977457 | 30579749 | 51005705 | 59091017 | 56883345 | 1 | 1 | 0.692390166 | 0.61638864 |
| Q9DBR1 | Xrn2 | 5'-3' exoribonuclease 2 Mus musculus | 1106.68 | 27.87 | 108687 | 21 | 21 | 15612037 | 14278676 | 19510667 | 14982429 | 17092421 | 17991569 | 16889809 | 18759791 | 21033045 | 15071528 | 20653423 | 21080833 | 19314541 | 21936897 | 22489725 | 7 | 1 | 1.101496007 | 1.294554485 |
| Q505B7 | Zbtb8os | Protein archease Mus musculus | 47.66 | 5.95 | 19727 | 1 | 1 | 1287060.12 | 1329972.12 | 1224755.25 | 1403056.38 | 1149620.25 | 1227718.38 | 922835.88 | 1382794.5 | 1972079 | 1107142.38 | 811937.5 | 1053506.75 | 908428.81 | 1220635.12 | 902848.06 | 11 | 1 | 1.034108569 | 0.765874379 |
| Q9Z2U2 | Zfp292 | Zinc finger protein 292 Mus musculus | 34.66 | 0.41 | 301048 | 1 | 1 | 923951.88 | 2765312 | 2547395 | 1599058 | 1505235.12 | 1128554.38 | 1181828 | 965151.19 | 2146554.25 | 836511.81 | 1 | 1 | 1081832.38 | 1 | 1 | 6 | 1 | 0.670017321 | 0.115816501 |
| Q9QUJ7 | Acsl4 | Long-chain-fatty-acid--CoA ligase 4 Mus musculus | 1179.65 | 41.63 | 79077 | 22 | 23 | 77413993 | 74544825 | 73041393 | 67446297 | 74989745 | 67852337 | 61157321 | 69063353 | 66458345 | 63639425 | 72232849 | 54891777 | 66980113 | 68631729 | 61568669 | 2 | 2 | 0.893136642 | 0.882616057 |
| Q9QZQ1 | Afdn | Afadin Mus musculus | 78.26 | 0.99 | 206499 | 1 | 1 | 997290.12 | 1019074 | 1066013.75 | 898114.5 | 978138.25 | 1157515.12 | 1050873.25 | 1269727.38 | 1333742.5 | 968917.56 | 1104009.25 | 1086212.12 | 1301657.12 | 1336512.25 | 1016038.62 | 5 | 2 | 1.165800854 | 1.178637775 |
| Q8C0I1 | Agps | Alkyldihydroxyacetonephosphate synthase, peroxisomal Mus musculus | 624.79 | 23.26 | 71684 | 11 | 11 | 6235871 | 5000361 | 8535396 | 3113150 | 5289479.5 | 6221845 | 10641165 | 10973204 | 10437807 | 7975788 | 10978843 | 8863578 | 8071466 | 7456885 | 10407553 | 3 | 2 | 1.641562657 | 1.62482809 |
| P47738 | Aldh2 | Aldehyde dehydrogenase, mitochondrial Mus musculus | 1506.28 | 68.21 | 56538 | 24 | 25 | 182985393 | 204331585 | 209732513 | 200265169 | 210116641 | 191271025 | 209545857 | 225216913 | 228857569 | 194503729 | 212343073 | 207853137 | 227798145 | 214104417 | 213676385 | 10 | 2 | 1.041654247 | 1.067839719 |
| Q80VQ0 | Aldh3b1 | Aldehyde dehydrogenase family 3 member B1 Mus musculus | 105.23 | 8.55 | 52292 | 2 | 2 | 2891052.25 | 3489360.75 | 2962967.5 | 2642890.75 | 2990492.75 | 2874865.25 | 2193427.5 | 2756263.75 | 3004328 | 2437974.5 | 1855160.38 | 2087835.12 | 2403081.75 | 2925832.25 | 1889801.88 | 6 | 2 | 0.885829476 | 0.745268563 |
| Q91X20 | Ash2l | Set1/Ash2 histone methyltransferase complex subunit ASH2 Mus musculus | 112.32 | 5.3 | 68250 | 2 | 2 | 690015.06 | 1 | 364462.91 | 1 | 613051.56 | 714846.12 | 1088333 | 796226.06 | 604197.88 | 560488.06 | 1613508 | 1584157.75 | 454052.47 | 722250 | 756632.12 | 4 | 2 | 2.257283327 | 3.076763616 |
| Q6PA06 | Atl2 | Atlastin-2 Mus musculus | 92.44 | 4.12 | 66224 | 2 | 2 | 733836.25 | 899531.75 | 923835.75 | 877421.19 | 803702.88 | 972467.75 | 1475015.75 | 963623.56 | 894900.94 | 1103005.38 | 906268.88 | 1224530.5 | 1152944.62 | 1186608.25 | 902368.69 | 2 | 2 | 1.276214019 | 1.267651104 |
| O35855 | Bcat2 | Branched-chain-amino-acid aminotransferase, mitochondrial Mus musculus | 290.18 | 15.01 | 44127 | 5 | 5 | 23033639 | 21263059 | 21014819 | 22546623 | 23722181 | 22603343 | 22739415 | 21394163 | 22079177 | 21918281 | 20837241 | 19071875 | 21769841 | 20771775 | 19864535 | 12 | 2 | 0.992418538 | 0.916965161 |
| P29594 | Casp2 | Caspase-2 Mus musculus | 45.19 | 1.99 | 50661 | 1 | 1 | 778569.94 | 877849.38 | 870675.62 | 927981.62 | 863769.14 | 1 | 1 | 1 | 1 | 1 | 1 | 1 | 798172.12 | 831248.62 | 1 | 0 | 2 | 1.15772E-06 | 0.377282231 |
| Q8VEH6 | Cbwd1 | COBW domain-containing protein 1 Mus musculus | 132.73 | 9.16 | 43772 | 2 | 2 | 1518167.38 | 361844.88 | 1746521.25 | 1475068.88 | 1537275.12 | 2163748.25 | 2115234.75 | 2072216 | 1924196.12 | 1723247.12 | 1667754.62 | 1989867.75 | 1655800.62 | 2028631.25 | 1834892.62 | 1 | 2 | 1.506074216 | 1.382303988 |
| P23198 | Cbx3 | Chromobox protein homolog 3 Mus musculus | 440.56 | 36.61 | 20855 | 5 | 7 | 36266237 | 42719905 | 35782081 | 40401865 | 45056341 | 41024453 | 44086485 | 41533629 | 40831349 | 45330669 | 46874357 | 45421365 | 45622977 | 40619385 | 46081829 | 7 | 2 | 1.062829648 | 1.121829491 |
| O08585 | Clta | Clathrin light chain A Mus musculus | 123.19 | 10.64 | 25604 | 3 | 3 | 22239505 | 23305617 | 20524421 | 27492887 | 23438983 | 21142393 | 16876155 | 20501821 | 19461057 | 20894157 | 21101131 | 17924757 | 21634389 | 18912855 | 16271817 | 2 | 2 | 0.845080247 | 0.819177705 |
| Q9QXT0 | Cnpy2 | Protein canopy homolog 2 Mus musculus | 147.19 | 15.93 | 20767 | 2 | 2 | 2663602.5 | 2918795.5 | 2321279.5 | 2802940.5 | 2482477.5 | 2093529.62 | 2550972.25 | 2076847 | 2359377.5 | 2287966.25 | 2098434.5 | 2131443.25 | 2587944.75 | 1867885.38 | 1932708.25 | 3 | 2 | 0.86197667 | 0.805090548 |
| Q8BXC6 | Commd2 | COMM domain-containing protein 2 Mus musculus | 74.28 | 9.55 | 22848 | 2 | 2 | 2578403.25 | 2830499.25 | 2162274.75 | 2516614.75 | 2383693 | 1939588.75 | 1954253.25 | 2307488.5 | 2412509.5 | 1878735.5 | 2128163.75 | 2031448.12 | 2356999.25 | 2231211.75 | 1805655.25 | 3 | 2 | 0.841325271 | 0.846208621 |
| Q8C166 | Cpne1 | Copine-1 Mus musculus | 608.36 | 28.36 | 58887 | 10 | 11 | 17995381 | 17383457 | 16692100 | 21040703 | 19500999 | 20580851 | 20763909 | 20206547 | 19734429 | 23906471 | 21675361 | 21673537 | 19350763 | 22531635 | 22290513 | 4 | 2 | 1.135829915 | 1.16098417 |
| P63154 | Crnkl1 | Crooked neck-like protein 1 Mus musculus | 108.38 | 4.49 | 83416 | 3 | 3 | 2482272.75 | 2071717.75 | 2441203.75 | 3190848 | 2019391.75 | 2310142.5 | 1256227.75 | 2066481.25 | 2559921 | 1654012.5 | 1472084.5 | 1935470 | 2119509.25 | 1949526.88 | 1801299.75 | 7 | 2 | 0.806754188 | 0.760144242 |
| P98078 | Dab2 | Disabled homolog 2 Mus musculus | 1579.37 | 45.04 | 82312 | 9 | 22 | 33908749 | 31361015 | 43650805 | 33180885 | 30821723 | 30817559 | 43306921 | 40136981 | 35062329 | 31336175 | 44797533 | 46747857 | 37451257 | 35115013 | 43923221 | 12 | 2 | 1.044741186 | 1.203047993 |
| Q91X52 | Dcxr | L-xylulose reductase Mus musculus | 65.94 | 7.38 | 25746 | 2 | 2 | 134480689 | 89065553 | 74447881 | 112225169 | 119284305 | 70548753 | 53814041 | 66494233 | 38887289 | 62245973 | 104602489 | 16328652 | 57226097 | 66423661 | 48135453 | 0 | 2 | 0.551441559 | 0.552812773 |
| O35286 | Dhx15 | Pre-mRNA-splicing factor ATP-dependent RNA helicase DHX15 Mus musculus | 1227.15 | 33.08 | 91007 | 21 | 23 | 40992317 | 41301829 | 43735069 | 41636641 | 43165537 | 44596437 | 44462501 | 42507173 | 43727801 | 43284265 | 46653053 | 44389185 | 43071265 | 43825093 | 43951017 | 4 | 2 | 1.036743978 | 1.052450538 |
| D4A2Z8 | Dhx36 | ATP-dependent DNA/RNA helicase DHX36 Rattus norvegicus | 261.69 | 7.4 | 113843 | 6 | 6 | 1602403.38 | 2228548.25 | 2125017.75 | 1845307 | 3894404.25 | 996273.81 | 1602195.75 | 2874390 | 1254084.38 | 1337136.88 | 1814774 | 1301755.88 | 1799316 | 1070283.62 | 1206505.25 | 4 | 2 | 0.689492222 | 0.614982144 |
| Q5M9G7 | Diexf | Digestive organ expansion factor homolog Rattus norvegicus | 160.09 | 4.59 | 87825 | 3 | 3 | 1 | 1 | 1 | 1 | 380879.97 | 1 | 264339.38 | 1 | 1 | 1 | 295015.78 | 524641.725 | 644858 | 786307.12 | 372386 | 12 | 2 | 0.694026005 | 6.887159428 |
| Q9D1M4 | Eef1e1 | Eukaryotic translation elongation factor 1 epsilon-1 Mus musculus | 98.65 | 20.11 | 19859 | 2 | 2 | 1216115.75 | 895468.0925 | 645879.75 | 441789.62 | 1278087.25 | 1062562.88 | 506088.62 | 490350.12 | 1983323 | 1276212.38 | 1913167.25 | 1425502.38 | 2052571.38 | 1163893.38 | 1987580.25 | 11 | 2 | 1.187878618 | 1.907988618 |
| Q9CZX9 | Emc4 | ER membrane protein complex subunit 4 Mus musculus | 53.73 | 8.74 | 20117 | 1 | 1 | 3121233 | 3703782.5 | 3606992.75 | 4100907.5 | 3399092.25 | 3726315.5 | 3168380.5 | 3552956.75 | 3091991.5 | 3497489.5 | 3015585 | 2461886.25 | 3068528.25 | 3331299.75 | 3229043 | 9 | 2 | 0.950096261 | 0.84242335 |
| P97481 | Epas1 | Endothelial PAS domain-containing protein 1 Mus musculus | 34.03 | 0.8 | 96712 | 1 | 1 | 2291132.25 | 1687931.88 | 1177006.25 | 2503481.5 | 2628611.75 | 1435919.25 | 1274467.38 | 1163758.75 | 488025.5 | 1864217.5 | 1543764.75 | 459869.75 | 1435132.62 | 876038 | 885240.25 | 4 | 2 | 0.605199198 | 0.505439606 |
| P84089 | Erh | Enhancer of rudimentary homolog Mus musculus | 247.72 | 45.19 | 12259 | 4 | 4 | 9151626 | 11230285 | 8897056 | 8986972 | 10046686 | 8500464 | 7166663 | 9157656 | 9922256 | 8503789 | 9081167 | 6445727 | 7422902 | 8507233 | 7052146 | 6 | 2 | 0.895228276 | 0.797083061 |
| Q3V1V3 | Esf1 | ESF1 homolog Mus musculus | 43.06 | 1.42 | 98048 | 1 | 1 | 295701.62 | 221579.8 | 296168.06 | 246615.015 | 173010.58 | 211978.69 | 308541.88 | 312044.56 | 452822.75 | 215109.88 | 306219.19 | 391339.5 | 285005.38 | 381102.28 | 420882.66 | 8 | 2 | 1.216874617 | 1.447234679 |
| Q5U2Q7 | Etf1 | Eukaryotic peptide chain release factor subunit 1 Rattus norvegicus | 1018.21 | 59.5 | 49031 | 18 | 18 | 16706434 | 18603379 | 22570753 | 18915315 | 16865905 | 21865413 | 19680507 | 24615825 | 30455587 | 19475593 | 20928809 | 23151867 | 19234061 | 25609017 | 23104641 | 3 | 2 | 1.239490831 | 1.196095011 |
| Q8CGI1 | Fam193a | Protein FAM193A Mus musculus | 35.38 | 0.81 | 136714 | 1 | 1 | 4412990 | 2779078.25 | 2425787 | 4287065.5 | 4312787 | 2320984.5 | 2365210.75 | 2306710.75 | 2495928.313 | 2990807.25 | 3153871.5 | 965172.31 | 1854834.88 | 2244710 | 2275938.5 | 3 | 2 | 0.685028091 | 0.576061892 |
| P16879 | Fes | Tyrosine-protein kinase Fes/Fps Mus musculus | 35.95 | 1.46 | 93779 | 1 | 1 | 2004771.88 | 1651227.75 | 1695138.62 | 1868781.12 | 1642265.75 | 869185.25 | 1557708.753 | 1608757.88 | 2010727.88 | 1742164 | 395239 | 1599948 | 1653506.62 | 1560964.75 | 1514985 | 8 | 2 | 0.878851396 | 0.758801952 |
| Q8QZS3 | Flcn | Folliculin Mus musculus | 43.05 | 2.25 | 64327 | 1 | 1 | 235372.88 | 227347.08 | 1 | 1 | 210333.64 | 162474.89 | 188774.22 | 192436.62 | 294409 | 242190.56 | 256815.69 | 278214.16 | 217110.31 | 252109.97 | 251062.5325 | 9 | 2 | 1.605046136 | 1.865095042 |
| Q6NWW9 | Fndc3b | Fibronectin type III domain-containing protein 3B Mus musculus | 137.45 | 2.57 | 132764 | 2 | 2 | 239133.61 | 324202 | 728307.06 | 279746.22 | 794567 | 846021.44 | 961582.19 | 321376.16 | 327865.84 | 908318.38 | 943348.5 | 898717.5 | 793548.56 | 775130.12 | 885799.94 | 5 | 2 | 1.422327451 | 1.815986781 |
| P97379 | G3bp2 | Ras GTPase-activating protein-binding protein 2 Mus musculus | 256.45 | 11 | 54088 | 3 | 5 | 4602877.5 | 6003054.5 | 5904047.5 | 5445513 | 6130925 | 6844776.5 | 4102425 | 5902282.5 | 7081814.5 | 5990039 | 4051858 | 4278537 | 3644684 | 3627956 | 5791589.5 | 10 | 2 | 1.065331223 | 0.761742736 |
| Q8BFQ8 | Gatd1 | Glutamine amidotransferase-like class 1 domain-containing protein 1 Mus musculus | 232.11 | 30 | 23277 | 4 | 4 | 3404697.5 | 3548624.75 | 3877741.25 | 3405962.75 | 3242598.75 | 2896195 | 3343239.25 | 4276396.5 | 4953308.5 | 3022099 | 3734045.5 | 4494037.5 | 4260417 | 3614464.75 | 4216592 | 11 | 2 | 1.057873853 | 1.162470977 |
| P50399 | Gdi2 | Rab GDP dissociation inhibitor beta Rattus norvegicus | 2045.24 | 74.38 | 50537 | 22 | 29 | 167608433 | 167957473 | 152872913 | 156400529 | 174835505 | 170658961 | 145581153 | 180769025 | 171308737 | 142153057 | 174200241 | 175019025 | 175844961 | 193122001 | 171391649 | 12 | 2 | 0.988771255 | 1.085281406 |
| Q8BMS1 | Hadha | Trifunctional enzyme subunit alpha, mitochondrial Mus musculus | 1110.22 | 38.66 | 82670 | 19 | 19 | 28916769 | 24112179 | 27524993 | 26538753 | 25760241 | 30190747 | 31342299 | 28258399 | 26115371 | 28527293 | 30012017 | 30177529 | 28270761 | 28579523 | 30398627 | 5 | 2 | 1.087172888 | 1.109786976 |
| O08528 | Hk2 | Hexokinase-2 Mus musculus | 1259.89 | 33.04 | 102535 | 18 | 23 | 36978961 | 40303117 | 41025301 | 33905165 | 36865553 | 38589873 | 41798797 | 42206665 | 50227873 | 40269713 | 47619449 | 47321605 | 45048685 | 53544405 | 39995781 | 4 | 2 | 1.127010079 | 1.235097712 |
| Q3TRM8 | Hk3 | Hexokinase-3 Mus musculus | 2099.18 | 46.85 | 100101 | 13 | 30 | 111634473 | 134472689 | 119574729 | 111517449 | 118247505 | 113514369 | 97629161 | 107615713 | 128792977 | 103279529 | 102817673 | 91471761 | 109456137 | 117566929 | 102365321 | 6 | 2 | 0.925072916 | 0.879470309 |
| P11499 | Hsp90ab1 | Heat shock protein HSP 90-beta Mus musculus | 4068.121 | 76.38 | 83281 | 38 | 61 | 858858113 | 907080065 | 860898369 | 795965633 | 908185857 | 972307969 | 886827201 | 960210113 | 966483137 | 887882817 | 1011949569 | 938138369 | 905636929 | 960531841 | 927077569 | 4 | 2 | 1.079132798 | 1.095208353 |
| P26772 | Hspe1 | 10 kDa heat shock protein, mitochondrial Rattus norvegicus | 215.66 | 42.16 | 10902 | 4 | 4 | 18476301 | 19934239 | 19126895 | 21050439 | 16966677 | 16631221 | 22463085 | 16221974 | 14268253 | 20142719 | 12452855 | 15622031 | 18734659 | 13708221 | 10608190 | 9 | 2 | 0.939015997 | 0.744349225 |
| Q8BU30 | Iars | Isoleucine--tRNA ligase, cytoplasmic Mus musculus | 622.99 | 13.55 | 144271 | 14 | 14 | 49474901 | 37118825 | 34908689 | 42228605 | 41116593 | 34525333 | 33683565 | 33059561 | 32200303 | 37844045 | 35971697 | 28782593 | 34869073 | 32809721 | 37057957 | 2 | 2 | 0.836293889 | 0.827400615 |
| P19182 | Ifrd1 | Interferon-related developmental regulator 1 Mus musculus | 141.02 | 5.35 | 49935 | 2 | 2 | 1 | 1 | 1 | 1 | 285720.09 | 1 | 1 | 1 | 1 | 1 | 342989.8075 | 258396.19 | 269131.66 | 481769.69 | 362661.69 | 10 | 2 | 1.74994E-05 | 6.002115669 |
| Q91YE6 | Ipo9 | Importin-9 Mus musculus | 1449.63 | 33.72 | 116052 | 22 | 22 | 43797889 | 48878905 | 45943877 | 43513609 | 41303201 | 42467081 | 36107033 | 50419781 | 54053473 | 40134745 | 59078193 | 51752917 | 45524673 | 55024733 | 49172077 | 11 | 2 | 0.998857094 | 1.166109606 |
| Q3UQ44 | Iqgap2 | Ras GTPase-activating-like protein IQGAP2 Mus musculus | 196.71 | 2.6 | 180528 | 1 | 5 | 22371009 | 24136241 | 22486175 | 24295237 | 22065695 | 21492253 | 29924151 | 21181037 | 20584471 | 26895545 | 27185649 | 25627359 | 26377093 | 23646155 | 30108579 | 10 | 2 | 1.040944271 | 1.152490798 |
| Q922S8 | Kif2c | Kinesin-like protein KIF2C Mus musculus | 95.48 | 4.72 | 81085 | 2 | 2 | 242709.95 | 1 | 275403.44 | 1 | 402230.97 | 641076.12 | 495552.03 | 824558.69 | 536093.19 | 496042.81 | 886416.38 | 885364.88 | 568168.5 | 839148.69 | 272486.38 | 0 | 2 | 3.252387329 | 3.750310731 |
| O88653 | Lamtor3 | Ragulator complex protein LAMTOR3 Mus musculus | 54.18 | 8.87 | 13553 | 1 | 1 | 332034.62 | 235556.84 | 417342.41 | 1 | 1 | 374260.66 | 279995.5 | 450683.44 | 862488.25 | 491856.9625 | 449411.56 | 401068.88 | 423667.36 | 437494.56 | 406694.44 | 3 | 2 | 2.496898415 | 2.150735763 |
| Q6P4Z6 | Lcmt1 | Leucine carboxyl methyltransferase 1 Rattus norvegicus | 44.66 | 2.71 | 38330 | 1 | 1 | 2115761.75 | 2183631.25 | 1710164.75 | 2253409.25 | 1937010.12 | 1561821.62 | 1717521 | 1571419.62 | 1810043.12 | 1668137.88 | 1979706.62 | 1382952.62 | 1519326.25 | 1621406.12 | 1396004.88 | 2 | 2 | 0.816564894 | 0.774452373 |
| Q61233 | Lcp1 | Plastin-2 Mus musculus | 4110.059 | 87.88 | 70149 | 48 | 62 | 1483341441 | 1456825729 | 1599929345 | 1449602433 | 1517976193 | 1574370177 | 1592926593 | 1567863937 | 1712278529 | 1415732481 | 1862544897 | 1695475073 | 1503721217 | 1615226625 | 1941363969 | 8 | 2 | 1.047351087 | 1.147936161 |
| Q9Z0M5 | Lipa | Lysosomal acid lipase/cholesteryl ester hydrolase Mus musculus | 683.19 | 22.17 | 45325 | 9 | 9 | 14897863 | 18597939 | 18244205 | 13282737 | 12877767 | 27761769 | 31115139 | 26800337 | 32122465 | 31368847 | 17982701 | 19490887 | 18888273 | 18895353 | 19370957 | 0 | 2 | 1.914859801 | 1.214731069 |
| Q498T9 | Lrrc8c | Volume-regulated anion channel subunit LRRC8C Rattus norvegicus | 181.54 | 3.86 | 92463 | 2 | 2 | 285118.25 | 180492.56 | 295830.62 | 227927.3775 | 150268.08 | 772115.88 | 275593.34 | 342910.34 | 415650.625 | 271982.94 | 235665.42 | 847373.25 | 373225.41 | 1374425 | 695306.62 | 4 | 2 | 1.823609913 | 3.093964173 |
| P70202 | Lxn | Latexin Mus musculus | 92.6 | 9.46 | 25492 | 2 | 2 | 4335441.5 | 5021072 | 4927896 | 4757538.5 | 4461408.5 | 3962391.5 | 2843572.75 | 4782727 | 5097369.5 | 3215122 | 4308377.5 | 3982099.25 | 3797591 | 4557076 | 3241313.25 | 8 | 2 | 0.846737901 | 0.846111363 |
| Q9QYL8 | Lypla2 | Acyl-protein thioesterase 2 Rattus norvegicus | 40.87 | 10.82 | 24807 | 1 | 1 | 875456.56 | 942451.81 | 740642.75 | 577799.25 | 760094.81 | 957185.1575 | 1123751.25 | 1057006.75 | 944334.38 | 703648.25 | 928682.81 | 945914.44 | 1022898.44 | 880852.88 | 1012071 | 4 | 2 | 1.228280026 | 1.229433329 |
| O09159 | Man2b1 | Lysosomal alpha-mannosidase Mus musculus | 636.26 | 14.61 | 114648 | 10 | 10 | 29760767 | 29134727 | 24821189 | 32504589 | 31066853 | 23837261 | 17736253 | 27431651 | 22412547 | 18264985 | 23631789 | 21328147 | 27183449 | 28367575 | 20803459 | 1 | 2 | 0.744681195 | 0.823653767 |
| P21708 | Mapk3 | Mitogen-activated protein kinase 3 Rattus norvegicus | 285.39 | 25.53 | 43081 | 3 | 6 | 14046033 | 13884697 | 13244656 | 12927679 | 14114616 | 14595198 | 18821231 | 14158102 | 13997421 | 16445258 | 18213003 | 14374974 | 15016812 | 13967979 | 16860421 | 2 | 2 | 1.143650867 | 1.149748685 |
| Q8K310 | Matr3 | Matrin-3 Mus musculus | 768.83 | 23.4 | 94630 | 13 | 13 | 13998072 | 13357893 | 12686952 | 13257240 | 14324118 | 12190444 | 16045667 | 13828481 | 13780901 | 14444488 | 12504033 | 12692393 | 12517168 | 12534885 | 12869282 | 9 | 2 | 1.039419365 | 0.933359522 |
| Q9DCS3 | Mecr | Enoyl-[acyl-carrier-protein] reductase, mitochondrial Mus musculus | 190.64 | 23.59 | 40343 | 3 | 3 | 200994.44 | 270153.69 | 361610.19 | 359813.31 | 298142.9075 | 358415.09 | 1234498.5 | 264972.34 | 181679.12 | 1584490.75 | 607389.19 | 478681.69 | 329424.97 | 441094.78 | 1150039.75 | 11 | 2 | 2.431086374 | 2.01690552 |
| Q9CPX7 | Mrps16 | 28S ribosomal protein S16, mitochondrial Mus musculus | 38.09 | 5.93 | 15192 | 1 | 1 | 1601795.12 | 1679947.62 | 1821793.75 | 2012624.5 | 1772623.25 | 1265519.12 | 1292718.12 | 1539882.25 | 1866163.12 | 1924255 | 1616489 | 1371493.5 | 1597628 | 1363025.62 | 1617792.38 | 8 | 2 | 0.887470929 | 0.851233228 |
| Q9D0G0 | Mrps30 | 28S ribosomal protein S30, mitochondrial Mus musculus | 214.45 | 16.52 | 49939 | 4 | 4 | 1011809.5 | 96723.46 | 1106397.88 | 1256107.75 | 1277453.75 | 2111868.25 | 2451351.5 | 1135040.25 | 1273020 | 2426968.5 | 2553890.25 | 1687515.62 | 2055870 | 1117331.88 | 2717801 | 3 | 2 | 1.979206836 | 2.133815962 |
| Q5I0K8 | Mrps7 | 28S ribosomal protein S7, mitochondrial Rattus norvegicus | 71.51 | 4.96 | 28197 | 1 | 1 | 2196894.25 | 593319.44 | 1789828.5 | 3287881 | 1966980.798 | 749228.88 | 2454863.25 | 1904289.25 | 1432760.25 | 2076166.5 | 1 | 1449954.38 | 1 | 1 | 1426669.5 | 11 | 2 | 0.876196467 | 0.292491608 |
| P22366 | Myd88 | Myeloid differentiation primary response protein MyD88 Mus musculus | 127.41 | 15.54 | 33753 | 3 | 3 | 241400.28 | 381308.16 | 334468.31 | 323696.16 | 173338.16 | 675455.5 | 338819.88 | 748254.38 | 648797.44 | 146790.27 | 437649.84 | 326204.19 | 500238.94 | 724273.5 | 385349.03 | 6 | 2 | 1.759110161 | 1.632304656 |
| P70670 | Naca | Nascent polypeptide-associated complex subunit alpha, muscle-specific form Mus musculus | 450.4 | 3.2 | 220499 | 6 | 6 | 16618589 | 2569047.5 | 20120981 | 21405269 | 23384415 | 21061647 | 27509769 | 20355949 | 19788319 | 26388099 | 24760885 | 27221315 | 24861289 | 20408983 | 29466805 | 7 | 2 | 1.368681423 | 1.50679948 |
| Q8VBW6 | Nae1 | NEDD8-activating enzyme E1 regulatory subunit Mus musculus | 551.95 | 32.21 | 60274 | 10 | 10 | 1727273 | 6163646 | 5096017.5 | 7515592.5 | 7532802 | 7891399 | 7319317 | 15850521 | 6879275 | 6958896 | 9237315 | 7704216 | 6289983 | 8334077 | 8297414.5 | 6 | 2 | 1.601529442 | 1.421884603 |
| Q5PPG6 | Nap1l5 | Nucleosome assembly protein 1-like 5 Rattus norvegicus 2 | 35.42 | 4.52 | 16858 | 1 | 1 | 2030543 | 1892679.25 | 1967458.75 | 2355257 | 1713276.25 | 1562184.62 | 1458416.5 | 1537442.88 | 1685425.25 | 1486442.25 | 1721551.62 | 1277105.88 | 1430691.25 | 1837019.5 | 1297264.25 | 0 | 2 | 0.776156764 | 0.759460768 |
| P09405 | Ncl | Nucleolin Mus musculus | 2657.22 | 41.58 | 76723 | 41 | 43 | 512448801 | 493881793 | 545296833 | 505632737 | 517477665 | 522771777 | 505673281 | 590702081 | 596260609 | 534518209 | 627965249 | 647741057 | 518995009 | 544230337 | 680891457 | 5 | 2 | 1.068041152 | 1.172866253 |
| Q9D1J1 | Necap2 | Adaptin ear-binding coat-associated protein 2 Mus musculus | 270.8 | 31.95 | 28598 | 2 | 5 | 3710688 | 4054495.5 | 3515519.5 | 3933436.5 | 4115200 | 5116532 | 5677140.5 | 4120635.25 | 3991092.5 | 5271131 | 4688011 | 5291598 | 4013848.5 | 4674000 | 5776335.5 | 2 | 2 | 1.250768618 | 1.264595358 |
| P97333 | Nrp1 | Neuropilin-1 Mus musculus | 105.47 | 3.58 | 103000 | 2 | 2 | 2158467 | 1629235.62 | 1624815.75 | 1570022 | 1405104.25 | 702659 | 1167366.5 | 1577390.88 | 696628.5 | 1222015.88 | 1256722.88 | 1540028.25 | 1073999.75 | 1566660.38 | 1148377 | 2 | 2 | 0.639757763 | 0.785177312 |
| Q9R1J0 | Nsdhl | Sterol-4-alpha-carboxylate 3-dehydrogenase, decarboxylating Mus musculus | 330.15 | 25.97 | 40686 | 6 | 6 | 4963761 | 2714816.5 | 4315897 | 3331419.5 | 3430446 | 3590332 | 6767532.5 | 3682288 | 5342688.5 | 6153319.5 | 4070346.25 | 6350070.5 | 5142092 | 4998720 | 6530081 | 4 | 2 | 1.361468202 | 1.44438146 |
| P09103 | P4hb | Protein disulfide-isomerase Mus musculus | 1769.22 | 55.6 | 57058 | 28 | 28 | 136559073 | 139142049 | 125258673 | 127324713 | 149719969 | 171607425 | 190946033 | 169422401 | 153140801 | 160991313 | 152096065 | 148470625 | 157284705 | 157814065 | 142346193 | 0 | 2 | 1.247938622 | 1.118003905 |
| P08003 | Pdia4 | Protein disulfide-isomerase A4 Mus musculus | 1869.14 | 53.61 | 71982 | 32 | 32 | 53638141 | 41823233 | 54634405 | 44436601 | 43733833 | 50848017 | 64155573 | 57014573 | 52986641 | 50704353 | 55308045 | 60637233 | 45910577 | 55917929 | 62759317 | 6 | 2 | 1.157147518 | 1.177393544 |
| Q9CWM4 | Pfdn1 | Prefoldin subunit 1 Mus musculus | 61.53 | 9.02 | 14255 | 1 | 1 | 2782856.25 | 2875558.25 | 2645507 | 3131745.75 | 2722431 | 2910236.5 | 2586206 | 2909724.5 | 1865114.25 | 3186527 | 2148616.75 | 2516925.25 | 2740792 | 2574439 | 1983105.62 | 12 | 2 | 0.950537849 | 0.845020172 |
| Q03958 | Pfdn6 | Prefoldin subunit 6 Mus musculus | 58.42 | 9.45 | 14455 | 1 | 1 | 1665957 | 1816728.12 | 1824478.53 | 2297962.75 | 1517266.25 | 1382403.25 | 1530436.88 | 1854146.88 | 1597139.065 | 1621569.25 | 1187155.38 | 1751231.38 | 1405580.38 | 1418356 | 933757.81 | 7 | 2 | 0.875394826 | 0.734026829 |
| Q2TBE6 | Pi4k2a | Phosphatidylinositol 4-kinase type 2-alpha Mus musculus | 169.01 | 6.26 | 54258 | 3 | 3 | 2433654.75 | 1953824.25 | 2286650 | 2070547.25 | 2256479 | 1964162 | 1641800 | 2090900.25 | 1991144.25 | 1854194.25 | 1619286.5 | 1906954.25 | 1762642.5 | 1953715.38 | 2187275.5 | 4 | 2 | 0.867381701 | 0.857171262 |
| Q8BTI9 | Pik3cb | Phosphatidylinositol 4,5-bisphosphate 3-kinase catalytic subunit beta isoform Mus musculus | 54.26 | 1.5 | 121711 | 1 | 1 | 528539.31 | 442406.34 | 1 | 1 | 1 | 700645.56 | 1 | 697379.31 | 691684.5 | 1 | 558780.56 | 583713.38 | 424283.34 | 940180.88 | 614179.69 | 7 | 2 | 2.152236753 | 3.214524115 |
| P27612 | Plaa | Phospholipase A-2-activating protein Mus musculus | 763.86 | 26.45 | 87221 | 14 | 14 | 6908505 | 8042811 | 8340288 | 9604593 | 11049423 | 11503360 | 11235243 | 11187209 | 8149116 | 13860019 | 12879563 | 12492611 | 11326172 | 9749513 | 10956633 | 3 | 2 | 1.272821888 | 1.306261966 |
| P43883 | Plin2 | Perilipin-2 Mus musculus | 38.59 | 3.76 | 46646 | 1 | 1 | 1331561.88 | 1921657.12 | 2026767.25 | 1684054.12 | 1694100.12 | 1281032.88 | 1414913.5 | 1548605.38 | 1822210.62 | 1405844.12 | 1271895.5 | 1028179.5 | 1673374.62 | 1635060.25 | 1130297.38 | 6 | 2 | 0.863072909 | 0.778320386 |
| Q9R0E2 | Plod1 | Procollagen-lysine,2-oxoglutarate 5-dioxygenase 1 Mus musculus | 218.15 | 8.79 | 83595 | 5 | 5 | 1740708.88 | 996182.12 | 2236154.5 | 505680.28 | 1145521.25 | 1213544.5 | 1967818 | 2080649.38 | 2748213.5 | 1073571 | 2368922.75 | 2646561.25 | 1286637.5 | 2373125.75 | 2656902.5 | 7 | 2 | 1.371294932 | 1.710707602 |
| Q6ZQB6 | Ppip5k2 | Inositol hexakisphosphate and diphosphoinositol-pentakisphosphate kinase 2 Mus musculus | 141.43 | 4.07 | 128427 | 3 | 3 | 1313955.12 | 1195377.88 | 756056.56 | 1340971.5 | 1055065.38 | 558345.19 | 984502.94 | 717093.25 | 233355.45 | 968674.94 | 895340.44 | 684509.46 | 769825.56 | 609484.75 | 463387.09 | 2 | 2 | 0.611501678 | 0.604537979 |
| Q4QQT4 | Ppp2r1b | Serine/threonine-protein phosphatase 2A 65 kDa regulatory subunit A beta isoform Rattus norvegicus | 933.86 | 29.78 | 66006 | 4 | 13 | 39195481 | 41542317 | 44276041 | 40827941 | 41004665 | 48668225 | 39258677 | 46801641 | 48819097 | 43445749 | 49151237 | 44037349 | 44210489 | 45610553 | 48188337 | 5 | 2 | 1.097400485 | 1.117727525 |
| Q5XIS9 | Prkd2 | Serine/threonine-protein kinase D2 Rattus norvegicus | 73.48 | 1.26 | 96497 | 1 | 1 | 1 | 143415.03 | 1 | 1 | 144991.11 | 161520.64 | 153071.89 | 237913.2 | 235054.06 | 145472.48 | 220041.52 | 193733.435 | 142557.83 | 218713.11 | 193621.28 | 0 | 2 | 3.235099519 | 3.3586563 |
| Q5RKH1 | Prpf4b | Serine/threonine-protein kinase PRP4 homolog Rattus norvegicus | 51.07 | 1.69 | 117007 | 1 | 1 | 1424606.88 | 1689704.62 | 1048682.38 | 1515060.25 | 1387228.25 | 1146401 | 830589 | 1064548.62 | 1440622.75 | 1162901.75 | 841158.12 | 543436.88 | 1413520.12 | 973611.88 | 782595.94 | 6 | 2 | 0.7989862 | 0.644605933 |
| Q9R1P1 | Psmb3 | Proteasome subunit beta type-3 Mus musculus | 493.69 | 53.17 | 22965 | 8 | 8 | 38088821 | 39792409 | 41585749 | 38853289 | 41475689 | 42663561 | 43969641 | 40239129 | 43768621 | 44703805 | 42588357 | 41988505 | 45713765 | 41178841 | 42573357 | 2 | 2 | 1.077823397 | 1.071307089 |
| O35295 | Purb | Transcriptional activator protein Pur-beta Mus musculus | 225.25 | 24.07 | 33901 | 5 | 5 | 3858990.5 | 4825599 | 3841536.75 | 4611350 | 4764654.5 | 5250101 | 6905895.5 | 5837604 | 3114281.5 | 4755829.5 | 6213444.5 | 5672742 | 4670730.5 | 4834612.5 | 8436056 | 7 | 2 | 1.1808765 | 1.361857704 |
| Q91V41 | Rab14 | Ras-related protein Rab-14 Mus musculus | 815.37 | 80 | 23897 | 12 | 14 | 25779797 | 25251989 | 26092057 | 26823629 | 27031935 | 26280155 | 29128297 | 25126925 | 28368967 | 28239017 | 30238873 | 27247043 | 27063093 | 26129909 | 30110257 | 7 | 2 | 1.047060482 | 1.074895499 |
| P09527 | Rab7a | Ras-related protein Rab-7a Rattus norvegicus | 1025.46 | 68.12 | 23504 | 14 | 15 | 97731393 | 109525561 | 103554041 | 103969705 | 107382417 | 102180417 | 89586025 | 105958585 | 107245825 | 97257081 | 83996561 | 95928697 | 102199545 | 103322585 | 97441481 | 7 | 2 | 0.961821922 | 0.92478548 |
| Q8BX09 | Rbbp5 | Retinoblastoma-binding protein 5 Mus musculus | 57.48 | 1.67 | 59098 | 1 | 1 | 1099447.25 | 1230452.38 | 1205071 | 1385181.62 | 1310407.38 | 1110323.62 | 755671.19 | 1103980.75 | 1415510.5 | 1023861.12 | 1105930.5 | 948955.94 | 908911.62 | 1153794.5 | 879509.94 | 7 | 2 | 0.868196037 | 0.802031085 |
| Q0VBL3 | Rbm15 | RNA-binding protein 15 Mus musculus | 179.31 | 5.2 | 105722 | 4 | 4 | 1143650.88 | 1253638.5 | 1074182.25 | 1482568.12 | 1370062.12 | 1652140.75 | 1114756.75 | 988258.5 | 650327.94 | 1688307.5 | 745620.5 | 1226603.75 | 644566.81 | 366201.38 | 997105.62 | 11 | 2 | 0.963582113 | 0.629353882 |
| P14115 | Rpl27a | 60S ribosomal protein L27a Mus musculus | 371 | 33.11 | 16605 | 7 | 7 | 40037089 | 42905105 | 45102537 | 44705289 | 43530089 | 39790725 | 44296481 | 40801053 | 41524761 | 42693853 | 27935301 | 38935337 | 39817385 | 39647617 | 43217513 | 6 | 2 | 0.966833584 | 0.876424346 |
| P07742 | Rrm1 | Ribonucleoside-diphosphate reductase large subunit Mus musculus | 647.75 | 26.64 | 90210 | 14 | 14 | 18246387 | 12284210 | 5937449 | 16716314 | 16616034 | 9535447 | 4218572.5 | 4721790 | 7043884.5 | 4911893.5 | 5650169 | 4992719 | 5052815.5 | 8698067 | 9365249 | 2 | 2 | 0.435980168 | 0.483650844 |
| Q9WTM5 | Ruvbl2 | RuvB-like 2 Mus musculus | 1156.47 | 54.64 | 51113 | 21 | 21 | 159727153 | 116194353 | 119336737 | 173795409 | 156698625 | 109794737 | 136161889 | 102835385 | 67733913 | 153292465 | 139777217 | 71474217 | 107363233 | 89141105 | 108427185 | 4 | 2 | 0.785141717 | 0.711238495 |
| P05964 | S100a6 | Protein S100-A6 Rattus norvegicus | 373.2 | 64.04 | 10035 | 5 | 5 | 147885137 | 153946993 | 109438433 | 153265377 | 135110529 | 130598289 | 117486065 | 120629009 | 97620825 | 132451969 | 108503449 | 87248649 | 130402329 | 117599337 | 84515473 | 4 | 2 | 0.855841033 | 0.755051673 |
| Q8CFE4 | Scyl2 | SCY1-like protein 2 Mus musculus | 80.93 | 2.69 | 103317 | 2 | 2 | 185693.36 | 540642.88 | 685600.88 | 205727.7 | 206164.47 | 597248.5 | 255295.67 | 1101223.5 | 1218457.12 | 206922.92 | 720390.12 | 973893.62 | 283078.81 | 854196.25 | 770933.62 | 5 | 2 | 1.852776314 | 1.975235533 |
| O08992 | Sdcbp | Syntenin-1 Mus musculus | 423.87 | 53.18 | 32379 | 7 | 7 | 1300222.12 | 1207490.5 | 2344484 | 801476.5 | 1769486.5 | 1932388.88 | 3215072 | 1519113.38 | 1112105.38 | 3408814 | 2434432 | 2705793.25 | 2373426.25 | 1328876 | 3378984 | 7 | 2 | 1.507106706 | 1.646402897 |
| Q99NB9 | Sf3b1 | Splicing factor 3B subunit 1 Mus musculus | 985.69 | 23.54 | 145816 | 21 | 21 | 34263065 | 14947607 | 13554555 | 16902905 | 15601250 | 12576204 | 11827910 | 12929530 | 12649411 | 16618713 | 15566746 | 12770150 | 12598195 | 12158316 | 12078774 | 3 | 2 | 0.699088906 | 0.684083172 |
| Q8R216 | Sirt4 | NAD-dependent protein lipoamidase sirtuin-4, mitochondrial Mus musculus | 51.98 | 3.6 | 37525 | 1 | 1 | 472245.59 | 448417.03 | 421806.75 | 477248.72 | 454929.5225 | 456790.41 | 511965.12 | 479036.81 | 571783.94 | 456906.81 | 235662.91 | 386782.44 | 370353.0325 | 426299.44 | 432667.34 | 4 | 2 | 1.088732636 | 0.814088808 |
| Q8CI08 | Slain2 | SLAIN motif-containing protein 2 Mus musculus | 48.31 | 2.07 | 62378 | 1 | 1 | 1065192.75 | 797507.75 | 527056.31 | 1317345.38 | 1272489.5 | 648170.75 | 492699.72 | 1 | 1 | 857958 | 792764.94 | 1 | 623578.38 | 1 | 1 | 3 | 2 | 0.401404491 | 0.284430212 |
| O09044 | Snap23 | Synaptosomal-associated protein 23 Mus musculus | 89.9 | 6.67 | 23261 | 1 | 1 | 2179954.25 | 2663049.25 | 2291606.75 | 2343632.25 | 1942696.5 | 2121699.25 | 1809910.25 | 2704765.25 | 2287853.5 | 2034623.75 | 1416420.25 | 1816306.25 | 2151266.25 | 1559292.38 | 2093848.62 | 9 | 2 | 0.959540367 | 0.791277648 |
| Q5BK61 | Snx20 | Sorting nexin-20 Rattus norvegicus | 45.89 | 3.83 | 35723 | 1 | 1 | 781292.69 | 719573.88 | 749767.56 | 606141.81 | 766262.44 | 501851.78 | 613557.06 | 646920.75 | 816417.94 | 751198.75 | 555807.88 | 516052.69 | 586433 | 729454.69 | 591070.69 | 10 | 2 | 0.919103231 | 0.822188075 |
| Q58A65 | Spag9 | C-Jun-amino-terminal kinase-interacting protein 4 Mus musculus | 403.93 | 9.01 | 146219 | 7 | 7 | 4373358.5 | 4123626 | 3886400 | 3164691 | 5128186 | 5395138.5 | 5981896 | 5632487.5 | 6745488 | 5882954 | 4880454 | 5267936.5 | 4656879 | 5219346.5 | 6306012.5 | 0 | 2 | 1.433429539 | 1.27347144 |
| P48428 | Tbca | Tubulin-specific chaperone A Mus musculus | 224.12 | 56.48 | 12758 | 5 | 5 | 9131711 | 9397210 | 8753151 | 10509868 | 8785775 | 7585664 | 6202783 | 8991456 | 8507587 | 8926501 | 9005037 | 7408565.5 | 8686178 | 8077935.5 | 7330137 | 4 | 2 | 0.863374062 | 0.869683131 |
| Q9CR02 | Tma16 | Translation machinery-associated protein 16 Mus musculus | 97.78 | 17.19 | 25782 | 2 | 2 | 2082810.12 | 1679686.38 | 1689868.88 | 2088082.88 | 2318952.25 | 1 | 1882131.25 | 709378.19 | 1 | 1 | 1672875.62 | 1105107.5 | 2073011.38 | 746404.31 | 1399349.703 | 2 | 2 | 0.262846857 | 0.70965253 |
| Q5I0E7 | Tmed9 | Transmembrane emp24 domain-containing protein 9 Rattus norvegicus | 225.67 | 20 | 27028 | 4 | 5 | 27894041 | 27263373 | 25066293 | 24434101 | 24617893 | 26948341 | 23535077 | 25655101 | 27429665 | 23196233 | 22401157 | 22268675 | 21987895 | 24936635 | 19726717 | 10 | 2 | 0.980574199 | 0.861113714 |
| B0BN86 | Tmem11 | Transmembrane protein 11, mitochondrial Rattus norvegicus 2 | 39.37 | 4.21 | 21342 | 1 | 1 | 1 | 841602.56 | 1191889.12 | 755534.25 | 881622.19 | 1110277.25 | 878418.56 | 1512629.88 | 1728173.62 | 871786.75 | 1276877.75 | 1314066.12 | 922817.44 | 1309639.88 | 910818.19 | 5 | 2 | 1.662181772 | 1.562181291 |
| Q61510 | Trim25 | E3 ubiquitin/ISG15 ligase TRIM25 Mus musculus | 42.96 | 1.89 | 71726 | 1 | 1 | 1870693.75 | 1544405.88 | 1901792.25 | 1853437.5 | 1749071.25 | 1716854.38 | 1605197.88 | 1723985.12 | 1649199.25 | 1760053.38 | 1399626.5 | 1408073.75 | 1631595 | 1201017 | 1578895.12 | 6 | 2 | 0.947966165 | 0.809382566 |
| Q9QZE7 | Tsnax | Translin-associated protein X Mus musculus | 385.33 | 42.41 | 32926 | 5 | 8 | 2525334 | 1737998.12 | 2549686 | 2309082 | 2140633.75 | 2335023.75 | 2242932.25 | 2116349.75 | 1973180.62 | 1934555.25 | 1695069.25 | 1564361.88 | 1779394.5 | 1663456.62 | 1869812 | 8 | 2 | 0.941338199 | 0.761102442 |
| Q3UX10 | Tubal3 | Tubulin alpha chain-like 3 Mus musculus 2 | 144.57 | 11.66 | 49988 | 1 | 3 | 18683813 | 24827005 | 17155457 | 15835367 | 20305229 | 23150319 | 30874673 | 18547001 | 28414065 | 23256077 | 32788669 | 20666781 | 25526539 | 30867749 | 24560469 | 5 | 2 | 1.283402032 | 1.388436643 |
| P26369 | U2af2 | Splicing factor U2AF 65 kDa subunit Mus musculus | 826.54 | 42.74 | 53517 | 12 | 12 | 25919089 | 30357559 | 35058081 | 34916045 | 29659301 | 28332425 | 33459787 | 35408569 | 40334525 | 36172917 | 36312177 | 42281941 | 34605001 | 35972701 | 37405653 | 6 | 2 | 1.114156497 | 1.19669927 |
| O08759 | Ube3a | Ubiquitin-protein ligase E3A Mus musculus | 266.1 | 7.47 | 99819 | 5 | 5 | 27386849 | 20405337 | 25101607 | 27857019 | 29841137 | 33784813 | 36036953 | 32921333 | 21070223 | 36375017 | 39138753 | 34748021 | 38388857 | 27940439 | 29675299 | 4 | 2 | 1.226632577 | 1.300932946 |
| Q9ES00 | Ube4b | Ubiquitin conjugation factor E4 B Mus musculus | 102.79 | 4.01 | 133318 | 3 | 3 | 439971 | 962346 | 1530228.12 | 872385.12 | 728183.56 | 1265487.38 | 804700.25 | 1332053.5 | 1598718.88 | 939348.5 | 1753108 | 1543893.62 | 1069739.5 | 1625496 | 1515047 | 7 | 2 | 1.310425631 | 1.656098755 |
| Q99KD5 | Unc45a | Protein unc-45 homolog A Mus musculus | 119.85 | 4.24 | 103447 | 2 | 2 | 1 | 1 | 271007.47 | 93428.48 | 1 | 1 | 63461.32 | 283832.47 | 228165.52 | 1 | 253575.97 | 355847.19 | 276605.485 | 295359.03 | 201639.75 | 10 | 2 | 1.579033498 | 3.794949538 |
| Q9EPU0 | Upf1 | Regulator of nonsense transcripts 1 Mus musculus | 769.85 | 16.64 | 123967 | 15 | 15 | 4513688.5 | 4762802 | 4437975 | 5356068 | 5269039.5 | 4200129.5 | 7651897.5 | 4579007 | 3328904.75 | 5869676.5 | 7074408 | 7934471 | 5645362 | 4980795.5 | 7717048 | 12 | 2 | 1.053001844 | 1.370282235 |
| Q8BH43 | Wasf2 | Wiskott-Aldrich syndrome protein family member 2 Mus musculus | 520.27 | 21.33 | 54074 | 9 | 9 | 7733303.5 | 8904244 | 9569390 | 7331849 | 11301141 | 10284164 | 11763803 | 8315744.5 | 11253333 | 12061459 | 13948751 | 10160071 | 12063375 | 10305633 | 11377461 | 5 | 2 | 1.197113967 | 1.290262813 |
| Q6NZF1 | Zc3h11a | Zinc finger CCCH domain-containing protein 11A Mus musculus | 217.47 | 8.33 | 86492 | 5 | 5 | 1478486.5 | 1571028 | 1751738.25 | 1529590 | 1550828.5 | 439925.28 | 1228319 | 501807.97 | 1153117.12 | 1342855.25 | 1137574.62 | 1445505.75 | 1545351.75 | 605430.94 | 1186179.25 | 0 | 2 | 0.592009546 | 0.751115103 |
| Q3TIV5 | Zc3h15 | Zinc finger CCCH domain-containing protein 15 Mus musculus | 202.4 | 9.62 | 48327 | 3 | 3 | 379547.72 | 478608.06 | 597786.88 | 451137.66 | 644188.06 | 1178483.62 | 950049.62 | 682897.75 | 816810.62 | 648497.94 | 1054115.88 | 724461.31 | 645336.62 | 532725.44 | 1222010.12 | 0 | 2 | 1.676318957 | 1.637871344 |
| Q80YR4 | Znf598 | E3 ubiquitin-protein ligase ZNF598 Mus musculus | 36.77 | 1.54 | 99192 | 1 | 1 | 669236.88 | 730088.88 | 643516.69 | 1071744.75 | 744059.88 | 495694.12 | 682799.31 | 683198.31 | 474349.41 | 693498.19 | 686416.5 | 494240.09 | 442985.97 | 536930.94 | 613733.69 | 6 | 2 | 0.785129937 | 0.718984435 |

Supplementary Table 3: Results of the pathway analysis by the David tool on the proteins modulated in response to PET beads

| Annotation Cluster 1 | Enrichment Score: 8.084268709806619 | | | | | | | |
|---|---|---|---|---|---|---|---|---|
| Category | Term | Count | % | PValue | Genes | Fold Enrichment | Benjamini | FDR |
| **GOTERM_CC_DIRECT** | **GO:0005739~mitochondrion** | **49** | **19.44444** | **1.096050598E-09** | **P29452, P19096, P38647, Q9R112, Q9CPW3, Q9DBL1, P36552, P62075, Q8BTZ7, Q8BGH2, Q9Z2Y8, P41216, Q8BH59, Q9CZR8, Q61207, Q8JZQ2, Q99MN1, Q9DCM0, Q07813, O55028, P06151, Q99KI0, Q9Z1T1, Q922Q1, Q9EQI8, Q811U4, P29594, Q9QXX4, Q62465, Q922D8, P97807, Q9CZD3, Q8BIJ6, Q9QUJ7, Q9JM90, Q8CGK3, Q9QYB1, P97450, Q9Z1Q9, P97333, Q791V5, P53395, Q8BYL4, Q99L13, P16332, Q78IK4, Q9JK81, P29758, P07742** | **2.5843090054** | **1.42E-07** | **1.36E-07** |
| UP_SEQ_FEATURE | TRANSIT:Mitochondrion | 24 | 9.52381 | 1.463698457E-09 | P97450, P53395, P38647, Q8JZQ2, Q9R112, Q9CPW3, Q8BYL4, Q99L13, Q9DBL1, P97807, Q9DCM0, P16332, P36552, O55028, Q9CZD3, Q8BIJ6, Q99KI0, Q78IK4, Q922Q1, P29758, Q9JK81, Q9CZR8, Q8CGK3, Q9EQI8 | 4.7520695695 | 1.18E-06 | 1.17E-06 |
| UP_KW_DOMAIN | KW-0809~Transit peptide | 24 | 9.52381 | 2.006978989E-09 | P97450, P53395, P38647, Q8JZQ2, Q9R112, Q9CPW3, Q8BYL4, Q99L13, Q9DBL1, P97807, Q9DCM0, P16332, P36552, O55028, Q9CZD3, Q8BIJ6, Q99KI0, Q78IK4, Q922Q1, P29758, Q9JK81, Q9CZR8, Q8CGK3, Q9EQI8 | 4.56368758 | 4.62E-08 | 4.42E-08 |
| UP_KW_CELLULAR_COMPONENT | KW-0496~Mitochondrion | 37 | 14.68254 | 7.536299405E-08 | Q811U4, P38647, Q9R112, Q9QXX4, Q9CPW3, Q62465, Q9DBL1, P97807, P36552, Q9CZD3, Q8BIJ6, P62075, Q9QUJ7, Q8BGH2, Q9JM90, P41216, Q8BH59, Q9CZR8, Q8CGK3, Q9QYB1, P97450, P97333, Q791V5, P53395, Q8JZQ2, Q8BYL4, Q99L13, Q9DCM0, P16332, Q07813, O55028, Q99KI0, Q78IK4, Q922Q1, P29758, Q9JK81, Q9EQI8 | 2.692606103 | 1.43E-06 | 1.39E-06 |
| GOTERM_CC_DIRECT | GO:0005759~mitochondrial matrix | 15 | 5.952381 | 1.561972067E-07 | P53395, P38647, Q8BYL4, Q9DBL1, P97807, Q9DCM0, P16332, O55028, Q8BIJ6, Q99KI0, P29758, Q9JK81, P24860, Q9CZR8, Q8CGK3 | 6.1874197689 | 1.52E-05 | 1.46E-05 |

| Annotation Cluster 2 | Enrichment Score: 5.205935381326194 | | | | | | | |
|---|---|---|---|---|---|---|---|---|
| Category | Term | Count | % | PValue | Genes | Fold Enrichment | Benjamini | FDR |
| GOTERM_MF_DIRECT | GO:0000166~nucleotide binding | 47 | 18.65079 | 7.859667861E-09 | Q811U4, Q6P9P6, Q8R4K2, Q61656, Q6PHZ2, Q9CY64, Q501J6, P38647, Q9R0N0, Q922D8, P12382, Q8VEH6, Q9JKR6, Q9CZD3, Q8BIJ6, Q62095, Q9QVP9, Q9QUJ7, P61027, Q8BTZ7, P42208, Q8BU30, P41216, Q8CGK3, Q9D0F6, Q9ESL4, Q9Z1Q9, Q8JZQ2, Q9JHG7, Q99MN1, Q6PA06, Q6ZQB6, Q8BYL4, Q9ESV0, Q05144, Q61024, Q3V300, P58252, O55028, P83741, Q5NC05, E9Q3L2, Q9WUA3, E9Q634, Q9DBY8, Q922S8, P07742 | 2.4930428991 | 2.11E-06 | 2.08E-06 |
| GOTERM_MF_DIRECT | GO:0005524~ATP binding | 38 | 15.07937 | 3.716203084E-06 | Q6P9P6, Q8R4K2, Q61656, Q6PHZ2, Q501J6, P38647, Q9R0N0, Q922D8, P12382, Q9JKR6, Q9CZD3, Q8BIJ6, Q62095, Q9QVP9, Q9QUJ7, Q8BU30, P41216, Q8CGK3, Q9D0F6, Q9ESL4, Q9Z1Q9, Q8JZQ2, Q9JHG7, Q99MN1, Q6ZQB6, Q8BYL4, Q9ESV0, Q61024, Q3V300, O55028, P83741, Q5NC05, E9Q3L2, Q9WUA3, E9Q634, Q9DBY8, Q922S8, P07742 | 2.2584222561 | 0.00062 | 0.000612 |
| UP_KW_LIGAND | KW-0547~Nucleotide-binding | 46 | 18.25397 | 9.943995452E-06 | Q811U4, Q6P9P6, Q8R4K2, Q61656, Q6PHZ2, Q501J6, P38647, Q9R0N0, Q922D8, P12382, Q8VEH6, Q9JKR6, Q9CZD3, Q8BIJ6, Q62095, Q9QVP9, Q9QUJ7, P61027, Q8BTZ7, P42208, Q8BU30, P41216, Q8CGK3, Q9D0F6, Q9ESL4, Q9Z1Q9, Q8JZQ2, Q9JHG7, Q99MN1, Q6PA06, Q6ZQB6, Q8BYL4, Q9ESV0, Q05144, Q61024, Q3V300, P58252, O55028, P83741, Q5NC05, E9Q3L2, Q9WUA3, E9Q634, Q9DBY8, Q922S8, P07742 | 1.7954978253 | 0.000229 | 0.000229 |

| | | | | | | | | |
|---|---|---|---|---|---|---|---|---|
| UP_KW_LIGAND | KW-0067~ATP-binding | 38 | 15.07937 | 2.909238643E-05 | Q6P9P6, Q8R4K2, Q61656, Q6PHZ2, Q501J6, P38647, Q9R0N0, Q922D8, P12382, Q9JKR6, Q9CZD3, Q8BIJ6, Q62095, Q9QVP9, Q9QUJ7, Q8BU30, P41216, Q8CGK3, Q9D0F6, Q9ESL4, Q9Z1Q9, Q8JZQ2, Q9JHG7, Q99MN1, Q6ZQB6, Q8BYL4, Q9ESV0, Q61024, Q3V300, O55028, P83741, Q5NC05, E9Q3L2, Q9WUA3, E9Q634, Q9DBY8, Q922S8, P07742 | 1.9148784411 | 0.000335 | 0.000335 |
| Annotation Cluster 3 | Enrichment Score: 3.286924098455261 | | | | | | | |
| Category | Term | Count | % | PValue | Genes | Fold Enrichment | Benjamini | FDR |
| GOTERM_MF_DIRECT | GO:0016874~ligase activity | 10 | 3.968254 | 2.200147548E-05 | Q9Z1Q9, Q9CZD3, Q8BIJ6, Q9QUJ7, Q99MN1, Q8BYL4, Q8BU30, P41216, Q61024, Q922D8 | 6.5674679768 | 0.002359 | 0.002328 |
| GOTERM_BP_DIRECT | GO:0006418~tRNA aminoacylation for protein translation | 6 | 2.380952 | 2.573571681E-05 | Q9Z1Q9, Q9CZD3, Q8BIJ6, Q99MN1, Q8BYL4, Q8BU30 | 16.843273232 | 0.042181 | 0.042104 |
| UP_KW_BIOLOGICAL_PROCESS | KW-0648~Protein biosynthesis | 10 | 3.968254 | 2.974183792E-05 | Q9Z1Q9, P58252, Q9CZD3, Q8BIJ6, Q99MN1, Q9CQJ6, Q8BYL4, Q8BU30, Q9CZR8, Q8BX17 | 6.2327665441 | 0.002409 | 0.002409 |
| UP_KW_MOLECULAR_FUNCTION | KW-0436~Ligase | 10 | 3.968254 | 3.81245603E-05 | Q9Z1Q9, Q9CZD3, Q8BIJ6, Q9QUJ7, Q99MN1, Q8BYL4, Q8BU30, P41216, Q61024, Q922D8 | 6.0582582885 | 0.002554 | 0.00244 |
| GOTERM_MF_DIRECT | GO:0004812~aminoacyl-tRNA ligase activity | 6 | 2.380952 | 8.981608684E-05 | Q9Z1Q9, Q9CZD3, Q8BIJ6, Q99MN1, Q8BYL4, Q8BU30 | 13.041114983 | 0.008024 | 0.007919 |
| UP_KW_MOLECULAR_FUNCTION | KW-0030~Aminoacyl-tRNA synthetase | 6 | 2.380952 | 0.0001002971606 | Q9Z1Q9, Q9CZD3, Q8BIJ6, Q99MN1, Q8BYL4, Q8BU30 | 12.631468531 | 0.003055 | 0.002918 |
| Annotation Cluster 4 | Enrichment Score: 2.8148018004467232 | | | | | | | |
| Category | Term | Count | % | PValue | Genes | Fold Enrichment | Benjamini | FDR |
| **KEGG_PATHWAY** | **mmu00052:Galactose metabolism** | **6** | **2.380952** | **8.875502898E-05** | **P12382, Q7TSV4, Q9R0N0, P15535, Q9WUA3, Q9D0F9** | **12.794444444** | **0.006952** | **0.006775** |
| Annotation Cluster 5 | Enrichment Score: 2.468622798837743 | | | | | | | |
| Category | Term | Count | % | PValue | Genes | Fold Enrichment | Benjamini | FDR |
| **KEGG_PATHWAY** | **mmu04142:Lysosome** | **11** | **4.365079** | **2.55225983E-05** | **O09159, O35657, O35643, Q9Z0M5, Q61207, Q8R0H9, O08585, P97821, Q9Z1T1, P17439, Q80V94** | **5.5600548697** | **0.003401** | **0.003314** |
| GOTERM_CC_DIRECT | GO:0043202~lysosomal lumen | 4 | 1.587302 | 0.0022863288204 | O09159, O35657, P17439, O35405 | 15.033138402 | 0.07943 | 0.076163 |
| Annotation Cluster 6 | Enrichment Score: 2.203279698489198 | | | | | | | |
| Category | Term | Count | % | PValue | Genes | Fold Enrichment | Benjamini | FDR |
| GOTERM_BP_DIRECT | GO:0008152~metabolic process | 10 | 3.968254 | 0.0001260217872 | O09159, P12382, O35657, P08905, P19096, P17439, Q9WUA3, Q8BHN3, P07742, Q922D8 | 5.2541404911 | 0.06885 | 0.068724 |
| Annotation Cluster 7 | Enrichment Score: 2.178086833780126 | | | | | | | |
| Category | Term | Count | % | PValue | Genes | Fold Enrichment | Benjamini | FDR |
| GOTERM_BP_DIRECT | GO:0006886~intracellular protein transport | 13 | 5.15873 | 6.938776954E-05 | Q61207, Q8R0H9, Q8K1E0, Q80V94, Q8VBT9, Q99LG2, O35643, Q9CWK8, A2AWA9, O08585, P61027, Q9Z1T1, O88983 | 4.1878083718 | 0.056863 | 0.056759 |
| Annotation Cluster 8 | Enrichment Score: 2.172411997062546 | | | | | | | |

| Category | Term | Count | % | PValue | Genes | Fold Enrichment | Benjamini | FDR |
|---|---|---|---|---|---|---|---|---|
| KEGG_PATHWAY | mmu00010:Glycolysis / Gluconeogenesis | 6 | 2.380952 | 0.0028355897471 | P12382, Q7TSV4, P06151, P17182, Q9WUA3, Q9D0F9 | 6.1107794362 | 0.095195 | 0.092764 |
| Annotation Cluster 9 | Enrichment Score: 2.164392962846174 | | | | | | | |
| Category | Term | Count | % | PValue | Genes | Fold Enrichment | Benjamini | FDR |
| GOTERM_CC_DIRECT | GO:0005783~endoplasmic reticulum | 31 | 12.30159 | 0.0011434516429 | Q6PHZ2, Q8BYI6, Q8R2E9, P70188, P97821, Q9CPU4, P70227, P45878, Q9JKR6, O88736, Q9WUD1, Q9QUJ7, P61027, Q9D0F3, Q8K1T1, P41216, Q6Y7W8, Q8BHN3, O35405, Q8K1E0, P35564, Q6PA06, P17439, Q99P72, Q8R180, O55242, Q07813, Q8BMK4, P09103, Q8CCJ3, Q80WJ7 | 1.8580532844 | 0.049423 | 0.04739 |
| Annotation Cluster 10 | Enrichment Score: 2.0325527140574406 | | | | | | | |
| Category | Term | Count | % | PValue | Genes | Fold Enrichment | Benjamini | FDR |
| GOTERM_BP_DIRECT | GO:0043488~regulation of mRNA stability | 5 | 1.984127 | 0.0003735249856 | Q91YT7, Q8BYK6, Q8BFV2, P35922, Q9WVR4 | 14.448886351 | 0.102035 | 0.101848 |
| GOTERM_BP_DIRECT | GO:0061157~mRNA destabilization | 4 | 1.587302 | 0.0013680122612 | Q91YT7, Q8BYK6, Q9WVR4, Q6Y7W8 | 17.86407767 | 0.24435 | 0.243903 |
| GOTERM_MF_DIRECT | GO:1990247~N6-methyladenosine-containing RNA binding | 3 | 1.190476 | 0.0060960131016 | Q91YT7, Q8BYK6, P35922 | 24.896674058 | 0.261846 | 0.258426 |
| GOTERM_CC_DIRECT | GO:0010494~cytoplasmic stress granule | 5 | 1.984127 | 0.0128418905374 | Q91YT7, Q8BYK6, P35922, Q9WVR4, Q6Y7W8 | 5.5148741419 | 0.227068 | 0.217728 |
| GOTERM_BP_DIRECT | GO:0034063~stress granule assembly | 3 | 1.190476 | 0.0303499304492 | Q91YT7, Q8BYK6, P35922 | 10.916936354 | 0.931594 | 0.929889 |
| GOTERM_CC_DIRECT | GO:0036464~cytoplasmic ribonucleoprotein granule | 4 | 1.587302 | 0.0541389005759 | Q91YT7, Q8BMK4, P35922, Q9WVR4 | 4.6654567453 | 0.5497 | 0.52709 |
| GOTERM_MF_DIRECT | GO:0003729~mRNA binding | 7 | 2.777778 | 0.0900283656799 | Q91YT7, Q8BYK6, Q9CQJ6, Q6NZF1, P35922, Q9WVR4, Q05CL8 | 2.2580022408 | 0.8617 | 0.850447 |
| Annotation Cluster 11 | Enrichment Score: 1.9697081061111987 | | | | | | | |
| Category | Term | Count | % | PValue | Genes | Fold Enrichment | Benjamini | FDR |
| KEGG_PATHWAY | mmu04141:Protein processing in endoplasmic reticulum | 10 | 3.968254 | 0.0010010670018 | Q9JKR6, Q07813, Q8BMK4, Q9WUD1, Q8R2E9, P09103, Q9D0F3, P35564, Q8R180, Q8BHN3 | 3.8992592593 | 0.058813 | 0.057311 |
| Annotation Cluster 12 | Enrichment Score: 1.6319566730587778 | | | | | | | |
| Category | Term | Count | % | PValue | Genes | Fold Enrichment | Benjamini | FDR |
| KEGG_PATHWAY | mmu00010:Glycolysis / Gluconeogenesis | 6 | 2.380952 | 0.0028355897471 | P12382, Q7TSV4, P06151, P17182, Q9WUA3, Q9D0F9 | 6.1107794362 | 0.095195 | 0.092764 |
| Annotation Cluster 13 | Enrichment Score: 1.5609646193649531 | | | | | | | |
| Category | Term | Count | % | PValue | Genes | Fold Enrichment | Benjamini | FDR |
| UP_KW_CELLULAR_COMPONENT | KW-1000~Mitochondrion outer membrane | 8 | 3.174603 | 0.0011034979764 | Q811U4, Q07813, Q791V5, Q9QUJ7, Q8BGH2, Q922Q1, P41216, Q62465 | 5.0083869164 | 0.013978 | 0.01361 |

| Annotation Cluster 14 | Enrichment Score: 1.32182905454915 | | | | | | | |
|---|---|---|---|---|---|---|---|---|
| Category | Term | Count | % | PValue | Genes | Fold Enrichment | Benjamini | FDR |
| GOTERM_CC_DIRECT | GO:0005743~mitochondrial inner membrane | 14 | 5.555556 | 0.0024502690195 | P97450, Q811U4, Q791V5, Q8JZQ2, Q9R112, Q9QXX4, Q9CPW3, P36552, P62075, Q8BGH2, Q78IK4, Q922Q1, Q8BH59, Q9EQI8 | 2.6603587621 | 0.07943 | 0.076163 |

| Annotation Cluster 15 | Enrichment Score: 1.308051188432693 | | | | | | | |
|---|---|---|---|---|---|---|---|---|
| Category | Term | Count | % | PValue | Genes | Fold Enrichment | Benjamini | FDR |
| GOTERM_MF_DIRECT | GO:0003723~RNA binding | 22 | 8.730159 | 0.0013271132984 | O89086, Q91YT7, Q9ESL4, Q61545, Q505F5, Q60973, Q61656, Q8BYK6, P47199, Q501J6, Q8BYL4, Q8BFV2, Q9ESV0, P35922, Q9WVR4, Q05CL8, Q8BX17, P58252, Q62095, Q6NS46, P17182, O35737 | 2.1548623469 | 0.071133 | 0.070204 |

Supplementary Table 5: Results of the pathway analysis by the David tool on the proteins modulated in response to PCL beads

Annotation Cluster 1 — Enrichment Score: 4.866251651544132

| Category | Term | Count | % | PValue | Genes | Fold Enrichment | Benjamini | FDR |
|---|---|---|---|---|---|---|---|---|
| **GOTERM_CC_DIRECT** | **GO:0005739~mitochondrion** | **45** | **16.01423** | **7E-07** | **Q9D0K2, P29594, Q9D0G0, Q9WTP7, P16858, O08528, P58059, Q9QXX4, Q8BHF7, Q9CZN7, P47738, Q9D7B6, O09044, P62075, Q8BTX9, Q9QUJ7, Q9D938, Q8BTZ7, Q9JM90, P45952, P56380, Q9DCT2, Q2TBE6, P97450, P57716, O70325, Q9Z1Q9, Q9CXV1, Q3TRM8, P97333, P11499, P59017, Q9CQ54, Q9CPX7, Q8C0I1, O35855, Q9DCM0, O55126, O09111, Q9Z2I9, P06151, Q9DCS3, Q8BMS1, P07742, Q8R216** | **2.2045738045738** | **7.55378E-05** | **7.3491E-05** |
| UP_KW_DOMAIN | KW-0809~Transit peptide | 20 | 7.117438 | 8E-07 | Q9D0K2, P97450, O70325, Q9CXV1, Q9CPX7, Q8C0I1, O35855, Q9DCM0, Q8BHF7, Q9CZN7, P47738, O09111, Q9D7B6, Q9Z2I9, Q9D938, P45952, Q9DCS3, Q8BMS1, Q9DCT2, Q8R216 | 3.88515369522564 | 1.26018E-05 | 1.1814E-05 |
| UP_SEQ_FEATURE | TRANSIT:Mitochondrion | 19 | 6.761566 | 7E-06 | Q9D0K2, P97450, O70325, Q9CXV1, Q9CPX7, O35855, Q9DCM0, Q8BHF7, Q9CZN7, P47738, O09111, Q9D7B6, Q9Z2I9, Q9D938, P45952, Q9DCS3, Q8BMS1, Q9DCT2, Q8R216 | 3.5756469415006 | 0.002817439 | 0.0028071 |
| UP_KW_CELLULAR_COMPONENT | KW-0496~Mitochondrion | 33 | 11.74377 | 2E-05 | Q9D0K2, Q9D0G0, Q9WTP7, O08528, P58059, Q9QXX4, Q8BHF7, Q9CZN7, P47738, Q9D7B6, P62075, Q8BTX9, Q9QUJ7, Q9D938, Q9JM90, P45952, Q9DCT2, Q2TBE6, P97450, O70325, Q9CXV1, P97333, P59017, Q9CQ54, Q9CPX7, O35855, Q9DCM0, O55126, O09111, Q9Z2I9, Q9DCS3, Q8BMS1, Q8R216 | 2.27389688913505 | 0.000248432 | 0.0002307 |
| KEGG_PATHWAY | mmu01100:Metabolic pathways | 44 | 15.65836 | 5E-05 | Q9D0K2, P23591, Q9D6Y9, Q8BTI9, Q9DCD0, Q80VQ0, P16858, Q9R0N0, O08528, Q9R1J0, Q8BHF7, Q9CZN7, P47738, Q9D7B6, Q9QUJ7, Q8BTZ7, Q9D0F9, P45952, P56380, O35405, Q9DCT2, Q91X52, Q2TBE6, P97450, O70325, P51432, P06745, Q9CXV1, Q3TRM8, Q9CQ54, Q8C0I1, O35855, Q9DCM0, Q9R0E2, O09111, Q9Z2I9, P06151, Q8BVW0, P17182, Q9WUA3, Q9DCS3, Q8BMS1, P07742, Q8R216 | 1.81418123550948 | 0.00429911 | 0.0041648 |
| GOTERM_CC_DIRECT | GO:0005743~mitochondrial inner membrane | 15 | 5.338078 | 0.0017 | P97450, O70325, Q9CXV1, Q9D0G0, P58059, Q9QXX4, Q9CQ54, Q9CPX7, Q9CZN7, O09111, P62075, Q9D938, Q8BMS1, Q9DCT2, Q8R216 | 2.64769038701623 | 0.062687999 | 0.0609896 |

Annotation Cluster 2 — Enrichment Score: 3.4206462695154976

| Category | Term | Count | % | PValue | Genes | Fold Enrichment | Benjamini | FDR |
|---|---|---|---|---|---|---|---|---|
| GOTERM_BP_DIRECT | GO:0006457~protein folding | 11 | 3.914591 | 6E-07 | Q9WUD1, Q9QYJ3, Q9CWM4, P08003, P09103, P08113, P11499, Q9ES00, Q03958, P48428, Q8R180 | 8.61043431179178 | 0.00089979 | 0.0008968 |
| KEGG_PATHWAY | mmu04141:Protein processing in endoplasmic reticulum | 10 | 3.558719 | 0.0012 | Q9WTX6, Q9WUD1, Q9QYJ3, P08003, P09103, P08113, P11499, Q9ES00, P27612, Q8R180 | 3.81449275362319 | 0.037477924 | 0.0363067 |
| GOTERM_BP_DIRECT | GO:0034976~response to endoplasmic reticulum stress | 4 | 1.423488 | 0.0769 | P08003, P09103, Q9ES00, Q8R180 | 4.02565760031823 | 1 | 0.9972973 |

Annotation Cluster 3 — Enrichment Score: 3.3555443876499904

| Category | Term | Count | % | PValue | Genes | Fold Enrichment | Benjamini | FDR |
|---|---|---|---|---|---|---|---|---|
| GOTERM_BP_DIRECT | GO:0006457~protein folding | 11 | 3.914591 | 6E-07 | Q9WUD1, Q9QYJ3, Q9CWM4, P08003, P09103, P08113, P11499, Q9ES00, Q03958, P48428, Q8R180 | 8.61043431179178 | 0.00089979 | 0.0008968 |
| UP_KW_MOLECULAR_FUNCTION | KW-0143~Chaperone | 13 | 4.626335 | 8E-06 | Q9WU84, P70670, P11499, Q8VEH6, Q91YE6, P62075, Q9QYJ3, Q9CWM4, P08113, P09103, Q03958, P48428, Q99KD5 | 5.16380789022298 | 0.000389413 | 0.0003827 |
| GOTERM_MF_DIRECT | GO:0051082~unfolded protein binding | 7 | 2.491103 | 0.0007 | P70670, Q9QYJ3, Q9CWM4, P08113, P11499, Q9ES00, Q03958 | 6.53519580942879 | 0.039468916 | 0.0389207 |

Annotation Cluster 4 — Enrichment Score: 3.308419629174293

| Category | Term | Count | % | PValue | Genes | Fold Enrichment | Benjamini | FDR |
|---|---|---|---|---|---|---|---|---|
| KEGG_PATHWAY | mmu01250:Biosynthesis of nucleotide sugars | 7 | 2.491103 | 2E-05 | P23591, P06745, Q3TRM8, Q9R0N0, O08528, Q8BTZ7, Q9D0F9 | 12.2967200610221 | 0.002316515 | 0.0022441 |
| GOTERM_BP_DIRECT | GO:0005975~carbohydrate metabolic process | 12 | 4.270463 | 2E-05 | O09159, Q9D6Y9, P06745, Q3TRM8, P06151, Q9DCD0, Q8BVW0, P16858, Q9R0N0, O08528, Q9D0F9, Q91X52 | 5.11164895296222 | 0.012152911 | 0.0121119 |
| KEGG_PATHWAY | mmu00052:Galactose metabolism | 6 | 2.135231 | 1E-04 | Q3TRM8, Q8BVW0, Q9R0N0, O08528, Q9WUA3, Q9D0F9 | 12.5163043478261 | 0.00429911 | 0.0041648 |
| KEGG_PATHWAY | mmu00520:Amino sugar and nucleotide sugar metabolism | 7 | 2.491103 | 0.0001 | P23591, P06745, Q3TRM8, Q9R0N0, O08528, Q8BTZ7, Q9D0F9 | 9.16226200625178 | 0.00429911 | 0.0041648 |
| KEGG_PATHWAY | mmu00500:Starch and sucrose metabolism | 6 | 2.135231 | 0.0001 | Q9D6Y9, P06745, Q3TRM8, Q8BVW0, O08528, Q9D0F9 | 11.7800511508951 | 0.004864266 | 0.0047123 |
| GOTERM_BP_DIRECT | GO:0006006~glucose metabolic process | 7 | 2.491103 | 0.0003 | Q9Z0M5, Q3TRM8, Q8BVW0, P16858, O08528, Q9D0F9, Q91X52 | 7.63197586726998 | 0.076306501 | 0.0760489 |
| KEGG_PATHWAY | mmu00051:Fructose and mannose metabolism | 5 | 1.779359 | 0.0019 | P23591, Q3TRM8, O08528, Q8BTZ7, Q9WUA3 | 9.27133655394525 | 0.044368862 | 0.0429823 |

Annotation Cluster 5 — Enrichment Score: 3.2450020805674726

| Category | Term | Count | % | PValue | Genes | Fold Enrichment | Benjamini | FDR |
|---|---|---|---|---|---|---|---|---|
| KEGG_PATHWAY | mmu00010:Glycolysis / Gluconeogenesis | 10 | 3.558719 | 6E-07 | P47738, P06745, Q3TRM8, P06151, Q80VQ0, P16858, O08528, P17182, Q9WUA3, Q9D0F9 | 9.9632273415531 | 0.000144117 | 0.0001396 |
| GOTERM_BP_DIRECT | GO:0006096~glycolytic process | 7 | 2.491103 | 6E-06 | Q9Z0M5, P06745, Q3TRM8, P16858, O08528, P17182, Q9WUA3 | 15.26395173454 | 0.004315881 | 0.0043013 |
| KEGG_PATHWAY | mmu01200:Carbon metabolism | 10 | 3.558719 | 8E-05 | Q9CZN7, Q9CXV1, P06745, Q3TRM8, Q9Z2I9, Q9DCD0, P16858, O08528, P17182, Q9WUA3 | 5.51682836267817 | 0.00429911 | 0.0041648 |
| UP_KW_BIOLOGICAL_PROCESS | KW-0324~Glycolysis | 6 | 2.135231 | 8E-05 | P06745, Q3TRM8, P16858, O08528, P17182, Q9WUA3 | 13.2048681541582 | 0.006270864 | 0.0062709 |

| Category | Term | Count | % | PValue | Genes | Fold Enrichment | Benjamini | FDR |
|---|---|---|---|---|---|---|---|---|
| KEGG_PATHWAY | mmu04066:HIF-1 signaling pathway | 8 | 2.846975 | 0.0015 | Q9D4H8, Q8BTI9, Q3TRM8, P06151, P16858, O08528, P17182, Q9WUA3 | 4.68446478515128 | 0.038980272 | 0.0377621 |
| Annotation Cluster 6 | Enrichment Score: 3.2272226351499116 | | | | | | | |
| Category | Term | Count | % | PValue | Genes | Fold Enrichment | Benjamini | FDR |
| UP_SEQ_FEATURE | CROSSLNK:Glycyl lysine isopeptide (Lys-Gly) (interchain with G-Cter in SUMO2) | 25 | 8.896797 | 0.0001 | Q9QXE7, Q6P9P6, Q99NB9, Q9QZE7, P09405, P23198, Q501J6, P16858, Q8BX09, Q8CGC6, P84089, Q0VBL3, P20152, Q8K310, P97379, Q9WTM5, O35286, Q6NZF1, Q8CGZ0, Q05CL8, Q3V300, Q9DBY8, E9PVX6, O88286, Q9CWK3 | 2.40197093645369 | 0.03234816 | 0.03223 |
| UP_KW_PTM | KW-0832~Ubl conjugation | 54 | 19.21708 | 0.001 | Q61510, Q9QXE7, P97461, Q9QZE7, P09405, P23198, Q8VBW6, Q9WTX6, P51943, P16858, Q8BX09, Q9D4H8, Q0VBL3, P20152, P07356, Q8K310, O35405, Q2TBE6, P06745, O35286, P11499, Q9WTM5, Q6NZF1, Q5DQR4, Q08509, P06151, O88286, Q9CWK3, P97481, Q6P9P6, Q99NB9, Q501J6, Q8CGC6, P84089, O88712, Q9WUD1, Q9D1H7, Q9WU84, P70670, P97379, P14426, Q8CGZ0, Q05CL8, Q3V300, P58252, P22366, P83741, P17182, Q03958, P24860, Q9DBY8, Q922S8, P26369, E9PVX6 | 1.52847541563495 | 0.006408662 | 0.0057678 |
| UP_KW_PTM | KW-1017~Isopeptide bond | 39 | 13.879 | 0.0018 | Q61510, Q9QXE7, P97461, Q6P9P6, Q99NB9, Q9QZE7, P09405, P23198, Q9WTX6, Q501J6, P16858, Q8BX09, Q9D4H8, Q8CGC6, P84089, Q0VBL3, O88712, Q9WUD1, P20152, P07356, Q8K310, Q9WU84, P97379, P70670, Q9WTM5, O35286, Q6NZF1, Q8CGZ0, Q05CL8, Q3V300, P58252, P06151, P17182, Q03958, Q9DBY8, E9PVX6, O88286, Q9CWK3, P26369 | 1.64836551440812 | 0.00915832 | 0.0082425 |
| Annotation Cluster 7 | Enrichment Score: 2.7475789894014313 | | | | | | | |
| Category | Term | Count | % | PValue | Genes | Fold Enrichment | Benjamini | FDR |
| GOTERM_MF_DIRECT | GO:0000166~nucleotide binding | 39 | 13.879 | 4E-05 | Q6P9P6, Q8BTI9, Q501J6, Q9WTP7, Q9R0N0, O08528, P16879, Q6P9L6, Q8VEH6, Q8BHF7, Q9EPU0, O54984, Q9QUJ7, Q8BTZ7, Q8BU30, Q99KH8, P56380, Q2TBE6, Q9ESL4, Q9Z1Q9, Q3TRM8, Q9WTM5, O35286, P11499, Q6PA06, Q6ZQB6, Q3UX10, Q3V300, P58252, Q9Z2I9, P08113, Q91V41, P83741, Q5NC05, Q9WUA3, Q9DBY8, Q922S8, E9PVX6, P07742 | 2.00986971643842 | 0.003907905 | 0.0038536 |
| GOTERM_MF_DIRECT | GO:0005524~ATP binding | 35 | 12.45552 | 0.0001 | Q6P9P6, Q8BTI9, Q501J6, Q9WTP7, Q9R0N0, O08528, P16879, Q9ES00, Q6P9L6, Q8BHF7, Q9EPU0, Q8CFE4, Q6A028, O54984, Q9QUJ7, Q8BU30, Q99KH8, Q2TBE6, Q9ESL4, Q9Z1Q9, Q3TRM8, Q9WTM5, O35286, P11499, Q6ZQB6, Q3V300, Q9Z2I9, P08113, P83741, Q5NC05, Q9WUA3, Q9DBY8, Q922S8, E9PVX6, P07742 | 2.02097526658768 | 0.008457628 | 0.0083402 |
| UP_KW_LIGAND | KW-0547~Nucleotide-binding | 40 | 14.23488 | 0.0029 | Q6P9P6, Q8BTI9, Q501J6, Q9WTP7, P16858, Q9R0N0, O08528, P16879, Q6P9L6, Q8VEH6, Q8BHF7, Q9EPU0, O54984, Q9QUJ7, Q8BTZ7, Q8BU30, Q99KH8, P56380, Q2TBE6, Q9ESL4, Q9Z1Q9, Q3TRM8, Q9WTM5, O35286, P11499, Q6PA06, Q6ZQB6, Q3UX10, Q3V300, P58252, Q9Z2I9, P08113, Q91V41, P83741, Q5NC05, Q9WUA3, Q9DBY8, Q922S8, E9PVX6, P07742 | 1.51301475195079 | 0.067668684 | 0.0676687 |
| UP_KW_LIGAND | KW-0067~ATP-binding | 31 | 11.03203 | 0.0125 | Q6P9P6, Q8BTI9, Q501J6, Q9R0N0, O08528, P16879, Q6P9L6, Q8BHF7, Q9EPU0, O54984, Q9QUJ7, Q8BU30, Q99KH8, Q2TBE6, Q9ESL4, Q9Z1Q9, Q3TRM8, Q9WTM5, O35286, P11499, Q6ZQB6, Q3V300, Q9Z2I9, P08113, P83741, Q5NC05, Q9WUA3, Q9DBY8, Q922S8, E9PVX6, P07742 | 1.51382413930026 | 0.095603798 | 0.0956038 |
| INTERPRO | IPR027417:P-loop containing nucleoside triphosphate hydrolase | 15 | 5.338078 | 0.1277 | Q6P9P6, Q501J6, Q9WTP7, Q9WTM5, O35286, Q6PA06, Q6P9L6, Q3V300, Q9EPU0, P58252, O54984, Q91V41, Q5NC05, Q9DBY8, Q922S8 | 1.49672072072072 | 1 | 1 |
| Annotation Cluster 8 | Enrichment Score: 2.66900536393511 | | | | | | | |
| Category | Term | Count | % | PValue | Genes | Fold Enrichment | Benjamini | FDR |
| GOTERM_MF_DIRECT | GO:0003723~RNA binding | 30 | 10.67616 | 6E-07 | Q9CWY8, P97461, Q99NB9, Q9QZE7, P09405, Q501J6, O35295, Q62189, Q9EPU0, Q8CGC6, Q3V1V3, Q0VBL3, P20152, P62313, Q9DBR1, Q8K310, Q9ESL4, Q9D554, Q921I9, P97379, O35286, P70279, Q8CGZ0, Q05CL8, P58252, P63154, Q8C166, P08113, P17182, P26369 | 2.85489087321766 | 0.000318364 | 0.0003139 |
| Annotation Cluster 9 | Enrichment Score: 2.6457562493371674 | | | | | | | |
| Category | Term | Count | % | PValue | Genes | Fold Enrichment | Benjamini | FDR |
| UP_KW_BIOLOGICAL_PROCESS | KW-0324~Glycolysis | 6 | 2.135231 | 8E-05 | P06745, Q3TRM8, P16858, O08528, P17182, Q9WUA3 | 13.2048681541582 | 0.006270864 | 0.0062709 |
| KEGG_PATHWAY | mmu00052:Galactose metabolism | 6 | 2.135231 | 1E-04 | Q3TRM8, Q8BVW0, Q9R0N0, O08528, Q9WUA3, Q9D0F9 | 12.5163043478261 | 0.00429911 | 0.0041648 |
| KEGG_PATHWAY | mmu00051:Fructose and mannose metabolism | 5 | 1.779359 | 0.0019 | P23591, Q3TRM8, O08528, Q8BTZ7, Q9WUA3 | 9.27133655394525 | 0.044368862 | 0.0429823 |
| Annotation Cluster 10 | Enrichment Score: 2.3141634554540302 | | | | | | | |
| Category | Term | Count | % | PValue | Genes | Fold Enrichment | Benjamini | FDR |
| GOTERM_MF_DIRECT | GO:0051287~NAD binding | 6 | 2.135231 | 0.0005 | P47738, P06151, O88712, P16858, Q8BMS1, Q8R216 | 9.17502859944435 | 0.034528274 | 0.0340487 |
| UP_KW_LIGAND | KW-0520~NAD | 9 | 3.202847 | 0.0083 | P47738, P06151, O88712, Q80VQ0, P16858, Q9R1J0, Q8BMS1, Q9DCT2, Q8R216 | 3.06554854593014 | 0.095603798 | 0.0956038 |
| Annotation Cluster 11 | Enrichment Score: 2.306164348276061 | | | | | | | |
| Category | Term | Count | % | PValue | Genes | Fold Enrichment | Benjamini | FDR |
| GOTERM_CC_DIRECT | GO:0005681~spliceosomal complex | 10 | 3.558719 | 0.0001 | P09405, Q99NB9, Q8CGC6, Q9D554, P63154, O35286, Q5NC05, Q62189, P62313, P26369 | 5.15069823922283 | 0.008579958 | 0.0083475 |

| | | | | | | | | |
|---|---|---|---|---|---|---|---|---|
| GOTERM_BP_DIRECT | GO:0008380~RNA splicing | 12 | 4.270463 | 0.0003 | Q99NB9, Q8CGC6, Q9D554, P63154, Q501J6, O35286, Q5NC05, Q62189, P62313, Q05CL8, Q9CWK3, P26369 | 3.89717916626552 | 0.076306501 | 0.0760489 |
| GOTERM_BP_DIRECT | GO:0006397~mRNA processing | 13 | 4.626335 | 0.0007 | Q99NB9, Q9D554, Q501J6, O35286, Q62189, Q05CL8, Q8CGC6, P63154, Q5NC05, P62313, Q9DBR1, P26369, Q9CWK3 | 3.23529411764706 | 0.142494073 | 0.142013 |
| UP_KW_BIOLOGICAL_PROCESS | KW-0508~mRNA splicing | 12 | 4.270463 | 0.0016 | Q99NB9, Q8CGC6, Q9D554, P63154, Q501J6, O35286, Q5NC05, Q62189, P62313, Q05CL8, Q9CWK3, P26369 | 3.11781609195402 | 0.062474979 | 0.062475 |
| UP_KW_BIOLOGICAL_PROCESS | KW-0507~mRNA processing | 13 | 4.626335 | 0.0034 | Q99NB9, Q9D554, Q501J6, O35286, Q62189, Q05CL8, Q8CGC6, P63154, Q5NC05, P62313, Q9DBR1, P26369, Q9CWK3 | 2.66509211147851 | 0.089788703 | 0.0897887 |
| UP_KW_CELLULAR_COMPONENT | KW-0747~Spliceosome | 7 | 2.491103 | 0.0073 | Q99NB9, Q8CGC6, Q9D554, P63154, Q5NC05, Q62189, P62313 | 4.093386865126 | 0.051431317 | 0.0477577 |

| Annotation Cluster 12 | Enrichment Score: 2.2772353104159118 | | | | | | | |
|---|---|---|---|---|---|---|---|---|
| Category | Term | Count | % | PValue | Genes | Fold Enrichment | Benjamini | FDR |
| UP_SEQ_FEATURE | MOTIF:Prevents secretion from ER | 6 | 2.135231 | 0.0005 | P45878, P08003, P09103, P08113, Q9QXT0, Q9R1J0 | 9.25904365904366 | 0.076921096 | 0.07664 |
| GOTERM_CC_DIRECT | GO:0042470~melanosome | 7 | 2.491103 | 0.0007 | P57716, O08992, P08003, P09103, P08113, P11499, P07356 | 6.59804444444445 | 0.030569374 | 0.0297411 |

| Annotation Cluster 13 | Enrichment Score: 1.874667419534498 | | | | | | | |
|---|---|---|---|---|---|---|---|---|
| Category | Term | Count | % | PValue | Genes | Fold Enrichment | Benjamini | FDR |
| UP_KW_BIOLOGICAL_PROCESS | KW-0324~Glycolysis | 6 | 2.135231 | 8E-05 | P06745, Q3TRM8, P16858, O08528, P17182, Q9WUA3 | 13.2048681541582 | 0.006270864 | 0.0062709 |
| KEGG_PATHWAY | mmu04066:HIF-1 signaling pathway | 8 | 2.846975 | 0.0015 | Q9D4H8, Q8BTI9, Q3TRM8, P06151, P16858, O08528, P17182, Q9WUA3 | 4.68446478515128 | 0.038980272 | 0.0377621 |

| Annotation Cluster 14 | Enrichment Score: 1.7366826602926042 | | | | | | | |
|---|---|---|---|---|---|---|---|---|
| Category | Term | Count | % | PValue | Genes | Fold Enrichment | Benjamini | FDR |
| GOTERM_CC_DIRECT | GO:0005874~microtubule | 13 | 4.626335 | 0.0006 | Q6P9P6, Q8VC03, Q8BUK6, Q3UX10, Q62433, Q6P9L6, Q3V300, Q3UQ44, P62482, Q8BTU1, Q6A065, P48428, Q922S8 | 3.33883136542537 | 0.028351246 | 0.0275831 |
| UP_KW_CELLULAR_COMPONENT | KW-0493~Microtubule | 11 | 3.914591 | 0.0015 | Q3V300, Q6P9P6, Q8VC03, Q8BUK6, Q8BTU1, Q3UX10, Q6A065, P48428, Q62433, Q6P9L6, Q922S8 | 3.40001725327812 | 0.01604902 | 0.0149027 |
