## Supplemental Tables 6-10 for "Effects of true to life polyethylene terephthalate and polycaprolactone nanoparticles on macrophages under a repeated exposure mode"

Supplementary Table 6: List of mitochondrial proteins modulated in response to PET or PCL beads
Color code: purple, proteins modulated in response to PET particles,
blue, proteins modulated in response to PCL particles or to both particles
black, unmodulated proteins related to the selected proteins

| accession | gene_name | description | U-PET | U-PCL | ratio PET | ratio PCL |
|---|---|---|---|---|---|---|
| Q920A7 | Afg3l1 | AFG3-like protein 1 | 10 | 10 | 1.080611 | 0.95299 |
| Q8JZQ2 | Afg3l2 | AFG3-like protein 2 | 0 | 7 | 1.258789 | 1.03946 |
| Q8R404 | Mic13 | MICOS complex subunit MIC13 | 9 | 11 | 1.083776 | 0.93517 |
| Q9CRB9 | Chchd3 | MICOS complex subunit Mic19 | 12 | 10 | 1.131371 | 1.03765 |
| Q91VN4 | Chchd6 | MICOS complex subunit Mic25 | 8 | 12 | 1.166974 | 1.04073 |
| Q9DCZ4 | Apoo | MICOS complex subunit Mic26 | 3 | 9 | 1.681299 | 1.14431 |
| Q78IK4 | Apool | MICOS complex subunit Mic27 | 2 | 8 | 1.230369 | 1.11793 |
| Q8CAQ8 | Immt | MICOS complex subunit Mic60 | 8 | 12 | 1.071826 | 0.99526 |
| Q811U4 | Mfn1 | Mitofusin-1 | 1 | 6 | 1.35714 | 1.16999 |
| P84817 | Fis1 | Mitochondrial fission 1 protein | 3 | 5 | 0.748247 | 0.75658 |
| O55125 | Nipsnap1 | Protein NipSnap homolog 1 | 9 | 12 | 0.999182 | 0.93192 |
| O55126 | Nipsnap2 | Protein NipSnap homolog 2 | 9 | 0 | 1.394934 | 1.63432 |
| P11157 | Rrm2 | Ribonucleoside-diphosphate reductase subunit M2 | 12 | 8 | 0.983153 | 1.07933 |
| P29452 | Casp1 | Caspase-1 | 0 | 7 | 0.847666 | 0.91812 |
| P29594 | Casp2 | Caspase-2 | 0 | 2 | 1.16E-06 | 0.37728 |
| P70677 | Casp3 | Caspase-3 | 12 | 12 | 0.996804 | 0.96303 |
| O08738 | Casp6 | Caspase-6 | 10 | 10 | 0.972151 | 0.93431 |
| P97864 | Casp7 | Caspase-7 | 12 | 10.5 | 1.205771 | 0.77886 |
| O89110 | Casp8 | Caspase-8 | 11 | 10 | 1.037108 | 1.01757 |
| Q8C3Q9 | Casp9 | Caspase-9 | 9 | 11 | 1.489823 | 1.25674 |
| P62075 | Timm13 | Mitochondrial import inner membrane translocase subunit Tim13 | 1 | 0 | 0.77383 | 0.64944 |
| O35092 | Timm17a | Mitochondrial import inner membrane translocase subunit Tim17-A | 9 | 11 | 0.755872 | 0.65579 |
| Q9Z0V7 | Timm17b | Mitochondrial import inner membrane translocase subunit Tim17-B | 10.5 | 7 | 3.256168 | 5.38374 |
| Q9WTQ8 | Timm23 | Mitochondrial import inner membrane translocase subunit Tim23 | 12 | 8 | 1.029786 | 0.79237 |
| O35857 | Timm44 | Mitochondrial import inner membrane translocase subunit TIM44 | 6 | 9 | 1.112183 | 1.04471 |
| Q9D880 | Timm50 | Mitochondrial import inner membrane translocase subunit TIM50 | 4 | 10 | 1.05334 | 1.03881 |
| P41216 | Acsl1 | Long-chain-fatty-acid--CoA ligase 1 | 1 | 12 | 1.215821 | 0.98193 |
| Q9CZW4 | Acsl3 | Long-chain-fatty-acid--CoA ligase 3 | 11 | 10 | 1.07693 | 1.03954 |
| Q9QUJ7 | Acsl4 | Long-chain-fatty-acid--CoA ligase 4 | 2 | 2 | 0.893137 | 0.88262 |
| Q8JZR0 | Acsl5 | Long-chain-fatty-acid--CoA ligase 5 | 3 | 11 | 1.108872 | 0.96327 |
| Q8BMS1 | Hadha | Trifunctional enzyme subunit alpha, mitochondrial | 5 | 2 | 1.087173 | 1.10979 |
| Q99JY0 | Hadhb | Trifunctional enzyme subunit beta, mitochondrial | 8 | 7 | 1.229037 | 1.05968 |
| Q99KI0 | Aco2 | Aconitate hydratase, mitochondrial | 0 | 4 | 1.163546 | 1.07945 |
| Q9CZU6 | Cs | Citrate synthase, mitochondrial | 9 | 10 | 1.064165 | 0.96207 |
| O08749 | Dld | Dihydrolipoyl dehydrogenase, mitochondrial | 12 | 7 | 1.008502 | 0.92636 |

| | | | | | | |
|---|---|---|---|---|---|---|
| Q01205 | Dlst | Dihydrolipoyllysine-residue succinyltransferase component of 2-oxoglutarate dehydrogenase complex, mitochondrial | 11 | 7 | 0.932431 | 0.82795 |
| P97807 | Fh | Fumarate hydratase, mitochondrial | 2 | 8 | 1.119781 | 1.02663 |
| Q9D6R2 | Idh3a | Isocitrate dehydrogenase [NAD] subunit alpha, mitochondrial | 4 | 10 | 1.061872 | 1.00728 |
| Q68FX0 | Idh3B | Isocitrate dehydrogenase [NAD] subunit beta, mitochondrial | 5 | 5 | 1.105628 | 1.07529 |
| P41565 | Idh3g | Isocitrate dehydrogenase [NAD] subunit gamma 1, mitochondrial | 1 | 0 | 1.132687 | 1.19195 |
| P08249 | Mdh2 | Malate dehydrogenase, mitochondrial | 11 | 6 | 0.991606 | 0.92487 |
| Q60597 | Ogdh | 2-oxoglutarate dehydrogenase, mitochondrial | 12 | 4 | 1.00083 | 0.88306 |
| Q63065 | Pdk1 | [Pyruvate dehydrogenase (acetyl-transferring)] kinase isozyme 1, mitochondrial | 1 | 4 | 1.536283 | 1.31264 |
| Q9Z2I9 | Sucla2 | Succinate--CoA ligase [ADP-forming] subunit beta, mitochondrial | 5 | 0 | 1.198712 | 1.23705 |
| Q9WUM5 | Suclg1 | Succinate--CoA ligase [ADP/GDP-forming] subunit alpha, mitochondrial | 10 | 6 | 1.298965 | 1.33272 |
| Q9Z2I8 | Suclg2 | Succinate--CoA ligase [GDP-forming] subunit beta, mitochondrial | 9 | 7 | 1.125794 | 1.12168 |
| Q99LC3 | Ndufa10 | NADH dehydrogenase [ubiquinone] 1 alpha subcomplex subunit 10, mitochondrial | 9 | 9 | 0.897155 | 0.98656 |
| Q9D8B4 | Ndufa11 | NADH dehydrogenase [ubiquinone] 1 alpha subcomplex subunit 11 | 11 | 12 | 0.961218 | 0.94596 |
| Q7TMF3 | Ndufa12 | NADH dehydrogenase [ubiquinone] 1 alpha subcomplex subunit 12 | 6 | 11 | 0.816017 | 1.05121 |
| Q9ERS2 | Ndufa13 | NADH dehydrogenase [ubiquinone] 1 alpha subcomplex subunit 13 | 11 | 10 | 0.90902 | 0.91622 |
| Q9CQ75 | Ndufa2 | NADH dehydrogenase [ubiquinone] 1 alpha subcomplex subunit 2 | 8 | 7 | 0.939162 | 0.95561 |
| Q62425 | Ndufa4 | Cytochrome c oxidase subunit NDUFA4 | 7 | 4 | 0.868894 | 0.79626 |
| Q9CPP6 | Ndufa5 | NADH dehydrogenase [ubiquinone] 1 alpha subcomplex subunit 5 | 10 | 6 | 1.684145 | 1.63 |
| Q9DCJ5 | Ndufa8 | NADH dehydrogenase [ubiquinone] 1 alpha subcomplex subunit 8 | 4 | 5 | 0.747703 | 0.85589 |
| Q9DC69 | Ndufa9 | NADH dehydrogenase [ubiquinone] 1 alpha subcomplex subunit 9, mitochondrial | 9 | 9 | 0.727004 | 0.83996 |
| Q9DCS9 | Ndufb10 | NADH dehydrogenase [ubiquinone] 1 beta subcomplex subunit 10 | 6 | 6 | 0.732293 | 0.84467 |
| O09111 | Ndufb11 | NADH dehydrogenase [ubiquinone] 1 beta subcomplex subunit 11, mitochondrial | 10 | 1 | 5.99E-06 | 6.48183 |
| Q9CQZ6 | Ndufb3 | NADH dehydrogenase [ubiquinone] 1 beta subcomplex subunit 3 | 9 | 11 | 0.555626 | 0.75179 |
| Q9CQH3 | Ndufb5 | NADH dehydrogenase [ubiquinone] 1 beta subcomplex subunit 5, mitochondrial | 10 | 12.5 | 66438.65 | 1 |
| Q9CQ54 | Ndufc2 | NADH dehydrogenase [ubiquinone] 1 subunit C2 | 12 | 0 | 0.942299 | 0.79265 |
| Q91VD9 | Ndufs1 | NADH-ubiquinone oxidoreductase 75 kDa subunit, mitochondrial | 9 | 12 | 1.040132 | 0.98687 |
| Q91WD5 | Ndufs2 | NADH dehydrogenase [ubiquinone] iron-sulfur protein 2, mitochondrial | 12 | 8 | 1.004338 | 0.92557 |
| Q9DCT2 | Ndufs3 | NADH dehydrogenase [ubiquinone] iron-sulfur protein 3, mitochondrial | 12 | 1 | 0.975475 | 0.83231 |
| Q9DC70 | Ndufs7 | NADH dehydrogenase [ubiquinone] iron-sulfur protein 7, mitochondrial | 5 | 4 | 1.042276 | 1.07159 |
| Q8K3J1 | Ndufs8 | NADH dehydrogenase [ubiquinone] iron-sulfur protein 8, mitochondrial | 6 | 8 | 0.701245 | 0.89781 |
| Q91YT0 | Ndufv1 | NADH dehydrogenase [ubiquinone] flavoprotein 1, mitochondrial | 10 | 9 | 0.874932 | 0.86626 |
| Q9D6J6 | Ndufv2 | NADH dehydrogenase [ubiquinone] flavoprotein 2, mitochondrial | 11 | 3 | 0.975228 | 0.85815 |

| | | | | | | |
|---|---|---|---|---|---|---|
| Q8K2B3 | Sdha | Succinate dehydrogenase [ubiquinone] flavoprotein subunit, mitochondrial | 7 | 8 | 1.050519 | 1.06078 |
| Q9CQA3 | Sdhb | Succinate dehydrogenase [ubiquinone] iron-sulfur subunit, mitochondrial | 5 | 6 | 1.063702 | 1.04547 |
| Q9CZB0 | Sdhc | Succinate dehydrogenase cytochrome b560 subunit, mitochondrial | 10 | 8 | 1.047129 | 0.88885 |
| Q9CXV1 | Sdhd | Succinate dehydrogenase [ubiquinone] cytochrome b small subunit, mitochondrial | 6 | 1 | 0.752464 | 0.59362 |
| Q9CZ13 | Uqcrc1 | Cytochrome b-c1 complex subunit 1, mitochondrial | 12 | 11 | 1.023929 | 1.02749 |
| Q9DB77 | Uqcrc2 | Cytochrome b-c1 complex subunit 2, mitochondrial | 11 | 12 | 0.948391 | 0.96776 |
| P99028 | Uqcrh | Cytochrome b-c1 complex subunit 6, mitochondrial | 9 | 3 | 0.857398 | 0.78724 |
| Q7TQ16 | Uqcrq | Cytochrome b-c1 complex subunit 8 | 1 | 4 | 4.312054 | 2.99275 |
| Q8R1I1 | Uqcr10 | Cytochrome b-c1 complex subunit 9 | 9 | 10 | 0.956969 | 0.81713 |
| Q9CR68 | Uqcrfs1 | Cytochrome b-c1 complex subunit Rieske, mitochondrial | 10 | 7 | 1.080178 | 1.12872 |
| P15999 | Atp5f1a | ATP synthase subunit alpha, mitochondrial | 12 | 12 | 1.069406 | 1.0081 |
| Q03265 | Atp5f1a | ATP synthase subunit alpha, mitochondrial | 11 | 9 | 1.069067 | 1.02632 |
| P56480 | Atp5f1b | ATP synthase subunit beta, mitochondrial | 4 | 5 | 1.100594 | 1.0957 |
| Q91VR2 | Atp5f1c | ATP synthase subunit gamma, mitochondrial | 3 | 4 | 0.905526 | 0.89858 |
| Q9D3D9 | Atp5f1d | ATP synthase subunit delta, mitochondrial | 12 | 11 | 1.010425 | 0.96147 |
| O35143 | ATP5IF1 | ATPase inhibitor, mitochondrial | 8 | 9 | 0.911641 | 0.94799 |
| P56383 | Atp5mc2 | ATP synthase F(0) complex subunit C2, mitochondrial | 12 | 6 | 1.058203 | 1.35626 |
| Q78IK2 | Atp5md | Up-regulated during skeletal muscle growth protein 5 | 11 | 6 | 0.883645 | 0.91062 |
| Q06185 | Atp5me | ATP synthase subunit e, mitochondrial | 11 | 11 | 0.810853 | 0.71916 |
| P56135 | Atp5mf | ATP synthase subunit f, mitochondrial | 12 | 8 | 1.029971 | 0.89484 |
| Q9CQQ7 | Atp5pb | ATP synthase F(0) complex subunit B1, mitochondrial | 10 | 9 | 0.952016 | 0.90892 |
| Q9DCX2 | Atp5pd | ATP synthase subunit d, mitochondrial | 11 | 6 | 0.948231 | 0.88524 |
| P97450 | Atp5pf | ATP synthase-coupling factor 6, mitochondrial | 0 | 0 | 0.657351 | 0.59807 |
| Q9DB20 | Atp5po | ATP synthase subunit O, mitochondrial | 4 | 11 | 0.920919 | 0.96149 |
| Q9D7B6 | Acad8 | Isobutyryl-CoA dehydrogenase, mitochondrial | 7 | 0 | 0.870262 | 0.80093 |
| P45952 | Acadm | Medium-chain specific acyl-CoA dehydrogenase, mitochondrial | 5 | 1 | 0.941499 | 0.87692 |
| Q9DBL1 | Acadsb | Short/branched chain specific acyl-CoA dehydrogenase, mitochondrial | 2 | 4 | 1.236735 | 1.18782 |
| Q8JZQ2 | Afg3l2 | AFG3-like protein 2 | 0 | 7 | 1.258789 | 1.03946 |
| Q9WTP7 | Ak3 | GTP:AMP phosphotransferase AK3, mitochondrial | 7 | 0 | 0.843324 | 0.77651 |
| P47738 | Aldh2 | Aldehyde dehydrogenase, mitochondrial | 10 | 2 | 1.041654 | 1.06784 |
| Q9Z1T1 | Ap3b1 | AP-3 complex subunit beta-1 | 2 | 3 | 0.822236 | 0.86935 |
| Q3KRE0 | Atad3 | ATPase family AAA domain-containing protein 3 | 0 | 0 | 6.910883 | 4.4941 |
| Q07813 | Bax | Apoptosis regulator BAX | 0 | 5 | 1.173514 | 1.08352 |
| O35855 | Bcat2 | Branched-chain-amino-acid aminotransferase, mitochondrial | 12 | 2 | 0.992419 | 0.91697 |
| O55028 | Bckdk | [3-methyl-2-oxobutanoate dehydrogenase [lipoamide]] kinase, mitochondrial | 2 | 6 | 1.326462 | 1.25419 |
| P59017 | Bcl2l13 | Bcl-2-like protein 13 | 5 | 0 | 1.236793 | 1.33715 |
| Q9QYB1 | Clic4 | Chloride intracellular channel protein 4 | 0 | 3 | 1.235342 | 1.14544 |
| P36552 | Cpox | Oxygen-dependent coproporphyrinogen-III oxidase, mitochondrial | 2 | 4 | 2.000694 | 1.84757 |

| | | | | | | |
|---|---|---|---|---|---|---|
| P53395 | Dbt | Lipoamide acyltransferase component of branched-chain alpha-keto acid dehydrogenase complex, mitochondrial | 0 | 11 | 1.331907 | 1.07914 |
| Q9DCM0 | Ethe1 | Persulfide dioxygenase ETHE1, mitochondrial | 1 | 1 | 0.8008 | 0.82926 |
| P19096 | Fasn | Fatty acid synthase | 1 | 6 | 0.908504 | 1.03708 |
| Q8BTZ7 | Gmppb | Mannose-1-phosphate guanyltransferase beta | 0 | 0 | 0.713261 | 0.57899 |
| O70325 | Gpx4 | Phospholipid hydroperoxide glutathione peroxidase | 9 | 1 | 0.942623 | 0.87575 |
| P97576 | Grpel1 | GrpE protein homolog 1, mitochondrial | 1 | 1 | 2.348022 | 2.05865 |
| Q99L13 | Hibadh | 3-hydroxyisobutyrate dehydrogenase, mitochondrial | 2 | 3 | 1.368612 | 1.28463 |
| Q8VH49 | Higd1a | HIG1 domain family member 1A, mitochondrial | 8 | 0 | 1.35525 | 1.58089 |
| Q8BTX9 | Hsdl1 | Inactive hydroxysteroid dehydrogenase-like protein 1 | 9 | 1 | 0.916798 | 0.82609 |
| P38647 | Hspa9 | Stress-70 protein, mitochondrial | 1 | 6 | 1.086427 | 0.97621 |
| P26772 | Hspe1 | 10 kDa heat shock protein, mitochondrial | 9 | 2 | 0.939016 | 0.74435 |
| Q8BIJ6 | Iars2 | Isoleucine--tRNA ligase, mitochondrial | 2 | 7 | 1.133793 | 1.04804 |
| Q8CGK3 | Lonp1 | Lon protease homolog, mitochondrial | 1 | 3 | 1.204599 | 1.12028 |
| Q922Q1 | Marc2 | Mitochondrial amidoxime reducing component 2 | 1 | 8 | 1.240946 | 1.12051 |
| Q9DCS3 | Mecr | Enoyl-[acyl-carrier-protein] reductase, mitochondrial | 11 | 2 | 2.431086 | 2.01691 |
| Q9EQI8 | Mrpl46 | 39S ribosomal protein L46, mitochondrial | 2 | 8 | 1.346841 | 1.06539 |
| Q9CPW3 | Mrpl54 | 39S ribosomal protein L54, mitochondrial | 1 | 4 | 0.171152 | 0.23938 |
| Q9CPX7 | Mrps16 | 28S ribosomal protein S16, mitochondrial | 8 | 2 | 0.887471 | 0.85123 |
| P58059 | Mrps21 | 28S ribosomal protein S21, mitochondrial | 10 | 0 | 1220790 | 4083027 |
| Q9D0G0 | Mrps30 | 28S ribosomal protein S30, mitochondrial | 3 | 2 | 1.979207 | 2.13382 |
| Q791V5 | Mtch2 | Mitochondrial carrier homolog 2 | 1 | 4 | 1.243205 | 1.09326 |
| P16332 | Mut | Methylmalonyl-CoA mutase, mitochondrial | 0 | 5 | 1.178573 | 0.89649 |
| Q9JK81 | Myg1 | UPF0160 protein MYG1, mitochondrial | 1 | 3 | 0.808369 | 0.79489 |
| P57716 | Ncstn | Nicastrin | 7 | 1 | 0.849643 | 0.82751 |
| P97333 | Nrp1 | Neuropilin-1 | 2 | 2 | 0.639758 | 0.78518 |
| P56380 | Nudt2 | Bis(5'-nucleosyl)-tetraphosphatase [asymmetrical] | 11 | 0 | 1.073931 | 0.48535 |
| P29758 | Oat | Ornithine aminotransferase, mitochondrial | 0 | 8 | 1.224597 | 1.06871 |
| Q9D0K2 | Oxct1 | Succinyl-CoA:3-ketoacid coenzyme A transferase 1, mitochondrial | 4 | 0 | 1.146905 | 1.18162 |
| Q8BHF7 | Pgs1 | CDP-diacylglycerol--glycerol-3-phosphate 3-phosphatidyltransferase, mitochondrial | 4 | 0 | 0.859968 | 0.4299 |
| Q2TBE6 | Pi4k2a | Phosphatidylinositol 4-kinase type 2-alpha | 4 | 2 | 0.867382 | 0.85717 |
| Q9Z2Y8 | Plpbp | Pyridoxal phosphate homeostasis protein | 2 | 7 | 0.898519 | 0.91761 |
| Q91YU8 | Ppan | Suppressor of SWI4 1 homolog | 5 | 0 | 1.420845 | 1.6367 |
| Q61207 | Psap | Prosaposin | 1 | 3 | 0.851105 | 0.78345 |
| P07742 | Rrm1 | Ribonucleoside-diphosphate reductase large subunit | 2 | 2 | 0.43598 | 0.48365 |
| Q8BGH2 | Samm50 | Sorting and assembly machinery component 50 homolog | 2 | 4 | 1.090996 | 1.05929 |
| Q6AYE2 | Sh3glb1 | Endophilin-B1 | 2 | 1 | 0.845924 | 0.78013 |
| Q9CZN7 | Shmt2 | Serine hydroxymethyltransferase, mitochondrial | 9 | 1 | 0.952585 | 0.88238 |
| Q8R216 | Sirt4 | NAD-dependent protein lipoamidase sirtuin-4, mitochondrial | 4 | 2 | 1.088733 | 0.81409 |
| Q8BH59 | Slc25a12 | Calcium-binding mitochondrial carrier protein Aralar1 | 1 | 4 | 1.122127 | 1.14339 |
| Q9QXX4 | Slc25a13 | Calcium-binding mitochondrial carrier protein Aralar2 | 0 | 0 | 1.228629 | 1.18939 |
| O09044 | Snap23 | Synaptosomal-associated protein 23 | 9 | 2 | 0.95954 | 0.79128 |
| Q9R112 | Sqor | Sulfide:quinone oxidoreductase, mitochondrial | 1 | 6 | 1.20952 | 0.90257 |
| Q9JM90 | Stap1 | Signal-transducing adaptor protein 1 | 1 | 1 | 1.454641 | 1.48956 |
| Q9WUD1 | Stub1 | STIP1 homology and U box-containing protein 1 | 0 | 0 | 0.800094 | 0.81634 |
| B0BN86 | Tmem11 | Transmembrane protein 11, mitochondrial | 5 | 2 | 1.662182 | 1.56218 |
| Q9D938 | Tmem160 | Transmembrane protein 160 | 7 | 0 | 0.94453 | 0.82729 |
| Q9CZR8 | Tsfm | Elongation factor Ts, mitochondrial | 2 | 11 | 1.105552 | 0.93823 |

| | | | | | | |
|---|---|---|---|---|---|---|
| Q62465 | Vat1 | Synaptic vesicle membrane protein VAT-1 homolog | 1 | 6 | 1.421408 | 1.27461 |
| Q8BYL4 | Yars2 | Tyrosine--tRNA ligase, mitochondrial | 0 | 8 | 1.53539 | 1.10869 |

Supplementary Table 7: List of lysosomal proteins modulated in response to PET or PCL beads
Color code: purple, proteins modulated in response to PET particles,
blue, proteins modulated in response to PCL particles or to both particles
black, unmodulated proteins related to the selected proteins

| accession | gene_name | description | U-PET | U-PCL | ratio PET | ratio PCL |
|---|---|---|---|---|---|---|
| B2RUP2 | Unc13d | Protein unc-13 homolog D | 2 | 5 | 0.474384793 | 0.487479974 |
| O08585 | Clta | Clathrin light chain A | 2 | 2 | 0.845080247 | 0.819177705 |
| O09159 | Man2b1 | Lysosomal alpha-mannosidase | 1 | 2 | 0.744681195 | 0.823653767 |
| O35643 | Ap1b1 | AP-1 complex subunit beta-1 | 2 | 1 | 1.218469657 | 1.297335418 |
| O35657 | Neu1 | Sialidase-1 | 0 | 10 | 297133.478 | 66227.412 |
| O88653 | Lamtor3 | Ragulator complex protein LAMTOR3 | 3 | 2 | 2.496898415 | 2.150735763 |
| O89001 | Cpd | Carboxypeptidase D | 0 | 8 | 1.428179619 | 1.08448 |
| P0C1X8 | Aak1 | AP2-associated protein kinase 1 | 3 | 1 | 1.934774549 | 2.052667043 |
| P17439 | Gba | Glucosylceramidase | 0 | 5 | 0.789816017 | 0.863882824 |
| P59438 | Hps5 | Hermansky-Pudlak syndrome 5 protein homolog | 4 | 1 | 0.638241902 | 0.493741056 |
| P97821 | Ctsc | Dipeptidyl peptidase 1 | 0 | 0 | 0.766897845 | 0.812396875 |
| P98078 | Dab2 | Disabled homolog 2 | 12 | 2 | 1.044741186 | 1.203047993 |
| Q58A65 | Spag9 | C-Jun-amino-terminal kinase-interacting protein 4 | 0 | 2 | 1.433429539 | 1.27347144 |
| Q61207 | Psap | Prosaposin | 1 | 3 | 0.851104995 | 0.783453497 |
| Q62885 | Unc119 | Protein unc-119 homolog A | 0 | 5 | 684030.78 | 405474.848 |
| Q80V94 | Ap4e1 | AP-4 complex subunit epsilon-1 | 1 | 1 | 0.363755297 | 0.380027332 |
| Q8BVW0 | Ganc | Neutral alpha-glucosidase C | 5 | 0 | 0.872759365 | 0.737697034 |
| Q8CFE4 | Scyl2 | SCY1-like protein 2 | 5 | 2 | 1.852776314 | 1.975235533 |
| Q8R0H9 | Gga1 | ADP-ribosylation factor-binding protein GGA1 | 2 | 8 | 1.304538364 | 1.085384048 |
| Q8R1T1 | Chmp7 | Charged multivesicular body protein 7 | 2 | 6 | 1.268432849 | 1.212162433 |
| Q91VZ6 | Smap1 | Stromal membrane-associated protein 1 | 4 | 1 | 2.585185288 | 3.039094765 |
| Q9CWK8 | Snx2 | Sorting nexin-2 | 2 | 9 | 0.889769423 | 0.975427282 |
| Q9D1J1 | Necap2 | Adaptin ear-binding coat-associated protein 2 | 2 | 2 | 1.250768618 | 1.264595358 |
| Q9Z0M5 | Lipa | Lysosomal acid lipase/cholesteryl ester hydrolase | 0 | 2 | 1.914859801 | 1.214731069 |
| Q9Z1T1 | Ap3b1 | AP-3 complex subunit beta-1 | 2 | 3 | 0.82223571 | 0.86934834 |

Supplementary Table 8: List of proteins imlicated in the immune response and modulated in response to PET or PCL beads
Color code: purple, proteins modulated in response to PET particles,
blue, proteins modulated in response to PCL particles or to both particles
black, unmodulated proteins related to the selected proteins

| accession | gene_name | description | U-PET | U-PCL | ratio PET | ratio PCL |
|---|---|---|---|---|---|---|
| Q66H39 | Abcf3 | ATP-binding cassette sub-family F member 3 | 2 | 4 | 0.352262682 | 0.394620289 |
| Q5SSL4 | Abr | Active breakpoint cluster region-related protein | 9 | 0 | 1.333625676 | 1.59445803 |
| P58366 | Ankh | Progressive ankylosis protein homolog | 1 | 8 | 0.35534458 | 0.681371758 |
| P98086 | C1qa | Complement C1q subcomponent subunit A | 0 | 10 | 0.259047764 | 0.959089809 |
| P30993 | C5ar1 | C5a anaphylatoxin chemotactic receptor 1 | 1 | 7 | 0.587312456 | 0.785079709 |
| P10810 | Cd14 | Monocyte differentiation antigen CD14 | 0 | 9 | 0.805617142 | 0.953300841 |
| P21855 | Cd72 | B-cell differentiation antigen CD72 | 0 | 8 | 1.427391559 | 0.970851552 |
| P13233 | Cnp | 2',3'-cyclic-nucleotide 3'-phosphodiesterase | 2 | 8 | 0.402805573 | 0.87581541 |
| O35286 | Dhx15 | Pre-mRNA-splicing factor ATP-dependent RNA helicase DHX15 | 4 | 2 | 1.036743978 | 1.052450538 |
| D4A2Z8 | Dhx36 | ATP-dependent DNA/RNA helicase DHX36 | 4 | 2 | 0.689492222 | 0.614982144 |
| P26151 | Fcgr1 | High affinity immunoglobulin gamma Fc receptor I | 0 | 8 | 0.741546034 | 0.928497751 |
| A0A0B4J1G0 | Fcgr4 | Low affinity immunoglobulin gamma Fc region receptor IV | 2 | 0 | 0.781891909 | 0.784101941 |
| P16879 | Fes | Tyrosine-protein kinase Fes/Fps | 8 | 2 | 0.878851396 | 0.758801952 |
| P97379 | G3bp2 | Ras GTPase-activating protein-binding protein 2 | 10 | 2 | 1.065331223 | 0.761742736 |
| P14426 | H2-D1 | H-2 class I histocompatibility antigen, D-K alpha chain | 11 | 0 | 1.021886393 | 0.829348237 |
| P01901 | H2-K1 | H-2 class I histocompatibility antigen, K-B alpha chain | 8 | 1 | 1.033555019 | 0.848710901 |
| Q9R002 | Ifi202 | Interferon-activable protein 202 | 5 | 0 | 0.534327593 | 0.288709616 |
| P19182 | Ifrd1 | Interferon-related developmental regulator 1 | 10 | 2 | 1.74994E-05 | 6.002115669 |
| Q8R4K2 | Irak4 | Interleukin-1 receptor-associated kinase 4 | 2 | 4 | 0.885349736 | 0.907116033 |
| Q8K1T1 | Lrrc25 | Leucine-rich repeat-containing protein 25 | 0 | 5 | 0.161127246 | 0.50963365 |
| P70202 | Lxn | Latexin | 8 | 2 | 0.846737901 | 0.846111363 |
| P08905 | Lyz2 | Lysozyme C-2 | 0 | 5 | 0.692962473 | 0.816514011 |
| Q8K310 | Matr3 | Matrin-3 | 9 | 2 | 1.039419365 | 0.933359522 |
| P34960 | Mmp12 | Macrophage metalloelastase | 0 | 6 | 3.772410843 | 1.624721991 |
| O35682 | Myadm | Myeloid-associated differentiation marker | 8 | 0 | 1.849866338 | 2.626659178 |
| P22366 | Myd88 | Myeloid differentiation primary response protein MyD88 | 6 | 2 | 1.759110161 | 1.632304656 |
| Q9JHG7 | Pik3cg | Phosphatidylinositol 4,5-bisphosphate 3-kinase catalytic subunit gamma isoform | 1 | 5 | 0.738138161 | 0.83933569 |
| O35405 | Pld3 | 5'-3' exonuclease PLD3 | 0 | 0 | 0.692442333 | 0.610357767 |
| Q6ZQB6 | Ppip5k2 | Inositol hexakisphosphate and diphosphoinositol-pentakisphosphate kinase 2 | 2 | 2 | 0.611501678 | 0.604537979 |
| Q5XIS9 | Prkd2 | Serine/threonine-protein kinase D2 | 0 | 2 | 3.235099519 | 3.3586563 |
| P97814 | Pstpip1 | Proline-serine-threonine phosphatase-interacting protein 1 | 0 | 0 | 0.851694614 | 0.824913873 |
| Q9QVP9 | Ptk2b | Protein-tyrosine kinase 2-beta | 2 | 6 | 1.196925433 | 1.112281283 |
| Q4G075 | Serpinb1a | Leukocyte elastase inhibitor A | 2 | 8 | 1.266558686 | 0.887388584 |
| P97797 | Sirpa | Tyrosine-protein phosphatase non-receptor type substrate 1 | 1 | 4 | 0.85049919 | 0.880949953 |
| Q6P9P0 | Slf2 | SMC5-SMC6 complex localization factor protein 2 | 8 | 1 | 0.750120108 | 0.323522715 |
| O88838 | Spsb2 | SPRY domain-containing SOCS box protein 2 | 1 | 1 | 0.785557952 | 0.778891987 |
| Q91YX0 | Themis2 | Protein THEMIS2 | 0 | 0 | 0.599566946 | 0.622222205 |
| B2RUP2 | Unc13d | Protein unc-13 homolog D | 2 | 5 | 0.474384793 | 0.487479974 |
| Q91YT7 | Ythdf2 | YTH domain-containing family protein 2 | 1 | 6 | 0.70043023 | 0.761101395 |
| P42082 | Cd86 | T-lymphocyte activation antigen CD86 | 4 | 6 | 1.11891693 | 1.089707166 |
| Q8K4Q8 | Colec12 | Collectin-12 | 9 | 10 | 0.778159606 | 0.913755886 |
| P30204 | Msr1 | Macrophage scavenger receptor types I and II | 5 | 12 | 0.838836334 | 0.999166223 |
| Q9QUN7 | Tlr2 | Toll-like receptor 2 | 3 | 3 | 0.753059515 | 0.76250414 |
| P58681 | Tlr7 | Toll-like receptor 7 | 3 | 9 | 0.856989788 | 1.118617145 |

Supplementary Table 9: List of mitochondrial proteins modulated in response to PET or PCL beads
Color code: purple, proteins modulated in response to PET particles,
blue, proteins modulated in response to PCL particles or to both particles
black, unmodulated proteins related to the selected proteins

| accession | gene_name | description | U-PET | U-PCL | ratio PET | ratio PCL |
|---|---|---|---|---|---|---|
| O08528 | Hk2 | Hexokinase-2 | 4 | 2 | 1.127010079 | 1.235097712 |
| O08730 | Gyg1 | Glycogenin-1 | 0 | 0 | 0.525052494 | 0.386349234 |
| P06151 | Ldha | L-lactate dehydrogenase A chain | 0 | 1 | 1.164587388 | 1.135338252 |
| P06745 | Gpi | Glucose-6-phosphate isomerase | 4 | 0 | 1.134611488 | 1.263128768 |
| P12382 | Pfkl | ATP-dependent 6-phosphofructokinase, liver type | 2 | 7 | 1.217213304 | 1.107699532 |
| P16858 | Gapdh | Glyceraldehyde-3-phosphate dehydrogenase | 5 | 1 | 1.293062535 | 1.287459261 |
| P17182 | Eno1 | Alpha-enolase | 0 | 0 | 1.351741063 | 1.257977291 |
| P54645 | Prkaa1 | 5'-AMP-activated protein kinase catalytic subunit alpha-1 | 11 | 1 | 1.094489698 | 1.257316728 |
| Q3TRM8 | Hk3 | Hexokinase-3 | 6 | 2 | 0.925072916 | 0.879470309 |
| Q7TSV4 | Pgm2 | Phosphoglucomutase-2 | 2 | 4 | 0.721375029 | 0.778027805 |
| Q80VQ0 | Aldh3b1 | Aldehyde dehydrogenase family 3 member B1 | 6 | 2 | 0.885829476 | 0.745268563 |
| Q91X52 | Dcxr | L-xylulose reductase | 0 | 2 | 0.551441559 | 0.552812773 |
| Q9D0F9 | Pgm1 | Phosphoglucomutase-1 | 2 | 1 | 1.197934066 | 1.158500016 |
| Q9DCD0 | Pgd | 6-phosphogluconate dehydrogenase, decarboxylating | 5 | 1 | 1.151942903 | 1.199962009 |
| Q9R0N0 | Galk1 | Galactokinase | 1 | 0 | 1.65626098 | 1.640265934 |
| Q9WUA3 | Pfkp | ATP-dependent 6-phosphofructokinase, platelet type | 0 | 0 | 1.274484215 | 1.156182378 |
| P17710 | Hk1 | Hexokinase-1 | 7 | 6 | 1.09115837 | 1.065232356 |
| P47857 | Pfkm | ATP-dependent 6-phosphofructokinase, muscle type | 12 | 10 | 1.002518447 | 0.955893112 |
| P17183 | Eno2 | Gamma-enolase | 7 | 5 | 1.329479544 | 1.189261073 |
| P15429 | Eno3 | Beta-enolase | 7 | 5 | 1.26098966 | 1.153710907 |
| Q93092 | Taldo1 | Transaldolase | 10 | 12 | 1.021490778 | 1.014933296 |
| P05064 | Aldoa | Fructose-bisphosphate aldolase A | 5 | 7 | 1.116960008 | 1.075353276 |
| P05063 | Aldoc | Fructose-bisphosphate aldolase C | 7 | 12 | 1.126192366 | 1.038962445 |
| P17751 | Tpi1 | Triosephosphate isomerase | 10 | 6 | 0.994715088 | 1.067399745 |
| P09411 | Pgk1 | Phosphoglycerate kinase 1 | 5 | 3 | 1.135188805 | 1.1396633 |
| P09041 | Pgk2 | Phosphoglycerate kinase 2 | 7 | 4 | 1.296020152 | 1.218522407 |
| Q00612 | G6pdx | Glucose-6-phosphate 1-dehydrogenase X | 10 | 5 | 1.048390502 | 1.046534361 |
| Q8VEE0 | Rpe | Ribulose-phosphate 3-epimerase | 6 | 11 | 0.788671203 | 0.945483238 |
| P47968 | Rpia | Ribose-5-phosphate isomerase | 9 | 12 | 0.961442219 | 0.981427563 |
| P52480 | Pkm | Pyruvate kinase PKM | 4 | 4 | 1.148712202 | 1.131410824 |

Supplementary Table 10: List of proteins implicated in the response to a chemical stress and modulated in response to PET or PCL beads

Color code: purple, proteins modulated in response to PET particles,
blue, proteins modulated in response to PCL particles or to both particles
black, unmodulated proteins related to the selected proteins

| accession | gene_name | description | U-PET | U-PCL | ratio PET | ratio PCL |
|---|---|---|---|---|---|---|
| O88653 | Lamtor3 | Ragulator complex protein LAMTOR3 | 3 | 2 | 2.496898415 | 2.150735763 |
| P10649 | Gstm1 | Glutathione S-transferase Mu 1 | 1 | 7 | 1.460251409 | 1.252742271 |
| P20291 | Alox5ap | Arachidonate 5-lipoxygenase-activating protein | 6 | 0 | 4.48832198 | 7.890807977 |
| P29594 | Casp2 | Caspase-2 | 0 | 2 | 1.15772E-06 | 0.377282231 |
| P47199 | Cryz | Quinone oxidoreductase | 0 | 3 | 1.315733833 | 1.512117278 |
| Q62433 | Ndrg1 | Protein NDRG1 | 4 | 0 | 1.290708607 | 1.376083758 |
| Q6AYE2 | Sh3glb1 | Endophilin-B1 | 2 | 1 | 0.845924324 | 0.780127767 |
| Q80VQ0 | Aldh3b1 | Aldehyde dehydrogenase family 3 member B1 | 6 | 2 | 0.885829476 | 0.745268563 |
| Q8CG45 | Akr7a2 | Aflatoxin B1 aldehyde reductase member 2 | 5 | 1 | 1.927222641 | 2.35954965 |
| Q8QZS3 | Flcn | Folliculin | 9 | 2 | 1.605046136 | 1.865095042 |
| Q8R180 | Ero1a | ERO1-like protein alpha | 0 | 0 | 1.559348995 | 1.225056774 |
| Q8R2E9 | Ero1b | ERO1-like protein beta | 0 | 5 | 1.651736102 | 1.190024756 |
| Q91Z53 | Grhpr | Glyoxylate reductase/hydroxypyruvate reductase | 0 | 6 | 1.293014843 | 1.101679275 |
| Q921W4 | Cryzl1 | Quinone oxidoreductase-like protein 1 | 10 | 1 | 1.159639835 | 2.481803098 |
| Q923V8 | Selenof | Selenoprotein F | 2 | 6 | 1.346149664 | 1.20288711 |
| Q99KH8 | Stk24 | Serine/threonine-protein kinase 24 | 5 | 1 | 1.358815841 | 1.520983117 |
| Q9CPU4 | Mgst3 | Microsomal glutathione S-transferase 3 | 2 | 10 | 1.290536185 | 1.156733275 |
| Q9QYJ3 | Dnajb1 | DnaJ homolog subfamily B member 1 | 11 | 1 | 0.991523016 | 1.399972024 |
| Q9WU84 | Ccs | Copper chaperone for superoxide dismutase | 2 | 0 | 8.276390442 | 11.15089553 |
| P47738 | Aldh2 | Aldehyde dehydrogenase, mitochondrial | 10 | 2 | 1.041654247 | 1.067839719 |
| P08113 | Hsp90b1 | Endoplasmin | 5 | 0 | 1.127354499 | 1.08859257 |
| P11499 | Hsp90ab1 | Heat shock protein HSP 90-beta | 4 | 2 | 1.079132798 | 1.095208353 |
| Q9CQM5 | Txndc17 | Thioredoxin domain-containing protein 17 | 5 | 1 | 1.107557377 | 1.139556399 |
| Q66HF8 | Aldh1b1 | Aldehyde dehydrogenase X, mitochondrial | 3 | 1 | 1.118175523 | 1.197826624 |
| Q91X52 | Dcxr | L-xylulose reductase | 0 | 2 | 0.551441559 | 0.552812773 |
| O55242 | Sigmar1 | Sigma non-opioid intracellular receptor 1 | 2 | 0 | 0.717406216 | 0.5519129 |
